# Exploring natural odour landscapes: A case study with implications for human-biting insects

**DOI:** 10.1101/2023.05.08.539789

**Authors:** Jessica L. Zung, Sumer M. Kotb, Carolyn S. McBride

## Abstract

The natural world is full of odours—blends of volatile chemicals emitted by potential sources of food, social partners, predators, and pathogens. Animals rely heavily on these signals for survival and reproduction. Yet we remain remarkably ignorant of the composition of the chemical world. How many compounds do natural odours typically contain? How often are those compounds shared across stimuli? What are the best statistical strategies for discrimination? Answering these questions will deliver crucial insight into how brains can most efficiently encode olfactory information. Here, we undertake the first large-scale survey of vertebrate body odours, a set of stimuli relevant to blood-feeding arthropods. We quantitatively characterize the odour of 64 vertebrate species (mostly mammals), representing 29 families and 13 orders. We confirm that these stimuli are complex blends of relatively common, shared compounds and show that they are much less likely to contain unique components than are floral odours—a finding with implications for olfactory coding in blood feeders and floral visitors. We also find that vertebrate body odours carry little phylogenetic information, yet show consistency within a species. Human odour is especially unique, even compared to the odour of other great apes. Finally, we use our newfound understanding of odour-space statistics to make specific predictions about olfactory coding, which align with known features of mosquito olfactory systems. Our work provides one of the first quantitative descriptions of a natural odour space and demonstrates how understanding the statistics of sensory environments can provide novel insight into sensory coding and evolution.

## Introduction

How naturalistic scenes are encoded by the brain is a longstanding fundamental question in sensory neuroscience. In order for animals to navigate complex environments, sensory systems must compress a flood of information, amplifying the salience of informative cues to direct appropriate behaviours. Information theory tells us that the best way to compress sensory information depends strongly on the structure of the stimulus space. Indeed, studying the statistics of natural images (Field 1987; Ruderman 1994; Simoncelli and Olshausen 2001) and sounds (Theunissen and Elie 2014) has provided key insight into how visual and auditory systems have evolved to process those stimuli optimally. Yet compared to visual and auditory scenes, we know very little about the statistics of natural *olfactory* scenes (Laurent 1999). This ignorance is partly due to our own perceptual limitations; most people find it extremely difficult to describe odours (Yeshurun and Sobel 2010; but see Majid and Burenhult 2014), and humans are very poor at identifying the individual components of odour blends (Jinks and Laing 2001; Marshall et al. 2006). We are therefore “blind” to the structure of odour space and have no intuitive understanding of how this space is organized.

Progress in understanding natural odours has also been hindered by the technical challenges of capturing and analysing odour, especially the complex blends that constitute most natural odours. While cameras and microphones allow us to capture visual and auditory information with relative ease, taking an “olfactory snapshot” is a complicated endeavour that involves trapping the odour, separating it into its constituent components (often dozens of them), then identifying and quantifying individual compounds. Each step in this sequence poses its own challenges that introduce biases into the type and amount of compounds sampled. Because of these challenges, it is difficult to compare data across studies, which predominantly focus on individual odours and often report only presence/absence data. Very few studies have quantitatively characterized broad odour spaces (Mansourian et al. 2016; Kantsa et al. 2018; Dweck et al. 2018; Khallaf et al. 2021), leaving us with only glimpses of the rich information available in entire olfactory scenes. To extend the visual analogy, what we know about the olfactory world is akin to having only close-up images of isolated objects, often in grainy black and white.

The dearth of quantitative odour data leaves us ignorant of basic features of odour spaces. Consider an animal navigating the world by smell. Out of the billions of theoretically possible small organic molecules (Ruddigkeit et al. 2012), how many does the animal encounter in its natural environment? What subset of those are produced by stimuli it needs to detect (e.g., food, predators, pathogens, and social partners)? Do these stimuli often produce unique or highly characteristic single compounds, or must the animal tune in to higher-order features of blends (e.g., compound combinations, correlations, or relative abundances) for recognition?

Answering these questions will deliver vital insight into how animals may most efficiently encode odours. For instance, if a compound is produced by nothing in the environment except the target stimulus, we might expect the evolution of a narrowly tuned olfactory receptor embedded in a dedicated olfactory pathway. Indeed, this is the case for many pheromones, which are often rare, distinctive compounds detected via labelled lines (Nakagawa et al. 2005; Ruta et al. 2010; Demir et al. 2020). In other cases, stimuli may produce a unique combination of ubiquitous compounds. In this case, no single compound is adequate for discrimination, and we might expect evolution to fashion a combinatorial pathway that ignores single compounds and is activated only by the presence of multiple coinciding cues (Bruce and Pickett 2011). The most efficient strategy depends critically on the olfactory discrimination tasks an animal must perform, which in turn depend on the structure of the odour space.

One group of organisms that may face a particularly difficult olfactory discrimination task is host-seeking mosquitoes and other blood-feeding arthropods. Although turbulent plumes of carbon dioxide reliably signal the presence of a live (breathing) animal (Gillies 1980), blood feeders also depend strongly on organic compounds in vertebrate body odours to zero in on a host (DeGennaro et al. 2013; Cardé 2015). Previous studies of a handful of species suggest that vertebrate body odours overlap extensively in composition, sharing many of the same compounds (Syed and Leal 2009; Jaleta et al. 2016; Verhulst et al. 2018; Zhao et al. 2022). Discriminating among these similar odours may be difficult. However, some blood feeders rise to the challenge, exhibiting a strong preference toward feeding on particular taxonomic groups (e.g., Gouck 1972; Dekker et al. 2002; Krasnov et al. 2002; Rayaisse et al. 2010). For example, a few species have evolved to specialize in biting humans and thus become dangerous disease vectors (Takken and Verhulst 2013; McBride 2016). Understanding the composition of vertebrate body odours, including how they vary across species and more generally how they differ from other natural odours, would aid in our understanding of how blood feeders encode host odour.

Here we undertake the first large-scale study of vertebrate odour space, including a quantitative description of the odour of 64 vertebrate species, with a special focus on mammals. We describe the general statistics of vertebrate odours and show how they differ markedly from the statistics of floral odours, with implications for olfactory coding in blood feeders and floral visitors. We also use our data to make specific predictions about how mosquitoes have evolved to encode host odour. Our work shows how descriptions of odour space, coupled with an understanding of the ecology and evolutionary history of organisms, enable new insights into olfactory coding.

## Results

### Odour sampling

We collected odour from 120 individual animals spanning 64 vertebrate species, 29 families, and 13 orders via dynamic headspace sampling (Fig. 1a). In this procedure, air is actively passed over the odour-emitting object and through an adsorbent filter that traps volatile compounds (Fig. 1b–d). Some of the odour extracts came from live animals sourced from the lab or at local farms and zoos, while others came from hair that was freshly shorn and promptly frozen—e.g., during veterinary procedures at participating zoos (Fig. 1e). The sampled species therefore included common domestic animals such as guinea pigs, chickens, and sheep as well as charismatic megafauna such as tigers, giraffes, and polar bears. We also worked with great-ape sanctuaries to specifically collect hair samples from our closest relatives, including chimpanzees, bonobos, and orangutans. Because a large majority of our odour extracts were obtained from hair, we achieved especially broad sampling among mammals. We collected most samples opportunistically, sampling 1–4 biological replicates per species, except for three species (humans, chickens, and sheep) where we made a special effort to sample multiple individuals (Fig. 1f).

**Figure 1.**
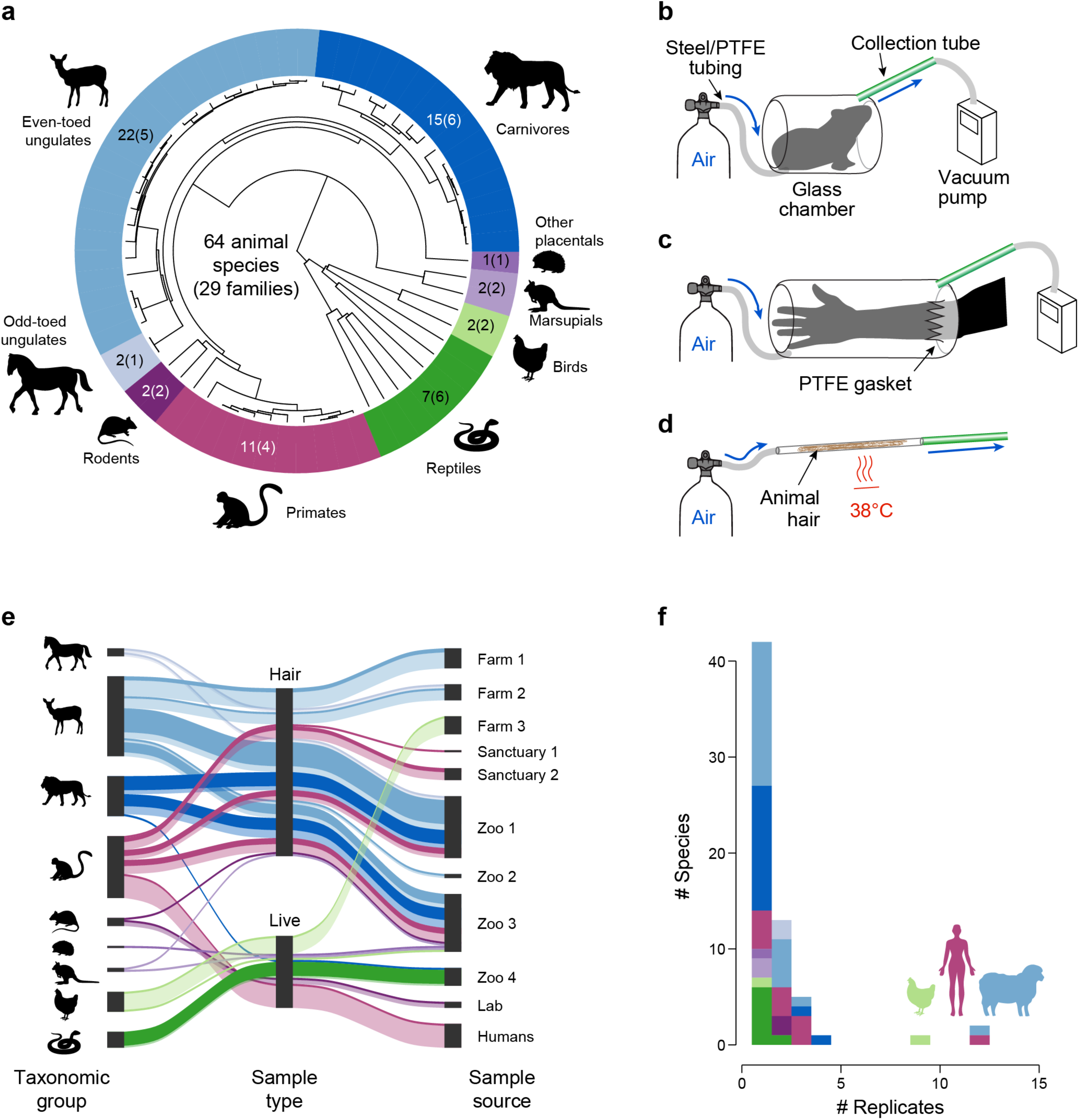
Odour-extraction methods and breadth of sampling. **(a)** Number and distribution of sampled animal species (and families) across phylogeny (time-calibrated phylogeny shown; see Methods for source data). **(b**–**d)** Three different types of glass chamber were used for odour sampling: **(b)** enclosed chambers of varying sizes for small and/or hairless animals, **(c)** a chamber with a Teflon gasket to accommodate a human arm, and **(d)** a narrow, heated tube containing tightly packed animal hair. Hair samples were heated to approximate body temperature (38°C) during odour extraction. **(e)** Sample types and sources. Coloured lines represent odour samples from different taxonomic groups (left) that were sampled from hair or live animals (centre) acquired from different sources (right). Line thickness reflects the number of unique species (dark shades) or within-species replicates (light shades). **(f)** Number of replicate individuals sampled for each species. Colours in (e,f) represent the same taxonomic groups listed in (a).

Hair odour is likely a reasonable proxy for mammalian body odour, but may lose some volatile compounds during storage. Live-animal odour, on the other hand, can be contaminated with compounds from faeces or urine occasionally excreted during sampling. We therefore looked for systematic differences in matched hair and live-animal extractions conducted on a small number of individuals from 3 mammal species (Fig. S1). Our results showed general correspondence between the two types of samples, which were then treated as equivalent in downstream analyses.

Odour trapped on the adsorbent-polymer collection tubes was analysed by TD-GC-MS (thermal desorption–gas chromatography–mass spectrometry). We used the metabolomics software XCMS (Smith et al. 2006) and a custom downstream pipeline to identify compounds and estimate their abundance (Fig. S2) (see Methods for discussion of abundance estimates, which correspond directly to total ion count and are not individually calibrated for each compound). Importantly, and unlike many prior studies, our data analysis was untargeted so as to obtain as unbiased and comprehensive a view of host-odour space as possible. However, one important class of compounds we undersample is the carboxylic acids, which are generally too polar or non-volatile to be analysed by gas chromatography without a special derivatization step (e.g., de Obaldia et al. 2022). We are also missing highly volatile compounds, which are difficult to capture (and therefore quantify) reliably.

### Basic statistics of vertebrate odour space

We identified a total of 116 compounds in our vertebrate odour extracts (Fig. 2a, S3, S4). These were generally non-polar or moderately polar compounds of medium volatility, most having 4–10 carbons and a molecular weight between 80 and 150 AMU (Fig. S3a). Aldehydes, ketones, alcohols, aromatics, terpenes, and hydrocarbons dominated the dataset (but see above re carboxylic acids). We identified between 13 and 61 compounds in each sample (median = 35; Fig. 2b). This finding is in line with previous estimates of the number of compounds found in human odour from other headspace- sampling studies (Dormont et al. 2013; Rankin- Turner and McMeniman 2022).

**Figure 2.**
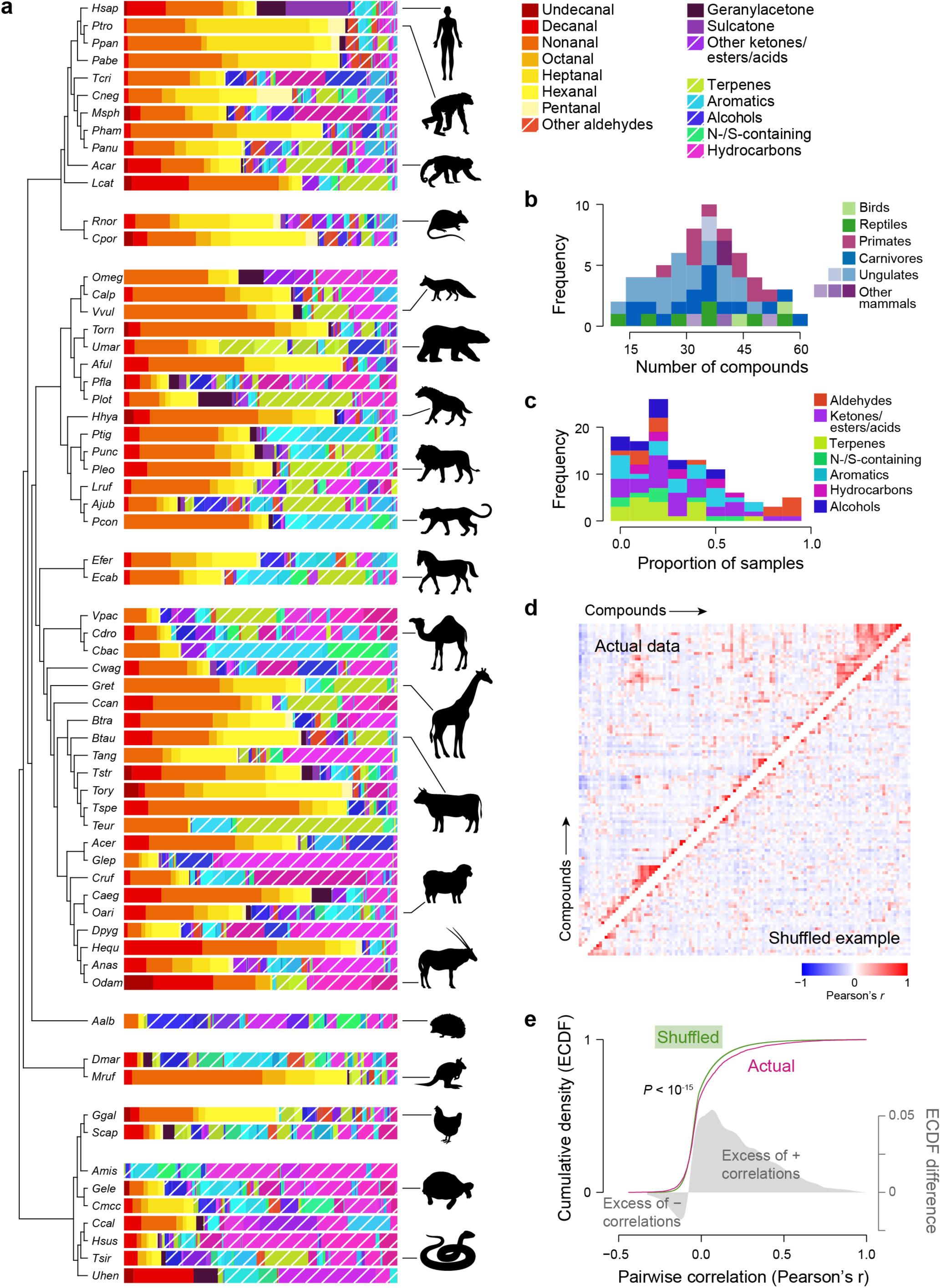
Odour of vertebrate animals. **(a)** Odour composition of one representative from each animal species sampled. The cladogram shows phylogenetic relationships (see Methods for source data). The length of each coloured bar shows the relative proportion of individual compounds in a sample. Also see Fig. S4 for an interactive plot showing all species replicates (including common names) and individual compound names. **(b)** Number of compounds found in each sample. **(c)** Frequency of compounds across samples (presence/absence). **(d)** Pairwise compound correlations across species for the real dataset (upper triangle) and a shuffled dataset (lower triangle; see Methods for details on shuffling). See Fig. S6 for compound labels. **(e)** Empirical cumulative distribution functions (ECDFs) of all pairwise compound correlations for the actual data (pink) and the median across 1000 shuffled datasets (green) (significantly different by Kolmogorov–Smirnov test, *P* < 10^-15^). For shuffled data, the narrow 95% credible interval (light-green shading) is barely visible behind the green line. Shaded density plots illustrate the difference between the two ECDFs on a 10×-magnified scale.

**Figure 3.**
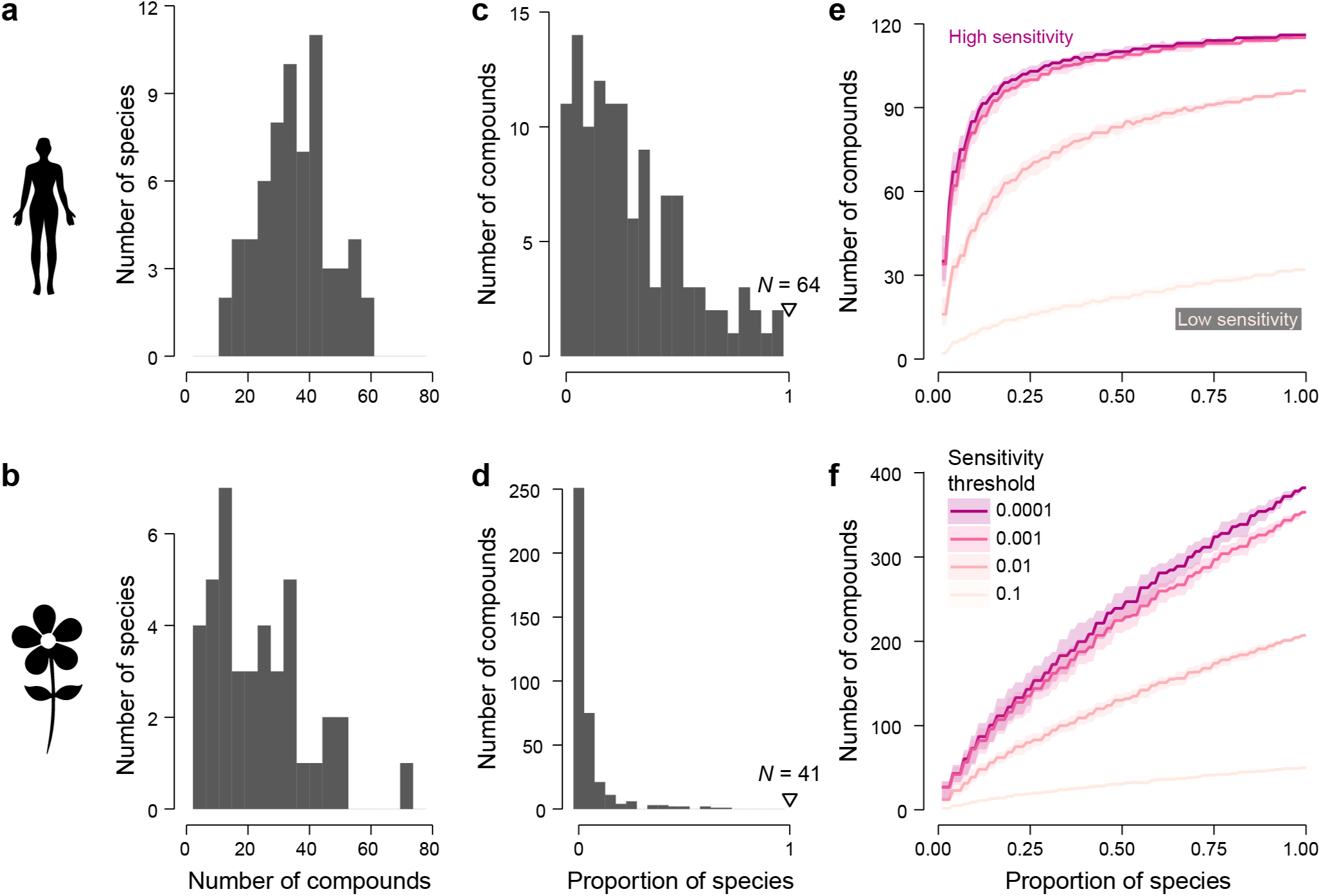
Comparison of vertebrate and floral odour-space. Top row: vertebrate odour space (this study), bottom row: floral odour space (Kantsa et al. 2018). **(a,b)** Number of compounds found in the odour of each species. **(c,d)** Proportion of species in which each compound is found. *N* = 64 and 41 for vertebrate and flower samples, respectively. **(e,f)** The total number of compounds one expects to encounter across a given proportion of all species (*x* axis) for a given sensitivity threshold (shades of pink). Lines and shading show medians and interquartile ranges for 100 randomly sampled sets containing the given number of odour samples. Lower sensitivity thresholds allow less abundant compounds in each species to contribute to the total.

**Figure 4.**
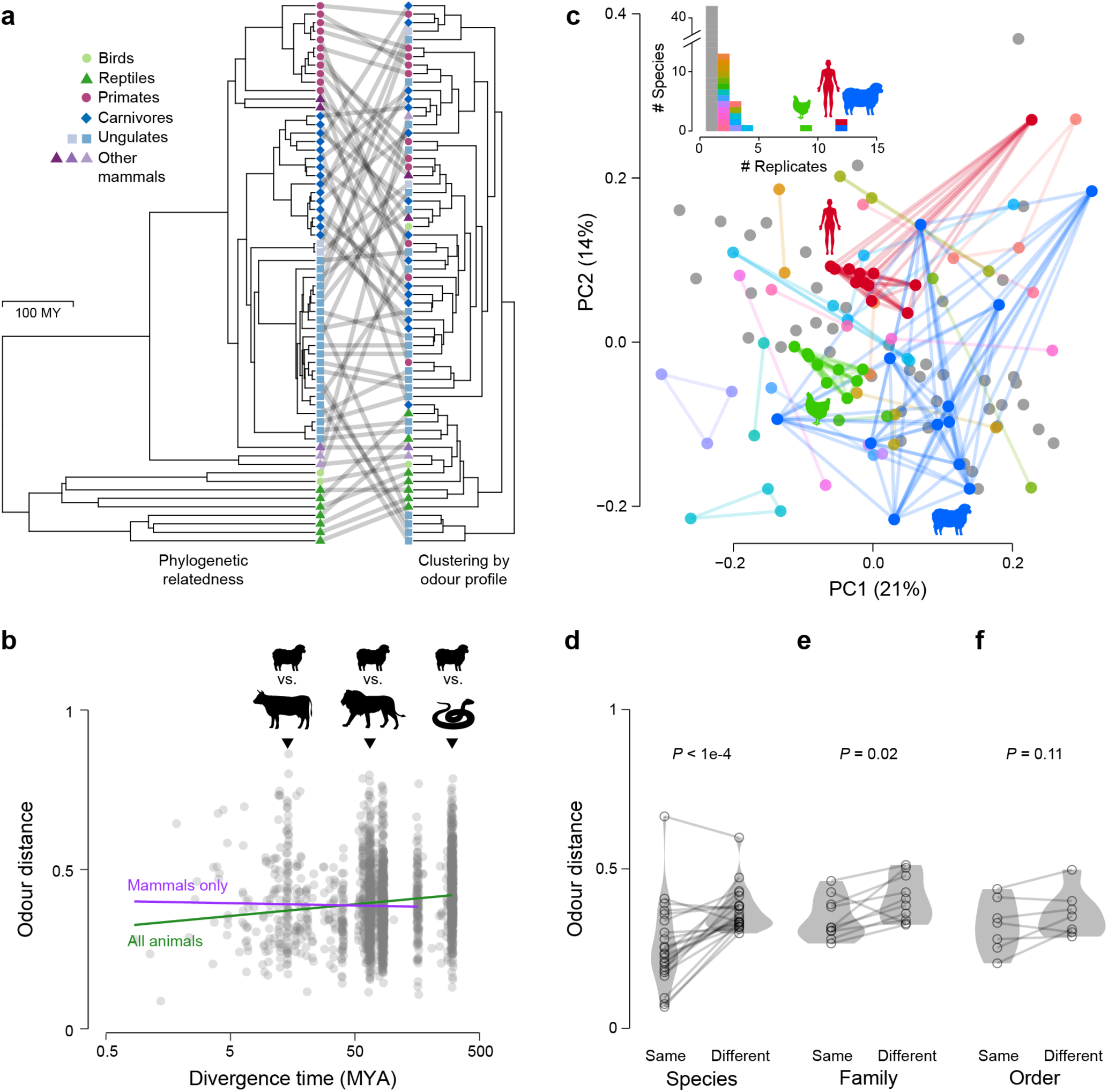
Phylogenetic signal in odour-blend composition. **(a)** Time-calibrated phylogenetic tree for sampled animal species (left) shown with linkages to a hierarchical clustering of odour profiles based on Euclidean distance (right). One representative sample from each species was chosen. Nodes have been rotated to optimize vertical matching of tips between trees. **(b)** Pairwise Euclidean distance (in *n*-dimensional odour space) between odour profiles (one from each species) as a function of divergence time. Mantel test *P* = 0.04 for all animals; *P* = 0.52 for mammals only. **(c)** Unscaled PCA of odour blends showing all biological replicates from all animal species. Coloured dots connected by lines represent replicates of the same species. Grey dots represent species that were only sampled once. Inset: Number of replicates sampled from each species. Same as Fig. 1f, but colours now match the PCA. **(d**–**f)** Median odour distance within and between species (*N* = 22), families (*N* = 10), and orders (*N* = 7) (see Methods for details). *P*-values shown for paired *t*-tests.

Compounds varied widely in their prevalence across species, from those found in only a few samples to those that were nearly ubiquitous (Fig. 2c, S5). However, over half were found in at least 15 samples and very few compounds were species-specific (only 7/116). The high prevalence of straight-chain, saturated aldehydes was especially remarkable. These compounds were found in virtually every sample. They also tended to be found at a high abundance within each odour blend, constituting on average 46% (±3% SE) of each odour extract (Fig. 2a, 2c, S5). Among reptiles, long (C12– C16) unbranched hydrocarbons dominated instead of aldehydes (Fig. 2a, S4). However, we caution that we have low confidence in the accuracy of the reptile samples; they tended to be less concentrated with a lower signal-to-noise ratio. There are potentially both biological and technical explanations: First, it is possible that reptiles truly emit less odour than warm-blooded vertebrates do, perhaps because of lower emission of endogenous volatiles or less microbial activity on colder and drier skin. Second, all reptiles were sampled as live hosts (Fig. 1e). Many of these extractions yielded lower odour abundances than typically achieved with hair samples, where air is forced through a dense matrix of hair instead of simply passing over an animal.

We next asked whether there were correlations among compounds across species. A correlation matrix revealed a small but significant excess of correlations compared to a matrix generated from a shuffled dataset (Fig. 2d,e; S6). Correlations were generally weak (|*r*| < 0.5); in other words, the abundance of one compound does not reliably predict the abundance of another. Taken together, these results indicate that vertebrate body odours are complex blends that overlap broadly in composition and show little correlational structure.

### Comparison to floral odour space

In order to better evaluate the properties of vertebrate odour space, we compared it to one of the only other quantitative, large-scale odour datasets we could find: a study of floral odour space that estimated the abundance of 382 compounds in 41 Mediterranean scrubland species (Kantsa et al. 2018). The floral odours tended to be slightly less complex than vertebrate odours, with fewer constituent compounds (median = 21 vs. 35 for vertebrates; Fig. 3a,b). They were also far more likely to contain unique compounds (Fig. 3c,d): strikingly, nearly two thirds of floral compounds were found in only a single species, compared to just 6% of vertebrate-odour compounds.

To explore these patterns more closely, we plotted rarefaction curves that show how the total number of compounds encountered across species increases with the number of included species (Fig. 3e,f). The number of compounds plateaued quickly for vertebrates; even a handful of vertebrate species together contained most of the compounds found in the entire dataset. For floral odours, this number reached its maximum more slowly; new compounds were still being encountered even when we had already examined a large number of flower species. This observation also held true when we restricted the analysis to abundant compounds by decreasing the within-sample sensitivity (lighter-pink curves): even floral compounds that are abundant in the odour of individual species may be rare across species. Overall, this analysis demonstrates that vertebrate odours share vastly more compounds than floral odours do.

### Phylogenetic signal

The broad range of taxa included in our survey allowed us to ask whether odour profiles carry any phylogenetic signal. We found little to no evidence that more closely related species had more similar odour profiles (Fig. 4a,b). A Mantel test comparing phylogenetic and odour distance matrices was marginally significant (*P* = 0.04), but the effect appears to be driven purely by the dissimilarity between the odours of reptiles and mammals; it disappeared when considering mammals only (*P* = 0.48, Fig. 4b). As discussed above, reptile extracts may have looked different for technical rather than biological reasons. Additional tests for phylogenetic signal among reptiles and birds only, or among samples within three mammalian subclades (ungulates, carnivores, and primates), likewise failed to identify any phylogenetic signal (Fig. S7).

Body-odour composition may be species-specific, even if phylogenetic signal is lacking at higher taxonomic levels. We tested this idea by leveraging the replicate samples we had collected for a subset of the species in our dataset. A principal-components analysis indicated strong within-species clustering in at least some cases (Fig. 4c). To confirm this impression, we conducted paired *t*-tests based on median odour distance within and between species (see Methods for details). This approach confirmed strong within-species similarity (Fig. 4d). Similar tests at the level of family and order reinforced our earlier findings that phylogenetic signal is weak or non-existent at higher taxonomic levels (Fig. 4e,f). We caution that many of our species replicates came from the same farm or zoo and are therefore confounded with sample origin (Fig. S4). Thus, although we are confident that at least some species in our dataset have consistent and characteristic odour profiles, the extent to which this finding applies broadly across animals deserves follow-up in a better-controlled experiment.

### Specific compounds as host-seeking cues

A quantitative description of odour space allows us to predict how animals might best discriminate specific stimuli of ecological importance. For example, from the perspective of a generalist blood feeder, we can ask which compounds or compound classes best separate the odour of live vertebrate hosts from other, non-host stimuli (Fig. 5a). To make this comparison, we collected and analysed odour from 23 diverse non-host stimuli, including faeces, urine, carcasses, soil, vegetation, and flowers (Fig. S8c).

**Figure 5.**
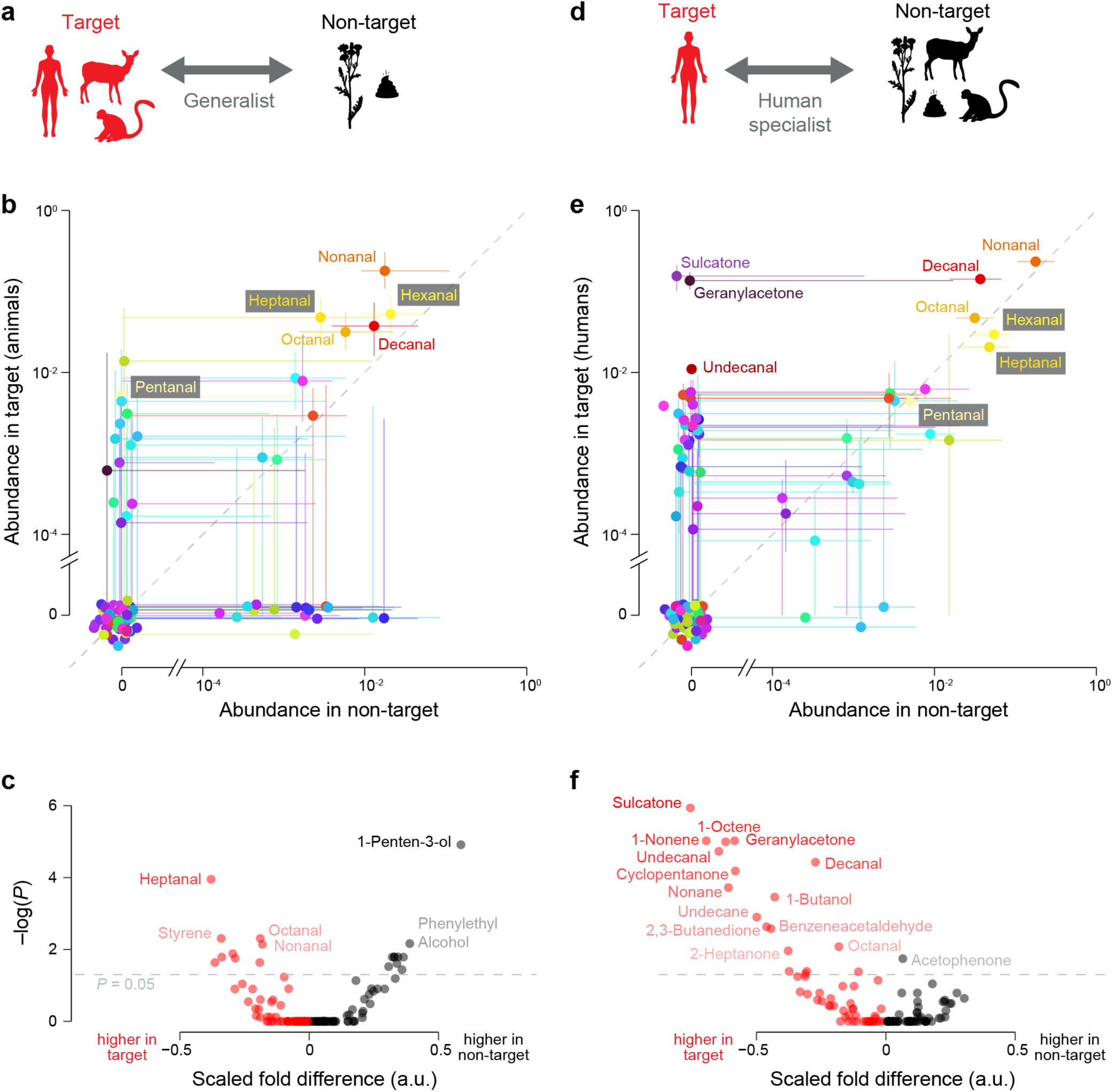
Compounds informative for host discrimination. **(a)** Odour-discrimination tasks for a generalist blood feeder: distinguishing between animals and non-animals. **(b)** Compound abundance (relative proportion within a sample) in target and non-target odours for a generalist. *N* = 64 animal samples (one per species) vs. 23 other samples. Dots and lines show medians and interquartile ranges. Points along the axes are jittered. Lines corresponding to points at the origin have been omitted for clarity. Compounds are coloured according to the scheme in Fig. 2a. **(c)** Volcano plot showing two metrics for each compound for the generalist task: (1) scaled fold difference between target and non-target (see Methods) and (2) significance level of that difference. *P*-values come from Kolmogorov–Smirnov tests followed by Benjamini–Hochberg multiple-test correction. Red compounds are those with higher abundance in the target; black for non-target. **(d–f)** Same as (a–c), but for a human specialist distinguishing between humans and all other odour blends. *N* = 12 humans vs. 63 other animals (one representative per species) and 23 other samples.

Non-host odours contained a set of compounds that overlapped almost completely with compounds found in vertebrate odour (Fig. S8d), but there were important quantitative differences. As noted above, straight-chain, saturated aldehydes with 5–10 carbons all stood out as abundant in vertebrate odours. Most non-host stimuli also emitted aldehydes, but at substantially lower levels on average (Fig. 5b, S8a,c: red, orange, and yellow compounds). The comparison in Fig. 5b emphasizes high-abundance compounds, but minor components may also hold strong predictive value. We therefore carried out an additional analysis based on scaled fold differences that is insensitive to absolute abundance (Fig. 5c). Three straight-chain aldehydes were again among the top hits. Heptanal was by far the most predictive of host odour, followed by octanal, nonanal, and several minor components. Straight-chain aldehydes therefore appear to be general features of vertebrate host odour that may represent valuable cues for blood feeders. 1-Penten-3-ol, a common alcohol released by damaged leaves (Fall et al. 2001), was strongly biased in the opposite direction, despite constituting on average just 0.5% of each non-host odour blend (Fig. S4).

A handful of mosquito species have evolved to specialize in biting humans and thereby become dangerously efficient vectors of human disease (Takken and Verhulst 2013; McBride 2016). Assuming the perspective of these human specialists, we also searched for compounds that differed consistently in abundance between the 12 human subjects in our survey and the 63 other vertebrate species, as well as the odour of all non-host stimuli (Fig. 5d, S8a–c). Two differences from the generalist analysis were immediately apparent: First, while the aldehydes with 5–9 carbons became uninformative in this comparison, the 10-carbon aldehyde decanal was even more human-biased (Fig. 5e) than it had been animal-biased in the generalist analysis (Fig. 5d). Second, several compounds emerged out of relative obscurity in the generalist task to become extremely strong predictors of human hosts. The most prominent of these were sulcatone, geranylacetone, and the 11-carbon aldehyde undecanal (Fig. 5e), in agreement with the trends shown in our earlier survey of just 6 species (Zhao et al. 2022).

Many additional compounds emerged as human-biased in the analysis of scaled fold differences (Fig. 5f), as expected from the high consistency in composition across human odour blends (Fig. S4, S8b). Some of these hits may result from shared contaminants or noise—e.g., from exogenous sources common in human environments or from sampling all humans in the same manner (Fig. 1c). However, we are confident that sulcatone, geranylacetone, decanal, and undecanal—which are endogenously produced from human sebum (Zung & McBride in prep)—represent real biological hallmarks of human odour. Importantly, this is true even when we compare humans to other great apes; the odours of chimpanzees, bonobos, and orangutans are strikingly similar to each other, with high amounts of heptanal, but little to none of the four most abundant, human-biased compounds (Fig. S4, S8). Thus, although human odour contains no unique compounds in our analysis (Fig. S8d), several quantitative differences clearly set it apart.

## Discussion

Much research has focused on how a combinatorial code allows olfactory systems to discriminate among a large number of compounds (Koulakov et al. 2007; Nara et al. 2011; Bushdid et al. 2014). However, olfactory systems have evolved not necessarily to discriminate among as many compounds as possible, but instead to discern odours of distinct ecological relevance, such as those of food, social partners, predators, and pathogens. Studying the composition of these ecologically relevant odours can therefore deliver fresh insight into olfactory coding and its evolution.

Here we have presented the first broad-scale, quantitative study of the odour of vertebrate animals. We confirm previous suggestions that vertebrate odours are blends of relatively common compounds that are frequently shared across species (Fig. 3c,e). In this way, vertebrate odours stand in stark contrast to floral odours, which often contain species-specific components (Fig. 3d,f). We suspect the difference is at least partly attributable to the distinct selective pressures acting on floral vs. vertebrate odour. Flowers function primarily as advertisements and may be under selection to signal on private channels to attract loyal pollinators—those with innate preference or those that have learned to recognize the most rewarding species at any given point in the season (Raguso 2008; Hopkins and Rausher 2012; Schiestl and Johnson 2013). Vertebrate body odours, in contrast, are more likely to reflect the byproducts of common metabolic processes intrinsic to skin physiology or microbial activity. More generally, odour spaces characteristic of mutualistic interactions where the emitter “wants” to be found may differ systematically from parasitic, predatory, or commensal ones. In addition, plants may simply have greater evolutionary lability in the type of volatile compounds they produce: a distinguishing feature of plant biochemistry is the terpenoid secondary metabolites, a class of compounds that are unconstrained by basic metabolic processes and endlessly variable (Tholl 2006).

Regardless of exactly why the statistics of these two odour spaces differ, the fact that they do has implications for olfactory coding in blood feeders and floral visitors (Atick 1992; Zwicker et al. 2016). For example, a blood feeder searching for a specific vertebrate host may need to integrate information about the relative abundances of many compounds, while floral visitors may employ a comparatively simplistic code. Indeed, specialist mosquitoes are notoriously unresponsive to single compounds aside from CO_2_ and require odour blends to detect hosts (Dormont et al. 2021). Honeybees, however, are sometimes able to generalize from an attractive mixture of compounds to its individual components (Laloi et al. 2000; Reinhard et al. 2010; Szyszka and Stierle 2014), implying that one or a few individual compounds may account for most of the behavioural response. These apparent differences may partly reflect the different odour spaces mosquitoes and bees inhabit.

Compounds in vertebrate odour are not only shared broadly across species, but also common in other natural odours (Fig. S8d). Their ubiquitous nature may at first glance pose a challenge for generalist blood feeders. However, our study suggests that straight-chain, saturated aldehydes are still useful indicators of vertebrate odour by virtue of their unusually high abundance (Fig. 5b,c). Indeed, these aldehydes have been shown to enhance host seeking in diverse mosquito taxa (Syed and Leal 2009; Tchouassi et al. 2013; Leal et al. 2017; Zhao et al. 2022), and even malaria parasites and orchids appear to deploy them as mosquito attractors (Emami et al. 2017; Robinson et al. 2018; De Moraes et al. 2018; Lahondère et al. 2020). Other compounds used by blood feeders—such as ammonia and some volatile carboxylic acids (Geier et al. 1999; Dekker et al. 2002; Smallegange et al. 2005; Dormont et al. 2021)—may also reliably indicate the presence of a vertebrate host, but do not emerge from our study because our approach was not designed to capture highly volatile or polar compounds.

Another notable feature of vertebrate odour space is the lack of phylogenetic signal—at least among mammals, which make up 55 of our 64 sampled species. This may help explain why few, if any, mosquitoes are known to exhibit preferences for taxonomic groups at the level of family or order (Clements 1999). Moreover, when mosquitoes specialize on even broader groups, such as frogs or birds, their preferences may be mediated by other sensory cues (e.g., frog calls (Borkent and Belton 2006)) or be a function of habitat use (e.g., frequenting trees and specializing on roosting birds (Janousek et al. 2014)). Despite the lack of phylogenetic signal at broad taxonomic scales, our work suggests high consistency in odour-blend composition *within* a species. This implies that *species-specific* host preference may be more easily mediated by olfactory cues (Gouck 1972; Osterkamp et al. 1999).

We focus in particular on odour features that may be valuable cues for human specialists. Four of the eight most abundant components of human odour are distinctive—sulcatone, geranylacetone, decanal, and undecanal (Fig. 5e,f). These findings align with our earlier study of just 6 species (Zhao et al. 2022), but also reveal additional insight. For example, of the two long-chain aldehydes, undecanal (11 carbons) turns out to be a more reliable indicator of human odour than decanal (10 carbons), despite its overall lower abundance (Fig. 5e,f; S8e). Undecanal was less common than decanal across non-human animals (Fig. S4; S8a,b). It also appears to be less common in nature more broadly (Fig. S4, S8c; Lemfack et al. 2018; VCF Online). For instance, undecanal has been reported in the floral odour of just 7% of plant families studied, as compared to decanal at 27% (Knudsen et al. 2006). Both compounds are enriched in human odour because they are oxidation products of unique lipids found on human skin, but undecanal is less likely to be produced by other biochemical pathways more common in nature (Zung & McBride in prep). The relative rarity of undecanal helps explain our previous findings on human-odour coding in *Ae. aegypti*. Females of this mosquito species have a group of neurons that are exquisitely tuned to long-chain aldehydes and that help drive host seeking (Zhao et al. 2022). But the neurons are substantially more sensitive to undecanal than to decanal, a feature we can now trace to the statistics of odour space.

The two ketones are just as or even more reliable indicators of “human-ness” than undecanal (Fig. 5e,f, S8e). Why, then, should human-specialist mosquitoes rely more heavily on the aldehydes? The answer may lie in the evolutionary history of this mosquito. The human-specialist subspecies of *Ae. aegypti* arose in just the last 5000 years from an ancestral generalist subspecies that still thrives across much of sub-Saharan Africa (Rose et al. 2020, 2023). Aldehydes are valuable cues for generalist blood feeders (Fig. 5b,c), and aldehyde-sensing neurons may be an important feature of host-seeking circuits in the ancestral generalists. If so, retuning these neurons to the *long-chain* aldehydes enriched in human odour may have provided a relatively simple and fast evolutionary path to human preference. Sulcatone and geranylacetone, in contrast, are not chemically similar to any compounds predicted to be important cues for generalists, posing more of a challenge for rapid sensory evolution. While there is some evidence that these mosquitoes are beginning to evolve greater sensitivity to sulcatone (McBride et al. 2014), full incorporation of ketone detection into the host-seeking circuit would likely take longer than a subtle retuning of aldehyde-sensing neurons.

Navigating chemical space is one of the oldest sensory challenges that organisms have faced (Jacobs 2012). Yet despite the widespread importance of olfaction across the animal kingdom, the neural transformation from odour to perception remains mysterious: which signals does a given olfactory system choose to amplify, and how is information about multiple compounds integrated to drive behaviour? Understanding the composition of natural odours is a critical first step in answering these questions. Our work adds to a very small number of studies that quantitatively characterize an odour space to discover the most statistically informative features that distinguish target from non-target stimuli. We can only understand why animal olfactory systems have evolved to encode odours in the way they do when we step into the shoes of our study organisms to appreciate the olfactory challenges they face.

## Acknowledgements

We thank the following for help with sample collection: Alex Ernst and the staff at Cape May County Zoo, Clyde Peeling’s Reptiland, San Diego Zoo and Safari Park, Pete Watson and the staff at Howell Living History Farm, the Center for Great Apes, Ape Initiative, Six Flags Great Adventure NJ, Mary McLaughlin, Liam Nygaard, and PJ the cat. We also thank James Thomson and members of the McBride lab for helpful discussion and Cassie Stoddard for comments on the manuscript. J.L.Z. was supported in part by a fellowship from the Natural Sciences and Engineering Research Council of Canada. This work was funded by grants to C.S.M. from the National Institutes of Health (NIAID: DP2AI144246) and the New York Stem Cell Foundation. C.S.M. is a New York Stem Cell Foundation – Robertson Investigator.

## Author contributions

J.L.Z. and C.S.M. conceived the project and designed and interpreted the experiments. S.M.K. helped with processing of hair samples. J.L.Z. carried out all other aspects of data collection and analysis. J.L.Z. and C.S.M. wrote the paper.

## Methods

### Ethics and regulatory information

The use of live non-human animals and non-human animal hair in odour extractions was approved and monitored by the Princeton University Institutional Animal Care and Use Committee (protocol #1999 for laboratory animals, #2136F for animals off site). The participation of human subjects in odour extractions was approved by the Princeton University Institutional Review Board (protocol #8170). All human subjects gave their informed consent to participate in work carried out at Princeton University.

### Sample collection

Hair samples were sourced from zoos, farms, and animal sanctuaries. Samples were generally collected opportunistically (e.g., when parts of an animal’s body needed to be shaved for a veterinary examination), but we asked staff to aim for a broad sampling across phylogeny. Staff were instructed to seal the hair in glass jars with Teflon-lined lids (Thermo Scientific V2200125) immediately after collection, then store samples at −20°C until shipping on ice overnight to our laboratory. Hair samples were generally stored at −20°C at the source facility and/or in our laboratory for 1–6 months before odour extraction, but a small number of samples were stored for up to 2 years (Supplementary Table 1).

To check whether the odour of hair was a reasonable proxy for the odour of live hosts, we collected both types of extracts from a small number of individuals and species: 4 humans, 2 rats, and 1 guinea pig (Fig. S1). Human hair samples were taken from the head, and rat and guinea-pig samples from the whole back and sides of the animal. For live-host sampling, we extracted odour from the entire animal (for rodents) or human arms, as shown in Fig. 1b,c. Except in two cases (one where the guinea pig urinated and defecated in the chamber and one where a human participant’s odour contained a large quantity of limonene, a probable contaminant from soap), we found that live-animal odour was generally similar to odour from hair (Fig. S1).

We visited two local zoos (Cape May County Zoo, Reptiland) and one privately owned farm for sampling odour from live hosts. Species were selected from among all available at a given location for physical size match to our extraction chambers, comfort level in a small enclosed space, and phylogenetic diversity. We also extracted odour from two species available in our lab: rats and guinea pigs.

Twelve human subjects participated in the experiment (6 male, 6 female, with ages ranging between 20 and 50 years). Participants were asked to avoid showering for at least 24h in advance of the study and to avoid using the sampled arm (Fig. 1c) to contact skin products or other strong-smelling substances the day of the experiment (e.g., soap, lotions, oily food, citrus fruits). Four human volunteers also provided hair samples (Fig. S1). These participants were asked to avoid showering for at least 24h in advance. Hair was collected from as close to the scalp as possible and stored immediately in glass jars with Teflon-lined lids at −20°C until odour extraction.

Non-host samples were collected in the vicinity of Princeton, NJ, USA—or, in the case of some faecal/urine samples, collected opportunistically during visits to local zoos for live-host odour extractions. We extracted odour from all non-host samples immediately after collection, with the exception of the two carcasses; these were discovered when freshly killed (by a cat or window strike) and allowed to decay outdoors (8–23°C) for 3–4 days before odour extraction. Storage containers were almost never reused. In the few cases when they were, containers were rinsed with a polar and non-polar solvent (methanol and hexane or ethanol and diethyl ether) and air-dried before reuse.

### Odour extractions

We collected odour by passing zero-grade air over hair or live animals (see below for details on extraction chambers) and through tubes filled with Tenax TA beads (Supelco 30133-U, Gerstel 020586- 010-00, or Markes International C1-CAXX-5003). All extractions were run for 5–80 minutes depending on the expected odour concentration of the sample. We usually collected odour on two tubes consecutively with different extraction durations to allow for a second GC-MS run at a different concentration and split ratio if required. All fittings and lines in the flow path upstream of the Tenax tube were made of glass, Teflon, Viton, or metal.

For non-human live-host extractions, we placed each animal in an appropriately sized glass chamber: a domed bell jar with a hard acrylic base (45 cm high × 30 cm I.D.; Chamber 1) or one of three cylindrical chambers with glass caps (21 cm long × 13 cm I.D., 30 cm long × 7.5 cm I.D., 37 cm long × 13 cm I.D.; Chambers 2–4). These chambers have threaded ports at either end for inflow/outflow of air. We supplied the chamber with zero-grade air (Airgas AI Z300) at 0.5–2L/min through one port. Odour-laden air was pulled onto a Tenax tube at the second port using a vacuum pump at a rate of 200mL/min.

Input flow rate was set higher than output flow rate to create positive pressure in the chamber that minimized the amount of room air entering via the leaky seal around the chamber entrance.

Human-arm-odour extractions were run using the 37 cm long × 13 cm I.D. cylindrical glass chamber. Instead of sealing the open end with its glass cap, we fashioned gaskets out of Teflon sheets that fit snugly around participants’ arms. Participants inserted their arms up to just above or below the elbow. Flow rates were the same as for other live-host extractions.

For non-host extractions, we used one of the three cylindrical chambers, a 250 mL glass gas-washing bottle (Chemglass CG-1112-02; Chamber 5), or a small glass jar (7 cm tall × 4.5 cm I.D.; Chamber 6) sealed with a Teflon-lined lid, which had input and output holes drilled into it. Flow rates were the same as for live-host extractions. Cut stems of plants were immersed in water whenever possible to reduce signal from wound-induced green-leaf volatiles.

Hair extractions were run in a thermal-desorption sample-collection system (part number 782050) manufactured by Scientific Instrument Services, Inc., Ringoes, NJ, USA (merged with Adaptas Solutions, Palmer, MA, USA). Hair (∼0.5–4g) was packed tightly into a 0.36” I.D. glass tube (Chamber 7) and heated to 38°C. We pushed zero-grade air through the tube at a flow rate of 50 mL/min. No vacuum pump was required because fittings were tight enough to hold pressure. In a small number of cases, hair from multiple individuals of the same species was combined for extraction due to the small amount of hair collected (see Supplementary Table 1).

We washed glass chambers between extractions by rinsing with both polar and non-polar solvents (methanol and hexane or ethanol and diethyl ether) and then setting them out to air-dry. If residue was visible, we added an initial scrubbing step using water and fragrance-free soap (Babyganics Foaming Dish & Bottle Soap) before the solvent rinse.

### TD-GC-MS analysis

Odour extracts on Tenax tubes were analysed as described in (Zhao et al. 2022). Briefly, we used an Agilent GC-MS system (Agilent Technologies, Santa Clara, CA, USA; GC 7890B, MS 5977B, high-efficiency source) outfitted with a DB-624 fused-silica capillary column (30 m long x 0.25 mm I.D., d.f. = 1.40 μm, Agilent 122-1334UI). Our instrument also has a backflush module installed, including a ∼0.5m no-phase column downstream of the main column. Tenax tubes were inserted into a Gerstel TD3.5+ thermal-desorption unit (Gerstel Inc., Linthicum, MD, USA) mounted on a PTV inlet (Gerstel CIS 4) with a glass-bead–packed liner. Tubes were heated in the TD unit from 50°C to 280°C at a rate of 400°C/min, then held at 280°C for 3 min. During the TD heating time, volatiles were swept splitless into the cold inlet (−120°C) under helium flow of 50 mL/min. After the tube was removed and the inlet repressurized, the inlet began heating at a rate of 720°C/min to a 3 min hold temperature of 275°C. The GC oven program began simultaneously with inlet heating, starting at an initial temperature of 40°C and ramping at a rate of 8°C/min to a 10 min hold temperature of 220°C. An appropriate inlet split ratio was chosen based on concentration of odour in the Tenax tube. If concentration was too low or too high, we ran a second tube from a replicate extraction, at a different split ratio if necessary. Carrier-gas flow rate was 1.2 mL/min. The MS was operated in EI mode, scanning from m/z 40 to 250 at a rate of 6.4 Hz.

### Analysis and curation of GC-MS data

#### Initial processing with XCMS

We used the program XCMS to process the raw chromatograms (Smith et al. 2006). This program performs peak detection, retention-time correction, and peak matching across samples to generate a matrix of components and their abundances across samples. Components are peaks at a specific m/z ratio and retention time; since our MS breaks compounds into many fragments with different m/z ratios, compound peaks are made of many components (Fig. S2a).

#### Blank subtraction

We conducted a series of blank extractions: tubes run on an empty glass chamber or in the thermal extractor used for hair samples, then analysed on the GC-MS with different split ratios. Each real sample was matched to a blank with the same type of extraction chamber and split ratio. Blank component peaks were subtracted from peaks in real samples to produce a clean matrix of component peaks attributable to the sample and not technical noise (Fig. S2a).

#### Creation of custom library

Next, we searched for matches between the NIST17 library and groups of components in each sample with similar retention time (within a sliding 4s window). These groups of components corresponded to putative compounds. We applied two filters to putative compounds: total raw ion abundance must exceed 5×10^6^, and no fewer than 4 components must account for 95% of the total abundance. Matches with a cosine similarity of >0.95 to the NIST library were individually reviewed by eye and added to our custom library. We did not review putative compound spectra that met these criteria but were deemed similar enough to ones that had already been manually reviewed (i.e., a cosine similarity of >0.9 and <0.5s retention-time difference).

We also manually checked the most prominent (generally top 5–10) peaks in each sample to ensure that they were included in the library. Those not caught by our initial automatic screen were identified using the NIST MS search software and manually added to the library.

#### Component distinctiveness

When two compounds in our library had similar retention times and overlapping components, it was sometimes difficult to distinguish them and estimate their separate abundances. To aid in this process (see below), we therefore calculated the “distinctiveness” of each component for each compound in our library. This metric describes how unique a given component is for the focal compound relative to other compounds with similar retention time (Fig. S2a). Specifically, distinctiveness of a given ion for a given focal compound was calculated as follows:

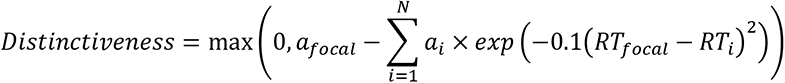

where

*a* is the abundance of a given ion in a compound’s mass spectrum, normalized to its highest peak; *N* is the number of compounds in the entire library, excluding the focal compound; and *RT* is the retention time of a compound in seconds.

#### Detection of compounds and abundance estimation

Custom library in hand, we next quantified the abundance of compounds present in our samples (Fig. S2b). For each compound in our custom library, we selected all XCMS components in the focal sample with a retention time within 2s of the focal compound. If the similarity between library and sample spectra met threshold (>0.7 cosine similarity), we counted a compound as “present” in that sample and estimated its abundance as follows. Each component gives an estimate of the total ion count for that compound (given by the ratio of component abundance to total ion count in the library spectrum). For each compound in each sample, we took the mean of all these estimates, weighted by the distinctiveness of each component for that compound. This weighted mean served as our estimate of compound abundance.

Although XCMS performs peak deconvolution, errors in this process may result in a failure to detect low-abundance compounds co-eluting with other compounds. To detect these compounds, we attempted to remove the signal corresponding to already identified compounds (Fig. S2b). Specifically, we used our abundance estimates of already identified compounds to back-calculate the true underlying component abundances, again using the ratio of component to compound abundance in library spectra. We subtracted these inferred component abundances from the original component peaks to generate a “residual sample”. We then followed the same procedure outlined in the previous paragraph to estimate the abundance of the compound represented by the residual sample.

Raw estimated abundances were converted to proportional abundance in each sample (i.e., the abundances of all compounds in a given sample sum to 1). These proportional abundances were used for all subsequent analyses.

#### Advantages and disadvantages of approach

We expect that our approach to analysing GC-MS data is more sensitive than existing automated, untargeted approaches. In existing methods, peak-detection and deconvolution errors often cause components to be missed or incorrectly grouped together, especially for low-abundance compounds. The raw total ion count in these cases is not a reliable measure of compound abundance, so normally the threshold for a match between library and sample spectra must be set higher to avoid including these compounds in the analysis. Instead of simply summing component peaks, we obtain reliable estimates of the “true” total ion count by averaging across estimates from all components, weighted such that more-distinctive components have more influence on the final estimate of total ion count (see Fig. S2b). This strategy allows us to set a relatively low match threshold, so that even compounds close to the detection threshold whose signal is partly obscured by noise can be quantified.

Because it was not feasible to run calibration curves for every compound in our library, our measure of compound abundance corresponds directly to the area under the total ion chromatogram. Our abundance measurements are therefore confounded with GC-MS sensitivity to each compound, and quantitative comparisons *across compounds* should be made with caution, especially in the case of chemically dissimilar compounds that are likely to have different GC-MS response factors. In contrast, our quantitative estimates of the same compound *across samples* are more reliable and thus form the basis of the most crucial analyses in this paper.

#### Excluded samples and compounds

Nine samples were excluded because of known issues at the sample-collection stage (improper storage/metadata collection, known contamination). An additional 49 samples were excluded because of low quality (low total odour content and/or suspected contamination characterized by large, late-eluting hydrocarbon peaks with very poor chromatographic separation that obscured all other signals).

We excluded compounds with retention time <500s since we could not be sure we had quantitatively captured these small, early-eluting compounds on our Tenax tubes. We also excluded two classes of compounds that were clearly non-biological in origin: siloxanes and halogenated hydrocarbons. We suspect that other contaminants remain in our dataset. For example, it is clear that human replicates #4 and #5 contain large anomalous peaks of limonene and 2-phenoxyethanol, respectively (Fig. S4). As these compounds are common additives to soaps and lotions, they almost certainly represent contaminants rather than real biological variation in human odour. Although we suspect that such contamination is common throughout the dataset, we cannot reliably distinguish it from biological variation, especially in animal species with only one sampled replicate. Therefore, we have taken the liberal approach of excluding only a handful of compounds that are clearly non-biological in origin (i.e., those that include atoms other than C, O, N, and S).

#### Species representative

Except where otherwise indicated, analyses were performed on a dataset that includes only one representative odour profile per species. The species representative was chosen at random—except that we avoided choosing extracts that were clearly lower in concentration or outliers in composition compared to extracts from conspecifics.

### Assignment of compounds to classes

We assigned compounds to each of the following classes: aldehyde, ketone, ester (incl. lactones), acid, ether, terpene/terpenoid, nitrogenous, sulphurous, aromatic (incl. benzenoids, pyrroles, and furans), hydrocarbon, and alcohol (Fig. S3b, Supplementary Table 2). For visualization purposes, we assigned compounds belonging to multiple classes to a main class in descending order of priority as follows: nitrogenous/sulphurous, terpene/terpenoid (only if *N*_carbons_ was a multiple of 5), aromatic, carbonyl-containing (aldehyde/ketone/ester/acid), alcohol, ether, hydrocarbon (Supplementary Table 2).

### Compound correlations (Fig. 2d,e; S6)

We calculated the correlation (Pearson’s *r*) in abundance across samples (one representative per species) for all pairs of compounds in the dataset. To shuffle the compound-by-samples matrix, we randomly permuted each row (abundance of one compound across samples), then renormalized the data so that each column (abundance of all compounds in one sample) summed to 1. The renormalization was essential to regenerate any negative correlations induced by dealing with proportional abundances. We generated 1000 shuffled matrices and calculated the pairwise compound correlation matrix for each one. The distributions of correlation coefficients for the actual and shuffled data were compared using a Kolmogorov–Smirnov test.

### Odour-space comparison (Fig. 3)

The floral-odour dataset was sourced from Kantsa et al. 2018. These authors calculated absolute emission rates; for better correspondence with our dataset, we scaled these abundances to sum to 1 within each species. Comparing odours from different studies merits caution, as sampling and analytical choices can drastically affect which compounds are detected. However, most of the usual caveats do not apply in this case because we focus on statistics that are agnostic to compound identity, such as the number of species across which compounds are found. The observed patterns could be inflated by other technical differences unrelated to compound identity, such as higher sensitivity to low-abundance compounds or noisier samples in our vertebrate dataset compared to the floral dataset. However, the patterns are apparent even when we consider only the higher-abundance compounds (Fig. 3e,f, lighter pink lines), instilling confidence in our main conclusions from this analysis.

The analysis underlying Fig. 3e,f was performed as follows. We randomly sampled a set of species from the dataset (with the size of the set given by the *x* axis) and counted the total number of compounds appearing in that set with an abundance greater than the given sensitivity threshold in any species. We performed this sampling 100 times for each set size.

### Phylogenetic analyses (Fig. 1a, 2a, 4, S7)

Most phylogenetic relationships and divergence-time estimates were adopted directly from estimates available at https://vertlife.org/ (Jetz et al. 2012; Tonini et al. 2016; Upham et al. 2019). The remaining few nodes were dated using data available at https://www.onezoom.org/ (Pleurodira/Cryptodira 209.2 MYA, Crocodilia/Aves 240.4 MYA, Testudines/Archosauria 244.2 MYA, Lepidosauria/Archelosauria 276 MYA, Synapsida/Sauropsida 301.7 MYA) (Rosindell and Harmon 2012). Odour distances were calculated as Euclidean distances in *n-*dimensional odour space, where *n* = the number of compounds. Hierarchical clustering of odours was performed using the complete-linkage method.

The analysis in Fig. 4d comparing odour distance within and between species was performed as follows. Each pair of dots with connecting line represents one species for which multiple replicate individuals are available in the dataset. A “same-species” dot represents the median Euclidean distance between all pairs of replicates for the given species. A “different-species” dot represents the median distance between a randomly sampled individual of that species and a randomly sampled individual of each of the other species included in the panel. Fig. 4e,f show analyses exactly analogous to Fig. 4d, but at the level of family and order instead of species.

### Analysis using normalized log-transformed compound abundances (Fig. 5c,f)

We wanted to test whether individual compounds were biased towards a focal group of samples (e.g., hosts vs. non-hosts or humans vs. non-human animals) irrespective of their abundance within a sample. A simple fold-change metric calculated from the raw abundances was unsatisfactory because of the many instances in which a compound was present in only a few samples, resulting in an infinite but non-significant fold change. Therefore, we adopted a different metric. We first log-transformed and normalized the abundances of compounds within a sample, as follows:

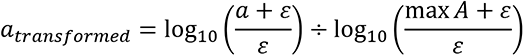

where

*A* denotes the set of abundances of a given compound across samples, *a* ∈ *A*, and *ε* = 10^−6^ (chosen to be smaller than the smallest non-zero value in the dataset: 3×10^−6^).

This transformation sets the highest observed abundance of a given compound at 1 and the estimated lower detection threshold at 0, then distributes values between these extremes on a log scale.

The significance of differences in compound abundance between target and non-target sample groups was calculated using Kolmogorov–Smirnov tests on the transformed abundances. We corrected *P*- values for multiple comparisons using the Benjamini–Hochberg method. The magnitude of differences was calculated as the mean difference in transformed abundances between sample groups. This value is analogous to fold change, but is scaled to the maximum observed abundance of each compound.

## Code availability

Analysis code is available at https://github.com/mcbridelab/Zung_2023_OdourSpace.

## Supplementary figures and tables

**Figure S1.**
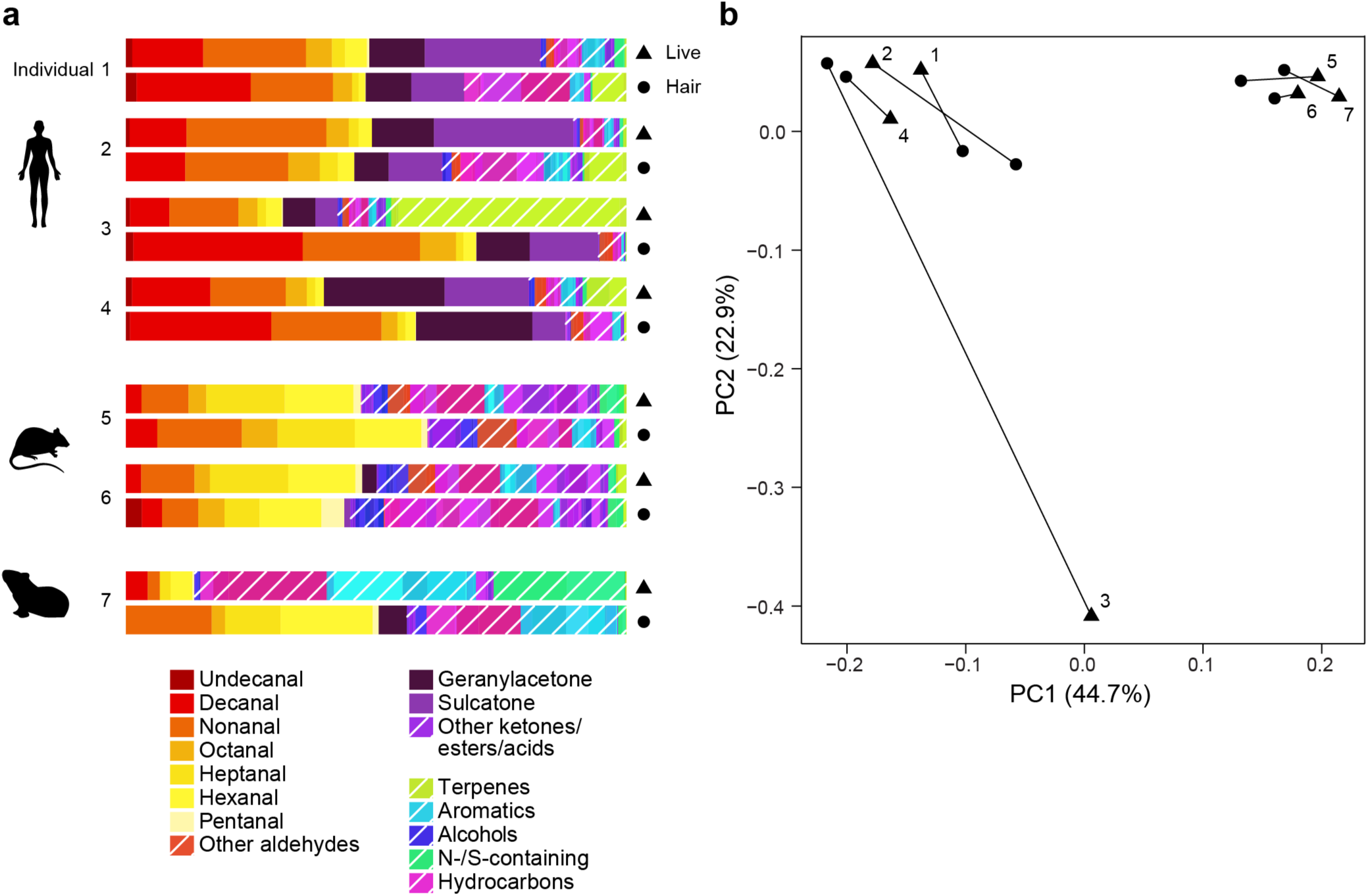
Comparison between odour extracted from live hosts vs. hair samples. **(a)** Odour composition of 7 individuals spanning 3 species. Human hair samples were taken from the head, and rat and guinea-pig samples from the whole back and sides of the animal. The guinea pig urinated and defecated during the live-host extraction. **(b)** PCA of the odour profiles in (a). Triangles and circles denote live-host and hair extractions, respectively. Line segments connect odour blends from the same individual. Note that the live-host sample from human subject 3 is an outlier in the PCA because it contains a high level of the terpene limonene, a common component of soaps and other skin products.

**Figure S2.**
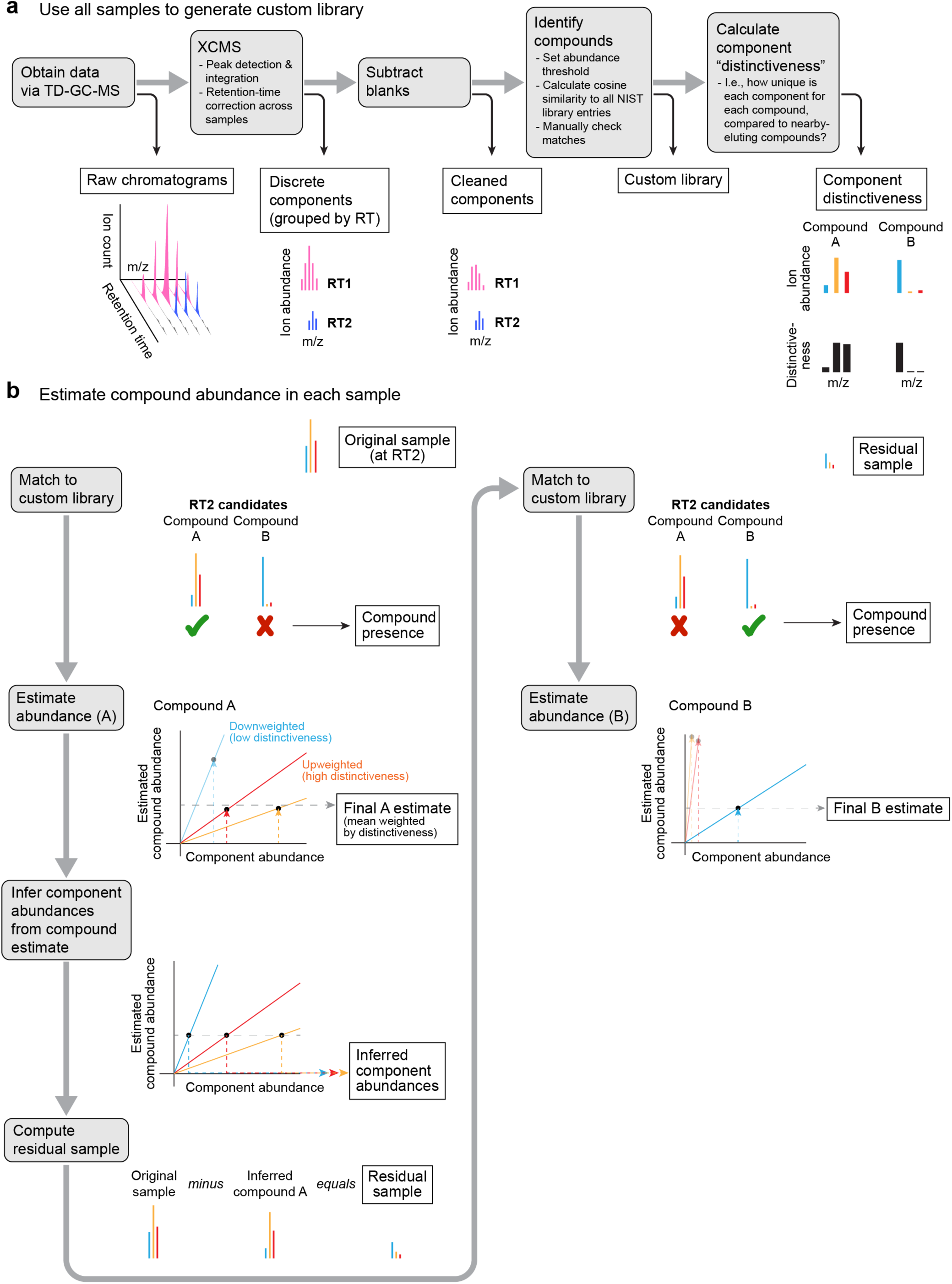
Odour-analysis pipeline. **(a)** Schematic of pipeline for identifying compounds present in the odour blends and creating a custom library of compounds. **(b)** Schematic of pipeline for deconvoluting compounds and estimating their abundance. See Methods for details.

**Figure S3.**
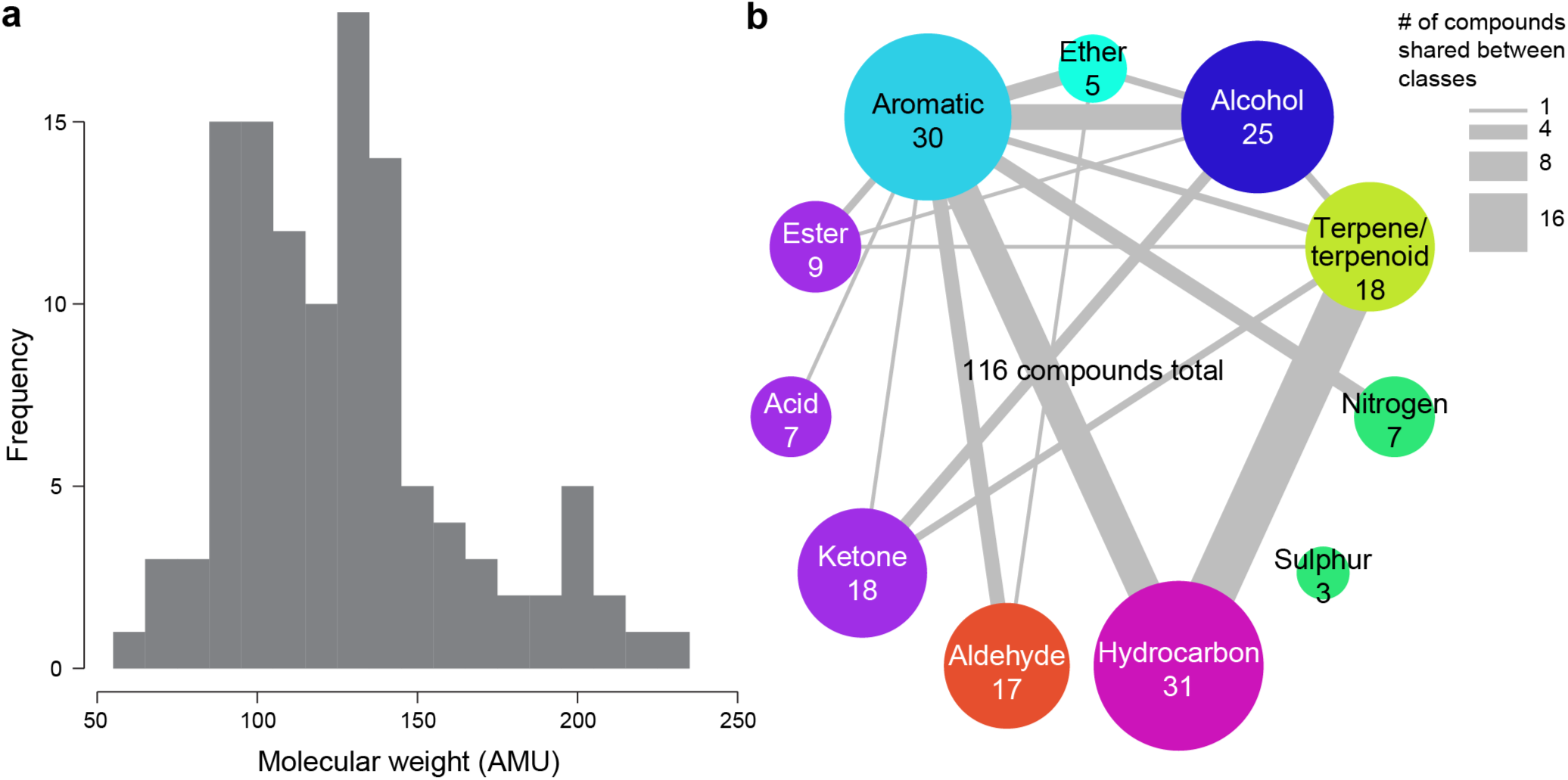
Additional compound information. **(a)** Molecular weight of compounds found in the dataset. Note that our methods undersample highly volatile (low-AMU) compounds. **(b)** Circle areas denote the number of compounds belonging to each class. Line widths show the number of compounds shared between two given classes. For visualization purposes in all other plots, compounds belonging to multiple classes were assigned a main class in descending order of priority as follows: nitrogenous/sulphurous, terpene/terpenoid (only if *N*_carbons_ was a multiple of 5), aromatic, carbonyl-containing (aldehyde/ketone/ester/acid), alcohol, ether, hydrocarbon.

**Figure S4.**
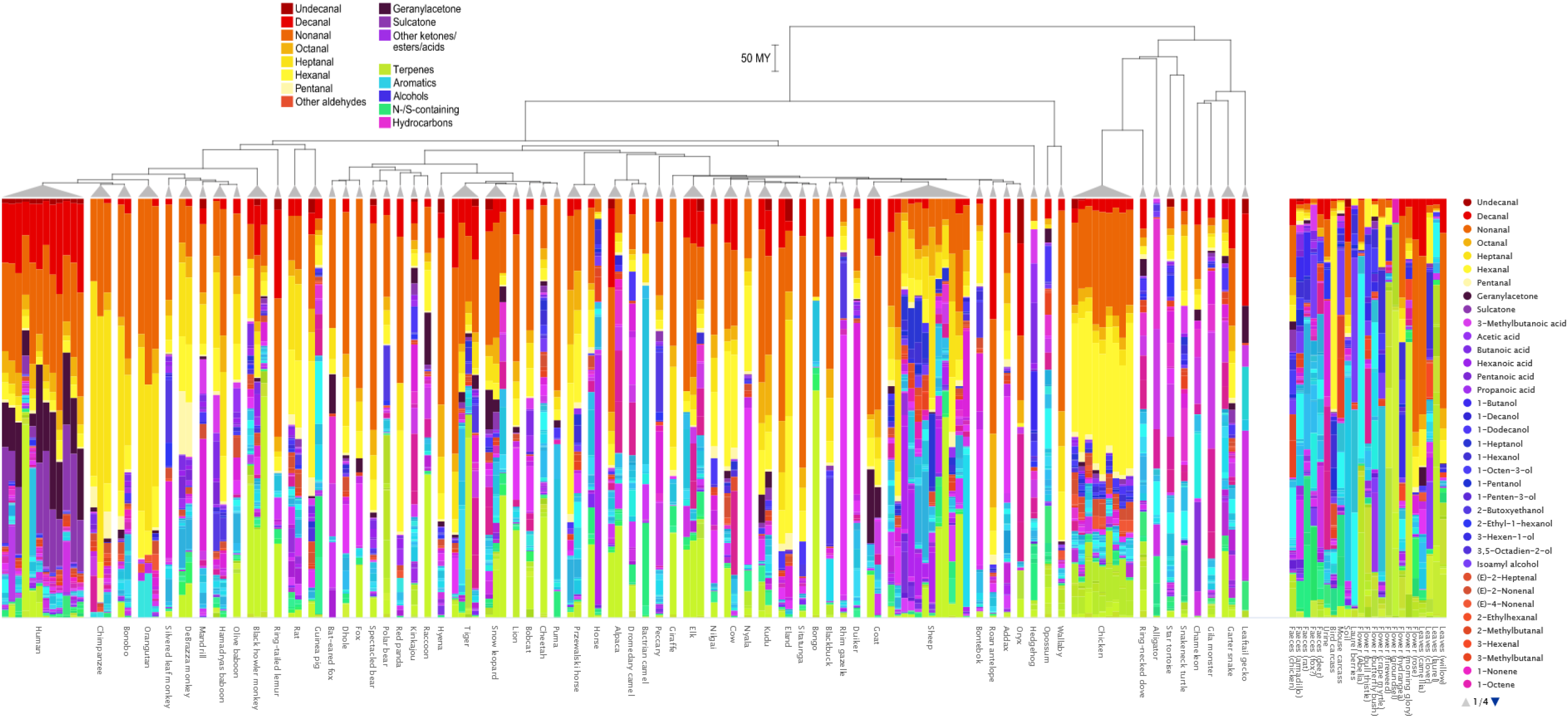
Interactive barplot showing odour profiles of all sampled individuals. Static screenshot shown above; please download the file for full functionality. A time-calibrated phylogeny is shown for animal samples. Colour scheme is the same as in Fig. 2a. Compounds are listed (in order by compound class, except for the first 9 compounds) in the scrollable legend on the right. Mousing over a compound in the legend highlights that compound in all samples. Mousing over a coloured bar in the barplot also highlights the given compound across all other samples and additionally shows a tooltip with more information about the focal sample, an image of the compound structure, and the exact numerical abundance of the compound in that sample.

**Figure S5.**
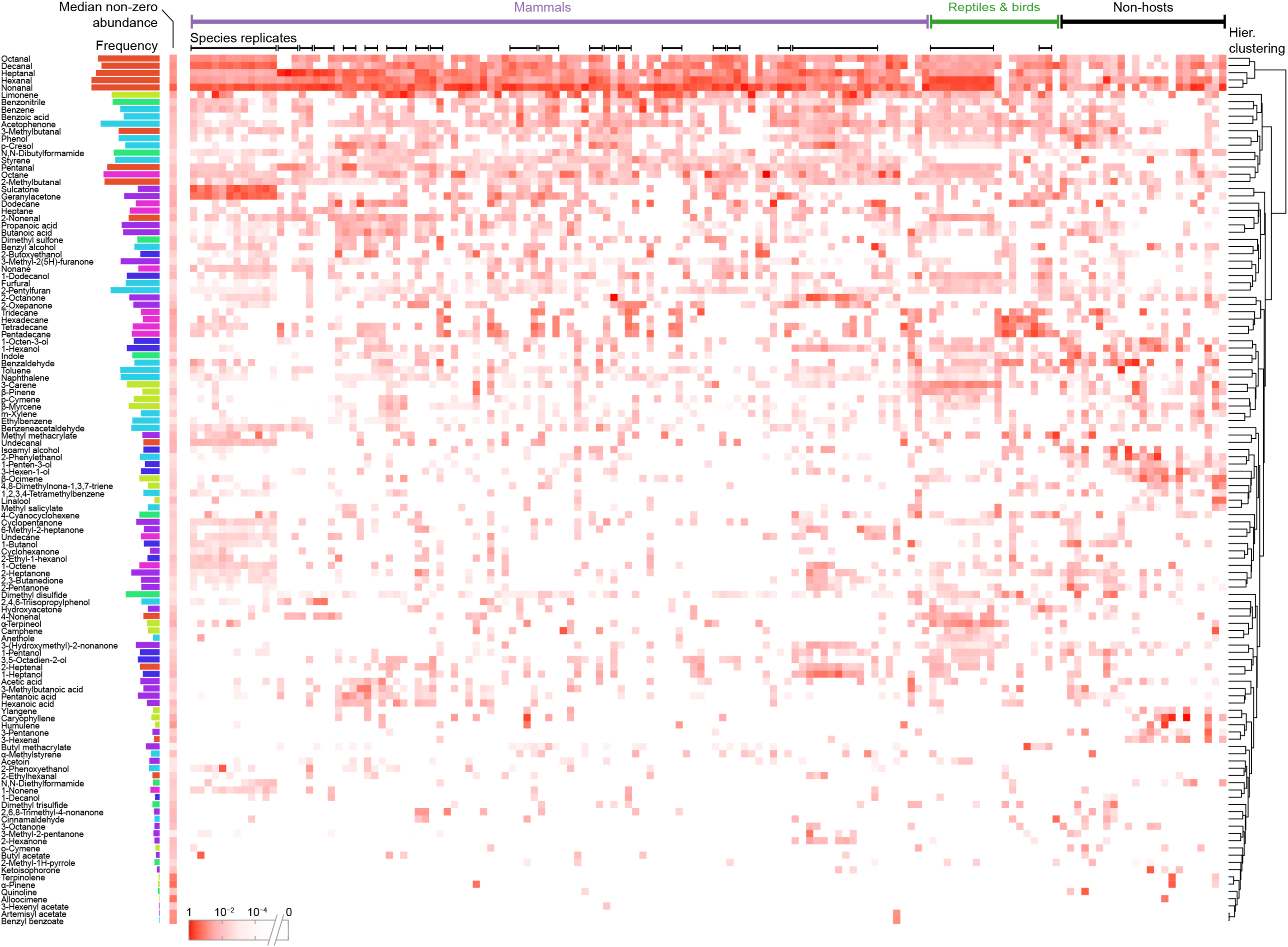
Compounds x samples heatmap. Heatmap shows compound abundance across all samples (including species replicates and non-host samples). Bars on the left show frequency (presence/absence) of compounds across all samples, coloured by compound class according to the colour scheme in Fig. S3b. Column immediately to the right of bars shows the median abundance of each compound across samples, excluding samples where the compound was not detected. Compounds are clustered by their abundance across samples (complete-linkage method), shown on the right. Samples are ordered as in Fig. S4.

**Figure S6.**
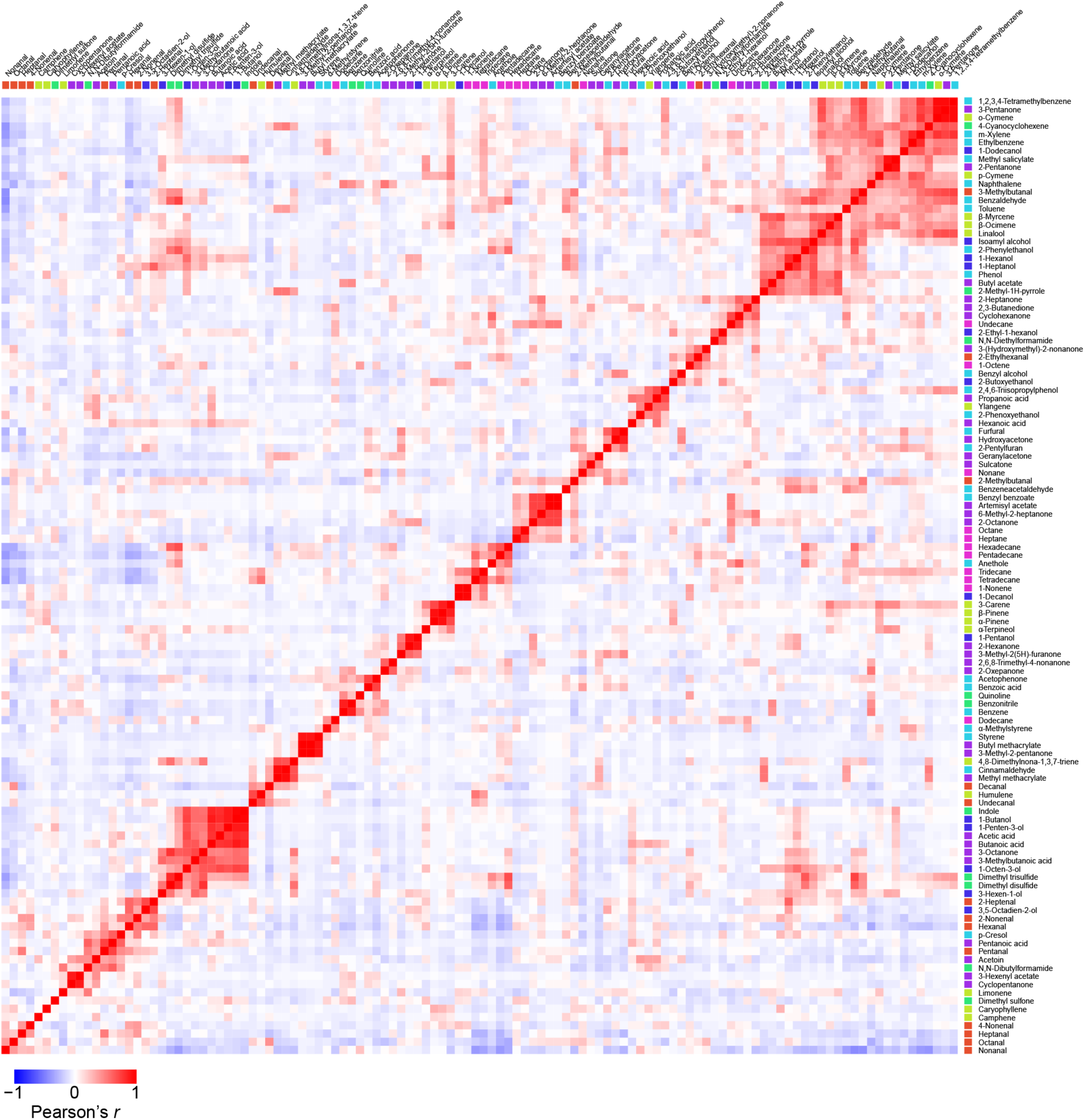
Pairwise compound correlations across species. Same correlation matrix as upper left in Fig. 2d, but with compound labels. Rows and columns are ordered by hierarchical clustering of compound correlations (complete-linkage method). Note that compound order differs from that in Fig. S5 because (1) the correlation analysis includes only one representative per species and (2) it does not consider non-host samples. Coloured boxes next to compound labels denote compound class according to the scheme in Fig. 2c, S3b.

**Figure S7.**
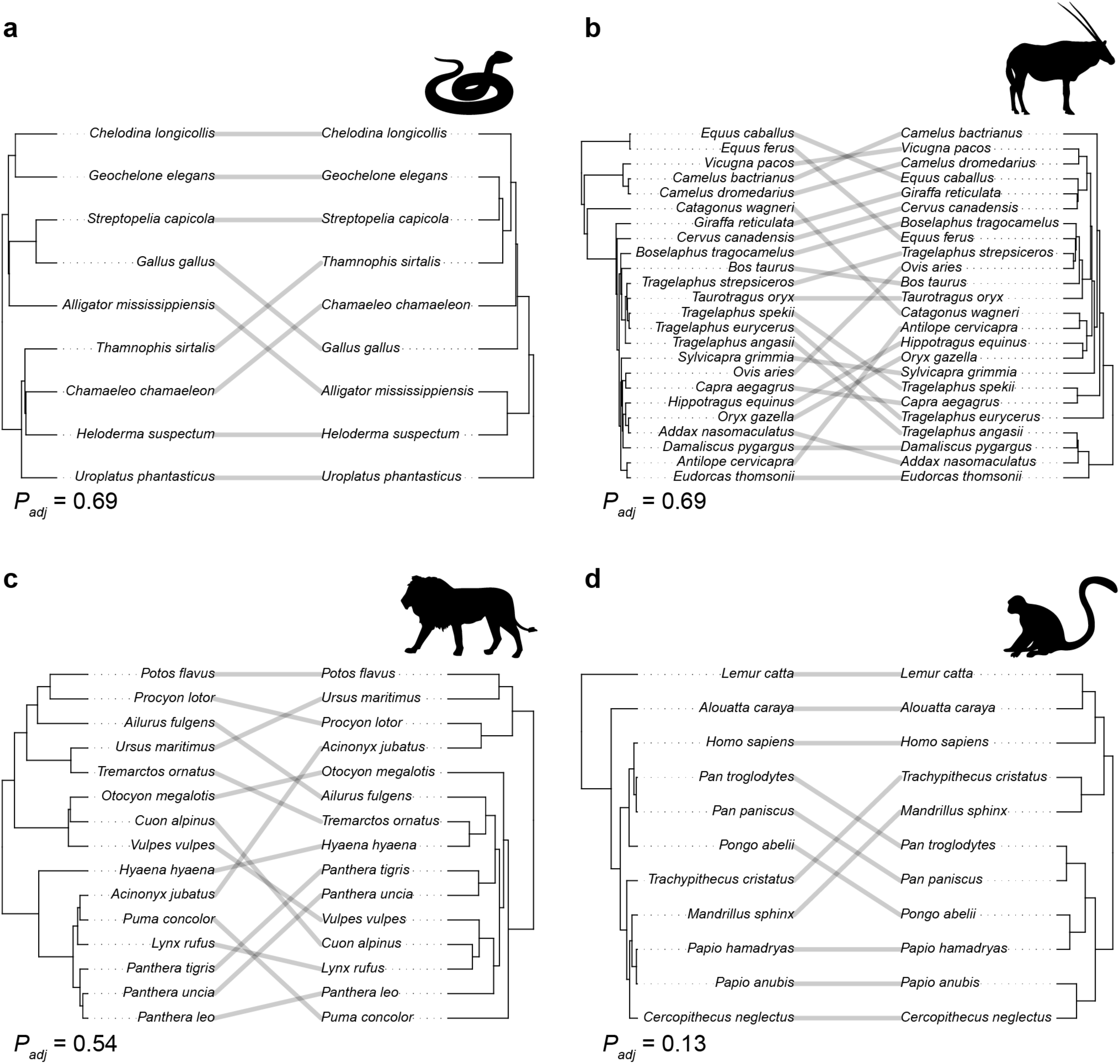
Association between phylogeny and odour profile for subclades. Same type of analysis as in Fig. 4a, but for four smaller clades: **(a)** reptiles and birds, **(b)** ungulates, **(c)** carnivores, and **(d)** primates. *P*-values shown are for Mantel tests, adjusted for multiple comparisons using the Benjamini–Hochberg method.

**Figure S8.**
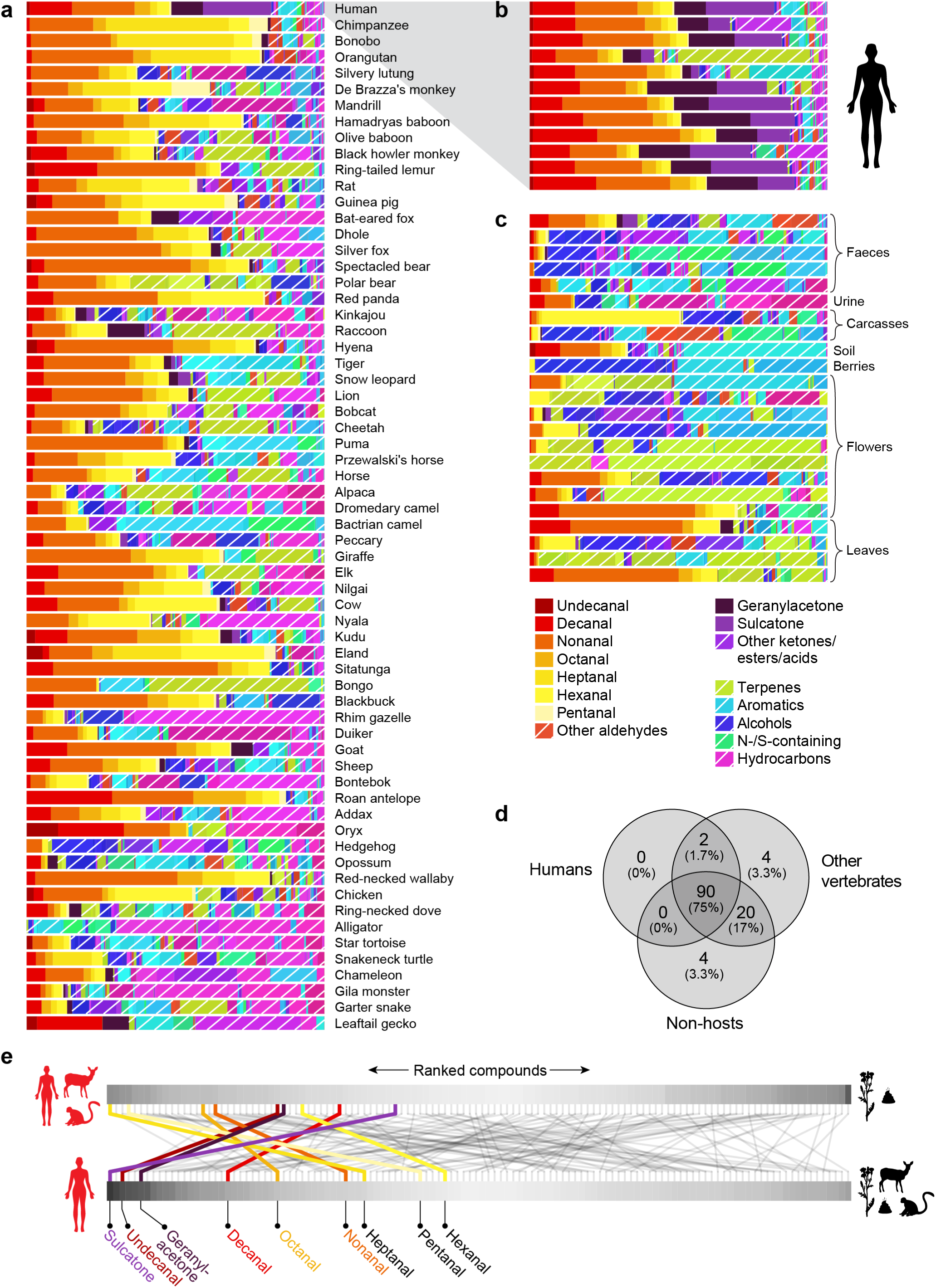
Composition of odour blends and host discrimination. **(a–c)** Overview of odour blends used in the analyses underlying Fig. 5. **(a)** Odour-blend composition of one representative from each animal species sampled. Same data as in Fig. 2a. **(b)** Odour blends from 12 individual humans. **(c)** Odour blends from 23 non-host extracts. The analyses in Fig. 5b,c compare the blends shown in (a) to the blends shown in (c). Analyses in Fig. 5e,f compare the human odour blends shown in (b) to the non-human blends shown in both (a) and (c). **(d)** Number of compounds (and percentage of the total) shared among groups of odour blends. **(e)** Rank change in informativeness of compounds for generalists vs. specialists. Grey bars show compounds ranked and shaded by scaled fold difference (see Fig. 5c,f) for the generalist (top) and specialist (bottom) tasks. Lines trace changes in rank between the tasks.

**Supplementary Table 1.** Information about the 150 odour extracts analysed in this study, including details about the individual animals studied, hair samples, extraction parameters, and taxonomic information. The “species_representative” column indicates whether each extract was included in the dataset that comprises only one extract from each species.

**Supplementary Table 2.** All compounds identified in this study. The “matching_NIST_entry” column shows the exact name of the entry in the NIST library whose mass spectrum best matched each compound. The “all_classes” column indicates the chemical classes to which each compound belongs, and the “main_class” column shows the class to which the compound was assigned for visualization purposes (see Methods).

**Supplementary Table 3.** Proportional abundance of compounds in each odour blend. This exact matrix is illustrated in Figure S4.

