## Supplementary material for "Exploring natural odour landscapes: A case study with implications for human-biting insects": Figure S4: highchart.js

```
HTMLWidgets.widget({
  
  name: 'highchart',

  type: 'output',

  initialize: function(el, width, height) {

    return {
      // TODO: add instance fields as required
    };

  },

  renderValue: function(el, x, instance) {
    
    if(x.debug) {
      window.xclone = JSON.parse(JSON.stringify(x));
      window.elclone = $(el);
      console.log(el);
      console.log("hc_opts", x.hc_opts);
      console.log("theme", x.theme);
      console.log("conf_opts", x.conf_opts);
    }

    if(x.fonts !== undefined) {
      
      x.fonts = ((typeof(x.fonts) == "string") ? [x.fonts] : x.fonts);
    
      x.fonts.forEach(function(s){
        /* http://stackoverflow.com/questions/4724606 */
        var urlfont = 'https://fonts.googleapis.com/css?family=' + s;
        if (!$("link[href='" + urlfont + "']").length) {
          $('<link href="' + urlfont + '" rel="stylesheet" type="text/css">').appendTo("head");
        }
        
      });
      
    }
    
    ResetHighchartsOptions();
    
    if(x.theme !== null) {
      
      if(x.debug) console.log("adding THEME");
      
      Highcharts.setOptions(x.theme);
      
    }
    
    if((x.theme && x.theme.chart.divBackgroundImage !== undefined) |
         (x.hc_opts.chart  && x.hc_opts.chart.divBackgroundImage !== undefined)) {
           
      if(x.debug) console.log("adding BackgroundImage");     
           
      var bkgrnd = x.theme.chart.divBackgroundImage || x.hc_opts.chart.divBackgroundImage;
      
      Highcharts.wrap(Highcharts.Chart.prototype, "getContainer", function (proceed) {
        
        proceed.call(this);
        
        $("#" + el.id).css("background-image", "url(" + bkgrnd + ")");
        $("#" + el.id).css("-webkit-background-size", "cover");
        $("#" + el.id).css("-moz-background-size", "cover");
        $("#" + el.id).css("-o-background-size", "cover");
        $("#" + el.id).css("background-size", "cover");
        
      });
      
    }
    
    Highcharts.setOptions(x.conf_opts);
    
    if(x.type == "chart") {
      if(x.debug) console.log("charting CHART");
      $("#" + el.id).highcharts(x.hc_opts);
    } else if (x.type == "stock") {
      if(x.debug) console.log("charting STOCK");
      $("#" + el.id).highcharts('StockChart', x.hc_opts);  
    } else if (x.type == "map"){
      if(x.debug) console.log("charting MAP");

      x.hc_opts.series = x.hc_opts.series.map(function(e){
        if(e.geojson === true) {
          if(x.debug) console.log("geojson\n\t", e.type, "\n\t", typeof(e.series));
          e.data = Highcharts.geojson(e.data, e.type);
        }
        return e;
      });
      
      $("#" + el.id).highcharts('Map', x.hc_opts); 
      
      if(x.hc_opts.mapNavigation !== undefined && x.hc_opts.mapNavigation.enabled === true){
        /* if have navigation option and enabled true: http://stackoverflow.com/questions/7600454 */
        $("#" + el.id).bind( 'mousewheel DOMMouseScroll', function ( e ) {
          var e0 = e.originalEvent,
          delta = e0.wheelDelta || -e0.detail;
          this.scrollTop += ( delta < 0 ? 1 : -1 ) * 30;
          e.preventDefault();

        });
        
      }
      
    } else if (x.type == "gantt") {
      
      if(x.debug) console.log("charting GANTT");
      
      chart = Highcharts.ganttChart(el.id, x.hc_opts);
      
    }
    
    if(x.hc_opts.motion !== undefined) {
      
      $("#" + el.id).css({"position" : "relative" });
      
      if(x.debug) console.log("setting MOTION options");
      
      var pc = $($("#" + el.id).find("#play-controls")[0]);
      
      var ct = x.theme.chart;
      
      if(ct.backgroundColor !== undefined) $(pc.find("#play-pause-button")[0]).css({backgroundColor : x.theme.chart.backgroundColor});
      if(ct.style !== undefined)  $(pc.find("#play-output")[0]).css(x.theme.chart.style);
      if(ct.style !== undefined && ct.style.color !== undefined) $(pc.find("#play-pause-button")[0]).css({color : x.theme.chart.style.color});
      
      
    } 
    
  },

  resize: function(el, width, height, instance) {
    
    /* http://stackoverflow.com/questions/18445784/ */
    var chart = $("#" +el.id).highcharts();
    
    if (chart && chart.options.chart.reflow === true) {  // _check for reflow option_
      var w = chart.renderTo.clientWidth; 
      var h = chart.renderTo.clientHeight; 
      chart.setSize(w, h); 
    }
    
  }

});

if (HTMLWidgets.shinyMode) {
  
  Shiny.addCustomMessageHandler('addSeries', function(msg) {
    
    var chart = $("#" + msg.id).highcharts();
    
    chart.addSeries(msg.series);
      
  });
  
  Shiny.addCustomMessageHandler('removeSeries', function(msg) {
    
    var chart = $("#" + msg.id).highcharts();
    
    chart.get(msg.idSeries).remove();
    
  });
  
  Shiny.addCustomMessageHandler('removeAllSeries', function(msg) {
    
    var chart = $("#" + msg.id).highcharts();
    
    while (chart.series.length) {
      chart.series[0].remove();
      }
    
  });
  
  Shiny.addCustomMessageHandler('updateChart', function(msg) {
    
    var chart = $("#" + msg.id).highcharts();
    
    chart.update(msg.options);
   
  });
  
  Shiny.addCustomMessageHandler('updateSeries', function(msg) {
    
    var chart = $("#" + msg.id).highcharts();
    
    chart.get(msg.idSeries).update(msg.options);
    
  });
  
  Shiny.addCustomMessageHandler('updatePoint', function(msg) {
    
    var chart = $("#" + msg.id).highcharts();
    
    chart.get(msg.idSeries).data[msg.idPoint].update(msg.options);
    
  });
  
  Shiny.addCustomMessageHandler('showLoading', function(msg) {
    
    var chart = $("#" + msg.id).highcharts();
    
    if (msg.showLoading) {
      
      chart.showLoading();
      
    } else {
      
      chart.hideLoading();
      
    }
      
  });
  
  Shiny.addCustomMessageHandler('addPoint', function(msg) {
    
    var chart = $("#" + msg.id).highcharts();
    
    chart.get(msg.idSeries).addPoint(msg.point, msg.redraw, msg.shift, msg.animation);
    
  });
  
  Shiny.addCustomMessageHandler('removePoint', function(msg) {
    
    var chart = $("#" + msg.id).highcharts();
    
    chart.get(msg.idSeries).removePoint(msg.i, msg.redraw);
    
  });
  
  Shiny.addCustomMessageHandler('setData', function(msg) {
    
    var chart = $("#" + msg.id).highcharts();
    
    if (typeof chart != 'undefined') {
      
      chart.series[msg.serie].setData(
        data = msg.data,
        redraw = msg.redraw,
        animation = msg.animation,
        updatePoints = msg.updatePoints
        );
      
    }
    
  });
  
  Shiny.addCustomMessageHandler('redraw', function(msg) {
    
    var chart = $("#" + msg.id).highcharts();
    
    if (typeof chart != 'undefined') {
      
      chart.redraw();
      
    }
      
  });
  
}
```
