## Supplementary material for "Exploring natural odour landscapes: A case study with implications for human-biting insects": Figure S4: appear.js

```
/**
* https://jsfiddle.net/gh/get/library/pure/highcharts/highcharts/tree/master/samples/highcharts/studies/appear/ 
* Highcharts plugin to defer initial series animation until the element has
* appeared.
*
  * Updated 2019-04-10
*
  * @todo
* - If the element is greater than the viewport (or a certain fraction of it),
*   show the series when it is partially visible.
*/
  (function (H) {
    
    var pendingRenders = [];
    
    // https://stackoverflow.com/questions/123999/how-to-tell-if-a-dom-element-is-visible-in-the-current-viewport/7557433#7557433
    function isElementInViewport(el) {
      
      var rect = el.getBoundingClientRect();
      
      return (
        rect.top >= 0 &&
          rect.left >= 0 &&
          rect.bottom <= (
            window.innerHeight ||
              document.documentElement.clientHeight
          ) &&
          rect.right <= (
            window.innerWidth ||
              document.documentElement.clientWidth
          )
      );
    }
    
    H.wrap(H.Series.prototype, 'render', function deferRender(proceed) {
      var series = this,
      renderTo = this.chart.container.parentNode;
      
      // It is appeared, render it
      if (isElementInViewport(renderTo) || !series.options.animation) {
        proceed.call(series);
        
        // It is not appeared, halt renering until appear
      } else  {
        pendingRenders.push({
          element: renderTo,
          appear: function () {
            proceed.call(series);
          }
        });
      }
    });
    
    function recalculate() {
      pendingRenders.forEach(function (item) {
        if (isElementInViewport(item.element)) {
          item.appear();
          H.erase(pendingRenders, item);
        }
      });
    }
    
    if (window.addEventListener) {
      ['DOMContentLoaded', 'load', 'scroll', 'resize']
      .forEach(function (eventType) {
        addEventListener(eventType, recalculate, false);
      });
    }
    
  }(Highcharts));
```
