## Supplementary material for "Exploring natural odour landscapes: A case study with implications for human-biting insects": Figure S4: delay-animation.js

```
/**
 * by @TorsteinHonsi and modified by @frank-eisen
 * returns a highcharts animation object with an own easing-function
 * @param {number} duration
 * @param {number} delay
 * @param {function} timingFunction
 * @return {number|object}
 */
function highchartsAnimation(duration, delay, timingFunction)
{
  if(typeof duration === 'number' && typeof delay === 'number' && typeof timingFunction === 'function') {
    var delay_percent = delay / (duration + delay),
        perc = 1 - delay_percent;
    
    return {
      duration: duration + delay,
      easing: function(pos) {
        if(pos <= delay_percent) {
          return 0;
        }
        else {
          return timingFunction((pos - delay_percent) / perc);
        }
      }
    };
  }
  else {
    return 750;
  }
}

/**
 * returns a timing function
 * @param name
 * @return {function}
 */
function getTimingFunction(name)
{
	var args = arguments;

	function needArgumentsCount(need_count)
	{
		if(args.length < need_count) {
			throw new Error('TimingFunction.' + name + ' needs ' + need_count + ' arguments');
		}
	}

	switch(name) {
		case 'pow':
		case 'power':
			needArgumentsCount(1);
			return function power(percent_done) {
				return Math.pow(percent_done, args[1]);
			};

		case 'quad':
			return getTimingFunction.apply(this, ['pow', 2]);

		case 'cubic':
			return getTimingFunction.apply(this, ['pow', 3]);

		case 'sqrt':
		case 'squareroot':
			return function sqrt(percent_done) {
				return Math.sqrt(percent_done);
			};

		case 'sine':
		case 'sinus':
			return function sinus(percent_done) {
				return Math.sin(percent_done * (Math.PI / 2));
			};

		case 'steps':
			needArgumentsCount(2);
			return function steps(percent_done) {
				return Math.floor(percent_done * (args[1])) / (args[1]);
			};

		case 'bezier':
			needArgumentsCount(5);

			var bezier = simpleBezier([0, 0], [args[1], args[2]], [args[3], args[4]], [1, 1]);

			return function bez(percent_done) {
				var coords = bezier(percent_done);
				
				return coords[1];
			};

		case 'linear':
		default:
			return function linear(percent_done) {
				return percent_done;
			};
	}


	function simpleBezier(p0, p1, p2, p3)
	{
		var x_c = getC(p0[0], p1[0]),
			x_b = getB(p1[0], p2[0], x_c),
			x_a = getA(p0[0], p3[0], x_b, x_c),
			y_c = getC(p0[1], p1[1]),
			y_b = getB(p1[1], p2[1], y_c),
			y_a = getA(p0[1], p3[1], y_b, y_c);

		function getC(p0, p1)
		{
			return 3 * (p1 - p0);
		}

		function getB(p1, p2, c)
		{
			return 3 * (p2 - p1) - c
		}

		function getA(p0, p3, b, c)
		{
			return p3 - p0 - c - b
		}

		function calculateBezier(a, b, c, t, position)
		{
			return (a * Math.pow(t, 3)) + (b * Math.pow(t, 2)) + (c * t) + position;
		}

		return function getPositionFor(t) {
			return [
				calculateBezier(x_a, x_b, x_c, t, p0[0]),
				calculateBezier(y_a, y_b, y_c, t, p0[1])
			];
		};
	}
}
```
