## Supplementary material for "Exploring natural odour landscapes: A case study with implications for human-biting insects": Figure S4: reset.js

```
var HCDefaults = $.extend(true, {}, Highcharts.getOptions(), {});

function ResetHighchartsOptions() {
    // Fortunately, Highcharts returns the reference to defaultOptions itself
    // We can manipulate this and delete all the properties
    var defaultOptions = Highcharts.getOptions();
    for (var prop in defaultOptions) {
        if (typeof defaultOptions[prop] !== 'function') delete defaultOptions[prop];
    }
    // Fall back to the defaults that we captured initially, this resets the theme
    Highcharts.setOptions(HCDefaults);
}
```
