## Supplementary material for "Exploring natural odour landscapes: A case study with implications for human-biting insects": Figure S4: symbols-extra.js

```
(function (H) {
  
  // cross http://stackoverflow.com/a/25379352/829971
  Highcharts.Renderer.prototype.symbols.cross = function (x, y, radius) {
    var r = radius * 0.4,
        e = r * 0.8,
        a = e / Math.sqrt(2),
        p = r / Math.sqrt(2);
    return [
        'M', x, y + a,
        'L', x + p, y + a + p,
        x, y + a + (2 * p),
        x + a, y + (2 * a) + (2 * p),
        x + a + p, y + (2 * a) + p,
        x + a + (2 * p), y + (2 * a) + (2 * p),
        x + (2 * a) + (2 * p), y + a + (2 * p),
        x + (2 * a) + p, y + a + p,
        x + (2 * a) + (2 * p), y + a,
        x + a + (2 * p), y,
        x + a + p, y + p,
        x + a, y,
        'Z'];
    };

  // plus http://stackoverflow.com/a/36270224/829971
  Highcharts.SVGRenderer.prototype.symbols.plus = function (x, y, w, h) {
    return [
        'M', x, y + (5 * h) / 8,
        'L', x, y + (3 * h) / 8,
        'L', x + (3 * w) / 8, y + (3 * h) / 8,
        'L', x + (3 * w) / 8, y,
        'L', x + (5 * w) / 8, y,
        'L', x + (5 * w) / 8, y + (3 * h) / 8,
        'L', x + w, y + (3 * h) / 8,
        'L', x + w, y + (5 * h) / 8,
        'L', x + (5 * w) / 8, y + (5 * h) / 8,
        'L', x + (5 * w) / 8, y + h,
        'L', x + (3 * w) / 8, y + h,
        'L', x + (3 * w) / 8, y + (5 * h) / 8,
        'L', x, y + (5 * h) / 8,
        'z'
    ];
  };
  
  
}(Highcharts));
```
