## Supplementary material for "Exploring natural odour landscapes: A case study with implications for human-biting insects": Figure S4: text-symbols.js

```
/* Highcharts plugin to handle text symbols */
(function (H) {
    function symbolWrap(proceed, symbol, x, y, w, h, options) {
        if (symbol.indexOf('text:') === 0) {
            var text = unescape(JSON.parse('"\\u' + symbol.split(':')[1] + '"')),
                svgElem = this.text(text, x, y)
                  .attr({
                    translateY: h,
                    translateX: -1
                  })
                  .css({
                    fontFamily: 'FontAwesome',
                    fontSize: h * 2
                  });
                
            if (svgElem.renderer.isVML) {
                svgElem.fillSetter = function (value, key, element) {
                    element.style.color = H.Color(value).get('rgb');
                };
            }
            return svgElem;
        }
        return proceed.apply(this, [].slice.call(arguments, 1));
    }
    H.wrap(H.SVGRenderer.prototype, 'symbol', symbolWrap);
    if (H.VMLRenderer) {
        H.wrap(H.VMLRenderer.prototype, 'symbol', symbolWrap);
    }
    
    // Load the font for SVG files also
    /*
    H.wrap(H.Chart.prototype, 'getSVG', function (proceed) {
        var svg = proceed.call(this);
        svg = '<?xml-stylesheet type="text/css" ' +
            'href="http://netdna.bootstrapcdn.com/font-awesome/4.1.0/css/font-awesome.min.css" ?>' + 
            svg;
        return svg;
    });
    */
}(Highcharts));
```
