## Supplementary material for "Exploring natural odour landscapes: A case study with implications for human-biting insects": Figure S4: tooltip-delay.js

```
(function(H) {

  let timerId = {};

  const generatePointsUniqueKey = (points) => {

    const generatePointKey = (point) => {
      return point.category + " " + point.series.name + ": " + point.x + " " + point.y;
    };

    const result = points.map(generatePointKey).join(', ');

    return result;
  }

  H.wrap(H.Tooltip.prototype, 'refresh', function(proceed) {
      let seriesName;

      if (Array.isArray(arguments[ 1 ])) {
        // Can be array in case that, it's shared tooltip
        seriesName = generatePointsUniqueKey(arguments[ 1 ]);
      } else {
        seriesName = arguments[ 1 ].series.name;
      }

      const delayForDisplay = this.chart.options.tooltip.delayForDisplay ? this.chart.options.tooltip.delayForDisplay : 1000;

      if (timerId[ seriesName ]) {
        clearTimeout(timerId[ seriesName ]);
        delete timerId[ seriesName ];
      }

      timerId[ seriesName ] = window.setTimeout(function() {
        let pointOrPoints = this.refreshArguments[ 0 ];

        if (pointOrPoints === this.chart.hoverPoint || $.inArray(this.chart.hoverPoint, pointOrPoints) > -1) {
          proceed.apply(this.tooltip, this.refreshArguments);
        }

      }.bind({
        refreshArguments: Array.prototype.slice.call(arguments, 1),
        chart: this.chart,
        tooltip: this
      }), delayForDisplay);
    
  });

}(Highcharts));
```
