## Supplementary material for "Exploring natural odour landscapes: A case study with implications for human-biting insects": Figure S4: highcharts-3d.js

```
/*
 Highcharts JS v9.3.1 (2021-11-05)

 3D features for Highcharts JS

 License: www.highcharts.com/license
*/
'use strict';(function(a){"object"===typeof module&&module.exports?(a["default"]=a,module.exports=a):"function"===typeof define&&define.amd?define("highcharts/highcharts-3d",["highcharts"],function(E){a(E);a.Highcharts=E;return a}):a("undefined"!==typeof Highcharts?Highcharts:void 0)})(function(a){function E(a,w,t,D){a.hasOwnProperty(w)||(a[w]=D.apply(null,t))}a=a?a._modules:{};E(a,"Extensions/Math3D.js",[a["Core/Globals.js"],a["Core/Utilities.js"]],function(a,w){function t(d,c,b){c=0<b&&b<Number.POSITIVE_INFINITY?
b/(d.z+c.z+b):1;return{x:d.x*c,y:d.y*c}}function F(d,c,b,f){var k=c.options.chart.options3d,a=x(f,b?c.inverted:!1),e={x:c.plotWidth/2,y:c.plotHeight/2,z:k.depth/2,vd:x(k.depth,1)*x(k.viewDistance,0)},n=c.scale3d||1;f=l*k.beta*(a?-1:1);k=l*k.alpha*(a?-1:1);var u=Math.cos(k),q=Math.cos(-f),p=Math.sin(k),C=Math.sin(-f);b||(e.x+=c.plotLeft,e.y+=c.plotTop);return d.map(function(b){var c=(a?b.y:b.x)-e.x;var d=(a?b.x:b.y)-e.y;b=(b.z||0)-e.z;c={x:q*c-C*b,y:-p*C*c+u*d-q*p*b,z:u*C*c+p*d+u*q*b};d=t(c,e,e.vd);
d.x=d.x*n+e.x;d.y=d.y*n+e.y;d.z=c.z*n+e.z;return{x:a?d.y:d.x,y:a?d.x:d.y,z:d.z}})}function f(d,c){var b=c.options.chart.options3d,f=c.plotWidth/2;c=c.plotHeight/2;b=x(b.depth,1)*x(b.viewDistance,0)+b.depth;return Math.sqrt(Math.pow(f-x(d.plotX,d.x),2)+Math.pow(c-x(d.plotY,d.y),2)+Math.pow(b-x(d.plotZ,d.z),2))}function z(d){var c=0,b;for(b=0;b<d.length;b++){var f=(b+1)%d.length;c+=d[b].x*d[f].y-d[f].x*d[b].y}return c/2}function v(d,c,b){return z(F(d,c,b))}var x=w.pick,l=a.deg2rad;a.perspective3D=t;
a.perspective=F;a.pointCameraDistance=f;a.shapeArea=z;a.shapeArea3d=v;return{perspective:F,perspective3D:t,pointCameraDistance:f,shapeArea:z,shapeArea3D:v}});E(a,"Core/Renderer/SVG/SVGElement3D.js",[a["Core/Color/Color.js"],a["Core/Renderer/SVG/SVGElement.js"],a["Core/Utilities.js"]],function(a,w,t){var F=a.parse,f=t.defined;a=t.merge;var z=t.objectEach,v=t.pick,x={base:{initArgs:function(f){var d=this,c=d.renderer,b=c[d.pathType+"Path"](f),a=b.zIndexes;d.parts.forEach(function(f){d[f]=c.path(b[f]).attr({"class":"highcharts-3d-"+
f,zIndex:a[f]||0}).add(d)});d.attr({"stroke-linejoin":"round",zIndex:a.group});d.originalDestroy=d.destroy;d.destroy=d.destroyParts;d.forcedSides=b.forcedSides},singleSetterForParts:function(f,d,c,b,a,k){var l={};b=[null,null,b||"attr",a,k];var e=c&&c.zIndexes;c?(e&&e.group&&this.attr({zIndex:e.group}),z(c,function(b,d){l[d]={};l[d][f]=b;e&&(l[d].zIndex=c.zIndexes[d]||0)}),b[1]=l):(l[f]=d,b[0]=l);return this.processParts.apply(this,b)},processParts:function(f,d,c,b,a){var l=this;l.parts.forEach(function(k){d&&
(f=v(d[k],!1));if(!1!==f)l[k][c](f,b,a)});return l},destroyParts:function(){this.processParts(null,null,"destroy");return this.originalDestroy()}}};x.cuboid=a(x.base,{parts:["front","top","side"],pathType:"cuboid",attr:function(a,d,c,b){if("string"===typeof a&&"undefined"!==typeof d){var l=a;a={};a[l]=d}return a.shapeArgs||f(a.x)?this.singleSetterForParts("d",null,this.renderer[this.pathType+"Path"](a.shapeArgs||a)):w.prototype.attr.call(this,a,void 0,c,b)},animate:function(a,d,c){if(f(a.x)&&f(a.y)){a=
this.renderer[this.pathType+"Path"](a);var b=a.forcedSides;this.singleSetterForParts("d",null,a,"animate",d,c);this.attr({zIndex:a.zIndexes.group});b!==this.forcedSides&&(this.forcedSides=b,x.cuboid.fillSetter.call(this,this.fill))}else w.prototype.animate.call(this,a,d,c);return this},fillSetter:function(a){this.forcedSides=this.forcedSides||[];this.singleSetterForParts("fill",null,{front:a,top:F(a).brighten(0<=this.forcedSides.indexOf("top")?0:.1).get(),side:F(a).brighten(0<=this.forcedSides.indexOf("side")?
0:-.1).get()});this.color=this.fill=a;return this}});return x});E(a,"Core/Renderer/SVG/SVGRenderer3D.js",[a["Core/Animation/AnimationUtilities.js"],a["Core/Color/Color.js"],a["Core/Globals.js"],a["Extensions/Math3D.js"],a["Core/Renderer/SVG/SVGElement.js"],a["Core/Renderer/SVG/SVGElement3D.js"],a["Core/Renderer/SVG/SVGRenderer.js"],a["Core/Utilities.js"]],function(a,w,t,D,f,z,v,x){var l=this&&this.__extends||function(){var b=function(e,d){b=Object.setPrototypeOf||{__proto__:[]}instanceof Array&&function(b,
h){b.__proto__=h}||function(b,h){for(var r in h)h.hasOwnProperty(r)&&(b[r]=h[r])};return b(e,d)};return function(e,d){function c(){this.constructor=e}b(e,d);e.prototype=null===d?Object.create(d):(c.prototype=d.prototype,new c)}}(),d=a.animObject,c=w.parse,b=t.charts,p=t.deg2rad,k=D.perspective,A=D.shapeArea,e=x.defined,n=x.extend,u=x.merge,q=x.pick,G=Math.cos,C=Math.sin,H=Math.PI,B=4*(Math.sqrt(2)-1)/3/(H/2);return function(a){function m(){return null!==a&&a.apply(this,arguments)||this}l(m,a);m.compose=
function(b){b=b.prototype;var e=m.prototype;b.elements3d=z;b.arc3d=e.arc3d;b.arc3dPath=e.arc3dPath;b.cuboid=e.cuboid;b.cuboidPath=e.cuboidPath;b.element3d=e.element3d;b.face3d=e.face3d;b.polyhedron=e.polyhedron;b.toLinePath=e.toLinePath;b.toLineSegments=e.toLineSegments};m.curveTo=function(b,e,h,r,g,y,I,d){var c=[],a=y-g;return y>g&&y-g>Math.PI/2+.0001?(c=c.concat(this.curveTo(b,e,h,r,g,g+Math.PI/2,I,d)),c=c.concat(this.curveTo(b,e,h,r,g+Math.PI/2,y,I,d))):y<g&&g-y>Math.PI/2+.0001?(c=c.concat(this.curveTo(b,
e,h,r,g,g-Math.PI/2,I,d)),c=c.concat(this.curveTo(b,e,h,r,g-Math.PI/2,y,I,d))):[["C",b+h*Math.cos(g)-h*B*a*Math.sin(g)+I,e+r*Math.sin(g)+r*B*a*Math.cos(g)+d,b+h*Math.cos(y)+h*B*a*Math.sin(y)+I,e+r*Math.sin(y)-r*B*a*Math.cos(y)+d,b+h*Math.cos(y)+I,e+r*Math.sin(y)+d]]};m.prototype.toLinePath=function(b,e){var h=[];b.forEach(function(b){h.push(["L",b.x,b.y])});b.length&&(h[0][0]="M",e&&h.push(["Z"]));return h};m.prototype.toLineSegments=function(b){var e=[],h=!0;b.forEach(function(b){e.push(h?["M",b.x,
b.y]:["L",b.x,b.y]);h=!h});return e};m.prototype.face3d=function(d){var c=this,h=this.createElement("path");h.vertexes=[];h.insidePlotArea=!1;h.enabled=!0;h.attr=function(h){if("object"===typeof h&&(e(h.enabled)||e(h.vertexes)||e(h.insidePlotArea))){this.enabled=q(h.enabled,this.enabled);this.vertexes=q(h.vertexes,this.vertexes);this.insidePlotArea=q(h.insidePlotArea,this.insidePlotArea);delete h.enabled;delete h.vertexes;delete h.insidePlotArea;var g=k(this.vertexes,b[c.chartIndex],this.insidePlotArea),
y=c.toLinePath(g,!0);g=A(g);h.d=y;h.visibility=this.enabled&&0<g?"visible":"hidden"}return f.prototype.attr.apply(this,arguments)};h.animate=function(h){if("object"===typeof h&&(e(h.enabled)||e(h.vertexes)||e(h.insidePlotArea))){this.enabled=q(h.enabled,this.enabled);this.vertexes=q(h.vertexes,this.vertexes);this.insidePlotArea=q(h.insidePlotArea,this.insidePlotArea);delete h.enabled;delete h.vertexes;delete h.insidePlotArea;var g=k(this.vertexes,b[c.chartIndex],this.insidePlotArea),y=c.toLinePath(g,
!0);g=A(g);g=this.enabled&&0<g?"visible":"hidden";h.d=y;this.attr("visibility",g)}return f.prototype.animate.apply(this,arguments)};return h.attr(d)};m.prototype.polyhedron=function(b){var d=this,h=this.g(),r=h.destroy;this.styledMode||h.attr({"stroke-linejoin":"round"});h.faces=[];h.destroy=function(){for(var b=0;b<h.faces.length;b++)h.faces[b].destroy();return r.call(this)};h.attr=function(b,y,r,c){if("object"===typeof b&&e(b.faces)){for(;h.faces.length>b.faces.length;)h.faces.pop().destroy();for(;h.faces.length<
b.faces.length;)h.faces.push(d.face3d().add(h));for(var g=0;g<b.faces.length;g++)d.styledMode&&delete b.faces[g].fill,h.faces[g].attr(b.faces[g],null,r,c);delete b.faces}return f.prototype.attr.apply(this,arguments)};h.animate=function(b,e,r){if(b&&b.faces){for(;h.faces.length>b.faces.length;)h.faces.pop().destroy();for(;h.faces.length<b.faces.length;)h.faces.push(d.face3d().add(h));for(var g=0;g<b.faces.length;g++)h.faces[g].animate(b.faces[g],e,r);delete b.faces}return f.prototype.animate.apply(this,
arguments)};return h.attr(b)};m.prototype.element3d=function(b,e){var h=this.g();n(h,this.elements3d[b]);h.initArgs(e);return h};m.prototype.cuboid=function(b){return this.element3d("cuboid",b)};m.prototype.cuboidPath=function(e){function d(b){return 0===c&&1<b&&6>b?{x:m[b].x,y:m[b].y+10,z:m[b].z}:m[0].x===m[7].x&&4<=b?{x:m[b].x+10,y:m[b].y,z:m[b].z}:0===n&&2>b||5<b?{x:m[b].x,y:m[b].y,z:m[b].z+10}:m[b]}function h(b){return m[b]}var r=e.x||0,g=e.y||0,y=e.z||0,c=e.height||0,a=e.width||0,n=e.depth||
0,f=b[this.chartIndex],q=f.options.chart.options3d.alpha,u=0,m=[{x:r,y:g,z:y},{x:r+a,y:g,z:y},{x:r+a,y:g+c,z:y},{x:r,y:g+c,z:y},{x:r,y:g+c,z:y+n},{x:r+a,y:g+c,z:y+n},{x:r+a,y:g,z:y+n},{x:r,y:g,z:y+n}],C=[];m=k(m,f,e.insidePlotArea);var J=function(b,g,e){var y=[[],-1],r=b.map(h),c=g.map(h);b=b.map(d);g=g.map(d);0>A(r)?y=[r,0]:0>A(c)?y=[c,1]:e&&(C.push(e),y=0>A(b)?[r,0]:0>A(g)?[c,1]:[r,0]);return y};var p=J([3,2,1,0],[7,6,5,4],"front");e=p[0];var G=p[1];p=J([1,6,7,0],[4,5,2,3],"top");a=p[0];var l=p[1];
p=J([1,2,5,6],[0,7,4,3],"side");J=p[0];p=p[1];1===p?u+=1E6*(f.plotWidth-r):p||(u+=1E6*r);u+=10*(!l||0<=q&&180>=q||360>q&&357.5<q?f.plotHeight-g:10+g);1===G?u+=100*y:G||(u+=100*(1E3-y));return{front:this.toLinePath(e,!0),top:this.toLinePath(a,!0),side:this.toLinePath(J,!0),zIndexes:{group:Math.round(u)},forcedSides:C,isFront:G,isTop:l}};m.prototype.arc3d=function(b){function e(b){var h=!1,e={},y;b=u(b);for(y in b)-1!==g.indexOf(y)&&(e[y]=b[y],delete b[y],h=!0);return h?[e,b]:!1}var h=this.g(),r=h.renderer,
g="x y r innerR start end depth".split(" ");b=u(b);b.alpha=(b.alpha||0)*p;b.beta=(b.beta||0)*p;h.top=r.path();h.side1=r.path();h.side2=r.path();h.inn=r.path();h.out=r.path();h.onAdd=function(){var b=h.parentGroup,e=h.attr("class");h.top.add(h);["out","inn","side1","side2"].forEach(function(g){h[g].attr({"class":e+" highcharts-3d-side"}).add(b)})};["addClass","removeClass"].forEach(function(b){h[b]=function(){var e=arguments;["top","out","inn","side1","side2"].forEach(function(g){h[g][b].apply(h[g],
e)})}});h.setPaths=function(b){var e=h.renderer.arc3dPath(b),g=100*e.zTop;h.attribs=b;h.top.attr({d:e.top,zIndex:e.zTop});h.inn.attr({d:e.inn,zIndex:e.zInn});h.out.attr({d:e.out,zIndex:e.zOut});h.side1.attr({d:e.side1,zIndex:e.zSide1});h.side2.attr({d:e.side2,zIndex:e.zSide2});h.zIndex=g;h.attr({zIndex:g});b.center&&(h.top.setRadialReference(b.center),delete b.center)};h.setPaths(b);h.fillSetter=function(b){var e=c(b).brighten(-.1).get();this.fill=b;this.side1.attr({fill:e});this.side2.attr({fill:e});
this.inn.attr({fill:e});this.out.attr({fill:e});this.top.attr({fill:b});return this};["opacity","translateX","translateY","visibility"].forEach(function(b){h[b+"Setter"]=function(b,e){h[e]=b;["out","inn","side1","side2","top"].forEach(function(g){h[g].attr(e,b)})}});h.attr=function(b){var g;if("object"===typeof b&&(g=e(b))){var y=g[0];arguments[0]=g[1];n(h.attribs,y);h.setPaths(h.attribs)}return f.prototype.attr.apply(h,arguments)};h.animate=function(b,g,r){var y=this.attribs,c="data-"+Math.random().toString(26).substring(2,
9);delete b.center;delete b.z;delete b.alpha;delete b.beta;var a=d(q(g,this.renderer.globalAnimation));if(a.duration){g=e(b);h[c]=0;b[c]=1;h[c+"Setter"]=t.noop;if(g){var n=g[0];a.step=function(b,e){function g(b){return y[b]+(q(n[b],y[b])-y[b])*e.pos}e.prop===c&&e.elem.setPaths(u(y,{x:g("x"),y:g("y"),r:g("r"),innerR:g("innerR"),start:g("start"),end:g("end"),depth:g("depth")}))}}g=a}return f.prototype.animate.call(this,b,g,r)};h.destroy=function(){this.top.destroy();this.out.destroy();this.inn.destroy();
this.side1.destroy();this.side2.destroy();return f.prototype.destroy.call(this)};h.hide=function(){this.top.hide();this.out.hide();this.inn.hide();this.side1.hide();this.side2.hide()};h.show=function(b){this.top.show(b);this.out.show(b);this.inn.show(b);this.side1.show(b);this.side2.show(b)};return h};m.prototype.arc3dPath=function(b){function e(b){b%=2*Math.PI;b>Math.PI&&(b=2*Math.PI-b);return b}var h=b.x||0,r=b.y||0,g=b.start||0,c=(b.end||0)-.00001,d=b.r||0,a=b.innerR||0,n=b.depth||0,f=b.alpha||
0,q=b.beta||0,u=Math.cos(g),p=Math.sin(g);b=Math.cos(c);var k=Math.sin(c),l=d*Math.cos(q);d*=Math.cos(f);var A=a*Math.cos(q),v=a*Math.cos(f);a=n*Math.sin(q);var B=n*Math.sin(f);n=[["M",h+l*u,r+d*p]];n=n.concat(m.curveTo(h,r,l,d,g,c,0,0));n.push(["L",h+A*b,r+v*k]);n=n.concat(m.curveTo(h,r,A,v,c,g,0,0));n.push(["Z"]);var z=0<q?Math.PI/2:0;q=0<f?0:Math.PI/2;z=g>-z?g:c>-z?-z:g;var t=c<H-q?c:g<H-q?H-q:c,x=2*H-q;f=[["M",h+l*G(z),r+d*C(z)]];f=f.concat(m.curveTo(h,r,l,d,z,t,0,0));c>x&&g<x?(f.push(["L",h+
l*G(t)+a,r+d*C(t)+B]),f=f.concat(m.curveTo(h,r,l,d,t,x,a,B)),f.push(["L",h+l*G(x),r+d*C(x)]),f=f.concat(m.curveTo(h,r,l,d,x,c,0,0)),f.push(["L",h+l*G(c)+a,r+d*C(c)+B]),f=f.concat(m.curveTo(h,r,l,d,c,x,a,B)),f.push(["L",h+l*G(x),r+d*C(x)]),f=f.concat(m.curveTo(h,r,l,d,x,t,0,0))):c>H-q&&g<H-q&&(f.push(["L",h+l*Math.cos(t)+a,r+d*Math.sin(t)+B]),f=f.concat(m.curveTo(h,r,l,d,t,c,a,B)),f.push(["L",h+l*Math.cos(c),r+d*Math.sin(c)]),f=f.concat(m.curveTo(h,r,l,d,c,t,0,0)));f.push(["L",h+l*Math.cos(t)+a,r+
d*Math.sin(t)+B]);f=f.concat(m.curveTo(h,r,l,d,t,z,a,B));f.push(["Z"]);q=[["M",h+A*u,r+v*p]];q=q.concat(m.curveTo(h,r,A,v,g,c,0,0));q.push(["L",h+A*Math.cos(c)+a,r+v*Math.sin(c)+B]);q=q.concat(m.curveTo(h,r,A,v,c,g,a,B));q.push(["Z"]);u=[["M",h+l*u,r+d*p],["L",h+l*u+a,r+d*p+B],["L",h+A*u+a,r+v*p+B],["L",h+A*u,r+v*p],["Z"]];h=[["M",h+l*b,r+d*k],["L",h+l*b+a,r+d*k+B],["L",h+A*b+a,r+v*k+B],["L",h+A*b,r+v*k],["Z"]];k=Math.atan2(B,-a);r=Math.abs(c+k);b=Math.abs(g+k);g=Math.abs((g+c)/2+k);r=e(r);b=e(b);
g=e(g);g*=1E5;c=1E5*b;r*=1E5;return{top:n,zTop:1E5*Math.PI+1,out:f,zOut:Math.max(g,c,r),inn:q,zInn:Math.max(g,c,r),side1:u,zSide1:.99*r,side2:h,zSide2:.99*c}};return m}(v)});E(a,"Core/Chart/Chart3D.js",[a["Core/Color/Color.js"],a["Extensions/Math3D.js"],a["Core/DefaultOptions.js"],a["Core/Utilities.js"]],function(a,w,t,D){var f=a.parse,z=w.perspective,v=w.shapeArea3D,x=t.defaultOptions,l=D.addEvent,d=D.isArray,c=D.merge,b=D.pick,p=D.wrap,k;(function(a){function e(b){this.is3d()&&"scatter"===b.options.type&&
(b.options.type="scatter3d")}function n(){if(this.chart3d&&this.is3d()){var b=this.renderer,e=this.options.chart.options3d,g=this.chart3d.get3dFrame(),c=this.plotLeft,d=this.plotLeft+this.plotWidth,a=this.plotTop,q=this.plotTop+this.plotHeight;e=e.depth;var n=c-(g.left.visible?g.left.size:0),u=d+(g.right.visible?g.right.size:0),m=a-(g.top.visible?g.top.size:0),k=q+(g.bottom.visible?g.bottom.size:0),p=0-(g.front.visible?g.front.size:0),l=e+(g.back.visible?g.back.size:0),C=this.hasRendered?"animate":
"attr";this.chart3d.frame3d=g;this.frameShapes||(this.frameShapes={bottom:b.polyhedron().add(),top:b.polyhedron().add(),left:b.polyhedron().add(),right:b.polyhedron().add(),back:b.polyhedron().add(),front:b.polyhedron().add()});this.frameShapes.bottom[C]({"class":"highcharts-3d-frame highcharts-3d-frame-bottom",zIndex:g.bottom.frontFacing?-1E3:1E3,faces:[{fill:f(g.bottom.color).brighten(.1).get(),vertexes:[{x:n,y:k,z:p},{x:u,y:k,z:p},{x:u,y:k,z:l},{x:n,y:k,z:l}],enabled:g.bottom.visible},{fill:f(g.bottom.color).brighten(.1).get(),
vertexes:[{x:c,y:q,z:e},{x:d,y:q,z:e},{x:d,y:q,z:0},{x:c,y:q,z:0}],enabled:g.bottom.visible},{fill:f(g.bottom.color).brighten(-.1).get(),vertexes:[{x:n,y:k,z:p},{x:n,y:k,z:l},{x:c,y:q,z:e},{x:c,y:q,z:0}],enabled:g.bottom.visible&&!g.left.visible},{fill:f(g.bottom.color).brighten(-.1).get(),vertexes:[{x:u,y:k,z:l},{x:u,y:k,z:p},{x:d,y:q,z:0},{x:d,y:q,z:e}],enabled:g.bottom.visible&&!g.right.visible},{fill:f(g.bottom.color).get(),vertexes:[{x:u,y:k,z:p},{x:n,y:k,z:p},{x:c,y:q,z:0},{x:d,y:q,z:0}],enabled:g.bottom.visible&&
!g.front.visible},{fill:f(g.bottom.color).get(),vertexes:[{x:n,y:k,z:l},{x:u,y:k,z:l},{x:d,y:q,z:e},{x:c,y:q,z:e}],enabled:g.bottom.visible&&!g.back.visible}]});this.frameShapes.top[C]({"class":"highcharts-3d-frame highcharts-3d-frame-top",zIndex:g.top.frontFacing?-1E3:1E3,faces:[{fill:f(g.top.color).brighten(.1).get(),vertexes:[{x:n,y:m,z:l},{x:u,y:m,z:l},{x:u,y:m,z:p},{x:n,y:m,z:p}],enabled:g.top.visible},{fill:f(g.top.color).brighten(.1).get(),vertexes:[{x:c,y:a,z:0},{x:d,y:a,z:0},{x:d,y:a,z:e},
{x:c,y:a,z:e}],enabled:g.top.visible},{fill:f(g.top.color).brighten(-.1).get(),vertexes:[{x:n,y:m,z:l},{x:n,y:m,z:p},{x:c,y:a,z:0},{x:c,y:a,z:e}],enabled:g.top.visible&&!g.left.visible},{fill:f(g.top.color).brighten(-.1).get(),vertexes:[{x:u,y:m,z:p},{x:u,y:m,z:l},{x:d,y:a,z:e},{x:d,y:a,z:0}],enabled:g.top.visible&&!g.right.visible},{fill:f(g.top.color).get(),vertexes:[{x:n,y:m,z:p},{x:u,y:m,z:p},{x:d,y:a,z:0},{x:c,y:a,z:0}],enabled:g.top.visible&&!g.front.visible},{fill:f(g.top.color).get(),vertexes:[{x:u,
y:m,z:l},{x:n,y:m,z:l},{x:c,y:a,z:e},{x:d,y:a,z:e}],enabled:g.top.visible&&!g.back.visible}]});this.frameShapes.left[C]({"class":"highcharts-3d-frame highcharts-3d-frame-left",zIndex:g.left.frontFacing?-1E3:1E3,faces:[{fill:f(g.left.color).brighten(.1).get(),vertexes:[{x:n,y:k,z:p},{x:c,y:q,z:0},{x:c,y:q,z:e},{x:n,y:k,z:l}],enabled:g.left.visible&&!g.bottom.visible},{fill:f(g.left.color).brighten(.1).get(),vertexes:[{x:n,y:m,z:l},{x:c,y:a,z:e},{x:c,y:a,z:0},{x:n,y:m,z:p}],enabled:g.left.visible&&
!g.top.visible},{fill:f(g.left.color).brighten(-.1).get(),vertexes:[{x:n,y:k,z:l},{x:n,y:m,z:l},{x:n,y:m,z:p},{x:n,y:k,z:p}],enabled:g.left.visible},{fill:f(g.left.color).brighten(-.1).get(),vertexes:[{x:c,y:a,z:e},{x:c,y:q,z:e},{x:c,y:q,z:0},{x:c,y:a,z:0}],enabled:g.left.visible},{fill:f(g.left.color).get(),vertexes:[{x:n,y:k,z:p},{x:n,y:m,z:p},{x:c,y:a,z:0},{x:c,y:q,z:0}],enabled:g.left.visible&&!g.front.visible},{fill:f(g.left.color).get(),vertexes:[{x:n,y:m,z:l},{x:n,y:k,z:l},{x:c,y:q,z:e},{x:c,
y:a,z:e}],enabled:g.left.visible&&!g.back.visible}]});this.frameShapes.right[C]({"class":"highcharts-3d-frame highcharts-3d-frame-right",zIndex:g.right.frontFacing?-1E3:1E3,faces:[{fill:f(g.right.color).brighten(.1).get(),vertexes:[{x:u,y:k,z:l},{x:d,y:q,z:e},{x:d,y:q,z:0},{x:u,y:k,z:p}],enabled:g.right.visible&&!g.bottom.visible},{fill:f(g.right.color).brighten(.1).get(),vertexes:[{x:u,y:m,z:p},{x:d,y:a,z:0},{x:d,y:a,z:e},{x:u,y:m,z:l}],enabled:g.right.visible&&!g.top.visible},{fill:f(g.right.color).brighten(-.1).get(),
vertexes:[{x:d,y:a,z:0},{x:d,y:q,z:0},{x:d,y:q,z:e},{x:d,y:a,z:e}],enabled:g.right.visible},{fill:f(g.right.color).brighten(-.1).get(),vertexes:[{x:u,y:k,z:p},{x:u,y:m,z:p},{x:u,y:m,z:l},{x:u,y:k,z:l}],enabled:g.right.visible},{fill:f(g.right.color).get(),vertexes:[{x:u,y:m,z:p},{x:u,y:k,z:p},{x:d,y:q,z:0},{x:d,y:a,z:0}],enabled:g.right.visible&&!g.front.visible},{fill:f(g.right.color).get(),vertexes:[{x:u,y:k,z:l},{x:u,y:m,z:l},{x:d,y:a,z:e},{x:d,y:q,z:e}],enabled:g.right.visible&&!g.back.visible}]});
this.frameShapes.back[C]({"class":"highcharts-3d-frame highcharts-3d-frame-back",zIndex:g.back.frontFacing?-1E3:1E3,faces:[{fill:f(g.back.color).brighten(.1).get(),vertexes:[{x:u,y:k,z:l},{x:n,y:k,z:l},{x:c,y:q,z:e},{x:d,y:q,z:e}],enabled:g.back.visible&&!g.bottom.visible},{fill:f(g.back.color).brighten(.1).get(),vertexes:[{x:n,y:m,z:l},{x:u,y:m,z:l},{x:d,y:a,z:e},{x:c,y:a,z:e}],enabled:g.back.visible&&!g.top.visible},{fill:f(g.back.color).brighten(-.1).get(),vertexes:[{x:n,y:k,z:l},{x:n,y:m,z:l},
{x:c,y:a,z:e},{x:c,y:q,z:e}],enabled:g.back.visible&&!g.left.visible},{fill:f(g.back.color).brighten(-.1).get(),vertexes:[{x:u,y:m,z:l},{x:u,y:k,z:l},{x:d,y:q,z:e},{x:d,y:a,z:e}],enabled:g.back.visible&&!g.right.visible},{fill:f(g.back.color).get(),vertexes:[{x:c,y:a,z:e},{x:d,y:a,z:e},{x:d,y:q,z:e},{x:c,y:q,z:e}],enabled:g.back.visible},{fill:f(g.back.color).get(),vertexes:[{x:n,y:k,z:l},{x:u,y:k,z:l},{x:u,y:m,z:l},{x:n,y:m,z:l}],enabled:g.back.visible}]});this.frameShapes.front[C]({"class":"highcharts-3d-frame highcharts-3d-frame-front",
zIndex:g.front.frontFacing?-1E3:1E3,faces:[{fill:f(g.front.color).brighten(.1).get(),vertexes:[{x:n,y:k,z:p},{x:u,y:k,z:p},{x:d,y:q,z:0},{x:c,y:q,z:0}],enabled:g.front.visible&&!g.bottom.visible},{fill:f(g.front.color).brighten(.1).get(),vertexes:[{x:u,y:m,z:p},{x:n,y:m,z:p},{x:c,y:a,z:0},{x:d,y:a,z:0}],enabled:g.front.visible&&!g.top.visible},{fill:f(g.front.color).brighten(-.1).get(),vertexes:[{x:n,y:m,z:p},{x:n,y:k,z:p},{x:c,y:q,z:0},{x:c,y:a,z:0}],enabled:g.front.visible&&!g.left.visible},{fill:f(g.front.color).brighten(-.1).get(),
vertexes:[{x:u,y:k,z:p},{x:u,y:m,z:p},{x:d,y:a,z:0},{x:d,y:q,z:0}],enabled:g.front.visible&&!g.right.visible},{fill:f(g.front.color).get(),vertexes:[{x:d,y:a,z:0},{x:c,y:a,z:0},{x:c,y:q,z:0},{x:d,y:q,z:0}],enabled:g.front.visible},{fill:f(g.front.color).get(),vertexes:[{x:u,y:k,z:p},{x:n,y:k,z:p},{x:n,y:m,z:p},{x:u,y:m,z:p}],enabled:g.front.visible}]})}}function u(){this.styledMode&&(this.renderer.definition({tagName:"style",textContent:".highcharts-3d-top{filter: url(#highcharts-brighter)}\n.highcharts-3d-side{filter: url(#highcharts-darker)}\n"}),
[{name:"darker",slope:.6},{name:"brighter",slope:1.4}].forEach(function(b){this.renderer.definition({tagName:"filter",attributes:{id:"highcharts-"+b.name},children:[{tagName:"feComponentTransfer",children:[{tagName:"feFuncR",attributes:{type:"linear",slope:b.slope}},{tagName:"feFuncG",attributes:{type:"linear",slope:b.slope}},{tagName:"feFuncB",attributes:{type:"linear",slope:b.slope}}]}]})},this))}function q(){var b=this.options;this.is3d()&&(b.series||[]).forEach(function(e){"scatter"===(e.type||
b.chart.type||b.chart.defaultSeriesType)&&(e.type="scatter3d")})}function k(){var b=this.options.chart.options3d;if(this.chart3d&&this.is3d()){b&&(b.alpha=b.alpha%360+(0<=b.alpha?0:360),b.beta=b.beta%360+(0<=b.beta?0:360));var e=this.inverted,g=this.clipBox,c=this.margin;g[e?"y":"x"]=-(c[3]||0);g[e?"x":"y"]=-(c[0]||0);g[e?"height":"width"]=this.chartWidth+(c[3]||0)+(c[1]||0);g[e?"width":"height"]=this.chartHeight+(c[0]||0)+(c[2]||0);this.scale3d=1;!0===b.fitToPlot&&(this.scale3d=this.chart3d.getScale(b.depth));
this.chart3d.frame3d=this.chart3d.get3dFrame()}}function C(){this.is3d()&&(this.isDirtyBox=!0)}function A(){this.chart3d&&this.is3d()&&(this.chart3d.frame3d=this.chart3d.get3dFrame())}function B(){this.chart3d||(this.chart3d=new F(this))}function m(b){return this.is3d()||b.apply(this,[].slice.call(arguments,1))}function t(b){var e=this.series.length;if(this.is3d())for(;e--;)b=this.series[e],b.translate(),b.render();else b.call(this)}function w(b){b.apply(this,[].slice.call(arguments,1));this.is3d()&&
(this.container.className+=" highcharts-3d-chart")}var F=function(){function e(b){this.frame3d=void 0;this.chart=b}e.prototype.get3dFrame=function(){var e=this.chart,g=e.options.chart.options3d,c=g.frame,d=e.plotLeft,a=e.plotLeft+e.plotWidth,h=e.plotTop,q=e.plotTop+e.plotHeight,n=g.depth,f=function(b){b=v(b,e);return.5<b?1:-.5>b?-1:0},u=f([{x:d,y:q,z:n},{x:a,y:q,z:n},{x:a,y:q,z:0},{x:d,y:q,z:0}]),m=f([{x:d,y:h,z:0},{x:a,y:h,z:0},{x:a,y:h,z:n},{x:d,y:h,z:n}]),k=f([{x:d,y:h,z:0},{x:d,y:h,z:n},{x:d,
y:q,z:n},{x:d,y:q,z:0}]),p=f([{x:a,y:h,z:n},{x:a,y:h,z:0},{x:a,y:q,z:0},{x:a,y:q,z:n}]),l=f([{x:d,y:q,z:0},{x:a,y:q,z:0},{x:a,y:h,z:0},{x:d,y:h,z:0}]);f=f([{x:d,y:h,z:n},{x:a,y:h,z:n},{x:a,y:q,z:n},{x:d,y:q,z:n}]);var C=!1,G=!1,B=!1,A=!1;[].concat(e.xAxis,e.yAxis,e.zAxis).forEach(function(b){b&&(b.horiz?b.opposite?G=!0:C=!0:b.opposite?A=!0:B=!0)});var t=function(e,c,d){for(var g=["size","color","visible"],a={},q=0;q<g.length;q++)for(var h=g[q],n=0;n<e.length;n++)if("object"===typeof e[n]){var f=e[n][h];
if("undefined"!==typeof f&&null!==f){a[h]=f;break}}e=d;!0===a.visible||!1===a.visible?e=a.visible:"auto"===a.visible&&(e=0<c);return{size:b(a.size,1),color:b(a.color,"none"),frontFacing:0<c,visible:e}};c={axes:{},bottom:t([c.bottom,c.top,c],u,C),top:t([c.top,c.bottom,c],m,G),left:t([c.left,c.right,c.side,c],k,B),right:t([c.right,c.left,c.side,c],p,A),back:t([c.back,c.front,c],f,!0),front:t([c.front,c.back,c],l,!1)};"auto"===g.axisLabelPosition?(p=function(b,e){return b.visible!==e.visible||b.visible&&
e.visible&&b.frontFacing!==e.frontFacing},g=[],p(c.left,c.front)&&g.push({y:(h+q)/2,x:d,z:0,xDir:{x:1,y:0,z:0}}),p(c.left,c.back)&&g.push({y:(h+q)/2,x:d,z:n,xDir:{x:0,y:0,z:-1}}),p(c.right,c.front)&&g.push({y:(h+q)/2,x:a,z:0,xDir:{x:0,y:0,z:1}}),p(c.right,c.back)&&g.push({y:(h+q)/2,x:a,z:n,xDir:{x:-1,y:0,z:0}}),u=[],p(c.bottom,c.front)&&u.push({x:(d+a)/2,y:q,z:0,xDir:{x:1,y:0,z:0}}),p(c.bottom,c.back)&&u.push({x:(d+a)/2,y:q,z:n,xDir:{x:-1,y:0,z:0}}),m=[],p(c.top,c.front)&&m.push({x:(d+a)/2,y:h,z:0,
xDir:{x:1,y:0,z:0}}),p(c.top,c.back)&&m.push({x:(d+a)/2,y:h,z:n,xDir:{x:-1,y:0,z:0}}),k=[],p(c.bottom,c.left)&&k.push({z:(0+n)/2,y:q,x:d,xDir:{x:0,y:0,z:-1}}),p(c.bottom,c.right)&&k.push({z:(0+n)/2,y:q,x:a,xDir:{x:0,y:0,z:1}}),q=[],p(c.top,c.left)&&q.push({z:(0+n)/2,y:h,x:d,xDir:{x:0,y:0,z:-1}}),p(c.top,c.right)&&q.push({z:(0+n)/2,y:h,x:a,xDir:{x:0,y:0,z:1}}),d=function(b,c,d){if(0===b.length)return null;if(1===b.length)return b[0];for(var g=z(b,e,!1),a=0,q=1;q<g.length;q++)d*g[q][c]>d*g[a][c]?a=
q:d*g[q][c]===d*g[a][c]&&g[q].z<g[a].z&&(a=q);return b[a]},c.axes={y:{left:d(g,"x",-1),right:d(g,"x",1)},x:{top:d(m,"y",-1),bottom:d(u,"y",1)},z:{top:d(q,"y",-1),bottom:d(k,"y",1)}}):c.axes={y:{left:{x:d,z:0,xDir:{x:1,y:0,z:0}},right:{x:a,z:0,xDir:{x:0,y:0,z:1}}},x:{top:{y:h,z:0,xDir:{x:1,y:0,z:0}},bottom:{y:q,z:0,xDir:{x:1,y:0,z:0}}},z:{top:{x:B?a:d,y:h,xDir:B?{x:0,y:0,z:1}:{x:0,y:0,z:-1}},bottom:{x:B?a:d,y:q,xDir:B?{x:0,y:0,z:1}:{x:0,y:0,z:-1}}}};return c};e.prototype.getScale=function(b){var e=
this.chart,c=e.plotLeft,d=e.plotWidth+c,a=e.plotTop,q=e.plotHeight+a,h=c+e.plotWidth/2,n=a+e.plotHeight/2,f=Number.MAX_VALUE,u=-Number.MAX_VALUE,m=Number.MAX_VALUE,p=-Number.MAX_VALUE,k=1;var l=[{x:c,y:a,z:0},{x:c,y:a,z:b}];[0,1].forEach(function(b){l.push({x:d,y:l[b].y,z:l[b].z})});[0,1,2,3].forEach(function(b){l.push({x:l[b].x,y:q,z:l[b].z})});l=z(l,e,!1);l.forEach(function(b){f=Math.min(f,b.x);u=Math.max(u,b.x);m=Math.min(m,b.y);p=Math.max(p,b.y)});c>f&&(k=Math.min(k,1-Math.abs((c+h)/(f+h))%1));
d<u&&(k=Math.min(k,(d-h)/(u-h)));a>m&&(k=0>m?Math.min(k,(a+n)/(-m+a+n)):Math.min(k,1-(a+n)/(m+n)%1));q<p&&(k=Math.min(k,Math.abs((q-n)/(p-n))));return k};return e}();a.Composition=F;a.defaultOptions={chart:{options3d:{enabled:!1,alpha:0,beta:0,depth:100,fitToPlot:!0,viewDistance:25,axisLabelPosition:null,frame:{visible:"default",size:1,bottom:{},top:{},left:{},right:{},back:{},front:{}}}}};a.compose=function(b,f){var g=b.prototype;f=f.prototype;g.is3d=function(){return!(!this.options.chart.options3d||
!this.options.chart.options3d.enabled)};g.propsRequireDirtyBox.push("chart.options3d");g.propsRequireUpdateSeries.push("chart.options3d");f.matrixSetter=function(){if(1>this.pos&&(d(this.start)||d(this.end))){var b=this.start||[1,0,0,1,0,0],e=this.end||[1,0,0,1,0,0];var c=[];for(var a=0;6>a;a++)c.push(this.pos*e[a]+(1-this.pos)*b[a])}else c=this.end;this.elem.attr(this.prop,c,null,!0)};c(!0,x,a.defaultOptions);l(b,"init",B);l(b,"addSeries",e);l(b,"afterDrawChartBox",n);l(b,"afterGetContainer",u);
l(b,"afterInit",q);l(b,"afterSetChartSize",k);l(b,"beforeRedraw",C);l(b,"beforeRender",A);p(g,"isInsidePlot",m);p(b,"renderSeries",t);p(b,"setClassName",w)}})(k||(k={}));"";return k});E(a,"Core/Axis/ZAxis.js",[a["Core/Axis/Axis.js"],a["Core/Utilities.js"]],function(a,w){var t=this&&this.__extends||function(){var a=function(c,b){a=Object.setPrototypeOf||{__proto__:[]}instanceof Array&&function(b,c){b.__proto__=c}||function(b,c){for(var a in c)c.hasOwnProperty(a)&&(b[a]=c[a])};return a(c,b)};return function(c,
b){function d(){this.constructor=c}a(c,b);c.prototype=null===b?Object.create(b):(d.prototype=b.prototype,new d)}}(),F=w.addEvent,f=w.merge,z=w.pick,v=w.splat,x=function(){function a(){}a.compose=function(c){F(c,"afterGetAxes",a.onAfterGetAxes);c=c.prototype;c.addZAxis=a.wrapAddZAxis;c.collectionsWithInit.zAxis=[c.addZAxis];c.collectionsWithUpdate.push("zAxis")};a.onAfterGetAxes=function(){var c=this,b=this.options;b=b.zAxis=v(b.zAxis||{});c.is3d()&&(c.zAxis=[],b.forEach(function(b,a){b.index=a;b.isX=
!0;c.addZAxis(b).setScale()}))};a.wrapAddZAxis=function(c){return new l(this,c)};return a}(),l=function(a){function c(b,c){b=a.call(this,b,c)||this;b.isZAxis=!0;return b}t(c,a);c.prototype.getSeriesExtremes=function(){var b=this,c=b.chart;b.hasVisibleSeries=!1;b.dataMin=b.dataMax=b.ignoreMinPadding=b.ignoreMaxPadding=void 0;b.stacking&&b.stacking.buildStacks();b.series.forEach(function(a){if(a.visible||!c.options.chart.ignoreHiddenSeries)b.hasVisibleSeries=!0,a=a.zData,a.length&&(b.dataMin=Math.min(z(b.dataMin,
a[0]),Math.min.apply(null,a)),b.dataMax=Math.max(z(b.dataMax,a[0]),Math.max.apply(null,a)))})};c.prototype.setAxisSize=function(){var b=this.chart;a.prototype.setAxisSize.call(this);this.width=this.len=b.options.chart.options3d&&b.options.chart.options3d.depth||0;this.right=b.chartWidth-this.width-this.left};c.prototype.setOptions=function(b){b=f({offset:0,lineWidth:0},b);this.isZAxis=!0;a.prototype.setOptions.call(this,b);this.coll="zAxis"};c.ZChartComposition=x;return c}(a);return l});E(a,"Core/Axis/Tick3D.js",
[a["Core/Utilities.js"]],function(a){var w=a.addEvent,t=a.extend,F=a.wrap;return function(){function a(){}a.compose=function(f){w(f,"afterGetLabelPosition",a.onAfterGetLabelPosition);F(f.prototype,"getMarkPath",a.wrapGetMarkPath)};a.onAfterGetLabelPosition=function(a){var f=this.axis.axis3D;f&&t(a.pos,f.fix3dPosition(a.pos))};a.wrapGetMarkPath=function(a){var f=this.axis.axis3D,t=a.apply(this,[].slice.call(arguments,1));if(f){var l=t[0],d=t[1];if("M"===l[0]&&"L"===d[0])return f=[f.fix3dPosition({x:l[1],
y:l[2],z:0}),f.fix3dPosition({x:d[1],y:d[2],z:0})],this.axis.chart.renderer.toLineSegments(f)}return t};return a}()});E(a,"Core/Axis/Axis3D.js",[a["Core/Globals.js"],a["Extensions/Math3D.js"],a["Core/Axis/Tick.js"],a["Core/Axis/Tick3D.js"],a["Core/Utilities.js"]],function(a,w,t,D,f){var z=a.deg2rad,v=w.perspective,x=w.perspective3D,l=w.shapeArea,d=f.addEvent,c=f.merge,b=f.pick,p=f.wrap,k=function(){function c(b){this.axis=b}c.prototype.fix3dPosition=function(e,c){var a=this.axis,d=a.chart;if("colorAxis"===
a.coll||!d.chart3d||!d.is3d())return e;var n=z*d.options.chart.options3d.alpha,f=z*d.options.chart.options3d.beta,p=b(c&&a.options.title.position3d,a.options.labels.position3d);c=b(c&&a.options.title.skew3d,a.options.labels.skew3d);var k=d.chart3d.frame3d,m=d.plotLeft,t=d.plotWidth+m,A=d.plotTop,x=d.plotHeight+A,h=d=0,r={x:0,y:1,z:0},g=!1;e=a.axis3D.swapZ({x:e.x,y:e.y,z:0});if(a.isZAxis)if(a.opposite){if(null===k.axes.z.top)return{};h=e.y-A;e.x=k.axes.z.top.x;e.y=k.axes.z.top.y;m=k.axes.z.top.xDir;
g=!k.top.frontFacing}else{if(null===k.axes.z.bottom)return{};h=e.y-x;e.x=k.axes.z.bottom.x;e.y=k.axes.z.bottom.y;m=k.axes.z.bottom.xDir;g=!k.bottom.frontFacing}else if(a.horiz)if(a.opposite){if(null===k.axes.x.top)return{};h=e.y-A;e.y=k.axes.x.top.y;e.z=k.axes.x.top.z;m=k.axes.x.top.xDir;g=!k.top.frontFacing}else{if(null===k.axes.x.bottom)return{};h=e.y-x;e.y=k.axes.x.bottom.y;e.z=k.axes.x.bottom.z;m=k.axes.x.bottom.xDir;g=!k.bottom.frontFacing}else if(a.opposite){if(null===k.axes.y.right)return{};
d=e.x-t;e.x=k.axes.y.right.x;e.z=k.axes.y.right.z;m=k.axes.y.right.xDir;m={x:m.z,y:m.y,z:-m.x}}else{if(null===k.axes.y.left)return{};d=e.x-m;e.x=k.axes.y.left.x;e.z=k.axes.y.left.z;m=k.axes.y.left.xDir}"chart"!==p&&("flap"===p?a.horiz?(f=Math.sin(n),n=Math.cos(n),a.opposite&&(f=-f),g&&(f=-f),r={x:m.z*f,y:n,z:-m.x*f}):m={x:Math.cos(f),y:0,z:Math.sin(f)}:"ortho"===p?a.horiz?(r=Math.cos(n),p=Math.sin(f)*r,n=-Math.sin(n),f=-r*Math.cos(f),r={x:m.y*f-m.z*n,y:m.z*p-m.x*f,z:m.x*n-m.y*p},n=1/Math.sqrt(r.x*
r.x+r.y*r.y+r.z*r.z),g&&(n=-n),r={x:n*r.x,y:n*r.y,z:n*r.z}):m={x:Math.cos(f),y:0,z:Math.sin(f)}:a.horiz?r={x:Math.sin(f)*Math.sin(n),y:Math.cos(n),z:-Math.cos(f)*Math.sin(n)}:m={x:Math.cos(f),y:0,z:Math.sin(f)});e.x+=d*m.x+h*r.x;e.y+=d*m.y+h*r.y;e.z+=d*m.z+h*r.z;d=v([e],a.chart)[0];c&&(0>l(v([e,{x:e.x+m.x,y:e.y+m.y,z:e.z+m.z},{x:e.x+r.x,y:e.y+r.y,z:e.z+r.z}],a.chart))&&(m={x:-m.x,y:-m.y,z:-m.z}),e=v([{x:e.x,y:e.y,z:e.z},{x:e.x+m.x,y:e.y+m.y,z:e.z+m.z},{x:e.x+r.x,y:e.y+r.y,z:e.z+r.z}],a.chart),d.matrix=
[e[1].x-e[0].x,e[1].y-e[0].y,e[2].x-e[0].x,e[2].y-e[0].y,d.x,d.y],d.matrix[4]-=d.x*d.matrix[0]+d.y*d.matrix[2],d.matrix[5]-=d.x*d.matrix[1]+d.y*d.matrix[3]);return d};c.prototype.swapZ=function(b,a){var e=this.axis;return e.isZAxis?(a=a?0:e.chart.plotLeft,{x:a+b.z,y:b.y,z:b.x-a}):b};return c}();return function(){function a(){}a.compose=function(b){c(!0,b.defaultOptions,a.defaultOptions);b.keepProps.push("axis3D");d(b,"init",a.onInit);d(b,"afterSetOptions",a.onAfterSetOptions);d(b,"drawCrosshair",
a.onDrawCrosshair);b=b.prototype;p(b,"getLinePath",a.wrapGetLinePath);p(b,"getPlotBandPath",a.wrapGetPlotBandPath);p(b,"getPlotLinePath",a.wrapGetPlotLinePath);p(b,"getSlotWidth",a.wrapGetSlotWidth);p(b,"getTitlePosition",a.wrapGetTitlePosition);D.compose(t)};a.onAfterSetOptions=function(){var a=this.chart,c=this.options;a.is3d&&a.is3d()&&"colorAxis"!==this.coll&&(c.tickWidth=b(c.tickWidth,0),c.gridLineWidth=b(c.gridLineWidth,1))};a.onDrawCrosshair=function(b){this.chart.is3d()&&"colorAxis"!==this.coll&&
b.point&&(b.point.crosshairPos=this.isXAxis?b.point.axisXpos:this.len-b.point.axisYpos)};a.onInit=function(){this.axis3D||(this.axis3D=new k(this))};a.wrapGetLinePath=function(b){return this.chart.is3d()&&"colorAxis"!==this.coll?[]:b.apply(this,[].slice.call(arguments,1))};a.wrapGetPlotBandPath=function(b){if(!this.chart.is3d()||"colorAxis"===this.coll)return b.apply(this,[].slice.call(arguments,1));var a=arguments,c=a[2],e=[];a=this.getPlotLinePath({value:a[1]});c=this.getPlotLinePath({value:c});
if(a&&c)for(var d=0;d<a.length;d+=2){var f=a[d],k=a[d+1],p=c[d],m=c[d+1];"M"===f[0]&&"L"===k[0]&&"M"===p[0]&&"L"===m[0]&&e.push(f,k,m,["L",p[1],p[2]],["Z"])}return e};a.wrapGetPlotLinePath=function(b){var a=this.axis3D,c=this.chart,e=b.apply(this,[].slice.call(arguments,1));if("colorAxis"===this.coll||!c.chart3d||!c.is3d()||null===e)return e;var d=c.options.chart.options3d,f=this.isZAxis?c.plotWidth:d.depth;d=c.chart3d.frame3d;var k=e[0],p=e[1];e=[];"M"===k[0]&&"L"===p[0]&&(a=[a.swapZ({x:k[1],y:k[2],
z:0}),a.swapZ({x:k[1],y:k[2],z:f}),a.swapZ({x:p[1],y:p[2],z:0}),a.swapZ({x:p[1],y:p[2],z:f})],this.horiz?(this.isZAxis?(d.left.visible&&e.push(a[0],a[2]),d.right.visible&&e.push(a[1],a[3])):(d.front.visible&&e.push(a[0],a[2]),d.back.visible&&e.push(a[1],a[3])),d.top.visible&&e.push(a[0],a[1]),d.bottom.visible&&e.push(a[2],a[3])):(d.front.visible&&e.push(a[0],a[2]),d.back.visible&&e.push(a[1],a[3]),d.left.visible&&e.push(a[0],a[1]),d.right.visible&&e.push(a[2],a[3])),e=v(e,this.chart,!1));return c.renderer.toLineSegments(e)};
a.wrapGetSlotWidth=function(a,c){var e=this.chart,d=this.ticks,f=this.gridGroup;if(this.categories&&e.frameShapes&&e.is3d()&&f&&c&&c.label){f=f.element.childNodes[0].getBBox();var n=e.frameShapes.left.getBBox(),k=e.options.chart.options3d;e={x:e.plotWidth/2,y:e.plotHeight/2,z:k.depth/2,vd:b(k.depth,1)*b(k.viewDistance,0)};k=c.pos;var p=d[k-1],m=d[k+1],l=d=void 0;0!==k&&p&&p.label&&p.label.xy&&(d=x({x:p.label.xy.x,y:p.label.xy.y,z:null},e,e.vd));m&&m.label&&m.label.xy&&(l=x({x:m.label.xy.x,y:m.label.xy.y,
z:null},e,e.vd));k={x:c.label.xy.x,y:c.label.xy.y,z:null};k=x(k,e,e.vd);return Math.abs(d?k.x-d.x:l?l.x-k.x:f.x-n.x)}return a.apply(this,[].slice.call(arguments,1))};a.wrapGetTitlePosition=function(b){var a=b.apply(this,[].slice.call(arguments,1));return this.axis3D?this.axis3D.fix3dPosition(a,!0):a};a.defaultOptions={labels:{position3d:"offset",skew3d:!1},title:{position3d:null,skew3d:null}};return a}()});E(a,"Core/Series/Series3D.js",[a["Extensions/Math3D.js"],a["Core/Series/Series.js"],a["Core/Utilities.js"]],
function(a,w,t){var F=this&&this.__extends||function(){var a=function(c,b){a=Object.setPrototypeOf||{__proto__:[]}instanceof Array&&function(b,a){b.__proto__=a}||function(b,a){for(var c in a)a.hasOwnProperty(c)&&(b[c]=a[c])};return a(c,b)};return function(c,b){function d(){this.constructor=c}a(c,b);c.prototype=null===b?Object.create(b):(d.prototype=b.prototype,new d)}}(),f=a.perspective;a=t.addEvent;var z=t.extend,v=t.merge,x=t.pick,l=t.isNumber;t=function(a){function c(){return null!==a&&a.apply(this,
arguments)||this}F(c,a);c.prototype.translate=function(){a.prototype.translate.apply(this,arguments);this.chart.is3d()&&this.translate3dPoints()};c.prototype.translate3dPoints=function(){var b=this.options,a=this.chart,c=x(this.zAxis,a.options.zAxis[0]),d=[],e,n=[];this.zPadding=(b.stacking?l(b.stack)?b.stack:0:this.index||0)*(b.depth||0+(b.groupZPadding||1));for(e=0;e<this.data.length;e++){b=this.data[e];if(c&&c.translate){var u=c.logarithmic&&c.val2lin?c.val2lin(b.z):b.z;b.plotZ=c.translate(u);
b.isInside=b.isInside?u>=c.min&&u<=c.max:!1}else b.plotZ=this.zPadding;b.axisXpos=b.plotX;b.axisYpos=b.plotY;b.axisZpos=b.plotZ;d.push({x:b.plotX,y:b.plotY,z:b.plotZ});n.push(b.plotX||0)}this.rawPointsX=n;a=f(d,a,!0);for(e=0;e<this.data.length;e++)b=this.data[e],c=a[e],b.plotX=c.x,b.plotY=c.y,b.plotZ=c.z};c.defaultOptions=v(w.defaultOptions);return c}(w);a(w,"afterTranslate",function(){this.chart.is3d()&&this.translate3dPoints()});z(w.prototype,{translate3dPoints:t.prototype.translate3dPoints});return t});
E(a,"Series/Column3D/Column3DComposition.js",[a["Series/Column/ColumnSeries.js"],a["Core/Globals.js"],a["Core/Series/Series.js"],a["Extensions/Math3D.js"],a["Core/Series/SeriesRegistry.js"],a["Extensions/Stacking.js"],a["Core/Utilities.js"]],function(a,w,t,D,f,z,v){function x(b,a){var c=b.series,e={totalStacks:0},d,f=1;c.forEach(function(b){d=A(b.options.stack,a?0:c.length-1-b.index);e[d]?e[d].series.push(b):(e[d]={series:[b],position:f},f++)});e.totalStacks=f+1;return e}function l(b){var a=b.apply(this,
[].slice.call(arguments,1));this.chart.is3d&&this.chart.is3d()&&(a.stroke=this.options.edgeColor||a.fill,a["stroke-width"]=A(this.options.edgeWidth,1));return a}function d(b,a,c){var e=this.chart.is3d&&this.chart.is3d();e&&(this.options.inactiveOtherPoints=!0);b.call(this,a,c);e&&(this.options.inactiveOtherPoints=!1)}function c(b){for(var a=[],c=1;c<arguments.length;c++)a[c-1]=arguments[c];return this.series.chart.is3d()?this.graphic&&"g"!==this.graphic.element.nodeName:b.apply(this,a)}var b=a.prototype,
p=w.svg,k=D.perspective;w=v.addEvent;var A=v.pick;v=v.wrap;v(b,"translate",function(b){b.apply(this,[].slice.call(arguments,1));this.chart.is3d()&&this.translate3dShapes()});v(t.prototype,"justifyDataLabel",function(b){return arguments[2].outside3dPlot?!1:b.apply(this,[].slice.call(arguments,1))});b.translate3dPoints=function(){};b.translate3dShapes=function(){var b=this,a=b.chart,c=b.options,d=c.depth,f=(c.stacking?c.stack||0:b.index)*(d+(c.groupZPadding||1)),p=b.borderWidth%2?.5:0,l;a.inverted&&
!b.yAxis.reversed&&(p*=-1);!1!==c.grouping&&(f=0);f+=c.groupZPadding||1;b.data.forEach(function(c){c.outside3dPlot=null;if(null!==c.y){var e=c.shapeArgs,n=c.tooltipPos,q;[["x","width"],["y","height"]].forEach(function(a){q=e[a[0]]-p;0>q&&(e[a[1]]+=e[a[0]]+p,e[a[0]]=-p,q=0);q+e[a[1]]>b[a[0]+"Axis"].len&&0!==e[a[1]]&&(e[a[1]]=b[a[0]+"Axis"].len-e[a[0]]);if(0!==e[a[1]]&&(e[a[0]]>=b[a[0]+"Axis"].len||e[a[0]]+e[a[1]]<=p)){for(var d in e)e[d]="y"===d?-9999:0;c.outside3dPlot=!0}});"rect"===c.shapeType&&
(c.shapeType="cuboid");e.z=f;e.depth=d;e.insidePlotArea=!0;l={x:e.x+e.width/2,y:e.y,z:f+d/2};a.inverted&&(l.x=e.height,l.y=c.clientX);c.plot3d=k([l],a,!0,!1)[0];n=k([{x:n[0],y:n[1],z:f+d/2}],a,!0,!1)[0];c.tooltipPos=[n.x,n.y]}});b.z=f};v(b,"animate",function(b){if(this.chart.is3d()){var a=arguments[1],c=this.yAxis,e=this,d=this.yAxis.reversed;p&&(a?e.data.forEach(function(b){null!==b.y&&(b.height=b.shapeArgs.height,b.shapey=b.shapeArgs.y,b.shapeArgs.height=1,d||(b.shapeArgs.y=b.stackY?b.plotY+c.translate(b.stackY):
b.plotY+(b.negative?-b.height:b.height)))}):(e.data.forEach(function(b){if(null!==b.y&&(b.shapeArgs.height=b.height,b.shapeArgs.y=b.shapey,b.graphic))b.graphic[b.outside3dPlot?"attr":"animate"](b.shapeArgs,e.options.animation)}),this.drawDataLabels()))}else b.apply(this,[].slice.call(arguments,1))});v(b,"plotGroup",function(b,a,c,d,f,k){"dataLabelsGroup"!==a&&this.chart.is3d()&&(this[a]&&delete this[a],k&&(this.chart.columnGroup||(this.chart.columnGroup=this.chart.renderer.g("columnGroup").add(k)),
this[a]=this.chart.columnGroup,this.chart.columnGroup.attr(this.getPlotBox()),this[a].survive=!0,"group"===a||"markerGroup"===a))&&(arguments[3]="visible");return b.apply(this,Array.prototype.slice.call(arguments,1))});v(b,"setVisible",function(b,a){var c=this;c.chart.is3d()&&c.data.forEach(function(b){b.visible=b.options.visible=a="undefined"===typeof a?!A(c.visible,b.visible):a;c.options.data[c.data.indexOf(b)]=b.options;b.graphic&&b.graphic.attr({visibility:a?"visible":"hidden"})});b.apply(this,
Array.prototype.slice.call(arguments,1))});w(a,"afterInit",function(){if(this.chart.is3d()){var b=this.options,a=b.grouping,c=b.stacking,d=this.yAxis.options.reversedStacks,f=0;if("undefined"===typeof a||a){a=x(this.chart,c);f=b.stack||0;for(c=0;c<a[f].series.length&&a[f].series[c]!==this;c++);f=10*(a.totalStacks-a[f].position)+(d?c:-c);this.xAxis.reversed||(f=10*a.totalStacks-f)}b.depth=b.depth||25;this.z=this.z||0;b.zIndex=f}});v(b,"pointAttribs",l);v(b,"setState",d);v(b.pointClass.prototype,"hasNewShapeType",
c);f.seriesTypes.columnRange&&(w=f.seriesTypes.columnrange.prototype,v(w,"pointAttribs",l),v(w,"setState",d),v(w.pointClass.prototype,"hasNewShapeType",c),w.plotGroup=b.plotGroup,w.setVisible=b.setVisible);v(t.prototype,"alignDataLabel",function(b,a,c,d,f){var e=this.chart;d.outside3dPlot=a.outside3dPlot;if(e.is3d()&&this.is("column")){var p=this.options,l=A(d.inside,!!this.options.stacking),m=e.options.chart.options3d,q=a.pointWidth/2||0;p={x:f.x+q,y:f.y,z:this.z+p.depth/2};e.inverted&&(l&&(f.width=
0,p.x+=a.shapeArgs.height/2),90<=m.alpha&&270>=m.alpha&&(p.y+=a.shapeArgs.width));p=k([p],e,!0,!1)[0];f.x=p.x-q;f.y=a.outside3dPlot?-9E9:p.y}b.apply(this,[].slice.call(arguments,1))});v(z.prototype,"getStackBox",function(b,a,c,d,p,l,t,v){var e=b.apply(this,[].slice.call(arguments,1));if(a.is3d()&&c.base){var q=+c.base.split(",")[0],n=a.series[q];q=a.options.chart.options3d;n&&n instanceof f.seriesTypes.column&&(n={x:e.x+(a.inverted?t:l/2),y:e.y,z:n.options.depth/2},a.inverted&&(e.width=0,90<=q.alpha&&
270>=q.alpha&&(n.y+=l)),n=k([n],a,!0,!1)[0],e.x=n.x-l/2,e.y=n.y)}return e});"";return a});E(a,"Series/Pie3D/Pie3DPoint.js",[a["Core/Series/SeriesRegistry.js"]],function(a){var w=this&&this.__extends||function(){var a=function(f,t){a=Object.setPrototypeOf||{__proto__:[]}instanceof Array&&function(a,f){a.__proto__=f}||function(a,f){for(var l in f)f.hasOwnProperty(l)&&(a[l]=f[l])};return a(f,t)};return function(f,t){function v(){this.constructor=f}a(f,t);f.prototype=null===t?Object.create(t):(v.prototype=
t.prototype,new v)}}();a=a.seriesTypes.pie.prototype.pointClass;var t=a.prototype.haloPath;return function(a){function f(){var f=null!==a&&a.apply(this,arguments)||this;f.series=void 0;return f}w(f,a);f.prototype.haloPath=function(){return this.series.chart.is3d()?[]:t.apply(this,arguments)};return f}(a)});E(a,"Series/Pie3D/Pie3DSeries.js",[a["Core/Globals.js"],a["Series/Pie3D/Pie3DPoint.js"],a["Core/Series/SeriesRegistry.js"],a["Core/Utilities.js"]],function(a,w,t,D){var f=this&&this.__extends||
function(){var a=function(d,c){a=Object.setPrototypeOf||{__proto__:[]}instanceof Array&&function(b,a){b.__proto__=a}||function(b,a){for(var c in a)a.hasOwnProperty(c)&&(b[c]=a[c])};return a(d,c)};return function(d,c){function b(){this.constructor=d}a(d,c);d.prototype=null===c?Object.create(c):(b.prototype=c.prototype,new b)}}(),z=a.deg2rad,v=a.svg;a=D.extend;var x=D.pick;t=function(a){function d(){return null!==a&&a.apply(this,arguments)||this}f(d,a);d.prototype.addPoint=function(){a.prototype.addPoint.apply(this,
arguments);this.chart.is3d()&&this.update(this.userOptions,!0)};d.prototype.animate=function(c){if(this.chart.is3d()){var b=this.options.animation;var d=this.center;var f=this.group,l=this.markerGroup;v&&(!0===b&&(b={}),c?(f.oldtranslateX=x(f.oldtranslateX,f.translateX),f.oldtranslateY=x(f.oldtranslateY,f.translateY),d={translateX:d[0],translateY:d[1],scaleX:.001,scaleY:.001},f.attr(d),l&&(l.attrSetters=f.attrSetters,l.attr(d))):(d={translateX:f.oldtranslateX,translateY:f.oldtranslateY,scaleX:1,scaleY:1},
f.animate(d,b),l&&l.animate(d,b)))}else a.prototype.animate.apply(this,arguments)};d.prototype.drawDataLabels=function(){if(this.chart.is3d()){var c=this.chart.options.chart.options3d;this.data.forEach(function(b){var a=b.shapeArgs,d=a.r,f=(a.start+a.end)/2;b=b.labelPosition;var e=b.connectorPosition,l=-d*(1-Math.cos((a.alpha||c.alpha)*z))*Math.sin(f),t=d*(Math.cos((a.beta||c.beta)*z)-1)*Math.cos(f);[b.natural,e.breakAt,e.touchingSliceAt].forEach(function(b){b.x+=t;b.y+=l})})}a.prototype.drawDataLabels.apply(this,
arguments)};d.prototype.pointAttribs=function(c){var b=a.prototype.pointAttribs.apply(this,arguments),d=this.options;this.chart.is3d()&&!this.chart.styledMode&&(b.stroke=d.edgeColor||c.color||this.color,b["stroke-width"]=x(d.edgeWidth,1));return b};d.prototype.translate=function(){a.prototype.translate.apply(this,arguments);if(this.chart.is3d()){var c=this,b=c.options,d=b.depth||0,f=c.chart.options.chart.options3d,l=f.alpha,e=f.beta,n=b.stacking?(b.stack||0)*d:c._i*d;n+=d/2;!1!==b.grouping&&(n=0);
c.data.forEach(function(a){var f=a.shapeArgs;a.shapeType="arc3d";f.z=n;f.depth=.75*d;f.alpha=l;f.beta=e;f.center=c.center;f=(f.end+f.start)/2;a.slicedTranslation={translateX:Math.round(Math.cos(f)*b.slicedOffset*Math.cos(l*z)),translateY:Math.round(Math.sin(f)*b.slicedOffset*Math.cos(l*z))}})}};return d}(t.seriesTypes.pie);a(t.prototype,{pointClass:w});"";return t});E(a,"Series/Pie3D/Pie3DComposition.js",[a["Series/Pie3D/Pie3DPoint.js"],a["Series/Pie3D/Pie3DSeries.js"],a["Core/Series/SeriesRegistry.js"]],
function(a,w,t){t.seriesTypes.pie.prototype.pointClass.prototype.haloPath=a.prototype.haloPath;t.seriesTypes.pie=w});E(a,"Series/Scatter3D/Scatter3DPoint.js",[a["Series/Scatter/ScatterSeries.js"],a["Core/Utilities.js"]],function(a,w){var t=this&&this.__extends||function(){var a=function(f,t){a=Object.setPrototypeOf||{__proto__:[]}instanceof Array&&function(a,f){a.__proto__=f}||function(a,f){for(var d in f)f.hasOwnProperty(d)&&(a[d]=f[d])};return a(f,t)};return function(f,t){function v(){this.constructor=
f}a(f,t);f.prototype=null===t?Object.create(t):(v.prototype=t.prototype,new v)}}(),F=w.defined;return function(a){function f(){var f=null!==a&&a.apply(this,arguments)||this;f.options=void 0;f.series=void 0;return f}t(f,a);f.prototype.applyOptions=function(){a.prototype.applyOptions.apply(this,arguments);F(this.z)||(this.z=0);return this};return f}(a.prototype.pointClass)});E(a,"Series/Scatter3D/Scatter3DSeries.js",[a["Extensions/Math3D.js"],a["Series/Scatter3D/Scatter3DPoint.js"],a["Series/Scatter/ScatterSeries.js"],
a["Core/Series/SeriesRegistry.js"],a["Core/Utilities.js"]],function(a,w,t,D,f){var z=this&&this.__extends||function(){var a=function(d,c){a=Object.setPrototypeOf||{__proto__:[]}instanceof Array&&function(a,c){a.__proto__=c}||function(a,c){for(var b in c)c.hasOwnProperty(b)&&(a[b]=c[b])};return a(d,c)};return function(d,c){function b(){this.constructor=d}a(d,c);d.prototype=null===c?Object.create(c):(b.prototype=c.prototype,new b)}}(),v=a.pointCameraDistance;a=f.extend;var x=f.merge;f=function(a){function d(){var c=
null!==a&&a.apply(this,arguments)||this;c.data=void 0;c.options=void 0;c.points=void 0;return c}z(d,a);d.prototype.pointAttribs=function(c){var b=a.prototype.pointAttribs.apply(this,arguments);this.chart.is3d()&&c&&(b.zIndex=v(c,this.chart));return b};d.defaultOptions=x(t.defaultOptions,{tooltip:{pointFormat:"x: <b>{point.x}</b><br/>y: <b>{point.y}</b><br/>z: <b>{point.z}</b><br/>"}});return d}(t);a(f.prototype,{axisTypes:["xAxis","yAxis","zAxis"],directTouch:!0,parallelArrays:["x","y","z"],pointArrayMap:["x",
"y","z"],pointClass:w});D.registerSeriesType("scatter3d",f);"";return f});E(a,"Series/Area3DSeries.js",[a["Extensions/Math3D.js"],a["Core/Series/SeriesRegistry.js"],a["Core/Utilities.js"]],function(a,w,t){var F=a.perspective;a=w.seriesTypes;var f=a.line,z=t.pick;t=t.wrap;t(a.area.prototype,"getGraphPath",function(a){var t=a.apply(this,[].slice.call(arguments,1));if(!this.chart.is3d())return t;var l=f.prototype.getGraphPath,d=this.options;var c=[];var b=[],p=z(d.connectNulls,"percent"===d.stacking),
k=Math.round(this.yAxis.getThreshold(d.threshold));if(this.rawPointsX)for(var v=0;v<this.points.length;v++)c.push({x:this.rawPointsX[v],y:d.stacking?this.points[v].yBottom:k,z:this.zPadding});d=this.chart.options.chart.options3d;c=F(c,this.chart,!0).map(function(a){return{plotX:a.x,plotY:a.y,plotZ:a.z}});this.group&&d&&d.depth&&d.beta&&(this.markerGroup&&(this.markerGroup.add(this.group),this.markerGroup.attr({translateX:0,translateY:0})),this.group.attr({zIndex:Math.max(1,270<d.beta||90>d.beta?d.depth-
Math.round(this.zPadding||0):Math.round(this.zPadding||0))}));c.reversed=!0;c=l.call(this,c,!0,!0);c[0]&&"M"===c[0][0]&&(c[0]=["L",c[0][1],c[0][2]]);this.areaPath&&(c=this.areaPath.splice(0,this.areaPath.length/2).concat(c),c.xMap=this.areaPath.xMap,this.areaPath=c,l.call(this,b,!1,p));return t})});E(a,"masters/highcharts-3d.src.js",[a["Core/Globals.js"],a["Core/Renderer/SVG/SVGRenderer3D.js"],a["Core/Chart/Chart3D.js"],a["Core/Axis/ZAxis.js"],a["Core/Axis/Axis3D.js"]],function(a,w,t,D,f){w.compose(a.SVGRenderer);
t.compose(a.Chart,a.Fx);D.ZChartComposition.compose(a.Chart);f.compose(a.Axis)})});
//# sourceMappingURL=highcharts-3d.js.map
```
