## Supplementary material for "Exploring natural odour landscapes: A case study with implications for human-biting insects": Figure S4: highcharts-more.js

```
/*
 Highcharts JS v9.3.1 (2021-11-05)

 (c) 2009-2021 Torstein Honsi

 License: www.highcharts.com/license
*/
'use strict';(function(d){"object"===typeof module&&module.exports?(d["default"]=d,module.exports=d):"function"===typeof define&&define.amd?define("highcharts/highcharts-more",["highcharts"],function(A){d(A);d.Highcharts=A;return d}):d("undefined"!==typeof Highcharts?Highcharts:void 0)})(function(d){function A(d,e,l,a){d.hasOwnProperty(e)||(d[e]=a.apply(null,l))}d=d?d._modules:{};A(d,"Extensions/Pane.js",[d["Core/Chart/Chart.js"],d["Series/CenteredUtilities.js"],d["Core/Globals.js"],d["Core/Pointer.js"],
d["Core/Utilities.js"]],function(d,e,l,a,c){function t(b,m,n){return Math.sqrt(Math.pow(b-n[0],2)+Math.pow(m-n[1],2))<=n[2]/2}var p=c.addEvent,k=c.extend,x=c.merge,w=c.pick,b=c.splat;d.prototype.collectionsWithUpdate.push("pane");c=function(){function g(b,g){this.options=this.chart=this.center=this.background=void 0;this.coll="pane";this.defaultOptions={center:["50%","50%"],size:"85%",innerSize:"0%",startAngle:0};this.defaultBackgroundOptions={shape:"circle",borderWidth:1,borderColor:"#cccccc",backgroundColor:{linearGradient:{x1:0,
y1:0,x2:0,y2:1},stops:[[0,"#ffffff"],[1,"#e6e6e6"]]},from:-Number.MAX_VALUE,innerRadius:0,to:Number.MAX_VALUE,outerRadius:"105%"};this.init(b,g)}g.prototype.init=function(b,g){this.chart=g;this.background=[];g.pane.push(this);this.setOptions(b)};g.prototype.setOptions=function(b){this.options=x(this.defaultOptions,this.chart.angular?{background:{}}:void 0,b)};g.prototype.render=function(){var g=this.options,n=this.options.background,q=this.chart.renderer;this.group||(this.group=q.g("pane-group").attr({zIndex:g.zIndex||
0}).add());this.updateCenter();if(n)for(n=b(n),g=Math.max(n.length,this.background.length||0),q=0;q<g;q++)n[q]&&this.axis?this.renderBackground(x(this.defaultBackgroundOptions,n[q]),q):this.background[q]&&(this.background[q]=this.background[q].destroy(),this.background.splice(q,1))};g.prototype.renderBackground=function(b,g){var n="animate",m={"class":"highcharts-pane "+(b.className||"")};this.chart.styledMode||k(m,{fill:b.backgroundColor,stroke:b.borderColor,"stroke-width":b.borderWidth});this.background[g]||
(this.background[g]=this.chart.renderer.path().add(this.group),n="attr");this.background[g][n]({d:this.axis.getPlotBandPath(b.from,b.to,b)}).attr(m)};g.prototype.updateCenter=function(b){this.center=(b||this.axis||{}).center=e.getCenter.call(this)};g.prototype.update=function(b,g){x(!0,this.options,b);this.setOptions(this.options);this.render();this.chart.axes.forEach(function(b){b.pane===this&&(b.pane=null,b.update({},g))},this)};return g}();d.prototype.getHoverPane=function(b){var g=this,n;b&&g.pane.forEach(function(q){var m=
b.chartX-g.plotLeft,a=b.chartY-g.plotTop;t(g.inverted?a:m,g.inverted?m:a,q.center)&&(n=q)});return n};p(d,"afterIsInsidePlot",function(b){this.polar&&(b.isInsidePlot=this.pane.some(function(g){return t(b.x,b.y,g.center)}))});p(a,"beforeGetHoverData",function(b){var g=this.chart;g.polar?(g.hoverPane=g.getHoverPane(b),b.filter=function(n){return n.visible&&!(!b.shared&&n.directTouch)&&w(n.options.enableMouseTracking,!0)&&(!g.hoverPane||n.xAxis.pane===g.hoverPane)}):g.hoverPane=void 0});p(a,"afterGetHoverData",
function(b){var g=this.chart;b.hoverPoint&&b.hoverPoint.plotX&&b.hoverPoint.plotY&&g.hoverPane&&!t(b.hoverPoint.plotX,b.hoverPoint.plotY,g.hoverPane.center)&&(b.hoverPoint=void 0)});l.Pane=c;return l.Pane});A(d,"Core/Axis/RadialAxis.js",[d["Core/Axis/AxisDefaults.js"],d["Core/DefaultOptions.js"],d["Core/Globals.js"],d["Core/Utilities.js"]],function(d,e,l,a){var c=e.defaultOptions,t=l.noop,p=a.addEvent,k=a.correctFloat,x=a.defined,w=a.extend,b=a.fireEvent,g=a.merge,m=a.pick,n=a.relativeLength,q=a.wrap,
H;(function(a){function e(){this.autoConnect=this.isCircular&&"undefined"===typeof m(this.userMax,this.options.max)&&k(this.endAngleRad-this.startAngleRad)===k(2*Math.PI);!this.isCircular&&this.chart.inverted&&this.max++;this.autoConnect&&(this.max+=this.categories&&1||this.pointRange||this.closestPointRange||0)}function y(){var h=this;return function(){if(h.isRadial&&h.tickPositions&&h.options.labels&&!0!==h.options.labels.allowOverlap)return h.tickPositions.map(function(f){return h.ticks[f]&&h.ticks[f].label}).filter(function(h){return!!h})}}
function z(){return t}function f(h,f,b){var r=this.pane.center,u=h.value;if(this.isCircular){if(x(u))h.point&&(g=h.point.shapeArgs||{},g.start&&(u=this.chart.inverted?this.translate(h.point.rectPlotY,!0):h.point.x));else{var g=h.chartX||0;var v=h.chartY||0;u=this.translate(Math.atan2(v-b,g-f)-this.startAngleRad,!0)}h=this.getPosition(u);g=h.x;v=h.y}else x(u)||(g=h.chartX,v=h.chartY),x(g)&&x(v)&&(b=r[1]+this.chart.plotTop,u=this.translate(Math.min(Math.sqrt(Math.pow(g-f,2)+Math.pow(v-b,2)),r[2]/2)-
r[3]/2,!0));return[u,g||0,v||0]}function h(h,f,b){h=this.pane.center;var r=this.chart,u=this.left||0,g=this.top||0,v=m(f,h[2]/2-this.offset);"undefined"===typeof b&&(b=this.horiz?0:this.center&&-this.center[3]/2);b&&(v+=b);this.isCircular||"undefined"!==typeof f?(f=this.chart.renderer.symbols.arc(u+h[0],g+h[1],v,v,{start:this.startAngleRad,end:this.endAngleRad,open:!0,innerR:0}),f.xBounds=[u+h[0]],f.yBounds=[g+h[1]-v]):(f=this.postTranslate(this.angleRad,v),f=[["M",this.center[0]+r.plotLeft,this.center[1]+
r.plotTop],["L",f.x,f.y]]);return f}function u(){this.constructor.prototype.getOffset.call(this);this.chart.axisOffset[this.side]=0}function r(h,f,b){var r=this.chart,u=function(h){if("string"===typeof h){var f=parseInt(h,10);y.test(h)&&(f=f*B/100);return f}return h},g=this.center,v=this.startAngleRad,B=g[2]/2,n=Math.min(this.offset,0),q=this.left||0,a=this.top||0,y=/%$/,z=this.isCircular,c=m(u(b.outerRadius),B),k=u(b.innerRadius);u=m(u(b.thickness),10);if("polygon"===this.options.gridLineInterpolation)n=
this.getPlotLinePath({value:h}).concat(this.getPlotLinePath({value:f,reverse:!0}));else{h=Math.max(h,this.min);f=Math.min(f,this.max);h=this.translate(h);f=this.translate(f);z||(c=h||0,k=f||0);if("circle"!==b.shape&&z)b=v+(h||0),v+=f||0;else{b=-Math.PI/2;v=1.5*Math.PI;var E=!0}c-=n;n=r.renderer.symbols.arc(q+g[0],a+g[1],c,c,{start:Math.min(b,v),end:Math.max(b,v),innerR:m(k,c-(u-n)),open:E});z&&(z=(v+b)/2,q=q+g[0]+g[2]/2*Math.cos(z),n.xBounds=z>-Math.PI/2&&z<Math.PI/2?[q,r.plotWidth]:[0,q],n.yBounds=
[a+g[1]+g[2]/2*Math.sin(z)],n.yBounds[0]+=z>-Math.PI&&0>z||z>Math.PI?-10:10)}return n}function B(h){var f=this,b=this.pane.center,r=this.chart,u=r.inverted,g=h.reverse,v=this.pane.options.background?this.pane.options.background[0]||this.pane.options.background:{},B=v.innerRadius||"0%",q=v.outerRadius||"100%",a=b[0]+r.plotLeft,z=b[1]+r.plotTop,c=this.height,y=h.isCrosshair;v=b[3]/2;var m=h.value,k;var E=this.getPosition(m);var e=E.x;E=E.y;y&&(E=this.getCrosshairPosition(h,a,z),m=E[0],e=E[1],E=E[2]);
if(this.isCircular)m=Math.sqrt(Math.pow(e-a,2)+Math.pow(E-z,2)),g="string"===typeof B?n(B,1):B/m,r="string"===typeof q?n(q,1):q/m,b&&v&&(v/=m,g<v&&(g=v),r<v&&(r=v)),b=[["M",a+g*(e-a),z-g*(z-E)],["L",e-(1-r)*(e-a),E+(1-r)*(z-E)]];else if((m=this.translate(m))&&(0>m||m>c)&&(m=0),"circle"===this.options.gridLineInterpolation)b=this.getLinePath(0,m,v);else if(b=[],r[u?"yAxis":"xAxis"].forEach(function(h){h.pane===f.pane&&(k=h)}),k)for(a=k.tickPositions,k.autoConnect&&(a=a.concat([a[0]])),g&&(a=a.slice().reverse()),
m&&(m+=v),z=0;z<a.length;z++)v=k.getPosition(a[z],m),b.push(z?["L",v.x,v.y]:["M",v.x,v.y]);return b}function v(h,f){h=this.translate(h);return this.postTranslate(this.isCircular?h:this.angleRad,m(this.isCircular?f:0>h?0:h,this.center[2]/2)-this.offset)}function E(){var h=this.center,f=this.chart,b=this.options.title;return{x:f.plotLeft+h[0]+(b.x||0),y:f.plotTop+h[1]-{high:.5,middle:.25,low:0}[b.align]*h[2]+(b.y||0)}}function l(b){b.beforeSetTickPositions=e;b.createLabelCollector=y;b.getCrosshairPosition=
f;b.getLinePath=h;b.getOffset=u;b.getPlotBandPath=r;b.getPlotLinePath=B;b.getPosition=v;b.getTitlePosition=E;b.postTranslate=O;b.setAxisSize=A;b.setAxisTranslation=P;b.setOptions=Q}function L(){var h=this.chart,f=this.options,b=this.pane,r=b&&b.options;h.angular&&this.isXAxis||!b||!h.angular&&!h.polar||(this.angleRad=(f.angle||0)*Math.PI/180,this.startAngleRad=(r.startAngle-90)*Math.PI/180,this.endAngleRad=(m(r.endAngle,r.startAngle+360)-90)*Math.PI/180,this.offset=f.offset||0)}function H(h){this.isRadial&&
(h.align=void 0,h.preventDefault())}function K(){if(this.chart&&this.chart.labelCollectors){var h=this.labelCollector?this.chart.labelCollectors.indexOf(this.labelCollector):-1;0<=h&&this.chart.labelCollectors.splice(h,1)}}function C(h){var f=this.chart,b=f.inverted,r=f.angular,u=f.polar,v=this.isXAxis,B=this.coll,n=r&&v,a=f.options;h=h.userOptions.pane||0;h=this.pane=f.pane&&f.pane[h];var q;if("colorAxis"===B)this.isRadial=!1;else{if(r){if(n?(this.isHidden=!0,this.createLabelCollector=z,this.getOffset=
t,this.render=this.redraw=G,this.setTitle=this.setCategories=this.setScale=t):l(this),q=!v)this.defaultPolarOptions=R}else u&&(l(this),this.defaultPolarOptions=(q=this.horiz)?S:g("xAxis"===B?d.defaultXAxisOptions:d.defaultYAxisOptions,T),b&&"yAxis"===B&&(this.defaultPolarOptions.stackLabels=d.defaultYAxisOptions.stackLabels,this.defaultPolarOptions.reversedStacks=!0));r||u?(this.isRadial=!0,a.chart.zoomType=null,this.labelCollector||(this.labelCollector=this.createLabelCollector()),this.labelCollector&&
f.labelCollectors.push(this.labelCollector)):this.isRadial=!1;h&&q&&(h.axis=this);this.isCircular=q}}function U(){this.isRadial&&this.beforeSetTickPositions()}function J(h){var f=this.label;if(f){var b=this.axis,r=f.getBBox(),u=b.options.labels,v=(b.translate(this.pos)+b.startAngleRad+Math.PI/2)/Math.PI*180%360,g=Math.round(v),B=x(u.y)?0:.3*-r.height,a=u.y,q=20,z=u.align,c="end",y=0>g?g+360:g,E=y,k=0,e=0;if(b.isRadial){var l=b.getPosition(this.pos,b.center[2]/2+n(m(u.distance,-25),b.center[2]/2,-b.center[2]/
2));"auto"===u.rotation?f.attr({rotation:v}):x(a)||(a=b.chart.renderer.fontMetrics(f.styles&&f.styles.fontSize).b-r.height/2);x(z)||(b.isCircular?(r.width>b.len*b.tickInterval/(b.max-b.min)&&(q=0),z=v>q&&v<180-q?"left":v>180+q&&v<360-q?"right":"center"):z="center",f.attr({align:z}));if("auto"===z&&2===b.tickPositions.length&&b.isCircular){90<y&&180>y?y=180-y:270<y&&360>=y&&(y=540-y);180<E&&360>=E&&(E=360-E);if(b.pane.options.startAngle===g||b.pane.options.startAngle===g+360||b.pane.options.startAngle===
g-360)c="start";z=-90<=g&&90>=g||-360<=g&&-270>=g||270<=g&&360>=g?"start"===c?"right":"left":"start"===c?"left":"right";70<E&&110>E&&(z="center");15>y||180<=y&&195>y?k=.3*r.height:15<=y&&35>=y?k="start"===c?0:.75*r.height:195<=y&&215>=y?k="start"===c?.75*r.height:0:35<y&&90>=y?k="start"===c?.25*-r.height:r.height:215<y&&270>=y&&(k="start"===c?r.height:.25*-r.height);15>E?e="start"===c?.15*-r.height:.15*r.height:165<E&&180>=E&&(e="start"===c?.15*r.height:.15*-r.height);f.attr({align:z});f.translate(e,
k+B)}h.pos.x=l.x+(u.x||0);h.pos.y=l.y+(a||0)}}}function V(h){this.axis.getPosition&&w(h.pos,this.axis.getPosition(this.pos))}function O(h,f){var b=this.chart,r=this.center;h=this.startAngleRad+h;return{x:b.plotLeft+r[0]+Math.cos(h)*f,y:b.plotTop+r[1]+Math.sin(h)*f}}function G(){this.isDirty=!1}function A(){this.constructor.prototype.setAxisSize.call(this);if(this.isRadial){this.pane.updateCenter(this);var h=this.center=this.pane.center.slice();if(this.isCircular)this.sector=this.endAngleRad-this.startAngleRad;
else{var f=this.postTranslate(this.angleRad,h[3]/2);h[0]=f.x-this.chart.plotLeft;h[1]=f.y-this.chart.plotTop}this.len=this.width=this.height=(h[2]-h[3])*m(this.sector,1)/2}}function P(){this.constructor.prototype.setAxisTranslation.call(this);this.center&&(this.transA=this.isCircular?(this.endAngleRad-this.startAngleRad)/(this.max-this.min||1):(this.center[2]-this.center[3])/2/(this.max-this.min||1),this.minPixelPadding=this.isXAxis?this.transA*this.minPointOffset:0)}function Q(h){h=this.options=
g(this.constructor.defaultOptions,this.defaultPolarOptions,c[this.coll],h);h.plotBands||(h.plotBands=[]);b(this,"afterSetOptions")}function W(h,f,b,r,u,g,v){var B=this.axis;B.isRadial?(h=B.getPosition(this.pos,B.center[2]/2+r),f=["M",f,b,"L",h.x,h.y]):f=h.call(this,f,b,r,u,g,v);return f}var N=[],S={gridLineWidth:1,labels:{align:void 0,distance:15,x:0,y:void 0,style:{textOverflow:"none"}},maxPadding:0,minPadding:0,showLastLabel:!1,tickLength:0},R={labels:{align:"center",x:0,y:void 0},minorGridLineWidth:0,
minorTickInterval:"auto",minorTickLength:10,minorTickPosition:"inside",minorTickWidth:1,tickLength:10,tickPosition:"inside",tickWidth:2,title:{rotation:0},zIndex:2},T={gridLineInterpolation:"circle",gridLineWidth:1,labels:{align:"right",x:-3,y:-2},showLastLabel:!1,title:{x:4,text:null,rotation:90}};a.compose=function(h,f){-1===N.indexOf(h)&&(N.push(h),p(h,"afterInit",L),p(h,"autoLabelAlign",H),p(h,"destroy",K),p(h,"init",C),p(h,"initialAxisTranslation",U));-1===N.indexOf(f)&&(N.push(f),p(f,"afterGetLabelPosition",
J),p(f,"afterGetPosition",V),q(f.prototype,"getMarkPath",W));return h}})(H||(H={}));return H});A(d,"Series/AreaRange/AreaRangePoint.js",[d["Series/Area/AreaSeries.js"],d["Core/Series/Point.js"],d["Core/Utilities.js"]],function(d,e,l){var a=this&&this.__extends||function(){var a=function(c,k){a=Object.setPrototypeOf||{__proto__:[]}instanceof Array&&function(b,g){b.__proto__=g}||function(b,g){for(var a in g)g.hasOwnProperty(a)&&(b[a]=g[a])};return a(c,k)};return function(c,k){function b(){this.constructor=
c}a(c,k);c.prototype=null===k?Object.create(k):(b.prototype=k.prototype,new b)}}(),c=e.prototype,t=l.defined,p=l.isNumber;return function(k){function e(){var a=null!==k&&k.apply(this,arguments)||this;a.high=void 0;a.low=void 0;a.options=void 0;a.plotHigh=void 0;a.plotLow=void 0;a.plotHighX=void 0;a.plotLowX=void 0;a.plotX=void 0;a.series=void 0;return a}a(e,k);e.prototype.setState=function(){var a=this.state,b=this.series,g=b.chart.polar;t(this.plotHigh)||(this.plotHigh=b.yAxis.toPixels(this.high,
!0));t(this.plotLow)||(this.plotLow=this.plotY=b.yAxis.toPixels(this.low,!0));b.stateMarkerGraphic&&(b.lowerStateMarkerGraphic=b.stateMarkerGraphic,b.stateMarkerGraphic=b.upperStateMarkerGraphic);this.graphic=this.upperGraphic;this.plotY=this.plotHigh;g&&(this.plotX=this.plotHighX);c.setState.apply(this,arguments);this.state=a;this.plotY=this.plotLow;this.graphic=this.lowerGraphic;g&&(this.plotX=this.plotLowX);b.stateMarkerGraphic&&(b.upperStateMarkerGraphic=b.stateMarkerGraphic,b.stateMarkerGraphic=
b.lowerStateMarkerGraphic,b.lowerStateMarkerGraphic=void 0);c.setState.apply(this,arguments)};e.prototype.haloPath=function(){var a=this.series.chart.polar,b=[];this.plotY=this.plotLow;a&&(this.plotX=this.plotLowX);this.isInside&&(b=c.haloPath.apply(this,arguments));this.plotY=this.plotHigh;a&&(this.plotX=this.plotHighX);this.isTopInside&&(b=b.concat(c.haloPath.apply(this,arguments)));return b};e.prototype.isValid=function(){return p(this.low)&&p(this.high)};return e}(d.prototype.pointClass)});A(d,
"Series/AreaRange/AreaRangeSeries.js",[d["Series/AreaRange/AreaRangePoint.js"],d["Series/Area/AreaSeries.js"],d["Series/Column/ColumnSeries.js"],d["Core/Globals.js"],d["Core/Series/Series.js"],d["Core/Series/SeriesRegistry.js"],d["Core/Utilities.js"]],function(d,e,l,a,c,t,p){var k=this&&this.__extends||function(){var b=function(g,a){b=Object.setPrototypeOf||{__proto__:[]}instanceof Array&&function(b,f){b.__proto__=f}||function(b,f){for(var h in f)f.hasOwnProperty(h)&&(b[h]=f[h])};return b(g,a)};return function(g,
a){function n(){this.constructor=g}b(g,a);g.prototype=null===a?Object.create(a):(n.prototype=a.prototype,new n)}}(),x=e.prototype,w=l.prototype;l=a.noop;var b=c.prototype,g=p.defined,m=p.extend,n=p.isArray,q=p.pick,H=p.merge;c=function(a){function c(){var b=null!==a&&a.apply(this,arguments)||this;b.data=void 0;b.options=void 0;b.points=void 0;b.lowerStateMarkerGraphic=void 0;b.xAxis=void 0;return b}k(c,a);c.prototype.toYData=function(b){return[b.low,b.high]};c.prototype.highToXY=function(b){var g=
this.chart,f=this.xAxis.postTranslate(b.rectPlotX||0,this.yAxis.len-b.plotHigh);b.plotHighX=f.x-g.plotLeft;b.plotHigh=f.y-g.plotTop;b.plotLowX=b.plotX};c.prototype.translate=function(){var b=this,g=b.yAxis;x.translate.apply(b);b.points.forEach(function(f){var h=f.high,u=f.plotY;f.isNull?f.plotY=null:(f.plotLow=u,f.plotHigh=g.translate(b.dataModify?b.dataModify.modifyValue(h):h,0,1,0,1),b.dataModify&&(f.yBottom=f.plotHigh))});this.chart.polar&&this.points.forEach(function(f){b.highToXY(f);f.tooltipPos=
[(f.plotHighX+f.plotLowX)/2,(f.plotHigh+f.plotLow)/2]})};c.prototype.getGraphPath=function(b){var g=[],f=[],h,u=x.getGraphPath;var r=this.options;var a=this.chart.polar,v=a&&!1!==r.connectEnds,n=r.connectNulls,c=r.step;b=b||this.points;for(h=b.length;h--;){var m=b[h];var k=a?{plotX:m.rectPlotX,plotY:m.yBottom,doCurve:!1}:{plotX:m.plotX,plotY:m.plotY,doCurve:!1};m.isNull||v||n||b[h+1]&&!b[h+1].isNull||f.push(k);var e={polarPlotY:m.polarPlotY,rectPlotX:m.rectPlotX,yBottom:m.yBottom,plotX:q(m.plotHighX,
m.plotX),plotY:m.plotHigh,isNull:m.isNull};f.push(e);g.push(e);m.isNull||v||n||b[h-1]&&!b[h-1].isNull||f.push(k)}b=u.call(this,b);c&&(!0===c&&(c="left"),r.step={left:"right",center:"center",right:"left"}[c]);g=u.call(this,g);f=u.call(this,f);r.step=c;r=[].concat(b,g);!this.chart.polar&&f[0]&&"M"===f[0][0]&&(f[0]=["L",f[0][1],f[0][2]]);this.graphPath=r;this.areaPath=b.concat(f);r.isArea=!0;r.xMap=b.xMap;this.areaPath.xMap=b.xMap;return r};c.prototype.drawDataLabels=function(){var g=this.points,a=g.length,
f,h=[],u=this.options.dataLabels,r,B=this.chart.inverted;if(u){if(n(u)){var v=u[0]||{enabled:!1};var c=u[1]||{enabled:!1}}else v=m({},u),v.x=u.xHigh,v.y=u.yHigh,c=m({},u),c.x=u.xLow,c.y=u.yLow;if(v.enabled||this._hasPointLabels){for(f=a;f--;)if(r=g[f]){var q=v.inside?r.plotHigh<r.plotLow:r.plotHigh>r.plotLow;r.y=r.high;r._plotY=r.plotY;r.plotY=r.plotHigh;h[f]=r.dataLabel;r.dataLabel=r.dataLabelUpper;r.below=q;B?v.align||(v.align=q?"right":"left"):v.verticalAlign||(v.verticalAlign=q?"top":"bottom")}this.options.dataLabels=
v;b.drawDataLabels&&b.drawDataLabels.apply(this,arguments);for(f=a;f--;)if(r=g[f])r.dataLabelUpper=r.dataLabel,r.dataLabel=h[f],delete r.dataLabels,r.y=r.low,r.plotY=r._plotY}if(c.enabled||this._hasPointLabels){for(f=a;f--;)if(r=g[f])q=c.inside?r.plotHigh<r.plotLow:r.plotHigh>r.plotLow,r.below=!q,B?c.align||(c.align=q?"left":"right"):c.verticalAlign||(c.verticalAlign=q?"bottom":"top");this.options.dataLabels=c;b.drawDataLabels&&b.drawDataLabels.apply(this,arguments)}if(v.enabled)for(f=a;f--;)if(r=
g[f])r.dataLabels=[r.dataLabelUpper,r.dataLabel].filter(function(h){return!!h});this.options.dataLabels=u}};c.prototype.alignDataLabel=function(){w.alignDataLabel.apply(this,arguments)};c.prototype.drawPoints=function(){var a=this.points.length,c;b.drawPoints.apply(this,arguments);for(c=0;c<a;){var f=this.points[c];f.origProps={plotY:f.plotY,plotX:f.plotX,isInside:f.isInside,negative:f.negative,zone:f.zone,y:f.y};f.lowerGraphic=f.graphic;f.graphic=f.upperGraphic;f.plotY=f.plotHigh;g(f.plotHighX)&&
(f.plotX=f.plotHighX);f.y=q(f.high,f.origProps.y);f.negative=f.y<(this.options.threshold||0);this.zones.length&&(f.zone=f.getZone());this.chart.polar||(f.isInside=f.isTopInside="undefined"!==typeof f.plotY&&0<=f.plotY&&f.plotY<=this.yAxis.len&&0<=f.plotX&&f.plotX<=this.xAxis.len);c++}b.drawPoints.apply(this,arguments);for(c=0;c<a;)f=this.points[c],f.upperGraphic=f.graphic,f.graphic=f.lowerGraphic,f.origProps&&(m(f,f.origProps),delete f.origProps),c++};c.defaultOptions=H(e.defaultOptions,{lineWidth:1,
threshold:null,tooltip:{pointFormat:'<span style="color:{series.color}">\u25cf</span> {series.name}: <b>{point.low}</b> - <b>{point.high}</b><br/>'},trackByArea:!0,dataLabels:{align:void 0,verticalAlign:void 0,xLow:0,xHigh:0,yLow:0,yHigh:0}});return c}(e);m(c.prototype,{pointArrayMap:["low","high"],pointValKey:"low",deferTranslatePolar:!0,pointClass:d,setStackedPoints:l});t.registerSeriesType("arearange",c);"";return c});A(d,"Series/AreaSplineRange/AreaSplineRangeSeries.js",[d["Series/AreaRange/AreaRangeSeries.js"],
d["Core/Series/SeriesRegistry.js"],d["Core/Utilities.js"]],function(d,e,l){var a=this&&this.__extends||function(){var a=function(c,k){a=Object.setPrototypeOf||{__proto__:[]}instanceof Array&&function(b,g){b.__proto__=g}||function(b,g){for(var a in g)g.hasOwnProperty(a)&&(b[a]=g[a])};return a(c,k)};return function(c,k){function b(){this.constructor=c}a(c,k);c.prototype=null===k?Object.create(k):(b.prototype=k.prototype,new b)}}(),c=e.seriesTypes.spline,t=l.merge;l=l.extend;var p=function(c){function k(){var a=
null!==c&&c.apply(this,arguments)||this;a.options=void 0;a.data=void 0;a.points=void 0;return a}a(k,c);k.defaultOptions=t(d.defaultOptions);return k}(d);l(p.prototype,{getPointSpline:c.prototype.getPointSpline});e.registerSeriesType("areasplinerange",p);"";return p});A(d,"Series/BoxPlot/BoxPlotSeries.js",[d["Series/Column/ColumnSeries.js"],d["Core/Globals.js"],d["Core/Series/SeriesRegistry.js"],d["Core/Utilities.js"]],function(d,e,l,a){var c=this&&this.__extends||function(){var a=function(c,b){a=
Object.setPrototypeOf||{__proto__:[]}instanceof Array&&function(b,a){b.__proto__=a}||function(b,a){for(var g in a)a.hasOwnProperty(g)&&(b[g]=a[g])};return a(c,b)};return function(c,b){function g(){this.constructor=c}a(c,b);c.prototype=null===b?Object.create(b):(g.prototype=b.prototype,new g)}}();e=e.noop;var t=a.extend,p=a.merge,k=a.pick;a=function(a){function e(){var b=null!==a&&a.apply(this,arguments)||this;b.data=void 0;b.options=void 0;b.points=void 0;return b}c(e,a);e.prototype.pointAttribs=
function(){return{}};e.prototype.translate=function(){var b=this.yAxis,g=this.pointArrayMap;a.prototype.translate.apply(this);this.points.forEach(function(a){g.forEach(function(g){null!==a[g]&&(a[g+"Plot"]=b.translate(a[g],0,1,0,1))});a.plotHigh=a.highPlot})};e.prototype.drawPoints=function(){var b=this,g=b.options,a=b.chart,c=a.renderer,q,e,l,p,y,z,f=0,h,u,r,B,v=!1!==b.doQuartiles,E,d=b.options.whiskerLength;b.points.forEach(function(n){var m=n.graphic,H=m?"animate":"attr",t=n.shapeArgs,x={},I={},
w={},L={},M=n.color||b.color;"undefined"!==typeof n.plotY&&(h=Math.round(t.width),u=Math.floor(t.x),r=u+h,B=Math.round(h/2),q=Math.floor(v?n.q1Plot:n.lowPlot),e=Math.floor(v?n.q3Plot:n.lowPlot),l=Math.floor(n.highPlot),p=Math.floor(n.lowPlot),m||(n.graphic=m=c.g("point").add(b.group),n.stem=c.path().addClass("highcharts-boxplot-stem").add(m),d&&(n.whiskers=c.path().addClass("highcharts-boxplot-whisker").add(m)),v&&(n.box=c.path(void 0).addClass("highcharts-boxplot-box").add(m)),n.medianShape=c.path(void 0).addClass("highcharts-boxplot-median").add(m)),
a.styledMode||(I.stroke=n.stemColor||g.stemColor||M,I["stroke-width"]=k(n.stemWidth,g.stemWidth,g.lineWidth),I.dashstyle=n.stemDashStyle||g.stemDashStyle||g.dashStyle,n.stem.attr(I),d&&(w.stroke=n.whiskerColor||g.whiskerColor||M,w["stroke-width"]=k(n.whiskerWidth,g.whiskerWidth,g.lineWidth),w.dashstyle=n.whiskerDashStyle||g.whiskerDashStyle||g.dashStyle,n.whiskers.attr(w)),v&&(x.fill=n.fillColor||g.fillColor||M,x.stroke=g.lineColor||M,x["stroke-width"]=g.lineWidth||0,x.dashstyle=n.boxDashStyle||g.boxDashStyle||
g.dashStyle,n.box.attr(x)),L.stroke=n.medianColor||g.medianColor||M,L["stroke-width"]=k(n.medianWidth,g.medianWidth,g.lineWidth),L.dashstyle=n.medianDashStyle||g.medianDashStyle||g.dashStyle,n.medianShape.attr(L)),z=n.stem.strokeWidth()%2/2,f=u+B+z,m=[["M",f,e],["L",f,l],["M",f,q],["L",f,p]],n.stem[H]({d:m}),v&&(z=n.box.strokeWidth()%2/2,q=Math.floor(q)+z,e=Math.floor(e)+z,u+=z,r+=z,m=[["M",u,e],["L",u,q],["L",r,q],["L",r,e],["L",u,e],["Z"]],n.box[H]({d:m})),d&&(z=n.whiskers.strokeWidth()%2/2,l+=
z,p+=z,E=/%$/.test(d)?B*parseFloat(d)/100:d/2,m=[["M",f-E,l],["L",f+E,l],["M",f-E,p],["L",f+E,p]],n.whiskers[H]({d:m})),y=Math.round(n.medianPlot),z=n.medianShape.strokeWidth()%2/2,y+=z,m=[["M",u,y],["L",r,y]],n.medianShape[H]({d:m}))})};e.prototype.toYData=function(b){return[b.low,b.q1,b.median,b.q3,b.high]};e.defaultOptions=p(d.defaultOptions,{threshold:null,tooltip:{pointFormat:'<span style="color:{point.color}">\u25cf</span> <b> {series.name}</b><br/>Maximum: {point.high}<br/>Upper quartile: {point.q3}<br/>Median: {point.median}<br/>Lower quartile: {point.q1}<br/>Minimum: {point.low}<br/>'},
whiskerLength:"50%",fillColor:"#ffffff",lineWidth:1,medianWidth:2,whiskerWidth:2});return e}(d);t(a.prototype,{pointArrayMap:["low","q1","median","q3","high"],pointValKey:"high",drawDataLabels:e,setStackedPoints:e});l.registerSeriesType("boxplot",a);"";return a});A(d,"Series/Bubble/BubbleLegendDefaults.js",[],function(){return{borderColor:void 0,borderWidth:2,className:void 0,color:void 0,connectorClassName:void 0,connectorColor:void 0,connectorDistance:60,connectorWidth:1,enabled:!1,labels:{className:void 0,
allowOverlap:!1,format:"",formatter:void 0,align:"right",style:{fontSize:"10px",color:"#000000"},x:0,y:0},maxSize:60,minSize:10,legendIndex:0,ranges:{value:void 0,borderColor:void 0,color:void 0,connectorColor:void 0},sizeBy:"area",sizeByAbsoluteValue:!1,zIndex:1,zThreshold:0}});A(d,"Series/Bubble/BubbleLegendItem.js",[d["Core/Color/Color.js"],d["Core/FormatUtilities.js"],d["Core/Globals.js"],d["Core/Utilities.js"]],function(d,e,l,a){var c=d.parse,t=l.noop,p=a.arrayMax,k=a.arrayMin,x=a.isNumber,w=
a.merge,b=a.pick,g=a.stableSort;"";return function(){function a(b,a){this.options=this.symbols=this.visible=this.selected=this.ranges=this.movementX=this.maxLabel=this.legendSymbol=this.legendItemWidth=this.legendItemHeight=this.legendItem=this.legendGroup=this.legend=this.fontMetrics=this.chart=void 0;this.setState=t;this.init(b,a)}a.prototype.init=function(b,a){this.options=b;this.visible=!0;this.chart=a.chart;this.legend=a};a.prototype.addToLegend=function(b){b.splice(this.options.legendIndex,
0,this)};a.prototype.drawLegendSymbol=function(a){var c=this.chart,n=this.options,e=b(a.options.itemDistance,20),k=n.ranges,m=n.connectorDistance;this.fontMetrics=c.renderer.fontMetrics(n.labels.style.fontSize);k&&k.length&&x(k[0].value)?(g(k,function(b,f){return f.value-b.value}),this.ranges=k,this.setOptions(),this.render(),a=this.getMaxLabelSize(),k=this.ranges[0].radius,c=2*k,m=m-k+a.width,m=0<m?m:0,this.maxLabel=a,this.movementX="left"===n.labels.align?m:0,this.legendItemWidth=c+m+e,this.legendItemHeight=
c+this.fontMetrics.h/2):a.options.bubbleLegend.autoRanges=!0};a.prototype.setOptions=function(){var a=this.ranges,g=this.options,k=this.chart.series[g.seriesIndex],e=this.legend.baseline,m={zIndex:g.zIndex,"stroke-width":g.borderWidth},l={zIndex:g.zIndex,"stroke-width":g.connectorWidth},z={align:this.legend.options.rtl||"left"===g.labels.align?"right":"left",zIndex:g.zIndex},f=k.options.marker.fillOpacity,h=this.chart.styledMode;a.forEach(function(u,r){h||(m.stroke=b(u.borderColor,g.borderColor,k.color),
m.fill=b(u.color,g.color,1!==f?c(k.color).setOpacity(f).get("rgba"):k.color),l.stroke=b(u.connectorColor,g.connectorColor,k.color));a[r].radius=this.getRangeRadius(u.value);a[r]=w(a[r],{center:a[0].radius-a[r].radius+e});h||w(!0,a[r],{bubbleAttribs:w(m),connectorAttribs:w(l),labelAttribs:z})},this)};a.prototype.getRangeRadius=function(b){var a=this.options;return this.chart.series[this.options.seriesIndex].getRadius.call(this,a.ranges[a.ranges.length-1].value,a.ranges[0].value,a.minSize,a.maxSize,
b)};a.prototype.render=function(){var b=this.chart.renderer,a=this.options.zThreshold;this.symbols||(this.symbols={connectors:[],bubbleItems:[],labels:[]});this.legendSymbol=b.g("bubble-legend");this.legendItem=b.g("bubble-legend-item");this.legendSymbol.translateX=0;this.legendSymbol.translateY=0;this.ranges.forEach(function(b){b.value>=a&&this.renderRange(b)},this);this.legendSymbol.add(this.legendItem);this.legendItem.add(this.legendGroup);this.hideOverlappingLabels()};a.prototype.renderRange=
function(b){var a=this.options,g=a.labels,c=this.chart,n=c.series[a.seriesIndex],k=c.renderer,e=this.symbols;c=e.labels;var f=b.center,h=Math.abs(b.radius),u=a.connectorDistance||0,r=g.align,B=a.connectorWidth,v=this.ranges[0].radius||0,m=f-h-a.borderWidth/2+B/2,l=this.fontMetrics;l=l.f/2-(l.h-l.f)/2;var d=k.styledMode;u=this.legend.options.rtl||"left"===r?-u:u;"center"===r&&(u=0,a.connectorDistance=0,b.labelAttribs.align="center");r=m+a.labels.y;var p=v+u+a.labels.x;e.bubbleItems.push(k.circle(v,
f+((m%1?1:.5)-(B%2?0:.5)),h).attr(d?{}:b.bubbleAttribs).addClass((d?"highcharts-color-"+n.colorIndex+" ":"")+"highcharts-bubble-legend-symbol "+(a.className||"")).add(this.legendSymbol));e.connectors.push(k.path(k.crispLine([["M",v,m],["L",v+u,m]],a.connectorWidth)).attr(d?{}:b.connectorAttribs).addClass((d?"highcharts-color-"+this.options.seriesIndex+" ":"")+"highcharts-bubble-legend-connectors "+(a.connectorClassName||"")).add(this.legendSymbol));b=k.text(this.formatLabel(b),p,r+l).attr(d?{}:b.labelAttribs).css(d?
{}:g.style).addClass("highcharts-bubble-legend-labels "+(a.labels.className||"")).add(this.legendSymbol);c.push(b);b.placed=!0;b.alignAttr={x:p,y:r+l}};a.prototype.getMaxLabelSize=function(){var b,a;this.symbols.labels.forEach(function(g){a=g.getBBox(!0);b=b?a.width>b.width?a:b:a});return b||{}};a.prototype.formatLabel=function(b){var a=this.options,g=a.labels.formatter;a=a.labels.format;var c=this.chart.numberFormatter;return a?e.format(a,b):g?g.call(b):c(b.value,1)};a.prototype.hideOverlappingLabels=
function(){var b=this.chart,a=this.symbols;!this.options.labels.allowOverlap&&a&&(b.hideOverlappingLabels(a.labels),a.labels.forEach(function(b,g){b.newOpacity?b.newOpacity!==b.oldOpacity&&a.connectors[g].show():a.connectors[g].hide()}))};a.prototype.getRanges=function(){var a=this.legend.bubbleLegend,g=a.options.ranges,c,e=Number.MAX_VALUE,m=-Number.MAX_VALUE;a.chart.series.forEach(function(a){a.isBubble&&!a.ignoreSeries&&(c=a.zData.filter(x),c.length&&(e=b(a.options.zMin,Math.min(e,Math.max(k(c),
!1===a.options.displayNegative?a.options.zThreshold:-Number.MAX_VALUE))),m=b(a.options.zMax,Math.max(m,p(c)))))});var l=e===m?[{value:m}]:[{value:e},{value:(e+m)/2},{value:m,autoRanges:!0}];g.length&&g[0].radius&&l.reverse();l.forEach(function(b,f){g&&g[f]&&(l[f]=w(g[f],b))});return l};a.prototype.predictBubbleSizes=function(){var b=this.chart,a=this.fontMetrics,g=b.legend.options,c=g.floating,k=(g="horizontal"===g.layout)?b.legend.lastLineHeight:0,e=b.plotSizeX,m=b.plotSizeY,f=b.series[this.options.seriesIndex],
h=f.getPxExtremes();b=Math.ceil(h.minPxSize);h=Math.ceil(h.maxPxSize);var u=Math.min(m,e);f=f.options.maxSize;if(c||!/%$/.test(f))a=h;else if(f=parseFloat(f),a=(u+k-a.h/2)*f/100/(f/100+1),g&&m-a>=e||!g&&e-a>=m)a=h;return[b,Math.ceil(a)]};a.prototype.updateRanges=function(b,a){var g=this.legend.options.bubbleLegend;g.minSize=b;g.maxSize=a;g.ranges=this.getRanges()};a.prototype.correctSizes=function(){var b=this.legend,a=this.chart.series[this.options.seriesIndex].getPxExtremes();1<Math.abs(Math.ceil(a.maxPxSize)-
this.options.maxSize)&&(this.updateRanges(this.options.minSize,a.maxPxSize),b.render())};return a}()});A(d,"Series/Bubble/BubbleLegendComposition.js",[d["Series/Bubble/BubbleLegendDefaults.js"],d["Series/Bubble/BubbleLegendItem.js"],d["Core/DefaultOptions.js"],d["Core/Utilities.js"]],function(d,e,l,a){var c=l.setOptions,t=a.addEvent,p=a.objectEach,k=a.wrap,x;(function(a){function b(b,a,c){var f=this.legend,h=0<=g(this);if(f&&f.options.enabled&&f.bubbleLegend&&f.options.bubbleLegend.autoRanges&&h){var u=
f.bubbleLegend.options;h=f.bubbleLegend.predictBubbleSizes();f.bubbleLegend.updateRanges(h[0],h[1]);u.placed||(f.group.placed=!1,f.allItems.forEach(function(h){h.legendGroup.translateY=null}));f.render();this.getMargins();this.axes.forEach(function(h){h.visible&&h.render();u.placed||(h.setScale(),h.updateNames(),p(h.ticks,function(h){h.isNew=!0;h.isNewLabel=!0}))});u.placed=!0;this.getMargins();b.call(this,a,c);f.bubbleLegend.correctSizes();x(f,m(f))}else b.call(this,a,c),f&&f.options.enabled&&f.bubbleLegend&&
(f.render(),x(f,m(f)))}function g(b){b=b.series;for(var a=0;a<b.length;){if(b[a]&&b[a].isBubble&&b[a].visible&&b[a].zData.length)return a;a++}return-1}function m(b){b=b.allItems;var a=[],g=b.length,f,h=0;for(f=0;f<g;f++)if(b[f].legendItemHeight&&(b[f].itemHeight=b[f].legendItemHeight),b[f]===b[g-1]||b[f+1]&&b[f]._legendItemPos[1]!==b[f+1]._legendItemPos[1]){a.push({height:0});var u=a[a.length-1];for(h;h<=f;h++)b[h].itemHeight>u.height&&(u.height=b[h].itemHeight);u.step=f}return a}function n(b){var a=
this.bubbleLegend,c=this.options,f=c.bubbleLegend,h=g(this.chart);a&&a.ranges&&a.ranges.length&&(f.ranges.length&&(f.autoRanges=!!f.ranges[0].autoRanges),this.destroyItem(a));0<=h&&c.enabled&&f.enabled&&(f.seriesIndex=h,this.bubbleLegend=new e(f,this),this.bubbleLegend.addToLegend(b.allItems))}function l(){var b=this.chart,a=this.visible,c=this.chart.legend;c&&c.bubbleLegend&&(this.visible=!a,this.ignoreSeries=a,b=0<=g(b),c.bubbleLegend.visible!==b&&(c.update({bubbleLegend:{enabled:b}}),c.bubbleLegend.visible=
b),this.visible=a)}function x(b,a){var g=b.options.rtl,f,h,u,r=0;b.allItems.forEach(function(b,c){f=b.legendGroup.translateX;h=b._legendItemPos[1];if((u=b.movementX)||g&&b.ranges)u=g?f-b.options.maxSize/2:f+u,b.legendGroup.attr({translateX:u});c>a[r].step&&r++;b.legendGroup.attr({translateY:Math.round(h+a[r].height/2)});b._legendItemPos[1]=h+a[r].height/2})}var w=[];a.compose=function(a,g,e){-1===w.indexOf(a)&&(w.push(a),c({legend:{bubbleLegend:d}}),k(a.prototype,"drawChartBox",b));-1===w.indexOf(g)&&
(w.push(g),t(g,"afterGetAllItems",n));-1===w.indexOf(e)&&(w.push(e),t(e,"legendItemClick",l))}})(x||(x={}));return x});A(d,"Series/Bubble/BubblePoint.js",[d["Core/Series/Point.js"],d["Core/Series/SeriesRegistry.js"],d["Core/Utilities.js"]],function(d,e,l){var a=this&&this.__extends||function(){var a=function(c,e){a=Object.setPrototypeOf||{__proto__:[]}instanceof Array&&function(a,c){a.__proto__=c}||function(a,c){for(var e in c)c.hasOwnProperty(e)&&(a[e]=c[e])};return a(c,e)};return function(c,e){function k(){this.constructor=
c}a(c,e);c.prototype=null===e?Object.create(e):(k.prototype=e.prototype,new k)}}();l=l.extend;e=function(c){function e(){var a=null!==c&&c.apply(this,arguments)||this;a.options=void 0;a.series=void 0;return a}a(e,c);e.prototype.haloPath=function(a){return d.prototype.haloPath.call(this,0===a?0:(this.marker?this.marker.radius||0:0)+a)};return e}(e.seriesTypes.scatter.prototype.pointClass);l(e.prototype,{ttBelow:!1});return e});A(d,"Series/Bubble/BubbleSeries.js",[d["Core/Axis/Axis.js"],d["Series/Bubble/BubbleLegendComposition.js"],
d["Series/Bubble/BubblePoint.js"],d["Core/Color/Color.js"],d["Core/Globals.js"],d["Core/Series/Series.js"],d["Core/Series/SeriesRegistry.js"],d["Core/Utilities.js"]],function(d,e,l,a,c,t,p,k){var x=this&&this.__extends||function(){var b=function(f,h){b=Object.setPrototypeOf||{__proto__:[]}instanceof Array&&function(b,h){b.__proto__=h}||function(b,h){for(var f in h)h.hasOwnProperty(f)&&(b[f]=h[f])};return b(f,h)};return function(f,h){function a(){this.constructor=f}b(f,h);f.prototype=null===h?Object.create(h):
(a.prototype=h.prototype,new a)}}(),w=a.parse;a=c.noop;var b=p.seriesTypes;c=b.column;var g=b.scatter;b=k.addEvent;var m=k.arrayMax,n=k.arrayMin,q=k.clamp,H=k.extend,K=k.isNumber,I=k.merge,y=k.pick;k=function(b){function f(){var h=null!==b&&b.apply(this,arguments)||this;h.data=void 0;h.maxPxSize=void 0;h.minPxSize=void 0;h.options=void 0;h.points=void 0;h.radii=void 0;h.yData=void 0;h.zData=void 0;return h}x(f,b);f.prototype.animate=function(b){!b&&this.points.length<this.options.animationLimit&&
this.points.forEach(function(b){var h=b.graphic;h&&h.width&&(this.hasRendered||h.attr({x:b.plotX,y:b.plotY,width:1,height:1}),h.animate(this.markerAttribs(b),this.options.animation))},this)};f.prototype.getRadii=function(){var b=this,f=this.zData,a=this.yData,g=[],c=this.chart.bubbleZExtremes;var e=this.getPxExtremes();var m=e.minPxSize,k=e.maxPxSize;if(!c){var n=Number.MAX_VALUE,l=-Number.MAX_VALUE,d;this.chart.series.forEach(function(h){h.bubblePadding&&(h.visible||!b.chart.options.chart.ignoreHiddenSeries)&&
(h=h.getZExtremes())&&(n=Math.min(n||h.zMin,h.zMin),l=Math.max(l||h.zMax,h.zMax),d=!0)});d?(c={zMin:n,zMax:l},this.chart.bubbleZExtremes=c):c={zMin:0,zMax:0}}var p=0;for(e=f.length;p<e;p++){var q=f[p];g.push(this.getRadius(c.zMin,c.zMax,m,k,q,a[p]))}this.radii=g};f.prototype.getRadius=function(b,f,a,g,c,e){var h=this.options,r="width"!==h.sizeBy,u=h.zThreshold,v=f-b,m=.5;if(null===e||null===c)return null;if(K(c)){h.sizeByAbsoluteValue&&(c=Math.abs(c-u),v=Math.max(f-u,Math.abs(b-u)),b=0);if(c<b)return a/
2-1;0<v&&(m=(c-b)/v)}r&&0<=m&&(m=Math.sqrt(m));return Math.ceil(a+m*(g-a))/2};f.prototype.hasData=function(){return!!this.processedXData.length};f.prototype.pointAttribs=function(b,f){var h=this.options.marker.fillOpacity;b=t.prototype.pointAttribs.call(this,b,f);1!==h&&(b.fill=w(b.fill).setOpacity(h).get("rgba"));return b};f.prototype.translate=function(){b.prototype.translate.call(this);this.getRadii();this.translateBubble()};f.prototype.translateBubble=function(){for(var b=this.data,f=this.radii,
a=this.getPxExtremes().minPxSize,g=b.length;g--;){var c=b[g],e=f?f[g]:0;K(e)&&e>=a/2?(c.marker=H(c.marker,{radius:e,width:2*e,height:2*e}),c.dlBox={x:c.plotX-e,y:c.plotY-e,width:2*e,height:2*e}):c.shapeArgs=c.plotY=c.dlBox=void 0}};f.prototype.getPxExtremes=function(){var b=Math.min(this.chart.plotWidth,this.chart.plotHeight),f=function(h){if("string"===typeof h){var f=/%$/.test(h);h=parseInt(h,10)}return f?b*h/100:h},a=f(y(this.options.minSize,8));f=Math.max(f(y(this.options.maxSize,"20%")),a);return{minPxSize:a,
maxPxSize:f}};f.prototype.getZExtremes=function(){var b=this.options,f=(this.zData||[]).filter(K);if(f.length){var a=y(b.zMin,q(n(f),!1===b.displayNegative?b.zThreshold||0:-Number.MAX_VALUE,Number.MAX_VALUE));b=y(b.zMax,m(f));if(K(a)&&K(b))return{zMin:a,zMax:b}}};f.compose=e.compose;f.defaultOptions=I(g.defaultOptions,{dataLabels:{formatter:function(){var b=this.series.chart.numberFormatter,f=this.point.z;return K(f)?b(f,-1):""},inside:!0,verticalAlign:"middle"},animationLimit:250,marker:{lineColor:null,
lineWidth:1,fillOpacity:.5,radius:null,states:{hover:{radiusPlus:0}},symbol:"circle"},minSize:8,maxSize:"20%",softThreshold:!1,states:{hover:{halo:{size:5}}},tooltip:{pointFormat:"({point.x}, {point.y}), Size: {point.z}"},turboThreshold:0,zThreshold:0,zoneAxis:"z"});return f}(g);H(k.prototype,{alignDataLabel:c.prototype.alignDataLabel,applyZones:a,bubblePadding:!0,buildKDTree:a,directTouch:!0,isBubble:!0,pointArrayMap:["y","z"],pointClass:l,parallelArrays:["x","y","z"],trackerGroups:["group","dataLabelsGroup"],
specialGroup:"group",zoneAxis:"z"});b(k,"updatedData",function(b){delete b.target.chart.bubbleZExtremes});d.prototype.beforePadding=function(){var b=this,f=this.len,h=this.chart,a=0,g=f,c=this.isXAxis,v=c?"xData":"yData",e=this.min,m=this.max-e,k=f/m,n;this.series.forEach(function(f){if(f.bubblePadding&&(f.visible||!h.options.chart.ignoreHiddenSeries)){n=b.allowZoomOutside=!0;var r=f[v];c&&f.getRadii(0,0,f);if(0<m)for(var u=r.length;u--;)if(K(r[u])&&b.dataMin<=r[u]&&r[u]<=b.max){var B=f.radii&&f.radii[u]||
0;a=Math.min((r[u]-e)*k-B,a);g=Math.max((r[u]-e)*k+B,g)}}});n&&0<m&&!this.logarithmic&&(g-=f,k*=(f+Math.max(0,a)-Math.min(g,f))/f,[["min","userMin",a],["max","userMax",g]].forEach(function(h){"undefined"===typeof y(b.options[h[0]],b[h[1]])&&(b[h[0]]+=h[2]/k)}))};p.registerSeriesType("bubble",k);"";"";return k});A(d,"Series/ColumnRange/ColumnRangePoint.js",[d["Core/Series/SeriesRegistry.js"],d["Core/Utilities.js"]],function(d,e){var l=this&&this.__extends||function(){var a=function(c,e){a=Object.setPrototypeOf||
{__proto__:[]}instanceof Array&&function(a,b){a.__proto__=b}||function(a,b){for(var g in b)b.hasOwnProperty(g)&&(a[g]=b[g])};return a(c,e)};return function(c,e){function k(){this.constructor=c}a(c,e);c.prototype=null===e?Object.create(e):(k.prototype=e.prototype,new k)}}(),a=d.seriesTypes;d=a.column.prototype.pointClass;var c=e.extend,t=e.isNumber;e=function(a){function c(){var c=null!==a&&a.apply(this,arguments)||this;c.series=void 0;c.options=void 0;c.barX=void 0;c.pointWidth=void 0;c.shapeType=
void 0;return c}l(c,a);c.prototype.isValid=function(){return t(this.low)};return c}(a.arearange.prototype.pointClass);c(e.prototype,{setState:d.prototype.setState});return e});A(d,"Series/ColumnRange/ColumnRangeSeries.js",[d["Series/ColumnRange/ColumnRangePoint.js"],d["Core/Globals.js"],d["Core/Series/SeriesRegistry.js"],d["Core/Utilities.js"]],function(d,e,l,a){var c=this&&this.__extends||function(){var b=function(a,g){b=Object.setPrototypeOf||{__proto__:[]}instanceof Array&&function(b,a){b.__proto__=
a}||function(b,a){for(var g in a)a.hasOwnProperty(g)&&(b[g]=a[g])};return b(a,g)};return function(a,g){function c(){this.constructor=a}b(a,g);a.prototype=null===g?Object.create(g):(c.prototype=g.prototype,new c)}}();e=e.noop;var t=l.seriesTypes,p=t.arearange,k=t.column,x=k.prototype,w=p.prototype,b=a.clamp,g=a.merge,m=a.pick;a=a.extend;var n={pointRange:null,marker:null,states:{hover:{halo:!1}}};t=function(a){function e(){var b=null!==a&&a.apply(this,arguments)||this;b.data=void 0;b.points=void 0;
b.options=void 0;return b}c(e,a);e.prototype.setOptions=function(){g(!0,arguments[0],{stacking:void 0});return w.setOptions.apply(this,arguments)};e.prototype.translate=function(){var a=this,g=a.yAxis,c=a.xAxis,e=c.startAngleRad,f,h=a.chart,u=a.xAxis.isRadial,r=Math.max(h.chartWidth,h.chartHeight)+999,k;x.translate.apply(a);a.points.forEach(function(v){var n=v.shapeArgs||{},B=a.options.minPointLength;v.plotHigh=k=b(g.translate(v.high,0,1,0,1),-r,r);v.plotLow=b(v.plotY,-r,r);var l=k;var d=m(v.rectPlotY,
v.plotY)-k;Math.abs(d)<B?(B-=d,d+=B,l-=B/2):0>d&&(d*=-1,l-=d);u?(f=v.barX+e,v.shapeType="arc",v.shapeArgs=a.polarArc(l+d,l,f,f+v.pointWidth)):(n.height=d,n.y=l,B=n.x,B=void 0===B?0:B,n=n.width,n=void 0===n?0:n,v.tooltipPos=h.inverted?[g.len+g.pos-h.plotLeft-l-d/2,c.len+c.pos-h.plotTop-B-n/2,d]:[c.left-h.plotLeft+B+n/2,g.pos-h.plotTop+l+d/2,d])})};e.prototype.crispCol=function(){return x.crispCol.apply(this,arguments)};e.prototype.drawPoints=function(){return x.drawPoints.apply(this,arguments)};e.prototype.drawTracker=
function(){return x.drawTracker.apply(this,arguments)};e.prototype.getColumnMetrics=function(){return x.getColumnMetrics.apply(this,arguments)};e.prototype.pointAttribs=function(){return x.pointAttribs.apply(this,arguments)};e.prototype.adjustForMissingColumns=function(){return x.adjustForMissingColumns.apply(this,arguments)};e.prototype.animate=function(){return x.animate.apply(this,arguments)};e.prototype.translate3dPoints=function(){return x.translate3dPoints.apply(this,arguments)};e.prototype.translate3dShapes=
function(){return x.translate3dShapes.apply(this,arguments)};e.defaultOptions=g(k.defaultOptions,p.defaultOptions,n);return e}(p);a(t.prototype,{directTouch:!0,trackerGroups:["group","dataLabelsGroup"],drawGraph:e,getSymbol:e,polarArc:function(){return x.polarArc.apply(this,arguments)},pointClass:d});l.registerSeriesType("columnrange",t);"";return t});A(d,"Series/ColumnPyramid/ColumnPyramidSeries.js",[d["Series/Column/ColumnSeries.js"],d["Core/Series/SeriesRegistry.js"],d["Core/Utilities.js"]],function(d,
e,l){var a=this&&this.__extends||function(){var a=function(c,b){a=Object.setPrototypeOf||{__proto__:[]}instanceof Array&&function(b,a){b.__proto__=a}||function(b,a){for(var g in a)a.hasOwnProperty(g)&&(b[g]=a[g])};return a(c,b)};return function(c,b){function g(){this.constructor=c}a(c,b);c.prototype=null===b?Object.create(b):(g.prototype=b.prototype,new g)}}(),c=d.prototype,t=l.clamp,p=l.merge,k=l.pick;l=function(e){function l(){var b=null!==e&&e.apply(this,arguments)||this;b.data=void 0;b.options=
void 0;b.points=void 0;return b}a(l,e);l.prototype.translate=function(){var b=this,a=b.chart,e=b.options,n=b.dense=2>b.closestPointRange*b.xAxis.transA;n=b.borderWidth=k(e.borderWidth,n?0:1);var d=b.yAxis,l=e.threshold,p=b.translatedThreshold=d.getThreshold(l),x=k(e.minPointLength,5),w=b.getColumnMetrics(),z=w.width,f=b.barW=Math.max(z,1+2*n),h=b.pointXOffset=w.offset;a.inverted&&(p-=.5);e.pointPadding&&(f=Math.ceil(f));c.translate.apply(b);b.points.forEach(function(g){var c=k(g.yBottom,p),u=999+
Math.abs(c),v=t(g.plotY,-u,d.len+u);u=g.plotX+h;var m=f/2,n=Math.min(v,c);c=Math.max(v,c)-n;var q;g.barX=u;g.pointWidth=z;g.tooltipPos=a.inverted?[d.len+d.pos-a.plotLeft-v,b.xAxis.len-u-m,c]:[u+m,v+d.pos-a.plotTop,c];v=l+(g.total||g.y);"percent"===e.stacking&&(v=l+(0>g.y)?-100:100);v=d.toPixels(v,!0);var w=(q=a.plotHeight-v-(a.plotHeight-p))?m*(n-v)/q:0;var y=q?m*(n+c-v)/q:0;q=u-w+m;w=u+w+m;var C=u+y+m;y=u-y+m;var H=n-x;var D=n+c;0>g.y&&(H=n,D=n+c+x);a.inverted&&(C=d.width-n,q=v-(d.width-p),w=m*(v-
C)/q,y=m*(v-(C-c))/q,q=u+m+w,w=q-2*w,C=u-y+m,y=u+y+m,H=n,D=n+c-x,0>g.y&&(D=n+c+x));g.shapeType="path";g.shapeArgs={x:q,y:H,width:w-q,height:c,d:[["M",q,H],["L",w,H],["L",C,D],["L",y,D],["Z"]]}})};l.defaultOptions=p(d.defaultOptions,{});return l}(d);e.registerSeriesType("columnpyramid",l);"";return l});A(d,"Series/ErrorBar/ErrorBarSeries.js",[d["Series/BoxPlot/BoxPlotSeries.js"],d["Series/Column/ColumnSeries.js"],d["Core/Series/SeriesRegistry.js"],d["Core/Utilities.js"]],function(d,e,l,a){var c=this&&
this.__extends||function(){var a=function(c,b){a=Object.setPrototypeOf||{__proto__:[]}instanceof Array&&function(b,a){b.__proto__=a}||function(b,a){for(var g in a)a.hasOwnProperty(g)&&(b[g]=a[g])};return a(c,b)};return function(c,b){function g(){this.constructor=c}a(c,b);c.prototype=null===b?Object.create(b):(g.prototype=b.prototype,new g)}}(),t=l.seriesTypes.arearange,p=a.merge;a=a.extend;var k=function(a){function k(){var b=null!==a&&a.apply(this,arguments)||this;b.data=void 0;b.options=void 0;
b.points=void 0;return b}c(k,a);k.prototype.getColumnMetrics=function(){return this.linkedParent&&this.linkedParent.columnMetrics||e.prototype.getColumnMetrics.call(this)};k.prototype.drawDataLabels=function(){var b=this.pointValKey;t&&(t.prototype.drawDataLabels.call(this),this.data.forEach(function(a){a.y=a[b]}))};k.prototype.toYData=function(b){return[b.low,b.high]};k.defaultOptions=p(d.defaultOptions,{color:"#000000",grouping:!1,linkedTo:":previous",tooltip:{pointFormat:'<span style="color:{point.color}">\u25cf</span> {series.name}: <b>{point.low}</b> - <b>{point.high}</b><br/>'},
whiskerWidth:null});return k}(d);a(k.prototype,{pointArrayMap:["low","high"],pointValKey:"high",doQuartiles:!1});l.registerSeriesType("errorbar",k);"";return k});A(d,"Series/Gauge/GaugePoint.js",[d["Core/Series/SeriesRegistry.js"]],function(d){var e=this&&this.__extends||function(){var e=function(a,c){e=Object.setPrototypeOf||{__proto__:[]}instanceof Array&&function(a,c){a.__proto__=c}||function(a,c){for(var e in c)c.hasOwnProperty(e)&&(a[e]=c[e])};return e(a,c)};return function(a,c){function d(){this.constructor=
a}e(a,c);a.prototype=null===c?Object.create(c):(d.prototype=c.prototype,new d)}}();return function(d){function a(){var a=null!==d&&d.apply(this,arguments)||this;a.options=void 0;a.series=void 0;a.shapeArgs=void 0;return a}e(a,d);a.prototype.setState=function(a){this.state=a};return a}(d.series.prototype.pointClass)});A(d,"Series/Gauge/GaugeSeries.js",[d["Series/Gauge/GaugePoint.js"],d["Core/Globals.js"],d["Core/Series/SeriesRegistry.js"],d["Core/Utilities.js"]],function(d,e,l,a){var c=this&&this.__extends||
function(){var b=function(a,g){b=Object.setPrototypeOf||{__proto__:[]}instanceof Array&&function(b,a){b.__proto__=a}||function(b,a){for(var g in a)a.hasOwnProperty(g)&&(b[g]=a[g])};return b(a,g)};return function(a,g){function c(){this.constructor=a}b(a,g);a.prototype=null===g?Object.create(g):(c.prototype=g.prototype,new c)}}();e=e.noop;var t=l.series,p=l.seriesTypes.column,k=a.clamp,x=a.isNumber,w=a.extend,b=a.merge,g=a.pick,m=a.pInt;a=function(a){function e(){var b=null!==a&&a.apply(this,arguments)||
this;b.data=void 0;b.points=void 0;b.options=void 0;b.yAxis=void 0;return b}c(e,a);e.prototype.translate=function(){var a=this.yAxis,c=this.options,e=a.center;this.generatePoints();this.points.forEach(function(d){var n=b(c.dial,d.dial),f=m(g(n.radius,"80%"))*e[2]/200,h=m(g(n.baseLength,"70%"))*f/100,u=m(g(n.rearLength,"10%"))*f/100,r=n.baseWidth||3,l=n.topWidth||1,v=c.overshoot,p=a.startAngleRad+a.translate(d.y,null,null,null,!0);if(x(v)||!1===c.wrap)v=x(v)?v/180*Math.PI:0,p=k(p,a.startAngleRad-v,
a.endAngleRad+v);p=180*p/Math.PI;d.shapeType="path";d.shapeArgs={d:n.path||[["M",-u,-r/2],["L",h,-r/2],["L",f,-l/2],["L",f,l/2],["L",h,r/2],["L",-u,r/2],["Z"]],translateX:e[0],translateY:e[1],rotation:p};d.plotX=e[0];d.plotY=e[1]})};e.prototype.drawPoints=function(){var a=this,c=a.chart,e=a.yAxis.center,d=a.pivot,m=a.options,f=m.pivot,h=c.renderer;a.points.forEach(function(f){var g=f.graphic,e=f.shapeArgs,u=e.d,d=b(m.dial,f.dial);g?(g.animate(e),e.d=u):f.graphic=h[f.shapeType](e).attr({rotation:e.rotation,
zIndex:1}).addClass("highcharts-dial").add(a.group);if(!c.styledMode)f.graphic[g?"animate":"attr"]({stroke:d.borderColor||"none","stroke-width":d.borderWidth||0,fill:d.backgroundColor||"#000000"})});d?d.animate({translateX:e[0],translateY:e[1]}):(a.pivot=h.circle(0,0,g(f.radius,5)).attr({zIndex:2}).addClass("highcharts-pivot").translate(e[0],e[1]).add(a.group),c.styledMode||a.pivot.attr({"stroke-width":f.borderWidth||0,stroke:f.borderColor||"#cccccc",fill:f.backgroundColor||"#000000"}))};e.prototype.animate=
function(b){var a=this;b||a.points.forEach(function(b){var g=b.graphic;g&&(g.attr({rotation:180*a.yAxis.startAngleRad/Math.PI}),g.animate({rotation:b.shapeArgs.rotation},a.options.animation))})};e.prototype.render=function(){this.group=this.plotGroup("group","series",this.visible?"visible":"hidden",this.options.zIndex,this.chart.seriesGroup);t.prototype.render.call(this);this.group.clip(this.chart.clipRect)};e.prototype.setData=function(b,a){t.prototype.setData.call(this,b,!1);this.processData();
this.generatePoints();g(a,!0)&&this.chart.redraw()};e.prototype.hasData=function(){return!!this.points.length};e.defaultOptions=b(t.defaultOptions,{dataLabels:{borderColor:"#cccccc",borderRadius:3,borderWidth:1,crop:!1,defer:!1,enabled:!0,verticalAlign:"top",y:15,zIndex:2},dial:{},pivot:{},tooltip:{headerFormat:""},showInLegend:!1});return e}(t);w(a.prototype,{angular:!0,directTouch:!0,drawGraph:e,drawTracker:p.prototype.drawTracker,fixedBox:!0,forceDL:!0,noSharedTooltip:!0,pointClass:d,trackerGroups:["group",
"dataLabelsGroup"]});l.registerSeriesType("gauge",a);"";return a});A(d,"Series/PackedBubble/PackedBubblePoint.js",[d["Core/Chart/Chart.js"],d["Core/Series/Point.js"],d["Core/Series/SeriesRegistry.js"]],function(d,e,l){var a=this&&this.__extends||function(){var a=function(c,e){a=Object.setPrototypeOf||{__proto__:[]}instanceof Array&&function(a,c){a.__proto__=c}||function(a,c){for(var e in c)c.hasOwnProperty(e)&&(a[e]=c[e])};return a(c,e)};return function(c,e){function d(){this.constructor=c}a(c,e);
c.prototype=null===e?Object.create(e):(d.prototype=e.prototype,new d)}}();return function(c){function l(){var a=null!==c&&c.apply(this,arguments)||this;a.degree=NaN;a.mass=NaN;a.radius=NaN;a.options=void 0;a.series=void 0;a.value=null;return a}a(l,c);l.prototype.destroy=function(){this.series.layout&&this.series.layout.removeElementFromCollection(this,this.series.layout.nodes);return e.prototype.destroy.apply(this,arguments)};l.prototype.firePointEvent=function(){var a=this.series.options;if(this.isParentNode&&
a.parentNode){var c=a.allowPointSelect;a.allowPointSelect=a.parentNode.allowPointSelect;e.prototype.firePointEvent.apply(this,arguments);a.allowPointSelect=c}else e.prototype.firePointEvent.apply(this,arguments)};l.prototype.select=function(){var a=this.series.chart;this.isParentNode?(a.getSelectedPoints=a.getSelectedParentNodes,e.prototype.select.apply(this,arguments),a.getSelectedPoints=d.prototype.getSelectedPoints):e.prototype.select.apply(this,arguments)};return l}(l.seriesTypes.bubble.prototype.pointClass)});
A(d,"Series/Networkgraph/DraggableNodes.js",[d["Core/Chart/Chart.js"],d["Core/Globals.js"],d["Core/Utilities.js"]],function(d,e,l){var a=l.addEvent;e.dragNodesMixin={onMouseDown:function(a,e){e=this.chart.pointer.normalize(e);a.fixedPosition={chartX:e.chartX,chartY:e.chartY,plotX:a.plotX,plotY:a.plotY};a.inDragMode=!0},onMouseMove:function(a,e){if(a.fixedPosition&&a.inDragMode){var c=this.chart,d=c.pointer.normalize(e);e=a.fixedPosition.chartX-d.chartX;d=a.fixedPosition.chartY-d.chartY;var l=void 0,
w=void 0,b=c.graphLayoutsLookup;if(5<Math.abs(e)||5<Math.abs(d))l=a.fixedPosition.plotX-e,w=a.fixedPosition.plotY-d,c.isInsidePlot(l,w)&&(a.plotX=l,a.plotY=w,a.hasDragged=!0,this.redrawHalo(a),b.forEach(function(b){b.restartSimulation()}))}},onMouseUp:function(a,e){a.fixedPosition&&(a.hasDragged&&(this.layout.enableSimulation?this.layout.start():this.chart.redraw()),a.inDragMode=a.hasDragged=!1,this.options.fixedDraggable||delete a.fixedPosition)},redrawHalo:function(a){a&&this.halo&&this.halo.attr({d:a.haloPath(this.options.states.hover.halo.size)})}};
a(d,"load",function(){var c=this,e,d,l;c.container&&(e=a(c.container,"mousedown",function(e){var k=c.hoverPoint;k&&k.series&&k.series.hasDraggableNodes&&k.series.options.draggable&&(k.series.onMouseDown(k,e),d=a(c.container,"mousemove",function(b){return k&&k.series&&k.series.onMouseMove(k,b)}),l=a(c.container.ownerDocument,"mouseup",function(b){d();l();return k&&k.series&&k.series.onMouseUp(k,b)}))}));a(c,"destroy",function(){e()})})});A(d,"Series/Networkgraph/Integrations.js",[d["Core/Globals.js"]],
function(d){d.networkgraphIntegrations={verlet:{attractiveForceFunction:function(e,d){return(d-e)/e},repulsiveForceFunction:function(e,d){return(d-e)/e*(d>e?1:0)},barycenter:function(){var e=this.options.gravitationalConstant,d=this.barycenter.xFactor,a=this.barycenter.yFactor;d=(d-(this.box.left+this.box.width)/2)*e;a=(a-(this.box.top+this.box.height)/2)*e;this.nodes.forEach(function(c){c.fixedPosition||(c.plotX-=d/c.mass/c.degree,c.plotY-=a/c.mass/c.degree)})},repulsive:function(e,d,a){d=d*this.diffTemperature/
e.mass/e.degree;e.fixedPosition||(e.plotX+=a.x*d,e.plotY+=a.y*d)},attractive:function(e,d,a){var c=e.getMass(),l=-a.x*d*this.diffTemperature;d=-a.y*d*this.diffTemperature;e.fromNode.fixedPosition||(e.fromNode.plotX-=l*c.fromNode/e.fromNode.degree,e.fromNode.plotY-=d*c.fromNode/e.fromNode.degree);e.toNode.fixedPosition||(e.toNode.plotX+=l*c.toNode/e.toNode.degree,e.toNode.plotY+=d*c.toNode/e.toNode.degree)},integrate:function(e,d){var a=-e.options.friction,c=e.options.maxSpeed,l=(d.plotX+d.dispX-d.prevX)*
a;a*=d.plotY+d.dispY-d.prevY;var p=Math.abs,k=p(l)/(l||1);p=p(a)/(a||1);l=k*Math.min(c,Math.abs(l));a=p*Math.min(c,Math.abs(a));d.prevX=d.plotX+d.dispX;d.prevY=d.plotY+d.dispY;d.plotX+=l;d.plotY+=a;d.temperature=e.vectorLength({x:l,y:a})},getK:function(e){return Math.pow(e.box.width*e.box.height/e.nodes.length,.5)}},euler:{attractiveForceFunction:function(e,d){return e*e/d},repulsiveForceFunction:function(e,d){return d*d/e},barycenter:function(){var e=this.options.gravitationalConstant,d=this.barycenter.xFactor,
a=this.barycenter.yFactor;this.nodes.forEach(function(c){if(!c.fixedPosition){var l=c.getDegree();l*=1+l/2;c.dispX+=(d-c.plotX)*e*l/c.degree;c.dispY+=(a-c.plotY)*e*l/c.degree}})},repulsive:function(e,d,a,c){e.dispX+=a.x/c*d/e.degree;e.dispY+=a.y/c*d/e.degree},attractive:function(e,d,a,c){var l=e.getMass(),p=a.x/c*d;d*=a.y/c;e.fromNode.fixedPosition||(e.fromNode.dispX-=p*l.fromNode/e.fromNode.degree,e.fromNode.dispY-=d*l.fromNode/e.fromNode.degree);e.toNode.fixedPosition||(e.toNode.dispX+=p*l.toNode/
e.toNode.degree,e.toNode.dispY+=d*l.toNode/e.toNode.degree)},integrate:function(e,d){d.dispX+=d.dispX*e.options.friction;d.dispY+=d.dispY*e.options.friction;var a=d.temperature=e.vectorLength({x:d.dispX,y:d.dispY});0!==a&&(d.plotX+=d.dispX/a*Math.min(Math.abs(d.dispX),e.temperature),d.plotY+=d.dispY/a*Math.min(Math.abs(d.dispY),e.temperature))},getK:function(e){return Math.pow(e.box.width*e.box.height/e.nodes.length,.3)}}}});A(d,"Series/Networkgraph/QuadTree.js",[d["Core/Globals.js"],d["Core/Utilities.js"]],
function(d,e){e=e.extend;var l=d.QuadTreeNode=function(a){this.box=a;this.boxSize=Math.min(a.width,a.height);this.nodes=[];this.body=this.isInternal=!1;this.isEmpty=!0};e(l.prototype,{insert:function(a,c){this.isInternal?this.nodes[this.getBoxPosition(a)].insert(a,c-1):(this.isEmpty=!1,this.body?c?(this.isInternal=!0,this.divideBox(),!0!==this.body&&(this.nodes[this.getBoxPosition(this.body)].insert(this.body,c-1),this.body=!0),this.nodes[this.getBoxPosition(a)].insert(a,c-1)):(c=new l({top:a.plotX,
left:a.plotY,width:.1,height:.1}),c.body=a,c.isInternal=!1,this.nodes.push(c)):(this.isInternal=!1,this.body=a))},updateMassAndCenter:function(){var a=0,c=0,e=0;this.isInternal?(this.nodes.forEach(function(d){d.isEmpty||(a+=d.mass,c+=d.plotX*d.mass,e+=d.plotY*d.mass)}),c/=a,e/=a):this.body&&(a=this.body.mass,c=this.body.plotX,e=this.body.plotY);this.mass=a;this.plotX=c;this.plotY=e},divideBox:function(){var a=this.box.width/2,c=this.box.height/2;this.nodes[0]=new l({left:this.box.left,top:this.box.top,
width:a,height:c});this.nodes[1]=new l({left:this.box.left+a,top:this.box.top,width:a,height:c});this.nodes[2]=new l({left:this.box.left+a,top:this.box.top+c,width:a,height:c});this.nodes[3]=new l({left:this.box.left,top:this.box.top+c,width:a,height:c})},getBoxPosition:function(a){var c=a.plotY<this.box.top+this.box.height/2;return a.plotX<this.box.left+this.box.width/2?c?0:3:c?1:2}});d=d.QuadTree=function(a,c,e,d){this.box={left:a,top:c,width:e,height:d};this.maxDepth=25;this.root=new l(this.box,
"0");this.root.isInternal=!0;this.root.isRoot=!0;this.root.divideBox()};e(d.prototype,{insertNodes:function(a){a.forEach(function(a){this.root.insert(a,this.maxDepth)},this)},visitNodeRecursive:function(a,c,e){var d;a||(a=this.root);a===this.root&&c&&(d=c(a));!1!==d&&(a.nodes.forEach(function(a){if(a.isInternal){c&&(d=c(a));if(!1===d)return;this.visitNodeRecursive(a,c,e)}else a.body&&c&&c(a.body);e&&e(a)},this),a===this.root&&e&&e(a))},calculateMassAndCenter:function(){this.visitNodeRecursive(null,
null,function(a){a.updateMassAndCenter()})}})});A(d,"Series/Networkgraph/Layouts.js",[d["Core/Chart/Chart.js"],d["Core/Animation/AnimationUtilities.js"],d["Core/Globals.js"],d["Core/Utilities.js"]],function(d,e,l,a){var c=e.setAnimation;e=a.addEvent;var t=a.clamp,p=a.defined,k=a.extend,x=a.isFunction,w=a.pick;l.layouts={"reingold-fruchterman":function(){}};k(l.layouts["reingold-fruchterman"].prototype,{init:function(b){this.options=b;this.nodes=[];this.links=[];this.series=[];this.box={x:0,y:0,width:0,
height:0};this.setInitialRendering(!0);this.integration=l.networkgraphIntegrations[b.integration];this.enableSimulation=b.enableSimulation;this.attractiveForce=w(b.attractiveForce,this.integration.attractiveForceFunction);this.repulsiveForce=w(b.repulsiveForce,this.integration.repulsiveForceFunction);this.approximation=b.approximation},updateSimulation:function(b){this.enableSimulation=w(b,this.options.enableSimulation)},start:function(){var b=this.series,a=this.options;this.currentStep=0;this.forces=
b[0]&&b[0].forces||[];this.chart=b[0]&&b[0].chart;this.initialRendering&&(this.initPositions(),b.forEach(function(b){b.finishedAnimating=!0;b.render()}));this.setK();this.resetSimulation(a);this.enableSimulation&&this.step()},step:function(){var b=this,a=this.series;b.currentStep++;"barnes-hut"===b.approximation&&(b.createQuadTree(),b.quadTree.calculateMassAndCenter());b.forces.forEach(function(a){b[a+"Forces"](b.temperature)});b.applyLimits(b.temperature);b.temperature=b.coolDown(b.startTemperature,
b.diffTemperature,b.currentStep);b.prevSystemTemperature=b.systemTemperature;b.systemTemperature=b.getSystemTemperature();b.enableSimulation&&(a.forEach(function(b){b.chart&&b.render()}),b.maxIterations--&&isFinite(b.temperature)&&!b.isStable()?(b.simulation&&l.win.cancelAnimationFrame(b.simulation),b.simulation=l.win.requestAnimationFrame(function(){b.step()})):b.simulation=!1)},stop:function(){this.simulation&&l.win.cancelAnimationFrame(this.simulation)},setArea:function(b,a,c,e){this.box={left:b,
top:a,width:c,height:e}},setK:function(){this.k=this.options.linkLength||this.integration.getK(this)},addElementsToCollection:function(b,a){b.forEach(function(b){-1===a.indexOf(b)&&a.push(b)})},removeElementFromCollection:function(b,a){b=a.indexOf(b);-1!==b&&a.splice(b,1)},clear:function(){this.nodes.length=0;this.links.length=0;this.series.length=0;this.resetSimulation()},resetSimulation:function(){this.forcedStop=!1;this.systemTemperature=0;this.setMaxIterations();this.setTemperature();this.setDiffTemperature()},
restartSimulation:function(){this.simulation?this.resetSimulation():(this.setInitialRendering(!1),this.enableSimulation?this.start():this.setMaxIterations(1),this.chart&&this.chart.redraw(),this.setInitialRendering(!0))},setMaxIterations:function(b){this.maxIterations=w(b,this.options.maxIterations)},setTemperature:function(){this.temperature=this.startTemperature=Math.sqrt(this.nodes.length)},setDiffTemperature:function(){this.diffTemperature=this.startTemperature/(this.options.maxIterations+1)},
setInitialRendering:function(b){this.initialRendering=b},createQuadTree:function(){this.quadTree=new l.QuadTree(this.box.left,this.box.top,this.box.width,this.box.height);this.quadTree.insertNodes(this.nodes)},initPositions:function(){var b=this.options.initialPositions;x(b)?(b.call(this),this.nodes.forEach(function(b){p(b.prevX)||(b.prevX=b.plotX);p(b.prevY)||(b.prevY=b.plotY);b.dispX=0;b.dispY=0})):"circle"===b?this.setCircularPositions():this.setRandomPositions()},setCircularPositions:function(){function b(a){a.linksFrom.forEach(function(a){l[a.toNode.id]||
(l[a.toNode.id]=!0,k.push(a.toNode),b(a.toNode))})}var a=this.box,c=this.nodes,e=2*Math.PI/(c.length+1),d=c.filter(function(b){return 0===b.linksTo.length}),k=[],l={},p=this.options.initialPositionRadius;d.forEach(function(a){k.push(a);b(a)});k.length?c.forEach(function(b){-1===k.indexOf(b)&&k.push(b)}):k=c;k.forEach(function(b,c){b.plotX=b.prevX=w(b.plotX,a.width/2+p*Math.cos(c*e));b.plotY=b.prevY=w(b.plotY,a.height/2+p*Math.sin(c*e));b.dispX=0;b.dispY=0})},setRandomPositions:function(){function b(b){b=
b*b/Math.PI;return b-=Math.floor(b)}var a=this.box,c=this.nodes,e=c.length+1;c.forEach(function(c,g){c.plotX=c.prevX=w(c.plotX,a.width*b(g));c.plotY=c.prevY=w(c.plotY,a.height*b(e+g));c.dispX=0;c.dispY=0})},force:function(b){this.integration[b].apply(this,Array.prototype.slice.call(arguments,1))},barycenterForces:function(){this.getBarycenter();this.force("barycenter")},getBarycenter:function(){var b=0,a=0,c=0;this.nodes.forEach(function(g){a+=g.plotX*g.mass;c+=g.plotY*g.mass;b+=g.mass});return this.barycenter=
{x:a,y:c,xFactor:a/b,yFactor:c/b}},barnesHutApproximation:function(b,a){var c=this.getDistXY(b,a),g=this.vectorLength(c);if(b!==a&&0!==g)if(a.isInternal)if(a.boxSize/g<this.options.theta&&0!==g){var e=this.repulsiveForce(g,this.k);this.force("repulsive",b,e*a.mass,c,g);var d=!1}else d=!0;else e=this.repulsiveForce(g,this.k),this.force("repulsive",b,e*a.mass,c,g);return d},repulsiveForces:function(){var b=this;"barnes-hut"===b.approximation?b.nodes.forEach(function(a){b.quadTree.visitNodeRecursive(null,
function(c){return b.barnesHutApproximation(a,c)})}):b.nodes.forEach(function(a){b.nodes.forEach(function(c){if(a!==c&&!a.fixedPosition){var g=b.getDistXY(a,c);var e=b.vectorLength(g);if(0!==e){var d=b.repulsiveForce(e,b.k);b.force("repulsive",a,d*c.mass,g,e)}}})})},attractiveForces:function(){var b=this,a,c,e;b.links.forEach(function(g){g.fromNode&&g.toNode&&(a=b.getDistXY(g.fromNode,g.toNode),c=b.vectorLength(a),0!==c&&(e=b.attractiveForce(c,b.k),b.force("attractive",g,e,a,c)))})},applyLimits:function(){var b=
this;b.nodes.forEach(function(a){a.fixedPosition||(b.integration.integrate(b,a),b.applyLimitBox(a,b.box),a.dispX=0,a.dispY=0)})},applyLimitBox:function(b,a){var c=b.radius;b.plotX=t(b.plotX,a.left+c,a.width-c);b.plotY=t(b.plotY,a.top+c,a.height-c)},coolDown:function(b,a,c){return b-a*c},isStable:function(){return.00001>Math.abs(this.systemTemperature-this.prevSystemTemperature)||0>=this.temperature},getSystemTemperature:function(){return this.nodes.reduce(function(b,a){return b+a.temperature},0)},
vectorLength:function(b){return Math.sqrt(b.x*b.x+b.y*b.y)},getDistR:function(b,a){b=this.getDistXY(b,a);return this.vectorLength(b)},getDistXY:function(b,a){var c=b.plotX-a.plotX;b=b.plotY-a.plotY;return{x:c,y:b,absX:Math.abs(c),absY:Math.abs(b)}}});e(d,"predraw",function(){this.graphLayoutsLookup&&this.graphLayoutsLookup.forEach(function(b){b.stop()})});e(d,"render",function(){function b(b){b.maxIterations--&&isFinite(b.temperature)&&!b.isStable()&&!b.enableSimulation&&(b.beforeStep&&b.beforeStep(),
b.step(),e=!1,a=!0)}var a=!1;if(this.graphLayoutsLookup){c(!1,this);for(this.graphLayoutsLookup.forEach(function(b){b.start()});!e;){var e=!0;this.graphLayoutsLookup.forEach(b)}a&&this.series.forEach(function(b){b&&b.layout&&b.render()})}});e(d,"beforePrint",function(){this.graphLayoutsLookup&&(this.graphLayoutsLookup.forEach(function(b){b.updateSimulation(!1)}),this.redraw())});e(d,"afterPrint",function(){this.graphLayoutsLookup&&this.graphLayoutsLookup.forEach(function(b){b.updateSimulation()});
this.redraw()})});A(d,"Series/PackedBubble/PackedBubbleComposition.js",[d["Core/Chart/Chart.js"],d["Core/Globals.js"],d["Core/Utilities.js"]],function(d,e,l){var a=e.layouts["reingold-fruchterman"],c=l.addEvent,t=l.extendClass,p=l.pick;d.prototype.getSelectedParentNodes=function(){var a=[];this.series.forEach(function(c){c.parentNode&&c.parentNode.selected&&a.push(c.parentNode)});return a};e.networkgraphIntegrations.packedbubble={repulsiveForceFunction:function(a,c,e,b){return Math.min(a,(e.marker.radius+
b.marker.radius)/2)},barycenter:function(){var a=this,c=a.options.gravitationalConstant,e=a.box,b=a.nodes,d,m;b.forEach(function(g){a.options.splitSeries&&!g.isParentNode?(d=g.series.parentNode.plotX,m=g.series.parentNode.plotY):(d=e.width/2,m=e.height/2);g.fixedPosition||(g.plotX-=(g.plotX-d)*c/(g.mass*Math.sqrt(b.length)),g.plotY-=(g.plotY-m)*c/(g.mass*Math.sqrt(b.length)))})},repulsive:function(a,c,e,b){var g=c*this.diffTemperature/a.mass/a.degree;c=e.x*g;e=e.y*g;a.fixedPosition||(a.plotX+=c,a.plotY+=
e);b.fixedPosition||(b.plotX-=c,b.plotY-=e)},integrate:e.networkgraphIntegrations.verlet.integrate,getK:e.noop};e.layouts.packedbubble=t(a,{beforeStep:function(){this.options.marker&&this.series.forEach(function(a){a&&a.calculateParentRadius()})},isStable:function(){var a=Math.abs(this.prevSystemTemperature-this.systemTemperature);return 1>Math.abs(10*this.systemTemperature/Math.sqrt(this.nodes.length))&&.00001>a||0>=this.temperature},setCircularPositions:function(){var a=this,c=a.box,e=a.nodes,b=
2*Math.PI/(e.length+1),g,d,n=a.options.initialPositionRadius;e.forEach(function(e,k){a.options.splitSeries&&!e.isParentNode?(g=e.series.parentNode.plotX,d=e.series.parentNode.plotY):(g=c.width/2,d=c.height/2);e.plotX=e.prevX=p(e.plotX,g+n*Math.cos(e.index||k*b));e.plotY=e.prevY=p(e.plotY,d+n*Math.sin(e.index||k*b));e.dispX=0;e.dispY=0})},repulsiveForces:function(){var a=this,c,e,b,g=a.options.bubblePadding;a.nodes.forEach(function(d){d.degree=d.mass;d.neighbours=0;a.nodes.forEach(function(k){c=0;
d===k||d.fixedPosition||!a.options.seriesInteraction&&d.series!==k.series||(b=a.getDistXY(d,k),e=a.vectorLength(b)-(d.marker.radius+k.marker.radius+g),0>e&&(d.degree+=.01,d.neighbours++,c=a.repulsiveForce(-e/Math.sqrt(d.neighbours),a.k,d,k)),a.force("repulsive",d,c*k.mass,b,k,e))})})},applyLimitBox:function(c){if(this.options.splitSeries&&!c.isParentNode&&this.options.parentNodeLimit){var e=this.getDistXY(c,c.series.parentNode);var d=c.series.parentNodeRadius-c.marker.radius-this.vectorLength(e);
0>d&&d>-2*c.marker.radius&&(c.plotX-=.01*e.x,c.plotY-=.01*e.y)}a.prototype.applyLimitBox.apply(this,arguments)}});c(d,"beforeRedraw",function(){this.allDataPoints&&delete this.allDataPoints})});A(d,"Series/PackedBubble/PackedBubbleSeries.js",[d["Core/Color/Color.js"],d["Core/Globals.js"],d["Series/PackedBubble/PackedBubblePoint.js"],d["Core/Series/SeriesRegistry.js"],d["Core/Utilities.js"]],function(d,e,l,a,c){var t=this&&this.__extends||function(){var a=function(b,h){a=Object.setPrototypeOf||{__proto__:[]}instanceof
Array&&function(a,b){a.__proto__=b}||function(a,b){for(var h in b)b.hasOwnProperty(h)&&(a[h]=b[h])};return a(b,h)};return function(b,h){function f(){this.constructor=b}a(b,h);b.prototype=null===h?Object.create(h):(f.prototype=h.prototype,new f)}}(),p=d.parse,k=a.series,x=a.seriesTypes.bubble,w=c.addEvent,b=c.clamp,g=c.defined,m=c.extend,n=c.fireEvent,q=c.isArray,H=c.isNumber,A=c.merge,I=c.pick,y=e.dragNodesMixin;d=function(a){function f(){var b=null!==a&&a.apply(this,arguments)||this;b.chart=void 0;
b.data=void 0;b.layout=void 0;b.options=void 0;b.points=void 0;b.xData=void 0;return b}t(f,a);f.prototype.accumulateAllPoints=function(a){var b=a.chart,h=[],f,c;for(f=0;f<b.series.length;f++)if(a=b.series[f],a.is("packedbubble")&&a.visible||!b.options.chart.ignoreHiddenSeries)for(c=0;c<a.yData.length;c++)h.push([null,null,a.yData[c],a.index,c,{id:c,marker:{radius:0}}]);return h};f.prototype.addLayout=function(){var a=this.options.layoutAlgorithm,b=this.chart.graphLayoutsStorage,f=this.chart.graphLayoutsLookup,
c=this.chart.options.chart;b||(this.chart.graphLayoutsStorage=b={},this.chart.graphLayoutsLookup=f=[]);var d=b[a.type];d||(a.enableSimulation=g(c.forExport)?!c.forExport:a.enableSimulation,b[a.type]=d=new e.layouts[a.type],d.init(a),f.splice(d.index,0,d));this.layout=d;this.points.forEach(function(a){a.mass=2;a.degree=1;a.collisionNmb=1});d.setArea(0,0,this.chart.plotWidth,this.chart.plotHeight);d.addElementsToCollection([this],d.series);d.addElementsToCollection(this.points,d.nodes)};f.prototype.addSeriesLayout=
function(){var a=this.options.layoutAlgorithm,b=this.chart.graphLayoutsStorage,f=this.chart.graphLayoutsLookup,c=A(a,a.parentNodeOptions,{enableSimulation:this.layout.options.enableSimulation});var d=b[a.type+"-series"];d||(b[a.type+"-series"]=d=new e.layouts[a.type],d.init(c),f.splice(d.index,0,d));this.parentNodeLayout=d;this.createParentNodes()};f.prototype.calculateParentRadius=function(){var a=this.seriesBox();this.parentNodeRadius=b(Math.sqrt(2*this.parentNodeMass/Math.PI)+20,20,a?Math.max(Math.sqrt(Math.pow(a.width,
2)+Math.pow(a.height,2))/2+20,20):Math.sqrt(2*this.parentNodeMass/Math.PI)+20);this.parentNode&&(this.parentNode.marker.radius=this.parentNode.radius=this.parentNodeRadius)};f.prototype.calculateZExtremes=function(){var a=this.options.zMin,b=this.options.zMax,f=Infinity,c=-Infinity;if(a&&b)return[a,b];this.chart.series.forEach(function(a){a.yData.forEach(function(a){g(a)&&(a>c&&(c=a),a<f&&(f=a))})});a=I(a,f);b=I(b,c);return[a,b]};f.prototype.checkOverlap=function(a,b){var h=a[0]-b[0],f=a[1]-b[1];
return-.001>Math.sqrt(h*h+f*f)-Math.abs(a[2]+b[2])};f.prototype.createParentNodes=function(){var a=this,b=a.chart,f=a.parentNodeLayout,c,e=a.parentNode,d=a.pointClass,g=a.layout.options,k={radius:a.parentNodeRadius,lineColor:a.color,fillColor:p(a.color).brighten(.4).get()};g.parentNodeOptions&&(k=A(g.parentNodeOptions.marker||{},k));a.parentNodeMass=0;a.points.forEach(function(b){a.parentNodeMass+=Math.PI*Math.pow(b.marker.radius,2)});a.calculateParentRadius();f.nodes.forEach(function(b){b.seriesIndex===
a.index&&(c=!0)});f.setArea(0,0,b.plotWidth,b.plotHeight);c||(e||(e=(new d).init(this,{mass:a.parentNodeRadius/2,marker:k,dataLabels:{inside:!1},states:{normal:{marker:k},hover:{marker:k}},dataLabelOnNull:!0,degree:a.parentNodeRadius,isParentNode:!0,seriesIndex:a.index})),a.parentNode&&(e.plotX=a.parentNode.plotX,e.plotY=a.parentNode.plotY),a.parentNode=e,f.addElementsToCollection([a],f.series),f.addElementsToCollection([e],f.nodes))};f.prototype.deferLayout=function(){var a=this.options.layoutAlgorithm;
this.visible&&(this.addLayout(),a.splitSeries&&this.addSeriesLayout())};f.prototype.destroy=function(){this.chart.graphLayoutsLookup&&this.chart.graphLayoutsLookup.forEach(function(a){a.removeElementFromCollection(this,a.series)},this);this.parentNode&&this.parentNodeLayout&&(this.parentNodeLayout.removeElementFromCollection(this.parentNode,this.parentNodeLayout.nodes),this.parentNode.dataLabel&&(this.parentNode.dataLabel=this.parentNode.dataLabel.destroy()));k.prototype.destroy.apply(this,arguments)};
f.prototype.drawDataLabels=function(){var a=this.options.dataLabels.textPath,b=this.points;k.prototype.drawDataLabels.apply(this,arguments);this.parentNode&&(this.parentNode.formatPrefix="parentNode",this.points=[this.parentNode],this.options.dataLabels.textPath=this.options.dataLabels.parentNodeTextPath,k.prototype.drawDataLabels.apply(this,arguments),this.points=b,this.options.dataLabels.textPath=a)};f.prototype.drawGraph=function(){if(this.layout&&this.layout.options.splitSeries){var a=this.chart;
var b=this.layout.options.parentNodeOptions.marker;var f={fill:b.fillColor||p(this.color).brighten(.4).get(),opacity:b.fillOpacity,stroke:b.lineColor||this.color,"stroke-width":I(b.lineWidth,this.options.lineWidth)};this.parentNodesGroup||(this.parentNodesGroup=this.plotGroup("parentNodesGroup","parentNode",this.visible?"inherit":"hidden",.1,a.seriesGroup),this.group.attr({zIndex:2}));this.calculateParentRadius();b=A({x:this.parentNode.plotX-this.parentNodeRadius,y:this.parentNode.plotY-this.parentNodeRadius,
width:2*this.parentNodeRadius,height:2*this.parentNodeRadius},f);this.parentNode.graphic||(this.graph=this.parentNode.graphic=a.renderer.symbol(f.symbol).add(this.parentNodesGroup));this.parentNode.graphic.attr(b)}};f.prototype.drawTracker=function(){var b=this.parentNode;a.prototype.drawTracker.call(this);if(b){var f=q(b.dataLabels)?b.dataLabels:b.dataLabel?[b.dataLabel]:[];b.graphic&&(b.graphic.element.point=b);f.forEach(function(a){a.div?a.div.point=b:a.element.point=b})}};f.prototype.getPointRadius=
function(){var a=this,f=a.chart,c=a.options,e=c.useSimulation,d=Math.min(f.plotWidth,f.plotHeight),g={},k=[],l=f.allDataPoints,n,m,p,z;["minSize","maxSize"].forEach(function(a){var b=parseInt(c[a],10),f=/%$/.test(c[a]);g[a]=f?d*b/100:b*Math.sqrt(l.length)});f.minRadius=n=g.minSize/Math.sqrt(l.length);f.maxRadius=m=g.maxSize/Math.sqrt(l.length);var q=e?a.calculateZExtremes():[n,m];(l||[]).forEach(function(f,h){p=e?b(f[2],q[0],q[1]):f[2];z=a.getRadius(q[0],q[1],n,m,p);0===z&&(z=null);l[h][2]=z;k.push(z)});
a.radii=k};f.prototype.init=function(){k.prototype.init.apply(this,arguments);this.eventsToUnbind.push(w(this,"updatedData",function(){this.chart.series.forEach(function(a){a.type===this.type&&(a.isDirty=!0)},this)}));return this};f.prototype.onMouseUp=function(a){if(a.fixedPosition&&!a.removed){var b,f,h=this.layout,c=this.parentNodeLayout;c&&h.options.dragBetweenSeries&&c.nodes.forEach(function(c){a&&a.marker&&c!==a.series.parentNode&&(b=h.getDistXY(a,c),f=h.vectorLength(b)-c.marker.radius-a.marker.radius,
0>f&&(c.series.addPoint(A(a.options,{plotX:a.plotX,plotY:a.plotY}),!1),h.removeElementFromCollection(a,h.nodes),a.remove()))});y.onMouseUp.apply(this,arguments)}};f.prototype.placeBubbles=function(a){var b=this.checkOverlap,f=this.positionBubble,c=[],h=1,e=0,d=0;var g=[];var k;a=a.sort(function(a,b){return b[2]-a[2]});if(a.length){c.push([[0,0,a[0][2],a[0][3],a[0][4]]]);if(1<a.length)for(c.push([[0,0-a[1][2]-a[0][2],a[1][2],a[1][3],a[1][4]]]),k=2;k<a.length;k++)a[k][2]=a[k][2]||1,g=f(c[h][e],c[h-
1][d],a[k]),b(g,c[h][0])?(c.push([]),d=0,c[h+1].push(f(c[h][e],c[h][0],a[k])),h++,e=0):1<h&&c[h-1][d+1]&&b(g,c[h-1][d+1])?(d++,c[h].push(f(c[h][e],c[h-1][d],a[k])),e++):(e++,c[h].push(g));this.chart.stages=c;this.chart.rawPositions=[].concat.apply([],c);this.resizeRadius();g=this.chart.rawPositions}return g};f.prototype.pointAttribs=function(a,b){var f=this.options,c=f.marker;a&&a.isParentNode&&f.layoutAlgorithm&&f.layoutAlgorithm.parentNodeOptions&&(c=f.layoutAlgorithm.parentNodeOptions.marker);
f=c.fillOpacity;a=k.prototype.pointAttribs.call(this,a,b);1!==f&&(a["fill-opacity"]=f);return a};f.prototype.positionBubble=function(a,b,f){var c=Math.sqrt,h=Math.asin,e=Math.acos,d=Math.pow,g=Math.abs;c=c(d(a[0]-b[0],2)+d(a[1]-b[1],2));e=e((d(c,2)+d(f[2]+b[2],2)-d(f[2]+a[2],2))/(2*(f[2]+b[2])*c));h=h(g(a[0]-b[0])/c);a=(0>a[1]-b[1]?0:Math.PI)+e+h*(0>(a[0]-b[0])*(a[1]-b[1])?1:-1);return[b[0]+(b[2]+f[2])*Math.sin(a),b[1]-(b[2]+f[2])*Math.cos(a),f[2],f[3],f[4]]};f.prototype.render=function(){var a=[];
k.prototype.render.apply(this,arguments);this.options.dataLabels.allowOverlap||(this.data.forEach(function(b){q(b.dataLabels)&&b.dataLabels.forEach(function(b){a.push(b)})}),this.options.useSimulation&&this.chart.hideOverlappingLabels(a))};f.prototype.resizeRadius=function(){var a=this.chart,b=a.rawPositions,f=Math.min,c=Math.max,e=a.plotLeft,d=a.plotTop,g=a.plotHeight,k=a.plotWidth,l,n,m;var p=l=Number.POSITIVE_INFINITY;var z=n=Number.NEGATIVE_INFINITY;for(m=0;m<b.length;m++){var q=b[m][2];p=f(p,
b[m][0]-q);z=c(z,b[m][0]+q);l=f(l,b[m][1]-q);n=c(n,b[m][1]+q)}m=[z-p,n-l];f=f.apply([],[(k-e)/m[0],(g-d)/m[1]]);if(1e-10<Math.abs(f-1)){for(m=0;m<b.length;m++)b[m][2]*=f;this.placeBubbles(b)}else a.diffY=g/2+d-l-(n-l)/2,a.diffX=k/2+e-p-(z-p)/2};f.prototype.seriesBox=function(){var a=this.chart,b=Math.max,f=Math.min,c,e=[a.plotLeft,a.plotLeft+a.plotWidth,a.plotTop,a.plotTop+a.plotHeight];this.data.forEach(function(a){g(a.plotX)&&g(a.plotY)&&a.marker.radius&&(c=a.marker.radius,e[0]=f(e[0],a.plotX-c),
e[1]=b(e[1],a.plotX+c),e[2]=f(e[2],a.plotY-c),e[3]=b(e[3],a.plotY+c))});return H(e.width/e.height)?e:null};f.prototype.setVisible=function(){var a=this;k.prototype.setVisible.apply(a,arguments);a.parentNodeLayout&&a.graph?a.visible?(a.graph.show(),a.parentNode.dataLabel&&a.parentNode.dataLabel.show()):(a.graph.hide(),a.parentNodeLayout.removeElementFromCollection(a.parentNode,a.parentNodeLayout.nodes),a.parentNode.dataLabel&&a.parentNode.dataLabel.hide()):a.layout&&(a.visible?a.layout.addElementsToCollection(a.points,
a.layout.nodes):a.points.forEach(function(b){a.layout.removeElementFromCollection(b,a.layout.nodes)}))};f.prototype.translate=function(){var a=this.chart,b=this.data,f=this.index,c,e=this.options.useSimulation;this.processedXData=this.xData;this.generatePoints();g(a.allDataPoints)||(a.allDataPoints=this.accumulateAllPoints(this),this.getPointRadius());if(e)var d=a.allDataPoints;else d=this.placeBubbles(a.allDataPoints),this.options.draggable=!1;for(c=0;c<d.length;c++)if(d[c][3]===f){var k=b[d[c][4]];
var l=I(d[c][2],void 0);e||(k.plotX=d[c][0]-a.plotLeft+a.diffX,k.plotY=d[c][1]-a.plotTop+a.diffY);H(l)&&(k.marker=m(k.marker,{radius:l,width:2*l,height:2*l}),k.radius=l)}e&&this.deferLayout();n(this,"afterTranslate")};f.defaultOptions=A(x.defaultOptions,{minSize:"10%",maxSize:"50%",sizeBy:"area",zoneAxis:"y",crisp:!1,tooltip:{pointFormat:"Value: {point.value}"},draggable:!0,useSimulation:!0,parentNode:{allowPointSelect:!1},dataLabels:{formatter:function(){var a=this.series.chart.numberFormatter,b=
this.point.value;return H(b)?a(b,-1):""},parentNodeFormatter:function(){return this.name},parentNodeTextPath:{enabled:!0},padding:0,style:{transition:"opacity 2000ms"}},layoutAlgorithm:{initialPositions:"circle",initialPositionRadius:20,bubblePadding:5,parentNodeLimit:!1,seriesInteraction:!0,dragBetweenSeries:!1,parentNodeOptions:{maxIterations:400,gravitationalConstant:.03,maxSpeed:50,initialPositionRadius:100,seriesInteraction:!0,marker:{fillColor:null,fillOpacity:1,lineWidth:null,lineColor:null,
symbol:"circle"}},enableSimulation:!0,type:"packedbubble",integration:"packedbubble",maxIterations:1E3,splitSeries:!1,maxSpeed:5,gravitationalConstant:.01,friction:-.981}});return f}(x);m(d.prototype,{alignDataLabel:k.prototype.alignDataLabel,axisTypes:[],directTouch:!0,forces:["barycenter","repulsive"],hasDraggableNodes:!0,isCartesian:!1,noSharedTooltip:!0,onMouseDown:y.onMouseDown,onMouseMove:y.onMouseMove,pointArrayMap:["value"],pointClass:l,pointValKey:"value",redrawHalo:y.redrawHalo,requireSorting:!1,
searchPoint:e.noop,trackerGroups:["group","dataLabelsGroup","parentNodesGroup"]});a.registerSeriesType("packedbubble",d);"";"";return d});A(d,"Series/Polygon/PolygonSeries.js",[d["Core/Globals.js"],d["Core/Legend/LegendSymbol.js"],d["Core/Series/SeriesRegistry.js"],d["Core/Utilities.js"]],function(d,e,l,a){var c=this&&this.__extends||function(){var a=function(b,c){a=Object.setPrototypeOf||{__proto__:[]}instanceof Array&&function(a,b){a.__proto__=b}||function(a,b){for(var c in b)b.hasOwnProperty(c)&&
(a[c]=b[c])};return a(b,c)};return function(b,c){function e(){this.constructor=b}a(b,c);b.prototype=null===c?Object.create(c):(e.prototype=c.prototype,new e)}}();d=d.noop;var t=l.series,p=l.seriesTypes,k=p.area,x=p.line,w=p.scatter;p=a.extend;var b=a.merge;a=function(a){function e(){var b=null!==a&&a.apply(this,arguments)||this;b.data=void 0;b.options=void 0;b.points=void 0;return b}c(e,a);e.prototype.getGraphPath=function(){for(var a=x.prototype.getGraphPath.call(this),b=a.length+1;b--;)(b===a.length||
"M"===a[b][0])&&0<b&&a.splice(b,0,["Z"]);return this.areaPath=a};e.prototype.drawGraph=function(){this.options.fillColor=this.color;k.prototype.drawGraph.call(this)};e.defaultOptions=b(w.defaultOptions,{marker:{enabled:!1,states:{hover:{enabled:!1}}},stickyTracking:!1,tooltip:{followPointer:!0,pointFormat:""},trackByArea:!0});return e}(w);p(a.prototype,{type:"polygon",drawLegendSymbol:e.drawRectangle,drawTracker:t.prototype.drawTracker,setStackedPoints:d});l.registerSeriesType("polygon",a);"";return a});
A(d,"Core/Axis/WaterfallAxis.js",[d["Extensions/Stacking.js"],d["Core/Utilities.js"]],function(d,e){var l=e.addEvent,a=e.objectEach,c;(function(c){function e(){var a=this.waterfall.stacks;a&&(a.changed=!1,delete a.alreadyChanged)}function k(){var a=this.options.stackLabels;a&&a.enabled&&this.waterfall.stacks&&this.waterfall.renderStackTotals()}function x(){for(var a=this.axes,b=this.series,c=b.length;c--;)b[c].options.stacking&&(a.forEach(function(a){a.isXAxis||(a.waterfall.stacks.changed=!0)}),c=
0)}function w(){this.waterfall||(this.waterfall=new b(this))}var b=function(){function b(a){this.axis=a;this.stacks={changed:!1}}b.prototype.renderStackTotals=function(){var b=this.axis,c=b.waterfall.stacks,e=b.stacking&&b.stacking.stackTotalGroup,g=new d(b,b.options.stackLabels,!1,0,void 0);this.dummyStackItem=g;a(c,function(b){a(b,function(a){g.total=a.stackTotal;a.label&&(g.label=a.label);d.prototype.render.call(g,e);a.label=g.label;delete g.label})});g.total=null};return b}();c.Composition=b;
c.compose=function(a,b){l(a,"init",w);l(a,"afterBuildStacks",e);l(a,"afterRender",k);l(b,"beforeRedraw",x)}})(c||(c={}));return c});A(d,"Series/Waterfall/WaterfallPoint.js",[d["Series/Column/ColumnSeries.js"],d["Core/Series/Point.js"],d["Core/Utilities.js"]],function(d,e,l){var a=this&&this.__extends||function(){var a=function(c,e){a=Object.setPrototypeOf||{__proto__:[]}instanceof Array&&function(a,c){a.__proto__=c}||function(a,c){for(var b in c)c.hasOwnProperty(b)&&(a[b]=c[b])};return a(c,e)};return function(c,
e){function d(){this.constructor=c}a(c,e);c.prototype=null===e?Object.create(e):(d.prototype=e.prototype,new d)}}(),c=l.isNumber;return function(d){function l(){var a=null!==d&&d.apply(this,arguments)||this;a.options=void 0;a.series=void 0;return a}a(l,d);l.prototype.getClassName=function(){var a=e.prototype.getClassName.call(this);this.isSum?a+=" highcharts-sum":this.isIntermediateSum&&(a+=" highcharts-intermediate-sum");return a};l.prototype.isValid=function(){return c(this.y)||this.isSum||!!this.isIntermediateSum};
return l}(d.prototype.pointClass)});A(d,"Series/Waterfall/WaterfallSeries.js",[d["Core/Axis/Axis.js"],d["Core/Chart/Chart.js"],d["Core/Series/SeriesRegistry.js"],d["Core/Utilities.js"],d["Core/Axis/WaterfallAxis.js"],d["Series/Waterfall/WaterfallPoint.js"]],function(d,e,l,a,c,t){var p=this&&this.__extends||function(){var a=function(b,c){a=Object.setPrototypeOf||{__proto__:[]}instanceof Array&&function(a,b){a.__proto__=b}||function(a,b){for(var c in b)b.hasOwnProperty(c)&&(a[c]=b[c])};return a(b,c)};
return function(b,c){function f(){this.constructor=b}a(b,c);b.prototype=null===c?Object.create(c):(f.prototype=c.prototype,new f)}}(),k=l.seriesTypes,x=k.column,w=k.line,b=a.arrayMax,g=a.arrayMin,m=a.correctFloat;k=a.extend;var n=a.isNumber,q=a.merge,A=a.objectEach,J=a.pick;a=function(a){function c(){var b=null!==a&&a.apply(this,arguments)||this;b.chart=void 0;b.data=void 0;b.options=void 0;b.points=void 0;b.stackedYNeg=void 0;b.stackedYPos=void 0;b.stackKey=void 0;b.xData=void 0;b.yAxis=void 0;b.yData=
void 0;return b}p(c,a);c.prototype.generatePoints=function(){var a;x.prototype.generatePoints.apply(this);var b=0;for(a=this.points.length;b<a;b++){var c=this.points[b];var e=this.processedYData[b];if(c.isIntermediateSum||c.isSum)c.y=m(e)}};c.prototype.translate=function(){var a=this.options,b=this.yAxis,c=J(a.minPointLength,5),e=c/2,d=a.threshold||0,g=d,k=d;a=a.stacking;var l=b.waterfall.stacks[this.stackKey];x.prototype.translate.apply(this);for(var m=this.points,p=0;p<m.length;p++){var q=m[p];
var w=this.processedYData[p];var C=q.shapeArgs;if(C&&n(w)){var t=[0,w];var D=q.y;if(a){if(l){t=l[p];if("overlap"===a){var y=t.stackState[t.stateIndex--];y=0<=D?y:y-D;Object.hasOwnProperty.call(t,"absolutePos")&&delete t.absolutePos;Object.hasOwnProperty.call(t,"absoluteNeg")&&delete t.absoluteNeg}else 0<=D?(y=t.threshold+t.posTotal,t.posTotal-=D):(y=t.threshold+t.negTotal,t.negTotal-=D,y-=D),!t.posTotal&&Object.hasOwnProperty.call(t,"absolutePos")&&(t.posTotal=t.absolutePos,delete t.absolutePos),
!t.negTotal&&Object.hasOwnProperty.call(t,"absoluteNeg")&&(t.negTotal=t.absoluteNeg,delete t.absoluteNeg);q.isSum||(t.connectorThreshold=t.threshold+t.stackTotal);b.reversed?(w=0<=D?y-D:y+D,D=y):(w=y,D=y-D);q.below=w<=d;C.y=b.translate(w,!1,!0,!1,!0)||0;C.height=Math.abs(C.y-(b.translate(D,!1,!0,!1,!0)||0));if(D=b.waterfall.dummyStackItem)D.x=p,D.label=l[p].label,D.setOffset(this.pointXOffset||0,this.barW||0,this.stackedYNeg[p],this.stackedYPos[p])}}else y=Math.max(g,g+D)+t[0],C.y=b.translate(y,!1,
!0,!1,!0)||0,q.isSum?(C.y=b.translate(t[1],!1,!0,!1,!0)||0,C.height=Math.min(b.translate(t[0],!1,!0,!1,!0)||0,b.len)-C.y,q.below=t[1]<=d):q.isIntermediateSum?(0<=D?(w=t[1]+k,D=k):(w=k,D=t[1]+k),b.reversed&&(w^=D,D^=w,w^=D),C.y=b.translate(w,!1,!0,!1,!0)||0,C.height=Math.abs(C.y-Math.min(b.translate(D,!1,!0,!1,!0)||0,b.len)),k+=t[1],q.below=w<=d):(C.height=0<w?(b.translate(g,!1,!0,!1,!0)||0)-C.y:(b.translate(g,!1,!0,!1,!0)||0)-(b.translate(g-w,!1,!0,!1,!0)||0),g+=w,q.below=g<d),0>C.height&&(C.y+=C.height,
C.height*=-1);q.plotY=C.y=Math.round(C.y||0)-this.borderWidth%2/2;C.height=Math.max(Math.round(C.height||0),.001);q.yBottom=C.y+C.height;C.height<=c&&!q.isNull?(C.height=c,C.y-=e,q.plotY=C.y,q.minPointLengthOffset=0>q.y?-e:e):(q.isNull&&(C.width=0),q.minPointLengthOffset=0);D=q.plotY+(q.negative?C.height:0);q.below&&(q.plotY+=C.height);q.tooltipPos&&(this.chart.inverted?q.tooltipPos[0]=b.len-D:q.tooltipPos[1]=D)}}};c.prototype.processData=function(b){var c=this.options,e=this.yData,d=c.data,g=e.length,
k=c.threshold||0,l,n,p,q,t;for(t=n=l=p=q=0;t<g;t++){var z=e[t];var w=d&&d[t]?d[t]:{};"sum"===z||w.isSum?e[t]=m(n):"intermediateSum"===z||w.isIntermediateSum?(e[t]=m(l),l=0):(n+=z,l+=z);p=Math.min(n,p);q=Math.max(n,q)}a.prototype.processData.call(this,b);c.stacking||(this.dataMin=p+k,this.dataMax=q)};c.prototype.toYData=function(a){return a.isSum?"sum":a.isIntermediateSum?"intermediateSum":a.y};c.prototype.updateParallelArrays=function(b,c){a.prototype.updateParallelArrays.call(this,b,c);if("sum"===
this.yData[0]||"intermediateSum"===this.yData[0])this.yData[0]=null};c.prototype.pointAttribs=function(a,b){var c=this.options.upColor;c&&!a.options.color&&(a.color=0<a.y?c:null);a=x.prototype.pointAttribs.call(this,a,b);delete a.dashstyle;return a};c.prototype.getGraphPath=function(){return[["M",0,0]]};c.prototype.getCrispPath=function(){var a=this.data,b=this.yAxis,c=a.length,e=Math.round(this.graph.strokeWidth())%2/2,d=Math.round(this.borderWidth)%2/2,g=this.xAxis.reversed,k=this.yAxis.reversed,
l=this.options.stacking,m=[],n;for(n=1;n<c;n++){var p=a[n].shapeArgs;var q=a[n-1];var t=a[n-1].shapeArgs;var w=b.waterfall.stacks[this.stackKey];var x=0<q.y?-t.height:0;w&&t&&p&&(w=w[n-1],l?(w=w.connectorThreshold,x=Math.round(b.translate(w,0,1,0,1)+(k?x:0))-e):x=t.y+q.minPointLengthOffset+d-e,m.push(["M",(t.x||0)+(g?0:t.width||0),x],["L",(p.x||0)+(g?p.width||0:0),x]));t&&m.length&&(!l&&0>q.y&&!k||0<q.y&&k)&&((q=m[m.length-2])&&"number"===typeof q[2]&&(q[2]+=t.height||0),(q=m[m.length-1])&&"number"===
typeof q[2]&&(q[2]+=t.height||0))}return m};c.prototype.drawGraph=function(){w.prototype.drawGraph.call(this);this.graph.attr({d:this.getCrispPath()})};c.prototype.setStackedPoints=function(){function a(a,b,c,f){if(J)for(c;c<J;c++)A.stackState[c]+=f;else A.stackState[0]=a,J=A.stackState.length;A.stackState.push(A.stackState[J-1]+b)}var b=this.options,c=this.yAxis.waterfall.stacks,e=b.threshold,d=e||0,g=d,k=this.stackKey,l=this.xData,m=l.length,n,p,q;this.yAxis.stacking.usePercentage=!1;var t=p=q=
d;if(this.visible||!this.chart.options.chart.ignoreHiddenSeries){var w=c.changed;(n=c.alreadyChanged)&&0>n.indexOf(k)&&(w=!0);c[k]||(c[k]={});n=c[k];for(var x=0;x<m;x++){var y=l[x];if(!n[y]||w)n[y]={negTotal:0,posTotal:0,stackTotal:0,threshold:0,stateIndex:0,stackState:[],label:w&&n[y]?n[y].label:void 0};var A=n[y];var G=this.yData[x];0<=G?A.posTotal+=G:A.negTotal+=G;var F=b.data[x];y=A.absolutePos=A.posTotal;var H=A.absoluteNeg=A.negTotal;A.stackTotal=y+H;var J=A.stackState.length;F&&F.isIntermediateSum?
(a(q,p,0,q),q=p,p=e,d^=g,g^=d,d^=g):F&&F.isSum?(a(e,t,J),d=e):(a(d,G,0,t),F&&(t+=G,p+=G));A.stateIndex++;A.threshold=d;d+=A.stackTotal}c.changed=!1;c.alreadyChanged||(c.alreadyChanged=[]);c.alreadyChanged.push(k)}};c.prototype.getExtremes=function(){var a=this.options.stacking;if(a){var c=this.yAxis;c=c.waterfall.stacks;var e=this.stackedYNeg=[];var d=this.stackedYPos=[];"overlap"===a?A(c[this.stackKey],function(a){e.push(g(a.stackState));d.push(b(a.stackState))}):A(c[this.stackKey],function(a){e.push(a.negTotal+
a.threshold);d.push(a.posTotal+a.threshold)});return{dataMin:g(e),dataMax:b(d)}}return{dataMin:this.dataMin,dataMax:this.dataMax}};c.defaultOptions=q(x.defaultOptions,{dataLabels:{inside:!0},lineWidth:1,lineColor:"#333333",dashStyle:"Dot",borderColor:"#333333",states:{hover:{lineWidthPlus:0}}});return c}(x);k(a.prototype,{getZonesGraphs:w.prototype.getZonesGraphs,pointValKey:"y",showLine:!0,pointClass:t});l.registerSeriesType("waterfall",a);c.compose(d,e);"";return a});A(d,"Extensions/Polar.js",[d["Core/Animation/AnimationUtilities.js"],
d["Core/Chart/Chart.js"],d["Core/Globals.js"],d["Extensions/Pane.js"],d["Core/Pointer.js"],d["Core/Series/Series.js"],d["Core/Series/SeriesRegistry.js"],d["Core/Renderer/SVG/SVGRenderer.js"],d["Core/Utilities.js"]],function(d,e,l,a,c,t,p,k,x){var w=d.animObject;p=p.seriesTypes;var b=x.addEvent,g=x.defined,m=x.find,n=x.isNumber,q=x.pick,A=x.splat,J=x.uniqueKey;d=x.wrap;var I=t.prototype;c=c.prototype;I.searchPointByAngle=function(a){var b=this.chart,c=this.xAxis.pane.center;return this.searchKDTree({clientX:180+
-180/Math.PI*Math.atan2(a.chartX-c[0]-b.plotLeft,a.chartY-c[1]-b.plotTop)})};I.getConnectors=function(a,b,c,e){var f=e?1:0;var d=0<=b&&b<=a.length-1?b:0>b?a.length-1+b:0;b=0>d-1?a.length-(1+f):d-1;f=d+1>a.length-1?f:d+1;var g=a[b];f=a[f];var h=g.plotX;g=g.plotY;var k=f.plotX;var l=f.plotY;f=a[d].plotX;d=a[d].plotY;h=(1.5*f+h)/2.5;g=(1.5*d+g)/2.5;k=(1.5*f+k)/2.5;var r=(1.5*d+l)/2.5;l=Math.sqrt(Math.pow(h-f,2)+Math.pow(g-d,2));var n=Math.sqrt(Math.pow(k-f,2)+Math.pow(r-d,2));h=Math.atan2(g-d,h-f);r=
Math.PI/2+(h+Math.atan2(r-d,k-f))/2;Math.abs(h-r)>Math.PI/2&&(r-=Math.PI);h=f+Math.cos(r)*l;g=d+Math.sin(r)*l;k=f+Math.cos(Math.PI+r)*n;r=d+Math.sin(Math.PI+r)*n;f={rightContX:k,rightContY:r,leftContX:h,leftContY:g,plotX:f,plotY:d};c&&(f.prevPointCont=this.getConnectors(a,b,!1,e));return f};I.toXY=function(a){var b=this.chart,c=this.xAxis,f=this.yAxis,d=a.plotX,e=a.plotY,g=a.series,k=b.inverted,l=a.y,m=k?d:f.len-e;k&&g&&!g.isRadialBar&&(a.plotY=e="number"===typeof l?f.translate(l)||0:0);a.rectPlotX=
d;a.rectPlotY=e;f.center&&(m+=f.center[3]/2);n(e)&&(f=k?f.postTranslate(e,m):c.postTranslate(d,m),a.plotX=a.polarPlotX=f.x-b.plotLeft,a.plotY=a.polarPlotY=f.y-b.plotTop);this.kdByAngle?(b=(d/Math.PI*180+c.pane.options.startAngle)%360,0>b&&(b+=360),a.clientX=b):a.clientX=a.plotX};p.spline&&(d(p.spline.prototype,"getPointSpline",function(a,b,c,d){this.chart.polar?d?(a=this.getConnectors(b,d,!0,this.connectEnds),b=a.prevPointCont&&a.prevPointCont.rightContX,c=a.prevPointCont&&a.prevPointCont.rightContY,
a=["C",n(b)?b:a.plotX,n(c)?c:a.plotY,n(a.leftContX)?a.leftContX:a.plotX,n(a.leftContY)?a.leftContY:a.plotY,a.plotX,a.plotY]):a=["M",c.plotX,c.plotY]:a=a.call(this,b,c,d);return a}),p.areasplinerange&&(p.areasplinerange.prototype.getPointSpline=p.spline.prototype.getPointSpline));b(t,"afterTranslate",function(){var a=this.chart;if(a.polar&&this.xAxis){(this.kdByAngle=a.tooltip&&a.tooltip.shared)?this.searchPoint=this.searchPointByAngle:this.options.findNearestPointBy="xy";if(!this.preventPostTranslate)for(var c=
this.points,d=c.length;d--;)this.toXY(c[d]),!a.hasParallelCoordinates&&!this.yAxis.reversed&&c[d].y<this.yAxis.min&&(c[d].isNull=!0);this.hasClipCircleSetter||(this.hasClipCircleSetter=!!this.eventsToUnbind.push(b(this,"afterRender",function(){if(a.polar){var b=this.yAxis.pane.center;this.clipCircle?this.clipCircle.animate({x:b[0],y:b[1],r:b[2]/2,innerR:b[3]/2}):this.clipCircle=a.renderer.clipCircle(b[0],b[1],b[2]/2,b[3]/2);this.group.clip(this.clipCircle);this.setClip=l.noop}})))}},{order:2});d(p.line.prototype,
"getGraphPath",function(a,b){var c=this,d;if(this.chart.polar){b=b||this.points;for(d=0;d<b.length;d++)if(!b[d].isNull){var e=d;break}if(!1!==this.options.connectEnds&&"undefined"!==typeof e){this.connectEnds=!0;b.splice(b.length,0,b[e]);var f=!0}b.forEach(function(a){"undefined"===typeof a.polarPlotY&&c.toXY(a)})}d=a.apply(this,[].slice.call(arguments,1));f&&b.pop();return d});var y=function(a,b){var c=this,d=this.chart,e=this.options.animation,f=this.group,g=this.markerGroup,h=this.xAxis&&this.xAxis.center,
k=d.plotLeft,n=d.plotTop,m,p,t,x;if(d.polar)if(c.isRadialBar)b||(c.startAngleRad=q(c.translatedThreshold,c.xAxis.startAngleRad),l.seriesTypes.pie.prototype.animate.call(c,b));else{if(d.renderer.isSVG)if(e=w(e),c.is("column")){if(!b){var y=h[3]/2;c.points.forEach(function(a){m=a.graphic;t=(p=a.shapeArgs)&&p.r;x=p&&p.innerR;m&&p&&(m.attr({r:y,innerR:y}),m.animate({r:t,innerR:x},c.options.animation))})}}else b?(a={translateX:h[0]+k,translateY:h[1]+n,scaleX:.001,scaleY:.001},f.attr(a),g&&g.attr(a)):(a=
{translateX:k,translateY:n,scaleX:1,scaleY:1},f.animate(a,e),g&&g.animate(a,e))}else a.call(this,b)};d(I,"animate",y);if(p.column){var z=p.arearange.prototype;p=p.column.prototype;p.polarArc=function(a,b,c,d){var e=this.xAxis.center,f=this.yAxis.len,g=e[3]/2;b=f-b+g;a=f-q(a,f)+g;this.yAxis.reversed&&(0>b&&(b=g),0>a&&(a=g));return{x:e[0],y:e[1],r:b,innerR:a,start:c,end:d}};d(p,"animate",y);d(p,"translate",function(a){var b=this.options,c=b.stacking,d=this.chart,e=this.xAxis,f=this.yAxis,k=f.reversed,
l=f.center,m=e.startAngleRad,p=e.endAngleRad-m;this.preventPostTranslate=!0;a.call(this);if(e.isRadial){a=this.points;e=a.length;var q=f.translate(f.min);var t=f.translate(f.max);b=b.threshold||0;if(d.inverted&&n(b)){var w=f.translate(b);g(w)&&(0>w?w=0:w>p&&(w=p),this.translatedThreshold=w+m)}for(;e--;){b=a[e];var y=b.barX;var z=b.x;var A=b.y;b.shapeType="arc";if(d.inverted){b.plotY=f.translate(A);if(c&&f.stacking){if(A=f.stacking.stacks[(0>A?"-":"")+this.stackKey],this.visible&&A&&A[z]&&!b.isNull){var G=
A[z].points[this.getStackIndicator(void 0,z,this.index).key];var F=f.translate(G[0]);G=f.translate(G[1]);g(F)&&(F=x.clamp(F,0,p))}}else F=w,G=b.plotY;F>G&&(G=[F,F=G][0]);if(!k)if(F<q)F=q;else if(G>t)G=t;else{if(G<q||F>t)F=G=0}else if(G>q)G=q;else if(F<t)F=t;else if(F>q||G<t)F=G=p;f.min>f.max&&(F=G=k?p:0);F+=m;G+=m;l&&(b.barX=y+=l[3]/2);z=Math.max(y,0);A=Math.max(y+b.pointWidth,0);b.shapeArgs={x:l&&l[0],y:l&&l[1],r:A,innerR:z,start:F,end:G};b.opacity=F===G?0:void 0;b.plotY=(g(this.translatedThreshold)&&
(F<this.translatedThreshold?F:G))-m}else F=y+m,b.shapeArgs=this.polarArc(b.yBottom,b.plotY,F,F+b.pointWidth);this.toXY(b);d.inverted?(y=f.postTranslate(b.rectPlotY,y+b.pointWidth/2),b.tooltipPos=[y.x-d.plotLeft,y.y-d.plotTop]):b.tooltipPos=[b.plotX,b.plotY];l&&(b.ttBelow=b.plotY>l[1])}}});p.findAlignments=function(a,b){null===b.align&&(b.align=20<a&&160>a?"left":200<a&&340>a?"right":"center");null===b.verticalAlign&&(b.verticalAlign=45>a||315<a?"bottom":135<a&&225>a?"top":"middle");return b};z&&(z.findAlignments=
p.findAlignments);d(p,"alignDataLabel",function(a,b,c,d,e,g){var f=this.chart,h=q(d.inside,!!this.options.stacking);f.polar?(a=b.rectPlotX/Math.PI*180,f.inverted?(this.forceDL=f.isInsidePlot(b.plotX,Math.round(b.plotY)),h&&b.shapeArgs?(e=b.shapeArgs,e=this.yAxis.postTranslate(((e.start||0)+(e.end||0))/2-this.xAxis.startAngleRad,b.barX+b.pointWidth/2),e={x:e.x-f.plotLeft,y:e.y-f.plotTop}):b.tooltipPos&&(e={x:b.tooltipPos[0],y:b.tooltipPos[1]}),d.align=q(d.align,"center"),d.verticalAlign=q(d.verticalAlign,
"middle")):this.findAlignments&&(d=this.findAlignments(a,d)),I.alignDataLabel.call(this,b,c,d,e,g),this.isRadialBar&&b.shapeArgs&&b.shapeArgs.start===b.shapeArgs.end&&c.hide(!0)):a.call(this,b,c,d,e,g)})}d(c,"getCoordinates",function(a,b){var c=this.chart,d={xAxis:[],yAxis:[]};c.polar?c.axes.forEach(function(a){var e=a.isXAxis,f=a.center;if("colorAxis"!==a.coll){var g=b.chartX-f[0]-c.plotLeft;f=b.chartY-f[1]-c.plotTop;d[e?"xAxis":"yAxis"].push({axis:a,value:a.translate(e?Math.PI-Math.atan2(g,f):Math.sqrt(Math.pow(g,
2)+Math.pow(f,2)),!0)})}}):d=a.call(this,b);return d});k.prototype.clipCircle=function(a,b,c,d){var e=J(),f=this.createElement("clipPath").attr({id:e}).add(this.defs);a=d?this.arc(a,b,c,d,0,2*Math.PI).add(f):this.circle(a,b,c).add(f);a.id=e;a.clipPath=f;return a};b(e,"getAxes",function(){this.pane||(this.pane=[]);this.options.pane=A(this.options.pane);this.options.pane.forEach(function(b){new a(b,this)},this)});b(e,"afterDrawChartBox",function(){this.pane.forEach(function(a){a.render()})});b(t,"afterInit",
function(){var a=this.chart;a.inverted&&a.polar&&(this.isRadialSeries=!0,this.is("column")&&(this.isRadialBar=!0))});d(e.prototype,"get",function(a,b){return m(this.pane||[],function(a){return a.options.id===b})||a.call(this,b)})});A(d,"masters/highcharts-more.src.js",[d["Core/Globals.js"],d["Core/Axis/RadialAxis.js"],d["Series/Bubble/BubbleSeries.js"]],function(d,e,l){e.compose(d.Axis,d.Tick);l.compose(d.Chart,d.Legend,d.Series)})});
//# sourceMappingURL=highcharts-more.js.map
```
