## Supplementary material for "Exploring natural odour landscapes: A case study with implications for human-biting insects": Figure S4: highcharts.js

```
/*
 Highcharts JS v9.3.1 (2021-11-05)

 (c) 2009-2021 Torstein Honsi

 License: www.highcharts.com/license
*/
'use strict';(function(Z,L){"object"===typeof module&&module.exports?(L["default"]=L,module.exports=Z.document?L(Z):L):"function"===typeof define&&define.amd?define("highcharts/highcharts",function(){return L(Z)}):(Z.Highcharts&&Z.Highcharts.error(16,!0),Z.Highcharts=L(Z))})("undefined"!==typeof window?window:this,function(Z){function L(u,a,A,G){u.hasOwnProperty(a)||(u[a]=G.apply(null,A))}var a={};L(a,"Core/Globals.js",[],function(){var u="undefined"!==typeof Z?Z:"undefined"!==typeof window?window:
{},a;(function(a){a.SVG_NS="http://www.w3.org/2000/svg";a.product="Highcharts";a.version="9.3.1";a.win=u;a.doc=a.win.document;a.svg=a.doc&&a.doc.createElementNS&&!!a.doc.createElementNS(a.SVG_NS,"svg").createSVGRect;a.userAgent=a.win.navigator&&a.win.navigator.userAgent||"";a.isChrome=-1!==a.userAgent.indexOf("Chrome");a.isFirefox=-1!==a.userAgent.indexOf("Firefox");a.isMS=/(edge|msie|trident)/i.test(a.userAgent)&&!a.win.opera;a.isSafari=!a.isChrome&&-1!==a.userAgent.indexOf("Safari");a.isTouchDevice=
/(Mobile|Android|Windows Phone)/.test(a.userAgent);a.isWebKit=-1!==a.userAgent.indexOf("AppleWebKit");a.deg2rad=2*Math.PI/360;a.hasBidiBug=a.isFirefox&&4>parseInt(a.userAgent.split("Firefox/")[1],10);a.hasTouch=!!a.win.TouchEvent;a.marginNames=["plotTop","marginRight","marginBottom","plotLeft"];a.noop=function(){};a.supportsPassiveEvents=function(){var u=!1;if(!a.isMS){var t=Object.defineProperty({},"passive",{get:function(){u=!0}});a.win.addEventListener&&a.win.removeEventListener&&(a.win.addEventListener("testPassive",
a.noop,t),a.win.removeEventListener("testPassive",a.noop,t))}return u}();a.charts=[];a.dateFormats={};a.seriesTypes={};a.symbolSizes={};a.chartCount=0})(a||(a={}));"";return a});L(a,"Core/Utilities.js",[a["Core/Globals.js"]],function(a){function u(e,g,c,k){var v=g?"Highcharts error":"Highcharts warning";32===e&&(e=v+": Deprecated member");var d=m(e),E=d?v+" #"+e+": www.highcharts.com/errors/"+e+"/":e.toString();if("undefined"!==typeof k){var J="";d&&(E+="?");r(k,function(b,e){J+="\n - "+e+": "+b;
d&&(E+=encodeURI(e)+"="+encodeURI(b))});E+=J}B(a,"displayError",{chart:c,code:e,message:E,params:k},function(){if(g)throw Error(E);b.console&&-1===u.messages.indexOf(E)&&console.warn(E)});u.messages.push(E)}function A(b,e){var v={};r(b,function(g,c){if(I(b[c],!0)&&!b.nodeType&&e[c])g=A(b[c],e[c]),Object.keys(g).length&&(v[c]=g);else if(I(b[c])||b[c]!==e[c])v[c]=b[c]});return v}function G(b,e){return parseInt(b,e||10)}function x(b){return"string"===typeof b}function C(b){b=Object.prototype.toString.call(b);
return"[object Array]"===b||"[object Array Iterator]"===b}function I(b,e){return!!b&&"object"===typeof b&&(!e||!C(b))}function z(b){return I(b)&&"number"===typeof b.nodeType}function q(b){var e=b&&b.constructor;return!(!I(b,!0)||z(b)||!e||!e.name||"Object"===e.name)}function m(b){return"number"===typeof b&&!isNaN(b)&&Infinity>b&&-Infinity<b}function h(b){return"undefined"!==typeof b&&null!==b}function d(b,e,g){var v;x(e)?h(g)?b.setAttribute(e,g):b&&b.getAttribute&&((v=b.getAttribute(e))||"class"!==
e||(v=b.getAttribute(e+"Name"))):r(e,function(e,v){h(e)?b.setAttribute(v,e):b.removeAttribute(v)});return v}function c(b,e){var v;b||(b={});for(v in e)b[v]=e[v];return b}function l(){for(var b=arguments,e=b.length,g=0;g<e;g++){var c=b[g];if("undefined"!==typeof c&&null!==c)return c}}function f(b,e){a.isMS&&!a.svg&&e&&"undefined"!==typeof e.opacity&&(e.filter="alpha(opacity="+100*e.opacity+")");c(b.style,e)}function w(b,e,g,d,n){b=k.createElement(b);e&&c(b,e);n&&f(b,{padding:"0",border:"none",margin:"0"});
g&&f(b,g);d&&d.appendChild(b);return b}function p(b,e){return 1E14<b?b:parseFloat(b.toPrecision(e||14))}function K(e,g,c){var v=a.getStyle||K;if("width"===g)return g=Math.min(e.offsetWidth,e.scrollWidth),c=e.getBoundingClientRect&&e.getBoundingClientRect().width,c<g&&c>=g-1&&(g=Math.floor(c)),Math.max(0,g-(v(e,"padding-left",!0)||0)-(v(e,"padding-right",!0)||0));if("height"===g)return Math.max(0,Math.min(e.offsetHeight,e.scrollHeight)-(v(e,"padding-top",!0)||0)-(v(e,"padding-bottom",!0)||0));b.getComputedStyle||
u(27,!0);if(e=b.getComputedStyle(e,void 0)){var k=e.getPropertyValue(g);l(c,"opacity"!==g)&&(k=G(k))}return k}function r(b,e,g){for(var v in b)Object.hasOwnProperty.call(b,v)&&e.call(g||b[v],b[v],v,b)}function y(b,e,g){function v(e,M){var J=b.removeEventListener||a.removeEventListenerPolyfill;J&&J.call(b,e,M,!1)}function c(J){var M;if(b.nodeName){if(e){var g={};g[e]=!0}else g=J;r(g,function(b,e){if(J[e])for(M=J[e].length;M--;)v(e,J[e][M].fn)})}}var k="function"===typeof b&&b.prototype||b;if(Object.hasOwnProperty.call(k,
"hcEvents")){var d=k.hcEvents;e?(k=d[e]||[],g?(d[e]=k.filter(function(b){return g!==b.fn}),v(e,g)):(c(d),d[e]=[])):(c(d),delete k.hcEvents)}}function B(b,e,g,d){g=g||{};if(k.createEvent&&(b.dispatchEvent||b.fireEvent&&b!==a)){var v=k.createEvent("Events");v.initEvent(e,!0,!0);g=c(v,g);b.dispatchEvent?b.dispatchEvent(g):b.fireEvent(e,g)}else if(b.hcEvents){g.target||c(g,{preventDefault:function(){g.defaultPrevented=!0},target:b,type:e});v=[];for(var n=b,p=!1;n.hcEvents;)Object.hasOwnProperty.call(n,
"hcEvents")&&n.hcEvents[e]&&(v.length&&(p=!0),v.unshift.apply(v,n.hcEvents[e])),n=Object.getPrototypeOf(n);p&&v.sort(function(b,e){return b.order-e.order});v.forEach(function(e){!1===e.fn.call(b,g)&&g.preventDefault()})}d&&!g.defaultPrevented&&d.call(b,g)}var n=a.charts,k=a.doc,b=a.win;(u||(u={})).messages=[];var g;Math.easeInOutSine=function(b){return-.5*(Math.cos(Math.PI*b)-1)};var e=Array.prototype.find?function(b,e){return b.find(e)}:function(b,e){var g,v=b.length;for(g=0;g<v;g++)if(e(b[g],g))return b[g]};
r({map:"map",each:"forEach",grep:"filter",reduce:"reduce",some:"some"},function(b,e){a[e]=function(g){var v;u(32,!1,void 0,(v={},v["Highcharts."+e]="use Array."+b,v));return Array.prototype[b].apply(g,[].slice.call(arguments,1))}});var D,H=function(){var b=Math.random().toString(36).substring(2,9)+"-",e=0;return function(){return"highcharts-"+(D?"":b)+e++}}();b.jQuery&&(b.jQuery.fn.highcharts=function(){var b=[].slice.call(arguments);if(this[0])return b[0]?(new (a[x(b[0])?b.shift():"Chart"])(this[0],
b[0],b[1]),this):n[d(this[0],"data-highcharts-chart")]});e={addEvent:function(b,e,g,c){void 0===c&&(c={});var v="function"===typeof b&&b.prototype||b;Object.hasOwnProperty.call(v,"hcEvents")||(v.hcEvents={});v=v.hcEvents;a.Point&&b instanceof a.Point&&b.series&&b.series.chart&&(b.series.chart.runTrackerClick=!0);var k=b.addEventListener||a.addEventListenerPolyfill;k&&k.call(b,e,g,a.supportsPassiveEvents?{passive:void 0===c.passive?-1!==e.indexOf("touch"):c.passive,capture:!1}:!1);v[e]||(v[e]=[]);
v[e].push({fn:g,order:"number"===typeof c.order?c.order:Infinity});v[e].sort(function(b,e){return b.order-e.order});return function(){y(b,e,g)}},arrayMax:function(b){for(var e=b.length,g=b[0];e--;)b[e]>g&&(g=b[e]);return g},arrayMin:function(b){for(var e=b.length,g=b[0];e--;)b[e]<g&&(g=b[e]);return g},attr:d,clamp:function(b,e,g){return b>e?b<g?b:g:e},cleanRecursively:A,clearTimeout:function(b){h(b)&&clearTimeout(b)},correctFloat:p,createElement:w,css:f,defined:h,destroyObjectProperties:function(b,
e){r(b,function(g,c){g&&g!==e&&g.destroy&&g.destroy();delete b[c]})},discardElement:function(b){g||(g=w("div"));b&&g.appendChild(b);g.innerHTML=""},erase:function(b,e){for(var g=b.length;g--;)if(b[g]===e){b.splice(g,1);break}},error:u,extend:c,extendClass:function(b,e){var g=function(){};g.prototype=new b;c(g.prototype,e);return g},find:e,fireEvent:B,getMagnitude:function(b){return Math.pow(10,Math.floor(Math.log(b)/Math.LN10))},getNestedProperty:function(e,g){for(e=e.split(".");e.length&&h(g);){var c=
e.shift();if("undefined"===typeof c||"__proto__"===c)return;g=g[c];if(!h(g)||"function"===typeof g||"number"===typeof g.nodeType||g===b)return}return g},getStyle:K,inArray:function(b,e,g){u(32,!1,void 0,{"Highcharts.inArray":"use Array.indexOf"});return e.indexOf(b,g)},isArray:C,isClass:q,isDOMElement:z,isFunction:function(b){return"function"===typeof b},isNumber:m,isObject:I,isString:x,keys:function(b){u(32,!1,void 0,{"Highcharts.keys":"use Object.keys"});return Object.keys(b)},merge:function(){var b,
e=arguments,g={},c=function(b,e){"object"!==typeof b&&(b={});r(e,function(J,g){"__proto__"!==g&&"constructor"!==g&&(!I(J,!0)||q(J)||z(J)?b[g]=e[g]:b[g]=c(b[g]||{},J))});return b};!0===e[0]&&(g=e[1],e=Array.prototype.slice.call(e,2));var k=e.length;for(b=0;b<k;b++)g=c(g,e[b]);return g},normalizeTickInterval:function(b,e,g,c,k){var d=b;g=l(g,1);var v=b/g;e||(e=k?[1,1.2,1.5,2,2.5,3,4,5,6,8,10]:[1,2,2.5,5,10],!1===c&&(1===g?e=e.filter(function(b){return 0===b%1}):.1>=g&&(e=[1/g])));for(c=0;c<e.length&&
!(d=e[c],k&&d*g>=b||!k&&v<=(e[c]+(e[c+1]||e[c]))/2);c++);return d=p(d*g,-Math.round(Math.log(.001)/Math.LN10))},objectEach:r,offset:function(e){var g=k.documentElement;e=e.parentElement||e.parentNode?e.getBoundingClientRect():{top:0,left:0,width:0,height:0};return{top:e.top+(b.pageYOffset||g.scrollTop)-(g.clientTop||0),left:e.left+(b.pageXOffset||g.scrollLeft)-(g.clientLeft||0),width:e.width,height:e.height}},pad:function(b,e,g){return Array((e||2)+1-String(b).replace("-","").length).join(g||"0")+
b},pick:l,pInt:G,relativeLength:function(b,e,g){return/%$/.test(b)?e*parseFloat(b)/100+(g||0):parseFloat(b)},removeEvent:y,splat:function(b){return C(b)?b:[b]},stableSort:function(b,e){var g=b.length,c,k;for(k=0;k<g;k++)b[k].safeI=k;b.sort(function(b,g){c=e(b,g);return 0===c?b.safeI-g.safeI:c});for(k=0;k<g;k++)delete b[k].safeI},syncTimeout:function(b,e,g){if(0<e)return setTimeout(b,e,g);b.call(0,g);return-1},timeUnits:{millisecond:1,second:1E3,minute:6E4,hour:36E5,day:864E5,week:6048E5,month:24192E5,
year:314496E5},uniqueKey:H,useSerialIds:function(b){return D=l(b,D)},wrap:function(b,e,g){var c=b[e];b[e]=function(){var b=Array.prototype.slice.call(arguments),e=arguments,k=this;k.proceed=function(){c.apply(k,arguments.length?arguments:e)};b.unshift(c);b=g.apply(this,b);k.proceed=null;return b}}};"";return e});L(a,"Core/Chart/ChartDefaults.js",[],function(){return{panning:{enabled:!1,type:"x"},styledMode:!1,borderRadius:0,colorCount:10,defaultSeriesType:"line",ignoreHiddenSeries:!0,spacing:[10,
10,15,10],resetZoomButton:{theme:{zIndex:6},position:{align:"right",x:-10,y:10}},zoomBySingleTouch:!1,width:null,height:null,borderColor:"#335cad",backgroundColor:"#ffffff",plotBorderColor:"#cccccc"}});L(a,"Core/Color/Color.js",[a["Core/Globals.js"],a["Core/Utilities.js"]],function(a,t){var u=t.isNumber,G=t.merge,x=t.pInt;t=function(){function t(u){this.rgba=[NaN,NaN,NaN,NaN];this.input=u;var z=a.Color;if(z&&z!==t)return new z(u);if(!(this instanceof t))return new t(u);this.init(u)}t.parse=function(a){return a?
new t(a):t.None};t.prototype.init=function(a){var z;if("object"===typeof a&&"undefined"!==typeof a.stops)this.stops=a.stops.map(function(d){return new t(d[1])});else if("string"===typeof a){this.input=a=t.names[a.toLowerCase()]||a;if("#"===a.charAt(0)){var q=a.length;var m=parseInt(a.substr(1),16);7===q?z=[(m&16711680)>>16,(m&65280)>>8,m&255,1]:4===q&&(z=[(m&3840)>>4|(m&3840)>>8,(m&240)>>4|m&240,(m&15)<<4|m&15,1])}if(!z)for(m=t.parsers.length;m--&&!z;){var h=t.parsers[m];(q=h.regex.exec(a))&&(z=h.parse(q))}}z&&
(this.rgba=z)};t.prototype.get=function(a){var z=this.input,q=this.rgba;if("object"===typeof z&&"undefined"!==typeof this.stops){var m=G(z);m.stops=[].slice.call(m.stops);this.stops.forEach(function(h,d){m.stops[d]=[m.stops[d][0],h.get(a)]});return m}return q&&u(q[0])?"rgb"===a||!a&&1===q[3]?"rgb("+q[0]+","+q[1]+","+q[2]+")":"a"===a?""+q[3]:"rgba("+q.join(",")+")":z};t.prototype.brighten=function(a){var z=this.rgba;if(this.stops)this.stops.forEach(function(m){m.brighten(a)});else if(u(a)&&0!==a)for(var q=
0;3>q;q++)z[q]+=x(255*a),0>z[q]&&(z[q]=0),255<z[q]&&(z[q]=255);return this};t.prototype.setOpacity=function(a){this.rgba[3]=a;return this};t.prototype.tweenTo=function(a,z){var q=this.rgba,m=a.rgba;if(!u(q[0])||!u(m[0]))return a.input||"none";a=1!==m[3]||1!==q[3];return(a?"rgba(":"rgb(")+Math.round(m[0]+(q[0]-m[0])*(1-z))+","+Math.round(m[1]+(q[1]-m[1])*(1-z))+","+Math.round(m[2]+(q[2]-m[2])*(1-z))+(a?","+(m[3]+(q[3]-m[3])*(1-z)):"")+")"};t.names={white:"#ffffff",black:"#000000"};t.parsers=[{regex:/rgba\(\s*([0-9]{1,3})\s*,\s*([0-9]{1,3})\s*,\s*([0-9]{1,3})\s*,\s*([0-9]?(?:\.[0-9]+)?)\s*\)/,
parse:function(a){return[x(a[1]),x(a[2]),x(a[3]),parseFloat(a[4],10)]}},{regex:/rgb\(\s*([0-9]{1,3})\s*,\s*([0-9]{1,3})\s*,\s*([0-9]{1,3})\s*\)/,parse:function(a){return[x(a[1]),x(a[2]),x(a[3]),1]}}];t.None=new t("");return t}();"";return t});L(a,"Core/Color/Palettes.js",[],function(){return{colors:"#7cb5ec #434348 #90ed7d #f7a35c #8085e9 #f15c80 #e4d354 #2b908f #f45b5b #91e8e1".split(" ")}});L(a,"Core/Time.js",[a["Core/Globals.js"],a["Core/Utilities.js"]],function(a,t){var u=a.win,G=t.defined,x=
t.error,C=t.extend,I=t.isObject,z=t.merge,q=t.objectEach,m=t.pad,h=t.pick,d=t.splat,c=t.timeUnits,l=a.isSafari&&u.Intl&&u.Intl.DateTimeFormat.prototype.formatRange,f=a.isSafari&&u.Intl&&!u.Intl.DateTimeFormat.prototype.formatRange;t=function(){function w(c){this.options={};this.variableTimezone=this.useUTC=!1;this.Date=u.Date;this.getTimezoneOffset=this.timezoneOffsetFunction();this.update(c)}w.prototype.get=function(c,d){if(this.variableTimezone||this.timezoneOffset){var p=d.getTime(),l=p-this.getTimezoneOffset(d);
d.setTime(l);c=d["getUTC"+c]();d.setTime(p);return c}return this.useUTC?d["getUTC"+c]():d["get"+c]()};w.prototype.set=function(c,d,r){if(this.variableTimezone||this.timezoneOffset){if("Milliseconds"===c||"Seconds"===c||"Minutes"===c&&0===this.getTimezoneOffset(d)%36E5)return d["setUTC"+c](r);var p=this.getTimezoneOffset(d);p=d.getTime()-p;d.setTime(p);d["setUTC"+c](r);c=this.getTimezoneOffset(d);p=d.getTime()+c;return d.setTime(p)}return this.useUTC||l&&"FullYear"===c?d["setUTC"+c](r):d["set"+c](r)};
w.prototype.update=function(c){var d=h(c&&c.useUTC,!0);this.options=c=z(!0,this.options||{},c);this.Date=c.Date||u.Date||Date;this.timezoneOffset=(this.useUTC=d)&&c.timezoneOffset;this.getTimezoneOffset=this.timezoneOffsetFunction();this.variableTimezone=d&&!(!c.getTimezoneOffset&&!c.timezone)};w.prototype.makeTime=function(c,d,r,l,B,n){if(this.useUTC){var k=this.Date.UTC.apply(0,arguments);var b=this.getTimezoneOffset(k);k+=b;var g=this.getTimezoneOffset(k);b!==g?k+=g-b:b-36E5!==this.getTimezoneOffset(k-
36E5)||f||(k-=36E5)}else k=(new this.Date(c,d,h(r,1),h(l,0),h(B,0),h(n,0))).getTime();return k};w.prototype.timezoneOffsetFunction=function(){var c=this,d=this.options,r=d.moment||u.moment;if(!this.useUTC)return function(c){return 6E4*(new Date(c.toString())).getTimezoneOffset()};if(d.timezone){if(r)return function(c){return 6E4*-r.tz(c,d.timezone).utcOffset()};x(25)}return this.useUTC&&d.getTimezoneOffset?function(c){return 6E4*d.getTimezoneOffset(c.valueOf())}:function(){return 6E4*(c.timezoneOffset||
0)}};w.prototype.dateFormat=function(c,d,r){if(!G(d)||isNaN(d))return a.defaultOptions.lang&&a.defaultOptions.lang.invalidDate||"";c=h(c,"%Y-%m-%d %H:%M:%S");var p=this,l=new this.Date(d),n=this.get("Hours",l),k=this.get("Day",l),b=this.get("Date",l),g=this.get("Month",l),e=this.get("FullYear",l),D=a.defaultOptions.lang,f=D&&D.weekdays,v=D&&D.shortWeekdays;l=C({a:v?v[k]:f[k].substr(0,3),A:f[k],d:m(b),e:m(b,2," "),w:k,b:D.shortMonths[g],B:D.months[g],m:m(g+1),o:g+1,y:e.toString().substr(2,2),Y:e,H:m(n),
k:n,I:m(n%12||12),l:n%12||12,M:m(this.get("Minutes",l)),p:12>n?"AM":"PM",P:12>n?"am":"pm",S:m(l.getSeconds()),L:m(Math.floor(d%1E3),3)},a.dateFormats);q(l,function(b,e){for(;-1!==c.indexOf("%"+e);)c=c.replace("%"+e,"function"===typeof b?b.call(p,d):b)});return r?c.substr(0,1).toUpperCase()+c.substr(1):c};w.prototype.resolveDTLFormat=function(c){return I(c,!0)?c:(c=d(c),{main:c[0],from:c[1],to:c[2]})};w.prototype.getTimeTicks=function(d,l,r,f){var p=this,n=[],k={},b=new p.Date(l),g=d.unitRange,e=d.count||
1,D;f=h(f,1);if(G(l)){p.set("Milliseconds",b,g>=c.second?0:e*Math.floor(p.get("Milliseconds",b)/e));g>=c.second&&p.set("Seconds",b,g>=c.minute?0:e*Math.floor(p.get("Seconds",b)/e));g>=c.minute&&p.set("Minutes",b,g>=c.hour?0:e*Math.floor(p.get("Minutes",b)/e));g>=c.hour&&p.set("Hours",b,g>=c.day?0:e*Math.floor(p.get("Hours",b)/e));g>=c.day&&p.set("Date",b,g>=c.month?1:Math.max(1,e*Math.floor(p.get("Date",b)/e)));if(g>=c.month){p.set("Month",b,g>=c.year?0:e*Math.floor(p.get("Month",b)/e));var H=p.get("FullYear",
b)}g>=c.year&&p.set("FullYear",b,H-H%e);g===c.week&&(H=p.get("Day",b),p.set("Date",b,p.get("Date",b)-H+f+(H<f?-7:0)));H=p.get("FullYear",b);f=p.get("Month",b);var v=p.get("Date",b),w=p.get("Hours",b);l=b.getTime();!p.variableTimezone&&p.useUTC||!G(r)||(D=r-l>4*c.month||p.getTimezoneOffset(l)!==p.getTimezoneOffset(r));l=b.getTime();for(b=1;l<r;)n.push(l),l=g===c.year?p.makeTime(H+b*e,0):g===c.month?p.makeTime(H,f+b*e):!D||g!==c.day&&g!==c.week?D&&g===c.hour&&1<e?p.makeTime(H,f,v,w+b*e):l+g*e:p.makeTime(H,
f,v+b*e*(g===c.day?1:7)),b++;n.push(l);g<=c.hour&&1E4>n.length&&n.forEach(function(b){0===b%18E5&&"000000000"===p.dateFormat("%H%M%S%L",b)&&(k[b]="day")})}n.info=C(d,{higherRanks:k,totalRange:g*e});return n};w.prototype.getDateFormat=function(d,l,f,h){var p=this.dateFormat("%m-%d %H:%M:%S.%L",l),n={millisecond:15,second:12,minute:9,hour:6,day:3},k="millisecond";for(b in c){if(d===c.week&&+this.dateFormat("%w",l)===f&&"00:00:00.000"===p.substr(6)){var b="week";break}if(c[b]>d){b=k;break}if(n[b]&&p.substr(n[b])!==
"01-01 00:00:00.000".substr(n[b]))break;"week"!==b&&(k=b)}if(b)var g=this.resolveDTLFormat(h[b]).main;return g};return w}();"";return t});L(a,"Core/DefaultOptions.js",[a["Core/Chart/ChartDefaults.js"],a["Core/Color/Color.js"],a["Core/Globals.js"],a["Core/Color/Palettes.js"],a["Core/Time.js"],a["Core/Utilities.js"]],function(a,t,A,G,x,C){t=t.parse;var u=C.merge,z={colors:G.colors,symbols:["circle","diamond","square","triangle","triangle-down"],lang:{loading:"Loading...",months:"January February March April May June July August September October November December".split(" "),
shortMonths:"Jan Feb Mar Apr May Jun Jul Aug Sep Oct Nov Dec".split(" "),weekdays:"Sunday Monday Tuesday Wednesday Thursday Friday Saturday".split(" "),decimalPoint:".",numericSymbols:"kMGTPE".split(""),resetZoom:"Reset zoom",resetZoomTitle:"Reset zoom level 1:1",thousandsSep:" "},global:{},time:{Date:void 0,getTimezoneOffset:void 0,timezone:void 0,timezoneOffset:0,useUTC:!0},chart:a,title:{text:"Chart title",align:"center",margin:15,widthAdjust:-44},subtitle:{text:"",align:"center",widthAdjust:-44},
caption:{margin:15,text:"",align:"left",verticalAlign:"bottom"},plotOptions:{},labels:{style:{position:"absolute",color:"#333333"}},legend:{enabled:!0,align:"center",alignColumns:!0,className:"highcharts-no-tooltip",layout:"horizontal",labelFormatter:function(){return this.name},borderColor:"#999999",borderRadius:0,navigation:{activeColor:"#003399",inactiveColor:"#cccccc"},itemStyle:{color:"#333333",cursor:"pointer",fontSize:"12px",fontWeight:"bold",textOverflow:"ellipsis"},itemHoverStyle:{color:"#000000"},
itemHiddenStyle:{color:"#cccccc"},shadow:!1,itemCheckboxStyle:{position:"absolute",width:"13px",height:"13px"},squareSymbol:!0,symbolPadding:5,verticalAlign:"bottom",x:0,y:0,title:{style:{fontWeight:"bold"}}},loading:{labelStyle:{fontWeight:"bold",position:"relative",top:"45%"},style:{position:"absolute",backgroundColor:"#ffffff",opacity:.5,textAlign:"center"}},tooltip:{enabled:!0,animation:A.svg,borderRadius:3,dateTimeLabelFormats:{millisecond:"%A, %b %e, %H:%M:%S.%L",second:"%A, %b %e, %H:%M:%S",
minute:"%A, %b %e, %H:%M",hour:"%A, %b %e, %H:%M",day:"%A, %b %e, %Y",week:"Week from %A, %b %e, %Y",month:"%B %Y",year:"%Y"},footerFormat:"",headerShape:"callout",hideDelay:500,padding:8,shape:"callout",shared:!1,snap:A.isTouchDevice?25:10,headerFormat:'<span style="font-size: 10px">{point.key}</span><br/>',pointFormat:'<span style="color:{point.color}">\u25cf</span> {series.name}: <b>{point.y}</b><br/>',backgroundColor:t("#f7f7f7").setOpacity(.85).get(),borderWidth:1,shadow:!0,stickOnContact:!1,
style:{color:"#333333",cursor:"default",fontSize:"12px",whiteSpace:"nowrap"},useHTML:!1},credits:{enabled:!0,href:"https://www.highcharts.com?credits",position:{align:"right",x:-10,verticalAlign:"bottom",y:-5},style:{cursor:"pointer",color:"#999999",fontSize:"9px"},text:"Highcharts.com"}};z.chart.styledMode=!1;"";var q=new x(u(z.global,z.time));a={defaultOptions:z,defaultTime:q,getOptions:function(){return z},setOptions:function(m){u(!0,z,m);if(m.time||m.global)A.time?A.time.update(u(z.global,z.time,
m.global,m.time)):A.time=q;return z}};"";return a});L(a,"Core/Animation/Fx.js",[a["Core/Color/Color.js"],a["Core/Globals.js"],a["Core/Utilities.js"]],function(a,t,A){var u=a.parse,x=t.win,C=A.isNumber,I=A.objectEach;return function(){function a(a,m,h){this.pos=NaN;this.options=m;this.elem=a;this.prop=h}a.prototype.dSetter=function(){var a=this.paths,m=a&&a[0];a=a&&a[1];var h=this.now||0,d=[];if(1!==h&&m&&a)if(m.length===a.length&&1>h)for(var c=0;c<a.length;c++){for(var l=m[c],f=a[c],w=[],p=0;p<f.length;p++){var K=
l[p],r=f[p];C(K)&&C(r)&&("A"!==f[0]||4!==p&&5!==p)?w[p]=K+h*(r-K):w[p]=r}d.push(w)}else d=a;else d=this.toD||[];this.elem.attr("d",d,void 0,!0)};a.prototype.update=function(){var a=this.elem,m=this.prop,h=this.now,d=this.options.step;if(this[m+"Setter"])this[m+"Setter"]();else a.attr?a.element&&a.attr(m,h,null,!0):a.style[m]=h+this.unit;d&&d.call(a,h,this)};a.prototype.run=function(q,m,h){var d=this,c=d.options,l=function(c){return l.stopped?!1:d.step(c)},f=x.requestAnimationFrame||function(c){setTimeout(c,
13)},w=function(){for(var c=0;c<a.timers.length;c++)a.timers[c]()||a.timers.splice(c--,1);a.timers.length&&f(w)};q!==m||this.elem["forceAnimate:"+this.prop]?(this.startTime=+new Date,this.start=q,this.end=m,this.unit=h,this.now=this.start,this.pos=0,l.elem=this.elem,l.prop=this.prop,l()&&1===a.timers.push(l)&&f(w)):(delete c.curAnim[this.prop],c.complete&&0===Object.keys(c.curAnim).length&&c.complete.call(this.elem))};a.prototype.step=function(a){var m=+new Date,h=this.options,d=this.elem,c=h.complete,
l=h.duration,f=h.curAnim;if(d.attr&&!d.element)a=!1;else if(a||m>=l+this.startTime){this.now=this.end;this.pos=1;this.update();var w=f[this.prop]=!0;I(f,function(c){!0!==c&&(w=!1)});w&&c&&c.call(d);a=!1}else this.pos=h.easing((m-this.startTime)/l),this.now=this.start+(this.end-this.start)*this.pos,this.update(),a=!0;return a};a.prototype.initPath=function(a,m,h){function d(c,d){for(;c.length<y;){var k=c[0],b=d[y-c.length];b&&"M"===k[0]&&(c[0]="C"===b[0]?["C",k[1],k[2],k[1],k[2],k[1],k[2]]:["L",k[1],
k[2]]);c.unshift(k);w&&(k=c.pop(),c.push(c[c.length-1],k))}}function c(c,d){for(;c.length<y;)if(d=c[Math.floor(c.length/p)-1].slice(),"C"===d[0]&&(d[1]=d[5],d[2]=d[6]),w){var k=c[Math.floor(c.length/p)].slice();c.splice(c.length/2,0,d,k)}else c.push(d)}var l=a.startX,f=a.endX;h=h.slice();var w=a.isArea,p=w?2:1;m=m&&m.slice();if(!m)return[h,h];if(l&&f&&f.length){for(a=0;a<l.length;a++)if(l[a]===f[0]){var K=a;break}else if(l[0]===f[f.length-l.length+a]){K=a;var r=!0;break}else if(l[l.length-1]===f[f.length-
l.length+a]){K=l.length-a;break}"undefined"===typeof K&&(m=[])}if(m.length&&C(K)){var y=h.length+K*p;r?(d(m,h),c(h,m)):(d(h,m),c(m,h))}return[m,h]};a.prototype.fillSetter=function(){a.prototype.strokeSetter.apply(this,arguments)};a.prototype.strokeSetter=function(){this.elem.attr(this.prop,u(this.start).tweenTo(u(this.end),this.pos),null,!0)};a.timers=[];return a}()});L(a,"Core/Animation/AnimationUtilities.js",[a["Core/Animation/Fx.js"],a["Core/Utilities.js"]],function(a,t){function u(c){return q(c)?
m({duration:500,defer:0},c):{duration:c?500:0,defer:0}}function G(c,d){for(var l=a.timers.length;l--;)a.timers[l].elem!==c||d&&d!==a.timers[l].prop||(a.timers[l].stopped=!0)}var x=t.defined,C=t.getStyle,I=t.isArray,z=t.isNumber,q=t.isObject,m=t.merge,h=t.objectEach,d=t.pick;return{animate:function(c,d,f){var l,p="",K,r;if(!q(f)){var y=arguments;f={duration:y[2],easing:y[3],complete:y[4]}}z(f.duration)||(f.duration=400);f.easing="function"===typeof f.easing?f.easing:Math[f.easing]||Math.easeInOutSine;
f.curAnim=m(d);h(d,function(h,n){G(c,n);r=new a(c,f,n);K=void 0;"d"===n&&I(d.d)?(r.paths=r.initPath(c,c.pathArray,d.d),r.toD=d.d,l=0,K=1):c.attr?l=c.attr(n):(l=parseFloat(C(c,n))||0,"opacity"!==n&&(p="px"));K||(K=h);"string"===typeof K&&K.match("px")&&(K=K.replace(/px/g,""));r.run(l,K,p)})},animObject:u,getDeferredAnimation:function(c,d,f){var l=u(d),p=0,a=0;(f?[f]:c.series).forEach(function(c){c=u(c.options.animation);p=d&&x(d.defer)?l.defer:Math.max(p,c.duration+c.defer);a=Math.min(l.duration,c.duration)});
c.renderer.forExport&&(p=0);return{defer:Math.max(0,p-a),duration:Math.min(p,a)}},setAnimation:function(c,l){l.renderer.globalAnimation=d(c,l.options.chart.animation,!0)},stop:G}});L(a,"Core/Renderer/HTML/AST.js",[a["Core/Globals.js"],a["Core/Utilities.js"]],function(a,t){var u=a.SVG_NS,G=t.attr,x=t.createElement,C=t.discardElement,I=t.error,z=t.isString,q=t.objectEach,m=t.splat;try{var h=!!(new DOMParser).parseFromString("","text/html")}catch(d){h=!1}t=function(){function d(c){this.nodes="string"===
typeof c?this.parseMarkup(c):c}d.filterUserAttributes=function(c){q(c,function(l,f){var a=!0;-1===d.allowedAttributes.indexOf(f)&&(a=!1);-1!==["background","dynsrc","href","lowsrc","src"].indexOf(f)&&(a=z(l)&&d.allowedReferences.some(function(c){return 0===l.indexOf(c)}));a||(I("Highcharts warning: Invalid attribute '"+f+"' in config"),delete c[f])});return c};d.setElementHTML=function(c,l){c.innerHTML="";l&&(new d(l)).addToDOM(c)};d.prototype.addToDOM=function(c){function l(c,h){var p;m(c).forEach(function(c){var f=
c.tagName,w=c.textContent?a.doc.createTextNode(c.textContent):void 0;if(f)if("#text"===f)var m=w;else if(-1!==d.allowedTags.indexOf(f)){f=a.doc.createElementNS("svg"===f?u:h.namespaceURI||u,f);var n=c.attributes||{};q(c,function(c,b){"tagName"!==b&&"attributes"!==b&&"children"!==b&&"textContent"!==b&&(n[b]=c)});G(f,d.filterUserAttributes(n));w&&f.appendChild(w);l(c.children||[],f);m=f}else I("Highcharts warning: Invalid tagName '"+f+"' in config");m&&h.appendChild(m);p=m});return p}return l(this.nodes,
c)};d.prototype.parseMarkup=function(c){var d=[];c=c.trim();if(h)c=(new DOMParser).parseFromString(c,"text/html");else{var f=x("div");f.innerHTML=c;c={body:f}}var a=function(c,d){var l=c.nodeName.toLowerCase(),p={tagName:l};"#text"===l&&(p.textContent=c.textContent||"");if(l=c.attributes){var f={};[].forEach.call(l,function(c){f[c.name]=c.value});p.attributes=f}if(c.childNodes.length){var n=[];[].forEach.call(c.childNodes,function(c){a(c,n)});n.length&&(p.children=n)}d.push(p)};[].forEach.call(c.body.childNodes,
function(c){return a(c,d)});f&&C(f);return d};d.allowedAttributes="aria-controls aria-describedby aria-expanded aria-haspopup aria-hidden aria-label aria-labelledby aria-live aria-pressed aria-readonly aria-roledescription aria-selected class clip-path color colspan cx cy d dx dy disabled fill height href id in markerHeight markerWidth offset opacity orient padding paddingLeft paddingRight patternUnits r refX refY role scope slope src startOffset stdDeviation stroke stroke-linecap stroke-width style tableValues result rowspan summary target tabindex text-align textAnchor textLength title type valign width x x1 x2 y y1 y2 zIndex".split(" ");
d.allowedReferences="https:// http:// mailto: / ../ ./ #".split(" ");d.allowedTags="a abbr b br button caption circle clipPath code dd defs div dl dt em feComponentTransfer feFuncA feFuncB feFuncG feFuncR feGaussianBlur feOffset feMerge feMergeNode filter h1 h2 h3 h4 h5 h6 hr i img li linearGradient marker ol p path pattern pre rect small span stop strong style sub sup svg table text thead tbody tspan td th tr u ul #text".split(" ");return d}();"";return t});L(a,"Core/FormatUtilities.js",[a["Core/DefaultOptions.js"],
a["Core/Utilities.js"]],function(a,t){function u(a,h,d,c){a=+a||0;h=+h;var l=G.lang,f=(a.toString().split(".")[1]||"").split("e")[0].length,w=a.toString().split("e"),p=h;if(-1===h)h=Math.min(f,20);else if(!I(h))h=2;else if(h&&w[1]&&0>w[1]){var m=h+ +w[1];0<=m?(w[0]=(+w[0]).toExponential(m).split("e")[0],h=m):(w[0]=w[0].split(".")[0]||0,a=20>h?(w[0]*Math.pow(10,w[1])).toFixed(h):0,w[1]=0)}m=(Math.abs(w[1]?w[0]:a)+Math.pow(10,-Math.max(h,f)-1)).toFixed(h);f=String(q(m));var r=3<f.length?f.length%3:
0;d=z(d,l.decimalPoint);c=z(c,l.thousandsSep);a=(0>a?"-":"")+(r?f.substr(0,r)+c:"");a=0>+w[1]&&!p?"0":a+f.substr(r).replace(/(\d{3})(?=\d)/g,"$1"+c);h&&(a+=d+m.slice(-h));w[1]&&0!==+a&&(a+="e"+w[1]);return a}var G=a.defaultOptions,x=a.defaultTime,C=t.getNestedProperty,I=t.isNumber,z=t.pick,q=t.pInt;return{dateFormat:function(a,h,d){return x.dateFormat(a,h,d)},format:function(a,h,d){var c="{",l=!1,f=/f$/,w=/\.([0-9])/,p=G.lang,m=d&&d.time||x;d=d&&d.numberFormatter||u;for(var r=[];a;){var y=a.indexOf(c);
if(-1===y)break;var B=a.slice(0,y);if(l){B=B.split(":");c=C(B.shift()||"",h);if(B.length&&"number"===typeof c)if(B=B.join(":"),f.test(B)){var n=parseInt((B.match(w)||["","-1"])[1],10);null!==c&&(c=d(c,n,p.decimalPoint,-1<B.indexOf(",")?p.thousandsSep:""))}else c=m.dateFormat(B,c);r.push(c)}else r.push(B);a=a.slice(y+1);c=(l=!l)?"}":"{"}r.push(a);return r.join("")},numberFormat:u}});L(a,"Core/Renderer/RendererUtilities.js",[a["Core/Utilities.js"]],function(a){var u=a.clamp,A=a.pick,G=a.stableSort,
x;(function(a){function t(a,q,m){var h=a,d=h.reducedLen||q,c=function(c,d){return(d.rank||0)-(c.rank||0)},l=function(c,d){return c.target-d.target},f,w=!0,p=[],K=0;for(f=a.length;f--;)K+=a[f].size;if(K>d){G(a,c);for(K=f=0;K<=d;)K+=a[f].size,f++;p=a.splice(f-1,a.length)}G(a,l);for(a=a.map(function(c){return{size:c.size,targets:[c.target],align:A(c.align,.5)}});w;){for(f=a.length;f--;)d=a[f],c=(Math.min.apply(0,d.targets)+Math.max.apply(0,d.targets))/2,d.pos=u(c-d.size*d.align,0,q-d.size);f=a.length;
for(w=!1;f--;)0<f&&a[f-1].pos+a[f-1].size>a[f].pos&&(a[f-1].size+=a[f].size,a[f-1].targets=a[f-1].targets.concat(a[f].targets),a[f-1].align=.5,a[f-1].pos+a[f-1].size>q&&(a[f-1].pos=q-a[f-1].size),a.splice(f,1),w=!0)}h.push.apply(h,p);f=0;a.some(function(c){var d=0;return(c.targets||[]).some(function(){h[f].pos=c.pos+d;if("undefined"!==typeof m&&Math.abs(h[f].pos-h[f].target)>m)return h.slice(0,f+1).forEach(function(c){return delete c.pos}),h.reducedLen=(h.reducedLen||q)-.1*q,h.reducedLen>.1*q&&t(h,
q,m),!0;d+=h[f].size;f++;return!1})});G(h,l);return h}a.distribute=t})(x||(x={}));return x});L(a,"Core/Renderer/SVG/SVGElement.js",[a["Core/Animation/AnimationUtilities.js"],a["Core/Renderer/HTML/AST.js"],a["Core/Color/Color.js"],a["Core/Globals.js"],a["Core/Utilities.js"]],function(a,t,A,G,x){var u=a.animate,I=a.animObject,z=a.stop,q=G.deg2rad,m=G.doc,h=G.noop,d=G.svg,c=G.SVG_NS,l=G.win,f=x.addEvent,w=x.attr,p=x.createElement,K=x.css,r=x.defined,y=x.erase,B=x.extend,n=x.fireEvent,k=x.isArray,b=x.isFunction,
g=x.isNumber,e=x.isString,D=x.merge,H=x.objectEach,v=x.pick,E=x.pInt,O=x.syncTimeout,S=x.uniqueKey;a=function(){function a(){this.element=void 0;this.onEvents={};this.opacity=1;this.renderer=void 0;this.SVG_NS=c;this.symbolCustomAttribs="x y width height r start end innerR anchorX anchorY rounded".split(" ")}a.prototype._defaultGetter=function(b){b=v(this[b+"Value"],this[b],this.element?this.element.getAttribute(b):null,0);/^[\-0-9\.]+$/.test(b)&&(b=parseFloat(b));return b};a.prototype._defaultSetter=
function(b,e,c){c.setAttribute(e,b)};a.prototype.add=function(b){var e=this.renderer,c=this.element;b&&(this.parentGroup=b);this.parentInverted=b&&b.inverted;"undefined"!==typeof this.textStr&&"text"===this.element.nodeName&&e.buildText(this);this.added=!0;if(!b||b.handleZ||this.zIndex)var g=this.zIndexSetter();g||(b?b.element:e.box).appendChild(c);if(this.onAdd)this.onAdd();return this};a.prototype.addClass=function(b,e){var c=e?"":this.attr("class")||"";b=(b||"").split(/ /g).reduce(function(b,e){-1===
c.indexOf(e)&&b.push(e);return b},c?[c]:[]).join(" ");b!==c&&this.attr("class",b);return this};a.prototype.afterSetters=function(){this.doTransform&&(this.updateTransform(),this.doTransform=!1)};a.prototype.align=function(b,c,g){var J={},d=this.renderer,a=d.alignedObjects,k,n,F;if(b){if(this.alignOptions=b,this.alignByTranslate=c,!g||e(g))this.alignTo=k=g||"renderer",y(a,this),a.push(this),g=void 0}else b=this.alignOptions,c=this.alignByTranslate,k=this.alignTo;g=v(g,d[k],"scrollablePlotBox"===k?
d.plotBox:void 0,d);k=b.align;var D=b.verticalAlign;d=(g.x||0)+(b.x||0);a=(g.y||0)+(b.y||0);"right"===k?n=1:"center"===k&&(n=2);n&&(d+=(g.width-(b.width||0))/n);J[c?"translateX":"x"]=Math.round(d);"bottom"===D?F=1:"middle"===D&&(F=2);F&&(a+=(g.height-(b.height||0))/F);J[c?"translateY":"y"]=Math.round(a);this[this.placed?"animate":"attr"](J);this.placed=!0;this.alignAttr=J;return this};a.prototype.alignSetter=function(b){var e={left:"start",center:"middle",right:"end"};e[b]&&(this.alignValue=b,this.element.setAttribute("text-anchor",
e[b]))};a.prototype.animate=function(b,e,c){var g=this,J=I(v(e,this.renderer.globalAnimation,!0));e=J.defer;v(m.hidden,m.msHidden,m.webkitHidden,!1)&&(J.duration=0);0!==J.duration?(c&&(J.complete=c),O(function(){g.element&&u(g,b,J)},e)):(this.attr(b,void 0,c),H(b,function(b,e){J.step&&J.step.call(this,b,{prop:e,pos:1,elem:this})},this));return this};a.prototype.applyTextOutline=function(b){var e=this.element;-1!==b.indexOf("contrast")&&(b=b.replace(/contrast/g,this.renderer.getContrast(e.style.fill)));
var g=b.split(" ");b=g[g.length-1];if((g=g[0])&&"none"!==g&&G.svg){this.fakeTS=!0;this.ySetter=this.xSetter;g=g.replace(/(^[\d\.]+)(.*?)$/g,function(b,e,c){return 2*Number(e)+c});this.removeTextOutline();var d=m.createElementNS(c,"tspan");w(d,{"class":"highcharts-text-outline",fill:b,stroke:b,"stroke-width":g,"stroke-linejoin":"round"});[].forEach.call(e.childNodes,function(b){var e=b.cloneNode(!0);e.removeAttribute&&["fill","stroke","stroke-width","stroke"].forEach(function(b){return e.removeAttribute(b)});
d.appendChild(e)});var a=m.createElementNS(c,"tspan");a.textContent="\u200b";["x","y"].forEach(function(b){var c=e.getAttribute(b);c&&a.setAttribute(b,c)});d.appendChild(a);e.insertBefore(d,e.firstChild)}};a.prototype.attr=function(b,e,c,g){var d=this.element,J=this.symbolCustomAttribs,a,M=this,F,k;if("string"===typeof b&&"undefined"!==typeof e){var n=b;b={};b[n]=e}"string"===typeof b?M=(this[b+"Getter"]||this._defaultGetter).call(this,b,d):(H(b,function(e,c){F=!1;g||z(this,c);this.symbolName&&-1!==
J.indexOf(c)&&(a||(this.symbolAttr(b),a=!0),F=!0);!this.rotation||"x"!==c&&"y"!==c||(this.doTransform=!0);F||(k=this[c+"Setter"]||this._defaultSetter,k.call(this,e,c,d),!this.styledMode&&this.shadows&&/^(width|height|visibility|x|y|d|transform|cx|cy|r)$/.test(c)&&this.updateShadows(c,e,k))},this),this.afterSetters());c&&c.call(this);return M};a.prototype.clip=function(b){return this.attr("clip-path",b?"url("+this.renderer.url+"#"+b.id+")":"none")};a.prototype.crisp=function(b,e){e=e||b.strokeWidth||
0;var c=Math.round(e)%2/2;b.x=Math.floor(b.x||this.x||0)+c;b.y=Math.floor(b.y||this.y||0)+c;b.width=Math.floor((b.width||this.width||0)-2*c);b.height=Math.floor((b.height||this.height||0)-2*c);r(b.strokeWidth)&&(b.strokeWidth=e);return b};a.prototype.complexColor=function(b,e,c){var g=this.renderer,d,J,a,v,F,l,p,f,h,w,E=[],m;n(this.renderer,"complexColor",{args:arguments},function(){b.radialGradient?J="radialGradient":b.linearGradient&&(J="linearGradient");if(J){a=b[J];F=g.gradients;l=b.stops;h=c.radialReference;
k(a)&&(b[J]=a={x1:a[0],y1:a[1],x2:a[2],y2:a[3],gradientUnits:"userSpaceOnUse"});"radialGradient"===J&&h&&!r(a.gradientUnits)&&(v=a,a=D(a,g.getRadialAttr(h,v),{gradientUnits:"userSpaceOnUse"}));H(a,function(b,e){"id"!==e&&E.push(e,b)});H(l,function(b){E.push(b)});E=E.join(",");if(F[E])w=F[E].attr("id");else{a.id=w=S();var M=F[E]=g.createElement(J).attr(a).add(g.defs);M.radAttr=v;M.stops=[];l.forEach(function(b){0===b[1].indexOf("rgba")?(d=A.parse(b[1]),p=d.get("rgb"),f=d.get("a")):(p=b[1],f=1);b=g.createElement("stop").attr({offset:b[0],
"stop-color":p,"stop-opacity":f}).add(M);M.stops.push(b)})}m="url("+g.url+"#"+w+")";c.setAttribute(e,m);c.gradient=E;b.toString=function(){return m}}})};a.prototype.css=function(b){var e=this.styles,c={},g=this.element,a=["textOutline","textOverflow","width"],k="",n=!e;b&&b.color&&(b.fill=b.color);e&&H(b,function(b,g){e&&e[g]!==b&&(c[g]=b,n=!0)});if(n){e&&(b=B(e,c));if(b)if(null===b.width||"auto"===b.width)delete this.textWidth;else if("text"===g.nodeName.toLowerCase()&&b.width)var v=this.textWidth=
E(b.width);this.styles=b;v&&!d&&this.renderer.forExport&&delete b.width;if(g.namespaceURI===this.SVG_NS){var F=function(b,e){return"-"+e.toLowerCase()};H(b,function(b,e){-1===a.indexOf(e)&&(k+=e.replace(/([A-Z])/g,F)+":"+b+";")});k&&w(g,"style",k)}else K(g,b);this.added&&("text"===this.element.nodeName&&this.renderer.buildText(this),b&&b.textOutline&&this.applyTextOutline(b.textOutline))}return this};a.prototype.dashstyleSetter=function(b){var e=this["stroke-width"];"inherit"===e&&(e=1);if(b=b&&b.toLowerCase()){var c=
b.replace("shortdashdotdot","3,1,1,1,1,1,").replace("shortdashdot","3,1,1,1").replace("shortdot","1,1,").replace("shortdash","3,1,").replace("longdash","8,3,").replace(/dot/g,"1,3,").replace("dash","4,3,").replace(/,$/,"").split(",");for(b=c.length;b--;)c[b]=""+E(c[b])*v(e,NaN);b=c.join(",").replace(/NaN/g,"none");this.element.setAttribute("stroke-dasharray",b)}};a.prototype.destroy=function(){var b=this,e=b.element||{},c=b.renderer,g=e.ownerSVGElement,d=c.isSVG&&"SPAN"===e.nodeName&&b.parentGroup||
void 0;e.onclick=e.onmouseout=e.onmouseover=e.onmousemove=e.point=null;z(b);if(b.clipPath&&g){var a=b.clipPath;[].forEach.call(g.querySelectorAll("[clip-path],[CLIP-PATH]"),function(b){-1<b.getAttribute("clip-path").indexOf(a.element.id)&&b.removeAttribute("clip-path")});b.clipPath=a.destroy()}if(b.stops){for(g=0;g<b.stops.length;g++)b.stops[g].destroy();b.stops.length=0;b.stops=void 0}b.safeRemoveChild(e);for(c.styledMode||b.destroyShadows();d&&d.div&&0===d.div.childNodes.length;)e=d.parentGroup,
b.safeRemoveChild(d.div),delete d.div,d=e;b.alignTo&&y(c.alignedObjects,b);H(b,function(e,c){b[c]&&b[c].parentGroup===b&&b[c].destroy&&b[c].destroy();delete b[c]})};a.prototype.destroyShadows=function(){(this.shadows||[]).forEach(function(b){this.safeRemoveChild(b)},this);this.shadows=void 0};a.prototype.destroyTextPath=function(b,e){var c=b.getElementsByTagName("text")[0];if(c){if(c.removeAttribute("dx"),c.removeAttribute("dy"),e.element.setAttribute("id",""),this.textPathWrapper&&c.getElementsByTagName("textPath").length){for(b=
this.textPathWrapper.element.childNodes;b.length;)c.appendChild(b[0]);c.removeChild(this.textPathWrapper.element)}}else if(b.getAttribute("dx")||b.getAttribute("dy"))b.removeAttribute("dx"),b.removeAttribute("dy");this.textPathWrapper&&(this.textPathWrapper=this.textPathWrapper.destroy())};a.prototype.dSetter=function(b,e,c){k(b)&&("string"===typeof b[0]&&(b=this.renderer.pathToSegments(b)),this.pathArray=b,b=b.reduce(function(b,e,c){return e&&e.join?(c?b+" ":"")+e.join(" "):(e||"").toString()},""));
/(NaN| {2}|^$)/.test(b)&&(b="M 0 0");this[e]!==b&&(c.setAttribute(e,b),this[e]=b)};a.prototype.fadeOut=function(b){var e=this;e.animate({opacity:0},{duration:v(b,150),complete:function(){e.attr({y:-9999}).hide()}})};a.prototype.fillSetter=function(b,e,c){"string"===typeof b?c.setAttribute(e,b):b&&this.complexColor(b,e,c)};a.prototype.getBBox=function(e,c){var g=this.renderer,d=this.element,k=this.styles,n=this.textStr,D=g.cache,l=g.cacheKeys,F=d.namespaceURI===this.SVG_NS;c=v(c,this.rotation,0);var p=
g.styledMode?d&&a.prototype.getStyle.call(d,"font-size"):k&&k.fontSize,f;if(r(n)){var H=n.toString();-1===H.indexOf("<")&&(H=H.replace(/[0-9]/g,"0"));H+=["",c,p,this.textWidth,k&&k.textOverflow,k&&k.fontWeight].join()}H&&!e&&(f=D[H]);if(!f){if(F||g.forExport){try{var h=this.fakeTS&&function(b){var e=d.querySelector(".highcharts-text-outline");e&&K(e,{display:b})};b(h)&&h("none");f=d.getBBox?B({},d.getBBox()):{width:d.offsetWidth,height:d.offsetHeight};b(h)&&h("")}catch(X){""}if(!f||0>f.width)f={width:0,
height:0}}else f=this.htmlGetBBox();g.isSVG&&(e=f.width,g=f.height,F&&(f.height=g={"11px,17":14,"13px,20":16}[k&&k.fontSize+","+Math.round(g)]||g),c&&(k=c*q,f.width=Math.abs(g*Math.sin(k))+Math.abs(e*Math.cos(k)),f.height=Math.abs(g*Math.cos(k))+Math.abs(e*Math.sin(k))));if(H&&(""===n||0<f.height)){for(;250<l.length;)delete D[l.shift()];D[H]||l.push(H);D[H]=f}}return f};a.prototype.getStyle=function(b){return l.getComputedStyle(this.element||this,"").getPropertyValue(b)};a.prototype.hasClass=function(b){return-1!==
(""+this.attr("class")).split(" ").indexOf(b)};a.prototype.hide=function(b){b?this.attr({y:-9999}):this.attr({visibility:"hidden"});return this};a.prototype.htmlGetBBox=function(){return{height:0,width:0,x:0,y:0}};a.prototype.init=function(b,e){this.element="span"===e?p(e):m.createElementNS(this.SVG_NS,e);this.renderer=b;n(this,"afterInit")};a.prototype.invert=function(b){this.inverted=b;this.updateTransform();return this};a.prototype.on=function(b,e){var c=this.onEvents;if(c[b])c[b]();c[b]=f(this.element,
b,e);return this};a.prototype.opacitySetter=function(b,e,c){this.opacity=b=Number(Number(b).toFixed(3));c.setAttribute(e,b)};a.prototype.removeClass=function(b){return this.attr("class",(""+this.attr("class")).replace(e(b)?new RegExp("(^| )"+b+"( |$)"):b," ").replace(/ +/g," ").trim())};a.prototype.removeTextOutline=function(){var b=this.element.querySelector("tspan.highcharts-text-outline");b&&this.safeRemoveChild(b)};a.prototype.safeRemoveChild=function(b){var e=b.parentNode;e&&e.removeChild(b)};
a.prototype.setRadialReference=function(b){var e=this.element.gradient&&this.renderer.gradients[this.element.gradient];this.element.radialReference=b;e&&e.radAttr&&e.animate(this.renderer.getRadialAttr(b,e.radAttr));return this};a.prototype.setTextPath=function(b,e){var c=this.element,d=this.text?this.text.element:c,a={textAnchor:"text-anchor"},k=!1,n=this.textPathWrapper,v=!n;e=D(!0,{enabled:!0,attributes:{dy:-5,startOffset:"50%",textAnchor:"middle"}},e);var F=t.filterUserAttributes(e.attributes);
if(b&&e&&e.enabled){n&&null===n.element.parentNode?(v=!0,n=n.destroy()):n&&this.removeTextOutline.call(n.parentGroup);this.options&&this.options.padding&&(F.dx=-this.options.padding);n||(this.textPathWrapper=n=this.renderer.createElement("textPath"),k=!0);var f=n.element;(e=b.element.getAttribute("id"))||b.element.setAttribute("id",e=S());if(v)for(d.setAttribute("y",0),g(F.dx)&&d.setAttribute("x",-F.dx),b=[].slice.call(d.childNodes),v=0;v<b.length;v++){var p=b[v];p.nodeType!==l.Node.TEXT_NODE&&"tspan"!==
p.nodeName||f.appendChild(p)}k&&n&&n.add({element:d});f.setAttributeNS("http://www.w3.org/1999/xlink","href",this.renderer.url+"#"+e);r(F.dy)&&(f.parentNode.setAttribute("dy",F.dy),delete F.dy);r(F.dx)&&(f.parentNode.setAttribute("dx",F.dx),delete F.dx);H(F,function(b,e){f.setAttribute(a[e]||e,b)});c.removeAttribute("transform");this.removeTextOutline.call(n);this.text&&!this.renderer.styledMode&&this.attr({fill:"none","stroke-width":0});this.applyTextOutline=this.updateTransform=h}else n&&(delete this.updateTransform,
delete this.applyTextOutline,this.destroyTextPath(c,b),this.updateTransform(),this.options&&this.options.rotation&&this.applyTextOutline(this.options.style.textOutline));return this};a.prototype.shadow=function(b,e,c){var g=[],d=this.element,a=this.oldShadowOptions,k={color:"#000000",offsetX:this.parentInverted?-1:1,offsetY:this.parentInverted?-1:1,opacity:.15,width:3},J=!1,F;!0===b?F=k:"object"===typeof b&&(F=B(k,b));F&&(F&&a&&H(F,function(b,e){b!==a[e]&&(J=!0)}),J&&this.destroyShadows(),this.oldShadowOptions=
F);if(!F)this.destroyShadows();else if(!this.shadows){var n=F.opacity/F.width;var v=this.parentInverted?"translate("+F.offsetY+", "+F.offsetX+")":"translate("+F.offsetX+", "+F.offsetY+")";for(k=1;k<=F.width;k++){var f=d.cloneNode(!1);var D=2*F.width+1-2*k;w(f,{stroke:b.color||"#000000","stroke-opacity":n*k,"stroke-width":D,transform:v,fill:"none"});f.setAttribute("class",(f.getAttribute("class")||"")+" highcharts-shadow");c&&(w(f,"height",Math.max(w(f,"height")-D,0)),f.cutHeight=D);e?e.element.appendChild(f):
d.parentNode&&d.parentNode.insertBefore(f,d);g.push(f)}this.shadows=g}return this};a.prototype.show=function(b){return this.attr({visibility:b?"inherit":"visible"})};a.prototype.strokeSetter=function(b,e,c){this[e]=b;this.stroke&&this["stroke-width"]?(a.prototype.fillSetter.call(this,this.stroke,"stroke",c),c.setAttribute("stroke-width",this["stroke-width"]),this.hasStroke=!0):"stroke-width"===e&&0===b&&this.hasStroke?(c.removeAttribute("stroke"),this.hasStroke=!1):this.renderer.styledMode&&this["stroke-width"]&&
(c.setAttribute("stroke-width",this["stroke-width"]),this.hasStroke=!0)};a.prototype.strokeWidth=function(){if(!this.renderer.styledMode)return this["stroke-width"]||0;var b=this.getStyle("stroke-width"),e=0;if(b.indexOf("px")===b.length-2)e=E(b);else if(""!==b){var g=m.createElementNS(c,"rect");w(g,{width:b,"stroke-width":0});this.element.parentNode.appendChild(g);e=g.getBBox().width;g.parentNode.removeChild(g)}return e};a.prototype.symbolAttr=function(b){var e=this;"x y r start end width height innerR anchorX anchorY clockwise".split(" ").forEach(function(c){e[c]=
v(b[c],e[c])});e.attr({d:e.renderer.symbols[e.symbolName](e.x,e.y,e.width,e.height,e)})};a.prototype.textSetter=function(b){b!==this.textStr&&(delete this.textPxLength,this.textStr=b,this.added&&this.renderer.buildText(this))};a.prototype.titleSetter=function(b){var e=this.element,c=e.getElementsByTagName("title")[0]||m.createElementNS(this.SVG_NS,"title");e.insertBefore?e.insertBefore(c,e.firstChild):e.appendChild(c);c.textContent=String(v(b,"")).replace(/<[^>]*>/g,"").replace(/&lt;/g,"<").replace(/&gt;/g,
">")};a.prototype.toFront=function(){var b=this.element;b.parentNode.appendChild(b);return this};a.prototype.translate=function(b,e){return this.attr({translateX:b,translateY:e})};a.prototype.updateShadows=function(b,e,c){var g=this.shadows;if(g)for(var d=g.length;d--;)c.call(g[d],"height"===b?Math.max(e-(g[d].cutHeight||0),0):"d"===b?this.d:e,b,g[d])};a.prototype.updateTransform=function(){var b=this.scaleX,e=this.scaleY,c=this.inverted,g=this.rotation,d=this.matrix,a=this.element,k=this.translateX||
0,n=this.translateY||0;c&&(k+=this.width,n+=this.height);k=["translate("+k+","+n+")"];r(d)&&k.push("matrix("+d.join(",")+")");c?k.push("rotate(90) scale(-1,1)"):g&&k.push("rotate("+g+" "+v(this.rotationOriginX,a.getAttribute("x"),0)+" "+v(this.rotationOriginY,a.getAttribute("y")||0)+")");(r(b)||r(e))&&k.push("scale("+v(b,1)+" "+v(e,1)+")");k.length&&a.setAttribute("transform",k.join(" "))};a.prototype.visibilitySetter=function(b,e,c){"inherit"===b?c.removeAttribute(e):this[e]!==b&&c.setAttribute(e,
b);this[e]=b};a.prototype.xGetter=function(b){"circle"===this.element.nodeName&&("x"===b?b="cx":"y"===b&&(b="cy"));return this._defaultGetter(b)};a.prototype.zIndexSetter=function(b,e){var c=this.renderer,g=this.parentGroup,d=(g||c).element||c.box,a=this.element;c=d===c.box;var k=!1;var n=this.added;var F;r(b)?(a.setAttribute("data-z-index",b),b=+b,this[e]===b&&(n=!1)):r(this[e])&&a.removeAttribute("data-z-index");this[e]=b;if(n){(b=this.zIndex)&&g&&(g.handleZ=!0);e=d.childNodes;for(F=e.length-1;0<=
F&&!k;F--){g=e[F];n=g.getAttribute("data-z-index");var v=!r(n);if(g!==a)if(0>b&&v&&!c&&!F)d.insertBefore(a,e[F]),k=!0;else if(E(n)<=b||v&&(!r(b)||0<=b))d.insertBefore(a,e[F+1]||null),k=!0}k||(d.insertBefore(a,e[c?3:0]||null),k=!0)}return k};return a}();a.prototype["stroke-widthSetter"]=a.prototype.strokeSetter;a.prototype.yGetter=a.prototype.xGetter;a.prototype.matrixSetter=a.prototype.rotationOriginXSetter=a.prototype.rotationOriginYSetter=a.prototype.rotationSetter=a.prototype.scaleXSetter=a.prototype.scaleYSetter=
a.prototype.translateXSetter=a.prototype.translateYSetter=a.prototype.verticalAlignSetter=function(b,e){this[e]=b;this.doTransform=!0};"";return a});L(a,"Core/Renderer/RendererRegistry.js",[a["Core/Globals.js"]],function(a){var u;(function(u){u.rendererTypes={};var t;u.getRendererType=function(a){void 0===a&&(a=t);return u.rendererTypes[a]||u.rendererTypes[t]};u.registerRendererType=function(x,A,I){u.rendererTypes[x]=A;if(!t||I)t=x,a.Renderer=A}})(u||(u={}));return u});L(a,"Core/Renderer/SVG/SVGLabel.js",
[a["Core/Renderer/SVG/SVGElement.js"],a["Core/Utilities.js"]],function(a,t){var u=this&&this.__extends||function(){var a=function(h,d){a=Object.setPrototypeOf||{__proto__:[]}instanceof Array&&function(c,d){c.__proto__=d}||function(c,d){for(var a in d)d.hasOwnProperty(a)&&(c[a]=d[a])};return a(h,d)};return function(h,d){function c(){this.constructor=h}a(h,d);h.prototype=null===d?Object.create(d):(c.prototype=d.prototype,new c)}}(),G=t.defined,x=t.extend,C=t.isNumber,I=t.merge,z=t.pick,q=t.removeEvent;
return function(m){function h(d,c,a,f,w,p,K,r,y,B){var n=m.call(this)||this;n.paddingLeftSetter=n.paddingSetter;n.paddingRightSetter=n.paddingSetter;n.init(d,"g");n.textStr=c;n.x=a;n.y=f;n.anchorX=p;n.anchorY=K;n.baseline=y;n.className=B;n.addClass("button"===B?"highcharts-no-tooltip":"highcharts-label");B&&n.addClass("highcharts-"+B);n.text=d.text(void 0,0,0,r).attr({zIndex:1});var k;"string"===typeof w&&((k=/^url\((.*?)\)$/.test(w))||n.renderer.symbols[w])&&(n.symbolKey=w);n.bBox=h.emptyBBox;n.padding=
3;n.baselineOffset=0;n.needsBox=d.styledMode||k;n.deferredAttr={};n.alignFactor=0;return n}u(h,m);h.prototype.alignSetter=function(d){d={left:0,center:.5,right:1}[d];d!==this.alignFactor&&(this.alignFactor=d,this.bBox&&C(this.xSetting)&&this.attr({x:this.xSetting}))};h.prototype.anchorXSetter=function(d,c){this.anchorX=d;this.boxAttr(c,Math.round(d)-this.getCrispAdjust()-this.xSetting)};h.prototype.anchorYSetter=function(d,c){this.anchorY=d;this.boxAttr(c,d-this.ySetting)};h.prototype.boxAttr=function(d,
c){this.box?this.box.attr(d,c):this.deferredAttr[d]=c};h.prototype.css=function(d){if(d){var c={};d=I(d);h.textProps.forEach(function(a){"undefined"!==typeof d[a]&&(c[a]=d[a],delete d[a])});this.text.css(c);var l="width"in c;"fontSize"in c||"fontWeight"in c?this.updateTextPadding():l&&this.updateBoxSize()}return a.prototype.css.call(this,d)};h.prototype.destroy=function(){q(this.element,"mouseenter");q(this.element,"mouseleave");this.text&&this.text.destroy();this.box&&(this.box=this.box.destroy());
a.prototype.destroy.call(this)};h.prototype.fillSetter=function(d,c){d&&(this.needsBox=!0);this.fill=d;this.boxAttr(c,d)};h.prototype.getBBox=function(){this.textStr&&0===this.bBox.width&&0===this.bBox.height&&this.updateBoxSize();var d=this.padding,c=z(this.paddingLeft,d);return{width:this.width,height:this.height,x:this.bBox.x-c,y:this.bBox.y-d}};h.prototype.getCrispAdjust=function(){return this.renderer.styledMode&&this.box?this.box.strokeWidth()%2/2:(this["stroke-width"]?parseInt(this["stroke-width"],
10):0)%2/2};h.prototype.heightSetter=function(d){this.heightSetting=d};h.prototype.onAdd=function(){var d=this.textStr;this.text.add(this);this.attr({text:G(d)?d:"",x:this.x,y:this.y});this.box&&G(this.anchorX)&&this.attr({anchorX:this.anchorX,anchorY:this.anchorY})};h.prototype.paddingSetter=function(d,c){C(d)?d!==this[c]&&(this[c]=d,this.updateTextPadding()):this[c]=void 0};h.prototype.rSetter=function(d,c){this.boxAttr(c,d)};h.prototype.shadow=function(d){d&&!this.renderer.styledMode&&(this.updateBoxSize(),
this.box&&this.box.shadow(d));return this};h.prototype.strokeSetter=function(d,c){this.stroke=d;this.boxAttr(c,d)};h.prototype["stroke-widthSetter"]=function(d,c){d&&(this.needsBox=!0);this["stroke-width"]=d;this.boxAttr(c,d)};h.prototype["text-alignSetter"]=function(d){this.textAlign=d};h.prototype.textSetter=function(d){"undefined"!==typeof d&&this.text.attr({text:d});this.updateTextPadding()};h.prototype.updateBoxSize=function(){var d=this.text.element.style,c={},a=this.padding,f=this.bBox=C(this.widthSetting)&&
C(this.heightSetting)&&!this.textAlign||!G(this.text.textStr)?h.emptyBBox:this.text.getBBox();this.width=this.getPaddedWidth();this.height=(this.heightSetting||f.height||0)+2*a;d=this.renderer.fontMetrics(d&&d.fontSize,this.text);this.baselineOffset=a+Math.min((this.text.firstLineMetrics||d).b,f.height||Infinity);this.heightSetting&&(this.baselineOffset+=(this.heightSetting-d.h)/2);this.needsBox&&(this.box||(a=this.box=this.symbolKey?this.renderer.symbol(this.symbolKey):this.renderer.rect(),a.addClass(("button"===
this.className?"":"highcharts-label-box")+(this.className?" highcharts-"+this.className+"-box":"")),a.add(this)),a=this.getCrispAdjust(),c.x=a,c.y=(this.baseline?-this.baselineOffset:0)+a,c.width=Math.round(this.width),c.height=Math.round(this.height),this.box.attr(x(c,this.deferredAttr)),this.deferredAttr={})};h.prototype.updateTextPadding=function(){var d=this.text;this.updateBoxSize();var c=this.baseline?0:this.baselineOffset,a=z(this.paddingLeft,this.padding);G(this.widthSetting)&&this.bBox&&
("center"===this.textAlign||"right"===this.textAlign)&&(a+={center:.5,right:1}[this.textAlign]*(this.widthSetting-this.bBox.width));if(a!==d.x||c!==d.y)d.attr("x",a),d.hasBoxWidthChanged&&(this.bBox=d.getBBox(!0)),"undefined"!==typeof c&&d.attr("y",c);d.x=a;d.y=c};h.prototype.widthSetter=function(d){this.widthSetting=C(d)?d:void 0};h.prototype.getPaddedWidth=function(){var d=this.padding,c=z(this.paddingLeft,d);d=z(this.paddingRight,d);return(this.widthSetting||this.bBox.width||0)+c+d};h.prototype.xSetter=
function(d){this.x=d;this.alignFactor&&(d-=this.alignFactor*this.getPaddedWidth(),this["forceAnimate:x"]=!0);this.xSetting=Math.round(d);this.attr("translateX",this.xSetting)};h.prototype.ySetter=function(d){this.ySetting=this.y=Math.round(d);this.attr("translateY",this.ySetting)};h.emptyBBox={width:0,height:0,x:0,y:0};h.textProps="color direction fontFamily fontSize fontStyle fontWeight lineHeight textAlign textDecoration textOutline textOverflow width".split(" ");return h}(a)});L(a,"Core/Renderer/SVG/Symbols.js",
[a["Core/Utilities.js"]],function(a){function u(a,q,m,h,d){var c=[];if(d){var l=d.start||0,f=I(d.r,m);m=I(d.r,h||m);var w=(d.end||0)-.001;h=d.innerR;var p=I(d.open,.001>Math.abs((d.end||0)-l-2*Math.PI)),K=Math.cos(l),r=Math.sin(l),y=Math.cos(w),B=Math.sin(w);l=I(d.longArc,.001>w-l-Math.PI?0:1);c.push(["M",a+f*K,q+m*r],["A",f,m,0,l,I(d.clockwise,1),a+f*y,q+m*B]);x(h)&&c.push(p?["M",a+h*y,q+h*B]:["L",a+h*y,q+h*B],["A",h,h,0,l,x(d.clockwise)?1-d.clockwise:0,a+h*K,q+h*r]);p||c.push(["Z"])}return c}function A(a,
q,m,h,d){return d&&d.r?G(a,q,m,h,d):[["M",a,q],["L",a+m,q],["L",a+m,q+h],["L",a,q+h],["Z"]]}function G(a,q,m,h,d){d=d&&d.r||0;return[["M",a+d,q],["L",a+m-d,q],["C",a+m,q,a+m,q,a+m,q+d],["L",a+m,q+h-d],["C",a+m,q+h,a+m,q+h,a+m-d,q+h],["L",a+d,q+h],["C",a,q+h,a,q+h,a,q+h-d],["L",a,q+d],["C",a,q,a,q,a+d,q]]}var x=a.defined,C=a.isNumber,I=a.pick;return{arc:u,callout:function(a,q,m,h,d){var c=Math.min(d&&d.r||0,m,h),l=c+6,f=d&&d.anchorX;d=d&&d.anchorY||0;var w=G(a,q,m,h,{r:c});if(!C(f))return w;a+f>=m?
d>q+l&&d<q+h-l?w.splice(3,1,["L",a+m,d-6],["L",a+m+6,d],["L",a+m,d+6],["L",a+m,q+h-c]):w.splice(3,1,["L",a+m,h/2],["L",f,d],["L",a+m,h/2],["L",a+m,q+h-c]):0>=a+f?d>q+l&&d<q+h-l?w.splice(7,1,["L",a,d+6],["L",a-6,d],["L",a,d-6],["L",a,q+c]):w.splice(7,1,["L",a,h/2],["L",f,d],["L",a,h/2],["L",a,q+c]):d&&d>h&&f>a+l&&f<a+m-l?w.splice(5,1,["L",f+6,q+h],["L",f,q+h+6],["L",f-6,q+h],["L",a+c,q+h]):d&&0>d&&f>a+l&&f<a+m-l&&w.splice(1,1,["L",f-6,q],["L",f,q-6],["L",f+6,q],["L",m-c,q]);return w},circle:function(a,
q,m,h){return u(a+m/2,q+h/2,m/2,h/2,{start:.5*Math.PI,end:2.5*Math.PI,open:!1})},diamond:function(a,q,m,h){return[["M",a+m/2,q],["L",a+m,q+h/2],["L",a+m/2,q+h],["L",a,q+h/2],["Z"]]},rect:A,roundedRect:G,square:A,triangle:function(a,q,m,h){return[["M",a+m/2,q],["L",a+m,q+h],["L",a,q+h],["Z"]]},"triangle-down":function(a,q,m,h){return[["M",a,q],["L",a+m,q],["L",a+m/2,q+h],["Z"]]}}});L(a,"Core/Renderer/SVG/TextBuilder.js",[a["Core/Renderer/HTML/AST.js"],a["Core/Globals.js"],a["Core/Utilities.js"]],function(a,
t,A){var u=t.doc,x=t.SVG_NS,C=t.win,I=A.attr,z=A.isString,q=A.objectEach,m=A.pick;return function(){function h(a){var c=a.styles;this.renderer=a.renderer;this.svgElement=a;this.width=a.textWidth;this.textLineHeight=c&&c.lineHeight;this.textOutline=c&&c.textOutline;this.ellipsis=!(!c||"ellipsis"!==c.textOverflow);this.noWrap=!(!c||"nowrap"!==c.whiteSpace);this.fontSize=c&&c.fontSize}h.prototype.buildSVG=function(){var d=this.svgElement,c=d.element,l=d.renderer,f=m(d.textStr,"").toString(),h=-1!==f.indexOf("<"),
p=c.childNodes;l=this.width&&!d.added&&l.box;var K=/<br.*?>/g,r=[f,this.ellipsis,this.noWrap,this.textLineHeight,this.textOutline,this.fontSize,this.width].join();if(r!==d.textCache){d.textCache=r;delete d.actualWidth;for(r=p.length;r--;)c.removeChild(p[r]);h||this.ellipsis||this.width||-1!==f.indexOf(" ")&&(!this.noWrap||K.test(f))?""!==f&&(l&&l.appendChild(c),f=new a(f),this.modifyTree(f.nodes),f.addToDOM(d.element),this.modifyDOM(),this.ellipsis&&-1!==(c.textContent||"").indexOf("\u2026")&&d.attr("title",
this.unescapeEntities(d.textStr||"",["&lt;","&gt;"])),l&&l.removeChild(c)):c.appendChild(u.createTextNode(this.unescapeEntities(f)));z(this.textOutline)&&d.applyTextOutline&&d.applyTextOutline(this.textOutline)}};h.prototype.modifyDOM=function(){var a=this,c=this.svgElement,l=I(c.element,"x");c.firstLineMetrics=void 0;for(var f;f=c.element.firstChild;)if(/^[\s\u200B]*$/.test(f.textContent||" "))c.element.removeChild(f);else break;[].forEach.call(c.element.querySelectorAll("tspan.highcharts-br"),function(d,
f){d.nextSibling&&d.previousSibling&&(0===f&&1===d.previousSibling.nodeType&&(c.firstLineMetrics=c.renderer.fontMetrics(void 0,d.previousSibling)),I(d,{dy:a.getLineHeight(d.nextSibling),x:l}))});var h=this.width||0;if(h){var p=function(d,f){var p=d.textContent||"",n=p.replace(/([^\^])-/g,"$1- ").split(" "),k=!a.noWrap&&(1<n.length||1<c.element.childNodes.length),b=a.getLineHeight(f),g=0,e=c.actualWidth;if(a.ellipsis)p&&a.truncate(d,p,void 0,0,Math.max(0,h-parseInt(a.fontSize||12,10)),function(b,e){return b.substring(0,
e)+"\u2026"});else if(k){p=[];for(k=[];f.firstChild&&f.firstChild!==d;)k.push(f.firstChild),f.removeChild(f.firstChild);for(;n.length;)n.length&&!a.noWrap&&0<g&&(p.push(d.textContent||""),d.textContent=n.join(" ").replace(/- /g,"-")),a.truncate(d,void 0,n,0===g?e||0:0,h,function(b,e){return n.slice(0,e).join(" ").replace(/- /g,"-")}),e=c.actualWidth,g++;k.forEach(function(b){f.insertBefore(b,d)});p.forEach(function(e){f.insertBefore(u.createTextNode(e),d);e=u.createElementNS(x,"tspan");e.textContent=
"\u200b";I(e,{dy:b,x:l});f.insertBefore(e,d)})}},m=function(a){[].slice.call(a.childNodes).forEach(function(d){d.nodeType===C.Node.TEXT_NODE?p(d,a):(-1!==d.className.baseVal.indexOf("highcharts-br")&&(c.actualWidth=0),m(d))})};m(c.element)}};h.prototype.getLineHeight=function(a){var c;a=a.nodeType===C.Node.TEXT_NODE?a.parentElement:a;this.renderer.styledMode||(c=a&&/(px|em)$/.test(a.style.fontSize)?a.style.fontSize:this.fontSize||this.renderer.style.fontSize||12);return this.textLineHeight?parseInt(this.textLineHeight.toString(),
10):this.renderer.fontMetrics(c,a||this.svgElement.element).h};h.prototype.modifyTree=function(a){var c=this,d=function(f,l){var p=f.tagName,h=c.renderer.styledMode,r=f.attributes||{};if("b"===p||"strong"===p)h?r["class"]="highcharts-strong":r.style="font-weight:bold;"+(r.style||"");else if("i"===p||"em"===p)h?r["class"]="highcharts-emphasized":r.style="font-style:italic;"+(r.style||"");z(r.style)&&(r.style=r.style.replace(/(;| |^)color([ :])/,"$1fill$2"));"br"===p&&(r["class"]="highcharts-br",f.textContent=
"\u200b",(l=a[l+1])&&l.textContent&&(l.textContent=l.textContent.replace(/^ +/gm,"")));"#text"!==p&&"a"!==p&&(f.tagName="tspan");f.attributes=r;f.children&&f.children.filter(function(c){return"#text"!==c.tagName}).forEach(d)};a.forEach(d)};h.prototype.truncate=function(a,c,l,f,h,p){var d=this.svgElement,r=d.renderer,m=d.rotation,w=[],n=l?1:0,k=(c||l||"").length,b=k,g,e=function(b,e){e=e||b;var g=a.parentNode;if(g&&"undefined"===typeof w[e])if(g.getSubStringLength)try{w[e]=f+g.getSubStringLength(0,
l?e+1:e)}catch(O){""}else r.getSpanWidth&&(a.textContent=p(c||l,b),w[e]=f+r.getSpanWidth(d,a));return w[e]};d.rotation=0;var D=e(a.textContent.length);if(f+D>h){for(;n<=k;)b=Math.ceil((n+k)/2),l&&(g=p(l,b)),D=e(b,g&&g.length-1),n===k?n=k+1:D>h?k=b-1:n=b;0===k?a.textContent="":c&&k===c.length-1||(a.textContent=g||p(c||l,b))}l&&l.splice(0,b);d.actualWidth=D;d.rotation=m};h.prototype.unescapeEntities=function(a,c){q(this.renderer.escapes,function(d,f){c&&-1!==c.indexOf(d)||(a=a.toString().replace(new RegExp(d,
"g"),f))});return a};return h}()});L(a,"Core/Renderer/SVG/SVGRenderer.js",[a["Core/Renderer/HTML/AST.js"],a["Core/Color/Color.js"],a["Core/Globals.js"],a["Core/Renderer/RendererRegistry.js"],a["Core/Renderer/SVG/SVGElement.js"],a["Core/Renderer/SVG/SVGLabel.js"],a["Core/Renderer/SVG/Symbols.js"],a["Core/Renderer/SVG/TextBuilder.js"],a["Core/Utilities.js"]],function(a,t,A,G,x,C,I,z,q){var m=A.charts,h=A.deg2rad,d=A.doc,c=A.isFirefox,l=A.isMS,f=A.isWebKit,w=A.noop,p=A.SVG_NS,K=A.symbolSizes,r=A.win,
y=q.addEvent,B=q.attr,n=q.createElement,k=q.css,b=q.defined,g=q.destroyObjectProperties,e=q.extend,D=q.isArray,H=q.isNumber,v=q.isObject,E=q.isString,O=q.merge,u=q.pick,P=q.pInt,U=q.uniqueKey,Y;A=function(){function J(b,e,c,a,g,d,k){this.width=this.url=this.style=this.isSVG=this.imgCount=this.height=this.gradients=this.globalAnimation=this.defs=this.chartIndex=this.cacheKeys=this.cache=this.boxWrapper=this.box=this.alignedObjects=void 0;this.init(b,e,c,a,g,d,k)}J.prototype.init=function(b,e,a,g,J,
F,n){var M=this.createElement("svg").attr({version:"1.1","class":"highcharts-root"}),f=M.element;n||M.css(this.getStyle(g));b.appendChild(f);B(b,"dir","ltr");-1===b.innerHTML.indexOf("xmlns")&&B(f,"xmlns",this.SVG_NS);this.isSVG=!0;this.box=f;this.boxWrapper=M;this.alignedObjects=[];this.url=this.getReferenceURL();this.createElement("desc").add().element.appendChild(d.createTextNode("Created with Highcharts 9.3.1"));this.defs=this.createElement("defs").add();this.allowHTML=F;this.forExport=J;this.styledMode=
n;this.gradients={};this.cache={};this.cacheKeys=[];this.imgCount=0;this.setSize(e,a,!1);var v;c&&b.getBoundingClientRect&&(e=function(){k(b,{left:0,top:0});v=b.getBoundingClientRect();k(b,{left:Math.ceil(v.left)-v.left+"px",top:Math.ceil(v.top)-v.top+"px"})},e(),this.unSubPixelFix=y(r,"resize",e))};J.prototype.definition=function(b){return(new a([b])).addToDOM(this.defs.element)};J.prototype.getReferenceURL=function(){if((c||f)&&d.getElementsByTagName("base").length){if(!b(Y)){var e=U();e=(new a([{tagName:"svg",
attributes:{width:8,height:8},children:[{tagName:"defs",children:[{tagName:"clipPath",attributes:{id:e},children:[{tagName:"rect",attributes:{width:4,height:4}}]}]},{tagName:"rect",attributes:{id:"hitme",width:8,height:8,"clip-path":"url(#"+e+")",fill:"rgba(0,0,0,0.001)"}}]}])).addToDOM(d.body);k(e,{position:"fixed",top:0,left:0,zIndex:9E5});var g=d.elementFromPoint(6,6);Y="hitme"===(g&&g.id);d.body.removeChild(e)}if(Y)return r.location.href.split("#")[0].replace(/<[^>]*>/g,"").replace(/([\('\)])/g,
"\\$1").replace(/ /g,"%20")}return""};J.prototype.getStyle=function(b){return this.style=e({fontFamily:'"Lucida Grande", "Lucida Sans Unicode", Arial, Helvetica, sans-serif',fontSize:"12px"},b)};J.prototype.setStyle=function(b){this.boxWrapper.css(this.getStyle(b))};J.prototype.isHidden=function(){return!this.boxWrapper.getBBox().width};J.prototype.destroy=function(){var b=this.defs;this.box=null;this.boxWrapper=this.boxWrapper.destroy();g(this.gradients||{});this.gradients=null;b&&(this.defs=b.destroy());
this.unSubPixelFix&&this.unSubPixelFix();return this.alignedObjects=null};J.prototype.createElement=function(b){var e=new this.Element;e.init(this,b);return e};J.prototype.getRadialAttr=function(b,e){return{cx:b[0]-b[2]/2+(e.cx||0)*b[2],cy:b[1]-b[2]/2+(e.cy||0)*b[2],r:(e.r||0)*b[2]}};J.prototype.buildText=function(b){(new z(b)).buildSVG()};J.prototype.getContrast=function(b){b=t.parse(b).rgba;b[0]*=1;b[1]*=1.2;b[2]*=.5;return 459<b[0]+b[1]+b[2]?"#000000":"#FFFFFF"};J.prototype.button=function(b,c,
g,d,k,F,J,n,v,f){var p=this.label(b,c,g,v,void 0,void 0,f,void 0,"button"),D=this.styledMode,M=0,h=k?O(k):{};b=h&&h.style||{};h=a.filterUserAttributes(h);p.attr(O({padding:8,r:2},h));if(!D){h=O({fill:"#f7f7f7",stroke:"#cccccc","stroke-width":1,style:{color:"#333333",cursor:"pointer",fontWeight:"normal"}},{style:b},h);var H=h.style;delete h.style;F=O(h,{fill:"#e6e6e6"},a.filterUserAttributes(F||{}));var N=F.style;delete F.style;J=O(h,{fill:"#e6ebf5",style:{color:"#000000",fontWeight:"bold"}},a.filterUserAttributes(J||
{}));var r=J.style;delete J.style;n=O(h,{style:{color:"#cccccc"}},a.filterUserAttributes(n||{}));var E=n.style;delete n.style}y(p.element,l?"mouseover":"mouseenter",function(){3!==M&&p.setState(1)});y(p.element,l?"mouseout":"mouseleave",function(){3!==M&&p.setState(M)});p.setState=function(b){1!==b&&(p.state=M=b);p.removeClass(/highcharts-button-(normal|hover|pressed|disabled)/).addClass("highcharts-button-"+["normal","hover","pressed","disabled"][b||0]);D||p.attr([h,F,J,n][b||0]).css([H,N,r,E][b||
0])};D||p.attr(h).css(e({cursor:"default"},H));return p.on("touchstart",function(b){return b.stopPropagation()}).on("click",function(b){3!==M&&d.call(p,b)})};J.prototype.crispLine=function(e,c,a){void 0===a&&(a="round");var g=e[0],d=e[1];b(g[1])&&g[1]===d[1]&&(g[1]=d[1]=Math[a](g[1])-c%2/2);b(g[2])&&g[2]===d[2]&&(g[2]=d[2]=Math[a](g[2])+c%2/2);return e};J.prototype.path=function(b){var c=this.styledMode?{}:{fill:"none"};D(b)?c.d=b:v(b)&&e(c,b);return this.createElement("path").attr(c)};J.prototype.circle=
function(b,e,c){b=v(b)?b:"undefined"===typeof b?{}:{x:b,y:e,r:c};e=this.createElement("circle");e.xSetter=e.ySetter=function(b,e,c){c.setAttribute("c"+e,b)};return e.attr(b)};J.prototype.arc=function(b,e,c,a,g,d){v(b)?(a=b,e=a.y,c=a.r,b=a.x):a={innerR:a,start:g,end:d};b=this.symbol("arc",b,e,c,c,a);b.r=c;return b};J.prototype.rect=function(b,e,c,a,g,d){g=v(b)?b.r:g;var k=this.createElement("rect");b=v(b)?b:"undefined"===typeof b?{}:{x:b,y:e,width:Math.max(c,0),height:Math.max(a,0)};this.styledMode||
("undefined"!==typeof d&&(b["stroke-width"]=d,b=k.crisp(b)),b.fill="none");g&&(b.r=g);k.rSetter=function(b,e,c){k.r=b;B(c,{rx:b,ry:b})};k.rGetter=function(){return k.r||0};return k.attr(b)};J.prototype.setSize=function(b,e,c){this.width=b;this.height=e;this.boxWrapper.animate({width:b,height:e},{step:function(){this.attr({viewBox:"0 0 "+this.attr("width")+" "+this.attr("height")})},duration:u(c,!0)?void 0:0});this.alignElements()};J.prototype.g=function(b){var e=this.createElement("g");return b?e.attr({"class":"highcharts-"+
b}):e};J.prototype.image=function(b,e,c,a,g,d){var k={preserveAspectRatio:"none"},F=function(b,e){b.setAttributeNS?b.setAttributeNS("http://www.w3.org/1999/xlink","href",e):b.setAttribute("hc-svg-href",e)};H(e)&&(k.x=e);H(c)&&(k.y=c);H(a)&&(k.width=a);H(g)&&(k.height=g);var J=this.createElement("image").attr(k);e=function(e){F(J.element,b);d.call(J,e)};d?(F(J.element,"data:image/gif;base64,R0lGODlhAQABAAAAACH5BAEKAAEALAAAAAABAAEAAAICTAEAOw=="),c=new r.Image,y(c,"load",e),c.src=b,c.complete&&e({})):
F(J.element,b);return J};J.prototype.symbol=function(c,a,g,J,v,F){var f=this,p=/^url\((.*?)\)$/,D=p.test(c),l=!D&&(this.symbols[c]?c:"circle"),h=l&&this.symbols[l],H;if(h){"number"===typeof a&&(H=h.call(this.symbols,Math.round(a||0),Math.round(g||0),J||0,v||0,F));var M=this.path(H);f.styledMode||M.attr("fill","none");e(M,{symbolName:l||void 0,x:a,y:g,width:J,height:v});F&&e(M,F)}else if(D){var r=c.match(p)[1];var E=M=this.image(r);E.imgwidth=u(K[r]&&K[r].width,F&&F.width);E.imgheight=u(K[r]&&K[r].height,
F&&F.height);var w=function(b){return b.attr({width:b.width,height:b.height})};["width","height"].forEach(function(e){E[e+"Setter"]=function(e,c){var a=this["img"+c];this[c]=e;b(a)&&(F&&"within"===F.backgroundSize&&this.width&&this.height&&(a=Math.round(a*Math.min(this.width/this.imgwidth,this.height/this.imgheight))),this.element&&this.element.setAttribute(c,a),this.alignByTranslate||(e=((this[c]||0)-a)/2,this.attr("width"===c?{translateX:e}:{translateY:e})))}});b(a)&&E.attr({x:a,y:g});E.isImg=!0;
b(E.imgwidth)&&b(E.imgheight)?w(E):(E.attr({width:0,height:0}),n("img",{onload:function(){var b=m[f.chartIndex];0===this.width&&(k(this,{position:"absolute",top:"-999em"}),d.body.appendChild(this));K[r]={width:this.width,height:this.height};E.imgwidth=this.width;E.imgheight=this.height;E.element&&w(E);this.parentNode&&this.parentNode.removeChild(this);f.imgCount--;if(!f.imgCount&&b&&!b.hasLoaded)b.onload()},src:r}),this.imgCount++)}return M};J.prototype.clipRect=function(b,e,c,a){var g=U()+"-",d=
this.createElement("clipPath").attr({id:g}).add(this.defs);b=this.rect(b,e,c,a,0).add(d);b.id=g;b.clipPath=d;b.count=0;return b};J.prototype.text=function(e,c,a,g){var d={};if(g&&(this.allowHTML||!this.forExport))return this.html(e,c,a);d.x=Math.round(c||0);a&&(d.y=Math.round(a));b(e)&&(d.text=e);e=this.createElement("text").attr(d);if(!g||this.forExport&&!this.allowHTML)e.xSetter=function(b,e,c){for(var a=c.getElementsByTagName("tspan"),g=c.getAttribute(e),d=0,k;d<a.length;d++)k=a[d],k.getAttribute(e)===
g&&k.setAttribute(e,b);c.setAttribute(e,b)};return e};J.prototype.fontMetrics=function(b,e){b=!this.styledMode&&/px/.test(b)||!r.getComputedStyle?b||e&&e.style&&e.style.fontSize||this.style&&this.style.fontSize:e&&x.prototype.getStyle.call(e,"font-size");b=/px/.test(b)?P(b):12;e=24>b?b+3:Math.round(1.2*b);return{h:e,b:Math.round(.8*e),f:b}};J.prototype.rotCorr=function(b,e,c){var a=b;e&&c&&(a=Math.max(a*Math.cos(e*h),4));return{x:-b/3*Math.sin(e*h),y:a}};J.prototype.pathToSegments=function(b){for(var e=
[],c=[],a={A:8,C:7,H:2,L:3,M:3,Q:5,S:5,T:3,V:2},g=0;g<b.length;g++)E(c[0])&&H(b[g])&&c.length===a[c[0].toUpperCase()]&&b.splice(g,0,c[0].replace("M","L").replace("m","l")),"string"===typeof b[g]&&(c.length&&e.push(c.slice(0)),c.length=0),c.push(b[g]);e.push(c.slice(0));return e};J.prototype.label=function(b,e,c,a,g,d,k,J,n){return new C(this,b,e,c,a,g,d,k,J,n)};J.prototype.alignElements=function(){this.alignedObjects.forEach(function(b){return b.align()})};return J}();e(A.prototype,{Element:x,SVG_NS:p,
escapes:{"&":"&amp;","<":"&lt;",">":"&gt;","'":"&#39;",'"':"&quot;"},symbols:I,draw:w});G.registerRendererType("svg",A,!0);"";return A});L(a,"Core/Renderer/HTML/HTMLElement.js",[a["Core/Globals.js"],a["Core/Renderer/SVG/SVGElement.js"],a["Core/Utilities.js"]],function(a,t,A){var u=this&&this.__extends||function(){var c=function(a,d){c=Object.setPrototypeOf||{__proto__:[]}instanceof Array&&function(c,a){c.__proto__=a}||function(c,a){for(var d in a)a.hasOwnProperty(d)&&(c[d]=a[d])};return c(a,d)};return function(a,
d){function f(){this.constructor=a}c(a,d);a.prototype=null===d?Object.create(d):(f.prototype=d.prototype,new f)}}(),x=a.isFirefox,C=a.isMS,I=a.isWebKit,z=a.win,q=A.css,m=A.defined,h=A.extend,d=A.pick,c=A.pInt;return function(a){function f(){return null!==a&&a.apply(this,arguments)||this}u(f,a);f.compose=function(c){if(-1===f.composedClasses.indexOf(c)){f.composedClasses.push(c);var a=f.prototype,d=c.prototype;d.getSpanCorrection=a.getSpanCorrection;d.htmlCss=a.htmlCss;d.htmlGetBBox=a.htmlGetBBox;
d.htmlUpdateTransform=a.htmlUpdateTransform;d.setSpanRotation=a.setSpanRotation}return c};f.prototype.getSpanCorrection=function(c,a,d){this.xCorr=-c*d;this.yCorr=-a};f.prototype.htmlCss=function(c){var a="SPAN"===this.element.tagName&&c&&"width"in c,f=d(a&&c.width,void 0);if(a){delete c.width;this.textWidth=f;var l=!0}c&&"ellipsis"===c.textOverflow&&(c.whiteSpace="nowrap",c.overflow="hidden");this.styles=h(this.styles,c);q(this.element,c);l&&this.htmlUpdateTransform();return this};f.prototype.htmlGetBBox=
function(){var c=this.element;return{x:c.offsetLeft,y:c.offsetTop,width:c.offsetWidth,height:c.offsetHeight}};f.prototype.htmlUpdateTransform=function(){if(this.added){var a=this.renderer,d=this.element,f=this.translateX||0,l=this.translateY||0,h=this.x||0,B=this.y||0,n=this.textAlign||"left",k={left:0,center:.5,right:1}[n],b=this.styles;b=b&&b.whiteSpace;q(d,{marginLeft:f,marginTop:l});!a.styledMode&&this.shadows&&this.shadows.forEach(function(b){q(b,{marginLeft:f+1,marginTop:l+1})});this.inverted&&
[].forEach.call(d.childNodes,function(b){a.invertChild(b,d)});if("SPAN"===d.tagName){var g=this.rotation,e=this.textWidth&&c(this.textWidth),D=[g,n,d.innerHTML,this.textWidth,this.textAlign].join(),H=void 0;H=!1;if(e!==this.oldTextWidth){if(this.textPxLength)var v=this.textPxLength;else q(d,{width:"",whiteSpace:b||"nowrap"}),v=d.offsetWidth;(e>this.oldTextWidth||v>e)&&(/[ \-]/.test(d.textContent||d.innerText)||"ellipsis"===d.style.textOverflow)&&(q(d,{width:v>e||g?e+"px":"auto",display:"block",whiteSpace:b||
"normal"}),this.oldTextWidth=e,H=!0)}this.hasBoxWidthChanged=H;D!==this.cTT&&(H=a.fontMetrics(d.style.fontSize,d).b,!m(g)||g===(this.oldRotation||0)&&n===this.oldAlign||this.setSpanRotation(g,k,H),this.getSpanCorrection(!m(g)&&this.textPxLength||d.offsetWidth,H,k,g,n));q(d,{left:h+(this.xCorr||0)+"px",top:B+(this.yCorr||0)+"px"});this.cTT=D;this.oldRotation=g;this.oldAlign=n}}else this.alignOnAdd=!0};f.prototype.setSpanRotation=function(c,a,d){var f={},p=C&&!/Edge/.test(z.navigator.userAgent)?"-ms-transform":
I?"-webkit-transform":x?"MozTransform":z.opera?"-o-transform":void 0;p&&(f[p]=f.transform="rotate("+c+"deg)",f[p+(x?"Origin":"-origin")]=f.transformOrigin=100*a+"% "+d+"px",q(this.element,f))};f.composedClasses=[];return f}(t)});L(a,"Core/Renderer/HTML/HTMLRenderer.js",[a["Core/Renderer/HTML/AST.js"],a["Core/Renderer/SVG/SVGElement.js"],a["Core/Renderer/SVG/SVGRenderer.js"],a["Core/Utilities.js"]],function(a,t,A,G){var u=this&&this.__extends||function(){var a=function(h,d){a=Object.setPrototypeOf||
{__proto__:[]}instanceof Array&&function(c,a){c.__proto__=a}||function(c,a){for(var d in a)a.hasOwnProperty(d)&&(c[d]=a[d])};return a(h,d)};return function(h,d){function c(){this.constructor=h}a(h,d);h.prototype=null===d?Object.create(d):(c.prototype=d.prototype,new c)}}(),C=G.attr,I=G.createElement,z=G.extend,q=G.pick;return function(m){function h(){return null!==m&&m.apply(this,arguments)||this}u(h,m);h.compose=function(a){-1===h.composedClasses.indexOf(a)&&(h.composedClasses.push(a),a.prototype.html=
h.prototype.html);return a};h.prototype.html=function(d,c,l){var f=this.createElement("span"),h=f.element,p=f.renderer,m=p.isSVG,r=function(c,a){["opacity","visibility"].forEach(function(d){c[d+"Setter"]=function(k,b,g){var e=c.div?c.div.style:a;t.prototype[d+"Setter"].call(this,k,b,g);e&&(e[b]=k)}});c.addedSetters=!0};f.textSetter=function(c){c!==this.textStr&&(delete this.bBox,delete this.oldTextWidth,a.setElementHTML(this.element,q(c,"")),this.textStr=c,f.doTransform=!0)};m&&r(f,f.element.style);
f.xSetter=f.ySetter=f.alignSetter=f.rotationSetter=function(c,a){"align"===a?f.alignValue=f.textAlign=c:f[a]=c;f.doTransform=!0};f.afterSetters=function(){this.doTransform&&(this.htmlUpdateTransform(),this.doTransform=!1)};f.attr({text:d,x:Math.round(c),y:Math.round(l)}).css({position:"absolute"});p.styledMode||f.css({fontFamily:this.style.fontFamily,fontSize:this.style.fontSize});h.style.whiteSpace="nowrap";f.css=f.htmlCss;m&&(f.add=function(c){var a=p.box.parentNode,d=[];if(this.parentGroup=c){var k=
c.div;if(!k){for(;c;)d.push(c),c=c.parentGroup;d.reverse().forEach(function(b){function c(e,c){b[c]=e;"translateX"===c?p.left=e+"px":p.top=e+"px";b.doTransform=!0}var e=C(b.element,"class"),n=b.styles||{};k=b.div=b.div||I("div",e?{className:e}:void 0,{position:"absolute",left:(b.translateX||0)+"px",top:(b.translateY||0)+"px",display:b.display,opacity:b.opacity,cursor:n.cursor,pointerEvents:n.pointerEvents,visibility:b.visibility},k||a);var p=k.style;z(b,{classSetter:function(b){return function(e){this.element.setAttribute("class",
e);b.className=e}}(k),on:function(){d[0].div&&f.on.apply({element:d[0].div,onEvents:b.onEvents},arguments);return b},translateXSetter:c,translateYSetter:c});b.addedSetters||r(b)})}}else k=a;k.appendChild(h);f.added=!0;f.alignOnAdd&&f.htmlUpdateTransform();return f});return f};h.composedClasses=[];return h}(A)});L(a,"Core/Axis/AxisDefaults.js",[],function(){var a;(function(a){a.defaultXAxisOptions={alignTicks:!0,allowDecimals:void 0,panningEnabled:!0,zIndex:2,zoomEnabled:!0,dateTimeLabelFormats:{millisecond:{main:"%H:%M:%S.%L",
range:!1},second:{main:"%H:%M:%S",range:!1},minute:{main:"%H:%M",range:!1},hour:{main:"%H:%M",range:!1},day:{main:"%e. %b"},week:{main:"%e. %b"},month:{main:"%b '%y"},year:{main:"%Y"}},endOnTick:!1,gridLineDashStyle:"Solid",gridZIndex:1,labels:{autoRotation:void 0,autoRotationLimit:80,distance:void 0,enabled:!0,indentation:10,overflow:"justify",padding:5,reserveSpace:void 0,rotation:void 0,staggerLines:0,step:0,useHTML:!1,x:0,zIndex:7,style:{color:"#666666",cursor:"default",fontSize:"11px"}},maxPadding:.01,
minorGridLineDashStyle:"Solid",minorTickLength:2,minorTickPosition:"outside",minPadding:.01,offset:void 0,opposite:!1,reversed:void 0,reversedStacks:!1,showEmpty:!0,showFirstLabel:!0,showLastLabel:!0,startOfWeek:1,startOnTick:!1,tickLength:10,tickPixelInterval:100,tickmarkPlacement:"between",tickPosition:"outside",title:{align:"middle",rotation:0,useHTML:!1,x:0,y:0,style:{color:"#666666"}},type:"linear",uniqueNames:!0,visible:!0,minorGridLineColor:"#f2f2f2",minorGridLineWidth:1,minorTickColor:"#999999",
lineColor:"#ccd6eb",lineWidth:1,gridLineColor:"#e6e6e6",gridLineWidth:void 0,tickColor:"#ccd6eb"};a.defaultYAxisOptions={reversedStacks:!0,endOnTick:!0,maxPadding:.05,minPadding:.05,tickPixelInterval:72,showLastLabel:!0,labels:{x:-8},startOnTick:!0,title:{rotation:270,text:"Values"},stackLabels:{animation:{},allowOverlap:!1,enabled:!1,crop:!0,overflow:"justify",formatter:function(){var a=this.axis.chart.numberFormatter;return a(this.total,-1)},style:{color:"#000000",fontSize:"11px",fontWeight:"bold",
textOutline:"1px contrast"}},gridLineWidth:1,lineWidth:0};a.defaultLeftAxisOptions={labels:{x:-15},title:{rotation:270}};a.defaultRightAxisOptions={labels:{x:15},title:{rotation:90}};a.defaultBottomAxisOptions={labels:{autoRotation:[-45],x:0},margin:15,title:{rotation:0}};a.defaultTopAxisOptions={labels:{autoRotation:[-45],x:0},margin:15,title:{rotation:0}}})(a||(a={}));return a});L(a,"Core/Foundation.js",[a["Core/Utilities.js"]],function(a){var u=a.addEvent,A=a.isFunction,G=a.objectEach,x=a.removeEvent,
C;(function(a){a.registerEventOptions=function(a,q){a.eventOptions=a.eventOptions||{};G(q.events,function(m,h){a.eventOptions[h]!==m&&(a.eventOptions[h]&&(x(a,h,a.eventOptions[h]),delete a.eventOptions[h]),A(m)&&(a.eventOptions[h]=m,u(a,h,m)))})}})(C||(C={}));return C});L(a,"Core/Axis/Tick.js",[a["Core/FormatUtilities.js"],a["Core/Globals.js"],a["Core/Utilities.js"]],function(a,t,A){var u=t.deg2rad,x=A.clamp,C=A.correctFloat,I=A.defined,z=A.destroyObjectProperties,q=A.extend,m=A.fireEvent,h=A.isNumber,
d=A.merge,c=A.objectEach,l=A.pick;t=function(){function f(c,a,d,f,h){this.isNewLabel=this.isNew=!0;this.axis=c;this.pos=a;this.type=d||"";this.parameters=h||{};this.tickmarkOffset=this.parameters.tickmarkOffset;this.options=this.parameters.options;m(this,"init");d||f||this.addLabel()}f.prototype.addLabel=function(){var c=this,d=c.axis,f=d.options,r=d.chart,y=d.categories,B=d.logarithmic,n=d.names,k=c.pos,b=l(c.options&&c.options.labels,f.labels),g=d.tickPositions,e=k===g[0],D=k===g[g.length-1],H=
(!b.step||1===b.step)&&1===d.tickInterval;g=g.info;var v=c.label,E;y=this.parameters.category||(y?l(y[k],n[k],k):k);B&&h(y)&&(y=C(B.lin2log(y)));if(d.dateTime)if(g){var O=r.time.resolveDTLFormat(f.dateTimeLabelFormats[!f.grid&&g.higherRanks[k]||g.unitName]);var u=O.main}else h(y)&&(u=d.dateTime.getXDateFormat(y,f.dateTimeLabelFormats||{}));c.isFirst=e;c.isLast=D;var P={axis:d,chart:r,dateTimeLabelFormat:u,isFirst:e,isLast:D,pos:k,tick:c,tickPositionInfo:g,value:y};m(this,"labelFormat",P);var t=function(e){return b.formatter?
b.formatter.call(e,e):b.format?(e.text=d.defaultLabelFormatter.call(e),a.format(b.format,e,r)):d.defaultLabelFormatter.call(e,e)};f=t.call(P,P);var z=O&&O.list;c.shortenLabel=z?function(){for(E=0;E<z.length;E++)if(q(P,{dateTimeLabelFormat:z[E]}),v.attr({text:t.call(P,P)}),v.getBBox().width<d.getSlotWidth(c)-2*b.padding)return;v.attr({text:""})}:void 0;H&&d._addedPlotLB&&c.moveLabel(f,b);I(v)||c.movedLabel?v&&v.textStr!==f&&!H&&(!v.textWidth||b.style.width||v.styles.width||v.css({width:null}),v.attr({text:f}),
v.textPxLength=v.getBBox().width):(c.label=v=c.createLabel({x:0,y:0},f,b),c.rotation=0)};f.prototype.createLabel=function(c,a,f){var h=this.axis,l=h.chart;if(c=I(a)&&f.enabled?l.renderer.text(a,c.x,c.y,f.useHTML).add(h.labelGroup):null)l.styledMode||c.css(d(f.style)),c.textPxLength=c.getBBox().width;return c};f.prototype.destroy=function(){z(this,this.axis)};f.prototype.getPosition=function(c,a,d,f){var h=this.axis,l=h.chart,n=f&&l.oldChartHeight||l.chartHeight;c={x:c?C(h.translate(a+d,null,null,
f)+h.transB):h.left+h.offset+(h.opposite?(f&&l.oldChartWidth||l.chartWidth)-h.right-h.left:0),y:c?n-h.bottom+h.offset-(h.opposite?h.height:0):C(n-h.translate(a+d,null,null,f)-h.transB)};c.y=x(c.y,-1E5,1E5);m(this,"afterGetPosition",{pos:c});return c};f.prototype.getLabelPosition=function(c,a,d,f,h,l,n,k){var b=this.axis,g=b.transA,e=b.isLinked&&b.linkedParent?b.linkedParent.reversed:b.reversed,D=b.staggerLines,p=b.tickRotCorr||{x:0,y:0},v=f||b.reserveSpaceDefault?0:-b.labelOffset*("center"===b.labelAlign?
.5:1),E={},r=h.y;I(r)||(r=0===b.side?d.rotation?-8:-d.getBBox().height:2===b.side?p.y+8:Math.cos(d.rotation*u)*(p.y-d.getBBox(!1,0).height/2));c=c+h.x+v+p.x-(l&&f?l*g*(e?-1:1):0);a=a+r-(l&&!f?l*g*(e?1:-1):0);D&&(d=n/(k||1)%D,b.opposite&&(d=D-d-1),a+=b.labelOffset/D*d);E.x=c;E.y=Math.round(a);m(this,"afterGetLabelPosition",{pos:E,tickmarkOffset:l,index:n});return E};f.prototype.getLabelSize=function(){return this.label?this.label.getBBox()[this.axis.horiz?"height":"width"]:0};f.prototype.getMarkPath=
function(c,a,d,f,h,l){return l.crispLine([["M",c,a],["L",c+(h?0:-d),a+(h?d:0)]],f)};f.prototype.handleOverflow=function(c){var a=this.axis,d=a.options.labels,f=c.x,h=a.chart.chartWidth,m=a.chart.spacing,n=l(a.labelLeft,Math.min(a.pos,m[3]));m=l(a.labelRight,Math.max(a.isRadial?0:a.pos+a.len,h-m[1]));var k=this.label,b=this.rotation,g={left:0,center:.5,right:1}[a.labelAlign||k.attr("align")],e=k.getBBox().width,D=a.getSlotWidth(this),H={},v=D,E=1,w;if(b||"justify"!==d.overflow)0>b&&f-g*e<n?w=Math.round(f/
Math.cos(b*u)-n):0<b&&f+g*e>m&&(w=Math.round((h-f)/Math.cos(b*u)));else if(h=f+(1-g)*e,f-g*e<n?v=c.x+v*(1-g)-n:h>m&&(v=m-c.x+v*g,E=-1),v=Math.min(D,v),v<D&&"center"===a.labelAlign&&(c.x+=E*(D-v-g*(D-Math.min(e,v)))),e>v||a.autoRotation&&(k.styles||{}).width)w=v;w&&(this.shortenLabel?this.shortenLabel():(H.width=Math.floor(w)+"px",(d.style||{}).textOverflow||(H.textOverflow="ellipsis"),k.css(H)))};f.prototype.moveLabel=function(a,d){var f=this,h=f.label,l=f.axis,p=l.reversed,n=!1;h&&h.textStr===a?
(f.movedLabel=h,n=!0,delete f.label):c(l.ticks,function(b){n||b.isNew||b===f||!b.label||b.label.textStr!==a||(f.movedLabel=b.label,n=!0,b.labelPos=f.movedLabel.xy,delete b.label)});if(!n&&(f.labelPos||h)){var k=f.labelPos||h.xy;h=l.horiz?p?0:l.width+l.left:k.x;l=l.horiz?k.y:p?l.width+l.left:0;f.movedLabel=f.createLabel({x:h,y:l},a,d);f.movedLabel&&f.movedLabel.attr({opacity:0})}};f.prototype.render=function(c,a,d){var f=this.axis,h=f.horiz,p=this.pos,n=l(this.tickmarkOffset,f.tickmarkOffset);p=this.getPosition(h,
p,n,a);n=p.x;var k=p.y;f=h&&n===f.pos+f.len||!h&&k===f.pos?-1:1;h=l(d,this.label&&this.label.newOpacity,1);d=l(d,1);this.isActive=!0;this.renderGridLine(a,d,f);this.renderMark(p,d,f);this.renderLabel(p,a,h,c);this.isNew=!1;m(this,"afterRender")};f.prototype.renderGridLine=function(c,a,d){var f=this.axis,h=f.options,p={},n=this.pos,k=this.type,b=l(this.tickmarkOffset,f.tickmarkOffset),g=f.chart.renderer,e=this.gridLine,D=h.gridLineWidth,H=h.gridLineColor,v=h.gridLineDashStyle;"minor"===this.type&&
(D=h.minorGridLineWidth,H=h.minorGridLineColor,v=h.minorGridLineDashStyle);e||(f.chart.styledMode||(p.stroke=H,p["stroke-width"]=D||0,p.dashstyle=v),k||(p.zIndex=1),c&&(a=0),this.gridLine=e=g.path().attr(p).addClass("highcharts-"+(k?k+"-":"")+"grid-line").add(f.gridGroup));if(e&&(d=f.getPlotLinePath({value:n+b,lineWidth:e.strokeWidth()*d,force:"pass",old:c})))e[c||this.isNew?"attr":"animate"]({d:d,opacity:a})};f.prototype.renderMark=function(c,a,d){var f=this.axis,h=f.options,p=f.chart.renderer,n=
this.type,k=f.tickSize(n?n+"Tick":"tick"),b=c.x;c=c.y;var g=l(h["minor"!==n?"tickWidth":"minorTickWidth"],!n&&f.isXAxis?1:0);h=h["minor"!==n?"tickColor":"minorTickColor"];var e=this.mark,D=!e;k&&(f.opposite&&(k[0]=-k[0]),e||(this.mark=e=p.path().addClass("highcharts-"+(n?n+"-":"")+"tick").add(f.axisGroup),f.chart.styledMode||e.attr({stroke:h,"stroke-width":g})),e[D?"attr":"animate"]({d:this.getMarkPath(b,c,k[0],e.strokeWidth()*d,f.horiz,p),opacity:a}))};f.prototype.renderLabel=function(c,a,d,f){var p=
this.axis,m=p.horiz,n=p.options,k=this.label,b=n.labels,g=b.step;p=l(this.tickmarkOffset,p.tickmarkOffset);var e=c.x;c=c.y;var D=!0;k&&h(e)&&(k.xy=c=this.getLabelPosition(e,c,k,m,b,p,f,g),this.isFirst&&!this.isLast&&!n.showFirstLabel||this.isLast&&!this.isFirst&&!n.showLastLabel?D=!1:!m||b.step||b.rotation||a||0===d||this.handleOverflow(c),g&&f%g&&(D=!1),D&&h(c.y)?(c.opacity=d,k[this.isNewLabel?"attr":"animate"](c),this.isNewLabel=!1):(k.attr("y",-9999),this.isNewLabel=!0))};f.prototype.replaceMovedLabel=
function(){var c=this.label,a=this.axis,d=a.reversed;if(c&&!this.isNew){var f=a.horiz?d?a.left:a.width+a.left:c.xy.x;d=a.horiz?c.xy.y:d?a.width+a.top:a.top;c.animate({x:f,y:d,opacity:0},void 0,c.destroy);delete this.label}a.isDirty=!0;this.label=this.movedLabel;delete this.movedLabel};return f}();"";return t});L(a,"Core/Axis/Axis.js",[a["Core/Animation/AnimationUtilities.js"],a["Core/Axis/AxisDefaults.js"],a["Core/Color/Color.js"],a["Core/DefaultOptions.js"],a["Core/Foundation.js"],a["Core/Globals.js"],
a["Core/Axis/Tick.js"],a["Core/Utilities.js"]],function(a,t,A,G,x,C,I,z){var q=a.animObject,m=G.defaultOptions,h=x.registerEventOptions,d=C.deg2rad,c=z.arrayMax,l=z.arrayMin,f=z.clamp,w=z.correctFloat,p=z.defined,K=z.destroyObjectProperties,r=z.erase,y=z.error,B=z.extend,n=z.fireEvent,k=z.getMagnitude,b=z.isArray,g=z.isNumber,e=z.isString,D=z.merge,H=z.normalizeTickInterval,v=z.objectEach,E=z.pick,O=z.relativeLength,u=z.removeEvent,P=z.splat,U=z.syncTimeout;a=function(){function a(b,e){this.zoomEnabled=
this.width=this.visible=this.userOptions=this.translationSlope=this.transB=this.transA=this.top=this.ticks=this.tickRotCorr=this.tickPositions=this.tickmarkOffset=this.tickInterval=this.tickAmount=this.side=this.series=this.right=this.positiveValuesOnly=this.pos=this.pointRangePadding=this.pointRange=this.plotLinesAndBandsGroups=this.plotLinesAndBands=this.paddedTicks=this.overlap=this.options=this.offset=this.names=this.minPixelPadding=this.minorTicks=this.minorTickInterval=this.min=this.maxLabelLength=
this.max=this.len=this.left=this.labelFormatter=this.labelEdge=this.isLinked=this.height=this.hasVisibleSeries=this.hasNames=this.eventOptions=this.coll=this.closestPointRange=this.chart=this.categories=this.bottom=this.alternateBands=void 0;this.init(b,e)}a.prototype.init=function(b,e){var a=e.isX;this.chart=b;this.horiz=b.inverted&&!this.isZAxis?!a:a;this.isXAxis=a;this.coll=this.coll||(a?"xAxis":"yAxis");n(this,"init",{userOptions:e});this.opposite=E(e.opposite,this.opposite);this.side=E(e.side,
this.side,this.horiz?this.opposite?0:2:this.opposite?1:3);this.setOptions(e);var c=this.options,d=c.labels,k=c.type;this.userOptions=e;this.minPixelPadding=0;this.reversed=E(c.reversed,this.reversed);this.visible=c.visible;this.zoomEnabled=c.zoomEnabled;this.hasNames="category"===k||!0===c.categories;this.categories=c.categories||this.hasNames;this.names||(this.names=[],this.names.keys={});this.plotLinesAndBandsGroups={};this.positiveValuesOnly=!!this.logarithmic;this.isLinked=p(c.linkedTo);this.ticks=
{};this.labelEdge=[];this.minorTicks={};this.plotLinesAndBands=[];this.alternateBands={};this.len=0;this.minRange=this.userMinRange=c.minRange||c.maxZoom;this.range=c.range;this.offset=c.offset||0;this.min=this.max=null;e=E(c.crosshair,P(b.options.tooltip.crosshairs)[a?0:1]);this.crosshair=!0===e?{}:e;-1===b.axes.indexOf(this)&&(a?b.axes.splice(b.xAxis.length,0,this):b.axes.push(this),b[this.coll].push(this));this.series=this.series||[];b.inverted&&!this.isZAxis&&a&&"undefined"===typeof this.reversed&&
(this.reversed=!0);this.labelRotation=g(d.rotation)?d.rotation:void 0;h(this,c);n(this,"afterInit")};a.prototype.setOptions=function(b){this.options=D(t.defaultXAxisOptions,"yAxis"===this.coll&&t.defaultYAxisOptions,[t.defaultTopAxisOptions,t.defaultRightAxisOptions,t.defaultBottomAxisOptions,t.defaultLeftAxisOptions][this.side],D(m[this.coll],b));n(this,"afterSetOptions",{userOptions:b})};a.prototype.defaultLabelFormatter=function(b){var e=this.axis;b=this.chart.numberFormatter;var a=g(this.value)?
this.value:NaN,c=e.chart.time,d=this.dateTimeLabelFormat,k=m.lang,f=k.numericSymbols;k=k.numericSymbolMagnitude||1E3;var n=e.logarithmic?Math.abs(a):e.tickInterval,J=f&&f.length;if(e.categories)var h=""+this.value;else if(d)h=c.dateFormat(d,a);else if(J&&1E3<=n)for(;J--&&"undefined"===typeof h;)e=Math.pow(k,J+1),n>=e&&0===10*a%e&&null!==f[J]&&0!==a&&(h=b(a/e,-1)+f[J]);"undefined"===typeof h&&(h=1E4<=Math.abs(a)?b(a,-1):b(a,-1,void 0,""));return h};a.prototype.getSeriesExtremes=function(){var b=this,
e=b.chart,a;n(this,"getSeriesExtremes",null,function(){b.hasVisibleSeries=!1;b.dataMin=b.dataMax=b.threshold=null;b.softThreshold=!b.isXAxis;b.stacking&&b.stacking.buildStacks();b.series.forEach(function(c){if(c.visible||!e.options.chart.ignoreHiddenSeries){var d=c.options,k=d.threshold;b.hasVisibleSeries=!0;b.positiveValuesOnly&&0>=k&&(k=null);if(b.isXAxis){if(d=c.xData,d.length){d=b.logarithmic?d.filter(b.validatePositiveValue):d;a=c.getXExtremes(d);var f=a.min;var n=a.max;g(f)||f instanceof Date||
(d=d.filter(g),a=c.getXExtremes(d),f=a.min,n=a.max);d.length&&(b.dataMin=Math.min(E(b.dataMin,f),f),b.dataMax=Math.max(E(b.dataMax,n),n))}}else if(c=c.applyExtremes(),g(c.dataMin)&&(f=c.dataMin,b.dataMin=Math.min(E(b.dataMin,f),f)),g(c.dataMax)&&(n=c.dataMax,b.dataMax=Math.max(E(b.dataMax,n),n)),p(k)&&(b.threshold=k),!d.softThreshold||b.positiveValuesOnly)b.softThreshold=!1}})});n(this,"afterGetSeriesExtremes")};a.prototype.translate=function(b,e,a,c,d,k){var f=this.linkedParent||this,n=c&&f.old?
f.old.min:f.min,h=f.minPixelPadding;d=(f.isOrdinal||f.brokenAxis&&f.brokenAxis.hasBreaks||f.logarithmic&&d)&&f.lin2val;var v=1,J=0;c=c&&f.old?f.old.transA:f.transA;c||(c=f.transA);a&&(v*=-1,J=f.len);f.reversed&&(v*=-1,J-=v*(f.sector||f.len));e?(b=(b*v+J-h)/c+n,d&&(b=f.lin2val(b))):(d&&(b=f.val2lin(b)),b=g(n)?v*(b-n)*c+J+v*h+(g(k)?c*k:0):void 0);return b};a.prototype.toPixels=function(b,e){return this.translate(b,!1,!this.horiz,null,!0)+(e?0:this.pos)};a.prototype.toValue=function(b,e){return this.translate(b-
(e?0:this.pos),!0,!this.horiz,null,!0)};a.prototype.getPlotLinePath=function(b){function e(b,e,a){if("pass"!==H&&b<e||b>a)H?b=f(b,e,a):O=!0;return b}var a=this,c=a.chart,d=a.left,k=a.top,F=b.old,h=b.value,v=b.lineWidth,J=F&&c.oldChartHeight||c.chartHeight,l=F&&c.oldChartWidth||c.chartWidth,D=a.transB,p=b.translatedValue,H=b.force,m,r,B,q,O;b={value:h,lineWidth:v,old:F,force:H,acrossPanes:b.acrossPanes,translatedValue:p};n(this,"getPlotLinePath",b,function(b){p=E(p,a.translate(h,null,null,F));p=f(p,
-1E5,1E5);m=B=Math.round(p+D);r=q=Math.round(J-p-D);g(p)?a.horiz?(r=k,q=J-a.bottom,m=B=e(m,d,d+a.width)):(m=d,B=l-a.right,r=q=e(r,k,k+a.height)):(O=!0,H=!1);b.path=O&&!H?null:c.renderer.crispLine([["M",m,r],["L",B,q]],v||1)});return b.path};a.prototype.getLinearTickPositions=function(b,e,a){var c=w(Math.floor(e/b)*b);a=w(Math.ceil(a/b)*b);var d=[],g;w(c+b)===c&&(g=20);if(this.single)return[e];for(e=c;e<=a;){d.push(e);e=w(e+b,g);if(e===k)break;var k=e}return d};a.prototype.getMinorTickInterval=function(){var b=
this.options;return!0===b.minorTicks?E(b.minorTickInterval,"auto"):!1===b.minorTicks?null:b.minorTickInterval};a.prototype.getMinorTickPositions=function(){var b=this.options,e=this.tickPositions,a=this.minorTickInterval,c=this.pointRangePadding||0,d=this.min-c;c=this.max+c;var g=c-d,k=[];if(g&&g/a<this.len/3){var f=this.logarithmic;if(f)this.paddedTicks.forEach(function(b,e,c){e&&k.push.apply(k,f.getLogTickPositions(a,c[e-1],c[e],!0))});else if(this.dateTime&&"auto"===this.getMinorTickInterval())k=
k.concat(this.getTimeTicks(this.dateTime.normalizeTimeTickInterval(a),d,c,b.startOfWeek));else for(b=d+(e[0]-d)%a;b<=c&&b!==k[0];b+=a)k.push(b)}0!==k.length&&this.trimTicks(k);return k};a.prototype.adjustForMinRange=function(){var b=this.options,e=this.logarithmic,a=this.min,d=this.max,g=0,k,f,n,h;this.isXAxis&&"undefined"===typeof this.minRange&&!e&&(p(b.min)||p(b.max)||p(b.floor)||p(b.ceiling)?this.minRange=null:(this.series.forEach(function(b){n=b.xData;h=b.xIncrement?1:n.length-1;if(1<n.length)for(k=
h;0<k;k--)if(f=n[k]-n[k-1],!g||f<g)g=f}),this.minRange=Math.min(5*g,this.dataMax-this.dataMin)));if(d-a<this.minRange){var v=this.dataMax-this.dataMin>=this.minRange;var D=this.minRange;var H=(D-d+a)/2;H=[a-H,E(b.min,a-H)];v&&(H[2]=this.logarithmic?this.logarithmic.log2lin(this.dataMin):this.dataMin);a=c(H);d=[a+D,E(b.max,a+D)];v&&(d[2]=e?e.log2lin(this.dataMax):this.dataMax);d=l(d);d-a<D&&(H[0]=d-D,H[1]=E(b.min,d-D),a=c(H))}this.min=a;this.max=d};a.prototype.getClosest=function(){var b;this.categories?
b=1:this.series.forEach(function(e){var a=e.closestPointRange,c=e.visible||!e.chart.options.chart.ignoreHiddenSeries;!e.noSharedTooltip&&p(a)&&c&&(b=p(b)?Math.min(b,a):a)});return b};a.prototype.nameToX=function(e){var a=b(this.categories),c=a?this.categories:this.names,d=e.options.x;e.series.requireSorting=!1;p(d)||(d=this.options.uniqueNames?a?c.indexOf(e.name):E(c.keys[e.name],-1):e.series.autoIncrement());if(-1===d){if(!a)var g=c.length}else g=d;"undefined"!==typeof g&&(this.names[g]=e.name,this.names.keys[e.name]=
g);return g};a.prototype.updateNames=function(){var b=this,e=this.names;0<e.length&&(Object.keys(e.keys).forEach(function(b){delete e.keys[b]}),e.length=0,this.minRange=this.userMinRange,(this.series||[]).forEach(function(e){e.xIncrement=null;if(!e.points||e.isDirtyData)b.max=Math.max(b.max,e.xData.length-1),e.processData(),e.generatePoints();e.data.forEach(function(a,c){if(a&&a.options&&"undefined"!==typeof a.name){var d=b.nameToX(a);"undefined"!==typeof d&&d!==a.x&&(a.x=d,e.xData[c]=d)}})}))};a.prototype.setAxisTranslation=
function(){var b=this,a=b.max-b.min,c=b.linkedParent,d=!!b.categories,g=b.isXAxis,k=b.axisPointRange||0,f=0,h=0,v=b.transA;if(g||d||k){var D=b.getClosest();c?(f=c.minPointOffset,h=c.pointRangePadding):b.series.forEach(function(a){var c=d?1:g?E(a.options.pointRange,D,0):b.axisPointRange||0,n=a.options.pointPlacement;k=Math.max(k,c);if(!b.single||d)a=a.is("xrange")?!g:g,f=Math.max(f,a&&e(n)?0:c/2),h=Math.max(h,a&&"on"===n?0:c)});c=b.ordinal&&b.ordinal.slope&&D?b.ordinal.slope/D:1;b.minPointOffset=f*=
c;b.pointRangePadding=h*=c;b.pointRange=Math.min(k,b.single&&d?1:a);g&&(b.closestPointRange=D)}b.translationSlope=b.transA=v=b.staticScale||b.len/(a+h||1);b.transB=b.horiz?b.left:b.bottom;b.minPixelPadding=v*f;n(this,"afterSetAxisTranslation")};a.prototype.minFromRange=function(){return this.max-this.range};a.prototype.setTickInterval=function(b){var e=this.chart,a=this.logarithmic,c=this.options,d=this.isXAxis,f=this.isLinked,h=c.tickPixelInterval,v=this.categories,D=this.softThreshold,l=c.maxPadding,
J=c.minPadding,m=g(c.tickInterval)&&0<=c.tickInterval?c.tickInterval:void 0,r=g(this.threshold)?this.threshold:null;this.dateTime||v||f||this.getTickAmount();var B=E(this.userMin,c.min);var q=E(this.userMax,c.max);if(f){this.linkedParent=e[this.coll][c.linkedTo];var O=this.linkedParent.getExtremes();this.min=E(O.min,O.dataMin);this.max=E(O.max,O.dataMax);c.type!==this.linkedParent.options.type&&y(11,1,e)}else{if(D&&p(r))if(this.dataMin>=r)O=r,J=0;else if(this.dataMax<=r){var K=r;l=0}this.min=E(B,
O,this.dataMin);this.max=E(q,K,this.dataMax)}a&&(this.positiveValuesOnly&&!b&&0>=Math.min(this.min,E(this.dataMin,this.min))&&y(10,1,e),this.min=w(a.log2lin(this.min),16),this.max=w(a.log2lin(this.max),16));this.range&&p(this.max)&&(this.userMin=this.min=B=Math.max(this.dataMin,this.minFromRange()),this.userMax=q=this.max,this.range=null);n(this,"foundExtremes");this.beforePadding&&this.beforePadding();this.adjustForMinRange();!(v||this.axisPointRange||this.stacking&&this.stacking.usePercentage||
f)&&p(this.min)&&p(this.max)&&(e=this.max-this.min)&&(!p(B)&&J&&(this.min-=e*J),!p(q)&&l&&(this.max+=e*l));g(this.userMin)||(g(c.softMin)&&c.softMin<this.min&&(this.min=B=c.softMin),g(c.floor)&&(this.min=Math.max(this.min,c.floor)));g(this.userMax)||(g(c.softMax)&&c.softMax>this.max&&(this.max=q=c.softMax),g(c.ceiling)&&(this.max=Math.min(this.max,c.ceiling)));D&&p(this.dataMin)&&(r=r||0,!p(B)&&this.min<r&&this.dataMin>=r?this.min=this.options.minRange?Math.min(r,this.max-this.minRange):r:!p(q)&&
this.max>r&&this.dataMax<=r&&(this.max=this.options.minRange?Math.max(r,this.min+this.minRange):r));g(this.min)&&g(this.max)&&!this.chart.polar&&this.min>this.max&&(p(this.options.min)?this.max=this.min:p(this.options.max)&&(this.min=this.max));this.tickInterval=this.min===this.max||"undefined"===typeof this.min||"undefined"===typeof this.max?1:f&&this.linkedParent&&!m&&h===this.linkedParent.options.tickPixelInterval?m=this.linkedParent.tickInterval:E(m,this.tickAmount?(this.max-this.min)/Math.max(this.tickAmount-
1,1):void 0,v?1:(this.max-this.min)*h/Math.max(this.len,h));if(d&&!b){var u=this.min!==(this.old&&this.old.min)||this.max!==(this.old&&this.old.max);this.series.forEach(function(b){b.forceCrop=b.forceCropping&&b.forceCropping();b.processData(u)});n(this,"postProcessData",{hasExtemesChanged:u})}this.setAxisTranslation();n(this,"initialAxisTranslation");this.pointRange&&!m&&(this.tickInterval=Math.max(this.pointRange,this.tickInterval));b=E(c.minTickInterval,this.dateTime&&!this.series.some(function(b){return b.noSharedTooltip})?
this.closestPointRange:0);!m&&this.tickInterval<b&&(this.tickInterval=b);this.dateTime||this.logarithmic||m||(this.tickInterval=H(this.tickInterval,void 0,k(this.tickInterval),E(c.allowDecimals,.5>this.tickInterval||void 0!==this.tickAmount),!!this.tickAmount));this.tickAmount||(this.tickInterval=this.unsquish());this.setTickPositions()};a.prototype.setTickPositions=function(){var b=this.options,e=b.tickPositions,a=this.getMinorTickInterval(),c=this.hasVerticalPanning(),d="colorAxis"===this.coll,
g=(d||!c)&&b.startOnTick;c=(d||!c)&&b.endOnTick;d=b.tickPositioner;this.tickmarkOffset=this.categories&&"between"===b.tickmarkPlacement&&1===this.tickInterval?.5:0;this.minorTickInterval="auto"===a&&this.tickInterval?this.tickInterval/5:a;this.single=this.min===this.max&&p(this.min)&&!this.tickAmount&&(parseInt(this.min,10)===this.min||!1!==b.allowDecimals);this.tickPositions=a=e&&e.slice();!a&&(this.ordinal&&this.ordinal.positions||!((this.max-this.min)/this.tickInterval>Math.max(2*this.len,200))?
a=this.dateTime?this.getTimeTicks(this.dateTime.normalizeTimeTickInterval(this.tickInterval,b.units),this.min,this.max,b.startOfWeek,this.ordinal&&this.ordinal.positions,this.closestPointRange,!0):this.logarithmic?this.logarithmic.getLogTickPositions(this.tickInterval,this.min,this.max):this.getLinearTickPositions(this.tickInterval,this.min,this.max):(a=[this.min,this.max],y(19,!1,this.chart)),a.length>this.len&&(a=[a[0],a.pop()],a[0]===a[1]&&(a.length=1)),this.tickPositions=a,d&&(d=d.apply(this,
[this.min,this.max])))&&(this.tickPositions=a=d);this.paddedTicks=a.slice(0);this.trimTicks(a,g,c);this.isLinked||(this.single&&2>a.length&&!this.categories&&!this.series.some(function(b){return b.is("heatmap")&&"between"===b.options.pointPlacement})&&(this.min-=.5,this.max+=.5),e||d||this.adjustTickAmount());n(this,"afterSetTickPositions")};a.prototype.trimTicks=function(b,e,a){var c=b[0],d=b[b.length-1],g=!this.isOrdinal&&this.minPointOffset||0;n(this,"trimTicks");if(!this.isLinked){if(e&&-Infinity!==
c)this.min=c;else for(;this.min-g>b[0];)b.shift();if(a)this.max=d;else for(;this.max+g<b[b.length-1];)b.pop();0===b.length&&p(c)&&!this.options.tickPositions&&b.push((d+c)/2)}};a.prototype.alignToOthers=function(){var b={},e=this.options,a;!1!==this.chart.options.chart.alignTicks&&e.alignTicks&&!1!==e.startOnTick&&!1!==e.endOnTick&&!this.logarithmic&&this.chart[this.coll].forEach(function(e){var c=e.options;c=[e.horiz?c.left:c.top,c.width,c.height,c.pane].join();e.series.length&&(b[c]?a=!0:b[c]=1)});
return a};a.prototype.getTickAmount=function(){var b=this.options,e=b.tickPixelInterval,a=b.tickAmount;!p(b.tickInterval)&&!a&&this.len<e&&!this.isRadial&&!this.logarithmic&&b.startOnTick&&b.endOnTick&&(a=2);!a&&this.alignToOthers()&&(a=Math.ceil(this.len/e)+1);4>a&&(this.finalTickAmt=a,a=5);this.tickAmount=a};a.prototype.adjustTickAmount=function(){var b=this.options,e=this.tickInterval,a=this.tickPositions,c=this.tickAmount,d=this.finalTickAmt,k=a&&a.length,f=E(this.threshold,this.softThreshold?
0:null);if(this.hasData()&&g(this.min)&&g(this.max)){if(k<c){for(;a.length<c;)a.length%2||this.min===f?a.push(w(a[a.length-1]+e)):a.unshift(w(a[0]-e));this.transA*=(k-1)/(c-1);this.min=b.startOnTick?a[0]:Math.min(this.min,a[0]);this.max=b.endOnTick?a[a.length-1]:Math.max(this.max,a[a.length-1])}else k>c&&(this.tickInterval*=2,this.setTickPositions());if(p(d)){for(e=b=a.length;e--;)(3===d&&1===e%2||2>=d&&0<e&&e<b-1)&&a.splice(e,1);this.finalTickAmt=void 0}}};a.prototype.setScale=function(){var b=!1,
e=!1;this.series.forEach(function(a){b=b||a.isDirtyData||a.isDirty;e=e||a.xAxis&&a.xAxis.isDirty||!1});this.setAxisSize();var a=this.len!==(this.old&&this.old.len);a||b||e||this.isLinked||this.forceRedraw||this.userMin!==(this.old&&this.old.userMin)||this.userMax!==(this.old&&this.old.userMax)||this.alignToOthers()?(this.stacking&&this.stacking.resetStacks(),this.forceRedraw=!1,this.getSeriesExtremes(),this.setTickInterval(),this.isDirty||(this.isDirty=a||this.min!==(this.old&&this.old.min)||this.max!==
(this.old&&this.old.max))):this.stacking&&this.stacking.cleanStacks();b&&this.panningState&&(this.panningState.isDirty=!0);n(this,"afterSetScale")};a.prototype.setExtremes=function(b,e,a,c,d){var g=this,k=g.chart;a=E(a,!0);g.series.forEach(function(b){delete b.kdTree});d=B(d,{min:b,max:e});n(g,"setExtremes",d,function(){g.userMin=b;g.userMax=e;g.eventArgs=d;a&&k.redraw(c)})};a.prototype.zoom=function(b,e){var a=this,c=this.dataMin,d=this.dataMax,g=this.options,k=Math.min(c,E(g.min,c)),f=Math.max(d,
E(g.max,d));b={newMin:b,newMax:e};n(this,"zoom",b,function(b){var e=b.newMin,g=b.newMax;if(e!==a.min||g!==a.max)a.allowZoomOutside||(p(c)&&(e<k&&(e=k),e>f&&(e=f)),p(d)&&(g<k&&(g=k),g>f&&(g=f))),a.displayBtn="undefined"!==typeof e||"undefined"!==typeof g,a.setExtremes(e,g,!1,void 0,{trigger:"zoom"});b.zoomed=!0});return b.zoomed};a.prototype.setAxisSize=function(){var b=this.chart,e=this.options,a=e.offsets||[0,0,0,0],c=this.horiz,d=this.width=Math.round(O(E(e.width,b.plotWidth-a[3]+a[1]),b.plotWidth)),
g=this.height=Math.round(O(E(e.height,b.plotHeight-a[0]+a[2]),b.plotHeight)),k=this.top=Math.round(O(E(e.top,b.plotTop+a[0]),b.plotHeight,b.plotTop));e=this.left=Math.round(O(E(e.left,b.plotLeft+a[3]),b.plotWidth,b.plotLeft));this.bottom=b.chartHeight-g-k;this.right=b.chartWidth-d-e;this.len=Math.max(c?d:g,0);this.pos=c?e:k};a.prototype.getExtremes=function(){var b=this.logarithmic;return{min:b?w(b.lin2log(this.min)):this.min,max:b?w(b.lin2log(this.max)):this.max,dataMin:this.dataMin,dataMax:this.dataMax,
userMin:this.userMin,userMax:this.userMax}};a.prototype.getThreshold=function(b){var e=this.logarithmic,a=e?e.lin2log(this.min):this.min;e=e?e.lin2log(this.max):this.max;null===b||-Infinity===b?b=a:Infinity===b?b=e:a>b?b=a:e<b&&(b=e);return this.translate(b,0,1,0,1)};a.prototype.autoLabelAlign=function(b){var e=(E(b,0)-90*this.side+720)%360;b={align:"center"};n(this,"autoLabelAlign",b,function(b){15<e&&165>e?b.align="right":195<e&&345>e&&(b.align="left")});return b.align};a.prototype.tickSize=function(b){var e=
this.options,a=E(e["tick"===b?"tickWidth":"minorTickWidth"],"tick"===b&&this.isXAxis&&!this.categories?1:0),c=e["tick"===b?"tickLength":"minorTickLength"];if(a&&c){"inside"===e[b+"Position"]&&(c=-c);var d=[c,a]}b={tickSize:d};n(this,"afterTickSize",b);return b.tickSize};a.prototype.labelMetrics=function(){var b=this.tickPositions&&this.tickPositions[0]||0;return this.chart.renderer.fontMetrics(this.options.labels.style.fontSize,this.ticks[b]&&this.ticks[b].label)};a.prototype.unsquish=function(){var b=
this.options.labels,e=this.horiz,a=this.tickInterval,c=this.len/(((this.categories?1:0)+this.max-this.min)/a),k=b.rotation,f=this.labelMetrics(),n=Math.max(this.max-this.min,0),h=function(b){var e=b/(c||1);e=1<e?Math.ceil(e):1;e*a>n&&Infinity!==b&&Infinity!==c&&n&&(e=Math.ceil(n/a));return w(e*a)},v=a,D,l,H=Number.MAX_VALUE;if(e){if(!b.staggerLines&&!b.step)if(g(k))var p=[k];else c<b.autoRotationLimit&&(p=b.autoRotation);p&&p.forEach(function(b){if(b===k||b&&-90<=b&&90>=b){l=h(Math.abs(f.h/Math.sin(d*
b)));var e=l+Math.abs(b/360);e<H&&(H=e,D=b,v=l)}})}else b.step||(v=h(f.h));this.autoRotation=p;this.labelRotation=E(D,g(k)?k:0);return v};a.prototype.getSlotWidth=function(b){var e=this.chart,a=this.horiz,c=this.options.labels,d=Math.max(this.tickPositions.length-(this.categories?0:1),1),k=e.margin[3];if(b&&g(b.slotWidth))return b.slotWidth;if(a&&2>c.step)return c.rotation?0:(this.staggerLines||1)*this.len/d;if(!a){b=c.style.width;if(void 0!==b)return parseInt(String(b),10);if(k)return k-e.spacing[3]}return.33*
e.chartWidth};a.prototype.renderUnsquish=function(){var b=this.chart,a=b.renderer,c=this.tickPositions,d=this.ticks,g=this.options.labels,k=g.style,f=this.horiz,n=this.getSlotWidth(),h=Math.max(1,Math.round(n-2*g.padding)),v={},D=this.labelMetrics(),l=k.textOverflow,H=0;e(g.rotation)||(v.rotation=g.rotation||0);c.forEach(function(b){b=d[b];b.movedLabel&&b.replaceMovedLabel();b&&b.label&&b.label.textPxLength>H&&(H=b.label.textPxLength)});this.maxLabelLength=H;if(this.autoRotation)H>h&&H>D.h?v.rotation=
this.labelRotation:this.labelRotation=0;else if(n){var p=h;if(!l){var m="clip";for(h=c.length;!f&&h--;){var E=c[h];if(E=d[E].label)E.styles&&"ellipsis"===E.styles.textOverflow?E.css({textOverflow:"clip"}):E.textPxLength>n&&E.css({width:n+"px"}),E.getBBox().height>this.len/c.length-(D.h-D.f)&&(E.specificTextOverflow="ellipsis")}}}v.rotation&&(p=H>.5*b.chartHeight?.33*b.chartHeight:H,l||(m="ellipsis"));if(this.labelAlign=g.align||this.autoLabelAlign(this.labelRotation))v.align=this.labelAlign;c.forEach(function(b){var e=
(b=d[b])&&b.label,a=k.width,c={};e&&(e.attr(v),b.shortenLabel?b.shortenLabel():p&&!a&&"nowrap"!==k.whiteSpace&&(p<e.textPxLength||"SPAN"===e.element.tagName)?(c.width=p+"px",l||(c.textOverflow=e.specificTextOverflow||m),e.css(c)):e.styles&&e.styles.width&&!c.width&&!a&&e.css({width:null}),delete e.specificTextOverflow,b.rotation=v.rotation)},this);this.tickRotCorr=a.rotCorr(D.b,this.labelRotation||0,0!==this.side)};a.prototype.hasData=function(){return this.series.some(function(b){return b.hasData()})||
this.options.showEmpty&&p(this.min)&&p(this.max)};a.prototype.addTitle=function(b){var e=this.chart.renderer,a=this.horiz,c=this.opposite,d=this.options.title,g=this.chart.styledMode,k;this.axisTitle||((k=d.textAlign)||(k=(a?{low:"left",middle:"center",high:"right"}:{low:c?"right":"left",middle:"center",high:c?"left":"right"})[d.align]),this.axisTitle=e.text(d.text||"",0,0,d.useHTML).attr({zIndex:7,rotation:d.rotation,align:k}).addClass("highcharts-axis-title"),g||this.axisTitle.css(D(d.style)),this.axisTitle.add(this.axisGroup),
this.axisTitle.isNew=!0);g||d.style.width||this.isRadial||this.axisTitle.css({width:this.len+"px"});this.axisTitle[b?"show":"hide"](b)};a.prototype.generateTick=function(b){var e=this.ticks;e[b]?e[b].addLabel():e[b]=new I(this,b)};a.prototype.getOffset=function(){var b=this,e=this,a=e.chart,c=e.horiz,d=e.options,g=e.side,k=e.ticks,f=e.tickPositions,h=e.coll,D=e.axisParent,l=a.renderer,H=a.inverted&&!e.isZAxis?[1,0,3,2][g]:g,m=e.hasData(),r=d.title,B=d.labels,q=a.axisOffset;a=a.clipOffset;var O=[-1,
1,1,-1][g],w=d.className,y,K=0,ja=0,ca=0;e.showAxis=y=m||d.showEmpty;e.staggerLines=e.horiz&&B.staggerLines||void 0;if(!e.axisGroup){var u=function(e,a,c){return l.g(e).attr({zIndex:c}).addClass("highcharts-"+h.toLowerCase()+a+" "+(b.isRadial?"highcharts-radial-axis"+a+" ":"")+(w||"")).add(D)};e.gridGroup=u("grid","-grid",d.gridZIndex);e.axisGroup=u("axis","",d.zIndex);e.labelGroup=u("axis-labels","-labels",B.zIndex)}m||e.isLinked?(f.forEach(function(b){e.generateTick(b)}),e.renderUnsquish(),e.reserveSpaceDefault=
0===g||2===g||{1:"left",3:"right"}[g]===e.labelAlign,E(B.reserveSpace,"center"===e.labelAlign?!0:null,e.reserveSpaceDefault)&&f.forEach(function(b){ca=Math.max(k[b].getLabelSize(),ca)}),e.staggerLines&&(ca*=e.staggerLines),e.labelOffset=ca*(e.opposite?-1:1)):v(k,function(b,e){b.destroy();delete k[e]});if(r&&r.text&&!1!==r.enabled&&(e.addTitle(y),y&&!1!==r.reserveSpace)){e.titleOffset=K=e.axisTitle.getBBox()[c?"height":"width"];var P=r.offset;ja=p(P)?0:E(r.margin,c?5:10)}e.renderLine();e.offset=O*
E(d.offset,q[g]?q[g]+(d.margin||0):0);e.tickRotCorr=e.tickRotCorr||{x:0,y:0};r=0===g?-e.labelMetrics().h:2===g?e.tickRotCorr.y:0;m=Math.abs(ca)+ja;ca&&(m=m-r+O*(c?E(B.y,e.tickRotCorr.y+8*O):B.x));e.axisTitleMargin=E(P,m);e.getMaxLabelDimensions&&(e.maxLabelDimensions=e.getMaxLabelDimensions(k,f));"colorAxis"!==h&&(c=this.tickSize("tick"),q[g]=Math.max(q[g],(e.axisTitleMargin||0)+K+O*e.offset,m,f&&f.length&&c?c[0]+O*e.offset:0),d=!e.axisLine||d.offset?0:2*Math.floor(e.axisLine.strokeWidth()/2),a[H]=
Math.max(a[H],d));n(this,"afterGetOffset")};a.prototype.getLinePath=function(b){var e=this.chart,a=this.opposite,c=this.offset,d=this.horiz,g=this.left+(a?this.width:0)+c;c=e.chartHeight-this.bottom-(a?this.height:0)+c;a&&(b*=-1);return e.renderer.crispLine([["M",d?this.left:g,d?c:this.top],["L",d?e.chartWidth-this.right:g,d?c:e.chartHeight-this.bottom]],b)};a.prototype.renderLine=function(){this.axisLine||(this.axisLine=this.chart.renderer.path().addClass("highcharts-axis-line").add(this.axisGroup),
this.chart.styledMode||this.axisLine.attr({stroke:this.options.lineColor,"stroke-width":this.options.lineWidth,zIndex:7}))};a.prototype.getTitlePosition=function(){var b=this.horiz,e=this.left,a=this.top,c=this.len,d=this.options.title,g=b?e:a,k=this.opposite,f=this.offset,h=d.x,v=d.y,D=this.axisTitle,l=this.chart.renderer.fontMetrics(d.style.fontSize,D);D=Math.max(D.getBBox(null,0).height-l.h-1,0);c={low:g+(b?0:c),middle:g+c/2,high:g+(b?c:0)}[d.align];e=(b?a+this.height:e)+(b?1:-1)*(k?-1:1)*this.axisTitleMargin+
[-D,D,l.f,-D][this.side];b={x:b?c+h:e+(k?this.width:0)+f+h,y:b?e+v-(k?this.height:0)+f:c+v};n(this,"afterGetTitlePosition",{titlePosition:b});return b};a.prototype.renderMinorTick=function(b,e){var a=this.minorTicks;a[b]||(a[b]=new I(this,b,"minor"));e&&a[b].isNew&&a[b].render(null,!0);a[b].render(null,!1,1)};a.prototype.renderTick=function(b,e,a){var c=this.ticks;if(!this.isLinked||b>=this.min&&b<=this.max||this.grid&&this.grid.isColumn)c[b]||(c[b]=new I(this,b)),a&&c[b].isNew&&c[b].render(e,!0,
-1),c[b].render(e)};a.prototype.render=function(){var b=this,e=b.chart,a=b.logarithmic,c=b.options,d=b.isLinked,k=b.tickPositions,f=b.axisTitle,h=b.ticks,D=b.minorTicks,l=b.alternateBands,H=c.stackLabels,p=c.alternateGridColor,m=b.tickmarkOffset,E=b.axisLine,r=b.showAxis,B=q(e.renderer.globalAnimation),O,w;b.labelEdge.length=0;b.overlap=!1;[h,D,l].forEach(function(b){v(b,function(b){b.isActive=!1})});if(b.hasData()||d){var y=b.chart.hasRendered&&b.old&&g(b.old.min);b.minorTickInterval&&!b.categories&&
b.getMinorTickPositions().forEach(function(e){b.renderMinorTick(e,y)});k.length&&(k.forEach(function(e,a){b.renderTick(e,a,y)}),m&&(0===b.min||b.single)&&(h[-1]||(h[-1]=new I(b,-1,null,!0)),h[-1].render(-1)));p&&k.forEach(function(c,d){w="undefined"!==typeof k[d+1]?k[d+1]+m:b.max-m;0===d%2&&c<b.max&&w<=b.max+(e.polar?-m:m)&&(l[c]||(l[c]=new C.PlotLineOrBand(b)),O=c+m,l[c].options={from:a?a.lin2log(O):O,to:a?a.lin2log(w):w,color:p,className:"highcharts-alternate-grid"},l[c].render(),l[c].isActive=
!0)});b._addedPlotLB||(b._addedPlotLB=!0,(c.plotLines||[]).concat(c.plotBands||[]).forEach(function(e){b.addPlotBandOrLine(e)}))}[h,D,l].forEach(function(b){var a=[],c=B.duration;v(b,function(b,e){b.isActive||(b.render(e,!1,0),b.isActive=!1,a.push(e))});U(function(){for(var e=a.length;e--;)b[a[e]]&&!b[a[e]].isActive&&(b[a[e]].destroy(),delete b[a[e]])},b!==l&&e.hasRendered&&c?c:0)});E&&(E[E.isPlaced?"animate":"attr"]({d:this.getLinePath(E.strokeWidth())}),E.isPlaced=!0,E[r?"show":"hide"](r));f&&r&&
(c=b.getTitlePosition(),g(c.y)?(f[f.isNew?"attr":"animate"](c),f.isNew=!1):(f.attr("y",-9999),f.isNew=!0));H&&H.enabled&&b.stacking&&b.stacking.renderStackTotals();b.old={len:b.len,max:b.max,min:b.min,transA:b.transA,userMax:b.userMax,userMin:b.userMin};b.isDirty=!1;n(this,"afterRender")};a.prototype.redraw=function(){this.visible&&(this.render(),this.plotLinesAndBands.forEach(function(b){b.render()}));this.series.forEach(function(b){b.isDirty=!0})};a.prototype.getKeepProps=function(){return this.keepProps||
a.keepProps};a.prototype.destroy=function(b){var e=this,a=e.plotLinesAndBands,c=this.eventOptions;n(this,"destroy",{keepEvents:b});b||u(e);[e.ticks,e.minorTicks,e.alternateBands].forEach(function(b){K(b)});if(a)for(b=a.length;b--;)a[b].destroy();"axisLine axisTitle axisGroup gridGroup labelGroup cross scrollbar".split(" ").forEach(function(b){e[b]&&(e[b]=e[b].destroy())});for(var d in e.plotLinesAndBandsGroups)e.plotLinesAndBandsGroups[d]=e.plotLinesAndBandsGroups[d].destroy();v(e,function(b,a){-1===
e.getKeepProps().indexOf(a)&&delete e[a]});this.eventOptions=c};a.prototype.drawCrosshair=function(b,e){var a=this.crosshair,c=E(a&&a.snap,!0),d=this.chart,g,k=this.cross;n(this,"drawCrosshair",{e:b,point:e});b||(b=this.cross&&this.cross.e);if(a&&!1!==(p(e)||!c)){c?p(e)&&(g=E("colorAxis"!==this.coll?e.crosshairPos:null,this.isXAxis?e.plotX:this.len-e.plotY)):g=b&&(this.horiz?b.chartX-this.pos:this.len-b.chartY+this.pos);if(p(g)){var f={value:e&&(this.isXAxis?e.x:E(e.stackY,e.y)),translatedValue:g};
d.polar&&B(f,{isCrosshair:!0,chartX:b&&b.chartX,chartY:b&&b.chartY,point:e});f=this.getPlotLinePath(f)||null}if(!p(f)){this.hideCrosshair();return}c=this.categories&&!this.isRadial;k||(this.cross=k=d.renderer.path().addClass("highcharts-crosshair highcharts-crosshair-"+(c?"category ":"thin ")+(a.className||"")).attr({zIndex:E(a.zIndex,2)}).add(),d.styledMode||(k.attr({stroke:a.color||(c?A.parse("#ccd6eb").setOpacity(.25).get():"#cccccc"),"stroke-width":E(a.width,1)}).css({"pointer-events":"none"}),
a.dashStyle&&k.attr({dashstyle:a.dashStyle})));k.show().attr({d:f});c&&!a.width&&k.attr({"stroke-width":this.transA});this.cross.e=b}else this.hideCrosshair();n(this,"afterDrawCrosshair",{e:b,point:e})};a.prototype.hideCrosshair=function(){this.cross&&this.cross.hide();n(this,"afterHideCrosshair")};a.prototype.hasVerticalPanning=function(){var b=this.chart.options.chart.panning;return!!(b&&b.enabled&&/y/.test(b.type))};a.prototype.validatePositiveValue=function(b){return g(b)&&0<b};a.prototype.update=
function(b,e){var a=this.chart;b=D(this.userOptions,b);this.destroy(!0);this.init(a,b);a.isDirtyBox=!0;E(e,!0)&&a.redraw()};a.prototype.remove=function(b){for(var e=this.chart,a=this.coll,c=this.series,d=c.length;d--;)c[d]&&c[d].remove(!1);r(e.axes,this);r(e[a],this);e[a].forEach(function(b,e){b.options.index=b.userOptions.index=e});this.destroy();e.isDirtyBox=!0;E(b,!0)&&e.redraw()};a.prototype.setTitle=function(b,e){this.update({title:b},e)};a.prototype.setCategories=function(b,e){this.update({categories:b},
e)};a.defaultOptions=t.defaultXAxisOptions;a.keepProps="extKey hcEvents names series userMax userMin".split(" ");return a}();"";return a});L(a,"Core/Axis/DateTimeAxis.js",[a["Core/Utilities.js"]],function(a){var u=a.addEvent,A=a.getMagnitude,G=a.normalizeTickInterval,x=a.timeUnits,C;(function(a){function t(){return this.chart.time.getTimeTicks.apply(this.chart.time,arguments)}function q(a){"datetime"!==a.userOptions.type?this.dateTime=void 0:this.dateTime||(this.dateTime=new h(this))}var m=[];a.compose=
function(a){-1===m.indexOf(a)&&(m.push(a),a.keepProps.push("dateTime"),a.prototype.getTimeTicks=t,u(a,"init",q));return a};var h=function(){function a(a){this.axis=a}a.prototype.normalizeTimeTickInterval=function(a,d){var c=d||[["millisecond",[1,2,5,10,20,25,50,100,200,500]],["second",[1,2,5,10,15,30]],["minute",[1,2,5,10,15,30]],["hour",[1,2,3,4,6,8,12]],["day",[1,2]],["week",[1,2]],["month",[1,2,3,4,6]],["year",null]];d=c[c.length-1];var h=x[d[0]],l=d[1],m;for(m=0;m<c.length&&!(d=c[m],h=x[d[0]],
l=d[1],c[m+1]&&a<=(h*l[l.length-1]+x[c[m+1][0]])/2);m++);h===x.year&&a<5*h&&(l=[1,2,5]);a=G(a/h,l,"year"===d[0]?Math.max(A(a/h),1):1);return{unitRange:h,count:a,unitName:d[0]}};a.prototype.getXDateFormat=function(a,d){var c=this.axis;return c.closestPointRange?c.chart.time.getDateFormat(c.closestPointRange,a,c.options.startOfWeek,d)||d.year:d.day};return a}();a.Additions=h})(C||(C={}));return C});L(a,"Core/Axis/LogarithmicAxis.js",[a["Core/Utilities.js"]],function(a){var u=a.addEvent,A=a.getMagnitude,
G=a.normalizeTickInterval,x=a.pick,C;(function(a){function t(a){var c=this.logarithmic;"logarithmic"!==a.userOptions.type?this.logarithmic=void 0:c||(this.logarithmic=new h(this))}function q(){var a=this.logarithmic;a&&(this.lin2val=function(c){return a.lin2log(c)},this.val2lin=function(c){return a.log2lin(c)})}var m=[];a.compose=function(a){-1===m.indexOf(a)&&(m.push(a),a.keepProps.push("logarithmic"),u(a,"init",t),u(a,"afterInit",q));return a};var h=function(){function a(a){this.axis=a}a.prototype.getLogTickPositions=
function(a,d,f,h){var c=this.axis,l=c.len,m=c.options,q=[];h||(this.minorAutoInterval=void 0);if(.5<=a)a=Math.round(a),q=c.getLinearTickPositions(a,d,f);else if(.08<=a){var B=Math.floor(d),n,k=m=void 0;for(l=.3<a?[1,2,4]:.15<a?[1,2,4,6,8]:[1,2,3,4,5,6,7,8,9];B<f+1&&!k;B++){var b=l.length;for(n=0;n<b&&!k;n++){var g=this.log2lin(this.lin2log(B)*l[n]);g>d&&(!h||m<=f)&&"undefined"!==typeof m&&q.push(m);m>f&&(k=!0);m=g}}}else d=this.lin2log(d),f=this.lin2log(f),a=h?c.getMinorTickInterval():m.tickInterval,
a=x("auto"===a?null:a,this.minorAutoInterval,m.tickPixelInterval/(h?5:1)*(f-d)/((h?l/c.tickPositions.length:l)||1)),a=G(a,void 0,A(a)),q=c.getLinearTickPositions(a,d,f).map(this.log2lin),h||(this.minorAutoInterval=a/5);h||(c.tickInterval=a);return q};a.prototype.lin2log=function(a){return Math.pow(10,a)};a.prototype.log2lin=function(a){return Math.log(a)/Math.LN10};return a}();a.Additions=h})(C||(C={}));return C});L(a,"Core/Axis/PlotLineOrBand/PlotLineOrBandAxis.js",[a["Core/Utilities.js"]],function(a){var u=
a.erase,A=a.extend,G=a.isNumber,x;(function(a){var t=[],z;a.compose=function(a,h){z||(z=a);-1===t.indexOf(h)&&(t.push(h),A(h.prototype,q.prototype));return h};var q=function(){function a(){}a.prototype.getPlotBandPath=function(a,d,c){void 0===c&&(c=this.options);var h=this.getPlotLinePath({value:d,force:!0,acrossPanes:c.acrossPanes}),f=[],m=this.horiz;d=!G(this.min)||!G(this.max)||a<this.min&&d<this.min||a>this.max&&d>this.max;a=this.getPlotLinePath({value:a,force:!0,acrossPanes:c.acrossPanes});c=
1;if(a&&h){if(d){var p=a.toString()===h.toString();c=0}for(d=0;d<a.length;d+=2){var q=a[d],r=a[d+1],y=h[d],B=h[d+1];"M"!==q[0]&&"L"!==q[0]||"M"!==r[0]&&"L"!==r[0]||"M"!==y[0]&&"L"!==y[0]||"M"!==B[0]&&"L"!==B[0]||(m&&y[1]===q[1]?(y[1]+=c,B[1]+=c):m||y[2]!==q[2]||(y[2]+=c,B[2]+=c),f.push(["M",q[1],q[2]],["L",r[1],r[2]],["L",B[1],B[2]],["L",y[1],y[2]],["Z"]));f.isFlat=p}}return f};a.prototype.addPlotBand=function(a){return this.addPlotBandOrLine(a,"plotBands")};a.prototype.addPlotLine=function(a){return this.addPlotBandOrLine(a,
"plotLines")};a.prototype.addPlotBandOrLine=function(a,d){var c=this,h=this.userOptions,f=new z(this,a);this.visible&&(f=f.render());if(f){this._addedPlotLB||(this._addedPlotLB=!0,(h.plotLines||[]).concat(h.plotBands||[]).forEach(function(a){c.addPlotBandOrLine(a)}));if(d){var m=h[d]||[];m.push(a);h[d]=m}this.plotLinesAndBands.push(f)}return f};a.prototype.removePlotBandOrLine=function(a){var d=this.plotLinesAndBands,c=this.options,h=this.userOptions;if(d){for(var f=d.length;f--;)d[f].id===a&&d[f].destroy();
[c.plotLines||[],h.plotLines||[],c.plotBands||[],h.plotBands||[]].forEach(function(c){for(f=c.length;f--;)(c[f]||{}).id===a&&u(c,c[f])})}};a.prototype.removePlotBand=function(a){this.removePlotBandOrLine(a)};a.prototype.removePlotLine=function(a){this.removePlotBandOrLine(a)};return a}()})(x||(x={}));return x});L(a,"Core/Axis/PlotLineOrBand/PlotLineOrBand.js",[a["Core/Axis/PlotLineOrBand/PlotLineOrBandAxis.js"],a["Core/Utilities.js"]],function(a,t){var u=t.arrayMax,G=t.arrayMin,x=t.defined,C=t.destroyObjectProperties,
I=t.erase,z=t.fireEvent,q=t.merge,m=t.objectEach,h=t.pick;t=function(){function d(a,d){this.axis=a;d&&(this.options=d,this.id=d.id)}d.compose=function(c){return a.compose(d,c)};d.prototype.render=function(){z(this,"render");var a=this,d=a.axis,f=d.horiz,w=d.logarithmic,p=a.options,u=p.color,r=h(p.zIndex,0),y=p.events,B={},n=d.chart.renderer,k=p.label,b=a.label,g=p.to,e=p.from,D=p.value,H=a.svgElem,v=[],E=x(e)&&x(g);v=x(D);var O=!H,S={"class":"highcharts-plot-"+(E?"band ":"line ")+(p.className||"")},
P=E?"bands":"lines";w&&(e=w.log2lin(e),g=w.log2lin(g),D=w.log2lin(D));d.chart.styledMode||(v?(S.stroke=u||"#999999",S["stroke-width"]=h(p.width,1),p.dashStyle&&(S.dashstyle=p.dashStyle)):E&&(S.fill=u||"#e6ebf5",p.borderWidth&&(S.stroke=p.borderColor,S["stroke-width"]=p.borderWidth)));B.zIndex=r;P+="-"+r;(w=d.plotLinesAndBandsGroups[P])||(d.plotLinesAndBandsGroups[P]=w=n.g("plot-"+P).attr(B).add());O&&(a.svgElem=H=n.path().attr(S).add(w));if(v)v=d.getPlotLinePath({value:D,lineWidth:H.strokeWidth(),
acrossPanes:p.acrossPanes});else if(E)v=d.getPlotBandPath(e,g,p);else return;!a.eventsAdded&&y&&(m(y,function(b,e){H.on(e,function(b){y[e].apply(a,[b])})}),a.eventsAdded=!0);(O||!H.d)&&v&&v.length?H.attr({d:v}):H&&(v?(H.show(!0),H.animate({d:v})):H.d&&(H.hide(),b&&(a.label=b=b.destroy())));k&&(x(k.text)||x(k.formatter))&&v&&v.length&&0<d.width&&0<d.height&&!v.isFlat?(k=q({align:f&&E&&"center",x:f?!E&&4:10,verticalAlign:!f&&E&&"middle",y:f?E?16:10:E?6:-4,rotation:f&&!E&&90},k),this.renderLabel(k,v,
E,r)):b&&b.hide();return a};d.prototype.renderLabel=function(a,d,f,h){var c=this.axis,l=c.chart.renderer,m=this.label;m||(this.label=m=l.text(this.getLabelText(a),0,0,a.useHTML).attr({align:a.textAlign||a.align,rotation:a.rotation,"class":"highcharts-plot-"+(f?"band":"line")+"-label "+(a.className||""),zIndex:h}).add(),c.chart.styledMode||m.css(q({textOverflow:"ellipsis"},a.style)));h=d.xBounds||[d[0][1],d[1][1],f?d[2][1]:d[0][1]];d=d.yBounds||[d[0][2],d[1][2],f?d[2][2]:d[0][2]];f=G(h);l=G(d);m.align(a,
!1,{x:f,y:l,width:u(h)-f,height:u(d)-l});m.alignValue&&"left"!==m.alignValue||m.css({width:(90===m.rotation?c.height-(m.alignAttr.y-c.top):c.width-(m.alignAttr.x-c.left))+"px"});m.show(!0)};d.prototype.getLabelText=function(a){return x(a.formatter)?a.formatter.call(this):a.text};d.prototype.destroy=function(){I(this.axis.plotLinesAndBands,this);delete this.axis;C(this)};return d}();"";"";return t});L(a,"Core/Tooltip.js",[a["Core/FormatUtilities.js"],a["Core/Globals.js"],a["Core/Renderer/RendererUtilities.js"],
a["Core/Renderer/RendererRegistry.js"],a["Core/Utilities.js"]],function(a,t,A,G,x){var u=a.format,I=t.doc,z=A.distribute,q=x.addEvent,m=x.clamp,h=x.css,d=x.defined,c=x.discardElement,l=x.extend,f=x.fireEvent,w=x.isArray,p=x.isNumber,K=x.isString,r=x.merge,y=x.pick,B=x.splat,n=x.syncTimeout;a=function(){function a(b,a){this.allowShared=!0;this.container=void 0;this.crosshairs=[];this.distance=0;this.isHidden=!0;this.isSticky=!1;this.now={};this.options={};this.outside=!1;this.chart=b;this.init(b,a)}
a.prototype.applyFilter=function(){var b=this.chart;b.renderer.definition({tagName:"filter",attributes:{id:"drop-shadow-"+b.index,opacity:.5},children:[{tagName:"feGaussianBlur",attributes:{"in":"SourceAlpha",stdDeviation:1}},{tagName:"feOffset",attributes:{dx:1,dy:1}},{tagName:"feComponentTransfer",children:[{tagName:"feFuncA",attributes:{type:"linear",slope:.3}}]},{tagName:"feMerge",children:[{tagName:"feMergeNode"},{tagName:"feMergeNode",attributes:{"in":"SourceGraphic"}}]}]})};a.prototype.bodyFormatter=
function(b){return b.map(function(b){var e=b.series.tooltipOptions;return(e[(b.point.formatPrefix||"point")+"Formatter"]||b.point.tooltipFormatter).call(b.point,e[(b.point.formatPrefix||"point")+"Format"]||"")})};a.prototype.cleanSplit=function(b){this.chart.series.forEach(function(a){var e=a&&a.tt;e&&(!e.isActive||b?a.tt=e.destroy():e.isActive=!1)})};a.prototype.defaultFormatter=function(b){var a=this.points||B(this);var e=[b.tooltipFooterHeaderFormatter(a[0])];e=e.concat(b.bodyFormatter(a));e.push(b.tooltipFooterHeaderFormatter(a[0],
!0));return e};a.prototype.destroy=function(){this.label&&(this.label=this.label.destroy());this.split&&this.tt&&(this.cleanSplit(!0),this.tt=this.tt.destroy());this.renderer&&(this.renderer=this.renderer.destroy(),c(this.container));x.clearTimeout(this.hideTimer);x.clearTimeout(this.tooltipTimeout)};a.prototype.getAnchor=function(b,a){var e=this.chart,c=e.pointer,d=e.inverted,g=e.plotTop,k=e.plotLeft,f,n,h=0,l=0;b=B(b);this.followPointer&&a?("undefined"===typeof a.chartX&&(a=c.normalize(a)),c=[a.chartX-
k,a.chartY-g]):b[0].tooltipPos?c=b[0].tooltipPos:(b.forEach(function(b){f=b.series.yAxis;n=b.series.xAxis;h+=b.plotX||0;l+=b.plotLow?(b.plotLow+(b.plotHigh||0))/2:b.plotY||0;n&&f&&(d?(h+=g+e.plotHeight-n.len-n.pos,l+=k+e.plotWidth-f.len-f.pos):(h+=n.pos-k,l+=f.pos-g))}),h/=b.length,l/=b.length,c=[d?e.plotWidth-l:h,d?e.plotHeight-h:l],this.shared&&1<b.length&&a&&(d?c[0]=a.chartX-k:c[1]=a.chartY-g));return c.map(Math.round)};a.prototype.getLabel=function(){var b=this,a=this.chart.styledMode,e=this.options,
c=this.split&&this.allowShared,k="tooltip"+(d(e.className)?" "+e.className:""),f=e.style.pointerEvents||(!this.followPointer&&e.stickOnContact?"auto":"none"),n=function(){b.inContact=!0},l=function(a){var e=b.chart.hoverSeries;b.inContact=b.shouldStickOnContact()&&b.chart.pointer.inClass(a.relatedTarget,"highcharts-tooltip");if(!b.inContact&&e&&e.onMouseOut)e.onMouseOut()},m,p=this.chart.renderer;if(b.label){var r=!b.label.hasClass("highcharts-label");(c&&!r||!c&&r)&&b.destroy()}if(!this.label){if(this.outside){r=
this.chart.options.chart.style;var B=G.getRendererType();this.container=m=t.doc.createElement("div");m.className="highcharts-tooltip-container";h(m,{position:"absolute",top:"1px",pointerEvents:f,zIndex:Math.max(this.options.style.zIndex||0,(r&&r.zIndex||0)+3)});q(m,"mouseenter",n);q(m,"mouseleave",l);t.doc.body.appendChild(m);this.renderer=p=new B(m,0,0,r,void 0,void 0,p.styledMode)}c?this.label=p.g(k):(this.label=p.label("",0,0,e.shape,void 0,void 0,e.useHTML,void 0,k).attr({padding:e.padding,r:e.borderRadius}),
a||this.label.attr({fill:e.backgroundColor,"stroke-width":e.borderWidth}).css(e.style).css({pointerEvents:f}).shadow(e.shadow));a&&e.shadow&&(this.applyFilter(),this.label.attr({filter:"url(#drop-shadow-"+this.chart.index+")"}));if(b.outside&&!b.split){var y=this.label,w=y.xSetter,u=y.ySetter;y.xSetter=function(a){w.call(y,b.distance);m.style.left=a+"px"};y.ySetter=function(a){u.call(y,b.distance);m.style.top=a+"px"}}this.label.on("mouseenter",n).on("mouseleave",l).attr({zIndex:8}).add()}return this.label};
a.prototype.getPosition=function(b,a,e){var c=this.chart,d=this.distance,g={},k=c.inverted&&e.h||0,f=this.outside,n=f?I.documentElement.clientWidth-2*d:c.chartWidth,h=f?Math.max(I.body.scrollHeight,I.documentElement.scrollHeight,I.body.offsetHeight,I.documentElement.offsetHeight,I.documentElement.clientHeight):c.chartHeight,l=c.pointer.getChartPosition(),m=function(g){var k="x"===g;return[g,k?n:h,k?b:a].concat(f?[k?b*l.scaleX:a*l.scaleY,k?l.left-d+(e.plotX+c.plotLeft)*l.scaleX:l.top-d+(e.plotY+c.plotTop)*
l.scaleY,0,k?n:h]:[k?b:a,k?e.plotX+c.plotLeft:e.plotY+c.plotTop,k?c.plotLeft:c.plotTop,k?c.plotLeft+c.plotWidth:c.plotTop+c.plotHeight])},p=m("y"),r=m("x"),B;m=!!e.negative;!c.polar&&c.hoverSeries&&c.hoverSeries.yAxis&&c.hoverSeries.yAxis.reversed&&(m=!m);var q=!this.followPointer&&y(e.ttBelow,!c.inverted===m),w=function(b,a,e,c,n,h,v){var D=f?"y"===b?d*l.scaleY:d*l.scaleX:d,m=(e-c)/2,F=c<n-d,H=n+d+c<a,p=n-D-e+m;n=n+D-m;if(q&&H)g[b]=n;else if(!q&&F)g[b]=p;else if(F)g[b]=Math.min(v-c,0>p-k?p:p-k);
else if(H)g[b]=Math.max(h,n+k+e>a?n:n+k);else return!1},u=function(b,a,e,c,k){var f;k<d||k>a-d?f=!1:g[b]=k<e/2?1:k>a-c/2?a-c-2:k-e/2;return f},F=function(b){var a=p;p=r;r=a;B=b},T=function(){!1!==w.apply(0,p)?!1!==u.apply(0,r)||B||(F(!0),T()):B?g.x=g.y=0:(F(!0),T())};(c.inverted||1<this.len)&&F();T();return g};a.prototype.hide=function(b){var a=this;x.clearTimeout(this.hideTimer);b=y(b,this.options.hideDelay);this.isHidden||(this.hideTimer=n(function(){a.getLabel().fadeOut(b?void 0:b);a.isHidden=
!0},b))};a.prototype.init=function(b,a){this.chart=b;this.options=a;this.crosshairs=[];this.now={x:0,y:0};this.isHidden=!0;this.split=a.split&&!b.inverted&&!b.polar;this.shared=a.shared||this.split;this.outside=y(a.outside,!(!b.scrollablePixelsX&&!b.scrollablePixelsY))};a.prototype.shouldStickOnContact=function(){return!(this.followPointer||!this.options.stickOnContact)};a.prototype.isStickyOnContact=function(){return!(!this.shouldStickOnContact()||!this.inContact)};a.prototype.move=function(b,a,
e,c){var d=this,g=d.now,k=!1!==d.options.animation&&!d.isHidden&&(1<Math.abs(b-g.x)||1<Math.abs(a-g.y)),f=d.followPointer||1<d.len;l(g,{x:k?(2*g.x+b)/3:b,y:k?(g.y+a)/2:a,anchorX:f?void 0:k?(2*g.anchorX+e)/3:e,anchorY:f?void 0:k?(g.anchorY+c)/2:c});d.getLabel().attr(g);d.drawTracker();k&&(x.clearTimeout(this.tooltipTimeout),this.tooltipTimeout=setTimeout(function(){d&&d.move(b,a,e,c)},32))};a.prototype.refresh=function(b,a){var e=this.chart,c=this.options,d=B(b),g=d[0],k=[],n=c.formatter||this.defaultFormatter,
h=this.shared,l=e.styledMode,m={};if(c.enabled){x.clearTimeout(this.hideTimer);this.allowShared=!(!w(b)&&b.series&&b.series.noSharedTooltip);this.followPointer=!this.split&&g.series.tooltipOptions.followPointer;b=this.getAnchor(b,a);var p=b[0],r=b[1];h&&this.allowShared?(e.pointer.applyInactiveState(d),d.forEach(function(b){b.setState("hover");k.push(b.getLabelConfig())}),m={x:g.category,y:g.y},m.points=k):m=g.getLabelConfig();this.len=k.length;n=n.call(m,this);h=g.series;this.distance=y(h.tooltipOptions.distance,
16);if(!1===n)this.hide();else{if(this.split&&this.allowShared)this.renderSplit(n,d);else{var q=p,u=r;a&&e.pointer.isDirectTouch&&(q=a.chartX-e.plotLeft,u=a.chartY-e.plotTop);if(e.polar||!1===h.options.clip||d.some(function(b){return b.series.shouldShowTooltip(q,u)}))a=this.getLabel(),c.style.width&&!l||a.css({width:this.chart.spacingBox.width+"px"}),a.attr({text:n&&n.join?n.join(""):n}),a.removeClass(/highcharts-color-[\d]+/g).addClass("highcharts-color-"+y(g.colorIndex,h.colorIndex)),l||a.attr({stroke:c.borderColor||
g.color||h.color||"#666666"}),this.updatePosition({plotX:p,plotY:r,negative:g.negative,ttBelow:g.ttBelow,h:b[2]||0});else{this.hide();return}}this.isHidden&&this.label&&this.label.attr({opacity:1}).show();this.isHidden=!1}f(this,"refresh")}};a.prototype.renderSplit=function(b,a){function e(b,a,e,d,g){void 0===g&&(g=!0);e?(a=X?0:C,b=m(b-d/2,N.left,N.right-d-(c.outside?R:0))):(a-=A,b=g?b-d-x:b+x,b=m(b,g?b:N.left,N.right));return{x:b,y:a}}var c=this,d=c.chart,g=c.chart,k=g.chartWidth,f=g.chartHeight,
n=g.plotHeight,h=g.plotLeft,p=g.plotTop,r=g.pointer,B=g.scrollablePixelsY;B=void 0===B?0:B;var q=g.scrollablePixelsX,w=g.scrollingContainer;w=void 0===w?{scrollLeft:0,scrollTop:0}:w;var u=w.scrollLeft;w=w.scrollTop;var t=g.styledMode,x=c.distance,F=c.options,T=c.options.positioner,N=c.outside&&"number"!==typeof q?I.documentElement.getBoundingClientRect():{left:u,right:u+k,top:w,bottom:w+f},V=c.getLabel(),W=this.renderer||d.renderer,X=!(!d.xAxis[0]||!d.xAxis[0].opposite);d=r.getChartPosition();var R=
d.left;d=d.top;var A=p+w,aa=0,C=n-B;K(b)&&(b=[!1,b]);b=b.slice(0,a.length+1).reduce(function(b,d,g){if(!1!==d&&""!==d){g=a[g-1]||{isHeader:!0,plotX:a[0].plotX,plotY:n,series:{}};var k=g.isHeader,f=k?c:g.series;d=d.toString();var v=f.tt,l=g.isHeader;var D=g.series;var H="highcharts-color-"+y(g.colorIndex,D.colorIndex,"none");v||(v={padding:F.padding,r:F.borderRadius},t||(v.fill=F.backgroundColor,v["stroke-width"]=F.borderWidth),v=W.label("",0,0,F[l?"headerShape":"shape"],void 0,void 0,F.useHTML).addClass((l?
"highcharts-tooltip-header ":"")+"highcharts-tooltip-box "+H).attr(v).add(V));v.isActive=!0;v.attr({text:d});t||v.css(F.style).shadow(F.shadow).attr({stroke:F.borderColor||g.color||D.color||"#333333"});f=f.tt=v;l=f.getBBox();d=l.width+f.strokeWidth();k&&(aa=l.height,C+=aa,X&&(A-=aa));D=g.plotX;D=void 0===D?0:D;H=g.plotY;H=void 0===H?0:H;v=g.series;if(g.isHeader){D=h+D;var E=p+n/2}else{var r=v.xAxis,B=v.yAxis;D=r.pos+m(D,-x,r.len+x);v.shouldShowTooltip(0,B.pos-p+H,{ignoreX:!0})&&(E=B.pos+H)}D=m(D,
N.left-x,N.right+x);"number"===typeof E?(l=l.height+1,H=T?T.call(c,d,l,g):e(D,E,k,d),b.push({align:T?0:void 0,anchorX:D,anchorY:E,boxWidth:d,point:g,rank:y(H.rank,k?1:0),size:l,target:H.y,tt:f,x:H.x})):f.isActive=!1}return b},[]);!T&&b.some(function(b){var a=(c.outside?R:0)+b.anchorX;return a<N.left&&a+b.boxWidth<N.right?!0:a<R-N.left+b.boxWidth&&N.right-a>a})&&(b=b.map(function(b){var a=e(b.anchorX,b.anchorY,b.point.isHeader,b.boxWidth,!1);return l(b,{target:a.y,x:a.x})}));c.cleanSplit();z(b,C);
var G=R,ba=R;b.forEach(function(b){var a=b.x,e=b.boxWidth;b=b.isHeader;b||(c.outside&&R+a<G&&(G=R+a),!b&&c.outside&&G+e>ba&&(ba=R+a))});b.forEach(function(b){var a=b.x,e=b.anchorX,d=b.pos,g=b.point.isHeader;d={visibility:"undefined"===typeof d?"hidden":"inherit",x:a,y:d+A,anchorX:e,anchorY:b.anchorY};if(c.outside&&a<e){var k=R-G;0<k&&(g||(d.x=a+k,d.anchorX=e+k),g&&(d.x=(ba-G)/2,d.anchorX=e+k))}b.tt.attr(d)});b=c.container;B=c.renderer;c.outside&&b&&B&&(g=V.getBBox(),B.setSize(g.width+g.x,g.height+
g.y,!1),b.style.left=G+"px",b.style.top=d+"px")};a.prototype.drawTracker=function(){if(this.followPointer||!this.options.stickOnContact)this.tracker&&this.tracker.destroy();else{var b=this.chart,a=this.label,e=this.shared?b.hoverPoints:b.hoverPoint;if(a&&e){var c={x:0,y:0,width:0,height:0};e=this.getAnchor(e);var d=a.getBBox();e[0]+=b.plotLeft-a.translateX;e[1]+=b.plotTop-a.translateY;c.x=Math.min(0,e[0]);c.y=Math.min(0,e[1]);c.width=0>e[0]?Math.max(Math.abs(e[0]),d.width-e[0]):Math.max(Math.abs(e[0]),
d.width);c.height=0>e[1]?Math.max(Math.abs(e[1]),d.height-Math.abs(e[1])):Math.max(Math.abs(e[1]),d.height);this.tracker?this.tracker.attr(c):(this.tracker=a.renderer.rect(c).addClass("highcharts-tracker").add(a),b.styledMode||this.tracker.attr({fill:"rgba(0,0,0,0)"}))}}};a.prototype.styledModeFormat=function(b){return b.replace('style="font-size: 10px"','class="highcharts-header"').replace(/style="color:{(point|series)\.color}"/g,'class="highcharts-color-{$1.colorIndex}"')};a.prototype.tooltipFooterHeaderFormatter=
function(b,a){var e=b.series,c=e.tooltipOptions,d=e.xAxis,g=d&&d.dateTime;d={isFooter:a,labelConfig:b};var k=c.xDateFormat,n=c[a?"footerFormat":"headerFormat"];f(this,"headerFormatter",d,function(a){g&&!k&&p(b.key)&&(k=g.getXDateFormat(b.key,c.dateTimeLabelFormats));g&&k&&(b.point&&b.point.tooltipDateKeys||["key"]).forEach(function(b){n=n.replace("{point."+b+"}","{point."+b+":"+k+"}")});e.chart.styledMode&&(n=this.styledModeFormat(n));a.text=u(n,{point:b,series:e},this.chart)});return d.text};a.prototype.update=
function(b){this.destroy();r(!0,this.chart.options.tooltip.userOptions,b);this.init(this.chart,r(!0,this.options,b))};a.prototype.updatePosition=function(b){var a=this.chart,e=this.options,c=a.pointer,d=this.getLabel();c=c.getChartPosition();var k=(e.positioner||this.getPosition).call(this,d.width,d.height,b),f=b.plotX+a.plotLeft;b=b.plotY+a.plotTop;if(this.outside){e=e.borderWidth+2*this.distance;this.renderer.setSize(d.width+e,d.height+e,!1);if(1!==c.scaleX||1!==c.scaleY)h(this.container,{transform:"scale("+
c.scaleX+", "+c.scaleY+")"}),f*=c.scaleX,b*=c.scaleY;f+=c.left-k.x;b+=c.top-k.y}this.move(Math.round(k.x),Math.round(k.y||0),f,b)};return a}();"";return a});L(a,"Core/Series/Point.js",[a["Core/Renderer/HTML/AST.js"],a["Core/Animation/AnimationUtilities.js"],a["Core/DefaultOptions.js"],a["Core/FormatUtilities.js"],a["Core/Utilities.js"]],function(a,t,A,G,x){var u=t.animObject,I=A.defaultOptions,z=G.format,q=x.addEvent,m=x.defined,h=x.erase,d=x.extend,c=x.fireEvent,l=x.getNestedProperty,f=x.isArray,
w=x.isFunction,p=x.isNumber,K=x.isObject,r=x.merge,y=x.objectEach,B=x.pick,n=x.syncTimeout,k=x.removeEvent,b=x.uniqueKey;t=function(){function g(){this.colorIndex=this.category=void 0;this.formatPrefix="point";this.id=void 0;this.isNull=!1;this.percentage=this.options=this.name=void 0;this.selected=!1;this.total=this.series=void 0;this.visible=!0;this.x=void 0}g.prototype.animateBeforeDestroy=function(){var b=this,a={x:b.startXPos,opacity:0},c=b.getGraphicalProps();c.singular.forEach(function(e){b[e]=
b[e].animate("dataLabel"===e?{x:b[e].startXPos,y:b[e].startYPos,opacity:0}:a)});c.plural.forEach(function(a){b[a].forEach(function(a){a.element&&a.animate(d({x:b.startXPos},a.startYPos?{x:a.startXPos,y:a.startYPos}:{}))})})};g.prototype.applyOptions=function(b,a){var e=this.series,c=e.options.pointValKey||e.pointValKey;b=g.prototype.optionsToObject.call(this,b);d(this,b);this.options=this.options?d(this.options,b):b;b.group&&delete this.group;b.dataLabels&&delete this.dataLabels;c&&(this.y=g.prototype.getNestedProperty.call(this,
c));this.formatPrefix=(this.isNull=B(this.isValid&&!this.isValid(),null===this.x||!p(this.y)))?"null":"point";this.selected&&(this.state="select");"name"in this&&"undefined"===typeof a&&e.xAxis&&e.xAxis.hasNames&&(this.x=e.xAxis.nameToX(this));"undefined"===typeof this.x&&e?this.x="undefined"===typeof a?e.autoIncrement():a:p(b.x)&&e.options.relativeXValue&&(this.x=e.autoIncrement(b.x));return this};g.prototype.destroy=function(){function b(){if(a.graphic||a.dataLabel||a.dataLabels)k(a),a.destroyElements();
for(l in a)a[l]=null}var a=this,c=a.series,d=c.chart;c=c.options.dataSorting;var g=d.hoverPoints,f=u(a.series.chart.renderer.globalAnimation),l;a.legendItem&&d.legend.destroyItem(a);g&&(a.setState(),h(g,a),g.length||(d.hoverPoints=null));if(a===d.hoverPoint)a.onMouseOut();c&&c.enabled?(this.animateBeforeDestroy(),n(b,f.duration)):b();d.pointCount--};g.prototype.destroyElements=function(b){var a=this;b=a.getGraphicalProps(b);b.singular.forEach(function(b){a[b]=a[b].destroy()});b.plural.forEach(function(b){a[b].forEach(function(b){b.element&&
b.destroy()});delete a[b]})};g.prototype.firePointEvent=function(b,a,d){var e=this,g=this.series.options;(g.point.events[b]||e.options&&e.options.events&&e.options.events[b])&&e.importEvents();"click"===b&&g.allowPointSelect&&(d=function(b){e.select&&e.select(null,b.ctrlKey||b.metaKey||b.shiftKey)});c(e,b,a,d)};g.prototype.getClassName=function(){return"highcharts-point"+(this.selected?" highcharts-point-select":"")+(this.negative?" highcharts-negative":"")+(this.isNull?" highcharts-null-point":"")+
("undefined"!==typeof this.colorIndex?" highcharts-color-"+this.colorIndex:"")+(this.options.className?" "+this.options.className:"")+(this.zone&&this.zone.className?" "+this.zone.className.replace("highcharts-negative",""):"")};g.prototype.getGraphicalProps=function(b){var a=this,e=[],c={singular:[],plural:[]},d;b=b||{graphic:1,dataLabel:1};b.graphic&&e.push("graphic","upperGraphic","shadowGroup");b.dataLabel&&e.push("dataLabel","dataLabelUpper","connector");for(d=e.length;d--;){var g=e[d];a[g]&&
c.singular.push(g)}["dataLabel","connector"].forEach(function(e){var d=e+"s";b[e]&&a[d]&&c.plural.push(d)});return c};g.prototype.getLabelConfig=function(){return{x:this.category,y:this.y,color:this.color,colorIndex:this.colorIndex,key:this.name||this.category,series:this.series,point:this,percentage:this.percentage,total:this.total||this.stackTotal}};g.prototype.getNestedProperty=function(b){if(b)return 0===b.indexOf("custom.")?l(b,this.options):this[b]};g.prototype.getZone=function(){var b=this.series,
a=b.zones;b=b.zoneAxis||"y";var c,d=0;for(c=a[d];this[b]>=c.value;)c=a[++d];this.nonZonedColor||(this.nonZonedColor=this.color);this.color=c&&c.color&&!this.options.color?c.color:this.nonZonedColor;return c};g.prototype.hasNewShapeType=function(){return(this.graphic&&(this.graphic.symbolName||this.graphic.element.nodeName))!==this.shapeType};g.prototype.init=function(a,d,g){this.series=a;this.applyOptions(d,g);this.id=m(this.id)?this.id:b();this.resolveColor();a.chart.pointCount++;c(this,"afterInit");
return this};g.prototype.optionsToObject=function(b){var a=this.series,e=a.options.keys,c=e||a.pointArrayMap||["y"],d=c.length,k={},n=0,h=0;if(p(b)||null===b)k[c[0]]=b;else if(f(b))for(!e&&b.length>d&&(a=typeof b[0],"string"===a?k.name=b[0]:"number"===a&&(k.x=b[0]),n++);h<d;)e&&"undefined"===typeof b[n]||(0<c[h].indexOf(".")?g.prototype.setNestedProperty(k,b[n],c[h]):k[c[h]]=b[n]),n++,h++;else"object"===typeof b&&(k=b,b.dataLabels&&(a._hasPointLabels=!0),b.marker&&(a._hasPointMarkers=!0));return k};
g.prototype.resolveColor=function(){var b=this.series,a=b.chart.styledMode;var c=b.chart.options.chart.colorCount;delete this.nonZonedColor;if(b.options.colorByPoint){if(!a){c=b.options.colors||b.chart.options.colors;var d=c[b.colorCounter];c=c.length}a=b.colorCounter;b.colorCounter++;b.colorCounter===c&&(b.colorCounter=0)}else a||(d=b.color),a=b.colorIndex;this.colorIndex=B(this.options.colorIndex,a);this.color=B(this.options.color,d)};g.prototype.setNestedProperty=function(b,a,c){c.split(".").reduce(function(b,
e,c,d){b[e]=d.length-1===c?a:K(b[e],!0)?b[e]:{};return b[e]},b);return b};g.prototype.tooltipFormatter=function(b){var a=this.series,e=a.tooltipOptions,c=B(e.valueDecimals,""),d=e.valuePrefix||"",g=e.valueSuffix||"";a.chart.styledMode&&(b=a.chart.tooltip.styledModeFormat(b));(a.pointArrayMap||["y"]).forEach(function(a){a="{point."+a;if(d||g)b=b.replace(RegExp(a+"}","g"),d+a+"}"+g);b=b.replace(RegExp(a+"}","g"),a+":,."+c+"f}")});return z(b,{point:this,series:this.series},a.chart)};g.prototype.update=
function(b,a,c,d){function e(){g.applyOptions(b);var e=f&&g.hasDummyGraphic;e=null===g.y?!e:e;f&&e&&(g.graphic=f.destroy(),delete g.hasDummyGraphic);K(b,!0)&&(f&&f.element&&b&&b.marker&&"undefined"!==typeof b.marker.symbol&&(g.graphic=f.destroy()),b&&b.dataLabels&&g.dataLabel&&(g.dataLabel=g.dataLabel.destroy()),g.connector&&(g.connector=g.connector.destroy()));l=g.index;k.updateParallelArrays(g,l);h.data[l]=K(h.data[l],!0)||K(b,!0)?g.options:B(b,h.data[l]);k.isDirty=k.isDirtyData=!0;!k.fixedBox&&
k.hasCartesianSeries&&(n.isDirtyBox=!0);"point"===h.legendType&&(n.isDirtyLegend=!0);a&&n.redraw(c)}var g=this,k=g.series,f=g.graphic,n=k.chart,h=k.options,l;a=B(a,!0);!1===d?e():g.firePointEvent("update",{options:b},e)};g.prototype.remove=function(b,a){this.series.removePoint(this.series.data.indexOf(this),b,a)};g.prototype.select=function(b,a){var e=this,c=e.series,d=c.chart;this.selectedStaging=b=B(b,!e.selected);e.firePointEvent(b?"select":"unselect",{accumulate:a},function(){e.selected=e.options.selected=
b;c.options.data[c.data.indexOf(e)]=e.options;e.setState(b&&"select");a||d.getSelectedPoints().forEach(function(b){var a=b.series;b.selected&&b!==e&&(b.selected=b.options.selected=!1,a.options.data[a.data.indexOf(b)]=b.options,b.setState(d.hoverPoints&&a.options.inactiveOtherPoints?"inactive":""),b.firePointEvent("unselect"))})});delete this.selectedStaging};g.prototype.onMouseOver=function(b){var a=this.series.chart,e=a.pointer;b=b?e.normalize(b):e.getChartCoordinatesFromPoint(this,a.inverted);e.runPointActions(b,
this)};g.prototype.onMouseOut=function(){var b=this.series.chart;this.firePointEvent("mouseOut");this.series.options.inactiveOtherPoints||(b.hoverPoints||[]).forEach(function(b){b.setState()});b.hoverPoints=b.hoverPoint=null};g.prototype.importEvents=function(){if(!this.hasImportedEvents){var b=this,a=r(b.series.options.point,b.options).events;b.events=a;y(a,function(a,e){w(a)&&q(b,e,a)});this.hasImportedEvents=!0}};g.prototype.setState=function(b,g){var e=this.series,k=this.state,f=e.options.states[b||
"normal"]||{},n=I.plotOptions[e.type].marker&&e.options.marker,h=n&&!1===n.enabled,l=n&&n.states&&n.states[b||"normal"]||{},m=!1===l.enabled,D=this.marker||{},r=e.chart,q=n&&e.markerAttribs,w=e.halo,y,u=e.stateMarkerGraphic;b=b||"";if(!(b===this.state&&!g||this.selected&&"select"!==b||!1===f.enabled||b&&(m||h&&!1===l.enabled)||b&&D.states&&D.states[b]&&!1===D.states[b].enabled)){this.state=b;q&&(y=e.markerAttribs(this,b));if(this.graphic&&!this.hasDummyGraphic){k&&this.graphic.removeClass("highcharts-point-"+
k);b&&this.graphic.addClass("highcharts-point-"+b);if(!r.styledMode){var K=e.pointAttribs(this,b);var F=B(r.options.chart.animation,f.animation);e.options.inactiveOtherPoints&&p(K.opacity)&&((this.dataLabels||[]).forEach(function(b){b&&b.animate({opacity:K.opacity},F)}),this.connector&&this.connector.animate({opacity:K.opacity},F));this.graphic.animate(K,F)}y&&this.graphic.animate(y,B(r.options.chart.animation,l.animation,n.animation));u&&u.hide()}else{if(b&&l){k=D.symbol||e.symbol;u&&u.currentSymbol!==
k&&(u=u.destroy());if(y)if(u)u[g?"animate":"attr"]({x:y.x,y:y.y});else k&&(e.stateMarkerGraphic=u=r.renderer.symbol(k,y.x,y.y,y.width,y.height).add(e.markerGroup),u.currentSymbol=k);!r.styledMode&&u&&"inactive"!==this.state&&u.attr(e.pointAttribs(this,b))}u&&(u[b&&this.isInside?"show":"hide"](),u.element.point=this,u.addClass(this.getClassName(),!0))}f=f.halo;y=(u=this.graphic||u)&&u.visibility||"inherit";f&&f.size&&u&&"hidden"!==y&&!this.isCluster?(w||(e.halo=w=r.renderer.path().add(u.parentGroup)),
w.show()[g?"animate":"attr"]({d:this.haloPath(f.size)}),w.attr({"class":"highcharts-halo highcharts-color-"+B(this.colorIndex,e.colorIndex)+(this.className?" "+this.className:""),visibility:y,zIndex:-1}),w.point=this,r.styledMode||w.attr(d({fill:this.color||e.color,"fill-opacity":f.opacity},a.filterUserAttributes(f.attributes||{})))):w&&w.point&&w.point.haloPath&&w.animate({d:w.point.haloPath(0)},null,w.hide);c(this,"afterSetState",{state:b})}};g.prototype.haloPath=function(b){return this.series.chart.renderer.symbols.circle(Math.floor(this.plotX)-
b,this.plotY-b,2*b,2*b)};return g}();"";return t});L(a,"Core/Pointer.js",[a["Core/Color/Color.js"],a["Core/Globals.js"],a["Core/Tooltip.js"],a["Core/Utilities.js"]],function(a,t,A,G){var u=a.parse,C=t.charts,I=t.noop,z=G.addEvent,q=G.attr,m=G.css,h=G.defined,d=G.extend,c=G.find,l=G.fireEvent,f=G.isNumber,w=G.isObject,p=G.objectEach,K=G.offset,r=G.pick,y=G.splat;a=function(){function a(a,c){this.lastValidTouch={};this.pinchDown=[];this.runChartClick=!1;this.eventsToUnbind=[];this.chart=a;this.hasDragged=
!1;this.options=c;this.init(a,c)}a.prototype.applyInactiveState=function(a){var c=[],b;(a||[]).forEach(function(a){b=a.series;c.push(b);b.linkedParent&&c.push(b.linkedParent);b.linkedSeries&&(c=c.concat(b.linkedSeries));b.navigatorSeries&&c.push(b.navigatorSeries)});this.chart.series.forEach(function(b){-1===c.indexOf(b)?b.setState("inactive",!0):b.options.inactiveOtherPoints&&b.setAllPointsToState("inactive")})};a.prototype.destroy=function(){var c=this;this.eventsToUnbind.forEach(function(a){return a()});
this.eventsToUnbind=[];t.chartCount||(a.unbindDocumentMouseUp&&(a.unbindDocumentMouseUp=a.unbindDocumentMouseUp()),a.unbindDocumentTouchEnd&&(a.unbindDocumentTouchEnd=a.unbindDocumentTouchEnd()));clearInterval(c.tooltipTimeout);p(c,function(a,b){c[b]=void 0})};a.prototype.drag=function(a){var c=this.chart,b=c.options.chart,d=this.zoomHor,e=this.zoomVert,f=c.plotLeft,n=c.plotTop,h=c.plotWidth,l=c.plotHeight,m=this.mouseDownX||0,p=this.mouseDownY||0,r=w(b.panning)?b.panning&&b.panning.enabled:b.panning,
q=b.panKey&&a[b.panKey+"Key"],B=a.chartX,y=a.chartY,K=this.selectionMarker;if(!K||!K.touch)if(B<f?B=f:B>f+h&&(B=f+h),y<n?y=n:y>n+l&&(y=n+l),this.hasDragged=Math.sqrt(Math.pow(m-B,2)+Math.pow(p-y,2)),10<this.hasDragged){var t=c.isInsidePlot(m-f,p-n,{visiblePlotOnly:!0});!c.hasCartesianSeries&&!c.mapView||!this.zoomX&&!this.zoomY||!t||q||K||(this.selectionMarker=K=c.renderer.rect(f,n,d?1:h,e?1:l,0).attr({"class":"highcharts-selection-marker",zIndex:7}).add(),c.styledMode||K.attr({fill:b.selectionMarkerFill||
u("#335cad").setOpacity(.25).get()}));K&&d&&(d=B-m,K.attr({width:Math.abs(d),x:(0<d?0:d)+m}));K&&e&&(d=y-p,K.attr({height:Math.abs(d),y:(0<d?0:d)+p}));t&&!K&&r&&c.pan(a,b.panning)}};a.prototype.dragStart=function(a){var c=this.chart;c.mouseIsDown=a.type;c.cancelClick=!1;c.mouseDownX=this.mouseDownX=a.chartX;c.mouseDownY=this.mouseDownY=a.chartY};a.prototype.drop=function(a){var c=this,b=this.chart,g=this.hasPinched;if(this.selectionMarker){var e=this.selectionMarker,n=e.attr?e.attr("x"):e.x,p=e.attr?
e.attr("y"):e.y,v=e.attr?e.attr("width"):e.width,r=e.attr?e.attr("height"):e.height,q={originalEvent:a,xAxis:[],yAxis:[],x:n,y:p,width:v,height:r},B=!!b.mapView;if(this.hasDragged||g)b.axes.forEach(function(b){if(b.zoomEnabled&&h(b.min)&&(g||c[{xAxis:"zoomX",yAxis:"zoomY"}[b.coll]])&&f(n)&&f(p)){var e=b.horiz,d="touchend"===a.type?b.minPixelPadding:0,k=b.toValue((e?n:p)+d);e=b.toValue((e?n+v:p+r)-d);q[b.coll].push({axis:b,min:Math.min(k,e),max:Math.max(k,e)});B=!0}}),B&&l(b,"selection",q,function(a){b.zoom(d(a,
g?{animation:!1}:null))});f(b.index)&&(this.selectionMarker=this.selectionMarker.destroy());g&&this.scaleGroups()}b&&f(b.index)&&(m(b.container,{cursor:b._cursor}),b.cancelClick=10<this.hasDragged,b.mouseIsDown=this.hasDragged=this.hasPinched=!1,this.pinchDown=[])};a.prototype.findNearestKDPoint=function(a,c,b){var d=this.chart,e=d.hoverPoint;d=d.tooltip;if(e&&d&&d.isStickyOnContact())return e;var k;a.forEach(function(a){var e=!(a.noSharedTooltip&&c)&&0>a.options.findNearestPointBy.indexOf("y");a=
a.searchPoint(b,e);if((e=w(a,!0)&&a.series)&&!(e=!w(k,!0))){e=k.distX-a.distX;var d=k.dist-a.dist,g=(a.series.group&&a.series.group.zIndex)-(k.series.group&&k.series.group.zIndex);e=0<(0!==e&&c?e:0!==d?d:0!==g?g:k.series.index>a.series.index?-1:1)}e&&(k=a)});return k};a.prototype.getChartCoordinatesFromPoint=function(a,c){var b=a.series,d=b.xAxis;b=b.yAxis;var e=a.shapeArgs;if(d&&b){var k=r(a.clientX,a.plotX),n=a.plotY||0;a.isNode&&e&&f(e.x)&&f(e.y)&&(k=e.x,n=e.y);return c?{chartX:b.len+b.pos-n,chartY:d.len+
d.pos-k}:{chartX:k+d.pos,chartY:n+b.pos}}if(e&&e.x&&e.y)return{chartX:e.x,chartY:e.y}};a.prototype.getChartPosition=function(){if(this.chartPosition)return this.chartPosition;var a=this.chart.container,c=K(a);this.chartPosition={left:c.left,top:c.top,scaleX:1,scaleY:1};var b=a.offsetWidth;a=a.offsetHeight;2<b&&2<a&&(this.chartPosition.scaleX=c.width/b,this.chartPosition.scaleY=c.height/a);return this.chartPosition};a.prototype.getCoordinates=function(a){var c={xAxis:[],yAxis:[]};this.chart.axes.forEach(function(b){c[b.isXAxis?
"xAxis":"yAxis"].push({axis:b,value:b.toValue(a[b.horiz?"chartX":"chartY"])})});return c};a.prototype.getHoverData=function(a,d,b,g,e,f){var k=[];g=!(!g||!a);var h={chartX:f?f.chartX:void 0,chartY:f?f.chartY:void 0,shared:e};l(this,"beforeGetHoverData",h);var n=d&&!d.stickyTracking?[d]:b.filter(function(b){return h.filter?h.filter(b):b.visible&&!(!e&&b.directTouch)&&r(b.options.enableMouseTracking,!0)&&b.stickyTracking});var m=g||!f?a:this.findNearestKDPoint(n,e,f);d=m&&m.series;m&&(e&&!d.noSharedTooltip?
(n=b.filter(function(b){return h.filter?h.filter(b):b.visible&&!(!e&&b.directTouch)&&r(b.options.enableMouseTracking,!0)&&!b.noSharedTooltip}),n.forEach(function(b){var a=c(b.points,function(b){return b.x===m.x&&!b.isNull});w(a)&&(b.chart.isBoosting&&(a=b.getPoint(a)),k.push(a))})):k.push(m));h={hoverPoint:m};l(this,"afterGetHoverData",h);return{hoverPoint:h.hoverPoint,hoverSeries:d,hoverPoints:k}};a.prototype.getPointFromEvent=function(a){a=a.target;for(var c;a&&!c;)c=a.point,a=a.parentNode;return c};
a.prototype.onTrackerMouseOut=function(a){a=a.relatedTarget||a.toElement;var c=this.chart.hoverSeries;this.isDirectTouch=!1;if(!(!c||!a||c.stickyTracking||this.inClass(a,"highcharts-tooltip")||this.inClass(a,"highcharts-series-"+c.index)&&this.inClass(a,"highcharts-tracker")))c.onMouseOut()};a.prototype.inClass=function(a,c){for(var b;a;){if(b=q(a,"class")){if(-1!==b.indexOf(c))return!0;if(-1!==b.indexOf("highcharts-container"))return!1}a=a.parentNode}};a.prototype.init=function(a,c){this.options=
c;this.chart=a;this.runChartClick=!(!c.chart.events||!c.chart.events.click);this.pinchDown=[];this.lastValidTouch={};A&&(a.tooltip=new A(a,c.tooltip),this.followTouchMove=r(c.tooltip.followTouchMove,!0));this.setDOMEvents()};a.prototype.normalize=function(a,c){var b=a.touches,g=b?b.length?b.item(0):r(b.changedTouches,a.changedTouches)[0]:a;c||(c=this.getChartPosition());b=g.pageX-c.left;g=g.pageY-c.top;b/=c.scaleX;g/=c.scaleY;return d(a,{chartX:Math.round(b),chartY:Math.round(g)})};a.prototype.onContainerClick=
function(a){var c=this.chart,b=c.hoverPoint;a=this.normalize(a);var g=c.plotLeft,e=c.plotTop;c.cancelClick||(b&&this.inClass(a.target,"highcharts-tracker")?(l(b.series,"click",d(a,{point:b})),c.hoverPoint&&b.firePointEvent("click",a)):(d(a,this.getCoordinates(a)),c.isInsidePlot(a.chartX-g,a.chartY-e,{visiblePlotOnly:!0})&&l(c,"click",a)))};a.prototype.onContainerMouseDown=function(a){var c=1===((a.buttons||a.button)&1);a=this.normalize(a);if(t.isFirefox&&0!==a.button)this.onContainerMouseMove(a);
if("undefined"===typeof a.button||c)this.zoomOption(a),c&&a.preventDefault&&a.preventDefault(),this.dragStart(a)};a.prototype.onContainerMouseLeave=function(c){var d=C[r(a.hoverChartIndex,-1)],b=this.chart.tooltip;b&&b.shouldStickOnContact()&&this.inClass(c.relatedTarget,"highcharts-tooltip-container")||(c=this.normalize(c),d&&(c.relatedTarget||c.toElement)&&(d.pointer.reset(),d.pointer.chartPosition=void 0),b&&!b.isHidden&&this.reset())};a.prototype.onContainerMouseEnter=function(a){delete this.chartPosition};
a.prototype.onContainerMouseMove=function(a){var c=this.chart;a=this.normalize(a);this.setHoverChartIndex();a.preventDefault||(a.returnValue=!1);("mousedown"===c.mouseIsDown||this.touchSelect(a))&&this.drag(a);c.openMenu||!this.inClass(a.target,"highcharts-tracker")&&!c.isInsidePlot(a.chartX-c.plotLeft,a.chartY-c.plotTop,{visiblePlotOnly:!0})||(this.inClass(a.target,"highcharts-no-tooltip")?this.reset(!1,0):this.runPointActions(a))};a.prototype.onDocumentTouchEnd=function(c){var d=C[r(a.hoverChartIndex,
-1)];d&&d.pointer.drop(c)};a.prototype.onContainerTouchMove=function(a){if(this.touchSelect(a))this.onContainerMouseMove(a);else this.touch(a)};a.prototype.onContainerTouchStart=function(a){if(this.touchSelect(a))this.onContainerMouseDown(a);else this.zoomOption(a),this.touch(a,!0)};a.prototype.onDocumentMouseMove=function(a){var c=this.chart,b=this.chartPosition;a=this.normalize(a,b);var d=c.tooltip;!b||d&&d.isStickyOnContact()||c.isInsidePlot(a.chartX-c.plotLeft,a.chartY-c.plotTop,{visiblePlotOnly:!0})||
this.inClass(a.target,"highcharts-tracker")||this.reset()};a.prototype.onDocumentMouseUp=function(c){var d=C[r(a.hoverChartIndex,-1)];d&&d.pointer.drop(c)};a.prototype.pinch=function(a){var c=this,b=c.chart,g=c.pinchDown,e=a.touches||[],f=e.length,h=c.lastValidTouch,n=c.hasZoom,m={},p=1===f&&(c.inClass(a.target,"highcharts-tracker")&&b.runTrackerClick||c.runChartClick),q={},B=c.selectionMarker;1<f?c.initiated=!0:1===f&&this.followTouchMove&&(c.initiated=!1);n&&c.initiated&&!p&&!1!==a.cancelable&&
a.preventDefault();[].map.call(e,function(b){return c.normalize(b)});"touchstart"===a.type?([].forEach.call(e,function(b,a){g[a]={chartX:b.chartX,chartY:b.chartY}}),h.x=[g[0].chartX,g[1]&&g[1].chartX],h.y=[g[0].chartY,g[1]&&g[1].chartY],b.axes.forEach(function(a){if(a.zoomEnabled){var c=b.bounds[a.horiz?"h":"v"],e=a.minPixelPadding,d=a.toPixels(Math.min(r(a.options.min,a.dataMin),a.dataMin)),g=a.toPixels(Math.max(r(a.options.max,a.dataMax),a.dataMax)),f=Math.max(d,g);c.min=Math.min(a.pos,Math.min(d,
g)-e);c.max=Math.max(a.pos+a.len,f+e)}}),c.res=!0):c.followTouchMove&&1===f?this.runPointActions(c.normalize(a)):g.length&&(l(b,"touchpan",{originalEvent:a},function(){B||(c.selectionMarker=B=d({destroy:I,touch:!0},b.plotBox));c.pinchTranslate(g,e,m,B,q,h);c.hasPinched=n;c.scaleGroups(m,q)}),c.res&&(c.res=!1,this.reset(!1,0)))};a.prototype.pinchTranslate=function(a,c,b,d,e,f){this.zoomHor&&this.pinchTranslateDirection(!0,a,c,b,d,e,f);this.zoomVert&&this.pinchTranslateDirection(!1,a,c,b,d,e,f)};a.prototype.pinchTranslateDirection=
function(a,c,b,d,e,f,h,l){var g=this.chart,k=a?"x":"y",n=a?"X":"Y",m="chart"+n,p=a?"width":"height",v=g["plot"+(a?"Left":"Top")],r=g.inverted,D=g.bounds[a?"h":"v"],q=1===c.length,B=c[0][m],y=!q&&c[1][m];c=function(){"number"===typeof u&&20<Math.abs(B-y)&&(w=l||Math.abs(N-u)/Math.abs(B-y));F=(v-N)/w+B;H=g["plot"+(a?"Width":"Height")]/w};var H,F,w=l||1,N=b[0][m],u=!q&&b[1][m];c();b=F;if(b<D.min){b=D.min;var K=!0}else b+H>D.max&&(b=D.max-H,K=!0);K?(N-=.8*(N-h[k][0]),"number"===typeof u&&(u-=.8*(u-h[k][1])),
c()):h[k]=[N,u];r||(f[k]=F-v,f[p]=H);f=r?1/w:w;e[p]=H;e[k]=b;d[r?a?"scaleY":"scaleX":"scale"+n]=w;d["translate"+n]=f*v+(N-f*B)};a.prototype.reset=function(a,c){var b=this.chart,d=b.hoverSeries,e=b.hoverPoint,f=b.hoverPoints,k=b.tooltip,h=k&&k.shared?f:e;a&&h&&y(h).forEach(function(b){b.series.isCartesian&&"undefined"===typeof b.plotX&&(a=!1)});if(a)k&&h&&y(h).length&&(k.refresh(h),k.shared&&f?f.forEach(function(b){b.setState(b.state,!0);b.series.isCartesian&&(b.series.xAxis.crosshair&&b.series.xAxis.drawCrosshair(null,
b),b.series.yAxis.crosshair&&b.series.yAxis.drawCrosshair(null,b))}):e&&(e.setState(e.state,!0),b.axes.forEach(function(b){b.crosshair&&e.series[b.coll]===b&&b.drawCrosshair(null,e)})));else{if(e)e.onMouseOut();f&&f.forEach(function(b){b.setState()});if(d)d.onMouseOut();k&&k.hide(c);this.unDocMouseMove&&(this.unDocMouseMove=this.unDocMouseMove());b.axes.forEach(function(b){b.hideCrosshair()});this.hoverX=b.hoverPoints=b.hoverPoint=null}};a.prototype.runPointActions=function(d,f){var b=this.chart,
g=b.tooltip&&b.tooltip.options.enabled?b.tooltip:void 0,e=g?g.shared:!1,k=f||b.hoverPoint,h=k&&k.series||b.hoverSeries;f=this.getHoverData(k,h,b.series,(!d||"touchmove"!==d.type)&&(!!f||h&&h.directTouch&&this.isDirectTouch),e,d);k=f.hoverPoint;h=f.hoverSeries;var l=f.hoverPoints;f=h&&h.tooltipOptions.followPointer&&!h.tooltipOptions.split;e=e&&h&&!h.noSharedTooltip;if(k&&(k!==b.hoverPoint||g&&g.isHidden)){(b.hoverPoints||[]).forEach(function(b){-1===l.indexOf(b)&&b.setState()});if(b.hoverSeries!==
h)h.onMouseOver();this.applyInactiveState(l);(l||[]).forEach(function(b){b.setState("hover")});b.hoverPoint&&b.hoverPoint.firePointEvent("mouseOut");if(!k.series)return;b.hoverPoints=l;b.hoverPoint=k;k.firePointEvent("mouseOver");g&&g.refresh(e?l:k,d)}else f&&g&&!g.isHidden&&(k=g.getAnchor([{}],d),b.isInsidePlot(k[0],k[1],{visiblePlotOnly:!0})&&g.updatePosition({plotX:k[0],plotY:k[1]}));this.unDocMouseMove||(this.unDocMouseMove=z(b.container.ownerDocument,"mousemove",function(b){var c=C[a.hoverChartIndex];
if(c)c.pointer.onDocumentMouseMove(b)}),this.eventsToUnbind.push(this.unDocMouseMove));b.axes.forEach(function(a){var e=r((a.crosshair||{}).snap,!0),g;e&&((g=b.hoverPoint)&&g.series[a.coll]===a||(g=c(l,function(b){return b.series[a.coll]===a})));g||!e?a.drawCrosshair(d,g):a.hideCrosshair()})};a.prototype.scaleGroups=function(a,c){var b=this.chart;b.series.forEach(function(d){var e=a||d.getPlotBox();d.group&&(d.xAxis&&d.xAxis.zoomEnabled||b.mapView)&&(d.group.attr(e),d.markerGroup&&(d.markerGroup.attr(e),
d.markerGroup.clip(c?b.clipRect:null)),d.dataLabelsGroup&&d.dataLabelsGroup.attr(e))});b.clipRect.attr(c||b.clipBox)};a.prototype.setDOMEvents=function(){var c=this,d=this.chart.container,b=d.ownerDocument;d.onmousedown=this.onContainerMouseDown.bind(this);d.onmousemove=this.onContainerMouseMove.bind(this);d.onclick=this.onContainerClick.bind(this);this.eventsToUnbind.push(z(d,"mouseenter",this.onContainerMouseEnter.bind(this)));this.eventsToUnbind.push(z(d,"mouseleave",this.onContainerMouseLeave.bind(this)));
a.unbindDocumentMouseUp||(a.unbindDocumentMouseUp=z(b,"mouseup",this.onDocumentMouseUp.bind(this)));for(var g=this.chart.renderTo.parentElement;g&&"BODY"!==g.tagName;)this.eventsToUnbind.push(z(g,"scroll",function(){delete c.chartPosition})),g=g.parentElement;t.hasTouch&&(this.eventsToUnbind.push(z(d,"touchstart",this.onContainerTouchStart.bind(this),{passive:!1})),this.eventsToUnbind.push(z(d,"touchmove",this.onContainerTouchMove.bind(this),{passive:!1})),a.unbindDocumentTouchEnd||(a.unbindDocumentTouchEnd=
z(b,"touchend",this.onDocumentTouchEnd.bind(this),{passive:!1})))};a.prototype.setHoverChartIndex=function(){var c=this.chart,d=t.charts[r(a.hoverChartIndex,-1)];if(d&&d!==c)d.pointer.onContainerMouseLeave({relatedTarget:!0});d&&d.mouseIsDown||(a.hoverChartIndex=c.index)};a.prototype.touch=function(a,c){var b=this.chart,d;this.setHoverChartIndex();if(1===a.touches.length)if(a=this.normalize(a),(d=b.isInsidePlot(a.chartX-b.plotLeft,a.chartY-b.plotTop,{visiblePlotOnly:!0}))&&!b.openMenu){c&&this.runPointActions(a);
if("touchmove"===a.type){c=this.pinchDown;var e=c[0]?4<=Math.sqrt(Math.pow(c[0].chartX-a.chartX,2)+Math.pow(c[0].chartY-a.chartY,2)):!1}r(e,!0)&&this.pinch(a)}else c&&this.reset();else 2===a.touches.length&&this.pinch(a)};a.prototype.touchSelect=function(a){return!(!this.chart.options.chart.zoomBySingleTouch||!a.touches||1!==a.touches.length)};a.prototype.zoomOption=function(a){var c=this.chart,b=c.options.chart;c=c.inverted;var d=b.zoomType||"";/touch/.test(a.type)&&(d=r(b.pinchType,d));this.zoomX=
a=/x/.test(d);this.zoomY=b=/y/.test(d);this.zoomHor=a&&!c||b&&c;this.zoomVert=b&&!c||a&&c;this.hasZoom=a||b};return a}();"";return a});L(a,"Core/MSPointer.js",[a["Core/Globals.js"],a["Core/Pointer.js"],a["Core/Utilities.js"]],function(a,t,A){function u(){var a=[];a.item=function(a){return this[a]};c(f,function(c){a.push({pageX:c.pageX,pageY:c.pageY,target:c.target})});return a}function x(a,c,d,f){var h=I[t.hoverChartIndex||NaN];"touch"!==a.pointerType&&a.pointerType!==a.MSPOINTER_TYPE_TOUCH||!h||
(h=h.pointer,f(a),h[c]({type:d,target:a.currentTarget,preventDefault:q,touches:u()}))}var C=this&&this.__extends||function(){var a=function(c,d){a=Object.setPrototypeOf||{__proto__:[]}instanceof Array&&function(a,c){a.__proto__=c}||function(a,c){for(var d in c)c.hasOwnProperty(d)&&(a[d]=c[d])};return a(c,d)};return function(c,d){function f(){this.constructor=c}a(c,d);c.prototype=null===d?Object.create(d):(f.prototype=d.prototype,new f)}}(),I=a.charts,z=a.doc,q=a.noop,m=a.win,h=A.addEvent,d=A.css,
c=A.objectEach,l=A.removeEvent,f={},w=!!m.PointerEvent;return function(c){function p(){return null!==c&&c.apply(this,arguments)||this}C(p,c);p.isRequired=function(){return!(a.hasTouch||!m.PointerEvent&&!m.MSPointerEvent)};p.prototype.batchMSEvents=function(a){a(this.chart.container,w?"pointerdown":"MSPointerDown",this.onContainerPointerDown);a(this.chart.container,w?"pointermove":"MSPointerMove",this.onContainerPointerMove);a(z,w?"pointerup":"MSPointerUp",this.onDocumentPointerUp)};p.prototype.destroy=
function(){this.batchMSEvents(l);c.prototype.destroy.call(this)};p.prototype.init=function(a,f){c.prototype.init.call(this,a,f);this.hasZoom&&d(a.container,{"-ms-touch-action":"none","touch-action":"none"})};p.prototype.onContainerPointerDown=function(a){x(a,"onContainerTouchStart","touchstart",function(a){f[a.pointerId]={pageX:a.pageX,pageY:a.pageY,target:a.currentTarget}})};p.prototype.onContainerPointerMove=function(a){x(a,"onContainerTouchMove","touchmove",function(a){f[a.pointerId]={pageX:a.pageX,
pageY:a.pageY};f[a.pointerId].target||(f[a.pointerId].target=a.currentTarget)})};p.prototype.onDocumentPointerUp=function(a){x(a,"onDocumentTouchEnd","touchend",function(a){delete f[a.pointerId]})};p.prototype.setDOMEvents=function(){c.prototype.setDOMEvents.call(this);(this.hasZoom||this.followTouchMove)&&this.batchMSEvents(h)};return p}(t)});L(a,"Core/Legend/Legend.js",[a["Core/Animation/AnimationUtilities.js"],a["Core/FormatUtilities.js"],a["Core/Globals.js"],a["Core/Series/Point.js"],a["Core/Renderer/RendererUtilities.js"],
a["Core/Utilities.js"]],function(a,t,A,G,x,C){var u=a.animObject,z=a.setAnimation,q=t.format;a=A.isFirefox;var m=A.marginNames;A=A.win;var h=x.distribute,d=C.addEvent,c=C.createElement,l=C.css,f=C.defined,w=C.discardElement,p=C.find,K=C.fireEvent,r=C.isNumber,y=C.merge,B=C.pick,n=C.relativeLength,k=C.stableSort,b=C.syncTimeout;x=C.wrap;C=function(){function a(b,a){this.allItems=[];this.contentGroup=this.box=void 0;this.display=!1;this.group=void 0;this.offsetWidth=this.maxLegendWidth=this.maxItemWidth=
this.legendWidth=this.legendHeight=this.lastLineHeight=this.lastItemY=this.itemY=this.itemX=this.itemMarginTop=this.itemMarginBottom=this.itemHeight=this.initialItemY=0;this.options={};this.padding=0;this.pages=[];this.proximate=!1;this.scrollGroup=void 0;this.widthOption=this.totalItemWidth=this.titleHeight=this.symbolWidth=this.symbolHeight=0;this.chart=b;this.init(b,a)}a.prototype.init=function(b,a){this.chart=b;this.setOptions(a);a.enabled&&(this.render(),d(this.chart,"endResize",function(){this.legend.positionCheckboxes()}),
this.proximate?this.unchartrender=d(this.chart,"render",function(){this.legend.proximatePositions();this.legend.positionItems()}):this.unchartrender&&this.unchartrender())};a.prototype.setOptions=function(b){var a=B(b.padding,8);this.options=b;this.chart.styledMode||(this.itemStyle=b.itemStyle,this.itemHiddenStyle=y(this.itemStyle,b.itemHiddenStyle));this.itemMarginTop=b.itemMarginTop||0;this.itemMarginBottom=b.itemMarginBottom||0;this.padding=a;this.initialItemY=a-5;this.symbolWidth=B(b.symbolWidth,
16);this.pages=[];this.proximate="proximate"===b.layout&&!this.chart.inverted;this.baseline=void 0};a.prototype.update=function(b,a){var c=this.chart;this.setOptions(y(!0,this.options,b));this.destroy();c.isDirtyLegend=c.isDirtyBox=!0;B(a,!0)&&c.redraw();K(this,"afterUpdate")};a.prototype.colorizeItem=function(b,a){b.legendGroup[a?"removeClass":"addClass"]("highcharts-legend-item-hidden");if(!this.chart.styledMode){var c=this.options,e=b.legendItem,d=b.legendLine,g=b.legendSymbol,f=this.itemHiddenStyle.color;
c=a?c.itemStyle.color:f;var k=a?b.color||f:f,h=b.options&&b.options.marker,l={fill:k};e&&e.css({fill:c,color:c});d&&d.attr({stroke:k});g&&(h&&g.isMarker&&(l=b.pointAttribs(),a||(l.stroke=l.fill=f)),g.attr(l))}K(this,"afterColorizeItem",{item:b,visible:a})};a.prototype.positionItems=function(){this.allItems.forEach(this.positionItem,this);this.chart.isResizing||this.positionCheckboxes()};a.prototype.positionItem=function(b){var a=this,c=this.options,e=c.symbolPadding,d=!c.rtl,g=b._legendItemPos;c=
g[0];g=g[1];var k=b.checkbox,h=b.legendGroup;h&&h.element&&(e={translateX:d?c:this.legendWidth-c-2*e-4,translateY:g},d=function(){K(a,"afterPositionItem",{item:b})},f(h.translateY)?h.animate(e,void 0,d):(h.attr(e),d()));k&&(k.x=c,k.y=g)};a.prototype.destroyItem=function(b){var a=b.checkbox;["legendItem","legendLine","legendSymbol","legendGroup"].forEach(function(a){b[a]&&(b[a]=b[a].destroy())});a&&w(b.checkbox)};a.prototype.destroy=function(){function b(b){this[b]&&(this[b]=this[b].destroy())}this.getAllItems().forEach(function(a){["legendItem",
"legendGroup"].forEach(b,a)});"clipRect up down pager nav box title group".split(" ").forEach(b,this);this.display=null};a.prototype.positionCheckboxes=function(){var b=this.group&&this.group.alignAttr,a=this.clipHeight||this.legendHeight,c=this.titleHeight;if(b){var d=b.translateY;this.allItems.forEach(function(e){var g=e.checkbox;if(g){var f=d+c+g.y+(this.scrollOffset||0)+3;l(g,{left:b.translateX+e.checkboxOffset+g.x-20+"px",top:f+"px",display:this.proximate||f>d-6&&f<d+a-6?"":"none"})}},this)}};
a.prototype.renderTitle=function(){var b=this.options,a=this.padding,c=b.title,d=0;c.text&&(this.title||(this.title=this.chart.renderer.label(c.text,a-3,a-4,null,null,null,b.useHTML,null,"legend-title").attr({zIndex:1}),this.chart.styledMode||this.title.css(c.style),this.title.add(this.group)),c.width||this.title.css({width:this.maxLegendWidth+"px"}),b=this.title.getBBox(),d=b.height,this.offsetWidth=b.width,this.contentGroup.attr({translateY:d}));this.titleHeight=d};a.prototype.setText=function(b){var a=
this.options;b.legendItem.attr({text:a.labelFormat?q(a.labelFormat,b,this.chart):a.labelFormatter.call(b)})};a.prototype.renderItem=function(b){var a=this.chart,c=a.renderer,e=this.options,d=this.symbolWidth,g=e.symbolPadding||0,f=this.itemStyle,k=this.itemHiddenStyle,h="horizontal"===e.layout?B(e.itemDistance,20):0,l=!e.rtl,m=!b.series,n=!m&&b.series.drawLegendSymbol?b.series:b,p=n.options,r=this.createCheckboxForItem&&p&&p.showCheckbox,q=e.useHTML,w=b.options.className,F=b.legendItem;p=d+g+h+(r?
20:0);F||(b.legendGroup=c.g("legend-item").addClass("highcharts-"+n.type+"-series highcharts-color-"+b.colorIndex+(w?" "+w:"")+(m?" highcharts-series-"+b.index:"")).attr({zIndex:1}).add(this.scrollGroup),b.legendItem=F=c.text("",l?d+g:-g,this.baseline||0,q),a.styledMode||F.css(y(b.visible?f:k)),F.attr({align:l?"left":"right",zIndex:2}).add(b.legendGroup),this.baseline||(this.fontMetrics=c.fontMetrics(a.styledMode?12:f.fontSize,F),this.baseline=this.fontMetrics.f+3+this.itemMarginTop,F.attr("y",this.baseline),
this.symbolHeight=e.symbolHeight||this.fontMetrics.f,e.squareSymbol&&(this.symbolWidth=B(e.symbolWidth,Math.max(this.symbolHeight,16)),p=this.symbolWidth+g+h+(r?20:0),l&&F.attr("x",this.symbolWidth+g))),n.drawLegendSymbol(this,b),this.setItemEvents&&this.setItemEvents(b,F,q));r&&!b.checkbox&&this.createCheckboxForItem&&this.createCheckboxForItem(b);this.colorizeItem(b,b.visible);!a.styledMode&&f.width||F.css({width:(e.itemWidth||this.widthOption||a.spacingBox.width)-p+"px"});this.setText(b);a=F.getBBox();
b.itemWidth=b.checkboxOffset=e.itemWidth||b.legendItemWidth||a.width+p;this.maxItemWidth=Math.max(this.maxItemWidth,b.itemWidth);this.totalItemWidth+=b.itemWidth;this.itemHeight=b.itemHeight=Math.round(b.legendItemHeight||a.height||this.symbolHeight)};a.prototype.layoutItem=function(b){var a=this.options,c=this.padding,e="horizontal"===a.layout,d=b.itemHeight,g=this.itemMarginBottom,f=this.itemMarginTop,k=e?B(a.itemDistance,20):0,h=this.maxLegendWidth;a=a.alignColumns&&this.totalItemWidth>h?this.maxItemWidth:
b.itemWidth;e&&this.itemX-c+a>h&&(this.itemX=c,this.lastLineHeight&&(this.itemY+=f+this.lastLineHeight+g),this.lastLineHeight=0);this.lastItemY=f+this.itemY+g;this.lastLineHeight=Math.max(d,this.lastLineHeight);b._legendItemPos=[this.itemX,this.itemY];e?this.itemX+=a:(this.itemY+=f+d+g,this.lastLineHeight=d);this.offsetWidth=this.widthOption||Math.max((e?this.itemX-c-(b.checkbox?0:k):a)+c,this.offsetWidth)};a.prototype.getAllItems=function(){var b=[];this.chart.series.forEach(function(a){var c=a&&
a.options;a&&B(c.showInLegend,f(c.linkedTo)?!1:void 0,!0)&&(b=b.concat(a.legendItems||("point"===c.legendType?a.data:a)))});K(this,"afterGetAllItems",{allItems:b});return b};a.prototype.getAlignment=function(){var b=this.options;return this.proximate?b.align.charAt(0)+"tv":b.floating?"":b.align.charAt(0)+b.verticalAlign.charAt(0)+b.layout.charAt(0)};a.prototype.adjustMargins=function(b,a){var c=this.chart,e=this.options,d=this.getAlignment();d&&[/(lth|ct|rth)/,/(rtv|rm|rbv)/,/(rbh|cb|lbh)/,/(lbv|lm|ltv)/].forEach(function(g,
k){g.test(d)&&!f(b[k])&&(c[m[k]]=Math.max(c[m[k]],c.legend[(k+1)%2?"legendHeight":"legendWidth"]+[1,-1,-1,1][k]*e[k%2?"x":"y"]+B(e.margin,12)+a[k]+(c.titleOffset[k]||0)))})};a.prototype.proximatePositions=function(){var b=this.chart,a=[],c="left"===this.options.align;this.allItems.forEach(function(e){var d;var g=c;if(e.yAxis){e.xAxis.options.reversed&&(g=!g);e.points&&(d=p(g?e.points:e.points.slice(0).reverse(),function(b){return r(b.plotY)}));g=this.itemMarginTop+e.legendItem.getBBox().height+this.itemMarginBottom;
var f=e.yAxis.top-b.plotTop;e.visible?(d=d?d.plotY:e.yAxis.height,d+=f-.3*g):d=f+e.yAxis.height;a.push({target:d,size:g,item:e})}},this);h(a,b.plotHeight).forEach(function(a){a.item._legendItemPos&&(a.item._legendItemPos[1]=b.plotTop-b.spacing[0]+a.pos)})};a.prototype.render=function(){var b=this.chart,a=b.renderer,c=this.options,d=this.padding,g=this.getAllItems(),f=this.group,h=this.box;this.itemX=d;this.itemY=this.initialItemY;this.lastItemY=this.offsetWidth=0;this.widthOption=n(c.width,b.spacingBox.width-
d);var l=b.spacingBox.width-2*d-c.x;-1<["rm","lm"].indexOf(this.getAlignment().substring(0,2))&&(l/=2);this.maxLegendWidth=this.widthOption||l;f||(this.group=f=a.g("legend").addClass(c.className||"").attr({zIndex:7}).add(),this.contentGroup=a.g().attr({zIndex:1}).add(f),this.scrollGroup=a.g().add(this.contentGroup));this.renderTitle();k(g,function(b,a){return(b.options&&b.options.legendIndex||0)-(a.options&&a.options.legendIndex||0)});c.reversed&&g.reverse();this.allItems=g;this.display=l=!!g.length;
this.itemHeight=this.totalItemWidth=this.maxItemWidth=this.lastLineHeight=0;g.forEach(this.renderItem,this);g.forEach(this.layoutItem,this);g=(this.widthOption||this.offsetWidth)+d;var m=this.lastItemY+this.lastLineHeight+this.titleHeight;m=this.handleOverflow(m);m+=d;h||(this.box=h=a.rect().addClass("highcharts-legend-box").attr({r:c.borderRadius}).add(f),h.isNew=!0);b.styledMode||h.attr({stroke:c.borderColor,"stroke-width":c.borderWidth||0,fill:c.backgroundColor||"none"}).shadow(c.shadow);0<g&&
0<m&&(h[h.isNew?"attr":"animate"](h.crisp.call({},{x:0,y:0,width:g,height:m},h.strokeWidth())),h.isNew=!1);h[l?"show":"hide"]();b.styledMode&&"none"===f.getStyle("display")&&(g=m=0);this.legendWidth=g;this.legendHeight=m;l&&this.align();this.proximate||this.positionItems();K(this,"afterRender")};a.prototype.align=function(b){void 0===b&&(b=this.chart.spacingBox);var a=this.chart,c=this.options,e=b.y;/(lth|ct|rth)/.test(this.getAlignment())&&0<a.titleOffset[0]?e+=a.titleOffset[0]:/(lbh|cb|rbh)/.test(this.getAlignment())&&
0<a.titleOffset[2]&&(e-=a.titleOffset[2]);e!==b.y&&(b=y(b,{y:e}));this.group.align(y(c,{width:this.legendWidth,height:this.legendHeight,verticalAlign:this.proximate?"top":c.verticalAlign}),!0,b)};a.prototype.handleOverflow=function(b){var a=this,c=this.chart,e=c.renderer,d=this.options,g=d.y,f="top"===d.verticalAlign,k=this.padding,h=d.maxHeight,l=d.navigation,m=B(l.animation,!0),n=l.arrowSize||12,p=this.pages,r=this.allItems,q=function(b){"number"===typeof b?y.attr({height:b}):y&&(a.clipRect=y.destroy(),
a.contentGroup.clip());a.contentGroup.div&&(a.contentGroup.div.style.clip=b?"rect("+k+"px,9999px,"+(k+b)+"px,0)":"auto")},w=function(b){a[b]=e.circle(0,0,1.3*n).translate(n/2,n/2).add(N);c.styledMode||a[b].attr("fill","rgba(0,0,0,0.0001)");return a[b]},F,u;g=c.spacingBox.height+(f?-g:g)-k;var N=this.nav,y=this.clipRect;"horizontal"!==d.layout||"middle"===d.verticalAlign||d.floating||(g/=2);h&&(g=Math.min(g,h));p.length=0;b&&0<g&&b>g&&!1!==l.enabled?(this.clipHeight=F=Math.max(g-20-this.titleHeight-
k,0),this.currentPage=B(this.currentPage,1),this.fullHeight=b,r.forEach(function(b,a){var c=b._legendItemPos[1],e=Math.round(b.legendItem.getBBox().height),d=p.length;if(!d||c-p[d-1]>F&&(u||c)!==p[d-1])p.push(u||c),d++;b.pageIx=d-1;u&&(r[a-1].pageIx=d-1);a===r.length-1&&c+e-p[d-1]>F&&e<=F&&(p.push(c),b.pageIx=d);c!==u&&(u=c)}),y||(y=a.clipRect=e.clipRect(0,k,9999,0),a.contentGroup.clip(y)),q(F),N||(this.nav=N=e.g().attr({zIndex:1}).add(this.group),this.up=e.symbol("triangle",0,0,n,n).add(N),w("upTracker").on("click",
function(){a.scroll(-1,m)}),this.pager=e.text("",15,10).addClass("highcharts-legend-navigation"),c.styledMode||this.pager.css(l.style),this.pager.add(N),this.down=e.symbol("triangle-down",0,0,n,n).add(N),w("downTracker").on("click",function(){a.scroll(1,m)})),a.scroll(0),b=g):N&&(q(),this.nav=N.destroy(),this.scrollGroup.attr({translateY:1}),this.clipHeight=0);return b};a.prototype.scroll=function(a,c){var e=this,d=this.chart,g=this.pages,f=g.length,k=this.clipHeight,h=this.options.navigation,l=this.pager,
m=this.padding,n=this.currentPage+a;n>f&&(n=f);0<n&&("undefined"!==typeof c&&z(c,d),this.nav.attr({translateX:m,translateY:k+this.padding+7+this.titleHeight,visibility:"visible"}),[this.up,this.upTracker].forEach(function(b){b.attr({"class":1===n?"highcharts-legend-nav-inactive":"highcharts-legend-nav-active"})}),l.attr({text:n+"/"+f}),[this.down,this.downTracker].forEach(function(b){b.attr({x:18+this.pager.getBBox().width,"class":n===f?"highcharts-legend-nav-inactive":"highcharts-legend-nav-active"})},
this),d.styledMode||(this.up.attr({fill:1===n?h.inactiveColor:h.activeColor}),this.upTracker.css({cursor:1===n?"default":"pointer"}),this.down.attr({fill:n===f?h.inactiveColor:h.activeColor}),this.downTracker.css({cursor:n===f?"default":"pointer"})),this.scrollOffset=-g[n-1]+this.initialItemY,this.scrollGroup.animate({translateY:this.scrollOffset}),this.currentPage=n,this.positionCheckboxes(),a=u(B(c,d.renderer.globalAnimation,!0)),b(function(){K(e,"afterScroll",{currentPage:n})},a.duration))};a.prototype.setItemEvents=
function(b,a,c){var e=this,d=e.chart.renderer.boxWrapper,g=b instanceof G,f="highcharts-legend-"+(g?"point":"series")+"-active",k=e.chart.styledMode,h=function(a){e.allItems.forEach(function(c){b!==c&&[c].concat(c.linkedSeries||[]).forEach(function(b){b.setState(a,!g)})})};(c?[a,b.legendSymbol]:[b.legendGroup]).forEach(function(c){if(c)c.on("mouseover",function(){b.visible&&h("inactive");b.setState("hover");b.visible&&d.addClass(f);k||a.css(e.options.itemHoverStyle)}).on("mouseout",function(){e.chart.styledMode||
a.css(y(b.visible?e.itemStyle:e.itemHiddenStyle));h("");d.removeClass(f);b.setState()}).on("click",function(a){var c=function(){b.setVisible&&b.setVisible();h(b.visible?"inactive":"")};d.removeClass(f);a={browserEvent:a};b.firePointEvent?b.firePointEvent("legendItemClick",a,c):K(b,"legendItemClick",a,c)})})};a.prototype.createCheckboxForItem=function(b){b.checkbox=c("input",{type:"checkbox",className:"highcharts-legend-checkbox",checked:b.selected,defaultChecked:b.selected},this.options.itemCheckboxStyle,
this.chart.container);d(b.checkbox,"click",function(a){K(b.series||b,"checkboxClick",{checked:a.target.checked,item:b},function(){b.select()})})};return a}();(/Trident\/7\.0/.test(A.navigator&&A.navigator.userAgent)||a)&&x(C.prototype,"positionItem",function(b,a){var c=this,e=function(){a._legendItemPos&&b.call(c,a)};e();c.bubbleLegend||setTimeout(e)});"";return C});L(a,"Core/Series/SeriesRegistry.js",[a["Core/Globals.js"],a["Core/DefaultOptions.js"],a["Core/Series/Point.js"],a["Core/Utilities.js"]],
function(a,t,A,G){var u=t.defaultOptions,C=G.error,I=G.extendClass,z=G.merge,q;(function(m){function h(a,c){var d=u.plotOptions||{},f=c.defaultOptions;c.prototype.pointClass||(c.prototype.pointClass=A);c.prototype.type=a;f&&(d[a]=f);m.seriesTypes[a]=c}m.seriesTypes=a.seriesTypes;m.getSeries=function(a,c){void 0===c&&(c={});var d=a.options.chart;d=c.type||d.type||d.defaultSeriesType||"";var f=m.seriesTypes[d];m||C(17,!0,a,{missingModuleFor:d});d=new f;"function"===typeof d.init&&d.init(a,c);return d};
m.registerSeriesType=h;m.seriesType=function(a,c,l,f,q){var d=u.plotOptions||{};c=c||"";d[a]=z(d[c],l);h(a,I(m.seriesTypes[c]||function(){},f));m.seriesTypes[a].prototype.type=a;q&&(m.seriesTypes[a].prototype.pointClass=I(A,q));return m.seriesTypes[a]}})(q||(q={}));return q});L(a,"Core/Chart/Chart.js",[a["Core/Animation/AnimationUtilities.js"],a["Core/Axis/Axis.js"],a["Core/FormatUtilities.js"],a["Core/Foundation.js"],a["Core/Globals.js"],a["Core/Legend/Legend.js"],a["Core/MSPointer.js"],a["Core/DefaultOptions.js"],
a["Core/Pointer.js"],a["Core/Renderer/RendererRegistry.js"],a["Core/Series/SeriesRegistry.js"],a["Core/Renderer/SVG/SVGRenderer.js"],a["Core/Time.js"],a["Core/Utilities.js"],a["Core/Renderer/HTML/AST.js"]],function(a,t,A,G,x,C,I,z,q,m,h,d,c,l,f){var w=a.animate,p=a.animObject,u=a.setAnimation,r=A.numberFormat,y=G.registerEventOptions,B=x.charts,n=x.doc,k=x.marginNames,b=x.svg,g=x.win,e=z.defaultOptions,D=z.defaultTime,H=h.seriesTypes,v=l.addEvent,E=l.attr,O=l.cleanRecursively,S=l.createElement,P=
l.css,U=l.defined,Y=l.discardElement,J=l.erase,M=l.error,L=l.extend,da=l.find,Q=l.fireEvent,ea=l.getStyle,F=l.isArray,T=l.isNumber,N=l.isObject,V=l.isString,W=l.merge,X=l.objectEach,R=l.pick,fa=l.pInt,aa=l.relativeLength,ia=l.removeEvent,ha=l.splat,ba=l.syncTimeout,ka=l.uniqueKey;a=function(){function a(b,a,c){this.series=this.renderTo=this.renderer=this.pointer=this.pointCount=this.plotWidth=this.plotTop=this.plotLeft=this.plotHeight=this.plotBox=this.options=this.numberFormatter=this.margin=this.legend=
this.labelCollectors=this.isResizing=this.index=this.eventOptions=this.container=this.colorCounter=this.clipBox=this.chartWidth=this.chartHeight=this.bounds=this.axisOffset=this.axes=void 0;this.sharedClips={};this.yAxis=this.xAxis=this.userOptions=this.titleOffset=this.time=this.symbolCounter=this.spacingBox=this.spacing=void 0;this.getArgs(b,a,c)}a.chart=function(b,c,e){return new a(b,c,e)};a.prototype.getArgs=function(b,a,c){V(b)||b.nodeName?(this.renderTo=b,this.init(a,c)):this.init(b,a)};a.prototype.init=
function(b,a){var d=b.plotOptions||{};Q(this,"init",{args:arguments},function(){var g=W(e,b),f=g.chart;X(g.plotOptions,function(b,a){N(b)&&(b.tooltip=d[a]&&W(d[a].tooltip)||void 0)});g.tooltip.userOptions=b.chart&&b.chart.forExport&&b.tooltip.userOptions||b.tooltip;this.userOptions=b;this.margin=[];this.spacing=[];this.bounds={h:{},v:{}};this.labelCollectors=[];this.callback=a;this.isResizing=0;this.options=g;this.axes=[];this.series=[];this.time=b.time&&Object.keys(b.time).length?new c(b.time):x.time;
this.numberFormatter=f.numberFormatter||r;this.styledMode=f.styledMode;this.hasCartesianSeries=f.showAxes;this.index=B.length;B.push(this);x.chartCount++;y(this,f);this.xAxis=[];this.yAxis=[];this.pointCount=this.colorCounter=this.symbolCounter=0;Q(this,"afterInit");this.firstRender()})};a.prototype.initSeries=function(b){var a=this.options.chart;a=b.type||a.type||a.defaultSeriesType;var c=H[a];c||M(17,!0,this,{missingModuleFor:a});a=new c;"function"===typeof a.init&&a.init(this,b);return a};a.prototype.setSeriesData=
function(){this.getSeriesOrderByLinks().forEach(function(b){b.points||b.data||!b.enabledDataSorting||b.setData(b.options.data,!1)})};a.prototype.getSeriesOrderByLinks=function(){return this.series.concat().sort(function(b,a){return b.linkedSeries.length||a.linkedSeries.length?a.linkedSeries.length-b.linkedSeries.length:0})};a.prototype.orderSeries=function(b){var a=this.series;b=b||0;for(var c=a.length;b<c;++b)a[b]&&(a[b].index=b,a[b].name=a[b].getName())};a.prototype.isInsidePlot=function(b,a,c){void 0===
c&&(c={});var e=this.inverted,d=this.plotBox,g=this.plotLeft,f=this.plotTop,k=this.scrollablePlotBox,h=0;var l=0;c.visiblePlotOnly&&this.scrollingContainer&&(l=this.scrollingContainer,h=l.scrollLeft,l=l.scrollTop);var m=c.series;d=c.visiblePlotOnly&&k||d;k=c.inverted?a:b;a=c.inverted?b:a;b={x:k,y:a,isInsidePlot:!0};if(!c.ignoreX){var n=m&&(e?m.yAxis:m.xAxis)||{pos:g,len:Infinity};k=c.paneCoordinates?n.pos+k:g+k;k>=Math.max(h+g,n.pos)&&k<=Math.min(h+g+d.width,n.pos+n.len)||(b.isInsidePlot=!1)}!c.ignoreY&&
b.isInsidePlot&&(e=m&&(e?m.xAxis:m.yAxis)||{pos:f,len:Infinity},c=c.paneCoordinates?e.pos+a:f+a,c>=Math.max(l+f,e.pos)&&c<=Math.min(l+f+d.height,e.pos+e.len)||(b.isInsidePlot=!1));Q(this,"afterIsInsidePlot",b);return b.isInsidePlot};a.prototype.redraw=function(b){Q(this,"beforeRedraw");var a=this.hasCartesianSeries?this.axes:this.colorAxis||[],c=this.series,e=this.pointer,d=this.legend,g=this.userOptions.legend,f=this.renderer,k=f.isHidden(),h=[],l=this.isDirtyBox,m=this.isDirtyLegend;this.setResponsive&&
this.setResponsive(!1);u(this.hasRendered?b:!1,this);k&&this.temporaryDisplay();this.layOutTitles();for(b=c.length;b--;){var n=c[b];if(n.options.stacking||n.options.centerInCategory){var p=!0;if(n.isDirty){var F=!0;break}}}if(F)for(b=c.length;b--;)n=c[b],n.options.stacking&&(n.isDirty=!0);c.forEach(function(b){b.isDirty&&("point"===b.options.legendType?("function"===typeof b.updateTotals&&b.updateTotals(),m=!0):g&&(g.labelFormatter||g.labelFormat)&&(m=!0));b.isDirtyData&&Q(b,"updatedData")});m&&d&&
d.options.enabled&&(d.render(),this.isDirtyLegend=!1);p&&this.getStacks();a.forEach(function(b){b.updateNames();b.setScale()});this.getMargins();a.forEach(function(b){b.isDirty&&(l=!0)});a.forEach(function(b){var a=b.min+","+b.max;b.extKey!==a&&(b.extKey=a,h.push(function(){Q(b,"afterSetExtremes",L(b.eventArgs,b.getExtremes()));delete b.eventArgs}));(l||p)&&b.redraw()});l&&this.drawChartBox();Q(this,"predraw");c.forEach(function(b){(l||b.isDirty)&&b.visible&&b.redraw();b.isDirtyData=!1});e&&e.reset(!0);
f.draw();Q(this,"redraw");Q(this,"render");k&&this.temporaryDisplay(!0);h.forEach(function(b){b.call()})};a.prototype.get=function(b){function a(a){return a.id===b||a.options&&a.options.id===b}for(var c=this.series,e=da(this.axes,a)||da(this.series,a),d=0;!e&&d<c.length;d++)e=da(c[d].points||[],a);return e};a.prototype.getAxes=function(){var b=this,a=this.options,c=a.xAxis=ha(a.xAxis||{});a=a.yAxis=ha(a.yAxis||{});Q(this,"getAxes");c.forEach(function(b,a){b.index=a;b.isX=!0});a.forEach(function(b,
a){b.index=a});c.concat(a).forEach(function(a){new t(b,a)});Q(this,"afterGetAxes")};a.prototype.getSelectedPoints=function(){return this.series.reduce(function(b,a){a.getPointsCollection().forEach(function(a){R(a.selectedStaging,a.selected)&&b.push(a)});return b},[])};a.prototype.getSelectedSeries=function(){return this.series.filter(function(b){return b.selected})};a.prototype.setTitle=function(b,a,c){this.applyDescription("title",b);this.applyDescription("subtitle",a);this.applyDescription("caption",
void 0);this.layOutTitles(c)};a.prototype.applyDescription=function(b,a){var c=this,e="title"===b?{color:"#333333",fontSize:this.options.isStock?"16px":"18px"}:{color:"#666666"};e=this.options[b]=W(!this.styledMode&&{style:e},this.options[b],a);var d=this[b];d&&a&&(this[b]=d=d.destroy());e&&!d&&(d=this.renderer.text(e.text,0,0,e.useHTML).attr({align:e.align,"class":"highcharts-"+b,zIndex:e.zIndex||4}).add(),d.update=function(a){c[{title:"setTitle",subtitle:"setSubtitle",caption:"setCaption"}[b]](a)},
this.styledMode||d.css(e.style),this[b]=d)};a.prototype.layOutTitles=function(b){var a=[0,0,0],c=this.renderer,e=this.spacingBox;["title","subtitle","caption"].forEach(function(b){var d=this[b],g=this.options[b],f=g.verticalAlign||"top";b="title"===b?"top"===f?-3:0:"top"===f?a[0]+2:0;var k;if(d){this.styledMode||(k=g.style&&g.style.fontSize);k=c.fontMetrics(k,d).b;d.css({width:(g.width||e.width+(g.widthAdjust||0))+"px"});var h=Math.round(d.getBBox(g.useHTML).height);d.align(L({y:"bottom"===f?k:b+
k,height:h},g),!1,"spacingBox");g.floating||("top"===f?a[0]=Math.ceil(a[0]+h):"bottom"===f&&(a[2]=Math.ceil(a[2]+h)))}},this);a[0]&&"top"===(this.options.title.verticalAlign||"top")&&(a[0]+=this.options.title.margin);a[2]&&"bottom"===this.options.caption.verticalAlign&&(a[2]+=this.options.caption.margin);var d=!this.titleOffset||this.titleOffset.join(",")!==a.join(",");this.titleOffset=a;Q(this,"afterLayOutTitles");!this.isDirtyBox&&d&&(this.isDirtyBox=this.isDirtyLegend=d,this.hasRendered&&R(b,!0)&&
this.isDirtyBox&&this.redraw())};a.prototype.getChartSize=function(){var b=this.options.chart,a=b.width;b=b.height;var c=this.renderTo;U(a)||(this.containerWidth=ea(c,"width"));U(b)||(this.containerHeight=ea(c,"height"));this.chartWidth=Math.max(0,a||this.containerWidth||600);this.chartHeight=Math.max(0,aa(b,this.chartWidth)||(1<this.containerHeight?this.containerHeight:400))};a.prototype.temporaryDisplay=function(b){var a=this.renderTo;if(b)for(;a&&a.style;)a.hcOrigStyle&&(P(a,a.hcOrigStyle),delete a.hcOrigStyle),
a.hcOrigDetached&&(n.body.removeChild(a),a.hcOrigDetached=!1),a=a.parentNode;else for(;a&&a.style;){n.body.contains(a)||a.parentNode||(a.hcOrigDetached=!0,n.body.appendChild(a));if("none"===ea(a,"display",!1)||a.hcOricDetached)a.hcOrigStyle={display:a.style.display,height:a.style.height,overflow:a.style.overflow},b={display:"block",overflow:"hidden"},a!==this.renderTo&&(b.height=0),P(a,b),a.offsetWidth||a.style.setProperty("display","block","important");a=a.parentNode;if(a===n.body)break}};a.prototype.setClassName=
function(b){this.container.className="highcharts-container "+(b||"")};a.prototype.getContainer=function(){var a=this.options,c=a.chart,e=ka(),g,f=this.renderTo;f||(this.renderTo=f=c.renderTo);V(f)&&(this.renderTo=f=n.getElementById(f));f||M(13,!0,this);var k=fa(E(f,"data-highcharts-chart"));T(k)&&B[k]&&B[k].hasRendered&&B[k].destroy();E(f,"data-highcharts-chart",this.index);f.innerHTML="";c.skipClone||f.offsetWidth||this.temporaryDisplay();this.getChartSize();k=this.chartWidth;var h=this.chartHeight;
P(f,{overflow:"hidden"});this.styledMode||(g=L({position:"relative",overflow:"hidden",width:k+"px",height:h+"px",textAlign:"left",lineHeight:"normal",zIndex:0,"-webkit-tap-highlight-color":"rgba(0,0,0,0)",userSelect:"none","touch-action":"manipulation",outline:"none"},c.style||{}));this.container=e=S("div",{id:e},g,f);this._cursor=e.style.cursor;this.renderer=new (c.renderer||!b?m.getRendererType(c.renderer):d)(e,k,h,void 0,c.forExport,a.exporting&&a.exporting.allowHTML,this.styledMode);u(void 0,
this);this.setClassName(c.className);if(this.styledMode)for(var l in a.defs)this.renderer.definition(a.defs[l]);else this.renderer.setStyle(c.style);this.renderer.chartIndex=this.index;Q(this,"afterGetContainer")};a.prototype.getMargins=function(b){var a=this.spacing,c=this.margin,e=this.titleOffset;this.resetMargins();e[0]&&!U(c[0])&&(this.plotTop=Math.max(this.plotTop,e[0]+a[0]));e[2]&&!U(c[2])&&(this.marginBottom=Math.max(this.marginBottom,e[2]+a[2]));this.legend&&this.legend.display&&this.legend.adjustMargins(c,
a);Q(this,"getMargins");b||this.getAxisMargins()};a.prototype.getAxisMargins=function(){var b=this,a=b.axisOffset=[0,0,0,0],c=b.colorAxis,e=b.margin,d=function(b){b.forEach(function(b){b.visible&&b.getOffset()})};b.hasCartesianSeries?d(b.axes):c&&c.length&&d(c);k.forEach(function(c,d){U(e[d])||(b[c]+=a[d])});b.setChartSize()};a.prototype.reflow=function(b){var a=this,c=a.options.chart,e=a.renderTo,d=U(c.width)&&U(c.height),f=c.width||ea(e,"width");c=c.height||ea(e,"height");e=b?b.target:g;delete a.pointer.chartPosition;
if(!d&&!a.isPrinting&&f&&c&&(e===g||e===n)){if(f!==a.containerWidth||c!==a.containerHeight)l.clearTimeout(a.reflowTimeout),a.reflowTimeout=ba(function(){a.container&&a.setSize(void 0,void 0,!1)},b?100:0);a.containerWidth=f;a.containerHeight=c}};a.prototype.setReflow=function(b){var a=this;!1===b||this.unbindReflow?!1===b&&this.unbindReflow&&(this.unbindReflow=this.unbindReflow()):(this.unbindReflow=v(g,"resize",function(b){a.options&&a.reflow(b)}),v(this,"destroy",this.unbindReflow))};a.prototype.setSize=
function(b,a,c){var e=this,d=e.renderer;e.isResizing+=1;u(c,e);c=d.globalAnimation;e.oldChartHeight=e.chartHeight;e.oldChartWidth=e.chartWidth;"undefined"!==typeof b&&(e.options.chart.width=b);"undefined"!==typeof a&&(e.options.chart.height=a);e.getChartSize();e.styledMode||(c?w:P)(e.container,{width:e.chartWidth+"px",height:e.chartHeight+"px"},c);e.setChartSize(!0);d.setSize(e.chartWidth,e.chartHeight,c);e.axes.forEach(function(b){b.isDirty=!0;b.setScale()});e.isDirtyLegend=!0;e.isDirtyBox=!0;e.layOutTitles();
e.getMargins();e.redraw(c);e.oldChartHeight=null;Q(e,"resize");ba(function(){e&&Q(e,"endResize",null,function(){--e.isResizing})},p(c).duration)};a.prototype.setChartSize=function(b){var a=this.inverted,c=this.renderer,e=this.chartWidth,d=this.chartHeight,g=this.options.chart,f=this.spacing,k=this.clipOffset,h,l,m,n;this.plotLeft=h=Math.round(this.plotLeft);this.plotTop=l=Math.round(this.plotTop);this.plotWidth=m=Math.max(0,Math.round(e-h-this.marginRight));this.plotHeight=n=Math.max(0,Math.round(d-
l-this.marginBottom));this.plotSizeX=a?n:m;this.plotSizeY=a?m:n;this.plotBorderWidth=g.plotBorderWidth||0;this.spacingBox=c.spacingBox={x:f[3],y:f[0],width:e-f[3]-f[1],height:d-f[0]-f[2]};this.plotBox=c.plotBox={x:h,y:l,width:m,height:n};a=2*Math.floor(this.plotBorderWidth/2);e=Math.ceil(Math.max(a,k[3])/2);d=Math.ceil(Math.max(a,k[0])/2);this.clipBox={x:e,y:d,width:Math.floor(this.plotSizeX-Math.max(a,k[1])/2-e),height:Math.max(0,Math.floor(this.plotSizeY-Math.max(a,k[2])/2-d))};b||(this.axes.forEach(function(b){b.setAxisSize();
b.setAxisTranslation()}),c.alignElements());Q(this,"afterSetChartSize",{skipAxes:b})};a.prototype.resetMargins=function(){Q(this,"resetMargins");var b=this,a=b.options.chart;["margin","spacing"].forEach(function(c){var e=a[c],d=N(e)?e:[e,e,e,e];["Top","Right","Bottom","Left"].forEach(function(e,g){b[c][g]=R(a[c+e],d[g])})});k.forEach(function(a,c){b[a]=R(b.margin[c],b.spacing[c])});b.axisOffset=[0,0,0,0];b.clipOffset=[0,0,0,0]};a.prototype.drawChartBox=function(){var b=this.options.chart,a=this.renderer,
c=this.chartWidth,e=this.chartHeight,d=this.styledMode,g=this.plotBGImage,f=b.backgroundColor,k=b.plotBackgroundColor,h=b.plotBackgroundImage,l=this.plotLeft,m=this.plotTop,n=this.plotWidth,p=this.plotHeight,F=this.plotBox,r=this.clipRect,N=this.clipBox,q=this.chartBackground,v=this.plotBackground,B=this.plotBorder,w,u="animate";q||(this.chartBackground=q=a.rect().addClass("highcharts-background").add(),u="attr");if(d)var y=w=q.strokeWidth();else{y=b.borderWidth||0;w=y+(b.shadow?8:0);f={fill:f||"none"};
if(y||q["stroke-width"])f.stroke=b.borderColor,f["stroke-width"]=y;q.attr(f).shadow(b.shadow)}q[u]({x:w/2,y:w/2,width:c-w-y%2,height:e-w-y%2,r:b.borderRadius});u="animate";v||(u="attr",this.plotBackground=v=a.rect().addClass("highcharts-plot-background").add());v[u](F);d||(v.attr({fill:k||"none"}).shadow(b.plotShadow),h&&(g?(h!==g.attr("href")&&g.attr("href",h),g.animate(F)):this.plotBGImage=a.image(h,l,m,n,p).add()));r?r.animate({width:N.width,height:N.height}):this.clipRect=a.clipRect(N);u="animate";
B||(u="attr",this.plotBorder=B=a.rect().addClass("highcharts-plot-border").attr({zIndex:1}).add());d||B.attr({stroke:b.plotBorderColor,"stroke-width":b.plotBorderWidth||0,fill:"none"});B[u](B.crisp({x:l,y:m,width:n,height:p},-B.strokeWidth()));this.isDirtyBox=!1;Q(this,"afterDrawChartBox")};a.prototype.propFromSeries=function(){var b=this,a=b.options.chart,c=b.options.series,e,d,g;["inverted","angular","polar"].forEach(function(f){d=H[a.type||a.defaultSeriesType];g=a[f]||d&&d.prototype[f];for(e=c&&
c.length;!g&&e--;)(d=H[c[e].type])&&d.prototype[f]&&(g=!0);b[f]=g})};a.prototype.linkSeries=function(){var b=this,a=b.series;a.forEach(function(b){b.linkedSeries.length=0});a.forEach(function(a){var c=a.options.linkedTo;V(c)&&(c=":previous"===c?b.series[a.index-1]:b.get(c))&&c.linkedParent!==a&&(c.linkedSeries.push(a),a.linkedParent=c,c.enabledDataSorting&&a.setDataSortingOptions(),a.visible=R(a.options.visible,c.options.visible,a.visible))});Q(this,"afterLinkSeries")};a.prototype.renderSeries=function(){this.series.forEach(function(b){b.translate();
b.render()})};a.prototype.renderLabels=function(){var b=this,a=b.options.labels;a.items&&a.items.forEach(function(c){var e=L(a.style,c.style),d=fa(e.left)+b.plotLeft,g=fa(e.top)+b.plotTop+12;delete e.left;delete e.top;b.renderer.text(c.html,d,g).attr({zIndex:2}).css(e).add()})};a.prototype.render=function(){var b=this.axes,a=this.colorAxis,c=this.renderer,e=this.options,d=function(b){b.forEach(function(b){b.visible&&b.render()})},g=0;this.setTitle();this.legend=new C(this,e.legend);this.getStacks&&
this.getStacks();this.getMargins(!0);this.setChartSize();e=this.plotWidth;b.some(function(b){if(b.horiz&&b.visible&&b.options.labels.enabled&&b.series.length)return g=21,!0});var f=this.plotHeight=Math.max(this.plotHeight-g,0);b.forEach(function(b){b.setScale()});this.getAxisMargins();var k=1.1<e/this.plotWidth,h=1.05<f/this.plotHeight;if(k||h)b.forEach(function(b){(b.horiz&&k||!b.horiz&&h)&&b.setTickInterval(!0)}),this.getMargins();this.drawChartBox();this.hasCartesianSeries?d(b):a&&a.length&&d(a);
this.seriesGroup||(this.seriesGroup=c.g("series-group").attr({zIndex:3}).add());this.renderSeries();this.renderLabels();this.addCredits();this.setResponsive&&this.setResponsive();this.hasRendered=!0};a.prototype.addCredits=function(b){var a=this,c=W(!0,this.options.credits,b);c.enabled&&!this.credits&&(this.credits=this.renderer.text(c.text+(this.mapCredits||""),0,0).addClass("highcharts-credits").on("click",function(){c.href&&(g.location.href=c.href)}).attr({align:c.position.align,zIndex:8}),a.styledMode||
this.credits.css(c.style),this.credits.add().align(c.position),this.credits.update=function(b){a.credits=a.credits.destroy();a.addCredits(b)})};a.prototype.destroy=function(){var b=this,a=b.axes,c=b.series,e=b.container,d=e&&e.parentNode,g;Q(b,"destroy");b.renderer.forExport?J(B,b):B[b.index]=void 0;x.chartCount--;b.renderTo.removeAttribute("data-highcharts-chart");ia(b);for(g=a.length;g--;)a[g]=a[g].destroy();this.scroller&&this.scroller.destroy&&this.scroller.destroy();for(g=c.length;g--;)c[g]=
c[g].destroy();"title subtitle chartBackground plotBackground plotBGImage plotBorder seriesGroup clipRect credits pointer rangeSelector legend resetZoomButton tooltip renderer".split(" ").forEach(function(a){var c=b[a];c&&c.destroy&&(b[a]=c.destroy())});e&&(e.innerHTML="",ia(e),d&&Y(e));X(b,function(a,c){delete b[c]})};a.prototype.firstRender=function(){var b=this,a=b.options;if(!b.isReadyToRender||b.isReadyToRender()){b.getContainer();b.resetMargins();b.setChartSize();b.propFromSeries();b.getAxes();
(F(a.series)?a.series:[]).forEach(function(a){b.initSeries(a)});b.linkSeries();b.setSeriesData();Q(b,"beforeRender");q&&(I.isRequired()?b.pointer=new I(b,a):b.pointer=new q(b,a));b.render();b.pointer.getChartPosition();if(!b.renderer.imgCount&&!b.hasLoaded)b.onload();b.temporaryDisplay(!0)}};a.prototype.onload=function(){this.callbacks.concat([this.callback]).forEach(function(b){b&&"undefined"!==typeof this.index&&b.apply(this,[this])},this);Q(this,"load");Q(this,"render");U(this.index)&&this.setReflow(this.options.chart.reflow);
this.hasLoaded=!0};a.prototype.addSeries=function(b,a,c){var e=this,d;b&&(a=R(a,!0),Q(e,"addSeries",{options:b},function(){d=e.initSeries(b);e.isDirtyLegend=!0;e.linkSeries();d.enabledDataSorting&&d.setData(b.data,!1);Q(e,"afterAddSeries",{series:d});a&&e.redraw(c)}));return d};a.prototype.addAxis=function(b,a,c,e){return this.createAxis(a?"xAxis":"yAxis",{axis:b,redraw:c,animation:e})};a.prototype.addColorAxis=function(b,a,c){return this.createAxis("colorAxis",{axis:b,redraw:a,animation:c})};a.prototype.createAxis=
function(b,a){b=new t(this,W(a.axis,{index:this[b].length,isX:"xAxis"===b}));R(a.redraw,!0)&&this.redraw(a.animation);return b};a.prototype.showLoading=function(b){var a=this,c=a.options,e=c.loading,d=function(){g&&P(g,{left:a.plotLeft+"px",top:a.plotTop+"px",width:a.plotWidth+"px",height:a.plotHeight+"px"})},g=a.loadingDiv,k=a.loadingSpan;g||(a.loadingDiv=g=S("div",{className:"highcharts-loading highcharts-loading-hidden"},null,a.container));k||(a.loadingSpan=k=S("span",{className:"highcharts-loading-inner"},
null,g),v(a,"redraw",d));g.className="highcharts-loading";f.setElementHTML(k,R(b,c.lang.loading,""));a.styledMode||(P(g,L(e.style,{zIndex:10})),P(k,e.labelStyle),a.loadingShown||(P(g,{opacity:0,display:""}),w(g,{opacity:e.style.opacity||.5},{duration:e.showDuration||0})));a.loadingShown=!0;d()};a.prototype.hideLoading=function(){var b=this.options,a=this.loadingDiv;a&&(a.className="highcharts-loading highcharts-loading-hidden",this.styledMode||w(a,{opacity:0},{duration:b.loading.hideDuration||100,
complete:function(){P(a,{display:"none"})}}));this.loadingShown=!1};a.prototype.update=function(b,a,e,d){var g=this,f={credits:"addCredits",title:"setTitle",subtitle:"setSubtitle",caption:"setCaption"},k=b.isResponsiveOptions,h=[],l,m;Q(g,"update",{options:b});k||g.setResponsive(!1,!0);b=O(b,g.options);g.userOptions=W(g.userOptions,b);var n=b.chart;if(n){W(!0,g.options.chart,n);"className"in n&&g.setClassName(n.className);"reflow"in n&&g.setReflow(n.reflow);if("inverted"in n||"polar"in n||"type"in
n){g.propFromSeries();var p=!0}"alignTicks"in n&&(p=!0);"events"in n&&y(this,n);X(n,function(b,a){-1!==g.propsRequireUpdateSeries.indexOf("chart."+a)&&(l=!0);-1!==g.propsRequireDirtyBox.indexOf(a)&&(g.isDirtyBox=!0);-1!==g.propsRequireReflow.indexOf(a)&&(k?g.isDirtyBox=!0:m=!0)});!g.styledMode&&n.style&&g.renderer.setStyle(g.options.chart.style||{})}!g.styledMode&&b.colors&&(this.options.colors=b.colors);b.time&&(this.time===D&&(this.time=new c(b.time)),W(!0,g.options.time,b.time));X(b,function(a,
c){if(g[c]&&"function"===typeof g[c].update)g[c].update(a,!1);else if("function"===typeof g[f[c]])g[f[c]](a);else"colors"!==c&&-1===g.collectionsWithUpdate.indexOf(c)&&W(!0,g.options[c],b[c]);"chart"!==c&&-1!==g.propsRequireUpdateSeries.indexOf(c)&&(l=!0)});this.collectionsWithUpdate.forEach(function(a){if(b[a]){var c=[];g[a].forEach(function(b,a){b.options.isInternal||c.push(R(b.options.index,a))});ha(b[a]).forEach(function(b,d){var f=U(b.id),k;f&&(k=g.get(b.id));!k&&g[a]&&(k=g[a][c?c[d]:d])&&f&&
U(k.options.id)&&(k=void 0);k&&k.coll===a&&(k.update(b,!1),e&&(k.touched=!0));!k&&e&&g.collectionsWithInit[a]&&(g.collectionsWithInit[a][0].apply(g,[b].concat(g.collectionsWithInit[a][1]||[]).concat([!1])).touched=!0)});e&&g[a].forEach(function(b){b.touched||b.options.isInternal?delete b.touched:h.push(b)})}});h.forEach(function(b){b.chart&&b.remove&&b.remove(!1)});p&&g.axes.forEach(function(b){b.update({},!1)});l&&g.getSeriesOrderByLinks().forEach(function(b){b.chart&&b.update({},!1)},this);p=n&&
n.width;n=n&&(V(n.height)?aa(n.height,p||g.chartWidth):n.height);m||T(p)&&p!==g.chartWidth||T(n)&&n!==g.chartHeight?g.setSize(p,n,d):R(a,!0)&&g.redraw(d);Q(g,"afterUpdate",{options:b,redraw:a,animation:d})};a.prototype.setSubtitle=function(b,a){this.applyDescription("subtitle",b);this.layOutTitles(a)};a.prototype.setCaption=function(b,a){this.applyDescription("caption",b);this.layOutTitles(a)};a.prototype.showResetZoom=function(){function b(){a.zoomOut()}var a=this,c=e.lang,d=a.options.chart.resetZoomButton,
g=d.theme,f=g.states,k="chart"===d.relativeTo||"spacingBox"===d.relativeTo?null:"scrollablePlotBox";Q(this,"beforeShowResetZoom",null,function(){a.resetZoomButton=a.renderer.button(c.resetZoom,null,null,b,g,f&&f.hover).attr({align:d.position.align,title:c.resetZoomTitle}).addClass("highcharts-reset-zoom").add().align(d.position,!1,k)});Q(this,"afterShowResetZoom")};a.prototype.zoomOut=function(){Q(this,"selection",{resetSelection:!0},this.zoom)};a.prototype.zoom=function(b){var a=this,c=a.pointer,
e=a.inverted?c.mouseDownX:c.mouseDownY,d=!1,g;!b||b.resetSelection?(a.axes.forEach(function(b){g=b.zoom()}),c.initiated=!1):b.xAxis.concat(b.yAxis).forEach(function(b){var f=b.axis,k=a.inverted?f.left:f.top,h=a.inverted?k+f.width:k+f.height,l=f.isXAxis,n=!1;if(!l&&e>=k&&e<=h||l||!U(e))n=!0;c[l?"zoomX":"zoomY"]&&n&&(g=f.zoom(b.min,b.max),f.displayBtn&&(d=!0))});var f=a.resetZoomButton;d&&!f?a.showResetZoom():!d&&N(f)&&(a.resetZoomButton=f.destroy());g&&a.redraw(R(a.options.chart.animation,b&&b.animation,
100>a.pointCount))};a.prototype.pan=function(b,a){var c=this,e=c.hoverPoints;a="object"===typeof a?a:{enabled:a,type:"x"};var d=c.options.chart,g=c.options.mapNavigation&&c.options.mapNavigation.enabled;d&&d.panning&&(d.panning=a);var f=a.type,k;Q(this,"pan",{originalEvent:b},function(){e&&e.forEach(function(b){b.setState()});var a=c.xAxis;"xy"===f?a=a.concat(c.yAxis):"y"===f&&(a=c.yAxis);var d={};a.forEach(function(a){if(a.options.panningEnabled&&!a.options.isInternal){var e=a.horiz,h=b[e?"chartX":
"chartY"];e=e?"mouseDownX":"mouseDownY";var l=c[e],n=a.minPointOffset||0,m=a.reversed&&!c.inverted||!a.reversed&&c.inverted?-1:1,p=a.getExtremes(),F=a.toValue(l-h,!0)+n*m,r=a.toValue(l+a.len-h,!0)-(n*m||a.isXAxis&&a.pointRangePadding||0),q=r<F;m=a.hasVerticalPanning();l=q?r:F;F=q?F:r;var N=a.panningState;!m||a.isXAxis||N&&!N.isDirty||a.series.forEach(function(b){var a=b.getProcessedData(!0);a=b.getExtremes(a.yData,!0);N||(N={startMin:Number.MAX_VALUE,startMax:-Number.MAX_VALUE});T(a.dataMin)&&T(a.dataMax)&&
(N.startMin=Math.min(R(b.options.threshold,Infinity),a.dataMin,N.startMin),N.startMax=Math.max(R(b.options.threshold,-Infinity),a.dataMax,N.startMax))});m=Math.min(R(N&&N.startMin,p.dataMin),n?p.min:a.toValue(a.toPixels(p.min)-a.minPixelPadding));r=Math.max(R(N&&N.startMax,p.dataMax),n?p.max:a.toValue(a.toPixels(p.max)+a.minPixelPadding));a.panningState=N;a.isOrdinal||(n=m-l,0<n&&(F+=n,l=m),n=F-r,0<n&&(F=r,l-=n),a.series.length&&l!==p.min&&F!==p.max&&l>=m&&F<=r&&(a.setExtremes(l,F,!1,!1,{trigger:"pan"}),
c.resetZoomButton||g||l===m||F===r||!f.match("y")||(c.showResetZoom(),a.displayBtn=!1),k=!0),d[e]=h)}});X(d,function(b,a){c[a]=b});k&&c.redraw(!1);P(c.container,{cursor:"move"})})};return a}();L(a.prototype,{callbacks:[],collectionsWithInit:{xAxis:[a.prototype.addAxis,[!0]],yAxis:[a.prototype.addAxis,[!1]],series:[a.prototype.addSeries]},collectionsWithUpdate:["xAxis","yAxis","series"],propsRequireDirtyBox:"backgroundColor borderColor borderWidth borderRadius plotBackgroundColor plotBackgroundImage plotBorderColor plotBorderWidth plotShadow shadow".split(" "),
propsRequireReflow:"margin marginTop marginRight marginBottom marginLeft spacing spacingTop spacingRight spacingBottom spacingLeft".split(" "),propsRequireUpdateSeries:"chart.inverted chart.polar chart.ignoreHiddenSeries chart.type colors plotOptions time tooltip".split(" ")});"";return a});L(a,"Core/Legend/LegendSymbol.js",[a["Core/Utilities.js"]],function(a){var u=a.merge,A=a.pick,G;(function(a){a.drawLineMarker=function(a){var t=this.options,z=a.symbolWidth,q=a.symbolHeight,m=q/2,h=this.chart.renderer,
d=this.legendGroup;a=a.baseline-Math.round(.3*a.fontMetrics.b);var c={},l=t.marker;this.chart.styledMode||(c={"stroke-width":t.lineWidth||0},t.dashStyle&&(c.dashstyle=t.dashStyle));this.legendLine=h.path([["M",0,a],["L",z,a]]).addClass("highcharts-graph").attr(c).add(d);l&&!1!==l.enabled&&z&&(t=Math.min(A(l.radius,m),m),0===this.symbol.indexOf("url")&&(l=u(l,{width:q,height:q}),t=0),this.legendSymbol=z=h.symbol(this.symbol,z/2-t,a-t,2*t,2*t,l).addClass("highcharts-point").add(d),z.isMarker=!0)};a.drawRectangle=
function(a,u){var t=a.symbolHeight,q=a.options.squareSymbol;u.legendSymbol=this.chart.renderer.rect(q?(a.symbolWidth-t)/2:0,a.baseline-t+1,q?t:a.symbolWidth,t,A(a.options.symbolRadius,t/2)).addClass("highcharts-point").attr({zIndex:3}).add(u.legendGroup)}})(G||(G={}));return G});L(a,"Core/Series/SeriesDefaults.js",[],function(){return{lineWidth:2,allowPointSelect:!1,crisp:!0,showCheckbox:!1,animation:{duration:1E3},events:{},marker:{enabledThreshold:2,lineColor:"#ffffff",lineWidth:0,radius:4,states:{normal:{animation:!0},
hover:{animation:{duration:50},enabled:!0,radiusPlus:2,lineWidthPlus:1},select:{fillColor:"#cccccc",lineColor:"#000000",lineWidth:2}}},point:{events:{}},dataLabels:{animation:{},align:"center",defer:!0,formatter:function(){var a=this.series.chart.numberFormatter;return"number"!==typeof this.y?"":a(this.y,-1)},padding:5,style:{fontSize:"11px",fontWeight:"bold",color:"contrast",textOutline:"1px contrast"},verticalAlign:"bottom",x:0,y:0},cropThreshold:300,opacity:1,pointRange:0,softThreshold:!0,states:{normal:{animation:!0},
hover:{animation:{duration:50},lineWidthPlus:1,marker:{},halo:{size:10,opacity:.25}},select:{animation:{duration:0}},inactive:{animation:{duration:50},opacity:.2}},stickyTracking:!0,turboThreshold:1E3,findNearestPointBy:"x"}});L(a,"Core/Series/Series.js",[a["Core/Animation/AnimationUtilities.js"],a["Core/DefaultOptions.js"],a["Core/Foundation.js"],a["Core/Globals.js"],a["Core/Legend/LegendSymbol.js"],a["Core/Series/Point.js"],a["Core/Series/SeriesDefaults.js"],a["Core/Series/SeriesRegistry.js"],a["Core/Renderer/SVG/SVGElement.js"],
a["Core/Utilities.js"]],function(a,t,A,G,x,C,I,z,q,m){var h=a.animObject,d=a.setAnimation,c=t.defaultOptions,l=A.registerEventOptions,f=G.hasTouch,w=G.svg,p=G.win,u=z.seriesTypes,r=m.addEvent,y=m.arrayMax,B=m.arrayMin,n=m.clamp,k=m.cleanRecursively,b=m.correctFloat,g=m.defined,e=m.erase,D=m.error,H=m.extend,v=m.find,E=m.fireEvent,O=m.getNestedProperty,S=m.isArray,P=m.isNumber,U=m.isString,Y=m.merge,J=m.objectEach,M=m.pick,L=m.removeEvent,da=m.splat,Q=m.syncTimeout;a=function(){function a(){this.zones=
this.yAxis=this.xAxis=this.userOptions=this.tooltipOptions=this.processedYData=this.processedXData=this.points=this.options=this.linkedSeries=this.index=this.eventsToUnbind=this.eventOptions=this.data=this.chart=this._i=void 0}a.prototype.init=function(b,a){E(this,"init",{options:a});var c=this,e=b.series;this.eventsToUnbind=[];c.chart=b;c.options=c.setOptions(a);a=c.options;c.linkedSeries=[];c.bindAxes();H(c,{name:a.name,state:"",visible:!1!==a.visible,selected:!0===a.selected});l(this,a);var d=
a.events;if(d&&d.click||a.point&&a.point.events&&a.point.events.click||a.allowPointSelect)b.runTrackerClick=!0;c.getColor();c.getSymbol();c.parallelArrays.forEach(function(b){c[b+"Data"]||(c[b+"Data"]=[])});c.isCartesian&&(b.hasCartesianSeries=!0);var g;e.length&&(g=e[e.length-1]);c._i=M(g&&g._i,-1)+1;c.opacity=c.options.opacity;b.orderSeries(this.insert(e));a.dataSorting&&a.dataSorting.enabled?c.setDataSortingOptions():c.points||c.data||c.setData(a.data,!1);E(this,"afterInit")};a.prototype.is=function(b){return u[b]&&
this instanceof u[b]};a.prototype.insert=function(b){var a=this.options.index,c;if(P(a)){for(c=b.length;c--;)if(a>=M(b[c].options.index,b[c]._i)){b.splice(c+1,0,this);break}-1===c&&b.unshift(this);c+=1}else b.push(this);return M(c,b.length-1)};a.prototype.bindAxes=function(){var b=this,a=b.options,c=b.chart,e;E(this,"bindAxes",null,function(){(b.axisTypes||[]).forEach(function(d){var g=0;c[d].forEach(function(c){e=c.options;if(a[d]===g&&!e.isInternal||"undefined"!==typeof a[d]&&a[d]===e.id||"undefined"===
typeof a[d]&&0===e.index)b.insert(c.series),b[d]=c,c.isDirty=!0;e.isInternal||g++});b[d]||b.optionalAxis===d||D(18,!0,c)})});E(this,"afterBindAxes")};a.prototype.updateParallelArrays=function(b,a){var c=b.series,e=arguments,d=P(a)?function(e){var d="y"===e&&c.toYData?c.toYData(b):b[e];c[e+"Data"][a]=d}:function(b){Array.prototype[a].apply(c[b+"Data"],Array.prototype.slice.call(e,2))};c.parallelArrays.forEach(d)};a.prototype.hasData=function(){return this.visible&&"undefined"!==typeof this.dataMax&&
"undefined"!==typeof this.dataMin||this.visible&&this.yData&&0<this.yData.length};a.prototype.autoIncrement=function(b){var a=this.options,c=a.pointIntervalUnit,e=a.relativeXValue,d=this.chart.time,g=this.xIncrement,f;g=M(g,a.pointStart,0);this.pointInterval=f=M(this.pointInterval,a.pointInterval,1);e&&P(b)&&(f*=b);c&&(a=new d.Date(g),"day"===c?d.set("Date",a,d.get("Date",a)+f):"month"===c?d.set("Month",a,d.get("Month",a)+f):"year"===c&&d.set("FullYear",a,d.get("FullYear",a)+f),f=a.getTime()-g);if(e&&
P(b))return g+f;this.xIncrement=g+f;return g};a.prototype.setDataSortingOptions=function(){var b=this.options;H(this,{requireSorting:!1,sorted:!1,enabledDataSorting:!0,allowDG:!1});g(b.pointRange)||(b.pointRange=1)};a.prototype.setOptions=function(b){var a=this.chart,e=a.options,d=e.plotOptions,f=a.userOptions||{};b=Y(b);a=a.styledMode;var k={plotOptions:d,userOptions:b};E(this,"setOptions",k);var h=k.plotOptions[this.type],l=f.plotOptions||{};this.userOptions=k.userOptions;f=Y(h,d.series,f.plotOptions&&
f.plotOptions[this.type],b);this.tooltipOptions=Y(c.tooltip,c.plotOptions.series&&c.plotOptions.series.tooltip,c.plotOptions[this.type].tooltip,e.tooltip.userOptions,d.series&&d.series.tooltip,d[this.type].tooltip,b.tooltip);this.stickyTracking=M(b.stickyTracking,l[this.type]&&l[this.type].stickyTracking,l.series&&l.series.stickyTracking,this.tooltipOptions.shared&&!this.noSharedTooltip?!0:f.stickyTracking);null===h.marker&&delete f.marker;this.zoneAxis=f.zoneAxis;d=this.zones=(f.zones||[]).slice();
!f.negativeColor&&!f.negativeFillColor||f.zones||(e={value:f[this.zoneAxis+"Threshold"]||f.threshold||0,className:"highcharts-negative"},a||(e.color=f.negativeColor,e.fillColor=f.negativeFillColor),d.push(e));d.length&&g(d[d.length-1].value)&&d.push(a?{}:{color:this.color,fillColor:this.fillColor});E(this,"afterSetOptions",{options:f});return f};a.prototype.getName=function(){return M(this.options.name,"Series "+(this.index+1))};a.prototype.getCyclic=function(b,a,c){var e=this.chart,d=this.userOptions,
f=b+"Index",k=b+"Counter",h=c?c.length:M(e.options.chart[b+"Count"],e[b+"Count"]);if(!a){var l=M(d[f],d["_"+f]);g(l)||(e.series.length||(e[k]=0),d["_"+f]=l=e[k]%h,e[k]+=1);c&&(a=c[l])}"undefined"!==typeof l&&(this[f]=l);this[b]=a};a.prototype.getColor=function(){this.chart.styledMode?this.getCyclic("color"):this.options.colorByPoint?this.color="#cccccc":this.getCyclic("color",this.options.color||c.plotOptions[this.type].color,this.chart.options.colors)};a.prototype.getPointsCollection=function(){return(this.hasGroupedData?
this.points:this.data)||[]};a.prototype.getSymbol=function(){this.getCyclic("symbol",this.options.marker.symbol,this.chart.options.symbols)};a.prototype.findPointIndex=function(b,a){var c=b.id,e=b.x,d=this.points,g=this.options.dataSorting,f,k;if(c)g=this.chart.get(c),g instanceof C&&(f=g);else if(this.linkedParent||this.enabledDataSorting||this.options.relativeXValue)if(f=function(a){return!a.touched&&a.index===b.index},g&&g.matchByName?f=function(a){return!a.touched&&a.name===b.name}:this.options.relativeXValue&&
(f=function(a){return!a.touched&&a.options.x===b.x}),f=v(d,f),!f)return;if(f){var h=f&&f.index;"undefined"!==typeof h&&(k=!0)}"undefined"===typeof h&&P(e)&&(h=this.xData.indexOf(e,a));-1!==h&&"undefined"!==typeof h&&this.cropped&&(h=h>=this.cropStart?h-this.cropStart:h);!k&&P(h)&&d[h]&&d[h].touched&&(h=void 0);return h};a.prototype.updateData=function(b,a){var c=this.options,e=c.dataSorting,d=this.points,f=[],k=this.requireSorting,h=b.length===d.length,l,n,m,p=!0;this.xIncrement=null;b.forEach(function(b,
a){var n=g(b)&&this.pointClass.prototype.optionsToObject.call({series:this},b)||{},p=n.x;if(n.id||P(p)){if(n=this.findPointIndex(n,m),-1===n||"undefined"===typeof n?f.push(b):d[n]&&b!==c.data[n]?(d[n].update(b,!1,null,!1),d[n].touched=!0,k&&(m=n+1)):d[n]&&(d[n].touched=!0),!h||a!==n||e&&e.enabled||this.hasDerivedData)l=!0}else f.push(b)},this);if(l)for(b=d.length;b--;)(n=d[b])&&!n.touched&&n.remove&&n.remove(!1,a);else!h||e&&e.enabled?p=!1:(b.forEach(function(b,a){b!==d[a].y&&d[a].update&&d[a].update(b,
!1,null,!1)}),f.length=0);d.forEach(function(b){b&&(b.touched=!1)});if(!p)return!1;f.forEach(function(b){this.addPoint(b,!1,null,null,!1)},this);null===this.xIncrement&&this.xData&&this.xData.length&&(this.xIncrement=y(this.xData),this.autoIncrement());return!0};a.prototype.setData=function(b,a,c,e){var d=this,g=d.points,f=g&&g.length||0,k=d.options,h=d.chart,l=k.dataSorting,n=d.xAxis,m=k.turboThreshold,p=this.xData,F=this.yData,r=d.pointArrayMap;r=r&&r.length;var q=k.keys,v,B=0,w=1,u=null;b=b||[];
var y=b.length;a=M(a,!0);l&&l.enabled&&(b=this.sortData(b));!1!==e&&y&&f&&!d.cropped&&!d.hasGroupedData&&d.visible&&!d.isSeriesBoosting&&(v=this.updateData(b,c));if(!v){d.xIncrement=null;d.colorCounter=0;this.parallelArrays.forEach(function(b){d[b+"Data"].length=0});if(m&&y>m)if(u=d.getFirstValidPoint(b),P(u))for(c=0;c<y;c++)p[c]=this.autoIncrement(),F[c]=b[c];else if(S(u))if(r)if(u.length===r)for(c=0;c<y;c++)p[c]=this.autoIncrement(),F[c]=b[c];else for(c=0;c<y;c++)e=b[c],p[c]=e[0],F[c]=e.slice(1,
r+1);else if(q&&(B=q.indexOf("x"),w=q.indexOf("y"),B=0<=B?B:0,w=0<=w?w:1),1===u.length&&(w=0),B===w)for(c=0;c<y;c++)p[c]=this.autoIncrement(),F[c]=b[c][w];else for(c=0;c<y;c++)e=b[c],p[c]=e[B],F[c]=e[w];else D(12,!1,h);else for(c=0;c<y;c++)"undefined"!==typeof b[c]&&(e={series:d},d.pointClass.prototype.applyOptions.apply(e,[b[c]]),d.updateParallelArrays(e,c));F&&U(F[0])&&D(14,!0,h);d.data=[];d.options.data=d.userOptions.data=b;for(c=f;c--;)g[c]&&g[c].destroy&&g[c].destroy();n&&(n.minRange=n.userMinRange);
d.isDirty=h.isDirtyBox=!0;d.isDirtyData=!!g;c=!1}"point"===k.legendType&&(this.processData(),this.generatePoints());a&&h.redraw(c)};a.prototype.sortData=function(b){var a=this,c=a.options.dataSorting.sortKey||"y",e=function(b,a){return g(a)&&b.pointClass.prototype.optionsToObject.call({series:b},a)||{}};b.forEach(function(c,d){b[d]=e(a,c);b[d].index=d},this);b.concat().sort(function(b,a){b=O(c,b);a=O(c,a);return a<b?-1:a>b?1:0}).forEach(function(b,a){b.x=a},this);a.linkedSeries&&a.linkedSeries.forEach(function(a){var c=
a.options,d=c.data;c.dataSorting&&c.dataSorting.enabled||!d||(d.forEach(function(c,g){d[g]=e(a,c);b[g]&&(d[g].x=b[g].x,d[g].index=g)}),a.setData(d,!1))});return b};a.prototype.getProcessedData=function(b){var a=this.xAxis,c=this.options,e=c.cropThreshold,d=b||this.getExtremesFromAll||c.getExtremesFromAll,g=this.isCartesian;b=a&&a.val2lin;c=!(!a||!a.logarithmic);var f=0,k=this.xData,h=this.yData,l=this.requireSorting;var n=!1;var m=k.length;if(a){n=a.getExtremes();var p=n.min;var r=n.max;n=a.categories&&
!a.names.length}if(g&&this.sorted&&!d&&(!e||m>e||this.forceCrop))if(k[m-1]<p||k[0]>r)k=[],h=[];else if(this.yData&&(k[0]<p||k[m-1]>r)){var F=this.cropData(this.xData,this.yData,p,r);k=F.xData;h=F.yData;f=F.start;F=!0}for(e=k.length||1;--e;)if(a=c?b(k[e])-b(k[e-1]):k[e]-k[e-1],0<a&&("undefined"===typeof q||a<q))var q=a;else 0>a&&l&&!n&&(D(15,!1,this.chart),l=!1);return{xData:k,yData:h,cropped:F,cropStart:f,closestPointRange:q}};a.prototype.processData=function(b){var a=this.xAxis;if(this.isCartesian&&
!this.isDirty&&!a.isDirty&&!this.yAxis.isDirty&&!b)return!1;b=this.getProcessedData();this.cropped=b.cropped;this.cropStart=b.cropStart;this.processedXData=b.xData;this.processedYData=b.yData;this.closestPointRange=this.basePointRange=b.closestPointRange;E(this,"afterProcessData")};a.prototype.cropData=function(b,a,c,e,d){var g=b.length,f,k=0,h=g;d=M(d,this.cropShoulder);for(f=0;f<g;f++)if(b[f]>=c){k=Math.max(0,f-d);break}for(c=f;c<g;c++)if(b[c]>e){h=c+d;break}return{xData:b.slice(k,h),yData:a.slice(k,
h),start:k,end:h}};a.prototype.generatePoints=function(){var b=this.options,a=b.data,c=this.processedXData,e=this.processedYData,d=this.pointClass,g=c.length,f=this.cropStart||0,k=this.hasGroupedData,h=b.keys,l=[];b=b.dataGrouping&&b.dataGrouping.groupAll?f:0;var n,m,p=this.data;if(!p&&!k){var r=[];r.length=a.length;p=this.data=r}h&&k&&(this.options.keys=!1);for(m=0;m<g;m++){r=f+m;if(k){var q=(new d).init(this,[c[m]].concat(da(e[m])));q.dataGroup=this.groupMap[b+m];q.dataGroup.options&&(q.options=
q.dataGroup.options,H(q,q.dataGroup.options),delete q.dataLabels)}else(q=p[r])||"undefined"===typeof a[r]||(p[r]=q=(new d).init(this,a[r],c[m]));q&&(q.index=k?b+m:r,l[m]=q)}this.options.keys=h;if(p&&(g!==(n=p.length)||k))for(m=0;m<n;m++)m!==f||k||(m+=g),p[m]&&(p[m].destroyElements(),p[m].plotX=void 0);this.data=p;this.points=l;E(this,"afterGeneratePoints")};a.prototype.getXExtremes=function(b){return{min:B(b),max:y(b)}};a.prototype.getExtremes=function(b,a){var c=this.xAxis,e=this.yAxis,d=this.processedXData||
this.xData,g=[],f=this.requireSorting?this.cropShoulder:0;e=e?e.positiveValuesOnly:!1;var k,h=0,l=0,n=0;b=b||this.stackedYData||this.processedYData||[];var m=b.length;if(c){var p=c.getExtremes();h=p.min;l=p.max}for(k=0;k<m;k++){var r=d[k];p=b[k];var q=(P(p)||S(p))&&(p.length||0<p||!e);r=a||this.getExtremesFromAll||this.options.getExtremesFromAll||this.cropped||!c||(d[k+f]||r)>=h&&(d[k-f]||r)<=l;if(q&&r)if(q=p.length)for(;q--;)P(p[q])&&(g[n++]=p[q]);else g[n++]=p}b={activeYData:g,dataMin:B(g),dataMax:y(g)};
E(this,"afterGetExtremes",{dataExtremes:b});return b};a.prototype.applyExtremes=function(){var b=this.getExtremes();this.dataMin=b.dataMin;this.dataMax=b.dataMax;return b};a.prototype.getFirstValidPoint=function(b){for(var a=b.length,c=0,e=null;null===e&&c<a;)e=b[c],c++;return e};a.prototype.translate=function(){this.processedXData||this.processData();this.generatePoints();var a=this.options,c=a.stacking,e=this.xAxis,d=e.categories,f=this.enabledDataSorting,k=this.yAxis,h=this.points,l=h.length,m=
this.pointPlacementToXValue(),p=!!m,r=a.threshold,q=a.startFromThreshold?r:0,v=this.zoneAxis||"y",B,w,u=Number.MAX_VALUE;for(B=0;B<l;B++){var y=h[B],t=y.x,D=void 0,K=void 0,H=y.y,z=y.low,x=c&&k.stacking&&k.stacking.stacks[(this.negStacks&&H<(q?0:r)?"-":"")+this.stackKey];if(k.positiveValuesOnly&&!k.validatePositiveValue(H)||e.positiveValuesOnly&&!e.validatePositiveValue(t))y.isNull=!0;y.plotX=w=b(n(e.translate(t,0,0,0,1,m,"flags"===this.type),-1E5,1E5));if(c&&this.visible&&x&&x[t]){var A=this.getStackIndicator(A,
t,this.index);y.isNull||(D=x[t],K=D.points[A.key])}S(K)&&(z=K[0],H=K[1],z===q&&A.key===x[t].base&&(z=M(P(r)&&r,k.min)),k.positiveValuesOnly&&0>=z&&(z=null),y.total=y.stackTotal=D.total,y.percentage=D.total&&y.y/D.total*100,y.stackY=H,this.irregularWidths||D.setOffset(this.pointXOffset||0,this.barW||0));y.yBottom=g(z)?n(k.translate(z,0,1,0,1),-1E5,1E5):null;this.dataModify&&(H=this.dataModify.modifyValue(H,B));y.plotY=void 0;P(H)&&(D=k.translate(H,!1,!0,!1,!0),"undefined"!==typeof D&&(y.plotY=n(D,
-1E5,1E5)));y.isInside=this.isPointInside(y);y.clientX=p?b(e.translate(t,0,0,0,1,m)):w;y.negative=y[v]<(a[v+"Threshold"]||r||0);y.category=d&&"undefined"!==typeof d[y.x]?d[y.x]:y.x;if(!y.isNull&&!1!==y.visible){"undefined"!==typeof G&&(u=Math.min(u,Math.abs(w-G)));var G=w}y.zone=this.zones.length?y.getZone():void 0;!y.graphic&&this.group&&f&&(y.isNew=!0)}this.closestPointRangePx=u;E(this,"afterTranslate")};a.prototype.getValidPoints=function(b,a,c){var e=this.chart;return(b||this.points||[]).filter(function(b){return a&&
!e.isInsidePlot(b.plotX,b.plotY,{inverted:e.inverted})?!1:!1!==b.visible&&(c||!b.isNull)})};a.prototype.getClipBox=function(){var b=this.chart,a=this.xAxis,c=this.yAxis,e=Y(b.clipBox);a&&a.len!==b.plotSizeX&&(e.width=a.len);c&&c.len!==b.plotSizeY&&(e.height=c.len);return e};a.prototype.getSharedClipKey=function(){return this.sharedClipKey=(this.options.xAxis||0)+","+(this.options.yAxis||0)};a.prototype.setClip=function(){var b=this.chart,a=this.group,c=this.markerGroup,e=b.sharedClips;b=b.renderer;
var d=this.getClipBox(),g=this.getSharedClipKey(),f=e[g];f?f.animate(d):e[g]=f=b.clipRect(d);a&&a.clip(!1===this.options.clip?void 0:f);c&&c.clip()};a.prototype.animate=function(b){var a=this.chart,c=this.group,e=this.markerGroup,d=a.inverted,g=h(this.options.animation),f=[this.getSharedClipKey(),g.duration,g.easing,g.defer].join(),k=a.sharedClips[f],l=a.sharedClips[f+"m"];if(b&&c)g=this.getClipBox(),k?k.attr("height",g.height):(g.width=0,d&&(g.x=a.plotHeight),k=a.renderer.clipRect(g),a.sharedClips[f]=
k,l=a.renderer.clipRect({x:d?(a.plotSizeX||0)+99:-99,y:d?-a.plotLeft:-a.plotTop,width:99,height:d?a.chartWidth:a.chartHeight}),a.sharedClips[f+"m"]=l),c.clip(k),e&&e.clip(l);else if(k&&!k.hasClass("highcharts-animating")){a=this.getClipBox();var n=g.step;e&&e.element.childNodes.length&&(g.step=function(b,a){n&&n.apply(a,arguments);l&&l.element&&l.attr(a.prop,"width"===a.prop?b+99:b)});k.addClass("highcharts-animating").animate(a,g)}};a.prototype.afterAnimate=function(){var b=this;this.setClip();J(this.chart.sharedClips,
function(a,c,e){a&&!b.chart.container.querySelector('[clip-path="url(#'+a.id+')"]')&&(a.destroy(),delete e[c])});this.finishedAnimating=!0;E(this,"afterAnimate")};a.prototype.drawPoints=function(){var b=this.points,a=this.chart,c=this.options.marker,e=this[this.specialGroup]||this.markerGroup,d=this.xAxis,g=M(c.enabled,!d||d.isRadial?!0:null,this.closestPointRangePx>=c.enabledThreshold*c.radius),f,k;if(!1!==c.enabled||this._hasPointMarkers)for(f=0;f<b.length;f++){var h=b[f];var l=(k=h.graphic)?"animate":
"attr";var n=h.marker||{};var m=!!h.marker;if((g&&"undefined"===typeof n.enabled||n.enabled)&&!h.isNull&&!1!==h.visible){var p=M(n.symbol,this.symbol,"rect");var r=this.markerAttribs(h,h.selected&&"select");this.enabledDataSorting&&(h.startXPos=d.reversed?-(r.width||0):d.width);var q=!1!==h.isInside;k?k[q?"show":"hide"](q).animate(r):q&&(0<(r.width||0)||h.hasImage)&&(h.graphic=k=a.renderer.symbol(p,r.x,r.y,r.width,r.height,m?n:c).add(e),this.enabledDataSorting&&a.hasRendered&&(k.attr({x:h.startXPos}),
l="animate"));k&&"animate"===l&&k[q?"show":"hide"](q).animate(r);if(k&&!a.styledMode)k[l](this.pointAttribs(h,h.selected&&"select"));k&&k.addClass(h.getClassName(),!0)}else k&&(h.graphic=k.destroy())}};a.prototype.markerAttribs=function(b,a){var c=this.options,e=c.marker,d=b.marker||{},g=d.symbol||e.symbol,f=M(d.radius,e.radius);a&&(e=e.states[a],a=d.states&&d.states[a],f=M(a&&a.radius,e&&e.radius,f+(e&&e.radiusPlus||0)));b.hasImage=g&&0===g.indexOf("url");b.hasImage&&(f=0);b={x:c.crisp?Math.floor(b.plotX-
f):b.plotX-f,y:b.plotY-f};f&&(b.width=b.height=2*f);return b};a.prototype.pointAttribs=function(b,a){var c=this.options.marker,e=b&&b.options,d=e&&e.marker||{},g=e&&e.color,f=b&&b.color,k=b&&b.zone&&b.zone.color,h=this.color;b=M(d.lineWidth,c.lineWidth);e=1;h=g||k||f||h;g=d.fillColor||c.fillColor||h;f=d.lineColor||c.lineColor||h;a=a||"normal";c=c.states[a]||{};a=d.states&&d.states[a]||{};b=M(a.lineWidth,c.lineWidth,b+M(a.lineWidthPlus,c.lineWidthPlus,0));g=a.fillColor||c.fillColor||g;f=a.lineColor||
c.lineColor||f;e=M(a.opacity,c.opacity,e);return{stroke:f,"stroke-width":b,fill:g,opacity:e}};a.prototype.destroy=function(b){var a=this,c=a.chart,d=/AppleWebKit\/533/.test(p.navigator.userAgent),g=a.data||[],f,k,h,l;E(a,"destroy");this.removeEvents(b);(a.axisTypes||[]).forEach(function(b){(l=a[b])&&l.series&&(e(l.series,a),l.isDirty=l.forceRedraw=!0)});a.legendItem&&a.chart.legend.destroyItem(a);for(k=g.length;k--;)(h=g[k])&&h.destroy&&h.destroy();a.clips&&a.clips.forEach(function(b){return b.destroy()});
m.clearTimeout(a.animationTimeout);J(a,function(b,a){b instanceof q&&!b.survive&&(f=d&&"group"===a?"hide":"destroy",b[f]())});c.hoverSeries===a&&(c.hoverSeries=void 0);e(c.series,a);c.orderSeries();J(a,function(c,e){b&&"hcEvents"===e||delete a[e]})};a.prototype.applyZones=function(){var b=this,a=this.chart,c=a.renderer,e=this.zones,d=this.clips||[],g=this.graph,f=this.area,k=Math.max(a.chartWidth,a.chartHeight),h=this[(this.zoneAxis||"y")+"Axis"],l=a.inverted,m,p,r,q,v,y,B,w,u=!1;if(e.length&&(g||
f)&&h&&"undefined"!==typeof h.min){var t=h.reversed;var D=h.horiz;g&&!this.showLine&&g.hide();f&&f.hide();var E=h.getExtremes();e.forEach(function(e,F){m=t?D?a.plotWidth:0:D?0:h.toPixels(E.min)||0;m=n(M(p,m),0,k);p=n(Math.round(h.toPixels(M(e.value,E.max),!0)||0),0,k);u&&(m=p=h.toPixels(E.max));q=Math.abs(m-p);v=Math.min(m,p);y=Math.max(m,p);h.isXAxis?(r={x:l?y:v,y:0,width:q,height:k},D||(r.x=a.plotHeight-r.x)):(r={x:0,y:l?y:v,width:k,height:q},D&&(r.y=a.plotWidth-r.y));l&&c.isVML&&(r=h.isXAxis?{x:0,
y:t?v:y,height:r.width,width:a.chartWidth}:{x:r.y-a.plotLeft-a.spacingBox.x,y:0,width:r.height,height:a.chartHeight});d[F]?d[F].animate(r):d[F]=c.clipRect(r);B=b["zone-area-"+F];w=b["zone-graph-"+F];g&&w&&w.clip(d[F]);f&&B&&B.clip(d[F]);u=e.value>E.max;b.resetZones&&0===p&&(p=void 0)});this.clips=d}else b.visible&&(g&&g.show(!0),f&&f.show(!0))};a.prototype.invertGroups=function(b){function a(){["group","markerGroup"].forEach(function(a){c[a]&&(e.renderer.isVML&&c[a].attr({width:c.yAxis.len,height:c.xAxis.len}),
c[a].width=c.yAxis.len,c[a].height=c.xAxis.len,c[a].invert(c.isRadialSeries?!1:b))})}var c=this,e=c.chart;c.xAxis&&(c.eventsToUnbind.push(r(e,"resize",a)),a(),c.invertGroups=a)};a.prototype.plotGroup=function(b,a,c,e,d){var f=this[b],k=!f;c={visibility:c,zIndex:e||.1};"undefined"===typeof this.opacity||this.chart.styledMode||"inactive"===this.state||(c.opacity=this.opacity);k&&(this[b]=f=this.chart.renderer.g().add(d));f.addClass("highcharts-"+a+" highcharts-series-"+this.index+" highcharts-"+this.type+
"-series "+(g(this.colorIndex)?"highcharts-color-"+this.colorIndex+" ":"")+(this.options.className||"")+(f.hasClass("highcharts-tracker")?" highcharts-tracker":""),!0);f.attr(c)[k?"attr":"animate"](this.getPlotBox());return f};a.prototype.getPlotBox=function(){var b=this.chart,a=this.xAxis,c=this.yAxis;b.inverted&&(a=c,c=this.xAxis);return{translateX:a?a.left:b.plotLeft,translateY:c?c.top:b.plotTop,scaleX:1,scaleY:1}};a.prototype.removeEvents=function(b){b||L(this);this.eventsToUnbind.length&&(this.eventsToUnbind.forEach(function(b){b()}),
this.eventsToUnbind.length=0)};a.prototype.render=function(){var b=this,a=b.chart,c=b.options,e=h(c.animation),d=b.visible?"inherit":"hidden",g=c.zIndex,f=b.hasRendered,k=a.seriesGroup,l=a.inverted;a=!b.finishedAnimating&&a.renderer.isSVG?e.duration:0;E(this,"render");var n=b.plotGroup("group","series",d,g,k);b.markerGroup=b.plotGroup("markerGroup","markers",d,g,k);!1!==c.clip&&b.setClip();b.animate&&a&&b.animate(!0);n.inverted=M(b.invertible,b.isCartesian)?l:!1;b.drawGraph&&(b.drawGraph(),b.applyZones());
b.visible&&b.drawPoints();b.drawDataLabels&&b.drawDataLabels();b.redrawPoints&&b.redrawPoints();b.drawTracker&&!1!==b.options.enableMouseTracking&&b.drawTracker();b.invertGroups(l);b.animate&&a&&b.animate();f||(a&&e.defer&&(a+=e.defer),b.animationTimeout=Q(function(){b.afterAnimate()},a||0));b.isDirty=!1;b.hasRendered=!0;E(b,"afterRender")};a.prototype.redraw=function(){var b=this.chart,a=this.isDirty||this.isDirtyData,c=this.group,e=this.xAxis,d=this.yAxis;c&&(b.inverted&&c.attr({width:b.plotWidth,
height:b.plotHeight}),c.animate({translateX:M(e&&e.left,b.plotLeft),translateY:M(d&&d.top,b.plotTop)}));this.translate();this.render();a&&delete this.kdTree};a.prototype.searchPoint=function(b,a){var c=this.xAxis,e=this.yAxis,d=this.chart.inverted;return this.searchKDTree({clientX:d?c.len-b.chartY+c.pos:b.chartX-c.pos,plotY:d?e.len-b.chartX+e.pos:b.chartY-e.pos},a,b)};a.prototype.buildKDTree=function(b){function a(b,e,d){var g=b&&b.length;if(g){var f=c.kdAxisArray[e%d];b.sort(function(b,a){return b[f]-
a[f]});g=Math.floor(g/2);return{point:b[g],left:a(b.slice(0,g),e+1,d),right:a(b.slice(g+1),e+1,d)}}}this.buildingKdTree=!0;var c=this,e=-1<c.options.findNearestPointBy.indexOf("y")?2:1;delete c.kdTree;Q(function(){c.kdTree=a(c.getValidPoints(null,!c.directTouch),e,e);c.buildingKdTree=!1},c.options.kdNow||b&&"touchstart"===b.type?0:1)};a.prototype.searchKDTree=function(b,a,c){function e(b,a,c,l){var n=a.point,m=d.kdAxisArray[c%l],p=n,r=g(b[f])&&g(n[f])?Math.pow(b[f]-n[f],2):null;var q=g(b[k])&&g(n[k])?
Math.pow(b[k]-n[k],2):null;q=(r||0)+(q||0);n.dist=g(q)?Math.sqrt(q):Number.MAX_VALUE;n.distX=g(r)?Math.sqrt(r):Number.MAX_VALUE;m=b[m]-n[m];q=0>m?"left":"right";r=0>m?"right":"left";a[q]&&(q=e(b,a[q],c+1,l),p=q[h]<p[h]?q:n);a[r]&&Math.sqrt(m*m)<p[h]&&(b=e(b,a[r],c+1,l),p=b[h]<p[h]?b:p);return p}var d=this,f=this.kdAxisArray[0],k=this.kdAxisArray[1],h=a?"distX":"dist";a=-1<d.options.findNearestPointBy.indexOf("y")?2:1;this.kdTree||this.buildingKdTree||this.buildKDTree(c);if(this.kdTree)return e(b,
this.kdTree,a,a)};a.prototype.pointPlacementToXValue=function(){var b=this.options,a=b.pointRange,c=this.xAxis;b=b.pointPlacement;"between"===b&&(b=c.reversed?-.5:.5);return P(b)?b*(a||c.pointRange):0};a.prototype.isPointInside=function(b){var a=this.chart,c=this.xAxis,e=this.yAxis;return"undefined"!==typeof b.plotY&&"undefined"!==typeof b.plotX&&0<=b.plotY&&b.plotY<=(e?e.len:a.plotHeight)&&0<=b.plotX&&b.plotX<=(c?c.len:a.plotWidth)};a.prototype.drawTracker=function(){var b=this,a=b.options,c=a.trackByArea,
e=[].concat(c?b.areaPath:b.graphPath),d=b.chart,g=d.pointer,k=d.renderer,h=d.options.tooltip.snap,l=b.tracker,n=function(a){if(d.hoverSeries!==b)b.onMouseOver()},m="rgba(192,192,192,"+(w?.0001:.002)+")";l?l.attr({d:e}):b.graph&&(b.tracker=k.path(e).attr({visibility:b.visible?"visible":"hidden",zIndex:2}).addClass(c?"highcharts-tracker-area":"highcharts-tracker-line").add(b.group),d.styledMode||b.tracker.attr({"stroke-linecap":"round","stroke-linejoin":"round",stroke:m,fill:c?m:"none","stroke-width":b.graph.strokeWidth()+
(c?0:2*h)}),[b.tracker,b.markerGroup,b.dataLabelsGroup].forEach(function(b){if(b&&(b.addClass("highcharts-tracker").on("mouseover",n).on("mouseout",function(b){g.onTrackerMouseOut(b)}),a.cursor&&!d.styledMode&&b.css({cursor:a.cursor}),f))b.on("touchstart",n)}));E(this,"afterDrawTracker")};a.prototype.addPoint=function(b,a,c,e,d){var g=this.options,f=this.data,k=this.chart,h=this.xAxis;h=h&&h.hasNames&&h.names;var l=g.data,n=this.xData,m;a=M(a,!0);var p={series:this};this.pointClass.prototype.applyOptions.apply(p,
[b]);var r=p.x;var q=n.length;if(this.requireSorting&&r<n[q-1])for(m=!0;q&&n[q-1]>r;)q--;this.updateParallelArrays(p,"splice",q,0,0);this.updateParallelArrays(p,q);h&&p.name&&(h[r]=p.name);l.splice(q,0,b);m&&(this.data.splice(q,0,null),this.processData());"point"===g.legendType&&this.generatePoints();c&&(f[0]&&f[0].remove?f[0].remove(!1):(f.shift(),this.updateParallelArrays(p,"shift"),l.shift()));!1!==d&&E(this,"addPoint",{point:p});this.isDirtyData=this.isDirty=!0;a&&k.redraw(e)};a.prototype.removePoint=
function(b,a,c){var e=this,g=e.data,f=g[b],k=e.points,h=e.chart,l=function(){k&&k.length===g.length&&k.splice(b,1);g.splice(b,1);e.options.data.splice(b,1);e.updateParallelArrays(f||{series:e},"splice",b,1);f&&f.destroy();e.isDirty=!0;e.isDirtyData=!0;a&&h.redraw()};d(c,h);a=M(a,!0);f?f.firePointEvent("remove",null,l):l()};a.prototype.remove=function(b,a,c,e){function d(){g.destroy(e);f.isDirtyLegend=f.isDirtyBox=!0;f.linkSeries();M(b,!0)&&f.redraw(a)}var g=this,f=g.chart;!1!==c?E(g,"remove",null,
d):d()};a.prototype.update=function(b,a){b=k(b,this.userOptions);E(this,"update",{options:b});var c=this,e=c.chart,d=c.userOptions,g=c.initialType||c.type,f=e.options.plotOptions,h=u[g].prototype,l=c.finishedAnimating&&{animation:!1},n={},m,p=["eventOptions","navigatorSeries","baseSeries"],r=b.type||d.type||e.options.chart.type,q=!(this.hasDerivedData||r&&r!==this.type||"undefined"!==typeof b.pointStart||"undefined"!==typeof b.pointInterval||"undefined"!==typeof b.relativeXValue||c.hasOptionChanged("dataGrouping")||
c.hasOptionChanged("pointStart")||c.hasOptionChanged("pointInterval")||c.hasOptionChanged("pointIntervalUnit")||c.hasOptionChanged("keys"));r=r||g;q&&(p.push("data","isDirtyData","points","processedXData","processedYData","xIncrement","cropped","_hasPointMarkers","_hasPointLabels","clips","nodes","layout","mapMap","mapData","minY","maxY","minX","maxX"),!1!==b.visible&&p.push("area","graph"),c.parallelArrays.forEach(function(b){p.push(b+"Data")}),b.data&&(b.dataSorting&&H(c.options.dataSorting,b.dataSorting),
this.setData(b.data,!1)));b=Y(d,l,{index:"undefined"===typeof d.index?c.index:d.index,pointStart:M(f&&f.series&&f.series.pointStart,d.pointStart,c.xData[0])},!q&&{data:c.options.data},b);q&&b.data&&(b.data=c.options.data);p=["group","markerGroup","dataLabelsGroup","transformGroup"].concat(p);p.forEach(function(b){p[b]=c[b];delete c[b]});f=!1;if(u[r]){if(f=r!==c.type,c.remove(!1,!1,!1,!0),f)if(Object.setPrototypeOf)Object.setPrototypeOf(c,u[r].prototype);else{l=Object.hasOwnProperty.call(c,"hcEvents")&&
c.hcEvents;for(m in h)c[m]=void 0;H(c,u[r].prototype);l?c.hcEvents=l:delete c.hcEvents}}else D(17,!0,e,{missingModuleFor:r});p.forEach(function(b){c[b]=p[b]});c.init(e,b);if(q&&this.points){var v=c.options;!1===v.visible?(n.graphic=1,n.dataLabel=1):c._hasPointLabels||(b=v.marker,h=v.dataLabels,!b||!1!==b.enabled&&(d.marker&&d.marker.symbol)===b.symbol||(n.graphic=1),h&&!1===h.enabled&&(n.dataLabel=1));this.points.forEach(function(b){b&&b.series&&(b.resolveColor(),Object.keys(n).length&&b.destroyElements(n),
!1===v.showInLegend&&b.legendItem&&e.legend.destroyItem(b))},this)}c.initialType=g;e.linkSeries();f&&c.linkedSeries.length&&(c.isDirtyData=!0);E(this,"afterUpdate");M(a,!0)&&e.redraw(q?void 0:!1)};a.prototype.setName=function(b){this.name=this.options.name=this.userOptions.name=b;this.chart.isDirtyLegend=!0};a.prototype.hasOptionChanged=function(b){var a=this.options[b],c=this.chart.options.plotOptions,e=this.userOptions[b];return e?a!==e:a!==M(c&&c[this.type]&&c[this.type][b],c&&c.series&&c.series[b],
a)};a.prototype.onMouseOver=function(){var b=this.chart,a=b.hoverSeries;b.pointer.setHoverChartIndex();if(a&&a!==this)a.onMouseOut();this.options.events.mouseOver&&E(this,"mouseOver");this.setState("hover");b.hoverSeries=this};a.prototype.onMouseOut=function(){var b=this.options,a=this.chart,c=a.tooltip,e=a.hoverPoint;a.hoverSeries=null;if(e)e.onMouseOut();this&&b.events.mouseOut&&E(this,"mouseOut");!c||this.stickyTracking||c.shared&&!this.noSharedTooltip||c.hide();a.series.forEach(function(b){b.setState("",
!0)})};a.prototype.setState=function(b,a){var c=this,e=c.options,d=c.graph,g=e.inactiveOtherPoints,f=e.states,k=M(f[b||"normal"]&&f[b||"normal"].animation,c.chart.options.chart.animation),h=e.lineWidth,l=0,n=e.opacity;b=b||"";if(c.state!==b&&([c.group,c.markerGroup,c.dataLabelsGroup].forEach(function(a){a&&(c.state&&a.removeClass("highcharts-series-"+c.state),b&&a.addClass("highcharts-series-"+b))}),c.state=b,!c.chart.styledMode)){if(f[b]&&!1===f[b].enabled)return;b&&(h=f[b].lineWidth||h+(f[b].lineWidthPlus||
0),n=M(f[b].opacity,n));if(d&&!d.dashstyle)for(e={"stroke-width":h},d.animate(e,k);c["zone-graph-"+l];)c["zone-graph-"+l].animate(e,k),l+=1;g||[c.group,c.markerGroup,c.dataLabelsGroup,c.labelBySeries].forEach(function(b){b&&b.animate({opacity:n},k)})}a&&g&&c.points&&c.setAllPointsToState(b||void 0)};a.prototype.setAllPointsToState=function(b){this.points.forEach(function(a){a.setState&&a.setState(b)})};a.prototype.setVisible=function(b,a){var c=this,e=c.chart,d=c.legendItem,g=e.options.chart.ignoreHiddenSeries,
f=c.visible,k=(c.visible=b=c.options.visible=c.userOptions.visible="undefined"===typeof b?!f:b)?"show":"hide";["group","dataLabelsGroup","markerGroup","tracker","tt"].forEach(function(b){if(c[b])c[b][k]()});if(e.hoverSeries===c||(e.hoverPoint&&e.hoverPoint.series)===c)c.onMouseOut();d&&e.legend.colorizeItem(c,b);c.isDirty=!0;c.options.stacking&&e.series.forEach(function(b){b.options.stacking&&b.visible&&(b.isDirty=!0)});c.linkedSeries.forEach(function(a){a.setVisible(b,!1)});g&&(e.isDirtyBox=!0);
E(c,k);!1!==a&&e.redraw()};a.prototype.show=function(){this.setVisible(!0)};a.prototype.hide=function(){this.setVisible(!1)};a.prototype.select=function(b){this.selected=b=this.options.selected="undefined"===typeof b?!this.selected:b;this.checkbox&&(this.checkbox.checked=b);E(this,b?"select":"unselect")};a.prototype.shouldShowTooltip=function(b,a,c){void 0===c&&(c={});c.series=this;c.visiblePlotOnly=!0;return this.chart.isInsidePlot(b,a,c)};a.defaultOptions=I;return a}();H(a.prototype,{axisTypes:["xAxis",
"yAxis"],coll:"series",colorCounter:0,cropShoulder:1,directTouch:!1,drawLegendSymbol:x.drawLineMarker,isCartesian:!0,kdAxisArray:["clientX","plotY"],parallelArrays:["x","y"],pointClass:C,requireSorting:!0,sorted:!0});z.series=a;"";"";return a});L(a,"Extensions/ScrollablePlotArea.js",[a["Core/Animation/AnimationUtilities.js"],a["Core/Axis/Axis.js"],a["Core/Chart/Chart.js"],a["Core/Series/Series.js"],a["Core/Renderer/RendererRegistry.js"],a["Core/Utilities.js"]],function(a,t,A,G,x,C){var u=a.stop,z=
C.addEvent,q=C.createElement,m=C.merge,h=C.pick;z(A,"afterSetChartSize",function(a){var c=this.options.chart.scrollablePlotArea,d=c&&c.minWidth;c=c&&c.minHeight;if(!this.renderer.forExport){if(d){if(this.scrollablePixelsX=d=Math.max(0,d-this.chartWidth)){this.scrollablePlotBox=this.renderer.scrollablePlotBox=m(this.plotBox);this.plotBox.width=this.plotWidth+=d;this.inverted?this.clipBox.height+=d:this.clipBox.width+=d;var f={1:{name:"right",value:d}}}}else c&&(this.scrollablePixelsY=d=Math.max(0,
c-this.chartHeight))&&(this.scrollablePlotBox=this.renderer.scrollablePlotBox=m(this.plotBox),this.plotBox.height=this.plotHeight+=d,this.inverted?this.clipBox.width+=d:this.clipBox.height+=d,f={2:{name:"bottom",value:d}});f&&!a.skipAxes&&this.axes.forEach(function(a){f[a.side]?a.getPlotLinePath=function(){var c=f[a.side].name,d=this[c];this[c]=d-f[a.side].value;var h=t.prototype.getPlotLinePath.apply(this,arguments);this[c]=d;return h}:(a.setAxisSize(),a.setAxisTranslation())})}});z(A,"render",function(){this.scrollablePixelsX||
this.scrollablePixelsY?(this.setUpScrolling&&this.setUpScrolling(),this.applyFixed()):this.fixedDiv&&this.applyFixed()});A.prototype.setUpScrolling=function(){var a=this,c={WebkitOverflowScrolling:"touch",overflowX:"hidden",overflowY:"hidden"};this.scrollablePixelsX&&(c.overflowX="auto");this.scrollablePixelsY&&(c.overflowY="auto");this.scrollingParent=q("div",{className:"highcharts-scrolling-parent"},{position:"relative"},this.renderTo);this.scrollingContainer=q("div",{className:"highcharts-scrolling"},
c,this.scrollingParent);z(this.scrollingContainer,"scroll",function(){a.pointer&&delete a.pointer.chartPosition});this.innerContainer=q("div",{className:"highcharts-inner-container"},null,this.scrollingContainer);this.innerContainer.appendChild(this.container);this.setUpScrolling=null};A.prototype.moveFixedElements=function(){var a=this.container,c=this.fixedRenderer,h=".highcharts-contextbutton .highcharts-credits .highcharts-legend .highcharts-legend-checkbox .highcharts-navigator-series .highcharts-navigator-xaxis .highcharts-navigator-yaxis .highcharts-navigator .highcharts-reset-zoom .highcharts-drillup-button .highcharts-scrollbar .highcharts-subtitle .highcharts-title".split(" "),
f;this.scrollablePixelsX&&!this.inverted?f=".highcharts-yaxis":this.scrollablePixelsX&&this.inverted?f=".highcharts-xaxis":this.scrollablePixelsY&&!this.inverted?f=".highcharts-xaxis":this.scrollablePixelsY&&this.inverted&&(f=".highcharts-yaxis");f&&h.push(f+":not(.highcharts-radial-axis)",f+"-labels:not(.highcharts-radial-axis-labels)");h.forEach(function(d){[].forEach.call(a.querySelectorAll(d),function(a){(a.namespaceURI===c.SVG_NS?c.box:c.box.parentNode).appendChild(a);a.style.pointerEvents="auto"})})};
A.prototype.applyFixed=function(){var a=!this.fixedDiv,c=this.options.chart,l=c.scrollablePlotArea,f=x.getRendererType();a?(this.fixedDiv=q("div",{className:"highcharts-fixed"},{position:"absolute",overflow:"hidden",pointerEvents:"none",zIndex:(c.style&&c.style.zIndex||0)+2,top:0},null,!0),this.scrollingContainer&&this.scrollingContainer.parentNode.insertBefore(this.fixedDiv,this.scrollingContainer),this.renderTo.style.overflow="visible",this.fixedRenderer=c=new f(this.fixedDiv,this.chartWidth,this.chartHeight,
this.options.chart.style),this.scrollableMask=c.path().attr({fill:this.options.chart.backgroundColor||"#fff","fill-opacity":h(l.opacity,.85),zIndex:-1}).addClass("highcharts-scrollable-mask").add(),z(this,"afterShowResetZoom",this.moveFixedElements),z(this,"afterDrilldown",this.moveFixedElements),z(this,"afterLayOutTitles",this.moveFixedElements)):this.fixedRenderer.setSize(this.chartWidth,this.chartHeight);if(this.scrollableDirty||a)this.scrollableDirty=!1,this.moveFixedElements();c=this.chartWidth+
(this.scrollablePixelsX||0);f=this.chartHeight+(this.scrollablePixelsY||0);u(this.container);this.container.style.width=c+"px";this.container.style.height=f+"px";this.renderer.boxWrapper.attr({width:c,height:f,viewBox:[0,0,c,f].join(" ")});this.chartBackground.attr({width:c,height:f});this.scrollingContainer.style.height=this.chartHeight+"px";a&&(l.scrollPositionX&&(this.scrollingContainer.scrollLeft=this.scrollablePixelsX*l.scrollPositionX),l.scrollPositionY&&(this.scrollingContainer.scrollTop=this.scrollablePixelsY*
l.scrollPositionY));f=this.axisOffset;a=this.plotTop-f[0]-1;l=this.plotLeft-f[3]-1;c=this.plotTop+this.plotHeight+f[2]+1;f=this.plotLeft+this.plotWidth+f[1]+1;var m=this.plotLeft+this.plotWidth-(this.scrollablePixelsX||0),p=this.plotTop+this.plotHeight-(this.scrollablePixelsY||0);a=this.scrollablePixelsX?[["M",0,a],["L",this.plotLeft-1,a],["L",this.plotLeft-1,c],["L",0,c],["Z"],["M",m,a],["L",this.chartWidth,a],["L",this.chartWidth,c],["L",m,c],["Z"]]:this.scrollablePixelsY?[["M",l,0],["L",l,this.plotTop-
1],["L",f,this.plotTop-1],["L",f,0],["Z"],["M",l,p],["L",l,this.chartHeight],["L",f,this.chartHeight],["L",f,p],["Z"]]:[["M",0,0]];"adjustHeight"!==this.redrawTrigger&&this.scrollableMask.attr({d:a})};z(t,"afterInit",function(){this.chart.scrollableDirty=!0});z(G,"show",function(){this.chart.scrollableDirty=!0});""});L(a,"Core/Axis/StackingAxis.js",[a["Core/Animation/AnimationUtilities.js"],a["Core/Axis/Axis.js"],a["Core/Utilities.js"]],function(a,t,A){var u=a.getDeferredAnimation,x=A.addEvent,C=
A.destroyObjectProperties,I=A.fireEvent,z=A.isNumber,q=A.objectEach,m;(function(a){function d(){var a=this.stacking;if(a){var c=a.stacks;q(c,function(a,d){C(a);c[d]=null});a&&a.stackTotalGroup&&a.stackTotalGroup.destroy()}}function c(){this.stacking||(this.stacking=new f(this))}var h=[];a.compose=function(a){-1===h.indexOf(a)&&(h.push(a),x(a,"init",c),x(a,"destroy",d));return a};var f=function(){function a(a){this.oldStacks={};this.stacks={};this.stacksTouched=0;this.axis=a}a.prototype.buildStacks=
function(){var a=this.axis,c=a.series,d=a.options.reversedStacks,f=c.length,h;if(!a.isXAxis){this.usePercentage=!1;for(h=f;h--;){var l=c[d?h:f-h-1];l.setStackedPoints();l.setGroupedPoints()}for(h=0;h<f;h++)c[h].modifyStacks();I(a,"afterBuildStacks")}};a.prototype.cleanStacks=function(){if(!this.axis.isXAxis){if(this.oldStacks)var a=this.stacks=this.oldStacks;q(a,function(a){q(a,function(a){a.cumulative=a.total})})}};a.prototype.resetStacks=function(){var a=this,c=a.stacks;a.axis.isXAxis||q(c,function(c){q(c,
function(d,f){z(d.touched)&&d.touched<a.stacksTouched?(d.destroy(),delete c[f]):(d.total=null,d.cumulative=null)})})};a.prototype.renderStackTotals=function(){var a=this.axis,c=a.chart,d=c.renderer,f=this.stacks;a=u(c,a.options.stackLabels&&a.options.stackLabels.animation||!1);var h=this.stackTotalGroup=this.stackTotalGroup||d.g("stack-labels").attr({visibility:"visible",zIndex:6,opacity:0}).add();h.translate(c.plotLeft,c.plotTop);q(f,function(a){q(a,function(a){a.render(h)})});h.animate({opacity:1},
a)};return a}();a.Additions=f})(m||(m={}));return m});L(a,"Extensions/Stacking.js",[a["Core/Axis/Axis.js"],a["Core/Chart/Chart.js"],a["Core/FormatUtilities.js"],a["Core/Globals.js"],a["Core/Series/Series.js"],a["Core/Axis/StackingAxis.js"],a["Core/Utilities.js"]],function(a,t,A,G,x,C,I){var u=A.format,q=I.correctFloat,m=I.defined,h=I.destroyObjectProperties,d=I.isArray,c=I.isNumber,l=I.objectEach,f=I.pick,w=function(){function a(a,c,d,f,h){var k=a.chart.inverted;this.axis=a;this.isNegative=d;this.options=
c=c||{};this.x=f;this.total=null;this.points={};this.hasValidPoints=!1;this.stack=h;this.rightCliff=this.leftCliff=0;this.alignOptions={align:c.align||(k?d?"left":"right":"center"),verticalAlign:c.verticalAlign||(k?"middle":d?"bottom":"top"),y:c.y,x:c.x};this.textAlign=c.textAlign||(k?d?"right":"left":"center")}a.prototype.destroy=function(){h(this,this.axis)};a.prototype.render=function(a){var c=this.axis.chart,d=this.options,h=d.format;h=h?u(h,this,c):d.formatter.call(this);this.label?this.label.attr({text:h,
visibility:"hidden"}):(this.label=c.renderer.label(h,null,null,d.shape,null,null,d.useHTML,!1,"stack-labels"),h={r:d.borderRadius||0,text:h,rotation:d.rotation,padding:f(d.padding,5),visibility:"hidden"},c.styledMode||(h.fill=d.backgroundColor,h.stroke=d.borderColor,h["stroke-width"]=d.borderWidth,this.label.css(d.style)),this.label.attr(h),this.label.added||this.label.add(a));this.label.labelrank=c.plotSizeY};a.prototype.setOffset=function(a,d,h,l,n){var k=this.axis,b=k.chart;l=k.translate(k.stacking.usePercentage?
100:l?l:this.total,0,0,0,1);h=k.translate(h?h:0);h=m(l)&&Math.abs(l-h);a=f(n,b.xAxis[0].translate(this.x))+a;k=m(l)&&this.getStackBox(b,this,a,l,d,h,k);d=this.label;h=this.isNegative;a="justify"===f(this.options.overflow,"justify");var g=this.textAlign;d&&k&&(n=d.getBBox(),l=d.padding,g="left"===g?b.inverted?-l:l:"right"===g?n.width:b.inverted&&"center"===g?n.width/2:b.inverted?h?n.width+l:-l:n.width/2,h=b.inverted?n.height/2:h?-l:n.height,this.alignOptions.x=f(this.options.x,0),this.alignOptions.y=
f(this.options.y,0),k.x-=g,k.y-=h,d.align(this.alignOptions,null,k),b.isInsidePlot(d.alignAttr.x+g-this.alignOptions.x,d.alignAttr.y+h-this.alignOptions.y)?d.show():(d.alignAttr.y=-9999,a=!1),a&&x.prototype.justifyDataLabel.call(this.axis,d,this.alignOptions,d.alignAttr,n,k),d.attr({x:d.alignAttr.x,y:d.alignAttr.y}),f(!a&&this.options.crop,!0)&&((b=c(d.x)&&c(d.y)&&b.isInsidePlot(d.x-l+d.width,d.y)&&b.isInsidePlot(d.x+l,d.y))||d.hide()))};a.prototype.getStackBox=function(a,c,d,f,h,k,b){var g=c.axis.reversed,
e=a.inverted,l=b.height+b.pos-(e?a.plotLeft:a.plotTop);c=c.isNegative&&!g||!c.isNegative&&g;return{x:e?c?f-b.right:f-k+b.pos-a.plotLeft:d+a.xAxis[0].transB-a.plotLeft,y:e?b.height-d-h:c?l-f-k:l-f,width:e?k:h,height:e?h:k}};return a}();t.prototype.getStacks=function(){var a=this,c=a.inverted;a.yAxis.forEach(function(a){a.stacking&&a.stacking.stacks&&a.hasVisibleSeries&&(a.stacking.oldStacks=a.stacking.stacks)});a.series.forEach(function(d){var h=d.xAxis&&d.xAxis.options||{};!d.options.stacking||!0!==
d.visible&&!1!==a.options.chart.ignoreHiddenSeries||(d.stackKey=[d.type,f(d.options.stack,""),c?h.top:h.left,c?h.height:h.width].join())})};C.compose(a);x.prototype.setGroupedPoints=function(){var a=this.yAxis.stacking;this.options.centerInCategory&&(this.is("column")||this.is("columnrange"))&&!this.options.stacking&&1<this.chart.series.length?x.prototype.setStackedPoints.call(this,"group"):a&&l(a.stacks,function(c,d){"group"===d.slice(-5)&&(l(c,function(a){return a.destroy()}),delete a.stacks[d])})};
x.prototype.setStackedPoints=function(a){var c=a||this.options.stacking;if(c&&(!0===this.visible||!1===this.chart.options.chart.ignoreHiddenSeries)){var h=this.processedXData,l=this.processedYData,p=[],n=l.length,k=this.options,b=k.threshold,g=f(k.startFromThreshold&&b,0);k=k.stack;a=a?this.type+","+c:this.stackKey;var e="-"+a,u=this.negStacks,t=this.yAxis,v=t.stacking.stacks,E=t.stacking.oldStacks,z,x;t.stacking.stacksTouched+=1;for(x=0;x<n;x++){var A=h[x];var G=l[x];var I=this.getStackIndicator(I,
A,this.index);var J=I.key;var C=(z=u&&G<(g?0:b))?e:a;v[C]||(v[C]={});v[C][A]||(E[C]&&E[C][A]?(v[C][A]=E[C][A],v[C][A].total=null):v[C][A]=new w(t,t.options.stackLabels,z,A,k));C=v[C][A];null!==G?(C.points[J]=C.points[this.index]=[f(C.cumulative,g)],m(C.cumulative)||(C.base=J),C.touched=t.stacking.stacksTouched,0<I.index&&!1===this.singleStacks&&(C.points[J][0]=C.points[this.index+","+A+",0"][0])):C.points[J]=C.points[this.index]=null;"percent"===c?(z=z?a:e,u&&v[z]&&v[z][A]?(z=v[z][A],C.total=z.total=
Math.max(z.total,C.total)+Math.abs(G)||0):C.total=q(C.total+(Math.abs(G)||0))):"group"===c?(d(G)&&(G=G[0]),null!==G&&(C.total=(C.total||0)+1)):C.total=q(C.total+(G||0));C.cumulative="group"===c?(C.total||1)-1:f(C.cumulative,g)+(G||0);null!==G&&(C.points[J].push(C.cumulative),p[x]=C.cumulative,C.hasValidPoints=!0)}"percent"===c&&(t.stacking.usePercentage=!0);"group"!==c&&(this.stackedYData=p);t.stacking.oldStacks={}}};x.prototype.modifyStacks=function(){var a=this,c=a.stackKey,d=a.yAxis.stacking.stacks,
f=a.processedXData,h,l=a.options.stacking;a[l+"Stacker"]&&[c,"-"+c].forEach(function(c){for(var b=f.length,g,e;b--;)if(g=f[b],h=a.getStackIndicator(h,g,a.index,c),e=(g=d[c]&&d[c][g])&&g.points[h.key])a[l+"Stacker"](e,g,b)})};x.prototype.percentStacker=function(a,c,d){c=c.total?100/c.total:0;a[0]=q(a[0]*c);a[1]=q(a[1]*c);this.stackedYData[d]=a[1]};x.prototype.getStackIndicator=function(a,c,d,f){!m(a)||a.x!==c||f&&a.key!==f?a={x:c,index:0,key:f}:a.index++;a.key=[d,c,a.index].join();return a};G.StackItem=
w;"";return G.StackItem});L(a,"Series/Line/LineSeries.js",[a["Core/Series/Series.js"],a["Core/Series/SeriesRegistry.js"],a["Core/Utilities.js"]],function(a,t,A){var u=this&&this.__extends||function(){var a=function(u,q){a=Object.setPrototypeOf||{__proto__:[]}instanceof Array&&function(a,h){a.__proto__=h}||function(a,h){for(var d in h)h.hasOwnProperty(d)&&(a[d]=h[d])};return a(u,q)};return function(u,q){function m(){this.constructor=u}a(u,q);u.prototype=null===q?Object.create(q):(m.prototype=q.prototype,
new m)}}(),x=A.defined,C=A.merge;A=function(t){function z(){var a=null!==t&&t.apply(this,arguments)||this;a.data=void 0;a.options=void 0;a.points=void 0;return a}u(z,t);z.prototype.drawGraph=function(){var a=this,m=this.options,h=(this.gappedPath||this.getGraphPath).call(this),d=this.chart.styledMode,c=[["graph","highcharts-graph"]];d||c[0].push(m.lineColor||this.color||"#cccccc",m.dashStyle);c=a.getZonesGraphs(c);c.forEach(function(c,f){var l=c[0],p=a[l],q=p?"animate":"attr";p?(p.endX=a.preventGraphAnimation?
null:h.xMap,p.animate({d:h})):h.length&&(a[l]=p=a.chart.renderer.path(h).addClass(c[1]).attr({zIndex:1}).add(a.group));p&&!d&&(l={stroke:c[2],"stroke-width":m.lineWidth,fill:a.fillGraph&&a.color||"none"},c[3]?l.dashstyle=c[3]:"square"!==m.linecap&&(l["stroke-linecap"]=l["stroke-linejoin"]="round"),p[q](l).shadow(2>f&&m.shadow));p&&(p.startX=h.xMap,p.isArea=h.isArea)})};z.prototype.getGraphPath=function(a,m,h){var d=this,c=d.options,l=[],f=[],q,p=c.step;a=a||d.points;var u=a.reversed;u&&a.reverse();
(p={right:1,center:2}[p]||p&&3)&&u&&(p=4-p);a=this.getValidPoints(a,!1,!(c.connectNulls&&!m&&!h));a.forEach(function(r,u){var w=r.plotX,n=r.plotY,k=a[u-1];(r.leftCliff||k&&k.rightCliff)&&!h&&(q=!0);r.isNull&&!x(m)&&0<u?q=!c.connectNulls:r.isNull&&!m?q=!0:(0===u||q?u=[["M",r.plotX,r.plotY]]:d.getPointSpline?u=[d.getPointSpline(a,r,u)]:p?(u=1===p?[["L",k.plotX,n]]:2===p?[["L",(k.plotX+w)/2,k.plotY],["L",(k.plotX+w)/2,n]]:[["L",w,k.plotY]],u.push(["L",w,n])):u=[["L",w,n]],f.push(r.x),p&&(f.push(r.x),
2===p&&f.push(r.x)),l.push.apply(l,u),q=!1)});l.xMap=f;return d.graphPath=l};z.prototype.getZonesGraphs=function(a){this.zones.forEach(function(m,h){h=["zone-graph-"+h,"highcharts-graph highcharts-zone-graph-"+h+" "+(m.className||"")];this.chart.styledMode||h.push(m.color||this.color,m.dashStyle||this.options.dashStyle);a.push(h)},this);return a};z.defaultOptions=C(a.defaultOptions,{});return z}(a);t.registerSeriesType("line",A);"";return A});L(a,"Series/Area/AreaSeries.js",[a["Core/Color/Color.js"],
a["Core/Legend/LegendSymbol.js"],a["Core/Series/SeriesRegistry.js"],a["Core/Utilities.js"]],function(a,t,A,G){var u=this&&this.__extends||function(){var a=function(d,c){a=Object.setPrototypeOf||{__proto__:[]}instanceof Array&&function(a,c){a.__proto__=c}||function(a,c){for(var d in c)c.hasOwnProperty(d)&&(a[d]=c[d])};return a(d,c)};return function(d,c){function h(){this.constructor=d}a(d,c);d.prototype=null===c?Object.create(c):(h.prototype=c.prototype,new h)}}(),C=a.parse,I=A.seriesTypes.line;a=
G.extend;var z=G.merge,q=G.objectEach,m=G.pick;G=function(a){function d(){var c=null!==a&&a.apply(this,arguments)||this;c.data=void 0;c.options=void 0;c.points=void 0;return c}u(d,a);d.prototype.drawGraph=function(){this.areaPath=[];a.prototype.drawGraph.apply(this);var c=this,d=this.areaPath,f=this.options,h=[["area","highcharts-area",this.color,f.fillColor]];this.zones.forEach(function(a,d){h.push(["zone-area-"+d,"highcharts-area highcharts-zone-area-"+d+" "+a.className,a.color||c.color,a.fillColor||
f.fillColor])});h.forEach(function(a){var h=a[0],l=c[h],p=l?"animate":"attr",q={};l?(l.endX=c.preventGraphAnimation?null:d.xMap,l.animate({d:d})):(q.zIndex=0,l=c[h]=c.chart.renderer.path(d).addClass(a[1]).add(c.group),l.isArea=!0);c.chart.styledMode||(q.fill=m(a[3],C(a[2]).setOpacity(m(f.fillOpacity,.75)).get()));l[p](q);l.startX=d.xMap;l.shiftUnit=f.step?2:1})};d.prototype.getGraphPath=function(a){var c=I.prototype.getGraphPath,d=this.options,h=d.stacking,p=this.yAxis,q,r=[],u=[],t=this.index,n=
p.stacking.stacks[this.stackKey],k=d.threshold,b=Math.round(p.getThreshold(d.threshold));d=m(d.connectNulls,"percent"===h);var g=function(c,e,d){var g=a[c];c=h&&n[g.x].points[t];var f=g[d+"Null"]||0;d=g[d+"Cliff"]||0;g=!0;if(d||f){var l=(f?c[0]:c[1])+d;var m=c[0]+d;g=!!f}else!h&&a[e]&&a[e].isNull&&(l=m=k);"undefined"!==typeof l&&(u.push({plotX:D,plotY:null===l?b:p.getThreshold(l),isNull:g,isCliff:!0}),r.push({plotX:D,plotY:null===m?b:p.getThreshold(m),doCurve:!1}))};a=a||this.points;h&&(a=this.getStackPoints(a));
for(q=0;q<a.length;q++){h||(a[q].leftCliff=a[q].rightCliff=a[q].leftNull=a[q].rightNull=void 0);var e=a[q].isNull;var D=m(a[q].rectPlotX,a[q].plotX);var H=h?m(a[q].yBottom,b):b;if(!e||d)d||g(q,q-1,"left"),e&&!h&&d||(u.push(a[q]),r.push({x:q,plotX:D,plotY:H})),d||g(q,q+1,"right")}q=c.call(this,u,!0,!0);r.reversed=!0;e=c.call(this,r,!0,!0);(H=e[0])&&"M"===H[0]&&(e[0]=["L",H[1],H[2]]);e=q.concat(e);e.length&&e.push(["Z"]);c=c.call(this,u,!1,d);e.xMap=q.xMap;this.areaPath=e;return c};d.prototype.getStackPoints=
function(a){var c=this,d=[],h=[],p=this.xAxis,u=this.yAxis,r=u.stacking.stacks[this.stackKey],t={},B=u.series,n=B.length,k=u.options.reversedStacks?1:-1,b=B.indexOf(c);a=a||this.points;if(this.options.stacking){for(var g=0;g<a.length;g++)a[g].leftNull=a[g].rightNull=void 0,t[a[g].x]=a[g];q(r,function(a,b){null!==a.total&&h.push(b)});h.sort(function(a,b){return a-b});var e=B.map(function(a){return a.visible});h.forEach(function(a,g){var f=0,l,q;if(t[a]&&!t[a].isNull)d.push(t[a]),[-1,1].forEach(function(d){var f=
1===d?"rightNull":"leftNull",m=0,p=r[h[g+d]];if(p)for(var u=b;0<=u&&u<n;){var v=B[u].index;l=p.points[v];l||(v===c.index?t[a][f]=!0:e[u]&&(q=r[a].points[v])&&(m-=q[1]-q[0]));u+=k}t[a][1===d?"rightCliff":"leftCliff"]=m});else{for(var w=b;0<=w&&w<n;){if(l=r[a].points[B[w].index]){f=l[1];break}w+=k}f=m(f,0);f=u.translate(f,0,1,0,1);d.push({isNull:!0,plotX:p.translate(a,0,0,0,1),x:a,plotY:f,yBottom:f})}})}return d};d.defaultOptions=z(I.defaultOptions,{threshold:0});return d}(I);a(G.prototype,{singleStacks:!1,
drawLegendSymbol:t.drawRectangle});A.registerSeriesType("area",G);"";return G});L(a,"Series/Spline/SplineSeries.js",[a["Core/Series/SeriesRegistry.js"],a["Core/Utilities.js"]],function(a,t){var u=this&&this.__extends||function(){var a=function(u,q){a=Object.setPrototypeOf||{__proto__:[]}instanceof Array&&function(a,h){a.__proto__=h}||function(a,h){for(var d in h)h.hasOwnProperty(d)&&(a[d]=h[d])};return a(u,q)};return function(u,q){function m(){this.constructor=u}a(u,q);u.prototype=null===q?Object.create(q):
(m.prototype=q.prototype,new m)}}(),G=a.seriesTypes.line,x=t.merge,C=t.pick;t=function(a){function t(){var q=null!==a&&a.apply(this,arguments)||this;q.data=void 0;q.options=void 0;q.points=void 0;return q}u(t,a);t.prototype.getPointSpline=function(a,m,h){var d=m.plotX||0,c=m.plotY||0,l=a[h-1];h=a[h+1];if(l&&!l.isNull&&!1!==l.doCurve&&!m.isCliff&&h&&!h.isNull&&!1!==h.doCurve&&!m.isCliff){a=l.plotY||0;var f=h.plotX||0;h=h.plotY||0;var q=0;var p=(1.5*d+(l.plotX||0))/2.5;var u=(1.5*c+a)/2.5;f=(1.5*d+
f)/2.5;var r=(1.5*c+h)/2.5;f!==p&&(q=(r-u)*(f-d)/(f-p)+c-r);u+=q;r+=q;u>a&&u>c?(u=Math.max(a,c),r=2*c-u):u<a&&u<c&&(u=Math.min(a,c),r=2*c-u);r>h&&r>c?(r=Math.max(h,c),u=2*c-r):r<h&&r<c&&(r=Math.min(h,c),u=2*c-r);m.rightContX=f;m.rightContY=r}m=["C",C(l.rightContX,l.plotX,0),C(l.rightContY,l.plotY,0),C(p,d,0),C(u,c,0),d,c];l.rightContX=l.rightContY=void 0;return m};t.defaultOptions=x(G.defaultOptions);return t}(G);a.registerSeriesType("spline",t);"";return t});L(a,"Series/AreaSpline/AreaSplineSeries.js",
[a["Series/Area/AreaSeries.js"],a["Series/Spline/SplineSeries.js"],a["Core/Legend/LegendSymbol.js"],a["Core/Series/SeriesRegistry.js"],a["Core/Utilities.js"]],function(a,t,A,G,x){var u=this&&this.__extends||function(){var a=function(h,d){a=Object.setPrototypeOf||{__proto__:[]}instanceof Array&&function(a,d){a.__proto__=d}||function(a,d){for(var c in d)d.hasOwnProperty(c)&&(a[c]=d[c])};return a(h,d)};return function(h,d){function c(){this.constructor=h}a(h,d);h.prototype=null===d?Object.create(d):
(c.prototype=d.prototype,new c)}}(),I=a.prototype,z=x.extend,q=x.merge;x=function(m){function h(){var a=null!==m&&m.apply(this,arguments)||this;a.data=void 0;a.points=void 0;a.options=void 0;return a}u(h,m);h.defaultOptions=q(t.defaultOptions,a.defaultOptions);return h}(t);z(x.prototype,{getGraphPath:I.getGraphPath,getStackPoints:I.getStackPoints,drawGraph:I.drawGraph,drawLegendSymbol:A.drawRectangle});G.registerSeriesType("areaspline",x);"";return x});L(a,"Series/Column/ColumnSeries.js",[a["Core/Animation/AnimationUtilities.js"],
a["Core/Color/Color.js"],a["Core/Globals.js"],a["Core/Legend/LegendSymbol.js"],a["Core/Series/Series.js"],a["Core/Series/SeriesRegistry.js"],a["Core/Utilities.js"]],function(a,t,A,G,x,C,I){var u=this&&this.__extends||function(){var a=function(c,b){a=Object.setPrototypeOf||{__proto__:[]}instanceof Array&&function(a,b){a.__proto__=b}||function(a,b){for(var c in b)b.hasOwnProperty(c)&&(a[c]=b[c])};return a(c,b)};return function(c,b){function d(){this.constructor=c}a(c,b);c.prototype=null===b?Object.create(b):
(d.prototype=b.prototype,new d)}}(),q=a.animObject,m=t.parse,h=A.hasTouch;a=A.noop;var d=I.clamp,c=I.css,l=I.defined,f=I.extend,w=I.fireEvent,p=I.isArray,K=I.isNumber,r=I.merge,y=I.pick,B=I.objectEach;I=function(a){function k(){var b=null!==a&&a.apply(this,arguments)||this;b.borderWidth=void 0;b.data=void 0;b.group=void 0;b.options=void 0;b.points=void 0;return b}u(k,a);k.prototype.animate=function(a){var b=this,c=this.yAxis,h=b.options,k=this.chart.inverted,l={},n=k?"translateX":"translateY";if(a)l.scaleY=
.001,a=d(c.toPixels(h.threshold),c.pos,c.pos+c.len),k?l.translateX=a-c.len:l.translateY=a,b.clipBox&&b.setClip(),b.group.attr(l);else{var m=Number(b.group.attr(n));b.group.animate({scaleY:1},f(q(b.options.animation),{step:function(a,e){b.group&&(l[n]=m+e.pos*(c.pos-m),b.group.attr(l))}}))}};k.prototype.init=function(b,c){a.prototype.init.apply(this,arguments);var e=this;b=e.chart;b.hasRendered&&b.series.forEach(function(a){a.type===e.type&&(a.isDirty=!0)})};k.prototype.getColumnMetrics=function(){var a=
this,c=a.options,e=a.xAxis,d=a.yAxis,f=e.options.reversedStacks;f=e.reversed&&!f||!e.reversed&&f;var h={},k,l=0;!1===c.grouping?l=1:a.chart.series.forEach(function(b){var c=b.yAxis,e=b.options;if(b.type===a.type&&(b.visible||!a.chart.options.chart.ignoreHiddenSeries)&&d.len===c.len&&d.pos===c.pos){if(e.stacking&&"group"!==e.stacking){k=b.stackKey;"undefined"===typeof h[k]&&(h[k]=l++);var g=h[k]}else!1!==e.grouping&&(g=l++);b.columnIndex=g}});var n=Math.min(Math.abs(e.transA)*(e.ordinal&&e.ordinal.slope||
c.pointRange||e.closestPointRange||e.tickInterval||1),e.len),m=n*c.groupPadding,p=(n-2*m)/(l||1);c=Math.min(c.maxPointWidth||e.len,y(c.pointWidth,p*(1-2*c.pointPadding)));a.columnMetrics={width:c,offset:(p-c)/2+(m+((a.columnIndex||0)+(f?1:0))*p-n/2)*(f?-1:1),paddedWidth:p,columnCount:l};return a.columnMetrics};k.prototype.crispCol=function(a,c,e,d){var b=this.chart,g=this.borderWidth,f=-(g%2?.5:0);g=g%2?.5:1;b.inverted&&b.renderer.isVML&&(g+=1);this.options.crisp&&(e=Math.round(a+e)+f,a=Math.round(a)+
f,e-=a);d=Math.round(c+d)+g;f=.5>=Math.abs(c)&&.5<d;c=Math.round(c)+g;d-=c;f&&d&&(--c,d+=1);return{x:a,y:c,width:e,height:d}};k.prototype.adjustForMissingColumns=function(a,c,e,d){var b=this,g=this.options.stacking;if(!e.isNull&&1<d.columnCount){var f=0,h=0;B(this.yAxis.stacking&&this.yAxis.stacking.stacks,function(a){if("number"===typeof e.x&&(a=a[e.x.toString()])){var c=a.points[b.index],d=a.total;g?(c&&(f=h),a.hasValidPoints&&h++):p(c)&&(f=c[1],h=d||0)}});a=(e.plotX||0)+((h-1)*d.paddedWidth+c)/
2-c-f*d.paddedWidth}return a};k.prototype.translate=function(){var a=this,c=a.chart,e=a.options,f=a.dense=2>a.closestPointRange*a.xAxis.transA;f=a.borderWidth=y(e.borderWidth,f?0:1);var h=a.xAxis,k=a.yAxis,n=e.threshold,m=a.translatedThreshold=k.getThreshold(n),p=y(e.minPointLength,5),q=a.getColumnMetrics(),r=q.width,u=a.pointXOffset=q.offset,t=a.dataMin,w=a.dataMax,B=a.barW=Math.max(r,1+2*f);c.inverted&&(m-=.5);e.pointPadding&&(B=Math.ceil(B));x.prototype.translate.apply(a);a.points.forEach(function(b){var g=
y(b.yBottom,m),f=999+Math.abs(g),v=b.plotX||0;f=d(b.plotY,-f,k.len+f);var E=Math.min(f,g),D=Math.max(f,g)-E,x=r,H=v+u,z=B;p&&Math.abs(D)<p&&(D=p,v=!k.reversed&&!b.negative||k.reversed&&b.negative,K(n)&&K(w)&&b.y===n&&w<=n&&(k.min||0)<n&&(t!==w||(k.max||0)<=n)&&(v=!v),E=Math.abs(E-m)>p?g-p:m-(v?p:0));l(b.options.pointWidth)&&(x=z=Math.ceil(b.options.pointWidth),H-=Math.round((x-r)/2));e.centerInCategory&&(H=a.adjustForMissingColumns(H,x,b,q));b.barX=H;b.pointWidth=x;b.tooltipPos=c.inverted?[d(k.len+
k.pos-c.plotLeft-f,k.pos-c.plotLeft,k.len+k.pos-c.plotLeft),h.len+h.pos-c.plotTop-H-z/2,D]:[h.left-c.plotLeft+H+z/2,d(f+k.pos-c.plotTop,k.pos-c.plotTop,k.len+k.pos-c.plotTop),D];b.shapeType=a.pointClass.prototype.shapeType||"rect";b.shapeArgs=a.crispCol.apply(a,b.isNull?[H,m,z,0]:[H,E,z,D])})};k.prototype.drawGraph=function(){this.group[this.dense?"addClass":"removeClass"]("highcharts-dense-data")};k.prototype.pointAttribs=function(a,c){var b=this.options,d=this.pointAttrToOptions||{},g=d.stroke||
"borderColor",f=d["stroke-width"]||"borderWidth",h=a&&a.color||this.color,k=a&&a[g]||b[g]||h;d=a&&a.options.dashStyle||b.dashStyle;var l=a&&a[f]||b[f]||this[f]||0,n=y(a&&a.opacity,b.opacity,1);if(a&&this.zones.length){var p=a.getZone();h=a.options.color||p&&(p.color||a.nonZonedColor)||this.color;p&&(k=p.borderColor||k,d=p.dashStyle||d,l=p.borderWidth||l)}c&&a&&(a=r(b.states[c],a.options.states&&a.options.states[c]||{}),c=a.brightness,h=a.color||"undefined"!==typeof c&&m(h).brighten(a.brightness).get()||
h,k=a[g]||k,l=a[f]||l,d=a.dashStyle||d,n=y(a.opacity,n));g={fill:h,stroke:k,"stroke-width":l,opacity:n};d&&(g.dashstyle=d);return g};k.prototype.drawPoints=function(){var a=this,c=this.chart,e=a.options,d=c.renderer,f=e.animationLimit||250,h;a.points.forEach(function(b){var g=b.graphic,k=!!g,l=g&&c.pointCount<f?"animate":"attr";if(K(b.plotY)&&null!==b.y){h=b.shapeArgs;g&&b.hasNewShapeType()&&(g=g.destroy());a.enabledDataSorting&&(b.startXPos=a.xAxis.reversed?-(h?h.width||0:0):a.xAxis.width);g||(b.graphic=
g=d[b.shapeType](h).add(b.group||a.group))&&a.enabledDataSorting&&c.hasRendered&&c.pointCount<f&&(g.attr({x:b.startXPos}),k=!0,l="animate");if(g&&k)g[l](r(h));if(e.borderRadius)g[l]({r:e.borderRadius});c.styledMode||g[l](a.pointAttribs(b,b.selected&&"select")).shadow(!1!==b.allowShadow&&e.shadow,null,e.stacking&&!e.borderRadius);g&&(g.addClass(b.getClassName(),!0),g.attr({visibility:b.visible?"inherit":"hidden"}))}else g&&(b.graphic=g.destroy())})};k.prototype.drawTracker=function(){var a=this,d=
a.chart,e=d.pointer,f=function(a){var b=e.getPointFromEvent(a);"undefined"!==typeof b&&(e.isDirectTouch=!0,b.onMouseOver(a))},k;a.points.forEach(function(a){k=p(a.dataLabels)?a.dataLabels:a.dataLabel?[a.dataLabel]:[];a.graphic&&(a.graphic.element.point=a);k.forEach(function(b){b.div?b.div.point=a:b.element.point=a})});a._hasTracking||(a.trackerGroups.forEach(function(b){if(a[b]){a[b].addClass("highcharts-tracker").on("mouseover",f).on("mouseout",function(a){e.onTrackerMouseOut(a)});if(h)a[b].on("touchstart",
f);!d.styledMode&&a.options.cursor&&a[b].css(c).css({cursor:a.options.cursor})}}),a._hasTracking=!0);w(this,"afterDrawTracker")};k.prototype.remove=function(){var a=this,c=a.chart;c.hasRendered&&c.series.forEach(function(b){b.type===a.type&&(b.isDirty=!0)});x.prototype.remove.apply(a,arguments)};k.defaultOptions=r(x.defaultOptions,{borderRadius:0,centerInCategory:!1,groupPadding:.2,marker:null,pointPadding:.1,minPointLength:0,cropThreshold:50,pointRange:null,states:{hover:{halo:!1,brightness:.1},
select:{color:"#cccccc",borderColor:"#000000"}},dataLabels:{align:void 0,verticalAlign:void 0,y:void 0},startFromThreshold:!0,stickyTracking:!1,tooltip:{distance:6},threshold:0,borderColor:"#ffffff"});return k}(x);f(I.prototype,{cropShoulder:0,directTouch:!0,drawLegendSymbol:G.drawRectangle,getSymbol:a,negStacks:!0,trackerGroups:["group","dataLabelsGroup"]});C.registerSeriesType("column",I);"";"";return I});L(a,"Core/Series/DataLabel.js",[a["Core/Animation/AnimationUtilities.js"],a["Core/FormatUtilities.js"],
a["Core/Utilities.js"]],function(a,t,A){var u=a.getDeferredAnimation,x=t.format,C=A.defined,I=A.extend,z=A.fireEvent,q=A.isArray,m=A.merge,h=A.objectEach,d=A.pick,c=A.splat,l;(function(a){function f(a,b,c,e,f){var g=this,h=this.chart,k=this.isCartesian&&h.inverted,l=this.enabledDataSorting,n=d(a.dlBox&&a.dlBox.centerX,a.plotX,-9999),m=d(a.plotY,-9999),p=b.getBBox(),q=c.rotation,r=c.align,u=h.isInsidePlot(n,Math.round(m),{inverted:k,paneCoordinates:!0,series:g}),t=function(c){l&&g.xAxis&&!w&&g.setDataLabelStartPos(a,
b,f,u,c)},w="justify"===d(c.overflow,l?"none":"justify"),B=this.visible&&!1!==a.visible&&(a.series.forceDL||l&&!w||u||d(c.inside,!!this.options.stacking)&&e&&h.isInsidePlot(n,k?e.x+1:e.y+e.height-1,{inverted:k,paneCoordinates:!0,series:g}));if(B){var y=h.renderer.fontMetrics(h.styledMode?void 0:c.style.fontSize,b).b;e=I({x:k?this.yAxis.len-m:n,y:Math.round(k?this.xAxis.len-n:m),width:0,height:0},e);I(c,{width:p.width,height:p.height});q?(w=!1,n=h.renderer.rotCorr(y,q),n={x:e.x+(c.x||0)+e.width/2+
n.x,y:e.y+(c.y||0)+{top:0,middle:.5,bottom:1}[c.verticalAlign]*e.height},t(n),b[f?"attr":"animate"](n).attr({align:r}),t=(q+720)%360,t=180<t&&360>t,"left"===r?n.y-=t?p.height:0:"center"===r?(n.x-=p.width/2,n.y-=p.height/2):"right"===r&&(n.x-=p.width,n.y-=t?0:p.height),b.placed=!0,b.alignAttr=n):(t(e),b.align(c,void 0,e),n=b.alignAttr);w&&0<=e.height?this.justifyDataLabel(b,c,n,p,e,f):d(c.crop,!0)&&(B=h.isInsidePlot(n.x,n.y,{paneCoordinates:!0,series:g})&&h.isInsidePlot(n.x+p.width,n.y+p.height,{paneCoordinates:!0,
series:g}));if(c.shape&&!q)b[f?"attr":"animate"]({anchorX:k?h.plotWidth-a.plotY:a.plotX,anchorY:k?h.plotHeight-a.plotX:a.plotY})}f&&l&&(b.placed=!1);B||l&&!w||(b.hide(!0),b.placed=!1)}function l(a,b){var c=b.filter;return c?(b=c.operator,a=a[c.property],c=c.value,">"===b&&a>c||"<"===b&&a<c||">="===b&&a>=c||"<="===b&&a<=c||"=="===b&&a==c||"==="===b&&a===c?!0:!1):!0}function t(){var a=this,b=a.chart,g=a.options,e=a.points,f=a.hasRendered||0,n=b.renderer,m=g.dataLabels,p,r=m.animation;r=m.defer?u(b,
r,a):{defer:0,duration:0};m=y(y(b.options.plotOptions&&b.options.plotOptions.series&&b.options.plotOptions.series.dataLabels,b.options.plotOptions&&b.options.plotOptions[a.type]&&b.options.plotOptions[a.type].dataLabels),m);z(this,"drawDataLabels");if(q(m)||m.enabled||a._hasPointLabels){var t=a.plotGroup("dataLabelsGroup","data-labels",f?"inherit":"hidden",m.zIndex||6);t.attr({opacity:+f});!f&&(f=a.dataLabelsGroup)&&(a.visible&&t.show(!0),f[g.animation?"animate":"attr"]({opacity:1},r));e.forEach(function(e){p=
c(y(m,e.dlOptions||e.options&&e.options.dataLabels));p.forEach(function(c,f){var k=c.enabled&&(!e.isNull||e.dataLabelOnNull)&&l(e,c),m=e.connectors?e.connectors[f]:e.connector,p=e.dataLabels?e.dataLabels[f]:e.dataLabel,q=d(c.distance,e.labelDistance),r=!p;if(k){var u=e.getLabelConfig();var w=d(c[e.formatPrefix+"Format"],c.format);u=C(w)?x(w,u,b):(c[e.formatPrefix+"Formatter"]||c.formatter).call(u,c);w=c.style;var B=c.rotation;b.styledMode||(w.color=d(c.color,w.color,a.color,"#000000"),"contrast"===
w.color?(e.contrastColor=n.getContrast(e.color||a.color),w.color=!C(q)&&c.inside||0>q||g.stacking?e.contrastColor:"#000000"):delete e.contrastColor,g.cursor&&(w.cursor=g.cursor));var v={r:c.borderRadius||0,rotation:B,padding:c.padding,zIndex:1};b.styledMode||(v.fill=c.backgroundColor,v.stroke=c.borderColor,v["stroke-width"]=c.borderWidth);h(v,function(a,b){"undefined"===typeof a&&delete v[b]})}!p||k&&C(u)?k&&C(u)&&(p?v.text=u:(e.dataLabels=e.dataLabels||[],p=e.dataLabels[f]=B?n.text(u,0,-9999,c.useHTML).addClass("highcharts-data-label"):
n.label(u,0,-9999,c.shape,null,null,c.useHTML,null,"data-label"),f||(e.dataLabel=p),p.addClass(" highcharts-data-label-color-"+e.colorIndex+" "+(c.className||"")+(c.useHTML?" highcharts-tracker":""))),p.options=c,p.attr(v),b.styledMode||p.css(w).shadow(c.shadow),p.added||p.add(t),c.textPath&&!c.useHTML&&(p.setTextPath(e.getDataLabelPath&&e.getDataLabelPath(p)||e.graphic,c.textPath),e.dataLabelPath&&!c.textPath.enabled&&(e.dataLabelPath=e.dataLabelPath.destroy())),a.alignDataLabel(e,p,c,null,r)):(e.dataLabel=
e.dataLabel&&e.dataLabel.destroy(),e.dataLabels&&(1===e.dataLabels.length?delete e.dataLabels:delete e.dataLabels[f]),f||delete e.dataLabel,m&&(e.connector=e.connector.destroy(),e.connectors&&(1===e.connectors.length?delete e.connectors:delete e.connectors[f])))})})}z(this,"afterDrawDataLabels")}function r(a,b,c,e,d,f){var g=this.chart,h=b.align,k=b.verticalAlign,l=a.box?0:a.padding||0,n=b.x;n=void 0===n?0:n;var m=b.y;m=void 0===m?0:m;var p=(c.x||0)+l;if(0>p){"right"===h&&0<=n?(b.align="left",b.inside=
!0):n-=p;var q=!0}p=(c.x||0)+e.width-l;p>g.plotWidth&&("left"===h&&0>=n?(b.align="right",b.inside=!0):n+=g.plotWidth-p,q=!0);p=c.y+l;0>p&&("bottom"===k&&0<=m?(b.verticalAlign="top",b.inside=!0):m-=p,q=!0);p=(c.y||0)+e.height-l;p>g.plotHeight&&("top"===k&&0>=m?(b.verticalAlign="bottom",b.inside=!0):m+=g.plotHeight-p,q=!0);q&&(b.x=n,b.y=m,a.placed=!f,a.align(b,void 0,d));return q}function y(a,b){var c=[],e;if(q(a)&&!q(b))c=a.map(function(a){return m(a,b)});else if(q(b)&&!q(a))c=b.map(function(b){return m(a,
b)});else if(q(a)||q(b))for(e=Math.max(a.length,b.length);e--;)c[e]=m(a[e],b[e]);else c=m(a,b);return c}function B(a,b,c,e,d){var f=this.chart,g=f.inverted,h=this.xAxis,k=h.reversed,l=g?b.height/2:b.width/2;a=(a=a.pointWidth)?a/2:0;b.startXPos=g?d.x:k?-l-a:h.width-l+a;b.startYPos=g?k?this.yAxis.height-l+a:-l-a:d.y;e?"hidden"===b.visibility&&(b.show(),b.attr({opacity:0}).animate({opacity:1})):b.attr({opacity:1}).animate({opacity:0},void 0,b.hide);f.hasRendered&&(c&&b.attr({x:b.startXPos,y:b.startYPos}),
b.placed=!0)}var n=[];a.compose=function(a){if(-1===n.indexOf(a)){var b=a.prototype;n.push(a);b.alignDataLabel=f;b.drawDataLabels=t;b.justifyDataLabel=r;b.setDataLabelStartPos=B}}})(l||(l={}));"";return l});L(a,"Series/Column/ColumnDataLabel.js",[a["Core/Series/DataLabel.js"],a["Core/Series/SeriesRegistry.js"],a["Core/Utilities.js"]],function(a,t,A){var u=t.series,x=A.merge,C=A.pick,I;(function(t){function q(a,d,c,l,f){var h=this.chart.inverted,m=a.series,q=(m.xAxis?m.xAxis.len:this.chart.plotSizeX)||
0;m=(m.yAxis?m.yAxis.len:this.chart.plotSizeY)||0;var r=a.dlBox||a.shapeArgs,t=C(a.below,a.plotY>C(this.translatedThreshold,m)),B=C(c.inside,!!this.options.stacking);r&&(l=x(r),0>l.y&&(l.height+=l.y,l.y=0),r=l.y+l.height-m,0<r&&r<l.height&&(l.height-=r),h&&(l={x:m-l.y-l.height,y:q-l.x-l.width,width:l.height,height:l.width}),B||(h?(l.x+=t?0:l.width,l.width=0):(l.y+=t?l.height:0,l.height=0)));c.align=C(c.align,!h||B?"center":t?"right":"left");c.verticalAlign=C(c.verticalAlign,h||B?"middle":t?"top":
"bottom");u.prototype.alignDataLabel.call(this,a,d,c,l,f);c.inside&&a.contrastColor&&d.css({color:a.contrastColor})}var m=[];t.compose=function(h){a.compose(u);-1===m.indexOf(h)&&(m.push(h),h.prototype.alignDataLabel=q)}})(I||(I={}));return I});L(a,"Series/Bar/BarSeries.js",[a["Series/Column/ColumnSeries.js"],a["Core/Series/SeriesRegistry.js"],a["Core/Utilities.js"]],function(a,t,A){var u=this&&this.__extends||function(){var a=function(u,q){a=Object.setPrototypeOf||{__proto__:[]}instanceof Array&&
function(a,h){a.__proto__=h}||function(a,h){for(var d in h)h.hasOwnProperty(d)&&(a[d]=h[d])};return a(u,q)};return function(u,q){function m(){this.constructor=u}a(u,q);u.prototype=null===q?Object.create(q):(m.prototype=q.prototype,new m)}}(),x=A.extend,C=A.merge;A=function(t){function x(){var a=null!==t&&t.apply(this,arguments)||this;a.data=void 0;a.options=void 0;a.points=void 0;return a}u(x,t);x.defaultOptions=C(a.defaultOptions,{});return x}(a);x(A.prototype,{inverted:!0});t.registerSeriesType("bar",
A);"";return A});L(a,"Series/Scatter/ScatterSeries.js",[a["Series/Column/ColumnSeries.js"],a["Series/Line/LineSeries.js"],a["Core/Series/SeriesRegistry.js"],a["Core/Utilities.js"]],function(a,t,A,G){var u=this&&this.__extends||function(){var a=function(m,h){a=Object.setPrototypeOf||{__proto__:[]}instanceof Array&&function(a,c){a.__proto__=c}||function(a,c){for(var d in c)c.hasOwnProperty(d)&&(a[d]=c[d])};return a(m,h)};return function(m,h){function d(){this.constructor=m}a(m,h);m.prototype=null===
h?Object.create(h):(d.prototype=h.prototype,new d)}}(),C=G.addEvent,I=G.extend,z=G.merge;G=function(a){function m(){var h=null!==a&&a.apply(this,arguments)||this;h.data=void 0;h.options=void 0;h.points=void 0;return h}u(m,a);m.prototype.applyJitter=function(){var a=this,d=this.options.jitter,c=this.points.length;d&&this.points.forEach(function(h,f){["x","y"].forEach(function(l,m){var p="plot"+l.toUpperCase();if(d[l]&&!h.isNull){var q=a[l+"Axis"];var u=d[l]*q.transA;if(q&&!q.isLog){var t=Math.max(0,
h[p]-u);q=Math.min(q.len,h[p]+u);m=1E4*Math.sin(f+m*c);h[p]=t+(q-t)*(m-Math.floor(m));"x"===l&&(h.clientX=h.plotX)}}})})};m.prototype.drawGraph=function(){this.options.lineWidth?a.prototype.drawGraph.call(this):this.graph&&(this.graph=this.graph.destroy())};m.defaultOptions=z(t.defaultOptions,{lineWidth:0,findNearestPointBy:"xy",jitter:{x:0,y:0},marker:{enabled:!0},tooltip:{headerFormat:'<span style="color:{point.color}">\u25cf</span> <span style="font-size: 10px"> {series.name}</span><br/>',pointFormat:"x: <b>{point.x}</b><br/>y: <b>{point.y}</b><br/>"}});
return m}(t);I(G.prototype,{drawTracker:a.prototype.drawTracker,sorted:!1,requireSorting:!1,noSharedTooltip:!0,trackerGroups:["group","markerGroup","dataLabelsGroup"],takeOrdinalPosition:!1});C(G,"afterTranslate",function(){this.applyJitter()});A.registerSeriesType("scatter",G);"";return G});L(a,"Series/CenteredUtilities.js",[a["Core/Globals.js"],a["Core/Series/Series.js"],a["Core/Utilities.js"]],function(a,t,A){var u=a.deg2rad,x=A.isNumber,C=A.pick,I=A.relativeLength,z;(function(a){a.getCenter=function(){var a=
this.options,h=this.chart,d=2*(a.slicedOffset||0),c=h.plotWidth-2*d,l=h.plotHeight-2*d,f=a.center,q=Math.min(c,l),p=a.size,u=a.innerSize||0;"string"===typeof p&&(p=parseFloat(p));"string"===typeof u&&(u=parseFloat(u));a=[C(f[0],"50%"),C(f[1],"50%"),C(p&&0>p?void 0:a.size,"100%"),C(u&&0>u?void 0:a.innerSize||0,"0%")];!h.angular||this instanceof t||(a[3]=0);for(f=0;4>f;++f)p=a[f],h=2>f||2===f&&/%$/.test(p),a[f]=I(p,[c,l,q,a[2]][f])+(h?d:0);a[3]>a[2]&&(a[3]=a[2]);return a};a.getStartAndEndRadians=function(a,
h){a=x(a)?a:0;h=x(h)&&h>a&&360>h-a?h:a+360;return{start:u*(a+-90),end:u*(h+-90)}}})(z||(z={}));"";return z});L(a,"Series/Pie/PiePoint.js",[a["Core/Animation/AnimationUtilities.js"],a["Core/Series/Point.js"],a["Core/Utilities.js"]],function(a,t,A){var u=this&&this.__extends||function(){var a=function(d,c){a=Object.setPrototypeOf||{__proto__:[]}instanceof Array&&function(a,c){a.__proto__=c}||function(a,c){for(var d in c)c.hasOwnProperty(d)&&(a[d]=c[d])};return a(d,c)};return function(d,c){function h(){this.constructor=
d}a(d,c);d.prototype=null===c?Object.create(c):(h.prototype=c.prototype,new h)}}(),x=a.setAnimation,C=A.addEvent,I=A.defined;a=A.extend;var z=A.isNumber,q=A.pick,m=A.relativeLength;t=function(a){function d(){var c=null!==a&&a.apply(this,arguments)||this;c.labelDistance=void 0;c.options=void 0;c.series=void 0;return c}u(d,a);d.prototype.getConnectorPath=function(){var a=this.labelPosition,d=this.series.options.dataLabels,f=this.connectorShapes,h=d.connectorShape;f[h]&&(h=f[h]);return h.call(this,{x:a.final.x,
y:a.final.y,alignment:a.alignment},a.connectorPosition,d)};d.prototype.getTranslate=function(){return this.sliced?this.slicedTranslation:{translateX:0,translateY:0}};d.prototype.haloPath=function(a){var c=this.shapeArgs;return this.sliced||!this.visible?[]:this.series.chart.renderer.symbols.arc(c.x,c.y,c.r+a,c.r+a,{innerR:c.r-1,start:c.start,end:c.end})};d.prototype.init=function(){var c=this;a.prototype.init.apply(this,arguments);this.name=q(this.name,"Slice");var d=function(a){c.slice("select"===
a.type)};C(this,"select",d);C(this,"unselect",d);return this};d.prototype.isValid=function(){return z(this.y)&&0<=this.y};d.prototype.setVisible=function(a,d){var c=this,h=this.series,l=h.chart,m=h.options.ignoreHiddenPoint;d=q(d,m);a!==this.visible&&(this.visible=this.options.visible=a="undefined"===typeof a?!this.visible:a,h.options.data[h.data.indexOf(this)]=this.options,["graphic","dataLabel","connector","shadowGroup"].forEach(function(d){if(c[d])c[d][a?"show":"hide"](a)}),this.legendItem&&l.legend.colorizeItem(this,
a),a||"hover"!==this.state||this.setState(""),m&&(h.isDirty=!0),d&&l.redraw())};d.prototype.slice=function(a,d,f){var c=this.series;x(f,c.chart);q(d,!0);this.sliced=this.options.sliced=I(a)?a:!this.sliced;c.options.data[c.data.indexOf(this)]=this.options;this.graphic&&this.graphic.animate(this.getTranslate());this.shadowGroup&&this.shadowGroup.animate(this.getTranslate())};return d}(t);a(t.prototype,{connectorShapes:{fixedOffset:function(a,d,c){var h=d.breakAt;d=d.touchingSliceAt;return[["M",a.x,
a.y],c.softConnector?["C",a.x+("left"===a.alignment?-5:5),a.y,2*h.x-d.x,2*h.y-d.y,h.x,h.y]:["L",h.x,h.y],["L",d.x,d.y]]},straight:function(a,d){d=d.touchingSliceAt;return[["M",a.x,a.y],["L",d.x,d.y]]},crookedLine:function(a,d,c){d=d.touchingSliceAt;var h=this.series,f=h.center[0],q=h.chart.plotWidth,p=h.chart.plotLeft;h=a.alignment;var u=this.shapeArgs.r;c=m(c.crookDistance,1);q="left"===h?f+u+(q+p-f-u)*(1-c):p+(f-u)*c;c=["L",q,a.y];f=!0;if("left"===h?q>a.x||q<d.x:q<a.x||q>d.x)f=!1;a=[["M",a.x,a.y]];
f&&a.push(c);a.push(["L",d.x,d.y]);return a}}});return t});L(a,"Series/Pie/PieSeries.js",[a["Series/CenteredUtilities.js"],a["Series/Column/ColumnSeries.js"],a["Core/Globals.js"],a["Core/Legend/LegendSymbol.js"],a["Series/Pie/PiePoint.js"],a["Core/Series/Series.js"],a["Core/Series/SeriesRegistry.js"],a["Core/Renderer/SVG/Symbols.js"],a["Core/Utilities.js"]],function(a,t,A,G,x,C,I,z,q){var m=this&&this.__extends||function(){var a=function(c,d){a=Object.setPrototypeOf||{__proto__:[]}instanceof Array&&
function(a,c){a.__proto__=c}||function(a,c){for(var d in c)c.hasOwnProperty(d)&&(a[d]=c[d])};return a(c,d)};return function(c,d){function f(){this.constructor=c}a(c,d);c.prototype=null===d?Object.create(d):(f.prototype=d.prototype,new f)}}(),h=a.getStartAndEndRadians;A=A.noop;var d=q.clamp,c=q.extend,l=q.fireEvent,f=q.merge,u=q.pick,p=q.relativeLength;q=function(a){function c(){var c=null!==a&&a.apply(this,arguments)||this;c.center=void 0;c.data=void 0;c.maxLabelDistance=void 0;c.options=void 0;c.points=
void 0;return c}m(c,a);c.prototype.animate=function(a){var c=this,d=c.points,f=c.startAngleRad;a||d.forEach(function(a){var b=a.graphic,e=a.shapeArgs;b&&e&&(b.attr({r:u(a.startR,c.center&&c.center[3]/2),start:f,end:f}),b.animate({r:e.r,start:e.start,end:e.end},c.options.animation))})};c.prototype.drawEmpty=function(){var a=this.startAngleRad,c=this.endAngleRad,d=this.options;if(0===this.total&&this.center){var f=this.center[0];var b=this.center[1];this.graph||(this.graph=this.chart.renderer.arc(f,
b,this.center[1]/2,0,a,c).addClass("highcharts-empty-series").add(this.group));this.graph.attr({d:z.arc(f,b,this.center[2]/2,0,{start:a,end:c,innerR:this.center[3]/2})});this.chart.styledMode||this.graph.attr({"stroke-width":d.borderWidth,fill:d.fillColor||"none",stroke:d.color||"#cccccc"})}else this.graph&&(this.graph=this.graph.destroy())};c.prototype.drawPoints=function(){var a=this.chart.renderer;this.points.forEach(function(c){c.graphic&&c.hasNewShapeType()&&(c.graphic=c.graphic.destroy());c.graphic||
(c.graphic=a[c.shapeType](c.shapeArgs).add(c.series.group),c.delayedRendering=!0)})};c.prototype.generatePoints=function(){a.prototype.generatePoints.call(this);this.updateTotals()};c.prototype.getX=function(a,c,f){var h=this.center,b=this.radii?this.radii[f.index]||0:h[2]/2;a=Math.asin(d((a-h[1])/(b+f.labelDistance),-1,1));return h[0]+(c?-1:1)*Math.cos(a)*(b+f.labelDistance)+(0<f.labelDistance?(c?-1:1)*this.options.dataLabels.padding:0)};c.prototype.hasData=function(){return!!this.processedXData.length};
c.prototype.redrawPoints=function(){var a=this,c=a.chart,d=c.renderer,h=a.options.shadow,b,g,e,l;this.drawEmpty();!h||a.shadowGroup||c.styledMode||(a.shadowGroup=d.g("shadow").attr({zIndex:-1}).add(a.group));a.points.forEach(function(k){var n={};g=k.graphic;if(!k.isNull&&g){var m=void 0;l=k.shapeArgs;b=k.getTranslate();c.styledMode||(m=k.shadowGroup,h&&!m&&(m=k.shadowGroup=d.g("shadow").add(a.shadowGroup)),m&&m.attr(b),e=a.pointAttribs(k,k.selected&&"select"));k.delayedRendering?(g.setRadialReference(a.center).attr(l).attr(b),
c.styledMode||g.attr(e).attr({"stroke-linejoin":"round"}).shadow(h,m),k.delayedRendering=!1):(g.setRadialReference(a.center),c.styledMode||f(!0,n,e),f(!0,n,l,b),g.animate(n));g.attr({visibility:k.visible?"inherit":"hidden"});g.addClass(k.getClassName(),!0)}else g&&(k.graphic=g.destroy())})};c.prototype.sortByAngle=function(a,c){a.sort(function(a,d){return"undefined"!==typeof a.angle&&(d.angle-a.angle)*c})};c.prototype.translate=function(a){this.generatePoints();var c=this.options,d=c.slicedOffset,
f=d+(c.borderWidth||0),b=h(c.startAngle,c.endAngle),g=this.startAngleRad=b.start;b=(this.endAngleRad=b.end)-g;var e=this.points,m=c.dataLabels.distance;c=c.ignoreHiddenPoint;var q=e.length,r,t=0;a||(this.center=a=this.getCenter());for(r=0;r<q;r++){var w=e[r];var x=g+t*b;!w.isValid()||c&&!w.visible||(t+=w.percentage/100);var y=g+t*b;var z={x:a[0],y:a[1],r:a[2]/2,innerR:a[3]/2,start:Math.round(1E3*x)/1E3,end:Math.round(1E3*y)/1E3};w.shapeType="arc";w.shapeArgs=z;w.labelDistance=u(w.options.dataLabels&&
w.options.dataLabels.distance,m);w.labelDistance=p(w.labelDistance,z.r);this.maxLabelDistance=Math.max(this.maxLabelDistance||0,w.labelDistance);y=(y+x)/2;y>1.5*Math.PI?y-=2*Math.PI:y<-Math.PI/2&&(y+=2*Math.PI);w.slicedTranslation={translateX:Math.round(Math.cos(y)*d),translateY:Math.round(Math.sin(y)*d)};z=Math.cos(y)*a[2]/2;var A=Math.sin(y)*a[2]/2;w.tooltipPos=[a[0]+.7*z,a[1]+.7*A];w.half=y<-Math.PI/2||y>Math.PI/2?1:0;w.angle=y;x=Math.min(f,w.labelDistance/5);w.labelPosition={natural:{x:a[0]+z+
Math.cos(y)*w.labelDistance,y:a[1]+A+Math.sin(y)*w.labelDistance},"final":{},alignment:0>w.labelDistance?"center":w.half?"right":"left",connectorPosition:{breakAt:{x:a[0]+z+Math.cos(y)*x,y:a[1]+A+Math.sin(y)*x},touchingSliceAt:{x:a[0]+z,y:a[1]+A}}}}l(this,"afterTranslate")};c.prototype.updateTotals=function(){var a=this.points,c=a.length,d=this.options.ignoreHiddenPoint,f,b=0;for(f=0;f<c;f++){var g=a[f];!g.isValid()||d&&!g.visible||(b+=g.y)}this.total=b;for(f=0;f<c;f++)g=a[f],g.percentage=0<b&&(g.visible||
!d)?g.y/b*100:0,g.total=b};c.defaultOptions=f(C.defaultOptions,{center:[null,null],clip:!1,colorByPoint:!0,dataLabels:{allowOverlap:!0,connectorPadding:5,connectorShape:"fixedOffset",crookDistance:"70%",distance:30,enabled:!0,formatter:function(){return this.point.isNull?void 0:this.point.name},softConnector:!0,x:0},fillColor:void 0,ignoreHiddenPoint:!0,inactiveOtherPoints:!0,legendType:"point",marker:null,size:null,showInLegend:!1,slicedOffset:10,stickyTracking:!1,tooltip:{followPointer:!0},borderColor:"#ffffff",
borderWidth:1,lineWidth:void 0,states:{hover:{brightness:.1}}});return c}(C);c(q.prototype,{axisTypes:[],directTouch:!0,drawGraph:void 0,drawLegendSymbol:G.drawRectangle,drawTracker:t.prototype.drawTracker,getCenter:a.getCenter,getSymbol:A,isCartesian:!1,noSharedTooltip:!0,pointAttribs:t.prototype.pointAttribs,pointClass:x,requireSorting:!1,searchPoint:A,trackerGroups:["group","dataLabelsGroup"]});I.registerSeriesType("pie",q);"";return q});L(a,"Series/Pie/PieDataLabel.js",[a["Core/Series/DataLabel.js"],
a["Core/Globals.js"],a["Core/Renderer/RendererUtilities.js"],a["Core/Series/SeriesRegistry.js"],a["Core/Utilities.js"]],function(a,t,A,G,x){var u=t.noop,I=A.distribute,z=G.series,q=x.arrayMax,m=x.clamp,h=x.defined,d=x.merge,c=x.pick,l=x.relativeLength,f;(function(f){function p(){var a=this,f=a.data,b=a.chart,g=a.options.dataLabels||{},e=g.connectorPadding,l=b.plotWidth,m=b.plotHeight,p=b.plotLeft,u=Math.round(b.chartWidth/3),r=a.center,t=r[2]/2,w=r[1],B=[[],[]],x=[0,0,0,0],y=a.dataLabelPositioners,
A,C,G,K,L,F,T,N,V,W,X,R;a.visible&&(g.enabled||a._hasPointLabels)&&(f.forEach(function(a){a.dataLabel&&a.visible&&a.dataLabel.shortened&&(a.dataLabel.attr({width:"auto"}).css({width:"auto",textOverflow:"clip"}),a.dataLabel.shortened=!1)}),z.prototype.drawDataLabels.apply(a),f.forEach(function(a){a.dataLabel&&(a.visible?(B[a.half].push(a),a.dataLabel._pos=null,!h(g.style.width)&&!h(a.options.dataLabels&&a.options.dataLabels.style&&a.options.dataLabels.style.width)&&a.dataLabel.getBBox().width>u&&(a.dataLabel.css({width:Math.round(.7*
u)+"px"}),a.dataLabel.shortened=!0)):(a.dataLabel=a.dataLabel.destroy(),a.dataLabels&&1===a.dataLabels.length&&delete a.dataLabels))}),B.forEach(function(d,f){var k=d.length,n=[],q;if(k){a.sortByAngle(d,f-.5);if(0<a.maxLabelDistance){var u=Math.max(0,w-t-a.maxLabelDistance);var v=Math.min(w+t+a.maxLabelDistance,b.plotHeight);d.forEach(function(a){0<a.labelDistance&&a.dataLabel&&(a.top=Math.max(0,w-t-a.labelDistance),a.bottom=Math.min(w+t+a.labelDistance,b.plotHeight),q=a.dataLabel.getBBox().height||
21,a.distributeBox={target:a.labelPosition.natural.y-a.top+q/2,size:q,rank:a.y},n.push(a.distributeBox))});u=v+q-u;I(n,u,u/5)}for(X=0;X<k;X++){A=d[X];F=A.labelPosition;K=A.dataLabel;W=!1===A.visible?"hidden":"inherit";V=u=F.natural.y;n&&h(A.distributeBox)&&("undefined"===typeof A.distributeBox.pos?W="hidden":(T=A.distributeBox.size,V=y.radialDistributionY(A)));delete A.positionIndex;if(g.justify)N=y.justify(A,t,r);else switch(g.alignTo){case "connectors":N=y.alignToConnectors(d,f,l,p);break;case "plotEdges":N=
y.alignToPlotEdges(K,f,l,p);break;default:N=y.radialDistributionX(a,A,V,u)}K._attr={visibility:W,align:F.alignment};R=A.options.dataLabels||{};K._pos={x:N+c(R.x,g.x)+({left:e,right:-e}[F.alignment]||0),y:V+c(R.y,g.y)-10};F.final.x=N;F.final.y=V;c(g.crop,!0)&&(L=K.getBBox().width,u=null,N-L<e&&1===f?(u=Math.round(L-N+e),x[3]=Math.max(u,x[3])):N+L>l-e&&0===f&&(u=Math.round(N+L-l+e),x[1]=Math.max(u,x[1])),0>V-T/2?x[0]=Math.max(Math.round(-V+T/2),x[0]):V+T/2>m&&(x[2]=Math.max(Math.round(V+T/2-m),x[2])),
K.sideOverflow=u)}}}),0===q(x)||this.verifyDataLabelOverflow(x))&&(this.placeDataLabels(),this.points.forEach(function(e){R=d(g,e.options.dataLabels);if(C=c(R.connectorWidth,1)){var f;G=e.connector;if((K=e.dataLabel)&&K._pos&&e.visible&&0<e.labelDistance){W=K._attr.visibility;if(f=!G)e.connector=G=b.renderer.path().addClass("highcharts-data-label-connector  highcharts-color-"+e.colorIndex+(e.className?" "+e.className:"")).add(a.dataLabelsGroup),b.styledMode||G.attr({"stroke-width":C,stroke:R.connectorColor||
e.color||"#666666"});G[f?"attr":"animate"]({d:e.getConnectorPath()});G.attr("visibility",W)}else G&&(e.connector=G.destroy())}}))}function t(){this.points.forEach(function(a){var c=a.dataLabel,b;c&&a.visible&&((b=c._pos)?(c.sideOverflow&&(c._attr.width=Math.max(c.getBBox().width-c.sideOverflow,0),c.css({width:c._attr.width+"px",textOverflow:(this.options.dataLabels.style||{}).textOverflow||"ellipsis"}),c.shortened=!0),c.attr(c._attr),c[c.moved?"animate":"attr"](b),c.moved=!0):c&&c.attr({y:-9999}));
delete a.distributeBox},this)}function r(a){var c=this.center,b=this.options,d=b.center,e=b.minSize||80,f=null!==b.size;if(!f){if(null!==d[0])var h=Math.max(c[2]-Math.max(a[1],a[3]),e);else h=Math.max(c[2]-a[1]-a[3],e),c[0]+=(a[3]-a[1])/2;null!==d[1]?h=m(h,e,c[2]-Math.max(a[0],a[2])):(h=m(h,e,c[2]-a[0]-a[2]),c[1]+=(a[0]-a[2])/2);h<c[2]?(c[2]=h,c[3]=Math.min(l(b.innerSize||0,h),h),this.translate(c),this.drawDataLabels&&this.drawDataLabels()):f=!0}return f}var w=[],B={radialDistributionY:function(a){return a.top+
a.distributeBox.pos},radialDistributionX:function(a,c,b,d){return a.getX(b<c.top+2||b>c.bottom-2?d:b,c.half,c)},justify:function(a,c,b){return b[0]+(a.half?-1:1)*(c+a.labelDistance)},alignToPlotEdges:function(a,c,b,d){a=a.getBBox().width;return c?a+d:b-a-d},alignToConnectors:function(a,c,b,d){var e=0,f;a.forEach(function(a){f=a.dataLabel.getBBox().width;f>e&&(e=f)});return c?e+d:b-e-d}};f.compose=function(c){a.compose(z);-1===w.indexOf(c)&&(w.push(c),c=c.prototype,c.dataLabelPositioners=B,c.alignDataLabel=
u,c.drawDataLabels=p,c.placeDataLabels=t,c.verifyDataLabelOverflow=r)}})(f||(f={}));return f});L(a,"Extensions/OverlappingDataLabels.js",[a["Core/Chart/Chart.js"],a["Core/Utilities.js"]],function(a,t){function u(a,h){var d=!1;if(a){var c=a.newOpacity;a.oldOpacity!==c&&(a.alignAttr&&a.placed?(a[c?"removeClass":"addClass"]("highcharts-data-label-hidden"),d=!0,a.alignAttr.opacity=c,a[a.isOld?"animate":"attr"](a.alignAttr,null,function(){h.styledMode||a.css({pointerEvents:c?"auto":"none"})}),x(h,"afterHideOverlappingLabel")):
a.attr({opacity:c}));a.isOld=!0}return d}var G=t.addEvent,x=t.fireEvent,C=t.isArray,I=t.isNumber,z=t.objectEach,q=t.pick;G(a,"render",function(){var a=this,h=[];(this.labelCollectors||[]).forEach(function(a){h=h.concat(a())});(this.yAxis||[]).forEach(function(a){a.stacking&&a.options.stackLabels&&!a.options.stackLabels.allowOverlap&&z(a.stacking.stacks,function(a){z(a,function(a){a.label&&"hidden"!==a.label.visibility&&h.push(a.label)})})});(this.series||[]).forEach(function(d){var c=d.options.dataLabels;
d.visible&&(!1!==c.enabled||d._hasPointLabels)&&(c=function(c){return c.forEach(function(c){c.visible&&(C(c.dataLabels)?c.dataLabels:c.dataLabel?[c.dataLabel]:[]).forEach(function(d){var f=d.options;d.labelrank=q(f.labelrank,c.labelrank,c.shapeArgs&&c.shapeArgs.height);f.allowOverlap?(d.oldOpacity=d.opacity,d.newOpacity=1,u(d,a)):h.push(d)})})},c(d.nodes||[]),c(d.points))});this.hideOverlappingLabels(h)});a.prototype.hideOverlappingLabels=function(a){var h=this,d=a.length,c=h.renderer,l,f,m,p=!1;
var q=function(a){var d,f=a.box?0:a.padding||0,b=d=0,g;if(a&&(!a.alignAttr||a.placed)){var e=a.alignAttr||{x:a.attr("x"),y:a.attr("y")};var h=a.parentGroup;a.width||(d=a.getBBox(),a.width=d.width,a.height=d.height,d=c.fontMetrics(null,a.element).h);var l=a.width-2*f;(g={left:"0",center:"0.5",right:"1"}[a.alignValue])?b=+g*l:I(a.x)&&Math.round(a.x)!==a.translateX&&(b=a.x-a.translateX);return{x:e.x+(h.translateX||0)+f-(b||0),y:e.y+(h.translateY||0)+f-d,width:a.width-2*f,height:a.height-2*f}}};for(f=
0;f<d;f++)if(l=a[f])l.oldOpacity=l.opacity,l.newOpacity=1,l.absoluteBox=q(l);a.sort(function(a,c){return(c.labelrank||0)-(a.labelrank||0)});for(f=0;f<d;f++){var r=(q=a[f])&&q.absoluteBox;for(l=f+1;l<d;++l){var t=(m=a[l])&&m.absoluteBox;!r||!t||q===m||0===q.newOpacity||0===m.newOpacity||t.x>=r.x+r.width||t.x+t.width<=r.x||t.y>=r.y+r.height||t.y+t.height<=r.y||((q.labelrank<m.labelrank?q:m).newOpacity=0)}}a.forEach(function(a){u(a,h)&&(p=!0)});p&&x(h,"afterHideAllOverlappingLabels")}});L(a,"Core/Responsive.js",
[a["Core/Utilities.js"]],function(a){var u=a.extend,A=a.find,G=a.isArray,x=a.isObject,C=a.merge,I=a.objectEach,z=a.pick,q=a.splat,m=a.uniqueKey,h;(function(a){var c=[];a.compose=function(a){-1===c.indexOf(a)&&(c.push(a),u(a.prototype,d.prototype));return a};var d=function(){function a(){}a.prototype.currentOptions=function(a){function c(a,f,h,k){var b;I(a,function(a,e){if(!k&&-1<d.collectionsWithUpdate.indexOf(e)&&f[e])for(a=q(a),h[e]=[],b=0;b<Math.max(a.length,f[e].length);b++)f[e][b]&&(void 0===
a[b]?h[e][b]=f[e][b]:(h[e][b]={},c(a[b],f[e][b],h[e][b],k+1)));else x(a)?(h[e]=G(a)?[]:{},c(a,f[e]||{},h[e],k+1)):h[e]="undefined"===typeof f[e]?null:f[e]})}var d=this,f={};c(a,this.options,f,0);return f};a.prototype.matchResponsiveRule=function(a,c){var d=a.condition;(d.callback||function(){return this.chartWidth<=z(d.maxWidth,Number.MAX_VALUE)&&this.chartHeight<=z(d.maxHeight,Number.MAX_VALUE)&&this.chartWidth>=z(d.minWidth,0)&&this.chartHeight>=z(d.minHeight,0)}).call(this)&&c.push(a._id)};a.prototype.setResponsive=
function(a,c){var d=this,f=this.options.responsive,h=this.currentResponsive,l=[];!c&&f&&f.rules&&f.rules.forEach(function(a){"undefined"===typeof a._id&&(a._id=m());d.matchResponsiveRule(a,l)},this);c=C.apply(void 0,l.map(function(a){return A((f||{}).rules||[],function(c){return c._id===a})}).map(function(a){return a&&a.chartOptions}));c.isResponsiveOptions=!0;l=l.toString()||void 0;l!==(h&&h.ruleIds)&&(h&&this.update(h.undoOptions,a,!0),l?(h=this.currentOptions(c),h.isResponsiveOptions=!0,this.currentResponsive=
{ruleIds:l,mergedOptions:c,undoOptions:h},this.update(c,a,!0)):this.currentResponsive=void 0)};return a}()})(h||(h={}));"";"";return h});L(a,"masters/highcharts.src.js",[a["Core/Globals.js"],a["Core/Utilities.js"],a["Core/DefaultOptions.js"],a["Core/Animation/Fx.js"],a["Core/Animation/AnimationUtilities.js"],a["Core/Renderer/HTML/AST.js"],a["Core/FormatUtilities.js"],a["Core/Renderer/RendererUtilities.js"],a["Core/Renderer/SVG/SVGElement.js"],a["Core/Renderer/SVG/SVGRenderer.js"],a["Core/Renderer/HTML/HTMLElement.js"],
a["Core/Renderer/HTML/HTMLRenderer.js"],a["Core/Axis/Axis.js"],a["Core/Axis/DateTimeAxis.js"],a["Core/Axis/LogarithmicAxis.js"],a["Core/Axis/PlotLineOrBand/PlotLineOrBand.js"],a["Core/Axis/Tick.js"],a["Core/Tooltip.js"],a["Core/Series/Point.js"],a["Core/Pointer.js"],a["Core/MSPointer.js"],a["Core/Legend/Legend.js"],a["Core/Chart/Chart.js"],a["Core/Series/Series.js"],a["Core/Series/SeriesRegistry.js"],a["Series/Column/ColumnSeries.js"],a["Series/Column/ColumnDataLabel.js"],a["Series/Pie/PieSeries.js"],
a["Series/Pie/PieDataLabel.js"],a["Core/Series/DataLabel.js"],a["Core/Responsive.js"],a["Core/Color/Color.js"],a["Core/Time.js"]],function(a,t,A,G,x,C,I,z,q,m,h,d,c,l,f,w,p,K,r,y,B,n,k,b,g,e,D,H,v,E,L,S,P){a.animate=x.animate;a.animObject=x.animObject;a.getDeferredAnimation=x.getDeferredAnimation;a.setAnimation=x.setAnimation;a.stop=x.stop;a.timers=G.timers;a.AST=C;a.Axis=c;a.Chart=k;a.chart=k.chart;a.Fx=G;a.Legend=n;a.PlotLineOrBand=w;a.Point=r;a.Pointer=B.isRequired()?B:y;a.Series=b;a.SVGElement=
q;a.SVGRenderer=m;a.Tick=p;a.Time=P;a.Tooltip=K;a.Color=S;a.color=S.parse;d.compose(m);h.compose(q);a.defaultOptions=A.defaultOptions;a.getOptions=A.getOptions;a.time=A.defaultTime;a.setOptions=A.setOptions;a.dateFormat=I.dateFormat;a.format=I.format;a.numberFormat=I.numberFormat;a.addEvent=t.addEvent;a.arrayMax=t.arrayMax;a.arrayMin=t.arrayMin;a.attr=t.attr;a.clearTimeout=t.clearTimeout;a.correctFloat=t.correctFloat;a.createElement=t.createElement;a.css=t.css;a.defined=t.defined;a.destroyObjectProperties=
t.destroyObjectProperties;a.discardElement=t.discardElement;a.distribute=z.distribute;a.erase=t.erase;a.error=t.error;a.extend=t.extend;a.extendClass=t.extendClass;a.find=t.find;a.fireEvent=t.fireEvent;a.getMagnitude=t.getMagnitude;a.getStyle=t.getStyle;a.inArray=t.inArray;a.isArray=t.isArray;a.isClass=t.isClass;a.isDOMElement=t.isDOMElement;a.isFunction=t.isFunction;a.isNumber=t.isNumber;a.isObject=t.isObject;a.isString=t.isString;a.keys=t.keys;a.merge=t.merge;a.normalizeTickInterval=t.normalizeTickInterval;
a.objectEach=t.objectEach;a.offset=t.offset;a.pad=t.pad;a.pick=t.pick;a.pInt=t.pInt;a.relativeLength=t.relativeLength;a.removeEvent=t.removeEvent;a.seriesType=g.seriesType;a.splat=t.splat;a.stableSort=t.stableSort;a.syncTimeout=t.syncTimeout;a.timeUnits=t.timeUnits;a.uniqueKey=t.uniqueKey;a.useSerialIds=t.useSerialIds;a.wrap=t.wrap;D.compose(e);E.compose(b);l.compose(c);f.compose(c);v.compose(H);w.compose(c);L.compose(k);return a});a["masters/highcharts.src.js"]._modules=a;return a["masters/highcharts.src.js"]});
//# sourceMappingURL=highcharts.js.map
```
