## Supplementary material for "Exploring natural odour landscapes: A case study with implications for human-biting insects": Figure S4: accessibility.js

```
/*
 Highcharts JS v9.3.1 (2021-11-05)

 Accessibility module

 (c) 2010-2021 Highsoft AS
 Author: Oystein Moseng

 License: www.highcharts.com/license
*/
'use strict';(function(a){"object"===typeof module&&module.exports?(a["default"]=a,module.exports=a):"function"===typeof define&&define.amd?define("highcharts/modules/accessibility",["highcharts"],function(v){a(v);a.Highcharts=v;return a}):a("undefined"!==typeof Highcharts?Highcharts:void 0)})(function(a){function v(a,h,m,u){a.hasOwnProperty(h)||(a[h]=u.apply(null,m))}a=a?a._modules:{};v(a,"Accessibility/A11yI18n.js",[a["Core/FormatUtilities.js"],a["Core/Utilities.js"]],function(a,h){var m=a.format,
u=h.pick,n;(function(a){function n(g,c){var e=g.indexOf("#each("),d=g.indexOf("#plural("),b=g.indexOf("["),f=g.indexOf("]");if(-1<e){f=g.slice(e).indexOf(")")+e;d=g.substring(0,e);b=g.substring(f+1);f=g.substring(e+6,f).split(",");e=Number(f[1]);g="";if(c=c[f[0]])for(e=isNaN(e)?c.length:e,e=0>e?c.length+e:Math.min(e,c.length),f=0;f<e;++f)g+=d+c[f]+b;return g.length?g:""}if(-1<d){b=g.slice(d).indexOf(")")+d;d=g.substring(d+8,b).split(",");switch(Number(c[d[0]])){case 0:g=u(d[4],d[1]);break;case 1:g=
u(d[2],d[1]);break;case 2:g=u(d[3],d[1]);break;default:g=d[1]}g?(c=g,c=c.trim&&c.trim()||c.replace(/^\s+|\s+$/g,"")):c="";return c}return-1<b?(d=g.substring(0,b),b=Number(g.substring(b+1,f)),g=void 0,c=c[d],!isNaN(b)&&c&&(0>b?(g=c[c.length+b],"undefined"===typeof g&&(g=c[0])):(g=c[b],"undefined"===typeof g&&(g=c[c.length-1]))),"undefined"!==typeof g?g:""):"{"+g+"}"}function t(g,c,e){var d=function(b,f){b=b.slice(f||0);var d=b.indexOf("{"),c=b.indexOf("}");if(-1<d&&c>d)return{statement:b.substring(d+
1,c),begin:f+d+1,end:f+c}},b=[],f=0;do{var A=d(g,f);var a=g.substring(f,A&&A.begin-1);a.length&&b.push({value:a,type:"constant"});A&&b.push({value:A.statement,type:"statement"});f=A?A.end+1:f+1}while(A);b.forEach(function(b){"statement"===b.type&&(b.value=n(b.value,c))});return m(b.reduce(function(b,f){return b+f.value},""),c,e)}function k(g,c){g=g.split(".");for(var e=this.options.lang,d=0;d<g.length;++d)e=e&&e[g[d]];return"string"===typeof e?t(e,c,this):""}var l=[];a.compose=function(g){-1===l.indexOf(g)&&
(l.push(g),g.prototype.langFormat=k);return g};a.i18nFormat=t})(n||(n={}));return n});v(a,"Accessibility/Utils/HTMLUtilities.js",[a["Core/Globals.js"],a["Core/Utilities.js"]],function(a,h){function m(a){if("function"===typeof p.MouseEvent)return new p.MouseEvent(a.type,a);if(n.createEvent){var t=n.createEvent("MouseEvent");if(t.initMouseEvent)return t.initMouseEvent(a.type,a.bubbles,a.cancelable,a.view||p,a.detail,a.screenX,a.screenY,a.clientX,a.clientY,a.ctrlKey,a.altKey,a.shiftKey,a.metaKey,a.button,
a.relatedTarget),t}return u(a.type)}function u(a,k){k=k||{x:0,y:0};if("function"===typeof p.MouseEvent)return new p.MouseEvent(a,{bubbles:!0,cancelable:!0,composed:!0,view:p,detail:"click"===a?1:0,screenX:k.x,screenY:k.y,clientX:k.x,clientY:k.y});if(n.createEvent){var l=n.createEvent("MouseEvent");if(l.initMouseEvent)return l.initMouseEvent(a,!0,!0,p,"click"===a?1:0,k.x,k.y,k.x,k.y,!1,!1,!1,!1,0,null),l}return{type:a}}var n=a.doc,p=a.win,w=h.css;return{addClass:function(a,k){a.classList?a.classList.add(k):
0>a.className.indexOf(k)&&(a.className+=" "+k)},cloneMouseEvent:m,cloneTouchEvent:function(a){var k=function(a){for(var g=[],c=0;c<a.length;++c){var e=a.item(c);e&&g.push(e)}return g};if("function"===typeof p.TouchEvent)return k=new p.TouchEvent(a.type,{touches:k(a.touches),targetTouches:k(a.targetTouches),changedTouches:k(a.changedTouches),ctrlKey:a.ctrlKey,shiftKey:a.shiftKey,altKey:a.altKey,metaKey:a.metaKey,bubbles:a.bubbles,cancelable:a.cancelable,composed:a.composed,detail:a.detail,view:a.view}),
a.defaultPrevented&&k.preventDefault(),k;k=m(a);k.touches=a.touches;k.changedTouches=a.changedTouches;k.targetTouches=a.targetTouches;return k},escapeStringForHTML:function(a){return a.replace(/&/g,"&amp;").replace(/</g,"&lt;").replace(/>/g,"&gt;").replace(/"/g,"&quot;").replace(/'/g,"&#x27;").replace(/\//g,"&#x2F;")},getElement:function(a){return n.getElementById(a)},getFakeMouseEvent:u,getHeadingTagNameForElement:function(a){var k=function(a){a=parseInt(a.slice(1),10);return"h"+Math.min(6,a+1)},
l=function(a){var c;a:{for(c=a;c=c.previousSibling;){var e=c.tagName||"";if(/H[1-6]/.test(e)){c=e;break a}}c=""}if(c)return k(c);a=a.parentElement;if(!a)return"p";c=a.tagName;return/H[1-6]/.test(c)?k(c):l(a)};return l(a)},removeChildNodes:function(a){for(;a.lastChild;)a.removeChild(a.lastChild)},removeClass:function(a,k){a.classList?a.classList.remove(k):a.className=a.className.replace(new RegExp(k,"g"),"")},removeElement:function(a){a&&a.parentNode&&a.parentNode.removeChild(a)},reverseChildNodes:function(a){for(var k=
a.childNodes.length;k--;)a.appendChild(a.childNodes[k])},stripHTMLTagsFromString:function(a){return"string"===typeof a?a.replace(/<\/?[^>]+(>|$)/g,""):a},visuallyHideElement:function(a){w(a,{position:"absolute",width:"1px",height:"1px",overflow:"hidden",whiteSpace:"nowrap",clip:"rect(1px, 1px, 1px, 1px)",marginTop:"-3px","-ms-filter":"progid:DXImageTransform.Microsoft.Alpha(Opacity=1)",filter:"alpha(opacity=1)",opacity:.01})}}});v(a,"Accessibility/Utils/ChartUtilities.js",[a["Core/Globals.js"],a["Accessibility/Utils/HTMLUtilities.js"],
a["Core/Utilities.js"]],function(a,h,m){function u(b,f){var c=f.type,a=b.hcEvents;l.createEvent&&(b.dispatchEvent||b.fireEvent)?b.dispatchEvent?b.dispatchEvent(f):b.fireEvent(c,f):a&&a[c]?d(b,c,f):b.element&&u(b.element,f)}function n(b){var f=b.chart,d={},c="Seconds";d.Seconds=((b.max||0)-(b.min||0))/1E3;d.Minutes=d.Seconds/60;d.Hours=d.Minutes/60;d.Days=d.Hours/24;["Minutes","Hours","Days"].forEach(function(b){2<d[b]&&(c=b)});var a=d[c].toFixed("Seconds"!==c&&"Minutes"!==c?1:0);return f.langFormat("accessibility.axis.timeRange"+
c,{chart:f,axis:b,range:a.replace(".0","")})}function p(b){var d=b.chart,c=d.options&&d.options.accessibility&&d.options.accessibility.screenReaderSection.axisRangeDateFormat||"",a=function(f){return b.dateTime?d.time.dateFormat(c,b[f]):b[f]};return d.langFormat("accessibility.axis.rangeFromTo",{chart:d,axis:b,rangeFrom:a("min"),rangeTo:a("max")})}function w(b){if(b.points&&b.points.length)return(b=e(b.points,function(b){return!!b.graphic}))&&b.graphic&&b.graphic.element}function t(b){var d=w(b);
return d&&d.parentNode||b.graph&&b.graph.element||b.group&&b.group.element}function k(b,d){d.setAttribute("aria-hidden",!1);d!==b.renderTo&&d.parentNode&&d.parentNode!==l.body&&(Array.prototype.forEach.call(d.parentNode.childNodes,function(b){b.hasAttribute("aria-hidden")||b.setAttribute("aria-hidden",!0)}),k(b,d.parentNode))}var l=a.doc,g=h.stripHTMLTagsFromString,c=m.defined,e=m.find,d=m.fireEvent;return{fireEventOnWrappedOrUnwrappedElement:u,getChartTitle:function(b){return g(b.options.title.text||
b.langFormat("accessibility.defaultChartTitle",{chart:b}))},getAxisDescription:function(b){return b&&(b.userOptions&&b.userOptions.accessibility&&b.userOptions.accessibility.description||b.axisTitle&&b.axisTitle.textStr||b.options.id||b.categories&&"categories"||b.dateTime&&"Time"||"values")},getAxisRangeDescription:function(b){var d=b.options||{};return d.accessibility&&"undefined"!==typeof d.accessibility.rangeDescription?d.accessibility.rangeDescription:b.categories?(d=b.chart,b=b.dataMax&&b.dataMin?
d.langFormat("accessibility.axis.rangeCategories",{chart:d,axis:b,numCategories:b.dataMax-b.dataMin+1}):"",b):!b.dateTime||0!==b.min&&0!==b.dataMin?p(b):n(b)},getPointFromXY:function(b,d,c){for(var f=b.length,a;f--;)if(a=e(b[f].points||[],function(b){return b.x===d&&b.y===c}))return a},getSeriesFirstPointElement:w,getSeriesFromName:function(b,d){return d?(b.series||[]).filter(function(b){return b.name===d}):b.series},getSeriesA11yElement:t,unhideChartElementFromAT:k,hideSeriesFromAT:function(b){(b=
t(b))&&b.setAttribute("aria-hidden",!0)},scrollToPoint:function(b){var f=b.series.xAxis,a=b.series.yAxis,e=f&&f.scrollbar?f:a;if((f=e&&e.scrollbar)&&c(f.to)&&c(f.from)){a=f.to-f.from;if(c(e.dataMin)&&c(e.dataMax)){var g=e.toPixels(e.dataMin),l=e.toPixels(e.dataMax);b=(e.toPixels(b["xAxis"===e.coll?"x":"y"]||0)-g)/(l-g)}else b=0;f.updatePosition(b-a/2,b+a/2);d(f,"changed",{from:f.from,to:f.to,trigger:"scrollbar",DOMEvent:null})}}}});v(a,"Accessibility/Utils/DOMElementProvider.js",[a["Core/Globals.js"],
a["Accessibility/Utils/HTMLUtilities.js"]],function(a,h){var m=a.doc,u=h.removeElement;return function(){function a(){this.elements=[]}a.prototype.createElement=function(){var a=m.createElement.apply(m,arguments);this.elements.push(a);return a};a.prototype.destroyCreatedElements=function(){this.elements.forEach(function(a){u(a)});this.elements=[]};return a}()});v(a,"Accessibility/Utils/EventProvider.js",[a["Core/Globals.js"],a["Core/Utilities.js"]],function(a,h){var m=h.addEvent;return function(){function h(){this.eventRemovers=
[]}h.prototype.addEvent=function(){var h=m.apply(a,arguments);this.eventRemovers.push(h);return h};h.prototype.removeAddedEvents=function(){this.eventRemovers.forEach(function(a){return a()});this.eventRemovers=[]};return h}()});v(a,"Accessibility/AccessibilityComponent.js",[a["Accessibility/Utils/ChartUtilities.js"],a["Accessibility/Utils/DOMElementProvider.js"],a["Accessibility/Utils/EventProvider.js"],a["Accessibility/Utils/HTMLUtilities.js"],a["Core/Utilities.js"]],function(a,h,m,u,n){var p=a.fireEventOnWrappedOrUnwrappedElement,
w=u.getFakeMouseEvent;a=n.extend;u=function(){function a(){this.proxyProvider=this.keyCodes=this.eventProvider=this.domElementProvider=this.chart=void 0}a.prototype.initBase=function(a,l){this.chart=a;this.eventProvider=new m;this.domElementProvider=new h;this.proxyProvider=l;this.keyCodes={left:37,right:39,up:38,down:40,enter:13,space:32,esc:27,tab:9,pageUp:33,pageDown:34,end:35,home:36}};a.prototype.addEvent=function(a,l,g,c){return this.eventProvider.addEvent(a,l,g,c)};a.prototype.createElement=
function(a,l){return this.domElementProvider.createElement(a,l)};a.prototype.fakeClickEvent=function(a){var l=w("click");p(a,l)};a.prototype.destroyBase=function(){this.domElementProvider.destroyCreatedElements();this.eventProvider.removeAddedEvents()};return a}();a(u.prototype,{init:function(){},getKeyboardNavigation:function(){},onChartUpdate:function(){},onChartRender:function(){},destroy:function(){}});return u});v(a,"Accessibility/KeyboardNavigationHandler.js",[a["Core/Utilities.js"]],function(a){var h=
a.find;a=function(){function a(a,h){this.chart=a;this.keyCodeMap=h.keyCodeMap||[];this.validate=h.validate;this.init=h.init;this.terminate=h.terminate;this.response={success:1,prev:2,next:3,noHandler:4,fail:5}}a.prototype.run=function(a){var n=a.which||a.keyCode,p=this.response.noHandler,m=h(this.keyCodeMap,function(a){return-1<a[0].indexOf(n)});m?p=m[1].call(this,n,a):9===n&&(p=this.response[a.shiftKey?"prev":"next"]);return p};return a}();"";return a});v(a,"Accessibility/Components/ContainerComponent.js",
[a["Accessibility/AccessibilityComponent.js"],a["Accessibility/KeyboardNavigationHandler.js"],a["Accessibility/Utils/ChartUtilities.js"],a["Core/Globals.js"],a["Accessibility/Utils/HTMLUtilities.js"]],function(a,h,m,u,n){var p=this&&this.__extends||function(){var a=function(c,e){a=Object.setPrototypeOf||{__proto__:[]}instanceof Array&&function(d,b){d.__proto__=b}||function(d,b){for(var a in b)b.hasOwnProperty(a)&&(d[a]=b[a])};return a(c,e)};return function(c,e){function d(){this.constructor=c}a(c,
e);c.prototype=null===e?Object.create(e):(d.prototype=e.prototype,new d)}}(),w=m.unhideChartElementFromAT,t=m.getChartTitle,k=u.doc,l=n.stripHTMLTagsFromString;return function(a){function c(){return null!==a&&a.apply(this,arguments)||this}p(c,a);c.prototype.onChartUpdate=function(){this.handleSVGTitleElement();this.setSVGContainerLabel();this.setGraphicContainerAttrs();this.setRenderToAttrs();this.makeCreditsAccessible()};c.prototype.handleSVGTitleElement=function(){var a=this.chart,d="highcharts-title-"+
a.index,b=l(a.langFormat("accessibility.svgContainerTitle",{chartTitle:t(a)}));if(b.length){var f=this.svgTitleElement=this.svgTitleElement||k.createElementNS("http://www.w3.org/2000/svg","title");f.textContent=b;f.id=d;a.renderTo.insertBefore(f,a.renderTo.firstChild)}};c.prototype.setSVGContainerLabel=function(){var a=this.chart,d=a.langFormat("accessibility.svgContainerLabel",{chartTitle:t(a)});a.renderer.box&&d.length&&a.renderer.box.setAttribute("aria-label",d)};c.prototype.setGraphicContainerAttrs=
function(){var a=this.chart,d=a.langFormat("accessibility.graphicContainerLabel",{chartTitle:t(a)});d.length&&a.container.setAttribute("aria-label",d)};c.prototype.setRenderToAttrs=function(){var a=this.chart;"disabled"!==a.options.accessibility.landmarkVerbosity?a.renderTo.setAttribute("role","region"):a.renderTo.removeAttribute("role");a.renderTo.setAttribute("aria-label",a.langFormat("accessibility.chartContainerLabel",{title:t(a),chart:a}))};c.prototype.makeCreditsAccessible=function(){var a=
this.chart,d=a.credits;d&&(d.textStr&&d.element.setAttribute("aria-label",a.langFormat("accessibility.credits",{creditsStr:l(d.textStr)})),w(a,d.element))};c.prototype.getKeyboardNavigation=function(){var a=this.chart;return new h(a,{keyCodeMap:[],validate:function(){return!0},init:function(){var d=a.accessibility;d&&d.keyboardNavigation.tabindexContainer.focus()}})};c.prototype.destroy=function(){this.chart.renderTo.setAttribute("aria-hidden",!0)};return c}(a)});v(a,"Accessibility/FocusBorder.js",
[a["Core/Renderer/SVG/SVGLabel.js"],a["Core/Utilities.js"]],function(a,h){var m=h.addEvent,u=h.pick,n;(function(h){function n(){var b=this.focusElement,a=this.options.accessibility.keyboardNavigation.focusBorder;b&&(b.removeFocusBorder(),a.enabled&&b.addFocusBorder(a.margin,{stroke:a.style.color,strokeWidth:a.style.lineWidth,r:a.style.borderRadius}))}function t(b,a){var d=this.options.accessibility.keyboardNavigation.focusBorder;(a=a||b.element)&&a.focus&&(a.hcEvents&&a.hcEvents.focusin||m(a,"focusin",
function(){}),a.focus(),d.hideBrowserFocusOutline&&(a.style.outline="none"));this.focusElement&&this.focusElement.removeFocusBorder();this.focusElement=b;this.renderFocusBorder()}function k(b){if(!b.focusBorderDestroyHook){var a=b.destroy;b.destroy=function(){b.focusBorder&&b.focusBorder.destroy&&b.focusBorder.destroy();return a.apply(b,arguments)};b.focusBorderDestroyHook=a}}function l(b,d){this.focusBorder&&this.removeFocusBorder();var c=this.getBBox(),f=u(b,3);c.x+=this.translateX?this.translateX:
0;c.y+=this.translateY?this.translateY:0;var e=c.x-f,A=c.y-f,x=c.width+2*f,l=c.height+2*f,h=this instanceof a;if("text"===this.element.nodeName||h){var n=!!this.rotation;if(h)var t={x:n?1:0,y:0};else{var m=t=0;"middle"===this.attr("text-anchor")?t=m=.5:this.rotation?t=.25:m=.75;t={x:t,y:m}}m=+this.attr("x");var q=+this.attr("y");isNaN(m)||(e=m-c.width*t.x-f);isNaN(q)||(A=q-c.height*t.y-f);h&&n&&(h=x,x=l,l=h,isNaN(m)||(e=m-c.height*t.x-f),isNaN(q)||(A=q-c.width*t.y-f))}this.focusBorder=this.renderer.rect(e,
A,x,l,parseInt((d&&d.r||0).toString(),10)).addClass("highcharts-focus-border").attr({zIndex:99}).add(this.parentGroup);this.renderer.styledMode||this.focusBorder.attr({stroke:d&&d.stroke,"stroke-width":d&&d.strokeWidth});g(this,b,d);k(this)}function g(a){for(var d=[],c=1;c<arguments.length;c++)d[c-1]=arguments[c];a.focusBorderUpdateHooks||(a.focusBorderUpdateHooks={},b.forEach(function(b){b+="Setter";var c=a[b]||a._defaultSetter;a.focusBorderUpdateHooks[b]=c;a[b]=function(){var b=c.apply(a,arguments);
a.addFocusBorder.apply(a,d);return b}}))}function c(){e(this);this.focusBorderDestroyHook&&(this.destroy=this.focusBorderDestroyHook,delete this.focusBorderDestroyHook);this.focusBorder&&(this.focusBorder.destroy(),delete this.focusBorder)}function e(b){b.focusBorderUpdateHooks&&(Object.keys(b.focusBorderUpdateHooks).forEach(function(a){var d=b.focusBorderUpdateHooks[a];d===b._defaultSetter?delete b[a]:b[a]=d}),delete b.focusBorderUpdateHooks)}var d=[],b="x y transform width height r d stroke-width".split(" ");
h.compose=function(b,a){-1===d.indexOf(b)&&(d.push(b),b=b.prototype,b.renderFocusBorder=n,b.setFocusToElement=t);-1===d.indexOf(a)&&(d.push(a),a=a.prototype,a.addFocusBorder=l,a.removeFocusBorder=c)}})(n||(n={}));return n});v(a,"Accessibility/Utils/Announcer.js",[a["Core/Renderer/HTML/AST.js"],a["Accessibility/Utils/DOMElementProvider.js"],a["Core/Globals.js"],a["Accessibility/Utils/HTMLUtilities.js"],a["Core/Utilities.js"]],function(a,h,m,u,n){var p=m.doc,w=u.addClass,t=u.visuallyHideElement,k=n.attr;
return function(){function l(a,c){this.chart=a;this.domElementProvider=new h;this.announceRegion=this.addAnnounceRegion(c)}l.prototype.destroy=function(){this.domElementProvider.destroyCreatedElements()};l.prototype.announce=function(g){var c=this;a.setElementHTML(this.announceRegion,g);this.clearAnnouncementRegionTimer&&clearTimeout(this.clearAnnouncementRegionTimer);this.clearAnnouncementRegionTimer=setTimeout(function(){c.announceRegion.innerHTML="";delete c.clearAnnouncementRegionTimer},1E3)};
l.prototype.addAnnounceRegion=function(a){var c=this.chart.announcerContainer||this.createAnnouncerContainer(),e=this.domElementProvider.createElement("div");k(e,{"aria-hidden":!1,"aria-live":a});this.chart.styledMode?w(e,"highcharts-visually-hidden"):t(e);c.appendChild(e);return e};l.prototype.createAnnouncerContainer=function(){var a=this.chart,c=p.createElement("div");k(c,{"aria-hidden":!1,"class":"highcharts-announcer-container"});c.style.position="relative";a.renderTo.insertBefore(c,a.renderTo.firstChild);
return a.announcerContainer=c};return l}()});v(a,"Accessibility/Components/AnnotationsA11y.js",[a["Accessibility/Utils/HTMLUtilities.js"]],function(a){function h(a){return(a.annotations||[]).reduce(function(a,l){l.options&&!1!==l.options.visible&&(a=a.concat(l.labels));return a},[])}function m(a){return a.options&&a.options.accessibility&&a.options.accessibility.description||a.graphic&&a.graphic.text&&a.graphic.text.textStr||""}function u(a){var k=a.options&&a.options.accessibility&&a.options.accessibility.description;
if(k)return k;k=a.chart;var l=m(a),g=a.points.filter(function(a){return!!a.graphic}).map(function(a){var b=a.accessibility&&a.accessibility.valueDescription||a.graphic&&a.graphic.element&&a.graphic.element.getAttribute("aria-label")||"";a=a&&a.series.name||"";return(a?a+", ":"")+"data point "+b}).filter(function(a){return!!a}),c=g.length,e="accessibility.screenReaderSection.annotations.description"+(1<c?"MultiplePoints":c?"SinglePoint":"NoPoints");a={annotationText:l,annotation:a,numPoints:c,annotationPoint:g[0],
additionalAnnotationPoints:g.slice(1)};return k.langFormat(e,a)}function n(a){return h(a).map(function(a){return(a=p(w(u(a))))?"<li>"+a+"</li>":""})}var p=a.escapeStringForHTML,w=a.stripHTMLTagsFromString;return{getAnnotationsInfoHTML:function(a){var k=a.annotations;return k&&k.length?'<ul style="list-style-type: none">'+n(a).join(" ")+"</ul>":""},getAnnotationLabelDescription:u,getAnnotationListItems:n,getPointAnnotationTexts:function(a){var k=h(a.series.chart).filter(function(l){return-1<l.points.indexOf(a)});
return k.length?k.map(function(a){return""+m(a)}):[]}}});v(a,"Accessibility/Components/InfoRegionsComponent.js",[a["Accessibility/A11yI18n.js"],a["Accessibility/AccessibilityComponent.js"],a["Accessibility/Utils/Announcer.js"],a["Accessibility/Components/AnnotationsA11y.js"],a["Core/Renderer/HTML/AST.js"],a["Accessibility/Utils/ChartUtilities.js"],a["Core/FormatUtilities.js"],a["Core/Globals.js"],a["Accessibility/Utils/HTMLUtilities.js"],a["Core/Utilities.js"]],function(a,h,m,u,n,p,E,t,k,l){function g(a,
b){var d=b[0],y=a.series&&a.series[0]||{};y={numSeries:a.series.length,numPoints:y.points&&y.points.length,chart:a,mapTitle:y.mapTitle};if(!d)return a.langFormat("accessibility.chartTypes.emptyChart",y);if("map"===d)return y.mapTitle?a.langFormat("accessibility.chartTypes.mapTypeDescription",y):a.langFormat("accessibility.chartTypes.unknownMap",y);if(1<a.types.length)return a.langFormat("accessibility.chartTypes.combinationChart",y);b=b[0];d=a.langFormat("accessibility.seriesTypeDescriptions."+b,
y);var c=a.series&&2>a.series.length?"Single":"Multiple";return(a.langFormat("accessibility.chartTypes."+b+c,y)||a.langFormat("accessibility.chartTypes.default"+c,y))+(d?" "+d:"")}var c=this&&this.__extends||function(){var a=function(b,d){a=Object.setPrototypeOf||{__proto__:[]}instanceof Array&&function(a,b){a.__proto__=b}||function(a,b){for(var d in b)b.hasOwnProperty(d)&&(a[d]=b[d])};return a(b,d)};return function(b,d){function r(){this.constructor=b}a(b,d);b.prototype=null===d?Object.create(d):
(r.prototype=d.prototype,new r)}}(),e=u.getAnnotationsInfoHTML,d=p.getAxisDescription,b=p.getAxisRangeDescription,f=p.getChartTitle,A=p.unhideChartElementFromAT,z=E.format,J=t.doc,C=k.addClass,F=k.getElement,x=k.getHeadingTagNameForElement,D=k.stripHTMLTagsFromString,w=k.visuallyHideElement,B=l.attr,G=l.pick;return function(l){function q(){var a=null!==l&&l.apply(this,arguments)||this;a.announcer=void 0;a.screenReaderSections={};return a}c(q,l);q.prototype.init=function(){var a=this.chart,b=this;
this.initRegionsDefinitions();this.addEvent(a,"aftergetTableAST",function(a){b.onDataTableCreated(a)});this.addEvent(a,"afterViewData",function(a){b.dataTableDiv=a;setTimeout(function(){b.focusDataTable()},300)});this.announcer=new m(a,"assertive")};q.prototype.initRegionsDefinitions=function(){var a=this;this.screenReaderSections={before:{element:null,buildContent:function(b){var d=b.options.accessibility.screenReaderSection.beforeChartFormatter;return d?d(b):a.defaultBeforeChartFormatter(b)},insertIntoDOM:function(a,
b){b.renderTo.insertBefore(a,b.renderTo.firstChild)},afterInserted:function(){"undefined"!==typeof a.sonifyButtonId&&a.initSonifyButton(a.sonifyButtonId);"undefined"!==typeof a.dataTableButtonId&&a.initDataTableButton(a.dataTableButtonId)}},after:{element:null,buildContent:function(b){var d=b.options.accessibility.screenReaderSection.afterChartFormatter;return d?d(b):a.defaultAfterChartFormatter()},insertIntoDOM:function(a,b){b.renderTo.insertBefore(a,b.container.nextSibling)},afterInserted:function(){a.chart.accessibility&&
a.chart.accessibility.keyboardNavigation.updateExitAnchor()}}}};q.prototype.onChartRender=function(){var a=this;this.linkedDescriptionElement=this.getLinkedDescriptionElement();this.setLinkedDescriptionAttrs();Object.keys(this.screenReaderSections).forEach(function(b){a.updateScreenReaderSection(b)})};q.prototype.getLinkedDescriptionElement=function(){var a=this.chart.options.accessibility.linkedDescription;if(a){if("string"!==typeof a)return a;a=z(a,this.chart);a=J.querySelectorAll(a);if(1===a.length)return a[0]}};
q.prototype.setLinkedDescriptionAttrs=function(){var a=this.linkedDescriptionElement;a&&(a.setAttribute("aria-hidden","true"),C(a,"highcharts-linked-description"))};q.prototype.updateScreenReaderSection=function(a){var b=this.chart,d=this.screenReaderSections[a],c=d.buildContent(b),r=d.element=d.element||this.createElement("div"),f=r.firstChild||this.createElement("div");c?(this.setScreenReaderSectionAttribs(r,a),n.setElementHTML(f,c),r.appendChild(f),d.insertIntoDOM(r,b),b.styledMode?C(f,"highcharts-visually-hidden"):
w(f),A(b,f),d.afterInserted&&d.afterInserted()):(r.parentNode&&r.parentNode.removeChild(r),delete d.element)};q.prototype.setScreenReaderSectionAttribs=function(a,b){var d=this.chart,c=d.langFormat("accessibility.screenReaderSection."+b+"RegionLabel",{chart:d,chartTitle:f(d)});B(a,{id:"highcharts-screen-reader-region-"+b+"-"+d.index,"aria-label":c});a.style.position="relative";"all"===d.options.accessibility.landmarkVerbosity&&c&&a.setAttribute("role","region")};q.prototype.defaultBeforeChartFormatter=
function(){var b=this.chart,d=b.options.accessibility.screenReaderSection.beforeChartFormat;if(!d)return"";var c=this.getAxesDescription(),L=b.sonify&&b.options.sonification&&b.options.sonification.enabled,H="highcharts-a11y-sonify-data-btn-"+b.index,q="hc-linkto-highcharts-data-table-"+b.index,g=e(b),A=b.langFormat("accessibility.screenReaderSection.annotations.heading",{chart:b});c={headingTagName:x(b.renderTo),chartTitle:f(b),typeDescription:this.getTypeDescriptionText(),chartSubtitle:this.getSubtitleText(),
chartLongdesc:this.getLongdescText(),xAxisDescription:c.xAxis,yAxisDescription:c.yAxis,playAsSoundButton:L?this.getSonifyButtonText(H):"",viewTableButton:b.getCSV?this.getDataTableButtonText(q):"",annotationsTitle:g?A:"",annotationsList:g};b=a.i18nFormat(d,c,b);this.dataTableButtonId=q;this.sonifyButtonId=H;return b.replace(/<(\w+)[^>]*?>\s*<\/\1>/g,"")};q.prototype.defaultAfterChartFormatter=function(){var b=this.chart,d=b.options.accessibility.screenReaderSection.afterChartFormat;if(!d)return"";
var c={endOfChartMarker:this.getEndOfChartMarkerText()};return a.i18nFormat(d,c,b).replace(/<(\w+)[^>]*?>\s*<\/\1>/g,"")};q.prototype.getLinkedDescription=function(){var a=this.linkedDescriptionElement;return D(a&&a.innerHTML||"")};q.prototype.getLongdescText=function(){var a=this.chart.options,b=a.caption;b=b&&b.text;var d=this.getLinkedDescription();return a.accessibility.description||d||b||""};q.prototype.getTypeDescriptionText=function(){var a=this.chart;return a.types?a.options.accessibility.typeDescription||
g(a,a.types):""};q.prototype.getDataTableButtonText=function(a){var b=this.chart;b=b.langFormat("accessibility.table.viewAsDataTableButtonText",{chart:b,chartTitle:f(b)});return'<button id="'+a+'">'+b+"</button>"};q.prototype.getSonifyButtonText=function(a){var b=this.chart;if(b.options.sonification&&!1===b.options.sonification.enabled)return"";b=b.langFormat("accessibility.sonification.playAsSoundButtonText",{chart:b,chartTitle:f(b)});return'<button id="'+a+'">'+b+"</button>"};q.prototype.getSubtitleText=
function(){var a=this.chart.options.subtitle;return D(a&&a.text||"")};q.prototype.getEndOfChartMarkerText=function(){var a=this.chart,b=a.langFormat("accessibility.screenReaderSection.endOfChartMarker",{chart:a});return'<div id="highcharts-end-of-chart-marker-'+a.index+'">'+b+"</div>"};q.prototype.onDataTableCreated=function(a){var b=this.chart;if(b.options.accessibility.enabled){this.viewDataTableButton&&this.viewDataTableButton.setAttribute("aria-expanded","true");var d=a.tree.attributes||{};d.tabindex=
-1;d.summary=b.langFormat("accessibility.table.tableSummary",{chart:b});a.tree.attributes=d}};q.prototype.focusDataTable=function(){var a=this.dataTableDiv;(a=a&&a.getElementsByTagName("table")[0])&&a.focus&&a.focus()};q.prototype.initSonifyButton=function(a){var b=this,d=this.sonifyButton=F(a),c=this.chart,f=function(a){d&&(d.setAttribute("aria-hidden","true"),d.setAttribute("aria-label",""));a.preventDefault();a.stopPropagation();a=c.langFormat("accessibility.sonification.playAsSoundClickAnnouncement",
{chart:c});b.announcer.announce(a);setTimeout(function(){d&&(d.removeAttribute("aria-hidden"),d.removeAttribute("aria-label"));c.sonify&&c.sonify()},1E3)};d&&c&&(d.setAttribute("tabindex",-1),d.onclick=function(a){(c.options.accessibility&&c.options.accessibility.screenReaderSection.onPlayAsSoundClick||f).call(this,a,c)})};q.prototype.initDataTableButton=function(a){var b=this.viewDataTableButton=F(a),d=this.chart;a=a.replace("hc-linkto-","");b&&(B(b,{tabindex:-1,"aria-expanded":!!F(a)}),b.onclick=
d.options.accessibility.screenReaderSection.onViewDataTableClick||function(){d.viewData()})};q.prototype.getAxesDescription=function(){var a=this.chart,b=function(b,d){b=a[b];return 1<b.length||b[0]&&G(b[0].options.accessibility&&b[0].options.accessibility.enabled,d)},d=!!a.types&&0>a.types.indexOf("map"),c=!!a.hasCartesianSeries,f=b("xAxis",!a.angular&&c&&d);b=b("yAxis",c&&d);d={};f&&(d.xAxis=this.getAxisDescriptionText("xAxis"));b&&(d.yAxis=this.getAxisDescriptionText("yAxis"));return d};q.prototype.getAxisDescriptionText=
function(a){var c=this.chart,f=c[a];return c.langFormat("accessibility.axis."+a+"Description"+(1<f.length?"Plural":"Singular"),{chart:c,names:f.map(function(a){return d(a)}),ranges:f.map(function(a){return b(a)}),numAxes:f.length})};q.prototype.destroy=function(){this.announcer&&this.announcer.destroy()};return q}(h)});v(a,"Accessibility/Components/MenuComponent.js",[a["Core/Chart/Chart.js"],a["Core/Utilities.js"],a["Accessibility/AccessibilityComponent.js"],a["Accessibility/KeyboardNavigationHandler.js"],
a["Accessibility/Utils/ChartUtilities.js"],a["Accessibility/Utils/HTMLUtilities.js"]],function(a,h,m,u,n,p){var w=this&&this.__extends||function(){var a=function(c,d){a=Object.setPrototypeOf||{__proto__:[]}instanceof Array&&function(a,d){a.__proto__=d}||function(a,d){for(var b in d)d.hasOwnProperty(b)&&(a[b]=d[b])};return a(c,d)};return function(c,d){function b(){this.constructor=c}a(c,d);c.prototype=null===d?Object.create(d):(b.prototype=d.prototype,new b)}}(),t=h.attr,k=n.getChartTitle,l=n.unhideChartElementFromAT,
g=p.getFakeMouseEvent;h=function(a){function c(){return null!==a&&a.apply(this,arguments)||this}w(c,a);c.prototype.init=function(){var a=this.chart,b=this;this.addEvent(a,"exportMenuShown",function(){b.onMenuShown()});this.addEvent(a,"exportMenuHidden",function(){b.onMenuHidden()});this.createProxyGroup()};c.prototype.onMenuHidden=function(){var a=this.chart.exportContextMenu;a&&a.setAttribute("aria-hidden","true");this.isExportMenuShown=!1;this.setExportButtonExpandedState("false")};c.prototype.onMenuShown=
function(){var a=this.chart,b=a.exportContextMenu;b&&(this.addAccessibleContextMenuAttribs(),l(a,b));this.isExportMenuShown=!0;this.setExportButtonExpandedState("true")};c.prototype.setExportButtonExpandedState=function(a){this.exportButtonProxy&&this.exportButtonProxy.buttonElement.setAttribute("aria-expanded",a)};c.prototype.onChartRender=function(){this.proxyProvider.clearGroup("chartMenu");this.proxyMenuButton()};c.prototype.proxyMenuButton=function(){var a=this.chart,b=this.proxyProvider,c=a.exportSVGElements&&
a.exportSVGElements[0],e=a.options.exporting,g=a.exportSVGElements&&a.exportSVGElements[0];e&&!1!==e.enabled&&e.accessibility&&e.accessibility.enabled&&g&&g.element&&c&&(this.exportButtonProxy=b.addProxyElement("chartMenu",{click:c},{"aria-label":a.langFormat("accessibility.exporting.menuButtonLabel",{chart:a,chartTitle:k(a)}),"aria-expanded":!1}))};c.prototype.createProxyGroup=function(){this.chart&&this.proxyProvider&&this.proxyProvider.addGroup("chartMenu","div")};c.prototype.addAccessibleContextMenuAttribs=
function(){var a=this.chart,b=a.exportDivElements;b&&b.length&&(b.forEach(function(a){a&&("LI"!==a.tagName||a.children&&a.children.length?a.setAttribute("aria-hidden","true"):a.setAttribute("tabindex",-1))}),(b=b[0]&&b[0].parentNode)&&t(b,{"aria-hidden":void 0,"aria-label":a.langFormat("accessibility.exporting.chartMenuLabel",{chart:a}),role:"list"}))};c.prototype.getKeyboardNavigation=function(){var a=this.keyCodes,b=this.chart,c=this;return new u(b,{keyCodeMap:[[[a.left,a.up],function(){return c.onKbdPrevious(this)}],
[[a.right,a.down],function(){return c.onKbdNext(this)}],[[a.enter,a.space],function(){return c.onKbdClick(this)}]],validate:function(){return!!b.exporting&&!1!==b.options.exporting.enabled&&!1!==b.options.exporting.accessibility.enabled},init:function(){var a=c.exportButtonProxy,d=c.chart.exportingGroup;a&&d&&b.setFocusToElement(d,a.buttonElement)},terminate:function(){b.hideExportMenu()}})};c.prototype.onKbdPrevious=function(a){var b=this.chart,d=b.options.accessibility;a=a.response;for(var c=b.highlightedExportItemIx||
0;c--;)if(b.highlightExportItem(c))return a.success;return d.keyboardNavigation.wrapAround?(b.highlightLastExportItem(),a.success):a.prev};c.prototype.onKbdNext=function(a){var b=this.chart,d=b.options.accessibility;a=a.response;for(var c=(b.highlightedExportItemIx||0)+1;c<b.exportDivElements.length;++c)if(b.highlightExportItem(c))return a.success;return d.keyboardNavigation.wrapAround?(b.highlightExportItem(0),a.success):a.next};c.prototype.onKbdClick=function(a){var b=this.chart,d=b.exportDivElements[b.highlightedExportItemIx],
c=(b.exportSVGElements&&b.exportSVGElements[0]).element;this.isExportMenuShown?this.fakeClickEvent(d):(this.fakeClickEvent(c),b.highlightExportItem(0));return a.response.success};return c}(m);(function(c){function e(){var a=this.exportSVGElements&&this.exportSVGElements[0];if(a&&(a=a.element,a.onclick))a.onclick(g("click"))}function d(){var a=this.exportDivElements;a&&this.exportContextMenu&&(a.forEach(function(a){if(a&&"highcharts-menu-item"===a.className&&a.onmouseout)a.onmouseout(g("mouseout"))}),
this.highlightedExportItemIx=0,this.exportContextMenu.hideMenu(),this.container.focus())}function b(a){var b=this.exportDivElements&&this.exportDivElements[a],d=this.exportDivElements&&this.exportDivElements[this.highlightedExportItemIx];if(b&&"LI"===b.tagName&&(!b.children||!b.children.length)){var c=!!(this.renderTo.getElementsByTagName("g")[0]||{}).focus;b.focus&&c&&b.focus();if(d&&d.onmouseout)d.onmouseout(g("mouseout"));if(b.onmouseover)b.onmouseover(g("mouseover"));this.highlightedExportItemIx=
a;return!0}return!1}function f(){if(this.exportDivElements)for(var a=this.exportDivElements.length;a--;)if(this.highlightExportItem(a))return!0;return!1}var l=[];c.compose=function(c){-1===l.indexOf(c)&&(l.push(c),c=a.prototype,c.hideExportMenu=d,c.highlightExportItem=b,c.highlightLastExportItem=f,c.showExportMenu=e)}})(h||(h={}));return h});v(a,"Accessibility/KeyboardNavigation.js",[a["Core/Globals.js"],a["Accessibility/Components/MenuComponent.js"],a["Core/Utilities.js"],a["Accessibility/Utils/EventProvider.js"],
a["Accessibility/Utils/HTMLUtilities.js"]],function(a,h,m,u,n){var p=a.doc,w=a.win,t=m.addEvent,k=m.fireEvent,l=n.getElement;m=function(){function a(a,e){this.components=this.chart=void 0;this.currentModuleIx=NaN;this.exitAnchor=this.eventProvider=void 0;this.modules=[];this.tabindexContainer=void 0;this.init(a,e)}a.prototype.init=function(a,e){var d=this,b=this.eventProvider=new u;this.chart=a;this.components=e;this.modules=[];this.currentModuleIx=0;this.update();b.addEvent(this.tabindexContainer,
"keydown",function(a){return d.onKeydown(a)});b.addEvent(this.tabindexContainer,"focus",function(a){return d.onFocus(a)});["mouseup","touchend"].forEach(function(a){return b.addEvent(p,a,function(){return d.onMouseUp()})});["mousedown","touchstart"].forEach(function(c){return b.addEvent(a.renderTo,c,function(){d.isClickingChart=!0})});b.addEvent(a.renderTo,"mouseover",function(){d.pointerIsOverChart=!0});b.addEvent(a.renderTo,"mouseout",function(){d.pointerIsOverChart=!1})};a.prototype.update=function(a){var c=
this.chart.options.accessibility;c=c&&c.keyboardNavigation;var d=this.components;this.updateContainerTabindex();c&&c.enabled&&a&&a.length?(this.modules=a.reduce(function(a,c){c=d[c].getKeyboardNavigation();return a.concat(c)},[]),this.updateExitAnchor()):(this.modules=[],this.currentModuleIx=0,this.removeExitAnchor())};a.prototype.onFocus=function(a){var c=this.chart;a=a.relatedTarget&&c.container.contains(a.relatedTarget);this.exiting||this.tabbingInBackwards||this.isClickingChart||a||!this.modules[0]||
this.modules[0].init(1);this.exiting=!1};a.prototype.onMouseUp=function(){delete this.isClickingChart;if(!this.keyboardReset&&!this.pointerIsOverChart){var a=this.chart,e=this.modules&&this.modules[this.currentModuleIx||0];e&&e.terminate&&e.terminate();a.focusElement&&a.focusElement.removeFocusBorder();this.currentModuleIx=0;this.keyboardReset=!0}};a.prototype.onKeydown=function(a){a=a||w.event;var c=this.modules&&this.modules.length&&this.modules[this.currentModuleIx],d;this.exiting=this.keyboardReset=
!1;if(c){var b=c.run(a);b===c.response.success?d=!0:b===c.response.prev?d=this.prev():b===c.response.next&&(d=this.next());d&&(a.preventDefault(),a.stopPropagation())}};a.prototype.prev=function(){return this.move(-1)};a.prototype.next=function(){return this.move(1)};a.prototype.move=function(a){var c=this.modules&&this.modules[this.currentModuleIx];c&&c.terminate&&c.terminate(a);this.chart.focusElement&&this.chart.focusElement.removeFocusBorder();this.currentModuleIx+=a;if(c=this.modules&&this.modules[this.currentModuleIx]){if(c.validate&&
!c.validate())return this.move(a);if(c.init)return c.init(a),!0}this.currentModuleIx=0;this.exiting=!0;0<a?this.exitAnchor.focus():this.tabindexContainer.focus();return!1};a.prototype.updateExitAnchor=function(){var a=l("highcharts-end-of-chart-marker-"+this.chart.index);this.removeExitAnchor();a?(this.makeElementAnExitAnchor(a),this.exitAnchor=a):this.createExitAnchor()};a.prototype.updateContainerTabindex=function(){var a=this.chart.options.accessibility;a=a&&a.keyboardNavigation;a=!(a&&!1===a.enabled);
var e=this.chart,d=e.container;e.renderTo.hasAttribute("tabindex")&&(d.removeAttribute("tabindex"),d=e.renderTo);this.tabindexContainer=d;var b=d.getAttribute("tabindex");a&&!b?d.setAttribute("tabindex","0"):a||e.container.removeAttribute("tabindex")};a.prototype.makeElementAnExitAnchor=function(a){var c=this.tabindexContainer.getAttribute("tabindex")||0;a.setAttribute("class","highcharts-exit-anchor");a.setAttribute("tabindex",c);a.setAttribute("aria-hidden",!1);this.addExitAnchorEventsToEl(a)};
a.prototype.createExitAnchor=function(){var a=this.chart,e=this.exitAnchor=p.createElement("div");a.renderTo.appendChild(e);this.makeElementAnExitAnchor(e)};a.prototype.removeExitAnchor=function(){this.exitAnchor&&this.exitAnchor.parentNode&&(this.exitAnchor.parentNode.removeChild(this.exitAnchor),delete this.exitAnchor)};a.prototype.addExitAnchorEventsToEl=function(a){var c=this.chart,d=this;this.eventProvider.addEvent(a,"focus",function(a){a=a||w.event;a.relatedTarget&&c.container.contains(a.relatedTarget)||
d.exiting?d.exiting=!1:(d.tabbingInBackwards=!0,d.tabindexContainer.focus(),delete d.tabbingInBackwards,a.preventDefault(),d.modules&&d.modules.length&&(d.currentModuleIx=d.modules.length-1,(a=d.modules[d.currentModuleIx])&&a.validate&&!a.validate()?d.prev():a&&a.init(-1)))})};a.prototype.destroy=function(){this.removeExitAnchor();this.eventProvider.removeAddedEvents();this.chart.container.removeAttribute("tabindex")};return a}();(function(g){function c(){var a=this;k(this,"dismissPopupContent",{},
function(){a.tooltip&&a.tooltip.hide(0);a.hideExportMenu()})}function e(b){27===(b.which||b.keyCode)&&a.charts&&a.charts.forEach(function(a){a&&a.dismissPopupContent&&a.dismissPopupContent()})}var d=[];g.compose=function(a){h.compose(a);-1===d.indexOf(a)&&(d.push(a),a.prototype.dismissPopupContent=c);-1===d.indexOf(p)&&(d.push(p),t(p,"keydown",e));return a}})(m||(m={}));return m});v(a,"Accessibility/Components/LegendComponent.js",[a["Core/Animation/AnimationUtilities.js"],a["Core/Chart/Chart.js"],
a["Core/Globals.js"],a["Core/Legend/Legend.js"],a["Core/Utilities.js"],a["Accessibility/AccessibilityComponent.js"],a["Accessibility/KeyboardNavigationHandler.js"],a["Accessibility/Utils/HTMLUtilities.js"],a["Accessibility/Utils/ChartUtilities.js"]],function(a,h,m,u,n,p,E,t,k){function l(a){var b=a.legend&&a.legend.allItems,d=a.options.legend.accessibility||{};return!(!b||!b.length||a.colorAxis&&a.colorAxis.length||!1===d.enabled)}var g=a.animObject,c=n.addEvent;a=n.extend;var e=n.fireEvent,d=n.isNumber,
b=n.pick,f=n.syncTimeout,A=t.stripHTMLTagsFromString,z=t.addClass,w=t.removeClass,C=k.getChartTitle;h.prototype.highlightLegendItem=function(a){var b=this.legend.allItems,c=this.accessibility&&this.accessibility.components.legend.highlightedLegendItemIx,f=b[a];return f?(d(c)&&b[c]&&e(b[c].legendGroup.element,"mouseout"),b=this.legend,a=b.allItems[a].pageIx,c=b.currentPage,"undefined"!==typeof a&&a+1!==c&&b.scroll(1+a-c),a=f.legendItem,b=f.a11yProxyElement&&f.a11yProxyElement.buttonElement,a&&a.element&&
b&&this.setFocusToElement(a,b),f.legendGroup&&e(f.legendGroup.element,"mouseover"),!0):!1};c(u,"afterColorizeItem",function(a){var b=a.item;this.chart.options.accessibility.enabled&&b&&b.a11yProxyElement&&b.a11yProxyElement.buttonElement.setAttribute("aria-pressed",a.visible?"true":"false")});h=function(){};h.prototype=new p;a(h.prototype,{init:function(){var a=this;this.recreateProxies();this.addEvent(u,"afterScroll",function(){this.chart===a.chart&&(a.proxyProvider.updateGroupProxyElementPositions("legend"),
a.updateLegendItemProxyVisibility(),-1<a.highlightedLegendItemIx&&this.chart.highlightLegendItem(a.highlightedLegendItemIx))});this.addEvent(u,"afterPositionItem",function(b){this.chart===a.chart&&this.chart.renderer&&a.updateProxyPositionForItem(b.item)});this.addEvent(u,"afterRender",function(){this.chart===a.chart&&this.chart.renderer&&a.recreateProxies()&&f(function(){return a.proxyProvider.updateGroupProxyElementPositions("legend")},g(b(this.chart.renderer.globalAnimation,!0)).duration)})},updateLegendItemProxyVisibility:function(){var a=
this.chart,b=a.legend,d=b.currentPage||1,c=b.clipHeight||0;(b.allItems||[]).forEach(function(f){if(f.a11yProxyElement){var e=f.a11yProxyElement.element,g=!1;if(b.pages&&b.pages.length){g=f.pageIx||0;var q=f._legendItemPos?f._legendItemPos[1]:0;f=f.legendItem?Math.round(f.legendItem.getBBox().height):0;g=q+f-b.pages[g]>c||g!==d-1}g?a.styledMode?z(e,"highcharts-a11y-invisible"):e.style.visibility="hidden":(w(e,"highcharts-a11y-invisible"),e.style.visibility="")}})},onChartRender:function(){l(this.chart)||
this.removeProxies()},highlightAdjacentLegendPage:function(a){var b=this.chart,d=b.legend;a=(d.currentPage||1)+a;var c=d.pages||[];if(0<a&&a<=c.length){c=d.allItems.length;for(var f=0;f<c;++f)if(d.allItems[f].pageIx+1===a){b.highlightLegendItem(f)&&(this.highlightedLegendItemIx=f);break}}},updateProxyPositionForItem:function(a){a.a11yProxyElement&&a.a11yProxyElement.refreshPosition()},recreateProxies:function(){this.removeProxies();return l(this.chart)?(this.addLegendProxyGroup(),this.proxyLegendItems(),
this.updateLegendItemProxyVisibility(),this.updateLegendTitle(),!0):!1},removeProxies:function(){this.proxyProvider.removeGroup("legend")},updateLegendTitle:function(){var a=this.chart,b=A((a.legend&&a.legend.options.title&&a.legend.options.title.text||"").replace(/<br ?\/?>/g," "));a=a.langFormat("accessibility.legend.legendLabel"+(b?"":"NoTitle"),{chart:a,legendTitle:b,chartTitle:C(a)});this.proxyProvider.updateGroupAttrs("legend",{"aria-label":a})},addLegendProxyGroup:function(){this.proxyProvider.addGroup("legend",
"ul",{"aria-label":"_placeholder_",role:"all"===this.chart.options.accessibility.landmarkVerbosity?"region":null})},proxyLegendItems:function(){var a=this;(this.chart.legend&&this.chart.legend.allItems||[]).forEach(function(b){b.legendItem&&b.legendItem.element&&a.proxyLegendItem(b)})},proxyLegendItem:function(a){if(a.legendItem&&a.legendGroup){var b=this.chart.langFormat("accessibility.legend.legendItem",{chart:this.chart,itemName:A(a.name),item:a});a.a11yProxyElement=this.proxyProvider.addProxyElement("legend",
{click:a.legendItem,visual:(a.legendGroup.div?a.legendItem:a.legendGroup).element},{tabindex:-1,"aria-pressed":a.visible,"aria-label":b})}},getKeyboardNavigation:function(){var a=this.keyCodes,b=this,d=this.chart;return new E(d,{keyCodeMap:[[[a.left,a.right,a.up,a.down],function(a){return b.onKbdArrowKey(this,a)}],[[a.enter,a.space],function(d){return m.isFirefox&&d===a.space?this.response.success:b.onKbdClick(this)}],[[a.pageDown,a.pageUp],function(d){b.highlightAdjacentLegendPage(d===a.pageDown?
1:-1);return this.response.success}]],validate:function(){return b.shouldHaveLegendNavigation()},init:function(a){return b.onKbdNavigationInit(a)},terminate:function(){b.highlightedLegendItemIx=-1;d.legend.allItems.forEach(function(a){return a.setState("",!0)})}})},onKbdArrowKey:function(a,b){var d=this.keyCodes,c=a.response,f=this.chart,e=f.options.accessibility,g=f.legend.allItems.length;b=b===d.left||b===d.up?-1:1;return f.highlightLegendItem(this.highlightedLegendItemIx+b)?(this.highlightedLegendItemIx+=
b,c.success):1<g&&e.keyboardNavigation.wrapAround?(a.init(b),c.success):c[0<b?"next":"prev"]},onKbdClick:function(a){var b=this.chart.legend.allItems[this.highlightedLegendItemIx];b&&b.a11yProxyElement&&b.a11yProxyElement.click();return a.response.success},shouldHaveLegendNavigation:function(){var a=this.chart,b=a.colorAxis&&a.colorAxis.length,d=(a.options.legend||{}).accessibility||{};return!!(a.legend&&a.legend.allItems&&a.legend.display&&!b&&d.enabled&&d.keyboardNavigation&&d.keyboardNavigation.enabled)},
onKbdNavigationInit:function(a){var b=this.chart,d=b.legend.allItems.length-1;a=0<a?0:d;b.highlightLegendItem(a);this.highlightedLegendItemIx=a}});return h});v(a,"Accessibility/Components/SeriesComponent/SeriesDescriber.js",[a["Accessibility/Components/AnnotationsA11y.js"],a["Accessibility/Utils/ChartUtilities.js"],a["Core/FormatUtilities.js"],a["Accessibility/Utils/HTMLUtilities.js"],a["Core/Utilities.js"]],function(a,h,m,u,n){function p(a){var b=a.index;return a.series&&a.series.data&&r(b)?v(a.series.data,
function(a){return!!(a&&"undefined"!==typeof a.index&&a.index>b&&a.graphic&&a.graphic.element)})||null:null}function w(a){var b=a.chart.options.accessibility.series.pointDescriptionEnabledThreshold;return!!(!1!==b&&a.points&&a.points.length>=b)}function t(a){var b=a.options.accessibility||{};return!w(a)&&!b.exposeAsGroupOnly}function k(a){var b=a.chart.options.accessibility.keyboardNavigation.seriesNavigation;return!(!a.points||!(a.points.length<b.pointNavigationEnabledThreshold||!1===b.pointNavigationEnabledThreshold))}
function l(a,b){var d=a.series,c=d.chart;a=c.options.accessibility.point||{};var f=d.options.accessibility&&d.options.accessibility.point||{};d=d.tooltipOptions||{};c=c.options.lang;return K(b)?D(b,f.valueDecimals||a.valueDecimals||d.valueDecimals||-1,c.decimalPoint,c.accessibility.thousandsSep||c.thousandsSep):b}function g(a){var b=(a.options.accessibility||{}).description;return b&&a.chart.langFormat("accessibility.series.description",{description:b,series:a})||""}function c(a,b){return a.chart.langFormat("accessibility.series."+
b+"Description",{name:z(a[b]),series:a})}function e(a,b,d){var c=b||"",f=d||"";return a.series.pointArrayMap.reduce(function(b,d){b+=b.length?", ":"";var r=l(a,q(a[d],a.options[d]));return b+(d+": "+c+r+f)},"")}function d(a){var b=a.series,d=b.chart,c=a.series;var f=c.chart;var y=c.options.accessibility;y=y&&y.point&&y.point.valueDescriptionFormat||f.options.accessibility.point.valueDescriptionFormat;c=q(c.xAxis&&c.xAxis.options.accessibility&&c.xAxis.options.accessibility.enabled,!f.angular);if(c){var g=
a.series;var k=g.chart;var h=g.options.accessibility&&g.options.accessibility.point||{},z=k.options.accessibility.point||{};(g=g.xAxis&&g.xAxis.dateTime)?(g=g.getXDateFormat(a.x||0,k.options.tooltip.dateTimeLabelFormats),h=h.dateFormatter&&h.dateFormatter(a)||z.dateFormatter&&z.dateFormatter(a)||h.dateFormat||z.dateFormat||g,k=k.time.dateFormat(h,a.x||0,void 0)):k=void 0;h=(a.series.xAxis||{}).categories&&r(a.category)&&(""+a.category).replace("<br/>"," ");z=a.id&&0>a.id.indexOf("highcharts-");g=
"x, "+a.x;k=a.name||k||h||(z?a.id:g)}else k="";h=r(a.index)?a.index+1:"";z=a.series;var m=z.chart.options.accessibility.point||{},n=z.chart.options.accessibility&&z.chart.options.accessibility.point||{},p=z.tooltipOptions||{};g=n.valuePrefix||m.valuePrefix||p.valuePrefix||"";m=n.valueSuffix||m.valueSuffix||p.valueSuffix||"";n=l(a,a["undefined"!==typeof a.value?"value":"y"]);z=a.isNull?z.chart.langFormat("accessibility.series.nullPointValue",{point:a}):z.pointArrayMap?e(a,g,m):g+n+m;f=x(y,{point:a,
index:h,xDescription:k,value:z,separator:c?", ":""},f);y=(y=a.options&&a.options.accessibility&&a.options.accessibility.description)?" "+y:"";b=1<d.series.length&&b.name?" "+b.name+".":"";d=a.series.chart;c=A(a);k={point:a,annotations:c};d=c.length?d.langFormat("accessibility.series.pointAnnotationsDescription",k):"";a.accessibility=a.accessibility||{};a.accessibility.valueDescription=f;return f+y+b+(d?" "+d:"")}function b(a){var b=t(a),c=k(a);(b||c)&&a.points.forEach(function(c){var f;if(!(f=c.graphic&&
c.graphic.element)&&(f=c.series&&c.series.is("sunburst"),f=c.isNull&&!f)){var r=c.series,e=p(c);r=(f=e&&e.graphic)?f.parentGroup:r.graph||r.group;e=e?{x:q(c.plotX,e.plotX,0),y:q(c.plotY,e.plotY,0)}:{x:q(c.plotX,0),y:q(c.plotY,0)};e=c.series.chart.renderer.rect(e.x,e.y,1,1);e.attr({"class":"highcharts-a11y-dummy-point",fill:"none",opacity:0,"fill-opacity":0,"stroke-opacity":0});r&&r.element?(c.graphic=e,c.hasDummyGraphic=!0,e.add(r),r.element.insertBefore(e.element,f?f.element:null),f=e.element):f=
void 0}r=c.options&&c.options.accessibility&&!1===c.options.accessibility.enabled;f&&(f.setAttribute("tabindex","-1"),a.chart.styledMode||(f.style.outline="none"),b&&!r?(e=c.series,r=e.chart.options.accessibility.point||{},e=e.options.accessibility&&e.options.accessibility.point||{},c=B(e.descriptionFormatter&&e.descriptionFormatter(c)||r.descriptionFormatter&&r.descriptionFormatter(c)||d(c)),f.setAttribute("role","img"),f.setAttribute("aria-label",c)):f.setAttribute("aria-hidden",!0))})}function f(a){var b=
a.chart,d=b.types||[],f=g(a),r=function(d){return b[d]&&1<b[d].length&&a[d]},e=c(a,"xAxis"),y=c(a,"yAxis"),q={name:a.name||"",ix:a.index+1,numSeries:b.series&&b.series.length,numPoints:a.points&&a.points.length,series:a};d=1<d.length?"Combination":"";return(b.langFormat("accessibility.series.summary."+a.type+d,q)||b.langFormat("accessibility.series.summary.default"+d,q))+(f?" "+f:"")+(r("yAxis")?" "+y:"")+(r("xAxis")?" "+e:"")}var A=a.getPointAnnotationTexts,z=h.getAxisDescription,J=h.getSeriesFirstPointElement,
C=h.getSeriesA11yElement,F=h.unhideChartElementFromAT,x=m.format,D=m.numberFormat,M=u.reverseChildNodes,B=u.stripHTMLTagsFromString,v=n.find,K=n.isNumber,q=n.pick,r=n.defined;return{defaultPointDescriptionFormatter:d,defaultSeriesDescriptionFormatter:f,describeSeries:function(a){var d=a.chart,c=J(a),r=C(a),e=d.is3d&&d.is3d();if(r){r.lastChild!==c||e||M(r);b(a);F(d,r);e=a.chart;d=e.options.chart;c=1<e.series.length;e=e.options.accessibility.series.describeSingleSeries;var g=(a.options.accessibility||
{}).exposeAsGroupOnly;d.options3d&&d.options3d.enabled&&c||!(c||e||g||w(a))?r.setAttribute("aria-label",""):(d=a.chart.options.accessibility,c=d.landmarkVerbosity,(a.options.accessibility||{}).exposeAsGroupOnly?r.setAttribute("role","img"):"all"===c&&r.setAttribute("role","region"),r.setAttribute("tabindex","-1"),a.chart.styledMode||(r.style.outline="none"),r.setAttribute("aria-label",B(d.series.descriptionFormatter&&d.series.descriptionFormatter(a)||f(a))))}}}});v(a,"Accessibility/Components/SeriesComponent/NewDataAnnouncer.js",
[a["Core/Globals.js"],a["Core/Utilities.js"],a["Accessibility/Utils/Announcer.js"],a["Accessibility/Utils/ChartUtilities.js"],a["Accessibility/Utils/EventProvider.js"],a["Accessibility/Components/SeriesComponent/SeriesDescriber.js"]],function(a,h,m,u,n,p){function w(a){var b=a.series.data.filter(function(b){return a.x===b.x&&a.y===b.y});return 1===b.length?b[0]:a}function t(a,b){var d=(a||[]).concat(b||[]).reduce(function(a,b){a[b.name+b.index]=b;return a},{});return Object.keys(d).map(function(a){return d[a]})}
var k=h.addEvent,l=h.defined,g=u.getChartTitle,c=p.defaultPointDescriptionFormatter,e=p.defaultSeriesDescriptionFormatter;h=function(){function d(a){this.announcer=void 0;this.dirty={allSeries:{}};this.eventProvider=void 0;this.lastAnnouncementTime=0;this.chart=a}d.prototype.init=function(){var a=this.chart,d=a.options.accessibility.announceNewData.interruptUser?"assertive":"polite";this.lastAnnouncementTime=0;this.dirty={allSeries:{}};this.eventProvider=new n;this.announcer=new m(a,d);this.addEventListeners()};
d.prototype.destroy=function(){this.eventProvider.removeAddedEvents();this.announcer.destroy()};d.prototype.addEventListeners=function(){var a=this,d=this.chart,c=this.eventProvider;c.addEvent(d,"afterDrilldown",function(){a.lastAnnouncementTime=0});c.addEvent(d,"afterAddSeries",function(b){a.onSeriesAdded(b.series)});c.addEvent(d,"redraw",function(){a.announceDirtyData()})};d.prototype.onSeriesAdded=function(a){this.chart.options.accessibility.announceNewData.enabled&&(this.dirty.hasDirty=!0,this.dirty.allSeries[a.name+
a.index]=a,this.dirty.newSeries=l(this.dirty.newSeries)?void 0:a)};d.prototype.announceDirtyData=function(){var a=this;if(this.chart.options.accessibility.announceNewData&&this.dirty.hasDirty){var d=this.dirty.newPoint;d&&(d=w(d));this.queueAnnouncement(Object.keys(this.dirty.allSeries).map(function(b){return a.dirty.allSeries[b]}),this.dirty.newSeries,d);this.dirty={allSeries:{}}}};d.prototype.queueAnnouncement=function(a,d,c){var b=this,f=this.chart.options.accessibility.announceNewData;if(f.enabled){var e=
+new Date;f=Math.max(0,f.minAnnounceInterval-(e-this.lastAnnouncementTime));a=t(this.queuedAnnouncement&&this.queuedAnnouncement.series,a);if(d=this.buildAnnouncementMessage(a,d,c))this.queuedAnnouncement&&clearTimeout(this.queuedAnnouncementTimer),this.queuedAnnouncement={time:e,message:d,series:a},this.queuedAnnouncementTimer=setTimeout(function(){b&&b.announcer&&(b.lastAnnouncementTime=+new Date,b.announcer.announce(b.queuedAnnouncement.message),delete b.queuedAnnouncement,delete b.queuedAnnouncementTimer)},
f)}};d.prototype.buildAnnouncementMessage=function(b,d,l){var f=this.chart,k=f.options.accessibility.announceNewData;if(k.announcementFormatter&&(b=k.announcementFormatter(b,d,l),!1!==b))return b.length?b:null;b=a.charts&&1<a.charts.length?"Multiple":"Single";b=d?"newSeriesAnnounce"+b:l?"newPointAnnounce"+b:"newDataAnnounce";k=g(f);return f.langFormat("accessibility.announceNewData."+b,{chartTitle:k,seriesDesc:d?e(d):null,pointDesc:l?c(l):null,point:l,series:d})};return d}();(function(a){function b(a){var b=
this.chart,d=this.newDataAnnouncer;d&&d.chart===b&&b.options.accessibility.announceNewData.enabled&&(d.dirty.newPoint=l(d.dirty.newPoint)?void 0:a.point)}function d(){var a=this.chart,b=this.newDataAnnouncer;b&&b.chart===a&&a.options.accessibility.announceNewData.enabled&&(b.dirty.hasDirty=!0,b.dirty.allSeries[this.name+this.index]=this)}a.composedClasses=[];a.compose=function(c){-1===a.composedClasses.indexOf(c)&&(a.composedClasses.push(c),k(c,"addPoint",b),k(c,"updatedData",d))}})(h||(h={}));return h});
v(a,"Accessibility/ProxyElement.js",[a["Core/Globals.js"],a["Core/Utilities.js"],a["Accessibility/Utils/EventProvider.js"],a["Accessibility/Utils/ChartUtilities.js"],a["Accessibility/Utils/HTMLUtilities.js"]],function(a,h,m,u,n){var p=a.doc,w=h.attr,t=h.css,k=h.merge,l=u.fireEventOnWrappedOrUnwrappedElement,g=n.cloneMouseEvent,c=n.cloneTouchEvent,e=n.getFakeMouseEvent,d=n.removeElement;return function(){function a(a,b,d,c){this.chart=a;this.target=b;this.groupType=d;this.eventProvider=new m;d="ul"===
d?p.createElement("li"):null;var f=this.buttonElement=p.createElement("button");a.styledMode||this.hideButtonVisually(f);d?(d.appendChild(f),this.element=d):this.element=f;this.updateTarget(b,c)}a.prototype.click=function(){var a=this.getTargetPosition();a.x+=a.width/2;a.y+=a.height/2;a=e("click",a);l(this.target.click,a)};a.prototype.updateTarget=function(a,b){this.target=a;this.updateCSSClassName();w(this.buttonElement,k({"aria-label":this.getTargetAttr(a.click,"aria-label")},b));this.eventProvider.removeAddedEvents();
this.addProxyEventsToButton(this.buttonElement,a.click);this.refreshPosition()};a.prototype.refreshPosition=function(){var a=this.getTargetPosition();t(this.buttonElement,{width:(a.width||1)+"px",height:(a.height||1)+"px",left:(Math.round(a.x)||0)+"px",top:(Math.round(a.y)||0)+"px"})};a.prototype.remove=function(){this.eventProvider.removeAddedEvents();d(this.element)};a.prototype.updateCSSClassName=function(){var a=this.chart.legend;a=a.group&&a.group.div;a=-1<(a&&a.className||"").indexOf("highcharts-no-tooltip");
var b=-1<(this.getTargetAttr(this.target.click,"class")||"").indexOf("highcharts-no-tooltip");this.buttonElement.className=a||b?"highcharts-a11y-proxy-button highcharts-no-tooltip":"highcharts-a11y-proxy-button"};a.prototype.addProxyEventsToButton=function(a,b){var d=this;"click touchstart touchend touchcancel touchmove mouseover mouseenter mouseleave mouseout".split(" ").forEach(function(f){var e=0===f.indexOf("touch");d.eventProvider.addEvent(a,f,function(a){var d=e?c(a):g(a);b&&l(b,d);a.stopPropagation();
e||a.preventDefault()},{passive:!1})})};a.prototype.hideButtonVisually=function(a){t(a,{borderWidth:0,backgroundColor:"transparent",cursor:"pointer",outline:"none",opacity:.001,filter:"alpha(opacity=1)",zIndex:999,overflow:"hidden",padding:0,margin:0,display:"block",position:"absolute","-ms-filter":"progid:DXImageTransform.Microsoft.Alpha(Opacity=1)"})};a.prototype.getTargetPosition=function(){var a=this.target.click;a=a.element?a.element:a;var b=this.target.visual||a;return(a=this.chart.renderTo)&&
b&&b.getBoundingClientRect?(b=b.getBoundingClientRect(),a=a.getBoundingClientRect(),{x:b.left-a.left,y:b.top-a.top,width:b.right-b.left,height:b.bottom-b.top}):{x:0,y:0,width:1,height:1}};a.prototype.getTargetAttr=function(a,b){return a.element?a.element.getAttribute(b):a.getAttribute(b)};return a}()});v(a,"Accessibility/ProxyProvider.js",[a["Core/Globals.js"],a["Core/Utilities.js"],a["Accessibility/Utils/ChartUtilities.js"],a["Accessibility/Utils/DOMElementProvider.js"],a["Accessibility/Utils/HTMLUtilities.js"],
a["Accessibility/ProxyElement.js"]],function(a,h,m,u,n,p){var w=a.doc,t=h.attr,k=h.css,l=m.unhideChartElementFromAT,g=n.removeElement,c=n.removeChildNodes;return function(){function a(a){this.chart=a;this.domElementProvider=new u;this.groups={};this.groupOrder=[];this.beforeChartProxyPosContainer=this.createProxyPosContainer("before");this.afterChartProxyPosContainer=this.createProxyPosContainer("after");this.update()}a.prototype.addProxyElement=function(a,b,c){var d=this.groups[a];if(!d)throw Error("ProxyProvider.addProxyElement: Invalid group key "+
a);a=new p(this.chart,b,d.type,c);d.proxyContainerElement.appendChild(a.element);d.proxyElements.push(a);return a};a.prototype.addGroup=function(a,b,c){if(!this.groups[a]){var d=this.domElementProvider.createElement(b);if(c&&c.role&&"div"!==b){var f=this.domElementProvider.createElement("div");f.appendChild(d)}else f=d;f.className="highcharts-a11y-proxy-group highcharts-a11y-proxy-group-"+a.replace(/\W/g,"-");this.groups[a]={proxyContainerElement:d,groupElement:f,type:b,proxyElements:[]};t(f,c||{});
"ul"===b&&(this.chart.styledMode||(d.style.listStyle="none"),d.setAttribute("role","list"));this.afterChartProxyPosContainer.appendChild(f);this.updateGroupOrder(this.groupOrder)}};a.prototype.updateGroupAttrs=function(a,b){var d=this.groups[a];if(!d)throw Error("ProxyProvider.updateGroupAttrs: Invalid group key "+a);t(d.groupElement,b)};a.prototype.updateGroupOrder=function(a){var b=this;this.groupOrder=a.slice();if(!this.isDOMOrderGroupOrder()){var d=a.indexOf("series"),e=-1<d?a.slice(0,d):a,g=
-1<d?a.slice(d+1):[];a=w.activeElement;["before","after"].forEach(function(a){var d=b["before"===a?"beforeChartProxyPosContainer":"afterChartProxyPosContainer"];a="before"===a?e:g;c(d);a.forEach(function(a){(a=b.groups[a])&&d.appendChild(a.groupElement)})});(this.beforeChartProxyPosContainer.contains(a)||this.afterChartProxyPosContainer.contains(a))&&a&&a.focus&&a.focus()}};a.prototype.clearGroup=function(a){var b=this.groups[a];if(!b)throw Error("ProxyProvider.clearGroup: Invalid group key "+a);
c(b.proxyContainerElement)};a.prototype.removeGroup=function(a){var b=this.groups[a];b&&(g(b.groupElement),delete this.groups[a])};a.prototype.update=function(){this.updatePosContainerPositions();this.updateGroupOrder(this.groupOrder);this.updateProxyElementPositions()};a.prototype.updateProxyElementPositions=function(){Object.keys(this.groups).forEach(this.updateGroupProxyElementPositions.bind(this))};a.prototype.updateGroupProxyElementPositions=function(a){(a=this.groups[a])&&a.proxyElements.forEach(function(a){return a.refreshPosition()})};
a.prototype.destroy=function(){this.domElementProvider.destroyCreatedElements()};a.prototype.createProxyPosContainer=function(a){var b=this.domElementProvider.createElement("div");b.setAttribute("aria-hidden","false");b.className="highcharts-a11y-proxy-container"+(a?"-"+a:"");k(b,{top:"0",left:"0"});this.chart.styledMode||(b.style.whiteSpace="nowrap",b.style.position="absolute");return b};a.prototype.getCurrentGroupOrderInDOM=function(){var a=this,b=function(b){var d=[];b=b.children;for(var c=0;c<
b.length;++c){a:{var e=b[c];for(var f=Object.keys(a.groups),g=f.length;g--;){var l=f[g],k=a.groups[l];if(k&&e===k.groupElement){e=l;break a}}e=void 0}e&&d.push(e)}return d},c=b(this.beforeChartProxyPosContainer);b=b(this.afterChartProxyPosContainer);c.push("series");return c.concat(b)};a.prototype.isDOMOrderGroupOrder=function(){var a=this,b=this.getCurrentGroupOrderInDOM(),c=this.groupOrder.filter(function(b){return"series"===b||!!a.groups[b]}),e=b.length;if(e!==c.length)return!1;for(;e--;)if(b[e]!==
c[e])return!1;return!0};a.prototype.updatePosContainerPositions=function(){var a=this.chart,b=a.renderer.box;a.container.insertBefore(this.afterChartProxyPosContainer,b.nextSibling);a.container.insertBefore(this.beforeChartProxyPosContainer,b);l(this.chart,this.afterChartProxyPosContainer);l(this.chart,this.beforeChartProxyPosContainer)};return a}()});v(a,"Extensions/RangeSelector.js",[a["Core/Axis/Axis.js"],a["Core/Chart/Chart.js"],a["Core/Globals.js"],a["Core/DefaultOptions.js"],a["Core/Renderer/SVG/SVGElement.js"],
a["Core/Utilities.js"]],function(a,h,m,u,n,p){function w(a){if(-1!==a.indexOf("%L"))return"text";var b="aAdewbBmoyY".split("").some(function(b){return-1!==a.indexOf("%"+b)}),d="HkIlMS".split("").some(function(b){return-1!==a.indexOf("%"+b)});return b&&d?"datetime-local":b?"date":d?"time":"text"}var t=u.defaultOptions,k=p.addEvent,l=p.createElement,g=p.css,c=p.defined,e=p.destroyObjectProperties,d=p.discardElement,b=p.extend,f=p.find,A=p.fireEvent,z=p.isNumber,v=p.merge,C=p.objectEach,F=p.pad,x=p.pick,
D=p.pInt,M=p.splat;b(t,{rangeSelector:{allButtonsEnabled:!1,buttons:void 0,buttonSpacing:5,dropdown:"responsive",enabled:void 0,verticalAlign:"top",buttonTheme:{width:28,height:18,padding:2,zIndex:7},floating:!1,x:0,y:0,height:void 0,inputBoxBorderColor:"none",inputBoxHeight:17,inputBoxWidth:void 0,inputDateFormat:"%b %e, %Y",inputDateParser:void 0,inputEditDateFormat:"%Y-%m-%d",inputEnabled:!0,inputPosition:{align:"right",x:0,y:0},inputSpacing:5,selected:void 0,buttonPosition:{align:"left",x:0,y:0},
inputStyle:{color:"#335cad",cursor:"pointer"},labelStyle:{color:"#666666"}}});b(t.lang,{rangeSelectorZoom:"Zoom",rangeSelectorFrom:"",rangeSelectorTo:"\u2192"});var B=function(){function f(a){this.buttons=void 0;this.buttonOptions=f.prototype.defaultButtons;this.initialButtonGroupWidth=0;this.options=void 0;this.chart=a;this.init(a)}f.prototype.clickButton=function(b,d){var f=this.chart,r=this.buttonOptions[b],e=f.xAxis[0],g=f.scroller&&f.scroller.getUnionExtremes()||e||{},y=g.dataMin,l=g.dataMax,
q=e&&Math.round(Math.min(e.max,x(l,e.max))),h=r.type;g=r._range;var m,n=r.dataGrouping;if(null!==y&&null!==l){f.fixedRange=g;this.setSelected(b);n&&(this.forcedDataGrouping=!0,a.prototype.setDataGrouping.call(e||{chart:this.chart},n,!1),this.frozenStates=r.preserveDataGrouping);if("month"===h||"year"===h)if(e){h={range:r,max:q,chart:f,dataMin:y,dataMax:l};var p=e.minFromRange.call(h);z(h.newMax)&&(q=h.newMax)}else g=r;else if(g)p=Math.max(q-g,y),q=Math.min(p+g,l);else if("ytd"===h)if(e)"undefined"===
typeof l&&(y=Number.MAX_VALUE,l=Number.MIN_VALUE,f.series.forEach(function(a){a=a.xData;y=Math.min(a[0],y);l=Math.max(a[a.length-1],l)}),d=!1),q=this.getYTDExtremes(l,y,f.time.useUTC),p=m=q.min,q=q.max;else{this.deferredYTDClick=b;return}else"all"===h&&e&&(f.navigator&&f.navigator.baseSeries[0]&&(f.navigator.baseSeries[0].xAxis.options.range=void 0),p=y,q=l);c(p)&&(p+=r._offsetMin);c(q)&&(q+=r._offsetMax);this.dropdown&&(this.dropdown.selectedIndex=b+1);if(e)e.setExtremes(p,q,x(d,!0),void 0,{trigger:"rangeSelectorButton",
rangeSelectorButton:r});else{var t=M(f.options.xAxis)[0];var u=t.range;t.range=g;var w=t.min;t.min=m;k(f,"load",function(){t.range=u;t.min=w})}A(this,"afterBtnClick")}};f.prototype.setSelected=function(a){this.selected=this.options.selected=a};f.prototype.init=function(a){var b=this,d=a.options.rangeSelector,c=d.buttons||b.defaultButtons.slice(),f=d.selected,e=function(){var a=b.minInput,d=b.maxInput;a&&a.blur&&A(a,"blur");d&&d.blur&&A(d,"blur")};b.chart=a;b.options=d;b.buttons=[];b.buttonOptions=
c;this.eventsToUnbind=[];this.eventsToUnbind.push(k(a.container,"mousedown",e));this.eventsToUnbind.push(k(a,"resize",e));c.forEach(b.computeButtonRange);"undefined"!==typeof f&&c[f]&&this.clickButton(f,!1);this.eventsToUnbind.push(k(a,"load",function(){a.xAxis&&a.xAxis[0]&&k(a.xAxis[0],"setExtremes",function(d){this.max-this.min!==a.fixedRange&&"rangeSelectorButton"!==d.trigger&&"updatedData"!==d.trigger&&b.forcedDataGrouping&&!b.frozenStates&&this.setDataGrouping(!1,!1)})}))};f.prototype.updateButtonStates=
function(){var a=this,b=this.chart,d=this.dropdown,c=b.xAxis[0],f=Math.round(c.max-c.min),e=!c.hasVisibleSeries,g=b.scroller&&b.scroller.getUnionExtremes()||c,l=g.dataMin,q=g.dataMax;b=a.getYTDExtremes(q,l,b.time.useUTC);var k=b.min,h=b.max,m=a.selected,n=z(m),p=a.options.allButtonsEnabled,x=a.buttons;a.buttonOptions.forEach(function(b,r){var g=b._range,y=b.type,H=b.count||1,L=x[r],I=0,N=b._offsetMax-b._offsetMin;b=r===m;var t=g>q-l,A=g<c.minRange,u=!1,z=!1;g=g===f;("month"===y||"year"===y)&&f+36E5>=
864E5*{month:28,year:365}[y]*H-N&&f-36E5<=864E5*{month:31,year:366}[y]*H+N?g=!0:"ytd"===y?(g=h-k+N===f,u=!b):"all"===y&&(g=c.max-c.min>=q-l,z=!b&&n&&g);y=!p&&(t||A||z||e);H=b&&g||g&&!n&&!u||b&&a.frozenStates;y?I=3:H&&(n=!0,I=2);L.state!==I&&(L.setState(I),d&&(d.options[r+1].disabled=y,2===I&&(d.selectedIndex=r+1)),0===I&&m===r&&a.setSelected())})};f.prototype.computeButtonRange=function(a){var b=a.type,d=a.count||1,c={millisecond:1,second:1E3,minute:6E4,hour:36E5,day:864E5,week:6048E5};if(c[b])a._range=
c[b]*d;else if("month"===b||"year"===b)a._range=864E5*{month:30,year:365}[b]*d;a._offsetMin=x(a.offsetMin,0);a._offsetMax=x(a.offsetMax,0);a._range+=a._offsetMax-a._offsetMin};f.prototype.getInputValue=function(a){a="min"===a?this.minInput:this.maxInput;var b=this.chart.options.rangeSelector,d=this.chart.time;return a?("text"===a.type&&b.inputDateParser||this.defaultInputDateParser)(a.value,d.useUTC,d):0};f.prototype.setInputValue=function(a,b){var d=this.options,f=this.chart.time,e="min"===a?this.minInput:
this.maxInput;a="min"===a?this.minDateBox:this.maxDateBox;if(e){var r=e.getAttribute("data-hc-time");r=c(r)?Number(r):void 0;c(b)&&(c(r)&&e.setAttribute("data-hc-time-previous",r),e.setAttribute("data-hc-time",b),r=b);e.value=f.dateFormat(this.inputTypeFormats[e.type]||d.inputEditDateFormat,r);a&&a.attr({text:f.dateFormat(d.inputDateFormat,r)})}};f.prototype.setInputExtremes=function(a,b,d){if(a="min"===a?this.minInput:this.maxInput){var c=this.inputTypeFormats[a.type],f=this.chart.time;c&&(b=f.dateFormat(c,
b),a.min!==b&&(a.min=b),d=f.dateFormat(c,d),a.max!==d&&(a.max=d))}};f.prototype.showInput=function(a){var b="min"===a?this.minDateBox:this.maxDateBox;if((a="min"===a?this.minInput:this.maxInput)&&b&&this.inputGroup){var d="text"===a.type,c=this.inputGroup,f=c.translateX;c=c.translateY;var e=this.options.inputBoxWidth;g(a,{width:d?b.width+(e?-2:20)+"px":"auto",height:d?b.height-2+"px":"auto",border:"2px solid silver"});d&&e?g(a,{left:f+b.x+"px",top:c+"px"}):g(a,{left:Math.min(Math.round(b.x+f-(a.offsetWidth-
b.width)/2),this.chart.chartWidth-a.offsetWidth)+"px",top:c-(a.offsetHeight-b.height)/2+"px"})}};f.prototype.hideInput=function(a){(a="min"===a?this.minInput:this.maxInput)&&g(a,{top:"-9999em",border:0,width:"1px",height:"1px"})};f.prototype.defaultInputDateParser=function(a,b,d){var c=a.split("/").join("-").split(" ").join("T");-1===c.indexOf("T")&&(c+="T00:00");if(b)c+="Z";else{var f;if(f=m.isSafari)f=c,f=!(6<f.length&&(f.lastIndexOf("-")===f.length-6||f.lastIndexOf("+")===f.length-6));f&&(f=(new Date(c)).getTimezoneOffset()/
60,c+=0>=f?"+"+F(-f)+":00":"-"+F(f)+":00")}c=Date.parse(c);z(c)||(a=a.split("-"),c=Date.UTC(D(a[0]),D(a[1])-1,D(a[2])));d&&b&&z(c)&&(c+=d.getTimezoneOffset(c));return c};f.prototype.drawInput=function(a){function d(){var b=r.getInputValue(a),d=c.xAxis[0],f=c.scroller&&c.scroller.xAxis?c.scroller.xAxis:d,e=f.dataMin;f=f.dataMax;var g=r.maxInput,l=r.minInput;b!==Number(x.getAttribute("data-hc-time-previous"))&&z(b)&&(x.setAttribute("data-hc-time-previous",b),n&&g&&z(e)?b>Number(g.getAttribute("data-hc-time"))?
b=void 0:b<e&&(b=e):l&&z(f)&&(b<Number(l.getAttribute("data-hc-time"))?b=void 0:b>f&&(b=f)),"undefined"!==typeof b&&d.setExtremes(n?b:d.min,n?d.max:b,void 0,void 0,{trigger:"rangeSelectorInput"}))}var c=this.chart,f=this.div,e=this.inputGroup,r=this,q=c.renderer.style||{},k=c.renderer,h=c.options.rangeSelector,n="min"===a,p=t.lang[n?"rangeSelectorFrom":"rangeSelectorTo"]||"";p=k.label(p,0).addClass("highcharts-range-label").attr({padding:p?2:0,height:p?h.inputBoxHeight:0}).add(e);k=k.label("",0).addClass("highcharts-range-input").attr({padding:2,
width:h.inputBoxWidth,height:h.inputBoxHeight,"text-align":"center"}).on("click",function(){r.showInput(a);r[a+"Input"].focus()});c.styledMode||k.attr({stroke:h.inputBoxBorderColor,"stroke-width":1});k.add(e);var x=l("input",{name:a,className:"highcharts-range-selector"},void 0,f);x.setAttribute("type",w(h.inputDateFormat||"%b %e, %Y"));c.styledMode||(p.css(v(q,h.labelStyle)),k.css(v({color:"#333333"},q,h.inputStyle)),g(x,b({position:"absolute",border:0,boxShadow:"0 0 15px rgba(0,0,0,0.3)",width:"1px",
height:"1px",padding:0,textAlign:"center",fontSize:q.fontSize,fontFamily:q.fontFamily,top:"-9999em"},h.inputStyle)));x.onfocus=function(){r.showInput(a)};x.onblur=function(){x===m.doc.activeElement&&d();r.hideInput(a);r.setInputValue(a);x.blur()};var A=!1;x.onchange=function(){A||(d(),r.hideInput(a),x.blur())};x.onkeypress=function(a){13===a.keyCode&&d()};x.onkeydown=function(a){A=!0;38!==a.keyCode&&40!==a.keyCode||d()};x.onkeyup=function(){A=!1};return{dateBox:k,input:x,label:p}};f.prototype.getPosition=
function(){var a=this.chart,b=a.options.rangeSelector;a="top"===b.verticalAlign?a.plotTop-a.axisOffset[0]:0;return{buttonTop:a+b.buttonPosition.y,inputTop:a+b.inputPosition.y-10}};f.prototype.getYTDExtremes=function(a,b,d){var c=this.chart.time,f=new c.Date(a),e=c.get("FullYear",f);d=d?c.Date.UTC(e,0,1):+new c.Date(e,0,1);b=Math.max(b,d);f=f.getTime();return{max:Math.min(a||f,f),min:b}};f.prototype.render=function(a,b){var d=this.chart,f=d.renderer,e=d.container,g=d.options,r=g.rangeSelector,q=x(g.chart.style&&
g.chart.style.zIndex,0)+1;g=r.inputEnabled;if(!1!==r.enabled){this.rendered||(this.group=f.g("range-selector-group").attr({zIndex:7}).add(),this.div=l("div",void 0,{position:"relative",height:0,zIndex:q}),this.buttonOptions.length&&this.renderButtons(),e.parentNode&&e.parentNode.insertBefore(this.div,e),g&&(this.inputGroup=f.g("input-group").add(this.group),f=this.drawInput("min"),this.minDateBox=f.dateBox,this.minLabel=f.label,this.minInput=f.input,f=this.drawInput("max"),this.maxDateBox=f.dateBox,
this.maxLabel=f.label,this.maxInput=f.input));if(g&&(this.setInputValue("min",a),this.setInputValue("max",b),a=d.scroller&&d.scroller.getUnionExtremes()||d.xAxis[0]||{},c(a.dataMin)&&c(a.dataMax)&&(d=d.xAxis[0].minRange||0,this.setInputExtremes("min",a.dataMin,Math.min(a.dataMax,this.getInputValue("max"))-d),this.setInputExtremes("max",Math.max(a.dataMin,this.getInputValue("min"))+d,a.dataMax)),this.inputGroup)){var k=0;[this.minLabel,this.minDateBox,this.maxLabel,this.maxDateBox].forEach(function(a){if(a){var b=
a.getBBox().width;b&&(a.attr({x:k}),k+=b+r.inputSpacing)}})}this.alignElements();this.rendered=!0}};f.prototype.renderButtons=function(){var a=this,b=this.buttons,d=this.options,c=t.lang,f=this.chart.renderer,e=v(d.buttonTheme),g=e&&e.states,q=e.width||28;delete e.width;delete e.states;this.buttonGroup=f.g("range-selector-buttons").add(this.group);var h=this.dropdown=l("select",void 0,{position:"absolute",width:"1px",height:"1px",padding:0,border:0,top:"-9999em",cursor:"pointer",opacity:.0001},this.div);
k(h,"touchstart",function(){h.style.fontSize="16px"});[[m.isMS?"mouseover":"mouseenter"],[m.isMS?"mouseout":"mouseleave"],["change","click"]].forEach(function(d){var c=d[0],f=d[1];k(h,c,function(){var d=b[a.currentButtonIndex()];d&&A(d.element,f||c)})});this.zoomText=f.label(c&&c.rangeSelectorZoom||"",0).attr({padding:d.buttonTheme.padding,height:d.buttonTheme.height,paddingLeft:0,paddingRight:0}).add(this.buttonGroup);this.chart.styledMode||(this.zoomText.css(d.labelStyle),e["stroke-width"]=x(e["stroke-width"],
0));l("option",{textContent:this.zoomText.textStr,disabled:!0},void 0,h);this.buttonOptions.forEach(function(d,c){l("option",{textContent:d.title||d.text},void 0,h);b[c]=f.button(d.text,0,0,function(b){var f=d.events&&d.events.click,e;f&&(e=f.call(d,b));!1!==e&&a.clickButton(c);a.isActive=!0},e,g&&g.hover,g&&g.select,g&&g.disabled).attr({"text-align":"center",width:q}).add(a.buttonGroup);d.title&&b[c].attr("title",d.title)})};f.prototype.alignElements=function(){var a=this,b=this.buttonGroup,d=this.buttons,
c=this.chart,f=this.group,e=this.inputGroup,g=this.options,l=this.zoomText,q=c.options,k=q.exporting&&!1!==q.exporting.enabled&&q.navigation&&q.navigation.buttonOptions;q=g.buttonPosition;var h=g.inputPosition,n=g.verticalAlign,m=function(b,d){return k&&a.titleCollision(c)&&"top"===n&&"right"===d.align&&d.y-b.getBBox().height-12<(k.y||0)+(k.height||0)+c.spacing[0]?-40:0},p=c.plotLeft;if(f&&q&&h){var A=q.x-c.spacing[3];if(b){this.positionButtons();if(!this.initialButtonGroupWidth){var t=0;l&&(t+=l.getBBox().width+
5);d.forEach(function(a,b){t+=a.width;b!==d.length-1&&(t+=g.buttonSpacing)});this.initialButtonGroupWidth=t}p-=c.spacing[3];this.updateButtonStates();l=m(b,q);this.alignButtonGroup(l);f.placed=b.placed=c.hasLoaded}b=0;e&&(b=m(e,h),"left"===h.align?A=p:"right"===h.align&&(A=-Math.max(c.axisOffset[1],-b)),e.align({y:h.y,width:e.getBBox().width,align:h.align,x:h.x+A-2},!0,c.spacingBox),e.placed=c.hasLoaded);this.handleCollision(b);f.align({verticalAlign:n},!0,c.spacingBox);e=f.alignAttr.translateY;b=
f.getBBox().height+20;m=0;"bottom"===n&&(m=(m=c.legend&&c.legend.options)&&"bottom"===m.verticalAlign&&m.enabled&&!m.floating?c.legend.legendHeight+x(m.margin,10):0,b=b+m-20,m=e-b-(g.floating?0:g.y)-(c.titleOffset?c.titleOffset[2]:0)-10);if("top"===n)g.floating&&(m=0),c.titleOffset&&c.titleOffset[0]&&(m=c.titleOffset[0]),m+=c.margin[0]-c.spacing[0]||0;else if("middle"===n)if(h.y===q.y)m=e;else if(h.y||q.y)m=0>h.y||0>q.y?m-Math.min(h.y,q.y):e-b;f.translate(g.x,g.y+Math.floor(m));q=this.minInput;h=
this.maxInput;e=this.dropdown;g.inputEnabled&&q&&h&&(q.style.marginTop=f.translateY+"px",h.style.marginTop=f.translateY+"px");e&&(e.style.marginTop=f.translateY+"px")}};f.prototype.alignButtonGroup=function(a,b){var d=this.chart,c=this.buttonGroup,f=this.options.buttonPosition,e=d.plotLeft-d.spacing[3],g=f.x-d.spacing[3];"right"===f.align?g+=a-e:"center"===f.align&&(g-=e/2);c&&c.align({y:f.y,width:x(b,this.initialButtonGroupWidth),align:f.align,x:g},!0,d.spacingBox)};f.prototype.positionButtons=function(){var a=
this.buttons,b=this.chart,d=this.options,c=this.zoomText,f=b.hasLoaded?"animate":"attr",e=d.buttonPosition,g=b.plotLeft,q=g;c&&"hidden"!==c.visibility&&(c[f]({x:x(g+e.x,g)}),q+=e.x+c.getBBox().width+5);this.buttonOptions.forEach(function(b,c){if("hidden"!==a[c].visibility)a[c][f]({x:q}),q+=a[c].width+d.buttonSpacing;else a[c][f]({x:g})})};f.prototype.handleCollision=function(a){var b=this,d=this.chart,c=this.buttonGroup,f=this.inputGroup,e=this.options,g=e.buttonPosition,q=e.dropdown,l=e.inputPosition;
e=function(){var a=0;b.buttons.forEach(function(b){b=b.getBBox();b.width>a&&(a=b.width)});return a};var r=function(b){if(f&&c){var d=f.alignAttr.translateX+f.alignOptions.x-a+f.getBBox().x+2,e=f.alignOptions.width,q=c.alignAttr.translateX+c.getBBox().x;return q+b>d&&d+e>q&&g.y<l.y+f.getBBox().height}return!1},h=function(){f&&c&&f.attr({translateX:f.alignAttr.translateX+(d.axisOffset[1]>=-a?0:-a),translateY:f.alignAttr.translateY+c.getBBox().height+10})};if(c){if("always"===q){this.collapseButtons(a);
r(e())&&h();return}"never"===q&&this.expandButtons()}f&&c?l.align===g.align||r(this.initialButtonGroupWidth+20)?"responsive"===q?(this.collapseButtons(a),r(e())&&h()):h():"responsive"===q&&this.expandButtons():c&&"responsive"===q&&(this.initialButtonGroupWidth>d.plotWidth?this.collapseButtons(a):this.expandButtons())};f.prototype.collapseButtons=function(a){var b=this.buttons,d=this.buttonOptions,c=this.chart,f=this.dropdown,e=this.options,g=this.zoomText,q=c.userOptions.rangeSelector&&c.userOptions.rangeSelector.buttonTheme||
{},l=function(a){return{text:a?a+" \u25be":"\u25be",width:"auto",paddingLeft:x(e.buttonTheme.paddingLeft,q.padding,8),paddingRight:x(e.buttonTheme.paddingRight,q.padding,8)}};g&&g.hide();var r=!1;d.forEach(function(a,d){d=b[d];2!==d.state?d.hide():(d.show(),d.attr(l(a.text)),r=!0)});r||(f&&(f.selectedIndex=0),b[0].show(),b[0].attr(l(this.zoomText&&this.zoomText.textStr)));d=e.buttonPosition.align;this.positionButtons();"right"!==d&&"center"!==d||this.alignButtonGroup(a,b[this.currentButtonIndex()].getBBox().width);
this.showDropdown()};f.prototype.expandButtons=function(){var a=this.buttons,b=this.buttonOptions,d=this.options,c=this.zoomText;this.hideDropdown();c&&c.show();b.forEach(function(b,c){c=a[c];c.show();c.attr({text:b.text,width:d.buttonTheme.width||28,paddingLeft:x(d.buttonTheme.paddingLeft,"unset"),paddingRight:x(d.buttonTheme.paddingRight,"unset")});2>c.state&&c.setState(0)});this.positionButtons()};f.prototype.currentButtonIndex=function(){var a=this.dropdown;return a&&0<a.selectedIndex?a.selectedIndex-
1:0};f.prototype.showDropdown=function(){var a=this.buttonGroup,b=this.buttons,d=this.chart,c=this.dropdown;if(a&&c){var f=a.translateX;a=a.translateY;b=b[this.currentButtonIndex()].getBBox();g(c,{left:d.plotLeft+f+"px",top:a+.5+"px",width:b.width+"px",height:b.height+"px"});this.hasVisibleDropdown=!0}};f.prototype.hideDropdown=function(){var a=this.dropdown;a&&(g(a,{top:"-9999em",width:"1px",height:"1px"}),this.hasVisibleDropdown=!1)};f.prototype.getHeight=function(){var a=this.options,b=this.group,
d=a.y,c=a.buttonPosition.y,f=a.inputPosition.y;if(a.height)return a.height;this.alignElements();a=b?b.getBBox(!0).height+13+d:0;b=Math.min(f,c);if(0>f&&0>c||0<f&&0<c)a+=Math.abs(b);return a};f.prototype.titleCollision=function(a){return!(a.options.title.text||a.options.subtitle.text)};f.prototype.update=function(a){var b=this.chart;v(!0,b.options.rangeSelector,a);this.destroy();this.init(b);this.render()};f.prototype.destroy=function(){var a=this,b=a.minInput,c=a.maxInput;a.eventsToUnbind&&(a.eventsToUnbind.forEach(function(a){return a()}),
a.eventsToUnbind=void 0);e(a.buttons);b&&(b.onfocus=b.onblur=b.onchange=null);c&&(c.onfocus=c.onblur=c.onchange=null);C(a,function(b,c){b&&"chart"!==c&&(b instanceof n?b.destroy():b instanceof window.HTMLElement&&d(b));b!==f.prototype[c]&&(a[c]=null)},this)};return f}();B.prototype.defaultButtons=[{type:"month",count:1,text:"1m",title:"View 1 month"},{type:"month",count:3,text:"3m",title:"View 3 months"},{type:"month",count:6,text:"6m",title:"View 6 months"},{type:"ytd",text:"YTD",title:"View year to date"},
{type:"year",count:1,text:"1y",title:"View 1 year"},{type:"all",text:"All",title:"View all"}];B.prototype.inputTypeFormats={"datetime-local":"%Y-%m-%dT%H:%M:%S",date:"%Y-%m-%d",time:"%H:%M:%S"};a.prototype.minFromRange=function(){var a=this.range,b=a.type,d=this.max,c=this.chart.time,f=function(a,d){var f="year"===b?"FullYear":"Month",e=new c.Date(a),g=c.get(f,e);c.set(f,e,g+d);g===c.get(f,e)&&c.set("Date",e,0);return e.getTime()-a};if(z(a)){var e=d-a;var g=a}else e=d+f(d,-a.count),this.chart&&(this.chart.fixedRange=
d-e);var l=x(this.dataMin,Number.MIN_VALUE);z(e)||(e=l);e<=l&&(e=l,"undefined"===typeof g&&(g=f(e,a.count)),this.newMax=Math.min(e+g,this.dataMax));z(d)||(e=void 0);return e};if(!m.RangeSelector){var G=[],K=function(a){function b(){c&&(d=a.xAxis[0].getExtremes(),e=a.legend,l=c&&c.options.verticalAlign,z(d.min)&&c.render(d.min,d.max),e.display&&"top"===l&&l===e.options.verticalAlign&&(g=v(a.spacingBox),g.y="vertical"===e.options.layout?a.plotTop:g.y+c.getHeight(),e.group.placed=!1,e.align(g)))}var d,
c=a.rangeSelector,e,g,l;c&&(f(G,function(b){return b[0]===a})||G.push([a,[k(a.xAxis[0],"afterSetExtremes",function(a){c&&c.render(a.min,a.max)}),k(a,"redraw",b)]]),b())};k(h,"afterGetContainer",function(){this.options.rangeSelector&&this.options.rangeSelector.enabled&&(this.rangeSelector=new B(this))});k(h,"beforeRender",function(){var a=this.axes,b=this.rangeSelector;b&&(z(b.deferredYTDClick)&&(b.clickButton(b.deferredYTDClick),delete b.deferredYTDClick),a.forEach(function(a){a.updateNames();a.setScale()}),
this.getAxisMargins(),b.render(),a=b.options.verticalAlign,b.options.floating||("bottom"===a?this.extraBottomMargin=!0:"middle"!==a&&(this.extraTopMargin=!0)))});k(h,"update",function(a){var b=a.options.rangeSelector;a=this.rangeSelector;var d=this.extraBottomMargin,f=this.extraTopMargin;b&&b.enabled&&!c(a)&&this.options.rangeSelector&&(this.options.rangeSelector.enabled=!0,this.rangeSelector=a=new B(this));this.extraTopMargin=this.extraBottomMargin=!1;a&&(K(this),b=b&&b.verticalAlign||a.options&&
a.options.verticalAlign,a.options.floating||("bottom"===b?this.extraBottomMargin=!0:"middle"!==b&&(this.extraTopMargin=!0)),this.extraBottomMargin!==d||this.extraTopMargin!==f)&&(this.isDirtyBox=!0)});k(h,"render",function(){var a=this.rangeSelector;a&&!a.options.floating&&(a.render(),a=a.options.verticalAlign,"bottom"===a?this.extraBottomMargin=!0:"middle"!==a&&(this.extraTopMargin=!0))});k(h,"getMargins",function(){var a=this.rangeSelector;a&&(a=a.getHeight(),this.extraTopMargin&&(this.plotTop+=
a),this.extraBottomMargin&&(this.marginBottom+=a))});h.prototype.callbacks.push(K);k(h,"destroy",function(){for(var a=0;a<G.length;a++){var b=G[a];if(b[0]===this){b[1].forEach(function(a){return a()});G.splice(a,1);break}}});m.RangeSelector=B}return B});v(a,"Accessibility/Components/RangeSelectorComponent.js",[a["Extensions/RangeSelector.js"],a["Accessibility/AccessibilityComponent.js"],a["Accessibility/Utils/ChartUtilities.js"],a["Accessibility/Utils/Announcer.js"],a["Accessibility/KeyboardNavigationHandler.js"],
a["Core/Utilities.js"]],function(a,h,m,u,n,p){var w=this&&this.__extends||function(){var a=function(c,d){a=Object.setPrototypeOf||{__proto__:[]}instanceof Array&&function(a,d){a.__proto__=d}||function(a,d){for(var b in d)d.hasOwnProperty(b)&&(a[b]=d[b])};return a(c,d)};return function(c,d){function b(){this.constructor=c}a(c,d);c.prototype=null===d?Object.create(d):(b.prototype=d.prototype,new b)}}(),t=m.unhideChartElementFromAT,k=m.getAxisRangeDescription,l=p.addEvent,g=p.attr;h=function(a){function c(){var d=
null!==a&&a.apply(this,arguments)||this;d.announcer=void 0;return d}w(c,a);c.prototype.init=function(){this.announcer=new u(this.chart,"polite")};c.prototype.onChartUpdate=function(){var a=this.chart,b=this,c=a.rangeSelector;c&&(this.updateSelectorVisibility(),this.setDropdownAttrs(),c.buttons&&c.buttons.length&&c.buttons.forEach(function(a){b.setRangeButtonAttrs(a)}),c.maxInput&&c.minInput&&["minInput","maxInput"].forEach(function(d,f){if(d=c[d])t(a,d),b.setRangeInputAttrs(d,"accessibility.rangeSelector."+
(f?"max":"min")+"InputLabel")}))};c.prototype.updateSelectorVisibility=function(){var a=this.chart,b=a.rangeSelector,c=b&&b.dropdown,e=b&&b.buttons||[];b&&b.hasVisibleDropdown&&c?(t(a,c),e.forEach(function(a){return a.element.setAttribute("aria-hidden",!0)})):(c&&c.setAttribute("aria-hidden",!0),e.forEach(function(b){return t(a,b.element)}))};c.prototype.setDropdownAttrs=function(){var a=this.chart,b=a.rangeSelector&&a.rangeSelector.dropdown;b&&(a=a.langFormat("accessibility.rangeSelector.dropdownLabel",
{rangeTitle:a.options.lang.rangeSelectorZoom}),b.setAttribute("aria-label",a),b.setAttribute("tabindex",-1))};c.prototype.setRangeButtonAttrs=function(a){g(a.element,{tabindex:-1,role:"button"})};c.prototype.setRangeInputAttrs=function(a,b){var c=this.chart;g(a,{tabindex:-1,"aria-label":c.langFormat(b,{chart:c})})};c.prototype.onButtonNavKbdArrowKey=function(a,b){var c=a.response,d=this.keyCodes,e=this.chart,g=e.options.accessibility.keyboardNavigation.wrapAround;b=b===d.left||b===d.up?-1:1;return e.highlightRangeSelectorButton(e.highlightedRangeSelectorItemIx+
b)?c.success:g?(a.init(b),c.success):c[0<b?"next":"prev"]};c.prototype.onButtonNavKbdClick=function(a){a=a.response;var b=this.chart;3!==b.oldRangeSelectorItemState&&this.fakeClickEvent(b.rangeSelector.buttons[b.highlightedRangeSelectorItemIx].element);return a.success};c.prototype.onAfterBtnClick=function(){var a=this.chart,b=k(a.xAxis[0]);(a=a.langFormat("accessibility.rangeSelector.clickButtonAnnouncement",{chart:a,axisRangeDescription:b}))&&this.announcer.announce(a)};c.prototype.onInputKbdMove=
function(a){var b=this.chart,c=b.rangeSelector,d=b.highlightedInputRangeIx=(b.highlightedInputRangeIx||0)+a;1<d||0>d?b.accessibility&&(b.accessibility.keyboardNavigation.tabindexContainer.focus(),b.accessibility.keyboardNavigation[0>a?"prev":"next"]()):c&&(a=c[d?"maxDateBox":"minDateBox"],c=c[d?"maxInput":"minInput"],a&&c&&b.setFocusToElement(a,c))};c.prototype.onInputNavInit=function(a){var b=this,c=this,d=this.chart,e=0<a?0:1,g=d.rangeSelector,h=g&&g[e?"maxDateBox":"minDateBox"];a=g&&g.minInput;
g=g&&g.maxInput;d.highlightedInputRangeIx=e;if(h&&a&&g){d.setFocusToElement(h,e?g:a);this.removeInputKeydownHandler&&this.removeInputKeydownHandler();d=function(a){(a.which||a.keyCode)===b.keyCodes.tab&&(a.preventDefault(),a.stopPropagation(),c.onInputKbdMove(a.shiftKey?-1:1))};var k=l(a,"keydown",d),m=l(g,"keydown",d);this.removeInputKeydownHandler=function(){k();m()}}};c.prototype.onInputNavTerminate=function(){var a=this.chart.rangeSelector||{};a.maxInput&&a.hideInput("max");a.minInput&&a.hideInput("min");
this.removeInputKeydownHandler&&(this.removeInputKeydownHandler(),delete this.removeInputKeydownHandler)};c.prototype.initDropdownNav=function(){var a=this,b=this.chart,c=b.rangeSelector,e=c&&c.dropdown;c&&e&&(b.setFocusToElement(c.buttonGroup,e),this.removeDropdownKeydownHandler&&this.removeDropdownKeydownHandler(),this.removeDropdownKeydownHandler=l(e,"keydown",function(c){(c.which||c.keyCode)===a.keyCodes.tab&&(c.preventDefault(),c.stopPropagation(),b.accessibility&&(b.accessibility.keyboardNavigation.tabindexContainer.focus(),
b.accessibility.keyboardNavigation[c.shiftKey?"prev":"next"]()))}))};c.prototype.getRangeSelectorButtonNavigation=function(){var a=this.chart,b=this.keyCodes,c=this;return new n(a,{keyCodeMap:[[[b.left,b.right,b.up,b.down],function(a){return c.onButtonNavKbdArrowKey(this,a)}],[[b.enter,b.space],function(){return c.onButtonNavKbdClick(this)}]],validate:function(){return!!(a.rangeSelector&&a.rangeSelector.buttons&&a.rangeSelector.buttons.length)},init:function(b){var d=a.rangeSelector;d&&d.hasVisibleDropdown?
c.initDropdownNav():d&&(d=d.buttons.length-1,a.highlightRangeSelectorButton(0<b?0:d))},terminate:function(){c.removeDropdownKeydownHandler&&(c.removeDropdownKeydownHandler(),delete c.removeDropdownKeydownHandler)}})};c.prototype.getRangeSelectorInputNavigation=function(){var a=this.chart,b=this;return new n(a,{keyCodeMap:[],validate:function(){return!!(a.rangeSelector&&a.rangeSelector.inputGroup&&"hidden"!==a.rangeSelector.inputGroup.element.style.visibility&&!1!==a.options.rangeSelector.inputEnabled&&
a.rangeSelector.minInput&&a.rangeSelector.maxInput)},init:function(a){b.onInputNavInit(a)},terminate:function(){b.onInputNavTerminate()}})};c.prototype.getKeyboardNavigation=function(){return[this.getRangeSelectorButtonNavigation(),this.getRangeSelectorInputNavigation()]};c.prototype.destroy=function(){this.removeDropdownKeydownHandler&&this.removeDropdownKeydownHandler();this.removeInputKeydownHandler&&this.removeInputKeydownHandler();this.announcer&&this.announcer.destroy()};return c}(h);(function(c){function e(a){var b=
this.rangeSelector&&this.rangeSelector.buttons||[],c=this.highlightedRangeSelectorItemIx,d=this.rangeSelector&&this.rangeSelector.selected;"undefined"!==typeof c&&b[c]&&c!==d&&b[c].setState(this.oldRangeSelectorItemState||0);this.highlightedRangeSelectorItemIx=a;return b[a]?(this.setFocusToElement(b[a].box,b[a].element),a!==d&&(this.oldRangeSelectorItemState=b[a].state,b[a].setState(1)),!0):!1}function d(){if(this.chart.accessibility&&this.chart.accessibility.components.rangeSelector)return this.chart.accessibility.components.rangeSelector.onAfterBtnClick()}
var b=[];c.compose=function(c,g){-1===b.indexOf(c)&&(b.push(c),c.prototype.highlightRangeSelectorButton=e);-1===b.indexOf(g)&&(b.push(g),l(a,"afterBtnClick",d))}})(h||(h={}));return h});v(a,"Accessibility/Components/SeriesComponent/ForcedMarkers.js",[a["Core/Utilities.js"]],function(a){var h=a.addEvent,m=a.merge,u;(function(a){function n(a){m(!0,a,{marker:{enabled:!0,states:{normal:{opacity:0}}}})}function u(a){return a.marker.states&&a.marker.states.normal&&a.marker.states.normal.opacity}function t(){if(this.chart.styledMode){if(this.markerGroup)this.markerGroup[this.a11yMarkersForced?
"addClass":"removeClass"]("highcharts-a11y-markers-hidden");this._hasPointMarkers&&this.points&&this.points.length&&this.points.forEach(function(a){a.graphic&&(a.graphic[a.hasForcedA11yMarker?"addClass":"removeClass"]("highcharts-a11y-marker-hidden"),a.graphic[!1===a.hasForcedA11yMarker?"addClass":"removeClass"]("highcharts-a11y-marker-visible"))})}}function k(a){this.resetA11yMarkerOptions=m(a.options.marker||{},this.userOptions.marker||{})}function l(){var a=this.options,e=!1!==(this.options.accessibility&&
this.options.accessibility.enabled);if(e=this.chart.options.accessibility.enabled&&e)e=this.chart.options.accessibility,e=this.points.length<e.series.pointDescriptionEnabledThreshold||!1===e.series.pointDescriptionEnabledThreshold;if(e){if(a.marker&&!1===a.marker.enabled&&(this.a11yMarkersForced=!0,n(this.options)),this._hasPointMarkers&&this.points&&this.points.length)for(a=this.points.length;a--;){e=this.points[a];var d=e.options,b=e.hasForcedA11yMarker;delete e.hasForcedA11yMarker;d.marker&&(b=
b&&0===u(d),d.marker.enabled&&!b?(m(!0,d.marker,{states:{normal:{opacity:u(d)||1}}}),e.hasForcedA11yMarker=!1):!1===d.marker.enabled&&(n(d),e.hasForcedA11yMarker=!0))}}else this.a11yMarkersForced&&(delete this.a11yMarkersForced,(a=this.resetA11yMarkerOptions)&&m(!0,this.options,{marker:{enabled:a.enabled,states:{normal:{opacity:a.states&&a.states.normal&&a.states.normal.opacity}}}}))}var g=[];a.compose=function(a){-1===g.indexOf(a)&&(g.push(a),h(a,"afterSetOptions",k),h(a,"render",l),h(a,"afterRender",
t))}})(u||(u={}));return u});v(a,"Accessibility/Components/SeriesComponent/SeriesKeyboardNavigation.js",[a["Core/Series/Point.js"],a["Core/Series/Series.js"],a["Core/Series/SeriesRegistry.js"],a["Core/Globals.js"],a["Core/Utilities.js"],a["Accessibility/KeyboardNavigationHandler.js"],a["Accessibility/Utils/EventProvider.js"],a["Accessibility/Utils/ChartUtilities.js"]],function(a,h,m,u,n,p,v,t){function k(a){var b=a.index,c=a.series.points,d=c.length;if(c[b]!==a)for(;d--;){if(c[d]===a)return d}else return b}
function l(a){var b=a.chart.options.accessibility.keyboardNavigation.seriesNavigation,c=a.options.accessibility||{},d=c.keyboardNavigation;return d&&!1===d.enabled||!1===c.enabled||!1===a.options.enableMouseTracking||!a.visible||b.pointNavigationEnabledThreshold&&b.pointNavigationEnabledThreshold<=a.points.length}function g(a){var b=a.series.chart.options.accessibility,c=a.options.accessibility&&!1===a.options.accessibility.enabled;return a.isNull&&b.keyboardNavigation.seriesNavigation.skipNullPoints||
!1===a.visible||!1===a.isInside||c||l(a.series)}function c(a){var b=!1;delete a.highlightedPoint;return b=a.series.reduce(function(a,b){return a||b.highlightFirstValidPoint()},!1)}var e=m.seriesTypes,d=u.doc,b=n.defined,f=n.fireEvent,w=t.getPointFromXY,z=t.getSeriesFromName,E=t.scrollToPoint;m=function(){function e(a,b){this.keyCodes=b;this.chart=a}e.prototype.init=function(){var b=this,e=this.chart,f=this.eventProvider=new v;f.addEvent(h,"destroy",function(){return b.onSeriesDestroy(this)});f.addEvent(e,
"afterDrilldown",function(){c(this);this.focusElement&&this.focusElement.removeFocusBorder()});f.addEvent(e,"drilldown",function(a){a=a.point;var c=a.series;b.lastDrilledDownPoint={x:a.x,y:a.y,seriesName:c?c.name:""}});f.addEvent(e,"drillupall",function(){setTimeout(function(){b.onDrillupAll()},10)});f.addEvent(a,"afterSetState",function(){var a=this.graphic&&this.graphic.element,b=d.activeElement,c=b&&b.getAttribute("class");c=c&&-1<c.indexOf("highcharts-a11y-proxy-button");e.highlightedPoint===
this&&b!==a&&!c&&a&&a.focus&&a.focus()})};e.prototype.onDrillupAll=function(){var a=this.lastDrilledDownPoint,c=this.chart,d=a&&z(c,a.seriesName),e;a&&d&&b(a.x)&&b(a.y)&&(e=w(d,a.x,a.y));c.container&&c.container.focus();e&&e.highlight&&e.highlight();c.focusElement&&c.focusElement.removeFocusBorder()};e.prototype.getKeyboardNavigationHandler=function(){var a=this,b=this.keyCodes,d=this.chart,e=d.inverted;return new p(d,{keyCodeMap:[[e?[b.up,b.down]:[b.left,b.right],function(b){return a.onKbdSideways(this,
b)}],[e?[b.left,b.right]:[b.up,b.down],function(b){return a.onKbdVertical(this,b)}],[[b.enter,b.space],function(a,b){if(a=d.highlightedPoint)b.point=a,f(a.series,"click",b),a.firePointEvent("click");return this.response.success}],[[b.home],function(){c(d);return this.response.success}],[[b.end],function(){for(var a=d.series.length,b;a--&&!(d.highlightedPoint=d.series[a].points[d.series[a].points.length-1],b=d.series[a].highlightFirstValidPoint()););return this.response.success}],[[b.pageDown,b.pageUp],
function(a){d.highlightAdjacentSeries(a===b.pageDown);return this.response.success}]],init:function(){c(d);return this.response.success},terminate:function(){return a.onHandlerTerminate()}})};e.prototype.onKbdSideways=function(a,b){var c=this.keyCodes;return this.attemptHighlightAdjacentPoint(a,b===c.right||b===c.down)};e.prototype.onKbdVertical=function(a,b){var c=this.chart,d=this.keyCodes;b=b===d.down||b===d.right;d=c.options.accessibility.keyboardNavigation.seriesNavigation;if(d.mode&&"serialize"===
d.mode)return this.attemptHighlightAdjacentPoint(a,b);c[c.highlightedPoint&&c.highlightedPoint.series.keyboardMoveVertical?"highlightAdjacentPointVertical":"highlightAdjacentSeries"](b);return a.response.success};e.prototype.onHandlerTerminate=function(){var a=this.chart;a.tooltip&&a.tooltip.hide(0);var b=a.highlightedPoint&&a.highlightedPoint.series;if(b&&b.onMouseOut)b.onMouseOut();if(a.highlightedPoint&&a.highlightedPoint.onMouseOut)a.highlightedPoint.onMouseOut();delete a.highlightedPoint};e.prototype.attemptHighlightAdjacentPoint=
function(a,b){var c=this.chart,d=c.options.accessibility.keyboardNavigation.wrapAround;return c.highlightAdjacentPoint(b)?a.response.success:d?a.init(b?1:-1):a.response[b?"next":"prev"]};e.prototype.onSeriesDestroy=function(a){var b=this.chart;b.highlightedPoint&&b.highlightedPoint.series===a&&(delete b.highlightedPoint,b.focusElement&&b.focusElement.removeFocusBorder())};e.prototype.destroy=function(){this.eventProvider.removeAddedEvents()};return e}();(function(a){function c(a){var b=this.series,
c=this.highlightedPoint,d=c&&k(c)||0,e=c&&c.series.points||[],f=this.series&&this.series[this.series.length-1];f=f&&f.points&&f.points[f.points.length-1];if(!b[0]||!b[0].points)return!1;if(c){if(b=b[c.series.index+(a?1:-1)],d=e[d+(a?1:-1)],!d&&b&&(d=b.points[a?0:b.points.length-1]),!d)return!1}else d=a?b[0].points[0]:f;return g(d)?(b=d.series,l(b)?this.highlightedPoint=a?b.points[b.points.length-1]:b.points[0]:this.highlightedPoint=d,this.highlightAdjacentPoint(a)):d.highlight()}function d(a){var c=
this.highlightedPoint,d=Infinity,e;if(!b(c.plotX)||!b(c.plotY))return!1;this.series.forEach(function(f){l(f)||f.points.forEach(function(l){if(b(l.plotY)&&b(l.plotX)&&l!==c){var h=l.plotY-c.plotY,k=Math.abs(l.plotX-c.plotX);k=Math.abs(h)*Math.abs(h)+k*k*4;f.yAxis&&f.yAxis.reversed&&(h*=-1);!(0>=h&&a||0<=h&&!a||5>k||g(l))&&k<d&&(d=k,e=l)}})});return e?e.highlight():!1}function f(a){var b=this.highlightedPoint,c=this.series&&this.series[this.series.length-1],d=c&&c.points&&c.points[c.points.length-1];
if(!this.highlightedPoint)return c=a?this.series&&this.series[0]:c,(d=a?c&&c.points&&c.points[0]:d)?d.highlight():!1;c=this.series[b.series.index+(a?-1:1)];if(!c)return!1;d=h(b,c,4);if(!d)return!1;if(l(c))return d.highlight(),a=this.highlightAdjacentSeries(a),a?a:(b.highlight(),!1);d.highlight();return d.series.highlightFirstValidPoint()}function h(a,c,d,e){var f=Infinity,g=c.points.length,l=function(a){return!(b(a.plotX)&&b(a.plotY))};if(!l(a)){for(;g--;){var h=c.points[g];if(!l(h)&&(h=(a.plotX-
h.plotX)*(a.plotX-h.plotX)*(d||1)+(a.plotY-h.plotY)*(a.plotY-h.plotY)*(e||1),h<f)){f=h;var k=g}}return b(k)?c.points[k]:void 0}}function m(){var a=this.series.chart;if(this.isNull)a.tooltip&&a.tooltip.hide(0);else this.onMouseOver();E(this);this.graphic&&a.setFocusToElement(this.graphic);a.highlightedPoint=this;return this}function n(){var a=this.chart.highlightedPoint,b=(a&&a.series)===this?k(a):0;a=this.points;var c=a.length;if(a&&c){for(var d=b;d<c;++d)if(!g(a[d]))return a[d].highlight();for(;0<=
b;--b)if(!g(a[b]))return a[b].highlight()}return!1}var p=[];a.compose=function(a,b,g){-1===p.indexOf(a)&&(p.push(a),a=a.prototype,a.highlightAdjacentPoint=c,a.highlightAdjacentPointVertical=d,a.highlightAdjacentSeries=f);-1===p.indexOf(b)&&(p.push(b),b.prototype.highlight=m);-1===p.indexOf(g)&&(p.push(g),b=g.prototype,b.keyboardMoveVertical=!0,["column","gantt","pie"].forEach(function(a){e[a]&&(e[a].prototype.keyboardMoveVertical=!1)}),b.highlightFirstValidPoint=n)}})(m||(m={}));return m});v(a,"Accessibility/Components/SeriesComponent/SeriesComponent.js",
[a["Accessibility/AccessibilityComponent.js"],a["Accessibility/Utils/ChartUtilities.js"],a["Accessibility/Components/SeriesComponent/ForcedMarkers.js"],a["Accessibility/Components/SeriesComponent/NewDataAnnouncer.js"],a["Accessibility/Components/SeriesComponent/SeriesDescriber.js"],a["Accessibility/Components/SeriesComponent/SeriesKeyboardNavigation.js"],a["Core/Tooltip.js"]],function(a,h,m,u,n,p,v){var t=this&&this.__extends||function(){var a=function(c,e){a=Object.setPrototypeOf||{__proto__:[]}instanceof
Array&&function(a,b){a.__proto__=b}||function(a,b){for(var c in b)b.hasOwnProperty(c)&&(a[c]=b[c])};return a(c,e)};return function(c,e){function d(){this.constructor=c}a(c,e);c.prototype=null===e?Object.create(e):(d.prototype=e.prototype,new d)}}(),k=h.hideSeriesFromAT,l=n.describeSeries;return function(a){function c(){return null!==a&&a.apply(this,arguments)||this}t(c,a);c.compose=function(a,c,b){m.compose(b);p.compose(a,c,b)};c.prototype.init=function(){this.newDataAnnouncer=new u(this.chart);this.newDataAnnouncer.init();
this.keyboardNavigation=new p(this.chart,this.keyCodes);this.keyboardNavigation.init();this.hideTooltipFromATWhenShown();this.hideSeriesLabelsFromATWhenShown()};c.prototype.hideTooltipFromATWhenShown=function(){var a=this;this.addEvent(v,"refresh",function(){this.chart===a.chart&&this.label&&this.label.element&&this.label.element.setAttribute("aria-hidden",!0)})};c.prototype.hideSeriesLabelsFromATWhenShown=function(){this.addEvent(this.chart,"afterDrawSeriesLabels",function(){this.series.forEach(function(a){a.labelBySeries&&
a.labelBySeries.attr("aria-hidden",!0)})})};c.prototype.onChartRender=function(){this.chart.series.forEach(function(a){!1!==(a.options.accessibility&&a.options.accessibility.enabled)&&a.visible?l(a):k(a)})};c.prototype.getKeyboardNavigation=function(){return this.keyboardNavigation.getKeyboardNavigationHandler()};c.prototype.destroy=function(){this.newDataAnnouncer.destroy();this.keyboardNavigation.destroy()};return c}(a)});v(a,"Accessibility/Components/ZoomComponent.js",[a["Accessibility/AccessibilityComponent.js"],
a["Accessibility/Utils/ChartUtilities.js"],a["Core/Globals.js"],a["Accessibility/KeyboardNavigationHandler.js"],a["Core/Utilities.js"]],function(a,h,m,u,n){var p=h.unhideChartElementFromAT;h=m.noop;var w=n.attr,t=n.extend,k=n.pick;m.Axis.prototype.panStep=function(a,g){var c=g||3;g=this.getExtremes();var e=(g.max-g.min)/c*a;c=g.max+e;e=g.min+e;var d=c-e;0>a&&e<g.dataMin?(e=g.dataMin,c=e+d):0<a&&c>g.dataMax&&(c=g.dataMax,e=c-d);this.setExtremes(e,c)};h.prototype=new a;t(h.prototype,{init:function(){var a=
this,g=this.chart;this.proxyProvider.addGroup("zoom","div");["afterShowResetZoom","afterDrilldown","drillupall"].forEach(function(c){a.addEvent(g,c,function(){a.updateProxyOverlays()})})},onChartUpdate:function(){var a=this.chart,g=this;a.mapNavButtons&&a.mapNavButtons.forEach(function(c,e){p(a,c.element);g.setMapNavButtonAttrs(c.element,"accessibility.zoom.mapZoom"+(e?"Out":"In"))})},setMapNavButtonAttrs:function(a,g){var c=this.chart;g=c.langFormat(g,{chart:c});w(a,{tabindex:-1,role:"button","aria-label":g})},
onChartRender:function(){this.updateProxyOverlays()},updateProxyOverlays:function(){var a=this.chart;this.proxyProvider.clearGroup("zoom");a.resetZoomButton&&this.createZoomProxyButton(a.resetZoomButton,"resetZoomProxyButton",a.langFormat("accessibility.zoom.resetZoomButton",{chart:a}));a.drillUpButton&&this.createZoomProxyButton(a.drillUpButton,"drillUpProxyButton",a.langFormat("accessibility.drillUpButton",{chart:a,buttonText:a.getDrilldownBackText()}))},createZoomProxyButton:function(a,g,c){this[g]=
this.proxyProvider.addProxyElement("zoom",{click:a},{"aria-label":c,tabindex:-1})},getMapZoomNavigation:function(){var a=this.keyCodes,g=this.chart,c=this;return new u(g,{keyCodeMap:[[[a.up,a.down,a.left,a.right],function(a){return c.onMapKbdArrow(this,a)}],[[a.tab],function(a,d){return c.onMapKbdTab(this,d)}],[[a.space,a.enter],function(){return c.onMapKbdClick(this)}]],validate:function(){return!!(g.mapZoom&&g.mapNavButtons&&g.mapNavButtons.length)},init:function(a){return c.onMapNavInit(a)}})},
onMapKbdArrow:function(a,g){var c=this.keyCodes;this.chart[g===c.up||g===c.down?"yAxis":"xAxis"][0].panStep(g===c.left||g===c.up?-1:1);return a.response.success},onMapKbdTab:function(a,g){var c=this.chart;a=a.response;var e=(g=g.shiftKey)&&!this.focusedMapNavButtonIx||!g&&this.focusedMapNavButtonIx;c.mapNavButtons[this.focusedMapNavButtonIx].setState(0);if(e)return c.mapZoom(),a[g?"prev":"next"];this.focusedMapNavButtonIx+=g?-1:1;g=c.mapNavButtons[this.focusedMapNavButtonIx];c.setFocusToElement(g.box,
g.element);g.setState(2);return a.success},onMapKbdClick:function(a){this.fakeClickEvent(this.chart.mapNavButtons[this.focusedMapNavButtonIx].element);return a.response.success},onMapNavInit:function(a){var g=this.chart,c=g.mapNavButtons[0],e=g.mapNavButtons[1];c=0<a?c:e;g.setFocusToElement(c.box,c.element);c.setState(2);this.focusedMapNavButtonIx=0<a?0:1},simpleButtonNavigation:function(a,g,c){var e=this.keyCodes,d=this,b=this.chart;return new u(b,{keyCodeMap:[[[e.tab,e.up,e.down,e.left,e.right],
function(a,b){return this.response[a===e.tab&&b.shiftKey||a===e.left||a===e.up?"prev":"next"]}],[[e.space,e.enter],function(){var a=c(this,b);return k(a,this.response.success)}]],validate:function(){return b[a]&&b[a].box&&d[g].buttonElement},init:function(){b.setFocusToElement(b[a].box,d[g].buttonElement)}})},getKeyboardNavigation:function(){return[this.simpleButtonNavigation("resetZoomButton","resetZoomProxyButton",function(a,g){g.zoomOut()}),this.simpleButtonNavigation("drillUpButton","drillUpProxyButton",
function(a,g){g.drillUp();return a.response.prev}),this.getMapZoomNavigation()]}});return h});v(a,"Accessibility/HighContrastMode.js",[a["Core/Globals.js"]],function(a){var h=a.doc,m=a.isMS,u=a.win;return{isHighContrastModeActive:function(){var a=/(Edg)/.test(u.navigator.userAgent);if(u.matchMedia&&a)return u.matchMedia("(-ms-high-contrast: active)").matches;if(m&&u.getComputedStyle){a=h.createElement("div");a.style.backgroundImage="url(data:image/gif;base64,R0lGODlhAQABAAAAACH5BAEKAAEALAAAAAABAAEAAAICTAEAOw==)";
h.body.appendChild(a);var p=(a.currentStyle||u.getComputedStyle(a)).backgroundImage;h.body.removeChild(a);return"none"===p}return!1},setHighContrastTheme:function(a){a.highContrastModeActive=!0;var h=a.options.accessibility.highContrastTheme;a.update(h,!1);a.series.forEach(function(a){var m=h.plotOptions[a.type]||{};a.update({color:m.color||"windowText",colors:[m.color||"windowText"],borderColor:m.borderColor||"window"});a.points.forEach(function(a){a.options&&a.options.color&&a.update({color:m.color||
"windowText",borderColor:m.borderColor||"window"},!1)})});a.redraw()}}});v(a,"Accessibility/HighContrastTheme.js",[],function(){return{chart:{backgroundColor:"window"},title:{style:{color:"windowText"}},subtitle:{style:{color:"windowText"}},colorAxis:{minColor:"windowText",maxColor:"windowText",stops:[]},colors:["windowText"],xAxis:{gridLineColor:"windowText",labels:{style:{color:"windowText"}},lineColor:"windowText",minorGridLineColor:"windowText",tickColor:"windowText",title:{style:{color:"windowText"}}},
yAxis:{gridLineColor:"windowText",labels:{style:{color:"windowText"}},lineColor:"windowText",minorGridLineColor:"windowText",tickColor:"windowText",title:{style:{color:"windowText"}}},tooltip:{backgroundColor:"window",borderColor:"windowText",style:{color:"windowText"}},plotOptions:{series:{lineColor:"windowText",fillColor:"window",borderColor:"windowText",edgeColor:"windowText",borderWidth:1,dataLabels:{connectorColor:"windowText",color:"windowText",style:{color:"windowText",textOutline:"none"}},
marker:{lineColor:"windowText",fillColor:"windowText"}},pie:{color:"window",colors:["window"],borderColor:"windowText",borderWidth:1},boxplot:{fillColor:"window"},candlestick:{lineColor:"windowText",fillColor:"window"},errorbar:{fillColor:"window"}},legend:{backgroundColor:"window",itemStyle:{color:"windowText"},itemHoverStyle:{color:"windowText"},itemHiddenStyle:{color:"#555"},title:{style:{color:"windowText"}}},credits:{style:{color:"windowText"}},labels:{style:{color:"windowText"}},drilldown:{activeAxisLabelStyle:{color:"windowText"},
activeDataLabelStyle:{color:"windowText"}},navigation:{buttonOptions:{symbolStroke:"windowText",theme:{fill:"window"}}},rangeSelector:{buttonTheme:{fill:"window",stroke:"windowText",style:{color:"windowText"},states:{hover:{fill:"window",stroke:"windowText",style:{color:"windowText"}},select:{fill:"#444",stroke:"windowText",style:{color:"windowText"}}}},inputBoxBorderColor:"windowText",inputStyle:{backgroundColor:"window",color:"windowText"},labelStyle:{color:"windowText"}},navigator:{handles:{backgroundColor:"window",
borderColor:"windowText"},outlineColor:"windowText",maskFill:"transparent",series:{color:"windowText",lineColor:"windowText"},xAxis:{gridLineColor:"windowText"}},scrollbar:{barBackgroundColor:"#444",barBorderColor:"windowText",buttonArrowColor:"windowText",buttonBackgroundColor:"window",buttonBorderColor:"windowText",rifleColor:"windowText",trackBackgroundColor:"window",trackBorderColor:"windowText"}}});v(a,"Accessibility/Options/Options.js",[],function(){return{accessibility:{enabled:!0,screenReaderSection:{beforeChartFormat:"<{headingTagName}>{chartTitle}</{headingTagName}><div>{typeDescription}</div><div>{chartSubtitle}</div><div>{chartLongdesc}</div><div>{playAsSoundButton}</div><div>{viewTableButton}</div><div>{xAxisDescription}</div><div>{yAxisDescription}</div><div>{annotationsTitle}{annotationsList}</div>",
afterChartFormat:"{endOfChartMarker}",axisRangeDateFormat:"%Y-%m-%d %H:%M:%S"},series:{describeSingleSeries:!1,pointDescriptionEnabledThreshold:200},point:{valueDescriptionFormat:"{index}. {xDescription}{separator}{value}."},landmarkVerbosity:"all",linkedDescription:'*[data-highcharts-chart="{index}"] + .highcharts-description',keyboardNavigation:{enabled:!0,focusBorder:{enabled:!0,hideBrowserFocusOutline:!0,style:{color:"#335cad",lineWidth:2,borderRadius:3},margin:2},order:["series","zoom","rangeSelector",
"legend","chartMenu"],wrapAround:!0,seriesNavigation:{skipNullPoints:!0,pointNavigationEnabledThreshold:!1}},announceNewData:{enabled:!1,minAnnounceInterval:5E3,interruptUser:!1}},legend:{accessibility:{enabled:!0,keyboardNavigation:{enabled:!0}}},exporting:{accessibility:{enabled:!0}}}});v(a,"Accessibility/Options/LangOptions.js",[],function(){return{accessibility:{defaultChartTitle:"Chart",chartContainerLabel:"{title}. Highcharts interactive chart.",svgContainerLabel:"Interactive chart",drillUpButton:"{buttonText}",
credits:"Chart credits: {creditsStr}",thousandsSep:",",svgContainerTitle:"",graphicContainerLabel:"",screenReaderSection:{beforeRegionLabel:"Chart screen reader information, {chartTitle}.",afterRegionLabel:"",annotations:{heading:"Chart annotations summary",descriptionSinglePoint:"{annotationText}. Related to {annotationPoint}",descriptionMultiplePoints:"{annotationText}. Related to {annotationPoint}{ Also related to, #each(additionalAnnotationPoints)}",descriptionNoPoints:"{annotationText}"},endOfChartMarker:"End of interactive chart."},
sonification:{playAsSoundButtonText:"Play as sound, {chartTitle}",playAsSoundClickAnnouncement:"Play"},legend:{legendLabelNoTitle:"Toggle series visibility, {chartTitle}",legendLabel:"Chart legend: {legendTitle}",legendItem:"Show {itemName}"},zoom:{mapZoomIn:"Zoom chart",mapZoomOut:"Zoom out chart",resetZoomButton:"Reset zoom"},rangeSelector:{dropdownLabel:"{rangeTitle}",minInputLabel:"Select start date.",maxInputLabel:"Select end date.",clickButtonAnnouncement:"Viewing {axisRangeDescription}"},table:{viewAsDataTableButtonText:"View as data table, {chartTitle}",
tableSummary:"Table representation of chart."},announceNewData:{newDataAnnounce:"Updated data for chart {chartTitle}",newSeriesAnnounceSingle:"New data series: {seriesDesc}",newPointAnnounceSingle:"New data point: {pointDesc}",newSeriesAnnounceMultiple:"New data series in chart {chartTitle}: {seriesDesc}",newPointAnnounceMultiple:"New data point in chart {chartTitle}: {pointDesc}"},seriesTypeDescriptions:{boxplot:"Box plot charts are typically used to display groups of statistical data. Each data point in the chart can have up to 5 values: minimum, lower quartile, median, upper quartile, and maximum.",
arearange:"Arearange charts are line charts displaying a range between a lower and higher value for each point.",areasplinerange:"These charts are line charts displaying a range between a lower and higher value for each point.",bubble:"Bubble charts are scatter charts where each data point also has a size value.",columnrange:"Columnrange charts are column charts displaying a range between a lower and higher value for each point.",errorbar:"Errorbar series are used to display the variability of the data.",
funnel:"Funnel charts are used to display reduction of data in stages.",pyramid:"Pyramid charts consist of a single pyramid with item heights corresponding to each point value.",waterfall:"A waterfall chart is a column chart where each column contributes towards a total end value."},chartTypes:{emptyChart:"Empty chart",mapTypeDescription:"Map of {mapTitle} with {numSeries} data series.",unknownMap:"Map of unspecified region with {numSeries} data series.",combinationChart:"Combination chart with {numSeries} data series.",
defaultSingle:"Chart with {numPoints} data {#plural(numPoints, points, point)}.",defaultMultiple:"Chart with {numSeries} data series.",splineSingle:"Line chart with {numPoints} data {#plural(numPoints, points, point)}.",splineMultiple:"Line chart with {numSeries} lines.",lineSingle:"Line chart with {numPoints} data {#plural(numPoints, points, point)}.",lineMultiple:"Line chart with {numSeries} lines.",columnSingle:"Bar chart with {numPoints} {#plural(numPoints, bars, bar)}.",columnMultiple:"Bar chart with {numSeries} data series.",
barSingle:"Bar chart with {numPoints} {#plural(numPoints, bars, bar)}.",barMultiple:"Bar chart with {numSeries} data series.",pieSingle:"Pie chart with {numPoints} {#plural(numPoints, slices, slice)}.",pieMultiple:"Pie chart with {numSeries} pies.",scatterSingle:"Scatter chart with {numPoints} {#plural(numPoints, points, point)}.",scatterMultiple:"Scatter chart with {numSeries} data series.",boxplotSingle:"Boxplot with {numPoints} {#plural(numPoints, boxes, box)}.",boxplotMultiple:"Boxplot with {numSeries} data series.",
bubbleSingle:"Bubble chart with {numPoints} {#plural(numPoints, bubbles, bubble)}.",bubbleMultiple:"Bubble chart with {numSeries} data series."},axis:{xAxisDescriptionSingular:"The chart has 1 X axis displaying {names[0]}. {ranges[0]}",xAxisDescriptionPlural:"The chart has {numAxes} X axes displaying {#each(names, -1) }and {names[-1]}.",yAxisDescriptionSingular:"The chart has 1 Y axis displaying {names[0]}. {ranges[0]}",yAxisDescriptionPlural:"The chart has {numAxes} Y axes displaying {#each(names, -1) }and {names[-1]}.",
timeRangeDays:"Range: {range} days.",timeRangeHours:"Range: {range} hours.",timeRangeMinutes:"Range: {range} minutes.",timeRangeSeconds:"Range: {range} seconds.",rangeFromTo:"Range: {rangeFrom} to {rangeTo}.",rangeCategories:"Range: {numCategories} categories."},exporting:{chartMenuLabel:"Chart menu",menuButtonLabel:"View chart menu, {chartTitle}"},series:{summary:{"default":"{name}, series {ix} of {numSeries} with {numPoints} data {#plural(numPoints, points, point)}.",defaultCombination:"{name}, series {ix} of {numSeries} with {numPoints} data {#plural(numPoints, points, point)}.",
line:"{name}, line {ix} of {numSeries} with {numPoints} data {#plural(numPoints, points, point)}.",lineCombination:"{name}, series {ix} of {numSeries}. Line with {numPoints} data {#plural(numPoints, points, point)}.",spline:"{name}, line {ix} of {numSeries} with {numPoints} data {#plural(numPoints, points, point)}.",splineCombination:"{name}, series {ix} of {numSeries}. Line with {numPoints} data {#plural(numPoints, points, point)}.",column:"{name}, bar series {ix} of {numSeries} with {numPoints} {#plural(numPoints, bars, bar)}.",
columnCombination:"{name}, series {ix} of {numSeries}. Bar series with {numPoints} {#plural(numPoints, bars, bar)}.",bar:"{name}, bar series {ix} of {numSeries} with {numPoints} {#plural(numPoints, bars, bar)}.",barCombination:"{name}, series {ix} of {numSeries}. Bar series with {numPoints} {#plural(numPoints, bars, bar)}.",pie:"{name}, pie {ix} of {numSeries} with {numPoints} {#plural(numPoints, slices, slice)}.",pieCombination:"{name}, series {ix} of {numSeries}. Pie with {numPoints} {#plural(numPoints, slices, slice)}.",
scatter:"{name}, scatter plot {ix} of {numSeries} with {numPoints} {#plural(numPoints, points, point)}.",scatterCombination:"{name}, series {ix} of {numSeries}, scatter plot with {numPoints} {#plural(numPoints, points, point)}.",boxplot:"{name}, boxplot {ix} of {numSeries} with {numPoints} {#plural(numPoints, boxes, box)}.",boxplotCombination:"{name}, series {ix} of {numSeries}. Boxplot with {numPoints} {#plural(numPoints, boxes, box)}.",bubble:"{name}, bubble series {ix} of {numSeries} with {numPoints} {#plural(numPoints, bubbles, bubble)}.",
bubbleCombination:"{name}, series {ix} of {numSeries}. Bubble series with {numPoints} {#plural(numPoints, bubbles, bubble)}.",map:"{name}, map {ix} of {numSeries} with {numPoints} {#plural(numPoints, areas, area)}.",mapCombination:"{name}, series {ix} of {numSeries}. Map with {numPoints} {#plural(numPoints, areas, area)}.",mapline:"{name}, line {ix} of {numSeries} with {numPoints} data {#plural(numPoints, points, point)}.",maplineCombination:"{name}, series {ix} of {numSeries}. Line with {numPoints} data {#plural(numPoints, points, point)}.",
mapbubble:"{name}, bubble series {ix} of {numSeries} with {numPoints} {#plural(numPoints, bubbles, bubble)}.",mapbubbleCombination:"{name}, series {ix} of {numSeries}. Bubble series with {numPoints} {#plural(numPoints, bubbles, bubble)}."},description:"{description}",xAxisDescription:"X axis, {name}",yAxisDescription:"Y axis, {name}",nullPointValue:"No value",pointAnnotationsDescription:"{Annotation: #each(annotations). }"}}}});v(a,"Accessibility/Options/DeprecatedOptions.js",[a["Core/Utilities.js"]],
function(a){function h(a,h,g){for(var c,e=0;e<h.length-1;++e)c=h[e],a=a[c]=t(a[c],{});a[h[h.length-1]]=g}function m(a,l,g,c){function e(a,b){return b.reduce(function(a,b){return a[b]},a)}var d=e(a.options,l),b=e(a.options,g);Object.keys(c).forEach(function(e){var f,k=d[e];"undefined"!==typeof k&&(h(b,c[e],k),w(32,!1,a,(f={},f[l.join(".")+"."+e]=g.join(".")+"."+c[e].join("."),f)))})}function u(a){var h=a.options.chart,g=a.options.accessibility||{};["description","typeDescription"].forEach(function(c){var e;
h[c]&&(g[c]=h[c],w(32,!1,a,(e={},e["chart."+c]="use accessibility."+c,e)))})}function n(a){a.axes.forEach(function(h){(h=h.options)&&h.description&&(h.accessibility=h.accessibility||{},h.accessibility.description=h.description,w(32,!1,a,{"axis.description":"use axis.accessibility.description"}))})}function p(a){var k={description:["accessibility","description"],exposeElementToA11y:["accessibility","exposeAsGroupOnly"],pointDescriptionFormatter:["accessibility","point","descriptionFormatter"],skipKeyboardNavigation:["accessibility",
"keyboardNavigation","enabled"],"accessibility.pointDescriptionFormatter":["accessibility","point","descriptionFormatter"]};a.series.forEach(function(g){Object.keys(k).forEach(function(c){var e,d=g.options[c];"accessibility.pointDescriptionFormatter"===c&&(d=g.options.accessibility&&g.options.accessibility.pointDescriptionFormatter);"undefined"!==typeof d&&(h(g.options,k[c],"skipKeyboardNavigation"===c?!d:d),w(32,!1,a,(e={},e["series."+c]="series."+k[c].join("."),e)))})})}var w=a.error,t=a.pick;return function(a){u(a);
n(a);a.series&&p(a);m(a,["accessibility"],["accessibility"],{pointDateFormat:["point","dateFormat"],pointDateFormatter:["point","dateFormatter"],pointDescriptionFormatter:["point","descriptionFormatter"],pointDescriptionThreshold:["series","pointDescriptionEnabledThreshold"],pointNavigationThreshold:["keyboardNavigation","seriesNavigation","pointNavigationEnabledThreshold"],pointValueDecimals:["point","valueDecimals"],pointValuePrefix:["point","valuePrefix"],pointValueSuffix:["point","valueSuffix"],
screenReaderSectionFormatter:["screenReaderSection","beforeChartFormatter"],describeSingleSeries:["series","describeSingleSeries"],seriesDescriptionFormatter:["series","descriptionFormatter"],onTableAnchorClick:["screenReaderSection","onViewDataTableClick"],axisRangeDateFormat:["screenReaderSection","axisRangeDateFormat"]});m(a,["accessibility","keyboardNavigation"],["accessibility","keyboardNavigation","seriesNavigation"],{skipNullPoints:["skipNullPoints"],mode:["mode"]});m(a,["lang","accessibility"],
["lang","accessibility"],{legendItem:["legend","legendItem"],legendLabel:["legend","legendLabel"],mapZoomIn:["zoom","mapZoomIn"],mapZoomOut:["zoom","mapZoomOut"],resetZoomButton:["zoom","resetZoomButton"],screenReaderRegionLabel:["screenReaderSection","beforeRegionLabel"],rangeSelectorButton:["rangeSelector","buttonText"],rangeSelectorMaxInput:["rangeSelector","maxInputLabel"],rangeSelectorMinInput:["rangeSelector","minInputLabel"],svgContainerEnd:["screenReaderSection","endOfChartMarker"],viewAsDataTable:["table",
"viewAsDataTableButtonText"],tableSummary:["table","tableSummary"]})}});v(a,"Accessibility/Accessibility.js",[a["Core/DefaultOptions.js"],a["Core/Globals.js"],a["Core/Utilities.js"],a["Accessibility/A11yI18n.js"],a["Accessibility/Components/ContainerComponent.js"],a["Accessibility/FocusBorder.js"],a["Accessibility/Components/InfoRegionsComponent.js"],a["Accessibility/KeyboardNavigation.js"],a["Accessibility/Components/LegendComponent.js"],a["Accessibility/Components/MenuComponent.js"],a["Accessibility/Components/SeriesComponent/NewDataAnnouncer.js"],
a["Accessibility/ProxyProvider.js"],a["Accessibility/Components/RangeSelectorComponent.js"],a["Accessibility/Components/SeriesComponent/SeriesComponent.js"],a["Accessibility/Components/ZoomComponent.js"],a["Accessibility/HighContrastMode.js"],a["Accessibility/HighContrastTheme.js"],a["Accessibility/Options/Options.js"],a["Accessibility/Options/LangOptions.js"],a["Accessibility/Options/DeprecatedOptions.js"]],function(a,h,m,u,n,p,v,t,k,l,g,c,e,d,b,f,A,z,J,C){a=a.defaultOptions;var w=h.doc,x=m.addEvent,
D=m.extend,E=m.fireEvent,B=m.merge;h=function(){function a(a){this.proxyProvider=this.keyboardNavigation=this.components=this.chart=void 0;this.init(a)}a.prototype.init=function(a){this.chart=a;w.addEventListener&&a.renderer.isSVG?(C(a),this.proxyProvider=new c(this.chart),this.initComponents(),this.keyboardNavigation=new t(a,this.components),this.update()):(this.zombie=!0,this.components={},a.renderTo.setAttribute("aria-hidden",!0))};a.prototype.initComponents=function(){var a=this.chart,c=this.proxyProvider,
f=a.options.accessibility;this.components={container:new n,infoRegions:new v,legend:new k,chartMenu:new l,rangeSelector:new e,series:new d,zoom:new b};f.customComponents&&D(this.components,f.customComponents);var g=this.components;this.getComponentOrder().forEach(function(b){g[b].initBase(a,c);g[b].init()})};a.prototype.getComponentOrder=function(){if(!this.components)return[];if(!this.components.series)return Object.keys(this.components);var a=Object.keys(this.components).filter(function(a){return"series"!==
a});return["series"].concat(a)};a.prototype.update=function(){var a=this.components,b=this.chart,c=b.options.accessibility;E(b,"beforeA11yUpdate");b.types=this.getChartTypes();c=c.keyboardNavigation.order;this.proxyProvider.updateGroupOrder(c);this.getComponentOrder().forEach(function(c){a[c].onChartUpdate();E(b,"afterA11yComponentUpdate",{name:c,component:a[c]})});this.keyboardNavigation.update(c);!b.highContrastModeActive&&f.isHighContrastModeActive()&&f.setHighContrastTheme(b);E(b,"afterA11yUpdate",
{accessibility:this})};a.prototype.destroy=function(){var a=this.chart||{},b=this.components;Object.keys(b).forEach(function(a){b[a].destroy();b[a].destroyBase()});this.proxyProvider&&this.proxyProvider.destroy();this.keyboardNavigation&&this.keyboardNavigation.destroy();a.renderTo&&a.renderTo.setAttribute("aria-hidden",!0);a.focusElement&&a.focusElement.removeFocusBorder()};a.prototype.getChartTypes=function(){var a={};this.chart.series.forEach(function(b){a[b.type]=1});return Object.keys(a)};return a}();
(function(a){function b(){this.accessibility&&this.accessibility.destroy()}function c(){this.a11yDirty&&this.renderTo&&(delete this.a11yDirty,this.updateA11yEnabled());var a=this.accessibility;a&&!a.zombie&&(a.proxyProvider.updateProxyElementPositions(),a.getComponentOrder().forEach(function(b){a.components[b].onChartRender()}))}function f(a){if(a=a.options.accessibility)a.customComponents&&(this.options.accessibility.customComponents=a.customComponents,delete a.customComponents),B(!0,this.options.accessibility,
a),this.accessibility&&this.accessibility.destroy&&(this.accessibility.destroy(),delete this.accessibility);this.a11yDirty=!0}function h(){var b=this.accessibility,c=this.options.accessibility;c&&c.enabled?b&&!b.zombie?b.update():this.accessibility=new a(this):b?(b.destroy&&b.destroy(),delete this.accessibility):this.renderTo.setAttribute("aria-hidden",!0)}function k(){this.series.chart.accessibility&&(this.series.chart.a11yDirty=!0)}var m=[];a.i18nFormat=u.i18nFormat;a.compose=function(a,n,q,r,w){u.compose(a);
p.compose(a,r);t.compose(a);l.compose(a);g.compose(q);d.compose(a,n,q);w&&e.compose(a,w);-1===m.indexOf(a)&&(m.push(a),a.prototype.updateA11yEnabled=h,x(a,"destroy",b),x(a,"render",c),x(a,"update",f),["addSeries","init"].forEach(function(b){x(a,b,function(){this.a11yDirty=!0})}),["afterDrilldown","drillupall"].forEach(function(b){x(a,b,function(){var a=this.accessibility;a&&!a.zombie&&a.update()})}));-1===m.indexOf(n)&&(m.push(n),x(n,"update",k));-1===m.indexOf(q)&&(m.push(q),["update","updatedData",
"remove"].forEach(function(a){x(q,a,function(){this.chart.accessibility&&(this.chart.a11yDirty=!0)})}))}})(h||(h={}));B(!0,a,z,{accessibility:{highContrastTheme:A},lang:J});return h});v(a,"masters/modules/accessibility.src.js",[a["Core/Globals.js"],a["Accessibility/Accessibility.js"],a["Accessibility/AccessibilityComponent.js"],a["Accessibility/Utils/ChartUtilities.js"],a["Accessibility/Utils/HTMLUtilities.js"],a["Accessibility/KeyboardNavigationHandler.js"],a["Accessibility/Components/SeriesComponent/SeriesDescriber.js"]],
function(a,h,m,u,n,p,v){a.i18nFormat=h.i18nFormat;a.A11yChartUtilities=u;a.A11yHTMLUtilities=n;a.AccessibilityComponent=m;a.KeyboardNavigationHandler=p;a.SeriesAccessibilityDescriber=v;h.compose(a.Chart,a.Point,a.Series,a.SVGElement,a.RangeSelector)})});
//# sourceMappingURL=accessibility.js.map
```
