## Supplementary material for "Exploring natural odour landscapes: A case study with implications for human-biting insects": Figure S4: annotations-advanced.js

```
/*
 Highcharts JS v9.3.1 (2021-11-05)

 Annotations module

 (c) 2009-2021 Torstein Honsi

 License: www.highcharts.com/license
*/
'use strict';(function(c){"object"===typeof module&&module.exports?(c["default"]=c,module.exports=c):"function"===typeof define&&define.amd?define("highcharts/modules/annotations-advanced",["highcharts"],function(v){c(v);c.Highcharts=v;return c}):c("undefined"!==typeof Highcharts?Highcharts:void 0)})(function(c){function v(g,c,n,k){g.hasOwnProperty(c)||(g[c]=k.apply(null,n))}c=c?c._modules:{};v(c,"Extensions/Annotations/Mixins/EventEmitterMixin.js",[c["Core/Globals.js"],c["Core/Utilities.js"]],function(g,
c){var l=c.addEvent,k=c.fireEvent,r=c.objectEach,m=c.pick,e=c.removeEvent;return{addEvents:function(){var b=this,a=function(a){l(a,g.isTouchDevice?"touchstart":"mousedown",function(a){b.onMouseDown(a)},{passive:!1})};a(this.graphic.element);(b.labels||[]).forEach(function(d){d.options.useHTML&&d.graphic.text&&a(d.graphic.text.element)});r(b.options.events,function(a,d){var p=function(p){"click"===d&&b.cancelClick||a.call(b,b.chart.pointer.normalize(p),b.target)};if(-1===(b.nonDOMEvents||[]).indexOf(d))b.graphic.on(d,
p);else l(b,d,p,{passive:!1})});if(b.options.draggable&&(l(b,"drag",b.onDrag),!b.graphic.renderer.styledMode)){var d={cursor:{x:"ew-resize",y:"ns-resize",xy:"move"}[b.options.draggable]};b.graphic.css(d);(b.labels||[]).forEach(function(a){a.options.useHTML&&a.graphic.text&&a.graphic.text.css(d)})}b.isUpdating||k(b,"add")},removeDocEvents:function(){this.removeDrag&&(this.removeDrag=this.removeDrag());this.removeMouseUp&&(this.removeMouseUp=this.removeMouseUp())},onMouseDown:function(b){var a=this,
d=a.chart.pointer;b.preventDefault&&b.preventDefault();if(2!==b.button){b=d.normalize(b);var h=b.chartX;var f=b.chartY;a.cancelClick=!1;a.chart.hasDraggedAnnotation=!0;a.removeDrag=l(g.doc,g.isTouchDevice?"touchmove":"mousemove",function(b){a.hasDragged=!0;b=d.normalize(b);b.prevChartX=h;b.prevChartY=f;k(a,"drag",b);h=b.chartX;f=b.chartY},g.isTouchDevice?{passive:!1}:void 0);a.removeMouseUp=l(g.doc,g.isTouchDevice?"touchend":"mouseup",function(d){var b=m(a.target&&a.target.annotation,a.target);b&&
(b.cancelClick=a.hasDragged);a.cancelClick=a.hasDragged;a.hasDragged=!1;a.chart.hasDraggedAnnotation=!1;k(m(b,a),"afterUpdate");a.onMouseUp(d)},g.isTouchDevice?{passive:!1}:void 0)}},onMouseUp:function(b){var a=this.chart;b=this.target||this;var d=a.options.annotations;a=a.annotations.indexOf(b);this.removeDocEvents();d[a]=b.options},onDrag:function(b){if(this.chart.isInsidePlot(b.chartX-this.chart.plotLeft,b.chartY-this.chart.plotTop,{visiblePlotOnly:!0})){var a=this.mouseMoveToTranslation(b);"x"===
this.options.draggable&&(a.y=0);"y"===this.options.draggable&&(a.x=0);this.points.length?this.translate(a.x,a.y):(this.shapes.forEach(function(d){d.translate(a.x,a.y)}),this.labels.forEach(function(d){d.translate(a.x,a.y)}));this.redraw(!1)}},mouseMoveToRadians:function(b,a,d){var h=b.prevChartY-d,f=b.prevChartX-a;d=b.chartY-d;b=b.chartX-a;this.chart.inverted&&(a=f,f=h,h=a,a=b,b=d,d=a);return Math.atan2(d,b)-Math.atan2(h,f)},mouseMoveToTranslation:function(b){var a=b.chartX-b.prevChartX;b=b.chartY-
b.prevChartY;if(this.chart.inverted){var d=b;b=a;a=d}return{x:a,y:b}},mouseMoveToScale:function(b,a,d){a=(b.chartX-a||1)/(b.prevChartX-a||1);b=(b.chartY-d||1)/(b.prevChartY-d||1);this.chart.inverted&&(d=b,b=a,a=d);return{x:a,y:b}},destroy:function(){this.removeDocEvents();e(this);this.hcEvents=null}}});v(c,"Extensions/Annotations/ControlPoint.js",[c["Core/Utilities.js"],c["Extensions/Annotations/Mixins/EventEmitterMixin.js"]],function(g,c){var l=g.merge,k=g.pick;return function(){function g(g,e,b,
a){this.addEvents=c.addEvents;this.graphic=void 0;this.mouseMoveToRadians=c.mouseMoveToRadians;this.mouseMoveToScale=c.mouseMoveToScale;this.mouseMoveToTranslation=c.mouseMoveToTranslation;this.onDrag=c.onDrag;this.onMouseDown=c.onMouseDown;this.onMouseUp=c.onMouseUp;this.removeDocEvents=c.removeDocEvents;this.nonDOMEvents=["drag"];this.chart=g;this.target=e;this.options=b;this.index=k(b.index,a)}g.prototype.setVisibility=function(g){this.graphic.attr("visibility",g?"visible":"hidden");this.options.visible=
g};g.prototype.render=function(){var g=this.chart,e=this.options;this.graphic=g.renderer.symbol(e.symbol,0,0,e.width,e.height).add(g.controlPointsGroup).css(e.style);this.setVisibility(e.visible);this.addEvents()};g.prototype.redraw=function(g){this.graphic[g?"animate":"attr"](this.options.positioner.call(this,this.target))};g.prototype.destroy=function(){c.destroy.call(this);this.graphic&&(this.graphic=this.graphic.destroy());this.options=this.target=this.chart=null};g.prototype.update=function(g){var e=
this.chart,b=this.target,a=this.index;g=l(!0,this.options,g);this.destroy();this.constructor(e,b,g,a);this.render(e.controlPointsGroup);this.redraw()};return g}()});v(c,"Extensions/Annotations/MockPoint.js",[c["Core/Series/Series.js"],c["Core/Utilities.js"],c["Core/Axis/Axis.js"]],function(g,c,n){var k=c.defined,l=c.fireEvent;return function(){function c(e,b,a){this.y=this.x=this.ttBelow=this.plotY=this.plotX=this.negative=this.isInside=void 0;this.mock=!0;this.series={visible:!0,chart:e,getPlotBox:g.prototype.getPlotBox};
this.target=b||null;this.options=a;this.applyOptions(this.getOptions())}c.fromPoint=function(e){return new c(e.series.chart,null,{x:e.x,y:e.y,xAxis:e.series.xAxis,yAxis:e.series.yAxis})};c.pointToPixels=function(e,b){var a=e.series,d=a.chart,h=e.plotX,f=e.plotY;d.inverted&&(e.mock?(h=e.plotY,f=e.plotX):(h=d.plotWidth-e.plotY,f=d.plotHeight-e.plotX));a&&!b&&(e=a.getPlotBox(),h+=e.translateX,f+=e.translateY);return{x:h,y:f}};c.pointToOptions=function(e){return{x:e.x,y:e.y,xAxis:e.series.xAxis,yAxis:e.series.yAxis}};
c.prototype.hasDynamicOptions=function(){return"function"===typeof this.options};c.prototype.getOptions=function(){return this.hasDynamicOptions()?this.options(this.target):this.options};c.prototype.applyOptions=function(e){this.command=e.command;this.setAxis(e,"x");this.setAxis(e,"y");this.refresh()};c.prototype.setAxis=function(e,b){b+="Axis";e=e[b];var a=this.series.chart;this.series[b]=e instanceof n?e:k(e)?a[b][e]||a.get(e):null};c.prototype.toAnchor=function(){var e=[this.plotX,this.plotY,0,
0];this.series.chart.inverted&&(e[0]=this.plotY,e[1]=this.plotX);return e};c.prototype.getLabelConfig=function(){return{x:this.x,y:this.y,point:this}};c.prototype.isInsidePlot=function(){var e=this.plotX,b=this.plotY,a=this.series.xAxis,d=this.series.yAxis,h={x:e,y:b,isInsidePlot:!0};a&&(h.isInsidePlot=k(e)&&0<=e&&e<=a.len);d&&(h.isInsidePlot=h.isInsidePlot&&k(b)&&0<=b&&b<=d.len);l(this.series.chart,"afterIsInsidePlot",h);return h.isInsidePlot};c.prototype.refresh=function(){var e=this.series,b=e.xAxis;
e=e.yAxis;var a=this.getOptions();b?(this.x=a.x,this.plotX=b.toPixels(a.x,!0)):(this.x=null,this.plotX=a.x);e?(this.y=a.y,this.plotY=e.toPixels(a.y,!0)):(this.y=null,this.plotY=a.y);this.isInside=this.isInsidePlot()};c.prototype.translate=function(e,b,a,d){this.hasDynamicOptions()||(this.plotX+=a,this.plotY+=d,this.refreshOptions())};c.prototype.scale=function(e,b,a,d){if(!this.hasDynamicOptions()){var h=this.plotY*d;this.plotX=(1-a)*e+this.plotX*a;this.plotY=(1-d)*b+h;this.refreshOptions()}};c.prototype.rotate=
function(e,b,a){if(!this.hasDynamicOptions()){var d=Math.cos(a);a=Math.sin(a);var h=this.plotX,f=this.plotY;h-=e;f-=b;this.plotX=h*d-f*a+e;this.plotY=h*a+f*d+b;this.refreshOptions()}};c.prototype.refreshOptions=function(){var e=this.series,b=e.xAxis;e=e.yAxis;this.x=this.options.x=b?this.options.x=b.toValue(this.plotX,!0):this.plotX;this.y=this.options.y=e?e.toValue(this.plotY,!0):this.plotY};return c}()});v(c,"Extensions/Annotations/Mixins/ControllableMixin.js",[c["Extensions/Annotations/ControlPoint.js"],
c["Extensions/Annotations/MockPoint.js"],c["Core/Tooltip.js"],c["Core/Utilities.js"]],function(g,c,n,k){var l=k.isObject,m=k.isString,e=k.merge,b=k.splat;return{init:function(a,d,b){this.annotation=a;this.chart=a.chart;this.options=d;this.points=[];this.controlPoints=[];this.index=b;this.linkPoints();this.addControlPoints()},attr:function(){this.graphic.attr.apply(this.graphic,arguments)},getPointsOptions:function(){var a=this.options;return a.points||a.point&&b(a.point)},attrsFromOptions:function(a){var d=
this.constructor.attrsMap,b={},f,p=this.chart.styledMode;for(f in a){var e=d[f];!e||p&&-1!==["fill","stroke","stroke-width"].indexOf(e)||(b[e]=a[f])}return b},anchor:function(a){var d=a.series.getPlotBox(),b=a.series.chart,f=a.mock?a.toAnchor():n.prototype.getAnchor.call({chart:a.series.chart},a);f={x:f[0]+(this.options.x||0),y:f[1]+(this.options.y||0),height:f[2]||0,width:f[3]||0};return{relativePosition:f,absolutePosition:e(f,{x:f.x+(a.mock?d.translateX:b.plotLeft),y:f.y+(a.mock?d.translateY:b.plotTop)})}},
point:function(a,d){if(a&&a.series)return a;d&&null!==d.series||(l(a)?d=new c(this.chart,this,a):m(a)?d=this.chart.get(a)||null:"function"===typeof a&&(d=a.call(d,this),d=d.series?d:new c(this.chart,this,a)));return d},linkPoints:function(){var a=this.getPointsOptions(),d=this.points,b=a&&a.length||0,f;for(f=0;f<b;f++){var p=this.point(a[f],d[f]);if(!p){d.length=0;return}p.mock&&p.refresh();d[f]=p}return d},addControlPoints:function(){var a=this.options.controlPoints;(a||[]).forEach(function(d,b){d=
e(this.options.controlPointOptions,d);d.index||(d.index=b);a[b]=d;this.controlPoints.push(new g(this.chart,this,d))},this)},shouldBeDrawn:function(){return!!this.points.length},render:function(a){this.controlPoints.forEach(function(a){a.render()})},redraw:function(a){this.controlPoints.forEach(function(d){d.redraw(a)})},transform:function(a,d,b,f,p){if(this.chart.inverted){var h=d;d=b;b=h}this.points.forEach(function(h,e){this.transformPoint(a,d,b,f,p,e)},this)},transformPoint:function(a,d,b,f,p,
e){var h=this.points[e];h.mock||(h=this.points[e]=c.fromPoint(h));h[a](d,b,f,p)},translate:function(a,d){this.transform("translate",null,null,a,d)},translatePoint:function(a,d,b){this.transformPoint("translate",null,null,a,d,b)},translateShape:function(a,d,b){var f=this.annotation.chart,p=this.annotation.userOptions,h=f.annotations.indexOf(this.annotation);f=f.options.annotations[h];this.translatePoint(a,d,0);b&&this.translatePoint(a,d,1);f[this.collection][this.index].point=this.options.point;p[this.collection][this.index].point=
this.options.point},rotate:function(a,d,b){this.transform("rotate",a,d,b)},scale:function(a,d,b,f){this.transform("scale",a,d,b,f)},setControlPointsVisibility:function(a){this.controlPoints.forEach(function(d){d.setVisibility(a)})},destroy:function(){this.graphic&&(this.graphic=this.graphic.destroy());this.tracker&&(this.tracker=this.tracker.destroy());this.controlPoints.forEach(function(a){a.destroy()});this.options=this.controlPoints=this.points=this.chart=null;this.annotation&&(this.annotation=
null)},update:function(a){var d=this.annotation;a=e(!0,this.options,a);var b=this.graphic.parentGroup;this.destroy();this.constructor(d,a,this.index);this.render(b);this.redraw()}}});v(c,"Extensions/Annotations/Mixins/MarkerMixin.js",[c["Core/Chart/Chart.js"],c["Core/Renderer/SVG/SVGRenderer.js"],c["Core/Utilities.js"]],function(g,c,n){function k(a){return function(d){this.attr(a,"url(#"+d+")")}}var l=n.addEvent,m=n.defined,e=n.merge,b=n.uniqueKey,a={arrow:{tagName:"marker",attributes:{id:"arrow",
refY:5,refX:9,markerWidth:10,markerHeight:10},children:[{tagName:"path",attributes:{d:"M 0 0 L 10 5 L 0 10 Z","stroke-width":0}}]},"reverse-arrow":{tagName:"marker",attributes:{id:"reverse-arrow",refY:5,refX:1,markerWidth:10,markerHeight:10},children:[{tagName:"path",attributes:{d:"M 0 5 L 10 0 L 10 10 Z","stroke-width":0}}]}};c.prototype.addMarker=function(a,b){var d={attributes:{id:a}},p={stroke:b.color||"none",fill:b.color||"rgba(0, 0, 0, 0.75)"};d.children=b.children&&b.children.map(function(a){return e(p,
a)});b=e(!0,{attributes:{markerWidth:20,markerHeight:20,refX:0,refY:0,orient:"auto"}},b,d);b=this.definition(b);b.id=a;return b};c={markerEndSetter:k("marker-end"),markerStartSetter:k("marker-start"),setItemMarkers:function(a){var d=a.options,f=a.chart,p=f.options.defs,t=d.fill,g=m(t)&&"none"!==t?t:d.stroke;["markerStart","markerEnd"].forEach(function(h){var t=d[h],c;if(t){for(c in p){var x=p[c];if((t===(x.attributes&&x.attributes.id)||t===x.id)&&"marker"===x.tagName){var A=x;break}}A&&(t=a[h]=f.renderer.addMarker((d.id||
b())+"-"+t,e(A,{color:g})),a.attr(h,t.getAttribute("id")))}})}};l(g,"afterGetContainer",function(){this.options.defs=e(a,this.options.defs||{})});return c});v(c,"Extensions/Annotations/Controllables/ControllablePath.js",[c["Extensions/Annotations/Mixins/ControllableMixin.js"],c["Core/Globals.js"],c["Extensions/Annotations/Mixins/MarkerMixin.js"],c["Core/Utilities.js"]],function(g,c,n,k){var l=k.extend,m="rgba(192,192,192,"+(c.svg?.0001:.002)+")";return function(){function e(b,a,d){this.addControlPoints=
g.addControlPoints;this.anchor=g.anchor;this.attr=g.attr;this.attrsFromOptions=g.attrsFromOptions;this.destroy=g.destroy;this.getPointsOptions=g.getPointsOptions;this.init=g.init;this.linkPoints=g.linkPoints;this.point=g.point;this.rotate=g.rotate;this.scale=g.scale;this.setControlPointsVisibility=g.setControlPointsVisibility;this.setMarkers=n.setItemMarkers;this.transform=g.transform;this.transformPoint=g.transformPoint;this.translate=g.translate;this.translatePoint=g.translatePoint;this.translateShape=
g.translateShape;this.update=g.update;this.type="path";this.init(b,a,d);this.collection="shapes"}e.prototype.toD=function(){var b=this.options.d;if(b)return"function"===typeof b?b.call(this):b;b=this.points;var a=b.length,d=a,h=b[0],f=d&&this.anchor(h).absolutePosition,p=0,e=[];if(f)for(e.push(["M",f.x,f.y]);++p<a&&d;)h=b[p],d=h.command||"L",f=this.anchor(h).absolutePosition,"M"===d?e.push([d,f.x,f.y]):"L"===d?e.push([d,f.x,f.y]):"Z"===d&&e.push([d]),d=h.series.visible;return d?this.chart.renderer.crispLine(e,
this.graphic.strokeWidth()):null};e.prototype.shouldBeDrawn=function(){return g.shouldBeDrawn.call(this)||!!this.options.d};e.prototype.render=function(b){var a=this.options,d=this.attrsFromOptions(a);this.graphic=this.annotation.chart.renderer.path([["M",0,0]]).attr(d).add(b);a.className&&this.graphic.addClass(a.className);this.tracker=this.annotation.chart.renderer.path([["M",0,0]]).addClass("highcharts-tracker-line").attr({zIndex:2}).add(b);this.annotation.chart.styledMode||this.tracker.attr({"stroke-linejoin":"round",
stroke:m,fill:m,"stroke-width":this.graphic.strokeWidth()+2*a.snap});g.render.call(this);l(this.graphic,{markerStartSetter:n.markerStartSetter,markerEndSetter:n.markerEndSetter});this.setMarkers(this)};e.prototype.redraw=function(b){var a=this.toD(),d=b?"animate":"attr";a?(this.graphic[d]({d:a}),this.tracker[d]({d:a})):(this.graphic.attr({d:"M 0 -9000000000"}),this.tracker.attr({d:"M 0 -9000000000"}));this.graphic.placed=this.tracker.placed=!!a;g.redraw.call(this,b)};e.attrsMap={dashStyle:"dashstyle",
strokeWidth:"stroke-width",stroke:"stroke",fill:"fill",zIndex:"zIndex"};return e}()});v(c,"Extensions/Annotations/Controllables/ControllableRect.js",[c["Extensions/Annotations/Mixins/ControllableMixin.js"],c["Extensions/Annotations/Controllables/ControllablePath.js"],c["Core/Utilities.js"]],function(g,c,n){var k=n.merge;return function(){function l(c,e,b){this.addControlPoints=g.addControlPoints;this.anchor=g.anchor;this.attr=g.attr;this.attrsFromOptions=g.attrsFromOptions;this.destroy=g.destroy;
this.getPointsOptions=g.getPointsOptions;this.init=g.init;this.linkPoints=g.linkPoints;this.point=g.point;this.rotate=g.rotate;this.scale=g.scale;this.setControlPointsVisibility=g.setControlPointsVisibility;this.shouldBeDrawn=g.shouldBeDrawn;this.transform=g.transform;this.transformPoint=g.transformPoint;this.translatePoint=g.translatePoint;this.translateShape=g.translateShape;this.update=g.update;this.type="rect";this.translate=g.translateShape;this.init(c,e,b);this.collection="shapes"}l.prototype.render=
function(c){var e=this.attrsFromOptions(this.options);this.graphic=this.annotation.chart.renderer.rect(0,-9E9,0,0).attr(e).add(c);g.render.call(this)};l.prototype.redraw=function(c){var e=this.anchor(this.points[0]).absolutePosition;if(e)this.graphic[c?"animate":"attr"]({x:e.x,y:e.y,width:this.options.width,height:this.options.height});else this.attr({x:0,y:-9E9});this.graphic.placed=!!e;g.redraw.call(this,c)};l.attrsMap=k(c.attrsMap,{width:"width",height:"height"});return l}()});v(c,"Extensions/Annotations/Controllables/ControllableCircle.js",
[c["Extensions/Annotations/Mixins/ControllableMixin.js"],c["Extensions/Annotations/Controllables/ControllablePath.js"],c["Core/Utilities.js"]],function(g,c,n){var k=n.merge;return function(){function l(c,e,b){this.addControlPoints=g.addControlPoints;this.anchor=g.anchor;this.attr=g.attr;this.attrsFromOptions=g.attrsFromOptions;this.destroy=g.destroy;this.getPointsOptions=g.getPointsOptions;this.init=g.init;this.linkPoints=g.linkPoints;this.point=g.point;this.rotate=g.rotate;this.scale=g.scale;this.setControlPointsVisibility=
g.setControlPointsVisibility;this.shouldBeDrawn=g.shouldBeDrawn;this.transform=g.transform;this.transformPoint=g.transformPoint;this.translatePoint=g.translatePoint;this.translateShape=g.translateShape;this.update=g.update;this.type="circle";this.translate=g.translateShape;this.init(c,e,b);this.collection="shapes"}l.prototype.render=function(c){var e=this.attrsFromOptions(this.options);this.graphic=this.annotation.chart.renderer.circle(0,-9E9,0).attr(e).add(c);g.render.call(this)};l.prototype.redraw=
function(c){var e=this.anchor(this.points[0]).absolutePosition;if(e)this.graphic[c?"animate":"attr"]({x:e.x,y:e.y,r:this.options.r});else this.graphic.attr({x:0,y:-9E9});this.graphic.placed=!!e;g.redraw.call(this,c)};l.prototype.setRadius=function(c){this.options.r=c};l.attrsMap=k(c.attrsMap,{r:"r"});return l}()});v(c,"Extensions/Annotations/Controllables/ControllableEllipse.js",[c["Extensions/Annotations/Mixins/ControllableMixin.js"],c["Extensions/Annotations/Controllables/ControllablePath.js"],
c["Core/Utilities.js"]],function(c,l,n){var g=n.merge,r=n.defined;return function(){function m(e,b,a){this.addControlPoints=c.addControlPoints;this.anchor=c.anchor;this.attr=c.attr;this.attrsFromOptions=c.attrsFromOptions;this.destroy=c.destroy;this.getPointsOptions=c.getPointsOptions;this.linkPoints=c.linkPoints;this.point=c.point;this.scale=c.scale;this.setControlPointsVisibility=c.setControlPointsVisibility;this.shouldBeDrawn=c.shouldBeDrawn;this.transform=c.transform;this.translatePoint=c.translatePoint;
this.transformPoint=c.transformPoint;this.type="ellipse";this.init(e,b,a);this.collection="shapes"}m.prototype.init=function(e,b,a){r(b.yAxis)&&b.points.forEach(function(a){a.yAxis=b.yAxis});r(b.xAxis)&&b.points.forEach(function(a){a.xAxis=b.xAxis});c.init.call(this,e,b,a)};m.prototype.render=function(e){this.graphic=this.annotation.chart.renderer.createElement("ellipse").attr(this.attrsFromOptions(this.options)).add(e);c.render.call(this)};m.prototype.translate=function(e,b){c.translateShape.call(this,
e,b,!0)};m.prototype.getDistanceFromLine=function(e,b,a,d){return Math.abs((b.y-e.y)*a-(b.x-e.x)*d+b.x*e.y-b.y*e.x)/Math.sqrt((b.y-e.y)*(b.y-e.y)+(b.x-e.x)*(b.x-e.x))};m.prototype.getAttrs=function(e,b){var a=e.x,d=e.y,h=b.x,f=b.y;b=(a+h)/2;e=(d+f)/2;var p=Math.sqrt((a-h)*(a-h)/4+(d-f)*(d-f)/4);d=180*Math.atan((f-d)/(h-a))/Math.PI;b<a&&(d+=180);a=this.getRY();return{cx:b,cy:e,rx:p,ry:a,angle:d}};m.prototype.getRY=function(){var e=this.getYAxis();return r(e)?Math.abs(e.toPixels(this.options.ry)-e.toPixels(0)):
this.options.ry};m.prototype.getYAxis=function(){return this.chart.yAxis[this.options.yAxis]};m.prototype.getAbsolutePosition=function(e){return this.anchor(e).absolutePosition};m.prototype.redraw=function(e){var b=this.getAbsolutePosition(this.points[0]),a=this.getAbsolutePosition(this.points[1]);a=this.getAttrs(b,a);if(b)this.graphic[e?"animate":"attr"]({cx:a.cx,cy:a.cy,rx:a.rx,ry:a.ry,rotation:a.angle,rotationOriginX:a.cx,rotationOriginY:a.cy});else this.graphic.attr({x:0,y:-9E9});this.graphic.placed=
!!b;c.redraw.call(this,e)};m.prototype.setYRadius=function(e){this.options.ry=e;this.annotation.userOptions.shapes[0].ry=e;this.annotation.options.shapes[0].ry=e};m.attrsMap=g(l.attrsMap,{ry:"ry"});return m}()});v(c,"Extensions/Annotations/Controllables/ControllableLabel.js",[c["Extensions/Annotations/Mixins/ControllableMixin.js"],c["Core/FormatUtilities.js"],c["Extensions/Annotations/MockPoint.js"],c["Core/Renderer/SVG/SVGRenderer.js"],c["Core/Tooltip.js"],c["Core/Utilities.js"]],function(c,l,n,
k,r,m){var e=l.format;l=k.prototype.symbols;var b=m.extend,a=m.isNumber,d=m.pick;m=function(){function a(a,b,d){this.addControlPoints=c.addControlPoints;this.attr=c.attr;this.attrsFromOptions=c.attrsFromOptions;this.destroy=c.destroy;this.getPointsOptions=c.getPointsOptions;this.init=c.init;this.linkPoints=c.linkPoints;this.point=c.point;this.rotate=c.rotate;this.scale=c.scale;this.setControlPointsVisibility=c.setControlPointsVisibility;this.shouldBeDrawn=c.shouldBeDrawn;this.transform=c.transform;
this.transformPoint=c.transformPoint;this.translateShape=c.translateShape;this.update=c.update;this.init(a,b,d);this.collection="labels"}a.alignedPosition=function(a,b){var d=a.align,p=a.verticalAlign,f=(b.x||0)+(a.x||0),h=(b.y||0)+(a.y||0),e,c;"right"===d?e=1:"center"===d&&(e=2);e&&(f+=(b.width-(a.width||0))/e);"bottom"===p?c=1:"middle"===p&&(c=2);c&&(h+=(b.height-(a.height||0))/c);return{x:Math.round(f),y:Math.round(h)}};a.justifiedOptions=function(a,b,d,h){var f=d.align,p=d.verticalAlign,e=b.box?
0:b.padding||0,c=b.getBBox();b={align:f,verticalAlign:p,x:d.x,y:d.y,width:b.width,height:b.height};d=(h.x||0)-a.plotLeft;h=(h.y||0)-a.plotTop;var t=d+e;0>t&&("right"===f?b.align="left":b.x=(b.x||0)-t);t=d+c.width-e;t>a.plotWidth&&("left"===f?b.align="right":b.x=(b.x||0)+a.plotWidth-t);t=h+e;0>t&&("bottom"===p?b.verticalAlign="top":b.y=(b.y||0)-t);t=h+c.height-e;t>a.plotHeight&&("top"===p?b.verticalAlign="bottom":b.y=(b.y||0)+a.plotHeight-t);return b};a.prototype.translatePoint=function(a,b){c.translatePoint.call(this,
a,b,0)};a.prototype.translate=function(a,b){var d=this.annotation.chart,f=this.annotation.userOptions,p=d.annotations.indexOf(this.annotation);p=d.options.annotations[p];d.inverted&&(d=a,a=b,b=d);this.options.x+=a;this.options.y+=b;p[this.collection][this.index].x=this.options.x;p[this.collection][this.index].y=this.options.y;f[this.collection][this.index].x=this.options.x;f[this.collection][this.index].y=this.options.y};a.prototype.render=function(b){var d=this.options,f=this.attrsFromOptions(d),
h=d.style;this.graphic=this.annotation.chart.renderer.label("",0,-9999,d.shape,null,null,d.useHTML,null,"annotation-label").attr(f).add(b);this.annotation.chart.styledMode||("contrast"===h.color&&(h.color=this.annotation.chart.renderer.getContrast(-1<a.shapesWithoutBackground.indexOf(d.shape)?"#FFFFFF":d.backgroundColor)),this.graphic.css(d.style).shadow(d.shadow));d.className&&this.graphic.addClass(d.className);this.graphic.labelrank=d.labelrank;c.render.call(this)};a.prototype.redraw=function(a){var b=
this.options,d=this.text||b.format||b.text,f=this.graphic,h=this.points[0];f.attr({text:d?e(d,h.getLabelConfig(),this.annotation.chart):b.formatter.call(h,this)});b=this.anchor(h);(d=this.position(b))?(f.alignAttr=d,d.anchorX=b.absolutePosition.x,d.anchorY=b.absolutePosition.y,f[a?"animate":"attr"](d)):f.attr({x:0,y:-9999});f.placed=!!d;c.redraw.call(this,a)};a.prototype.anchor=function(a){var b=c.anchor.apply(this,arguments),d=this.options.x||0,f=this.options.y||0;b.absolutePosition.x-=d;b.absolutePosition.y-=
f;b.relativePosition.x-=d;b.relativePosition.y-=f;return b};a.prototype.position=function(f){var h=this.graphic,e=this.annotation.chart,c=this.points[0],g=this.options,m=f.absolutePosition,l=f.relativePosition,k=c.series.visible&&n.prototype.isInsidePlot.call(c);f=h.width;f=void 0===f?0:f;var y=h.height;y=void 0===y?0:y;if(k){if(g.distance)var q=r.prototype.getPosition.call({chart:e,distance:d(g.distance,16)},f,y,{plotX:l.x,plotY:l.y,negative:c.negative,ttBelow:c.ttBelow,h:l.height||l.width});else g.positioner?
q=g.positioner.call(this):(c={x:m.x,y:m.y,width:0,height:0},q=a.alignedPosition(b(g,{width:f,height:y}),c),"justify"===this.options.overflow&&(q=a.alignedPosition(a.justifiedOptions(e,h,g,q),c)));g.crop&&(h=q.x-e.plotLeft,g=q.y-e.plotTop,k=e.isInsidePlot(h,g)&&e.isInsidePlot(h+f,g+y))}return k?q:null};a.attrsMap={backgroundColor:"fill",borderColor:"stroke",borderWidth:"stroke-width",zIndex:"zIndex",borderRadius:"r",padding:"padding"};a.shapesWithoutBackground=["connector"];return a}();l.connector=
function(b,d,p,e,c){var f=c&&c.anchorX;c=c&&c.anchorY;var h=p/2;if(a(f)&&a(c)){var t=[["M",f,c]];var g=d-c;0>g&&(g=-e-g);g<p&&(h=f<b+p/2?g:p-g);c>d+e?t.push(["L",b+h,d+e]):c<d?t.push(["L",b+h,d]):f<b?t.push(["L",b,d+e/2]):f>b+p&&t.push(["L",b+p,d+e/2])}return t||[]};return m});v(c,"Extensions/Annotations/Controllables/ControllableImage.js",[c["Extensions/Annotations/Controllables/ControllableLabel.js"],c["Extensions/Annotations/Mixins/ControllableMixin.js"]],function(c,l){return function(){function g(c,
g,m){this.addControlPoints=l.addControlPoints;this.anchor=l.anchor;this.attr=l.attr;this.attrsFromOptions=l.attrsFromOptions;this.destroy=l.destroy;this.getPointsOptions=l.getPointsOptions;this.init=l.init;this.linkPoints=l.linkPoints;this.point=l.point;this.rotate=l.rotate;this.scale=l.scale;this.setControlPointsVisibility=l.setControlPointsVisibility;this.shouldBeDrawn=l.shouldBeDrawn;this.transform=l.transform;this.transformPoint=l.transformPoint;this.translatePoint=l.translatePoint;this.translateShape=
l.translateShape;this.update=l.update;this.type="image";this.translate=l.translateShape;this.init(c,g,m);this.collection="shapes"}g.prototype.render=function(c){var g=this.attrsFromOptions(this.options),m=this.options;this.graphic=this.annotation.chart.renderer.image(m.src,0,-9E9,m.width,m.height).attr(g).add(c);this.graphic.width=m.width;this.graphic.height=m.height;l.render.call(this)};g.prototype.redraw=function(g){var k=this.anchor(this.points[0]);if(k=c.prototype.position.call(this,k))this.graphic[g?
"animate":"attr"]({x:k.x,y:k.y});else this.graphic.attr({x:0,y:-9E9});this.graphic.placed=!!k;l.redraw.call(this,g)};g.attrsMap={width:"width",height:"height",zIndex:"zIndex"};return g}()});v(c,"Extensions/Annotations/Annotations.js",[c["Core/Animation/AnimationUtilities.js"],c["Core/Chart/Chart.js"],c["Extensions/Annotations/Mixins/ControllableMixin.js"],c["Extensions/Annotations/Controllables/ControllableRect.js"],c["Extensions/Annotations/Controllables/ControllableCircle.js"],c["Extensions/Annotations/Controllables/ControllableEllipse.js"],
c["Extensions/Annotations/Controllables/ControllablePath.js"],c["Extensions/Annotations/Controllables/ControllableImage.js"],c["Extensions/Annotations/Controllables/ControllableLabel.js"],c["Extensions/Annotations/ControlPoint.js"],c["Extensions/Annotations/Mixins/EventEmitterMixin.js"],c["Core/Globals.js"],c["Extensions/Annotations/MockPoint.js"],c["Core/Pointer.js"],c["Core/Utilities.js"]],function(c,l,n,k,r,m,e,b,a,d,h,f,p,t,x){var g=c.getDeferredAnimation;c=l.prototype;var E=x.addEvent,D=x.defined,
z=x.destroyObjectProperties,y=x.erase,q=x.extend,w=x.find,B=x.fireEvent,u=x.merge,C=x.pick,G=x.splat;x=x.wrap;var F=function(){function f(a,b){this.annotation=void 0;this.coll="annotations";this.shapesGroup=this.labelsGroup=this.labelCollector=this.group=this.graphic=this.animationConfig=this.collection=void 0;this.chart=a;this.points=[];this.controlPoints=[];this.coll="annotations";this.labels=[];this.shapes=[];this.options=u(this.defaultOptions,b);this.userOptions=b;b=this.getLabelsAndShapesOptions(this.options,
b);this.options.labels=b.labels;this.options.shapes=b.shapes;this.init(a,this.options)}f.prototype.init=function(){var a=this.chart,b=this.options.animation;this.linkPoints();this.addControlPoints();this.addShapes();this.addLabels();this.setLabelCollector();this.animationConfig=g(a,b)};f.prototype.getLabelsAndShapesOptions=function(a,b){var d={};["labels","shapes"].forEach(function(f){a[f]&&(d[f]=b[f]?G(b[f]).map(function(b,d){return u(a[f][d],b)}):a[f])});return d};f.prototype.addShapes=function(){(this.options.shapes||
[]).forEach(function(a,b){a=this.initShape(a,b);u(!0,this.options.shapes[b],a.options)},this)};f.prototype.addLabels=function(){(this.options.labels||[]).forEach(function(a,b){a=this.initLabel(a,b);u(!0,this.options.labels[b],a.options)},this)};f.prototype.addClipPaths=function(){this.setClipAxes();this.clipXAxis&&this.clipYAxis&&this.options.crop&&(this.clipRect=this.chart.renderer.clipRect(this.getClipBox()))};f.prototype.setClipAxes=function(){var a=this.chart.xAxis,b=this.chart.yAxis,d=(this.options.labels||
[]).concat(this.options.shapes||[]).reduce(function(d,f){f=f&&(f.point||f.points&&f.points[0]);return[a[f&&f.xAxis]||d[0],b[f&&f.yAxis]||d[1]]},[]);this.clipXAxis=d[0];this.clipYAxis=d[1]};f.prototype.getClipBox=function(){if(this.clipXAxis&&this.clipYAxis)return{x:this.clipXAxis.left,y:this.clipYAxis.top,width:this.clipXAxis.width,height:this.clipYAxis.height}};f.prototype.setLabelCollector=function(){var a=this;a.labelCollector=function(){return a.labels.reduce(function(a,b){b.options.allowOverlap||
a.push(b.graphic);return a},[])};a.chart.labelCollectors.push(a.labelCollector)};f.prototype.setOptions=function(a){this.options=u(this.defaultOptions,a)};f.prototype.redraw=function(a){this.linkPoints();this.graphic||this.render();this.clipRect&&this.clipRect.animate(this.getClipBox());this.redrawItems(this.shapes,a);this.redrawItems(this.labels,a);n.redraw.call(this,a)};f.prototype.redrawItems=function(a,b){for(var d=a.length;d--;)this.redrawItem(a[d],b)};f.prototype.renderItems=function(a){for(var b=
a.length;b--;)this.renderItem(a[b])};f.prototype.render=function(){var a=this.chart.renderer;this.graphic=a.g("annotation").attr({opacity:0,zIndex:this.options.zIndex,visibility:this.options.visible?"visible":"hidden"}).add();this.shapesGroup=a.g("annotation-shapes").add(this.graphic);this.options.crop&&this.shapesGroup.clip(this.chart.plotBoxClip);this.labelsGroup=a.g("annotation-labels").attr({translateX:0,translateY:0}).add(this.graphic);this.addClipPaths();this.clipRect&&this.graphic.clip(this.clipRect);
this.renderItems(this.shapes);this.renderItems(this.labels);this.addEvents();n.render.call(this)};f.prototype.setVisibility=function(a){var b=this.options,d=this.chart.navigationBindings;a=C(a,!b.visible);this.graphic.attr("visibility",a?"visible":"hidden");a||(this.setControlPointsVisibility(!1),d.activeAnnotation===this&&d.popup&&"annotation-toolbar"===d.popup.formType&&B(d,"closePopup"));b.visible=a};f.prototype.setControlPointsVisibility=function(a){var b=function(b){b.setControlPointsVisibility(a)};
n.setControlPointsVisibility.call(this,a);this.shapes.forEach(b);this.labels.forEach(b)};f.prototype.destroy=function(){var a=this.chart,b=function(a){a.destroy()};this.labels.forEach(b);this.shapes.forEach(b);this.clipYAxis=this.clipXAxis=null;y(a.labelCollectors,this.labelCollector);h.destroy.call(this);n.destroy.call(this);z(this,a)};f.prototype.remove=function(){return this.chart.removeAnnotation(this)};f.prototype.update=function(a,b){var d=this.chart,f=this.getLabelsAndShapesOptions(this.userOptions,
a),h=d.annotations.indexOf(this);a=u(!0,this.userOptions,a);a.labels=f.labels;a.shapes=f.shapes;this.destroy();this.constructor(d,a);d.options.annotations[h]=a;this.isUpdating=!0;C(b,!0)&&d.redraw();B(this,"afterUpdate");this.isUpdating=!1};f.prototype.initShape=function(a,b){a=u(this.options.shapeOptions,{controlPointOptions:this.options.controlPointOptions},a);b=new f.shapesMap[a.type](this,a,b);b.itemType="shape";this.shapes.push(b);return b};f.prototype.initLabel=function(b,d){b=u(this.options.labelOptions,
{controlPointOptions:this.options.controlPointOptions},b);d=new a(this,b,d);d.itemType="label";this.labels.push(d);return d};f.prototype.redrawItem=function(a,b){a.linkPoints();a.shouldBeDrawn()?(a.graphic||this.renderItem(a),a.redraw(C(b,!0)&&a.graphic.placed),a.points.length&&this.adjustVisibility(a)):this.destroyItem(a)};f.prototype.adjustVisibility=function(a){var b=!1,d=a.graphic;a.points.forEach(function(a){!1!==a.series.visible&&!1!==a.visible&&(b=!0)});b?"hidden"===d.visibility&&d.show():
d.hide()};f.prototype.destroyItem=function(a){y(this[a.itemType+"s"],a);a.destroy()};f.prototype.renderItem=function(a){a.render("label"===a.itemType?this.labelsGroup:this.shapesGroup)};f.ControlPoint=d;f.MockPoint=p;f.shapesMap={rect:k,circle:r,ellipse:m,path:e,image:b};f.types={};return f}();u(!0,F.prototype,n,h,u(F.prototype,{nonDOMEvents:["add","afterUpdate","drag","remove"],defaultOptions:{visible:!0,animation:{},crop:!0,draggable:"xy",labelOptions:{align:"center",allowOverlap:!1,backgroundColor:"rgba(0, 0, 0, 0.75)",
borderColor:"#000000",borderRadius:3,borderWidth:1,className:"highcharts-no-tooltip",crop:!1,formatter:function(){return D(this.y)?this.y:"Annotation label"},includeInDataExport:!0,overflow:"justify",padding:5,shadow:!1,shape:"callout",style:{fontSize:"11px",fontWeight:"normal",color:"contrast"},useHTML:!1,verticalAlign:"bottom",x:0,y:-16},shapeOptions:{stroke:"rgba(0, 0, 0, 0.75)",strokeWidth:1,fill:"rgba(0, 0, 0, 0.75)",r:0,snap:2},controlPointOptions:{symbol:"circle",width:10,height:10,style:{cursor:"pointer",
fill:"#ffffff",stroke:"#000000","stroke-width":2},visible:!1,events:{}},events:{},zIndex:6}}));f.extendAnnotation=function(a,b,d,f){b=b||F;q(a.prototype,u(b.prototype,d));a.prototype.defaultOptions=u(a.prototype.defaultOptions,f||{})};q(c,{initAnnotation:function(a){a=new (F.types[a.type]||F)(this,a);this.annotations.push(a);return a},addAnnotation:function(a,b){a=this.initAnnotation(a);this.options.annotations.push(a.options);C(b,!0)&&(a.redraw(),a.graphic.attr({opacity:1}));return a},removeAnnotation:function(a){var b=
this.annotations,d="annotations"===a.coll?a:w(b,function(b){return b.options.id===a});d&&(B(d,"remove"),y(this.options.annotations,d.options),y(b,d),d.destroy())},drawAnnotations:function(){this.plotBoxClip.attr(this.plotBox);this.annotations.forEach(function(a){a.redraw();a.graphic.animate({opacity:1},a.animationConfig)})}});c.collectionsWithUpdate.push("annotations");c.collectionsWithInit.annotations=[c.addAnnotation];E(l,"afterInit",function(){this.annotations=[];this.options.annotations||(this.options.annotations=
[])});c.callbacks.push(function(a){a.plotBoxClip=this.renderer.clipRect(this.plotBox);a.controlPointsGroup=a.renderer.g("control-points").attr({zIndex:99}).clip(a.plotBoxClip).add();a.options.annotations.forEach(function(b,d){if(!a.annotations.some(function(a){return a.options===b})){var f=a.initAnnotation(b);a.options.annotations[d]=f.options}});a.drawAnnotations();E(a,"redraw",a.drawAnnotations);E(a,"destroy",function(){a.plotBoxClip.destroy();a.controlPointsGroup.destroy()});E(a,"exportData",function(b){var d=
(this.options.exporting&&this.options.exporting.csv||{}).columnHeaderFormatter,f=!b.dataRows[1].xValues,h=a.options.lang&&a.options.lang.exportData&&a.options.lang.exportData.annotationHeader,c=function(a){if(d){var b=d(a);if(!1!==b)return b}b=h+" "+a;return f?{columnTitle:b,topLevelColumnTitle:b}:b},e=b.dataRows[0].length,p=a.options.exporting&&a.options.exporting.csv&&a.options.exporting.csv.annotations&&a.options.exporting.csv.annotations.itemDelimiter,t=a.options.exporting&&a.options.exporting.csv&&
a.options.exporting.csv.annotations&&a.options.exporting.csv.annotations.join;a.annotations.forEach(function(a){a.options.labelOptions.includeInDataExport&&a.labels.forEach(function(a){if(a.options.text){var d=a.options.text;a.points.forEach(function(a){var f=a.x,h=a.series.xAxis?a.series.xAxis.options.index:-1,c=!1;if(-1===h){a=b.dataRows[0].length;for(var u=Array(a),g=0;g<a;++g)u[g]="";u.push(d);u.xValues=[];u.xValues[h]=f;b.dataRows.push(u);c=!0}c||b.dataRows.forEach(function(a,b){!c&&a.xValues&&
void 0!==h&&f===a.xValues[h]&&(t&&a.length>e?a[a.length-1]+=p+d:a.push(d),c=!0)});if(!c){a=b.dataRows[0].length;u=Array(a);for(g=0;g<a;++g)u[g]="";u[0]=f;u.push(d);u.xValues=[];void 0!==h&&(u.xValues[h]=f);b.dataRows.push(u)}})}})});var u=0;b.dataRows.forEach(function(a){u=Math.max(u,a.length)});for(var C=u-b.dataRows[0].length,g=0;g<C;g++){var G=c(g+1);f?(b.dataRows[0].push(G.topLevelColumnTitle),b.dataRows[1].push(G.columnTitle)):b.dataRows[0].push(G)}})});x(t.prototype,"onContainerMouseDown",function(a){this.chart.hasDraggedAnnotation||
a.apply(this,Array.prototype.slice.call(arguments,1))});f.Annotation=F;"";return F});v(c,"Extensions/Annotations/Types/BasicAnnotation.js",[c["Extensions/Annotations/Annotations.js"],c["Extensions/Annotations/MockPoint.js"],c["Core/Utilities.js"]],function(c,l,n){var g=this&&this.__extends||function(){var c=function(e,b){c=Object.setPrototypeOf||{__proto__:[]}instanceof Array&&function(a,b){a.__proto__=b}||function(a,b){for(var d in b)b.hasOwnProperty(d)&&(a[d]=b[d])};return c(e,b)};return function(e,
b){function a(){this.constructor=e}c(e,b);e.prototype=null===b?Object.create(b):(a.prototype=b.prototype,new a)}}();n=n.merge;var r=function(m){function e(b,a){return m.call(this,b,a)||this}g(e,m);e.prototype.addControlPoints=function(){var b=this.options,a=e.basicControlPoints,d=this.basicType;(b.labels||b.shapes).forEach(function(b){b.controlPoints=a[d]})};e.prototype.init=function(){var b=this.options;b.shapes?(delete b.labelOptions,this.basicType=b.shapes[0].type?b.shapes[0].type:"rectangle"):
(delete b.shapes,this.basicType="label");c.prototype.init.apply(this,arguments)};e.basicControlPoints={label:[{symbol:"triangle-down",positioner:function(b){if(!b.graphic.placed)return{x:0,y:-9E7};b=l.pointToPixels(b.points[0]);return{x:b.x-this.graphic.width/2,y:b.y-this.graphic.height/2}},events:{drag:function(b,a){b=this.mouseMoveToTranslation(b);a.translatePoint(b.x,b.y);a.annotation.userOptions.labels[0].point=a.options.point;a.redraw(!1)}}},{symbol:"square",positioner:function(b){return b.graphic.placed?
{x:b.graphic.alignAttr.x-this.graphic.width/2,y:b.graphic.alignAttr.y-this.graphic.height/2}:{x:0,y:-9E7}},events:{drag:function(b,a){b=this.mouseMoveToTranslation(b);a.translate(b.x,b.y);a.annotation.userOptions.labels[0].point=a.options.point;a.redraw(!1)}}}],rectangle:[{positioner:function(b){b=l.pointToPixels(b.points[2]);return{x:b.x-4,y:b.y-4}},events:{drag:function(b,a){var d=a.annotation,c=this.chart.pointer.getCoordinates(b);b=c.xAxis[0].value;c=c.yAxis[0].value;var f=a.options.points;f[1].x=
b;f[2].x=b;f[2].y=c;f[3].y=c;d.userOptions.shapes[0].points=a.options.points;d.redraw(!1)}}}],circle:[{positioner:function(b){var a=l.pointToPixels(b.points[0]);b=b.options.r;return{x:a.x+b*Math.cos(Math.PI/4)-this.graphic.width/2,y:a.y+b*Math.sin(Math.PI/4)-this.graphic.height/2}},events:{drag:function(b,a){var d=a.annotation;b=this.mouseMoveToTranslation(b);a.setRadius(Math.max(a.options.r+b.y/Math.sin(Math.PI/4),5));d.userOptions.shapes[0].r=a.options.r;d.userOptions.shapes[0].point=a.options.point;
a.redraw(!1)}}}],ellipse:[{positioner:function(b){b=b.getAbsolutePosition(b.points[0]);return{x:b.x-this.graphic.width/2,y:b.y-this.graphic.height/2}},events:{drag:function(b,a){var d=a.getAbsolutePosition(a.points[0]);a.translatePoint(b.chartX-d.x,b.chartY-d.y,0);a.redraw(!1)}}},{positioner:function(b){b=b.getAbsolutePosition(b.points[1]);return{x:b.x-this.graphic.width/2,y:b.y-this.graphic.height/2}},events:{drag:function(b,a){var d=a.getAbsolutePosition(a.points[1]);a.translatePoint(b.chartX-d.x,
b.chartY-d.y,1);a.redraw(!1)}}},{positioner:function(b){var a=b.getAbsolutePosition(b.points[0]),d=b.getAbsolutePosition(b.points[1]);b=b.getAttrs(a,d);return{x:b.cx-this.graphic.width/2+b.ry*Math.sin(b.angle*Math.PI/180),y:b.cy-this.graphic.height/2-b.ry*Math.cos(b.angle*Math.PI/180)}},events:{drag:function(b,a){var d=a.getAbsolutePosition(a.points[0]),c=a.getAbsolutePosition(a.points[1]);b=a.getDistanceFromLine(d,c,b.chartX,b.chartY);d=a.getYAxis();b=Math.abs(d.toValue(0)-d.toValue(b));a.setYRadius(b);
a.redraw(!1)}}}]};return e}(c);r.prototype.defaultOptions=n(c.prototype.defaultOptions,{});return c.types.basicAnnotation=r});v(c,"Extensions/Annotations/Types/CrookedLine.js",[c["Extensions/Annotations/Annotations.js"],c["Extensions/Annotations/ControlPoint.js"],c["Extensions/Annotations/MockPoint.js"],c["Core/Utilities.js"]],function(c,l,n,k){var g=this&&this.__extends||function(){var c=function(b,a){c=Object.setPrototypeOf||{__proto__:[]}instanceof Array&&function(a,b){a.__proto__=b}||function(a,
b){for(var d in b)b.hasOwnProperty(d)&&(a[d]=b[d])};return c(b,a)};return function(b,a){function d(){this.constructor=b}c(b,a);b.prototype=null===a?Object.create(a):(d.prototype=a.prototype,new d)}}(),m=k.merge;k=function(c){function b(a,b){return c.call(this,a,b)||this}g(b,c);b.prototype.setClipAxes=function(){this.clipXAxis=this.chart.xAxis[this.options.typeOptions.xAxis];this.clipYAxis=this.chart.yAxis[this.options.typeOptions.yAxis]};b.prototype.getPointsOptions=function(){var a=this.options.typeOptions;
return(a.points||[]).map(function(b){b.xAxis=a.xAxis;b.yAxis=a.yAxis;return b})};b.prototype.getControlPointsOptions=function(){return this.getPointsOptions()};b.prototype.addControlPoints=function(){this.getControlPointsOptions().forEach(function(a,b){b=new l(this.chart,this,m(this.options.controlPointOptions,a.controlPoint),b);this.controlPoints.push(b);a.controlPoint=b.options},this)};b.prototype.addShapes=function(){var a=this.options.typeOptions,b=this.initShape(m(a.line,{type:"path",points:this.points.map(function(a,
b){return function(a){return a.annotation.points[b]}})}),0);a.line=b.options};return b}(c);k.prototype.defaultOptions=m(c.prototype.defaultOptions,{typeOptions:{xAxis:0,yAxis:0,line:{fill:"none"}},controlPointOptions:{positioner:function(c){var b=this.graphic;c=n.pointToPixels(c.points[this.index]);return{x:c.x-b.width/2,y:c.y-b.height/2}},events:{drag:function(c,b){b.chart.isInsidePlot(c.chartX-b.chart.plotLeft,c.chartY-b.chart.plotTop,{visiblePlotOnly:!0})&&(c=this.mouseMoveToTranslation(c),b.translatePoint(c.x,
c.y,this.index),b.options.typeOptions.points[this.index].x=b.points[this.index].x,b.options.typeOptions.points[this.index].y=b.points[this.index].y,b.redraw(!1))}}}});return c.types.crookedLine=k});v(c,"Extensions/Annotations/Types/ElliottWave.js",[c["Extensions/Annotations/Annotations.js"],c["Extensions/Annotations/Types/CrookedLine.js"],c["Core/Utilities.js"]],function(c,l,n){var g=this&&this.__extends||function(){var c=function(e,b){c=Object.setPrototypeOf||{__proto__:[]}instanceof Array&&function(a,
b){a.__proto__=b}||function(a,b){for(var d in b)b.hasOwnProperty(d)&&(a[d]=b[d])};return c(e,b)};return function(e,b){function a(){this.constructor=e}c(e,b);e.prototype=null===b?Object.create(b):(a.prototype=b.prototype,new a)}}(),r=n.merge;n=function(c){function e(b,a){return c.call(this,b,a)||this}g(e,c);e.prototype.addLabels=function(){this.getPointsOptions().forEach(function(b,a){var d=this.initLabel(r(b.label,{text:this.options.typeOptions.labels[a],point:function(b){return b.annotation.points[a]}}),
!1);b.label=d.options},this)};return e}(l);n.prototype.defaultOptions=r(l.prototype.defaultOptions,{typeOptions:{labels:"(0) (A) (B) (C) (D) (E)".split(" "),line:{strokeWidth:1}},labelOptions:{align:"center",allowOverlap:!0,crop:!0,overflow:"none",type:"rect",backgroundColor:"none",borderWidth:0,y:-5}});return c.types.elliottWave=n});v(c,"Extensions/Annotations/Types/Tunnel.js",[c["Extensions/Annotations/Annotations.js"],c["Extensions/Annotations/ControlPoint.js"],c["Extensions/Annotations/Types/CrookedLine.js"],
c["Extensions/Annotations/MockPoint.js"],c["Core/Utilities.js"]],function(c,l,n,k,r){var g=this&&this.__extends||function(){var b=function(a,d){b=Object.setPrototypeOf||{__proto__:[]}instanceof Array&&function(a,b){a.__proto__=b}||function(a,b){for(var d in b)b.hasOwnProperty(d)&&(a[d]=b[d])};return b(a,d)};return function(a,d){function c(){this.constructor=a}b(a,d);a.prototype=null===d?Object.create(d):(c.prototype=d.prototype,new c)}}(),e=r.merge;r=function(b){function a(a,c){return b.call(this,
a,c)||this}g(a,b);a.prototype.getPointsOptions=function(){var a=n.prototype.getPointsOptions.call(this);a[2]=this.heightPointOptions(a[1]);a[3]=this.heightPointOptions(a[0]);return a};a.prototype.getControlPointsOptions=function(){return this.getPointsOptions().slice(0,2)};a.prototype.heightPointOptions=function(a){a=e(a);a.y+=this.options.typeOptions.height;return a};a.prototype.addControlPoints=function(){n.prototype.addControlPoints.call(this);var a=this.options,b=a.typeOptions;a=new l(this.chart,
this,e(a.controlPointOptions,b.heightControlPoint),2);this.controlPoints.push(a);b.heightControlPoint=a.options};a.prototype.addShapes=function(){this.addLine();this.addBackground()};a.prototype.addLine=function(){var a=this.initShape(e(this.options.typeOptions.line,{type:"path",points:[this.points[0],this.points[1],function(a){a=k.pointToOptions(a.annotation.points[2]);a.command="M";return a},this.points[3]]}),0);this.options.typeOptions.line=a.options};a.prototype.addBackground=function(){var a=
this.initShape(e(this.options.typeOptions.background,{type:"path",points:this.points.slice()}),1);this.options.typeOptions.background=a.options};a.prototype.translateSide=function(a,b,f){f=Number(f);var d=0===f?3:2;this.translatePoint(a,b,f);this.translatePoint(a,b,d)};a.prototype.translateHeight=function(a){this.translatePoint(0,a,2);this.translatePoint(0,a,3);this.options.typeOptions.height=this.points[3].y-this.points[0].y};return a}(n);r.prototype.defaultOptions=e(n.prototype.defaultOptions,{typeOptions:{background:{fill:"rgba(130, 170, 255, 0.4)",
strokeWidth:0},line:{strokeWidth:1},height:-2,heightControlPoint:{positioner:function(b){var a=k.pointToPixels(b.points[2]);b=k.pointToPixels(b.points[3]);var d=(a.x+b.x)/2;return{x:d-this.graphic.width/2,y:(b.y-a.y)/(b.x-a.x)*(d-a.x)+a.y-this.graphic.height/2}},events:{drag:function(b,a){a.chart.isInsidePlot(b.chartX-a.chart.plotLeft,b.chartY-a.chart.plotTop,{visiblePlotOnly:!0})&&(a.translateHeight(this.mouseMoveToTranslation(b).y),a.redraw(!1))}}}},controlPointOptions:{events:{drag:function(b,
a){a.chart.isInsidePlot(b.chartX-a.chart.plotLeft,b.chartY-a.chart.plotTop,{visiblePlotOnly:!0})&&(b=this.mouseMoveToTranslation(b),a.translateSide(b.x,b.y,!!this.index),a.redraw(!1))}}}});return c.types.tunnel=r});v(c,"Extensions/Annotations/Types/InfinityLine.js",[c["Extensions/Annotations/Annotations.js"],c["Extensions/Annotations/Types/CrookedLine.js"],c["Extensions/Annotations/MockPoint.js"],c["Core/Utilities.js"]],function(c,l,n,k){var g=this&&this.__extends||function(){var c=function(b,a){c=
Object.setPrototypeOf||{__proto__:[]}instanceof Array&&function(a,b){a.__proto__=b}||function(a,b){for(var d in b)b.hasOwnProperty(d)&&(a[d]=b[d])};return c(b,a)};return function(b,a){function d(){this.constructor=b}c(b,a);b.prototype=null===a?Object.create(a):(d.prototype=a.prototype,new d)}}(),m=k.merge;k=function(c){function b(a,b){return c.call(this,a,b)||this}g(b,c);b.edgePoint=function(a,d){return function(c){c=c.annotation;var f=c.points,p=c.options.typeOptions.type;if("horizontalLine"===p||
"verticalLine"===p)f=[f[0],new n(c.chart,f[0].target,{x:f[0].x+ +("horizontalLine"===p),y:f[0].y+ +("verticalLine"===p),xAxis:f[0].options.xAxis,yAxis:f[0].options.yAxis})];return b.findEdgePoint(f[a],f[d])}};b.findEdgeCoordinate=function(a,b,c,f){var d="x"===c?"y":"x";return(b[c]-a[c])*(f-a[d])/(b[d]-a[d])+a[c]};b.findEdgePoint=function(a,d){var c=a.series.chart,f=a.series.xAxis,p=d.series.yAxis,e=n.pointToPixels(a);d=n.pointToPixels(d);var g=d.x-e.x,l=d.y-e.y,k=f.left,m=k+f.width;f=p.top;p=f+p.height;
m=0>g?k:m;var r=0>l?f:p;k={x:0===g?e.x:m,y:0===l?e.y:r};0!==g&&0!==l&&(g=b.findEdgeCoordinate(e,d,"y",m),e=b.findEdgeCoordinate(e,d,"x",r),g>=f&&g<=p?(k.x=m,k.y=g):(k.x=e,k.y=r));k.x-=c.plotLeft;k.y-=c.plotTop;a.series.chart.inverted&&(a=k.x,k.x=k.y,k.y=a);return k};b.prototype.addShapes=function(){var a=this.options.typeOptions,d=[this.points[0],b.endEdgePoint];a.type.match(/line/gi)&&(d[0]=b.startEdgePoint);d=this.initShape(m(a.line,{type:"path",points:d}),0);a.line=d.options};b.endEdgePoint=b.edgePoint(0,
1);b.startEdgePoint=b.edgePoint(1,0);return b}(l);k.prototype.defaultOptions=m(l.prototype.defaultOptions,{});c.types.infinityLine=k;"";return k});v(c,"Extensions/Annotations/Types/TimeCycles.js",[c["Extensions/Annotations/Annotations.js"],c["Extensions/Annotations/Types/CrookedLine.js"],c["Extensions/Annotations/ControlPoint.js"],c["Core/Utilities.js"]],function(c,l,n,k){function g(a,b,c,e){for(var d=[],f=1;f<=b;f++)d.push(["A",a/2,a/2,0,1,1,c+f*a,e]);return d}var m=this&&this.__extends||function(){var a=
function(b,d){a=Object.setPrototypeOf||{__proto__:[]}instanceof Array&&function(a,b){a.__proto__=b}||function(a,b){for(var d in b)b.hasOwnProperty(d)&&(a[d]=b[d])};return a(b,d)};return function(b,d){function c(){this.constructor=b}a(b,d);b.prototype=null===d?Object.create(d):(c.prototype=d.prototype,new c)}}(),e=k.merge,b=k.isNumber,a=k.defined;k=function(d){function c(){return null!==d&&d.apply(this,arguments)||this}m(c,d);c.prototype.init=function(b,c,e){a(c.yAxis)&&c.points.forEach(function(a){a.yAxis=
c.yAxis});a(c.xAxis)&&c.points.forEach(function(a){a.xAxis=c.xAxis});d.prototype.init.call(this,b,c,e)};c.prototype.setPath=function(){this.shapes[0].options.d=this.getPath()};c.prototype.getPath=function(){return[["M",this.startX,this.y]].concat(g(this.pixelInterval,this.numberOfCircles,this.startX,this.y))};c.prototype.addShapes=function(){var a=this.options.typeOptions;this.setPathProperties();var b=this.initShape(e(a.line,{type:"path",d:this.getPath(),points:this.options.points}),0);a.line=b.options};
c.prototype.addControlPoints=function(){var a=this,b=this.options,d=b.typeOptions;b.controlPointOptions.style.cursor=this.chart.inverted?"ns-resize":"ew-resize";d.controlPointOptions.forEach(function(d){d=e(b.controlPointOptions,d);d=new n(a.chart,a,d,0);a.controlPoints.push(d)})};c.prototype.setPathProperties=function(){var a=this.options.typeOptions,d=a.points;if(d){var c=d[0],e=this.chart.xAxis[a.xAxis||0],h=this.chart.yAxis[a.yAxis||0];a=c.x;var g=c.y;c=d[1].x;a&&c&&(d=b(g)?h.toPixels(g):h.top+
h.height,a=b(a)?e.toPixels(a):e.left,h=b(c)?e.toPixels(c):e.left+30,h=Math.round(Math.max(Math.abs(h-a),2)),c=Math.floor(e.len/h)+2,this.startX=a-(Math.floor((a-e.left)/h)+1)*h,this.y=d,this.pixelInterval=h,this.numberOfCircles=c)}};c.prototype.redraw=function(a){this.setPathProperties();this.setPath();d.prototype.redraw.call(this,a)};return c}(l);k.prototype.defaultOptions=e(l.prototype.defaultOptions,{typeOptions:{controlPointOptions:[{positioner:function(a){return{x:a.anchor(a.points[0]).absolutePosition.x-
this.graphic.width/2,y:a.y-this.graphic.height}},events:{drag:function(a,b){var d=b.anchor(b.points[0]).absolutePosition;b.translatePoint(a.chartX-d.x,0,0);b.redraw(!1)}}},{positioner:function(a){return{x:a.anchor(a.points[1]).absolutePosition.x-this.graphic.width/2,y:a.y-this.graphic.height}},events:{drag:function(a,b){var d=b.anchor(b.points[1]).absolutePosition;b.translatePoint(a.chartX-d.x,0,1);b.redraw(!1)}}}]}});c.types.timeCycles=k;"";return k});v(c,"Extensions/Annotations/Types/Fibonacci.js",
[c["Extensions/Annotations/Annotations.js"],c["Extensions/Annotations/MockPoint.js"],c["Extensions/Annotations/Types/Tunnel.js"],c["Core/Utilities.js"]],function(c,l,n,k){var g=this&&this.__extends||function(){var b=function(a,d){b=Object.setPrototypeOf||{__proto__:[]}instanceof Array&&function(a,b){a.__proto__=b}||function(a,b){for(var d in b)b.hasOwnProperty(d)&&(a[d]=b[d])};return b(a,d)};return function(a,d){function c(){this.constructor=a}b(a,d);a.prototype=null===d?Object.create(d):(c.prototype=
d.prototype,new c)}}(),m=k.merge,e=function(b,a){return function(){var d=this.annotation;if(!d.startRetracements||!d.endRetracements)return[];var c=this.anchor(d.startRetracements[b]).absolutePosition,f=this.anchor(d.endRetracements[b]).absolutePosition;c=[["M",Math.round(c.x),Math.round(c.y)],["L",Math.round(f.x),Math.round(f.y)]];a&&(f=this.anchor(d.endRetracements[b-1]).absolutePosition,d=this.anchor(d.startRetracements[b-1]).absolutePosition,c.push(["L",Math.round(f.x),Math.round(f.y)],["L",Math.round(d.x),
Math.round(d.y)]));return c}};k=function(b){function a(a,c){return b.call(this,a,c)||this}g(a,b);a.prototype.linkPoints=function(){b.prototype.linkPoints.call(this);this.linkRetracementsPoints()};a.prototype.linkRetracementsPoints=function(){var b=this.points,c=b[0].y-b[3].y,f=b[1].y-b[2].y,e=b[0].x,g=b[1].x;a.levels.forEach(function(a,d){var p=b[0].y-c*a;a=b[1].y-f*a;this.startRetracements=this.startRetracements||[];this.endRetracements=this.endRetracements||[];this.linkRetracementPoint(d,e,p,this.startRetracements);
this.linkRetracementPoint(d,g,a,this.endRetracements)},this)};a.prototype.linkRetracementPoint=function(a,b,c,e){var d=e[a],f=this.options.typeOptions;d?(d.options.x=b,d.options.y=c,d.refresh()):e[a]=new l(this.chart,this,{x:b,y:c,xAxis:f.xAxis,yAxis:f.yAxis})};a.prototype.addShapes=function(){a.levels.forEach(function(a,b){var d=this.options.typeOptions;a=d.backgroundColors;var c=d.lineColor;d=d.lineColors;this.initShape({type:"path",d:e(b),stroke:d[b]||c},b);0<b&&this.initShape({type:"path",fill:a[b-
1],strokeWidth:0,d:e(b,!0)})},this)};a.prototype.addLabels=function(){a.levels.forEach(function(a,b){var d=this.options.typeOptions;a=this.initLabel(m(d.labels[b],{point:function(a){return l.pointToOptions(a.annotation.startRetracements[b])},text:a.toString()}));d.labels[b]=a.options},this)};a.levels=[0,.236,.382,.5,.618,.786,1];return a}(n);k.prototype.defaultOptions=m(n.prototype.defaultOptions,{typeOptions:{height:2,backgroundColors:"rgba(130, 170, 255, 0.4);rgba(139, 191, 216, 0.4);rgba(150, 216, 192, 0.4);rgba(156, 229, 161, 0.4);rgba(162, 241, 130, 0.4);rgba(169, 255, 101, 0.4)".split(";"),
lineColor:"#999999",lineColors:[],labels:[]},labelOptions:{allowOverlap:!0,align:"right",backgroundColor:"none",borderWidth:0,crop:!1,overflow:"none",shape:"rect",style:{color:"grey"},verticalAlign:"middle",y:0}});return c.types.fibonacci=k});v(c,"Extensions/Annotations/Types/FibonacciTimeZones.js",[c["Extensions/Annotations/Annotations.js"],c["Extensions/Annotations/ControlPoint.js"],c["Extensions/Annotations/Types/CrookedLine.js"],c["Extensions/Annotations/Types/InfinityLine.js"],c["Extensions/Annotations/MockPoint.js"],
c["Core/Utilities.js"]],function(c,l,n,k,r,m){var e=this&&this.__extends||function(){var a=function(b,c){a=Object.setPrototypeOf||{__proto__:[]}instanceof Array&&function(a,b){a.__proto__=b}||function(a,b){for(var d in b)b.hasOwnProperty(d)&&(a[d]=b[d])};return a(b,c)};return function(b,c){function d(){this.constructor=b}a(b,c);b.prototype=null===c?Object.create(c):(d.prototype=c.prototype,new d)}}(),b=m.merge;m=function(a){function d(){return null!==a&&a.apply(this,arguments)||this}e(d,a);d.prototype.edgePoint=
function(a,b,d){return function(c){var f=c.annotation.chart;c=c.annotation.points;var e=c[0].series.xAxis.toValue(c[0].plotX+(f.inverted?f.plotTop:f.plotLeft)+d*(1<c.length?c[1].plotX-c[0].plotX:0));c=[new r(f,c[0].target,{x:e,y:0,xAxis:c[0].options.xAxis,yAxis:c[0].options.yAxis}),new r(f,c[0].target,{x:e,y:1,xAxis:c[0].options.xAxis,yAxis:c[0].options.yAxis})];return k.findEdgePoint(c[a],c[b])}};d.prototype.addShapes=function(){for(var a=1,d=1,c=0;11>c;c++){var e=c?a:0;e=[this.edgePoint(1,0,e),
this.edgePoint(0,1,e)];d=a+d;a=d-a;1===c&&(this.secondLineEdgePoints=[e[0],e[1]]);this.initShape(b(this.options.typeOptions.line,{type:"path",points:e}),c)}};d.prototype.addControlPoints=function(){var a=this.options,d=a.typeOptions;a=new l(this.chart,this,b(a.controlPointOptions,d.controlPointOptions),0);this.controlPoints.push(a);d.controlPointOptions=a.options};return d}(n);m.prototype.defaultOptions=b(n.prototype.defaultOptions,{typeOptions:{line:{stroke:"rgba(0, 0, 0, 0.75)",strokeWidth:1,fill:void 0},
controlPointOptions:{positioner:function(){var a=this.target,b=this.graphic,c=a.secondLineEdgePoints,f={annotation:a};var e=c[0](f).y;var g=c[1](f).y;a=this.chart.plotLeft;var k=this.chart.plotTop;c=c[0](f).x;e=(e+g)/2;this.chart.inverted&&(e=[e,c],c=e[0],e=e[1]);return{x:a+c-b.width/2,y:k+e-b.height/2}},events:{drag:function(a,b){b.chart.isInsidePlot(a.chartX-b.chart.plotLeft,a.chartY-b.chart.plotTop,{visiblePlotOnly:!0})&&(a=this.mouseMoveToTranslation(a),b.translatePoint(a.x,0,1),b.redraw(!1))}}}}});
c.types.fibonacciTimeZones=m;"";return m});v(c,"Extensions/Annotations/Types/Pitchfork.js",[c["Extensions/Annotations/Annotations.js"],c["Extensions/Annotations/Types/InfinityLine.js"],c["Extensions/Annotations/MockPoint.js"],c["Core/Utilities.js"]],function(c,l,n,k){var g=this&&this.__extends||function(){var c=function(b,a){c=Object.setPrototypeOf||{__proto__:[]}instanceof Array&&function(a,b){a.__proto__=b}||function(a,b){for(var d in b)b.hasOwnProperty(d)&&(a[d]=b[d])};return c(b,a)};return function(b,
a){function d(){this.constructor=b}c(b,a);b.prototype=null===a?Object.create(a):(d.prototype=a.prototype,new d)}}(),m=k.merge;k=function(c){function b(a,b){return c.call(this,a,b)||this}g(b,c);b.outerLineEdgePoint=function(a){return function(d){var c=d.annotation,f=c.points;return b.findEdgePoint(f[a],f[0],new n(c.chart,d,c.midPointOptions()))}};b.findEdgePoint=function(a,b,c){b=Math.atan2(c.plotY-b.plotY,c.plotX-b.plotX);return{x:a.plotX+1E7*Math.cos(b),y:a.plotY+1E7*Math.sin(b)}};b.middleLineEdgePoint=
function(a){var b=a.annotation;return l.findEdgePoint(b.points[0],new n(b.chart,a,b.midPointOptions()))};b.prototype.midPointOptions=function(){var a=this.points;return{x:(a[1].x+a[2].x)/2,y:(a[1].y+a[2].y)/2,xAxis:a[0].series.xAxis,yAxis:a[0].series.yAxis}};b.prototype.addShapes=function(){this.addLines();this.addBackgrounds()};b.prototype.addLines=function(){this.initShape({type:"path",points:[this.points[0],b.middleLineEdgePoint]},0);this.initShape({type:"path",points:[this.points[1],b.topLineEdgePoint]},
1);this.initShape({type:"path",points:[this.points[2],b.bottomLineEdgePoint]},2)};b.prototype.addBackgrounds=function(){var a=this.shapes,b=this.options.typeOptions,c=this.initShape(m(b.innerBackground,{type:"path",points:[function(a){var b=a.annotation;a=b.points;b=b.midPointOptions();return{x:(a[1].x+b.x)/2,y:(a[1].y+b.y)/2,xAxis:b.xAxis,yAxis:b.yAxis}},a[1].points[1],a[2].points[1],function(a){var b=a.annotation;a=b.points;b=b.midPointOptions();return{x:(b.x+a[2].x)/2,y:(b.y+a[2].y)/2,xAxis:b.xAxis,
yAxis:b.yAxis}}]}),3);a=this.initShape(m(b.outerBackground,{type:"path",points:[this.points[1],a[1].points[1],a[2].points[1],this.points[2]]}),4);b.innerBackground=c.options;b.outerBackground=a.options};b.topLineEdgePoint=b.outerLineEdgePoint(1);b.bottomLineEdgePoint=b.outerLineEdgePoint(0);return b}(l);k.prototype.defaultOptions=m(l.prototype.defaultOptions,{typeOptions:{innerBackground:{fill:"rgba(130, 170, 255, 0.4)",strokeWidth:0},outerBackground:{fill:"rgba(156, 229, 161, 0.4)",strokeWidth:0}}});
return c.types.pitchfork=k});v(c,"Extensions/Annotations/Types/VerticalLine.js",[c["Extensions/Annotations/Annotations.js"],c["Extensions/Annotations/MockPoint.js"],c["Core/Utilities.js"]],function(c,l,n){var g=this&&this.__extends||function(){var c=function(b,a){c=Object.setPrototypeOf||{__proto__:[]}instanceof Array&&function(a,b){a.__proto__=b}||function(a,b){for(var c in b)b.hasOwnProperty(c)&&(a[c]=b[c])};return c(b,a)};return function(b,a){function d(){this.constructor=b}c(b,a);b.prototype=
null===a?Object.create(a):(d.prototype=a.prototype,new d)}}(),r=n.merge,m=n.pick;n=function(c){function b(a,b){return c.call(this,a,b)||this}g(b,c);b.connectorFirstPoint=function(a){var b=a.annotation;a=b.chart;var c=a.inverted,f=b.points[0],e=m(f.series.yAxis&&f.series.yAxis.left,0),g=m(f.series.yAxis&&f.series.yAxis.top,0);b=b.options.typeOptions.label.offset;var k=l.pointToPixels(f,!0)[c?"x":"y"];return{x:f.x,xAxis:f.series.xAxis,y:k+b+(c?e-a.plotLeft:g-a.plotTop)}};b.connectorSecondPoint=function(a){var b=
a.annotation;a=b.chart;var c=a.inverted,e=b.options.typeOptions;b=b.points[0];var g=m(b.series.yAxis&&b.series.yAxis.left,0),t=m(b.series.yAxis&&b.series.yAxis.top,0),k=e.yOffset,n=l.pointToPixels(b,!0)[c?"x":"y"];0>e.label.offset&&(k*=-1);return{x:b.x,xAxis:b.series.xAxis,y:n+k+(c?g-a.plotLeft:t-a.plotTop)}};b.prototype.getPointsOptions=function(){return[this.options.typeOptions.point]};b.prototype.addShapes=function(){var a=this.options.typeOptions,c=this.initShape(r(a.connector,{type:"path",points:[b.connectorFirstPoint,
b.connectorSecondPoint]}),0);a.connector=c.options};b.prototype.addLabels=function(){var a=this.options.typeOptions,b=a.label,c=0,e=b.offset,g=0>b.offset?"bottom":"top",t="center";this.chart.inverted&&(c=b.offset,e=0,g="middle",t=0>b.offset?"right":"left");b=this.initLabel(r(b,{verticalAlign:g,align:t,x:c,y:e}));a.label=b.options};return b}(c);n.prototype.defaultOptions=r(c.prototype.defaultOptions,{typeOptions:{yOffset:10,label:{offset:-40,point:function(c){return c.annotation.points[0]},allowOverlap:!0,
backgroundColor:"none",borderWidth:0,crop:!0,overflow:"none",shape:"rect",text:"{y:.2f}"},connector:{strokeWidth:1,markerEnd:"arrow"}}});return c.types.verticalLine=n});v(c,"Extensions/Annotations/Types/Measure.js",[c["Extensions/Annotations/Annotations.js"],c["Extensions/Annotations/ControlPoint.js"],c["Core/Utilities.js"]],function(c,l,n){var g=this&&this.__extends||function(){var a=function(b,c){a=Object.setPrototypeOf||{__proto__:[]}instanceof Array&&function(a,b){a.__proto__=b}||function(a,b){for(var c in b)b.hasOwnProperty(c)&&
(a[c]=b[c])};return a(b,c)};return function(b,c){function d(){this.constructor=b}a(b,c);b.prototype=null===c?Object.create(c):(d.prototype=c.prototype,new d)}}(),r=n.defined,m=n.extend,e=n.isNumber,b=n.merge,a=n.pick,d=function(d){function f(a,b){return d.call(this,a,b)||this}g(f,d);f.prototype.init=function(a,b,d){c.prototype.init.call(this,a,b,d);this.resizeY=this.resizeX=this.offsetY=this.offsetX=0;f.calculations.init.call(this);this.addValues();this.addShapes()};f.prototype.setClipAxes=function(){this.clipXAxis=
this.chart.xAxis[this.options.typeOptions.xAxis];this.clipYAxis=this.chart.yAxis[this.options.typeOptions.yAxis]};f.prototype.pointsOptions=function(){return this.options.points};f.prototype.shapePointsOptions=function(){var a=this.options.typeOptions,b=a.xAxis;a=a.yAxis;return[{x:this.xAxisMin,y:this.yAxisMin,xAxis:b,yAxis:a},{x:this.xAxisMax,y:this.yAxisMin,xAxis:b,yAxis:a},{x:this.xAxisMax,y:this.yAxisMax,xAxis:b,yAxis:a},{x:this.xAxisMin,y:this.yAxisMax,xAxis:b,yAxis:a}]};f.prototype.addControlPoints=
function(){var a=this.chart.inverted,b=this.options.controlPointOptions,c=this.options.typeOptions.selectType;r(this.userOptions.controlPointOptions&&this.userOptions.controlPointOptions.style.cursor)||("x"===c?b.style.cursor=a?"ns-resize":"ew-resize":"y"===c&&(b.style.cursor=a?"ew-resize":"ns-resize"));a=new l(this.chart,this,this.options.controlPointOptions,0);this.controlPoints.push(a);"xy"!==c&&(a=new l(this.chart,this,this.options.controlPointOptions,1),this.controlPoints.push(a))};f.prototype.addValues=
function(b){var c=this.options.typeOptions,d=c.label.formatter;f.calculations.recalculate.call(this,b);c.label.enabled&&(0<this.labels.length?this.labels[0].text=d&&d.call(this)||f.calculations.defaultFormatter.call(this):this.initLabel(m({shape:"rect",backgroundColor:"none",color:"black",borderWidth:0,dashStyle:"Dash",overflow:"allow",align:"left",y:0,x:0,verticalAlign:"top",crop:!0,xAxis:0,yAxis:0,point:function(b){var d=b.annotation;b=b.options;return{x:d.xAxisMin,y:d.yAxisMin,xAxis:a(c.xAxis,
b.xAxis),yAxis:a(c.yAxis,b.yAxis)}},text:d&&d.call(this)||f.calculations.defaultFormatter.call(this)},c.label),void 0))};f.prototype.addShapes=function(){this.addCrosshairs();this.addBackground()};f.prototype.addBackground=function(){"undefined"!==typeof this.shapePointsOptions()[0].x&&this.initShape(m({type:"path",points:this.shapePointsOptions()},this.options.typeOptions.background),2)};f.prototype.addCrosshairs=function(){var a=this.chart,c=this.options.typeOptions,d=this.options.typeOptions.point,
e=a.xAxis[c.xAxis],f=a.yAxis[c.yAxis],g=a.inverted;a=e.toPixels(this.xAxisMin);e=e.toPixels(this.xAxisMax);var h=f.toPixels(this.yAxisMin),k=f.toPixels(this.yAxisMax),q={point:d,type:"path"};d=[];f=[];g&&(g=a,a=h,h=g,g=e,e=k,k=g);c.crosshairX.enabled&&(d=[["M",a,h+(k-h)/2],["L",e,h+(k-h)/2]]);c.crosshairY.enabled&&(f=[["M",a+(e-a)/2,h],["L",a+(e-a)/2,k]]);0<this.shapes.length?(this.shapes[0].options.d=d,this.shapes[1].options.d=f):(a=b(q,c.crosshairX),c=b(q,c.crosshairY),this.initShape(m({d:d},a),
0),this.initShape(m({d:f},c),1))};f.prototype.onDrag=function(a){var b=this.mouseMoveToTranslation(a),c=this.options.typeOptions.selectType;a="y"===c?0:b.x;b="x"===c?0:b.y;this.translate(a,b);this.offsetX+=a;this.offsetY+=b;this.redraw(!1,!1,!0)};f.prototype.resize=function(a,b,c,d){var e=this.shapes[2];"x"===d?0===c?(e.translatePoint(a,0,0),e.translatePoint(a,b,3)):(e.translatePoint(a,0,1),e.translatePoint(a,b,2)):"y"===d?0===c?(e.translatePoint(0,b,0),e.translatePoint(0,b,1)):(e.translatePoint(0,
b,2),e.translatePoint(0,b,3)):(e.translatePoint(a,0,1),e.translatePoint(a,b,2),e.translatePoint(0,b,3));f.calculations.updateStartPoints.call(this,!1,!0,c,a,b);this.options.typeOptions.background.height=Math.abs(this.startYMax-this.startYMin);this.options.typeOptions.background.width=Math.abs(this.startXMax-this.startXMin)};f.prototype.redraw=function(a,b,c){this.linkPoints();this.graphic||this.render();c&&f.calculations.updateStartPoints.call(this,!0,!1);this.clipRect&&this.clipRect.animate(this.getClipBox());
this.addValues(b);this.addCrosshairs();this.redrawItems(this.shapes,a);this.redrawItems(this.labels,a);this.controlPoints.forEach(function(a){a.redraw()})};f.prototype.translate=function(a,b){this.shapes.forEach(function(c){c.translate(a,b)});this.options.typeOptions.point.x=this.startXMin;this.options.typeOptions.point.y=this.startYMin};f.calculations={init:function(){var a=this.options.typeOptions,b=this.chart,c=f.calculations.getPointPos,d=b.inverted,g=b.xAxis[a.xAxis];b=b.yAxis[a.yAxis];var h=
a.background,k=d?h.height:h.width;h=d?h.width:h.height;var l=a.selectType,q=d?g.left:b.top;d=d?b.top:g.left;this.startXMin=a.point.x;this.startYMin=a.point.y;e(k)?this.startXMax=this.startXMin+k:this.startXMax=c(g,this.startXMin,parseFloat(k));e(h)?this.startYMax=this.startYMin-h:this.startYMax=c(b,this.startYMin,parseFloat(h));"x"===l?(this.startYMin=b.toValue(q),this.startYMax=b.toValue(q+b.len)):"y"===l&&(this.startXMin=g.toValue(d),this.startXMax=g.toValue(d+g.len))},recalculate:function(a){var b=
f.calculations,c=this.options.typeOptions,d=this.chart.xAxis[c.xAxis];c=this.chart.yAxis[c.yAxis];var e=f.calculations.getPointPos,g=this.offsetX,h=this.offsetY;this.xAxisMin=e(d,this.startXMin,g);this.xAxisMax=e(d,this.startXMax,g);this.yAxisMin=e(c,this.startYMin,h);this.yAxisMax=e(c,this.startYMax,h);this.min=b.min.call(this);this.max=b.max.call(this);this.average=b.average.call(this);this.bins=b.bins.call(this);a&&this.resize(0,0)},getPointPos:function(a,b,c){return a.toValue(a.toPixels(b)+c)},
updateStartPoints:function(a,b,c,d,e){var g=this.options.typeOptions,h=g.selectType,k=this.chart.xAxis[g.xAxis];g=this.chart.yAxis[g.yAxis];var q=f.calculations.getPointPos,w=this.startXMin,l=this.startXMax,u=this.startYMin,C=this.startYMax,G=this.offsetX,F=this.offsetY;b&&("x"===h?0===c?this.startXMin=q(k,w,d):this.startXMax=q(k,l,d):"y"===h?0===c?this.startYMin=q(g,u,e):this.startYMax=q(g,C,e):(this.startXMax=q(k,l,d),this.startYMax=q(g,C,e)));a&&(this.startXMin=q(k,w,G),this.startXMax=q(k,l,G),
this.startYMin=q(g,u,F),this.startYMax=q(g,C,F),this.offsetY=this.offsetX=0)},defaultFormatter:function(){return"Min: "+this.min+"<br>Max: "+this.max+"<br>Average: "+this.average+"<br>Bins: "+this.bins},getExtremes:function(a,b,c,d){return{xAxisMin:Math.min(b,a),xAxisMax:Math.max(b,a),yAxisMin:Math.min(d,c),yAxisMax:Math.max(d,c)}},min:function(){var a=Infinity,b=this.chart.series,c=f.calculations.getExtremes(this.xAxisMin,this.xAxisMax,this.yAxisMin,this.yAxisMax),d=!1;b.forEach(function(b){b.visible&&
"highcharts-navigator-series"!==b.options.id&&b.points.forEach(function(b){!b.isNull&&b.y<a&&b.x>c.xAxisMin&&b.x<=c.xAxisMax&&b.y>c.yAxisMin&&b.y<=c.yAxisMax&&(a=b.y,d=!0)})});d||(a="");return a},max:function(){var a=-Infinity,b=this.chart.series,c=f.calculations.getExtremes(this.xAxisMin,this.xAxisMax,this.yAxisMin,this.yAxisMax),d=!1;b.forEach(function(b){b.visible&&"highcharts-navigator-series"!==b.options.id&&b.points.forEach(function(b){!b.isNull&&b.y>a&&b.x>c.xAxisMin&&b.x<=c.xAxisMax&&b.y>
c.yAxisMin&&b.y<=c.yAxisMax&&(a=b.y,d=!0)})});d||(a="");return a},average:function(){var a="";""!==this.max&&""!==this.min&&(a=(this.max+this.min)/2);return a},bins:function(){var a=0,b=this.chart.series,c=f.calculations.getExtremes(this.xAxisMin,this.xAxisMax,this.yAxisMin,this.yAxisMax),d=!1;b.forEach(function(b){b.visible&&"highcharts-navigator-series"!==b.options.id&&b.points.forEach(function(b){!b.isNull&&b.x>c.xAxisMin&&b.x<=c.xAxisMax&&b.y>c.yAxisMin&&b.y<=c.yAxisMax&&(a++,d=!0)})});d||(a=
"");return a}};return f}(c);d.prototype.defaultOptions=b(c.prototype.defaultOptions,{typeOptions:{selectType:"xy",xAxis:0,yAxis:0,background:{fill:"rgba(130, 170, 255, 0.4)",strokeWidth:0,stroke:void 0},crosshairX:{enabled:!0,zIndex:6,dashStyle:"Dash",markerEnd:"arrow"},crosshairY:{enabled:!0,zIndex:6,dashStyle:"Dash",markerEnd:"arrow"},label:{enabled:!0,style:{fontSize:"11px",color:"#666666"},formatter:void 0}},controlPointOptions:{positioner:function(a){var b=this.index,c=a.chart,e=a.options,g=
e.typeOptions,h=g.selectType;e=e.controlPointOptions;var k=c.inverted,l=c.xAxis[g.xAxis];c=c.yAxis[g.yAxis];g=a.xAxisMax;var m=a.yAxisMax,n=d.calculations.getExtremes(a.xAxisMin,a.xAxisMax,a.yAxisMin,a.yAxisMax);"x"===h&&(m=(n.yAxisMax-n.yAxisMin)/2,0===b&&(g=a.xAxisMin));"y"===h&&(g=n.xAxisMin+(n.xAxisMax-n.xAxisMin)/2,0===b&&(m=a.yAxisMin));k?(a=c.toPixels(m),b=l.toPixels(g)):(a=l.toPixels(g),b=c.toPixels(m));return{x:a-e.width/2,y:b-e.height/2}},events:{drag:function(a,b){var c=this.mouseMoveToTranslation(a);
a=b.options.typeOptions.selectType;var d="y"===a?0:c.x;c="x"===a?0:c.y;b.resize(d,c,this.index,a);b.resizeX+=d;b.resizeY+=c;b.redraw(!1,!0)}}}});return c.types.measure=d});v(c,"Core/Chart/ChartNavigationComposition.js",[],function(){var c;(function(c){c.compose=function(c){c.navigation||(c.navigation=new g(c));return c};var g=function(){function c(c){this.updates=[];this.chart=c}c.prototype.addUpdate=function(c){this.chart.navigation.updates.push(c)};c.prototype.update=function(c,g){var e=this;this.updates.forEach(function(b){b.call(e.chart,
c,g)})};return c}();c.Additions=g})(c||(c={}));return c});v(c,"Extensions/Annotations/NavigationBindings.js",[c["Extensions/Annotations/Annotations.js"],c["Core/Chart/Chart.js"],c["Core/Chart/ChartNavigationComposition.js"],c["Core/FormatUtilities.js"],c["Core/Globals.js"],c["Core/DefaultOptions.js"],c["Core/Utilities.js"]],function(c,l,n,k,r,m,e){function b(a,b){var c=w.Element.prototype,d=c.matches||c.msMatchesSelector||c.webkitMatchesSelector,e=null;if(c.closest)e=c.closest.call(a,b);else{do{if(d.call(a,
b))return a;a=a.parentElement||a.parentNode}while(null!==a&&1===a.nodeType)}return e}function a(a){var b=a.prototype.defaultOptions.events&&a.prototype.defaultOptions.events.click;D(!0,a.prototype.defaultOptions.events,{click:function(a){var c=this,d=c.chart.navigationBindings,e=d.activeAnnotation;b&&b.call(c,a);e!==c?(d.deselectAnnotation(),d.activeAnnotation=c,c.setControlPointsVisibility(!0),p(d,"showPopup",{annotation:c,formType:"annotation-toolbar",options:d.annotationToFields(c),onSubmit:function(a){var b=
{};"remove"===a.actionType?(d.activeAnnotation=!1,d.chart.removeAnnotation(c)):(d.fieldsToOptions(a.fields,b),d.deselectAnnotation(),a=b.typeOptions,"measure"===c.options.type&&(a.crosshairY.enabled=0!==a.crosshairY.strokeWidth,a.crosshairX.enabled=0!==a.crosshairX.strokeWidth),c.update(b))}})):p(d,"closePopup");a.activeAnnotation=!0}})}var d=k.format;k=m.setOptions;var g=e.addEvent,f=e.attr,p=e.fireEvent,t=e.isArray,v=e.isFunction,A=e.isNumber,E=e.isObject,D=e.merge,z=e.objectEach,y=e.pick,q=r.doc,
w=r.win,B=function(){function a(a,b){this.selectedButton=this.boundClassNames=void 0;this.chart=a;this.options=b;this.eventsToUnbind=[];this.container=q.getElementsByClassName(this.options.bindingsClassName||"")}a.prototype.initEvents=function(){var a=this,b=a.chart,c=a.container,d=a.options;a.boundClassNames={};z(d.bindings||{},function(b){a.boundClassNames[b.className]=b});[].forEach.call(c,function(b){a.eventsToUnbind.push(g(b,"click",function(c){var d=a.getButtonEvents(b,c);d&&-1===d.button.className.indexOf("highcharts-disabled-btn")&&
a.bindingsButtonClick(d.button,d.events,c)}))});z(d.events||{},function(b,c){v(b)&&a.eventsToUnbind.push(g(a,c,b,{passive:!1}))});a.eventsToUnbind.push(g(b.container,"click",function(c){!b.cancelClick&&b.isInsidePlot(c.chartX-b.plotLeft,c.chartY-b.plotTop,{visiblePlotOnly:!0})&&a.bindingsChartClick(this,c)}));a.eventsToUnbind.push(g(b.container,r.isTouchDevice?"touchmove":"mousemove",function(b){a.bindingsContainerMouseMove(this,b)},r.isTouchDevice?{passive:!1}:void 0))};a.prototype.initUpdate=function(){var a=
this;n.compose(this.chart).navigation.addUpdate(function(b){a.update(b)})};a.prototype.bindingsButtonClick=function(a,b,c){var d=this.chart,e=d.renderer.boxWrapper,f=!0;this.selectedButtonElement&&(this.selectedButtonElement.classList===a.classList&&(f=!1),p(this,"deselectButton",{button:this.selectedButtonElement}),this.nextEvent&&(this.currentUserDetails&&"annotations"===this.currentUserDetails.coll&&d.removeAnnotation(this.currentUserDetails),this.mouseMoveEvent=this.nextEvent=!1));f?(this.selectedButton=
b,this.selectedButtonElement=a,p(this,"selectButton",{button:a}),b.init&&b.init.call(this,a,c),(b.start||b.steps)&&d.renderer.boxWrapper.addClass("highcharts-draw-mode")):(d.stockTools&&d.stockTools.toggleButtonAciveClass(a),e.removeClass("highcharts-draw-mode"),this.mouseMoveEvent=this.nextEvent=!1,this.selectedButton=null)};a.prototype.bindingsChartClick=function(a,c){a=this.chart;var d=this.activeAnnotation,e=this.selectedButton;a=a.renderer.boxWrapper;d&&(d.cancelClick||c.activeAnnotation||!c.target.parentNode||
b(c.target,".highcharts-popup")?d.cancelClick&&setTimeout(function(){d.cancelClick=!1},0):p(this,"closePopup"));e&&e.start&&(this.nextEvent?(this.nextEvent(c,this.currentUserDetails),this.steps&&(this.stepIndex++,e.steps[this.stepIndex]?this.mouseMoveEvent=this.nextEvent=e.steps[this.stepIndex]:(p(this,"deselectButton",{button:this.selectedButtonElement}),a.removeClass("highcharts-draw-mode"),e.end&&e.end.call(this,c,this.currentUserDetails),this.mouseMoveEvent=this.nextEvent=!1,this.selectedButton=
null))):(this.currentUserDetails=e.start.call(this,c))&&e.steps?(this.stepIndex=0,this.steps=!0,this.mouseMoveEvent=this.nextEvent=e.steps[this.stepIndex]):(p(this,"deselectButton",{button:this.selectedButtonElement}),a.removeClass("highcharts-draw-mode"),this.steps=!1,this.selectedButton=null,e.end&&e.end.call(this,c,this.currentUserDetails)))};a.prototype.bindingsContainerMouseMove=function(a,b){this.mouseMoveEvent&&this.mouseMoveEvent(b,this.currentUserDetails)};a.prototype.fieldsToOptions=function(a,
b){z(a,function(a,c){var d=parseFloat(a),e=c.split("."),f=b,g=e.length-1;!A(d)||a.match(/px/g)||c.match(/format/g)||(a=d);""!==a&&"undefined"!==a&&e.forEach(function(b,c){var d=y(e[c+1],"");g===c?f[b]=a:(f[b]||(f[b]=d.match(/\d/g)?[]:{}),f=f[b])})});return b};a.prototype.deselectAnnotation=function(){this.activeAnnotation&&(this.activeAnnotation.setControlPointsVisibility(!1),this.activeAnnotation=!1)};a.prototype.annotationToFields=function(b){function c(a,e,f,q){if(f&&a&&-1===C.indexOf(e)&&(0<=
(f.indexOf&&f.indexOf(e))||f[e]||!0===f))if(t(a))q[e]=[],a.forEach(function(a,b){E(a)?(q[e][b]={},z(a,function(a,d){c(a,d,g[e],q[e][b])})):c(a,0,g[e],q[e])});else if(E(a)){var w={};t(q)?(q.push(w),w[e]={},w=w[e]):q[e]=w;z(a,function(a,b){c(a,b,0===e?f:g[e],w)})}else"format"===e?q[e]=[d(a,b.labels[0].points[0]).toString(),"text"]:t(q)?q.push([a,u(a)]):q[e]=[a,u(a)]}var e=b.options,f=a.annotationsEditable,g=f.nestedOptions,u=this.utils.getFieldType,q=y(e.type,e.shapes&&e.shapes[0]&&e.shapes[0].type,
e.labels&&e.labels[0]&&e.labels[0].itemType,"label"),C=a.annotationsNonEditable[e.langKey]||[],w={langKey:e.langKey,type:q};z(e,function(a,b){"typeOptions"===b?(w[b]={},z(e[b],function(a,d){c(a,d,g,w[b],!0)})):c(a,b,f[q],w)});return w};a.prototype.getClickedClassNames=function(a,b){var c=b.target;b=[];for(var d;c&&((d=f(c,"class"))&&(b=b.concat(d.split(" ").map(function(a){return[a,c]}))),c=c.parentNode,c!==a););return b};a.prototype.getButtonEvents=function(a,b){var c=this,d;this.getClickedClassNames(a,
b).forEach(function(a){c.boundClassNames[a[0]]&&!d&&(d={events:c.boundClassNames[a[0]],button:a[1]})});return d};a.prototype.update=function(a){this.options=D(!0,this.options,a);this.removeEvents();this.initEvents()};a.prototype.removeEvents=function(){this.eventsToUnbind.forEach(function(a){a()})};a.prototype.destroy=function(){this.removeEvents()};a.annotationsEditable={nestedOptions:{labelOptions:["style","format","backgroundColor"],labels:["style"],label:["style"],style:["fontSize","color"],background:["fill",
"strokeWidth","stroke"],innerBackground:["fill","strokeWidth","stroke"],outerBackground:["fill","strokeWidth","stroke"],shapeOptions:["fill","strokeWidth","stroke"],shapes:["fill","strokeWidth","stroke"],line:["strokeWidth","stroke"],backgroundColors:[!0],connector:["fill","strokeWidth","stroke"],crosshairX:["strokeWidth","stroke"],crosshairY:["strokeWidth","stroke"]},circle:["shapes"],ellipse:["shapes"],verticalLine:[],label:["labelOptions"],measure:["background","crosshairY","crosshairX"],fibonacci:[],
tunnel:["background","line","height"],pitchfork:["innerBackground","outerBackground"],rect:["shapes"],crookedLine:[],basicAnnotation:["shapes","labelOptions"]};a.annotationsNonEditable={rectangle:["crosshairX","crosshairY","labelOptions"],ellipse:["labelOptions"],circle:["labelOptions"]};return a}();B.prototype.utils={getFieldType:function(a){return{string:"text",number:"number","boolean":"checkbox"}[typeof a]},updateRectSize:function(a,b){var c=b.chart,d=b.options.typeOptions,e=A(d.xAxis)&&c.xAxis[d.xAxis],
f=A(d.yAxis)&&c.yAxis[d.yAxis];e&&f&&(e=e.toValue(a[e.horiz?"chartX":"chartY"]),a=f.toValue(a[f.horiz?"chartX":"chartY"]),f=e-d.point.x,d=d.point.y-a,b.update({typeOptions:{background:{width:c.inverted?d:f,height:c.inverted?f:d}}}))},getAssignedAxis:function(a){return a.filter(function(a){var b=a.axis.getExtremes(),c=b.min;b=b.max;var d=y(a.axis.minPointOffset,0);return A(c)&&A(b)&&a.value>=c-d&&a.value<=b+d&&!a.axis.options.isInternal})[0]}};l.prototype.initNavigationBindings=function(){var a=this.options;
a&&a.navigation&&a.navigation.bindings&&(this.navigationBindings=new B(this,a.navigation),this.navigationBindings.initEvents(),this.navigationBindings.initUpdate())};g(l,"load",function(){this.initNavigationBindings()});g(l,"destroy",function(){this.navigationBindings&&this.navigationBindings.destroy()});g(B,"deselectButton",function(){this.selectedButtonElement=null});g(c,"remove",function(){this.chart.navigationBindings&&this.chart.navigationBindings.deselectAnnotation()});r.Annotation&&(a(c),z(c.types,
function(b){a(b)}));k({lang:{navigation:{popup:{simpleShapes:"Simple shapes",lines:"Lines",circle:"Circle",ellipse:"Ellipse",rectangle:"Rectangle",label:"Label",shapeOptions:"Shape options",typeOptions:"Details",fill:"Fill",format:"Text",strokeWidth:"Line width",stroke:"Line color",title:"Title",name:"Name",labelOptions:"Label options",labels:"Labels",backgroundColor:"Background color",backgroundColors:"Background colors",borderColor:"Border color",borderRadius:"Border radius",borderWidth:"Border width",
style:"Style",padding:"Padding",fontSize:"Font size",color:"Color",height:"Height",shapes:"Shape options"}}},navigation:{bindingsClassName:"highcharts-bindings-container",bindings:{circleAnnotation:{className:"highcharts-circle-annotation",start:function(a){var b=this.chart.pointer.getCoordinates(a);a=this.utils.getAssignedAxis(b.xAxis);b=this.utils.getAssignedAxis(b.yAxis);var c=this.chart.options.navigation;if(a&&b)return this.chart.addAnnotation(D({langKey:"circle",type:"basicAnnotation",shapes:[{type:"circle",
point:{x:a.value,y:b.value,xAxis:a.axis.options.index,yAxis:b.axis.options.index},r:5}]},c.annotationsOptions,c.bindings.circleAnnotation.annotationsOptions))},steps:[function(a,b){var c=b.options.shapes[0].point;if(A(c.xAxis)&&A(c.yAxis)){var d=this.chart.inverted;var e=this.chart.xAxis[c.xAxis].toPixels(c.x);c=this.chart.yAxis[c.yAxis].toPixels(c.y);d=Math.max(Math.sqrt(Math.pow(d?c-a.chartX:e-a.chartX,2)+Math.pow(d?e-a.chartY:c-a.chartY,2)),5)}b.update({shapes:[{r:d}]})}]},ellipseAnnotation:{className:"highcharts-ellipse-annotation",
start:function(a){var b=this.chart.pointer.getCoordinates(a);a=this.utils.getAssignedAxis(b.xAxis);b=this.utils.getAssignedAxis(b.yAxis);var c=this.chart.options.navigation;if(a&&b)return this.chart.addAnnotation(D({langKey:"ellipse",type:"basicAnnotation",shapes:[{type:"ellipse",xAxis:a.axis.options.index,yAxis:b.axis.options.index,points:[{x:a.value,y:b.value},{x:a.value,y:b.value}],ry:1}]},c.annotationsOptions,c.bindings.ellipseAnnotation.annotationOptions))},steps:[function(a,b){b=b.shapes[0];
var c=b.getAbsolutePosition(b.points[1]);b.translatePoint(a.chartX-c.x,a.chartY-c.y,1);b.redraw(!1)},function(a,b){b=b.shapes[0];var c=b.getAbsolutePosition(b.points[0]),d=b.getAbsolutePosition(b.points[1]);a=b.getDistanceFromLine(c,d,a.chartX,a.chartY);c=b.getYAxis();a=Math.abs(c.toValue(0)-c.toValue(a));b.setYRadius(a);b.redraw(!1)}]},rectangleAnnotation:{className:"highcharts-rectangle-annotation",start:function(a){a=this.chart.pointer.getCoordinates(a);var b=this.utils.getAssignedAxis(a.xAxis),
c=this.utils.getAssignedAxis(a.yAxis);if(b&&c){a=b.value;var d=c.value;b=b.axis.options.index;c=c.axis.options.index;var e=this.chart.options.navigation;return this.chart.addAnnotation(D({langKey:"rectangle",type:"basicAnnotation",shapes:[{type:"path",points:[{xAxis:b,yAxis:c,x:a,y:d},{xAxis:b,yAxis:c,x:a,y:d},{xAxis:b,yAxis:c,x:a,y:d},{xAxis:b,yAxis:c,x:a,y:d},{command:"Z"}]}]},e.annotationsOptions,e.bindings.rectangleAnnotation.annotationsOptions))}},steps:[function(a,b){var c=b.options.shapes[0].points,
d=this.chart.pointer.getCoordinates(a);a=this.utils.getAssignedAxis(d.xAxis);d=this.utils.getAssignedAxis(d.yAxis);a&&d&&(a=a.value,d=d.value,c[1].x=a,c[2].x=a,c[2].y=d,c[3].y=d,b.update({shapes:[{points:c}]}))}]},labelAnnotation:{className:"highcharts-label-annotation",start:function(a){var b=this.chart.pointer.getCoordinates(a);a=this.utils.getAssignedAxis(b.xAxis);b=this.utils.getAssignedAxis(b.yAxis);var c=this.chart.options.navigation;if(a&&b)return this.chart.addAnnotation(D({langKey:"label",
type:"basicAnnotation",labelOptions:{format:"{y:.2f}"},labels:[{point:{xAxis:a.axis.options.index,yAxis:b.axis.options.index,x:a.value,y:b.value},overflow:"none",crop:!0}]},c.annotationsOptions,c.bindings.labelAnnotation.annotationsOptions))}}},events:{},annotationsOptions:{animation:{defer:0}}}});g(l,"render",function(){var a=this,b=a.navigationBindings;if(a&&b){var c=!1;a.series.forEach(function(a){!a.options.isInternal&&a.visible&&(c=!0)});z(b.boundClassNames,function(b,d){if(a.navigationBindings&&
a.navigationBindings.container&&a.navigationBindings.container[0]&&(d=a.navigationBindings.container[0].querySelectorAll("."+d)))for(var e=0;e<d.length;e++){var f=d[e];"normal"===b.noDataState?-1!==f.className.indexOf("highcharts-disabled-btn")&&f.classList.remove("highcharts-disabled-btn"):c?-1!==f.className.indexOf("highcharts-disabled-btn")&&f.classList.remove("highcharts-disabled-btn"):-1===f.className.indexOf("highcharts-disabled-btn")&&(f.className+=" highcharts-disabled-btn")}})}});g(B,"closePopup",
function(){this.deselectAnnotation()});return B});v(c,"Extensions/Annotations/Popup.js",[c["Core/Globals.js"],c["Extensions/Annotations/NavigationBindings.js"],c["Core/DefaultOptions.js"],c["Core/Pointer.js"],c["Core/Utilities.js"]],function(c,l,n,k,r){var g=c.doc,e=c.isFirefox,b=n.getOptions,a=r.addEvent,d=r.createElement,h=r.defined,f=r.fireEvent,p=r.isArray,t=r.isObject,v=r.objectEach,A=r.pick,E=r.stableSort;n=r.wrap;var D=/\d/g,z;(function(a){a[a["params.algorithm"]=0]="params.algorithm";a[a["params.average"]=
1]="params.average"})(z||(z={}));var y={"algorithm-pivotpoints":["standard","fibonacci","camarilla"],"average-disparityindex":["sma","ema","dema","tema","wma"]};n(k.prototype,"onContainerMouseDown",function(a,b){this.inClass(b.target,"highcharts-popup")||a.apply(this,Array.prototype.slice.call(arguments,1))});c.Popup=function(a,b,c){this.init(a,b,c)};c.Popup.prototype={init:function(b,e,f){this.chart=f;this.container=d("div",{className:"highcharts-popup highcharts-no-tooltip"},void 0,b);a(this.container,
"mousedown",function(){var b=f&&f.navigationBindings&&f.navigationBindings.activeAnnotation;if(b){b.cancelClick=!0;var d=a(c.doc,"click",function(){setTimeout(function(){b.cancelClick=!1},0);d()})}});this.lang=this.getLangpack();this.iconsURL=e;this.addCloseBtn()},addCloseBtn:function(){var b=this,c=this.iconsURL;var e=d("div",{className:"highcharts-popup-close"},void 0,this.container);e.style["background-image"]="url("+(c.match(/png|svg|jpeg|jpg|gif/ig)?c:c+"close.svg")+")";["click","touchstart"].forEach(function(c){a(e,
c,function(){b.chart?f(b.chart.navigationBindings,"closePopup"):b.closePopup()})})},addColsContainer:function(a){var b=d("div",{className:"highcharts-popup-lhs-col"},void 0,a);a=d("div",{className:"highcharts-popup-rhs-col"},void 0,a);d("div",{className:"highcharts-popup-rhs-col-wrapper"},void 0,a);return{lhsCol:b,rhsCol:a}},addInput:function(a,b,c,e){var f=a.split(".");f=f[f.length-1];var q=this.lang;b="highcharts-"+b+"-"+A(e.htmlFor,f);b.match(D)||d("label",{htmlFor:b,className:e.labelClassName},
void 0,c).appendChild(g.createTextNode(q[f]||f));c=d("input",{name:b,value:e.value,type:e.type,className:"highcharts-popup-field"},void 0,c);c.setAttribute("highcharts-data-name",a);return c},addButton:function(b,c,e,f,h){var q=this,w=this.closePopup,k=this.getFields;var l=d("button",void 0,void 0,b);l.appendChild(g.createTextNode(c));h&&["click","touchstart"].forEach(function(b){a(l,b,function(){w.call(q);return h(k(f,e))})});return l},getFields:function(a,b){var c=Array.prototype.slice.call(a.querySelectorAll("input")),
d=Array.prototype.slice.call(a.querySelectorAll("select")),e=a.querySelectorAll("#highcharts-select-series > option:checked")[0];a=a.querySelectorAll("#highcharts-select-volume > option:checked")[0];var f={actionType:b,linkedTo:e&&e.getAttribute("value")||"",fields:{}};c.forEach(function(a){var b=a.getAttribute("highcharts-data-name");a.getAttribute("highcharts-data-series-id")?f.seriesId=a.value:b?f.fields[b]=a.value:f.type=a.value});d.forEach(function(a){var b=a.id;"highcharts-select-series"!==
b&&"highcharts-select-volume"!==b&&(b=b.split("highcharts-select-")[1],f.fields[b]=a.value)});a&&(f.fields["params.volumeSeriesID"]=a.getAttribute("value")||"");return f},showPopup:function(){var a=this.container,b=a.querySelectorAll(".highcharts-popup-close")[0];this.formType=void 0;a.innerHTML="";0<=a.className.indexOf("highcharts-annotation-toolbar")&&(a.classList.remove("highcharts-annotation-toolbar"),a.removeAttribute("style"));a.appendChild(b);a.style.display="block";a.style.height=""},closePopup:function(){A(this.popup&&
this.popup.container,this.container).style.display="none"},showForm:function(a,b,c,d){b&&(this.popup=b.navigationBindings.popup,this.showPopup(),"indicators"===a&&this.indicators.addForm.call(this,b,c,d),"annotation-toolbar"===a&&this.annotations.addToolbar.call(this,b,c,d),"annotation-edit"===a&&this.annotations.addForm.call(this,b,c,d),"flag"===a&&this.annotations.addForm.call(this,b,c,d,!0),this.formType=a,this.container.style.height=this.container.offsetHeight+"px")},getLangpack:function(){return b().lang.navigation.popup},
annotations:{addToolbar:function(a,b,c){var e=this,f=this.lang,h=this.popup.container,k=this.showForm;-1===h.className.indexOf("highcharts-annotation-toolbar")&&(h.className+=" highcharts-annotation-toolbar");a&&(h.style.top=a.plotTop+10+"px");d("span",void 0,void 0,h).appendChild(g.createTextNode(A(f[b.langKey]||b.langKey,b.shapes&&b.shapes[0].type)));var q=this.addButton(h,f.removeButton||"remove","remove",h,c);q.className+=" highcharts-annotation-remove-button";q.style["background-image"]="url("+
this.iconsURL+"destroy.svg)";q=this.addButton(h,f.editButton||"edit","edit",h,function(){k.call(e,"annotation-edit",a,b,c)});q.className+=" highcharts-annotation-edit-button";q.style["background-image"]="url("+this.iconsURL+"edit.svg)"},addForm:function(a,b,c,e){var f=this.popup.container,h=this.lang;if(a){var q=d("h2",{className:"highcharts-popup-main-title"},void 0,f);q.appendChild(g.createTextNode(h[b.langKey]||b.langKey||""));q=d("div",{className:"highcharts-popup-lhs-col highcharts-popup-lhs-full"},
void 0,f);var k=d("div",{className:"highcharts-popup-bottom-row"},void 0,f);this.annotations.addFormFields.call(this,q,a,"",b,[],!0);this.addButton(k,e?h.addButton||"add":h.saveButton||"save",e?"add":"save",f,c)}},addFormFields:function(a,b,c,f,h,k){var q=this,l=this.annotations.addFormFields,w=this.addInput,u=this.lang,m,n;b&&(v(f,function(d,e){m=""!==c?c+"."+e:e;t(d)&&(!p(d)||p(d)&&t(d[0])?(n=u[e]||e,n.match(D)||h.push([!0,n,a]),l.call(q,a,b,m,d,h,!1)):h.push([q,m,"annotation",a,d]))}),k&&(E(h,
function(a){return a[1].match(/format/g)?-1:1}),e&&h.reverse(),h.forEach(function(a){!0===a[0]?d("span",{className:"highcharts-annotation-title"},void 0,a[2]).appendChild(g.createTextNode(a[1])):(a[4]={value:a[4][0],type:a[4][1]},w.apply(a[0],a.splice(1)))})))}},indicators:{addForm:function(a,b,c){var d=this.indicators,e=this.lang;if(a){this.tabs.init.call(this,a);b=this.popup.container.querySelectorAll(".highcharts-tab-item-content");this.addColsContainer(b[0]);d.addSearchBox.call(this,a,b[0]);d.addIndicatorList.call(this,
a,b[0],"add");var f=b[0].querySelectorAll(".highcharts-popup-rhs-col")[0];this.addButton(f,e.addButton||"add","add",f,c);this.addColsContainer(b[1]);d.addIndicatorList.call(this,a,b[1],"edit");f=b[1].querySelectorAll(".highcharts-popup-rhs-col")[0];this.addButton(f,e.saveButton||"save","edit",f,c);this.addButton(f,e.removeButton||"remove","remove",f,c)}},filterSeries:function(a,b){var c=this.indicators,d=this.chart&&this.chart.options.lang,e=d&&d.navigation&&d.navigation.popup&&d.navigation.popup.indicatorAliases,
f=[],g;v(a,function(a,d){var h=a.options;if(a.params||h&&h.params)if(h=c.getNameType(a,d),d=h.indicatorFullName,h=h.indicatorType,b){var k=b.replace(/[.*+?^${}()|[\]\\]/g,"\\$&");k=new RegExp(k,"i");var l=e&&e[h]&&e[h].join(" ")||"";if(d.match(k)||l.match(k))g={indicatorFullName:d,indicatorType:h,series:a},f.push(g)}else g={indicatorFullName:d,indicatorType:h,series:a},f.push(g)});return f},filterSeriesArray:function(a){var b=[],c;a.forEach(function(a){a.is("sma")&&(c={indicatorFullName:a.name,indicatorType:a.type,
series:a},b.push(c))});return b},addIndicatorList:function(b,c,e,f){var h=this,k=h.indicators,l=h.lang,q=c.querySelectorAll(".highcharts-popup-lhs-col")[0];c=c.querySelectorAll(".highcharts-popup-rhs-col")[0];var m="edit"===e,n=this.indicators.addFormFields;e=m?b.series:b.options.plotOptions||{};if(b||!e){var w,u=[];m||p(e)?p(e)&&(u=k.filterSeriesArray.call(this,e)):u=k.filterSeries.call(this,e,f);E(u,function(a,b){a=a.indicatorFullName.toLowerCase();b=b.indicatorFullName.toLowerCase();return a<b?
-1:a>b?1:0});q.children[1]&&q.children[1].remove();var B=d("ul",{className:"highcharts-indicator-list"},void 0,q);var r=c.querySelectorAll(".highcharts-popup-rhs-col-wrapper")[0];u.forEach(function(c){var e=c.indicatorFullName,f=c.indicatorType,k=c.series;w=d("li",{className:"highcharts-indicator-list"},void 0,B);w.appendChild(g.createTextNode(e));["click","touchstart"].forEach(function(c){a(w,c,function(){var a=r.parentNode.children[1];n.call(h,b,k,f,r);a&&(a.style.display="block");m&&k.options&&
d("input",{type:"hidden",name:"highcharts-id-"+f,value:k.options.id},void 0,r).setAttribute("highcharts-data-series-id",k.options.id)})})});0<B.childNodes.length?B.childNodes[0].click():m||(r.parentNode.children[0].innerHTML=l.noFilterMatch||"",r.parentNode.children[1].style.display="none")}},addSearchBox:function(b,c){var e=this,f=c.querySelectorAll(".highcharts-popup-lhs-col")[0];c=this.lang.clearFilter;f=d("div",{className:"highcharts-input-wrapper"},void 0,f);var g=function(a){e.indicators.addIndicatorList.call(e,
b,e.container,"add",a)},h=this.addInput("searchIndicators","input",f,{value:"",type:"text",htmlFor:"search-indicators",labelClassName:"highcharts-input-search-indicators-label"}),k=d("a",{textContent:c},void 0,f);h.classList.add("highcharts-input-search-indicators");k.classList.add("clear-filter-button");a(h,"input",function(a){g(this.value);k.style.display=this.value.length?"inline-block":"none"});["click","touchstart"].forEach(function(b){a(k,b,function(){h.value="";g("");k.style.display="none"})})},
addSelection:function(a,b,c){var e=b.split(".");e=e[e.length-1];a="highcharts-"+b+"-type-"+a;var f=this.lang;d("label",{htmlFor:a},null,c).appendChild(g.createTextNode(f[e]||b));c=d("select",{name:a,className:"highcharts-popup-field",id:"highcharts-select-"+b},null,c);c.setAttribute("id","highcharts-select-"+b);return c},addSelectionOptions:function(a,b,c,e,f,k,l){"series"===b||"volume"===b?a.series.forEach(function(a){var e=a.options,f=e.name||e.params?a.name:e.id||"";"highcharts-navigator-series"!==
e.id&&e.id!==(l&&l.options&&l.options.id)&&(h(k)||"volume"!==b||"column"!==a.type||(k=e.id),d("option",{value:e.id},void 0,c).appendChild(g.createTextNode(f)))}):e&&f&&y[f+"-"+e].forEach(function(a){d("option",{value:a},void 0,c).appendChild(g.createTextNode(a))});h(k)&&(c.value=k)},getNameType:function(a,b){var d=a.options,e=c.seriesTypes;e=e[b]&&e[b].prototype.nameBase||b.toUpperCase();d&&d.type&&(b=a.options.type,e=a.name);return{indicatorFullName:e,indicatorType:b}},listAllSeries:function(a,b,
c,d,e,f){var g=this.indicators;c&&(a=g.addSelection.call(this,a,b,d),g.addSelectionOptions.call(this,c,b,a,void 0,void 0,void 0,e),h(f)&&(a.value=f))},addFormFields:function(a,b,c,e){var f=b.params||b.options.params,h=this.indicators.getNameType;e.innerHTML="";d("h3",{className:"highcharts-indicator-title"},void 0,e).appendChild(g.createTextNode(h(b,c).indicatorFullName));d("input",{type:"hidden",name:"highcharts-type-"+c,value:c},void 0,e);this.indicators.listAllSeries.call(this,c,"series",a,e,b,
b.linkedParent&&b.linkedParent.options.id);f.volumeSeriesID&&this.indicators.listAllSeries.call(this,c,"volume",a,e,b,b.linkedParent&&f.volumeSeriesID);this.indicators.addParamInputs.call(this,a,"params",f,c,e)},addParamInputs:function(a,b,c,d,e){var f=this,g=f.indicators,k=this.indicators.addParamInputs,l=this.addInput,m;a&&v(c,function(c,q){m=b+"."+q;if(h(c)&&m)if(t(c)&&(l.call(f,m,d,e,{}),k.call(f,a,m,c,d,e)),m in z){var n=g.addSelection.call(f,d,m,e);g.addSelectionOptions.call(f,a,b,n,d,q,c)}else"params.volumeSeriesID"===
m||p(c)||l.call(f,m,d,e,{value:c,type:"text"})})},getAmount:function(){var a=0;this.series.forEach(function(b){var c=b.options;(b.params||c&&c.params)&&a++});return a}},tabs:{init:function(a){var b=this.tabs,c=this.indicators.getAmount.call(a);a&&(a=b.addMenuItem.call(this,"add"),b.addMenuItem.call(this,"edit",c),b.addContentItem.call(this,"add"),b.addContentItem.call(this,"edit"),b.switchTabs.call(this,c),b.selectTab.call(this,a,0))},addMenuItem:function(a,b){var c=this.popup.container,e="highcharts-tab-item",
f=this.lang;0===b&&(e+=" highcharts-tab-disabled");b=d("span",{className:e},void 0,c);b.appendChild(g.createTextNode(f[a+"Button"]||a));b.setAttribute("highcharts-data-tab-type",a);return b},addContentItem:function(){return d("div",{className:"highcharts-tab-item-content highcharts-no-mousewheel"},void 0,this.popup.container)},switchTabs:function(b){var c=this,d;this.popup.container.querySelectorAll(".highcharts-tab-item").forEach(function(e,f){d=e.getAttribute("highcharts-data-tab-type");"edit"===
d&&0===b||["click","touchstart"].forEach(function(b){a(e,b,function(){c.tabs.deselectAll.call(c);c.tabs.selectTab.call(c,this,f)})})})},selectTab:function(a,b){var c=this.popup.container.querySelectorAll(".highcharts-tab-item-content");a.className+=" highcharts-tab-item-active";c[b].className+=" highcharts-tab-item-show"},deselectAll:function(){var a=this.popup.container,b=a.querySelectorAll(".highcharts-tab-item");a=a.querySelectorAll(".highcharts-tab-item-content");var c;for(c=0;c<b.length;c++)b[c].classList.remove("highcharts-tab-item-active"),
a[c].classList.remove("highcharts-tab-item-show")}}};a(l,"showPopup",function(a){this.popup||(this.popup=new c.Popup(this.chart.container,this.chart.options.navigation.iconsURL||this.chart.options.stockTools&&this.chart.options.stockTools.gui.iconsURL||"https://code.highcharts.com/9.3.1/gfx/stock-icons/",this.chart));this.popup.showForm(a.formType,this.chart,a.options,a.onSubmit)});a(l,"closePopup",function(){this.popup&&this.popup.closePopup()});return c.Popup});v(c,"masters/modules/annotations-advanced.src.js",
[],function(){})});
//# sourceMappingURL=annotations-advanced.js.map
```
