## Supplementary material for "Exploring natural odour landscapes: A case study with implications for human-biting insects": Figure S4: annotations.js

```
/*
 Highcharts JS v9.3.1 (2021-11-05)

 Annotations module

 (c) 2009-2021 Torstein Honsi

 License: www.highcharts.com/license
*/
'use strict';(function(a){"object"===typeof module&&module.exports?(a["default"]=a,module.exports=a):"function"===typeof define&&define.amd?define("highcharts/modules/annotations",["highcharts"],function(r){a(r);a.Highcharts=r;return a}):a("undefined"!==typeof Highcharts?Highcharts:void 0)})(function(a){function r(c,a,n,x){c.hasOwnProperty(a)||(c[a]=x.apply(null,n))}a=a?a._modules:{};r(a,"Extensions/Annotations/Mixins/EventEmitterMixin.js",[a["Core/Globals.js"],a["Core/Utilities.js"]],function(c,
a){var h=a.addEvent,x=a.fireEvent,m=a.objectEach,g=a.pick,b=a.removeEvent;return{addEvents:function(){var b=this,d=function(d){h(d,c.isTouchDevice?"touchstart":"mousedown",function(d){b.onMouseDown(d)},{passive:!1})};d(this.graphic.element);(b.labels||[]).forEach(function(b){b.options.useHTML&&b.graphic.text&&d(b.graphic.text.element)});m(b.options.events,function(d,e){var c=function(c){"click"===e&&b.cancelClick||d.call(b,b.chart.pointer.normalize(c),b.target)};if(-1===(b.nonDOMEvents||[]).indexOf(e))b.graphic.on(e,
c);else h(b,e,c,{passive:!1})});if(b.options.draggable&&(h(b,"drag",b.onDrag),!b.graphic.renderer.styledMode)){var e={cursor:{x:"ew-resize",y:"ns-resize",xy:"move"}[b.options.draggable]};b.graphic.css(e);(b.labels||[]).forEach(function(d){d.options.useHTML&&d.graphic.text&&d.graphic.text.css(e)})}b.isUpdating||x(b,"add")},removeDocEvents:function(){this.removeDrag&&(this.removeDrag=this.removeDrag());this.removeMouseUp&&(this.removeMouseUp=this.removeMouseUp())},onMouseDown:function(b){var d=this,
e=d.chart.pointer;b.preventDefault&&b.preventDefault();if(2!==b.button){b=e.normalize(b);var B=b.chartX;var f=b.chartY;d.cancelClick=!1;d.chart.hasDraggedAnnotation=!0;d.removeDrag=h(c.doc,c.isTouchDevice?"touchmove":"mousemove",function(b){d.hasDragged=!0;b=e.normalize(b);b.prevChartX=B;b.prevChartY=f;x(d,"drag",b);B=b.chartX;f=b.chartY},c.isTouchDevice?{passive:!1}:void 0);d.removeMouseUp=h(c.doc,c.isTouchDevice?"touchend":"mouseup",function(b){var e=g(d.target&&d.target.annotation,d.target);e&&
(e.cancelClick=d.hasDragged);d.cancelClick=d.hasDragged;d.hasDragged=!1;d.chart.hasDraggedAnnotation=!1;x(g(e,d),"afterUpdate");d.onMouseUp(b)},c.isTouchDevice?{passive:!1}:void 0)}},onMouseUp:function(b){var d=this.chart;b=this.target||this;var e=d.options.annotations;d=d.annotations.indexOf(b);this.removeDocEvents();e[d]=b.options},onDrag:function(b){if(this.chart.isInsidePlot(b.chartX-this.chart.plotLeft,b.chartY-this.chart.plotTop,{visiblePlotOnly:!0})){var d=this.mouseMoveToTranslation(b);"x"===
this.options.draggable&&(d.y=0);"y"===this.options.draggable&&(d.x=0);this.points.length?this.translate(d.x,d.y):(this.shapes.forEach(function(b){b.translate(d.x,d.y)}),this.labels.forEach(function(b){b.translate(d.x,d.y)}));this.redraw(!1)}},mouseMoveToRadians:function(b,d,e){var c=b.prevChartY-e,f=b.prevChartX-d;e=b.chartY-e;b=b.chartX-d;this.chart.inverted&&(d=f,f=c,c=d,d=b,b=e,e=d);return Math.atan2(e,b)-Math.atan2(c,f)},mouseMoveToTranslation:function(b){var d=b.chartX-b.prevChartX;b=b.chartY-
b.prevChartY;if(this.chart.inverted){var e=b;b=d;d=e}return{x:d,y:b}},mouseMoveToScale:function(b,d,e){d=(b.chartX-d||1)/(b.prevChartX-d||1);b=(b.chartY-e||1)/(b.prevChartY-e||1);this.chart.inverted&&(e=b,b=d,d=e);return{x:d,y:b}},destroy:function(){this.removeDocEvents();b(this);this.hcEvents=null}}});r(a,"Extensions/Annotations/ControlPoint.js",[a["Core/Utilities.js"],a["Extensions/Annotations/Mixins/EventEmitterMixin.js"]],function(c,a){var h=c.merge,x=c.pick;return function(){function c(c,b,B,
d){this.addEvents=a.addEvents;this.graphic=void 0;this.mouseMoveToRadians=a.mouseMoveToRadians;this.mouseMoveToScale=a.mouseMoveToScale;this.mouseMoveToTranslation=a.mouseMoveToTranslation;this.onDrag=a.onDrag;this.onMouseDown=a.onMouseDown;this.onMouseUp=a.onMouseUp;this.removeDocEvents=a.removeDocEvents;this.nonDOMEvents=["drag"];this.chart=c;this.target=b;this.options=B;this.index=x(B.index,d)}c.prototype.setVisibility=function(c){this.graphic.attr("visibility",c?"visible":"hidden");this.options.visible=
c};c.prototype.render=function(){var c=this.chart,b=this.options;this.graphic=c.renderer.symbol(b.symbol,0,0,b.width,b.height).add(c.controlPointsGroup).css(b.style);this.setVisibility(b.visible);this.addEvents()};c.prototype.redraw=function(c){this.graphic[c?"animate":"attr"](this.options.positioner.call(this,this.target))};c.prototype.destroy=function(){a.destroy.call(this);this.graphic&&(this.graphic=this.graphic.destroy());this.options=this.target=this.chart=null};c.prototype.update=function(c){var b=
this.chart,a=this.target,d=this.index;c=h(!0,this.options,c);this.destroy();this.constructor(b,a,c,d);this.render(b.controlPointsGroup);this.redraw()};return c}()});r(a,"Extensions/Annotations/MockPoint.js",[a["Core/Series/Series.js"],a["Core/Utilities.js"],a["Core/Axis/Axis.js"]],function(c,a,n){var h=a.defined,m=a.fireEvent;return function(){function a(b,a,d){this.y=this.x=this.ttBelow=this.plotY=this.plotX=this.negative=this.isInside=void 0;this.mock=!0;this.series={visible:!0,chart:b,getPlotBox:c.prototype.getPlotBox};
this.target=a||null;this.options=d;this.applyOptions(this.getOptions())}a.fromPoint=function(b){return new a(b.series.chart,null,{x:b.x,y:b.y,xAxis:b.series.xAxis,yAxis:b.series.yAxis})};a.pointToPixels=function(b,c){var d=b.series,e=d.chart,a=b.plotX,f=b.plotY;e.inverted&&(b.mock?(a=b.plotY,f=b.plotX):(a=e.plotWidth-b.plotY,f=e.plotHeight-b.plotX));d&&!c&&(b=d.getPlotBox(),a+=b.translateX,f+=b.translateY);return{x:a,y:f}};a.pointToOptions=function(b){return{x:b.x,y:b.y,xAxis:b.series.xAxis,yAxis:b.series.yAxis}};
a.prototype.hasDynamicOptions=function(){return"function"===typeof this.options};a.prototype.getOptions=function(){return this.hasDynamicOptions()?this.options(this.target):this.options};a.prototype.applyOptions=function(b){this.command=b.command;this.setAxis(b,"x");this.setAxis(b,"y");this.refresh()};a.prototype.setAxis=function(b,c){c+="Axis";b=b[c];var d=this.series.chart;this.series[c]=b instanceof n?b:h(b)?d[c][b]||d.get(b):null};a.prototype.toAnchor=function(){var b=[this.plotX,this.plotY,0,
0];this.series.chart.inverted&&(b[0]=this.plotY,b[1]=this.plotX);return b};a.prototype.getLabelConfig=function(){return{x:this.x,y:this.y,point:this}};a.prototype.isInsidePlot=function(){var b=this.plotX,c=this.plotY,d=this.series.xAxis,e=this.series.yAxis,a={x:b,y:c,isInsidePlot:!0};d&&(a.isInsidePlot=h(b)&&0<=b&&b<=d.len);e&&(a.isInsidePlot=a.isInsidePlot&&h(c)&&0<=c&&c<=e.len);m(this.series.chart,"afterIsInsidePlot",a);return a.isInsidePlot};a.prototype.refresh=function(){var b=this.series,c=b.xAxis;
b=b.yAxis;var d=this.getOptions();c?(this.x=d.x,this.plotX=c.toPixels(d.x,!0)):(this.x=null,this.plotX=d.x);b?(this.y=d.y,this.plotY=b.toPixels(d.y,!0)):(this.y=null,this.plotY=d.y);this.isInside=this.isInsidePlot()};a.prototype.translate=function(b,c,d,e){this.hasDynamicOptions()||(this.plotX+=d,this.plotY+=e,this.refreshOptions())};a.prototype.scale=function(b,c,d,e){if(!this.hasDynamicOptions()){var a=this.plotY*e;this.plotX=(1-d)*b+this.plotX*d;this.plotY=(1-e)*c+a;this.refreshOptions()}};a.prototype.rotate=
function(b,c,d){if(!this.hasDynamicOptions()){var e=Math.cos(d);d=Math.sin(d);var a=this.plotX,f=this.plotY;a-=b;f-=c;this.plotX=a*e-f*d+b;this.plotY=a*d+f*e+c;this.refreshOptions()}};a.prototype.refreshOptions=function(){var b=this.series,c=b.xAxis;b=b.yAxis;this.x=this.options.x=c?this.options.x=c.toValue(this.plotX,!0):this.plotX;this.y=this.options.y=b?b.toValue(this.plotY,!0):this.plotY};return a}()});r(a,"Extensions/Annotations/Mixins/ControllableMixin.js",[a["Extensions/Annotations/ControlPoint.js"],
a["Extensions/Annotations/MockPoint.js"],a["Core/Tooltip.js"],a["Core/Utilities.js"]],function(c,a,n,x){var h=x.isObject,g=x.isString,b=x.merge,B=x.splat;return{init:function(b,c,a){this.annotation=b;this.chart=b.chart;this.options=c;this.points=[];this.controlPoints=[];this.index=a;this.linkPoints();this.addControlPoints()},attr:function(){this.graphic.attr.apply(this.graphic,arguments)},getPointsOptions:function(){var b=this.options;return b.points||b.point&&B(b.point)},attrsFromOptions:function(b){var d=
this.constructor.attrsMap,c={},a,p=this.chart.styledMode;for(a in b){var u=d[a];!u||p&&-1!==["fill","stroke","stroke-width"].indexOf(u)||(c[u]=b[a])}return c},anchor:function(c){var d=c.series.getPlotBox(),a=c.series.chart,f=c.mock?c.toAnchor():n.prototype.getAnchor.call({chart:c.series.chart},c);f={x:f[0]+(this.options.x||0),y:f[1]+(this.options.y||0),height:f[2]||0,width:f[3]||0};return{relativePosition:f,absolutePosition:b(f,{x:f.x+(c.mock?d.translateX:a.plotLeft),y:f.y+(c.mock?d.translateY:a.plotTop)})}},
point:function(b,c){if(b&&b.series)return b;c&&null!==c.series||(h(b)?c=new a(this.chart,this,b):g(b)?c=this.chart.get(b)||null:"function"===typeof b&&(c=b.call(c,this),c=c.series?c:new a(this.chart,this,b)));return c},linkPoints:function(){var b=this.getPointsOptions(),c=this.points,a=b&&b.length||0,f;for(f=0;f<a;f++){var p=this.point(b[f],c[f]);if(!p){c.length=0;return}p.mock&&p.refresh();c[f]=p}return c},addControlPoints:function(){var d=this.options.controlPoints;(d||[]).forEach(function(a,v){a=
b(this.options.controlPointOptions,a);a.index||(a.index=v);d[v]=a;this.controlPoints.push(new c(this.chart,this,a))},this)},shouldBeDrawn:function(){return!!this.points.length},render:function(b){this.controlPoints.forEach(function(b){b.render()})},redraw:function(b){this.controlPoints.forEach(function(c){c.redraw(b)})},transform:function(b,c,a,f,p){if(this.chart.inverted){var d=c;c=a;a=d}this.points.forEach(function(d,e){this.transformPoint(b,c,a,f,p,e)},this)},transformPoint:function(b,c,v,f,p,
u){var d=this.points[u];d.mock||(d=this.points[u]=a.fromPoint(d));d[b](c,v,f,p)},translate:function(b,c){this.transform("translate",null,null,b,c)},translatePoint:function(b,c,a){this.transformPoint("translate",null,null,b,c,a)},translateShape:function(b,c,a){var d=this.annotation.chart,e=this.annotation.userOptions,v=d.annotations.indexOf(this.annotation);d=d.options.annotations[v];this.translatePoint(b,c,0);a&&this.translatePoint(b,c,1);d[this.collection][this.index].point=this.options.point;e[this.collection][this.index].point=
this.options.point},rotate:function(b,c,a){this.transform("rotate",b,c,a)},scale:function(b,c,a,f){this.transform("scale",b,c,a,f)},setControlPointsVisibility:function(b){this.controlPoints.forEach(function(c){c.setVisibility(b)})},destroy:function(){this.graphic&&(this.graphic=this.graphic.destroy());this.tracker&&(this.tracker=this.tracker.destroy());this.controlPoints.forEach(function(b){b.destroy()});this.options=this.controlPoints=this.points=this.chart=null;this.annotation&&(this.annotation=
null)},update:function(c){var a=this.annotation;c=b(!0,this.options,c);var d=this.graphic.parentGroup;this.destroy();this.constructor(a,c,this.index);this.render(d);this.redraw()}}});r(a,"Extensions/Annotations/Mixins/MarkerMixin.js",[a["Core/Chart/Chart.js"],a["Core/Renderer/SVG/SVGRenderer.js"],a["Core/Utilities.js"]],function(c,a,n){function h(b){return function(c){this.attr(b,"url(#"+c+")")}}var m=n.addEvent,g=n.defined,b=n.merge,B=n.uniqueKey,d={arrow:{tagName:"marker",attributes:{id:"arrow",
refY:5,refX:9,markerWidth:10,markerHeight:10},children:[{tagName:"path",attributes:{d:"M 0 0 L 10 5 L 0 10 Z","stroke-width":0}}]},"reverse-arrow":{tagName:"marker",attributes:{id:"reverse-arrow",refY:5,refX:1,markerWidth:10,markerHeight:10},children:[{tagName:"path",attributes:{d:"M 0 5 L 10 0 L 10 10 Z","stroke-width":0}}]}};a.prototype.addMarker=function(c,a){var d={attributes:{id:c}},e={stroke:a.color||"none",fill:a.color||"rgba(0, 0, 0, 0.75)"};d.children=a.children&&a.children.map(function(c){return b(e,
c)});a=b(!0,{attributes:{markerWidth:20,markerHeight:20,refX:0,refY:0,orient:"auto"}},a,d);a=this.definition(a);a.id=c;return a};a={markerEndSetter:h("marker-end"),markerStartSetter:h("marker-start"),setItemMarkers:function(c){var a=c.options,d=c.chart,e=d.options.defs,u=a.fill,h=g(u)&&"none"!==u?u:a.stroke;["markerStart","markerEnd"].forEach(function(p){var f=a[p],u;if(f){for(u in e){var g=e[u];if((f===(g.attributes&&g.attributes.id)||f===g.id)&&"marker"===g.tagName){var y=g;break}}y&&(f=c[p]=d.renderer.addMarker((a.id||
B())+"-"+f,b(y,{color:h})),c.attr(p,f.getAttribute("id")))}})}};m(c,"afterGetContainer",function(){this.options.defs=b(d,this.options.defs||{})});return a});r(a,"Extensions/Annotations/Controllables/ControllablePath.js",[a["Extensions/Annotations/Mixins/ControllableMixin.js"],a["Core/Globals.js"],a["Extensions/Annotations/Mixins/MarkerMixin.js"],a["Core/Utilities.js"]],function(c,a,n,x){var h=x.extend,g="rgba(192,192,192,"+(a.svg?.0001:.002)+")";return function(){function b(b,a,e){this.addControlPoints=
c.addControlPoints;this.anchor=c.anchor;this.attr=c.attr;this.attrsFromOptions=c.attrsFromOptions;this.destroy=c.destroy;this.getPointsOptions=c.getPointsOptions;this.init=c.init;this.linkPoints=c.linkPoints;this.point=c.point;this.rotate=c.rotate;this.scale=c.scale;this.setControlPointsVisibility=c.setControlPointsVisibility;this.setMarkers=n.setItemMarkers;this.transform=c.transform;this.transformPoint=c.transformPoint;this.translate=c.translate;this.translatePoint=c.translatePoint;this.translateShape=
c.translateShape;this.update=c.update;this.type="path";this.init(b,a,e);this.collection="shapes"}b.prototype.toD=function(){var b=this.options.d;if(b)return"function"===typeof b?b.call(this):b;b=this.points;var c=b.length,a=c,g=b[0],f=a&&this.anchor(g).absolutePosition,p=0,u=[];if(f)for(u.push(["M",f.x,f.y]);++p<c&&a;)g=b[p],a=g.command||"L",f=this.anchor(g).absolutePosition,"M"===a?u.push([a,f.x,f.y]):"L"===a?u.push([a,f.x,f.y]):"Z"===a&&u.push([a]),a=g.series.visible;return a?this.chart.renderer.crispLine(u,
this.graphic.strokeWidth()):null};b.prototype.shouldBeDrawn=function(){return c.shouldBeDrawn.call(this)||!!this.options.d};b.prototype.render=function(b){var a=this.options,e=this.attrsFromOptions(a);this.graphic=this.annotation.chart.renderer.path([["M",0,0]]).attr(e).add(b);a.className&&this.graphic.addClass(a.className);this.tracker=this.annotation.chart.renderer.path([["M",0,0]]).addClass("highcharts-tracker-line").attr({zIndex:2}).add(b);this.annotation.chart.styledMode||this.tracker.attr({"stroke-linejoin":"round",
stroke:g,fill:g,"stroke-width":this.graphic.strokeWidth()+2*a.snap});c.render.call(this);h(this.graphic,{markerStartSetter:n.markerStartSetter,markerEndSetter:n.markerEndSetter});this.setMarkers(this)};b.prototype.redraw=function(b){var a=this.toD(),e=b?"animate":"attr";a?(this.graphic[e]({d:a}),this.tracker[e]({d:a})):(this.graphic.attr({d:"M 0 -9000000000"}),this.tracker.attr({d:"M 0 -9000000000"}));this.graphic.placed=this.tracker.placed=!!a;c.redraw.call(this,b)};b.attrsMap={dashStyle:"dashstyle",
strokeWidth:"stroke-width",stroke:"stroke",fill:"fill",zIndex:"zIndex"};return b}()});r(a,"Extensions/Annotations/Controllables/ControllableRect.js",[a["Extensions/Annotations/Mixins/ControllableMixin.js"],a["Extensions/Annotations/Controllables/ControllablePath.js"],a["Core/Utilities.js"]],function(c,a,n){var h=n.merge;return function(){function m(a,b,h){this.addControlPoints=c.addControlPoints;this.anchor=c.anchor;this.attr=c.attr;this.attrsFromOptions=c.attrsFromOptions;this.destroy=c.destroy;
this.getPointsOptions=c.getPointsOptions;this.init=c.init;this.linkPoints=c.linkPoints;this.point=c.point;this.rotate=c.rotate;this.scale=c.scale;this.setControlPointsVisibility=c.setControlPointsVisibility;this.shouldBeDrawn=c.shouldBeDrawn;this.transform=c.transform;this.transformPoint=c.transformPoint;this.translatePoint=c.translatePoint;this.translateShape=c.translateShape;this.update=c.update;this.type="rect";this.translate=c.translateShape;this.init(a,b,h);this.collection="shapes"}m.prototype.render=
function(a){var b=this.attrsFromOptions(this.options);this.graphic=this.annotation.chart.renderer.rect(0,-9E9,0,0).attr(b).add(a);c.render.call(this)};m.prototype.redraw=function(a){var b=this.anchor(this.points[0]).absolutePosition;if(b)this.graphic[a?"animate":"attr"]({x:b.x,y:b.y,width:this.options.width,height:this.options.height});else this.attr({x:0,y:-9E9});this.graphic.placed=!!b;c.redraw.call(this,a)};m.attrsMap=h(a.attrsMap,{width:"width",height:"height"});return m}()});r(a,"Extensions/Annotations/Controllables/ControllableCircle.js",
[a["Extensions/Annotations/Mixins/ControllableMixin.js"],a["Extensions/Annotations/Controllables/ControllablePath.js"],a["Core/Utilities.js"]],function(c,a,n){var h=n.merge;return function(){function m(a,b,h){this.addControlPoints=c.addControlPoints;this.anchor=c.anchor;this.attr=c.attr;this.attrsFromOptions=c.attrsFromOptions;this.destroy=c.destroy;this.getPointsOptions=c.getPointsOptions;this.init=c.init;this.linkPoints=c.linkPoints;this.point=c.point;this.rotate=c.rotate;this.scale=c.scale;this.setControlPointsVisibility=
c.setControlPointsVisibility;this.shouldBeDrawn=c.shouldBeDrawn;this.transform=c.transform;this.transformPoint=c.transformPoint;this.translatePoint=c.translatePoint;this.translateShape=c.translateShape;this.update=c.update;this.type="circle";this.translate=c.translateShape;this.init(a,b,h);this.collection="shapes"}m.prototype.render=function(a){var b=this.attrsFromOptions(this.options);this.graphic=this.annotation.chart.renderer.circle(0,-9E9,0).attr(b).add(a);c.render.call(this)};m.prototype.redraw=
function(a){var b=this.anchor(this.points[0]).absolutePosition;if(b)this.graphic[a?"animate":"attr"]({x:b.x,y:b.y,r:this.options.r});else this.graphic.attr({x:0,y:-9E9});this.graphic.placed=!!b;c.redraw.call(this,a)};m.prototype.setRadius=function(c){this.options.r=c};m.attrsMap=h(a.attrsMap,{r:"r"});return m}()});r(a,"Extensions/Annotations/Controllables/ControllableEllipse.js",[a["Extensions/Annotations/Mixins/ControllableMixin.js"],a["Extensions/Annotations/Controllables/ControllablePath.js"],
a["Core/Utilities.js"]],function(c,a,n){var h=n.merge,m=n.defined;return function(){function g(b,a,d){this.addControlPoints=c.addControlPoints;this.anchor=c.anchor;this.attr=c.attr;this.attrsFromOptions=c.attrsFromOptions;this.destroy=c.destroy;this.getPointsOptions=c.getPointsOptions;this.linkPoints=c.linkPoints;this.point=c.point;this.scale=c.scale;this.setControlPointsVisibility=c.setControlPointsVisibility;this.shouldBeDrawn=c.shouldBeDrawn;this.transform=c.transform;this.translatePoint=c.translatePoint;
this.transformPoint=c.transformPoint;this.type="ellipse";this.init(b,a,d);this.collection="shapes"}g.prototype.init=function(b,a,d){m(a.yAxis)&&a.points.forEach(function(b){b.yAxis=a.yAxis});m(a.xAxis)&&a.points.forEach(function(b){b.xAxis=a.xAxis});c.init.call(this,b,a,d)};g.prototype.render=function(b){this.graphic=this.annotation.chart.renderer.createElement("ellipse").attr(this.attrsFromOptions(this.options)).add(b);c.render.call(this)};g.prototype.translate=function(b,a){c.translateShape.call(this,
b,a,!0)};g.prototype.getDistanceFromLine=function(b,c,a,e){return Math.abs((c.y-b.y)*a-(c.x-b.x)*e+c.x*b.y-c.y*b.x)/Math.sqrt((c.y-b.y)*(c.y-b.y)+(c.x-b.x)*(c.x-b.x))};g.prototype.getAttrs=function(b,c){var a=b.x,e=b.y,h=c.x,f=c.y;c=(a+h)/2;b=(e+f)/2;var p=Math.sqrt((a-h)*(a-h)/4+(e-f)*(e-f)/4);e=180*Math.atan((f-e)/(h-a))/Math.PI;c<a&&(e+=180);a=this.getRY();return{cx:c,cy:b,rx:p,ry:a,angle:e}};g.prototype.getRY=function(){var b=this.getYAxis();return m(b)?Math.abs(b.toPixels(this.options.ry)-b.toPixels(0)):
this.options.ry};g.prototype.getYAxis=function(){return this.chart.yAxis[this.options.yAxis]};g.prototype.getAbsolutePosition=function(b){return this.anchor(b).absolutePosition};g.prototype.redraw=function(b){var a=this.getAbsolutePosition(this.points[0]),d=this.getAbsolutePosition(this.points[1]);d=this.getAttrs(a,d);if(a)this.graphic[b?"animate":"attr"]({cx:d.cx,cy:d.cy,rx:d.rx,ry:d.ry,rotation:d.angle,rotationOriginX:d.cx,rotationOriginY:d.cy});else this.graphic.attr({x:0,y:-9E9});this.graphic.placed=
!!a;c.redraw.call(this,b)};g.prototype.setYRadius=function(b){this.options.ry=b;this.annotation.userOptions.shapes[0].ry=b;this.annotation.options.shapes[0].ry=b};g.attrsMap=h(a.attrsMap,{ry:"ry"});return g}()});r(a,"Extensions/Annotations/Controllables/ControllableLabel.js",[a["Extensions/Annotations/Mixins/ControllableMixin.js"],a["Core/FormatUtilities.js"],a["Extensions/Annotations/MockPoint.js"],a["Core/Renderer/SVG/SVGRenderer.js"],a["Core/Tooltip.js"],a["Core/Utilities.js"]],function(c,a,n,
x,m,g){var b=a.format;a=x.prototype.symbols;var h=g.extend,d=g.isNumber,e=g.pick;g=function(){function a(b,a,d){this.addControlPoints=c.addControlPoints;this.attr=c.attr;this.attrsFromOptions=c.attrsFromOptions;this.destroy=c.destroy;this.getPointsOptions=c.getPointsOptions;this.init=c.init;this.linkPoints=c.linkPoints;this.point=c.point;this.rotate=c.rotate;this.scale=c.scale;this.setControlPointsVisibility=c.setControlPointsVisibility;this.shouldBeDrawn=c.shouldBeDrawn;this.transform=c.transform;
this.transformPoint=c.transformPoint;this.translateShape=c.translateShape;this.update=c.update;this.init(b,a,d);this.collection="labels"}a.alignedPosition=function(b,a){var c=b.align,d=b.verticalAlign,p=(a.x||0)+(b.x||0),e=(a.y||0)+(b.y||0),f,h;"right"===c?f=1:"center"===c&&(f=2);f&&(p+=(a.width-(b.width||0))/f);"bottom"===d?h=1:"middle"===d&&(h=2);h&&(e+=(a.height-(b.height||0))/h);return{x:Math.round(p),y:Math.round(e)}};a.justifiedOptions=function(b,a,c,d){var e=c.align,f=c.verticalAlign,p=a.box?
0:a.padding||0,h=a.getBBox();a={align:e,verticalAlign:f,x:c.x,y:c.y,width:a.width,height:a.height};c=(d.x||0)-b.plotLeft;d=(d.y||0)-b.plotTop;var g=c+p;0>g&&("right"===e?a.align="left":a.x=(a.x||0)-g);g=c+h.width-p;g>b.plotWidth&&("left"===e?a.align="right":a.x=(a.x||0)+b.plotWidth-g);g=d+p;0>g&&("bottom"===f?a.verticalAlign="top":a.y=(a.y||0)-g);g=d+h.height-p;g>b.plotHeight&&("top"===f?a.verticalAlign="bottom":a.y=(a.y||0)+b.plotHeight-g);return a};a.prototype.translatePoint=function(b,a){c.translatePoint.call(this,
b,a,0)};a.prototype.translate=function(b,a){var c=this.annotation.chart,d=this.annotation.userOptions,e=c.annotations.indexOf(this.annotation);e=c.options.annotations[e];c.inverted&&(c=b,b=a,a=c);this.options.x+=b;this.options.y+=a;e[this.collection][this.index].x=this.options.x;e[this.collection][this.index].y=this.options.y;d[this.collection][this.index].x=this.options.x;d[this.collection][this.index].y=this.options.y};a.prototype.render=function(b){var d=this.options,e=this.attrsFromOptions(d),
f=d.style;this.graphic=this.annotation.chart.renderer.label("",0,-9999,d.shape,null,null,d.useHTML,null,"annotation-label").attr(e).add(b);this.annotation.chart.styledMode||("contrast"===f.color&&(f.color=this.annotation.chart.renderer.getContrast(-1<a.shapesWithoutBackground.indexOf(d.shape)?"#FFFFFF":d.backgroundColor)),this.graphic.css(d.style).shadow(d.shadow));d.className&&this.graphic.addClass(d.className);this.graphic.labelrank=d.labelrank;c.render.call(this)};a.prototype.redraw=function(a){var d=
this.options,e=this.text||d.format||d.text,f=this.graphic,h=this.points[0];f.attr({text:e?b(e,h.getLabelConfig(),this.annotation.chart):d.formatter.call(h,this)});d=this.anchor(h);(e=this.position(d))?(f.alignAttr=e,e.anchorX=d.absolutePosition.x,e.anchorY=d.absolutePosition.y,f[a?"animate":"attr"](e)):f.attr({x:0,y:-9999});f.placed=!!e;c.redraw.call(this,a)};a.prototype.anchor=function(b){var a=c.anchor.apply(this,arguments),d=this.options.x||0,e=this.options.y||0;a.absolutePosition.x-=d;a.absolutePosition.y-=
e;a.relativePosition.x-=d;a.relativePosition.y-=e;return a};a.prototype.position=function(b){var c=this.graphic,d=this.annotation.chart,f=this.points[0],g=this.options,x=b.absolutePosition,v=b.relativePosition,z=f.series.visible&&n.prototype.isInsidePlot.call(f);b=c.width;b=void 0===b?0:b;var r=c.height;r=void 0===r?0:r;if(z){if(g.distance)var w=m.prototype.getPosition.call({chart:d,distance:e(g.distance,16)},b,r,{plotX:v.x,plotY:v.y,negative:f.negative,ttBelow:f.ttBelow,h:v.height||v.width});else g.positioner?
w=g.positioner.call(this):(f={x:x.x,y:x.y,width:0,height:0},w=a.alignedPosition(h(g,{width:b,height:r}),f),"justify"===this.options.overflow&&(w=a.alignedPosition(a.justifiedOptions(d,c,g,w),f)));g.crop&&(c=w.x-d.plotLeft,g=w.y-d.plotTop,z=d.isInsidePlot(c,g)&&d.isInsidePlot(c+b,g+r))}return z?w:null};a.attrsMap={backgroundColor:"fill",borderColor:"stroke",borderWidth:"stroke-width",zIndex:"zIndex",borderRadius:"r",padding:"padding"};a.shapesWithoutBackground=["connector"];return a}();a.connector=
function(b,a,c,e,g){var f=g&&g.anchorX;g=g&&g.anchorY;var h=c/2;if(d(f)&&d(g)){var m=[["M",f,g]];var n=a-g;0>n&&(n=-e-n);n<c&&(h=f<b+c/2?n:c-n);g>a+e?m.push(["L",b+h,a+e]):g<a?m.push(["L",b+h,a]):f<b?m.push(["L",b,a+e/2]):f>b+c&&m.push(["L",b+c,a+e/2])}return m||[]};return g});r(a,"Extensions/Annotations/Controllables/ControllableImage.js",[a["Extensions/Annotations/Controllables/ControllableLabel.js"],a["Extensions/Annotations/Mixins/ControllableMixin.js"]],function(a,h){return function(){function c(a,
c,g){this.addControlPoints=h.addControlPoints;this.anchor=h.anchor;this.attr=h.attr;this.attrsFromOptions=h.attrsFromOptions;this.destroy=h.destroy;this.getPointsOptions=h.getPointsOptions;this.init=h.init;this.linkPoints=h.linkPoints;this.point=h.point;this.rotate=h.rotate;this.scale=h.scale;this.setControlPointsVisibility=h.setControlPointsVisibility;this.shouldBeDrawn=h.shouldBeDrawn;this.transform=h.transform;this.transformPoint=h.transformPoint;this.translatePoint=h.translatePoint;this.translateShape=
h.translateShape;this.update=h.update;this.type="image";this.translate=h.translateShape;this.init(a,c,g);this.collection="shapes"}c.prototype.render=function(a){var c=this.attrsFromOptions(this.options),g=this.options;this.graphic=this.annotation.chart.renderer.image(g.src,0,-9E9,g.width,g.height).attr(c).add(a);this.graphic.width=g.width;this.graphic.height=g.height;h.render.call(this)};c.prototype.redraw=function(c){var m=this.anchor(this.points[0]);if(m=a.prototype.position.call(this,m))this.graphic[c?
"animate":"attr"]({x:m.x,y:m.y});else this.graphic.attr({x:0,y:-9E9});this.graphic.placed=!!m;h.redraw.call(this,c)};c.attrsMap={width:"width",height:"height",zIndex:"zIndex"};return c}()});r(a,"Extensions/Annotations/Annotations.js",[a["Core/Animation/AnimationUtilities.js"],a["Core/Chart/Chart.js"],a["Extensions/Annotations/Mixins/ControllableMixin.js"],a["Extensions/Annotations/Controllables/ControllableRect.js"],a["Extensions/Annotations/Controllables/ControllableCircle.js"],a["Extensions/Annotations/Controllables/ControllableEllipse.js"],
a["Extensions/Annotations/Controllables/ControllablePath.js"],a["Extensions/Annotations/Controllables/ControllableImage.js"],a["Extensions/Annotations/Controllables/ControllableLabel.js"],a["Extensions/Annotations/ControlPoint.js"],a["Extensions/Annotations/Mixins/EventEmitterMixin.js"],a["Core/Globals.js"],a["Extensions/Annotations/MockPoint.js"],a["Core/Pointer.js"],a["Core/Utilities.js"]],function(a,h,n,x,m,g,b,r,d,e,v,f,p,u,y){var c=a.getDeferredAnimation;a=h.prototype;var B=y.addEvent,D=y.defined,
z=y.destroyObjectProperties,E=y.erase,w=y.extend,l=y.find,q=y.fireEvent,k=y.merge,t=y.pick,A=y.splat;y=y.wrap;var F=function(){function a(a,b){this.annotation=void 0;this.coll="annotations";this.shapesGroup=this.labelsGroup=this.labelCollector=this.group=this.graphic=this.animationConfig=this.collection=void 0;this.chart=a;this.points=[];this.controlPoints=[];this.coll="annotations";this.labels=[];this.shapes=[];this.options=k(this.defaultOptions,b);this.userOptions=b;b=this.getLabelsAndShapesOptions(this.options,
b);this.options.labels=b.labels;this.options.shapes=b.shapes;this.init(a,this.options)}a.prototype.init=function(){var a=this.chart,b=this.options.animation;this.linkPoints();this.addControlPoints();this.addShapes();this.addLabels();this.setLabelCollector();this.animationConfig=c(a,b)};a.prototype.getLabelsAndShapesOptions=function(a,b){var c={};["labels","shapes"].forEach(function(d){a[d]&&(c[d]=b[d]?A(b[d]).map(function(b,c){return k(a[d][c],b)}):a[d])});return c};a.prototype.addShapes=function(){(this.options.shapes||
[]).forEach(function(a,b){a=this.initShape(a,b);k(!0,this.options.shapes[b],a.options)},this)};a.prototype.addLabels=function(){(this.options.labels||[]).forEach(function(a,b){a=this.initLabel(a,b);k(!0,this.options.labels[b],a.options)},this)};a.prototype.addClipPaths=function(){this.setClipAxes();this.clipXAxis&&this.clipYAxis&&this.options.crop&&(this.clipRect=this.chart.renderer.clipRect(this.getClipBox()))};a.prototype.setClipAxes=function(){var a=this.chart.xAxis,b=this.chart.yAxis,c=(this.options.labels||
[]).concat(this.options.shapes||[]).reduce(function(c,d){d=d&&(d.point||d.points&&d.points[0]);return[a[d&&d.xAxis]||c[0],b[d&&d.yAxis]||c[1]]},[]);this.clipXAxis=c[0];this.clipYAxis=c[1]};a.prototype.getClipBox=function(){if(this.clipXAxis&&this.clipYAxis)return{x:this.clipXAxis.left,y:this.clipYAxis.top,width:this.clipXAxis.width,height:this.clipYAxis.height}};a.prototype.setLabelCollector=function(){var a=this;a.labelCollector=function(){return a.labels.reduce(function(a,b){b.options.allowOverlap||
a.push(b.graphic);return a},[])};a.chart.labelCollectors.push(a.labelCollector)};a.prototype.setOptions=function(a){this.options=k(this.defaultOptions,a)};a.prototype.redraw=function(a){this.linkPoints();this.graphic||this.render();this.clipRect&&this.clipRect.animate(this.getClipBox());this.redrawItems(this.shapes,a);this.redrawItems(this.labels,a);n.redraw.call(this,a)};a.prototype.redrawItems=function(a,b){for(var c=a.length;c--;)this.redrawItem(a[c],b)};a.prototype.renderItems=function(a){for(var b=
a.length;b--;)this.renderItem(a[b])};a.prototype.render=function(){var a=this.chart.renderer;this.graphic=a.g("annotation").attr({opacity:0,zIndex:this.options.zIndex,visibility:this.options.visible?"visible":"hidden"}).add();this.shapesGroup=a.g("annotation-shapes").add(this.graphic);this.options.crop&&this.shapesGroup.clip(this.chart.plotBoxClip);this.labelsGroup=a.g("annotation-labels").attr({translateX:0,translateY:0}).add(this.graphic);this.addClipPaths();this.clipRect&&this.graphic.clip(this.clipRect);
this.renderItems(this.shapes);this.renderItems(this.labels);this.addEvents();n.render.call(this)};a.prototype.setVisibility=function(a){var b=this.options,c=this.chart.navigationBindings;a=t(a,!b.visible);this.graphic.attr("visibility",a?"visible":"hidden");a||(this.setControlPointsVisibility(!1),c.activeAnnotation===this&&c.popup&&"annotation-toolbar"===c.popup.formType&&q(c,"closePopup"));b.visible=a};a.prototype.setControlPointsVisibility=function(a){var b=function(b){b.setControlPointsVisibility(a)};
n.setControlPointsVisibility.call(this,a);this.shapes.forEach(b);this.labels.forEach(b)};a.prototype.destroy=function(){var a=this.chart,b=function(a){a.destroy()};this.labels.forEach(b);this.shapes.forEach(b);this.clipYAxis=this.clipXAxis=null;E(a.labelCollectors,this.labelCollector);v.destroy.call(this);n.destroy.call(this);z(this,a)};a.prototype.remove=function(){return this.chart.removeAnnotation(this)};a.prototype.update=function(a,b){var c=this.chart,d=this.getLabelsAndShapesOptions(this.userOptions,
a),A=c.annotations.indexOf(this);a=k(!0,this.userOptions,a);a.labels=d.labels;a.shapes=d.shapes;this.destroy();this.constructor(c,a);c.options.annotations[A]=a;this.isUpdating=!0;t(b,!0)&&c.redraw();q(this,"afterUpdate");this.isUpdating=!1};a.prototype.initShape=function(b,c){b=k(this.options.shapeOptions,{controlPointOptions:this.options.controlPointOptions},b);c=new a.shapesMap[b.type](this,b,c);c.itemType="shape";this.shapes.push(c);return c};a.prototype.initLabel=function(a,b){a=k(this.options.labelOptions,
{controlPointOptions:this.options.controlPointOptions},a);b=new d(this,a,b);b.itemType="label";this.labels.push(b);return b};a.prototype.redrawItem=function(a,b){a.linkPoints();a.shouldBeDrawn()?(a.graphic||this.renderItem(a),a.redraw(t(b,!0)&&a.graphic.placed),a.points.length&&this.adjustVisibility(a)):this.destroyItem(a)};a.prototype.adjustVisibility=function(a){var b=!1,c=a.graphic;a.points.forEach(function(a){!1!==a.series.visible&&!1!==a.visible&&(b=!0)});b?"hidden"===c.visibility&&c.show():
c.hide()};a.prototype.destroyItem=function(a){E(this[a.itemType+"s"],a);a.destroy()};a.prototype.renderItem=function(a){a.render("label"===a.itemType?this.labelsGroup:this.shapesGroup)};a.ControlPoint=e;a.MockPoint=p;a.shapesMap={rect:x,circle:m,ellipse:g,path:b,image:r};a.types={};return a}();k(!0,F.prototype,n,v,k(F.prototype,{nonDOMEvents:["add","afterUpdate","drag","remove"],defaultOptions:{visible:!0,animation:{},crop:!0,draggable:"xy",labelOptions:{align:"center",allowOverlap:!1,backgroundColor:"rgba(0, 0, 0, 0.75)",
borderColor:"#000000",borderRadius:3,borderWidth:1,className:"highcharts-no-tooltip",crop:!1,formatter:function(){return D(this.y)?this.y:"Annotation label"},includeInDataExport:!0,overflow:"justify",padding:5,shadow:!1,shape:"callout",style:{fontSize:"11px",fontWeight:"normal",color:"contrast"},useHTML:!1,verticalAlign:"bottom",x:0,y:-16},shapeOptions:{stroke:"rgba(0, 0, 0, 0.75)",strokeWidth:1,fill:"rgba(0, 0, 0, 0.75)",r:0,snap:2},controlPointOptions:{symbol:"circle",width:10,height:10,style:{cursor:"pointer",
fill:"#ffffff",stroke:"#000000","stroke-width":2},visible:!1,events:{}},events:{},zIndex:6}}));f.extendAnnotation=function(a,b,c,d){b=b||F;w(a.prototype,k(b.prototype,c));a.prototype.defaultOptions=k(a.prototype.defaultOptions,d||{})};w(a,{initAnnotation:function(a){a=new (F.types[a.type]||F)(this,a);this.annotations.push(a);return a},addAnnotation:function(a,b){a=this.initAnnotation(a);this.options.annotations.push(a.options);t(b,!0)&&(a.redraw(),a.graphic.attr({opacity:1}));return a},removeAnnotation:function(a){var b=
this.annotations,c="annotations"===a.coll?a:l(b,function(b){return b.options.id===a});c&&(q(c,"remove"),E(this.options.annotations,c.options),E(b,c),c.destroy())},drawAnnotations:function(){this.plotBoxClip.attr(this.plotBox);this.annotations.forEach(function(a){a.redraw();a.graphic.animate({opacity:1},a.animationConfig)})}});a.collectionsWithUpdate.push("annotations");a.collectionsWithInit.annotations=[a.addAnnotation];B(h,"afterInit",function(){this.annotations=[];this.options.annotations||(this.options.annotations=
[])});a.callbacks.push(function(a){a.plotBoxClip=this.renderer.clipRect(this.plotBox);a.controlPointsGroup=a.renderer.g("control-points").attr({zIndex:99}).clip(a.plotBoxClip).add();a.options.annotations.forEach(function(b,c){if(!a.annotations.some(function(a){return a.options===b})){var d=a.initAnnotation(b);a.options.annotations[c]=d.options}});a.drawAnnotations();B(a,"redraw",a.drawAnnotations);B(a,"destroy",function(){a.plotBoxClip.destroy();a.controlPointsGroup.destroy()});B(a,"exportData",function(b){var c=
(this.options.exporting&&this.options.exporting.csv||{}).columnHeaderFormatter,d=!b.dataRows[1].xValues,k=a.options.lang&&a.options.lang.exportData&&a.options.lang.exportData.annotationHeader,t=function(a){if(c){var b=c(a);if(!1!==b)return b}b=k+" "+a;return d?{columnTitle:b,topLevelColumnTitle:b}:b},A=b.dataRows[0].length,w=a.options.exporting&&a.options.exporting.csv&&a.options.exporting.csv.annotations&&a.options.exporting.csv.annotations.itemDelimiter,F=a.options.exporting&&a.options.exporting.csv&&
a.options.exporting.csv.annotations&&a.options.exporting.csv.annotations.join;a.annotations.forEach(function(a){a.options.labelOptions.includeInDataExport&&a.labels.forEach(function(a){if(a.options.text){var c=a.options.text;a.points.forEach(function(a){var d=a.x,k=a.series.xAxis?a.series.xAxis.options.index:-1,t=!1;if(-1===k){a=b.dataRows[0].length;for(var e=Array(a),l=0;l<a;++l)e[l]="";e.push(c);e.xValues=[];e.xValues[k]=d;b.dataRows.push(e);t=!0}t||b.dataRows.forEach(function(a,b){!t&&a.xValues&&
void 0!==k&&d===a.xValues[k]&&(F&&a.length>A?a[a.length-1]+=w+c:a.push(c),t=!0)});if(!t){a=b.dataRows[0].length;e=Array(a);for(l=0;l<a;++l)e[l]="";e[0]=d;e.push(c);e.xValues=[];void 0!==k&&(e.xValues[k]=d);b.dataRows.push(e)}})}})});var e=0;b.dataRows.forEach(function(a){e=Math.max(e,a.length)});for(var l=e-b.dataRows[0].length,H=0;H<l;H++){var q=t(H+1);d?(b.dataRows[0].push(q.topLevelColumnTitle),b.dataRows[1].push(q.columnTitle)):b.dataRows[0].push(q)}})});y(u.prototype,"onContainerMouseDown",function(a){this.chart.hasDraggedAnnotation||
a.apply(this,Array.prototype.slice.call(arguments,1))});f.Annotation=F;"";return F});r(a,"Core/Chart/ChartNavigationComposition.js",[],function(){var a;(function(a){a.compose=function(a){a.navigation||(a.navigation=new c(a));return a};var c=function(){function a(a){this.updates=[];this.chart=a}a.prototype.addUpdate=function(a){this.chart.navigation.updates.push(a)};a.prototype.update=function(a,c){var b=this;this.updates.forEach(function(g){g.call(b.chart,a,c)})};return a}();a.Additions=c})(a||(a=
{}));return a});r(a,"Extensions/Annotations/NavigationBindings.js",[a["Extensions/Annotations/Annotations.js"],a["Core/Chart/Chart.js"],a["Core/Chart/ChartNavigationComposition.js"],a["Core/FormatUtilities.js"],a["Core/Globals.js"],a["Core/DefaultOptions.js"],a["Core/Utilities.js"]],function(a,h,n,r,m,g,b){function c(a,b){var c=l.Element.prototype,d=c.matches||c.msMatchesSelector||c.webkitMatchesSelector,k=null;if(c.closest)k=c.closest.call(a,b);else{do{if(d.call(a,b))return a;a=a.parentElement||
a.parentNode}while(null!==a&&1===a.nodeType)}return k}function d(a){var b=a.prototype.defaultOptions.events&&a.prototype.defaultOptions.events.click;D(!0,a.prototype.defaultOptions.events,{click:function(a){var c=this,d=c.chart.navigationBindings,k=d.activeAnnotation;b&&b.call(c,a);k!==c?(d.deselectAnnotation(),d.activeAnnotation=c,c.setControlPointsVisibility(!0),p(d,"showPopup",{annotation:c,formType:"annotation-toolbar",options:d.annotationToFields(c),onSubmit:function(a){var b={};"remove"===a.actionType?
(d.activeAnnotation=!1,d.chart.removeAnnotation(c)):(d.fieldsToOptions(a.fields,b),d.deselectAnnotation(),a=b.typeOptions,"measure"===c.options.type&&(a.crosshairY.enabled=0!==a.crosshairY.strokeWidth,a.crosshairX.enabled=0!==a.crosshairX.strokeWidth),c.update(b))}})):p(d,"closePopup");a.activeAnnotation=!0}})}var e=r.format;r=g.setOptions;var v=b.addEvent,f=b.attr,p=b.fireEvent,u=b.isArray,x=b.isFunction,C=b.isNumber,G=b.isObject,D=b.merge,z=b.objectEach,E=b.pick,w=m.doc,l=m.win,q=function(){function a(a,
b){this.selectedButton=this.boundClassNames=void 0;this.chart=a;this.options=b;this.eventsToUnbind=[];this.container=w.getElementsByClassName(this.options.bindingsClassName||"")}a.prototype.initEvents=function(){var a=this,b=a.chart,c=a.container,d=a.options;a.boundClassNames={};z(d.bindings||{},function(b){a.boundClassNames[b.className]=b});[].forEach.call(c,function(b){a.eventsToUnbind.push(v(b,"click",function(c){var d=a.getButtonEvents(b,c);d&&-1===d.button.className.indexOf("highcharts-disabled-btn")&&
a.bindingsButtonClick(d.button,d.events,c)}))});z(d.events||{},function(b,c){x(b)&&a.eventsToUnbind.push(v(a,c,b,{passive:!1}))});a.eventsToUnbind.push(v(b.container,"click",function(c){!b.cancelClick&&b.isInsidePlot(c.chartX-b.plotLeft,c.chartY-b.plotTop,{visiblePlotOnly:!0})&&a.bindingsChartClick(this,c)}));a.eventsToUnbind.push(v(b.container,m.isTouchDevice?"touchmove":"mousemove",function(b){a.bindingsContainerMouseMove(this,b)},m.isTouchDevice?{passive:!1}:void 0))};a.prototype.initUpdate=function(){var a=
this;n.compose(this.chart).navigation.addUpdate(function(b){a.update(b)})};a.prototype.bindingsButtonClick=function(a,b,c){var d=this.chart,k=d.renderer.boxWrapper,t=!0;this.selectedButtonElement&&(this.selectedButtonElement.classList===a.classList&&(t=!1),p(this,"deselectButton",{button:this.selectedButtonElement}),this.nextEvent&&(this.currentUserDetails&&"annotations"===this.currentUserDetails.coll&&d.removeAnnotation(this.currentUserDetails),this.mouseMoveEvent=this.nextEvent=!1));t?(this.selectedButton=
b,this.selectedButtonElement=a,p(this,"selectButton",{button:a}),b.init&&b.init.call(this,a,c),(b.start||b.steps)&&d.renderer.boxWrapper.addClass("highcharts-draw-mode")):(d.stockTools&&d.stockTools.toggleButtonAciveClass(a),k.removeClass("highcharts-draw-mode"),this.mouseMoveEvent=this.nextEvent=!1,this.selectedButton=null)};a.prototype.bindingsChartClick=function(a,b){a=this.chart;var d=this.activeAnnotation,k=this.selectedButton;a=a.renderer.boxWrapper;d&&(d.cancelClick||b.activeAnnotation||!b.target.parentNode||
c(b.target,".highcharts-popup")?d.cancelClick&&setTimeout(function(){d.cancelClick=!1},0):p(this,"closePopup"));k&&k.start&&(this.nextEvent?(this.nextEvent(b,this.currentUserDetails),this.steps&&(this.stepIndex++,k.steps[this.stepIndex]?this.mouseMoveEvent=this.nextEvent=k.steps[this.stepIndex]:(p(this,"deselectButton",{button:this.selectedButtonElement}),a.removeClass("highcharts-draw-mode"),k.end&&k.end.call(this,b,this.currentUserDetails),this.mouseMoveEvent=this.nextEvent=!1,this.selectedButton=
null))):(this.currentUserDetails=k.start.call(this,b))&&k.steps?(this.stepIndex=0,this.steps=!0,this.mouseMoveEvent=this.nextEvent=k.steps[this.stepIndex]):(p(this,"deselectButton",{button:this.selectedButtonElement}),a.removeClass("highcharts-draw-mode"),this.steps=!1,this.selectedButton=null,k.end&&k.end.call(this,b,this.currentUserDetails)))};a.prototype.bindingsContainerMouseMove=function(a,b){this.mouseMoveEvent&&this.mouseMoveEvent(b,this.currentUserDetails)};a.prototype.fieldsToOptions=function(a,
b){z(a,function(a,c){var d=parseFloat(a),k=c.split("."),t=b,e=k.length-1;!C(d)||a.match(/px/g)||c.match(/format/g)||(a=d);""!==a&&"undefined"!==a&&k.forEach(function(b,c){var d=E(k[c+1],"");e===c?t[b]=a:(t[b]||(t[b]=d.match(/\d/g)?[]:{}),t=t[b])})});return b};a.prototype.deselectAnnotation=function(){this.activeAnnotation&&(this.activeAnnotation.setControlPointsVisibility(!1),this.activeAnnotation=!1)};a.prototype.annotationToFields=function(b){function c(a,d,k,l){if(k&&a&&-1===q.indexOf(d)&&(0<=
(k.indexOf&&k.indexOf(d))||k[d]||!0===k))if(u(a))l[d]=[],a.forEach(function(a,b){G(a)?(l[d][b]={},z(a,function(a,k){c(a,k,t[d],l[d][b])})):c(a,0,t[d],l[d])});else if(G(a)){var A={};u(l)?(l.push(A),A[d]={},A=A[d]):l[d]=A;z(a,function(a,b){c(a,b,0===d?k:t[d],A)})}else"format"===d?l[d]=[e(a,b.labels[0].points[0]).toString(),"text"]:u(l)?l.push([a,w(a)]):l[d]=[a,w(a)]}var d=b.options,k=a.annotationsEditable,t=k.nestedOptions,w=this.utils.getFieldType,l=E(d.type,d.shapes&&d.shapes[0]&&d.shapes[0].type,
d.labels&&d.labels[0]&&d.labels[0].itemType,"label"),q=a.annotationsNonEditable[d.langKey]||[],g={langKey:d.langKey,type:l};z(d,function(a,b){"typeOptions"===b?(g[b]={},z(d[b],function(a,d){c(a,d,t,g[b],!0)})):c(a,b,k[l],g)});return g};a.prototype.getClickedClassNames=function(a,b){var c=b.target;b=[];for(var d;c&&((d=f(c,"class"))&&(b=b.concat(d.split(" ").map(function(a){return[a,c]}))),c=c.parentNode,c!==a););return b};a.prototype.getButtonEvents=function(a,b){var c=this,d;this.getClickedClassNames(a,
b).forEach(function(a){c.boundClassNames[a[0]]&&!d&&(d={events:c.boundClassNames[a[0]],button:a[1]})});return d};a.prototype.update=function(a){this.options=D(!0,this.options,a);this.removeEvents();this.initEvents()};a.prototype.removeEvents=function(){this.eventsToUnbind.forEach(function(a){a()})};a.prototype.destroy=function(){this.removeEvents()};a.annotationsEditable={nestedOptions:{labelOptions:["style","format","backgroundColor"],labels:["style"],label:["style"],style:["fontSize","color"],background:["fill",
"strokeWidth","stroke"],innerBackground:["fill","strokeWidth","stroke"],outerBackground:["fill","strokeWidth","stroke"],shapeOptions:["fill","strokeWidth","stroke"],shapes:["fill","strokeWidth","stroke"],line:["strokeWidth","stroke"],backgroundColors:[!0],connector:["fill","strokeWidth","stroke"],crosshairX:["strokeWidth","stroke"],crosshairY:["strokeWidth","stroke"]},circle:["shapes"],ellipse:["shapes"],verticalLine:[],label:["labelOptions"],measure:["background","crosshairY","crosshairX"],fibonacci:[],
tunnel:["background","line","height"],pitchfork:["innerBackground","outerBackground"],rect:["shapes"],crookedLine:[],basicAnnotation:["shapes","labelOptions"]};a.annotationsNonEditable={rectangle:["crosshairX","crosshairY","labelOptions"],ellipse:["labelOptions"],circle:["labelOptions"]};return a}();q.prototype.utils={getFieldType:function(a){return{string:"text",number:"number","boolean":"checkbox"}[typeof a]},updateRectSize:function(a,b){var c=b.chart,d=b.options.typeOptions,k=C(d.xAxis)&&c.xAxis[d.xAxis],
t=C(d.yAxis)&&c.yAxis[d.yAxis];k&&t&&(k=k.toValue(a[k.horiz?"chartX":"chartY"]),a=t.toValue(a[t.horiz?"chartX":"chartY"]),t=k-d.point.x,d=d.point.y-a,b.update({typeOptions:{background:{width:c.inverted?d:t,height:c.inverted?t:d}}}))},getAssignedAxis:function(a){return a.filter(function(a){var b=a.axis.getExtremes(),c=b.min;b=b.max;var d=E(a.axis.minPointOffset,0);return C(c)&&C(b)&&a.value>=c-d&&a.value<=b+d&&!a.axis.options.isInternal})[0]}};h.prototype.initNavigationBindings=function(){var a=this.options;
a&&a.navigation&&a.navigation.bindings&&(this.navigationBindings=new q(this,a.navigation),this.navigationBindings.initEvents(),this.navigationBindings.initUpdate())};v(h,"load",function(){this.initNavigationBindings()});v(h,"destroy",function(){this.navigationBindings&&this.navigationBindings.destroy()});v(q,"deselectButton",function(){this.selectedButtonElement=null});v(a,"remove",function(){this.chart.navigationBindings&&this.chart.navigationBindings.deselectAnnotation()});m.Annotation&&(d(a),z(a.types,
function(a){d(a)}));r({lang:{navigation:{popup:{simpleShapes:"Simple shapes",lines:"Lines",circle:"Circle",ellipse:"Ellipse",rectangle:"Rectangle",label:"Label",shapeOptions:"Shape options",typeOptions:"Details",fill:"Fill",format:"Text",strokeWidth:"Line width",stroke:"Line color",title:"Title",name:"Name",labelOptions:"Label options",labels:"Labels",backgroundColor:"Background color",backgroundColors:"Background colors",borderColor:"Border color",borderRadius:"Border radius",borderWidth:"Border width",
style:"Style",padding:"Padding",fontSize:"Font size",color:"Color",height:"Height",shapes:"Shape options"}}},navigation:{bindingsClassName:"highcharts-bindings-container",bindings:{circleAnnotation:{className:"highcharts-circle-annotation",start:function(a){var b=this.chart.pointer.getCoordinates(a);a=this.utils.getAssignedAxis(b.xAxis);b=this.utils.getAssignedAxis(b.yAxis);var c=this.chart.options.navigation;if(a&&b)return this.chart.addAnnotation(D({langKey:"circle",type:"basicAnnotation",shapes:[{type:"circle",
point:{x:a.value,y:b.value,xAxis:a.axis.options.index,yAxis:b.axis.options.index},r:5}]},c.annotationsOptions,c.bindings.circleAnnotation.annotationsOptions))},steps:[function(a,b){var c=b.options.shapes[0].point;if(C(c.xAxis)&&C(c.yAxis)){var d=this.chart.inverted;var k=this.chart.xAxis[c.xAxis].toPixels(c.x);c=this.chart.yAxis[c.yAxis].toPixels(c.y);d=Math.max(Math.sqrt(Math.pow(d?c-a.chartX:k-a.chartX,2)+Math.pow(d?k-a.chartY:c-a.chartY,2)),5)}b.update({shapes:[{r:d}]})}]},ellipseAnnotation:{className:"highcharts-ellipse-annotation",
start:function(a){var b=this.chart.pointer.getCoordinates(a);a=this.utils.getAssignedAxis(b.xAxis);b=this.utils.getAssignedAxis(b.yAxis);var c=this.chart.options.navigation;if(a&&b)return this.chart.addAnnotation(D({langKey:"ellipse",type:"basicAnnotation",shapes:[{type:"ellipse",xAxis:a.axis.options.index,yAxis:b.axis.options.index,points:[{x:a.value,y:b.value},{x:a.value,y:b.value}],ry:1}]},c.annotationsOptions,c.bindings.ellipseAnnotation.annotationOptions))},steps:[function(a,b){b=b.shapes[0];
var c=b.getAbsolutePosition(b.points[1]);b.translatePoint(a.chartX-c.x,a.chartY-c.y,1);b.redraw(!1)},function(a,b){b=b.shapes[0];var c=b.getAbsolutePosition(b.points[0]),d=b.getAbsolutePosition(b.points[1]);a=b.getDistanceFromLine(c,d,a.chartX,a.chartY);c=b.getYAxis();a=Math.abs(c.toValue(0)-c.toValue(a));b.setYRadius(a);b.redraw(!1)}]},rectangleAnnotation:{className:"highcharts-rectangle-annotation",start:function(a){a=this.chart.pointer.getCoordinates(a);var b=this.utils.getAssignedAxis(a.xAxis),
c=this.utils.getAssignedAxis(a.yAxis);if(b&&c){a=b.value;var d=c.value;b=b.axis.options.index;c=c.axis.options.index;var k=this.chart.options.navigation;return this.chart.addAnnotation(D({langKey:"rectangle",type:"basicAnnotation",shapes:[{type:"path",points:[{xAxis:b,yAxis:c,x:a,y:d},{xAxis:b,yAxis:c,x:a,y:d},{xAxis:b,yAxis:c,x:a,y:d},{xAxis:b,yAxis:c,x:a,y:d},{command:"Z"}]}]},k.annotationsOptions,k.bindings.rectangleAnnotation.annotationsOptions))}},steps:[function(a,b){var c=b.options.shapes[0].points,
d=this.chart.pointer.getCoordinates(a);a=this.utils.getAssignedAxis(d.xAxis);d=this.utils.getAssignedAxis(d.yAxis);a&&d&&(a=a.value,d=d.value,c[1].x=a,c[2].x=a,c[2].y=d,c[3].y=d,b.update({shapes:[{points:c}]}))}]},labelAnnotation:{className:"highcharts-label-annotation",start:function(a){var b=this.chart.pointer.getCoordinates(a);a=this.utils.getAssignedAxis(b.xAxis);b=this.utils.getAssignedAxis(b.yAxis);var c=this.chart.options.navigation;if(a&&b)return this.chart.addAnnotation(D({langKey:"label",
type:"basicAnnotation",labelOptions:{format:"{y:.2f}"},labels:[{point:{xAxis:a.axis.options.index,yAxis:b.axis.options.index,x:a.value,y:b.value},overflow:"none",crop:!0}]},c.annotationsOptions,c.bindings.labelAnnotation.annotationsOptions))}}},events:{},annotationsOptions:{animation:{defer:0}}}});v(h,"render",function(){var a=this,b=a.navigationBindings;if(a&&b){var c=!1;a.series.forEach(function(a){!a.options.isInternal&&a.visible&&(c=!0)});z(b.boundClassNames,function(b,d){if(a.navigationBindings&&
a.navigationBindings.container&&a.navigationBindings.container[0]&&(d=a.navigationBindings.container[0].querySelectorAll("."+d)))for(var e=0;e<d.length;e++){var l=d[e];"normal"===b.noDataState?-1!==l.className.indexOf("highcharts-disabled-btn")&&l.classList.remove("highcharts-disabled-btn"):c?-1!==l.className.indexOf("highcharts-disabled-btn")&&l.classList.remove("highcharts-disabled-btn"):-1===l.className.indexOf("highcharts-disabled-btn")&&(l.className+=" highcharts-disabled-btn")}})}});v(q,"closePopup",
function(){this.deselectAnnotation()});return q});r(a,"Extensions/Annotations/Popup.js",[a["Core/Globals.js"],a["Extensions/Annotations/NavigationBindings.js"],a["Core/DefaultOptions.js"],a["Core/Pointer.js"],a["Core/Utilities.js"]],function(a,h,n,r,m){var c=a.doc,b=a.isFirefox,x=n.getOptions,d=m.addEvent,e=m.createElement,v=m.defined,f=m.fireEvent,p=m.isArray,u=m.isObject,y=m.objectEach,C=m.pick,G=m.stableSort;n=m.wrap;var D=/\d/g,z;(function(a){a[a["params.algorithm"]=0]="params.algorithm";a[a["params.average"]=
1]="params.average"})(z||(z={}));var E={"algorithm-pivotpoints":["standard","fibonacci","camarilla"],"average-disparityindex":["sma","ema","dema","tema","wma"]};n(r.prototype,"onContainerMouseDown",function(a,b){this.inClass(b.target,"highcharts-popup")||a.apply(this,Array.prototype.slice.call(arguments,1))});a.Popup=function(a,b,c){this.init(a,b,c)};a.Popup.prototype={init:function(b,c,q){this.chart=q;this.container=e("div",{className:"highcharts-popup highcharts-no-tooltip"},void 0,b);d(this.container,
"mousedown",function(){var b=q&&q.navigationBindings&&q.navigationBindings.activeAnnotation;if(b){b.cancelClick=!0;var c=d(a.doc,"click",function(){setTimeout(function(){b.cancelClick=!1},0);c()})}});this.lang=this.getLangpack();this.iconsURL=c;this.addCloseBtn()},addCloseBtn:function(){var a=this,b=this.iconsURL;var c=e("div",{className:"highcharts-popup-close"},void 0,this.container);c.style["background-image"]="url("+(b.match(/png|svg|jpeg|jpg|gif/ig)?b:b+"close.svg")+")";["click","touchstart"].forEach(function(b){d(c,
b,function(){a.chart?f(a.chart.navigationBindings,"closePopup"):a.closePopup()})})},addColsContainer:function(a){var b=e("div",{className:"highcharts-popup-lhs-col"},void 0,a);a=e("div",{className:"highcharts-popup-rhs-col"},void 0,a);e("div",{className:"highcharts-popup-rhs-col-wrapper"},void 0,a);return{lhsCol:b,rhsCol:a}},addInput:function(a,b,d,k){var l=a.split(".");l=l[l.length-1];var w=this.lang;b="highcharts-"+b+"-"+C(k.htmlFor,l);b.match(D)||e("label",{htmlFor:b,className:k.labelClassName},
void 0,d).appendChild(c.createTextNode(w[l]||l));d=e("input",{name:b,value:k.value,type:k.type,className:"highcharts-popup-field"},void 0,d);d.setAttribute("highcharts-data-name",a);return d},addButton:function(a,b,q,k,f){var l=this,w=this.closePopup,g=this.getFields;var t=e("button",void 0,void 0,a);t.appendChild(c.createTextNode(b));f&&["click","touchstart"].forEach(function(a){d(t,a,function(){w.call(l);return f(g(k,q))})});return t},getFields:function(a,b){var c=Array.prototype.slice.call(a.querySelectorAll("input")),
d=Array.prototype.slice.call(a.querySelectorAll("select")),e=a.querySelectorAll("#highcharts-select-series > option:checked")[0];a=a.querySelectorAll("#highcharts-select-volume > option:checked")[0];var l={actionType:b,linkedTo:e&&e.getAttribute("value")||"",fields:{}};c.forEach(function(a){var b=a.getAttribute("highcharts-data-name");a.getAttribute("highcharts-data-series-id")?l.seriesId=a.value:b?l.fields[b]=a.value:l.type=a.value});d.forEach(function(a){var b=a.id;"highcharts-select-series"!==
b&&"highcharts-select-volume"!==b&&(b=b.split("highcharts-select-")[1],l.fields[b]=a.value)});a&&(l.fields["params.volumeSeriesID"]=a.getAttribute("value")||"");return l},showPopup:function(){var a=this.container,b=a.querySelectorAll(".highcharts-popup-close")[0];this.formType=void 0;a.innerHTML="";0<=a.className.indexOf("highcharts-annotation-toolbar")&&(a.classList.remove("highcharts-annotation-toolbar"),a.removeAttribute("style"));a.appendChild(b);a.style.display="block";a.style.height=""},closePopup:function(){C(this.popup&&
this.popup.container,this.container).style.display="none"},showForm:function(a,b,c,d){b&&(this.popup=b.navigationBindings.popup,this.showPopup(),"indicators"===a&&this.indicators.addForm.call(this,b,c,d),"annotation-toolbar"===a&&this.annotations.addToolbar.call(this,b,c,d),"annotation-edit"===a&&this.annotations.addForm.call(this,b,c,d),"flag"===a&&this.annotations.addForm.call(this,b,c,d,!0),this.formType=a,this.container.style.height=this.container.offsetHeight+"px")},getLangpack:function(){return x().lang.navigation.popup},
annotations:{addToolbar:function(a,b,d){var l=this,f=this.lang,g=this.popup.container,w=this.showForm;-1===g.className.indexOf("highcharts-annotation-toolbar")&&(g.className+=" highcharts-annotation-toolbar");a&&(g.style.top=a.plotTop+10+"px");e("span",void 0,void 0,g).appendChild(c.createTextNode(C(f[b.langKey]||b.langKey,b.shapes&&b.shapes[0].type)));var q=this.addButton(g,f.removeButton||"remove","remove",g,d);q.className+=" highcharts-annotation-remove-button";q.style["background-image"]="url("+
this.iconsURL+"destroy.svg)";q=this.addButton(g,f.editButton||"edit","edit",g,function(){w.call(l,"annotation-edit",a,b,d)});q.className+=" highcharts-annotation-edit-button";q.style["background-image"]="url("+this.iconsURL+"edit.svg)"},addForm:function(a,b,d,k){var l=this.popup.container,g=this.lang;if(a){var f=e("h2",{className:"highcharts-popup-main-title"},void 0,l);f.appendChild(c.createTextNode(g[b.langKey]||b.langKey||""));f=e("div",{className:"highcharts-popup-lhs-col highcharts-popup-lhs-full"},
void 0,l);var q=e("div",{className:"highcharts-popup-bottom-row"},void 0,l);this.annotations.addFormFields.call(this,f,a,"",b,[],!0);this.addButton(q,k?g.addButton||"add":g.saveButton||"save",k?"add":"save",l,d)}},addFormFields:function(a,d,g,k,f,h){var l=this,q=this.annotations.addFormFields,w=this.addInput,t=this.lang,m,A;d&&(y(k,function(b,c){m=""!==g?g+"."+c:c;u(b)&&(!p(b)||p(b)&&u(b[0])?(A=t[c]||c,A.match(D)||f.push([!0,A,a]),q.call(l,a,d,m,b,f,!1)):f.push([l,m,"annotation",a,b]))}),h&&(G(f,
function(a){return a[1].match(/format/g)?-1:1}),b&&f.reverse(),f.forEach(function(a){!0===a[0]?e("span",{className:"highcharts-annotation-title"},void 0,a[2]).appendChild(c.createTextNode(a[1])):(a[4]={value:a[4][0],type:a[4][1]},w.apply(a[0],a.splice(1)))})))}},indicators:{addForm:function(a,b,c){var d=this.indicators,e=this.lang;if(a){this.tabs.init.call(this,a);b=this.popup.container.querySelectorAll(".highcharts-tab-item-content");this.addColsContainer(b[0]);d.addSearchBox.call(this,a,b[0]);d.addIndicatorList.call(this,
a,b[0],"add");var l=b[0].querySelectorAll(".highcharts-popup-rhs-col")[0];this.addButton(l,e.addButton||"add","add",l,c);this.addColsContainer(b[1]);d.addIndicatorList.call(this,a,b[1],"edit");l=b[1].querySelectorAll(".highcharts-popup-rhs-col")[0];this.addButton(l,e.saveButton||"save","edit",l,c);this.addButton(l,e.removeButton||"remove","remove",l,c)}},filterSeries:function(a,b){var c=this.indicators,d=this.chart&&this.chart.options.lang,e=d&&d.navigation&&d.navigation.popup&&d.navigation.popup.indicatorAliases,
l=[],f;y(a,function(a,d){var k=a.options;if(a.params||k&&k.params)if(k=c.getNameType(a,d),d=k.indicatorFullName,k=k.indicatorType,b){var g=b.replace(/[.*+?^${}()|[\]\\]/g,"\\$&");g=new RegExp(g,"i");var q=e&&e[k]&&e[k].join(" ")||"";if(d.match(g)||q.match(g))f={indicatorFullName:d,indicatorType:k,series:a},l.push(f)}else f={indicatorFullName:d,indicatorType:k,series:a},l.push(f)});return l},filterSeriesArray:function(a){var b=[],c;a.forEach(function(a){a.is("sma")&&(c={indicatorFullName:a.name,indicatorType:a.type,
series:a},b.push(c))});return b},addIndicatorList:function(a,b,f,k){var l=this,g=l.indicators,h=l.lang,q=b.querySelectorAll(".highcharts-popup-lhs-col")[0];b=b.querySelectorAll(".highcharts-popup-rhs-col")[0];var w="edit"===f,m=this.indicators.addFormFields;f=w?a.series:a.options.plotOptions||{};if(a||!f){var n,r=[];w||p(f)?p(f)&&(r=g.filterSeriesArray.call(this,f)):r=g.filterSeries.call(this,f,k);G(r,function(a,b){a=a.indicatorFullName.toLowerCase();b=b.indicatorFullName.toLowerCase();return a<b?
-1:a>b?1:0});q.children[1]&&q.children[1].remove();var v=e("ul",{className:"highcharts-indicator-list"},void 0,q);var u=b.querySelectorAll(".highcharts-popup-rhs-col-wrapper")[0];r.forEach(function(b){var k=b.indicatorFullName,f=b.indicatorType,g=b.series;n=e("li",{className:"highcharts-indicator-list"},void 0,v);n.appendChild(c.createTextNode(k));["click","touchstart"].forEach(function(b){d(n,b,function(){var b=u.parentNode.children[1];m.call(l,a,g,f,u);b&&(b.style.display="block");w&&g.options&&
e("input",{type:"hidden",name:"highcharts-id-"+f,value:g.options.id},void 0,u).setAttribute("highcharts-data-series-id",g.options.id)})})});0<v.childNodes.length?v.childNodes[0].click():w||(u.parentNode.children[0].innerHTML=h.noFilterMatch||"",u.parentNode.children[1].style.display="none")}},addSearchBox:function(a,b){var c=this,l=b.querySelectorAll(".highcharts-popup-lhs-col")[0];b=this.lang.clearFilter;l=e("div",{className:"highcharts-input-wrapper"},void 0,l);var f=function(b){c.indicators.addIndicatorList.call(c,
a,c.container,"add",b)},g=this.addInput("searchIndicators","input",l,{value:"",type:"text",htmlFor:"search-indicators",labelClassName:"highcharts-input-search-indicators-label"}),h=e("a",{textContent:b},void 0,l);g.classList.add("highcharts-input-search-indicators");h.classList.add("clear-filter-button");d(g,"input",function(a){f(this.value);h.style.display=this.value.length?"inline-block":"none"});["click","touchstart"].forEach(function(a){d(h,a,function(){g.value="";f("");h.style.display="none"})})},
addSelection:function(a,b,d){var f=b.split(".");f=f[f.length-1];a="highcharts-"+b+"-type-"+a;var l=this.lang;e("label",{htmlFor:a},null,d).appendChild(c.createTextNode(l[f]||b));d=e("select",{name:a,className:"highcharts-popup-field",id:"highcharts-select-"+b},null,d);d.setAttribute("id","highcharts-select-"+b);return d},addSelectionOptions:function(a,b,d,f,g,h,m){"series"===b||"volume"===b?a.series.forEach(function(a){var f=a.options,l=f.name||f.params?a.name:f.id||"";"highcharts-navigator-series"!==
f.id&&f.id!==(m&&m.options&&m.options.id)&&(v(h)||"volume"!==b||"column"!==a.type||(h=f.id),e("option",{value:f.id},void 0,d).appendChild(c.createTextNode(l)))}):f&&g&&E[g+"-"+f].forEach(function(a){e("option",{value:a},void 0,d).appendChild(c.createTextNode(a))});v(h)&&(d.value=h)},getNameType:function(b,c){var d=b.options,e=a.seriesTypes;e=e[c]&&e[c].prototype.nameBase||c.toUpperCase();d&&d.type&&(c=b.options.type,e=b.name);return{indicatorFullName:e,indicatorType:c}},listAllSeries:function(a,b,
c,d,e,f){var g=this.indicators;c&&(a=g.addSelection.call(this,a,b,d),g.addSelectionOptions.call(this,c,b,a,void 0,void 0,void 0,e),v(f)&&(a.value=f))},addFormFields:function(a,b,d,f){var g=b.params||b.options.params,l=this.indicators.getNameType;f.innerHTML="";e("h3",{className:"highcharts-indicator-title"},void 0,f).appendChild(c.createTextNode(l(b,d).indicatorFullName));e("input",{type:"hidden",name:"highcharts-type-"+d,value:d},void 0,f);this.indicators.listAllSeries.call(this,d,"series",a,f,b,
b.linkedParent&&b.linkedParent.options.id);g.volumeSeriesID&&this.indicators.listAllSeries.call(this,d,"volume",a,f,b,b.linkedParent&&g.volumeSeriesID);this.indicators.addParamInputs.call(this,a,"params",g,d,f)},addParamInputs:function(a,b,c,d,e){var f=this,g=f.indicators,l=this.indicators.addParamInputs,k=this.addInput,h;a&&y(c,function(c,m){h=b+"."+m;if(v(c)&&h)if(u(c)&&(k.call(f,h,d,e,{}),l.call(f,a,h,c,d,e)),h in z){var q=g.addSelection.call(f,d,h,e);g.addSelectionOptions.call(f,a,b,q,d,m,c)}else"params.volumeSeriesID"===
h||p(c)||k.call(f,h,d,e,{value:c,type:"text"})})},getAmount:function(){var a=0;this.series.forEach(function(b){var c=b.options;(b.params||c&&c.params)&&a++});return a}},tabs:{init:function(a){var b=this.tabs,c=this.indicators.getAmount.call(a);a&&(a=b.addMenuItem.call(this,"add"),b.addMenuItem.call(this,"edit",c),b.addContentItem.call(this,"add"),b.addContentItem.call(this,"edit"),b.switchTabs.call(this,c),b.selectTab.call(this,a,0))},addMenuItem:function(a,b){var d=this.popup.container,f="highcharts-tab-item",
g=this.lang;0===b&&(f+=" highcharts-tab-disabled");b=e("span",{className:f},void 0,d);b.appendChild(c.createTextNode(g[a+"Button"]||a));b.setAttribute("highcharts-data-tab-type",a);return b},addContentItem:function(){return e("div",{className:"highcharts-tab-item-content highcharts-no-mousewheel"},void 0,this.popup.container)},switchTabs:function(a){var b=this,c;this.popup.container.querySelectorAll(".highcharts-tab-item").forEach(function(e,f){c=e.getAttribute("highcharts-data-tab-type");"edit"===
c&&0===a||["click","touchstart"].forEach(function(a){d(e,a,function(){b.tabs.deselectAll.call(b);b.tabs.selectTab.call(b,this,f)})})})},selectTab:function(a,b){var c=this.popup.container.querySelectorAll(".highcharts-tab-item-content");a.className+=" highcharts-tab-item-active";c[b].className+=" highcharts-tab-item-show"},deselectAll:function(){var a=this.popup.container,b=a.querySelectorAll(".highcharts-tab-item");a=a.querySelectorAll(".highcharts-tab-item-content");var c;for(c=0;c<b.length;c++)b[c].classList.remove("highcharts-tab-item-active"),
a[c].classList.remove("highcharts-tab-item-show")}}};d(h,"showPopup",function(b){this.popup||(this.popup=new a.Popup(this.chart.container,this.chart.options.navigation.iconsURL||this.chart.options.stockTools&&this.chart.options.stockTools.gui.iconsURL||"https://code.highcharts.com/9.3.1/gfx/stock-icons/",this.chart));this.popup.showForm(b.formType,this.chart,b.options,b.onSubmit)});d(h,"closePopup",function(){this.popup&&this.popup.closePopup()});return a.Popup});r(a,"masters/modules/annotations.src.js",
[],function(){})});
//# sourceMappingURL=annotations.js.map
```
