## Supplementary material for "Exploring natural odour landscapes: A case study with implications for human-biting insects": Figure S4: arrow-symbols.js

```
/*
 Highcharts JS v9.3.1 (2021-11-05)

 Arrow Symbols

 (c) 2017-2021 Lars A. V. Cabrera

 License: www.highcharts.com/license
*/
'use strict';(function(a){"object"===typeof module&&module.exports?(a["default"]=a,module.exports=a):"function"===typeof define&&define.amd?define("highcharts/modules/arrow-symbols",["highcharts"],function(b){a(b);a.Highcharts=b;return a}):a("undefined"!==typeof Highcharts?Highcharts:void 0)})(function(a){function b(a,b,f,g){a.hasOwnProperty(b)||(a[b]=g.apply(null,f))}a=a?a._modules:{};b(a,"Extensions/ArrowSymbols.js",[a["Core/Renderer/SVG/SVGRenderer.js"]],function(a){function b(a,d,b,c){return[["M",
a,d+c/2],["L",a+b,d],["L",a,d+c/2],["L",a+b,d+c]]}function f(a,b,e,c){return[["M",a+e,b],["L",a,b+c/2],["L",a+e,b+c],["Z"]]}function g(a,b,e,c){return f(a,b,e/2,c)}a=a.prototype.symbols;a.arrow=b;a["arrow-filled"]=f;a["arrow-filled-half"]=g;a["arrow-half"]=function(a,d,e,c){return b(a,d,e/2,c)};a["triangle-left"]=f;a["triangle-left-half"]=g;return a});b(a,"masters/modules/arrow-symbols.src.js",[],function(){})});
//# sourceMappingURL=arrow-symbols.js.map
```
