## Supplementary material for "Exploring natural odour landscapes: A case study with implications for human-biting insects": Figure S4: boost-canvas.js

```
/*
 Highcharts JS v9.3.1 (2021-11-05)

 Boost module

 (c) 2010-2021 Highsoft AS
 Author: Torstein Honsi

 License: www.highcharts.com/license
*/
'use strict';(function(f){"object"===typeof module&&module.exports?(f["default"]=f,module.exports=f):"function"===typeof define&&define.amd?define("highcharts/modules/boost-canvas",["highcharts"],function(g){f(g);f.Highcharts=g;return f}):f("undefined"!==typeof Highcharts?Highcharts:void 0)})(function(f){function g(f,g,v,y){f.hasOwnProperty(g)||(f[g]=y.apply(null,v))}f=f?f._modules:{};g(f,"Extensions/BoostCanvas.js",[f["Core/Chart/Chart.js"],f["Core/Color/Color.js"],f["Core/Globals.js"],f["Core/Series/Series.js"],
f["Core/Series/SeriesRegistry.js"],f["Core/Utilities.js"]],function(f,g,v,y,C,r){var da=g.parse,D=v.doc,ea=v.noop,m=C.seriesTypes,E=r.addEvent,z=r.extend,fa=r.fireEvent,ha=r.isNumber,ia=r.merge,ja=r.pick,A=r.wrap,O;return function(){v.seriesTypes.heatmap&&A(v.seriesTypes.heatmap.prototype,"drawPoints",function(){var a=this.chart,b=this.getContext(),e=this.chart.inverted,f=this.xAxis,c=this.yAxis;b?(this.points.forEach(function(d){var k=d.plotY;if("undefined"!==typeof k&&!isNaN(k)&&null!==d.y&&b){var l=
d.shapeArgs||{};k=l.x;k=void 0===k?0:k;var g=l.y;g=void 0===g?0:g;var u=l.width;u=void 0===u?0:u;l=l.height;l=void 0===l?0:l;d=a.styledMode?d.series.colorAttribs(d):d.series.pointAttribs(d);b.fillStyle=d.fill;e?b.fillRect(c.len-g+f.left,f.len-k+c.top,-l,-u):b.fillRect(k+f.left,g+c.top,u,l)}}),this.canvasToSVG()):this.chart.showLoading("Your browser doesn't support HTML5 canvas, <br>please use a modern browser")});z(y.prototype,{getContext:function(){var a=this.chart,b=a.chartWidth,e=a.chartHeight,
f=a.seriesGroup||this.group,c=this,g=function(a,c,e,b,f,d,g){a.call(this,e,c,b,f,d,g)};a.isChartSeriesBoosting()&&(c=a,f=a.seriesGroup);var k=c.ctx;c.canvas||(c.canvas=D.createElement("canvas"),c.renderTarget=a.renderer.image("",0,0,b,e).addClass("highcharts-boost-canvas").add(f),c.ctx=k=c.canvas.getContext("2d"),a.inverted&&["moveTo","lineTo","rect","arc"].forEach(function(a){A(k,a,g)}),c.boostCopy=function(){c.renderTarget.attr({href:c.canvas.toDataURL("image/png")})},c.boostClear=function(){k.clearRect(0,
0,c.canvas.width,c.canvas.height);c===this&&c.renderTarget.attr({href:""})},c.boostClipRect=a.renderer.clipRect(),c.renderTarget.clip(c.boostClipRect));c.canvas.width!==b&&(c.canvas.width=b);c.canvas.height!==e&&(c.canvas.height=e);c.renderTarget.attr({x:0,y:0,width:b,height:e,style:"pointer-events: none",href:""});c.boostClipRect.attr(a.getBoostClipRect(c));return k},canvasToSVG:function(){this.chart.isChartSeriesBoosting()?this.boostClear&&this.boostClear():(this.boostCopy||this.chart.boostCopy)&&
(this.boostCopy||this.chart.boostCopy)()},cvsLineTo:function(a,b,e){a.lineTo(b,e)},renderCanvas:function(){var a=this,b=a.options,e=a.chart,f=this.xAxis,c=this.yAxis,m=(e.options.boost||{}).timeRendering||!1,k=0,l=a.processedXData,y=a.processedYData,u=b.data,n=f.getExtremes(),F=n.min,G=n.max;n=c.getExtremes();var A=n.min,C=n.max,P={},H,D=!!a.sampling,I=b.marker&&b.marker.radius,Q=this.cvsDrawPoint,J=b.lineWidth?this.cvsLineTo:void 0,R=I&&1>=I?this.cvsMarkerSquare:this.cvsMarkerCircle,ka=this.cvsStrokeBatch||
1E3,la=!1!==b.enableMouseTracking,S;n=b.threshold;var w=c.getThreshold(n),T=ha(n),U=w,ma=this.fill,V=a.pointArrayMap&&"low,high"===a.pointArrayMap.join(","),W=!!b.stacking,X=a.cropStart||0;n=e.options.loading;var na=a.requireSorting,Y,oa=b.connectNulls,Z=!l,K,L,x,B,M,t=W?a.data:l||u,pa=a.fillOpacity?g.parse(a.color).setOpacity(ja(b.fillOpacity,.75)).get():a.color,aa=function(){ma?(p.fillStyle=pa,p.fill()):(p.strokeStyle=a.color,p.lineWidth=b.lineWidth,p.stroke())},ba=function(c,b,f,d){0===k&&(p.beginPath(),
J&&(p.lineJoin="round"));e.scroller&&"highcharts-navigator-series"===a.options.className?(b+=e.scroller.top,f&&(f+=e.scroller.top)):b+=e.plotTop;c+=e.plotLeft;Y?p.moveTo(c,b):Q?Q(p,c,b,f,S):J?J(p,c,b):R&&R.call(a,p,c,b,I,d);k+=1;k===ka&&(aa(),k=0);S={clientX:c,plotY:b,yBottom:f}},qa="x"===b.findNearestPointBy,ca=this.xData||this.options.xData||this.processedXData||!1,N=function(a,b,d){M=qa?a:a+","+b;la&&!P[M]&&(P[M]=!0,e.inverted&&(a=f.len-a,b=c.len-b),ra.push({x:ca?ca[X+d]:!1,clientX:a,plotX:a,plotY:b,
i:X+d}))};this.renderTarget&&this.renderTarget.attr({href:""});(this.points||this.graph)&&this.destroyGraphics();a.plotGroup("group","series",a.visible?"visible":"hidden",b.zIndex,e.seriesGroup);a.markerGroup=a.group;E(a,"destroy",function(){a.markerGroup=null});var ra=this.points=[];var p=this.getContext();a.buildKDTree=ea;this.boostClear&&this.boostClear();this.visible&&(99999<u.length&&(e.options.loading=ia(n,{labelStyle:{backgroundColor:da("#ffffff").setOpacity(.75).get(),padding:"1em",borderRadius:"0.5em"},
style:{backgroundColor:"none",opacity:1}}),r.clearTimeout(O),e.showLoading("Drawing..."),e.options.loading=n),m&&console.time("canvas rendering"),v.eachAsync(t,function(b,d){var g=!1,k=!1,l=!1,m=!1,n="undefined"===typeof e.index,p=!0;if(!n){if(Z){var q=b[0];var h=b[1];t[d+1]&&(l=t[d+1][0]);t[d-1]&&(m=t[d-1][0])}else q=b,h=y[d],t[d+1]&&(l=t[d+1]),t[d-1]&&(m=t[d-1]);l&&l>=F&&l<=G&&(g=!0);m&&m>=F&&m<=G&&(k=!0);if(V){Z&&(h=b.slice(1,3));var r=h[0];h=h[1]}else W&&(q=b.x,h=b.stackY,r=h-b.y);b=null===h;
na||(p=h>=A&&h<=C);if(!b&&(q>=F&&q<=G&&p||g||k))if(q=Math.round(f.toPixels(q,!0)),D){if("undefined"===typeof x||q===H){V||(r=h);if("undefined"===typeof B||h>L)L=h,B=d;if("undefined"===typeof x||r<K)K=r,x=d}q!==H&&("undefined"!==typeof x&&(h=c.toPixels(L,!0),w=c.toPixels(K,!0),ba(q,T?Math.min(h,U):h,T?Math.max(w,U):w,d),N(q,h,B),w!==h&&N(q,w,x)),x=B=void 0,H=q)}else h=Math.round(c.toPixels(h,!0)),ba(q,h,w,d),N(q,h,d);Y=b&&!oa;0===d%5E4&&(a.boostCopy||a.chart.boostCopy)&&(a.boostCopy||a.chart.boostCopy)()}return!n},
function(){var b=e.loadingDiv,c=e.loadingShown;aa();a.canvasToSVG();m&&console.timeEnd("canvas rendering");fa(a,"renderedCanvas");c&&(z(b.style,{transition:"opacity 250ms",opacity:0}),e.loadingShown=!1,O=setTimeout(function(){b.parentNode&&b.parentNode.removeChild(b);e.loadingDiv=e.loadingSpan=null},250));delete a.buildKDTree;a.buildKDTree()},e.renderer.forExport?Number.MAX_VALUE:void 0))}});m.scatter.prototype.cvsMarkerCircle=function(a,b,e,d){a.moveTo(b,e);a.arc(b,e,d,0,2*Math.PI,!1)};m.scatter.prototype.cvsMarkerSquare=
function(a,b,e,d){a.rect(b-d,e-d,2*d,2*d)};m.scatter.prototype.fill=!0;m.bubble&&(m.bubble.prototype.cvsMarkerCircle=function(a,b,e,d,c){a.moveTo(b,e);a.arc(b,e,this.radii&&this.radii[c],0,2*Math.PI,!1)},m.bubble.prototype.cvsStrokeBatch=1);z(m.area.prototype,{cvsDrawPoint:function(a,b,e,d,c){c&&b!==c.clientX&&(a.moveTo(c.clientX,c.yBottom),a.lineTo(c.clientX,c.plotY),a.lineTo(b,e),a.lineTo(b,d))},fill:!0,fillOpacity:!0,sampling:!0});z(m.column.prototype,{cvsDrawPoint:function(a,b,e,d){a.rect(b-1,
e,1,d-e)},fill:!0,sampling:!0});f.prototype.callbacks.push(function(a){E(a,"predraw",function(){a.renderTarget&&a.renderTarget.attr({href:""});a.canvas&&a.canvas.getContext("2d").clearRect(0,0,a.canvas.width,a.canvas.height)});E(a,"render",function(){a.boostCopy&&a.boostCopy()})})}});g(f,"masters/modules/boost-canvas.src.js",[],function(){})});
//# sourceMappingURL=boost-canvas.js.map
```
