## Supplementary material for "Exploring natural odour landscapes: A case study with implications for human-biting insects": Figure S4: boost.js

```
/*
 Highcharts JS v9.3.1 (2021-11-05)

 Boost module

 (c) 2010-2021 Highsoft AS
 Author: Torstein Honsi

 License: www.highcharts.com/license

*/
'use strict';(function(a){"object"===typeof module&&module.exports?(a["default"]=a,module.exports=a):"function"===typeof define&&define.amd?define("highcharts/modules/boost",["highcharts"],function(u){a(u);a.Highcharts=u;return a}):a("undefined"!==typeof Highcharts?Highcharts:void 0)})(function(a){function u(a,z,v,k){a.hasOwnProperty(z)||(a[z]=k.apply(null,v))}a=a?a._modules:{};u(a,"Extensions/Boost/Boostables.js",[],function(){return"area arearange column columnrange bar line scatter heatmap bubble treemap".split(" ")});
u(a,"Extensions/Boost/BoostableMap.js",[a["Extensions/Boost/Boostables.js"]],function(a){var q={};a.forEach(function(a){q[a]=1});return q});u(a,"Extensions/Boost/WGLShader.js",[a["Core/Utilities.js"]],function(a){var q=a.clamp,v=a.error,k=a.pick;return function(d){function a(){t.length&&v("[highcharts boost] shader error - "+t.join("\n"))}function C(b,c){var h=d.createShader("vertex"===c?d.VERTEX_SHADER:d.FRAGMENT_SHADER);d.shaderSource(h,b);d.compileShader(h);return d.getShaderParameter(h,d.COMPILE_STATUS)?
h:(t.push("when compiling "+c+" shader:\n"+d.getShaderInfoLog(h)),!1)}function p(){function b(b){return d.getUniformLocation(f,b)}var O=C("#version 100\n#define LN10 2.302585092994046\nprecision highp float;\nattribute vec4 aVertexPosition;\nattribute vec4 aColor;\nvarying highp vec2 position;\nvarying highp vec4 vColor;\nuniform mat4 uPMatrix;\nuniform float pSize;\nuniform float translatedThreshold;\nuniform bool hasThreshold;\nuniform bool skipTranslation;\nuniform float xAxisTrans;\nuniform float xAxisMin;\nuniform float xAxisMinPad;\nuniform float xAxisPointRange;\nuniform float xAxisLen;\nuniform bool  xAxisPostTranslate;\nuniform float xAxisOrdinalSlope;\nuniform float xAxisOrdinalOffset;\nuniform float xAxisPos;\nuniform bool  xAxisCVSCoord;\nuniform bool  xAxisIsLog;\nuniform bool  xAxisReversed;\nuniform float yAxisTrans;\nuniform float yAxisMin;\nuniform float yAxisMinPad;\nuniform float yAxisPointRange;\nuniform float yAxisLen;\nuniform bool  yAxisPostTranslate;\nuniform float yAxisOrdinalSlope;\nuniform float yAxisOrdinalOffset;\nuniform float yAxisPos;\nuniform bool  yAxisCVSCoord;\nuniform bool  yAxisIsLog;\nuniform bool  yAxisReversed;\nuniform bool  isBubble;\nuniform bool  bubbleSizeByArea;\nuniform float bubbleZMin;\nuniform float bubbleZMax;\nuniform float bubbleZThreshold;\nuniform float bubbleMinSize;\nuniform float bubbleMaxSize;\nuniform bool  bubbleSizeAbs;\nuniform bool  isInverted;\nfloat bubbleRadius(){\nfloat value = aVertexPosition.w;\nfloat zMax = bubbleZMax;\nfloat zMin = bubbleZMin;\nfloat radius = 0.0;\nfloat pos = 0.0;\nfloat zRange = zMax - zMin;\nif (bubbleSizeAbs){\nvalue = value - bubbleZThreshold;\nzMax = max(zMax - bubbleZThreshold, zMin - bubbleZThreshold);\nzMin = 0.0;\n}\nif (value < zMin){\nradius = bubbleZMin / 2.0 - 1.0;\n} else {\npos = zRange > 0.0 ? (value - zMin) / zRange : 0.5;\nif (bubbleSizeByArea && pos > 0.0){\npos = sqrt(pos);\n}\nradius = ceil(bubbleMinSize + pos * (bubbleMaxSize - bubbleMinSize)) / 2.0;\n}\nreturn radius * 2.0;\n}\nfloat translate(float val,\nfloat pointPlacement,\nfloat localA,\nfloat localMin,\nfloat minPixelPadding,\nfloat pointRange,\nfloat len,\nbool  cvsCoord,\nbool  isLog,\nbool  reversed\n){\nfloat sign = 1.0;\nfloat cvsOffset = 0.0;\nif (cvsCoord) {\nsign *= -1.0;\ncvsOffset = len;\n}\nif (isLog) {\nval = log(val) / LN10;\n}\nif (reversed) {\nsign *= -1.0;\ncvsOffset -= sign * len;\n}\nreturn sign * (val - localMin) * localA + cvsOffset + \n(sign * minPixelPadding);\n}\nfloat xToPixels(float value) {\nif (skipTranslation){\nreturn value;// + xAxisPos;\n}\nreturn translate(value, 0.0, xAxisTrans, xAxisMin, xAxisMinPad, xAxisPointRange, xAxisLen, xAxisCVSCoord, xAxisIsLog, xAxisReversed);// + xAxisPos;\n}\nfloat yToPixels(float value, float checkTreshold) {\nfloat v;\nif (skipTranslation){\nv = value;// + yAxisPos;\n} else {\nv = translate(value, 0.0, yAxisTrans, yAxisMin, yAxisMinPad, yAxisPointRange, yAxisLen, yAxisCVSCoord, yAxisIsLog, yAxisReversed);// + yAxisPos;\nif (v > yAxisLen) {\nv = yAxisLen;\n}\n}\nif (checkTreshold > 0.0 && hasThreshold) {\nv = min(v, translatedThreshold);\n}\nreturn v;\n}\nvoid main(void) {\nif (isBubble){\ngl_PointSize = bubbleRadius();\n} else {\ngl_PointSize = pSize;\n}\nvColor = aColor;\nif (skipTranslation && isInverted) {\ngl_Position = uPMatrix * vec4(aVertexPosition.y + yAxisPos, aVertexPosition.x + xAxisPos, 0.0, 1.0);\n} else if (isInverted) {\ngl_Position = uPMatrix * vec4(yToPixels(aVertexPosition.y, aVertexPosition.z) + yAxisPos, xToPixels(aVertexPosition.x) + xAxisPos, 0.0, 1.0);\n} else {\ngl_Position = uPMatrix * vec4(xToPixels(aVertexPosition.x) + xAxisPos, yToPixels(aVertexPosition.y, aVertexPosition.z) + yAxisPos, 0.0, 1.0);\n}\n}",
"vertex"),p=C("precision highp float;\nuniform vec4 fillColor;\nvarying highp vec2 position;\nvarying highp vec4 vColor;\nuniform sampler2D uSampler;\nuniform bool isCircle;\nuniform bool hasColor;\nvoid main(void) {\nvec4 col = fillColor;\nvec4 tcol = texture2D(uSampler, gl_PointCoord.st);\nif (hasColor) {\ncol = vColor;\n}\nif (isCircle) {\ncol *= tcol;\nif (tcol.r < 0.0) {\ndiscard;\n} else {\ngl_FragColor = col;\n}\n} else {\ngl_FragColor = col;\n}\n}","fragment");if(!O||!p)return f=!1,a(),!1;
f=d.createProgram();d.attachShader(f,O);d.attachShader(f,p);d.linkProgram(f);if(!d.getProgramParameter(f,d.LINK_STATUS))return t.push(d.getProgramInfoLog(f)),a(),f=!1;d.useProgram(f);d.bindAttribLocation(f,0,"aVertexPosition");e=b("uPMatrix");m=b("pSize");l=b("fillColor");K=b("isBubble");L=b("bubbleSizeAbs");D=b("bubbleSizeByArea");M=b("uSampler");H=b("skipTranslation");c=b("isCircle");h=b("isInverted");return!0}function y(b,c){d&&f&&(b=n[b]=n[b]||d.getUniformLocation(f,b),d.uniform1f(b,c))}var n=
{},f,e,m,l,K,L,D,H,c,h,t=[],M;return d&&!p()?!1:{psUniform:function(){return m},pUniform:function(){return e},fillColorUniform:function(){return l},setBubbleUniforms:function(b,h,e){var g=b.options,t=Number.MAX_VALUE,a=-Number.MAX_VALUE;if(d&&f&&b.is("bubble")){var M=b.getPxExtremes();t=k(g.zMin,q(h,!1===g.displayNegative?g.zThreshold:-Number.MAX_VALUE,t));a=k(g.zMax,Math.max(a,e));d.uniform1i(K,1);d.uniform1i(c,1);d.uniform1i(D,"width"!==b.options.sizeBy);d.uniform1i(L,b.options.sizeByAbsoluteValue);
y("bubbleZMin",t);y("bubbleZMax",a);y("bubbleZThreshold",b.options.zThreshold);y("bubbleMinSize",M.minPxSize);y("bubbleMaxSize",M.maxPxSize)}},bind:function(){d&&f&&d.useProgram(f)},program:function(){return f},create:p,setUniform:y,setPMatrix:function(b){d&&f&&d.uniformMatrix4fv(e,!1,b)},setColor:function(b){d&&f&&d.uniform4f(l,b[0]/255,b[1]/255,b[2]/255,b[3])},setPointSize:function(b){d&&f&&d.uniform1f(m,b)},setSkipTranslation:function(b){d&&f&&d.uniform1i(H,!0===b?1:0)},setTexture:function(b){d&&
f&&d.uniform1i(M,b)},setDrawAsCircle:function(b){d&&f&&d.uniform1i(c,b?1:0)},reset:function(){d&&f&&(d.uniform1i(K,0),d.uniform1i(c,0))},setInverted:function(b){d&&f&&d.uniform1i(h,b)},destroy:function(){d&&f&&(d.deleteProgram(f),f=!1)}}}});u(a,"Extensions/Boost/WGLVBuffer.js",[],function(){return function(a,z,v){function k(){d&&(a.deleteBuffer(d),r=d=!1);y=0;q=v||2;n=[]}var d=!1,r=!1,q=v||2,p=!1,y=0,n;return{destroy:k,bind:function(){if(!d)return!1;a.vertexAttribPointer(r,q,a.FLOAT,!1,0,0)},data:n,
build:function(f,e,m){var l;n=f||[];if(!(n&&0!==n.length||p))return k(),!1;q=m||q;d&&a.deleteBuffer(d);p||(l=new Float32Array(n));d=a.createBuffer();a.bindBuffer(a.ARRAY_BUFFER,d);a.bufferData(a.ARRAY_BUFFER,p||l,a.STATIC_DRAW);r=a.getAttribLocation(z.program(),e);a.enableVertexAttribArray(r);return!0},render:function(f,e,m){var l=p?p.length:n.length;if(!d||!l)return!1;if(!f||f>l||0>f)f=0;if(!e||e>l)e=l;if(f>=e)return!1;a.drawArrays(a[(m||"points").toUpperCase()],f/q,(e-f)/q);return!0},allocate:function(a){y=
-1;p=new Float32Array(4*a)},push:function(a,e,d,l){p&&(p[++y]=a,p[++y]=e,p[++y]=d,p[++y]=l)}}}});u(a,"Extensions/Boost/WGLRenderer.js",[a["Core/Color/Color.js"],a["Extensions/Boost/WGLShader.js"],a["Extensions/Boost/WGLVBuffer.js"],a["Core/Globals.js"],a["Core/Utilities.js"]],function(a,z,v,k,d){var r=a.parse,q=k.doc,p=d.isNumber,y=d.isObject,n=d.merge,f=d.objectEach,e=d.pick;return function(d){function m(b){if(b.isSeriesBoosting){var c=!!b.options.stacking;var h=b.xData||b.options.xData||b.processedXData;
c=(c?b.data:h||b.options.data).length;"treemap"===b.type?c*=12:"heatmap"===b.type?c*=6:ca[b.type]&&(c*=2);return c}return 0}function K(){g.clear(g.COLOR_BUFFER_BIT|g.DEPTH_BUFFER_BIT)}function L(b,c){function h(b){b&&(c.colorData.push(b[0]),c.colorData.push(b[1]),c.colorData.push(b[2]),c.colorData.push(b[3]))}function a(b,c,a,g,e){h(e);w.usePreallocated?(O.push(b,c,a?1:0,g||1),qa+=4):(B.push(b),B.push(c),B.push(a?1:0),B.push(g||1))}function g(){c.segments.length&&(c.segments[c.segments.length-1].to=
B.length||qa)}function e(){c.segments.length&&c.segments[c.segments.length-1].from===(B.length||qa)||(g(),c.segments.push({from:B.length||qa}))}function d(b,c,g,e,d){h(d);a(b+g,c);h(d);a(b,c);h(d);a(b,c+e);h(d);a(b,c+e);h(d);a(b+g,c+e);h(d);a(b+g,c)}function t(b,h){w.useGPUTranslations||(c.skipTranslation=!0,b.x=Q.toPixels(b.x,!0),b.y=v.toPixels(b.y,!0));h?B=[b.x,b.y,0,2].concat(B):a(b.x,b.y,0,2)}var M=b.pointArrayMap&&"low,high"===b.pointArrayMap.join(","),m=b.chart,x=b.options,S=!!x.stacking,f=
x.data,l=b.xAxis.getExtremes(),p=l.min,n=l.max;l=b.yAxis.getExtremes();var k=l.min,K=l.max,q=b.xData||x.xData||b.processedXData,D=b.yData||x.yData||b.processedYData,L=b.zData||x.zData||b.processedZData,v=b.yAxis,Q=b.xAxis,H=b.chart.plotWidth,R=!q||0===q.length,z=x.connectNulls;l=b.points||!1;var P=!1,u=!1,C,G,E=S?b.data:q||f,ma={x:Number.MAX_VALUE,y:0},na={x:-Number.MAX_VALUE,y:0},Ia=0,Y=!1,F,A,N,U,J=-1,aa=!1,ha=!1,X,Ra="undefined"===typeof m.index,ba=!1,Ba=!1,I=!1,Pa=ca[b.type],Ca=!1,Ja=!0,Ka=!0,
Qa=x.zoneAxis||"y",oa=x.zones||!1,T=!1,La=x.threshold,Da=!1;if(!(x.boostData&&0<x.boostData.length)){x.gapSize&&(Da="value"!==x.gapUnit?x.gapSize*b.closestPointRange:x.gapSize);if(oa){var pa=[];oa.forEach(function(b,c){if(b.color){var h=r(b.color).rgba;h[0]/=255;h[1]/=255;h[2]/=255;pa[c]=h;T||"undefined"!==typeof b.value||(T=h)}});T||(q=b.pointAttribs&&b.pointAttribs().fill||b.color,T=r(q).rgba,T[0]/=255,T[1]/=255,T[2]/=255)}m.inverted&&(H=b.chart.plotHeight);b.closestPointRangePx=Number.MAX_VALUE;
e();if(l&&0<l.length)c.skipTranslation=!0,c.drawMode="triangles",l[0].node&&l[0].node.levelDynamic&&l.sort(function(b,c){if(b.node){if(b.node.levelDynamic>c.node.levelDynamic)return 1;if(b.node.levelDynamic<c.node.levelDynamic)return-1}return 0}),l.forEach(function(c){var h=c.plotY;if("undefined"!==typeof h&&!isNaN(h)&&null!==c.y&&c.shapeArgs){var a=c.shapeArgs;h=a.x;h=void 0===h?0:h;var g=a.y;g=void 0===g?0:g;var e=a.width;e=void 0===e?0:e;a=a.height;a=void 0===a?0:a;var t=m.styledMode?c.series.colorAttribs(c):
t=c.series.pointAttribs(c);c=t["stroke-width"]||0;I=r(t.fill).rgba;I[0]/=255;I[1]/=255;I[2]/=255;"treemap"===b.type&&(c=c||1,G=r(t.stroke).rgba,G[0]/=255,G[1]/=255,G[2]/=255,d(h,g,e,a,G),c/=2);"heatmap"===b.type&&m.inverted&&(h=Q.len-h,g=v.len-g,e=-e,a=-a);d(h+c,g+c,e-2*c,a-2*c,I)}});else{for(l=function(){N=E[++J];if("undefined"===typeof N)return"continue";if(Ra)return"break";var h=f&&f[J];!R&&y(h,!0)&&h.color&&(I=r(h.color).rgba,I[0]/=255,I[1]/=255,I[2]/=255);R?(F=N[0],A=N[1],E[J+1]&&(ha=E[J+1][0]),
E[J-1]&&(aa=E[J-1][0]),3<=N.length&&(U=N[2],N[2]>c.zMax&&(c.zMax=N[2]),N[2]<c.zMin&&(c.zMin=N[2]))):(F=N,A=D[J],E[J+1]&&(ha=E[J+1]),E[J-1]&&(aa=E[J-1]),L&&L.length&&(U=L[J],L[J]>c.zMax&&(c.zMax=L[J]),L[J]<c.zMin&&(c.zMin=L[J])));if(!z&&(null===F||null===A))return e(),"continue";ha&&ha>=p&&ha<=n&&(ba=!0);aa&&aa>=p&&aa<=n&&(Ba=!0);M?(R&&(A=N.slice(1,3)),X=A[0],A=A[1]):S&&(F=N.x,A=N.stackY,X=A-N.y);null!==k&&"undefined"!==typeof k&&null!==K&&"undefined"!==typeof K&&(Ja=A>=k&&A<=K);F>n&&na.x<n&&(na.x=
F,na.y=A);F<p&&ma.x>p&&(ma.x=F,ma.y=A);if(null===A&&z)return"continue";if(null===A||!Ja&&!ba&&!Ba)return e(),"continue";(ha>=p||F>=p)&&(aa<=n||F<=n)&&(Ca=!0);if(!Ca&&!ba&&!Ba)return"continue";Da&&F-aa>Da&&e();if(oa){var g;oa.some(function(b,c){var h=oa[c-1];return"x"===Qa?"undefined"!==typeof b.value&&F<=b.value?(pa[c]&&(!h||F>=h.value)&&(g=pa[c]),!0):!1:"undefined"!==typeof b.value&&A<=b.value?(pa[c]&&(!h||A>=h.value)&&(g=pa[c]),!0):!1});I=g||T||I}if(!w.useGPUTranslations&&(c.skipTranslation=!0,
F=Q.toPixels(F,!0),A=v.toPixels(A,!0),F>H&&"points"===c.drawMode))return"continue";c.hasMarkers&&Ca&&!1!==P&&(b.closestPointRangePx=Math.min(b.closestPointRangePx,Math.abs(F-P)));if(!w.useGPUTranslations&&!w.usePreallocated&&P&&1>Math.abs(F-P)&&u&&1>Math.abs(A-u))return w.debug.showSkipSummary&&++Ia,"continue";if(Pa){C=X;if(!1===X||"undefined"===typeof X)C=0>A?A:0;M||S||(C=Math.max(null===La?k:La,k));w.useGPUTranslations||(C=v.toPixels(C,!0));a(F,C,0,0,I)}x.step&&!Ka&&a(F,u,0,2,I);a(F,A,0,"bubble"===
b.type?U||1:2,I);P=F;u=A;Y=!0;Ka=!1};J<E.length-1&&"break"!==l(););w.debug.showSkipSummary&&console.log("skipped points:",Ia);Y||!1===z||"line_strip"!==b.drawMode||(ma.x<Number.MAX_VALUE&&t(ma,!0),na.x>-Number.MAX_VALUE&&t(na))}g()}}function D(){G=[];C.data=B=[];u=[];O&&O.destroy()}function H(c){b&&(b.setUniform("xAxisTrans",c.transA),b.setUniform("xAxisMin",c.min),b.setUniform("xAxisMinPad",c.minPixelPadding),b.setUniform("xAxisPointRange",c.pointRange),b.setUniform("xAxisLen",c.len),b.setUniform("xAxisPos",
c.pos),b.setUniform("xAxisCVSCoord",!c.horiz),b.setUniform("xAxisIsLog",!!c.logarithmic),b.setUniform("xAxisReversed",!!c.reversed))}function c(c){b&&(b.setUniform("yAxisTrans",c.transA),b.setUniform("yAxisMin",c.min),b.setUniform("yAxisMinPad",c.minPixelPadding),b.setUniform("yAxisPointRange",c.pointRange),b.setUniform("yAxisLen",c.len),b.setUniform("yAxisPos",c.pos),b.setUniform("yAxisCVSCoord",!c.horiz),b.setUniform("yAxisIsLog",!!c.logarithmic),b.setUniform("yAxisReversed",!!c.reversed))}function h(c,
h){b.setUniform("hasThreshold",c);b.setUniform("translatedThreshold",h)}function t(t){if(t)R=t.chartWidth||800,P=t.chartHeight||400;else return!1;if(!(g&&R&&P&&b))return!1;w.debug.timeRendering&&console.time("gl rendering");g.canvas.width=R;g.canvas.height=P;b.bind();g.viewport(0,0,R,P);b.setPMatrix([2/R,0,0,0,0,-(2/P),0,0,0,0,-2,0,-1,1,-1,1]);1<w.lineWidth&&!k.isMS&&g.lineWidth(w.lineWidth);O.build(C.data,"aVertexPosition",4);O.bind();b.setInverted(t.inverted);G.forEach(function(d,M){var m=d.series.options,
l=m.marker;var f="undefined"!==typeof m.lineWidth?m.lineWidth:1;var x=m.threshold,n=p(x),k=d.series.yAxis.getThreshold(x);x=e(m.marker?m.marker.enabled:null,d.series.xAxis.isRadial?!0:null,d.series.closestPointRangePx>2*((m.marker?m.marker.radius:10)||10));l=U[l&&l.symbol||d.series.symbol]||U.circle;if(0!==d.segments.length&&d.segments[0].from!==d.segments[0].to){l.isReady&&(g.bindTexture(g.TEXTURE_2D,l.handle),b.setTexture(l.handle));t.styledMode?l=d.series.markerGroup&&d.series.markerGroup.getStyle("fill"):
(l="points"===d.drawMode&&d.series.pointAttribs&&d.series.pointAttribs().fill||d.series.color,m.colorByPoint&&(l=d.series.chart.options.colors[M]));d.series.fillOpacity&&m.fillOpacity&&(l=(new a(l)).setOpacity(e(m.fillOpacity,1)).get());l=r(l).rgba;w.useAlpha||(l[3]=1);"lines"===d.drawMode&&w.useAlpha&&1>l[3]&&(l[3]/=10);"add"===m.boostBlending?(g.blendFunc(g.SRC_ALPHA,g.ONE),g.blendEquation(g.FUNC_ADD)):"mult"===m.boostBlending||"multiply"===m.boostBlending?g.blendFunc(g.DST_COLOR,g.ZERO):"darken"===
m.boostBlending?(g.blendFunc(g.ONE,g.ONE),g.blendEquation(g.FUNC_MIN)):g.blendFuncSeparate(g.SRC_ALPHA,g.ONE_MINUS_SRC_ALPHA,g.ONE,g.ONE_MINUS_SRC_ALPHA);b.reset();0<d.colorData.length?(b.setUniform("hasColor",1),M=v(g,b),M.build(d.colorData,"aColor",4),M.bind()):g.disableVertexAttribArray(g.getAttribLocation(b.program(),"aColor"));b.setColor(l);H(d.series.xAxis);c(d.series.yAxis);h(n,k);"points"===d.drawMode&&(m.marker&&p(m.marker.radius)?b.setPointSize(2*m.marker.radius):b.setPointSize(1));b.setSkipTranslation(d.skipTranslation);
"bubble"===d.series.type&&b.setBubbleUniforms(d.series,d.zMin,d.zMax);b.setDrawAsCircle(ba[d.series.type]||!1);if(0<f||"line_strip"!==d.drawMode)for(f=0;f<d.segments.length;f++)O.render(d.segments[f].from,d.segments[f].to,d.drawMode);if(d.hasMarkers&&x)for(m.marker&&p(m.marker.radius)?b.setPointSize(2*m.marker.radius):b.setPointSize(10),b.setDrawAsCircle(!0),f=0;f<d.segments.length;f++)O.render(d.segments[f].from,d.segments[f].to,"POINTS")}});w.debug.timeRendering&&console.timeEnd("gl rendering");
d&&d();D()}function M(b){K();if(b.renderer.forExport)return t(b);Y?t(b):setTimeout(function(){M(b)},1)}var b=!1,O=!1,qa=0,g=!1,R=0,P=0,B=!1,u=!1,C={},Y=!1,G=[],U={},ca={column:!0,columnrange:!0,bar:!0,area:!0,arearange:!0},ba={scatter:!0,bubble:!0},w={pointSize:1,lineWidth:1,fillColor:"#AA00AA",useAlpha:!0,usePreallocated:!1,useGPUTranslations:!1,debug:{timeRendering:!1,timeSeriesProcessing:!1,timeSetup:!1,timeBufferCopy:!1,timeKDTree:!1,showSkipSummary:!1}};return C={allocateBufferForSingleSeries:function(b){var c=
0;w.usePreallocated&&(b.isSeriesBoosting&&(c=m(b)),O.allocate(c))},pushSeries:function(b){0<G.length&&G[G.length-1].hasMarkers&&(G[G.length-1].markerTo=u.length);w.debug.timeSeriesProcessing&&console.time("building "+b.type+" series");var c={segments:[],markerFrom:u.length,colorData:[],series:b,zMin:Number.MAX_VALUE,zMax:-Number.MAX_VALUE,hasMarkers:b.options.marker?!1!==b.options.marker.enabled:!1,showMarkers:!0,drawMode:{area:"lines",arearange:"lines",areaspline:"line_strip",column:"lines",columnrange:"lines",
bar:"lines",line:"line_strip",scatter:"points",heatmap:"triangles",treemap:"triangles",bubble:"points"}[b.type]||"line_strip"};b.index>=G.length?G.push(c):G[b.index]=c;L(b,c);w.debug.timeSeriesProcessing&&console.timeEnd("building "+b.type+" series")},setSize:function(c,h){R===c&&P===h||!b||(R=c,P=h,b.bind(),b.setPMatrix([2/R,0,0,0,0,-(2/P),0,0,0,0,-2,0,-1,1,-1,1]))},inited:function(){return Y},setThreshold:h,init:function(c,h){function d(b,c){var h={isReady:!1,texture:q.createElement("canvas"),handle:g.createTexture()},
d=h.texture.getContext("2d");U[b]=h;h.texture.width=512;h.texture.height=512;d.mozImageSmoothingEnabled=!1;d.webkitImageSmoothingEnabled=!1;d.msImageSmoothingEnabled=!1;d.imageSmoothingEnabled=!1;d.strokeStyle="rgba(255, 255, 255, 0)";d.fillStyle="#FFF";c(d);try{g.activeTexture(g.TEXTURE0),g.bindTexture(g.TEXTURE_2D,h.handle),g.texImage2D(g.TEXTURE_2D,0,g.RGBA,g.RGBA,g.UNSIGNED_BYTE,h.texture),g.texParameteri(g.TEXTURE_2D,g.TEXTURE_WRAP_S,g.CLAMP_TO_EDGE),g.texParameteri(g.TEXTURE_2D,g.TEXTURE_WRAP_T,
g.CLAMP_TO_EDGE),g.texParameteri(g.TEXTURE_2D,g.TEXTURE_MAG_FILTER,g.LINEAR),g.texParameteri(g.TEXTURE_2D,g.TEXTURE_MIN_FILTER,g.LINEAR),g.bindTexture(g.TEXTURE_2D,null),h.isReady=!0}catch(W){}}var a=0,e=["webgl","experimental-webgl","moz-webgl","webkit-3d"];Y=!1;if(!c)return!1;for(w.debug.timeSetup&&console.time("gl setup");a<e.length&&!(g=c.getContext(e[a],{}));a++);if(g)h||D();else return!1;g.enable(g.BLEND);g.blendFunc(g.SRC_ALPHA,g.ONE_MINUS_SRC_ALPHA);g.disable(g.DEPTH_TEST);g.depthFunc(g.LESS);
b=z(g);if(!b)return!1;O=v(g,b);d("circle",function(b){b.beginPath();b.arc(256,256,256,0,2*Math.PI);b.stroke();b.fill()});d("square",function(b){b.fillRect(0,0,512,512)});d("diamond",function(b){b.beginPath();b.moveTo(256,0);b.lineTo(512,256);b.lineTo(256,512);b.lineTo(0,256);b.lineTo(256,0);b.fill()});d("triangle",function(b){b.beginPath();b.moveTo(0,512);b.lineTo(256,0);b.lineTo(512,512);b.lineTo(0,512);b.fill()});d("triangle-down",function(b){b.beginPath();b.moveTo(0,0);b.lineTo(256,512);b.lineTo(512,
0);b.lineTo(0,0);b.fill()});Y=!0;w.debug.timeSetup&&console.timeEnd("gl setup");return!0},render:M,settings:w,valid:function(){return!1!==g},clear:K,flush:D,setXAxis:H,setYAxis:c,data:B,gl:function(){return g},allocateBuffer:function(b){var c=0;w.usePreallocated&&(b.series.forEach(function(b){b.isSeriesBoosting&&(c+=m(b))}),O.allocate(c))},destroy:function(){D();O.destroy();b.destroy();g&&(f(U,function(b){b.handle&&g.deleteTexture(b.handle)}),g.canvas.width=1,g.canvas.height=1)},setOptions:function(b){n(!0,
w,b)}}}});u(a,"Extensions/Boost/BoostAttach.js",[a["Core/Chart/Chart.js"],a["Extensions/Boost/WGLRenderer.js"],a["Core/Globals.js"],a["Core/Utilities.js"]],function(a,z,v,k){var d=v.doc,r=k.error,q;return function(p,k){var n=p.chartWidth,f=p.chartHeight,e=p,m=p.seriesGroup||k.group,l=d.implementation.hasFeature("www.http://w3.org/TR/SVG11/feature#Extensibility","1.1");e=p.isChartSeriesBoosting()?p:k;l=!1;q||(q=d.createElement("canvas"));e.renderTarget||(e.canvas=q,p.renderer.forExport||!l?(e.renderTarget=
p.renderer.image("",0,0,n,f).addClass("highcharts-boost-canvas").add(m),e.boostClear=function(){e.renderTarget.attr({href:""})},e.boostCopy=function(){e.boostResizeTarget();e.renderTarget.attr({href:e.canvas.toDataURL("image/png")})}):(e.renderTargetFo=p.renderer.createElement("foreignObject").add(m),e.renderTarget=d.createElement("canvas"),e.renderTargetCtx=e.renderTarget.getContext("2d"),e.renderTargetFo.element.appendChild(e.renderTarget),e.boostClear=function(){e.renderTarget.width=e.canvas.width;
e.renderTarget.height=e.canvas.height},e.boostCopy=function(){e.renderTarget.width=e.canvas.width;e.renderTarget.height=e.canvas.height;e.renderTargetCtx.drawImage(e.canvas,0,0)}),e.boostResizeTarget=function(){n=p.chartWidth;f=p.chartHeight;(e.renderTargetFo||e.renderTarget).attr({x:0,y:0,width:n,height:f}).css({pointerEvents:"none",mixedBlendMode:"normal",opacity:1});e instanceof a&&e.markerGroup.translate(p.plotLeft,p.plotTop)},e.boostClipRect=p.renderer.clipRect(),(e.renderTargetFo||e.renderTarget).clip(e.boostClipRect),
e instanceof a&&(e.markerGroup=e.renderer.g().add(m),e.markerGroup.translate(k.xAxis.pos,k.yAxis.pos)));e.canvas.width=n;e.canvas.height=f;e.boostClipRect.attr(p.getBoostClipRect(e));e.boostResizeTarget();e.boostClear();e.ogl||(e.ogl=z(function(){e.ogl.settings.debug.timeBufferCopy&&console.time("buffer copy");e.boostCopy();e.ogl.settings.debug.timeBufferCopy&&console.timeEnd("buffer copy")}),e.ogl.init(e.canvas)||r("[highcharts boost] - unable to init WebGL renderer"),e.ogl.setOptions(p.options.boost||
{}),e instanceof a&&e.ogl.allocateBuffer(p));e.ogl.setSize(n,f);return e.ogl}});u(a,"Extensions/Boost/BoostUtils.js",[a["Core/Globals.js"],a["Extensions/Boost/BoostableMap.js"],a["Extensions/Boost/BoostAttach.js"],a["Core/Utilities.js"]],function(a,z,v,k){function d(){for(var d=[],a=0;a<arguments.length;a++)d[a]=arguments[a];var e=-Number.MAX_VALUE;d.forEach(function(d){if("undefined"!==typeof d&&null!==d&&"undefined"!==typeof d.length&&0<d.length)return e=d.length,!0});return e}function r(d,a,e){d&&
a.renderTarget&&a.canvas&&!(e||a.chart).isChartSeriesBoosting()&&d.render(e||a.chart)}function q(d,a){d&&a.renderTarget&&a.canvas&&!a.chart.isChartSeriesBoosting()&&d.allocateBufferForSingleSeries(a)}function p(d,a,e,f,k,q){k=k||0;f=f||3E3;for(var c=k+f,h=!0;h&&k<c&&k<d.length;)h=a(d[k],k),++k;h&&(k<d.length?q?p(d,a,e,f,k,q):n.requestAnimationFrame?n.requestAnimationFrame(function(){p(d,a,e,f,k)}):setTimeout(function(){p(d,a,e,f,k)}):e&&e())}function y(){var d=0,a,e=["webgl","experimental-webgl",
"moz-webgl","webkit-3d"],k=!1;if("undefined"!==typeof n.WebGLRenderingContext)for(a=f.createElement("canvas");d<e.length;d++)try{if(k=a.getContext(e[d]),"undefined"!==typeof k&&null!==k)return!0}catch(D){}return!1}var n=a.win,f=a.doc,e=k.pick;k={patientMax:d,boostEnabled:function(d){return e(d&&d.options&&d.options.boost&&d.options.boost.enabled,!0)},shouldForceChartSeriesBoosting:function(a){var f=0,k=0,m=e(a.options.boost&&a.options.boost.allowForce,!0);if("undefined"!==typeof a.boostForceChartBoost)return a.boostForceChartBoost;
if(1<a.series.length)for(var p=0;p<a.series.length;p++){var n=a.series[p];0!==n.options.boostThreshold&&!1!==n.visible&&"heatmap"!==n.type&&(z[n.type]&&++k,d(n.processedXData,n.options.data,n.points)>=(n.options.boostThreshold||Number.MAX_VALUE)&&++f)}a.boostForceChartBoost=m&&(k===a.series.length&&0<f||5<f);return a.boostForceChartBoost},renderIfNotSeriesBoosting:r,allocateIfNotSeriesBoosting:q,eachAsync:p,hasWebGLSupport:y,pointDrawHandler:function(d){var a=!0;this.chart.options&&this.chart.options.boost&&
(a="undefined"===typeof this.chart.options.boost.enabled?!0:this.chart.options.boost.enabled);if(!a||!this.isSeriesBoosting)return d.call(this);this.chart.isBoosting=!0;if(d=v(this.chart,this))q(d,this),d.pushSeries(this);r(d,this)}};a.hasWebGLSupport=y;return k});u(a,"Extensions/Boost/BoostInit.js",[a["Core/Chart/Chart.js"],a["Core/Globals.js"],a["Core/Series/Series.js"],a["Core/Series/SeriesRegistry.js"],a["Core/Utilities.js"],a["Extensions/Boost/BoostUtils.js"],a["Extensions/Boost/BoostAttach.js"]],
function(a,z,v,k,d,r,u){var p=z.noop,q=k.seriesTypes,n=d.addEvent,f=d.extend,e=d.fireEvent,m=d.wrap,l=r.eachAsync,K=r.pointDrawHandler,L=r.allocateIfNotSeriesBoosting,C=r.renderIfNotSeriesBoosting,H=r.shouldForceChartSeriesBoosting,c;return function(){f(v.prototype,{renderCanvas:function(){function h(b,c){var h=!1,a="undefined"===typeof k.index,d=!0;if("undefined"===typeof b)return!0;if(!a){if(V){var e=b[0];var f=b[1]}else e=b,f=q[c];ia?(V&&(f=b.slice(1,3)),h=f[0],f=f[1]):ra&&(e=b.x,f=b.stackY,h=
f-b.y);ta||(d=f>=y&&f<=K);if(null!==f&&e>=v&&e<=z&&d)if(b=g.toPixels(e,!0),w){if("undefined"===typeof W||b===D){ia||(h=f);if("undefined"===typeof Z||f>ea)ea=f,Z=c;if("undefined"===typeof W||h<da)da=h,W=c}b!==D&&("undefined"!==typeof W&&(f=n.toPixels(ea,!0),S=n.toPixels(da,!0),ka(b,f,Z),S!==f&&ka(b,S,W)),W=Z=void 0,D=b)}else f=Math.ceil(n.toPixels(f,!0)),ka(b,f,c)}return!a}function d(){e(a,"renderedCanvas");delete a.buildKDTree;a.buildKDTree();la.debug.timeKDTree&&console.timeEnd("kd tree building")}
var a=this,b=a.options||{},f=!1,k=a.chart,g=this.xAxis,n=this.yAxis,m=b.xData||a.processedXData,q=b.yData||a.processedYData,r=b.data;f=g.getExtremes();var v=f.min,z=f.max;f=n.getExtremes();var y=f.min,K=f.max,H={},D,w=!!a.sampling,x=!1!==b.enableMouseTracking,S=n.getThreshold(b.threshold),ia=a.pointArrayMap&&"low,high"===a.pointArrayMap.join(","),ra=!!b.stacking,sa=a.cropStart||0,ta=a.requireSorting,V=!m,da,ea,W,Z,ua="x"===b.findNearestPointBy,ja=this.xData||this.options.xData||this.processedXData||
!1,ka=function(b,h,a){b=Math.ceil(b);c=ua?b:b+","+h;x&&!H[c]&&(H[c]=!0,k.inverted&&(b=g.len-b,h=n.len-h),xa.push({x:ja?ja[sa+a]:!1,clientX:b,plotX:b,plotY:h,i:sa+a}))};f=u(k,a);k.isBoosting=!0;var la=f.settings;if(this.visible){(this.points||this.graph)&&this.destroyGraphics();k.isChartSeriesBoosting()?(this.markerGroup&&this.markerGroup!==k.markerGroup&&this.markerGroup.destroy(),this.markerGroup=k.markerGroup,this.renderTarget&&(this.renderTarget=this.renderTarget.destroy())):(this.markerGroup===
k.markerGroup&&(this.markerGroup=void 0),this.markerGroup=a.plotGroup("markerGroup","markers",!0,1,k.seriesGroup));var xa=this.points=[];a.buildKDTree=p;f&&(L(f,this),f.pushSeries(a),C(f,this,k));k.renderer.forExport||(la.debug.timeKDTree&&console.time("kd tree building"),l(ra?a.data:m||r,h,d))}}});["heatmap","treemap"].forEach(function(c){q[c]&&m(q[c].prototype,"drawPoints",K)});q.bubble&&(delete q.bubble.prototype.buildKDTree,m(q.bubble.prototype,"markerAttribs",function(c){return this.isSeriesBoosting?
!1:c.apply(this,[].slice.call(arguments,1))}));q.scatter.prototype.fill=!0;f(q.area.prototype,{fill:!0,fillOpacity:!0,sampling:!0});f(q.column.prototype,{fill:!0,sampling:!0});a.prototype.propsRequireUpdateSeries.push("boost");a.prototype.callbacks.push(function(c){n(c,"predraw",function(){c.boostForceChartBoost=void 0;c.boostForceChartBoost=H(c);c.isBoosting=!1;!c.isChartSeriesBoosting()&&c.didBoost&&(c.didBoost=!1);c.boostClear&&c.boostClear();c.canvas&&c.ogl&&c.isChartSeriesBoosting()&&(c.didBoost=
!0,c.ogl.allocateBuffer(c));c.markerGroup&&c.xAxis&&0<c.xAxis.length&&c.yAxis&&0<c.yAxis.length&&c.markerGroup.translate(c.xAxis[0].pos,c.yAxis[0].pos)});n(c,"render",function(){c.ogl&&c.isChartSeriesBoosting()&&c.ogl.render(c)});var a=-1,d=-1;n(c.pointer,"afterGetHoverData",function(){var b=c.hoverSeries;if(c.markerGroup&&b){var h=c.inverted?b.yAxis:b.xAxis;b=c.inverted?b.xAxis:b.yAxis;if(h&&h.pos!==a||b&&b.pos!==d)c.markerGroup.translate(h.pos,b.pos),a=h.pos,d=b.pos}})})}});u(a,"Extensions/BoostCanvas.js",
[a["Core/Chart/Chart.js"],a["Core/Color/Color.js"],a["Core/Globals.js"],a["Core/Series/Series.js"],a["Core/Series/SeriesRegistry.js"],a["Core/Utilities.js"]],function(a,z,v,k,d,r){var q=z.parse,p=v.doc,u=v.noop,n=d.seriesTypes,f=r.addEvent,e=r.extend,m=r.fireEvent,l=r.isNumber,K=r.merge,L=r.pick,D=r.wrap,H;return function(){v.seriesTypes.heatmap&&D(v.seriesTypes.heatmap.prototype,"drawPoints",function(){var c=this.chart,a=this.getContext(),d=this.chart.inverted,e=this.xAxis,b=this.yAxis;a?(this.points.forEach(function(h){var f=
h.plotY;if("undefined"!==typeof f&&!isNaN(f)&&null!==h.y&&a){var g=h.shapeArgs||{};f=g.x;f=void 0===f?0:f;var t=g.y;t=void 0===t?0:t;var k=g.width;k=void 0===k?0:k;g=g.height;g=void 0===g?0:g;h=c.styledMode?h.series.colorAttribs(h):h.series.pointAttribs(h);a.fillStyle=h.fill;d?a.fillRect(b.len-t+e.left,e.len-f+b.top,-g,-k):a.fillRect(f+e.left,t+b.top,k,g)}}),this.canvasToSVG()):this.chart.showLoading("Your browser doesn't support HTML5 canvas, <br>please use a modern browser")});e(k.prototype,{getContext:function(){var c=
this.chart,a=c.chartWidth,d=c.chartHeight,e=c.seriesGroup||this.group,b=this,f=function(b,c,a,d,h,e,f){b.call(this,a,c,d,h,e,f)};c.isChartSeriesBoosting()&&(b=c,e=c.seriesGroup);var k=b.ctx;b.canvas||(b.canvas=p.createElement("canvas"),b.renderTarget=c.renderer.image("",0,0,a,d).addClass("highcharts-boost-canvas").add(e),b.ctx=k=b.canvas.getContext("2d"),c.inverted&&["moveTo","lineTo","rect","arc"].forEach(function(b){D(k,b,f)}),b.boostCopy=function(){b.renderTarget.attr({href:b.canvas.toDataURL("image/png")})},
b.boostClear=function(){k.clearRect(0,0,b.canvas.width,b.canvas.height);b===this&&b.renderTarget.attr({href:""})},b.boostClipRect=c.renderer.clipRect(),b.renderTarget.clip(b.boostClipRect));b.canvas.width!==a&&(b.canvas.width=a);b.canvas.height!==d&&(b.canvas.height=d);b.renderTarget.attr({x:0,y:0,width:a,height:d,style:"pointer-events: none",href:""});b.boostClipRect.attr(c.getBoostClipRect(b));return k},canvasToSVG:function(){this.chart.isChartSeriesBoosting()?this.boostClear&&this.boostClear():
(this.boostCopy||this.chart.boostCopy)&&(this.boostCopy||this.chart.boostCopy)()},cvsLineTo:function(c,a,d){c.lineTo(a,d)},renderCanvas:function(){var c=this,a=c.options,d=c.chart,k=this.xAxis,b=this.yAxis,n=(d.options.boost||{}).timeRendering||!1,p=0,g=c.processedXData,y=c.processedYData,D=a.data,B=k.getExtremes(),C=B.min,X=B.max;B=b.getExtremes();var Y=B.min,G=B.max,U={},ca,ba=!!c.sampling,w=a.marker&&a.marker.radius,x=this.cvsDrawPoint,S=a.lineWidth?this.cvsLineTo:void 0,ia=w&&1>=w?this.cvsMarkerSquare:
this.cvsMarkerCircle,ra=this.cvsStrokeBatch||1E3,sa=!1!==a.enableMouseTracking,ta;B=a.threshold;var V=b.getThreshold(B),da=l(B),ea=V,W=this.fill,Z=c.pointArrayMap&&"low,high"===c.pointArrayMap.join(","),ua=!!a.stacking,ja=c.cropStart||0;B=d.options.loading;var ka=c.requireSorting,la,xa=a.connectNulls,Ea=!g,ya,za,fa,va,wa,Q=ua?c.data:g||D,Ma=c.fillOpacity?z.parse(c.color).setOpacity(L(a.fillOpacity,.75)).get():c.color,Fa=function(){W?(E.fillStyle=Ma,E.fill()):(E.strokeStyle=c.color,E.lineWidth=a.lineWidth,
E.stroke())},Ga=function(b,a,h,e){0===p&&(E.beginPath(),S&&(E.lineJoin="round"));d.scroller&&"highcharts-navigator-series"===c.options.className?(a+=d.scroller.top,h&&(h+=d.scroller.top)):a+=d.plotTop;b+=d.plotLeft;la?E.moveTo(b,a):x?x(E,b,a,h,ta):S?S(E,b,a):ia&&ia.call(c,E,b,a,w,e);p+=1;p===ra&&(Fa(),p=0);ta={clientX:b,plotY:a,yBottom:h}},Na="x"===a.findNearestPointBy,Ha=this.xData||this.options.xData||this.processedXData||!1,Aa=function(c,a,h){wa=Na?c:c+","+a;sa&&!U[wa]&&(U[wa]=!0,d.inverted&&(c=
k.len-c,a=b.len-a),Oa.push({x:Ha?Ha[ja+h]:!1,clientX:c,plotX:c,plotY:a,i:ja+h}))};this.renderTarget&&this.renderTarget.attr({href:""});(this.points||this.graph)&&this.destroyGraphics();c.plotGroup("group","series",c.visible?"visible":"hidden",a.zIndex,d.seriesGroup);c.markerGroup=c.group;f(c,"destroy",function(){c.markerGroup=null});var Oa=this.points=[];var E=this.getContext();c.buildKDTree=u;this.boostClear&&this.boostClear();this.visible&&(99999<D.length&&(d.options.loading=K(B,{labelStyle:{backgroundColor:q("#ffffff").setOpacity(.75).get(),
padding:"1em",borderRadius:"0.5em"},style:{backgroundColor:"none",opacity:1}}),r.clearTimeout(H),d.showLoading("Drawing..."),d.options.loading=B),n&&console.time("canvas rendering"),v.eachAsync(Q,function(a,h){var e=!1,f=!1,g=!1,t=!1,n="undefined"===typeof d.index,p=!0;if(!n){if(Ea){var m=a[0];var l=a[1];Q[h+1]&&(g=Q[h+1][0]);Q[h-1]&&(t=Q[h-1][0])}else m=a,l=y[h],Q[h+1]&&(g=Q[h+1]),Q[h-1]&&(t=Q[h-1]);g&&g>=C&&g<=X&&(e=!0);t&&t>=C&&t<=X&&(f=!0);if(Z){Ea&&(l=a.slice(1,3));var q=l[0];l=l[1]}else ua&&
(m=a.x,l=a.stackY,q=l-a.y);a=null===l;ka||(p=l>=Y&&l<=G);if(!a&&(m>=C&&m<=X&&p||e||f))if(m=Math.round(k.toPixels(m,!0)),ba){if("undefined"===typeof fa||m===ca){Z||(q=l);if("undefined"===typeof va||l>za)za=l,va=h;if("undefined"===typeof fa||q<ya)ya=q,fa=h}m!==ca&&("undefined"!==typeof fa&&(l=b.toPixels(za,!0),V=b.toPixels(ya,!0),Ga(m,da?Math.min(l,ea):l,da?Math.max(V,ea):V,h),Aa(m,l,va),V!==l&&Aa(m,V,fa)),fa=va=void 0,ca=m)}else l=Math.round(b.toPixels(l,!0)),Ga(m,l,V,h),Aa(m,l,h);la=a&&!xa;0===h%
5E4&&(c.boostCopy||c.chart.boostCopy)&&(c.boostCopy||c.chart.boostCopy)()}return!n},function(){var b=d.loadingDiv,a=d.loadingShown;Fa();c.canvasToSVG();n&&console.timeEnd("canvas rendering");m(c,"renderedCanvas");a&&(e(b.style,{transition:"opacity 250ms",opacity:0}),d.loadingShown=!1,H=setTimeout(function(){b.parentNode&&b.parentNode.removeChild(b);d.loadingDiv=d.loadingSpan=null},250));delete c.buildKDTree;c.buildKDTree()},d.renderer.forExport?Number.MAX_VALUE:void 0))}});n.scatter.prototype.cvsMarkerCircle=
function(c,a,d,e){c.moveTo(a,d);c.arc(a,d,e,0,2*Math.PI,!1)};n.scatter.prototype.cvsMarkerSquare=function(c,a,d,e){c.rect(a-e,d-e,2*e,2*e)};n.scatter.prototype.fill=!0;n.bubble&&(n.bubble.prototype.cvsMarkerCircle=function(c,a,d,e,b){c.moveTo(a,d);c.arc(a,d,this.radii&&this.radii[b],0,2*Math.PI,!1)},n.bubble.prototype.cvsStrokeBatch=1);e(n.area.prototype,{cvsDrawPoint:function(c,a,d,e,b){b&&a!==b.clientX&&(c.moveTo(b.clientX,b.yBottom),c.lineTo(b.clientX,b.plotY),c.lineTo(a,d),c.lineTo(a,e))},fill:!0,
fillOpacity:!0,sampling:!0});e(n.column.prototype,{cvsDrawPoint:function(c,a,d,e){c.rect(a-1,d,1,e-d)},fill:!0,sampling:!0});a.prototype.callbacks.push(function(c){f(c,"predraw",function(){c.renderTarget&&c.renderTarget.attr({href:""});c.canvas&&c.canvas.getContext("2d").clearRect(0,0,c.canvas.width,c.canvas.height)});f(c,"render",function(){c.boostCopy&&c.boostCopy()})})}});u(a,"Extensions/Boost/BoostOverrides.js",[a["Core/Chart/Chart.js"],a["Core/DefaultOptions.js"],a["Core/Series/Point.js"],a["Core/Series/Series.js"],
a["Core/Series/SeriesRegistry.js"],a["Core/Utilities.js"],a["Extensions/Boost/BoostUtils.js"],a["Extensions/Boost/Boostables.js"],a["Extensions/Boost/BoostableMap.js"]],function(a,u,v,k,d,r,C,p,y){u=u.getOptions;var n=d.seriesTypes;d=r.addEvent;var f=r.error,e=r.isArray,m=r.isNumber,l=r.pick,q=r.wrap,z=C.boostEnabled,D=C.shouldForceChartSeriesBoosting,H=u().plotOptions;a.prototype.isChartSeriesBoosting=function(){return l(this.options.boost&&this.options.boost.seriesThreshold,50)<=this.series.length||
D(this)};a.prototype.getBoostClipRect=function(c){var a={x:this.plotLeft,y:this.plotTop,width:this.plotWidth,height:this.plotHeight};c===this&&(c=this.inverted?this.xAxis:this.yAxis,1>=c.length?(a.y=Math.min(c[0].pos,a.y),a.height=c[0].pos-this.plotTop+c[0].len):a.height=this.plotHeight);return a};k.prototype.getPoint=function(c){var a=c,d=this.xData||this.options.xData||this.processedXData||!1;!c||c instanceof this.pointClass||(a=(new this.pointClass).init(this,this.options.data[c.i],d?d[c.i]:void 0),
a.category=l(this.xAxis.categories?this.xAxis.categories[a.x]:a.x,a.x),a.dist=c.dist,a.distX=c.distX,a.plotX=c.plotX,a.plotY=c.plotY,a.index=c.i,a.isInside=this.isPointInside(c));return a};q(k.prototype,"searchPoint",function(c){return this.getPoint(c.apply(this,[].slice.call(arguments,1)))});q(v.prototype,"haloPath",function(c){var a=this.series,d=this.plotX,e=this.plotY,b=a.chart.inverted;a.isSeriesBoosting&&b&&(this.plotX=a.yAxis.len-e,this.plotY=a.xAxis.len-d);var f=c.apply(this,Array.prototype.slice.call(arguments,
1));a.isSeriesBoosting&&b&&(this.plotX=d,this.plotY=e);return f});q(k.prototype,"markerAttribs",function(c,a){var d=a.plotX,e=a.plotY,b=this.chart.inverted;this.isSeriesBoosting&&b&&(a.plotX=this.yAxis.len-e,a.plotY=this.xAxis.len-d);var h=c.apply(this,Array.prototype.slice.call(arguments,1));this.isSeriesBoosting&&b&&(a.plotX=d,a.plotY=e);return h});d(k,"destroy",function(){var c=this,a=c.chart;a.markerGroup===c.markerGroup&&(c.markerGroup=null);a.hoverPoints&&(a.hoverPoints=a.hoverPoints.filter(function(a){return a.series===
c}));a.hoverPoint&&a.hoverPoint.series===c&&(a.hoverPoint=null)});q(k.prototype,"getExtremes",function(a){return this.isSeriesBoosting&&this.hasExtremes&&this.hasExtremes()?{}:a.apply(this,Array.prototype.slice.call(arguments,1))});["translate","generatePoints","drawTracker","drawPoints","render"].forEach(function(a){function c(c){var d=this.options.stacking&&("translate"===a||"generatePoints"===a);if(!this.isSeriesBoosting||d||!z(this.chart)||"heatmap"===this.type||"treemap"===this.type||!y[this.type]||
0===this.options.boostThreshold)c.call(this);else if(this[a+"Canvas"])this[a+"Canvas"]()}q(k.prototype,a,c);"translate"===a&&"column bar arearange columnrange heatmap treemap".split(" ").forEach(function(d){n[d]&&q(n[d].prototype,a,c)})});q(k.prototype,"processData",function(a){function c(b){return d.forceCrop?!1:d.chart.isChartSeriesBoosting()||(b?b.length:0)>=(d.options.boostThreshold||Number.MAX_VALUE)}var d=this,k=this.options.data;z(this.chart)&&y[this.type]?(c(k)&&"heatmap"!==this.type&&"treemap"!==
this.type&&!this.options.stacking&&this.hasExtremes&&this.hasExtremes(!0)||(a.apply(this,Array.prototype.slice.call(arguments,1)),k=this.processedXData),(this.isSeriesBoosting=c(k))?(k=void 0,this.options.data&&this.options.data.length&&(k=this.getFirstValidPoint(this.options.data),m(k)||e(k)||f(12,!1,this.chart)),this.enterBoost()):this.exitBoost&&this.exitBoost()):a.apply(this,Array.prototype.slice.call(arguments,1))});d(k,"hide",function(){this.canvas&&this.renderTarget&&(this.ogl&&this.ogl.clear(),
this.boostClear())});k.prototype.enterBoost=function(){this.alteredByBoost=[];["allowDG","directTouch","stickyTracking"].forEach(function(a){this.alteredByBoost.push({prop:a,val:this[a],own:Object.hasOwnProperty.call(this,a)})},this);this.directTouch=this.allowDG=!1;this.finishedAnimating=this.stickyTracking=!0;this.labelBySeries&&(this.labelBySeries=this.labelBySeries.destroy())};k.prototype.exitBoost=function(){(this.alteredByBoost||[]).forEach(function(a){a.own?this[a.prop]=a.val:delete this[a.prop]},
this);this.boostClear&&this.boostClear()};k.prototype.hasExtremes=function(a){var c=this.options,d=this.xAxis&&this.xAxis.options,e=this.yAxis&&this.yAxis.options,b=this.colorAxis&&this.colorAxis.options;return c.data.length>(c.boostThreshold||Number.MAX_VALUE)&&m(e.min)&&m(e.max)&&(!a||m(d.min)&&m(d.max))&&(!b||m(b.min)&&m(b.max))};k.prototype.destroyGraphics=function(){var a=this,d=this,e=this.points,f,b;if(e)for(b=0;b<e.length;b+=1)(f=e[b])&&f.destroyElements&&f.destroyElements();["graph","area",
"tracker"].forEach(function(b){d[b]&&(d[b]=d[b].destroy())});this.getZonesGraphs&&this.getZonesGraphs([["graph","highcharts-graph"]]).forEach(function(b){var c=a[b[0]];c&&(a[b[0]]=c.destroy())})};p.forEach(function(a){H[a]&&(H[a].boostThreshold=5E3,H[a].boostData=[],n[a].prototype.fillOpacity=!0)})});u(a,"Extensions/Boost/NamedColors.js",[a["Core/Color/Color.js"]],function(a){var q={aliceblue:"#f0f8ff",antiquewhite:"#faebd7",aqua:"#00ffff",aquamarine:"#7fffd4",azure:"#f0ffff",beige:"#f5f5dc",bisque:"#ffe4c4",
black:"#000000",blanchedalmond:"#ffebcd",blue:"#0000ff",blueviolet:"#8a2be2",brown:"#a52a2a",burlywood:"#deb887",cadetblue:"#5f9ea0",chartreuse:"#7fff00",chocolate:"#d2691e",coral:"#ff7f50",cornflowerblue:"#6495ed",cornsilk:"#fff8dc",crimson:"#dc143c",cyan:"#00ffff",darkblue:"#00008b",darkcyan:"#008b8b",darkgoldenrod:"#b8860b",darkgray:"#a9a9a9",darkgreen:"#006400",darkkhaki:"#bdb76b",darkmagenta:"#8b008b",darkolivegreen:"#556b2f",darkorange:"#ff8c00",darkorchid:"#9932cc",darkred:"#8b0000",darksalmon:"#e9967a",
darkseagreen:"#8fbc8f",darkslateblue:"#483d8b",darkslategray:"#2f4f4f",darkturquoise:"#00ced1",darkviolet:"#9400d3",deeppink:"#ff1493",deepskyblue:"#00bfff",dimgray:"#696969",dodgerblue:"#1e90ff",feldspar:"#d19275",firebrick:"#b22222",floralwhite:"#fffaf0",forestgreen:"#228b22",fuchsia:"#ff00ff",gainsboro:"#dcdcdc",ghostwhite:"#f8f8ff",gold:"#ffd700",goldenrod:"#daa520",gray:"#808080",green:"#008000",greenyellow:"#adff2f",honeydew:"#f0fff0",hotpink:"#ff69b4",indianred:"#cd5c5c",indigo:"#4b0082",ivory:"#fffff0",
khaki:"#f0e68c",lavender:"#e6e6fa",lavenderblush:"#fff0f5",lawngreen:"#7cfc00",lemonchiffon:"#fffacd",lightblue:"#add8e6",lightcoral:"#f08080",lightcyan:"#e0ffff",lightgoldenrodyellow:"#fafad2",lightgrey:"#d3d3d3",lightgreen:"#90ee90",lightpink:"#ffb6c1",lightsalmon:"#ffa07a",lightseagreen:"#20b2aa",lightskyblue:"#87cefa",lightslateblue:"#8470ff",lightslategray:"#778899",lightsteelblue:"#b0c4de",lightyellow:"#ffffe0",lime:"#00ff00",limegreen:"#32cd32",linen:"#faf0e6",magenta:"#ff00ff",maroon:"#800000",
mediumaquamarine:"#66cdaa",mediumblue:"#0000cd",mediumorchid:"#ba55d3",mediumpurple:"#9370d8",mediumseagreen:"#3cb371",mediumslateblue:"#7b68ee",mediumspringgreen:"#00fa9a",mediumturquoise:"#48d1cc",mediumvioletred:"#c71585",midnightblue:"#191970",mintcream:"#f5fffa",mistyrose:"#ffe4e1",moccasin:"#ffe4b5",navajowhite:"#ffdead",navy:"#000080",oldlace:"#fdf5e6",olive:"#808000",olivedrab:"#6b8e23",orange:"#ffa500",orangered:"#ff4500",orchid:"#da70d6",palegoldenrod:"#eee8aa",palegreen:"#98fb98",paleturquoise:"#afeeee",
palevioletred:"#d87093",papayawhip:"#ffefd5",peachpuff:"#ffdab9",peru:"#cd853f",pink:"#ffc0cb",plum:"#dda0dd",powderblue:"#b0e0e6",purple:"#800080",red:"#ff0000",rosybrown:"#bc8f8f",royalblue:"#4169e1",saddlebrown:"#8b4513",salmon:"#fa8072",sandybrown:"#f4a460",seagreen:"#2e8b57",seashell:"#fff5ee",sienna:"#a0522d",silver:"#c0c0c0",skyblue:"#87ceeb",slateblue:"#6a5acd",slategray:"#708090",snow:"#fffafa",springgreen:"#00ff7f",steelblue:"#4682b4",tan:"#d2b48c",teal:"#008080",thistle:"#d8bfd8",tomato:"#ff6347",
turquoise:"#40e0d0",violet:"#ee82ee",violetred:"#d02090",wheat:"#f5deb3",white:"#ffffff",whitesmoke:"#f5f5f5",yellow:"#ffff00",yellowgreen:"#9acd32"};return a.names=q});u(a,"Extensions/Boost/Boost.js",[a["Extensions/Boost/BoostUtils.js"],a["Extensions/Boost/BoostInit.js"],a["Extensions/BoostCanvas.js"],a["Core/Utilities.js"]],function(a,u,v,k){k=k.error;a=a.hasWebGLSupport;a()?u():"undefined"!==typeof v?v():k(26)});u(a,"masters/modules/boost.src.js",[],function(){})});
//# sourceMappingURL=boost.js.map
```
