## Supplementary material for "Exploring natural odour landscapes: A case study with implications for human-biting insects": Figure S4: broken-axis.js

```
/*
 Highcharts JS v9.3.1 (2021-11-05)

 (c) 2009-2021 Torstein Honsi

 License: www.highcharts.com/license
*/
'use strict';(function(f){"object"===typeof module&&module.exports?(f["default"]=f,module.exports=f):"function"===typeof define&&define.amd?define("highcharts/modules/broken-axis",["highcharts"],function(l){f(l);f.Highcharts=l;return f}):f("undefined"!==typeof Highcharts?Highcharts:void 0)})(function(f){function l(f,h,n,l){f.hasOwnProperty(h)||(f[h]=l.apply(null,n))}f=f?f._modules:{};l(f,"Core/Axis/BrokenAxis.js",[f["Extensions/Stacking.js"],f["Core/Utilities.js"]],function(f,h){var n=h.addEvent,
l=h.find,x=h.fireEvent,z=h.isArray,m=h.isNumber,u=h.pick,v;(function(h){function v(){"undefined"!==typeof this.brokenAxis&&this.brokenAxis.setBreaks(this.options.breaks,!1)}function A(){this.brokenAxis&&this.brokenAxis.hasBreaks&&(this.options.ordinal=!1)}function B(){var c=this.brokenAxis;if(c&&c.hasBreaks){for(var b=this.tickPositions,g=this.tickPositions.info,e=[],a=0;a<b.length;a++)c.isInAnyBreak(b[a])||e.push(b[a]);this.tickPositions=e;this.tickPositions.info=g}}function C(){this.brokenAxis||
(this.brokenAxis=new y(this))}function D(){var c=this.options.connectNulls,b=this.points,g=this.xAxis,e=this.yAxis;if(this.isDirty)for(var a=b.length;a--;){var d=b[a],f=!(null===d.y&&!1===c)&&(g&&g.brokenAxis&&g.brokenAxis.isInAnyBreak(d.x,!0)||e&&e.brokenAxis&&e.brokenAxis.isInAnyBreak(d.y,!0));d.visible=f?!1:!1!==d.options.visible}}function E(){this.drawBreaks(this.xAxis,["x"]);this.drawBreaks(this.yAxis,u(this.pointArrayMap,["y"]))}function F(c,b){var g=this,e=g.points,a,d,f,k;if(c&&c.brokenAxis&&
c.brokenAxis.hasBreaks){var h=c.brokenAxis;b.forEach(function(b){a=h&&h.breakArray||[];d=c.isXAxis?c.min:u(g.options.threshold,c.min);e.forEach(function(e){k=u(e["stack"+b.toUpperCase()],e[b]);a.forEach(function(a){if(m(d)&&m(k)){f=!1;if(d<a.from&&k>a.to||d>a.from&&k<a.from)f="pointBreak";else if(d<a.from&&k>a.from&&k<a.to||d>a.from&&k>a.to&&k<a.from)f="pointInBreak";f&&x(c,f,{point:e,brk:a})}})})})}}function G(){var c=this.currentDataGrouping,b=c&&c.gapSize;c=this.points.slice();var g=this.yAxis,
e=this.options.gapSize,a=c.length-1,d;if(e&&0<a)for("value"!==this.options.gapUnit&&(e*=this.basePointRange),b&&b>e&&b>=this.basePointRange&&(e=b),d=void 0;a--;)d&&!1!==d.visible||(d=c[a+1]),b=c[a],!1!==d.visible&&!1!==b.visible&&(d.x-b.x>e&&(d=(b.x+d.x)/2,c.splice(a+1,0,{isNull:!0,x:d}),g.stacking&&this.options.stacking&&(d=g.stacking.stacks[this.stackKey][d]=new f(g,g.options.stackLabels,!1,d,this.stack),d.total=0)),d=b);return this.getGraphPath(c)}var w=[];h.compose=function(c,b){-1===w.indexOf(c)&&
(w.push(c),c.keepProps.push("brokenAxis"),n(c,"init",C),n(c,"afterInit",v),n(c,"afterSetTickPositions",B),n(c,"afterSetOptions",A));if(-1===w.indexOf(b)){w.push(b);var g=b.prototype;g.drawBreaks=F;g.gappedPath=G;n(b,"afterGeneratePoints",D);n(b,"afterRender",E)}return c};var y=function(){function c(b){this.hasBreaks=!1;this.axis=b}c.isInBreak=function(b,c){var e=b.repeat||Infinity,a=b.from,d=b.to-b.from;c=c>=a?(c-a)%e:e-(a-c)%e;return b.inclusive?c<=d:c<d&&0!==c};c.lin2Val=function(b){var g=this.brokenAxis;
g=g&&g.breakArray;if(!g||!m(b))return b;var e;for(e=0;e<g.length;e++){var a=g[e];if(a.from>=b)break;else a.to<b?b+=a.len:c.isInBreak(a,b)&&(b+=a.len)}return b};c.val2Lin=function(b){var g=this.brokenAxis;g=g&&g.breakArray;if(!g||!m(b))return b;var e=b,a;for(a=0;a<g.length;a++){var d=g[a];if(d.to<=b)e-=d.len;else if(d.from>=b)break;else if(c.isInBreak(d,b)){e-=b-d.from;break}}return e};c.prototype.findBreakAt=function(b,c){return l(c,function(c){return c.from<b&&b<c.to})};c.prototype.isInAnyBreak=
function(b,g){var e=this.axis,a=e.options.breaks||[],d=a.length,f;if(d&&m(b)){for(;d--;)if(c.isInBreak(a[d],b)){var k=!0;f||(f=u(a[d].showPoints,!e.isXAxis))}var h=k&&g?k&&!f:k}return h};c.prototype.setBreaks=function(b,f){var e=this,a=e.axis,d=z(b)&&!!b.length;a.isDirty=e.hasBreaks!==d;e.hasBreaks=d;a.options.breaks=a.userOptions.breaks=b;a.forceRedraw=!0;a.series.forEach(function(a){a.isDirty=!0});d||a.val2lin!==c.val2Lin||(delete a.val2lin,delete a.lin2val);d&&(a.userOptions.ordinal=!1,a.lin2val=
c.lin2Val,a.val2lin=c.val2Lin,a.setExtremes=function(b,c,d,f,g){if(e.hasBreaks){for(var h=this.options.breaks||[],k;k=e.findBreakAt(b,h);)b=k.to;for(;k=e.findBreakAt(c,h);)c=k.from;c<b&&(c=b)}a.constructor.prototype.setExtremes.call(this,b,c,d,f,g)},a.setAxisTranslation=function(){a.constructor.prototype.setAxisTranslation.call(this);e.unitLength=void 0;if(e.hasBreaks){var b=a.options.breaks||[],d=[],f=[],g=u(a.pointRangePadding,0),h=0,r,p=a.userMin||a.min,t=a.userMax||a.max,l;b.forEach(function(a){r=
a.repeat||Infinity;m(p)&&m(t)&&(c.isInBreak(a,p)&&(p+=a.to%r-p%r),c.isInBreak(a,t)&&(t-=t%r-a.from%r))});b.forEach(function(a){q=a.from;r=a.repeat||Infinity;if(m(p)&&m(t)){for(;q-r>p;)q-=r;for(;q<p;)q+=r;for(l=q;l<t;l+=r)d.push({value:l,move:"in"}),d.push({value:l+a.to-a.from,move:"out",size:a.breakSize})}});d.sort(function(a,b){return a.value===b.value?("in"===a.move?0:1)-("in"===b.move?0:1):a.value-b.value});var n=0;var q=p;d.forEach(function(a){n+="in"===a.move?1:-1;1===n&&"in"===a.move&&(q=a.value);
0===n&&m(q)&&(f.push({from:q,to:a.value,len:a.value-q-(a.size||0)}),h+=a.value-q-(a.size||0))});e.breakArray=f;m(p)&&m(t)&&m(a.min)&&(e.unitLength=t-p-h+g,x(a,"afterBreaks"),a.staticScale?a.transA=a.staticScale:e.unitLength&&(a.transA*=(t-a.min+g)/e.unitLength),g&&(a.minPixelPadding=a.transA*(a.minPointOffset||0)),a.min=p,a.max=t)}});u(f,!0)&&a.chart.redraw()};return c}();h.Additions=y})(v||(v={}));return v});l(f,"masters/modules/broken-axis.src.js",[f["Core/Globals.js"],f["Core/Axis/BrokenAxis.js"]],
function(f,h){h.compose(f.Axis,f.Series)})});
//# sourceMappingURL=broken-axis.js.map
```
