## Supplementary material for "Exploring natural odour landscapes: A case study with implications for human-biting insects": Figure S4: bullet.js

```
/*
 Highcharts JS v9.3.1 (2021-11-05)

 Bullet graph series type for Highcharts

 (c) 2010-2021 Kacper Madej

 License: www.highcharts.com/license
*/
'use strict';(function(b){"object"===typeof module&&module.exports?(b["default"]=b,module.exports=b):"function"===typeof define&&define.amd?define("highcharts/modules/bullet",["highcharts"],function(d){b(d);b.Highcharts=d;return b}):b("undefined"!==typeof Highcharts?Highcharts:void 0)})(function(b){function d(b,f,a,r){b.hasOwnProperty(f)||(b[f]=r.apply(null,a))}b=b?b._modules:{};d(b,"Series/Bullet/BulletPoint.js",[b["Series/Column/ColumnSeries.js"]],function(b){var f=this&&this.__extends||function(){var b=
function(a,c){b=Object.setPrototypeOf||{__proto__:[]}instanceof Array&&function(b,a){b.__proto__=a}||function(b,a){for(var c in a)a.hasOwnProperty(c)&&(b[c]=a[c])};return b(a,c)};return function(a,c){function f(){this.constructor=a}b(a,c);a.prototype=null===c?Object.create(c):(f.prototype=c.prototype,new f)}}();return function(b){function a(){var a=null!==b&&b.apply(this,arguments)||this;a.options=void 0;a.series=void 0;return a}f(a,b);a.prototype.destroy=function(){this.targetGraphic&&(this.targetGraphic=
this.targetGraphic.destroy());b.prototype.destroy.apply(this,arguments)};return a}(b.prototype.pointClass)});d(b,"Series/Bullet/BulletSeries.js",[b["Series/Bullet/BulletPoint.js"],b["Core/Series/SeriesRegistry.js"],b["Core/Utilities.js"]],function(b,f,a){var d=this&&this.__extends||function(){var b=function(a,e){b=Object.setPrototypeOf||{__proto__:[]}instanceof Array&&function(b,a){b.__proto__=a}||function(b,a){for(var e in a)a.hasOwnProperty(e)&&(b[e]=a[e])};return b(a,e)};return function(a,e){function k(){this.constructor=
a}b(a,e);a.prototype=null===e?Object.create(e):(k.prototype=e.prototype,new k)}}(),c=f.seriesTypes.column,t=a.extend,l=a.isNumber,u=a.merge,p=a.pick,v=a.relativeLength;a=function(b){function a(){var a=null!==b&&b.apply(this,arguments)||this;a.data=void 0;a.options=void 0;a.points=void 0;a.targetData=void 0;return a}d(a,b);a.prototype.drawPoints=function(){var a=this,k=a.chart,c=a.options,f=c.animationLimit||250;b.prototype.drawPoints.apply(this,arguments);a.points.forEach(function(b){var e=b.options,
d=b.target,q=b.y,g=b.targetGraphic;if(l(d)&&null!==d){var h=u(c.targetOptions,e.targetOptions);var r=h.height;var m=b.shapeArgs;b.dlBox&&m&&!l(m.width)&&(m=b.dlBox);var n=v(h.width,m.width);var t=a.yAxis.translate(d,!1,!0,!1,!0)-h.height/2-.5;n=a.crispCol.apply({chart:k,borderWidth:h.borderWidth,options:{crisp:c.crisp}},[m.x+m.width/2-n/2,t,n,r]);g?(g[k.pointCount<f?"animate":"attr"](n),l(q)&&null!==q?g.element.point=b:g.element.point=void 0):b.targetGraphic=g=k.renderer.rect().attr(n).add(a.group);
k.styledMode||g.attr({fill:p(h.color,e.color,a.zones.length&&(b.getZone.call({series:a,x:b.x,y:d,options:{}}).color||a.color)||void 0,b.color,a.color),stroke:p(h.borderColor,b.borderColor,a.options.borderColor),"stroke-width":h.borderWidth,r:h.borderRadius});l(q)&&null!==q&&(g.element.point=b);g.addClass(b.getClassName()+" highcharts-bullet-target",!0)}else g&&(b.targetGraphic=g.destroy())})};a.prototype.getExtremes=function(a){a=b.prototype.getExtremes.call(this,a);var c=this.targetData;c&&c.length&&
(c=b.prototype.getExtremes.call(this,c),l(c.dataMin)&&(a.dataMin=Math.min(p(a.dataMin,Infinity),c.dataMin)),l(c.dataMax)&&(a.dataMax=Math.max(p(a.dataMax,-Infinity),c.dataMax)));return a};a.defaultOptions=u(c.defaultOptions,{targetOptions:{width:"140%",height:3,borderWidth:0,borderRadius:0},tooltip:{pointFormat:'<span style="color:{series.color}">\u25cf</span> {series.name}: <b>{point.y}</b>. Target: <b>{point.target}</b><br/>'}});return a}(c);t(a.prototype,{parallelArrays:["x","y","target"],pointArrayMap:["y",
"target"]});a.prototype.pointClass=b;f.registerSeriesType("bullet",a);"";return a});d(b,"masters/modules/bullet.src.js",[],function(){})});
//# sourceMappingURL=bullet.js.map
```
