## Supplementary material for "Exploring natural odour landscapes: A case study with implications for human-biting insects": Figure S4: coloraxis.js

```
/*
 Highcharts JS v9.3.1 (2021-11-05)

 ColorAxis module

 (c) 2012-2021 Pawel Potaczek

 License: www.highcharts.com/license
*/
'use strict';(function(c){"object"===typeof module&&module.exports?(c["default"]=c,module.exports=c):"function"===typeof define&&define.amd?define("highcharts/modules/color-axis",["highcharts"],function(n){c(n);c.Highcharts=n;return c}):c("undefined"!==typeof Highcharts?Highcharts:void 0)})(function(c){function n(c,m,t,p){c.hasOwnProperty(m)||(c[m]=p.apply(null,t))}c=c?c._modules:{};n(c,"Core/Axis/Color/ColorAxisComposition.js",[c["Core/Color/Color.js"],c["Core/Utilities.js"]],function(c,m){var l=
c.parse,p=m.addEvent,n=m.extend,y=m.merge,z=m.pick,r=m.splat,u;(function(c){function m(){var b=this,a=this.options;this.colorAxis=[];a.colorAxis&&(a.colorAxis=r(a.colorAxis),a.colorAxis.forEach(function(a,d){a.index=d;new h(b,a)}))}function u(a){var b=this,d=function(d){d=a.allItems.indexOf(d);-1!==d&&(b.destroyItem(a.allItems[d]),a.allItems.splice(d,1))},f=[],g,h;(this.chart.colorAxis||[]).forEach(function(a){(g=a.options)&&g.showInLegend&&(g.dataClasses&&g.visible?f=f.concat(a.getDataClassLegendSymbols()):
g.visible&&f.push(a),a.series.forEach(function(a){if(!a.options.showInLegend||g.dataClasses)"point"===a.options.legendType?a.points.forEach(function(a){d(a)}):d(a)}))});for(h=f.length;h--;)a.allItems.unshift(f[h])}function t(a){a.visible&&a.item.legendColor&&a.item.legendSymbol.attr({fill:a.item.legendColor})}function v(){var a=this.chart.colorAxis;a&&a.forEach(function(a,d,b){a.update({},b)})}function x(){(this.chart.colorAxis&&this.chart.colorAxis.length||this.colorAttribs)&&this.translateColors()}
function q(){var a=this.axisTypes;a?-1===a.indexOf("colorAxis")&&a.push("colorAxis"):this.axisTypes=["colorAxis"]}function B(a){var d=this,b=a?"show":"hide";d.visible=d.options.visible=!!a;["graphic","dataLabel"].forEach(function(a){if(d[a])d[a][b]()});this.series.buildKDTree()}function e(){var a=this,d=this.options.nullColor,b=this.colorAxis,f=this.colorKey;(this.data.length?this.data:this.points).forEach(function(h){var g=h.getNestedProperty(f);(g=h.options.color||(h.isNull||null===h.value?d:b&&
"undefined"!==typeof g?b.toColor(g,h):h.color||a.color))&&h.color!==g&&(h.color=g,"point"===a.options.legendType&&h.legendItem&&a.chart.legend.colorizeItem(h,h.visible))})}function b(a){var d=a.prototype.createAxis;a.prototype.createAxis=function(a,b){if("colorAxis"!==a)return d.apply(this,arguments);var f=new h(this,y(b.axis,{index:this[a].length,isX:!1}));this.isDirtyLegend=!0;this.axes.forEach(function(a){a.series=[]});this.series.forEach(function(a){a.bindAxes();a.isDirtyData=!0});z(b.redraw,
!0)&&this.redraw(b.animation);return f}}function d(){this.elem.attr("fill",l(this.start).tweenTo(l(this.end),this.pos),void 0,!0)}function f(){this.elem.attr("stroke",l(this.start).tweenTo(l(this.end),this.pos),void 0,!0)}var a=[],h;c.compose=function(g,k,c,w,l){h||(h=g);-1===a.indexOf(k)&&(a.push(k),g=k.prototype,g.collectionsWithUpdate.push("colorAxis"),g.collectionsWithInit.colorAxis=[g.addColorAxis],p(k,"afterGetAxes",m),b(k));-1===a.indexOf(c)&&(a.push(c),k=c.prototype,k.fillSetter=d,k.strokeSetter=
f);-1===a.indexOf(w)&&(a.push(w),p(w,"afterGetAllItems",u),p(w,"afterColorizeItem",t),p(w,"afterUpdate",v));-1===a.indexOf(l)&&(a.push(l),n(l.prototype,{optionalAxis:"colorAxis",translateColors:e}),n(l.prototype.pointClass.prototype,{setVisible:B}),p(l,"afterTranslate",x),p(l,"bindAxes",q))};c.pointSetVisible=B})(u||(u={}));return u});n(c,"Core/Axis/Color/ColorAxisDefaults.js",[],function(){return{lineWidth:0,minPadding:0,maxPadding:0,gridLineWidth:1,tickPixelInterval:72,startOnTick:!0,endOnTick:!0,
offset:0,marker:{animation:{duration:50},width:.01,color:"#999999"},labels:{overflow:"justify",rotation:0},minColor:"#e6ebf5",maxColor:"#003399",tickLength:5,showInLegend:!0}});n(c,"Core/Axis/Color/ColorAxis.js",[c["Core/Axis/Axis.js"],c["Core/Color/Color.js"],c["Core/Axis/Color/ColorAxisComposition.js"],c["Core/Axis/Color/ColorAxisDefaults.js"],c["Core/Globals.js"],c["Core/Legend/LegendSymbol.js"],c["Core/Series/SeriesRegistry.js"],c["Core/Utilities.js"]],function(c,m,n,p,C,y,z,r){var l=this&&this.__extends||
function(){var c=function(e,b){c=Object.setPrototypeOf||{__proto__:[]}instanceof Array&&function(d,b){d.__proto__=b}||function(b,f){for(var a in f)f.hasOwnProperty(a)&&(b[a]=f[a])};return c(e,b)};return function(e,b){function d(){this.constructor=e}c(e,b);e.prototype=null===b?Object.create(b):(d.prototype=b.prototype,new d)}}(),v=m.parse,t=C.noop,A=z.series,D=r.extend,E=r.isNumber,x=r.merge,q=r.pick;m=function(c){function e(b,d){var f=c.call(this,b,d)||this;f.beforePadding=!1;f.chart=void 0;f.coll=
"colorAxis";f.dataClasses=void 0;f.legendItem=void 0;f.legendItems=void 0;f.name="";f.options=void 0;f.stops=void 0;f.visible=!0;f.init(b,d);return f}l(e,c);e.compose=function(b,d,f,a){n.compose(e,b,d,f,a)};e.prototype.init=function(b,d){var f=b.options.legend||{},a=d.layout?"vertical"!==d.layout:"vertical"!==f.layout,h=d.visible;f=x(e.defaultColorAxisOptions,d,{showEmpty:!1,title:null,visible:f.enabled&&!1!==h});this.coll="colorAxis";this.side=d.side||a?2:1;this.reversed=d.reversed||!a;this.opposite=
!a;c.prototype.init.call(this,b,f);this.userOptions.visible=h;d.dataClasses&&this.initDataClasses(d);this.initStops();this.horiz=a;this.zoomEnabled=!1};e.prototype.initDataClasses=function(b){var d=this.chart,f=this.options,a=b.dataClasses.length,h,c=0,e=d.options.chart.colorCount;this.dataClasses=h=[];this.legendItems=[];(b.dataClasses||[]).forEach(function(b,g){b=x(b);h.push(b);if(d.styledMode||!b.color)"category"===f.dataClassColor?(d.styledMode||(g=d.options.colors,e=g.length,b.color=g[c]),b.colorIndex=
c,c++,c===e&&(c=0)):b.color=v(f.minColor).tweenTo(v(f.maxColor),2>a?.5:g/(a-1))})};e.prototype.hasData=function(){return!!(this.tickPositions||[]).length};e.prototype.setTickPositions=function(){if(!this.dataClasses)return c.prototype.setTickPositions.call(this)};e.prototype.initStops=function(){this.stops=this.options.stops||[[0,this.options.minColor],[1,this.options.maxColor]];this.stops.forEach(function(b){b.color=v(b[1])})};e.prototype.setOptions=function(b){c.prototype.setOptions.call(this,b);
this.options.crosshair=this.options.marker};e.prototype.setAxisSize=function(){var b=this.legendSymbol,d=this.chart,f=d.options.legend||{},a,c;b?(this.left=f=b.attr("x"),this.top=a=b.attr("y"),this.width=c=b.attr("width"),this.height=b=b.attr("height"),this.right=d.chartWidth-f-c,this.bottom=d.chartHeight-a-b,this.len=this.horiz?c:b,this.pos=this.horiz?f:a):this.len=(this.horiz?f.symbolWidth:f.symbolHeight)||e.defaultLegendLength};e.prototype.normalizedValue=function(b){this.logarithmic&&(b=this.logarithmic.log2lin(b));
return 1-(this.max-b)/(this.max-this.min||1)};e.prototype.toColor=function(b,d){var f=this.dataClasses,a=this.stops,c;if(f)for(c=f.length;c--;){var e=f[c];var k=e.from;a=e.to;if(("undefined"===typeof k||b>=k)&&("undefined"===typeof a||b<=a)){var l=e.color;d&&(d.dataClass=c,d.colorIndex=e.colorIndex);break}}else{b=this.normalizedValue(b);for(c=a.length;c--&&!(b>a[c][0]););k=a[c]||a[c+1];a=a[c+1]||k;b=1-(a[0]-b)/(a[0]-k[0]||1);l=k.color.tweenTo(a.color,b)}return l};e.prototype.getOffset=function(){var b=
this.legendGroup,d=this.chart.axisOffset[this.side];b&&(this.axisParent=b,c.prototype.getOffset.call(this),this.added||(this.added=!0,this.labelLeft=0,this.labelRight=this.width),this.chart.axisOffset[this.side]=d)};e.prototype.setLegendColor=function(){var b=this.reversed,d=b?1:0;b=b?0:1;d=this.horiz?[d,0,b,0]:[0,b,0,d];this.legendColor={linearGradient:{x1:d[0],y1:d[1],x2:d[2],y2:d[3]},stops:this.stops}};e.prototype.drawLegendSymbol=function(b,d){var c=b.padding,a=b.options,h=this.horiz,g=q(a.symbolWidth,
h?e.defaultLegendLength:12),k=q(a.symbolHeight,h?12:e.defaultLegendLength),l=q(a.labelPadding,h?16:30);a=q(a.itemDistance,10);this.setLegendColor();d.legendSymbol=this.chart.renderer.rect(0,b.baseline-11,g,k).attr({zIndex:1}).add(d.legendGroup);this.legendItemWidth=g+c+(h?a:l);this.legendItemHeight=k+c+(h?l:0)};e.prototype.setState=function(b){this.series.forEach(function(d){d.setState(b)})};e.prototype.setVisible=function(){};e.prototype.getSeriesExtremes=function(){var b=this.series,d=b.length,
c;this.dataMin=Infinity;for(this.dataMax=-Infinity;d--;){var a=b[d];var e=a.colorKey=q(a.options.colorKey,a.colorKey,a.pointValKey,a.zoneAxis,"y");var g=a.pointArrayMap;var k=a[e+"Min"]&&a[e+"Max"];if(a[e+"Data"])var l=a[e+"Data"];else if(g){l=[];g=g.indexOf(e);var m=a.yData;if(0<=g&&m)for(c=0;c<m.length;c++)l.push(q(m[c][g],m[c]))}else l=a.yData;k?(a.minColorValue=a[e+"Min"],a.maxColorValue=a[e+"Max"]):(l=A.prototype.getExtremes.call(a,l),a.minColorValue=l.dataMin,a.maxColorValue=l.dataMax);"undefined"!==
typeof a.minColorValue&&(this.dataMin=Math.min(this.dataMin,a.minColorValue),this.dataMax=Math.max(this.dataMax,a.maxColorValue));k||A.prototype.applyExtremes.call(a)}};e.prototype.drawCrosshair=function(b,d){var f=d&&d.plotX,a=d&&d.plotY,e=this.pos,g=this.len;if(d){var k=this.toPixels(d.getNestedProperty(d.series.colorKey));k<e?k=e-2:k>e+g&&(k=e+g+2);d.plotX=k;d.plotY=this.len-k;c.prototype.drawCrosshair.call(this,b,d);d.plotX=f;d.plotY=a;this.cross&&!this.cross.addedToColorAxis&&this.legendGroup&&
(this.cross.addClass("highcharts-coloraxis-marker").add(this.legendGroup),this.cross.addedToColorAxis=!0,this.chart.styledMode||"object"!==typeof this.crosshair||this.cross.attr({fill:this.crosshair.color}))}};e.prototype.getPlotLinePath=function(b){var d=this.left,f=b.translatedValue,a=this.top;return E(f)?this.horiz?[["M",f-4,a-6],["L",f+4,a-6],["L",f,a],["Z"]]:[["M",d,f],["L",d-6,f+6],["L",d-6,f-6],["Z"]]:c.prototype.getPlotLinePath.call(this,b)};e.prototype.update=function(b,d){var f=this.chart.legend;
this.series.forEach(function(a){a.isDirtyData=!0});(b.dataClasses&&f.allItems||this.dataClasses)&&this.destroyItems();c.prototype.update.call(this,b,d);this.legendItem&&(this.setLegendColor(),f.colorizeItem(this,!0))};e.prototype.destroyItems=function(){var b=this.chart;this.legendItem?b.legend.destroyItem(this):this.legendItems&&this.legendItems.forEach(function(d){b.legend.destroyItem(d)});b.isDirtyLegend=!0};e.prototype.destroy=function(){this.chart.isDirtyLegend=!0;this.destroyItems();c.prototype.destroy.apply(this,
[].slice.call(arguments))};e.prototype.remove=function(b){this.destroyItems();c.prototype.remove.call(this,b)};e.prototype.getDataClassLegendSymbols=function(){var b=this,d=b.chart,c=b.legendItems,a=d.options.legend,e=a.valueDecimals,g=a.valueSuffix||"",k;c.length||b.dataClasses.forEach(function(a,f){var l=a.from,h=a.to,m=d.numberFormatter,n=!0;k="";"undefined"===typeof l?k="< ":"undefined"===typeof h&&(k="> ");"undefined"!==typeof l&&(k+=m(l,e)+g);"undefined"!==typeof l&&"undefined"!==typeof h&&
(k+=" - ");"undefined"!==typeof h&&(k+=m(h,e)+g);c.push(D({chart:d,name:k,options:{},drawLegendSymbol:y.drawRectangle,visible:!0,setState:t,isDataClass:!0,setVisible:function(){n=b.visible=!n;b.series.forEach(function(a){a.points.forEach(function(a){a.dataClass===f&&a.setVisible(n)})});d.legend.colorizeItem(this,n)}},a))});return c};e.defaultColorAxisOptions=p;e.defaultLegendLength=200;e.keepProps=["legendGroup","legendItemHeight","legendItemWidth","legendItem","legendSymbol"];return e}(c);Array.prototype.push.apply(c.keepProps,
m.keepProps);"";return m});n(c,"masters/modules/coloraxis.src.js",[c["Core/Globals.js"],c["Core/Axis/Color/ColorAxis.js"]],function(c,m){c.ColorAxis=m;m.compose(c.Chart,c.Fx,c.Legend,c.Series)})});
//# sourceMappingURL=coloraxis.js.map
```
