## Supplementary material for "Exploring natural odour landscapes: A case study with implications for human-biting insects": Figure S4: current-date-indicator.js

```
/*
 Highcharts Gantt JS v9.3.1 (2021-11-05)

 CurrentDateIndicator

 (c) 2010-2021 Lars A. V. Cabrera

 License: www.highcharts.com/license
*/
'use strict';(function(a){"object"===typeof module&&module.exports?(a["default"]=a,module.exports=a):"function"===typeof define&&define.amd?define("highcharts/modules/current-date-indicator",["highcharts"],function(b){a(b);a.Highcharts=b;return a}):a("undefined"!==typeof Highcharts?Highcharts:void 0)})(function(a){function b(a,b,d,e){a.hasOwnProperty(b)||(a[b]=e.apply(null,d))}a=a?a._modules:{};b(a,"Extensions/CurrentDateIndication.js",[a["Core/Axis/Axis.js"],a["Core/Axis/PlotLineOrBand/PlotLineOrBand.js"],
a["Core/Utilities.js"]],function(a,b,d){var e=d.addEvent,f=d.merge;d=d.wrap;var g={color:"#ccd6eb",width:2,label:{format:"%a, %b %d %Y, %H:%M",formatter:function(a,b){return this.axis.chart.time.dateFormat(b||"",a)},rotation:0,style:{fontSize:"10px"}}};e(a,"afterSetOptions",function(){var a=this.options,b=a.currentDateIndicator;b&&(b="object"===typeof b?f(g,b):f(g),b.value=Date.now(),b.className="highcharts-current-date-indicator",a.plotLines||(a.plotLines=[]),a.plotLines.push(b))});e(b,"render",
function(){this.label&&this.label.attr({text:this.getLabelText(this.options.label)})});d(b.prototype,"getLabelText",function(a,b){var c=this.options;return c&&c.className&&-1!==c.className.indexOf("highcharts-current-date-indicator")&&c.label&&"function"===typeof c.label.formatter?(c.value=Date.now(),c.label.formatter.call(this,c.value,c.label.format)):a.call(this,b)})});b(a,"masters/modules/current-date-indicator.src.js",[],function(){})});
//# sourceMappingURL=current-date-indicator.js.map
```
