## Supplementary material for "Exploring natural odour landscapes: A case study with implications for human-biting insects": Figure S4: cylinder.js

```
/*
 Highcharts JS v9.3.1 (2021-11-05)

 Highcharts cylinder module

 (c) 2010-2021 Kacper Madej

 License: www.highcharts.com/license
*/
'use strict';(function(a){"object"===typeof module&&module.exports?(a["default"]=a,module.exports=a):"function"===typeof define&&define.amd?define("highcharts/modules/cylinder",["highcharts","highcharts/highcharts-3d"],function(h){a(h);a.Highcharts=h;return a}):a("undefined"!==typeof Highcharts?Highcharts:void 0)})(function(a){function h(a,e,l,b){a.hasOwnProperty(e)||(a[e]=b.apply(null,l))}a=a?a._modules:{};h(a,"Series/Cylinder/CylinderPoint.js",[a["Core/Series/SeriesRegistry.js"],a["Core/Utilities.js"]],
function(a,e){var l=this&&this.__extends||function(){var a=function(b,g){a=Object.setPrototypeOf||{__proto__:[]}instanceof Array&&function(a,b){a.__proto__=b}||function(a,b){for(var g in b)b.hasOwnProperty(g)&&(a[g]=b[g])};return a(b,g)};return function(b,g){function e(){this.constructor=b}a(b,g);b.prototype=null===g?Object.create(g):(e.prototype=g.prototype,new e)}}();e=e.extend;a=function(a){function b(){var b=null!==a&&a.apply(this,arguments)||this;b.options=void 0;b.series=void 0;return b}l(b,
a);return b}(a.seriesTypes.column.prototype.pointClass);e(a.prototype,{shapeType:"cylinder"});return a});h(a,"Series/Cylinder/CylinderComposition.js",[a["Core/Color/Color.js"],a["Core/Globals.js"],a["Extensions/Math3D.js"],a["Core/Renderer/RendererRegistry.js"],a["Core/Utilities.js"]],function(a,e,l,b,h){var g=a.parse,m=e.charts,u=e.deg2rad,v=l.perspective;a=h.merge;var w=h.pick;b=b.getRendererType().prototype;var n=b.cuboidPath,p=function(c){return!c.some(function(c){return"C"===c[0]})};h=a(b.elements3d.cuboid,
{parts:["top","bottom","front","back"],pathType:"cylinder",fillSetter:function(c){this.singleSetterForParts("fill",null,{front:c,back:c,top:g(c).brighten(.1).get(),bottom:g(c).brighten(-.1).get()});this.color=this.fill=c;return this}});b.elements3d.cylinder=h;b.cylinder=function(c){return this.element3d("cylinder",c)};b.cylinderPath=function(c){var a=m[this.chartIndex],d=n.call(this,c),f=!d.isTop,b=!d.isFront,e=this.getCylinderEnd(a,c);c=this.getCylinderEnd(a,c,!0);return{front:this.getCylinderFront(e,
c),back:this.getCylinderBack(e,c),top:e,bottom:c,zIndexes:{top:f?3:0,bottom:f?0:3,front:b?2:1,back:b?1:2,group:d.zIndexes.group}}};b.getCylinderFront=function(c,a){c=c.slice(0,3);if(p(a)){var d=a[0];"M"===d[0]&&(c.push(a[2]),c.push(a[1]),c.push(["L",d[1],d[2]]))}else{d=a[0];var f=a[1];a=a[2];"M"===d[0]&&"C"===f[0]&&"C"===a[0]&&(c.push(["L",a[5],a[6]]),c.push(["C",a[3],a[4],a[1],a[2],f[5],f[6]]),c.push(["C",f[3],f[4],f[1],f[2],d[1],d[2]]))}c.push(["Z"]);return c};b.getCylinderBack=function(c,a){var d=
[];if(p(c)){var f=c[0],b=c[2];"M"===f[0]&&"L"===b[0]&&(d.push(["M",b[1],b[2]]),d.push(c[3]),d.push(["L",f[1],f[2]]))}else"C"===c[2][0]&&d.push(["M",c[2][5],c[2][6]]),d.push(c[3],c[4]);p(a)?(f=a[0],"M"===f[0]&&(d.push(["L",f[1],f[2]]),d.push(a[3]),d.push(a[2]))):(c=a[2],f=a[3],a=a[4],"C"===c[0]&&"C"===f[0]&&"C"===a[0]&&(d.push(["L",a[5],a[6]]),d.push(["C",a[3],a[4],a[1],a[2],f[5],f[6]]),d.push(["C",f[3],f[4],f[1],f[2],c[5],c[6]])));d.push(["Z"]);return d};b.getCylinderEnd=function(a,b,d){var c=b.width;
c=void 0===c?0:c;var e=b.height,g=void 0===e?0:e;e=b.alphaCorrection;var h=void 0===e?0:e;e=w(b.depth,c,0);var k=Math.min(c,e)/2;h=u*(a.options.chart.options3d.beta-90+h);d=(b.y||0)+(d?g:0);g=.5519*k;var l=c/2+(b.x||0),p=e/2+(b.z||0),n=[{x:0,y:d,z:k},{x:g,y:d,z:k},{x:k,y:d,z:g},{x:k,y:d,z:0},{x:k,y:d,z:-g},{x:g,y:d,z:-k},{x:0,y:d,z:-k},{x:-g,y:d,z:-k},{x:-k,y:d,z:-g},{x:-k,y:d,z:0},{x:-k,y:d,z:g},{x:-g,y:d,z:k},{x:0,y:d,z:k}],m=Math.cos(h),t=Math.sin(h),q,r;n.forEach(function(a,c){q=a.x;r=a.z;n[c].x=
q*m-r*t+l;n[c].z=r*m+q*t+p});a=v(n,a,!0);return 2.5>Math.abs(a[3].y-a[9].y)&&2.5>Math.abs(a[0].y-a[6].y)?this.toLinePath([a[0],a[3],a[6],a[9]],!0):this.getCurvedPath(a)};b.getCurvedPath=function(a){var c=[["M",a[0].x,a[0].y]],b=a.length-2,f;for(f=1;f<b;f+=3)c.push(["C",a[f].x,a[f].y,a[f+1].x,a[f+1].y,a[f+2].x,a[f+2].y]);return c}});h(a,"Series/Cylinder/CylinderSeries.js",[a["Series/Cylinder/CylinderPoint.js"],a["Core/Series/SeriesRegistry.js"],a["Core/Utilities.js"]],function(a,e,h){var b=this&&this.__extends||
function(){var a=function(b,e){a=Object.setPrototypeOf||{__proto__:[]}instanceof Array&&function(a,b){a.__proto__=b}||function(a,b){for(var c in b)b.hasOwnProperty(c)&&(a[c]=b[c])};return a(b,e)};return function(b,e){function g(){this.constructor=b}a(b,e);b.prototype=null===e?Object.create(e):(g.prototype=e.prototype,new g)}}(),l=e.seriesTypes.column,g=h.extend,m=h.merge;h=function(a){function e(){var b=null!==a&&a.apply(this,arguments)||this;b.data=void 0;b.options=void 0;b.points=void 0;return b}
b(e,a);e.defaultOptions=m(l.defaultOptions);return e}(l);g(h.prototype,{pointClass:a});e.registerSeriesType("cylinder",h);"";return h});h(a,"masters/modules/cylinder.src.js",[],function(){})});
//# sourceMappingURL=cylinder.js.map
```
