## Supplementary material for "Exploring natural odour landscapes: A case study with implications for human-biting insects": Figure S4: data.js

```
/*
 Highcharts JS v9.3.1 (2021-11-05)

 Data module

 (c) 2012-2021 Torstein Honsi

 License: www.highcharts.com/license
*/
'use strict';(function(b){"object"===typeof module&&module.exports?(b["default"]=b,module.exports=b):"function"===typeof define&&define.amd?define("highcharts/modules/data",["highcharts"],function(p){b(p);b.Highcharts=p;return b}):b("undefined"!==typeof Highcharts?Highcharts:void 0)})(function(b){function p(b,e,v,p){b.hasOwnProperty(e)||(b[e]=p.apply(null,v))}b=b?b._modules:{};p(b,"Core/HttpUtilities.js",[b["Core/Globals.js"],b["Core/Utilities.js"]],function(b,e){var v=b.doc,p=e.createElement,u=e.discardElement,
q=e.merge,D=e.objectEach,w={ajax:function(b){var l=q(!0,{url:!1,type:"get",dataType:"json",success:!1,error:!1,data:!1,headers:{}},b);b={json:"application/json",xml:"application/xml",text:"text/plain",octet:"application/octet-stream"};var e=new XMLHttpRequest;if(!l.url)return!1;e.open(l.type.toUpperCase(),l.url,!0);l.headers["Content-Type"]||e.setRequestHeader("Content-Type",b[l.dataType]||b.text);D(l.headers,function(b,l){e.setRequestHeader(l,b)});e.onreadystatechange=function(){if(4===e.readyState){if(200===
e.status){var b=e.responseText;if("json"===l.dataType)try{b=JSON.parse(b)}catch(A){l.error&&l.error(e,A);return}return l.success&&l.success(b)}l.error&&l.error(e,e.responseText)}};try{l.data=JSON.stringify(l.data)}catch(E){}e.send(l.data||!0)},getJSON:function(b,e){w.ajax({url:b,success:e,dataType:"json",headers:{"Content-Type":"text/plain"}})},post:function(b,e,w){var l=p("form",q({method:"post",action:b,enctype:"multipart/form-data"},w),{display:"none"},v.body);D(e,function(b,e){p("input",{type:"hidden",
name:e,value:b},null,l)});l.submit();u(l)}};"";return w});p(b,"Extensions/Data.js",[b["Core/Chart/Chart.js"],b["Core/Globals.js"],b["Core/HttpUtilities.js"],b["Core/Series/Point.js"],b["Core/Series/SeriesRegistry.js"],b["Core/Utilities.js"]],function(b,e,p,H,I,q){var u=e.doc,w=p.ajax,v=I.seriesTypes;p=q.addEvent;var l=q.defined,J=q.extend,E=q.fireEvent,A=q.isNumber,B=q.merge,K=q.objectEach,F=q.pick,L=q.splat;q=function(){function b(a,c,h){this.options=this.rawColumns=this.firstRowAsNames=this.chartOptions=
this.chart=void 0;this.dateFormats={"YYYY/mm/dd":{regex:/^([0-9]{4})[\-\/\.]([0-9]{1,2})[\-\/\.]([0-9]{1,2})$/,parser:function(a){return a?Date.UTC(+a[1],a[2]-1,+a[3]):NaN}},"dd/mm/YYYY":{regex:/^([0-9]{1,2})[\-\/\.]([0-9]{1,2})[\-\/\.]([0-9]{4})$/,parser:function(a){return a?Date.UTC(+a[3],a[2]-1,+a[1]):NaN},alternative:"mm/dd/YYYY"},"mm/dd/YYYY":{regex:/^([0-9]{1,2})[\-\/\.]([0-9]{1,2})[\-\/\.]([0-9]{4})$/,parser:function(a){return a?Date.UTC(+a[3],a[1]-1,+a[2]):NaN}},"dd/mm/YY":{regex:/^([0-9]{1,2})[\-\/\.]([0-9]{1,2})[\-\/\.]([0-9]{2})$/,
parser:function(a){if(!a)return NaN;var c=+a[3];c=c>(new Date).getFullYear()-2E3?c+1900:c+2E3;return Date.UTC(c,a[2]-1,+a[1])},alternative:"mm/dd/YY"},"mm/dd/YY":{regex:/^([0-9]{1,2})[\-\/\.]([0-9]{1,2})[\-\/\.]([0-9]{2})$/,parser:function(a){return a?Date.UTC(+a[3]+2E3,a[1]-1,+a[2]):NaN}}};this.init(a,c,h)}b.prototype.init=function(a,c,h){var d=a.decimalPoint;c&&(this.chartOptions=c);h&&(this.chart=h);"."!==d&&","!==d&&(d=void 0);this.options=a;this.columns=a.columns||this.rowsToColumns(a.rows)||
[];this.firstRowAsNames=F(a.firstRowAsNames,this.firstRowAsNames,!0);this.decimalRegex=d&&new RegExp("^(-?[0-9]+)"+d+"([0-9]+)$");this.rawColumns=[];if(this.columns.length){this.dataFound();var g=!0}this.hasURLOption(a)&&(clearTimeout(this.liveDataTimeout),g=!1);g||(g=this.fetchLiveData());g||(g=!!this.parseCSV().length);g||(g=!!this.parseTable().length);g||(g=this.parseGoogleSpreadsheet());!g&&a.afterComplete&&a.afterComplete()};b.prototype.hasURLOption=function(a){return!(!a||!(a.rowsURL||a.csvURL||
a.columnsURL))};b.prototype.getColumnDistribution=function(){var a=this.chartOptions,c=this.options,h=[],d=function(a){return(v[a||"line"].prototype.pointArrayMap||[0]).length},g=a&&a.chart&&a.chart.type,b=[],m=[],k=0;c=c&&c.seriesMapping||a&&a.series&&a.series.map(function(){return{x:0}})||[];var f;(a&&a.series||[]).forEach(function(a){b.push(d(a.type||g))});c.forEach(function(a){h.push(a.x||0)});0===h.length&&h.push(0);c.forEach(function(c){var h=new G,t=b[k]||d(g),e=(a&&a.series||[])[k]||{},C=
v[e.type||g||"line"].prototype.pointArrayMap,p=C||["y"];(l(c.x)||e.isCartesian||!C)&&h.addColumnReader(c.x,"x");K(c,function(a,c){"x"!==c&&h.addColumnReader(a,c)});for(f=0;f<t;f++)h.hasReader(p[f])||h.addColumnReader(void 0,p[f]);m.push(h);k++});c=v[g||"line"].prototype.pointArrayMap;"undefined"===typeof c&&(c=["y"]);this.valueCount={global:d(g),xColumns:h,individual:b,seriesBuilders:m,globalPointArrayMap:c}};b.prototype.dataFound=function(){this.options.switchRowsAndColumns&&(this.columns=this.rowsToColumns(this.columns));
this.getColumnDistribution();this.parseTypes();!1!==this.parsed()&&this.complete()};b.prototype.parseCSV=function(a){function c(a,c,h,d){function b(c){k=a[c];t=a[c-1];n=a[c+1]}function g(a){l.length<x+1&&l.push([a]);l[x][l[x].length-1]!==a&&l[x].push(a)}function e(){f>r||r>C?(++r,m=""):(!isNaN(parseFloat(m))&&isFinite(m)?(m=parseFloat(m),g("number")):isNaN(Date.parse(m))?g("string"):(m=m.replace(/\//g,"-"),g("date")),p.length<x+1&&p.push([]),h||(p[x][c]=m),m="",++x,++r)}var y=0,k="",t="",n="",m="",
r=0,x=0;if(a.trim().length&&"#"!==a.trim()[0]){for(;y<a.length;y++)if(b(y),'"'===k)for(b(++y);y<a.length&&('"'!==k||'"'===t||'"'===n);){if('"'!==k||'"'===k&&'"'!==t)m+=k;b(++y)}else d&&d[k]?d[k](k,m)&&e():k===q?e():m+=k;e()}}function h(a){var c=0,h=0,d=!1;a.some(function(a,d){var b=!1,g="";if(13<d)return!0;for(var k=0;k<a.length;k++){d=a[k];var e=a[k+1];var t=a[k-1];if("#"===d)break;if('"'===d)if(b){if('"'!==t&&'"'!==e){for(;" "===e&&k<a.length;)e=a[++k];"undefined"!==typeof r[e]&&r[e]++;b=!1}}else b=
!0;else"undefined"!==typeof r[d]?(g=g.trim(),isNaN(Date.parse(g))?!isNaN(g)&&isFinite(g)||r[d]++:r[d]++,g=""):g+=d;","===d&&h++;"."===d&&c++}});d=r[";"]>r[","]?";":",";b.decimalPoint||(b.decimalPoint=c>h?".":",",g.decimalRegex=new RegExp("^(-?[0-9]+)"+b.decimalPoint+"([0-9]+)$"));return d}function d(a,c){var d=[],h=0,k=!1,e=[],t=[],f;if(!c||c>a.length)c=a.length;for(;h<c;h++)if("undefined"!==typeof a[h]&&a[h]&&a[h].length){var n=a[h].trim().replace(/\//g," ").replace(/\-/g," ").replace(/\./g," ").split(" ");
d=["","",""];for(f=0;f<n.length;f++)f<d.length&&(n[f]=parseInt(n[f],10),n[f]&&(t[f]=!t[f]||t[f]<n[f]?n[f]:t[f],"undefined"!==typeof e[f]?e[f]!==n[f]&&(e[f]=!1):e[f]=n[f],31<n[f]?d[f]=100>n[f]?"YY":"YYYY":12<n[f]&&31>=n[f]?(d[f]="dd",k=!0):d[f].length||(d[f]="mm")))}if(k){for(f=0;f<e.length;f++)!1!==e[f]?12<t[f]&&"YY"!==d[f]&&"YYYY"!==d[f]&&(d[f]="YY"):12<t[f]&&"mm"===d[f]&&(d[f]="dd");3===d.length&&"dd"===d[1]&&"dd"===d[2]&&(d[2]="YY");a=d.join("/");return(b.dateFormats||g.dateFormats)[a]?a:(E("deduceDateFailed"),
"YYYY/mm/dd")}return"YYYY/mm/dd"}var g=this,b=a||this.options,e=b.csv;a="undefined"!==typeof b.startRow&&b.startRow?b.startRow:0;var k=b.endRow||Number.MAX_VALUE,f="undefined"!==typeof b.startColumn&&b.startColumn?b.startColumn:0,C=b.endColumn||Number.MAX_VALUE,n=0,l=[],r={",":0,";":0,"\t":0};var p=this.columns=[];e&&b.beforeParse&&(e=b.beforeParse.call(this,e));if(e){e=e.replace(/\r\n/g,"\n").replace(/\r/g,"\n").split(b.lineDelimiter||"\n");if(!a||0>a)a=0;if(!k||k>=e.length)k=e.length-1;if(b.itemDelimiter)var q=
b.itemDelimiter;else q=null,q=h(e);var z=0;for(n=a;n<=k;n++)"#"===e[n][0]?z++:c(e[n],n-a-z);b.columnTypes&&0!==b.columnTypes.length||!l.length||!l[0].length||"date"!==l[0][1]||b.dateFormat||(b.dateFormat=d(p[0]));this.dataFound()}return p};b.prototype.parseTable=function(){var a=this.options,c=a.table,h=this.columns||[],d=a.startRow||0,b=a.endRow||Number.MAX_VALUE,e=a.startColumn||0,m=a.endColumn||Number.MAX_VALUE;c&&("string"===typeof c&&(c=u.getElementById(c)),[].forEach.call(c.getElementsByTagName("tr"),
function(a,c){c>=d&&c<=b&&[].forEach.call(a.children,function(a,b){var g=h[b-e],f=1;if(("TD"===a.tagName||"TH"===a.tagName)&&b>=e&&b<=m)for(h[b-e]||(h[b-e]=[]),h[b-e][c-d]=a.innerHTML;c-d>=f&&void 0===g[c-d-f];)g[c-d-f]=null,f++})}),this.dataFound());return h};b.prototype.fetchLiveData=function(){function a(g){function f(f,k,t){function n(){e&&h.liveDataURL===f&&(c.liveDataTimeout=setTimeout(a,m))}if(!f||!/^(http|\/|\.\/|\.\.\/)/.test(f))return f&&d.error&&d.error("Invalid URL"),!1;g&&(clearTimeout(c.liveDataTimeout),
h.liveDataURL=f);w({url:f,dataType:t||"json",success:function(a){h&&h.series&&k(a);n()},error:function(a,c){3>++b&&n();return d.error&&d.error(c,a)}});return!0}f(k.csvURL,function(a){h.update({data:{csv:a}})},"text")||f(k.rowsURL,function(a){h.update({data:{rows:a}})})||f(k.columnsURL,function(a){h.update({data:{columns:a}})})}var c=this,h=this.chart,d=this.options,b=0,e=d.enablePolling,m=1E3*(d.dataRefreshRate||2),k=B(d);if(!this.hasURLOption(d))return!1;1E3>m&&(m=1E3);delete d.csvURL;delete d.rowsURL;
delete d.columnsURL;a(!0);return this.hasURLOption(d)};b.prototype.parseGoogleSpreadsheet=function(){function a(c){var b=["https://sheets.googleapis.com/v4/spreadsheets",d,"values",m(),"?alt=json&majorDimension=COLUMNS&valueRenderOption=UNFORMATTED_VALUE&dateTimeRenderOption=FORMATTED_STRING&key="+h.googleAPIKey].join("/");w({url:b,dataType:"json",success:function(d){c(d);h.enablePolling&&setTimeout(function(){a(c)},e)},error:function(a,c){return h.error&&h.error(c,a)}})}var c=this,h=this.options,
d=h.googleSpreadsheetKey,b=this.chart,e=Math.max(1E3*(h.dataRefreshRate||2),4E3),m=function(){if(h.googleSpreadsheetRange)return h.googleSpreadsheetRange;var a=("ABCDEFGHIJKLMNOPQRSTUVWXYZ".charAt(h.startColumn||0)||"A")+((h.startRow||0)+1),c="ABCDEFGHIJKLMNOPQRSTUVWXYZ".charAt(F(h.endColumn,-1))||"ZZ";l(h.endRow)&&(c+=h.endRow+1);return a+":"+c};d&&(delete h.googleSpreadsheetKey,a(function(a){a=a.values;if(!a||0===a.length)return!1;var d=a.reduce(function(a,c){return Math.max(a,c.length)},0);a.forEach(function(a){for(var c=
0;c<d;c++)"undefined"===typeof a[c]&&(a[c]=null)});b&&b.series?b.update({data:{columns:a}}):(c.columns=a,c.dataFound())}));return!1};b.prototype.trim=function(a,c){"string"===typeof a&&(a=a.replace(/^\s+|\s+$/g,""),c&&/^[0-9\s]+$/.test(a)&&(a=a.replace(/\s/g,"")),this.decimalRegex&&(a=a.replace(this.decimalRegex,"$1.$2")));return a};b.prototype.parseTypes=function(){for(var a=this.columns,c=a.length;c--;)this.parseColumn(a[c],c)};b.prototype.parseColumn=function(a,c){var b=this.rawColumns,d=this.columns,
g=a.length,e=this.firstRowAsNames,m=-1!==this.valueCount.xColumns.indexOf(c),k,f=[],l=this.chartOptions,n,p=(this.options.columnTypes||[])[c];l=m&&(l&&l.xAxis&&"category"===L(l.xAxis)[0].type||"string"===p);for(b[c]||(b[c]=[]);g--;){var r=f[g]||a[g];var q=this.trim(r);var z=this.trim(r,!0);var u=parseFloat(z);"undefined"===typeof b[c][g]&&(b[c][g]=q);l||0===g&&e?a[g]=""+q:+z===u?(a[g]=u,31536E6<u&&"float"!==p?a.isDatetime=!0:a.isNumeric=!0,"undefined"!==typeof a[g+1]&&(n=u>a[g+1])):(q&&q.length&&
(k=this.parseDate(r)),m&&A(k)&&"float"!==p?(f[g]=r,a[g]=k,a.isDatetime=!0,"undefined"!==typeof a[g+1]&&(r=k>a[g+1],r!==n&&"undefined"!==typeof n&&(this.alternativeFormat?(this.dateFormat=this.alternativeFormat,g=a.length,this.alternativeFormat=this.dateFormats[this.dateFormat].alternative):a.unsorted=!0),n=r)):(a[g]=""===q?null:q,0!==g&&(a.isDatetime||a.isNumeric)&&(a.mixed=!0)))}m&&a.mixed&&(d[c]=b[c]);if(m&&n&&this.options.sort)for(c=0;c<d.length;c++)d[c].reverse(),e&&d[c].unshift(d[c].pop())};
b.prototype.parseDate=function(a){var c=this.options.parseDate,b,d=this.options.dateFormat||this.dateFormat,g;if(c)var e=c(a);else if("string"===typeof a){if(d)(c=this.dateFormats[d])||(c=this.dateFormats["YYYY/mm/dd"]),(g=a.match(c.regex))&&(e=c.parser(g));else for(b in this.dateFormats)if(c=this.dateFormats[b],g=a.match(c.regex)){this.dateFormat=b;this.alternativeFormat=c.alternative;e=c.parser(g);break}g||(a.match(/:.+(GMT|UTC|[Z+-])/)&&(a=a.replace(/\s*(?:GMT|UTC)?([+-])(\d\d)(\d\d)$/,"$1$2:$3").replace(/(?:\s+|GMT|UTC)([+-])/,
"$1").replace(/(\d)\s*(?:GMT|UTC|Z)$/,"$1+00:00")),g=Date.parse(a),"object"===typeof g&&null!==g&&g.getTime?e=g.getTime()-6E4*g.getTimezoneOffset():A(g)&&(e=g-6E4*(new Date(g)).getTimezoneOffset()))}return e};b.prototype.rowsToColumns=function(a){var c,b;if(a){var d=[];var g=a.length;for(c=0;c<g;c++){var e=a[c].length;for(b=0;b<e;b++)d[b]||(d[b]=[]),d[b][c]=a[c][b]}}return d};b.prototype.getData=function(){if(this.columns)return this.rowsToColumns(this.columns).slice(1)};b.prototype.parsed=function(){if(this.options.parsed)return this.options.parsed.call(this,
this.columns)};b.prototype.getFreeIndexes=function(a,c){var b,d=[],g=[];for(b=0;b<a;b+=1)d.push(!0);for(a=0;a<c.length;a+=1){var e=c[a].getReferencedColumnIndexes();for(b=0;b<e.length;b+=1)d[e[b]]=!1}for(b=0;b<d.length;b+=1)d[b]&&g.push(b);return g};b.prototype.complete=function(){var a=this.columns,c,b=this.options,d,e,l=[];if(b.complete||b.afterComplete){if(this.firstRowAsNames)for(d=0;d<a.length;d++)a[d].name=a[d].shift();var m=[];var k=this.getFreeIndexes(a.length,this.valueCount.seriesBuilders);
for(d=0;d<this.valueCount.seriesBuilders.length;d++){var f=this.valueCount.seriesBuilders[d];f.populateColumns(k)&&l.push(f)}for(;0<k.length;){f=new G;f.addColumnReader(0,"x");d=k.indexOf(0);-1!==d&&k.splice(d,1);for(d=0;d<this.valueCount.global;d++)f.addColumnReader(void 0,this.valueCount.globalPointArrayMap[d]);f.populateColumns(k)&&l.push(f)}0<l.length&&0<l[0].readers.length&&(f=a[l[0].readers[0].columnIndex],"undefined"!==typeof f&&(f.isDatetime?c="datetime":f.isNumeric||(c="category")));if("category"===
c)for(d=0;d<l.length;d++)for(f=l[d],k=0;k<f.readers.length;k++)"x"===f.readers[k].configName&&(f.readers[k].configName="name");for(d=0;d<l.length;d++){f=l[d];k=[];for(e=0;e<a[0].length;e++)k[e]=f.read(a,e);m[d]={data:k};f.name&&(m[d].name=f.name);"category"===c&&(m[d].turboThreshold=0)}a={series:m};c&&(a.xAxis={type:c},"category"===c&&(a.xAxis.uniqueNames=!1));b.complete&&b.complete(a);b.afterComplete&&b.afterComplete(a)}};b.prototype.update=function(a,c){var b=this.chart;a&&(a.afterComplete=function(a){a&&
(a.xAxis&&b.xAxis[0]&&a.xAxis.type===b.xAxis[0].options.type&&delete a.xAxis,b.update(a,c,!0))},B(!0,b.options.data,a),this.init(b.options.data))};return b}();e.data=function(b,a,c){return new e.Data(b,a,c)};p(b,"init",function(b){var a=this,c=b.args[0]||{},h=b.args[1];c&&c.data&&!a.hasDataDef&&(a.hasDataDef=!0,a.data=new e.Data(J(c.data,{afterComplete:function(b){var d;if(Object.hasOwnProperty.call(c,"series"))if("object"===typeof c.series)for(d=Math.max(c.series.length,b&&b.series?b.series.length:
0);d--;){var e=c.series[d]||{};c.series[d]=B(e,b&&b.series?b.series[d]:{})}else delete c.series;c=B(b,c);a.init(c,h)}}),c,a),b.preventDefault())});var G=function(){function b(){this.readers=[];this.pointIsArray=!0;this.name=void 0}b.prototype.populateColumns=function(a){var b=!0;this.readers.forEach(function(b){"undefined"===typeof b.columnIndex&&(b.columnIndex=a.shift())});this.readers.forEach(function(a){"undefined"===typeof a.columnIndex&&(b=!1)});return b};b.prototype.read=function(a,b){var c=
this.pointIsArray,d=c?[]:{};this.readers.forEach(function(e){var g=a[e.columnIndex][b];c?d.push(g):0<e.configName.indexOf(".")?H.prototype.setNestedProperty(d,g,e.configName):d[e.configName]=g});if("undefined"===typeof this.name&&2<=this.readers.length){var e=this.getReferencedColumnIndexes();2<=e.length&&(e.shift(),e.sort(function(a,b){return a-b}),this.name=a[e.shift()].name)}return d};b.prototype.addColumnReader=function(a,b){this.readers.push({columnIndex:a,configName:b});"x"!==b&&"y"!==b&&"undefined"!==
typeof b&&(this.pointIsArray=!1)};b.prototype.getReferencedColumnIndexes=function(){var a,b=[];for(a=0;a<this.readers.length;a+=1){var e=this.readers[a];"undefined"!==typeof e.columnIndex&&b.push(e.columnIndex)}return b};b.prototype.hasReader=function(a){var b;for(b=0;b<this.readers.length;b+=1){var e=this.readers[b];if(e.configName===a)return!0}};return b}();e.Data=q;return e.Data});p(b,"masters/modules/data.src.js",[b["Core/Globals.js"],b["Core/HttpUtilities.js"],b["Extensions/Data.js"]],function(b,
e,p){b.ajax=e.ajax;b.getJSON=e.getJSON;b.post=e.post;b.Data=p;b.HttpUtilities=e})});
//# sourceMappingURL=data.js.map
```
