## Supplementary material for "Exploring natural odour landscapes: A case study with implications for human-biting insects": Figure S4: datagrouping.js

```
/*
 Highstock JS v9.3.1 (2021-11-05)

 Data grouping module

 (c) 2010-2021 Torstein Hnsi

 License: www.highcharts.com/license
*/
'use strict';(function(d){"object"===typeof module&&module.exports?(d["default"]=d,module.exports=d):"function"===typeof define&&define.amd?define("highcharts/modules/datagrouping",["highcharts"],function(r){d(r);d.Highcharts=r;return d}):d("undefined"!==typeof Highcharts?Highcharts:void 0)})(function(d){function r(d,r,n,z){d.hasOwnProperty(r)||(d[r]=z.apply(null,n))}d=d?d._modules:{};r(d,"Extensions/DataGrouping.js",[d["Core/Axis/Axis.js"],d["Core/Axis/DateTimeAxis.js"],d["Core/FormatUtilities.js"],
d["Core/Globals.js"],d["Core/Series/Point.js"],d["Core/Series/Series.js"],d["Core/Tooltip.js"],d["Core/DefaultOptions.js"],d["Core/Utilities.js"]],function(d,r,n,z,L,E,M,N,l){var K=n.format,x=E.prototype;n=l.addEvent;var O=l.arrayMax,P=l.arrayMin,Q=l.correctFloat,D=l.defined,G=l.error,R=l.extend,p=l.isNumber,F=l.merge,A=l.pick;"";var g=z.approximations={sum:function(a){var b=a.length;if(!b&&a.hasNulls)var c=null;else if(b)for(c=0;b--;)c+=a[b];return c},average:function(a){var b=a.length;a=g.sum(a);
p(a)&&b&&(a=Q(a/b));return a},averages:function(){var a=[];[].forEach.call(arguments,function(b){a.push(g.average(b))});return"undefined"===typeof a[0]?void 0:a},open:function(a){return a.length?a[0]:a.hasNulls?null:void 0},high:function(a){return a.length?O(a):a.hasNulls?null:void 0},low:function(a){return a.length?P(a):a.hasNulls?null:void 0},close:function(a){return a.length?a[a.length-1]:a.hasNulls?null:void 0},hlc:function(a,b,c){a=g.high(a);b=g.low(b);c=g.close(c);if(p(a)||p(b)||p(c))return[a,
b,c]},ohlc:function(a,b,c,h){a=g.open(a);b=g.high(b);c=g.low(c);h=g.close(h);if(p(a)||p(b)||p(c)||p(h))return[a,b,c,h]},range:function(a,b){a=g.low(a);b=g.high(b);if(p(a)||p(b))return[a,b];if(null===a&&null===b)return null}};l=function(a,b,c,h){var e=this,d=e.data,l=e.options&&e.options.data,f=[],H=[],B=[],t=a.length,k=!!b,m=[],u=e.pointArrayMap,r=u&&u.length,v=["x"].concat(u||["y"]),n=this.options.dataGrouping&&this.options.dataGrouping.groupAll,C=0,x=0,w;h="function"===typeof h?h:g[h]?g[h]:g[e.getDGApproximation&&
e.getDGApproximation()||"average"];r?u.forEach(function(){m.push([])}):m.push([]);var z=r||1;for(w=0;w<=t&&!(a[w]>=c[0]);w++);for(w;w<=t;w++){for(;"undefined"!==typeof c[C+1]&&a[w]>=c[C+1]||w===t;){var q=c[C];e.dataGroupInfo={start:n?x:e.cropStart+x,length:m[0].length};var A=h.apply(e,m);e.pointClass&&!D(e.dataGroupInfo.options)&&(e.dataGroupInfo.options=F(e.pointClass.prototype.optionsToObject.call({series:e},e.options.data[e.cropStart+x])),v.forEach(function(a){delete e.dataGroupInfo.options[a]}));
"undefined"!==typeof A&&(f.push(q),H.push(A),B.push(e.dataGroupInfo));x=w;for(q=0;q<z;q++)m[q].length=0,m[q].hasNulls=!1;C+=1;if(w===t)break}if(w===t)break;if(u){q=e.options.dataGrouping&&e.options.dataGrouping.groupAll?w:e.cropStart+w;A=d&&d[q]||e.pointClass.prototype.applyOptions.apply({series:e},[l[q]]);var y=void 0;for(q=0;q<r;q++)y=A[u[q]],p(y)?m[q].push(y):null===y&&(m[q].hasNulls=!0)}else q=k?b[w]:null,p(q)?m[0].push(q):null===q&&(m[0].hasNulls=!0)}return{groupedXData:f,groupedYData:H,groupMap:B}};
var I={approximations:g,groupData:l},S=x.generatePoints,y={groupPixelWidth:2,dateTimeLabelFormats:{millisecond:["%A, %b %e, %H:%M:%S.%L","%A, %b %e, %H:%M:%S.%L","-%H:%M:%S.%L"],second:["%A, %b %e, %H:%M:%S","%A, %b %e, %H:%M:%S","-%H:%M:%S"],minute:["%A, %b %e, %H:%M","%A, %b %e, %H:%M","-%H:%M"],hour:["%A, %b %e, %H:%M","%A, %b %e, %H:%M","-%H:%M"],day:["%A, %b %e, %Y","%A, %b %e","-%A, %b %e, %Y"],week:["Week from %A, %b %e, %Y","%A, %b %e","-%A, %b %e, %Y"],month:["%B %Y","%B","-%B %Y"],year:["%Y",
"%Y","-%Y"]}},J={line:{},spline:{},area:{},areaspline:{},arearange:{},column:{groupPixelWidth:10},columnrange:{groupPixelWidth:10},candlestick:{groupPixelWidth:10},ohlc:{groupPixelWidth:5},hlc:{groupPixelWidth:5},heikinashi:{groupPixelWidth:10}},T=z.defaultDataGroupingUnits=[["millisecond",[1,2,5,10,20,25,50,100,200,500]],["second",[1,2,5,10,15,30]],["minute",[1,2,5,10,15,30]],["hour",[1,2,3,4,6,8,12]],["day",[1]],["week",[1]],["month",[1,3,6]],["year",null]];x.getDGApproximation=function(){return this.is("arearange")?
"range":this.is("ohlc")?"ohlc":this.is("hlc")?"hlc":this.is("column")?"sum":"average"};x.groupData=l;x.applyGrouping=function(a){var b=this.chart,c=this.options.dataGrouping,h=!1!==this.allowDG&&c&&A(c.enabled,b.options.isStock),e=this.visible||!b.options.chart.ignoreHiddenSeries,d,l=this.currentDataGrouping,f=!1;h&&!this.requireSorting&&(this.requireSorting=f=!0);a=!1===!(this.isCartesian&&!this.isDirty&&!this.xAxis.isDirty&&!this.yAxis.isDirty&&!a)||!h;f&&(this.requireSorting=!1);if(!a){this.destroyGroupedData();
h=c.groupAll?this.xData:this.processedXData;var g=c.groupAll?this.yData:this.processedYData;a=b.plotSizeX;f=this.xAxis;var B=f.options.ordinal,t=this.groupPixelWidth;if(t&&h&&h.length){this.isDirty=d=!0;this.points=null;var k=f.getExtremes();var m=k.min;k=k.max;B=B&&f.ordinal&&f.ordinal.getGroupIntervalFactor(m,k,this)||1;a=f.getTimeTicks(r.Additions.prototype.normalizeTimeTickInterval(t*(k-m)/a*B,c.units||T),Math.min(m,h[0]),Math.max(k,h[h.length-1]),f.options.startOfWeek,h,this.closestPointRange);
t=x.groupData.apply(this,[h,g,a,c.approximation]);h=t.groupedXData;g=t.groupedYData;B=0;c&&c.smoothed&&h.length&&(c.firstAnchor="firstPoint",c.anchor="middle",c.lastAnchor="lastPoint",G(32,!1,b,{"dataGrouping.smoothed":"use dataGrouping.anchor"}));b=h;var u=this.options.dataGrouping;m=this.currentDataGrouping&&this.currentDataGrouping.gapSize;if(u&&this.xData&&m&&this.groupMap){var n=b.length-1;var v=u.anchor;var y=A(u.firstAnchor,v);u=A(u.lastAnchor,v);if(v&&"start"!==v){var C=m*{middle:.5,end:1}[v];
for(v=b.length-1;v--&&0<v;)b[v]+=C}if(y&&"start"!==y&&this.xData[0]>=b[0]){v=this.groupMap[0].start;C=this.groupMap[0].length;var z=void 0;p(v)&&p(C)&&(z=v+(C-1));b[0]={middle:b[0]+.5*m,end:b[0]+m,firstPoint:this.xData[0],lastPoint:z&&this.xData[z]}[y]}u&&"start"!==u&&m&&b[n]>=k-m&&(k=this.groupMap[this.groupMap.length-1].start,b[n]={middle:b[n]+.5*m,end:b[n]+m,firstPoint:k&&this.xData[k],lastPoint:this.xData[this.xData.length-1]}[u])}for(k=1;k<a.length;k++)a.info.segmentStarts&&-1!==a.info.segmentStarts.indexOf(k)||
(B=Math.max(a[k]-a[k-1],B));k=a.info;k.gapSize=B;this.closestPointRange=a.info.totalRange;this.groupMap=t.groupMap;if(e){e=h;if(D(e[0])&&p(f.min)&&p(f.dataMin)&&e[0]<f.min){if(!D(f.options.min)&&f.min<=f.dataMin||f.min===f.dataMin)f.min=Math.min(e[0],f.min);f.dataMin=Math.min(e[0],f.dataMin)}if(D(e[e.length-1])&&p(f.max)&&p(f.dataMax)&&e[e.length-1]>f.max){if(!D(f.options.max)&&p(f.dataMax)&&f.max>=f.dataMax||f.max===f.dataMax)f.max=Math.max(e[e.length-1],f.max);f.dataMax=Math.max(e[e.length-1],f.dataMax)}}c.groupAll&&
(this.allGroupedData=g,c=this.cropData(h,g,f.min,f.max,1),h=c.xData,g=c.yData,this.cropStart=c.start);this.processedXData=h;this.processedYData=g}else this.groupMap=null;this.hasGroupedData=d;this.currentDataGrouping=k;this.preventGraphAnimation=(l&&l.totalRange)!==(k&&k.totalRange)}};x.destroyGroupedData=function(){this.groupedData&&(this.groupedData.forEach(function(a,b){a&&(this.groupedData[b]=a.destroy?a.destroy():null)},this),this.groupedData.length=0)};x.generatePoints=function(){S.apply(this);
this.destroyGroupedData();this.groupedData=this.hasGroupedData?this.points:null};d.prototype.applyGrouping=function(a){var b=this;b.series.forEach(function(c){c.groupPixelWidth=void 0;c.groupPixelWidth=b.getGroupPixelWidth&&b.getGroupPixelWidth();c.groupPixelWidth&&(c.hasProcessed=!0);c.applyGrouping(!!a.hasExtemesChanged)})};d.prototype.getGroupPixelWidth=function(){var a=this.series,b=a.length,c,h=0,e=!1,d;for(c=b;c--;)(d=a[c].options.dataGrouping)&&(h=Math.max(h,A(d.groupPixelWidth,y.groupPixelWidth)));
for(c=b;c--;)if(d=a[c].options.dataGrouping)if(b=(a[c].processedXData||a[c].data).length,a[c].groupPixelWidth||b>this.chart.plotSizeX/h||b&&d.forced)e=!0;return e?h:0};d.prototype.setDataGrouping=function(a,b){var c;b=A(b,!0);a||(a={forced:!1,units:null});if(this instanceof d)for(c=this.series.length;c--;)this.series[c].update({dataGrouping:a},!1);else this.chart.options.series.forEach(function(b){b.dataGrouping=a},!1);this.ordinal&&(this.ordinal.slope=void 0);b&&this.chart.redraw()};n(d,"postProcessData",
d.prototype.applyGrouping);n(L,"update",function(){if(this.dataGroup)return G(24,!1,this.series.chart),!1});n(M,"headerFormatter",function(a){var b=this.chart,c=b.time,d=a.labelConfig,e=d.series,g=e.tooltipOptions,l=e.options.dataGrouping,f=g.xDateFormat,n=e.xAxis,r=g[a.isFooter?"footerFormat":"headerFormat"];if(n&&"datetime"===n.options.type&&l&&p(d.key)){var t=e.currentDataGrouping;l=l.dateTimeLabelFormats||y.dateTimeLabelFormats;if(t)if(g=l[t.unitName],1===t.count)f=g[0];else{f=g[1];var k=g[2]}else!f&&
l&&n.dateTime&&(f=n.dateTime.getXDateFormat(d.x,g.dateTimeLabelFormats));f=c.dateFormat(f,d.key);k&&(f+=c.dateFormat(k,d.key+t.totalRange-1));e.chart.styledMode&&(r=this.styledModeFormat(r));a.text=K(r,{point:R(d.point,{key:f}),series:e},b);a.preventDefault()}});n(E,"destroy",x.destroyGroupedData);n(E,"afterSetOptions",function(a){a=a.options;var b=this.type,c=this.chart.options.plotOptions,d=N.defaultOptions.plotOptions[b].dataGrouping,e=this.useCommonDataGrouping&&y;if(J[b]||e){d||(d=F(y,J[b]));
var g=this.chart.rangeSelector;a.dataGrouping=F(e,d,c.series&&c.series.dataGrouping,c[b].dataGrouping,this.userOptions.dataGrouping,!a.isInternal&&g&&p(g.selected)&&g.buttonOptions[g.selected].dataGrouping)}});n(d,"afterSetScale",function(){this.series.forEach(function(a){a.hasProcessed=!1})});z.dataGrouping=I;"";return I});r(d,"masters/modules/datagrouping.src.js",[d["Extensions/DataGrouping.js"]],function(d){return d})});
//# sourceMappingURL=datagrouping.js.map
```
