## Supplementary material for "Exploring natural odour landscapes: A case study with implications for human-biting insects": Figure S4: debugger.js

```
/*
 Highcharts JS v9.3.1 (2021-11-05)

 Debugger module

 (c) 2012-2021 Torstein Honsi

 License: www.highcharts.com/license
*/
'use strict';(function(a){"object"===typeof module&&module.exports?(a["default"]=a,module.exports=a):"function"===typeof define&&define.amd?define("highcharts/modules/debugger",["highcharts"],function(b){a(b);a.Highcharts=b;return a}):a("undefined"!==typeof Highcharts?Highcharts:void 0)})(function(a){function b(a,b,g,d){a.hasOwnProperty(b)||(a[b]=d.apply(null,g))}a=a?a._modules:{};b(a,"Extensions/Debugger/ErrorMessages.js",[],function(){return{10:{title:"Can't plot zero or subzero values on a logarithmic axis",
text:'<h1>Can\'t plot zero or subzero values on a logarithmic axis</h1><p>This error occurs in the following situations: </p><ul><li>If a zero or subzero data value is added to a logarithmic axis</li><li>If the minimum of a logarithimic axis is set to 0 or less</li><li>If the threshold is set to 0 or less</li></ul><p>Note: As of Highcharts 5.0.8 it\'s possible to bypass this error message by setting <code>Axis.prototype.allowNegativeLog</code> to true, and add custom conversion functions. <a href="https://jsfiddle.net/gh/get/library/pure/highcharts/highcharts/tree/master/samples/highcharts/yaxis/type-log-negative/">View live demo</a>. It is also possible to use a similar workaround for colorAxis. <a href="https://jsfiddle.net/gh/get/library/pure/highcharts/highcharts/tree/master/samples/highcharts/coloraxis/logarithmic-with-emulate-negative-values/">View live demo</a>.</p>',
enduser:'<h1>Can\'t plot zero or subzero values on a logarithmic axis</h1><p>This error occurs in the following situations:</p><ul><li>If a zero or subzero data value is added to a logarithmic axis</li><li>If the minimum of a logarithimic axis is set to 0 or less</li><li>If the threshold is set to 0 or less</li></ul><p>As of Highcharts 5.0.8 it\'s possible to bypass this error message by setting <code>Axis.prototype.allowNegativeLog</code> to <code>true</code> and add custom conversion functions. <a href="http://jsfiddle.net/gh/get/library/pure/highcharts/highcharts/tree/master/samples/highcharts/yaxis/type-log-negative/">">View Live Demo</a>. It is also possible to use a similar workaround for colorAxis. <a href="https://jsfiddle.net/gh/get/library/pure/highcharts/highcharts/tree/master/samples/highcharts/coloraxis/logarithmic-with-emulate-negative-values/">View live demo</a>.</p>'},
11:{title:"Can't link axes of different type",text:"<h1>Can't link axes of different type</h1><p>This error occurs when using the <code>linkedTo</code> option to link two axes of different types, for example a logarithmic axis to a linear axis. Highcharts can't link these because the calculation of ticks, extremes, padding etc. is different.</p>"},12:{title:"Highcharts expects point configuration to be numbers or arrays in turbo mode",text:'<h1>Highcharts expects point configuration to be numbers or arrays in turbo mode</h1><p>This error occurs if the <code>series.data</code> option contains object configurations and the number of points exceeds the turboThreshold. It can be fixed by either setting <code>turboThreshold</code> to a higher value, or changing the point configurations to numbers or arrays.</p><p>In boost mode, turbo mode is always on, which means only array of numbers or two dimensional arrays are allowed.</p><p>See <a href="https://api.highcharts.com/highcharts#plotOptions.series.turboThreshold">plotOptions.series.turboThreshold</a></p>'},
13:{title:"Rendering div not found",text:'<h1>Rendering div not found</h1><p>This error occurs if the <a href="https://api.highcharts.com/highcharts#chart.renderTo">chart.renderTo</a> option is misconfigured so that Highcharts is unable to find the HTML element to render the chart in.</p><p>If using a DOM ID when creating the chart, make sure a node with the same ID exists somewhere in the DOM.</p>'},14:{title:"String value sent to series.data, expected Number",text:"<h1>String value sent to series.data, expected Number</h1><p>This happens if using a string as a data point, for example in a setup like this: </p><pre>series: [{\n    data: [&quot;3&quot;, &quot;5&quot;, &quot;1&quot;, &quot;6&quot;]\n}]</pre><p>Highcharts expects numerical data values.</p><p>The most common reason for this error this is that data is parsed from CSV or from a XML source, and the implementer forgot to run <code>parseFloat</code> on the parsed value.</p><p>Note: For performance reasons internal type casting is not performed, and only the first value is checked (since 2.3).</p>"},
15:{title:"Highcharts expects data to be sorted",text:"<h1>Highcharts expects data to be sorted</h1><p>This happens when creating a line series or a stock chart where the data is not sorted in ascending X order.</p><p>For performance reasons, Highcharts does not sort the data, instead it requires that the implementer pre-sorts the data.</p>"},16:{title:"Highcharts already defined in the page",text:"<h1>Highcharts already defined in the page</h1><p>This error happens if the <code>Highcharts</code> namespace already exists when loading Highcharts or Highstock.</p><p>This is caused by including Highcharts or Highstock more than once.</p><p>Keep in mind that the <code>Highcharts.Chart</code> constructor and all features of Highcharts are included in Highstock, so if using the <code>Chart</code> and <code>StockChart</code> constructors in combination, only the <code>highstock.js</code> file is required.</p>"},
17:{title:"The requested series type does not exist",text:"<h1>The requested series type does not exist</h1><p>This error happens when setting <code>chart.type</code> or <code>series.type</code> to a series type that isn't defined in Highcharts. A typical reason may be that the module or extension where the series type is defined isn't included.</p><p>For example in order to create an <code>arearange</code> series, the <code>highcharts-more.js</code> file must be loaded.</p>"},18:{title:"The requested axis does not exist",
text:"<h1>The requested axis does not exist</h1><p>This error happens when setting a series' <code>xAxis</code> or <code>yAxis</code> property to point to an axis that does not exist.</p>"},19:{title:"Too many ticks",text:"<h1>Too many ticks</h1><p>This error happens when applying too many ticks to an axis, specifically when adding more ticks than the axis pixel length.</p><p>With default value this won't happen, but there are edge cases, for example when setting axis categories and <code>xAxis.labels.step</code> in combination with a long data range, when the axis is instructed to create a great number of ticks.</p>"},
20:{title:"Can't add object point configuration to a long data series",text:"<h1>Can't add object point configuration to a long data series</h1><p>In Highstock, when trying to add a point using the object literal configuration syntax, it will only work when the number of data points is below the series' <a href=\"https://api.highcharts.com/highstock#plotOptions.series.turboThreshold\">turboThreshold</a>. Instead of the object syntax, use the Array syntax.</p>"},21:{title:"Can't find Proj4js library",
text:'<h1>Can\'t find Proj4js library</h1><p>Using latitude/longitude functionality in Highmaps requires the <a href="http://proj4js.org">Proj4js</a> library to be loaded.</p>'},22:{title:"Map does not support latitude/longitude",text:'<h1>Map does not support latitude/longitude</h1><p>The loaded map does not support latitude/longitude functionality. This is only supported with maps from the <a href="https://code.highcharts.com/mapdata">official Highmaps map collection</a> from version 1.1.0 onwards. If you are using a custom map, consider using the <a href="https://proj4js.org">Proj4js</a> library to convert between projections.</p>'},
23:{title:"Unsupported color format used for color interpolation",text:"<h1>Unsupported color format used for color interpolation</h1><p>Highcharts supports three color formats primarily: hex (<code>#FFFFFF</code>), rgb (<code>rgba(255,255,255)</code>) and rgba (<code>rgba(255,255,255,1)</code>). If any other format, like 3-digit colors (<code>#FFF</code>), named colors (<code>white</code>) or gradient structures are used in for example a heatmap, Highcharts will fail to interpolate and will instead use the end-color with no interpolation applied.</p><p>We've chosen to preserve this limitation in order to keep the weight of the implementation at a minimum.</p>"},
24:{title:"Cannot run Point.update on a grouped point",text:"<h1>Cannot run Point.update on a grouped point</h1><p>Running <code>Point.update</code> in Highstock when a point is grouped by data grouping is not supported.</p><p>This is not supported because when data grouping is enabled, there won't be any references to the raw points, which is required by the <code>Point.update</code> function.</p>"},25:{title:"Can't find Moment.js library",text:'<h1>Can\'t find Moment.js library</h1><p>Using the global.timezone option requires the <a href="https://momentjs.com/">Moment.js</a> library to be loaded.</p>'},
26:{title:"WebGL not supported, and no fallback module included",text:"<h1>WebGL not supported, and no fallback module included</h1><p>This happens when the browser doesn't support WebGL,<b>and</b> the canvas fallback module (<code>boost-canvas.js</code>) hasn't been included OR if the fallback module was included<b>after</b> the boost module.</p><p>If a fallback is required, make sure to include <code>boost-canvas.js</code>, and that it's included before <code>boost.js</code>.</p><p>Please note that the fallback module is not intended as a fully-featured one. Rather, it's a minimal implementation of the WebGL counterpart.</p>"},
27:{title:"This browser does not support SVG",text:"<h1>This browser does not support SVG</h1><p>This happens in old IE when the <code>oldie.js</code> module isn't loaded.</p><p>If compatibility with IE versions 6, 7 and 8 is required, add the module after loading <code>highcharts.js</code>. In a website context, it's a good idea to load it in a conditional comment to avoid traffic overhead and dead code in modern browsers: </p><pre>&lt;!--[if lt IE 9]&gt;\n    &lt;script src='https://code.highcharts.com/modules/oldie.js'&gt;&lt;/script&gt;\n&lt;![endif]--&gt;</pre>"},
28:{title:"Fallback to export server disabled",text:'<h1>Fallback to export server disabled</h1><p>This happens when the offline export module encounters a chart that it can\'t export successfully, and the fallback to the online export server is disabled. The offline exporting module will fail for certain browsers, and certain features (e.g. <a href="https://api.highcharts.com/highcharts/exporting.allowHTML">exporting.allowHTML</a> ), depending on the type of image exporting to. For a compatibility overview, see <a href="https://www.highcharts.com/docs/export-module/client-side-export">Client Side Export</a>.</p><p>For very complex charts, it\'s possible that exporting fail in browsers that don\'t support Blob objects, due to data URL length limits. It\'s always recommended to define the <a href="https://api.highcharts.com/highcharts/exporting.error">exporting.error</a> callback when disabling the fallback, so that details can be provided to the end-user if offline export isn\'t working for them.</p>'},
29:{title:"Browser does not support WebAudio",text:"<h1>Browser does not support WebAudio</h1><p>This happens when you attempt to use the sonification module on a chart in a browser or environment that does not support the WebAudio API. This API is supported on all modern browsers, including Microsoft Edge, Google Chrome and Mozilla Firefox.</p>"},30:{title:"Invalid instrument",text:"<h1>Invalid instrument</h1><p>This happens when you try to use a sonification instrument that is not valid. If you are using a predefined instrument, make sure your spelling is correct.</p>"},
31:{title:"Non-unique point or node id",text:"<h1>Non-unique point or node id</h1><p>This error occurs when using the same <code>id</code> for two or more points or nodes.</p>"},32:{title:"Deprecated function or property",text:'<h1>Deprecated function or property</h1><p>This error occurs when using a deprecated function or property. Consult the <a href="https://api.highcharts.com/">API documentation</a> for alternatives, if no replacement is mentioned by the error itself.</p>'}}});b(a,"Extensions/Debugger/Debugger.js",
[a["Core/Chart/Chart.js"],a["Extensions/Debugger/ErrorMessages.js"],a["Core/Globals.js"],a["Core/DefaultOptions.js"],a["Core/Utilities.js"]],function(a,b,g,d,h){var l=g.charts;d=d.setOptions;var k=h.addEvent,m=h.find,n=h.isNumber;d({chart:{displayErrors:!0}});k(g,"displayError",function(a){var c=a.chart||m(l.slice().reverse(),function(a){return!!a});if(c){var e=a.code,f=c.options.chart;a=c.renderer;c.errorElements&&c.errorElements.forEach(function(a){a&&a.destroy()});if(f&&f.displayErrors&&a){c.errorElements=
[];e=n(e)?"Highcharts error #"+e+": "+b[e].text:e;f=c.chartWidth;var d=c.chartHeight;e=e.replace(/<h1>(.*)<\/h1>/g,'<br><span style="font-size: 24px">$1</span><br>').replace(/<p>/g,"").replace(/<\/p>/g,"<br>");c.errorElements[0]=a.rect(2,2,f-4,d-4).attr({"stroke-width":4,stroke:"#ff0000",zIndex:3}).add();c.errorElements[1]=a.label(e,0,0,"rect",void 0,void 0,void 0,void 0,"debugger").css({color:"#ffffff",width:f-16+"px",padding:0}).attr({fill:"rgba(255, 0, 0, 0.9)",width:f,padding:8,zIndex:10}).add();
c.errorElements[1].attr({y:d-c.errorElements[1].getBBox().height})}}});k(a,"beforeRedraw",function(){var a=this.errorElements;a&&a.length&&a.forEach(function(a){a.destroy()});delete this.errorElements})});b(a,"masters/modules/debugger.src.js",[a["Core/Globals.js"],a["Extensions/Debugger/ErrorMessages.js"]],function(a,b){a.errorMessages=b})});
//# sourceMappingURL=debugger.js.map
```
