## Supplementary material for "Exploring natural odour landscapes: A case study with implications for human-biting insects": Figure S4: dependency-wheel.js

```
/*
 Highcharts JS v9.3.1 (2021-11-05)

 Dependency wheel module

 (c) 2010-2021 Torstein Honsi

 License: www.highcharts.com/license
*/
'use strict';(function(b){"object"===typeof module&&module.exports?(b["default"]=b,module.exports=b):"function"===typeof define&&define.amd?define("highcharts/modules/dependency-wheel",["highcharts","highcharts/modules/sankey"],function(f){b(f);b.Highcharts=f;return b}):b("undefined"!==typeof Highcharts?Highcharts:void 0)})(function(b){function f(b,f,e,k){b.hasOwnProperty(f)||(b[f]=k.apply(null,e))}b=b?b._modules:{};f(b,"Series/DependencyWheel/DependencyWheelPoint.js",[b["Core/Series/SeriesRegistry.js"],
b["Core/Utilities.js"]],function(b,f){var e=this&&this.__extends||function(){var b=function(a,c){b=Object.setPrototypeOf||{__proto__:[]}instanceof Array&&function(b,c){b.__proto__=c}||function(b,c){for(var a in c)c.hasOwnProperty(a)&&(b[a]=c[a])};return b(a,c)};return function(a,c){function f(){this.constructor=a}b(a,c);a.prototype=null===c?Object.create(c):(f.prototype=c.prototype,new f)}}();return function(b){function a(){var c=null!==b&&b.apply(this,arguments)||this;c.angle=void 0;c.fromNode=void 0;
c.index=void 0;c.linksFrom=void 0;c.linksTo=void 0;c.options=void 0;c.series=void 0;c.shapeArgs=void 0;c.toNode=void 0;return c}e(a,b);a.prototype.getDataLabelPath=function(b){var c=this.series.chart.renderer,a=this.shapeArgs,f=0>this.angle||this.angle>Math.PI,e=a.start||0,p=a.end||0;this.dataLabelPath||(this.dataLabelPath=c.arc({open:!0,longArc:Math.abs(Math.abs(e)-Math.abs(p))<Math.PI?0:1}).add(b));this.dataLabelPath.attr({x:a.x,y:a.y,r:a.r+(this.dataLabel.options.distance||0),start:f?e:p,end:f?
p:e,clockwise:+f});return this.dataLabelPath};a.prototype.isValid=function(){return!0};return a}(b.seriesTypes.sankey.prototype.pointClass)});f(b,"Series/DependencyWheel/DependencyWheelSeries.js",[b["Core/Animation/AnimationUtilities.js"],b["Series/DependencyWheel/DependencyWheelPoint.js"],b["Core/Globals.js"],b["Core/Series/SeriesRegistry.js"],b["Core/Utilities.js"]],function(b,f,e,k,a){var c=this&&this.__extends||function(){var b=function(a,c){b=Object.setPrototypeOf||{__proto__:[]}instanceof Array&&
function(b,a){b.__proto__=a}||function(b,a){for(var c in a)a.hasOwnProperty(c)&&(b[c]=a[c])};return b(a,c)};return function(a,c){function l(){this.constructor=a}b(a,c);a.prototype=null===c?Object.create(c):(l.prototype=c.prototype,new l)}}(),q=b.animObject,r=e.deg2rad;e=k.seriesTypes;b=e.pie;var n=e.sankey;e=a.extend;var t=a.merge;a=function(b){function a(){var a=null!==b&&b.apply(this,arguments)||this;a.data=void 0;a.options=void 0;a.nodeColumns=void 0;a.nodes=void 0;a.points=void 0;return a}c(a,
b);a.prototype.animate=function(b){if(!b){var a=q(this.options.animation).duration/2/this.nodes.length;this.nodes.forEach(function(b,c){var l=b.graphic;l&&(l.attr({opacity:0}),setTimeout(function(){b.graphic&&b.graphic.animate({opacity:1},{duration:a})},a*c))},this);this.points.forEach(function(b){var a=b.graphic;!b.isNode&&a&&a.attr({opacity:0}).animate({opacity:1},this.options.animation)},this)}};a.prototype.createNode=function(b){var a=n.prototype.createNode.call(this,b);a.index=this.nodes.length-
1;a.getSum=function(){return a.linksFrom.concat(a.linksTo).reduce(function(a,b){return a+b.weight},0)};a.offset=function(b){function c(b){return b.fromNode===a?b.toNode:b.fromNode}var f=0,d,g=a.linksFrom.concat(a.linksTo);g.sort(function(b,a){return c(b).index-c(a).index});for(d=0;d<g.length;d++)if(c(g[d]).index>a.index){g=g.slice(0,d).reverse().concat(g.slice(d).reverse());var e=!0;break}e||g.reverse();for(d=0;d<g.length;d++){if(g[d]===b)return f;f+=g[d].weight}};return a};a.prototype.createNodeColumns=
function(){var b=[this.createNodeColumn()];this.nodes.forEach(function(a){a.column=0;b[0].push(a)});return b};a.prototype.getNodePadding=function(){return this.options.nodePadding/Math.PI};a.prototype.translate=function(){var b=this.options,a=2*Math.PI/(this.chart.plotHeight+this.getNodePadding()),c=this.getCenter(),f=(b.startAngle-90)*r;n.prototype.translate.call(this);this.nodeColumns[0].forEach(function(e){if(e.sum){var d=e.shapeArgs,g=c[0],l=c[1],k=c[2]/2,h=k-b.nodeWidth,m=f+a*(d.y||0);d=f+a*
((d.y||0)+(d.height||0));e.angle=m+(d-m)/2;e.shapeType="arc";e.shapeArgs={x:g,y:l,r:k,innerR:h,start:m,end:d};e.dlBox={x:g+Math.cos((m+d)/2)*(k+h)/2,y:l+Math.sin((m+d)/2)*(k+h)/2,width:1,height:1};e.linksFrom.forEach(function(c){if(c.linkBase){var e,d=c.linkBase.map(function(d,k){d*=a;var m=Math.cos(f+d)*(h+1),n=Math.sin(f+d)*(h+1),p=b.curveFactor;e=Math.abs(c.linkBase[3-k]*a-d);e>Math.PI&&(e=2*Math.PI-e);e*=h;e<h&&(p*=e/h);return{x:g+m,y:l+n,cpX:g+(1-p)*m,cpY:l+(1-p)*n}});c.shapeArgs={d:[["M",d[0].x,
d[0].y],["A",h,h,0,0,1,d[1].x,d[1].y],["C",d[1].cpX,d[1].cpY,d[2].cpX,d[2].cpY,d[2].x,d[2].y],["A",h,h,0,0,1,d[3].x,d[3].y],["C",d[3].cpX,d[3].cpY,d[0].cpX,d[0].cpY,d[0].x,d[0].y]]}}})}})};a.defaultOptions=t(n.defaultOptions,{center:[null,null],curveFactor:.6,startAngle:0});return a}(n);e(a.prototype,{orderNodes:!1,getCenter:b.prototype.getCenter});a.prototype.pointClass=f;k.registerSeriesType("dependencywheel",a);"";return a});f(b,"masters/modules/dependency-wheel.src.js",[],function(){})});
//# sourceMappingURL=dependency-wheel.js.map
```
