## Supplementary material for "Exploring natural odour landscapes: A case study with implications for human-biting insects": Figure S4: dotplot.js

```
/*
 Highcharts JS v9.3.1 (2021-11-05)

 Dot plot series type for Highcharts

 (c) 2010-2021 Torstein Honsi

 License: www.highcharts.com/license
*/
'use strict';(function(a){"object"===typeof module&&module.exports?(a["default"]=a,module.exports=a):"function"===typeof define&&define.amd?define("highcharts/modules/dotplot",["highcharts"],function(b){a(b);a.Highcharts=b;return a}):a("undefined"!==typeof Highcharts?Highcharts:void 0)})(function(a){function b(a,b,c,f){a.hasOwnProperty(b)||(a[b]=f.apply(null,c))}a=a?a._modules:{};b(a,"Series/DotPlot/DotPlotSeries.js",[a["Series/Column/ColumnSeries.js"],a["Core/Series/SeriesRegistry.js"],a["Core/Utilities.js"]],
function(a,b,c){var f=this&&this.__extends||function(){var a=function(b,e){a=Object.setPrototypeOf||{__proto__:[]}instanceof Array&&function(a,e){a.__proto__=e}||function(a,e){for(var b in e)e.hasOwnProperty(b)&&(a[b]=e[b])};return a(b,e)};return function(b,e){function c(){this.constructor=b}a(b,e);b.prototype=null===e?Object.create(e):(c.prototype=e.prototype,new c)}}(),q=c.extend,m=c.merge,x=c.objectEach,r=c.pick;c=function(b){function c(){var a=null!==b&&b.apply(this,arguments)||this;a.data=void 0;
a.options=void 0;a.points=void 0;return a}f(c,b);c.prototype.drawPoints=function(){var a=this,b=a.chart.renderer,c=this.options.marker,f=this.yAxis.transA*a.options.itemPadding,t=this.borderWidth%2?.5:1;this.points.forEach(function(d){var e;var g=d.marker||{};var u=g.symbol||c.symbol,m=r(g.radius,c.radius),v="rect"!==u;d.graphics=e=d.graphics||{};var l=d.pointAttr?d.pointAttr[d.selected?"selected":""]||a.pointAttr[""]:a.pointAttribs(d,d.selected&&"select");delete l.r;a.chart.styledMode&&(delete l.stroke,
delete l["stroke-width"]);if(null!==d.y){d.graphic||(d.graphic=b.g("point").add(a.group));var k=d.y;var w=r(d.stackY,d.y);var n=Math.min(d.pointWidth,a.yAxis.transA-f);for(g=w;g>w-d.y;g--){var h=d.barX+(v?d.pointWidth/2-n/2:0);var p=a.yAxis.toPixels(g,!0)+f/2;a.options.crisp&&(h=Math.round(h)-t,p=Math.round(p)+t);h={x:h,y:p,width:Math.round(v?n:d.pointWidth),height:Math.round(n),r:m};e[k]?e[k].animate(h):e[k]=b.symbol(u).attr(q(h,l)).add(d.graphic);e[k].isActive=!0;k--}}x(e,function(a,b){a.isActive?
a.isActive=!1:(a.destroy(),delete a[b])})})};c.defaultOptions=m(a.defaultOptions,{itemPadding:.2,marker:{symbol:"circle",states:{hover:{},select:{}}}});return c}(a);q(c.prototype,{markerAttribs:void 0});b.registerSeriesType("dotplot",c);return c});b(a,"masters/modules/dotplot.src.js",[],function(){})});
//# sourceMappingURL=dotplot.js.map
```
