## Supplementary material for "Exploring natural odour landscapes: A case study with implications for human-biting insects": Figure S4: drag-panes.js

```
/*
 Highstock JS v9.3.1 (2021-11-05)

 Drag-panes module

 (c) 2010-2021 Highsoft AS
 Author: Kacper Madej

 License: www.highcharts.com/license
*/
'use strict';(function(a){"object"===typeof module&&module.exports?(a["default"]=a,module.exports=a):"function"===typeof define&&define.amd?define("highcharts/modules/drag-panes",["highcharts","highcharts/modules/stock"],function(b){a(b);a.Highcharts=b;return a}):a("undefined"!==typeof Highcharts?Highcharts:void 0)})(function(a){function b(a,m,b,n){a.hasOwnProperty(m)||(a[m]=n.apply(null,b))}a=a?a._modules:{};b(a,"Extensions/DragPanes.js",[a["Core/Globals.js"],a["Core/Axis/Axis.js"],a["Core/Axis/AxisDefaults.js"],
a["Core/Pointer.js"],a["Core/Utilities.js"]],function(a,b,x,n,e){var m=a.hasTouch,f=e.addEvent,t=e.clamp,y=e.isNumber,z=e.merge,A=e.objectEach,v=e.relativeLength;e=e.wrap;var u=function(){function a(c){this.options=this.lastPos=this.controlLine=this.axis=void 0;this.init(c)}a.prototype.init=function(c,a){this.axis=c;this.options=c.options.resize;this.render();a||this.addMouseEvents()};a.prototype.render=function(){var c=this.axis,a=c.chart,d=this.options,w=d.x||0,b=d.y,k=t(c.top+c.height+b,a.plotTop,
a.plotTop+a.plotHeight),l={};a.styledMode||(l={cursor:d.cursor,stroke:d.lineColor,"stroke-width":d.lineWidth,dashstyle:d.lineDashStyle});this.lastPos=k-b;this.controlLine||(this.controlLine=a.renderer.path().addClass("highcharts-axis-resizer"));this.controlLine.add(c.axisGroup);d=a.styledMode?this.controlLine.strokeWidth():d.lineWidth;l.d=a.renderer.crispLine([["M",c.left+w,k],["L",c.left+c.width+w,k]],d);this.controlLine.attr(l)};a.prototype.addMouseEvents=function(){var c=this,a=c.controlLine.element,
d=c.axis.chart.container,b=[],e,k,l;c.mouseMoveHandler=e=function(a){c.onMouseMove(a)};c.mouseUpHandler=k=function(a){c.onMouseUp(a)};c.mouseDownHandler=l=function(a){c.onMouseDown(a)};b.push(f(d,"mousemove",e),f(d.ownerDocument,"mouseup",k),f(a,"mousedown",l));m&&b.push(f(d,"touchmove",e),f(d.ownerDocument,"touchend",k),f(a,"touchstart",l));c.eventsToUnbind=b};a.prototype.onMouseMove=function(a){a.touches&&0===a.touches[0].pageX||!this.grabbed||(this.hasDragged=!0,this.updateAxes(this.axis.chart.pointer.normalize(a).chartY-
this.options.y))};a.prototype.onMouseUp=function(a){this.hasDragged&&this.updateAxes(this.axis.chart.pointer.normalize(a).chartY-this.options.y);this.grabbed=this.hasDragged=this.axis.chart.activeResizer=null};a.prototype.onMouseDown=function(a){this.axis.chart.pointer.reset(!1,0);this.grabbed=this.axis.chart.activeResizer=!0};a.prototype.updateAxes=function(a){var c=this,d=c.axis.chart,b=c.options.controlledAxis,e=0===b.next.length?[d.yAxis.indexOf(c.axis)+1]:b.next;b=[c.axis].concat(b.prev);var k=
[],l=!1,q=d.plotTop,f=d.plotHeight,m=q+f;a=t(a,q,m);var r=a-c.lastPos;1>r*r||([b,e].forEach(function(b,e){b.forEach(function(b,h){var g=(b=y(b)?d.yAxis[b]:e||h?d.get(b):b)&&b.options;if(g&&"navigator-y-axis"!==g.id){h=b.top;var n=Math.round(v(g.minLength,f));var p=Math.round(v(g.maxLength,f));e?(r=a-c.lastPos,g=Math.round(t(b.len-r,n,p)),h=b.top+r,h+g>m&&(p=m-g-h,a+=p,h+=p),h<q&&(h=q,h+g>m&&(g=f)),g===n&&(l=!0),k.push({axis:b,options:{top:100*(h-q)/f+"%",height:100*g/f+"%"}})):(g=Math.round(t(a-h,
n,p)),g===p&&(l=!0),a=h+g,k.push({axis:b,options:{height:100*g/f+"%"}}))}})}),l||(k.forEach(function(a){a.axis.update(a.options,!1)}),d.redraw(!1)))};a.prototype.destroy=function(){var a=this;delete a.axis.resizer;this.eventsToUnbind&&this.eventsToUnbind.forEach(function(a){a()});a.controlLine.destroy();A(a,function(b,c){a[c]=null})};a.resizerOptions={minLength:"10%",maxLength:"100%",resize:{controlledAxis:{next:[],prev:[]},enabled:!1,cursor:"ns-resize",lineColor:"#cccccc",lineDashStyle:"Solid",lineWidth:4,
x:0,y:0}};return a}();b.keepProps.push("resizer");f(b,"afterRender",function(){var a=this.resizer,b=this.options.resize;b&&(b=!1!==b.enabled,a?b?a.init(this,!0):a.destroy():b&&(this.resizer=new u(this)))});f(b,"destroy",function(a){!a.keepEvents&&this.resizer&&this.resizer.destroy()});e(n.prototype,"runPointActions",function(a){this.chart.activeResizer||a.apply(this,Array.prototype.slice.call(arguments,1))});e(n.prototype,"drag",function(a){this.chart.activeResizer||a.apply(this,Array.prototype.slice.call(arguments,
1))});z(!0,x.defaultYAxisOptions,u.resizerOptions);a.AxisResizer=u;return a.AxisResizer});b(a,"masters/modules/drag-panes.src.js",[],function(){})});
//# sourceMappingURL=drag-panes.js.map
```
