## Supplementary material for "Exploring natural odour landscapes: A case study with implications for human-biting insects": Figure S4: draggable-points.js

```
/*
 Highcharts JS v9.3.1 (2021-11-05)

 (c) 2009-2021 Torstein Honsi

 License: www.highcharts.com/license
*/
'use strict';(function(e){"object"===typeof module&&module.exports?(e["default"]=e,module.exports=e):"function"===typeof define&&define.amd?define("highcharts/modules/draggable-points",["highcharts"],function(l){e(l);e.Highcharts=l;return e}):e("undefined"!==typeof Highcharts?Highcharts:void 0)})(function(e){function l(e,z,l,v){e.hasOwnProperty(z)||(e[z]=v.apply(null,l))}e=e?e._modules:{};l(e,"Extensions/DraggablePoints.js",[e["Core/Animation/AnimationUtilities.js"],e["Core/Chart/Chart.js"],e["Core/Globals.js"],
e["Core/Series/Point.js"],e["Core/Series/Series.js"],e["Core/Series/SeriesRegistry.js"],e["Core/Utilities.js"]],function(e,l,L,v,E,w,p){function z(a){return{left:"right",right:"left",top:"bottom",bottom:"top"}[a]}function M(a){var b=["draggableX","draggableY"],c;n(a.dragDropProps,function(a){a.optionName&&b.push(a.optionName)});for(c=b.length;c--;)if(a.options.dragDrop[b[c]])return!0}function N(a){var b=a.series?a.series.length:0;if(a.hasCartesianSeries&&!a.polar)for(;b--;)if(a.series[b].options.dragDrop&&
M(a.series[b]))return!0}function O(a){var b=a.series,c=b.options.dragDrop||{};a=a.options&&a.options.dragDrop;var d,f;n(b.dragDropProps,function(a){"x"===a.axis&&a.move?d=!0:"y"===a.axis&&a.move&&(f=!0)});return(c.draggableX&&d||c.draggableY&&f)&&!(a&&!1===a.draggableX&&!1===a.draggableY)&&b.yAxis&&b.xAxis}function A(a,b){return"undefined"===typeof a.chartX||"undefined"===typeof a.chartY?b.pointer.normalize(a):a}function B(a,b,c,d){var f=b.map(function(b){return r(a,b,c,d)});return function(){f.forEach(function(a){a()})}}
function P(a,b,c){var d=b.dragDropData.origin;b=d.chartX;d=d.chartY;var f=a.chartX;a=a.chartY;return Math.sqrt((f-b)*(f-b)+(a-d)*(a-d))>c}function Q(a,b,c){var d={chartX:a.chartX,chartY:a.chartY,guideBox:c&&{x:c.attr("x"),y:c.attr("y"),width:c.attr("width"),height:c.attr("height")},points:{}};b.forEach(function(b){var c={};n(b.series.dragDropProps,function(d,f){d=b.series[d.axis+"Axis"];c[f]=b[f];c[f+"Offset"]=d.toPixels(b[f])-(d.horiz?a.chartX:a.chartY)});c.point=b;d.points[b.id]=c});return d}function R(a){var b=
a.series,c=[],d=b.options.dragDrop.groupBy;b.isSeriesBoosting?b.options.data.forEach(function(a,d){c.push((new b.pointClass).init(b,a));c[c.length-1].index=d}):c=b.points;return a.options[d]?c.filter(function(b){return b.options[d]===a.options[d]}):[a]}function F(a,b){var c=R(b),d=b.series,f=d.chart,k;y(d.options.dragDrop&&d.options.dragDrop.liveRedraw,!0)||(f.dragGuideBox=k=d.getGuideBox(c),f.setGuideBoxState("default",d.options.dragDrop.guideBox).add(d.group));f.dragDropData={origin:Q(a,c,k),point:b,
groupedPoints:c,isDragging:!0}}function S(a,b){var c=a.point,d=q(c.series.options.dragDrop,c.options.dragDrop),f={},k=a.updateProp,e={};n(c.series.dragDropProps,function(a,b){if(!k||k===b&&a.resize&&(!a.optionName||!1!==d[a.optionName]))if(k||a.move&&("x"===a.axis&&d.draggableX||"y"===a.axis&&d.draggableY))f[b]=a});(k?[c]:a.groupedPoints).forEach(function(c){e[c.id]={point:c,newValues:c.getDropValues(a.origin,b,f)}});return e}function G(a,b){var c=a.dragDropData.newPoints;b=T(b);a.isDragDropAnimating=
!0;n(c,function(a){a.point.update(a.newValues,!1)});a.redraw(b);setTimeout(function(){delete a.isDragDropAnimating;a.hoverPoint&&!a.dragHandles&&a.hoverPoint.showDragHandles()},b.duration)}function H(a){var b=a.series&&a.series.chart,c=b&&b.dragDropData;!b||!b.dragHandles||c&&(c.isDragging&&c.draggedPastSensitivity||c.isHoveringHandle===a.id)||b.hideDragHandles()}function I(a){var b=0,c;for(c in a)Object.hasOwnProperty.call(a,c)&&b++;return b}function J(a){for(var b in a)if(Object.hasOwnProperty.call(a,
b))return a[b]}function U(a,b){if(!b.zoomOrPanKeyPressed(a)){var c=b.dragDropData;var d=0;if(c&&c.isDragging&&c.point.series){var f=c.point;d=f.series.options.dragDrop;a.preventDefault();c.draggedPastSensitivity||(c.draggedPastSensitivity=P(a,b,y(f.options.dragDrop&&f.options.dragDrop.dragSensitivity,d&&d.dragSensitivity,2)));c.draggedPastSensitivity&&(c.newPoints=S(c,a),b=c.newPoints,d=I(b),b=1===d?J(b):null,f.firePointEvent("drag",{origin:c.origin,newPoints:c.newPoints,newPoint:b&&b.newValues,newPointId:b&&
b.point.id,numNewPoints:d,chartX:a.chartX,chartY:a.chartY},function(){var b=f.series,c=b.chart,d=c.dragDropData,e=q(b.options.dragDrop,f.options.dragDrop),g=e.draggableX,m=e.draggableY;b=d.origin;var h=a.chartX-b.chartX,C=a.chartY-b.chartY,u=h;d=d.updateProp;c.inverted&&(h=-C,C=-u);if(y(e.liveRedraw,!0))G(c,!1),f.showDragHandles();else if(d){g=h;c=C;u=f.series;m=u.chart;d=m.dragDropData;e=u.dragDropProps[d.updateProp];var l=d.newPoints[f.id].newValues;var p="function"===typeof e.resizeSide?e.resizeSide(l,
f):e.resizeSide;e.beforeResize&&e.beforeResize(m.dragGuideBox,l,f);m=m.dragGuideBox;u="x"===e.axis&&u.xAxis.reversed||"y"===e.axis&&u.yAxis.reversed?z(p):p;g="x"===e.axis?g-(d.origin.prevdX||0):0;c="y"===e.axis?c-(d.origin.prevdY||0):0;switch(u){case "left":var n={x:m.attr("x")+g,width:Math.max(1,m.attr("width")-g)};break;case "right":n={width:Math.max(1,m.attr("width")+g)};break;case "top":n={y:m.attr("y")+c,height:Math.max(1,m.attr("height")-c)};break;case "bottom":n={height:Math.max(1,m.attr("height")+
c)}}m.attr(n)}else c.dragGuideBox.translate(g?h:0,m?C:0);b.prevdX=h;b.prevdY=C}))}}}function D(a,b){var c=b.dragDropData;if(c&&c.isDragging&&c.draggedPastSensitivity&&c.point.series){var d=c.point,f=c.newPoints,e=I(f),g=1===e?J(f):null;b.dragHandles&&b.hideDragHandles();a.preventDefault();b.cancelClick=!0;d.firePointEvent("drop",{origin:c.origin,chartX:a.chartX,chartY:a.chartY,newPoints:f,numNewPoints:e,newPoint:g&&g.newValues,newPointId:g&&g.point.id},function(){G(b)})}delete b.dragDropData;b.dragGuideBox&&
(b.dragGuideBox.destroy(),delete b.dragGuideBox)}function V(a){var b=a.container,c=L.doc;N(a)&&(B(b,["mousedown","touchstart"],function(b){b=A(b,a);var c=a.hoverPoint,d=q(c&&c.series.options.dragDrop,c&&c.options.dragDrop),e=d.draggableX||!1;d=d.draggableY||!1;a.cancelClick=!1;!e&&!d||a.zoomOrPanKeyPressed(b)||a.hasDraggedAnnotation||(a.dragDropData&&a.dragDropData.isDragging?D(b,a):c&&O(c)&&(a.mouseIsDown=!1,F(b,c),c.firePointEvent("dragStart",b)))}),B(b,["mousemove","touchmove"],function(b){U(A(b,
a),a)},{passive:!1}),r(b,"mouseleave",function(b){D(A(b,a),a)}),a.unbindDragDropMouseUp=B(c,["mouseup","touchend"],function(b){D(A(b,a),a)},{passive:!1}),a.hasAddedDragDropEvents=!0,r(a,"destroy",function(){a.unbindDragDropMouseUp&&a.unbindDragDropMouseUp()}))}var T=e.animObject,g=w.seriesTypes,r=p.addEvent,W=p.clamp,X=p.isNumber,q=p.merge,n=p.objectEach,y=p.pick;e=function(a){a=a.shapeArgs||a.graphic.getBBox();var b=a.r||0,c=a.height/2;return[["M",0,b],["L",0,c-5],["A",1,1,0,0,0,0,c+5],["A",1,1,
0,0,0,0,c-5],["M",0,c+5],["L",0,a.height-b]]};w=E.prototype.dragDropProps={x:{axis:"x",move:!0},y:{axis:"y",move:!0}};g.flags&&(g.flags.prototype.dragDropProps=w);var h=g.column.prototype.dragDropProps={x:{axis:"x",move:!0},y:{axis:"y",move:!1,resize:!0,beforeResize:function(a,b,c){var d=c.series.translatedThreshold,f=a.attr("y");b.y>=c.series.options.threshold?(b=a.attr("height"),a.attr({height:Math.max(0,Math.round(b+(d?d-(f+b):0)))})):a.attr({y:Math.round(f+(d?d-f:0))})},resizeSide:function(a,
b){var c=b.series.chart.dragHandles;a=a.y>=(b.series.options.threshold||0)?"top":"bottom";b=z(a);c[b]&&(c[b].destroy(),delete c[b]);return a},handlePositioner:function(a){var b=a.shapeArgs||a.graphic&&a.graphic.getBBox()||{},c=a.series.yAxis.reversed,d=a.series.options.threshold||0;a=a.y||0;return{x:b.x||0,y:!c&&a>=d||c&&a<d?b.y||0:(b.y||0)+(b.height||0)}},handleFormatter:function(a){var b=a.shapeArgs||{};a=b.r||0;b=b.width||0;var c=b/2;return[["M",a,0],["L",c-5,0],["A",1,1,0,0,0,c+5,0],["A",1,1,
0,0,0,c-5,0],["M",c+5,0],["L",b-a,0]]}}};g.bullet&&(g.bullet.prototype.dragDropProps={x:h.x,y:h.y,target:{optionName:"draggableTarget",axis:"y",move:!0,resize:!0,resizeSide:"top",handlePositioner:function(a){var b=a.targetGraphic.getBBox();return{x:a.barX,y:b.y+b.height/2}},handleFormatter:h.y.handleFormatter}});g.columnrange&&(g.columnrange.prototype.dragDropProps={x:{axis:"x",move:!0},low:{optionName:"draggableLow",axis:"y",move:!0,resize:!0,resizeSide:"bottom",handlePositioner:function(a){a=a.shapeArgs||
a.graphic.getBBox();return{x:a.x||0,y:(a.y||0)+(a.height||0)}},handleFormatter:h.y.handleFormatter,propValidate:function(a,b){return a<=b.high}},high:{optionName:"draggableHigh",axis:"y",move:!0,resize:!0,resizeSide:"top",handlePositioner:function(a){a=a.shapeArgs||a.graphic.getBBox();return{x:a.x||0,y:a.y||0}},handleFormatter:h.y.handleFormatter,propValidate:function(a,b){return a>=b.low}}});g.boxplot&&(g.boxplot.prototype.dragDropProps={x:h.x,low:{optionName:"draggableLow",axis:"y",move:!0,resize:!0,
resizeSide:"bottom",handlePositioner:function(a){return{x:a.shapeArgs.x||0,y:a.lowPlot}},handleFormatter:h.y.handleFormatter,propValidate:function(a,b){return a<=b.q1}},q1:{optionName:"draggableQ1",axis:"y",move:!0,resize:!0,resizeSide:"bottom",handlePositioner:function(a){return{x:a.shapeArgs.x||0,y:a.q1Plot}},handleFormatter:h.y.handleFormatter,propValidate:function(a,b){return a<=b.median&&a>=b.low}},median:{axis:"y",move:!0},q3:{optionName:"draggableQ3",axis:"y",move:!0,resize:!0,resizeSide:"top",
handlePositioner:function(a){return{x:a.shapeArgs.x||0,y:a.q3Plot}},handleFormatter:h.y.handleFormatter,propValidate:function(a,b){return a<=b.high&&a>=b.median}},high:{optionName:"draggableHigh",axis:"y",move:!0,resize:!0,resizeSide:"top",handlePositioner:function(a){return{x:a.shapeArgs.x||0,y:a.highPlot}},handleFormatter:h.y.handleFormatter,propValidate:function(a,b){return a>=b.q3}}});g.ohlc&&(g.ohlc.prototype.dragDropProps={x:h.x,low:{optionName:"draggableLow",axis:"y",move:!0,resize:!0,resizeSide:"bottom",
handlePositioner:function(a){return{x:a.shapeArgs.x,y:a.plotLow}},handleFormatter:h.y.handleFormatter,propValidate:function(a,b){return a<=b.open&&a<=b.close}},high:{optionName:"draggableHigh",axis:"y",move:!0,resize:!0,resizeSide:"top",handlePositioner:function(a){return{x:a.shapeArgs.x,y:a.plotHigh}},handleFormatter:h.y.handleFormatter,propValidate:function(a,b){return a>=b.open&&a>=b.close}},open:{optionName:"draggableOpen",axis:"y",move:!0,resize:!0,resizeSide:function(a){return a.open>=a.close?
"top":"bottom"},handlePositioner:function(a){return{x:a.shapeArgs.x,y:a.plotOpen}},handleFormatter:h.y.handleFormatter,propValidate:function(a,b){return a<=b.high&&a>=b.low}},close:{optionName:"draggableClose",axis:"y",move:!0,resize:!0,resizeSide:function(a){return a.open>=a.close?"bottom":"top"},handlePositioner:function(a){return{x:a.shapeArgs.x,y:a.plotClose}},handleFormatter:h.y.handleFormatter,propValidate:function(a,b){return a<=b.high&&a>=b.low}}});g.arearange&&(w=g.columnrange.prototype.dragDropProps,
p=function(a){a=a.graphic?a.graphic.getBBox().width/2+1:4;return[["M",0-a,0],["a",a,a,0,1,0,2*a,0],["a",a,a,0,1,0,-2*a,0]]},g.arearange.prototype.dragDropProps={x:w.x,low:{optionName:"draggableLow",axis:"y",move:!0,resize:!0,resizeSide:"bottom",handlePositioner:function(a){return(a=a.lowerGraphic&&a.lowerGraphic.getBBox())?{x:a.x+a.width/2,y:a.y+a.height/2}:{x:-999,y:-999}},handleFormatter:p,propValidate:w.low.propValidate},high:{optionName:"draggableHigh",axis:"y",move:!0,resize:!0,resizeSide:"top",
handlePositioner:function(a){return(a=a.upperGraphic&&a.upperGraphic.getBBox())?{x:a.x+a.width/2,y:a.y+a.height/2}:{x:-999,y:-999}},handleFormatter:p,propValidate:w.high.propValidate}});g.waterfall&&(g.waterfall.prototype.dragDropProps={x:h.x,y:q(h.y,{handleFormatter:function(a){return a.isSum||a.isIntermediateSum?null:h.y.handleFormatter(a)}})});if(g.xrange){var K=function(a,b){var c=a.series,d=c.xAxis,f=c.yAxis,e=c.chart.inverted;b=d.toPixels(a[b],!0);var g=f.toPixels(a.y,!0);a=c.columnMetrics?
c.columnMetrics.offset:-a.shapeArgs.height/2;e&&(b=d.len-b,g=f.len-g);return{x:Math.round(b),y:Math.round(g+a)}};e=g.xrange.prototype.dragDropProps={y:{axis:"y",move:!0},x:{optionName:"draggableX1",axis:"x",move:!0,resize:!0,resizeSide:"left",handlePositioner:function(a){return K(a,"x")},handleFormatter:e,propValidate:function(a,b){return a<=b.x2}},x2:{optionName:"draggableX2",axis:"x",move:!0,resize:!0,resizeSide:"right",handlePositioner:function(a){return K(a,"x2")},handleFormatter:e,propValidate:function(a,
b){return a>=b.x}}};g.gantt&&(g.gantt.prototype.dragDropProps={y:e.y,start:q(e.x,{optionName:"draggableStart",validateIndividualDrag:function(a){return!a.milestone}}),end:q(e.x2,{optionName:"draggableEnd",validateIndividualDrag:function(a){return!a.milestone}})})}"gauge pie sunburst wordcloud sankey histogram pareto vector windbarb treemap bellcurve sma map mapline".split(" ").forEach(function(a){g[a]&&(g[a].prototype.dragDropProps=null)});var Y={"default":{className:"highcharts-drag-box-default",
lineWidth:1,lineColor:"#888",color:"rgba(0, 0, 0, 0.1)",cursor:"move",zIndex:900}},Z={className:"highcharts-drag-handle",color:"#fff",lineColor:"rgba(0, 0, 0, 0.6)",lineWidth:1,zIndex:901};l.prototype.setGuideBoxState=function(a,b){var c=this.dragGuideBox;b=q(Y,b);a=q(b["default"],b[a]);return c.attr({className:a.className,stroke:a.lineColor,strokeWidth:a.lineWidth,fill:a.color,cursor:a.cursor,zIndex:a.zIndex}).css({pointerEvents:"none"})};v.prototype.getDropValues=function(a,b,c){var d=this,f=d.series,
e=q(f.options.dragDrop,d.options.dragDrop),g={},h=a.points[d.id],t;for(t in c)if(Object.hasOwnProperty.call(c,t)){if("undefined"!==typeof l){var l=!1;break}l=!0}n(c,function(a,c){var m=h[c],k=f[a.axis+"Axis"];k=k.toValue((k.horiz?b.chartX:b.chartY)+h[c+"Offset"]);var x=a.axis.toUpperCase(),t=f[x.toLowerCase()+"Axis"].categories?1:0;t=y(e["dragPrecision"+x],t);var n=y(e["dragMin"+x],-Infinity);x=y(e["dragMax"+x],Infinity);t&&(k=Math.round(k/t)*t);k=W(k,n,x);l&&a.propValidate&&!a.propValidate(k,d)||
"undefined"===typeof m||(g[c]=k)});return g};E.prototype.getGuideBox=function(a){var b=this.chart,c=Infinity,d=-Infinity,e=Infinity,g=-Infinity,h;a.forEach(function(a){var b=a.graphic&&a.graphic.getBBox()||a.shapeArgs;if(b){var f=void 0,k=a.x2;X(k)&&(f=a.series.xAxis.translate(k,!1,!1,!1,!0));k=!(b.width||b.height||b.x||b.y);h=!0;c=Math.min(a.plotX||0,f||0,k?Infinity:b.x||0,c);d=Math.max(a.plotX||0,f||0,(b.x||0)+(b.width||0),d);e=Math.min(a.plotY||0,k?Infinity:b.y||0,e);g=Math.max((b.y||0)+(b.height||
0),g)}});return h?b.renderer.rect(c,e,d-c,g-e):b.renderer.g()};v.prototype.showDragHandles=function(){var a=this,b=a.series,c=b.chart,d=c.renderer,e=q(b.options.dragDrop,a.options.dragDrop);n(b.dragDropProps,function(f,g){var h=q(Z,f.handleOptions,e.dragHandle),k={"class":h.className,"stroke-width":h.lineWidth,fill:h.color,stroke:h.lineColor},l=h.pathFormatter||f.handleFormatter,m=f.handlePositioner,n=f.validateIndividualDrag?f.validateIndividualDrag(a):!0;f.resize&&n&&f.resizeSide&&l&&(e["draggable"+
f.axis.toUpperCase()]||e[f.optionName])&&!1!==e[f.optionName]&&(c.dragHandles||(c.dragHandles={group:d.g("drag-drop-handles").add(b.markerGroup||b.group)}),c.dragHandles.point=a.id,m=m(a),k.d=l=l(a),!l||0>m.x||0>m.y||(k.cursor=h.cursor||"x"===f.axis!==!!c.inverted?"ew-resize":"ns-resize",(h=c.dragHandles[f.optionName])||(h=c.dragHandles[f.optionName]=d.path().add(c.dragHandles.group)),h.translate(m.x,m.y).attr(k),B(h.element,["touchstart","mousedown"],function(b){b=A(b,c);var d=a.series.chart;d.zoomOrPanKeyPressed(b)||
(d.mouseIsDown=!1,F(b,a),d.dragDropData.updateProp=b.updateProp=g,a.firePointEvent("dragStart",b),b.stopPropagation(),b.preventDefault())},{passive:!1}),r(c.dragHandles.group.element,"mouseover",function(){c.dragDropData=c.dragDropData||{};c.dragDropData.isHoveringHandle=a.id}),B(c.dragHandles.group.element,["touchend","mouseout"],function(){var b=a.series.chart;b.dragDropData&&a.id===b.dragDropData.isHoveringHandle&&delete b.dragDropData.isHoveringHandle;b.hoverPoint||H(a)})))})};l.prototype.hideDragHandles=
function(){this.dragHandles&&(n(this.dragHandles,function(a,b){"group"!==b&&a.destroy&&a.destroy()}),this.dragHandles.group&&this.dragHandles.group.destroy&&this.dragHandles.group.destroy(),delete this.dragHandles)};r(v,"mouseOver",function(){var a=this;setTimeout(function(){var b=a.series,c=b&&b.chart,d=c&&c.dragDropData,e=c&&c.is3d&&c.is3d();!c||d&&d.isDragging&&d.draggedPastSensitivity||c.isDragDropAnimating||!b.options.dragDrop||e||(c.dragHandles&&c.hideDragHandles(),a.showDragHandles())},12)});
r(v,"mouseOut",function(){var a=this;setTimeout(function(){a.series&&H(a)},10)});r(v,"remove",function(){var a=this.series.chart,b=a.dragHandles;b&&b.point===this.id&&a.hideDragHandles()});l.prototype.zoomOrPanKeyPressed=function(a){var b=this.userOptions.chart||{},c=b.panKey&&b.panKey+"Key";return a[b.zoomKey&&b.zoomKey+"Key"]||a[c]};r(l,"render",function(){this.hasAddedDragDropEvents||V(this)});""});l(e,"masters/modules/draggable-points.src.js",[],function(){})});
//# sourceMappingURL=draggable-points.js.map
```
