## Supplementary material for "Exploring natural odour landscapes: A case study with implications for human-biting insects": Figure S4: drilldown.js

```
/*
 Highcharts JS v9.3.1 (2021-11-05)

 Highcharts Drilldown module

 Author: Torstein Honsi
 License: www.highcharts.com/license

*/
'use strict';(function(c){"object"===typeof module&&module.exports?(c["default"]=c,module.exports=c):"function"===typeof define&&define.amd?define("highcharts/modules/drilldown",["highcharts"],function(n){c(n);c.Highcharts=n;return c}):c("undefined"!==typeof Highcharts?Highcharts:void 0)})(function(c){function n(c,n,k,z){c.hasOwnProperty(n)||(c[n]=z.apply(null,k))}c=c?c._modules:{};n(c,"Extensions/Drilldown.js",[c["Core/Animation/AnimationUtilities.js"],c["Core/Axis/Axis.js"],c["Core/Chart/Chart.js"],
c["Core/Color/Color.js"],c["Series/Column/ColumnSeries.js"],c["Core/FormatUtilities.js"],c["Core/Globals.js"],c["Core/DefaultOptions.js"],c["Core/Series/Point.js"],c["Core/Series/Series.js"],c["Core/Series/SeriesRegistry.js"],c["Core/Renderer/SVG/SVGRenderer.js"],c["Core/Axis/Tick.js"],c["Core/Utilities.js"]],function(c,n,k,z,t,G,H,I,u,B,x,J,C,p){var D=c.animObject,K=G.format,L=H.noop;c=I.defaultOptions;var m=p.addEvent,M=p.removeEvent,r=p.extend,y=p.fireEvent,v=p.merge,N=p.objectEach,w=p.pick,O=
p.syncTimeout;x=x.seriesTypes.pie;var E=1;r(c.lang,{drillUpText:"\u25c1 Back to {series.name}"});c.drilldown={activeAxisLabelStyle:{cursor:"pointer",color:"#003399",fontWeight:"bold",textDecoration:"underline"},activeDataLabelStyle:{cursor:"pointer",color:"#003399",fontWeight:"bold",textDecoration:"underline"},animation:{duration:500},drillUpButton:{position:{align:"right",x:-10,y:10}}};J.prototype.Element.prototype.fadeIn=function(a){this.attr({opacity:.1,visibility:"inherit"}).animate({opacity:w(this.newOpacity,
1)},a||{duration:250})};k.prototype.addSeriesAsDrilldown=function(a,b){this.addSingleSeriesAsDrilldown(a,b);this.applyDrilldown()};k.prototype.addSingleSeriesAsDrilldown=function(a,b){var d=a.series,g=d.xAxis,f=d.yAxis,e=[],q=[],h;var l=this.styledMode?{colorIndex:w(a.colorIndex,d.colorIndex)}:{color:a.color||d.color};this.drilldownLevels||(this.drilldownLevels=[]);var c=d.options._levelNumber||0;(h=this.drilldownLevels[this.drilldownLevels.length-1])&&h.levelNumber!==c&&(h=void 0);b=r(r({_ddSeriesId:E++},
l),b);var k=d.points.indexOf(a);d.chart.series.forEach(function(a){a.xAxis!==g||a.isDrilling||(a.options._ddSeriesId=a.options._ddSeriesId||E++,a.options._colorIndex=a.userOptions._colorIndex,a.options._levelNumber=a.options._levelNumber||c,h?(e=h.levelSeries,q=h.levelSeriesOptions):(e.push(a),a.purgedOptions=v({_ddSeriesId:a.options._ddSeriesId,_levelNumber:a.options._levelNumber,selected:a.options.selected},a.userOptions),q.push(a.purgedOptions)))});a=r({levelNumber:c,seriesOptions:d.options,seriesPurgedOptions:d.purgedOptions,
levelSeriesOptions:q,levelSeries:e,shapeArgs:a.shapeArgs,bBox:a.graphic?a.graphic.getBBox():{},color:a.isNull?z.parse(l.color).setOpacity(0).get():l.color,lowerSeriesOptions:b,pointOptions:d.options.data[k],pointIndex:k,oldExtremes:{xMin:g&&g.userMin,xMax:g&&g.userMax,yMin:f&&f.userMin,yMax:f&&f.userMax},resetZoomButton:this.resetZoomButton},l);this.drilldownLevels.push(a);g&&g.names&&(g.names.length=0);b=a.lowerSeries=this.addSeries(b,!1);b.options._levelNumber=c+1;g&&(g.oldPos=g.pos,g.userMin=g.userMax=
null,f.userMin=f.userMax=null);d.type===b.type&&(b.animate=b.animateDrilldown||L,b.options.animation=!0)};k.prototype.applyDrilldown=function(){var a=this.drilldownLevels;if(a&&0<a.length){var b=a[a.length-1].levelNumber;this.drilldownLevels.forEach(function(a){a.levelNumber===b&&a.levelSeries.forEach(function(a){a.options&&a.options._levelNumber===b&&a.remove(!1)})})}this.resetZoomButton&&(this.resetZoomButton.hide(),delete this.resetZoomButton);this.pointer.reset();this.redraw();this.showDrillUpButton();
y(this,"afterDrilldown")};k.prototype.getDrilldownBackText=function(){var a=this.drilldownLevels;if(a&&0<a.length)return a=a[a.length-1],a.series=a.seriesOptions,K(this.options.lang.drillUpText||"",a)};k.prototype.showDrillUpButton=function(){var a=this,b=this.getDrilldownBackText(),d=a.options.drilldown.drillUpButton,g,f="chart"===d.relativeTo||"spacingBox"===d.relativeTo?null:"scrollablePlotBox";if(this.drillUpButton)this.drillUpButton.attr({text:b}).align();else{var e=(g=d.theme)&&g.states;this.drillUpButton=
this.renderer.button(b,null,null,function(){a.drillUp()},g,e&&e.hover,e&&e.select).addClass("highcharts-drillup-button").attr({align:d.position.align,zIndex:7}).add().align(d.position,!1,f)}};k.prototype.drillUp=function(){if(this.drilldownLevels&&0!==this.drilldownLevels.length){for(var a=this,b=a.drilldownLevels,d=b[b.length-1].levelNumber,g=b.length,f=a.series,e,c,h,l,k=function(b){f.forEach(function(a){a.options._ddSeriesId===b._ddSeriesId&&(d=a)});var d=d||a.addSeries(b,!1);d.type===h.type&&
d.animateDrillupTo&&(d.animate=d.animateDrillupTo);b===c.seriesPurgedOptions&&(l=d)};g--;)if(c=b[g],c.levelNumber===d){b.pop();h=c.lowerSeries;if(!h.chart)for(e=f.length;e--;)if(f[e].options.id===c.lowerSeriesOptions.id&&f[e].options._levelNumber===d+1){h=f[e];break}h.xData=[];c.levelSeriesOptions.forEach(k);y(a,"drillup",{seriesOptions:c.seriesPurgedOptions||c.seriesOptions});this.resetZoomButton&&this.resetZoomButton.destroy();l.type===h.type&&(l.drilldownLevel=c,l.options.animation=a.options.drilldown.animation,
h.animateDrillupFrom&&h.chart&&h.animateDrillupFrom(c));l.options._levelNumber=d;h.remove(!1);l.xAxis&&(e=c.oldExtremes,l.xAxis.setExtremes(e.xMin,e.xMax,!1),l.yAxis.setExtremes(e.yMin,e.yMax,!1));c.resetZoomButton&&(a.resetZoomButton=c.resetZoomButton,a.resetZoomButton.show())}this.redraw();0===this.drilldownLevels.length?this.drillUpButton=this.drillUpButton.destroy():this.drillUpButton.attr({text:this.getDrilldownBackText()}).align();this.ddDupes.length=[];y(a,"drillupall")}};m(k,"afterInit",function(){var a=
this;a.drilldown={update:function(b,d){v(!0,a.options.drilldown,b);w(d,!0)&&a.redraw()}}});m(k,"afterShowResetZoom",function(){var a=this.resetZoomButton&&this.resetZoomButton.getBBox(),b=this.options.drilldown&&this.options.drilldown.drillUpButton;this.drillUpButton&&a&&b&&b.position&&b.position.x&&this.drillUpButton.align({x:b.position.x-a.width-10,y:b.position.y,align:b.position.align},!1,b.relativeTo||"plotBox")});m(k,"render",function(){(this.xAxis||[]).forEach(function(a){a.ddPoints={};a.series.forEach(function(b){var d,
g=b.xData||[],f=b.points;for(d=0;d<g.length;d++){var e=b.options.data[d];"number"!==typeof e&&(e=b.pointClass.prototype.optionsToObject.call({series:b},e),e.drilldown&&(a.ddPoints[g[d]]||(a.ddPoints[g[d]]=[]),e=d-(b.cropStart||0),a.ddPoints[g[d]].push(f&&0<=e&&e<f.length?f[e]:!0)))}});N(a.ticks,C.prototype.drillable)})});t.prototype.animateDrillupTo=function(a){if(!a){var b=this,d=b.drilldownLevel;this.points.forEach(function(a){var b=a.dataLabel;a.graphic&&a.graphic.hide();b&&(b.hidden="hidden"===
b.attr("visibility"),b.hidden||(b.hide(),a.connector&&a.connector.hide()))});O(function(){if(b.points){var a=[];b.data.forEach(function(b){a.push(b)});b.nodes&&(a=a.concat(b.nodes));a.forEach(function(a,b){b=b===(d&&d.pointIndex)?"show":"fadeIn";var g="show"===b?!0:void 0,c=a.dataLabel;if(a.graphic)a.graphic[b](g);c&&!c.hidden&&(c.fadeIn(),a.connector&&a.connector.fadeIn())})}},Math.max(this.chart.options.drilldown.animation.duration-50,0));delete this.animate}};t.prototype.animateDrilldown=function(a){var b=
this,d=this.chart,c=d.drilldownLevels,f,e=D(d.options.drilldown.animation),q=this.xAxis,h=d.styledMode;a||(c.forEach(function(a){b.options._ddSeriesId===a.lowerSeriesOptions._ddSeriesId&&(f=a.shapeArgs,h||(f.fill=a.color))}),f.x+=w(q.oldPos,q.pos)-q.pos,this.points.forEach(function(a){var d=a.shapeArgs;h||(d.fill=a.color);a.graphic&&a.graphic.attr(f).animate(r(a.shapeArgs,{fill:a.color||b.color}),e);a.dataLabel&&a.dataLabel.fadeIn(e)}),delete this.animate)};t.prototype.animateDrillupFrom=function(a){var b=
D(this.chart.options.drilldown.animation),d=this.group,c=d!==this.chart.columnGroup,f=this;f.trackerGroups.forEach(function(a){if(f[a])f[a].on("mouseover")});c&&delete this.group;this.points.forEach(function(g){var e=g.graphic,h=a.shapeArgs,l=function(){e.destroy();d&&c&&(d=d.destroy())};e&&h&&(delete g.graphic,f.chart.styledMode||(h.fill=a.color),b.duration?e.animate(h,v(b,{complete:l})):(e.attr(h),l()))})};x&&r(x.prototype,{animateDrillupTo:t.prototype.animateDrillupTo,animateDrillupFrom:t.prototype.animateDrillupFrom,
animateDrilldown:function(a){var b=this.chart.drilldownLevels[this.chart.drilldownLevels.length-1],d=this.chart.options.drilldown.animation;this.is("item")&&(d.duration=0);if(this.center){var c=b.shapeArgs,f=c.start,e=(c.end-f)/this.points.length,k=this.chart.styledMode;a||(this.points.forEach(function(a,g){var h=a.shapeArgs;k||(c.fill=b.color,h.fill=a.color);if(a.graphic)a.graphic.attr(v(c,{start:f+g*e,end:f+(g+1)*e}))[d?"animate":"attr"](h,d)}),delete this.animate)}}});u.prototype.doDrilldown=function(){this.runDrilldown()};
u.prototype.runDrilldown=function(a,b,d){var c=this.series.chart,f=c.options.drilldown,e=(f.series||[]).length;c.ddDupes||(c.ddDupes=[]);for(;e--&&!k;)if(f.series[e].id===this.drilldown&&-1===c.ddDupes.indexOf(this.drilldown)){var k=f.series[e];c.ddDupes.push(this.drilldown)}y(c,"drilldown",{point:this,seriesOptions:k,category:b,originalEvent:d,points:"undefined"!==typeof b&&this.series.xAxis.getDDPoints(b).slice(0)},function(b){var d=b.point.series&&b.point.series.chart,c=b.seriesOptions;d&&c&&(a?
d.addSingleSeriesAsDrilldown(b.point,c):d.addSeriesAsDrilldown(b.point,c))})};n.prototype.drilldownCategory=function(a,b){this.getDDPoints(a).forEach(function(d){d&&d.series&&d.series.visible&&d.runDrilldown&&d.runDrilldown(!0,a,b)});this.chart.applyDrilldown()};n.prototype.getDDPoints=function(a){return this.ddPoints&&this.ddPoints[a]||[]};C.prototype.drillable=function(){var a=this.pos,b=this.label,d=this.axis,c="xAxis"===d.coll&&d.getDDPoints,f=c&&d.getDDPoints(a),e=d.chart.styledMode;c&&(b&&f&&
f.length?(b.drillable=!0,b.basicStyles||e||(b.basicStyles=v(b.styles)),b.addClass("highcharts-drilldown-axis-label"),b.removeOnDrillableClick&&M(b.element,"click"),b.removeOnDrillableClick=m(b.element,"click",function(b){b.preventDefault();d.drilldownCategory(a,b)}),e||b.css(d.chart.options.drilldown.activeAxisLabelStyle)):b&&b.drillable&&b.removeOnDrillableClick&&(e||(b.styles={},b.css(b.basicStyles)),b.removeOnDrillableClick(),b.removeClass("highcharts-drilldown-axis-label")))};m(u,"afterInit",
function(){this.drilldown&&!this.unbindDrilldownClick&&(this.unbindDrilldownClick=m(this,"click",F));return this});m(u,"update",function(a){a=a.options||{};a.drilldown&&!this.unbindDrilldownClick?this.unbindDrilldownClick=m(this,"click",F):!a.drilldown&&void 0!==a.drilldown&&this.unbindDrilldownClick&&(this.unbindDrilldownClick=this.unbindDrilldownClick())});var F=function(a){var b=this.series;b.xAxis&&!1===b.chart.options.drilldown.allowPointDrilldown?b.xAxis.drilldownCategory(this.x,a):this.runDrilldown(void 0,
void 0,a)};m(B,"afterDrawDataLabels",function(){var a=this.chart.options.drilldown.activeDataLabelStyle,b=this.chart.renderer,c=this.chart.styledMode;this.points.forEach(function(d){var f=d.options.dataLabels,e=w(d.dlOptions,f&&f.style,{});d.drilldown&&d.dataLabel&&("contrast"!==a.color||c||(e.color=b.getContrast(d.color||this.color)),f&&f.color&&(e.color=f.color),d.dataLabel.addClass("highcharts-drilldown-data-label"),c||d.dataLabel.css(a).css(e))},this)});var A=function(a,b,d,c){a[d?"addClass":
"removeClass"]("highcharts-drilldown-point");c||a.css({cursor:b})};m(B,"afterDrawTracker",function(){var a=this.chart.styledMode;this.points.forEach(function(b){b.drilldown&&b.graphic&&A(b.graphic,"pointer",!0,a)})});m(u,"afterSetState",function(){var a=this.series.chart.styledMode;this.drilldown&&this.series.halo&&"hover"===this.state?A(this.series.halo,"pointer",!0,a):this.series.halo&&A(this.series.halo,"auto",!1,a)});m(k,"selection",function(a){!0===a.resetSelection&&this.drillUpButton&&(a=this.options.drilldown&&
this.options.drilldown.drillUpButton)&&a.position&&this.drillUpButton.align({x:a.position.x,y:a.position.y,align:a.position.align},!1,a.relativeTo||"plotBox")});m(k,"drillup",function(){this.resetZoomButton&&(this.resetZoomButton=this.resetZoomButton.destroy())})});n(c,"masters/modules/drilldown.src.js",[],function(){})});
//# sourceMappingURL=drilldown.js.map
```
