## Supplementary material for "Exploring natural odour landscapes: A case study with implications for human-biting insects": Figure S4: dumbbell.js

```
/*
 Highcharts JS v9.3.1 (2021-11-05)

 (c) 2009-2021 Sebastian Bochan, Rafal Sebestjanski

 License: www.highcharts.com/license
*/
'use strict';(function(a){"object"===typeof module&&module.exports?(a["default"]=a,module.exports=a):"function"===typeof define&&define.amd?define("highcharts/modules/dumbbell",["highcharts"],function(n){a(n);a.Highcharts=n;return a}):a("undefined"!==typeof Highcharts?Highcharts:void 0)})(function(a){function n(a,g,h,k){a.hasOwnProperty(g)||(a[g]=k.apply(null,h))}a=a?a._modules:{};n(a,"Series/AreaRange/AreaRangePoint.js",[a["Series/Area/AreaSeries.js"],a["Core/Series/Point.js"],a["Core/Utilities.js"]],
function(a,g,h){var k=this&&this.__extends||function(){var a=function(e,d){a=Object.setPrototypeOf||{__proto__:[]}instanceof Array&&function(b,a){b.__proto__=a}||function(a,d){for(var b in d)d.hasOwnProperty(b)&&(a[b]=d[b])};return a(e,d)};return function(e,d){function b(){this.constructor=e}a(e,d);e.prototype=null===d?Object.create(d):(b.prototype=d.prototype,new b)}}(),m=g.prototype,r=h.defined,c=h.isNumber;return function(a){function e(){var d=null!==a&&a.apply(this,arguments)||this;d.high=void 0;
d.low=void 0;d.options=void 0;d.plotHigh=void 0;d.plotLow=void 0;d.plotHighX=void 0;d.plotLowX=void 0;d.plotX=void 0;d.series=void 0;return d}k(e,a);e.prototype.setState=function(){var a=this.state,b=this.series,e=b.chart.polar;r(this.plotHigh)||(this.plotHigh=b.yAxis.toPixels(this.high,!0));r(this.plotLow)||(this.plotLow=this.plotY=b.yAxis.toPixels(this.low,!0));b.stateMarkerGraphic&&(b.lowerStateMarkerGraphic=b.stateMarkerGraphic,b.stateMarkerGraphic=b.upperStateMarkerGraphic);this.graphic=this.upperGraphic;
this.plotY=this.plotHigh;e&&(this.plotX=this.plotHighX);m.setState.apply(this,arguments);this.state=a;this.plotY=this.plotLow;this.graphic=this.lowerGraphic;e&&(this.plotX=this.plotLowX);b.stateMarkerGraphic&&(b.upperStateMarkerGraphic=b.stateMarkerGraphic,b.stateMarkerGraphic=b.lowerStateMarkerGraphic,b.lowerStateMarkerGraphic=void 0);m.setState.apply(this,arguments)};e.prototype.haloPath=function(){var a=this.series.chart.polar,b=[];this.plotY=this.plotLow;a&&(this.plotX=this.plotLowX);this.isInside&&
(b=m.haloPath.apply(this,arguments));this.plotY=this.plotHigh;a&&(this.plotX=this.plotHighX);this.isTopInside&&(b=b.concat(m.haloPath.apply(this,arguments)));return b};e.prototype.isValid=function(){return c(this.low)&&c(this.high)};return e}(a.prototype.pointClass)});n(a,"Series/Dumbbell/DumbbellPoint.js",[a["Series/AreaRange/AreaRangePoint.js"],a["Core/Utilities.js"]],function(a,g){var h=this&&this.__extends||function(){var a=function(c,l){a=Object.setPrototypeOf||{__proto__:[]}instanceof Array&&
function(a,d){a.__proto__=d}||function(a,d){for(var b in d)d.hasOwnProperty(b)&&(a[b]=d[b])};return a(c,l)};return function(c,l){function e(){this.constructor=c}a(c,l);c.prototype=null===l?Object.create(l):(e.prototype=l.prototype,new e)}}(),k=g.extend,m=g.pick;g=function(a){function c(){var c=null!==a&&a.apply(this,arguments)||this;c.series=void 0;c.options=void 0;c.connector=void 0;c.pointWidth=void 0;return c}h(c,a);c.prototype.setState=function(){var a=this.series,c=a.chart,d=a.options.marker,
b=this.options,g=m(b.lowColor,a.options.lowColor,b.color,this.zone&&this.zone.color,this.color,a.color),h="attr";this.pointSetState.apply(this,arguments);this.state||(h="animate",this.lowerGraphic&&!c.styledMode&&(this.lowerGraphic.attr({fill:g}),this.upperGraphic&&(c={y:this.y,zone:this.zone},this.y=this.high,this.zone=this.zone?this.getZone():void 0,d=m(this.marker?this.marker.fillColor:void 0,d?d.fillColor:void 0,b.color,this.zone?this.zone.color:void 0,this.color),this.upperGraphic.attr({fill:d}),
k(this,c))));this.connector[h](a.getConnectorAttribs(this))};c.prototype.destroy=function(){this.graphic||(this.graphic=this.connector,this.connector=void 0);return a.prototype.destroy.call(this)};return c}(a);k(g.prototype,{pointSetState:a.prototype.setState});return g});n(a,"Series/Dumbbell/DumbbellSeries.js",[a["Series/Column/ColumnSeries.js"],a["Series/Dumbbell/DumbbellPoint.js"],a["Core/Globals.js"],a["Core/Series/Series.js"],a["Core/Series/SeriesRegistry.js"],a["Core/Renderer/SVG/SVGRenderer.js"],
a["Core/Utilities.js"]],function(a,g,h,k,m,n,c){var l=this&&this.__extends||function(){var a=function(b,f){a=Object.setPrototypeOf||{__proto__:[]}instanceof Array&&function(a,f){a.__proto__=f}||function(a,f){for(var b in f)f.hasOwnProperty(b)&&(a[b]=f[b])};return a(b,f)};return function(b,f){function d(){this.constructor=b}a(b,f);b.prototype=null===f?Object.create(f):(d.prototype=f.prototype,new d)}}(),e=a.prototype;h=h.noop;var d=k.prototype;k=m.seriesTypes;var b=k.arearange;k=k.columnrange.prototype;
var t=b.prototype,r=c.extend,u=c.merge,p=c.pick;c=function(a){function c(){var f=null!==a&&a.apply(this,arguments)||this;f.data=void 0;f.options=void 0;f.points=void 0;f.columnMetrics=void 0;return f}l(c,a);c.prototype.getConnectorAttribs=function(a){var b=this.chart,f=a.options,c=this.options,d=this.xAxis,e=this.yAxis,g=p(f.connectorWidth,c.connectorWidth),h=p(f.connectorColor,c.connectorColor,f.color,a.zone?a.zone.color:void 0,a.color),k=p(c.states&&c.states.hover&&c.states.hover.connectorWidthPlus,
1),m=p(f.dashStyle,c.dashStyle),l=p(a.plotLow,a.plotY),q=e.toPixels(c.threshold||0,!0);q=p(a.plotHigh,b.inverted?e.len-q:q);a.state&&(g+=k);0>l?l=0:l>=e.len&&(l=e.len);0>q?q=0:q>=e.len&&(q=e.len);if(0>a.plotX||a.plotX>d.len)g=0;a.upperGraphic&&(d={y:a.y,zone:a.zone},a.y=a.high,a.zone=a.zone?a.getZone():void 0,h=p(f.connectorColor,c.connectorColor,f.color,a.zone?a.zone.color:void 0,a.color),r(a,d));a={d:n.prototype.crispLine([["M",a.plotX,l],["L",a.plotX,q]],g,"ceil")};b.styledMode||(a.stroke=h,a["stroke-width"]=
g,m&&(a.dashstyle=m));return a};c.prototype.drawConnector=function(a){var b=p(this.options.animationLimit,250);b=a.connector&&this.chart.pointCount<b?"animate":"attr";a.connector||(a.connector=this.chart.renderer.path().addClass("highcharts-lollipop-stem").attr({zIndex:-1}).add(this.markerGroup));a.connector[b](this.getConnectorAttribs(a))};c.prototype.getColumnMetrics=function(){var a=e.getColumnMetrics.apply(this,arguments);a.offset+=a.width/2;return a};c.prototype.translate=function(){this.setShapeArgs.apply(this);
this.translatePoint.apply(this,arguments);this.points.forEach(function(a){var b=a.shapeArgs,c=a.pointWidth;a.plotX=b.x;b.x=a.plotX-c/2;a.tooltipPos=null});this.columnMetrics.offset-=this.columnMetrics.width/2};c.prototype.drawPoints=function(){var a=this.chart,b=this.points.length,c=this.lowColor=this.options.lowColor,d=0;for(this.seriesDrawPoints.apply(this,arguments);d<b;){var e=this.points[d];this.drawConnector(e);e.upperGraphic&&(e.upperGraphic.element.point=e,e.upperGraphic.addClass("highcharts-lollipop-high"));
e.connector.element.point=e;if(e.lowerGraphic){var g=e.zone&&e.zone.color;g=p(e.options.lowColor,c,e.options.color,g,e.color,this.color);a.styledMode||e.lowerGraphic.attr({fill:g});e.lowerGraphic.addClass("highcharts-lollipop-low")}d++}};c.prototype.markerAttribs=function(){var a=t.markerAttribs.apply(this,arguments);a.x=Math.floor(a.x||0);a.y=Math.floor(a.y||0);return a};c.prototype.pointAttribs=function(a,b){var c=d.pointAttribs.apply(this,arguments);"hover"===b&&delete c.fill;return c};c.defaultOptions=
u(b.defaultOptions,{trackByArea:!1,fillColor:"none",lineWidth:0,pointRange:1,connectorWidth:1,stickyTracking:!1,groupPadding:.2,crisp:!1,pointPadding:.1,lowColor:"#333333",states:{hover:{lineWidthPlus:0,connectorWidthPlus:1,halo:!1}}});return c}(b);r(c.prototype,{crispCol:e.crispCol,drawGraph:h,drawTracker:a.prototype.drawTracker,pointClass:g,setShapeArgs:k.translate,seriesDrawPoints:t.drawPoints,trackerGroups:["group","markerGroup","dataLabelsGroup"],translatePoint:t.translate});m.registerSeriesType("dumbbell",
c);"";return c});n(a,"masters/modules/dumbbell.src.js",[],function(){})});
//# sourceMappingURL=dumbbell.js.map
```
