## Supplementary material for "Exploring natural odour landscapes: A case study with implications for human-biting insects": Figure S4: export-data.js

```
/*
 Highcharts JS v9.3.1 (2021-11-05)

 Exporting module

 (c) 2010-2021 Torstein Honsi

 License: www.highcharts.com/license
*/
'use strict';(function(a){"object"===typeof module&&module.exports?(a["default"]=a,module.exports=a):"function"===typeof define&&define.amd?define("highcharts/modules/export-data",["highcharts","highcharts/modules/exporting"],function(k){a(k);a.Highcharts=k;return a}):a("undefined"!==typeof Highcharts?Highcharts:void 0)})(function(a){function k(a,f,d,p){a.hasOwnProperty(f)||(a[f]=p.apply(null,d))}a=a?a._modules:{};k(a,"Extensions/DownloadURL.js",[a["Core/Globals.js"]],function(a){var f=a.isSafari,
d=a.win,p=d.document,l=d.URL||d.webkitURL||d,r=a.dataURLtoBlob=function(a){if((a=a.replace(/filename=.*;/,"").match(/data:([^;]*)(;base64)?,([0-9A-Za-z+/]+)/))&&3<a.length&&d.atob&&d.ArrayBuffer&&d.Uint8Array&&d.Blob&&l.createObjectURL){var f=d.atob(a[3]),n=new d.ArrayBuffer(f.length);n=new d.Uint8Array(n);for(var c=0;c<n.length;++c)n[c]=f.charCodeAt(c);a=new d.Blob([n],{type:a[1]});return l.createObjectURL(a)}};a=a.downloadURL=function(a,l){var n=d.navigator,c=p.createElement("a");if("string"===
typeof a||a instanceof String||!n.msSaveOrOpenBlob){a=""+a;n=/Edge\/\d+/.test(n.userAgent);if(f&&"string"===typeof a&&0===a.indexOf("data:application/pdf")||n||2E6<a.length)if(a=r(a)||"",!a)throw Error("Failed to convert to blob");if("undefined"!==typeof c.download)c.href=a,c.download=l,p.body.appendChild(c),c.click(),p.body.removeChild(c);else try{var g=d.open(a,"chart");if("undefined"===typeof g||null===g)throw Error("Failed to open window");}catch(E){d.location.href=a}}else n.msSaveOrOpenBlob(a,
l)};return{dataURLtoBlob:r,downloadURL:a}});k(a,"Extensions/ExportData.js",[a["Core/Axis/Axis.js"],a["Core/Chart/Chart.js"],a["Core/Renderer/HTML/AST.js"],a["Core/Globals.js"],a["Core/DefaultOptions.js"],a["Core/Utilities.js"],a["Extensions/DownloadURL.js"]],function(a,f,d,p,l,r,n){function k(a,m){var b=g.navigator,c=-1<b.userAgent.indexOf("WebKit")&&0>b.userAgent.indexOf("Chrome"),d=g.URL||g.webkitURL||g;try{if(b.msSaveOrOpenBlob&&g.MSBlobBuilder){var t=new g.MSBlobBuilder;t.append(a);return t.getBlob("image/svg+xml")}if(!c)return d.createObjectURL(new g.Blob(["\ufeff"+
a],{type:m}))}catch(N){}}var I=p.doc,c=p.seriesTypes,g=p.win;p=l.getOptions;l=l.setOptions;var E=r.addEvent,J=r.defined,F=r.extend,K=r.find,C=r.fireEvent,L=r.isNumber,v=r.pick,G=n.downloadURL;l({exporting:{csv:{annotations:{itemDelimiter:"; ",join:!1},columnHeaderFormatter:null,dateFormat:"%Y-%m-%d %H:%M:%S",decimalPoint:null,itemDelimiter:null,lineDelimiter:"\n"},showTable:!1,useMultiLevelHeaders:!0,useRowspanHeaders:!0},lang:{downloadCSV:"Download CSV",downloadXLS:"Download XLS",exportData:{annotationHeader:"Annotations",
categoryHeader:"Category",categoryDatetimeHeader:"DateTime"},viewData:"View data table",hideData:"Hide data table"}});E(f,"render",function(){this.options&&this.options.exporting&&this.options.exporting.showTable&&!this.options.chart.forExport&&!this.dataTableDiv&&this.viewData()});f.prototype.setUpKeyToAxis=function(){c.arearange&&(c.arearange.prototype.keyToAxis={low:"y",high:"y"});c.gantt&&(c.gantt.prototype.keyToAxis={start:"x",end:"x"})};f.prototype.getDataRows=function(b){var m=this.hasParallelCoordinates,
y=this.time,c=this.options.exporting&&this.options.exporting.csv||{},d=this.xAxis,t={},f=[],n=[],p=[],z;var g=this.options.lang.exportData;var l=g.categoryHeader,M=g.categoryDatetimeHeader,w=function(q,e,m){if(c.columnHeaderFormatter){var d=c.columnHeaderFormatter(q,e,m);if(!1!==d)return d}return q?q instanceof a?q.options.title&&q.options.title.text||(q.dateTime?M:l):b?{columnTitle:1<m?e:q.name,topLevelColumnTitle:q.name}:q.name+(1<m?" ("+e+")":""):l},H=function(a,b,e){var q={},m={};b.forEach(function(b){var c=
(a.keyToAxis&&a.keyToAxis[b]||b)+"Axis";c=L(e)?a.chart[c][e]:a[c];q[b]=c&&c.categories||[];m[b]=c&&c.dateTime});return{categoryMap:q,dateTimeValueAxisMap:m}},r=function(a,b){return a.data.filter(function(a){return"undefined"!==typeof a.y&&a.name}).length&&b&&!b.categories&&!a.keyToAxis?a.pointArrayMap&&a.pointArrayMap.filter(function(a){return"x"===a}).length?(a.pointArrayMap.unshift("x"),a.pointArrayMap):["x","y"]:a.pointArrayMap||["y"]},h=[];var x=0;this.setUpKeyToAxis();this.series.forEach(function(a){var e=
a.xAxis,q=a.options.keys||r(a,e),f=q.length,g=!a.requireSorting&&{},k=d.indexOf(e),B=H(a,q),l;if(!1!==a.options.includeInDataExport&&!a.options.isInternal&&!1!==a.visible){K(h,function(a){return a[0]===k})||h.push([k,x]);for(l=0;l<f;)z=w(a,q[l],q.length),p.push(z.columnTitle||z),b&&n.push(z.topLevelColumnTitle||z),l++;var A={chart:a.chart,autoIncrement:a.autoIncrement,options:a.options,pointArrayMap:a.pointArrayMap};a.options.data.forEach(function(b,d){m&&(B=H(a,q,d));var w={series:A};a.pointClass.prototype.applyOptions.apply(w,
[b]);b=w.x;var h=a.data[d]&&a.data[d].name;l=0;if(!e||"name"===a.exportKey||!m&&e&&e.hasNames&&h)b=h;g&&(g[b]&&(b+="|"+d),g[b]=!0);t[b]||(t[b]=[],t[b].xValues=[]);t[b].x=w.x;t[b].name=h;for(t[b].xValues[k]=w.x;l<f;)d=q[l],h=w[d],t[b][x+l]=v(B.categoryMap[d][h],B.dateTimeValueAxisMap[d]?y.dateFormat(c.dateFormat,h):null,h),l++});x+=l}});for(e in t)Object.hasOwnProperty.call(t,e)&&f.push(t[e]);var e=b?[n,p]:[p];for(x=h.length;x--;){var A=h[x][0];var D=h[x][1];var k=d[A];f.sort(function(a,b){return a.xValues[A]-
b.xValues[A]});g=w(k);e[0].splice(D,0,g);b&&e[1]&&e[1].splice(D,0,g);f.forEach(function(a){var b=a.name;k&&!J(b)&&(k.dateTime?(a.x instanceof Date&&(a.x=a.x.getTime()),b=y.dateFormat(c.dateFormat,a.x)):b=k.categories?v(k.names[a.x],k.categories[a.x],a.x):a.x);a.splice(D,0,b)})}e=e.concat(f);C(this,"exportData",{dataRows:e});return e};f.prototype.getCSV=function(a){var b="",d=this.getDataRows(),c=this.options.exporting.csv,f=v(c.decimalPoint,","!==c.itemDelimiter&&a?(1.1).toLocaleString()[1]:"."),
l=v(c.itemDelimiter,","===f?";":","),g=c.lineDelimiter;d.forEach(function(a,c){for(var m,y=a.length;y--;)m=a[y],"string"===typeof m&&(m='"'+m+'"'),"number"===typeof m&&"."!==f&&(m=m.toString().replace(".",f)),a[y]=m;b+=a.join(l);c<d.length-1&&(b+=g)});return b};f.prototype.getTable=function(a){var b=function(a){if(!a.tagName||"#text"===a.tagName)return a.textContent||"";var c=a.attributes,d="<"+a.tagName;c&&Object.keys(c).forEach(function(a){d+=" "+a+'="'+c[a]+'"'});d+=">";d+=a.textContent||"";(a.children||
[]).forEach(function(a){d+=b(a)});return d+="</"+a.tagName+">"};a=this.getTableAST(a);return b(a)};f.prototype.getTableAST=function(a){var b=0,c=[],d=this.options,f=a?(1.1).toLocaleString()[1]:".",l=v(d.exporting.useMultiLevelHeaders,!0);a=this.getDataRows(l);var g=l?a.shift():null,k=a.shift(),n=function(a,b,c,d){var h=v(d,"");b="text"+(b?" "+b:"");"number"===typeof h?(h=h.toString(),","===f&&(h=h.replace(".",f)),b="number"):d||(b="empty");c=F({"class":b},c);return{tagName:a,attributes:c,textContent:h}};
!1!==d.exporting.tableCaption&&c.push({tagName:"caption",attributes:{"class":"highcharts-table-caption"},textContent:v(d.exporting.tableCaption,d.title.text?d.title.text:"Chart")});for(var p=0,r=a.length;p<r;++p)a[p].length>b&&(b=a[p].length);c.push(function(a,b,c){var f=[],h=0;c=c||b&&b.length;var m=0,e;if(e=l&&a&&b){a:if(e=a.length,b.length===e){for(;e--;)if(a[e]!==b[e]){e=!1;break a}e=!0}else e=!1;e=!e}if(e){for(e=[];h<c;++h){var g=a[h];var k=a[h+1];g===k?++m:m?(e.push(n("th","highcharts-table-topheading",
{scope:"col",colspan:m+1},g)),m=0):(g===b[h]?d.exporting.useRowspanHeaders?(k=2,delete b[h]):(k=1,b[h]=""):k=1,g=n("th","highcharts-table-topheading",{scope:"col"},g),1<k&&g.attributes&&(g.attributes.valign="top",g.attributes.rowspan=k),e.push(g))}f.push({tagName:"tr",children:e})}if(b){e=[];h=0;for(c=b.length;h<c;++h)"undefined"!==typeof b[h]&&e.push(n("th",null,{scope:"col"},b[h]));f.push({tagName:"tr",children:e})}return{tagName:"thead",children:f}}(g,k,Math.max(b,k.length)));var u=[];a.forEach(function(a){for(var c=
[],d=0;d<b;d++)c.push(n(d?"td":"th",null,d?{}:{scope:"row"},a[d]));u.push({tagName:"tr",children:c})});c.push({tagName:"tbody",children:u});c={tree:{tagName:"table",id:"highcharts-data-table-"+this.index,children:c}};C(this,"aftergetTableAST",c);return c.tree};f.prototype.downloadCSV=function(){var a=this.getCSV(!0);G(k(a,"text/csv")||"data:text/csv,\ufeff"+encodeURIComponent(a),this.getFilename()+".csv")};f.prototype.downloadXLS=function(){var a='<html xmlns:o="urn:schemas-microsoft-com:office:office" xmlns:x="urn:schemas-microsoft-com:office:excel" xmlns="http://www.w3.org/TR/REC-html40"><head>\x3c!--[if gte mso 9]><xml><x:ExcelWorkbook><x:ExcelWorksheets><x:ExcelWorksheet><x:Name>Ark1</x:Name><x:WorksheetOptions><x:DisplayGridlines/></x:WorksheetOptions></x:ExcelWorksheet></x:ExcelWorksheets></x:ExcelWorkbook></xml><![endif]--\x3e<style>td{border:none;font-family: Calibri, sans-serif;} .number{mso-number-format:"0.00";} .text{ mso-number-format:"@";}</style><meta name=ProgId content=Excel.Sheet><meta charset=UTF-8></head><body>'+
this.getTable(!0)+"</body></html>";G(k(a,"application/vnd.ms-excel")||"data:application/vnd.ms-excel;base64,"+g.btoa(unescape(encodeURIComponent(a))),this.getFilename()+".xls")};f.prototype.viewData=function(){this.toggleDataTable(!0)};f.prototype.hideData=function(){this.toggleDataTable(!1)};f.prototype.toggleDataTable=function(a){(a=v(a,!this.isDataTableVisible))&&!this.dataTableDiv&&(this.dataTableDiv=I.createElement("div"),this.dataTableDiv.className="highcharts-data-table",this.renderTo.parentNode.insertBefore(this.dataTableDiv,
this.renderTo.nextSibling));this.dataTableDiv&&(this.dataTableDiv.style.display=a?"block":"none",a&&(this.dataTableDiv.innerHTML="",(new d([this.getTableAST()])).addToDOM(this.dataTableDiv),C(this,"afterViewData",this.dataTableDiv)));this.isDataTableVisible=a;a=this.exportDivElements;var b=this.options.exporting,c=b&&b.buttons&&b.buttons.contextButton.menuItems;b=this.options.lang;u&&u.menuItemDefinitions&&b&&b.viewData&&b.hideData&&c&&a&&(a=a[c.indexOf("viewData")])&&d.setElementHTML(a,this.isDataTableVisible?
b.hideData:b.viewData)};var u=p().exporting;u&&(F(u.menuItemDefinitions,{downloadCSV:{textKey:"downloadCSV",onclick:function(){this.downloadCSV()}},downloadXLS:{textKey:"downloadXLS",onclick:function(){this.downloadXLS()}},viewData:{textKey:"viewData",onclick:function(){this.toggleDataTable()}}}),u.buttons&&u.buttons.contextButton.menuItems.push("separator","downloadCSV","downloadXLS","viewData"));c.map&&(c.map.prototype.exportKey="name");c.mapbubble&&(c.mapbubble.prototype.exportKey="name");c.treemap&&
(c.treemap.prototype.exportKey="name")});k(a,"masters/modules/export-data.src.js",[],function(){})});
//# sourceMappingURL=export-data.js.map
```
