## Supplementary material for "Exploring natural odour landscapes: A case study with implications for human-biting insects": Figure S4: exporting.js

```
/*
 Highcharts JS v9.3.1 (2021-11-05)

 Exporting module

 (c) 2010-2021 Torstein Honsi

 License: www.highcharts.com/license
*/
'use strict';(function(a){"object"===typeof module&&module.exports?(a["default"]=a,module.exports=a):"function"===typeof define&&define.amd?define("highcharts/modules/exporting",["highcharts"],function(g){a(g);a.Highcharts=g;return a}):a("undefined"!==typeof Highcharts?Highcharts:void 0)})(function(a){function g(a,e,r,t){a.hasOwnProperty(e)||(a[e]=t.apply(null,r))}a=a?a._modules:{};g(a,"Extensions/FullScreen.js",[a["Core/Chart/Chart.js"],a["Core/Globals.js"],a["Core/Renderer/HTML/AST.js"],a["Core/Utilities.js"]],
function(a,e,r,t){var l=t.addEvent;t=function(){function a(d){this.chart=d;this.isOpen=!1;d=d.renderTo;this.browserProps||("function"===typeof d.requestFullscreen?this.browserProps={fullscreenChange:"fullscreenchange",requestFullscreen:"requestFullscreen",exitFullscreen:"exitFullscreen"}:d.mozRequestFullScreen?this.browserProps={fullscreenChange:"mozfullscreenchange",requestFullscreen:"mozRequestFullScreen",exitFullscreen:"mozCancelFullScreen"}:d.webkitRequestFullScreen?this.browserProps={fullscreenChange:"webkitfullscreenchange",
requestFullscreen:"webkitRequestFullScreen",exitFullscreen:"webkitExitFullscreen"}:d.msRequestFullscreen&&(this.browserProps={fullscreenChange:"MSFullscreenChange",requestFullscreen:"msRequestFullscreen",exitFullscreen:"msExitFullscreen"}))}a.prototype.close=function(){var d=this.chart,a=d.options.chart;if(this.isOpen&&this.browserProps&&d.container.ownerDocument instanceof Document)d.container.ownerDocument[this.browserProps.exitFullscreen]();this.unbindFullscreenEvent&&(this.unbindFullscreenEvent=
this.unbindFullscreenEvent());d.setSize(this.origWidth,this.origHeight,!1);this.origHeight=this.origWidth=void 0;a.width=this.origWidthOption;a.height=this.origHeightOption;this.origHeightOption=this.origWidthOption=void 0;this.isOpen=!1;this.setButtonText()};a.prototype.open=function(){var d=this,a=d.chart,b=a.options.chart;b&&(d.origWidthOption=b.width,d.origHeightOption=b.height);d.origWidth=a.chartWidth;d.origHeight=a.chartHeight;if(d.browserProps){var k=l(a.container.ownerDocument,d.browserProps.fullscreenChange,
function(){d.isOpen?(d.isOpen=!1,d.close()):(a.setSize(null,null,!1),d.isOpen=!0,d.setButtonText())}),e=l(a,"destroy",k);d.unbindFullscreenEvent=function(){k();e()};if(b=a.renderTo[d.browserProps.requestFullscreen]())b["catch"](function(){alert("Full screen is not supported inside a frame.")})}};a.prototype.setButtonText=function(){var d=this.chart,a=d.exportDivElements,b=d.options.exporting,k=b&&b.buttons&&b.buttons.contextButton.menuItems;d=d.options.lang;b&&b.menuItemDefinitions&&d&&d.exitFullscreen&&
d.viewFullscreen&&k&&a&&(a=a[k.indexOf("viewFullscreen")])&&r.setElementHTML(a,this.isOpen?d.exitFullscreen:b.menuItemDefinitions.viewFullscreen.text||d.viewFullscreen)};a.prototype.toggle=function(){this.isOpen?this.close():this.open()};return a}();e.Fullscreen=t;l(a,"beforeRender",function(){this.fullscreen=new e.Fullscreen(this)});return e.Fullscreen});g(a,"Core/Chart/ChartNavigationComposition.js",[],function(){var a;(function(a){a.compose=function(a){a.navigation||(a.navigation=new e(a));return a};
var e=function(){function a(a){this.updates=[];this.chart=a}a.prototype.addUpdate=function(a){this.chart.navigation.updates.push(a)};a.prototype.update=function(a,k){var d=this;this.updates.forEach(function(e){e.call(d.chart,a,k)})};return a}();a.Additions=e})(a||(a={}));return a});g(a,"Extensions/Exporting/ExportingDefaults.js",[a["Core/Globals.js"]],function(a){return{exporting:{type:"image/png",url:"https://export.highcharts.com/",printMaxWidth:780,scale:2,buttons:{contextButton:{className:"highcharts-contextbutton",
menuClassName:"highcharts-contextmenu",symbol:"menu",titleKey:"contextButtonTitle",menuItems:"viewFullscreen printChart separator downloadPNG downloadJPEG downloadPDF downloadSVG".split(" ")}},menuItemDefinitions:{viewFullscreen:{textKey:"viewFullscreen",onclick:function(){this.fullscreen.toggle()}},printChart:{textKey:"printChart",onclick:function(){this.print()}},separator:{separator:!0},downloadPNG:{textKey:"downloadPNG",onclick:function(){this.exportChart()}},downloadJPEG:{textKey:"downloadJPEG",
onclick:function(){this.exportChart({type:"image/jpeg"})}},downloadPDF:{textKey:"downloadPDF",onclick:function(){this.exportChart({type:"application/pdf"})}},downloadSVG:{textKey:"downloadSVG",onclick:function(){this.exportChart({type:"image/svg+xml"})}}}},lang:{viewFullscreen:"View in full screen",exitFullscreen:"Exit from full screen",printChart:"Print chart",downloadPNG:"Download PNG image",downloadJPEG:"Download JPEG image",downloadPDF:"Download PDF document",downloadSVG:"Download SVG vector image",
contextButtonTitle:"Chart context menu"},navigation:{buttonOptions:{symbolSize:14,symbolX:12.5,symbolY:10.5,align:"right",buttonSpacing:3,height:22,verticalAlign:"top",width:24,symbolFill:"#666666",symbolStroke:"#666666",symbolStrokeWidth:3,theme:{padding:5}},menuStyle:{border:"1px solid #999999",background:"#ffffff",padding:"5px 0"},menuItemStyle:{padding:"0.5em 1em",color:"#333333",background:"none",fontSize:a.isTouchDevice?"14px":"11px",transition:"background 250ms, color 250ms"},menuItemHoverStyle:{background:"#335cad",
color:"#ffffff"}}}});g(a,"Extensions/Exporting/ExportingSymbols.js",[],function(){var a;(function(a){function e(a,d,e,b){return[["M",a,d+2.5],["L",a+e,d+2.5],["M",a,d+b/2+.5],["L",a+e,d+b/2+.5],["M",a,d+b-1.5],["L",a+e,d+b-1.5]]}function l(a,d,e,b){a=b/3-2;b=[];return b=b.concat(this.circle(e-a,d,a,a),this.circle(e-a,d+a+4,a,a),this.circle(e-a,d+2*(a+4),a,a))}var g=[];a.compose=function(a){-1===g.indexOf(a)&&(g.push(a),a=a.prototype.symbols,a.menu=e,a.menuball=l.bind(a))}})(a||(a={}));return a});
g(a,"Core/HttpUtilities.js",[a["Core/Globals.js"],a["Core/Utilities.js"]],function(a,e){var l=a.doc,g=e.createElement,z=e.discardElement,k=e.merge,d=e.objectEach,J={ajax:function(a){var b=k(!0,{url:!1,type:"get",dataType:"json",success:!1,error:!1,data:!1,headers:{}},a);a={json:"application/json",xml:"application/xml",text:"text/plain",octet:"application/octet-stream"};var e=new XMLHttpRequest;if(!b.url)return!1;e.open(b.type.toUpperCase(),b.url,!0);b.headers["Content-Type"]||e.setRequestHeader("Content-Type",
a[b.dataType]||a.text);d(b.headers,function(a,d){e.setRequestHeader(d,a)});e.onreadystatechange=function(){if(4===e.readyState){if(200===e.status){var a=e.responseText;if("json"===b.dataType)try{a=JSON.parse(a)}catch(q){b.error&&b.error(e,q);return}return b.success&&b.success(a)}b.error&&b.error(e,e.responseText)}};try{b.data=JSON.stringify(b.data)}catch(x){}e.send(b.data||!0)},getJSON:function(a,d){J.ajax({url:a,success:d,dataType:"json",headers:{"Content-Type":"text/plain"}})},post:function(a,e,
r){var b=g("form",k({method:"post",action:a,enctype:"multipart/form-data"},r),{display:"none"},l.body);d(e,function(a,d){g("input",{type:"hidden",name:d,value:a},null,b)});b.submit();z(b)}};"";return J});g(a,"Extensions/Exporting/Exporting.js",[a["Core/Renderer/HTML/AST.js"],a["Core/Chart/Chart.js"],a["Core/Chart/ChartNavigationComposition.js"],a["Core/DefaultOptions.js"],a["Extensions/Exporting/ExportingDefaults.js"],a["Extensions/Exporting/ExportingSymbols.js"],a["Core/Globals.js"],a["Core/HttpUtilities.js"],
a["Core/Utilities.js"]],function(a,e,g,t,z,k,d,J,b){e=t.defaultOptions;var l=d.doc,r=d.win,x=b.addEvent,q=b.css,E=b.createElement,K=b.discardElement,F=b.extend,P=b.find,G=b.fireEvent,Q=b.isObject,m=b.merge,L=b.objectEach,w=b.pick,R=b.removeEvent,S=b.uniqueKey,H;(function(e){function t(a){var c=this,d=c.renderer,b=m(c.options.navigation.buttonOptions,a),e=b.onclick,B=b.menuItems,n=b.symbolSize||12;c.btnCount||(c.btnCount=0);c.exportDivElements||(c.exportDivElements=[],c.exportSVGElements=[]);if(!1!==
b.enabled&&b.theme){var f=b.theme,C=f.states,l=C&&C.hover;C=C&&C.select;var D;c.styledMode||(f.fill=w(f.fill,"#ffffff"),f.stroke=w(f.stroke,"none"));delete f.states;e?D=function(a){a&&a.stopPropagation();e.call(c,a)}:B&&(D=function(a){a&&a.stopPropagation();c.contextMenu(p.menuClassName,B,p.translateX,p.translateY,p.width,p.height,p);p.setState(2)});b.text&&b.symbol?f.paddingLeft=w(f.paddingLeft,30):b.text||F(f,{width:b.width,height:b.height,padding:0});c.styledMode||(f["stroke-linecap"]="round",
f.fill=w(f.fill,"#ffffff"),f.stroke=w(f.stroke,"none"));var p=d.button(b.text,0,0,D,f,l,C).addClass(a.className).attr({title:w(c.options.lang[b._titleKey||b.titleKey],"")});p.menuClassName=a.menuClassName||"highcharts-menu-"+c.btnCount++;if(b.symbol){var g=d.symbol(b.symbol,b.symbolX-n/2,b.symbolY-n/2,n,n,{width:n,height:n}).addClass("highcharts-button-symbol").attr({zIndex:1}).add(p);c.styledMode||g.attr({stroke:b.symbolStroke,fill:b.symbolFill,"stroke-width":b.symbolStrokeWidth||1})}p.add(c.exportingGroup).align(F(b,
{width:p.width,x:w(b.x,c.buttonOffset)}),!0,"spacingBox");c.buttonOffset+=(p.width+b.buttonSpacing)*("right"===b.align?-1:1);c.exportSVGElements.push(p,g)}}function z(){if(this.printReverseInfo){var a=this.printReverseInfo,b=a.childNodes,d=a.origDisplay;a=a.resetParams;this.moveContainers(this.renderTo);[].forEach.call(b,function(a,c){1===a.nodeType&&(a.style.display=d[c]||"")});this.isPrinting=!1;a&&this.setSize.apply(this,a);delete this.printReverseInfo;I=void 0;G(this,"afterPrint")}}function H(){var a=
l.body,b=this.options.exporting.printMaxWidth,d={childNodes:a.childNodes,origDisplay:[],resetParams:void 0};this.isPrinting=!0;this.pointer.reset(null,0);G(this,"beforePrint");b&&this.chartWidth>b&&(d.resetParams=[this.options.chart.width,void 0,!1],this.setSize(b,void 0,!1));[].forEach.call(d.childNodes,function(a,c){1===a.nodeType&&(d.origDisplay[c]=a.style.display,a.style.display="none")});this.moveContainers(a);this.printReverseInfo=d}function T(a){a.renderExporting();x(a,"redraw",a.renderExporting);
x(a,"destroy",a.destroyExport)}function U(c,d,e,y,g,B,n){var f=this,u=f.options.navigation,A=f.chartWidth,D=f.chartHeight,p="cache-"+c,v=Math.max(g,B),h=f[p];if(!h){f.exportContextMenu=f[p]=h=E("div",{className:c},{position:"absolute",zIndex:1E3,padding:v+"px",pointerEvents:"auto"},f.fixedDiv||f.container);var m=E("ul",{className:"highcharts-menu"},{listStyle:"none",margin:0,padding:0},h);f.styledMode||q(m,F({MozBoxShadow:"3px 3px 10px #888",WebkitBoxShadow:"3px 3px 10px #888",boxShadow:"3px 3px 10px #888"},
u.menuStyle));h.hideMenu=function(){q(h,{display:"none"});n&&n.setState(0);f.openMenu=!1;q(f.renderTo,{overflow:"hidden"});q(f.container,{overflow:"hidden"});b.clearTimeout(h.hideTimer);G(f,"exportMenuHidden")};f.exportEvents.push(x(h,"mouseleave",function(){h.hideTimer=r.setTimeout(h.hideMenu,500)}),x(h,"mouseenter",function(){b.clearTimeout(h.hideTimer)}),x(l,"mouseup",function(a){f.pointer.inClass(a.target,c)||h.hideMenu()}),x(h,"click",function(){f.openMenu&&h.hideMenu()}));d.forEach(function(c){"string"===
typeof c&&(c=f.options.exporting.menuItemDefinitions[c]);if(Q(c,!0)){var b=void 0;c.separator?b=E("hr",void 0,void 0,m):("viewData"===c.textKey&&f.isDataTableVisible&&(c.textKey="hideData"),b=E("li",{className:"highcharts-menu-item",onclick:function(a){a&&a.stopPropagation();h.hideMenu();c.onclick&&c.onclick.apply(f,arguments)}},void 0,m),a.setElementHTML(b,c.text||f.options.lang[c.textKey]),f.styledMode||(b.onmouseover=function(){q(this,u.menuItemHoverStyle)},b.onmouseout=function(){q(this,u.menuItemStyle)},
q(b,F({cursor:"pointer"},u.menuItemStyle))));f.exportDivElements.push(b)}});f.exportDivElements.push(m,h);f.exportMenuWidth=h.offsetWidth;f.exportMenuHeight=h.offsetHeight}d={display:"block"};e+f.exportMenuWidth>A?d.right=A-e-g-v+"px":d.left=e-v+"px";y+B+f.exportMenuHeight>D&&"top"!==n.alignOptions.verticalAlign?d.bottom=D-y-v+"px":d.top=y+B-v+"px";q(h,d);q(f.renderTo,{overflow:""});q(f.container,{overflow:""});f.openMenu=!0;G(f,"exportMenuShown")}function V(a){var c=a?a.target:this,d=c.exportSVGElements,
e=c.exportDivElements;a=c.exportEvents;var g;d&&(d.forEach(function(a,b){a&&(a.onclick=a.ontouchstart=null,g="cache-"+a.menuClassName,c[g]&&delete c[g],d[b]=a.destroy())}),d.length=0);c.exportingGroup&&(c.exportingGroup.destroy(),delete c.exportingGroup);e&&(e.forEach(function(a,c){a&&(b.clearTimeout(a.hideTimer),R(a,"mouseleave"),e[c]=a.onmouseout=a.onmouseover=a.ontouchstart=a.onclick=null,K(a))}),e.length=0);a&&(a.forEach(function(a){a()}),a.length=0)}function W(a,b){b=this.getSVGForExport(a,b);
a=m(this.options.exporting,a);J.post(a.url,{filename:a.filename?a.filename.replace(/\//g,"-"):this.getFilename(),type:a.type,width:a.width||0,scale:a.scale,svg:b},a.formAttributes)}function X(){this.styledMode&&this.inlineStyles();return this.container.innerHTML}function Y(){var a=this.userOptions.title&&this.userOptions.title.text,b=this.options.exporting.filename;if(b)return b.replace(/\//g,"-");"string"===typeof a&&(b=a.toLowerCase().replace(/<\/?[^>]+(>|$)/g,"").replace(/[\s_]+/g,"-").replace(/[^a-z0-9\-]/g,
"").replace(/^[\-]+/g,"").replace(/[\-]+/g,"-").substr(0,24).replace(/[\-]+$/g,""));if(!b||5>b.length)b="chart";return b}function Z(a){var b,c=m(this.options,a);c.plotOptions=m(this.userOptions.plotOptions,a&&a.plotOptions);c.time=m(this.userOptions.time,a&&a.time);var d=E("div",null,{position:"absolute",top:"-9999em",width:this.chartWidth+"px",height:this.chartHeight+"px"},l.body),e=this.renderTo.style.width;var g=this.renderTo.style.height;e=c.exporting.sourceWidth||c.chart.width||/px$/.test(e)&&
parseInt(e,10)||(c.isGantt?800:600);g=c.exporting.sourceHeight||c.chart.height||/px$/.test(g)&&parseInt(g,10)||400;F(c.chart,{animation:!1,renderTo:d,forExport:!0,renderer:"SVGRenderer",width:e,height:g});c.exporting.enabled=!1;delete c.data;c.series=[];this.series.forEach(function(a){b=m(a.userOptions,{animation:!1,enableMouseTracking:!1,showCheckbox:!1,visible:a.visible});b.isInternal||c.series.push(b)});var n={};this.axes.forEach(function(a){a.userOptions.internalKey||(a.userOptions.internalKey=
S());a.options.isInternal||(n[a.coll]||(n[a.coll]=!0,c[a.coll]=[]),c[a.coll].push(m(a.userOptions,{visible:a.visible})))});var f=new this.constructor(c,this.callback);a&&["xAxis","yAxis","series"].forEach(function(c){var b={};a[c]&&(b[c]=a[c],f.update(b))});this.axes.forEach(function(a){var c=P(f.axes,function(c){return c.options.internalKey===a.userOptions.internalKey}),b=a.getExtremes(),d=b.userMin;b=b.userMax;c&&("undefined"!==typeof d&&d!==c.min||"undefined"!==typeof b&&b!==c.max)&&c.setExtremes(d,
b,!0,!1)});g=f.getChartHTML();G(this,"getSVG",{chartCopy:f});g=this.sanitizeSVG(g,c);c=null;f.destroy();K(d);return g}function aa(a,b){var c=this.options.exporting;return this.getSVG(m({chart:{borderRadius:0}},c.chartOptions,b,{exporting:{sourceWidth:a&&a.sourceWidth||c.sourceWidth,sourceHeight:a&&a.sourceHeight||c.sourceHeight}}))}function M(a){return a.replace(/([A-Z])/g,function(a,c){return"-"+c.toLowerCase()})}function ba(){function a(c){function e(a,d){u=l=!1;if(g.length){for(k=g.length;k--&&
!l;)l=g[k].test(d);u=!l}"transform"===d&&"none"===a&&(u=!0);for(k=b.length;k--&&!u;)u=b[k].test(d)||"function"===typeof a;u||q[d]===a&&"svg"!==c.nodeName||y[c.nodeName][d]===a||(N&&-1===N.indexOf(d)?f+=M(d)+":"+a+";":a&&c.setAttribute(M(d),a))}var f="",u,l,k;if(1===c.nodeType&&-1===ca.indexOf(c.nodeName)){var h=r.getComputedStyle(c,null);var q="svg"===c.nodeName?{}:r.getComputedStyle(c.parentNode,null);if(!y[c.nodeName]){v=n.getElementsByTagName("svg")[0];var A=n.createElementNS(c.namespaceURI,c.nodeName);
v.appendChild(A);y[c.nodeName]=m(r.getComputedStyle(A,null));"text"===c.nodeName&&delete y.text.fill;v.removeChild(A)}if(d.isFirefox||d.isMS)for(var t in h)e(h[t],t);else L(h,e);f&&(h=c.getAttribute("style"),c.setAttribute("style",(h?h+";":"")+f));"svg"===c.nodeName&&c.setAttribute("stroke-width","1px");"text"!==c.nodeName&&[].forEach.call(c.children||c.childNodes,a)}}var b=da,g=e.inlineWhitelist,y={},v,k=l.createElement("iframe");q(k,{width:"1px",height:"1px",visibility:"hidden"});l.body.appendChild(k);
var n=k.contentWindow.document;n.open();n.write('<svg xmlns="http://www.w3.org/2000/svg"></svg>');n.close();a(this.container.querySelector("svg"));v.parentNode.removeChild(v);k.parentNode.removeChild(k)}function ea(a){(this.fixedDiv?[this.fixedDiv,this.scrollingContainer]:[this.container]).forEach(function(c){a.appendChild(c)})}function fa(){var a=this;a.exporting={update:function(c,b){a.isDirtyExporting=!0;m(!0,a.options.exporting,c);w(b,!0)&&a.redraw()}};g.compose(a).navigation.addUpdate(function(c,
b){a.isDirtyExporting=!0;m(!0,a.options.navigation,c);w(b,!0)&&a.redraw()})}function ha(){var a=this;a.isPrinting||(I=a,d.isSafari||a.beforePrint(),setTimeout(function(){r.focus();r.print();d.isSafari||setTimeout(function(){a.afterPrint()},1E3)},1))}function ia(){var a=this,b=a.options.exporting,d=b.buttons,e=a.isDirtyExporting||!a.exportSVGElements;a.buttonOffset=0;a.isDirtyExporting&&a.destroyExport();e&&!1!==b.enabled&&(a.exportEvents=[],a.exportingGroup=a.exportingGroup||a.renderer.g("exporting-group").attr({zIndex:3}).add(),
L(d,function(b){a.addButton(b)}),a.isDirtyExporting=!1)}function ja(a,b){var c=a.indexOf("</svg>")+6,d=a.substr(c);a=a.substr(0,c);b&&b.exporting&&b.exporting.allowHTML&&d&&(d='<foreignObject x="0" y="0" width="'+b.chart.width+'" height="'+b.chart.height+'"><body xmlns="http://www.w3.org/1999/xhtml">'+d.replace(/(<(?:img|br).*?(?=>))>/g,"$1 />")+"</body></foreignObject>",a=a.replace("</svg>",d+"</svg>"));a=a.replace(/zIndex="[^"]+"/g,"").replace(/symbolName="[^"]+"/g,"").replace(/jQuery[0-9]+="[^"]+"/g,
"").replace(/url\(("|&quot;)(.*?)("|&quot;);?\)/g,"url($2)").replace(/url\([^#]+#/g,"url(#").replace(/<svg /,'<svg xmlns:xlink="http://www.w3.org/1999/xlink" ').replace(/ (|NS[0-9]+:)href=/g," xlink:href=").replace(/\n/," ").replace(/(fill|stroke)="rgba\(([ 0-9]+,[ 0-9]+,[ 0-9]+),([ 0-9\.]+)\)"/g,'$1="rgb($2)" $1-opacity="$3"').replace(/&nbsp;/g,"\u00a0").replace(/&shy;/g,"\u00ad");this.ieSanitizeSVG&&(a=this.ieSanitizeSVG(a));return a}var O=[],da=[/-/,/^(clipPath|cssText|d|height|width)$/,/^font$/,
/[lL]ogical(Width|Height)$/,/perspective/,/TapHighlightColor/,/^transition/,/^length$/],N="fill stroke strokeLinecap strokeLinejoin strokeWidth textAnchor x y".split(" ");e.inlineWhitelist=[];var ca=["clipPath","defs","desc"],I;e.compose=function(a,b){k.compose(b);-1===O.indexOf(a)&&(O.push(a),b=a.prototype,b.afterPrint=z,b.exportChart=W,b.inlineStyles=ba,b.print=ha,b.sanitizeSVG=ja,b.getChartHTML=X,b.getSVG=Z,b.getSVGForExport=aa,b.getFilename=Y,b.moveContainers=ea,b.beforePrint=H,b.contextMenu=
U,b.addButton=t,b.destroyExport=V,b.renderExporting=ia,b.callbacks.push(T),x(a,"init",fa),d.isSafari&&d.win.matchMedia("print").addListener(function(a){I&&(a.matches?I.beforePrint():I.afterPrint())}))}})(H||(H={}));e.exporting=m(z.exporting,e.exporting);e.lang=m(z.lang,e.lang);e.navigation=m(z.navigation,e.navigation);"";"";return H});g(a,"masters/modules/exporting.src.js",[a["Core/Globals.js"],a["Extensions/Exporting/Exporting.js"],a["Core/HttpUtilities.js"]],function(a,e,g){a.HttpUtilities=g;a.ajax=
g.ajax;a.getJSON=g.getJSON;a.post=g.post;e.compose(a.Chart,a.Renderer)})});
//# sourceMappingURL=exporting.js.map
```
