## Supplementary material for "Exploring natural odour landscapes: A case study with implications for human-biting insects": Figure S4: full-screen.js

```
/*
 Highstock JS v9.3.1 (2021-11-05)

 Advanced Highcharts Stock tools

 (c) 2010-2021 Highsoft AS
 Author: Torstein Honsi

 License: www.highcharts.com/license
*/
'use strict';(function(a){"object"===typeof module&&module.exports?(a["default"]=a,module.exports=a):"function"===typeof define&&define.amd?define("highcharts/modules/full-screen",["highcharts"],function(c){a(c);a.Highcharts=c;return a}):a("undefined"!==typeof Highcharts?Highcharts:void 0)})(function(a){function c(a,g,c,e){a.hasOwnProperty(g)||(a[g]=e.apply(null,c))}a=a?a._modules:{};c(a,"Extensions/FullScreen.js",[a["Core/Chart/Chart.js"],a["Core/Globals.js"],a["Core/Renderer/HTML/AST.js"],a["Core/Utilities.js"]],
function(a,c,h,e){var f=e.addEvent;e=function(){function a(b){this.chart=b;this.isOpen=!1;b=b.renderTo;this.browserProps||("function"===typeof b.requestFullscreen?this.browserProps={fullscreenChange:"fullscreenchange",requestFullscreen:"requestFullscreen",exitFullscreen:"exitFullscreen"}:b.mozRequestFullScreen?this.browserProps={fullscreenChange:"mozfullscreenchange",requestFullscreen:"mozRequestFullScreen",exitFullscreen:"mozCancelFullScreen"}:b.webkitRequestFullScreen?this.browserProps={fullscreenChange:"webkitfullscreenchange",
requestFullscreen:"webkitRequestFullScreen",exitFullscreen:"webkitExitFullscreen"}:b.msRequestFullscreen&&(this.browserProps={fullscreenChange:"MSFullscreenChange",requestFullscreen:"msRequestFullscreen",exitFullscreen:"msExitFullscreen"}))}a.prototype.close=function(){var b=this.chart,a=b.options.chart;if(this.isOpen&&this.browserProps&&b.container.ownerDocument instanceof Document)b.container.ownerDocument[this.browserProps.exitFullscreen]();this.unbindFullscreenEvent&&(this.unbindFullscreenEvent=
this.unbindFullscreenEvent());b.setSize(this.origWidth,this.origHeight,!1);this.origHeight=this.origWidth=void 0;a.width=this.origWidthOption;a.height=this.origHeightOption;this.origHeightOption=this.origWidthOption=void 0;this.isOpen=!1;this.setButtonText()};a.prototype.open=function(){var b=this,a=b.chart,d=a.options.chart;d&&(b.origWidthOption=d.width,b.origHeightOption=d.height);b.origWidth=a.chartWidth;b.origHeight=a.chartHeight;if(b.browserProps){var c=f(a.container.ownerDocument,b.browserProps.fullscreenChange,
function(){b.isOpen?(b.isOpen=!1,b.close()):(a.setSize(null,null,!1),b.isOpen=!0,b.setButtonText())}),e=f(a,"destroy",c);b.unbindFullscreenEvent=function(){c();e()};if(d=a.renderTo[b.browserProps.requestFullscreen]())d["catch"](function(){alert("Full screen is not supported inside a frame.")})}};a.prototype.setButtonText=function(){var a=this.chart,c=a.exportDivElements,d=a.options.exporting,e=d&&d.buttons&&d.buttons.contextButton.menuItems;a=a.options.lang;d&&d.menuItemDefinitions&&a&&a.exitFullscreen&&
a.viewFullscreen&&e&&c&&(c=c[e.indexOf("viewFullscreen")])&&h.setElementHTML(c,this.isOpen?a.exitFullscreen:d.menuItemDefinitions.viewFullscreen.text||a.viewFullscreen)};a.prototype.toggle=function(){this.isOpen?this.close():this.open()};return a}();c.Fullscreen=e;f(a,"beforeRender",function(){this.fullscreen=new c.Fullscreen(this)});return c.Fullscreen});c(a,"masters/modules/full-screen.src.js",[],function(){})});
//# sourceMappingURL=full-screen.js.map
```
