## Supplementary material for "Exploring natural odour landscapes: A case study with implications for human-biting insects": Figure S4: funnel3d.js

```
/*
 Highcharts JS v9.3.1 (2021-11-05)

 Highcharts funnel module

 (c) 2010-2021 Kacper Madej

 License: www.highcharts.com/license
*/
'use strict';(function(a){"object"===typeof module&&module.exports?(a["default"]=a,module.exports=a):"function"===typeof define&&define.amd?define("highcharts/modules/funnel3d",["highcharts","highcharts/highcharts-3d","highcharts/modules/cylinder"],function(q){a(q);a.Highcharts=q;return a}):a("undefined"!==typeof Highcharts?Highcharts:void 0)})(function(a){function q(a,k,F,f){a.hasOwnProperty(k)||(a[k]=f.apply(null,F))}a=a?a._modules:{};q(a,"Series/Funnel3D/Funnel3DComposition.js",[a["Core/Color/Color.js"],
a["Core/Globals.js"],a["Core/Renderer/SVG/SVGRenderer3D.js"],a["Core/Utilities.js"]],function(a,k,F,f){var d=a.parse,w=k.charts,N=f.error,O=f.extend,m=f.merge,l;(function(a){function f(a){a.funnel3d=m(a.cuboid,{parts:"top bottom frontUpper backUpper frontLower backLower rightUpper rightLower".split(" "),mainParts:["top","bottom"],sideGroups:["upperGroup","lowerGroup"],sideParts:{upperGroup:["frontUpper","backUpper","rightUpper"],lowerGroup:["frontLower","backLower","rightLower"]},pathType:"funnel3d",
opacitySetter:function(a){var b=this,c=b.parts,h=k.charts[b.renderer.chartIndex],e="group-opacity-"+a+"-"+h.index;b.parts=b.mainParts;b.singleSetterForParts("opacity",a);b.parts=c;h.renderer.filterId||(h.renderer.definition({tagName:"filter",attributes:{id:e},children:[{tagName:"feComponentTransfer",children:[{tagName:"feFuncA",attributes:{type:"table",tableValues:"0 "+a}}]}]}),b.sideGroups.forEach(function(c){b[c].attr({filter:"url(#"+e+")"})}),b.renderer.styledMode&&(h.renderer.definition({tagName:"style",
textContent:".highcharts-"+e+" {filter:url(#"+e+")}"}),b.sideGroups.forEach(function(b){b.addClass("highcharts-"+e)})));return b},fillSetter:function(a){var b=this,c=d(a),h=c.rgba[3],e={top:d(a).brighten(.1).get(),bottom:d(a).brighten(-.2).get()};1>h?(c.rgba[3]=1,c=c.get("rgb"),b.attr({opacity:h})):c=a;c.linearGradient||c.radialGradient||!b.gradientForSides||(c={linearGradient:{x1:0,x2:1,y1:1,y2:1},stops:[[0,d(a).brighten(-.2).get()],[.5,a],[1,d(a).brighten(-.2).get()]]});c.linearGradient?b.sideGroups.forEach(function(a){var h=
b[a].gradientBox,z=c.linearGradient,f=m(c,{linearGradient:{x1:h.x+z.x1*h.width,y1:h.y+z.y1*h.height,x2:h.x+z.x2*h.width,y2:h.y+z.y2*h.height}});b.sideParts[a].forEach(function(b){e[b]=f})}):(m(!0,e,{frontUpper:c,backUpper:c,rightUpper:c,frontLower:c,backLower:c,rightLower:c}),c.radialGradient&&b.sideGroups.forEach(function(c){var a=b[c].gradientBox,e=a.x+a.width/2,h=a.y+a.height/2,z=Math.min(a.width,a.height);b.sideParts[c].forEach(function(c){b[c].setRadialReference([e,h,z])})}));b.singleSetterForParts("fill",
null,e);b.color=b.fill=a;c.linearGradient&&[b.frontLower,b.frontUpper].forEach(function(c){(c=(c=c.element)&&b.renderer.gradients[c.gradient])&&"userSpaceOnUse"!==c.attr("gradientUnits")&&c.attr({gradientUnits:"userSpaceOnUse"})});return b},adjustForGradient:function(){var a=this,b;a.sideGroups.forEach(function(c){var h={x:Number.MAX_VALUE,y:Number.MAX_VALUE},e={x:-Number.MAX_VALUE,y:-Number.MAX_VALUE};a.sideParts[c].forEach(function(c){b=a[c].getBBox(!0);h={x:Math.min(h.x,b.x),y:Math.min(h.y,b.y)};
e={x:Math.max(e.x,b.x+b.width),y:Math.max(e.y,b.y+b.height)}});a[c].gradientBox={x:h.x,width:e.x-h.x,y:h.y,height:e.y-h.y}})},zIndexSetter:function(){this.finishedOnAdd&&this.adjustForGradient();return this.renderer.Element.prototype.zIndexSetter.apply(this,arguments)},onAdd:function(){this.adjustForGradient();this.finishedOnAdd=!0}})}function l(a){var f=a.prototype;O(f,{funnel3d:function(b){var c=this.element3d("funnel3d",b),a=this.styledMode,e={"stroke-width":1,stroke:"none"};c.upperGroup=this.g("funnel3d-upper-group").attr({zIndex:c.frontUpper.zIndex}).add(c);
[c.frontUpper,c.backUpper,c.rightUpper].forEach(function(b){a||b.attr(e);b.add(c.upperGroup)});c.lowerGroup=this.g("funnel3d-lower-group").attr({zIndex:c.frontLower.zIndex}).add(c);[c.frontLower,c.backLower,c.rightLower].forEach(function(b){a||b.attr(e);b.add(c.lowerGroup)});c.gradientForSides=b.gradientForSides;return c},funnel3dPath:function(b){this.getCylinderEnd||N("A required Highcharts module is missing: cylinder.js",!0,w[this.chartIndex]);var c=w[this.chartIndex],a=b.alphaCorrection=90-Math.abs(c.options.chart.options3d.alpha%
180-90),e=f.cuboidPath.call(this,m(b,{depth:b.width,width:(b.width+b.bottom.width)/2})),z=e.isTop,d=!e.isFront,l=!!b.middle,A=this.getCylinderEnd(c,m(b,{x:b.x-b.width/2,z:b.z-b.width/2,alphaCorrection:a})),g=b.bottom.width,k=m(b,{width:g,x:b.x-g/2,z:b.z-g/2,alphaCorrection:a}),v=this.getCylinderEnd(c,k,!0),n=g,u=k,r=v,x=v;l&&(n=b.middle.width,u=m(b,{y:b.y+b.middle.fraction*b.height,width:n,x:b.x-n/2,z:b.z-n/2}),r=this.getCylinderEnd(c,u,!1),x=this.getCylinderEnd(c,u,!1));e={top:A,bottom:v,frontUpper:this.getCylinderFront(A,
r),zIndexes:{group:e.zIndexes.group,top:0!==z?0:3,bottom:1!==z?0:3,frontUpper:d?2:1,backUpper:d?1:2,rightUpper:d?2:1}};e.backUpper=this.getCylinderBack(A,r);A=1!==Math.min(n,b.width)/Math.max(n,b.width);e.rightUpper=this.getCylinderFront(this.getCylinderEnd(c,m(b,{x:b.x-b.width/2,z:b.z-b.width/2,alphaCorrection:A?-a:0}),!1),this.getCylinderEnd(c,m(u,{alphaCorrection:A?-a:0}),!l));l&&(A=1!==Math.min(n,g)/Math.max(n,g),m(!0,e,{frontLower:this.getCylinderFront(x,v),backLower:this.getCylinderBack(x,v),
rightLower:this.getCylinderFront(this.getCylinderEnd(c,m(k,{alphaCorrection:A?-a:0}),!0),this.getCylinderEnd(c,m(u,{alphaCorrection:A?-a:0}),!1)),zIndexes:{frontLower:d?2:1,backLower:d?1:2,rightLower:d?1:2}}));return e}})}a.compose=function(a){F.compose(a);f(a.prototype.elements3d);l(a)}})(l||(l={}));return l});q(a,"Series/Funnel3D/Funnel3DPoint.js",[a["Core/Series/SeriesRegistry.js"],a["Core/Utilities.js"]],function(a,k){var l=this&&this.__extends||function(){var a=function(d,f){a=Object.setPrototypeOf||
{__proto__:[]}instanceof Array&&function(a,d){a.__proto__=d}||function(a,d){for(var f in d)d.hasOwnProperty(f)&&(a[f]=d[f])};return a(d,f)};return function(d,f){function k(){this.constructor=d}a(d,f);d.prototype=null===f?Object.create(f):(k.prototype=f.prototype,new k)}}();k=k.extend;a=function(a){function d(){var d=null!==a&&a.apply(this,arguments)||this;d.dlBoxRaw=void 0;d.options=void 0;d.series=void 0;d.y=void 0;return d}l(d,a);return d}(a.seriesTypes.column.prototype.pointClass);k(a.prototype,
{shapeType:"funnel3d"});return a});q(a,"Series/Funnel3D/Funnel3DSeries.js",[a["Series/Funnel3D/Funnel3DComposition.js"],a["Series/Funnel3D/Funnel3DPoint.js"],a["Core/Globals.js"],a["Extensions/Math3D.js"],a["Core/Series/SeriesRegistry.js"],a["Core/Utilities.js"]],function(a,k,q,f,d,w){var l=this&&this.__extends||function(){var a=function(b,c){a=Object.setPrototypeOf||{__proto__:[]}instanceof Array&&function(a,b){a.__proto__=b}||function(a,b){for(var c in b)b.hasOwnProperty(c)&&(a[c]=b[c])};return a(b,
c)};return function(b,c){function d(){this.constructor=b}a(b,c);b.prototype=null===c?Object.create(c):(d.prototype=c.prototype,new d)}}();q=q.noop;var F=f.perspective,m=d.series,B=d.seriesTypes.column,I=w.extend,M=w.merge,G=w.pick,u=w.relativeLength;f=function(d){function b(){var a=null!==d&&d.apply(this,arguments)||this;a.center=void 0;a.data=void 0;a.options=void 0;a.points=void 0;return a}l(b,d);b.prototype.alignDataLabel=function(a,b,e){var c=a.dlBoxRaw,d=this.chart.inverted,h=a.plotY>G(this.translatedThreshold,
this.yAxis.len),f=G(e.inside,!!this.options.stacking),g={x:c.x,y:c.y,height:0};e.align=G(e.align,!d||f?"center":h?"right":"left");e.verticalAlign=G(e.verticalAlign,d||f?"middle":h?"top":"bottom");"top"!==e.verticalAlign&&(g.y+=c.bottom/("bottom"===e.verticalAlign?1:2));g.width=this.getWidthAt(g.y);this.options.reversed&&(g.width=c.fullWidth-g.width);f?g.x-=g.width/2:"left"===e.align?(e.align="right",g.x-=1.5*g.width):"right"===e.align?(e.align="left",g.x+=g.width/2):g.x-=g.width/2;a.dlBox=g;B.prototype.alignDataLabel.apply(this,
arguments)};b.prototype.bindAxes=function(){m.prototype.bindAxes.apply(this,arguments);I(this.xAxis.options,{gridLineWidth:0,lineWidth:0,title:void 0,tickPositions:[]});M(!0,this.yAxis.options,{gridLineWidth:0,title:void 0,labels:{enabled:!1}})};b.prototype.translate=function(){m.prototype.translate.apply(this,arguments);var a=0,b=this.chart,e=this.options,d=e.reversed,f=e.ignoreHiddenPoint,k=b.plotWidth,l=b.plotHeight,g=0,q=e.center,v=u(q[0],k),n=u(q[1],l),w=u(e.width,k),r,x,t=u(e.height,l),B=u(e.neckWidth,
k),J=u(e.neckHeight,l),E=n-t/2+t-J;k=this.data;var C,K,y,D,L,H,p;this.getWidthAt=x=function(a){var b=n-t/2;return a>E||t===J?B:B+(w-B)*(1-(a-b)/(t-J))};this.center=[v,n,t];this.centerX=v;k.forEach(function(b){f&&!1===b.visible||(a+=b.y)});k.forEach(function(c){L=null;C=a?c.y/a:0;y=n-t/2+g*t;D=y+C*t;r=x(y);H=D-y;p={gradientForSides:G(c.options.gradientForSides,e.gradientForSides),x:v,y:y,height:H,width:r,z:1,top:{width:r}};r=x(D);p.bottom={fraction:C,width:r};y>=E?p.isCylinder=!0:D>E&&(L=D,r=x(E),
D=E,p.bottom.width=r,p.middle={fraction:H?(E-y)/H:0,width:r});d&&(p.y=y=n+t/2-(g+C)*t,p.middle&&(p.middle.fraction=1-(H?p.middle.fraction:0)),r=p.width,p.width=p.bottom.width,p.bottom.width=r);c.shapeArgs=I(c.shapeArgs,p);c.percentage=100*C;c.plotX=v;c.plotY=d?n+t/2-(g+C/2)*t:(y+(L||D))/2;K=F([{x:v,y:c.plotY,z:d?-(w-x(c.plotY))/2:-x(c.plotY)/2}],b,!0)[0];c.tooltipPos=[K.x,K.y];c.dlBoxRaw={x:v,width:x(c.plotY),y:y,bottom:p.height||0,fullWidth:w};f&&!1===c.visible||(g+=C)})};b.compose=a.compose;b.defaultOptions=
M(B.defaultOptions,{center:["50%","50%"],width:"90%",neckWidth:"30%",height:"100%",neckHeight:"25%",reversed:!1,gradientForSides:!0,animation:!1,edgeWidth:0,colorByPoint:!0,showInLegend:!1,dataLabels:{align:"right",crop:!1,inside:!1,overflow:"allow"}});return b}(B);I(f.prototype,{pointClass:k,translate3dShapes:q});d.registerSeriesType("funnel3d",f);"";return f});q(a,"masters/modules/funnel3d.src.js",[a["Core/Renderer/RendererRegistry.js"],a["Series/Funnel3D/Funnel3DSeries.js"]],function(a,k){k.compose(a.getRendererType());
return k})});
//# sourceMappingURL=funnel3d.js.map
```
