## Supplementary material for "Exploring natural odour landscapes: A case study with implications for human-biting insects": Figure S4: funnel.js

```
/*
 Highcharts JS v9.3.1 (2021-11-05)

 Highcharts funnel module

 (c) 2010-2021 Torstein Honsi

 License: www.highcharts.com/license
*/
'use strict';(function(a){"object"===typeof module&&module.exports?(a["default"]=a,module.exports=a):"function"===typeof define&&define.amd?define("highcharts/modules/funnel",["highcharts"],function(l){a(l);a.Highcharts=l;return a}):a("undefined"!==typeof Highcharts?Highcharts:void 0)})(function(a){function l(a,r,f,d){a.hasOwnProperty(r)||(a[r]=d.apply(null,f))}a=a?a._modules:{};l(a,"Series/Funnel/FunnelSeries.js",[a["Core/Chart/Chart.js"],a["Core/Globals.js"],a["Core/Series/SeriesRegistry.js"],a["Core/Utilities.js"]],
function(a,r,f,d){var l=this&&this.__extends||function(){var a=function(b,g){a=Object.setPrototypeOf||{__proto__:[]}instanceof Array&&function(g,e){g.__proto__=e}||function(g,e){for(var a in e)e.hasOwnProperty(a)&&(g[a]=e[a])};return a(b,g)};return function(b,g){function D(){this.constructor=b}a(b,g);b.prototype=null===g?Object.create(g):(D.prototype=g.prototype,new D)}}(),J=r.noop,y=f.series,c=f.seriesTypes.pie;r=d.addEvent;var z=d.extend,L=d.fireEvent,A=d.isArray,B=d.merge,I=d.pick;d=function(a){function b(){var g=
null!==a&&a.apply(this,arguments)||this;g.data=void 0;g.options=void 0;g.points=void 0;return g}l(b,a);b.prototype.alignDataLabel=function(g,a,e,b,k){var u=g.series;b=u.options.reversed;var h=g.dlBox||g.shapeArgs,d=e.align,c=e.verticalAlign,f=((u.options||{}).dataLabels||{}).inside,m=u.center[1];u=u.getWidthAt((b?2*m-g.plotY:g.plotY)-h.height/2+a.height);u="middle"===c?(h.topWidth-h.bottomWidth)/4:(u-h.bottomWidth)/2;m=h.y;var n=h.x,D=I(a.height,a.getBBox().height);"middle"===c?m=h.y-h.height/2+D/
2:"top"===c&&(m=h.y-h.height+D+e.padding);if("top"===c&&!b||"bottom"===c&&b||"middle"===c)"right"===d?n=h.x-e.padding+u:"left"===d&&(n=h.x+e.padding-u);b={x:n,y:b?m-h.height:m,width:h.bottomWidth,height:h.height};e.verticalAlign="bottom";f&&!g.visible||y.prototype.alignDataLabel.call(this,g,a,e,b,k);f&&(!g.visible&&g.dataLabel&&(g.dataLabel.placed=!1),g.contrastColor&&a.css({color:g.contrastColor}))};b.prototype.drawDataLabels=function(){var a=this.data,b=this.options.dataLabels.distance,e,c=a.length;
for(this.center[2]-=2*b;c--;){var k=a[c];var d=(e=k.half)?1:-1;var h=k.plotY;k.labelDistance=I(k.options.dataLabels&&k.options.dataLabels.distance,b);this.maxLabelDistance=Math.max(k.labelDistance,this.maxLabelDistance||0);var y=this.getX(h,e,k);k.labelPosition={natural:{x:0,y:h},"final":{},alignment:e?"right":"left",connectorPosition:{breakAt:{x:y+(k.labelDistance-5)*d,y:h},touchingSliceAt:{x:y+k.labelDistance*d,y:h}}}}f.seriesTypes[this.options.dataLabels.inside?"column":"pie"].prototype.drawDataLabels.call(this)};
b.prototype.translate=function(){function a(a,b){return/%$/.test(a)?b*parseInt(a,10)/100:parseInt(a,10)}var b=0,e=this,c=e.chart,k=e.options,d=k.reversed,h=k.ignoreHiddenPoint,f=c.plotWidth;c=c.plotHeight;var y=0,l=k.center,m=a(l[0],f),n=a(l[1],c),r=a(k.width,f),v,w=a(k.height,c),z=a(k.neckWidth,f),H=a(k.neckHeight,c),A=n-w/2+w-H;f=e.data;var B,F,K="left"===k.dataLabels.position?1:0,E,p,G,x,q,C,t;e.getWidthAt=function(a){var b=n-w/2;return a>A||w===H?z:z+(r-z)*(1-(a-b)/(w-H))};e.getX=function(a,b,
c){return m+(b?-1:1)*(e.getWidthAt(d?2*n-a:a)/2+c.labelDistance)};e.center=[m,n,w];e.centerX=m;f.forEach(function(a){h&&!1===a.visible||(b+=a.y)});f.forEach(function(a){t=null;F=b?a.y/b:0;p=n-w/2+y*w;q=p+F*w;v=e.getWidthAt(p);E=m-v/2;G=E+v;v=e.getWidthAt(q);x=m-v/2;C=x+v;p>A?(E=x=m-z/2,G=C=m+z/2):q>A&&(t=q,v=e.getWidthAt(A),x=m-v/2,C=x+v,q=A);d&&(p=2*n-p,q=2*n-q,null!==t&&(t=2*n-t));B=[["M",E,p],["L",G,p],["L",C,q]];null!==t&&B.push(["L",C,t],["L",x,t]);B.push(["L",x,q],["Z"]);a.shapeType="path";
a.shapeArgs={d:B};a.percentage=100*F;a.plotX=m;a.plotY=(p+(t||q))/2;a.tooltipPos=[m,a.plotY];a.dlBox={x:x,y:p,topWidth:G-E,bottomWidth:C-x,height:Math.abs(I(t,q)-p),width:NaN};a.slice=J;a.half=K;h&&!1===a.visible||(y+=F)});L(e,"afterTranslate")};b.prototype.sortByAngle=function(a){a.sort(function(a,b){return a.plotY-b.plotY})};b.defaultOptions=B(c.defaultOptions,{animation:!1,center:["50%","50%"],width:"90%",neckWidth:"30%",height:"100%",neckHeight:"25%",reversed:!1,size:!0,dataLabels:{connectorWidth:1,
verticalAlign:"middle"},states:{select:{color:"#cccccc",borderColor:"#000000"}}});return b}(c);z(d.prototype,{animate:J});r(a,"afterHideAllOverlappingLabels",function(){this.series.forEach(function(a){var b=a.options&&a.options.dataLabels;A(b)&&(b=b[0]);a.is("pie")&&a.placeDataLabels&&b&&!b.inside&&a.placeDataLabels()})});f.registerSeriesType("funnel",d);"";return d});l(a,"Series/Pyramid/PyramidSeries.js",[a["Series/Funnel/FunnelSeries.js"],a["Core/Series/SeriesRegistry.js"],a["Core/Utilities.js"]],
function(a,l,f){var d=this&&this.__extends||function(){var a=function(d,c){a=Object.setPrototypeOf||{__proto__:[]}instanceof Array&&function(a,c){a.__proto__=c}||function(a,c){for(var d in c)c.hasOwnProperty(d)&&(a[d]=c[d])};return a(d,c)};return function(d,c){function f(){this.constructor=d}a(d,c);d.prototype=null===c?Object.create(c):(f.prototype=c.prototype,new f)}}(),r=f.merge;f=function(f){function l(){var a=null!==f&&f.apply(this,arguments)||this;a.data=void 0;a.options=void 0;a.points=void 0;
return a}d(l,f);l.defaultOptions=r(a.defaultOptions,{neckWidth:"0%",neckHeight:"0%",reversed:!0});return l}(a);l.registerSeriesType("pyramid",f);"";return f});l(a,"masters/modules/funnel.src.js",[],function(){})});
//# sourceMappingURL=funnel.js.map
```
