## Supplementary material for "Exploring natural odour landscapes: A case study with implications for human-biting insects": Figure S4: gantt.js

```
/*
 Highcharts Gantt JS v9.3.1 (2021-11-05)

 Gantt series

 (c) 2016-2021 Lars A. V. Cabrera

 License: www.highcharts.com/license
*/
'use strict';(function(b){"object"===typeof module&&module.exports?(b["default"]=b,module.exports=b):"function"===typeof define&&define.amd?define("highcharts/modules/gantt",["highcharts"],function(J){b(J);b.Highcharts=J;return b}):b("undefined"!==typeof Highcharts?Highcharts:void 0)})(function(b){function J(b,t,v,u){b.hasOwnProperty(t)||(b[t]=u.apply(null,v))}b=b?b._modules:{};J(b,"Series/XRange/XRangePoint.js",[b["Core/Series/Point.js"],b["Core/Series/SeriesRegistry.js"],b["Core/Utilities.js"]],
function(b,t,v){var u=this&&this.__extends||function(){var b=function(l,h){b=Object.setPrototypeOf||{__proto__:[]}instanceof Array&&function(h,d){h.__proto__=d}||function(h,d){for(var p in d)d.hasOwnProperty(p)&&(h[p]=d[p])};return b(l,h)};return function(l,h){function x(){this.constructor=l}b(l,h);l.prototype=null===h?Object.create(h):(x.prototype=h.prototype,new x)}}();v=v.extend;t=function(t){function l(){var h=null!==t&&t.apply(this,arguments)||this;h.options=void 0;h.series=void 0;return h}u(l,
t);l.getColorByCategory=function(h,l){var d=h.options.colors||h.chart.options.colors;h=l.y%(d?d.length:h.chart.options.chart.colorCount);return{colorIndex:h,color:d&&d[h]}};l.prototype.resolveColor=function(){var h=this.series;if(h.options.colorByPoint&&!this.options.color){var x=l.getColorByCategory(h,this);h.chart.styledMode||(this.color=x.color);this.options.colorIndex||(this.colorIndex=x.colorIndex)}else this.color||(this.color=h.color)};l.prototype.init=function(){b.prototype.init.apply(this,
arguments);this.y||(this.y=0);return this};l.prototype.setState=function(){b.prototype.setState.apply(this,arguments);this.series.drawPoint(this,this.series.getAnimationVerb())};l.prototype.getLabelConfig=function(){var h=b.prototype.getLabelConfig.call(this),l=this.series.yAxis.categories;h.x2=this.x2;h.yCategory=this.yCategory=l&&l[this.y];return h};l.prototype.isValid=function(){return"number"===typeof this.x&&"number"===typeof this.x2};return l}(t.seriesTypes.column.prototype.pointClass);v(t.prototype,
{ttBelow:!1,tooltipDateKeys:["x","x2"]});return t});J(b,"Series/XRange/XRangeComposition.js",[b["Core/Axis/Axis.js"],b["Core/Utilities.js"]],function(b,t){var v=t.addEvent,u=t.pick;v(b,"afterGetSeriesExtremes",function(){var b=this.series,l;if(this.isXAxis){var h=u(this.dataMax,-Number.MAX_VALUE);b.forEach(function(b){b.x2Data&&b.x2Data.forEach(function(d){d>h&&(h=d,l=!0)})});l&&(this.dataMax=h)}})});J(b,"Series/XRange/XRangeSeries.js",[b["Core/Globals.js"],b["Core/Color/Color.js"],b["Core/Series/SeriesRegistry.js"],
b["Core/Utilities.js"],b["Series/XRange/XRangePoint.js"]],function(b,t,v,u,B){var l=this&&this.__extends||function(){var a=function(c,m){a=Object.setPrototypeOf||{__proto__:[]}instanceof Array&&function(m,a){m.__proto__=a}||function(m,a){for(var c in a)a.hasOwnProperty(c)&&(m[c]=a[c])};return a(c,m)};return function(c,m){function C(){this.constructor=c}a(c,m);c.prototype=null===m?Object.create(m):(C.prototype=m.prototype,new C)}}(),h=t.parse,x=v.series,d=v.seriesTypes.column,p=d.prototype,g=u.clamp,
A=u.correctFloat,D=u.defined;t=u.extend;var H=u.find,e=u.isNumber,a=u.isObject,c=u.merge,r=u.pick;u=function(F){function G(){var a=null!==F&&F.apply(this,arguments)||this;a.data=void 0;a.options=void 0;a.points=void 0;return a}l(G,F);G.prototype.init=function(){d.prototype.init.apply(this,arguments);this.options.stacking=void 0};G.prototype.getColumnMetrics=function(){function a(){c.series.forEach(function(a){var m=a.xAxis;a.xAxis=a.yAxis;a.yAxis=m})}var c=this.chart;a();var r=p.getColumnMetrics.call(this);
a();return r};G.prototype.cropData=function(a,c,r,e){c=x.prototype.cropData.call(this,this.x2Data,c,r,e);c.xData=a.slice(c.start,c.end);return c};G.prototype.findPointIndex=function(a){var m=this.cropped,c=this.cropStart,r=this.points,g=a.id;if(g)var d=(d=H(r,function(a){return a.id===g}))?d.index:void 0;"undefined"===typeof d&&(d=(d=H(r,function(m){return m.x===a.x&&m.x2===a.x2&&!m.touched}))?d.index:void 0);m&&e(d)&&e(c)&&d>=c&&(d-=c);return d};G.prototype.translatePoint=function(m){var C=this.xAxis,
d=this.yAxis,F=this.columnMetrics,E=this.options,h=E.minPointLength||0,p=(m.shapeArgs&&m.shapeArgs.width||0)/2,z=this.pointXOffset=F.offset,n=m.plotX,y=r(m.x2,m.x+(m.len||0)),q=C.translate(y,0,0,0,1);y=Math.abs(q-n);var f=this.chart.inverted,k=r(E.borderWidth,1)%2/2,w=F.offset,L=Math.round(F.width);h&&(h-=y,0>h&&(h=0),n-=h/2,q+=h/2);n=Math.max(n,-10);q=g(q,-10,C.len+10);D(m.options.pointWidth)&&(w-=(Math.ceil(m.options.pointWidth)-L)/2,L=Math.ceil(m.options.pointWidth));E.pointPlacement&&e(m.plotY)&&
d.categories&&(m.plotY=d.translate(m.y,0,1,0,1,E.pointPlacement));E=Math.floor(Math.min(n,q))+k;E={x:E,y:Math.floor(m.plotY+w)+k,width:Math.floor(Math.max(n,q))+k-E,height:L,r:this.options.borderRadius};m.shapeArgs=E;f?m.tooltipPos[1]+=z+p:m.tooltipPos[0]-=p+z-E.width/2;p=E.x;z=p+E.width;0>p||z>C.len?(p=g(p,0,C.len),z=g(z,0,C.len),q=z-p,m.dlBox=c(E,{x:p,width:z-p,centerX:q?q/2:null})):m.dlBox=null;p=m.tooltipPos;z=f?1:0;q=f?0:1;F=this.columnMetrics?this.columnMetrics.offset:-F.width/2;p[z]=f?p[z]+
E.width/2:p[z]+(C.reversed?-1:0)*E.width;p[q]=g(p[q]+(f?-1:1)*F,0,d.len-1);if(d=m.partialFill)a(d)&&(d=d.amount),e(d)||(d=0),m.partShapeArgs=c(E,{r:this.options.borderRadius}),n=Math.max(Math.round(y*d+m.plotX-n),0),m.clipRectArgs={x:C.reversed?E.x+y-n:E.x,y:E.y,width:n,height:E.height}};G.prototype.translate=function(){p.translate.apply(this,arguments);this.points.forEach(function(a){this.translatePoint(a)},this)};G.prototype.drawPoint=function(m,C){var e=this.options,d=this.chart.renderer,g=m.graphic,
p=m.shapeType,F=m.shapeArgs,z=m.partShapeArgs,n=m.clipRectArgs,y=m.partialFill,q=e.stacking&&!e.borderRadius,f=m.state,k=e.states[f||"normal"]||{},w="undefined"===typeof f?"attr":C;f=this.pointAttribs(m,f);k=r(this.chart.options.chart.animation,k.animation);if(m.isNull||!1===m.visible)g&&(m.graphic=g.destroy());else{if(g)g.rect[C](F);else m.graphic=g=d.g("point").addClass(m.getClassName()).add(m.group||this.group),g.rect=d[p](c(F)).addClass(m.getClassName()).addClass("highcharts-partfill-original").add(g);
z&&(g.partRect?(g.partRect[C](c(z)),g.partialClipRect[C](c(n))):(g.partialClipRect=d.clipRect(n.x,n.y,n.width,n.height),g.partRect=d[p](z).addClass("highcharts-partfill-overlay").add(g).clip(g.partialClipRect)));this.chart.styledMode||(g.rect[C](f,k).shadow(e.shadow,null,q),z&&(a(y)||(y={}),a(e.partialFill)&&(y=c(e.partialFill,y)),m=y.fill||h(f.fill).brighten(-.3).get()||h(m.color||this.color).brighten(-.3).get(),f.fill=m,g.partRect[w](f,k).shadow(e.shadow,null,q)))}};G.prototype.drawPoints=function(){var a=
this,c=a.getAnimationVerb();a.points.forEach(function(m){a.drawPoint(m,c)})};G.prototype.getAnimationVerb=function(){return this.chart.pointCount<(this.options.animationLimit||250)?"animate":"attr"};G.prototype.isPointInside=function(a){var c=a.shapeArgs,m=a.plotX,e=a.plotY;return c?"undefined"!==typeof m&&"undefined"!==typeof e&&0<=e&&e<=this.yAxis.len&&0<=(c.x||0)+(c.width||0)&&m<=this.xAxis.len:F.prototype.isPointInside.apply(this,arguments)};G.defaultOptions=c(d.defaultOptions,{colorByPoint:!0,
dataLabels:{formatter:function(){var c=this.point.partialFill;a(c)&&(c=c.amount);if(e(c)&&0<c)return A(100*c)+"%"},inside:!0,verticalAlign:"middle"},tooltip:{headerFormat:'<span style="font-size: 10px">{point.x} - {point.x2}</span><br/>',pointFormat:'<span style="color:{point.color}">\u25cf</span> {series.name}: <b>{point.yCategory}</b><br/>'},borderRadius:3,pointRange:0});return G}(d);t(u.prototype,{type:"xrange",parallelArrays:["x","x2","y"],requireSorting:!1,animate:x.prototype.animate,cropShoulder:1,
getExtremesFromAll:!0,autoIncrement:b.noop,buildKDTree:b.noop,pointClass:B});v.registerSeriesType("xrange",u);"";return u});J(b,"Series/Gantt/GanttPoint.js",[b["Core/Series/SeriesRegistry.js"],b["Core/Utilities.js"]],function(b,t){var v=this&&this.__extends||function(){var b=function(l,h){b=Object.setPrototypeOf||{__proto__:[]}instanceof Array&&function(h,d){h.__proto__=d}||function(h,d){for(var p in d)d.hasOwnProperty(p)&&(h[p]=d[p])};return b(l,h)};return function(l,h){function x(){this.constructor=
l}b(l,h);l.prototype=null===h?Object.create(h):(x.prototype=h.prototype,new x)}}(),u=t.pick;return function(b){function l(){var h=null!==b&&b.apply(this,arguments)||this;h.options=void 0;h.series=void 0;return h}v(l,b);l.setGanttPointAliases=function(h){function b(d,p){"undefined"!==typeof p&&(h[d]=p)}b("x",u(h.start,h.x));b("x2",u(h.end,h.x2));b("partialFill",u(h.completed,h.partialFill))};l.prototype.applyOptions=function(h,x){h=b.prototype.applyOptions.call(this,h,x);l.setGanttPointAliases(h);
return h};l.prototype.isValid=function(){return("number"===typeof this.start||"number"===typeof this.x)&&("number"===typeof this.end||"number"===typeof this.x2||this.milestone)};return l}(b.seriesTypes.xrange.prototype.pointClass)});J(b,"Core/Axis/BrokenAxis.js",[b["Extensions/Stacking.js"],b["Core/Utilities.js"]],function(b,t){var v=t.addEvent,u=t.find,B=t.fireEvent,l=t.isArray,h=t.isNumber,x=t.pick,d;(function(d){function g(){"undefined"!==typeof this.brokenAxis&&this.brokenAxis.setBreaks(this.options.breaks,
!1)}function p(){this.brokenAxis&&this.brokenAxis.hasBreaks&&(this.options.ordinal=!1)}function D(){var a=this.brokenAxis;if(a&&a.hasBreaks){for(var c=this.tickPositions,e=this.tickPositions.info,r=[],g=0;g<c.length;g++)a.isInAnyBreak(c[g])||r.push(c[g]);this.tickPositions=r;this.tickPositions.info=e}}function H(){this.brokenAxis||(this.brokenAxis=new G(this))}function e(){var a=this.options.connectNulls,c=this.points,e=this.xAxis,g=this.yAxis;if(this.isDirty)for(var r=c.length;r--;){var d=c[r],p=
!(null===d.y&&!1===a)&&(e&&e.brokenAxis&&e.brokenAxis.isInAnyBreak(d.x,!0)||g&&g.brokenAxis&&g.brokenAxis.isInAnyBreak(d.y,!0));d.visible=p?!1:!1!==d.options.visible}}function a(){this.drawBreaks(this.xAxis,["x"]);this.drawBreaks(this.yAxis,x(this.pointArrayMap,["y"]))}function c(a,c){var m=this,e=m.points,g,r,d,z;if(a&&a.brokenAxis&&a.brokenAxis.hasBreaks){var n=a.brokenAxis;c.forEach(function(c){g=n&&n.breakArray||[];r=a.isXAxis?a.min:x(m.options.threshold,a.min);e.forEach(function(n){z=x(n["stack"+
c.toUpperCase()],n[c]);g.forEach(function(f){if(h(r)&&h(z)){d=!1;if(r<f.from&&z>f.to||r>f.from&&z<f.from)d="pointBreak";else if(r<f.from&&z>f.from&&z<f.to||r>f.from&&z>f.to&&z<f.from)d="pointInBreak";d&&B(a,d,{point:n,brk:f})}})})})}}function r(){var a=this.currentDataGrouping,c=a&&a.gapSize;a=this.points.slice();var e=this.yAxis,r=this.options.gapSize,g=a.length-1,d;if(r&&0<g)for("value"!==this.options.gapUnit&&(r*=this.basePointRange),c&&c>r&&c>=this.basePointRange&&(r=c),d=void 0;g--;)d&&!1!==
d.visible||(d=a[g+1]),c=a[g],!1!==d.visible&&!1!==c.visible&&(d.x-c.x>r&&(d=(c.x+d.x)/2,a.splice(g+1,0,{isNull:!0,x:d}),e.stacking&&this.options.stacking&&(d=e.stacking.stacks[this.stackKey][d]=new b(e,e.options.stackLabels,!1,d,this.stack),d.total=0)),d=c);return this.getGraphPath(a)}var F=[];d.compose=function(m,d){-1===F.indexOf(m)&&(F.push(m),m.keepProps.push("brokenAxis"),v(m,"init",H),v(m,"afterInit",g),v(m,"afterSetTickPositions",D),v(m,"afterSetOptions",p));if(-1===F.indexOf(d)){F.push(d);
var C=d.prototype;C.drawBreaks=c;C.gappedPath=r;v(d,"afterGeneratePoints",e);v(d,"afterRender",a)}return m};var G=function(){function a(a){this.hasBreaks=!1;this.axis=a}a.isInBreak=function(a,c){var d=a.repeat||Infinity,r=a.from,e=a.to-a.from;c=c>=r?(c-r)%d:d-(r-c)%d;return a.inclusive?c<=e:c<e&&0!==c};a.lin2Val=function(c){var d=this.brokenAxis;d=d&&d.breakArray;if(!d||!h(c))return c;var r;for(r=0;r<d.length;r++){var e=d[r];if(e.from>=c)break;else e.to<c?c+=e.len:a.isInBreak(e,c)&&(c+=e.len)}return c};
a.val2Lin=function(c){var d=this.brokenAxis;d=d&&d.breakArray;if(!d||!h(c))return c;var r=c,e;for(e=0;e<d.length;e++){var g=d[e];if(g.to<=c)r-=g.len;else if(g.from>=c)break;else if(a.isInBreak(g,c)){r-=c-g.from;break}}return r};a.prototype.findBreakAt=function(a,c){return u(c,function(c){return c.from<a&&a<c.to})};a.prototype.isInAnyBreak=function(c,d){var e=this.axis,r=e.options.breaks||[],g=r.length,m;if(g&&h(c)){for(;g--;)if(a.isInBreak(r[g],c)){var z=!0;m||(m=x(r[g].showPoints,!e.isXAxis))}var n=
z&&d?z&&!m:z}return n};a.prototype.setBreaks=function(c,d){var r=this,e=r.axis,g=l(c)&&!!c.length;e.isDirty=r.hasBreaks!==g;r.hasBreaks=g;e.options.breaks=e.userOptions.breaks=c;e.forceRedraw=!0;e.series.forEach(function(a){a.isDirty=!0});g||e.val2lin!==a.val2Lin||(delete e.val2lin,delete e.lin2val);g&&(e.userOptions.ordinal=!1,e.lin2val=a.lin2Val,e.val2lin=a.val2Lin,e.setExtremes=function(a,c,n,y,q){if(r.hasBreaks){for(var f=this.options.breaks||[],k;k=r.findBreakAt(a,f);)a=k.to;for(;k=r.findBreakAt(c,
f);)c=k.from;c<a&&(c=a)}e.constructor.prototype.setExtremes.call(this,a,c,n,y,q)},e.setAxisTranslation=function(){e.constructor.prototype.setAxisTranslation.call(this);r.unitLength=void 0;if(r.hasBreaks){var c=e.options.breaks||[],d=[],n=[],y=x(e.pointRangePadding,0),q=0,f,k=e.userMin||e.min,w=e.userMax||e.max,g;c.forEach(function(c){f=c.repeat||Infinity;h(k)&&h(w)&&(a.isInBreak(c,k)&&(k+=c.to%f-k%f),a.isInBreak(c,w)&&(w-=w%f-c.from%f))});c.forEach(function(a){O=a.from;f=a.repeat||Infinity;if(h(k)&&
h(w)){for(;O-f>k;)O-=f;for(;O<k;)O+=f;for(g=O;g<w;g+=f)d.push({value:g,move:"in"}),d.push({value:g+a.to-a.from,move:"out",size:a.breakSize})}});d.sort(function(f,k){return f.value===k.value?("in"===f.move?0:1)-("in"===k.move?0:1):f.value-k.value});var N=0;var O=k;d.forEach(function(f){N+="in"===f.move?1:-1;1===N&&"in"===f.move&&(O=f.value);0===N&&h(O)&&(n.push({from:O,to:f.value,len:f.value-O-(f.size||0)}),q+=f.value-O-(f.size||0))});r.breakArray=n;h(k)&&h(w)&&h(e.min)&&(r.unitLength=w-k-q+y,B(e,
"afterBreaks"),e.staticScale?e.transA=e.staticScale:r.unitLength&&(e.transA*=(w-e.min+y)/r.unitLength),y&&(e.minPixelPadding=e.transA*(e.minPointOffset||0)),e.min=k,e.max=w)}});x(d,!0)&&e.chart.redraw()};return a}();d.Additions=G})(d||(d={}));return d});J(b,"Core/Axis/GridAxis.js",[b["Core/Axis/Axis.js"],b["Core/Axis/AxisDefaults.js"],b["Core/Globals.js"],b["Core/Utilities.js"]],function(b,t,v,u){var B=v.dateFormats,l=u.addEvent,h=u.defined,x=u.erase,d=u.find,p=u.isArray,g=u.isNumber,A=u.merge,D=
u.pick,H=u.timeUnits,e=u.wrap,a;(function(a){function c(f,k){var a={width:0,height:0};k.forEach(function(k){k=f[k];if(u.isObject(k,!0)){var c=u.isObject(k.label,!0)?k.label:{};k=c.getBBox?c.getBBox().height:0;c.textStr&&!g(c.textPxLength)&&(c.textPxLength=c.getBBox().width);var n=g(c.textPxLength)?Math.round(c.textPxLength):0;c.textStr&&(n=Math.round(c.getBBox().width));a.height=Math.max(k,a.height);a.width=Math.max(n,a.width)}});"treegrid"===this.options.type&&this.treeGrid&&this.treeGrid.mapOfPosToGridNode&&
(a.width+=this.options.labels.indentation*((this.treeGrid.mapOfPosToGridNode[-1].height||0)-1));return a}function F(){var f=this.grid;(f&&f.columns||[]).forEach(function(f){f.getOffset()})}function G(f){if(!0===(this.options.grid||{}).enabled){var k=this.axisTitle,c=this.height,n=this.horiz,q=this.left,e=this.offset,N=this.opposite,w=this.options,y=this.top,d=this.width,r=this.tickSize(),g=k&&k.getBBox().width,m=w.title.x,z=w.title.y,L=D(w.title.margin,n?5:10);k=this.chart.renderer.fontMetrics(w.title.style.fontSize,
k).f;r=(n?y+c:q)+(n?1:-1)*(N?-1:1)*(r?r[0]/2:0)+(this.side===a.Side.bottom?k:0);f.titlePosition.x=n?q-(g||0)/2-L+m:r+(N?d:0)+e+m;f.titlePosition.y=n?r-(N?c:0)+(N?k:-k)/2+e+z:y-L+z}}function m(){var f=this.chart,k=this.options.grid;k=void 0===k?{}:k;var a=this.userOptions;if(k.enabled){var c=this.options;c.labels.align=D(c.labels.align,"center");this.categories||(c.showLastLabel=!1);this.labelRotation=0;c.labels.rotation=0}if(k.columns){c=this.grid.columns=[];for(var n=this.grid.columnIndex=0;++n<
k.columns.length;){var q=A(a,k.columns[k.columns.length-n-1],{linkedTo:0,type:"category",scrollbar:{enabled:!1}});delete q.grid.columns;q=new b(this.chart,q);q.grid.isColumn=!0;q.grid.columnIndex=n;x(f.axes,q);x(f[this.coll],q);c.push(q)}}}function C(){var f=this.grid,k=this.options;if(!0===(k.grid||{}).enabled){var c=this.min||0,n=this.max||0;this.maxLabelDimensions=this.getMaxLabelDimensions(this.ticks,this.tickPositions);this.rightWall&&this.rightWall.destroy();if(this.grid&&this.grid.isOuterAxis()&&
this.axisLine){var q=k.lineWidth;if(q){q=this.getLinePath(q);var e=q[0],w=q[1],y=((this.tickSize("tick")||[1])[0]-1)*(this.side===a.Side.top||this.side===a.Side.left?-1:1);"M"===e[0]&&"L"===w[0]&&(this.horiz?(e[2]+=y,w[2]+=y):(e[1]+=y,w[1]+=y));!this.horiz&&this.chart.marginRight&&(e=[e,["L",this.left,e[2]||0]],y=["L",this.chart.chartWidth-this.chart.marginRight,this.toPixels(n+this.tickmarkOffset)],w=[["M",w[1]||0,this.toPixels(n+this.tickmarkOffset)],y],this.grid.upperBorder||0===c%1||(this.grid.upperBorder=
this.grid.renderBorder(e)),this.grid.upperBorder&&(this.grid.upperBorder.attr({stroke:k.lineColor,"stroke-width":k.lineWidth}),this.grid.upperBorder.animate({d:e})),this.grid.lowerBorder||0===n%1||(this.grid.lowerBorder=this.grid.renderBorder(w)),this.grid.lowerBorder&&(this.grid.lowerBorder.attr({stroke:k.lineColor,"stroke-width":k.lineWidth}),this.grid.lowerBorder.animate({d:w})));this.grid.axisLineExtra?(this.grid.axisLineExtra.attr({stroke:k.lineColor,"stroke-width":k.lineWidth}),this.grid.axisLineExtra.animate({d:q})):
this.grid.axisLineExtra=this.grid.renderBorder(q);this.axisLine[this.showAxis?"show":"hide"](!0)}}(f&&f.columns||[]).forEach(function(f){f.render()});if(!this.horiz&&this.chart.hasRendered&&(this.scrollbar||this.linkedParent&&this.linkedParent.scrollbar)){f=this.tickmarkOffset;k=this.tickPositions[this.tickPositions.length-1];q=this.tickPositions[0];for(w=void 0;(w=this.hiddenLabels.pop())&&w.element;)w.show();(w=this.ticks[q].label)&&(c-q>f?this.hiddenLabels.push(w.hide()):w.show());(w=this.ticks[k].label)&&
(k-n>f?this.hiddenLabels.push(w.hide()):w.show());(c=this.ticks[k].mark)&&(k-n<f&&0<k-n&&this.ticks[k].isLast?c.hide():this.ticks[k-1]&&c.show())}}}function Q(){var f=this.tickPositions&&this.tickPositions.info,k=this.options,a=this.userOptions.labels||{};(k.grid||{}).enabled&&(this.horiz?(this.series.forEach(function(f){f.options.pointRange=0}),f&&k.dateTimeLabelFormats&&k.labels&&!h(a.align)&&(!1===k.dateTimeLabelFormats[f.unitName].range||1<f.count)&&(k.labels.align="left",h(a.x)||(k.labels.x=
3))):"treegrid"!==this.options.type&&this.grid&&this.grid.columns&&(this.minPointOffset=this.tickInterval))}function M(f){var k=this.options;f=f.userOptions;var a=k&&u.isObject(k.grid,!0)?k.grid:{};if(!0===a.enabled){var c=A(!0,{className:"highcharts-grid-axis "+(f.className||""),dateTimeLabelFormats:{hour:{list:["%H:%M","%H"]},day:{list:["%A, %e. %B","%a, %e. %b","%E"]},week:{list:["Week %W","W%W"]},month:{list:["%B","%b","%o"]}},grid:{borderWidth:1},labels:{padding:2,style:{fontSize:"13px"}},margin:0,
title:{text:null,reserveSpace:!1,rotation:0},units:[["millisecond",[1,10,100]],["second",[1,10]],["minute",[1,5,15]],["hour",[1,6]],["day",[1]],["week",[1]],["month",[1]],["year",null]]},f);"xAxis"===this.coll&&(h(f.linkedTo)&&!h(f.tickPixelInterval)&&(c.tickPixelInterval=350),h(f.tickPixelInterval)||!h(f.linkedTo)||h(f.tickPositioner)||h(f.tickInterval)||(c.tickPositioner=function(f,k){var a=this.linkedParent&&this.linkedParent.tickPositions&&this.linkedParent.tickPositions.info;if(a){for(var n=
c.units||[],q=void 0,w=void 0,e=void 0,y=0;y<n.length;y++)if(n[y][0]===a.unitName){q=y;break}n[q+1]?(e=n[q+1][0],w=(n[q+1][1]||[1])[0]):"year"===a.unitName&&(e="year",w=10*a.count);a=H[e];this.tickInterval=a*w;return this.getTimeTicks({unitRange:a,count:w,unitName:e},f,k,this.options.startOfWeek)}}));A(!0,this.options,c);this.horiz&&(k.minPadding=D(f.minPadding,0),k.maxPadding=D(f.maxPadding,0));g(k.grid.borderWidth)&&(k.tickWidth=k.lineWidth=a.borderWidth)}}function E(f){f=(f=f.userOptions)&&f.grid||
{};var k=f.columns;f.enabled&&k&&A(!0,this.options,k[k.length-1])}function K(){(this.grid.columns||[]).forEach(function(f){f.setScale()})}function I(f){var k=t.defaultLeftAxisOptions,a=this.horiz,c=this.maxLabelDimensions,n=this.options.grid;n=void 0===n?{}:n;n.enabled&&c&&(k=2*Math.abs(k.labels.x),a=a?n.cellHeight||k+c.height:k+c.width,p(f.tickSize)?f.tickSize[0]=a:f.tickSize=[a,0])}function z(){this.axes.forEach(function(f){(f.grid&&f.grid.columns||[]).forEach(function(f){f.setAxisSize();f.setAxisTranslation()})})}
function n(f){var k=this.grid;(k.columns||[]).forEach(function(k){k.destroy(f.keepEvents)});k.columns=void 0}function y(f){f=f.userOptions||{};var k=f.grid||{};k.enabled&&h(k.borderColor)&&(f.tickColor=f.lineColor=k.borderColor);this.grid||(this.grid=new L(this));this.hiddenLabels=[]}function q(f){var k=this.label,c=this.axis,n=c.reversed,q=c.chart,w=c.options.grid||{},e=c.options.labels,y=e.align,d=a.Side[c.side],r=f.tickmarkOffset,m=c.tickPositions,N=this.pos-r;m=g(m[f.index+1])?m[f.index+1]-r:
(c.max||0)+r;var z=c.tickSize("tick");r=z?z[0]:0;z=z?z[1]/2:0;if(!0===w.enabled){if("top"===d){w=c.top+c.offset;var L=w-r}else"bottom"===d?(L=q.chartHeight-c.bottom+c.offset,w=L+r):(w=c.top+c.len-(c.translate(n?m:N)||0),L=c.top+c.len-(c.translate(n?N:m)||0));"right"===d?(d=q.chartWidth-c.right+c.offset,n=d+r):"left"===d?(n=c.left+c.offset,d=n-r):(d=Math.round(c.left+(c.translate(n?m:N)||0))-z,n=Math.min(Math.round(c.left+(c.translate(n?N:m)||0))-z,c.left+c.len));this.slotWidth=n-d;f.pos.x="left"===
y?d:"right"===y?n:d+(n-d)/2;f.pos.y=L+(w-L)/2;q=q.renderer.fontMetrics(e.style.fontSize,k&&k.element);k=k?k.getBBox().height:0;e.useHTML?f.pos.y+=q.b+-(k/2):(k=Math.round(k/q.h),f.pos.y+=(q.b-(q.h-q.f))/2+-((k-1)*q.h/2));f.pos.x+=c.horiz&&e.x||0}}function f(f){var k=f.axis,a=f.value;if(k.options.grid&&k.options.grid.enabled){var c=k.tickPositions,n=(k.linkedParent||k).series[0],q=a===c[0];c=a===c[c.length-1];var w=n&&d(n.options.data,function(f){return f[k.isXAxis?"x":"y"]===a}),e=void 0;w&&n.is("gantt")&&
(e=A(w),v.seriesTypes.gantt.prototype.pointClass.setGanttPointAliases(e));f.isFirst=q;f.isLast=c;f.point=e}}function k(){var f=this.options,k=this.categories,a=this.tickPositions,c=a[0],n=a[a.length-1],q=this.linkedParent&&this.linkedParent.min||this.min,w=this.linkedParent&&this.linkedParent.max||this.max,e=this.tickInterval;!0!==(f.grid||{}).enabled||k||!this.horiz&&!this.isLinked||(c<q&&c+e>q&&!f.startOnTick&&(a[0]=q),n>w&&n-e<w&&!f.endOnTick&&(a[a.length-1]=w))}function w(f){var k=this.options.grid;
return!0===(void 0===k?{}:k).enabled&&this.categories?this.tickInterval:f.apply(this,Array.prototype.slice.call(arguments,1))}(function(f){f[f.top=0]="top";f[f.right=1]="right";f[f.bottom=2]="bottom";f[f.left=3]="left"})(a.Side||(a.Side={}));a.compose=function(a,d,r){-1===a.keepProps.indexOf("grid")&&(a.keepProps.push("grid"),a.prototype.getMaxLabelDimensions=c,e(a.prototype,"unsquish",w),l(a,"init",y),l(a,"afterGetOffset",F),l(a,"afterGetTitlePosition",G),l(a,"afterInit",m),l(a,"afterRender",C),
l(a,"afterSetAxisTranslation",Q),l(a,"afterSetOptions",M),l(a,"afterSetOptions",E),l(a,"afterSetScale",K),l(a,"afterTickSize",I),l(a,"trimTicks",k),l(a,"destroy",n));l(d,"afterSetChartSize",z);l(r,"afterGetLabelPosition",q);l(r,"labelFormat",f);return a};var L=function(){function f(f){this.axis=f}f.prototype.isOuterAxis=function(){var f=this.axis,k=f.grid.columnIndex,a=f.linkedParent&&f.linkedParent.grid.columns||f.grid.columns,c=k?f.linkedParent:f,n=-1,q=0;f.chart[f.coll].forEach(function(k,a){k.side!==
f.side||k.options.isInternal||(q=a,k===c&&(n=a))});return q===n&&(g(k)?a.length===k:!0)};f.prototype.renderBorder=function(f){var k=this.axis,a=k.chart.renderer,c=k.options;f=a.path(f).addClass("highcharts-axis-line").add(k.axisBorder);a.styledMode||f.attr({stroke:c.lineColor,"stroke-width":c.lineWidth,zIndex:7});return f};return f}();a.Additions=L})(a||(a={}));B.E=function(a){return this.dateFormat("%a",a,!0).charAt(0)};B.W=function(a){a=new this.Date(a);var c=(this.get("Day",a)+6)%7,e=new this.Date(a.valueOf());
this.set("Date",e,this.get("Date",a)-c+3);c=new this.Date(this.get("FullYear",e),0,1);4!==this.get("Day",c)&&(this.set("Month",a,0),this.set("Date",a,1+(11-this.get("Day",c))%7));return(1+Math.floor((e.valueOf()-c.valueOf())/6048E5)).toString()};"";return a});J(b,"Gantt/Tree.js",[b["Core/Utilities.js"]],function(b){var t=b.extend,v=b.isNumber,u=b.pick,B=function(h,b){var d=h.reduce(function(d,g){var p=u(g.parent,"");"undefined"===typeof d[p]&&(d[p]=[]);d[p].push(g);return d},{});Object.keys(d).forEach(function(p,
g){var h=d[p];""!==p&&-1===b.indexOf(p)&&(h.forEach(function(d){g[""].push(d)}),delete g[p])});return d},l=function(h,b,d,p,g,A){var D=0,H=0,e=A&&A.after,a=A&&A.before;b={data:p,depth:d-1,id:h,level:d,parent:b};var c,r;"function"===typeof a&&a(b,A);a=(g[h]||[]).map(function(a){var e=l(a.id,h,d+1,a,g,A),m=a.start;a=!0===a.milestone?m:a.end;c=!v(c)||m<c?m:c;r=!v(r)||a>r?a:r;D=D+1+e.descendants;H=Math.max(e.height+1,H);return e});p&&(p.start=u(p.start,c),p.end=u(p.end,r));t(b,{children:a,descendants:D,
height:H});"function"===typeof e&&e(b,A);return b};return{getListOfParents:B,getNode:l,getTree:function(h,b){var d=h.map(function(d){return d.id});h=B(h,d);return l("",null,1,null,h,b)}}});J(b,"Core/Axis/TreeGridTick.js",[b["Core/Utilities.js"]],function(b){var t=b.addEvent,v=b.isObject,u=b.isNumber,B=b.pick,l=b.wrap,h;(function(b){function d(){this.treeGrid||(this.treeGrid=new H(this))}function p(e,a){e=e.treeGrid;var c=!e.labelIcon,d=a.renderer,g=a.xy,b=a.options,m=b.width||0,p=b.height||0,h=g.x-
m/2-(b.padding||0);g=g.y-p/2;var l=a.collapsed?90:180,A=a.show&&u(g),D=e.labelIcon;D||(e.labelIcon=D=d.path(d.symbols[b.type](b.x||0,b.y||0,m,p)).addClass("highcharts-label-icon").add(a.group));D.attr({y:A?0:-9999});d.styledMode||D.attr({cursor:"pointer",fill:B(a.color,"#666666"),"stroke-width":1,stroke:b.lineColor,strokeWidth:b.lineWidth||0});D[c?"attr":"animate"]({translateX:h,translateY:g,rotation:l})}function g(e,a,c,d,g,b,m,p,h){var r=B(this.options&&this.options.labels,b);b=this.pos;var C=this.axis,
F="treegrid"===C.options.type;e=e.apply(this,[a,c,d,g,r,m,p,h]);F&&(a=r&&v(r.symbol,!0)?r.symbol:{},r=r&&u(r.indentation)?r.indentation:0,b=(b=(C=C.treeGrid.mapOfPosToGridNode)&&C[b])&&b.depth||1,e.x+=(a.width||0)+2*(a.padding||0)+(b-1)*r);return e}function h(e){var a=this,c=a.pos,d=a.axis,g=a.label,b=d.treeGrid.mapOfPosToGridNode,m=d.options,h=B(a.options&&a.options.labels,m&&m.labels),l=h&&v(h.symbol,!0)?h.symbol:{},A=(b=b&&b[c])&&b.depth;m="treegrid"===m.type;var D=-1<d.tickPositions.indexOf(c);
c=d.chart.styledMode;m&&b&&g&&g.element&&g.addClass("highcharts-treegrid-node-level-"+A);e.apply(a,Array.prototype.slice.call(arguments,1));m&&g&&g.element&&b&&b.descendants&&0<b.descendants&&(d=d.treeGrid.isCollapsed(b),p(a,{color:!c&&g.styles&&g.styles.color||"",collapsed:d,group:g.parentGroup,options:l,renderer:g.renderer,show:D,xy:g.xy}),l="highcharts-treegrid-node-"+(d?"expanded":"collapsed"),g.addClass("highcharts-treegrid-node-"+(d?"collapsed":"expanded")).removeClass(l),c||g.css({cursor:"pointer"}),
[g,a.treeGrid.labelIcon].forEach(function(c){c&&!c.attachedTreeGridEvents&&(t(c.element,"mouseover",function(){g.addClass("highcharts-treegrid-node-active");g.renderer.styledMode||g.css({textDecoration:"underline"})}),t(c.element,"mouseout",function(){var a=v(h.style)?h.style:{};g.removeClass("highcharts-treegrid-node-active");g.renderer.styledMode||g.css({textDecoration:a.textDecoration})}),t(c.element,"click",function(){a.treeGrid.toggleCollapse()}),c.attachedTreeGridEvents=!0)}))}var D=!1;b.compose=
function(e){D||(t(e,"init",d),l(e.prototype,"getLabelPosition",g),l(e.prototype,"renderLabel",h),e.prototype.collapse=function(a){this.treeGrid.collapse(a)},e.prototype.expand=function(a){this.treeGrid.expand(a)},e.prototype.toggleCollapse=function(a){this.treeGrid.toggleCollapse(a)},D=!0)};var H=function(){function e(a){this.tick=a}e.prototype.collapse=function(a){var c=this.tick,e=c.axis,d=e.brokenAxis;d&&e.treeGrid.mapOfPosToGridNode&&(c=e.treeGrid.collapse(e.treeGrid.mapOfPosToGridNode[c.pos]),
d.setBreaks(c,B(a,!0)))};e.prototype.expand=function(a){var c=this.tick,e=c.axis,d=e.brokenAxis;d&&e.treeGrid.mapOfPosToGridNode&&(c=e.treeGrid.expand(e.treeGrid.mapOfPosToGridNode[c.pos]),d.setBreaks(c,B(a,!0)))};e.prototype.toggleCollapse=function(a){var c=this.tick,e=c.axis,d=e.brokenAxis;d&&e.treeGrid.mapOfPosToGridNode&&(c=e.treeGrid.toggleCollapse(e.treeGrid.mapOfPosToGridNode[c.pos]),d.setBreaks(c,B(a,!0)))};return e}();b.Additions=H})(h||(h={}));return h});J(b,"Series/TreeUtilities.js",[b["Core/Color/Color.js"],
b["Core/Utilities.js"]],function(b,t){function v(b,g){var h=g.before,p=g.idRoot,l=g.mapIdToNode[p],e=g.points[b.i],a=e&&e.options||{},c=[],r=0;b.levelDynamic=b.level-(!1!==g.levelIsConstant?0:l.level);b.name=d(e&&e.name,"");b.visible=p===b.id||!0===g.visible;"function"===typeof h&&(b=h(b,g));b.children.forEach(function(a,e){var d=u({},g);u(d,{index:e,siblings:b.children.length,visible:b.visible});a=v(a,d);c.push(a);a.visible&&(r+=a.val)});h=d(a.value,r);b.visible=0<=h&&(0<r||b.visible);b.children=
c;b.childrenTotal=r;b.isLeaf=b.visible&&!r;b.val=h;return b}var u=t.extend,B=t.isArray,l=t.isNumber,h=t.isObject,x=t.merge,d=t.pick;return{getColor:function(h,g){var l=g.index,p=g.mapOptionsToLevel,H=g.parentColor,e=g.parentColorIndex,a=g.series,c=g.colors,r=g.siblings,F=a.points,G=a.chart.options.chart,m;if(h){F=F[h.i];h=p[h.level]||{};if(p=F&&h.colorByPoint){var C=F.index%(c?c.length:G.colorCount);var x=c&&c[C]}if(!a.chart.styledMode){c=F&&F.options.color;G=h&&h.color;if(m=H)m=(m=h&&h.colorVariation)&&
"brightness"===m.key&&l&&r?b.parse(H).brighten(l/r*m.to).get():H;m=d(c,G,x,m,a.color)}var t=d(F&&F.options.colorIndex,h&&h.colorIndex,C,e,g.colorIndex)}return{color:m,colorIndex:t}},getLevelOptions:function(b){var g=null;if(h(b)){g={};var A=l(b.from)?b.from:1;var p=b.levels;var H={};var e=h(b.defaults)?b.defaults:{};B(p)&&(H=p.reduce(function(a,c){if(h(c)&&l(c.level)){var g=x({},c);var b=d(g.levelIsConstant,e.levelIsConstant);delete g.levelIsConstant;delete g.level;c=c.level+(b?0:A-1);h(a[c])?x(!0,
a[c],g):a[c]=g}return a},{}));p=l(b.to)?b.to:1;for(b=0;b<=p;b++)g[b]=x({},e,h(H[b])?H[b]:{})}return g},setTreeValues:v,updateRootId:function(b){if(h(b)){var g=h(b.options)?b.options:{};g=d(b.rootNode,g.rootId,"");h(b.userOptions)&&(b.userOptions.rootId=g);b.rootNode=g}return g}}});J(b,"Core/Axis/TreeGridAxis.js",[b["Core/Axis/BrokenAxis.js"],b["Core/Axis/GridAxis.js"],b["Gantt/Tree.js"],b["Core/Axis/TreeGridTick.js"],b["Series/TreeUtilities.js"],b["Core/Utilities.js"]],function(b,t,v,u,B,l){var h=
B.getLevelOptions,x=l.addEvent,d=l.find,p=l.fireEvent,g=l.isArray,A=l.isObject,D=l.isString,H=l.merge,e=l.pick,a=l.wrap,c;(function(c){function r(a,c){var n=a.collapseEnd||0;a=a.collapseStart||0;n>=c&&(a-=.5);return{from:a,to:n,showPoints:!1}}function l(a,c,n){var e=[],q=[],f={},k="boolean"===typeof c?c:!1,w={},g=-1;a=v.getTree(a,{after:function(f){f=w[f.pos];var k=0,a=0;f.children.forEach(function(f){a+=(f.descendants||0)+1;k=Math.max((f.height||0)+1,k)});f.descendants=a;f.height=k;f.collapsed&&
q.push(f)},before:function(a){var c=A(a.data,!0)?a.data:{},n=D(c.name)?c.name:"",q=f[a.parent];q=A(q,!0)?w[q.pos]:null;var y=function(f){return f.name===n},b;k&&A(q,!0)&&(b=d(q.children,y))?(y=b.pos,b.nodes.push(a)):y=g++;w[y]||(w[y]=b={depth:q?q.depth+1:0,name:n,id:c.id,nodes:[a],children:[],pos:y},-1!==y&&e.push(n),A(q,!0)&&q.children.push(b));D(a.id)&&(f[a.id]=a);b&&!0===c.collapsed&&(b.collapsed=!0);a.pos=y}});w=function(f,k){var a=function(f,c,n){var q=c+(-1===c?0:k-1),e=(q-c)/2,w=c+e;f.nodes.forEach(function(f){var k=
f.data;A(k,!0)&&(k.y=c+(k.seriesIndex||0),delete k.seriesIndex);f.pos=w});n[w]=f;f.pos=w;f.tickmarkOffset=e+.5;f.collapseStart=q+.5;f.children.forEach(function(f){a(f,q+1,n);q=(f.collapseEnd||0)-.5});f.collapseEnd=q+.5;return n};return a(f["-1"],-1,{})}(w,n);return{categories:e,mapOfIdToNode:f,mapOfPosToGridNode:w,collapsedNodes:q,tree:a}}function m(a){a.target.axes.filter(function(a){return"treegrid"===a.options.type}).forEach(function(c){var n=c.options||{},e=n.labels,q=n.uniqueNames;n=n.max;var f=
0;if(!c.treeGrid.mapOfPosToGridNode||c.series.some(function(f){return!f.hasRendered||f.isDirtyData||f.isDirty})){var k=c.series.reduce(function(k,a){a.visible&&((a.options.data||[]).forEach(function(c){a.options.keys&&a.options.keys.length&&(c=a.pointClass.prototype.optionsToObject.call({series:a},c),a.pointClass.setGanttPointAliases(c));A(c,!0)&&(c.seriesIndex=f,k.push(c))}),!0===q&&f++);return k},[]);if(n&&k.length<n)for(var w=k.length;w<=n;w++)k.push({name:w+"\u200b"});n=l(k,q||!1,!0===q?f:1);
c.categories=n.categories;c.treeGrid.mapOfPosToGridNode=n.mapOfPosToGridNode;c.hasNames=!0;c.treeGrid.tree=n.tree;c.series.forEach(function(f){var a=(f.options.data||[]).map(function(a){g(a)&&f.options.keys&&f.options.keys.length&&k.forEach(function(f){0<=a.indexOf(f.x)&&0<=a.indexOf(f.x2)&&(a=f)});return A(a,!0)?H(a):a});f.visible&&f.setData(a,!1)});c.treeGrid.mapOptionsToLevel=h({defaults:e,from:1,levels:e&&e.levels,to:c.treeGrid.tree&&c.treeGrid.tree.height});"beforeRender"===a.type&&(c.treeGrid.collapsedNodes=
n.collapsedNodes)}})}function C(a,c){var n=this.treeGrid.mapOptionsToLevel||{},e=this.ticks,q=e[c],f;if("treegrid"===this.options.type&&this.treeGrid.mapOfPosToGridNode){var k=this.treeGrid.mapOfPosToGridNode[c];(n=n[k.depth])&&(f={labels:n});!q&&E?e[c]=new E(this,c,void 0,void 0,{category:k.name,tickmarkOffset:k.tickmarkOffset,options:f}):(q.parameters.category=k.name,q.options=f,q.addLabel())}else a.apply(this,Array.prototype.slice.call(arguments,1))}function Q(a,c,n){var e=this,q="treegrid"===
n.type;e.treeGrid||(e.treeGrid=new K(e));q&&(x(c,"beforeRender",m),x(c,"beforeRedraw",m),x(c,"addSeries",function(f){f.options.data&&(f=l(f.options.data,n.uniqueNames||!1,1),e.treeGrid.collapsedNodes=(e.treeGrid.collapsedNodes||[]).concat(f.collapsedNodes))}),x(e,"foundExtremes",function(){e.treeGrid.collapsedNodes&&e.treeGrid.collapsedNodes.forEach(function(f){var k=e.treeGrid.collapse(f);e.brokenAxis&&(e.brokenAxis.setBreaks(k,!1),e.treeGrid.collapsedNodes&&(e.treeGrid.collapsedNodes=e.treeGrid.collapsedNodes.filter(function(k){return f.collapseStart!==
k.collapseStart||f.collapseEnd!==k.collapseEnd})))})}),x(e,"afterBreaks",function(){"yAxis"===e.coll&&!e.staticScale&&e.chart.options.chart.height&&(e.isDirty=!0)}),n=H({grid:{enabled:!0},labels:{align:"left",levels:[{level:void 0},{level:1,style:{fontWeight:"bold"}}],symbol:{type:"triangle",x:-5,y:-5,height:10,width:10,padding:5}},uniqueNames:!1},n,{reversed:!0,grid:{columns:void 0}}));a.apply(e,[c,n]);q&&(e.hasNames=!0,e.options.showLastLabel=!0)}function B(a){var c=this.options;"treegrid"===c.type?
(this.min=e(this.userMin,c.min,this.dataMin),this.max=e(this.userMax,c.max,this.dataMax),p(this,"foundExtremes"),this.setAxisTranslation(),this.tickmarkOffset=.5,this.tickInterval=1,this.tickPositions=this.treeGrid.mapOfPosToGridNode?this.treeGrid.getTickPositions():[]):a.apply(this,Array.prototype.slice.call(arguments,1))}var E;c.compose=function(c,e,n,d){-1===c.keepProps.indexOf("treeGrid")&&(c.keepProps.push("treeGrid"),E=d,a(c.prototype,"generateTick",C),a(c.prototype,"init",Q),a(c.prototype,
"setTickInterval",B),c.prototype.utils={getNode:v.getNode},t.compose(c,e,d),b.compose(c,n),u.compose(d));return c};var K=function(){function a(a){this.axis=a}a.prototype.setCollapsedStatus=function(a){var c=this.axis,e=c.chart;c.series.forEach(function(c){var f=c.options.data;if(a.id&&f){var k=e.get(a.id);c=f[c.data.indexOf(k)];k&&c&&(k.collapsed=a.collapsed,c.collapsed=a.collapsed)}})};a.prototype.collapse=function(a){var c=this.axis,e=c.options.breaks||[],q=r(a,c.max);e.push(q);a.collapsed=!0;c.treeGrid.setCollapsedStatus(a);
return e};a.prototype.expand=function(a){var c=this.axis,e=c.options.breaks||[],q=r(a,c.max);a.collapsed=!1;c.treeGrid.setCollapsedStatus(a);return e.reduce(function(f,a){a.to===q.to&&a.from===q.from||f.push(a);return f},[])};a.prototype.getTickPositions=function(){var a=this.axis,c=Math.floor(a.min/a.tickInterval)*a.tickInterval,e=Math.ceil(a.max/a.tickInterval)*a.tickInterval;return Object.keys(a.treeGrid.mapOfPosToGridNode||{}).reduce(function(n,f){f=+f;!(f>=c&&f<=e)||a.brokenAxis&&a.brokenAxis.isInAnyBreak(f)||
n.push(f);return n},[])};a.prototype.isCollapsed=function(a){var c=this.axis,e=c.options.breaks||[],q=r(a,c.max);return e.some(function(f){return f.from===q.from&&f.to===q.to})};a.prototype.toggleCollapse=function(a){return this.isCollapsed(a)?this.expand(a):this.collapse(a)};return a}();c.Additions=K})(c||(c={}));return c});J(b,"Extensions/CurrentDateIndication.js",[b["Core/Axis/Axis.js"],b["Core/Axis/PlotLineOrBand/PlotLineOrBand.js"],b["Core/Utilities.js"]],function(b,t,v){var u=v.addEvent,B=v.merge;
v=v.wrap;var l={color:"#ccd6eb",width:2,label:{format:"%a, %b %d %Y, %H:%M",formatter:function(b,l){return this.axis.chart.time.dateFormat(l||"",b)},rotation:0,style:{fontSize:"10px"}}};u(b,"afterSetOptions",function(){var b=this.options,x=b.currentDateIndicator;x&&(x="object"===typeof x?B(l,x):B(l),x.value=Date.now(),x.className="highcharts-current-date-indicator",b.plotLines||(b.plotLines=[]),b.plotLines.push(x))});u(t,"render",function(){this.label&&this.label.attr({text:this.getLabelText(this.options.label)})});
v(t.prototype,"getLabelText",function(b,l){var d=this.options;return d&&d.className&&-1!==d.className.indexOf("highcharts-current-date-indicator")&&d.label&&"function"===typeof d.label.formatter?(d.value=Date.now(),d.label.formatter.call(this,d.value,d.label.format)):b.call(this,l)})});J(b,"Extensions/StaticScale.js",[b["Core/Axis/Axis.js"],b["Core/Chart/Chart.js"],b["Core/Utilities.js"]],function(b,t,v){var u=v.addEvent,B=v.defined,l=v.isNumber,h=v.pick;u(b,"afterSetOptions",function(){var b=this.chart.options.chart;
!this.horiz&&l(this.options.staticScale)&&(!b.height||b.scrollablePlotArea&&b.scrollablePlotArea.minHeight)&&(this.staticScale=this.options.staticScale)});t.prototype.adjustHeight=function(){"adjustHeight"!==this.redrawTrigger&&((this.axes||[]).forEach(function(b){var d=b.chart,l=!!d.initiatedScale&&d.options.animation,g=b.options.staticScale;if(b.staticScale&&B(b.min)){var A=h(b.brokenAxis&&b.brokenAxis.unitLength,b.max+b.tickInterval-b.min)*g;A=Math.max(A,g);g=A-d.plotHeight;!d.scrollablePixelsY&&
1<=Math.abs(g)&&(d.plotHeight=A,d.redrawTrigger="adjustHeight",d.setSize(void 0,d.chartHeight+g,l));b.series.forEach(function(g){(g=g.sharedClipKey&&d.sharedClips[g.sharedClipKey])&&g.attr(d.inverted?{width:d.plotHeight}:{height:d.plotHeight})})}}),this.initiatedScale=!0);this.redrawTrigger=null};u(t,"render",t.prototype.adjustHeight)});J(b,"Extensions/ArrowSymbols.js",[b["Core/Renderer/SVG/SVGRenderer.js"]],function(b){function t(b,l,h,t){return[["M",b,l+t/2],["L",b+h,l],["L",b,l+t/2],["L",b+h,l+
t]]}function v(b,l,h,t){return[["M",b+h,l],["L",b,l+t/2],["L",b+h,l+t],["Z"]]}function u(b,l,h,t){return v(b,l,h/2,t)}b=b.prototype.symbols;b.arrow=t;b["arrow-filled"]=v;b["arrow-filled-half"]=u;b["arrow-half"]=function(b,l,h,u){return t(b,l,h/2,u)};b["triangle-left"]=v;b["triangle-left-half"]=u;return b});J(b,"Gantt/Connection.js",[b["Core/Globals.js"],b["Core/DefaultOptions.js"],b["Core/Series/Point.js"],b["Core/Utilities.js"]],function(b,t,v,u){function B(d){var e=d.shapeArgs;return e?{xMin:e.x||
0,xMax:(e.x||0)+(e.width||0),yMin:e.y||0,yMax:(e.y||0)+(e.height||0)}:(e=d.graphic&&d.graphic.getBBox())?{xMin:d.plotX-e.width/2,xMax:d.plotX+e.width/2,yMin:d.plotY-e.height/2,yMax:d.plotY+e.height/2}:null}"";var l=u.defined,h=u.error,x=u.extend,d=u.merge,p=u.objectEach,g=b.deg2rad,A=Math.max,D=Math.min;x(t.defaultOptions,{connectors:{type:"straight",lineWidth:1,marker:{enabled:!1,align:"center",verticalAlign:"middle",inside:!1,lineWidth:1},startMarker:{symbol:"diamond"},endMarker:{symbol:"arrow-filled"}}});
t=function(){function b(e,a,c){this.toPoint=this.pathfinder=this.graphics=this.fromPoint=this.chart=void 0;this.init(e,a,c)}b.prototype.init=function(e,a,c){this.fromPoint=e;this.toPoint=a;this.options=c;this.chart=e.series.chart;this.pathfinder=this.chart.pathfinder};b.prototype.renderPath=function(e,a,c){var d=this.chart,b=d.styledMode,g=d.pathfinder,m=!d.options.chart.forExport&&!1!==c,h=this.graphics&&this.graphics.path;g.group||(g.group=d.renderer.g().addClass("highcharts-pathfinder-group").attr({zIndex:-1}).add(d.seriesGroup));
g.group.translate(d.plotLeft,d.plotTop);h&&h.renderer||(h=d.renderer.path().add(g.group),b||h.attr({opacity:0}));h.attr(a);e={d:e};b||(e.opacity=1);h[m?"animate":"attr"](e,c);this.graphics=this.graphics||{};this.graphics.path=h};b.prototype.addMarker=function(e,a,c){var d=this.fromPoint.series.chart,b=d.pathfinder;d=d.renderer;var h="start"===e?this.fromPoint:this.toPoint,m=h.getPathfinderAnchorPoint(a);if(a.enabled&&((c="start"===e?c[1]:c[c.length-2])&&"M"===c[0]||"L"===c[0])){c={x:c[1],y:c[2]};
c=h.getRadiansToVector(c,m);m=h.getMarkerVector(c,a.radius,m);c=-c/g;if(a.width&&a.height){var l=a.width;var A=a.height}else l=A=2*a.radius;this.graphics=this.graphics||{};m={x:m.x-l/2,y:m.y-A/2,width:l,height:A,rotation:c,rotationOriginX:m.x,rotationOriginY:m.y};this.graphics[e]?this.graphics[e].animate(m):(this.graphics[e]=d.symbol(a.symbol).addClass("highcharts-point-connecting-path-"+e+"-marker").attr(m).add(b.group),d.styledMode||this.graphics[e].attr({fill:a.color||this.fromPoint.color,stroke:a.lineColor,
"stroke-width":a.lineWidth,opacity:0}).animate({opacity:1},h.series.options.animation))}};b.prototype.getPath=function(e){var a=this.pathfinder,c=this.chart,b=a.algorithms[e.type],g=a.chartObstacles;if("function"!==typeof b)return h('"'+e.type+'" is not a Pathfinder algorithm.'),{path:[],obstacles:[]};b.requiresObstacles&&!g&&(g=a.chartObstacles=a.getChartObstacles(e),c.options.connectors.algorithmMargin=e.algorithmMargin,a.chartObstacleMetrics=a.getObstacleMetrics(g));return b(this.fromPoint.getPathfinderAnchorPoint(e.startMarker),
this.toPoint.getPathfinderAnchorPoint(e.endMarker),d({chartObstacles:g,lineObstacles:a.lineObstacles||[],obstacleMetrics:a.chartObstacleMetrics,hardBounds:{xMin:0,xMax:c.plotWidth,yMin:0,yMax:c.plotHeight},obstacleOptions:{margin:e.algorithmMargin},startDirectionX:a.getAlgorithmStartDirection(e.startMarker)},e))};b.prototype.render=function(){var e=this.fromPoint,a=e.series,c=a.chart,b=c.pathfinder,g=d(c.options.connectors,a.options.connectors,e.options.connectors,this.options),h={};c.styledMode||
(h.stroke=g.lineColor||e.color,h["stroke-width"]=g.lineWidth,g.dashStyle&&(h.dashstyle=g.dashStyle));h["class"]="highcharts-point-connecting-path highcharts-color-"+e.colorIndex;g=d(h,g);l(g.marker.radius)||(g.marker.radius=D(A(Math.ceil((g.algorithmMargin||8)/2)-1,1),5));e=this.getPath(g);c=e.path;e.obstacles&&(b.lineObstacles=b.lineObstacles||[],b.lineObstacles=b.lineObstacles.concat(e.obstacles));this.renderPath(c,h,a.options.animation);this.addMarker("start",d(g.marker,g.startMarker),c);this.addMarker("end",
d(g.marker,g.endMarker),c)};b.prototype.destroy=function(){this.graphics&&(p(this.graphics,function(e){e.destroy()}),delete this.graphics)};return b}();b.Connection=t;x(v.prototype,{getPathfinderAnchorPoint:function(d){var e=B(this);switch(d.align){case "right":var a="xMax";break;case "left":a="xMin"}switch(d.verticalAlign){case "top":var c="yMin";break;case "bottom":c="yMax"}return{x:a?e[a]:(e.xMin+e.xMax)/2,y:c?e[c]:(e.yMin+e.yMax)/2}},getRadiansToVector:function(d,e){var a;l(e)||(a=B(this))&&(e=
{x:(a.xMin+a.xMax)/2,y:(a.yMin+a.yMax)/2});return Math.atan2(e.y-d.y,d.x-e.x)},getMarkerVector:function(d,e,a){var c=2*Math.PI,b=B(this),g=b.xMax-b.xMin,h=b.yMax-b.yMin,m=Math.atan2(h,g),l=!1;g/=2;var A=h/2,p=b.xMin+g;b=b.yMin+A;for(var D=p,t=b,u=1,z=1;d<-Math.PI;)d+=c;for(;d>Math.PI;)d-=c;c=Math.tan(d);d>-m&&d<=m?(z=-1,l=!0):d>m&&d<=Math.PI-m?z=-1:d>Math.PI-m||d<=-(Math.PI-m)?(u=-1,l=!0):u=-1;l?(D+=u*g,t+=z*g*c):(D+=h/(2*c)*u,t+=z*A);a.x!==p&&(D=a.x);a.y!==b&&(t=a.y);return{x:D+e*Math.cos(d),y:t-
e*Math.sin(d)}}});return t});J(b,"Gantt/PathfinderAlgorithms.js",[b["Core/Utilities.js"]],function(b){function t(d,b,h){h=h||0;var g=d.length-1;b-=1e-7;for(var e,a;h<=g;)if(e=g+h>>1,a=b-d[e].xMin,0<a)h=e+1;else if(0>a)g=e-1;else return e;return 0<h?h-1:0}function v(d,b){for(var g=t(d,b.x+1)+1;g--;){var h;if(h=d[g].xMax>=b.x)h=d[g],h=b.x<=h.xMax&&b.x>=h.xMin&&b.y<=h.yMax&&b.y>=h.yMin;if(h)return g}return-1}function u(d){var b=[];if(d.length){b.push(["M",d[0].start.x,d[0].start.y]);for(var g=0;g<d.length;++g)b.push(["L",
d[g].end.x,d[g].end.y])}return b}function B(d,b){d.yMin=x(d.yMin,b.yMin);d.yMax=h(d.yMax,b.yMax);d.xMin=x(d.xMin,b.xMin);d.xMax=h(d.xMax,b.xMax)}var l=b.pick,h=Math.min,x=Math.max,d=Math.abs;b=function(b,h,p){function g(a,c,e,d,b){a={x:a.x,y:a.y};a[c]=e[d||c]+(b||0);return a}function e(a,c,e){var b=d(c[e]-a[e+"Min"])>d(c[e]-a[e+"Max"]);return g(c,e,a,e+(b?"Max":"Min"),b?1:-1)}var a=[],c=l(p.startDirectionX,d(h.x-b.x)>d(h.y-b.y))?"x":"y",r=p.chartObstacles,A=v(r,b);p=v(r,h);if(-1<p){var t=r[p];p=e(t,
h,c);t={start:p,end:h};var m=p}else m=h;-1<A&&(r=r[A],p=e(r,b,c),a.push({start:b,end:p}),p[c]>=b[c]===p[c]>=m[c]&&(c="y"===c?"x":"y",h=b[c]<h[c],a.push({start:p,end:g(p,c,r,c+(h?"Max":"Min"),h?1:-1)}),c="y"===c?"x":"y"));b=a.length?a[a.length-1].end:b;p=g(b,c,m);a.push({start:b,end:p});c=g(p,"y"===c?"x":"y",m);a.push({start:p,end:c});a.push(t);return{path:u(a),obstacles:a}};b.requiresObstacles=!0;var p=function(b,p,D){function g(a,c,f){var k,b=a.x<c.x?1:-1;if(a.x<c.x){var e=a;var d=c}else e=c,d=a;
if(a.y<c.y){var n=a;var q=c}else n=c,q=a;for(k=0>b?h(t(I,d.x),I.length-1):0;I[k]&&(0<b&&I[k].xMin<=d.x||0>b&&I[k].xMax>=e.x);){if(I[k].xMin<=d.x&&I[k].xMax>=e.x&&I[k].yMin<=q.y&&I[k].yMax>=n.y)return f?{y:a.y,x:a.x<c.x?I[k].xMin-1:I[k].xMax+1,obstacle:I[k]}:{x:a.x,y:a.y<c.y?I[k].yMin-1:I[k].yMax+1,obstacle:I[k]};k+=b}return c}function e(a,c,f,k,b){var e=b.soft,n=b.hard,q=k?"x":"y",w={x:c.x,y:c.y},h={x:c.x,y:c.y};b=a[q+"Max"]>=e[q+"Max"];e=a[q+"Min"]<=e[q+"Min"];var m=a[q+"Max"]>=n[q+"Max"];n=a[q+
"Min"]<=n[q+"Min"];var y=d(a[q+"Min"]-c[q]),r=d(a[q+"Max"]-c[q]);f=10>d(y-r)?c[q]<f[q]:r<y;h[q]=a[q+"Min"];w[q]=a[q+"Max"];a=g(c,h,k)[q]!==h[q];c=g(c,w,k)[q]!==w[q];f=a?c?f:!0:c?!1:f;f=e?b?f:!0:b?!1:f;return n?m?f:!0:m?!1:f}function a(c,b,f){if(c.x===b.x&&c.y===b.y)return[];var k=f?"x":"y",d=D.obstacleOptions.margin;var n={soft:{xMin:Q,xMax:M,yMin:E,yMax:K},hard:D.hardBounds};var q=v(I,c);if(-1<q){q=I[q];n=e(q,c,b,f,n);B(q,D.hardBounds);var r=f?{y:c.y,x:q[n?"xMax":"xMin"]+(n?1:-1)}:{x:c.x,y:q[n?"yMax":
"yMin"]+(n?1:-1)};var y=v(I,r);-1<y&&(y=I[y],B(y,D.hardBounds),r[k]=n?x(q[k+"Max"]-d+1,(y[k+"Min"]+q[k+"Max"])/2):h(q[k+"Min"]+d-1,(y[k+"Max"]+q[k+"Min"])/2),c.x===r.x&&c.y===r.y?(m&&(r[k]=n?x(q[k+"Max"],y[k+"Max"])+1:h(q[k+"Min"],y[k+"Min"])-1),m=!m):m=!1);c=[{start:c,end:r}]}else k=g(c,{x:f?b.x:c.x,y:f?c.y:b.y},f),c=[{start:c,end:{x:k.x,y:k.y}}],k[f?"x":"y"]!==b[f?"x":"y"]&&(n=e(k.obstacle,k,b,!f,n),B(k.obstacle,D.hardBounds),n={x:f?k.x:k.obstacle[n?"xMax":"xMin"]+(n?1:-1),y:f?k.obstacle[n?"yMax":
"yMin"]+(n?1:-1):k.y},f=!f,c=c.concat(a({x:k.x,y:k.y},n,f)));return c=c.concat(a(c[c.length-1].end,b,!f))}function c(a,c,f){var k=h(a.xMax-c.x,c.x-a.xMin)<h(a.yMax-c.y,c.y-a.yMin);f=e(a,c,f,k,{soft:D.hardBounds,hard:D.hardBounds});return k?{y:c.y,x:a[f?"xMax":"xMin"]+(f?1:-1)}:{x:c.x,y:a[f?"yMax":"yMin"]+(f?1:-1)}}var r=l(D.startDirectionX,d(p.x-b.x)>d(p.y-b.y)),A=r?"x":"y",G=[],m=!1,C=D.obstacleMetrics,Q=h(b.x,p.x)-C.maxWidth-10,M=x(b.x,p.x)+C.maxWidth+10,E=h(b.y,p.y)-C.maxHeight-10,K=x(b.y,p.y)+
C.maxHeight+10,I=D.chartObstacles;var z=t(I,Q);C=t(I,M);I=I.slice(z,C+1);if(-1<(C=v(I,p))){var n=c(I[C],p,b);G.push({end:p,start:n});p=n}for(;-1<(C=v(I,p));)z=0>p[A]-b[A],n={x:p.x,y:p.y},n[A]=I[C][z?A+"Max":A+"Min"]+(z?1:-1),G.push({end:p,start:n}),p=n;b=a(b,p,r);b=b.concat(G.reverse());return{path:u(b),obstacles:b}};p.requiresObstacles=!0;return{fastAvoid:p,straight:function(b,d){return{path:[["M",b.x,b.y],["L",d.x,d.y]],obstacles:[{start:b,end:d}]}},simpleConnect:b}});J(b,"Gantt/Pathfinder.js",
[b["Gantt/Connection.js"],b["Core/Chart/Chart.js"],b["Core/Globals.js"],b["Core/DefaultOptions.js"],b["Core/Series/Point.js"],b["Core/Utilities.js"],b["Gantt/PathfinderAlgorithms.js"]],function(b,t,v,u,B,l,h){function x(a){var c=a.shapeArgs;return c?{xMin:c.x||0,xMax:(c.x||0)+(c.width||0),yMin:c.y||0,yMax:(c.y||0)+(c.height||0)}:(c=a.graphic&&a.graphic.getBBox())?{xMin:a.plotX-c.width/2,xMax:a.plotX+c.width/2,yMin:a.plotY-c.height/2,yMax:a.plotY+c.height/2}:null}function d(c){for(var b=c.length,e=
0,d,g,h=[],m=function(c,b,e){e=a(e,10);var d=c.yMax+e>b.yMin-e&&c.yMin-e<b.yMax+e,f=c.xMax+e>b.xMin-e&&c.xMin-e<b.xMax+e,k=d?c.xMin>b.xMax?c.xMin-b.xMax:b.xMin-c.xMax:Infinity,n=f?c.yMin>b.yMax?c.yMin-b.yMax:b.yMin-c.yMax:Infinity;return f&&d?e?m(c,b,Math.floor(e/2)):Infinity:F(k,n)};e<b;++e)for(d=e+1;d<b;++d)g=m(c[e],c[d]),80>g&&h.push(g);h.push(80);return r(Math.floor(h.sort(function(a,c){return a-c})[Math.floor(h.length/10)]/2-1),1)}function p(a){if(a.options.pathfinder||a.series.reduce(function(a,
c){c.options&&e(!0,c.options.connectors=c.options.connectors||{},c.options.pathfinder);return a||c.options&&c.options.pathfinder},!1))e(!0,a.options.connectors=a.options.connectors||{},a.options.pathfinder),D('WARNING: Pathfinder options have been renamed. Use "chart.connectors" or "series.connectors" instead.')}"";var g=l.addEvent,A=l.defined,D=l.error,H=l.extend,e=l.merge,a=l.pick,c=l.splat,r=Math.max,F=Math.min;H(u.defaultOptions,{connectors:{type:"straight",lineWidth:1,marker:{enabled:!1,align:"center",
verticalAlign:"middle",inside:!1,lineWidth:1},startMarker:{symbol:"diamond"},endMarker:{symbol:"arrow-filled"}}});var G=function(){function e(a){this.lineObstacles=this.group=this.connections=this.chartObstacleMetrics=this.chartObstacles=this.chart=void 0;this.init(a)}e.prototype.init=function(a){this.chart=a;this.connections=[];g(a,"redraw",function(){this.pathfinder.update()})};e.prototype.update=function(a){var e=this.chart,d=this,g=d.connections;d.connections=[];e.series.forEach(function(a){a.visible&&
!a.options.isInternal&&a.points.forEach(function(f){var a=f.options;a&&a.dependency&&(a.connect=a.dependency);var n;a=f.options&&f.options.connect&&c(f.options.connect);f.visible&&!1!==f.isInside&&a&&a.forEach(function(a){n=e.get("string"===typeof a?a:a.to);n instanceof B&&n.series.visible&&n.visible&&!1!==n.isInside&&d.connections.push(new b(f,n,"string"===typeof a?{}:a))})})});for(var h=0,r=void 0,m=void 0,n=g.length,y=d.connections.length;h<n;++h){m=!1;for(r=0;r<y;++r)if(g[h].fromPoint===d.connections[r].fromPoint&&
g[h].toPoint===d.connections[r].toPoint){d.connections[r].graphics=g[h].graphics;m=!0;break}m||g[h].destroy()}delete this.chartObstacles;delete this.lineObstacles;d.renderConnections(a)};e.prototype.renderConnections=function(a){a?this.chart.series.forEach(function(a){var c=function(){var c=a.chart.pathfinder;(c&&c.connections||[]).forEach(function(c){c.fromPoint&&c.fromPoint.series===a&&c.render()});a.pathfinderRemoveRenderEvent&&(a.pathfinderRemoveRenderEvent(),delete a.pathfinderRemoveRenderEvent)};
!1===a.options.animation?c():a.pathfinderRemoveRenderEvent=g(a,"afterAnimate",c)}):this.connections.forEach(function(a){a.render()})};e.prototype.getChartObstacles=function(c){for(var b=[],e=this.chart.series,g=a(c.algorithmMargin,0),h,r=0,m=e.length;r<m;++r)if(e[r].visible&&!e[r].options.isInternal){var n=0,y=e[r].points.length,q=void 0;for(q=void 0;n<y;++n)q=e[r].points[n],q.visible&&(q=x(q))&&b.push({xMin:q.xMin-g,xMax:q.xMax+g,yMin:q.yMin-g,yMax:q.yMax+g})}b=b.sort(function(f,a){return f.xMin-
a.xMin});A(c.algorithmMargin)||(h=c.algorithmMargin=d(b),b.forEach(function(f){f.xMin-=h;f.xMax+=h;f.yMin-=h;f.yMax+=h}));return b};e.prototype.getObstacleMetrics=function(a){for(var c=0,b=0,e,d,g=a.length;g--;)e=a[g].xMax-a[g].xMin,d=a[g].yMax-a[g].yMin,c<e&&(c=e),b<d&&(b=d);return{maxHeight:b,maxWidth:c}};e.prototype.getAlgorithmStartDirection=function(a){var c="top"!==a.verticalAlign&&"bottom"!==a.verticalAlign;return"left"!==a.align&&"right"!==a.align?c?void 0:!1:c?!0:void 0};return e}();G.prototype.algorithms=
h;v.Pathfinder=G;H(B.prototype,{getPathfinderAnchorPoint:function(a){var c=x(this);switch(a.align){case "right":var b="xMax";break;case "left":b="xMin"}switch(a.verticalAlign){case "top":var e="yMin";break;case "bottom":e="yMax"}return{x:b?c[b]:(c.xMin+c.xMax)/2,y:e?c[e]:(c.yMin+c.yMax)/2}},getRadiansToVector:function(a,c){var b;A(c)||(b=x(this))&&(c={x:(b.xMin+b.xMax)/2,y:(b.yMin+b.yMax)/2});return Math.atan2(c.y-a.y,a.x-c.x)},getMarkerVector:function(a,c,b){var e=2*Math.PI,d=x(this),g=d.xMax-d.xMin,
h=d.yMax-d.yMin,r=Math.atan2(h,g),n=!1;g/=2;var m=h/2,q=d.xMin+g;d=d.yMin+m;for(var f=q,k=d,w=1,L=1;a<-Math.PI;)a+=e;for(;a>Math.PI;)a-=e;e=Math.tan(a);a>-r&&a<=r?(L=-1,n=!0):a>r&&a<=Math.PI-r?L=-1:a>Math.PI-r||a<=-(Math.PI-r)?(w=-1,n=!0):w=-1;n?(f+=w*g,k+=L*g*e):(f+=h/(2*e)*w,k+=L*m);b.x!==q&&(f=b.x);b.y!==d&&(k=b.y);return{x:f+c*Math.cos(a),y:k-c*Math.sin(a)}}});t.prototype.callbacks.push(function(a){!1!==a.options.connectors.enabled&&(p(a),this.pathfinder=new G(this),this.pathfinder.update(!0))});
return G});J(b,"Series/Gantt/GanttSeries.js",[b["Core/Axis/Axis.js"],b["Core/Chart/Chart.js"],b["Series/Gantt/GanttPoint.js"],b["Core/Series/SeriesRegistry.js"],b["Core/Axis/Tick.js"],b["Core/Utilities.js"],b["Core/Axis/TreeGridAxis.js"]],function(b,t,v,u,B,l,h){var x=this&&this.__extends||function(){var b=function(e,a){b=Object.setPrototypeOf||{__proto__:[]}instanceof Array&&function(a,b){a.__proto__=b}||function(a,b){for(var c in b)b.hasOwnProperty(c)&&(a[c]=b[c])};return b(e,a)};return function(e,
a){function c(){this.constructor=e}b(e,a);e.prototype=null===a?Object.create(a):(c.prototype=a.prototype,new c)}}(),d=u.series,p=u.seriesTypes.xrange,g=l.extend,A=l.isNumber,D=l.merge;h.compose(b,t,d,B);b=function(b){function e(){var a=null!==b&&b.apply(this,arguments)||this;a.data=void 0;a.options=void 0;a.points=void 0;return a}x(e,b);e.prototype.drawPoint=function(a,c){var b=this.options,e=this.chart.renderer,d=a.shapeArgs,g=a.plotY,h=a.graphic,l=a.selected&&"select",t=b.stacking&&!b.borderRadius;
if(a.options.milestone)if(A(g)&&null!==a.y&&!1!==a.visible){d=e.symbols.diamond(d.x||0,d.y||0,d.width||0,d.height||0);if(h)h[c]({d:d});else a.graphic=e.path(d).addClass(a.getClassName(),!0).add(a.group||this.group);this.chart.styledMode||a.graphic.attr(this.pointAttribs(a,l)).shadow(b.shadow,null,t)}else h&&(a.graphic=h.destroy());else p.prototype.drawPoint.call(this,a,c)};e.prototype.translatePoint=function(a){p.prototype.translatePoint.call(this,a);if(a.options.milestone){var c=a.shapeArgs;var b=
c.height||0;a.shapeArgs={x:(c.x||0)-b/2,y:c.y,width:b,height:b}}};e.defaultOptions=D(p.defaultOptions,{grouping:!1,dataLabels:{enabled:!0},tooltip:{headerFormat:'<span style="font-size: 10px">{series.name}</span><br/>',pointFormat:null,pointFormatter:function(){var a=this.series,c=a.xAxis,b=a.tooltipOptions.dateTimeLabelFormats,e=c.options.startOfWeek,d=a.tooltipOptions,g=d.xDateFormat,h=this.options.milestone,l="<b>"+(this.name||this.yCategory)+"</b>";if(d.pointFormat)return this.tooltipFormatter(d.pointFormat);
!g&&A(this.start)&&(g=a.chart.time.getDateFormat(c.closestPointRange,this.start,e,b||{}));c=a.chart.time.dateFormat(g,this.start);a=a.chart.time.dateFormat(g,this.end);l+="<br/>";return h?l+(c+"<br/>"):l+("Start: "+c+"<br/>End: ")+(a+"<br/>")}},connectors:{type:"simpleConnect",animation:{reversed:!0},startMarker:{enabled:!0,symbol:"arrow-filled",radius:4,fill:"#fa0",align:"left"},endMarker:{enabled:!1,align:"right"}}});return e}(p);g(b.prototype,{pointArrayMap:["start","end","y"],pointClass:v,setData:d.prototype.setData});
u.registerSeriesType("gantt",b);"";return b});J(b,"Core/Chart/GanttChart.js",[b["Core/Chart/Chart.js"],b["Core/DefaultOptions.js"],b["Core/Utilities.js"]],function(b,t,v){var u=this&&this.__extends||function(){var b=function(d,g){b=Object.setPrototypeOf||{__proto__:[]}instanceof Array&&function(b,d){b.__proto__=d}||function(b,d){for(var g in d)d.hasOwnProperty(g)&&(b[g]=d[g])};return b(d,g)};return function(d,g){function h(){this.constructor=d}b(d,g);d.prototype=null===g?Object.create(g):(h.prototype=
g.prototype,new h)}}(),B=t.getOptions,l=v.isArray,h=v.merge,x=v.splat;b=function(b){function d(){return null!==b&&b.apply(this,arguments)||this}u(d,b);d.prototype.init=function(d,p){var g=B(),t=d.xAxis,e=d.yAxis,a;d.xAxis=d.yAxis=void 0;var c=h(!0,{chart:{type:"gantt"},title:{text:null},legend:{enabled:!1},navigator:{series:{type:"gantt"},yAxis:{type:"category"}}},d,{isGantt:!0});d.xAxis=t;d.yAxis=e;c.xAxis=(l(d.xAxis)?d.xAxis:[d.xAxis||{},{}]).map(function(c,b){1===b&&(a=0);return h(g.xAxis,{grid:{enabled:!0},
opposite:!0,linkedTo:a},c,{type:"datetime"})});c.yAxis=x(d.yAxis||{}).map(function(a){return h(g.yAxis,{grid:{enabled:!0},staticScale:50,reversed:!0,type:a.categories?a.type:"treegrid"},a)});b.prototype.init.call(this,c,p)};return d}(b);(function(b){b.ganttChart=function(d,g,h){return new b(d,g,h)}})(b||(b={}));return b});J(b,"Core/Axis/ScrollbarAxis.js",[b["Core/Utilities.js"]],function(b){var t=b.addEvent,v=b.defined,u=b.pick;return function(){function b(){}b.compose=function(l,h){if(-1===b.composed.indexOf(l))b.composed.push(l);
else return l;var x=function(b){var d=u(b.options&&b.options.min,b.min),g=u(b.options&&b.options.max,b.max);return{axisMin:d,axisMax:g,scrollMin:v(b.dataMin)?Math.min(d,b.min,b.dataMin,u(b.threshold,Infinity)):d,scrollMax:v(b.dataMax)?Math.max(g,b.max,b.dataMax,u(b.threshold,-Infinity)):g}};t(l,"afterInit",function(){var b=this;b.options&&b.options.scrollbar&&b.options.scrollbar.enabled&&(b.options.scrollbar.vertical=!b.horiz,b.options.startOnTick=b.options.endOnTick=!1,b.scrollbar=new h(b.chart.renderer,
b.options.scrollbar,b.chart),t(b.scrollbar,"changed",function(d){var g=x(b),h=g.axisMax,l=g.scrollMin,p=g.scrollMax-l;v(g.axisMin)&&v(h)&&(b.horiz&&!b.reversed||!b.horiz&&b.reversed?(g=l+p*this.to,l+=p*this.from):(g=l+p*(1-this.from),l+=p*(1-this.to)),this.shouldUpdateExtremes(d.DOMType)?b.setExtremes(l,g,!0,"mousemove"!==d.DOMType&&"touchmove"!==d.DOMType,d):this.setRange(this.from,this.to))}))});t(l,"afterRender",function(){var b=x(this),h=b.scrollMin,g=b.scrollMax;b=this.scrollbar;var l=this.axisTitleMargin+
(this.titleOffset||0),t=this.chart.scrollbarsOffsets,u=this.options.margin||0;b&&(this.horiz?(this.opposite||(t[1]+=l),b.position(this.left,this.top+this.height+2+t[1]-(this.opposite?u:0),this.width,this.height),this.opposite||(t[1]+=u),l=1):(this.opposite&&(t[0]+=l),b.position(b.options.opposite?this.left+this.width+2+t[0]-(this.opposite?0:u):this.opposite?0:u,this.top,this.width,this.height),this.opposite&&(t[0]+=u),l=0),t[l]+=b.size+b.options.margin,isNaN(h)||isNaN(g)||!v(this.min)||!v(this.max)||
this.min===this.max?b.setRange(0,1):(t=(this.min-h)/(g-h),h=(this.max-h)/(g-h),this.horiz&&!this.reversed||!this.horiz&&this.reversed?b.setRange(t,h):b.setRange(1-h,1-t)))});t(l,"afterGetOffset",function(){var b=this.scrollbar&&!this.scrollbar.options.opposite;b=this.horiz?2:b?3:1;var h=this.scrollbar;h&&(this.chart.scrollbarsOffsets=[0,0],this.chart.axisOffset[b]+=h.size+h.options.margin)});return l};b.composed=[];return b}()});J(b,"Core/ScrollbarDefaults.js",[b["Core/Globals.js"]],function(b){return{height:b.isTouchDevice?
20:14,barBorderRadius:0,buttonBorderRadius:0,liveRedraw:void 0,margin:10,minWidth:6,opposite:!0,step:.2,zIndex:3,barBackgroundColor:"#cccccc",barBorderWidth:1,barBorderColor:"#cccccc",buttonArrowColor:"#333333",buttonBackgroundColor:"#e6e6e6",buttonBorderColor:"#cccccc",buttonBorderWidth:1,rifleColor:"#333333",trackBackgroundColor:"#f2f2f2",trackBorderColor:"#f2f2f2",trackBorderWidth:1}});J(b,"Core/Scrollbar.js",[b["Core/DefaultOptions.js"],b["Core/Globals.js"],b["Core/Axis/ScrollbarAxis.js"],b["Core/ScrollbarDefaults.js"],
b["Core/Utilities.js"]],function(b,t,v,u,B){var l=b.defaultOptions,h=B.addEvent,x=B.correctFloat,d=B.defined,p=B.destroyObjectProperties,g=B.fireEvent,A=B.merge,D=B.pick,H=B.removeEvent;b=function(){function b(a,c,b){this._events=[];this.chart=void 0;this.from=this.chartY=this.chartX=0;this.scrollbar=this.renderer=this.options=this.group=void 0;this.scrollbarButtons=[];this.scrollbarGroup=void 0;this.scrollbarLeft=0;this.scrollbarRifles=void 0;this.scrollbarStrokeWidth=1;this.to=this.size=this.scrollbarTop=
0;this.track=void 0;this.trackBorderWidth=1;this.userOptions=void 0;this.y=this.x=0;this.init(a,c,b)}b.compose=function(a){v.compose(a,b)};b.swapXY=function(a,c){c&&a.forEach(function(a){for(var c=a.length,b,e=0;e<c;e+=2)b=a[e+1],"number"===typeof b&&(a[e+1]=a[e+2],a[e+2]=b)});return a};b.prototype.addEvents=function(){var a=this.options.inverted?[1,0]:[0,1],c=this.scrollbarButtons,b=this.scrollbarGroup.element,e=this.track.element,d=this.mouseDownHandler.bind(this),g=this.mouseMoveHandler.bind(this),
l=this.mouseUpHandler.bind(this);a=[[c[a[0]].element,"click",this.buttonToMinClick.bind(this)],[c[a[1]].element,"click",this.buttonToMaxClick.bind(this)],[e,"click",this.trackClick.bind(this)],[b,"mousedown",d],[b.ownerDocument,"mousemove",g],[b.ownerDocument,"mouseup",l]];t.hasTouch&&a.push([b,"touchstart",d],[b.ownerDocument,"touchmove",g],[b.ownerDocument,"touchend",l]);a.forEach(function(a){h.apply(null,a)});this._events=a};b.prototype.buttonToMaxClick=function(a){var c=(this.to-this.from)*D(this.options.step,
.2);this.updatePosition(this.from+c,this.to+c);g(this,"changed",{from:this.from,to:this.to,trigger:"scrollbar",DOMEvent:a})};b.prototype.buttonToMinClick=function(a){var c=x(this.to-this.from)*D(this.options.step,.2);this.updatePosition(x(this.from-c),x(this.to-c));g(this,"changed",{from:this.from,to:this.to,trigger:"scrollbar",DOMEvent:a})};b.prototype.cursorToScrollbarPosition=function(a){var c=this.options;c=c.minWidth>this.calculatedWidth?c.minWidth:0;return{chartX:(a.chartX-this.x-this.xOffset)/
(this.barWidth-c),chartY:(a.chartY-this.y-this.yOffset)/(this.barWidth-c)}};b.prototype.destroy=function(){var a=this,c=a.chart.scroller;a.removeEvents();["track","scrollbarRifles","scrollbar","scrollbarGroup","group"].forEach(function(c){a[c]&&a[c].destroy&&(a[c]=a[c].destroy())});c&&a===c.scrollbar&&(c.scrollbar=null,p(c.scrollbarButtons))};b.prototype.drawScrollbarButton=function(a){var c=this.renderer,e=this.scrollbarButtons,d=this.options,g=this.size,h=c.g().add(this.group);e.push(h);h=c.rect().addClass("highcharts-scrollbar-button").add(h);
this.chart.styledMode||h.attr({stroke:d.buttonBorderColor,"stroke-width":d.buttonBorderWidth,fill:d.buttonBackgroundColor});h.attr(h.crisp({x:-.5,y:-.5,width:g+1,height:g+1,r:d.buttonBorderRadius},h.strokeWidth()));h=c.path(b.swapXY([["M",g/2+(a?-1:1),g/2-3],["L",g/2+(a?-1:1),g/2+3],["L",g/2+(a?2:-2),g/2]],d.vertical)).addClass("highcharts-scrollbar-arrow").add(e[a]);this.chart.styledMode||h.attr({fill:d.buttonArrowColor})};b.prototype.init=function(a,c,b){this.scrollbarButtons=[];this.renderer=a;
this.userOptions=c;this.options=A(u,l.scrollbar,c);this.chart=b;this.size=D(this.options.size,this.options.height);c.enabled&&(this.render(),this.addEvents())};b.prototype.mouseDownHandler=function(a){a=this.chart.pointer.normalize(a);a=this.cursorToScrollbarPosition(a);this.chartX=a.chartX;this.chartY=a.chartY;this.initPositions=[this.from,this.to];this.grabbedCenter=!0};b.prototype.mouseMoveHandler=function(a){var c=this.chart.pointer.normalize(a),b=this.options.vertical?"chartY":"chartX",e=this.initPositions||
[];!this.grabbedCenter||a.touches&&0===a.touches[0][b]||(c=this.cursorToScrollbarPosition(c)[b],b=this[b],b=c-b,this.hasDragged=!0,this.updatePosition(e[0]+b,e[1]+b),this.hasDragged&&g(this,"changed",{from:this.from,to:this.to,trigger:"scrollbar",DOMType:a.type,DOMEvent:a}))};b.prototype.mouseUpHandler=function(a){this.hasDragged&&g(this,"changed",{from:this.from,to:this.to,trigger:"scrollbar",DOMType:a.type,DOMEvent:a});this.grabbedCenter=this.hasDragged=this.chartX=this.chartY=null};b.prototype.position=
function(a,c,b,e){var d=this.options.vertical,g=this.rendered?"animate":"attr",h=0;this.x=a;this.y=c+this.trackBorderWidth;this.width=b;this.xOffset=this.height=e;this.yOffset=h;d?(this.width=this.yOffset=b=h=this.size,this.xOffset=c=0,this.barWidth=e-2*b,this.x=a+=this.options.margin):(this.height=this.xOffset=e=c=this.size,this.barWidth=b-2*e,this.y+=this.options.margin);this.group[g]({translateX:a,translateY:this.y});this.track[g]({width:b,height:e});this.scrollbarButtons[1][g]({translateX:d?0:
b-c,translateY:d?e-h:0})};b.prototype.removeEvents=function(){this._events.forEach(function(a){H.apply(null,a)});this._events.length=0};b.prototype.render=function(){var a=this.renderer,c=this.options,e=this.size,d=this.chart.styledMode,g=a.g("scrollbar").attr({zIndex:c.zIndex,translateY:-99999}).add();this.group=g;this.track=a.rect().addClass("highcharts-scrollbar-track").attr({x:0,r:c.trackBorderRadius||0,height:e,width:e}).add(g);d||this.track.attr({fill:c.trackBackgroundColor,stroke:c.trackBorderColor,
"stroke-width":c.trackBorderWidth});this.trackBorderWidth=this.track.strokeWidth();this.track.attr({y:-this.trackBorderWidth%2/2});this.scrollbarGroup=a.g().add(g);this.scrollbar=a.rect().addClass("highcharts-scrollbar-thumb").attr({height:e,width:e,r:c.barBorderRadius||0}).add(this.scrollbarGroup);this.scrollbarRifles=a.path(b.swapXY([["M",-3,e/4],["L",-3,2*e/3],["M",0,e/4],["L",0,2*e/3],["M",3,e/4],["L",3,2*e/3]],c.vertical)).addClass("highcharts-scrollbar-rifles").add(this.scrollbarGroup);d||(this.scrollbar.attr({fill:c.barBackgroundColor,
stroke:c.barBorderColor,"stroke-width":c.barBorderWidth}),this.scrollbarRifles.attr({stroke:c.rifleColor,"stroke-width":1}));this.scrollbarStrokeWidth=this.scrollbar.strokeWidth();this.scrollbarGroup.translate(-this.scrollbarStrokeWidth%2/2,-this.scrollbarStrokeWidth%2/2);this.drawScrollbarButton(0);this.drawScrollbarButton(1)};b.prototype.setRange=function(a,c){var b=this.options,e=b.vertical,g=b.minWidth,h=this.barWidth,l=!this.rendered||this.hasDragged||this.chart.navigator&&this.chart.navigator.hasDragged?
"attr":"animate";if(d(h)){var p=h*Math.min(c,1);a=Math.max(a,0);var t=Math.ceil(h*a);this.calculatedWidth=p=x(p-t);p<g&&(t=(h-g+p)*a,p=g);g=Math.floor(t+this.xOffset+this.yOffset);h=p/2-.5;this.from=a;this.to=c;e?(this.scrollbarGroup[l]({translateY:g}),this.scrollbar[l]({height:p}),this.scrollbarRifles[l]({translateY:h}),this.scrollbarTop=g,this.scrollbarLeft=0):(this.scrollbarGroup[l]({translateX:g}),this.scrollbar[l]({width:p}),this.scrollbarRifles[l]({translateX:h}),this.scrollbarLeft=g,this.scrollbarTop=
0);12>=p?this.scrollbarRifles.hide():this.scrollbarRifles.show(!0);!1===b.showFull&&(0>=a&&1<=c?this.group.hide():this.group.show());this.rendered=!0}};b.prototype.shouldUpdateExtremes=function(a){return D(this.options.liveRedraw,t.svg&&!t.isTouchDevice&&!this.chart.isBoosting)||"mouseup"===a||"touchend"===a||!d(a)};b.prototype.trackClick=function(a){var c=this.chart.pointer.normalize(a),b=this.to-this.from,e=this.y+this.scrollbarTop,d=this.x+this.scrollbarLeft;this.options.vertical&&c.chartY>e||
!this.options.vertical&&c.chartX>d?this.updatePosition(this.from+b,this.to+b):this.updatePosition(this.from-b,this.to-b);g(this,"changed",{from:this.from,to:this.to,trigger:"scrollbar",DOMEvent:a})};b.prototype.update=function(a){this.destroy();this.init(this.chart.renderer,A(!0,this.options,a),this.chart)};b.prototype.updatePosition=function(a,c){1<c&&(a=x(1-x(c-a)),c=1);0>a&&(c=x(c-a),a=0);this.from=a;this.to=c};b.defaultOptions=u;return b}();l.scrollbar=A(!0,b.defaultOptions,l.scrollbar);return b});
J(b,"Extensions/RangeSelector.js",[b["Core/Axis/Axis.js"],b["Core/Chart/Chart.js"],b["Core/Globals.js"],b["Core/DefaultOptions.js"],b["Core/Renderer/SVG/SVGElement.js"],b["Core/Utilities.js"]],function(b,t,v,u,B,l){function h(a){if(-1!==a.indexOf("%L"))return"text";var c="aAdewbBmoyY".split("").some(function(c){return-1!==a.indexOf("%"+c)}),b="HkIlMS".split("").some(function(c){return-1!==a.indexOf("%"+c)});return c&&b?"datetime-local":c?"date":b?"time":"text"}var x=u.defaultOptions,d=l.addEvent,
p=l.createElement,g=l.css,A=l.defined,D=l.destroyObjectProperties,H=l.discardElement,e=l.extend,a=l.find,c=l.fireEvent,r=l.isNumber,F=l.merge,G=l.objectEach,m=l.pad,C=l.pick,J=l.pInt,M=l.splat;e(x,{rangeSelector:{allButtonsEnabled:!1,buttons:void 0,buttonSpacing:5,dropdown:"responsive",enabled:void 0,verticalAlign:"top",buttonTheme:{width:28,height:18,padding:2,zIndex:7},floating:!1,x:0,y:0,height:void 0,inputBoxBorderColor:"none",inputBoxHeight:17,inputBoxWidth:void 0,inputDateFormat:"%b %e, %Y",
inputDateParser:void 0,inputEditDateFormat:"%Y-%m-%d",inputEnabled:!0,inputPosition:{align:"right",x:0,y:0},inputSpacing:5,selected:void 0,buttonPosition:{align:"left",x:0,y:0},inputStyle:{color:"#335cad",cursor:"pointer"},labelStyle:{color:"#666666"}}});e(x.lang,{rangeSelectorZoom:"Zoom",rangeSelectorFrom:"",rangeSelectorTo:"\u2192"});var E=function(){function a(c){this.buttons=void 0;this.buttonOptions=a.prototype.defaultButtons;this.initialButtonGroupWidth=0;this.options=void 0;this.chart=c;this.init(c)}
a.prototype.clickButton=function(a,e){var n=this.chart,f=this.buttonOptions[a],k=n.xAxis[0],g=n.scroller&&n.scroller.getUnionExtremes()||k||{},h=g.dataMin,l=g.dataMax,y=k&&Math.round(Math.min(k.max,C(l,k.max))),m=f.type;g=f._range;var p,t=f.dataGrouping;if(null!==h&&null!==l){n.fixedRange=g;this.setSelected(a);t&&(this.forcedDataGrouping=!0,b.prototype.setDataGrouping.call(k||{chart:this.chart},t,!1),this.frozenStates=f.preserveDataGrouping);if("month"===m||"year"===m)if(k){m={range:f,max:y,chart:n,
dataMin:h,dataMax:l};var u=k.minFromRange.call(m);r(m.newMax)&&(y=m.newMax)}else g=f;else if(g)u=Math.max(y-g,h),y=Math.min(u+g,l);else if("ytd"===m)if(k)"undefined"===typeof l&&(h=Number.MAX_VALUE,l=Number.MIN_VALUE,n.series.forEach(function(a){a=a.xData;h=Math.min(a[0],h);l=Math.max(a[a.length-1],l)}),e=!1),y=this.getYTDExtremes(l,h,n.time.useUTC),u=p=y.min,y=y.max;else{this.deferredYTDClick=a;return}else"all"===m&&k&&(n.navigator&&n.navigator.baseSeries[0]&&(n.navigator.baseSeries[0].xAxis.options.range=
void 0),u=h,y=l);A(u)&&(u+=f._offsetMin);A(y)&&(y+=f._offsetMax);this.dropdown&&(this.dropdown.selectedIndex=a+1);if(k)k.setExtremes(u,y,C(e,!0),void 0,{trigger:"rangeSelectorButton",rangeSelectorButton:f});else{var z=M(n.options.xAxis)[0];var v=z.range;z.range=g;var x=z.min;z.min=p;d(n,"load",function(){z.range=v;z.min=x})}c(this,"afterBtnClick")}};a.prototype.setSelected=function(a){this.selected=this.options.selected=a};a.prototype.init=function(a){var b=this,e=a.options.rangeSelector,f=e.buttons||
b.defaultButtons.slice(),k=e.selected,g=function(){var a=b.minInput,f=b.maxInput;a&&a.blur&&c(a,"blur");f&&f.blur&&c(f,"blur")};b.chart=a;b.options=e;b.buttons=[];b.buttonOptions=f;this.eventsToUnbind=[];this.eventsToUnbind.push(d(a.container,"mousedown",g));this.eventsToUnbind.push(d(a,"resize",g));f.forEach(b.computeButtonRange);"undefined"!==typeof k&&f[k]&&this.clickButton(k,!1);this.eventsToUnbind.push(d(a,"load",function(){a.xAxis&&a.xAxis[0]&&d(a.xAxis[0],"setExtremes",function(f){this.max-
this.min!==a.fixedRange&&"rangeSelectorButton"!==f.trigger&&"updatedData"!==f.trigger&&b.forcedDataGrouping&&!b.frozenStates&&this.setDataGrouping(!1,!1)})}))};a.prototype.updateButtonStates=function(){var a=this,b=this.chart,c=this.dropdown,f=b.xAxis[0],k=Math.round(f.max-f.min),e=!f.hasVisibleSeries,d=b.scroller&&b.scroller.getUnionExtremes()||f,g=d.dataMin,h=d.dataMax;b=a.getYTDExtremes(h,g,b.time.useUTC);var l=b.min,m=b.max,p=a.selected,t=r(p),u=a.options.allButtonsEnabled,z=a.buttons;a.buttonOptions.forEach(function(b,
d){var n=b._range,q=b.type,w=b.count||1,y=z[d],r=0,L=b._offsetMax-b._offsetMin;b=d===p;var P=n>h-g,O=n<f.minRange,N=!1,R=!1;n=n===k;("month"===q||"year"===q)&&k+36E5>=864E5*{month:28,year:365}[q]*w-L&&k-36E5<=864E5*{month:31,year:366}[q]*w+L?n=!0:"ytd"===q?(n=m-l+L===k,N=!b):"all"===q&&(n=f.max-f.min>=h-g,R=!b&&t&&n);q=!u&&(P||O||R||e);w=b&&n||n&&!t&&!N||b&&a.frozenStates;q?r=3:w&&(t=!0,r=2);y.state!==r&&(y.setState(r),c&&(c.options[d+1].disabled=q,2===r&&(c.selectedIndex=d+1)),0===r&&p===d&&a.setSelected())})};
a.prototype.computeButtonRange=function(a){var b=a.type,c=a.count||1,f={millisecond:1,second:1E3,minute:6E4,hour:36E5,day:864E5,week:6048E5};if(f[b])a._range=f[b]*c;else if("month"===b||"year"===b)a._range=864E5*{month:30,year:365}[b]*c;a._offsetMin=C(a.offsetMin,0);a._offsetMax=C(a.offsetMax,0);a._range+=a._offsetMax-a._offsetMin};a.prototype.getInputValue=function(a){a="min"===a?this.minInput:this.maxInput;var b=this.chart.options.rangeSelector,c=this.chart.time;return a?("text"===a.type&&b.inputDateParser||
this.defaultInputDateParser)(a.value,c.useUTC,c):0};a.prototype.setInputValue=function(a,b){var c=this.options,f=this.chart.time,k="min"===a?this.minInput:this.maxInput;a="min"===a?this.minDateBox:this.maxDateBox;if(k){var e=k.getAttribute("data-hc-time");e=A(e)?Number(e):void 0;A(b)&&(A(e)&&k.setAttribute("data-hc-time-previous",e),k.setAttribute("data-hc-time",b),e=b);k.value=f.dateFormat(this.inputTypeFormats[k.type]||c.inputEditDateFormat,e);a&&a.attr({text:f.dateFormat(c.inputDateFormat,e)})}};
a.prototype.setInputExtremes=function(a,b,c){if(a="min"===a?this.minInput:this.maxInput){var f=this.inputTypeFormats[a.type],k=this.chart.time;f&&(b=k.dateFormat(f,b),a.min!==b&&(a.min=b),c=k.dateFormat(f,c),a.max!==c&&(a.max=c))}};a.prototype.showInput=function(a){var b="min"===a?this.minDateBox:this.maxDateBox;if((a="min"===a?this.minInput:this.maxInput)&&b&&this.inputGroup){var c="text"===a.type,f=this.inputGroup,k=f.translateX;f=f.translateY;var e=this.options.inputBoxWidth;g(a,{width:c?b.width+
(e?-2:20)+"px":"auto",height:c?b.height-2+"px":"auto",border:"2px solid silver"});c&&e?g(a,{left:k+b.x+"px",top:f+"px"}):g(a,{left:Math.min(Math.round(b.x+k-(a.offsetWidth-b.width)/2),this.chart.chartWidth-a.offsetWidth)+"px",top:f-(a.offsetHeight-b.height)/2+"px"})}};a.prototype.hideInput=function(a){(a="min"===a?this.minInput:this.maxInput)&&g(a,{top:"-9999em",border:0,width:"1px",height:"1px"})};a.prototype.defaultInputDateParser=function(a,b,c){var f=a.split("/").join("-").split(" ").join("T");
-1===f.indexOf("T")&&(f+="T00:00");if(b)f+="Z";else{var k;if(k=v.isSafari)k=f,k=!(6<k.length&&(k.lastIndexOf("-")===k.length-6||k.lastIndexOf("+")===k.length-6));k&&(k=(new Date(f)).getTimezoneOffset()/60,f+=0>=k?"+"+m(-k)+":00":"-"+m(k)+":00")}f=Date.parse(f);r(f)||(a=a.split("-"),f=Date.UTC(J(a[0]),J(a[1])-1,J(a[2])));c&&b&&r(f)&&(f+=c.getTimezoneOffset(f));return f};a.prototype.drawInput=function(a){function b(){var b=d.getInputValue(a),f=c.xAxis[0],k=c.scroller&&c.scroller.xAxis?c.scroller.xAxis:
f,e=k.dataMin;k=k.dataMax;var g=d.maxInput,h=d.minInput;b!==Number(u.getAttribute("data-hc-time-previous"))&&r(b)&&(u.setAttribute("data-hc-time-previous",b),t&&g&&r(e)?b>Number(g.getAttribute("data-hc-time"))?b=void 0:b<e&&(b=e):h&&r(k)&&(b<Number(h.getAttribute("data-hc-time"))?b=void 0:b>k&&(b=k)),"undefined"!==typeof b&&f.setExtremes(t?b:f.min,t?f.max:b,void 0,void 0,{trigger:"rangeSelectorInput"}))}var c=this.chart,f=this.div,k=this.inputGroup,d=this,n=c.renderer.style||{},l=c.renderer,m=c.options.rangeSelector,
t="min"===a,P=x.lang[t?"rangeSelectorFrom":"rangeSelectorTo"]||"";P=l.label(P,0).addClass("highcharts-range-label").attr({padding:P?2:0,height:P?m.inputBoxHeight:0}).add(k);l=l.label("",0).addClass("highcharts-range-input").attr({padding:2,width:m.inputBoxWidth,height:m.inputBoxHeight,"text-align":"center"}).on("click",function(){d.showInput(a);d[a+"Input"].focus()});c.styledMode||l.attr({stroke:m.inputBoxBorderColor,"stroke-width":1});l.add(k);var u=p("input",{name:a,className:"highcharts-range-selector"},
void 0,f);u.setAttribute("type",h(m.inputDateFormat||"%b %e, %Y"));c.styledMode||(P.css(F(n,m.labelStyle)),l.css(F({color:"#333333"},n,m.inputStyle)),g(u,e({position:"absolute",border:0,boxShadow:"0 0 15px rgba(0,0,0,0.3)",width:"1px",height:"1px",padding:0,textAlign:"center",fontSize:n.fontSize,fontFamily:n.fontFamily,top:"-9999em"},m.inputStyle)));u.onfocus=function(){d.showInput(a)};u.onblur=function(){u===v.doc.activeElement&&b();d.hideInput(a);d.setInputValue(a);u.blur()};var z=!1;u.onchange=
function(){z||(b(),d.hideInput(a),u.blur())};u.onkeypress=function(a){13===a.keyCode&&b()};u.onkeydown=function(a){z=!0;38!==a.keyCode&&40!==a.keyCode||b()};u.onkeyup=function(){z=!1};return{dateBox:l,input:u,label:P}};a.prototype.getPosition=function(){var a=this.chart,b=a.options.rangeSelector;a="top"===b.verticalAlign?a.plotTop-a.axisOffset[0]:0;return{buttonTop:a+b.buttonPosition.y,inputTop:a+b.inputPosition.y-10}};a.prototype.getYTDExtremes=function(a,b,c){var f=this.chart.time,k=new f.Date(a),
e=f.get("FullYear",k);c=c?f.Date.UTC(e,0,1):+new f.Date(e,0,1);b=Math.max(b,c);k=k.getTime();return{max:Math.min(a||k,k),min:b}};a.prototype.render=function(a,b){var c=this.chart,f=c.renderer,k=c.container,e=c.options,d=e.rangeSelector,g=C(e.chart.style&&e.chart.style.zIndex,0)+1;e=d.inputEnabled;if(!1!==d.enabled){this.rendered||(this.group=f.g("range-selector-group").attr({zIndex:7}).add(),this.div=p("div",void 0,{position:"relative",height:0,zIndex:g}),this.buttonOptions.length&&this.renderButtons(),
k.parentNode&&k.parentNode.insertBefore(this.div,k),e&&(this.inputGroup=f.g("input-group").add(this.group),f=this.drawInput("min"),this.minDateBox=f.dateBox,this.minLabel=f.label,this.minInput=f.input,f=this.drawInput("max"),this.maxDateBox=f.dateBox,this.maxLabel=f.label,this.maxInput=f.input));if(e&&(this.setInputValue("min",a),this.setInputValue("max",b),a=c.scroller&&c.scroller.getUnionExtremes()||c.xAxis[0]||{},A(a.dataMin)&&A(a.dataMax)&&(c=c.xAxis[0].minRange||0,this.setInputExtremes("min",
a.dataMin,Math.min(a.dataMax,this.getInputValue("max"))-c),this.setInputExtremes("max",Math.max(a.dataMin,this.getInputValue("min"))+c,a.dataMax)),this.inputGroup)){var h=0;[this.minLabel,this.minDateBox,this.maxLabel,this.maxDateBox].forEach(function(a){if(a){var b=a.getBBox().width;b&&(a.attr({x:h}),h+=b+d.inputSpacing)}})}this.alignElements();this.rendered=!0}};a.prototype.renderButtons=function(){var a=this,b=this.buttons,e=this.options,f=x.lang,k=this.chart.renderer,g=F(e.buttonTheme),h=g&&g.states,
l=g.width||28;delete g.width;delete g.states;this.buttonGroup=k.g("range-selector-buttons").add(this.group);var m=this.dropdown=p("select",void 0,{position:"absolute",width:"1px",height:"1px",padding:0,border:0,top:"-9999em",cursor:"pointer",opacity:.0001},this.div);d(m,"touchstart",function(){m.style.fontSize="16px"});[[v.isMS?"mouseover":"mouseenter"],[v.isMS?"mouseout":"mouseleave"],["change","click"]].forEach(function(f){var k=f[0],e=f[1];d(m,k,function(){var f=b[a.currentButtonIndex()];f&&c(f.element,
e||k)})});this.zoomText=k.label(f&&f.rangeSelectorZoom||"",0).attr({padding:e.buttonTheme.padding,height:e.buttonTheme.height,paddingLeft:0,paddingRight:0}).add(this.buttonGroup);this.chart.styledMode||(this.zoomText.css(e.labelStyle),g["stroke-width"]=C(g["stroke-width"],0));p("option",{textContent:this.zoomText.textStr,disabled:!0},void 0,m);this.buttonOptions.forEach(function(c,f){p("option",{textContent:c.title||c.text},void 0,m);b[f]=k.button(c.text,0,0,function(b){var k=c.events&&c.events.click,
e;k&&(e=k.call(c,b));!1!==e&&a.clickButton(f);a.isActive=!0},g,h&&h.hover,h&&h.select,h&&h.disabled).attr({"text-align":"center",width:l}).add(a.buttonGroup);c.title&&b[f].attr("title",c.title)})};a.prototype.alignElements=function(){var a=this,b=this.buttonGroup,c=this.buttons,f=this.chart,k=this.group,e=this.inputGroup,d=this.options,g=this.zoomText,h=f.options,l=h.exporting&&!1!==h.exporting.enabled&&h.navigation&&h.navigation.buttonOptions;h=d.buttonPosition;var m=d.inputPosition,r=d.verticalAlign,
p=function(b,c){return l&&a.titleCollision(f)&&"top"===r&&"right"===c.align&&c.y-b.getBBox().height-12<(l.y||0)+(l.height||0)+f.spacing[0]?-40:0},t=f.plotLeft;if(k&&h&&m){var u=h.x-f.spacing[3];if(b){this.positionButtons();if(!this.initialButtonGroupWidth){var z=0;g&&(z+=g.getBBox().width+5);c.forEach(function(a,b){z+=a.width;b!==c.length-1&&(z+=d.buttonSpacing)});this.initialButtonGroupWidth=z}t-=f.spacing[3];this.updateButtonStates();g=p(b,h);this.alignButtonGroup(g);k.placed=b.placed=f.hasLoaded}b=
0;e&&(b=p(e,m),"left"===m.align?u=t:"right"===m.align&&(u=-Math.max(f.axisOffset[1],-b)),e.align({y:m.y,width:e.getBBox().width,align:m.align,x:m.x+u-2},!0,f.spacingBox),e.placed=f.hasLoaded);this.handleCollision(b);k.align({verticalAlign:r},!0,f.spacingBox);e=k.alignAttr.translateY;b=k.getBBox().height+20;p=0;"bottom"===r&&(p=(p=f.legend&&f.legend.options)&&"bottom"===p.verticalAlign&&p.enabled&&!p.floating?f.legend.legendHeight+C(p.margin,10):0,b=b+p-20,p=e-b-(d.floating?0:d.y)-(f.titleOffset?f.titleOffset[2]:
0)-10);if("top"===r)d.floating&&(p=0),f.titleOffset&&f.titleOffset[0]&&(p=f.titleOffset[0]),p+=f.margin[0]-f.spacing[0]||0;else if("middle"===r)if(m.y===h.y)p=e;else if(m.y||h.y)p=0>m.y||0>h.y?p-Math.min(m.y,h.y):e-b;k.translate(d.x,d.y+Math.floor(p));h=this.minInput;m=this.maxInput;e=this.dropdown;d.inputEnabled&&h&&m&&(h.style.marginTop=k.translateY+"px",m.style.marginTop=k.translateY+"px");e&&(e.style.marginTop=k.translateY+"px")}};a.prototype.alignButtonGroup=function(a,b){var c=this.chart,f=
this.buttonGroup,k=this.options.buttonPosition,e=c.plotLeft-c.spacing[3],d=k.x-c.spacing[3];"right"===k.align?d+=a-e:"center"===k.align&&(d-=e/2);f&&f.align({y:k.y,width:C(b,this.initialButtonGroupWidth),align:k.align,x:d},!0,c.spacingBox)};a.prototype.positionButtons=function(){var a=this.buttons,b=this.chart,c=this.options,f=this.zoomText,k=b.hasLoaded?"animate":"attr",e=c.buttonPosition,d=b.plotLeft,g=d;f&&"hidden"!==f.visibility&&(f[k]({x:C(d+e.x,d)}),g+=e.x+f.getBBox().width+5);this.buttonOptions.forEach(function(b,
f){if("hidden"!==a[f].visibility)a[f][k]({x:g}),g+=a[f].width+c.buttonSpacing;else a[f][k]({x:d})})};a.prototype.handleCollision=function(a){var b=this,c=this.chart,f=this.buttonGroup,k=this.inputGroup,e=this.options,d=e.buttonPosition,g=e.dropdown,h=e.inputPosition;e=function(){var a=0;b.buttons.forEach(function(b){b=b.getBBox();b.width>a&&(a=b.width)});return a};var n=function(b){if(k&&f){var c=k.alignAttr.translateX+k.alignOptions.x-a+k.getBBox().x+2,e=k.alignOptions.width,g=f.alignAttr.translateX+
f.getBBox().x;return g+b>c&&c+e>g&&d.y<h.y+k.getBBox().height}return!1},l=function(){k&&f&&k.attr({translateX:k.alignAttr.translateX+(c.axisOffset[1]>=-a?0:-a),translateY:k.alignAttr.translateY+f.getBBox().height+10})};if(f){if("always"===g){this.collapseButtons(a);n(e())&&l();return}"never"===g&&this.expandButtons()}k&&f?h.align===d.align||n(this.initialButtonGroupWidth+20)?"responsive"===g?(this.collapseButtons(a),n(e())&&l()):l():"responsive"===g&&this.expandButtons():f&&"responsive"===g&&(this.initialButtonGroupWidth>
c.plotWidth?this.collapseButtons(a):this.expandButtons())};a.prototype.collapseButtons=function(a){var b=this.buttons,c=this.buttonOptions,f=this.chart,e=this.dropdown,d=this.options,g=this.zoomText,h=f.userOptions.rangeSelector&&f.userOptions.rangeSelector.buttonTheme||{},n=function(a){return{text:a?a+" \u25be":"\u25be",width:"auto",paddingLeft:C(d.buttonTheme.paddingLeft,h.padding,8),paddingRight:C(d.buttonTheme.paddingRight,h.padding,8)}};g&&g.hide();var l=!1;c.forEach(function(a,c){c=b[c];2!==
c.state?c.hide():(c.show(),c.attr(n(a.text)),l=!0)});l||(e&&(e.selectedIndex=0),b[0].show(),b[0].attr(n(this.zoomText&&this.zoomText.textStr)));c=d.buttonPosition.align;this.positionButtons();"right"!==c&&"center"!==c||this.alignButtonGroup(a,b[this.currentButtonIndex()].getBBox().width);this.showDropdown()};a.prototype.expandButtons=function(){var a=this.buttons,b=this.buttonOptions,c=this.options,f=this.zoomText;this.hideDropdown();f&&f.show();b.forEach(function(b,f){f=a[f];f.show();f.attr({text:b.text,
width:c.buttonTheme.width||28,paddingLeft:C(c.buttonTheme.paddingLeft,"unset"),paddingRight:C(c.buttonTheme.paddingRight,"unset")});2>f.state&&f.setState(0)});this.positionButtons()};a.prototype.currentButtonIndex=function(){var a=this.dropdown;return a&&0<a.selectedIndex?a.selectedIndex-1:0};a.prototype.showDropdown=function(){var a=this.buttonGroup,b=this.buttons,c=this.chart,f=this.dropdown;if(a&&f){var e=a.translateX;a=a.translateY;b=b[this.currentButtonIndex()].getBBox();g(f,{left:c.plotLeft+
e+"px",top:a+.5+"px",width:b.width+"px",height:b.height+"px"});this.hasVisibleDropdown=!0}};a.prototype.hideDropdown=function(){var a=this.dropdown;a&&(g(a,{top:"-9999em",width:"1px",height:"1px"}),this.hasVisibleDropdown=!1)};a.prototype.getHeight=function(){var a=this.options,b=this.group,c=a.y,f=a.buttonPosition.y,e=a.inputPosition.y;if(a.height)return a.height;this.alignElements();a=b?b.getBBox(!0).height+13+c:0;b=Math.min(e,f);if(0>e&&0>f||0<e&&0<f)a+=Math.abs(b);return a};a.prototype.titleCollision=
function(a){return!(a.options.title.text||a.options.subtitle.text)};a.prototype.update=function(a){var b=this.chart;F(!0,b.options.rangeSelector,a);this.destroy();this.init(b);this.render()};a.prototype.destroy=function(){var b=this,c=b.minInput,e=b.maxInput;b.eventsToUnbind&&(b.eventsToUnbind.forEach(function(a){return a()}),b.eventsToUnbind=void 0);D(b.buttons);c&&(c.onfocus=c.onblur=c.onchange=null);e&&(e.onfocus=e.onblur=e.onchange=null);G(b,function(c,e){c&&"chart"!==e&&(c instanceof B?c.destroy():
c instanceof window.HTMLElement&&H(c));c!==a.prototype[e]&&(b[e]=null)},this)};return a}();E.prototype.defaultButtons=[{type:"month",count:1,text:"1m",title:"View 1 month"},{type:"month",count:3,text:"3m",title:"View 3 months"},{type:"month",count:6,text:"6m",title:"View 6 months"},{type:"ytd",text:"YTD",title:"View year to date"},{type:"year",count:1,text:"1y",title:"View 1 year"},{type:"all",text:"All",title:"View all"}];E.prototype.inputTypeFormats={"datetime-local":"%Y-%m-%dT%H:%M:%S",date:"%Y-%m-%d",
time:"%H:%M:%S"};b.prototype.minFromRange=function(){var a=this.range,b=a.type,c=this.max,e=this.chart.time,f=function(a,c){var f="year"===b?"FullYear":"Month",k=new e.Date(a),d=e.get(f,k);e.set(f,k,d+c);d===e.get(f,k)&&e.set("Date",k,0);return k.getTime()-a};if(r(a)){var k=c-a;var d=a}else k=c+f(c,-a.count),this.chart&&(this.chart.fixedRange=c-k);var g=C(this.dataMin,Number.MIN_VALUE);r(k)||(k=g);k<=g&&(k=g,"undefined"===typeof d&&(d=f(k,a.count)),this.newMax=Math.min(k+d,this.dataMax));r(c)||(k=
void 0);return k};if(!v.RangeSelector){var K=[],I=function(b){function c(){g&&(e=b.xAxis[0].getExtremes(),f=b.legend,h=g&&g.options.verticalAlign,r(e.min)&&g.render(e.min,e.max),f.display&&"top"===h&&h===f.options.verticalAlign&&(k=F(b.spacingBox),k.y="vertical"===f.options.layout?b.plotTop:k.y+g.getHeight(),f.group.placed=!1,f.align(k)))}var e,g=b.rangeSelector,f,k,h;g&&(a(K,function(a){return a[0]===b})||K.push([b,[d(b.xAxis[0],"afterSetExtremes",function(a){g&&g.render(a.min,a.max)}),d(b,"redraw",
c)]]),c())};d(t,"afterGetContainer",function(){this.options.rangeSelector&&this.options.rangeSelector.enabled&&(this.rangeSelector=new E(this))});d(t,"beforeRender",function(){var a=this.axes,b=this.rangeSelector;b&&(r(b.deferredYTDClick)&&(b.clickButton(b.deferredYTDClick),delete b.deferredYTDClick),a.forEach(function(a){a.updateNames();a.setScale()}),this.getAxisMargins(),b.render(),a=b.options.verticalAlign,b.options.floating||("bottom"===a?this.extraBottomMargin=!0:"middle"!==a&&(this.extraTopMargin=
!0)))});d(t,"update",function(a){var b=a.options.rangeSelector;a=this.rangeSelector;var c=this.extraBottomMargin,e=this.extraTopMargin;b&&b.enabled&&!A(a)&&this.options.rangeSelector&&(this.options.rangeSelector.enabled=!0,this.rangeSelector=a=new E(this));this.extraTopMargin=this.extraBottomMargin=!1;a&&(I(this),b=b&&b.verticalAlign||a.options&&a.options.verticalAlign,a.options.floating||("bottom"===b?this.extraBottomMargin=!0:"middle"!==b&&(this.extraTopMargin=!0)),this.extraBottomMargin!==c||this.extraTopMargin!==
e)&&(this.isDirtyBox=!0)});d(t,"render",function(){var a=this.rangeSelector;a&&!a.options.floating&&(a.render(),a=a.options.verticalAlign,"bottom"===a?this.extraBottomMargin=!0:"middle"!==a&&(this.extraTopMargin=!0))});d(t,"getMargins",function(){var a=this.rangeSelector;a&&(a=a.getHeight(),this.extraTopMargin&&(this.plotTop+=a),this.extraBottomMargin&&(this.marginBottom+=a))});t.prototype.callbacks.push(I);d(t,"destroy",function(){for(var a=0;a<K.length;a++){var b=K[a];if(b[0]===this){b[1].forEach(function(a){return a()});
K.splice(a,1);break}}});v.RangeSelector=E}return E});J(b,"Core/Axis/NavigatorAxis.js",[b["Core/Globals.js"],b["Core/Utilities.js"]],function(b,t){var v=b.isTouchDevice,u=t.addEvent,B=t.correctFloat,l=t.defined,h=t.isNumber,x=t.pick,d=function(){function b(b){this.axis=b}b.prototype.destroy=function(){this.axis=void 0};b.prototype.toFixedRange=function(b,d,p,t){var e=this.axis,a=e.chart;a=a&&a.fixedRange;var c=(e.pointRange||0)/2;b=x(p,e.translate(b,!0,!e.horiz));d=x(t,e.translate(d,!0,!e.horiz));
e=a&&(d-b)/a;l(p)||(b=B(b+c));l(t)||(d=B(d-c));.7<e&&1.3>e&&(t?b=d-a:d=b+a);h(b)&&h(d)||(b=d=void 0);return{min:b,max:d}};return b}();return function(){function b(){}b.compose=function(b){b.keepProps.push("navigatorAxis");u(b,"init",function(){this.navigatorAxis||(this.navigatorAxis=new d(this))});u(b,"zoom",function(b){var d=this.chart.options,g=d.navigator,e=this.navigatorAxis,a=d.chart.pinchType,c=d.rangeSelector;d=d.chart.zoomType;this.isXAxis&&(g&&g.enabled||c&&c.enabled)&&("y"===d?b.zoomed=
!1:(!v&&"xy"===d||v&&"xy"===a)&&this.options.range&&(g=e.previousZoom,l(b.newMin)?e.previousZoom=[this.min,this.max]:g&&(b.newMin=g[0],b.newMax=g[1],e.previousZoom=void 0)));"undefined"!==typeof b.zoomed&&b.preventDefault()})};b.AdditionsClass=d;return b}()});J(b,"Core/Navigator.js",[b["Core/Axis/Axis.js"],b["Core/Chart/Chart.js"],b["Core/Color/Color.js"],b["Core/Globals.js"],b["Core/Axis/NavigatorAxis.js"],b["Core/DefaultOptions.js"],b["Core/Renderer/RendererRegistry.js"],b["Core/Scrollbar.js"],
b["Core/Series/Series.js"],b["Core/Series/SeriesRegistry.js"],b["Core/Utilities.js"]],function(b,t,v,u,B,l,h,x,d,p,g){v=v.parse;var A=u.hasTouch,D=u.isTouchDevice,H=l.defaultOptions,e=g.addEvent,a=g.clamp,c=g.correctFloat,r=g.defined,F=g.destroyObjectProperties,G=g.erase,m=g.extend,C=g.find,J=g.isArray,M=g.isNumber,E=g.merge,K=g.pick,I=g.removeEvent,z=g.splat,n=function(a){for(var b=[],c=1;c<arguments.length;c++)b[c-1]=arguments[c];b=[].filter.call(b,M);if(b.length)return Math[a].apply(0,b)};l="undefined"===
typeof p.seriesTypes.areaspline?"line":"areaspline";m(H,{navigator:{height:40,margin:25,maskInside:!0,handles:{width:7,height:15,symbols:["navigator-handle","navigator-handle"],enabled:!0,lineWidth:1,backgroundColor:"#f2f2f2",borderColor:"#999999"},maskFill:v("#6685c2").setOpacity(.3).get(),outlineColor:"#cccccc",outlineWidth:1,series:{type:l,fillOpacity:.05,lineWidth:1,compare:null,dataGrouping:{approximation:"average",enabled:!0,groupPixelWidth:2,firstAnchor:"firstPoint",anchor:"middle",lastAnchor:"lastPoint",
units:[["millisecond",[1,2,5,10,20,25,50,100,200,500]],["second",[1,2,5,10,15,30]],["minute",[1,2,5,10,15,30]],["hour",[1,2,3,4,6,8,12]],["day",[1,2,3,4]],["week",[1,2,3]],["month",[1,3,6]],["year",null]]},dataLabels:{enabled:!1,zIndex:2},id:"highcharts-navigator-series",className:"highcharts-navigator-series",lineColor:null,marker:{enabled:!1},threshold:null},xAxis:{overscroll:0,className:"highcharts-navigator-xaxis",tickLength:0,lineWidth:0,gridLineColor:"#e6e6e6",gridLineWidth:1,tickPixelInterval:200,
labels:{align:"left",style:{color:"#999999"},x:3,y:-4},crosshair:!1},yAxis:{className:"highcharts-navigator-yaxis",gridLineWidth:0,startOnTick:!1,endOnTick:!1,minPadding:.1,maxPadding:.1,labels:{enabled:!1},crosshair:!1,title:{text:null},tickLength:0,tickWidth:0}}});h.getRendererType().prototype.symbols["navigator-handle"]=function(a,b,c,e,d){a=(d&&d.width||0)/2;b=Math.round(a/3)+.5;d=d&&d.height||0;return[["M",-a-1,.5],["L",a,.5],["L",a,d+.5],["L",-a-1,d+.5],["L",-a-1,.5],["M",-b,4],["L",-b,d-3],
["M",b-1,4],["L",b-1,d-3]]};var y=function(){function d(a){this.zoomedMin=this.zoomedMax=this.yAxis=this.xAxis=this.top=this.size=this.shades=this.rendered=this.range=this.outlineHeight=this.outline=this.opposite=this.navigatorSize=this.navigatorSeries=this.navigatorOptions=this.navigatorGroup=this.navigatorEnabled=this.left=this.height=this.handles=this.chart=this.baseSeries=void 0;this.init(a)}d.prototype.drawHandle=function(a,b,c,e){var f=this.navigatorOptions.handles.height;this.handles[b][e](c?
{translateX:Math.round(this.left+this.height/2),translateY:Math.round(this.top+parseInt(a,10)+.5-f)}:{translateX:Math.round(this.left+parseInt(a,10)),translateY:Math.round(this.top+this.height/2-f/2-1)})};d.prototype.drawOutline=function(a,b,c,e){var f=this.navigatorOptions.maskInside,d=this.outline.strokeWidth(),k=d/2,g=d%2/2;d=this.outlineHeight;var h=this.scrollbarHeight||0,l=this.size,n=this.left-h,m=this.top;c?(n-=k,c=m+b+g,b=m+a+g,g=[["M",n+d,m-h-g],["L",n+d,c],["L",n,c],["L",n,b],["L",n+d,
b],["L",n+d,m+l+h]],f&&g.push(["M",n+d,c-k],["L",n+d,b+k])):(a+=n+h-g,b+=n+h-g,m+=k,g=[["M",n,m],["L",a,m],["L",a,m+d],["L",b,m+d],["L",b,m],["L",n+l+2*h,m]],f&&g.push(["M",a-k,m],["L",b+k,m]));this.outline[e]({d:g})};d.prototype.drawMasks=function(a,b,c,e){var f=this.left,d=this.top,k=this.height;if(c){var g=[f,f,f];var h=[d,d+a,d+b];var l=[k,k,k];var n=[a,b-a,this.size-b]}else g=[f,f+a,f+b],h=[d,d,d],l=[a,b-a,this.size-b],n=[k,k,k];this.shades.forEach(function(a,b){a[e]({x:g[b],y:h[b],width:l[b],
height:n[b]})})};d.prototype.renderElements=function(){var a=this,b=a.navigatorOptions,c=b.maskInside,e=a.chart,d=e.renderer,g,h={cursor:e.inverted?"ns-resize":"ew-resize"};a.navigatorGroup=g=d.g("navigator").attr({zIndex:8,visibility:"hidden"}).add();[!c,c,!c].forEach(function(c,f){a.shades[f]=d.rect().addClass("highcharts-navigator-mask"+(1===f?"-inside":"-outside")).add(g);e.styledMode||a.shades[f].attr({fill:c?b.maskFill:"rgba(0,0,0,0)"}).css(1===f&&h)});a.outline=d.path().addClass("highcharts-navigator-outline").add(g);
e.styledMode||a.outline.attr({"stroke-width":b.outlineWidth,stroke:b.outlineColor});b.handles.enabled&&[0,1].forEach(function(c){b.handles.inverted=e.inverted;a.handles[c]=d.symbol(b.handles.symbols[c],-b.handles.width/2-1,0,b.handles.width,b.handles.height,b.handles);a.handles[c].attr({zIndex:7-c}).addClass("highcharts-navigator-handle highcharts-navigator-handle-"+["left","right"][c]).add(g);if(!e.styledMode){var f=b.handles;a.handles[c].attr({fill:f.backgroundColor,stroke:f.borderColor,"stroke-width":f.lineWidth}).css(h)}})};
d.prototype.update=function(a){(this.series||[]).forEach(function(a){a.baseSeries&&delete a.baseSeries.navigatorSeries});this.destroy();E(!0,this.chart.options.navigator,this.options,a);this.init(this.chart)};d.prototype.render=function(b,e,d,g){var f=this.chart,k=this.scrollbarHeight,h,l=this.xAxis,n=l.pointRange||0;var m=l.navigatorAxis.fake?f.xAxis[0]:l;var q=this.navigatorEnabled,w,p=this.rendered;var t=f.inverted;var u=f.xAxis[0].minRange,v=f.xAxis[0].options.maxRange;if(!this.hasDragged||r(d)){b=
c(b-n/2);e=c(e+n/2);if(!M(b)||!M(e))if(p)d=0,g=K(l.width,m.width);else return;this.left=K(l.left,f.plotLeft+k+(t?f.plotWidth:0));this.size=w=h=K(l.len,(t?f.plotHeight:f.plotWidth)-2*k);f=t?k:h+2*k;d=K(d,l.toPixels(b,!0));g=K(g,l.toPixels(e,!0));M(d)&&Infinity!==Math.abs(d)||(d=0,g=f);b=l.toValue(d,!0);e=l.toValue(g,!0);var x=Math.abs(c(e-b));x<u?this.grabbedLeft?d=l.toPixels(e-u-n,!0):this.grabbedRight&&(g=l.toPixels(b+u+n,!0)):r(v)&&c(x-n)>v&&(this.grabbedLeft?d=l.toPixels(e-v-n,!0):this.grabbedRight&&
(g=l.toPixels(b+v+n,!0)));this.zoomedMax=a(Math.max(d,g),0,w);this.zoomedMin=a(this.fixedWidth?this.zoomedMax-this.fixedWidth:Math.min(d,g),0,w);this.range=this.zoomedMax-this.zoomedMin;w=Math.round(this.zoomedMax);d=Math.round(this.zoomedMin);q&&(this.navigatorGroup.attr({visibility:"visible"}),p=p&&!this.hasDragged?"animate":"attr",this.drawMasks(d,w,t,p),this.drawOutline(d,w,t,p),this.navigatorOptions.handles.enabled&&(this.drawHandle(d,0,t,p),this.drawHandle(w,1,t,p)));this.scrollbar&&(t?(t=this.top-
k,m=this.left-k+(q||!m.opposite?0:(m.titleOffset||0)+m.axisTitleMargin),k=h+2*k):(t=this.top+(q?this.height:-k),m=this.left-k),this.scrollbar.position(m,t,f,k),this.scrollbar.setRange(this.zoomedMin/(h||1),this.zoomedMax/(h||1)));this.rendered=!0}};d.prototype.addMouseEvents=function(){var a=this,b=a.chart,c=b.container,d=[],g,h;a.mouseMoveHandler=g=function(b){a.onMouseMove(b)};a.mouseUpHandler=h=function(b){a.onMouseUp(b)};d=a.getPartsEvents("mousedown");d.push(e(b.renderTo,"mousemove",g),e(c.ownerDocument,
"mouseup",h));A&&(d.push(e(b.renderTo,"touchmove",g),e(c.ownerDocument,"touchend",h)),d.concat(a.getPartsEvents("touchstart")));a.eventsToUnbind=d;a.series&&a.series[0]&&d.push(e(a.series[0].xAxis,"foundExtremes",function(){b.navigator.modifyNavigatorAxisExtremes()}))};d.prototype.getPartsEvents=function(a){var b=this,c=[];["shades","handles"].forEach(function(f){b[f].forEach(function(d,k){c.push(e(d.element,a,function(a){b[f+"Mousedown"](a,k)}))})});return c};d.prototype.shadesMousedown=function(a,
b){a=this.chart.pointer.normalize(a);var c=this.chart,f=this.xAxis,d=this.zoomedMin,e=this.left,k=this.size,g=this.range,h=a.chartX;c.inverted&&(h=a.chartY,e=this.top);if(1===b)this.grabbedCenter=h,this.fixedWidth=g,this.dragOffset=h-d;else{a=h-e-g/2;if(0===b)a=Math.max(0,a);else if(2===b&&a+g>=k)if(a=k-g,this.reversedExtremes){a-=g;var l=this.getUnionExtremes().dataMin}else var n=this.getUnionExtremes().dataMax;a!==d&&(this.fixedWidth=g,b=f.navigatorAxis.toFixedRange(a,a+g,l,n),r(b.min)&&c.xAxis[0].setExtremes(Math.min(b.min,
b.max),Math.max(b.min,b.max),!0,null,{trigger:"navigator"}))}};d.prototype.handlesMousedown=function(a,b){this.chart.pointer.normalize(a);a=this.chart;var c=a.xAxis[0],f=this.reversedExtremes;0===b?(this.grabbedLeft=!0,this.otherHandlePos=this.zoomedMax,this.fixedExtreme=f?c.min:c.max):(this.grabbedRight=!0,this.otherHandlePos=this.zoomedMin,this.fixedExtreme=f?c.max:c.min);a.fixedRange=null};d.prototype.onMouseMove=function(a){var b=this,c=b.chart,f=b.left,d=b.navigatorSize,e=b.range,g=b.dragOffset,
h=c.inverted;a.touches&&0===a.touches[0].pageX||(a=c.pointer.normalize(a),c=a.chartX,h&&(f=b.top,c=a.chartY),b.grabbedLeft?(b.hasDragged=!0,b.render(0,0,c-f,b.otherHandlePos)):b.grabbedRight?(b.hasDragged=!0,b.render(0,0,b.otherHandlePos,c-f)):b.grabbedCenter&&(b.hasDragged=!0,c<g?c=g:c>d+g-e&&(c=d+g-e),b.render(0,0,c-g,c-g+e)),b.hasDragged&&b.scrollbar&&K(b.scrollbar.options.liveRedraw,u.svg&&!D&&!this.chart.isBoosting)&&(a.DOMType=a.type,setTimeout(function(){b.onMouseUp(a)},0)))};d.prototype.onMouseUp=
function(a){var b=this.chart,c=this.xAxis,f=this.scrollbar,d=a.DOMEvent||a,e=b.inverted,g=this.rendered&&!this.hasDragged?"animate":"attr";if(this.hasDragged&&(!f||!f.hasDragged)||"scrollbar"===a.trigger){f=this.getUnionExtremes();if(this.zoomedMin===this.otherHandlePos)var h=this.fixedExtreme;else if(this.zoomedMax===this.otherHandlePos)var l=this.fixedExtreme;this.zoomedMax===this.size&&(l=this.reversedExtremes?f.dataMin:f.dataMax);0===this.zoomedMin&&(h=this.reversedExtremes?f.dataMax:f.dataMin);
c=c.navigatorAxis.toFixedRange(this.zoomedMin,this.zoomedMax,h,l);r(c.min)&&b.xAxis[0].setExtremes(Math.min(c.min,c.max),Math.max(c.min,c.max),!0,this.hasDragged?!1:null,{trigger:"navigator",triggerOp:"navigator-drag",DOMEvent:d})}"mousemove"!==a.DOMType&&"touchmove"!==a.DOMType&&(this.grabbedLeft=this.grabbedRight=this.grabbedCenter=this.fixedWidth=this.fixedExtreme=this.otherHandlePos=this.hasDragged=this.dragOffset=null);this.navigatorEnabled&&M(this.zoomedMin)&&M(this.zoomedMax)&&(b=Math.round(this.zoomedMin),
a=Math.round(this.zoomedMax),this.shades&&this.drawMasks(b,a,e,g),this.outline&&this.drawOutline(b,a,e,g),this.navigatorOptions.handles.enabled&&Object.keys(this.handles).length===this.handles.length&&(this.drawHandle(b,0,e,g),this.drawHandle(a,1,e,g)))};d.prototype.removeEvents=function(){this.eventsToUnbind&&(this.eventsToUnbind.forEach(function(a){a()}),this.eventsToUnbind=void 0);this.removeBaseSeriesEvents()};d.prototype.removeBaseSeriesEvents=function(){var a=this.baseSeries||[];this.navigatorEnabled&&
a[0]&&(!1!==this.navigatorOptions.adaptToUpdatedData&&a.forEach(function(a){I(a,"updatedData",this.updatedDataHandler)},this),a[0].xAxis&&I(a[0].xAxis,"foundExtremes",this.modifyBaseAxisExtremes))};d.prototype.init=function(a){var c=a.options,f=c.navigator,d=f.enabled,g=c.scrollbar,h=g.enabled;c=d?f.height:0;var l=h?g.height:0;this.handles=[];this.shades=[];this.chart=a;this.setBaseSeries();this.height=c;this.scrollbarHeight=l;this.scrollbarEnabled=h;this.navigatorEnabled=d;this.navigatorOptions=
f;this.scrollbarOptions=g;this.outlineHeight=c+l;this.opposite=K(f.opposite,!(d||!a.inverted));var m=this;d=m.baseSeries;g=a.xAxis.length;h=a.yAxis.length;var q=d&&d[0]&&d[0].xAxis||a.xAxis[0]||{options:{}};a.isDirtyBox=!0;m.navigatorEnabled?(m.xAxis=new b(a,E({breaks:q.options.breaks,ordinal:q.options.ordinal},f.xAxis,{id:"navigator-x-axis",yAxis:"navigator-y-axis",isX:!0,type:"datetime",index:g,isInternal:!0,offset:0,keepOrdinalPadding:!0,startOnTick:!1,endOnTick:!1,minPadding:0,maxPadding:0,zoomEnabled:!1},
a.inverted?{offsets:[l,0,-l,0],width:c}:{offsets:[0,-l,0,l],height:c})),m.yAxis=new b(a,E(f.yAxis,{id:"navigator-y-axis",alignTicks:!1,offset:0,index:h,isInternal:!0,reversed:K(f.yAxis&&f.yAxis.reversed,a.yAxis[0]&&a.yAxis[0].reversed,!1),zoomEnabled:!1},a.inverted?{width:c}:{height:c})),d||f.series.data?m.updateNavigatorSeries(!1):0===a.series.length&&(m.unbindRedraw=e(a,"beforeRedraw",function(){0<a.series.length&&!m.series&&(m.setBaseSeries(),m.unbindRedraw())})),m.reversedExtremes=a.inverted&&
!m.xAxis.reversed||!a.inverted&&m.xAxis.reversed,m.renderElements(),m.addMouseEvents()):(m.xAxis={chart:a,navigatorAxis:{fake:!0},translate:function(b,c){var f=a.xAxis[0],d=f.getExtremes(),e=f.len-2*l,g=n("min",f.options.min,d.dataMin);f=n("max",f.options.max,d.dataMax)-g;return c?b*f/e+g:e*(b-g)/f},toPixels:function(a){return this.translate(a)},toValue:function(a){return this.translate(a,!0)}},m.xAxis.navigatorAxis.axis=m.xAxis,m.xAxis.navigatorAxis.toFixedRange=B.AdditionsClass.prototype.toFixedRange.bind(m.xAxis.navigatorAxis));
a.options.scrollbar.enabled&&(a.scrollbar=m.scrollbar=new x(a.renderer,E(a.options.scrollbar,{margin:m.navigatorEnabled?0:10,vertical:a.inverted}),a),e(m.scrollbar,"changed",function(a){var b=m.size,c=b*this.to;b*=this.from;m.hasDragged=m.scrollbar.hasDragged;m.render(0,0,b,c);this.shouldUpdateExtremes(a.DOMType)&&setTimeout(function(){m.onMouseUp(a)})}));m.addBaseSeriesEvents();m.addChartEvents()};d.prototype.getUnionExtremes=function(a){var b=this.chart.xAxis[0],c=this.xAxis,f=c.options,d=b.options,
e;a&&null===b.dataMin||(e={dataMin:K(f&&f.min,n("min",d.min,b.dataMin,c.dataMin,c.min)),dataMax:K(f&&f.max,n("max",d.max,b.dataMax,c.dataMax,c.max))});return e};d.prototype.setBaseSeries=function(a,b){var c=this.chart,f=this.baseSeries=[];a=a||c.options&&c.options.navigator.baseSeries||(c.series.length?C(c.series,function(a){return!a.options.isInternal}).index:0);(c.series||[]).forEach(function(b,c){b.options.isInternal||!b.options.showInNavigator&&(c!==a&&b.options.id!==a||!1===b.options.showInNavigator)||
f.push(b)});this.xAxis&&!this.xAxis.navigatorAxis.fake&&this.updateNavigatorSeries(!0,b)};d.prototype.updateNavigatorSeries=function(a,b){var c=this,f=c.chart,d=c.baseSeries,e,g,k=c.navigatorOptions.series,h,l={enableMouseTracking:!1,index:null,linkedTo:null,group:"nav",padXAxis:!1,xAxis:"navigator-x-axis",yAxis:"navigator-y-axis",showInLegend:!1,stacking:void 0,isInternal:!0,states:{inactive:{opacity:1}}},n=c.series=(c.series||[]).filter(function(a){var b=a.baseSeries;return 0>d.indexOf(b)?(b&&(I(b,
"updatedData",c.updatedDataHandler),delete b.navigatorSeries),a.chart&&a.destroy(),!1):!0});d&&d.length&&d.forEach(function(a){var q=a.navigatorSeries,p=m({color:a.color,visible:a.visible},J(k)?H.navigator.series:k);q&&!1===c.navigatorOptions.adaptToUpdatedData||(l.name="Navigator "+d.length,e=a.options||{},h=e.navigatorOptions||{},p.dataLabels=z(p.dataLabels),g=E(e,l,p,h),g.pointRange=K(p.pointRange,h.pointRange,H.plotOptions[g.type||"line"].pointRange),p=h.data||p.data,c.hasNavigatorData=c.hasNavigatorData||
!!p,g.data=p||e.data&&e.data.slice(0),q&&q.options?q.update(g,b):(a.navigatorSeries=f.initSeries(g),a.navigatorSeries.baseSeries=a,n.push(a.navigatorSeries)))});if(k.data&&(!d||!d.length)||J(k))c.hasNavigatorData=!1,k=z(k),k.forEach(function(a,b){l.name="Navigator "+(n.length+1);g=E(H.navigator.series,{color:f.series[b]&&!f.series[b].options.isInternal&&f.series[b].color||f.options.colors[b]||f.options.colors[0]},l,a);g.data=a.data;g.data&&(c.hasNavigatorData=!0,n.push(f.initSeries(g)))});a&&this.addBaseSeriesEvents()};
d.prototype.addBaseSeriesEvents=function(){var a=this,b=a.baseSeries||[];b[0]&&b[0].xAxis&&b[0].eventsToUnbind.push(e(b[0].xAxis,"foundExtremes",this.modifyBaseAxisExtremes));b.forEach(function(b){b.eventsToUnbind.push(e(b,"show",function(){this.navigatorSeries&&this.navigatorSeries.setVisible(!0,!1)}));b.eventsToUnbind.push(e(b,"hide",function(){this.navigatorSeries&&this.navigatorSeries.setVisible(!1,!1)}));!1!==this.navigatorOptions.adaptToUpdatedData&&b.xAxis&&b.eventsToUnbind.push(e(b,"updatedData",
this.updatedDataHandler));b.eventsToUnbind.push(e(b,"remove",function(){this.navigatorSeries&&(G(a.series,this.navigatorSeries),r(this.navigatorSeries.options)&&this.navigatorSeries.remove(!1),delete this.navigatorSeries)}))},this)};d.prototype.getBaseSeriesMin=function(a){return this.baseSeries.reduce(function(a,b){return Math.min(a,b.xData?b.xData[0]:a)},a)};d.prototype.modifyNavigatorAxisExtremes=function(){var a=this.xAxis,b;"undefined"!==typeof a.getExtremes&&(!(b=this.getUnionExtremes(!0))||
b.dataMin===a.min&&b.dataMax===a.max||(a.min=b.dataMin,a.max=b.dataMax))};d.prototype.modifyBaseAxisExtremes=function(){var a=this.chart.navigator,b=this.getExtremes(),c=b.dataMin,d=b.dataMax;b=b.max-b.min;var e=a.stickToMin,g=a.stickToMax,h=K(this.options.overscroll,0),l=a.series&&a.series[0],m=!!this.setExtremes;if(!this.eventArgs||"rangeSelectorButton"!==this.eventArgs.trigger){if(e){var n=c;var q=n+b}g&&(q=d+h,e||(n=Math.max(c,q-b,a.getBaseSeriesMin(l&&l.xData?l.xData[0]:-Number.MAX_VALUE))));
m&&(e||g)&&M(n)&&(this.min=this.userMin=n,this.max=this.userMax=q)}a.stickToMin=a.stickToMax=null};d.prototype.updatedDataHandler=function(){var a=this.chart.navigator,b=this.navigatorSeries;a.stickToMax=a.reversedExtremes?0===Math.round(a.zoomedMin):Math.round(a.zoomedMax)>=Math.round(a.size);a.stickToMin=a.shouldStickToMin(this,a);b&&!a.hasNavigatorData&&(b.options.pointStart=this.xData[0],b.setData(this.options.data,!1,null,!1))};d.prototype.shouldStickToMin=function(a,b){b=b.getBaseSeriesMin(a.xData[0]);
var c=a.xAxis;a=c.max;var d=c.min;c=c.options.range;return M(a)&&M(d)?c&&0<a-b?a-b<c:d<=b:!1};d.prototype.addChartEvents=function(){this.eventsToUnbind||(this.eventsToUnbind=[]);this.eventsToUnbind.push(e(this.chart,"redraw",function(){var a=this.navigator,b=a&&(a.baseSeries&&a.baseSeries[0]&&a.baseSeries[0].xAxis||this.xAxis[0]);b&&a.render(b.min,b.max)}),e(this.chart,"getMargins",function(){var a=this.navigator,b=a.opposite?"plotTop":"marginBottom";this.inverted&&(b=a.opposite?"marginRight":"plotLeft");
this[b]=(this[b]||0)+(a.navigatorEnabled||!this.inverted?a.outlineHeight:0)+a.navigatorOptions.margin}))};d.prototype.destroy=function(){this.removeEvents();this.xAxis&&(G(this.chart.xAxis,this.xAxis),G(this.chart.axes,this.xAxis));this.yAxis&&(G(this.chart.yAxis,this.yAxis),G(this.chart.axes,this.yAxis));(this.series||[]).forEach(function(a){a.destroy&&a.destroy()});"series xAxis yAxis shades outline scrollbarTrack scrollbarRifles scrollbarGroup scrollbar navigatorGroup rendered".split(" ").forEach(function(a){this[a]&&
this[a].destroy&&this[a].destroy();this[a]=null},this);[this.handles].forEach(function(a){F(a)},this)};return d}();u.Navigator||(u.Navigator=y,B.compose(b),e(t,"beforeShowResetZoom",function(){var a=this.options,b=a.navigator,c=a.rangeSelector;if((b&&b.enabled||c&&c.enabled)&&(!D&&"x"===a.chart.zoomType||D&&"x"===a.chart.pinchType))return!1}),e(t,"beforeRender",function(){var a=this.options;if(a.navigator.enabled||a.scrollbar.enabled)this.scroller=this.navigator=new y(this)}),e(t,"afterSetChartSize",
function(){var a=this.legend,b=this.navigator;if(b){var c=a&&a.options;var d=b.xAxis;var e=b.yAxis;var g=b.scrollbarHeight;this.inverted?(b.left=b.opposite?this.chartWidth-g-b.height:this.spacing[3]+g,b.top=this.plotTop+g):(b.left=K(d.left,this.plotLeft+g),b.top=b.navigatorOptions.top||this.chartHeight-b.height-g-this.spacing[2]-(this.rangeSelector&&this.extraBottomMargin?this.rangeSelector.getHeight():0)-(c&&"bottom"===c.verticalAlign&&"proximate"!==c.layout&&c.enabled&&!c.floating?a.legendHeight+
K(c.margin,10):0)-(this.titleOffset?this.titleOffset[2]:0));d&&e&&(this.inverted?d.options.left=e.options.left=b.left:d.options.top=e.options.top=b.top,d.setAxisSize(),e.setAxisSize())}}),e(t,"update",function(a){var b=a.options.navigator||{},c=a.options.scrollbar||{};this.navigator||this.scroller||!b.enabled&&!c.enabled||(E(!0,this.options.navigator,b),E(!0,this.options.scrollbar,c),delete a.options.navigator,delete a.options.scrollbar)}),e(t,"afterUpdate",function(a){this.navigator||this.scroller||
!this.options.navigator.enabled&&!this.options.scrollbar.enabled||(this.scroller=this.navigator=new y(this),K(a.redraw,!0)&&this.redraw(a.animation))}),e(t,"afterAddSeries",function(){this.navigator&&this.navigator.setBaseSeries(null,!1)}),e(d,"afterUpdate",function(){this.chart.navigator&&!this.options.isInternal&&this.chart.navigator.setBaseSeries(null,!1)}),t.prototype.callbacks.push(function(a){var b=a.navigator;b&&a.xAxis[0]&&(a=a.xAxis[0].getExtremes(),b.render(a.min,a.max))}));u.Navigator=
y;return u.Navigator});J(b,"masters/modules/gantt.src.js",[b["Core/Globals.js"],b["Core/Chart/GanttChart.js"],b["Core/Scrollbar.js"]],function(b,t,v){b.Scrollbar=v;b.GanttChart=t;b.ganttChart=t.ganttChart;v.compose(b.Axis)})});
//# sourceMappingURL=gantt.js.map
```
