## Supplementary material for "Exploring natural odour landscapes: A case study with implications for human-biting insects": Figure S4: grid-axis.js

```
/*
 Highcharts Gantt JS v9.3.1 (2021-11-05)

 GridAxis

 (c) 2016-2021 Lars A. V. Cabrera

 License: www.highcharts.com/license
*/
'use strict';(function(g){"object"===typeof module&&module.exports?(g["default"]=g,module.exports=g):"function"===typeof define&&define.amd?define("highcharts/modules/grid-axis",["highcharts"],function(n){g(n);g.Highcharts=n;return g}):g("undefined"!==typeof Highcharts?Highcharts:void 0)})(function(g){function n(g,m,n,l){g.hasOwnProperty(m)||(g[m]=l.apply(null,n))}g=g?g._modules:{};n(g,"Core/Axis/GridAxis.js",[g["Core/Axis/Axis.js"],g["Core/Axis/AxisDefaults.js"],g["Core/Globals.js"],g["Core/Utilities.js"]],
function(g,n,z,l){var m=z.dateFormats,e=l.addEvent,p=l.defined,y=l.erase,C=l.find,D=l.isArray,u=l.isNumber,v=l.merge,x=l.pick,E=l.timeUnits,F=l.wrap,w;(function(q){function m(a,b){var c={width:0,height:0};b.forEach(function(b){b=a[b];if(l.isObject(b,!0)){var d=l.isObject(b.label,!0)?b.label:{};b=d.getBBox?d.getBBox().height:0;d.textStr&&!u(d.textPxLength)&&(d.textPxLength=d.getBBox().width);var f=u(d.textPxLength)?Math.round(d.textPxLength):0;d.textStr&&(f=Math.round(d.getBBox().width));c.height=
Math.max(b,c.height);c.width=Math.max(f,c.width)}});"treegrid"===this.options.type&&this.treeGrid&&this.treeGrid.mapOfPosToGridNode&&(c.width+=this.options.labels.indentation*((this.treeGrid.mapOfPosToGridNode[-1].height||0)-1));return c}function G(){var a=this.grid;(a&&a.columns||[]).forEach(function(b){b.getOffset()})}function w(a){if(!0===(this.options.grid||{}).enabled){var b=this.axisTitle,c=this.height,f=this.horiz,d=this.left,h=this.offset,k=this.opposite,t=this.options,A=this.top,g=this.width,
r=this.tickSize(),l=b&&b.getBBox().width,e=t.title.x,n=t.title.y,m=x(t.title.margin,f?5:10);b=this.chart.renderer.fontMetrics(t.title.style.fontSize,b).f;r=(f?A+c:d)+(f?1:-1)*(k?-1:1)*(r?r[0]/2:0)+(this.side===q.Side.bottom?b:0);a.titlePosition.x=f?d-(l||0)/2-m+e:r+(k?g:0)+h+e;a.titlePosition.y=f?r-(k?c:0)+(k?b:-b)/2+h+n:A-m+n}}function H(){var a=this.chart,b=this.options.grid;b=void 0===b?{}:b;var c=this.userOptions;if(b.enabled){var f=this.options;f.labels.align=x(f.labels.align,"center");this.categories||
(f.showLastLabel=!1);this.labelRotation=0;f.labels.rotation=0}if(b.columns){f=this.grid.columns=[];for(var d=this.grid.columnIndex=0;++d<b.columns.length;){var h=v(c,b.columns[b.columns.length-d-1],{linkedTo:0,type:"category",scrollbar:{enabled:!1}});delete h.grid.columns;h=new g(this.chart,h);h.grid.isColumn=!0;h.grid.columnIndex=d;y(a.axes,h);y(a[this.coll],h);f.push(h)}}}function I(){var a=this.grid,b=this.options;if(!0===(b.grid||{}).enabled){var c=this.min||0,f=this.max||0;this.maxLabelDimensions=
this.getMaxLabelDimensions(this.ticks,this.tickPositions);this.rightWall&&this.rightWall.destroy();if(this.grid&&this.grid.isOuterAxis()&&this.axisLine){var d=b.lineWidth;if(d){d=this.getLinePath(d);var h=d[0],k=d[1],t=((this.tickSize("tick")||[1])[0]-1)*(this.side===q.Side.top||this.side===q.Side.left?-1:1);"M"===h[0]&&"L"===k[0]&&(this.horiz?(h[2]+=t,k[2]+=t):(h[1]+=t,k[1]+=t));!this.horiz&&this.chart.marginRight&&(h=[h,["L",this.left,h[2]||0]],t=["L",this.chart.chartWidth-this.chart.marginRight,
this.toPixels(f+this.tickmarkOffset)],k=[["M",k[1]||0,this.toPixels(f+this.tickmarkOffset)],t],this.grid.upperBorder||0===c%1||(this.grid.upperBorder=this.grid.renderBorder(h)),this.grid.upperBorder&&(this.grid.upperBorder.attr({stroke:b.lineColor,"stroke-width":b.lineWidth}),this.grid.upperBorder.animate({d:h})),this.grid.lowerBorder||0===f%1||(this.grid.lowerBorder=this.grid.renderBorder(k)),this.grid.lowerBorder&&(this.grid.lowerBorder.attr({stroke:b.lineColor,"stroke-width":b.lineWidth}),this.grid.lowerBorder.animate({d:k})));
this.grid.axisLineExtra?(this.grid.axisLineExtra.attr({stroke:b.lineColor,"stroke-width":b.lineWidth}),this.grid.axisLineExtra.animate({d:d})):this.grid.axisLineExtra=this.grid.renderBorder(d);this.axisLine[this.showAxis?"show":"hide"](!0)}}(a&&a.columns||[]).forEach(function(b){b.render()});if(!this.horiz&&this.chart.hasRendered&&(this.scrollbar||this.linkedParent&&this.linkedParent.scrollbar)){a=this.tickmarkOffset;b=this.tickPositions[this.tickPositions.length-1];d=this.tickPositions[0];for(k=
void 0;(k=this.hiddenLabels.pop())&&k.element;)k.show();(k=this.ticks[d].label)&&(c-d>a?this.hiddenLabels.push(k.hide()):k.show());(k=this.ticks[b].label)&&(b-f>a?this.hiddenLabels.push(k.hide()):k.show());(c=this.ticks[b].mark)&&(b-f<a&&0<b-f&&this.ticks[b].isLast?c.hide():this.ticks[b-1]&&c.show())}}}function J(){var a=this.tickPositions&&this.tickPositions.info,b=this.options,c=this.userOptions.labels||{};(b.grid||{}).enabled&&(this.horiz?(this.series.forEach(function(b){b.options.pointRange=0}),
a&&b.dateTimeLabelFormats&&b.labels&&!p(c.align)&&(!1===b.dateTimeLabelFormats[a.unitName].range||1<a.count)&&(b.labels.align="left",p(c.x)||(b.labels.x=3))):"treegrid"!==this.options.type&&this.grid&&this.grid.columns&&(this.minPointOffset=this.tickInterval))}function K(a){var b=this.options;a=a.userOptions;var c=b&&l.isObject(b.grid,!0)?b.grid:{};if(!0===c.enabled){var f=v(!0,{className:"highcharts-grid-axis "+(a.className||""),dateTimeLabelFormats:{hour:{list:["%H:%M","%H"]},day:{list:["%A, %e. %B",
"%a, %e. %b","%E"]},week:{list:["Week %W","W%W"]},month:{list:["%B","%b","%o"]}},grid:{borderWidth:1},labels:{padding:2,style:{fontSize:"13px"}},margin:0,title:{text:null,reserveSpace:!1,rotation:0},units:[["millisecond",[1,10,100]],["second",[1,10]],["minute",[1,5,15]],["hour",[1,6]],["day",[1]],["week",[1]],["month",[1]],["year",null]]},a);"xAxis"===this.coll&&(p(a.linkedTo)&&!p(a.tickPixelInterval)&&(f.tickPixelInterval=350),p(a.tickPixelInterval)||!p(a.linkedTo)||p(a.tickPositioner)||p(a.tickInterval)||
(f.tickPositioner=function(b,a){var c=this.linkedParent&&this.linkedParent.tickPositions&&this.linkedParent.tickPositions.info;if(c){for(var d=f.units||[],h=void 0,g=void 0,e=void 0,q=0;q<d.length;q++)if(d[q][0]===c.unitName){h=q;break}d[h+1]?(e=d[h+1][0],g=(d[h+1][1]||[1])[0]):"year"===c.unitName&&(e="year",g=10*c.count);c=E[e];this.tickInterval=c*g;return this.getTimeTicks({unitRange:c,count:g,unitName:e},b,a,this.options.startOfWeek)}}));v(!0,this.options,f);this.horiz&&(b.minPadding=x(a.minPadding,
0),b.maxPadding=x(a.maxPadding,0));u(b.grid.borderWidth)&&(b.tickWidth=b.lineWidth=c.borderWidth)}}function L(a){a=(a=a.userOptions)&&a.grid||{};var b=a.columns;a.enabled&&b&&v(!0,this.options,b[b.length-1])}function M(){(this.grid.columns||[]).forEach(function(a){a.setScale()})}function N(a){var b=n.defaultLeftAxisOptions,c=this.horiz,f=this.maxLabelDimensions,d=this.options.grid;d=void 0===d?{}:d;d.enabled&&f&&(b=2*Math.abs(b.labels.x),c=c?d.cellHeight||b+f.height:b+f.width,D(a.tickSize)?a.tickSize[0]=
c:a.tickSize=[c,0])}function O(){this.axes.forEach(function(a){(a.grid&&a.grid.columns||[]).forEach(function(b){b.setAxisSize();b.setAxisTranslation()})})}function P(a){var b=this.grid;(b.columns||[]).forEach(function(b){b.destroy(a.keepEvents)});b.columns=void 0}function Q(a){a=a.userOptions||{};var b=a.grid||{};b.enabled&&p(b.borderColor)&&(a.tickColor=a.lineColor=b.borderColor);this.grid||(this.grid=new B(this));this.hiddenLabels=[]}function R(a){var b=this.label,c=this.axis,f=c.reversed,d=c.chart,
h=c.options.grid||{},k=c.options.labels,g=k.align,e=q.Side[c.side],l=a.tickmarkOffset,r=c.tickPositions,n=this.pos-l;r=u(r[a.index+1])?r[a.index+1]-l:(c.max||0)+l;var m=c.tickSize("tick");l=m?m[0]:0;m=m?m[1]/2:0;if(!0===h.enabled){if("top"===e){h=c.top+c.offset;var p=h-l}else"bottom"===e?(p=d.chartHeight-c.bottom+c.offset,h=p+l):(h=c.top+c.len-(c.translate(f?r:n)||0),p=c.top+c.len-(c.translate(f?n:r)||0));"right"===e?(e=d.chartWidth-c.right+c.offset,f=e+l):"left"===e?(f=c.left+c.offset,e=f-l):(e=
Math.round(c.left+(c.translate(f?r:n)||0))-m,f=Math.min(Math.round(c.left+(c.translate(f?n:r)||0))-m,c.left+c.len));this.slotWidth=f-e;a.pos.x="left"===g?e:"right"===g?f:e+(f-e)/2;a.pos.y=p+(h-p)/2;d=d.renderer.fontMetrics(k.style.fontSize,b&&b.element);b=b?b.getBBox().height:0;k.useHTML?a.pos.y+=d.b+-(b/2):(b=Math.round(b/d.h),a.pos.y+=(d.b-(d.h-d.f))/2+-((b-1)*d.h/2));a.pos.x+=c.horiz&&k.x||0}}function S(a){var b=a.axis,c=a.value;if(b.options.grid&&b.options.grid.enabled){var f=b.tickPositions,
d=(b.linkedParent||b).series[0],h=c===f[0];f=c===f[f.length-1];var k=d&&C(d.options.data,function(a){return a[b.isXAxis?"x":"y"]===c}),e=void 0;k&&d.is("gantt")&&(e=v(k),z.seriesTypes.gantt.prototype.pointClass.setGanttPointAliases(e));a.isFirst=h;a.isLast=f;a.point=e}}function T(){var a=this.options,b=this.categories,c=this.tickPositions,f=c[0],d=c[c.length-1],h=this.linkedParent&&this.linkedParent.min||this.min,e=this.linkedParent&&this.linkedParent.max||this.max,g=this.tickInterval;!0!==(a.grid||
{}).enabled||b||!this.horiz&&!this.isLinked||(f<h&&f+g>h&&!a.startOnTick&&(c[0]=h),d>e&&d-g<e&&!a.endOnTick&&(c[c.length-1]=e))}function U(a){var b=this.options.grid;return!0===(void 0===b?{}:b).enabled&&this.categories?this.tickInterval:a.apply(this,Array.prototype.slice.call(arguments,1))}(function(a){a[a.top=0]="top";a[a.right=1]="right";a[a.bottom=2]="bottom";a[a.left=3]="left"})(q.Side||(q.Side={}));q.compose=function(a,b,c){-1===a.keepProps.indexOf("grid")&&(a.keepProps.push("grid"),a.prototype.getMaxLabelDimensions=
m,F(a.prototype,"unsquish",U),e(a,"init",Q),e(a,"afterGetOffset",G),e(a,"afterGetTitlePosition",w),e(a,"afterInit",H),e(a,"afterRender",I),e(a,"afterSetAxisTranslation",J),e(a,"afterSetOptions",K),e(a,"afterSetOptions",L),e(a,"afterSetScale",M),e(a,"afterTickSize",N),e(a,"trimTicks",T),e(a,"destroy",P));e(b,"afterSetChartSize",O);e(c,"afterGetLabelPosition",R);e(c,"labelFormat",S);return a};var B=function(){function a(b){this.axis=b}a.prototype.isOuterAxis=function(){var b=this.axis,a=b.grid.columnIndex,
f=b.linkedParent&&b.linkedParent.grid.columns||b.grid.columns,d=a?b.linkedParent:b,e=-1,g=0;b.chart[b.coll].forEach(function(a,c){a.side!==b.side||a.options.isInternal||(g=c,a===d&&(e=c))});return g===e&&(u(a)?f.length===a:!0)};a.prototype.renderBorder=function(b){var a=this.axis,f=a.chart.renderer,d=a.options;b=f.path(b).addClass("highcharts-axis-line").add(a.axisBorder);f.styledMode||b.attr({stroke:d.lineColor,"stroke-width":d.lineWidth,zIndex:7});return b};return a}();q.Additions=B})(w||(w={}));
m.E=function(e){return this.dateFormat("%a",e,!0).charAt(0)};m.W=function(e){e=new this.Date(e);var g=(this.get("Day",e)+6)%7,l=new this.Date(e.valueOf());this.set("Date",l,this.get("Date",e)-g+3);g=new this.Date(this.get("FullYear",l),0,1);4!==this.get("Day",g)&&(this.set("Month",e,0),this.set("Date",e,1+(11-this.get("Day",g))%7));return(1+Math.floor((l.valueOf()-g.valueOf())/6048E5)).toString()};"";return w});n(g,"masters/modules/grid-axis.src.js",[g["Core/Globals.js"],g["Core/Axis/GridAxis.js"]],
function(g,m){m.compose(g.Axis,g.Chart,g.Tick)})});
//# sourceMappingURL=grid-axis.js.map
```
