## Supplementary material for "Exploring natural odour landscapes: A case study with implications for human-biting insects": Figure S4: heatmap.js

```
/*
 Highmaps JS v9.3.1 (2021-11-05)

 (c) 2009-2021 Torstein Honsi

 License: www.highcharts.com/license
*/
'use strict';(function(b){"object"===typeof module&&module.exports?(b["default"]=b,module.exports=b):"function"===typeof define&&define.amd?define("highcharts/modules/heatmap",["highcharts"],function(u){b(u);b.Highcharts=u;return b}):b("undefined"!==typeof Highcharts?Highcharts:void 0)})(function(b){function u(b,l,G,r){b.hasOwnProperty(l)||(b[l]=r.apply(null,G))}b=b?b._modules:{};u(b,"Core/Axis/Color/ColorAxisComposition.js",[b["Core/Color/Color.js"],b["Core/Utilities.js"]],function(b,l){var p=b.parse,
r=l.addEvent,q=l.extend,y=l.merge,x=l.pick,z=l.splat,e;(function(e){function b(){var a=this,d=this.options;this.colorAxis=[];d.colorAxis&&(d.colorAxis=z(d.colorAxis),d.colorAxis.forEach(function(d,f){d.index=f;new g(a,d)}))}function k(a){var d=this,f=function(g){g=a.allItems.indexOf(g);-1!==g&&(d.destroyItem(a.allItems[g]),a.allItems.splice(g,1))},g=[],c,m;(this.chart.colorAxis||[]).forEach(function(a){(c=a.options)&&c.showInLegend&&(c.dataClasses&&c.visible?g=g.concat(a.getDataClassLegendSymbols()):
c.visible&&g.push(a),a.series.forEach(function(a){if(!a.options.showInLegend||c.dataClasses)"point"===a.options.legendType?a.points.forEach(function(a){f(a)}):f(a)}))});for(m=g.length;m--;)a.allItems.unshift(g[m])}function l(a){a.visible&&a.item.legendColor&&a.item.legendSymbol.attr({fill:a.item.legendColor})}function v(){var a=this.chart.colorAxis;a&&a.forEach(function(a,g,d){a.update({},d)})}function C(){(this.chart.colorAxis&&this.chart.colorAxis.length||this.colorAttribs)&&this.translateColors()}
function w(){var a=this.axisTypes;a?-1===a.indexOf("colorAxis")&&a.push("colorAxis"):this.axisTypes=["colorAxis"]}function t(a){var g=this,d=a?"show":"hide";g.visible=g.options.visible=!!a;["graphic","dataLabel"].forEach(function(a){if(g[a])g[a][d]()});this.series.buildKDTree()}function h(){var a=this,g=this.options.nullColor,d=this.colorAxis,f=this.colorKey;(this.data.length?this.data:this.points).forEach(function(c){var m=c.getNestedProperty(f);(m=c.options.color||(c.isNull||null===c.value?g:d&&
"undefined"!==typeof m?d.toColor(m,c):c.color||a.color))&&c.color!==m&&(c.color=m,"point"===a.options.legendType&&c.legendItem&&a.chart.legend.colorizeItem(c,c.visible))})}function c(a){var d=a.prototype.createAxis;a.prototype.createAxis=function(a,c){if("colorAxis"!==a)return d.apply(this,arguments);var f=new g(this,y(c.axis,{index:this[a].length,isX:!1}));this.isDirtyLegend=!0;this.axes.forEach(function(a){a.series=[]});this.series.forEach(function(a){a.bindAxes();a.isDirtyData=!0});x(c.redraw,
!0)&&this.redraw(c.animation);return f}}function d(){this.elem.attr("fill",p(this.start).tweenTo(p(this.end),this.pos),void 0,!0)}function f(){this.elem.attr("stroke",p(this.start).tweenTo(p(this.end),this.pos),void 0,!0)}var a=[],g;e.compose=function(m,A,e,B,D){g||(g=m);-1===a.indexOf(A)&&(a.push(A),m=A.prototype,m.collectionsWithUpdate.push("colorAxis"),m.collectionsWithInit.colorAxis=[m.addColorAxis],r(A,"afterGetAxes",b),c(A));-1===a.indexOf(e)&&(a.push(e),A=e.prototype,A.fillSetter=d,A.strokeSetter=
f);-1===a.indexOf(B)&&(a.push(B),r(B,"afterGetAllItems",k),r(B,"afterColorizeItem",l),r(B,"afterUpdate",v));-1===a.indexOf(D)&&(a.push(D),q(D.prototype,{optionalAxis:"colorAxis",translateColors:h}),q(D.prototype.pointClass.prototype,{setVisible:t}),r(D,"afterTranslate",C),r(D,"bindAxes",w))};e.pointSetVisible=t})(e||(e={}));return e});u(b,"Core/Axis/Color/ColorAxisDefaults.js",[],function(){return{lineWidth:0,minPadding:0,maxPadding:0,gridLineWidth:1,tickPixelInterval:72,startOnTick:!0,endOnTick:!0,
offset:0,marker:{animation:{duration:50},width:.01,color:"#999999"},labels:{overflow:"justify",rotation:0},minColor:"#e6ebf5",maxColor:"#003399",tickLength:5,showInLegend:!0}});u(b,"Core/Axis/Color/ColorAxis.js",[b["Core/Axis/Axis.js"],b["Core/Color/Color.js"],b["Core/Axis/Color/ColorAxisComposition.js"],b["Core/Axis/Color/ColorAxisDefaults.js"],b["Core/Globals.js"],b["Core/Legend/LegendSymbol.js"],b["Core/Series/SeriesRegistry.js"],b["Core/Utilities.js"]],function(b,l,u,r,q,y,x,z){var e=this&&this.__extends||
function(){var e=function(h,c){e=Object.setPrototypeOf||{__proto__:[]}instanceof Array&&function(c,f){c.__proto__=f}||function(c,f){for(var a in f)f.hasOwnProperty(a)&&(c[a]=f[a])};return e(h,c)};return function(h,c){function d(){this.constructor=h}e(h,c);h.prototype=null===c?Object.create(c):(d.prototype=c.prototype,new d)}}(),k=l.parse,E=q.noop,n=x.series,p=z.extend,v=z.isNumber,C=z.merge,w=z.pick;l=function(b){function h(c,d){var f=b.call(this,c,d)||this;f.beforePadding=!1;f.chart=void 0;f.coll=
"colorAxis";f.dataClasses=void 0;f.legendItem=void 0;f.legendItems=void 0;f.name="";f.options=void 0;f.stops=void 0;f.visible=!0;f.init(c,d);return f}e(h,b);h.compose=function(c,d,f,a){u.compose(h,c,d,f,a)};h.prototype.init=function(c,d){var f=c.options.legend||{},a=d.layout?"vertical"!==d.layout:"vertical"!==f.layout,g=d.visible;f=C(h.defaultColorAxisOptions,d,{showEmpty:!1,title:null,visible:f.enabled&&!1!==g});this.coll="colorAxis";this.side=d.side||a?2:1;this.reversed=d.reversed||!a;this.opposite=
!a;b.prototype.init.call(this,c,f);this.userOptions.visible=g;d.dataClasses&&this.initDataClasses(d);this.initStops();this.horiz=a;this.zoomEnabled=!1};h.prototype.initDataClasses=function(c){var d=this.chart,f=this.options,a=c.dataClasses.length,g,m=0,b=d.options.chart.colorCount;this.dataClasses=g=[];this.legendItems=[];(c.dataClasses||[]).forEach(function(c,e){c=C(c);g.push(c);if(d.styledMode||!c.color)"category"===f.dataClassColor?(d.styledMode||(e=d.options.colors,b=e.length,c.color=e[m]),c.colorIndex=
m,m++,m===b&&(m=0)):c.color=k(f.minColor).tweenTo(k(f.maxColor),2>a?.5:e/(a-1))})};h.prototype.hasData=function(){return!!(this.tickPositions||[]).length};h.prototype.setTickPositions=function(){if(!this.dataClasses)return b.prototype.setTickPositions.call(this)};h.prototype.initStops=function(){this.stops=this.options.stops||[[0,this.options.minColor],[1,this.options.maxColor]];this.stops.forEach(function(c){c.color=k(c[1])})};h.prototype.setOptions=function(c){b.prototype.setOptions.call(this,c);
this.options.crosshair=this.options.marker};h.prototype.setAxisSize=function(){var c=this.legendSymbol,d=this.chart,f=d.options.legend||{},a,g;c?(this.left=f=c.attr("x"),this.top=a=c.attr("y"),this.width=g=c.attr("width"),this.height=c=c.attr("height"),this.right=d.chartWidth-f-g,this.bottom=d.chartHeight-a-c,this.len=this.horiz?g:c,this.pos=this.horiz?f:a):this.len=(this.horiz?f.symbolWidth:f.symbolHeight)||h.defaultLegendLength};h.prototype.normalizedValue=function(c){this.logarithmic&&(c=this.logarithmic.log2lin(c));
return 1-(this.max-c)/(this.max-this.min||1)};h.prototype.toColor=function(c,d){var f=this.dataClasses,a=this.stops,g;if(f)for(g=f.length;g--;){var e=f[g];var b=e.from;a=e.to;if(("undefined"===typeof b||c>=b)&&("undefined"===typeof a||c<=a)){var h=e.color;d&&(d.dataClass=g,d.colorIndex=e.colorIndex);break}}else{c=this.normalizedValue(c);for(g=a.length;g--&&!(c>a[g][0]););b=a[g]||a[g+1];a=a[g+1]||b;c=1-(a[0]-c)/(a[0]-b[0]||1);h=b.color.tweenTo(a.color,c)}return h};h.prototype.getOffset=function(){var c=
this.legendGroup,d=this.chart.axisOffset[this.side];c&&(this.axisParent=c,b.prototype.getOffset.call(this),this.added||(this.added=!0,this.labelLeft=0,this.labelRight=this.width),this.chart.axisOffset[this.side]=d)};h.prototype.setLegendColor=function(){var c=this.reversed,d=c?1:0;c=c?0:1;d=this.horiz?[d,0,c,0]:[0,c,0,d];this.legendColor={linearGradient:{x1:d[0],y1:d[1],x2:d[2],y2:d[3]},stops:this.stops}};h.prototype.drawLegendSymbol=function(c,d){var f=c.padding,a=c.options,g=this.horiz,b=w(a.symbolWidth,
g?h.defaultLegendLength:12),e=w(a.symbolHeight,g?12:h.defaultLegendLength),k=w(a.labelPadding,g?16:30);a=w(a.itemDistance,10);this.setLegendColor();d.legendSymbol=this.chart.renderer.rect(0,c.baseline-11,b,e).attr({zIndex:1}).add(d.legendGroup);this.legendItemWidth=b+f+(g?a:k);this.legendItemHeight=e+f+(g?k:0)};h.prototype.setState=function(c){this.series.forEach(function(d){d.setState(c)})};h.prototype.setVisible=function(){};h.prototype.getSeriesExtremes=function(){var c=this.series,d=c.length,
f;this.dataMin=Infinity;for(this.dataMax=-Infinity;d--;){var a=c[d];var g=a.colorKey=w(a.options.colorKey,a.colorKey,a.pointValKey,a.zoneAxis,"y");var b=a.pointArrayMap;var e=a[g+"Min"]&&a[g+"Max"];if(a[g+"Data"])var h=a[g+"Data"];else if(b){h=[];b=b.indexOf(g);var k=a.yData;if(0<=b&&k)for(f=0;f<k.length;f++)h.push(w(k[f][b],k[f]))}else h=a.yData;e?(a.minColorValue=a[g+"Min"],a.maxColorValue=a[g+"Max"]):(h=n.prototype.getExtremes.call(a,h),a.minColorValue=h.dataMin,a.maxColorValue=h.dataMax);"undefined"!==
typeof a.minColorValue&&(this.dataMin=Math.min(this.dataMin,a.minColorValue),this.dataMax=Math.max(this.dataMax,a.maxColorValue));e||n.prototype.applyExtremes.call(a)}};h.prototype.drawCrosshair=function(c,d){var f=d&&d.plotX,a=d&&d.plotY,g=this.pos,e=this.len;if(d){var h=this.toPixels(d.getNestedProperty(d.series.colorKey));h<g?h=g-2:h>g+e&&(h=g+e+2);d.plotX=h;d.plotY=this.len-h;b.prototype.drawCrosshair.call(this,c,d);d.plotX=f;d.plotY=a;this.cross&&!this.cross.addedToColorAxis&&this.legendGroup&&
(this.cross.addClass("highcharts-coloraxis-marker").add(this.legendGroup),this.cross.addedToColorAxis=!0,this.chart.styledMode||"object"!==typeof this.crosshair||this.cross.attr({fill:this.crosshair.color}))}};h.prototype.getPlotLinePath=function(c){var d=this.left,f=c.translatedValue,a=this.top;return v(f)?this.horiz?[["M",f-4,a-6],["L",f+4,a-6],["L",f,a],["Z"]]:[["M",d,f],["L",d-6,f+6],["L",d-6,f-6],["Z"]]:b.prototype.getPlotLinePath.call(this,c)};h.prototype.update=function(c,d){var f=this.chart.legend;
this.series.forEach(function(a){a.isDirtyData=!0});(c.dataClasses&&f.allItems||this.dataClasses)&&this.destroyItems();b.prototype.update.call(this,c,d);this.legendItem&&(this.setLegendColor(),f.colorizeItem(this,!0))};h.prototype.destroyItems=function(){var c=this.chart;this.legendItem?c.legend.destroyItem(this):this.legendItems&&this.legendItems.forEach(function(d){c.legend.destroyItem(d)});c.isDirtyLegend=!0};h.prototype.destroy=function(){this.chart.isDirtyLegend=!0;this.destroyItems();b.prototype.destroy.apply(this,
[].slice.call(arguments))};h.prototype.remove=function(c){this.destroyItems();b.prototype.remove.call(this,c)};h.prototype.getDataClassLegendSymbols=function(){var c=this,d=c.chart,f=c.legendItems,a=d.options.legend,g=a.valueDecimals,b=a.valueSuffix||"",e;f.length||c.dataClasses.forEach(function(a,h){var k=a.from,m=a.to,B=d.numberFormatter,n=!0;e="";"undefined"===typeof k?e="< ":"undefined"===typeof m&&(e="> ");"undefined"!==typeof k&&(e+=B(k,g)+b);"undefined"!==typeof k&&"undefined"!==typeof m&&
(e+=" - ");"undefined"!==typeof m&&(e+=B(m,g)+b);f.push(p({chart:d,name:e,options:{},drawLegendSymbol:y.drawRectangle,visible:!0,setState:E,isDataClass:!0,setVisible:function(){n=c.visible=!n;c.series.forEach(function(a){a.points.forEach(function(a){a.dataClass===h&&a.setVisible(n)})});d.legend.colorizeItem(this,n)}},a))});return f};h.defaultColorAxisOptions=r;h.defaultLegendLength=200;h.keepProps=["legendGroup","legendItemHeight","legendItemWidth","legendItem","legendSymbol"];return h}(b);Array.prototype.push.apply(b.keepProps,
l.keepProps);"";return l});u(b,"Series/ColorMapComposition.js",[b["Core/Utilities.js"]],function(b){var l=b.defined,p=b.wrap,r={pointArrayMap:["value"],axisTypes:["xAxis","yAxis","colorAxis"],trackerGroups:["group","markerGroup","dataLabelsGroup"],parallelArrays:["x","y","value"],colorKey:"value"},q;(function(b){function x(){return null!==this.value&&Infinity!==this.value&&-Infinity!==this.value}function z(b){var e={};!l(b.color)||b.state&&"normal"!==b.state||(e[this.colorProp||"fill"]=b.color);return e}
function e(b,e,k){b=b.call(this,e,k);e.moveToTopOnHover&&(b.zIndex="hover"===k?1:0);return b}b.colorMapSeriesMixin=r;var k=[];b.compose=function(b,n){n&&-1===k.indexOf(n)&&(k.push(n),n=n.prototype,n.dataLabelOnNull=!0,n.moveToTopOnHover=!0,n.isValid=x);-1===k.indexOf(b)&&(k.push(b),n=b.prototype,n.colorAttribs=z,p(n,"pointAttribs",e));return b};b.seriesColorAttribs=z})(q||(q={}));return q});u(b,"Series/Heatmap/HeatmapPoint.js",[b["Core/Series/SeriesRegistry.js"],b["Core/Utilities.js"]],function(b,
l){var p=this&&this.__extends||function(){var b=function(l,e){b=Object.setPrototypeOf||{__proto__:[]}instanceof Array&&function(b,e){b.__proto__=e}||function(b,e){for(var k in e)e.hasOwnProperty(k)&&(b[k]=e[k])};return b(l,e)};return function(l,e){function k(){this.constructor=l}b(l,e);l.prototype=null===e?Object.create(e):(k.prototype=e.prototype,new k)}}(),r=l.clamp,q=l.extend,u=l.pick;b=function(b){function l(){var e=null!==b&&b.apply(this,arguments)||this;e.options=void 0;e.series=void 0;e.value=
void 0;e.x=void 0;e.y=void 0;return e}p(l,b);l.prototype.applyOptions=function(e,k){e=b.prototype.applyOptions.call(this,e,k);e.formatPrefix=e.isNull||null===e.value?"null":"point";return e};l.prototype.getCellAttributes=function(){var b=this.series,k=b.options,l=(k.colsize||1)/2,n=(k.rowsize||1)/2,p=b.xAxis,v=b.yAxis,q=this.options.marker||b.options.marker;b=b.pointPlacementToXValue();var w=u(this.pointPadding,k.pointPadding,0),t={x1:r(Math.round(p.len-(p.translate(this.x-l,!1,!0,!1,!0,-b)||0)),
-p.len,2*p.len),x2:r(Math.round(p.len-(p.translate(this.x+l,!1,!0,!1,!0,-b)||0)),-p.len,2*p.len),y1:r(Math.round(v.translate(this.y-n,!1,!0,!1,!0)||0),-v.len,2*v.len),y2:r(Math.round(v.translate(this.y+n,!1,!0,!1,!0)||0),-v.len,2*v.len)};[["width","x"],["height","y"]].forEach(function(b){var c=b[0];b=b[1];var d=b+"1",f=b+"2",a=Math.abs(t[d]-t[f]),g=q&&q.lineWidth||0,e=Math.abs(t[d]+t[f])/2;q[c]&&q[c]<a&&(t[d]=e-q[c]/2-g/2,t[f]=e+q[c]/2+g/2);w&&("y"===b&&(d=f,f=b+"1"),t[d]+=w,t[f]-=w)});return t};
l.prototype.haloPath=function(b){if(!b)return[];var e=this.shapeArgs;return["M",e.x-b,e.y-b,"L",e.x-b,e.y+e.height+b,e.x+e.width+b,e.y+e.height+b,e.x+e.width+b,e.y-b,"Z"]};l.prototype.isValid=function(){return Infinity!==this.value&&-Infinity!==this.value};return l}(b.seriesTypes.scatter.prototype.pointClass);q(b.prototype,{dataLabelOnNull:!0,moveToTopOnHover:!0,ttBelow:!1});return b});u(b,"Series/Heatmap/HeatmapSeries.js",[b["Core/Color/Color.js"],b["Series/ColorMapComposition.js"],b["Series/Heatmap/HeatmapPoint.js"],
b["Core/Legend/LegendSymbol.js"],b["Core/Series/SeriesRegistry.js"],b["Core/Renderer/SVG/SVGRenderer.js"],b["Core/Utilities.js"]],function(b,l,u,r,q,y,x){var p=this&&this.__extends||function(){var b=function(c,a){b=Object.setPrototypeOf||{__proto__:[]}instanceof Array&&function(a,b){a.__proto__=b}||function(a,b){for(var c in b)b.hasOwnProperty(c)&&(a[c]=b[c])};return b(c,a)};return function(c,a){function g(){this.constructor=c}b(c,a);c.prototype=null===a?Object.create(a):(g.prototype=a.prototype,
new g)}}(),e=l.colorMapSeriesMixin,k=q.series,E=q.seriesTypes,n=E.column,F=E.scatter,v=y.prototype.symbols,C=x.extend,w=x.fireEvent,t=x.isNumber,h=x.merge,c=x.pick;y=function(d){function e(){var a=null!==d&&d.apply(this,arguments)||this;a.colorAxis=void 0;a.data=void 0;a.options=void 0;a.points=void 0;a.valueMax=NaN;a.valueMin=NaN;return a}p(e,d);e.prototype.drawPoints=function(){var a=this;if((this.options.marker||{}).enabled||this._hasPointMarkers)k.prototype.drawPoints.call(this),this.points.forEach(function(b){b.graphic&&
(b.graphic[a.chart.styledMode?"css":"animate"](a.colorAttribs(b)),a.options.borderRadius&&b.graphic.attr({r:a.options.borderRadius}),(b.shapeArgs||{}).r=a.options.borderRadius,(b.shapeArgs||{}).d=b.graphic.pathArray,null===b.value&&b.graphic.addClass("highcharts-null-point"))})};e.prototype.getExtremes=function(){var a=k.prototype.getExtremes.call(this,this.valueData),b=a.dataMin;a=a.dataMax;t(b)&&(this.valueMin=b);t(a)&&(this.valueMax=a);return k.prototype.getExtremes.call(this)};e.prototype.getValidPoints=
function(a,b){return k.prototype.getValidPoints.call(this,a,b,!0)};e.prototype.hasData=function(){return!!this.processedXData.length};e.prototype.init=function(){k.prototype.init.apply(this,arguments);var a=this.options;a.pointRange=c(a.pointRange,a.colsize||1);this.yAxis.axisPointRange=a.rowsize||1;v.ellipse=v.circle};e.prototype.markerAttribs=function(a,b){var c=a.marker||{},e=this.options.marker||{},d=a.shapeArgs||{},g={};if(a.hasImage)return{x:a.plotX,y:a.plotY};if(b){var f=e.states[b]||{};var h=
c.states&&c.states[b]||{};[["width","x"],["height","y"]].forEach(function(a){g[a[0]]=(h[a[0]]||f[a[0]]||d[a[0]])+(h[a[0]+"Plus"]||f[a[0]+"Plus"]||0);g[a[1]]=d[a[1]]+(d[a[0]]-g[a[0]])/2})}return b?g:d};e.prototype.pointAttribs=function(a,c){var e=k.prototype.pointAttribs.call(this,a,c),d=this.options||{},g=this.chart.options.plotOptions||{},f=g.series||{},l=g.heatmap||{};g=a&&a.options.borderColor||d.borderColor||l.borderColor||f.borderColor;f=a&&a.options.borderWidth||d.borderWidth||l.borderWidth||
f.borderWidth||e["stroke-width"];e.stroke=a&&a.marker&&a.marker.lineColor||d.marker&&d.marker.lineColor||g||this.color;e["stroke-width"]=f;c&&(a=h(d.states[c],d.marker&&d.marker.states[c],a&&a.options.states&&a.options.states[c]||{}),c=a.brightness,e.fill=a.color||b.parse(e.fill).brighten(c||0).get(),e.stroke=a.lineColor);return e};e.prototype.setClip=function(a){var b=this.chart;k.prototype.setClip.apply(this,arguments);(!1!==this.options.clip||a)&&this.markerGroup.clip((a||this.clipBox)&&this.sharedClipKey?
b.sharedClips[this.sharedClipKey]:b.clipRect)};e.prototype.translate=function(){var a=this.options,b=a.marker&&a.marker.symbol||"rect",c=v[b]?b:"rect",e=-1!==["circle","square"].indexOf(c);this.generatePoints();this.points.forEach(function(a){var d=a.getCellAttributes(),f={};f.x=Math.min(d.x1,d.x2);f.y=Math.min(d.y1,d.y2);f.width=Math.max(Math.abs(d.x2-d.x1),0);f.height=Math.max(Math.abs(d.y2-d.y1),0);var g=a.hasImage=0===(a.marker&&a.marker.symbol||b||"").indexOf("url");if(e){var k=Math.abs(f.width-
f.height);f.x=Math.min(d.x1,d.x2)+(f.width<f.height?0:k/2);f.y=Math.min(d.y1,d.y2)+(f.width<f.height?k/2:0);f.width=f.height=Math.min(f.width,f.height)}k={plotX:(d.x1+d.x2)/2,plotY:(d.y1+d.y2)/2,clientX:(d.x1+d.x2)/2,shapeType:"path",shapeArgs:h(!0,f,{d:v[c](f.x,f.y,f.width,f.height)})};g&&(a.marker={width:f.width,height:f.height});C(a,k)});w(this,"afterTranslate")};e.defaultOptions=h(F.defaultOptions,{animation:!1,borderRadius:0,borderWidth:0,nullColor:"#f7f7f7",dataLabels:{formatter:function(){var a=
this.series.chart.numberFormatter,b=this.point.value;return t(b)?a(b,-1):""},inside:!0,verticalAlign:"middle",crop:!1,overflow:!1,padding:0},marker:{symbol:"rect",radius:0,lineColor:void 0,states:{hover:{lineWidthPlus:0},select:{}}},clip:!0,pointRange:null,tooltip:{pointFormat:"{point.x}, {point.y}: {point.value}<br/>"},states:{hover:{halo:!1,brightness:.2}}});return e}(F);C(y.prototype,{alignDataLabel:n.prototype.alignDataLabel,axisTypes:e.axisTypes,colorKey:"value",directTouch:!0,drawLegendSymbol:r.drawRectangle,
getExtremesFromAll:!0,getSymbol:k.prototype.getSymbol,parallelArrays:e.parallelArrays,pointArrayMap:["y","value"],pointClass:u,trackerGroups:e.trackerGroups});l.compose(y);q.registerSeriesType("heatmap",y);"";"";return y});u(b,"masters/modules/heatmap.src.js",[b["Core/Globals.js"],b["Core/Axis/Color/ColorAxis.js"]],function(b,l){b.ColorAxis=l;l.compose(b.Chart,b.Fx,b.Legend,b.Series)})});
//# sourceMappingURL=heatmap.js.map
```
