## Supplementary material for "Exploring natural odour landscapes: A case study with implications for human-biting insects": Figure S4: heikinashi.js

```
/*
 Highstock JS v9.3.1 (2021-11-05)

 HeikinAshi series type for Highcharts Stock

 (c) 2010-2021 Karol Kolodziej

 License: www.highcharts.com/license
*/
'use strict';(function(b){"object"===typeof module&&module.exports?(b["default"]=b,module.exports=b):"function"===typeof define&&define.amd?define("highcharts/modules/heikinashi",["highcharts","highcharts/modules/stock"],function(g){b(g);b.Highcharts=g;return b}):b("undefined"!==typeof Highcharts?Highcharts:void 0)})(function(b){function g(b,e,a,h){b.hasOwnProperty(e)||(b[e]=h.apply(null,a))}b=b?b._modules:{};g(b,"Series/HeikinAshi/HeikinAshiPoint.js",[b["Core/Series/SeriesRegistry.js"]],function(b){var e=
this&&this.__extends||function(){var b=function(a,c){b=Object.setPrototypeOf||{__proto__:[]}instanceof Array&&function(b,a){b.__proto__=a}||function(b,a){for(var c in a)a.hasOwnProperty(c)&&(b[c]=a[c])};return b(a,c)};return function(a,c){function e(){this.constructor=a}b(a,c);a.prototype=null===c?Object.create(c):(e.prototype=c.prototype,new e)}}();return function(b){function a(){var a=null!==b&&b.apply(this,arguments)||this;a.series=void 0;return a}e(a,b);return a}(b.seriesTypes.candlestick.prototype.pointClass)});
g(b,"Series/HeikinAshi/HeikinAshiSeries.js",[b["Series/HeikinAshi/HeikinAshiPoint.js"],b["Core/Series/SeriesRegistry.js"],b["Core/Utilities.js"],b["Core/Axis/Axis.js"]],function(b,e,a,g){var c=this&&this.__extends||function(){var b=function(a,f){b=Object.setPrototypeOf||{__proto__:[]}instanceof Array&&function(b,f){b.__proto__=f}||function(b,f){for(var a in f)f.hasOwnProperty(a)&&(b[a]=f[a])};return b(a,f)};return function(a,f){function m(){this.constructor=a}b(a,f);a.prototype=null===f?Object.create(f):
(m.prototype=f.prototype,new m)}}(),l=e.seriesTypes.candlestick,k=a.addEvent,h=a.merge;a=function(b){function a(){var a=null!==b&&b.apply(this,arguments)||this;a.data=void 0;a.heikiashiData=[];a.options=void 0;a.points=void 0;a.yData=void 0;a.processedYData=void 0;return a}c(a,b);a.prototype.getHeikinashiData=function(){var a=this.allGroupedData||this.yData,b=this.heikiashiData;if(!b.length&&a&&a.length){this.modifyFirstPointValue(a[0]);for(var d=1;d<a.length;d++)this.modifyDataPoint(a[d],b[d-1])}this.heikiashiData=
b};a.prototype.init=function(){b.prototype.init.apply(this,arguments);this.heikiashiData=[]};a.prototype.modifyFirstPointValue=function(a){this.heikiashiData.push([(a[0]+a[1]+a[2]+a[3])/4,a[1],a[2],(a[0]+a[3])/2])};a.prototype.modifyDataPoint=function(a,b){b=(b[0]+b[3])/2;var d=(a[0]+a[1]+a[2]+a[3])/4;this.heikiashiData.push([b,Math.max(a[1],d,b),Math.min(a[2],d,b),d])};a.defaultOptions=h(l.defaultOptions,{dataGrouping:{groupAll:!0}});return a}(l);k(a,"afterTranslate",function(){for(var a=this.points,
b=this.heikiashiData,f=this.cropStart||0,c=this.processedYData.length=0;c<a.length;c++){var d=a[c],e=b[c+f];d.open=e[0];d.high=e[1];d.low=e[2];d.close=e[3];this.processedYData.push([d.open,d.high,d.low,d.close])}});k(a,"updatedData",function(){this.heikiashiData.length&&(this.heikiashiData.length=0)});k(g,"postProcessData",function(){this.series.forEach(function(a){a.is("heikinashi")&&(a.heikiashiData.length=0,a.getHeikinashiData())})});a.prototype.pointClass=b;e.registerSeriesType("heikinashi",a);
"";return a});g(b,"masters/modules/heikinashi.src.js",[],function(){})});
//# sourceMappingURL=heikinashi.js.map
```
