## Supplementary material for "Exploring natural odour landscapes: A case study with implications for human-biting insects": Figure S4: histogram-bellcurve.js

```
/*
 Highcharts JS v9.3.1 (2021-11-05)

 (c) 2010-2021 Highsoft AS
 Author: Sebastian Domas

 License: www.highcharts.com/license
*/
'use strict';(function(a){"object"===typeof module&&module.exports?(a["default"]=a,module.exports=a):"function"===typeof define&&define.amd?define("highcharts/modules/histogram-bellcurve",["highcharts"],function(e){a(e);a.Highcharts=e;return a}):a("undefined"!==typeof Highcharts?Highcharts:void 0)})(function(a){function e(a,h,b,e){a.hasOwnProperty(h)||(a[h]=e.apply(null,b))}a=a?a._modules:{};e(a,"Series/DerivedComposition.js",[a["Core/Globals.js"],a["Core/Series/Series.js"],a["Core/Utilities.js"]],
function(a,h,b){var e=a.noop,g=b.addEvent,n=b.defined,p;(function(a){function b(){h.prototype.init.apply(this,arguments);this.initialised=!1;this.baseSeries=null;this.eventRemovers=[];this.addEvents()}function m(){var c=this.chart,a=this.options.baseSeries;this.baseSeries=n(a)&&(c.series[a]||c.get(a))||null}function d(){var c=this;this.eventRemovers.push(g(this.chart,"afterLinkSeries",function(){c.setBaseSeries();c.baseSeries&&!c.initialised&&(c.setDerivedData(),c.addBaseSeriesEvents(),c.initialised=
!0)}))}function f(){var c=this;this.eventRemovers.push(g(this.baseSeries,"updatedData",function(){c.setDerivedData()}),g(this.baseSeries,"destroy",function(){c.baseSeries=null;c.initialised=!1}))}function q(){this.eventRemovers.forEach(function(c){c()});h.prototype.destroy.apply(this,arguments)}var k=[];a.hasDerivedData=!0;a.setDerivedData=e;a.compose=function(c){if(-1===k.indexOf(c)){k.push(c);var a=c.prototype;a.addBaseSeriesEvents=f;a.addEvents=d;a.destroy=q;a.init=b;a.setBaseSeries=m}return c};
a.init=b;a.setBaseSeries=m;a.addEvents=d;a.addBaseSeriesEvents=f;a.destroy=q})(p||(p={}));return p});e(a,"Series/Histogram/HistogramSeries.js",[a["Series/DerivedComposition.js"],a["Core/Series/SeriesRegistry.js"],a["Core/Utilities.js"]],function(a,e,b){function h(a){return function(c){for(var d=1;a[d]<=c;)d++;return a[--d]}}var g=this&&this.__extends||function(){var a=function(c,d){a=Object.setPrototypeOf||{__proto__:[]}instanceof Array&&function(a,c){a.__proto__=c}||function(a,c){for(var d in c)c.hasOwnProperty(d)&&
(a[d]=c[d])};return a(c,d)};return function(c,d){function b(){this.constructor=c}a(c,d);c.prototype=null===d?Object.create(d):(b.prototype=d.prototype,new b)}}(),n=e.seriesTypes.column,p=b.arrayMax,r=b.arrayMin,l=b.correctFloat,m=b.extend,d=b.isNumber,f=b.merge,q=b.objectEach,k={"square-root":function(a){return Math.ceil(Math.sqrt(a.options.data.length))},sturges:function(a){return Math.ceil(Math.log(a.options.data.length)*Math.LOG2E)},rice:function(a){return Math.ceil(2*Math.pow(a.options.data.length,
1/3))}};b=function(a){function c(){var d=null!==a&&a.apply(this,arguments)||this;d.data=void 0;d.options=void 0;d.points=void 0;d.userOptions=void 0;return d}g(c,a);c.prototype.binsNumber=function(){var a=this.options.binsNumber,c=k[a]||"function"===typeof a&&a;return Math.ceil(c&&c(this.baseSeries)||(d(a)?a:k["square-root"](this.baseSeries)))};c.prototype.derivedData=function(a,c,b){var m=l(p(a)),f=l(r(a)),k=[],e={},g=[];b=this.binWidth=l(d(b)?b||1:(m-f)/c);this.options.pointRange=Math.max(b,0);
for(c=f;c<m&&(this.userOptions.binWidth||l(m-c)>=b||0>=l(l(f+k.length*b)-c));c=l(c+b))k.push(c),e[c]=0;0!==e[f]&&(k.push(f),e[f]=0);var t=h(k.map(function(a){return parseFloat(a)}));a.forEach(function(a){a=l(t(a));e[a]++});q(e,function(a,c){g.push({x:Number(c),y:a,x2:l(Number(c)+b)})});g.sort(function(a,c){return a.x-c.x});g[g.length-1].x2=m;return g};c.prototype.setDerivedData=function(){var a=this.baseSeries.yData;a.length?(a=this.derivedData(a,this.binsNumber(),this.options.binWidth),this.setData(a,
!1)):this.setData([])};c.defaultOptions=f(n.defaultOptions,{binsNumber:"square-root",binWidth:void 0,pointPadding:0,groupPadding:0,grouping:!1,pointPlacement:"between",tooltip:{headerFormat:"",pointFormat:'<span style="font-size: 10px">{point.x} - {point.x2}</span><br/><span style="color:{point.color}">\u25cf</span> {series.name} <b>{point.y}</b><br/>'}});return c}(n);m(b.prototype,{hasDerivedData:a.hasDerivedData});a.compose(b);e.registerSeriesType("histogram",b);"";return b});e(a,"Series/Bellcurve/BellcurveSeries.js",
[a["Series/DerivedComposition.js"],a["Core/Series/SeriesRegistry.js"],a["Core/Utilities.js"]],function(a,e,b){var h=this&&this.__extends||function(){var a=function(b,d){a=Object.setPrototypeOf||{__proto__:[]}instanceof Array&&function(a,d){a.__proto__=d}||function(a,d){for(var b in d)d.hasOwnProperty(b)&&(a[b]=d[b])};return a(b,d)};return function(b,d){function f(){this.constructor=b}a(b,d);b.prototype=null===d?Object.create(d):(f.prototype=d.prototype,new f)}}(),g=e.seriesTypes.areaspline,n=b.correctFloat,
p=b.isNumber,r=b.merge;b=function(a){function b(){var b=null!==a&&a.apply(this,arguments)||this;b.data=void 0;b.options=void 0;b.points=void 0;return b}h(b,a);b.mean=function(a){var b=a.length;a=a.reduce(function(a,b){return a+b},0);return 0<b&&a/b};b.standardDeviation=function(a,f){var d=a.length;f=p(f)?f:b.mean(a);a=a.reduce(function(a,b){b-=f;return a+b*b},0);return 1<d&&Math.sqrt(a/(d-1))};b.normalDensity=function(a,b,e){a-=b;return Math.exp(-(a*a)/(2*e*e))/(e*Math.sqrt(2*Math.PI))};b.prototype.derivedData=
function(a,e){var d=this.options.intervals,f=this.options.pointsInInterval,c=a-d*e;d=d*f*2+1;f=e/f;var g=[],h;for(h=0;h<d;h++)g.push([c,b.normalDensity(c,a,e)]),c+=f;return g};b.prototype.setDerivedData=function(){1<this.baseSeries.yData.length&&(this.setMean(),this.setStandardDeviation(),this.setData(this.derivedData(this.mean,this.standardDeviation),!1))};b.prototype.setMean=function(){this.mean=n(b.mean(this.baseSeries.yData))};b.prototype.setStandardDeviation=function(){this.standardDeviation=
n(b.standardDeviation(this.baseSeries.yData,this.mean))};b.defaultOptions=r(g.defaultOptions,{intervals:3,pointsInInterval:3,marker:{enabled:!1}});return b}(g);a.compose(b);e.registerSeriesType("bellcurve",b);"";return b});e(a,"masters/modules/histogram-bellcurve.src.js",[],function(){})});
//# sourceMappingURL=histogram-bellcurve.js.map
```
