## Supplementary material for "Exploring natural odour landscapes: A case study with implications for human-biting insects": Figure S4: hollowcandlestick.js

```
/*
 Highstock JS v9.3.1 (2021-11-05)

 Hollow Candlestick series type for Highcharts Stock

 (c) 2010-2021 Karol Kolodziej

 License: www.highcharts.com/license
*/
'use strict';(function(b){"object"===typeof module&&module.exports?(b["default"]=b,module.exports=b):"function"===typeof define&&define.amd?define("highcharts/modules/hollowcandlestick",["highcharts","highcharts/modules/stock"],function(f){b(f);b.Highcharts=f;return b}):b("undefined"!==typeof Highcharts?Highcharts:void 0)})(function(b){function f(b,h,a,k){b.hasOwnProperty(h)||(b[h]=k.apply(null,a))}b=b?b._modules:{};f(b,"Series/HollowCandlestick/HollowCandlestickPoint.js",[b["Core/Series/SeriesRegistry.js"]],
function(b){var h=this&&this.__extends||function(){var b=function(a,c){b=Object.setPrototypeOf||{__proto__:[]}instanceof Array&&function(b,a){b.__proto__=a}||function(b,a){for(var c in a)a.hasOwnProperty(c)&&(b[c]=a[c])};return b(a,c)};return function(a,c){function g(){this.constructor=a}b(a,c);a.prototype=null===c?Object.create(c):(g.prototype=c.prototype,new g)}}();return function(b){function a(){var a=null!==b&&b.apply(this,arguments)||this;a.series=void 0;return a}h(a,b);a.prototype.getClassName=
function(){var a=b.prototype.getClassName.apply(this),g=this.series.hollowCandlestickData[this.index];g.isBullish||"up"!==g.trendDirection||(a+="-bearish-up");return a};return a}(b.seriesTypes.candlestick.prototype.pointClass)});f(b,"Series/HollowCandlestick/HollowCandlestickSeries.js",[b["Series/HollowCandlestick/HollowCandlestickPoint.js"],b["Core/Series/SeriesRegistry.js"],b["Core/Utilities.js"],b["Core/Axis/Axis.js"]],function(b,h,a,f){var c=this&&this.__extends||function(){var b=function(a,e){b=
Object.setPrototypeOf||{__proto__:[]}instanceof Array&&function(b,a){b.__proto__=a}||function(b,a){for(var e in a)a.hasOwnProperty(e)&&(b[e]=a[e])};return b(a,e)};return function(a,e){function m(){this.constructor=a}b(a,e);a.prototype=null===e?Object.create(e):(m.prototype=e.prototype,new m)}}(),g=h.seriesTypes.candlestick,l=a.addEvent,k=a.merge;a=function(b){function a(){var a=null!==b&&b.apply(this,arguments)||this;a.data=void 0;a.hollowCandlestickData=[];a.options=void 0;a.points=void 0;return a}
c(a,b);a.prototype.getPriceMovement=function(){var a=this.allGroupedData||this.yData,b=this.hollowCandlestickData;if(!b.length&&a&&a.length){b.push({isBullish:!0,trendDirection:"up"});for(var d=1;d<a.length;d++)b.push(this.isBullish(a[d],a[d-1]))}};a.prototype.getLineColor=function(a){return"up"===a?this.options.upColor||"#06b535":this.options.color||"#f21313"};a.prototype.getPointFill=function(a){return a.isBullish?"transparent":"up"===a.trendDirection?this.options.upColor||"#06b535":this.options.color||
"#f21313"};a.prototype.init=function(){b.prototype.init.apply(this,arguments);this.hollowCandlestickData=[]};a.prototype.isBullish=function(a,b){return{isBullish:a[0]<=a[3],trendDirection:a[3]<b[3]?"down":"up"}};a.prototype.pointAttribs=function(a,c){var d=b.prototype.pointAttribs.call(this,a,c);a=this.hollowCandlestickData[a.index];d.fill=this.getPointFill(a)||d.fill;d.stroke=this.getLineColor(a.trendDirection)||d.stroke;c&&(c=this.options.states[c],d.fill=c.color||d.fill,d.stroke=c.lineColor||d.stroke,
d["stroke-width"]=c.lineWidth||d["stroke-width"]);return d};a.defaultOptions=k(g.defaultOptions,{color:"#f21313",dataGrouping:{groupAll:!0,groupPixelWidth:10},lineColor:"#f21313",upColor:"#06b535",upLineColor:"#06b535"});return a}(g);l(a,"updatedData",function(){this.hollowCandlestickData.length&&(this.hollowCandlestickData.length=0)});l(f,"postProcessData",function(){this.series.forEach(function(a){a.is("hollowcandlestick")&&a.getPriceMovement()})});a.prototype.pointClass=b;h.registerSeriesType("hollowcandlestick",
a);"";return a});f(b,"masters/modules/hollowcandlestick.src.js",[],function(){})});
//# sourceMappingURL=hollowcandlestick.js.map
```
