## Supplementary material for "Exploring natural odour landscapes: A case study with implications for human-biting insects": Figure S4: item-series.js

```
/*
 Highcharts JS v9.3.1 (2021-11-05)

 Item series type for Highcharts

 (c) 2019 Torstein Honsi

 License: www.highcharts.com/license
*/
'use strict';(function(a){"object"===typeof module&&module.exports?(a["default"]=a,module.exports=a):"function"===typeof define&&define.amd?define("highcharts/modules/item-series",["highcharts"],function(b){a(b);a.Highcharts=b;return a}):a("undefined"!==typeof Highcharts?Highcharts:void 0)})(function(a){function b(a,g,c,b){a.hasOwnProperty(g)||(a[g]=b.apply(null,c))}a=a?a._modules:{};b(a,"Series/Item/ItemPoint.js",[a["Core/Series/SeriesRegistry.js"],a["Core/Utilities.js"]],function(a,b){var c=this&&
this.__extends||function(){var a=function(b,e){a=Object.setPrototypeOf||{__proto__:[]}instanceof Array&&function(a,b){a.__proto__=b}||function(a,b){for(var e in b)b.hasOwnProperty(e)&&(a[e]=b[e])};return a(b,e)};return function(b,e){function f(){this.constructor=b}a(b,e);b.prototype=null===e?Object.create(e):(f.prototype=e.prototype,new f)}}(),g=a.series;b=b.extend;a=function(a){function b(){var b=null!==a&&a.apply(this,arguments)||this;b.graphics=void 0;b.options=void 0;b.series=void 0;return b}
c(b,a);return b}(a.seriesTypes.pie.prototype.pointClass);b(a.prototype,{haloPath:g.prototype.pointClass.prototype.haloPath});return a});b(a,"Series/Item/ItemSeries.js",[a["Core/Globals.js"],a["Series/Item/ItemPoint.js"],a["Core/DefaultOptions.js"],a["Core/Series/SeriesRegistry.js"],a["Core/Utilities.js"]],function(a,b,c,w,f){var g=this&&this.__extends||function(){var a=function(b,d){a=Object.setPrototypeOf||{__proto__:[]}instanceof Array&&function(a,d){a.__proto__=d}||function(a,d){for(var b in d)d.hasOwnProperty(b)&&
(a[b]=d[b])};return a(b,d)};return function(b,d){function z(){this.constructor=b}a(b,d);b.prototype=null===d?Object.create(d):(z.prototype=d.prototype,new z)}}(),e=c.defaultOptions,x=w.seriesTypes.pie,I=f.defined,y=f.extend,J=f.fireEvent,A=f.isNumber,B=f.merge,K=f.objectEach,L=f.pick;c=function(b){function c(){var a=null!==b&&b.apply(this,arguments)||this;a.data=void 0;a.options=void 0;a.points=void 0;return a}g(c,b);c.prototype.animate=function(a){a?this.group.attr({opacity:0}):this.group.animate({opacity:1},
this.options.animation)};c.prototype.drawDataLabels=function(){this.center&&this.slots?a.seriesTypes.pie.prototype.drawDataLabels.call(this):this.points.forEach(function(a){a.destroyElements({dataLabel:1})})};c.prototype.drawPoints=function(){var a=this,b=this.options,c=a.chart.renderer,e=b.marker,f=this.borderWidth%2?.5:1,C=0,r=this.getRows(),g=Math.ceil(this.total/r),t=this.chart.plotWidth/g,u=this.chart.plotHeight/r,v=this.itemSize||Math.min(t,u);this.points.forEach(function(d){var n,z,l=d.marker||
{},p=l.symbol||e.symbol;l=L(l.radius,e.radius);var G=I(l)?2*l:v,q=G*b.itemPadding,H;d.graphics=n=d.graphics||{};a.chart.styledMode||(z=a.pointAttribs(d,d.selected&&"select"));if(!d.isNull&&d.visible){d.graphic||(d.graphic=c.g("point").add(a.group));for(var h=0;h<d.y;h++){if(a.center&&a.slots){var m=a.slots.shift();var k=m.x-v/2;m=m.y-v/2}else"horizontal"===b.layout?(k=C%g*t,m=u*Math.floor(C/g)):(k=t*Math.floor(C/r),m=C%r*u);k+=q;m+=q;var M=H=Math.round(G-2*q);a.options.crisp&&(k=Math.round(k)-f,m=
Math.round(m)+f);k={x:k,y:m,width:H,height:M};"undefined"!==typeof l&&(k.r=l);n[h]?n[h].animate(k):n[h]=c.symbol(p,null,null,null,null,{backgroundSize:"within"}).attr(y(k,z)).add(d.graphic);n[h].isActive=!0;C++}}K(n,function(a,b){a.isActive?a.isActive=!1:(a.destroy(),delete n[b])})})};c.prototype.getRows=function(){var a=this.options.rows;if(!a){var b=this.chart.plotWidth/this.chart.plotHeight;a=Math.sqrt(this.total);if(1<b)for(a=Math.ceil(a);0<a;){var c=this.total/a;if(c/a>b)break;a--}else for(a=
Math.floor(a);a<this.total;){c=this.total/a;if(c/a<b)break;a++}}return a};c.prototype.getSlots=function(){function a(a){0<E&&(a.row.colCount--,E--)}for(var b=this.center,c=b[2],e=b[3],f,g=this.slots,r,w,t,u,v,x,n,D,l=0,p,F=this.endAngleRad-this.startAngleRad,q=Number.MAX_VALUE,y,h,m,k=this.options.rows,A=(c-e)/c,B=0===F%(2*Math.PI);q>this.total+(h&&B?h.length:0);)for(y=q,q=g.length=0,h=m,m=[],l++,p=c/l/2,k?(e=(p-k)/p*c,0<=e?p=k:(e=0,A=1)):p=Math.floor(p*A),f=p;0<f;f--)t=(e+f/p*(c-e-l))/2,u=F*t,v=
Math.ceil(u/l),m.push({rowRadius:t,rowLength:u,colCount:v}),q+=v+1;if(h){for(var E=y-this.total-(B?h.length:0);0<E;)h.map(function(a){return{angle:a.colCount/a.rowLength,row:a}}).sort(function(a,b){return b.angle-a.angle}).slice(0,Math.min(E,Math.ceil(h.length/2))).forEach(a);h.forEach(function(a){var c=a.rowRadius;x=(a=a.colCount)?F/a:0;for(D=0;D<=a;D+=1)n=this.startAngleRad+D*x,r=b[0]+Math.cos(n)*c,w=b[1]+Math.sin(n)*c,g.push({x:r,y:w,angle:n})},this);g.sort(function(a,b){return a.angle-b.angle});
this.itemSize=l;return g}};c.prototype.translate=function(b){0===this.total&&(this.center=this.getCenter());this.slots||(this.slots=[]);A(this.options.startAngle)&&A(this.options.endAngle)?(a.seriesTypes.pie.prototype.translate.apply(this,arguments),this.slots=this.getSlots()):(this.generatePoints(),J(this,"afterTranslate"))};c.defaultOptions=B(x.defaultOptions,{endAngle:void 0,innerSize:"40%",itemPadding:.1,layout:"vertical",marker:B(e.plotOptions.line.marker,{radius:null}),rows:void 0,crisp:!1,
showInLegend:!0,startAngle:void 0});return c}(x);y(c.prototype,{markerAttribs:void 0});c.prototype.pointClass=b;w.registerSeriesType("item",c);"";return c});b(a,"masters/modules/item-series.src.js",[],function(){})});
//# sourceMappingURL=item-series.js.map
```
