## Supplementary material for "Exploring natural odour landscapes: A case study with implications for human-biting insects": Figure S4: lollipop.js

```
/*
 Highcharts JS v9.3.1 (2021-11-05)

 (c) 2009-2021 Sebastian Bochan, Rafal Sebestjanski

 License: www.highcharts.com/license
*/
'use strict';(function(b){"object"===typeof module&&module.exports?(b["default"]=b,module.exports=b):"function"===typeof define&&define.amd?define("highcharts/modules/lollipop",["highcharts"],function(c){b(c);b.Highcharts=c;return b}):b("undefined"!==typeof Highcharts?Highcharts:void 0)})(function(b){function c(b,e,f,c){b.hasOwnProperty(e)||(b[e]=c.apply(null,f))}b=b?b._modules:{};c(b,"Series/Lollipop/LollipopPoint.js",[b["Core/Series/SeriesRegistry.js"],b["Core/Utilities.js"]],function(b,e){var f=
this&&this.__extends||function(){var b=function(d,a){b=Object.setPrototypeOf||{__proto__:[]}instanceof Array&&function(b,d){b.__proto__=d}||function(b,d){for(var a in d)d.hasOwnProperty(a)&&(b[a]=d[a])};return b(d,a)};return function(d,a){function c(){this.constructor=d}b(d,a);d.prototype=null===a?Object.create(a):(c.prototype=a.prototype,new c)}}(),c=b.series.prototype.pointClass,a=b.seriesTypes;b=a.area.prototype;var g=e.isObject;e=e.extend;a=function(b){function a(){var a=null!==b&&b.apply(this,
arguments)||this;a.series=void 0;a.options=void 0;return a}f(a,b);return a}(a.dumbbell.prototype.pointClass);e(a.prototype,{pointSetState:b.pointClass.prototype.setState,isValid:c.prototype.isValid,init:function(b,a,e){g(a)&&"low"in a&&(a.y=a.low,delete a.low);return c.prototype.init.apply(this,arguments)}});return a});c(b,"Series/Lollipop/LollipopSeries.js",[b["Series/Lollipop/LollipopPoint.js"],b["Core/Series/SeriesRegistry.js"],b["Core/Utilities.js"]],function(b,c,f){var e=this&&this.__extends||
function(){var b=function(a,c){b=Object.setPrototypeOf||{__proto__:[]}instanceof Array&&function(b,a){b.__proto__=a}||function(b,a){for(var c in a)a.hasOwnProperty(c)&&(b[c]=a[c])};return b(a,c)};return function(a,c){function d(){this.constructor=a}b(a,c);a.prototype=null===c?Object.create(c):(d.prototype=c.prototype,new d)}}(),a=c.seriesTypes,g=a.area.prototype,h=a.column.prototype,d=a.dumbbell,k=f.pick,l=f.merge;f=f.extend;a=function(b){function a(){var a=null!==b&&b.apply(this,arguments)||this;
a.data=void 0;a.options=void 0;a.points=void 0;return a}e(a,b);a.prototype.toYData=function(a){return[k(a.y,a.low)]};a.defaultOptions=l(d.defaultOptions,{lowColor:void 0,threshold:0,connectorWidth:1,groupPadding:.2,pointPadding:.1,states:{hover:{lineWidthPlus:0,connectorWidthPlus:1,halo:!1}},tooltip:{pointFormat:'<span style="color:{series.color}">\u25cf</span> {series.name}: <b>{point.y}</b><br/>'}});return a}(d);f(a.prototype,{pointArrayMap:["y"],pointValKey:"y",translatePoint:g.translate,drawPoint:g.drawPoints,
drawDataLabels:h.drawDataLabels,setShapeArgs:h.translate,pointClass:b});c.registerSeriesType("lollipop",a);"";return a});c(b,"masters/modules/lollipop.src.js",[],function(){})});
//# sourceMappingURL=lollipop.js.map
```
