## Supplementary material for "Exploring natural odour landscapes: A case study with implications for human-biting insects": Figure S4: map.js

```
/*
 Highmaps JS v9.3.1 (2021-11-05)

 Highmaps as a plugin for Highcharts or Highcharts Stock.

 (c) 2011-2021 Torstein Honsi

 License: www.highcharts.com/license
*/
'use strict';(function(c){"object"===typeof module&&module.exports?(c["default"]=c,module.exports=c):"function"===typeof define&&define.amd?define("highcharts/modules/map",["highcharts"],function(x){c(x);c.Highcharts=x;return c}):c("undefined"!==typeof Highcharts?Highcharts:void 0)})(function(c){function x(c,p,q,l){c.hasOwnProperty(p)||(c[p]=l.apply(null,q))}c=c?c._modules:{};x(c,"Core/Axis/Color/ColorAxisComposition.js",[c["Core/Color/Color.js"],c["Core/Utilities.js"]],function(c,p){var q=c.parse,
l=p.addEvent,n=p.extend,h=p.merge,b=p.pick,e=p.splat,g;(function(g){function k(){var a=this,d=this.options;this.colorAxis=[];d.colorAxis&&(d.colorAxis=e(d.colorAxis),d.colorAxis.forEach(function(d,m){d.index=m;new A(a,d)}))}function a(a){var d=this,m=function(m){m=a.allItems.indexOf(m);-1!==m&&(d.destroyItem(a.allItems[m]),a.allItems.splice(m,1))},w=[],f,v;(this.chart.colorAxis||[]).forEach(function(a){(f=a.options)&&f.showInLegend&&(f.dataClasses&&f.visible?w=w.concat(a.getDataClassLegendSymbols()):
f.visible&&w.push(a),a.series.forEach(function(a){if(!a.options.showInLegend||f.dataClasses)"point"===a.options.legendType?a.points.forEach(function(a){m(a)}):m(a)}))});for(v=w.length;v--;)a.allItems.unshift(w[v])}function d(a){a.visible&&a.item.legendColor&&a.item.legendSymbol.attr({fill:a.item.legendColor})}function f(){var a=this.chart.colorAxis;a&&a.forEach(function(a,d,m){a.update({},m)})}function r(){(this.chart.colorAxis&&this.chart.colorAxis.length||this.colorAttribs)&&this.translateColors()}
function y(){var a=this.axisTypes;a?-1===a.indexOf("colorAxis")&&a.push("colorAxis"):this.axisTypes=["colorAxis"]}function t(a){var m=this,d=a?"show":"hide";m.visible=m.options.visible=!!a;["graphic","dataLabel"].forEach(function(a){if(m[a])m[a][d]()});this.series.buildKDTree()}function u(){var a=this,m=this.options.nullColor,d=this.colorAxis,w=this.colorKey;(this.data.length?this.data:this.points).forEach(function(f){var v=f.getNestedProperty(w);(v=f.options.color||(f.isNull||null===f.value?m:d&&
"undefined"!==typeof v?d.toColor(v,f):f.color||a.color))&&f.color!==v&&(f.color=v,"point"===a.options.legendType&&f.legendItem&&a.chart.legend.colorizeItem(f,f.visible))})}function v(a){var m=a.prototype.createAxis;a.prototype.createAxis=function(a,d){if("colorAxis"!==a)return m.apply(this,arguments);var f=new A(this,h(d.axis,{index:this[a].length,isX:!1}));this.isDirtyLegend=!0;this.axes.forEach(function(a){a.series=[]});this.series.forEach(function(a){a.bindAxes();a.isDirtyData=!0});b(d.redraw,
!0)&&this.redraw(d.animation);return f}}function B(){this.elem.attr("fill",q(this.start).tweenTo(q(this.end),this.pos),void 0,!0)}function w(){this.elem.attr("stroke",q(this.start).tweenTo(q(this.end),this.pos),void 0,!0)}var m=[],A;g.compose=function(e,b,g,h,z){A||(A=e);-1===m.indexOf(b)&&(m.push(b),e=b.prototype,e.collectionsWithUpdate.push("colorAxis"),e.collectionsWithInit.colorAxis=[e.addColorAxis],l(b,"afterGetAxes",k),v(b));-1===m.indexOf(g)&&(m.push(g),b=g.prototype,b.fillSetter=B,b.strokeSetter=
w);-1===m.indexOf(h)&&(m.push(h),l(h,"afterGetAllItems",a),l(h,"afterColorizeItem",d),l(h,"afterUpdate",f));-1===m.indexOf(z)&&(m.push(z),n(z.prototype,{optionalAxis:"colorAxis",translateColors:u}),n(z.prototype.pointClass.prototype,{setVisible:t}),l(z,"afterTranslate",r),l(z,"bindAxes",y))};g.pointSetVisible=t})(g||(g={}));return g});x(c,"Core/Axis/Color/ColorAxisDefaults.js",[],function(){return{lineWidth:0,minPadding:0,maxPadding:0,gridLineWidth:1,tickPixelInterval:72,startOnTick:!0,endOnTick:!0,
offset:0,marker:{animation:{duration:50},width:.01,color:"#999999"},labels:{overflow:"justify",rotation:0},minColor:"#e6ebf5",maxColor:"#003399",tickLength:5,showInLegend:!0}});x(c,"Core/Axis/Color/ColorAxis.js",[c["Core/Axis/Axis.js"],c["Core/Color/Color.js"],c["Core/Axis/Color/ColorAxisComposition.js"],c["Core/Axis/Color/ColorAxisDefaults.js"],c["Core/Globals.js"],c["Core/Legend/LegendSymbol.js"],c["Core/Series/SeriesRegistry.js"],c["Core/Utilities.js"]],function(c,p,q,l,n,h,b,e){var g=this&&this.__extends||
function(){var a=function(d,f){a=Object.setPrototypeOf||{__proto__:[]}instanceof Array&&function(a,d){a.__proto__=d}||function(a,d){for(var m in d)d.hasOwnProperty(m)&&(a[m]=d[m])};return a(d,f)};return function(d,f){function v(){this.constructor=d}a(d,f);d.prototype=null===f?Object.create(f):(v.prototype=f.prototype,new v)}}(),t=p.parse,k=n.noop,a=b.series,d=e.extend,f=e.isNumber,r=e.merge,y=e.pick;p=function(e){function b(a,d){var f=e.call(this,a,d)||this;f.beforePadding=!1;f.chart=void 0;f.coll=
"colorAxis";f.dataClasses=void 0;f.legendItem=void 0;f.legendItems=void 0;f.name="";f.options=void 0;f.stops=void 0;f.visible=!0;f.init(a,d);return f}g(b,e);b.compose=function(a,d,f,m){q.compose(b,a,d,f,m)};b.prototype.init=function(a,d){var f=a.options.legend||{},m=d.layout?"vertical"!==d.layout:"vertical"!==f.layout,v=d.visible;f=r(b.defaultColorAxisOptions,d,{showEmpty:!1,title:null,visible:f.enabled&&!1!==v});this.coll="colorAxis";this.side=d.side||m?2:1;this.reversed=d.reversed||!m;this.opposite=
!m;e.prototype.init.call(this,a,f);this.userOptions.visible=v;d.dataClasses&&this.initDataClasses(d);this.initStops();this.horiz=m;this.zoomEnabled=!1};b.prototype.initDataClasses=function(a){var d=this.chart,f=this.options,m=a.dataClasses.length,e,b=0,v=d.options.chart.colorCount;this.dataClasses=e=[];this.legendItems=[];(a.dataClasses||[]).forEach(function(a,w){a=r(a);e.push(a);if(d.styledMode||!a.color)"category"===f.dataClassColor?(d.styledMode||(w=d.options.colors,v=w.length,a.color=w[b]),a.colorIndex=
b,b++,b===v&&(b=0)):a.color=t(f.minColor).tweenTo(t(f.maxColor),2>m?.5:w/(m-1))})};b.prototype.hasData=function(){return!!(this.tickPositions||[]).length};b.prototype.setTickPositions=function(){if(!this.dataClasses)return e.prototype.setTickPositions.call(this)};b.prototype.initStops=function(){this.stops=this.options.stops||[[0,this.options.minColor],[1,this.options.maxColor]];this.stops.forEach(function(a){a.color=t(a[1])})};b.prototype.setOptions=function(a){e.prototype.setOptions.call(this,a);
this.options.crosshair=this.options.marker};b.prototype.setAxisSize=function(){var a=this.legendSymbol,d=this.chart,f=d.options.legend||{},m,e;a?(this.left=f=a.attr("x"),this.top=m=a.attr("y"),this.width=e=a.attr("width"),this.height=a=a.attr("height"),this.right=d.chartWidth-f-e,this.bottom=d.chartHeight-m-a,this.len=this.horiz?e:a,this.pos=this.horiz?f:m):this.len=(this.horiz?f.symbolWidth:f.symbolHeight)||b.defaultLegendLength};b.prototype.normalizedValue=function(a){this.logarithmic&&(a=this.logarithmic.log2lin(a));
return 1-(this.max-a)/(this.max-this.min||1)};b.prototype.toColor=function(a,d){var f=this.dataClasses,m=this.stops,e;if(f)for(e=f.length;e--;){var b=f[e];var v=b.from;m=b.to;if(("undefined"===typeof v||a>=v)&&("undefined"===typeof m||a<=m)){var r=b.color;d&&(d.dataClass=e,d.colorIndex=b.colorIndex);break}}else{a=this.normalizedValue(a);for(e=m.length;e--&&!(a>m[e][0]););v=m[e]||m[e+1];m=m[e+1]||v;a=1-(m[0]-a)/(m[0]-v[0]||1);r=v.color.tweenTo(m.color,a)}return r};b.prototype.getOffset=function(){var a=
this.legendGroup,d=this.chart.axisOffset[this.side];a&&(this.axisParent=a,e.prototype.getOffset.call(this),this.added||(this.added=!0,this.labelLeft=0,this.labelRight=this.width),this.chart.axisOffset[this.side]=d)};b.prototype.setLegendColor=function(){var a=this.reversed,d=a?1:0;a=a?0:1;d=this.horiz?[d,0,a,0]:[0,a,0,d];this.legendColor={linearGradient:{x1:d[0],y1:d[1],x2:d[2],y2:d[3]},stops:this.stops}};b.prototype.drawLegendSymbol=function(a,d){var f=a.padding,m=a.options,e=this.horiz,r=y(m.symbolWidth,
e?b.defaultLegendLength:12),v=y(m.symbolHeight,e?12:b.defaultLegendLength),g=y(m.labelPadding,e?16:30);m=y(m.itemDistance,10);this.setLegendColor();d.legendSymbol=this.chart.renderer.rect(0,a.baseline-11,r,v).attr({zIndex:1}).add(d.legendGroup);this.legendItemWidth=r+f+(e?m:g);this.legendItemHeight=v+f+(e?g:0)};b.prototype.setState=function(a){this.series.forEach(function(d){d.setState(a)})};b.prototype.setVisible=function(){};b.prototype.getSeriesExtremes=function(){var d=this.series,f=d.length,
e;this.dataMin=Infinity;for(this.dataMax=-Infinity;f--;){var m=d[f];var b=m.colorKey=y(m.options.colorKey,m.colorKey,m.pointValKey,m.zoneAxis,"y");var r=m.pointArrayMap;var g=m[b+"Min"]&&m[b+"Max"];if(m[b+"Data"])var k=m[b+"Data"];else if(r){k=[];r=r.indexOf(b);var u=m.yData;if(0<=r&&u)for(e=0;e<u.length;e++)k.push(y(u[e][r],u[e]))}else k=m.yData;g?(m.minColorValue=m[b+"Min"],m.maxColorValue=m[b+"Max"]):(k=a.prototype.getExtremes.call(m,k),m.minColorValue=k.dataMin,m.maxColorValue=k.dataMax);"undefined"!==
typeof m.minColorValue&&(this.dataMin=Math.min(this.dataMin,m.minColorValue),this.dataMax=Math.max(this.dataMax,m.maxColorValue));g||a.prototype.applyExtremes.call(m)}};b.prototype.drawCrosshair=function(a,d){var f=d&&d.plotX,m=d&&d.plotY,b=this.pos,r=this.len;if(d){var g=this.toPixels(d.getNestedProperty(d.series.colorKey));g<b?g=b-2:g>b+r&&(g=b+r+2);d.plotX=g;d.plotY=this.len-g;e.prototype.drawCrosshair.call(this,a,d);d.plotX=f;d.plotY=m;this.cross&&!this.cross.addedToColorAxis&&this.legendGroup&&
(this.cross.addClass("highcharts-coloraxis-marker").add(this.legendGroup),this.cross.addedToColorAxis=!0,this.chart.styledMode||"object"!==typeof this.crosshair||this.cross.attr({fill:this.crosshair.color}))}};b.prototype.getPlotLinePath=function(a){var d=this.left,b=a.translatedValue,m=this.top;return f(b)?this.horiz?[["M",b-4,m-6],["L",b+4,m-6],["L",b,m],["Z"]]:[["M",d,b],["L",d-6,b+6],["L",d-6,b-6],["Z"]]:e.prototype.getPlotLinePath.call(this,a)};b.prototype.update=function(a,d){var f=this.chart.legend;
this.series.forEach(function(a){a.isDirtyData=!0});(a.dataClasses&&f.allItems||this.dataClasses)&&this.destroyItems();e.prototype.update.call(this,a,d);this.legendItem&&(this.setLegendColor(),f.colorizeItem(this,!0))};b.prototype.destroyItems=function(){var a=this.chart;this.legendItem?a.legend.destroyItem(this):this.legendItems&&this.legendItems.forEach(function(d){a.legend.destroyItem(d)});a.isDirtyLegend=!0};b.prototype.destroy=function(){this.chart.isDirtyLegend=!0;this.destroyItems();e.prototype.destroy.apply(this,
[].slice.call(arguments))};b.prototype.remove=function(a){this.destroyItems();e.prototype.remove.call(this,a)};b.prototype.getDataClassLegendSymbols=function(){var a=this,f=a.chart,e=a.legendItems,m=f.options.legend,b=m.valueDecimals,r=m.valueSuffix||"",g;e.length||a.dataClasses.forEach(function(m,w){var v=m.from,u=m.to,y=f.numberFormatter,t=!0;g="";"undefined"===typeof v?g="< ":"undefined"===typeof u&&(g="> ");"undefined"!==typeof v&&(g+=y(v,b)+r);"undefined"!==typeof v&&"undefined"!==typeof u&&
(g+=" - ");"undefined"!==typeof u&&(g+=y(u,b)+r);e.push(d({chart:f,name:g,options:{},drawLegendSymbol:h.drawRectangle,visible:!0,setState:k,isDataClass:!0,setVisible:function(){t=a.visible=!t;a.series.forEach(function(a){a.points.forEach(function(a){a.dataClass===w&&a.setVisible(t)})});f.legend.colorizeItem(this,t)}},m))});return e};b.defaultColorAxisOptions=l;b.defaultLegendLength=200;b.keepProps=["legendGroup","legendItemHeight","legendItemWidth","legendItem","legendSymbol"];return b}(c);Array.prototype.push.apply(c.keepProps,
p.keepProps);"";return p});x(c,"Maps/MapNavigationOptionsDefault.js",[c["Core/DefaultOptions.js"],c["Core/Utilities.js"]],function(c,p){p=p.extend;var q={buttonOptions:{alignTo:"plotBox",align:"left",verticalAlign:"top",x:0,width:18,height:18,padding:5,style:{fontSize:"15px",fontWeight:"bold"},theme:{"stroke-width":1,"text-align":"center"}},buttons:{zoomIn:{onclick:function(){this.mapZoom(.5)},text:"+",y:0},zoomOut:{onclick:function(){this.mapZoom(2)},text:"-",y:28}},mouseWheelSensitivity:1.1};p(c.defaultOptions.lang,
{zoomIn:"Zoom in",zoomOut:"Zoom out"});return c.defaultOptions.mapNavigation=q});x(c,"Maps/MapNavigation.js",[c["Core/Chart/Chart.js"],c["Core/Globals.js"],c["Core/Utilities.js"]],function(c,p,q){function l(a){a&&(a.preventDefault&&a.preventDefault(),a.stopPropagation&&a.stopPropagation(),a.cancelBubble=!0)}function n(a){this.init(a)}var h=p.doc,b=q.addEvent,e=q.extend,g=q.merge,t=q.objectEach,k=q.pick;n.prototype.init=function(a){this.chart=a;a.mapNavButtons=[]};n.prototype.update=function(a){var d=
this.chart,f=d.options.mapNavigation,r,y,h,u,v=function(a){this.handler.call(d,a);l(a)},c=d.mapNavButtons;a&&(f=d.options.mapNavigation=g(d.options.mapNavigation,a));for(;c.length;)c.pop().destroy();k(f.enableButtons,f.enabled)&&!d.renderer.forExport&&t(f.buttons,function(a,m){a=g(f.buttonOptions,a);!d.styledMode&&a.theme&&(r=a.theme,r.style=g(a.theme.style,a.style),h=(y=r.states)&&y.hover,u=y&&y.select,delete r.states);var k=d.renderer.button(a.text||"",0,0,v,r,h,u,void 0,"zoomIn"===m?"topbutton":
"bottombutton").addClass("highcharts-map-navigation highcharts-"+{zoomIn:"zoom-in",zoomOut:"zoom-out"}[m]).attr({width:a.width,height:a.height,title:d.options.lang[m],padding:a.padding,zIndex:5}).add();k.handler=a.onclick;b(k.element,"dblclick",l);c.push(k);e(a,{width:k.width,height:2*k.height});if(d.hasLoaded)k.align(a,!1,a.alignTo);else var w=b(d,"load",function(){k.element&&k.align(a,!1,a.alignTo);w()})});this.updateEvents(f)};n.prototype.updateEvents=function(a){var d=this.chart;k(a.enableDoubleClickZoom,
a.enabled)||a.enableDoubleClickZoomTo?this.unbindDblClick=this.unbindDblClick||b(d.container,"dblclick",function(a){d.pointer.onContainerDblClick(a)}):this.unbindDblClick&&(this.unbindDblClick=this.unbindDblClick());k(a.enableMouseWheelZoom,a.enabled)?this.unbindMouseWheel=this.unbindMouseWheel||b(d.container,void 0!==h.onwheel?"wheel":void 0!==h.onmousewheel?"mousewheel":"DOMMouseScroll",function(a){d.pointer.inClass(a.target,"highcharts-no-mousewheel")||(d.pointer.onContainerMouseWheel(a),l(a));
return!1}):this.unbindMouseWheel&&(this.unbindMouseWheel=this.unbindMouseWheel())};e(c.prototype,{fitToBox:function(a,d){[["x","width"],["y","height"]].forEach(function(f){var e=f[0];f=f[1];a[e]+a[f]>d[e]+d[f]&&(a[f]>d[f]?(a[f]=d[f],a[e]=d[e]):a[e]=d[e]+d[f]-a[f]);a[f]>d[f]&&(a[f]=d[f]);a[e]<d[e]&&(a[e]=d[e])});return a},mapZoom:function(a,d,f,e,b){this.mapView&&("number"===typeof a&&(a=Math.log(a)/Math.log(.5)),this.mapView.zoomBy(a,"number"===typeof d&&"number"===typeof f?this.mapView.projection.inverse([d,
f]):void 0,"number"===typeof e&&"number"===typeof b?[e,b]:void 0))}});b(c,"beforeRender",function(){this.mapNavigation=new n(this);this.mapNavigation.update()});p.MapNavigation=n});x(c,"Maps/MapPointer.js",[c["Core/Pointer.js"],c["Core/Utilities.js"]],function(c,p){var q=p.defined,l=p.extend,n=p.pick;p=p.wrap;var h=0,b;l(c.prototype,{onContainerDblClick:function(e){var b=this.chart;e=this.normalize(e);b.options.mapNavigation.enableDoubleClickZoomTo?b.pointer.inClass(e.target,"highcharts-tracker")&&
b.hoverPoint&&b.hoverPoint.zoomTo():b.isInsidePlot(e.chartX-b.plotLeft,e.chartY-b.plotTop)&&b.mapZoom(.5,void 0,void 0,e.chartX,e.chartY)},onContainerMouseWheel:function(e){var g=this.chart;e=this.normalize(e);var t=q(e.wheelDelta)&&-e.wheelDelta/120||e.deltaY||e.detail;1<=Math.abs(t)&&(h+=Math.abs(t),b&&clearTimeout(b),b=setTimeout(function(){h=0},50));10>h&&g.isInsidePlot(e.chartX-g.plotLeft,e.chartY-g.plotTop)&&g.mapView&&g.mapView.zoomBy((g.options.mapNavigation.mouseWheelSensitivity-1)*-t,void 0,
[e.chartX,e.chartY],1>Math.abs(t)?!1:void 0)}});p(c.prototype,"zoomOption",function(e){var b=this.chart.options.mapNavigation;n(b.enableTouchZoom,b.enabled)&&(this.chart.options.chart.pinchType="xy");e.apply(this,[].slice.call(arguments,1))});p(c.prototype,"pinchTranslate",function(e,b,t,k,a,d,f){e.call(this,b,t,k,a,d,f);"map"===this.chart.options.chart.type&&this.hasZoom&&(e=k.scaleX>k.scaleY,this.pinchTranslateDirection(!e,b,t,k,a,d,f,e?k.scaleX:k.scaleY))})});x(c,"Series/ColorMapComposition.js",
[c["Core/Utilities.js"]],function(c){var p=c.defined,q=c.wrap,l={pointArrayMap:["value"],axisTypes:["xAxis","yAxis","colorAxis"],trackerGroups:["group","markerGroup","dataLabelsGroup"],parallelArrays:["x","y","value"],colorKey:"value"},n;(function(c){function b(){return null!==this.value&&Infinity!==this.value&&-Infinity!==this.value}function e(b){var a={};!p(b.color)||b.state&&"normal"!==b.state||(a[this.colorProp||"fill"]=b.color);return a}function g(b,a,d){b=b.call(this,a,d);a.moveToTopOnHover&&
(b.zIndex="hover"===d?1:0);return b}c.colorMapSeriesMixin=l;var t=[];c.compose=function(k,a){a&&-1===t.indexOf(a)&&(t.push(a),a=a.prototype,a.dataLabelOnNull=!0,a.moveToTopOnHover=!0,a.isValid=b);-1===t.indexOf(k)&&(t.push(k),a=k.prototype,a.colorAttribs=e,q(a,"pointAttribs",g));return k};c.seriesColorAttribs=e})(n||(n={}));return n});x(c,"Maps/MapViewOptionsDefault.js",[],function(){return{center:[0,0],maxZoom:void 0,padding:0,projection:void 0,zoom:void 0}});x(c,"Maps/Projections/LambertConformalConic.js",
[],function(){var c=Math.sign||function(c){return 0===c?0:0<c?1:-1},p=Math.PI/180,q=Math.PI/2,l=0,n=0;return{init:function(h){var b=(h.parallels||[]).map(function(b){return b*p});h=b[0]||0;b=b[1]||h;var e=Math.cos(h);l=h===b?Math.sin(h):Math.log(e/Math.cos(b))/Math.log(Math.tan((q+b)/2)/Math.tan((q+h)/2));1e-10>Math.abs(l)&&(l=1e-10*(c(l)||1));n=e*Math.pow(Math.tan((q+h)/2),l)/l},forward:function(c){var b=c[0]*p;c=c[1]*p;0<n?c<-q+.000001&&(c=-q+.000001):c>q-.000001&&(c=q-.000001);c=n/Math.pow(Math.tan((q+
c)/2),l);return[c*Math.sin(l*b)*63.78137,63.78137*(n-c*Math.cos(l*b))]},inverse:function(h){var b=h[0]/63.78137;h=n-h[1]/63.78137;var e=c(l)*Math.sqrt(b*b+h*h),g=Math.atan2(b,Math.abs(h))*c(h);0>h*l&&(g-=Math.PI*c(b)*c(h));return[g/l/p,(2*Math.atan(Math.pow(n/e,1/l))-q)/p]}}});x(c,"Maps/Projections/EqualEarth.js",[],function(){var c=Math.sqrt(3)/2;return{forward:function(p){var q=Math.PI/180,l=Math.asin(c*Math.sin(p[1]*q)),n=l*l,h=n*n*n;return[p[0]*q*Math.cos(l)*74.03120656864502/(c*(1.340264+3*-.081106*
n+h*(7*.000893+.034164*n))),74.03120656864502*l*(1.340264+-.081106*n+h*(.000893+.003796*n))]},inverse:function(p){var q=p[0]/74.03120656864502;p=p[1]/74.03120656864502;var l=180/Math.PI,n=p,h;for(h=0;12>h;++h){var b=n*n;var e=b*b*b;var g=n*(1.340264+-.081106*b+e*(.000893+.003796*b))-p;b=1.340264+3*-.081106*b+e*(7*.000893+.034164*b);n-=g/=b;if(1e-9>Math.abs(g))break}b=n*n;return[l*c*q*(1.340264+3*-.081106*b+b*b*b*(7*.000893+.034164*b))/Math.cos(n),l*Math.asin(Math.sin(n)/c)]}}});x(c,"Maps/Projections/Miller.js",
[],function(){var c=Math.PI/4,p=Math.PI/180;return{forward:function(q){return[q[0]*p*63.78137,79.7267125*Math.log(Math.tan(c+.4*q[1]*p))]},inverse:function(q){return[q[0]/63.78137/p,2.5*(Math.atan(Math.exp(q[1]/63.78137*.8))-c)/p]}}});x(c,"Maps/Projections/Orthographic.js",[],function(){var c=Math.PI/180;return{forward:function(p){var q=p[0];if(-90>q||90<q)return[NaN,NaN];p=p[1]*c;return[Math.cos(p)*Math.sin(q*c)*63.78460826781007,63.78460826781007*Math.sin(p)]},inverse:function(p){var q=p[0]/63.78460826781007;
p=p[1]/63.78460826781007;var l=Math.sqrt(q*q+p*p),n=Math.asin(l),h=Math.sin(n);return[Math.atan2(q*h,l*Math.cos(n))/c,Math.asin(l&&p*h/l)/c]}}});x(c,"Maps/Projections/WebMercator.js",[],function(){var c=Math.PI/180;return{forward:function(p){if(85.0511287798<Math.abs(p[1]))return[NaN,NaN];var q=Math.sin(p[1]*c);return[63.78137*p[0]*c,63.78137*Math.log((1+q)/(1-q))/2]},inverse:function(p){return[p[0]/(63.78137*c),(2*Math.atan(Math.exp(p[1]/63.78137))-Math.PI/2)/c]},maxLatitude:85.0511287798}});x(c,
"Maps/Projections/ProjectionRegistry.js",[c["Maps/Projections/LambertConformalConic.js"],c["Maps/Projections/EqualEarth.js"],c["Maps/Projections/Miller.js"],c["Maps/Projections/Orthographic.js"],c["Maps/Projections/WebMercator.js"]],function(c,p,q,l,n){return{EqualEarth:p,LambertConformalConic:c,Miller:q,Orthographic:l,WebMercator:n}});x(c,"Maps/Projection.js",[c["Maps/Projections/ProjectionRegistry.js"],c["Core/Utilities.js"]],function(c,p){var q=this&&this.__spreadArrays||function(){for(var b=0,
e=0,c=arguments.length;e<c;e++)b+=arguments[e].length;b=Array(b);var t=0;for(e=0;e<c;e++)for(var k=arguments[e],a=0,d=k.length;a<d;a++,t++)b[t]=k[a];return b},l=p.erase,n=2*Math.PI/360,h=function(b){-180>b&&(b+=360);180<b&&(b-=360);return b};return function(){function b(e){void 0===e&&(e={});this.hasGeoProjection=this.hasCoordinates=!1;this.maxLatitude=90;this.options=e;var c=e.name,t=e.rotation;this.rotator=t?this.getRotator(t):void 0;var k=this.def=c?b.registry[c]:void 0,a=this.rotator;k&&(k.init&&
k.init(e),this.maxLatitude=k.maxLatitude||90,this.hasGeoProjection=!0);a&&k?(this.forward=function(d){d=a.forward(d);return k.forward(d)},this.inverse=function(d){d=k.inverse(d);return a.inverse(d)}):k?(this.forward=k.forward,this.inverse=k.inverse):a&&(this.forward=a.forward,this.inverse=a.inverse)}b.add=function(e,c){b.registry[e]=c};b.greatCircle=function(b,c,t){var e=Math.atan2,a=Math.cos,d=Math.sin,f=Math.sqrt,r=b[1]*n,g=b[0]*n,h=c[1]*n,u=c[0]*n,v=h-r,B=u-g;v=d(v/2)*d(v/2)+a(r)*a(h)*d(B/2)*d(B/
2);v=2*e(f(v),f(1-v));var w=Math.round(6371E3*v/5E5);B=[];t&&B.push(b);if(1<w)for(w=b=1/w;.999>w;w+=b){var m=d((1-w)*v)/d(v),A=d(w*v)/d(v),l=m*a(r)*a(g)+A*a(h)*a(u),q=m*a(r)*d(g)+A*a(h)*d(u);m=m*d(r)+A*d(h);m=e(m,f(l*l+q*q));l=e(q,l);B.push([l/n,m/n])}t&&B.push(c);return B};b.insertGreatCircles=function(e){for(var c=e.length-1;c--;)if(10<Math.max(Math.abs(e[c][0]-e[c+1][0]),Math.abs(e[c][1]-e[c+1][1]))){var t=b.greatCircle(e[c],e[c+1]);t.length&&e.splice.apply(e,q([c+1,0],t))}};b.toString=function(b){b=
b||{};var e=b.rotation;return[b.name,e&&e.join(",")].join(";")};b.prototype.getRotator=function(b){var e=b[0]*n,c=(b[1]||0)*n;b=(b[2]||0)*n;var k=Math.cos(c),a=Math.sin(c),d=Math.cos(b),f=Math.sin(b);if(0!==e||0!==c||0!==b)return{forward:function(b){var c=b[0]*n+e,r=b[1]*n,u=Math.cos(r);b=Math.cos(c)*u;c=Math.sin(c)*u;r=Math.sin(r);u=r*k+b*a;return[Math.atan2(c*d-u*f,b*k-r*a)/n,Math.asin(u*d+c*f)/n]},inverse:function(b){var c=b[0]*n,r=b[1]*n,u=Math.cos(r);b=Math.cos(c)*u;c=Math.sin(c)*u;r=Math.sin(r);
u=r*d-c*f;return[(Math.atan2(c*d+r*f,b*k+u*a)-e)/n,Math.asin(u*k-b*a)/n]}}};b.prototype.forward=function(b){return b};b.prototype.inverse=function(b){return b};b.prototype.clipOnAntimeridian=function(e,c){var g=[],k=[e];e.forEach(function(a,d){var b=e[d-1];if(!d){if(!c)return;b=e[e.length-1]}var f=b[0],m=a[0];(-90>f||90<f)&&(-90>m||90<m)&&0<f!==0<m&&g.push({i:d,lat:b[1]+(180-b[0])/(a[0]-b[0])*(a[1]-b[1]),direction:0>f?1:-1,previousLonLat:b,lonLat:a})});if(g.length)if(c){if(1===g.length%2){var a=g.slice().sort(function(a,
d){return Math.abs(d.lat)-Math.abs(a.lat)})[0];l(g,a)}for(var d=g.length-2;0<=d;){var f=g[d].i,r=h(180+.000001*g[d].direction),y=h(180-.000001*g[d].direction);f=e.splice.apply(e,q([f,g[d+1].i-f],b.greatCircle([r,g[d].lat],[r,g[d+1].lat],!0)));f.push.apply(f,b.greatCircle([y,g[d+1].lat],[y,g[d].lat],!0));k.push(f);d-=2}if(a)for(f=0;f<k.length;f++)if(d=k[f],y=d.indexOf(a.lonLat),-1<y){f=(0>a.lat?-1:1)*this.maxLatitude;r=h(180+.000001*a.direction);var n=h(180-.000001*a.direction);a=b.greatCircle([r,
a.lat],[r,f],!0).concat(b.greatCircle([n,f],[n,a.lat],!0));d.splice.apply(d,q([y,0],a));break}}else for(d=g.length;d--;)f=g[d].i,f=e.splice(f,e.length,[h(180+.000001*g[d].direction),g[d].lat]),f.unshift([h(180-.000001*g[d].direction),g[d].lat]),k.push(f);return k};b.prototype.path=function(e){var c=this,h=this.def,k=this.rotator,a=[],d="Polygon"===e.type||"MultiPolygon"===e.type,f=this.hasGeoProjection,r="Orthographic"!==this.options.name,y=r?k:void 0,l=r?h||this:this,u=function(e){e=e.map(function(a){if(r){y&&
(a=y.forward(a));var d=a[0];.000001>Math.abs(d-180)&&(d=180>d?179.999999:180.000001);a=[d,a[1]]}return a});var k=[e];f&&(b.insertGreatCircles(e),r&&(k=c.clipOnAntimeridian(e,d)));k.forEach(function(e){if(!(2>e.length))for(var m=!1,r,k,g=!1,v=function(d){m?a.push(["L",d[0],d[1]]):(a.push(["M",d[0],d[1]]),m=!0)},u=0;u<e.length;u++){var w=e[u],y=l.forward(w);isNaN(y[0])||isNaN(y[1])||f&&!(w[1]<=c.maxLatitude&&w[1]>=-c.maxLatitude)?g=!0:(d&&!r&&(r=w,e.push(w)),g&&k&&(d&&f?b.greatCircle(k,w).forEach(function(a){return v(l.forward(a))}):
m=!1),v(y),k=w,g=!1)}})};"LineString"===e.type?u(e.coordinates):"MultiLineString"===e.type?e.coordinates.forEach(function(a){return u(a)}):"Polygon"===e.type?(e.coordinates.forEach(function(a){return u(a)}),a.length&&a.push(["Z"])):"MultiPolygon"===e.type&&(e.coordinates.forEach(function(a){a.forEach(function(a){return u(a)})}),a.length&&a.push(["Z"]));return a};b.registry=c;return b}()});x(c,"Maps/MapView.js",[c["Maps/MapViewOptionsDefault.js"],c["Maps/Projection.js"],c["Core/Utilities.js"]],function(c,
p,q){var l=q.addEvent,n=q.clamp,h=q.fireEvent,b=q.isNumber,e=q.merge,g=q.pick,t=q.relativeLength;return function(){function k(a,d){var f=this;this.userOptions=d||{};d=e(c,d);this.chart=a;this.center=d.center;this.options=d;this.projection=new p(d.projection);this.zoom=d.zoom||0;l(a,"afterSetChartSize",function(){if(void 0===f.minZoom||f.minZoom===f.zoom)f.fitToBounds(void 0,void 0,!1),b(f.userOptions.zoom)&&(f.zoom=f.userOptions.zoom),f.userOptions.center&&e(!0,f.center,f.userOptions.center)});var r,
k,g;d=function(d){var b=a.pointer.pinchDown,e=a.mouseDownX,c=a.mouseDownY;1===b.length&&(e=b[0].chartX,c=b[0].chartY);if("number"===typeof e&&"number"===typeof c){var m=e+","+c,u=d.originalEvent;b=u.chartX;u=u.chartY;m!==k&&(k=m,r=f.projection.forward(f.center),g=(f.projection.options.rotation||[0,0]).slice());"Orthographic"===f.projection.options.name&&3>(f.minZoom||Infinity)?(m=440/(f.getScale()*Math.min(a.plotWidth,a.plotHeight)),g&&(e=(e-b)*m-g[0],c=n(-g[1]-(c-u)*m,-80,80),f.update({projection:{rotation:[-e,
-c]},center:[e,c],zoom:f.zoom},!0,!1))):(m=f.getScale(),c=f.projection.inverse([r[0]+(e-b)/m,r[1]-(c-u)/m]),f.setView(c,void 0,!0,!1));d.preventDefault()}};l(a,"pan",d);l(a,"touchpan",d);l(a,"selection",function(d){if(d.resetSelection)f.zoomBy();else{var b=d.x-a.plotLeft,e=d.y-a.plotTop,c=f.pixelsToProjectedUnits({x:b,y:e}),m=c.y;c=c.x;b=f.pixelsToProjectedUnits({x:b+d.width,y:e+d.height});f.fitToBounds({x1:c,y1:m,x2:b.x,y2:b.y},void 0,!0,d.originalEvent.touches?!1:void 0);/^touch/.test(d.originalEvent.type)||
a.showResetZoom();d.preventDefault()}})}k.prototype.fitToBounds=function(a,d,b,e){void 0===b&&(b=!0);var f=a||this.getProjectedBounds();if(f){var c=this.chart,r=c.plotWidth;c=c.plotHeight;var k=g(d,a?0:this.options.padding);d=t(k,r);k=t(k,c);r=Math.log(400.979322/Math.max((f.x2-f.x1)/((r-d)/256),(f.y2-f.y1)/((c-k)/256)))/Math.log(2);a||(this.minZoom=r);a=this.projection.inverse([(f.x2+f.x1)/2,(f.y2+f.y1)/2]);this.setView(a,r,b,e)}};k.prototype.getProjectedBounds=function(){var a=this.chart.series.reduce(function(a,
b){(b=b.getProjectedBounds&&b.getProjectedBounds())&&a.push(b);return a},[]);return k.compositeBounds(a)};k.prototype.getScale=function(){return 256/400.979322*Math.pow(2,this.zoom)};k.prototype.redraw=function(a){this.chart.series.forEach(function(a){a.useMapGeometry&&(a.isDirty=!0)});this.chart.redraw(a)};k.prototype.setView=function(a,d,b,e){void 0===b&&(b=!0);var f=!1;a&&(this.center=a);"number"===typeof d&&("number"===typeof this.minZoom&&(d=Math.max(d,this.minZoom)),"number"===typeof this.options.maxZoom&&
(d=Math.min(d,this.options.maxZoom)),f=d>this.zoom,this.zoom=d);var c=this.getProjectedBounds();if(c&&!f){f=this.projection.forward(this.center);d=this.chart;a=d.plotWidth;d=d.plotHeight;var r=this.getScale(),k=this.projectedUnitsToPixels({x:c.x1,y:c.y1}),g=this.projectedUnitsToPixels({x:c.x2,y:c.y2});c=[(c.x1+c.x2)/2,(c.y1+c.y2)/2];var w=k.x,m=g.y;g=g.x;k=k.y;g-w<a?f[0]=c[0]:0>w&&g<a?f[0]+=Math.max(w,g-a)/r:g>a&&0<w&&(f[0]+=Math.min(g-a,w)/r);k-m<d?f[1]=c[1]:0>m&&k<d?f[1]-=Math.max(m,k-d)/r:k>d&&
0<m&&(f[1]-=Math.min(k-d,m)/r);this.center=this.projection.inverse(f)}h(this,"afterSetView");b&&this.redraw(e)};k.prototype.projectedUnitsToPixels=function(a){var d=this.getScale(),b=this.projection.forward(this.center);return{x:this.chart.plotWidth/2-d*(b[0]-a.x),y:this.chart.plotHeight/2+d*(b[1]-a.y)}};k.prototype.pixelsToProjectedUnits=function(a){var d=a.x;a=a.y;var b=this.getScale(),e=this.projection.forward(this.center);return{x:e[0]+(d-this.chart.plotWidth/2)/b,y:e[1]-(a-this.chart.plotHeight/
2)/b}};k.prototype.update=function(a,d,f){void 0===d&&(d=!0);var c=a.projection;c=c&&p.toString(c)!==p.toString(this.options.projection);e(!0,this.userOptions,a);e(!0,this.options,a);c&&(this.chart.series.forEach(function(a){a.clearBounds&&a.clearBounds();a.isDirty=!0;a.isDirtyData=!0}),this.projection=new p(this.options.projection),a.center||b(a.zoom)||this.fitToBounds(void 0,void 0,!1));(a.center||b(a.zoom))&&this.setView(this.options.center,a.zoom,!1);d&&this.chart.redraw(f)};k.prototype.zoomBy=
function(a,d,b,e){var f=this.chart,c=this.projection.forward(this.center);d=d?this.projection.forward(d):[];var k=d[0],g=d[1];"number"===typeof a?(a=this.zoom+a,d=void 0,b&&(k=b[0],g=b[1],b=this.getScale(),k=k-f.plotLeft-f.plotWidth/2,f=g-f.plotTop-f.plotHeight/2,k=c[0]+k/b,g=c[1]+f/b),"number"===typeof k&&"number"===typeof g&&(b=1-Math.pow(2,this.zoom)/Math.pow(2,a),k=c[0]-k,f=c[1]-g,c[0]-=k*b,c[1]+=f*b,d=this.projection.inverse(c)),this.setView(d,a,void 0,e)):this.fitToBounds(void 0,void 0,void 0,
e)};k.compositeBounds=function(a){if(a.length)return a.slice(1).reduce(function(a,b){a.x1=Math.min(a.x1,b.x1);a.y1=Math.min(a.y1,b.y1);a.x2=Math.max(a.x2,b.x2);a.y2=Math.max(a.y2,b.y2);return a},e(a[0]))};return k}()});x(c,"Maps/MapSymbols.js",[c["Core/Renderer/SVG/SVGRenderer.js"]],function(c){function p(c,l,n,h,b,e,g,t){return[["M",c+b,l],["L",c+n-e,l],["C",c+n-e/2,l,c+n,l+e/2,c+n,l+e],["L",c+n,l+h-g],["C",c+n,l+h-g/2,c+n-g/2,l+h,c+n-g,l+h],["L",c+t,l+h],["C",c+t/2,l+h,c,l+h-t/2,c,l+h-t],["L",c,
l+b],["C",c,l+b/2,c+b/2,l,c+b,l],["Z"]]}c=c.prototype.symbols;c.bottombutton=function(c,l,n,h,b){b=b&&b.r||0;return p(c-1,l-1,n,h,0,0,b,b)};c.topbutton=function(c,l,n,h,b){b=b&&b.r||0;return p(c-1,l-1,n,h,b,b,0,0)};return c});x(c,"Core/Chart/MapChart.js",[c["Core/Chart/Chart.js"],c["Core/DefaultOptions.js"],c["Maps/MapView.js"],c["Core/Renderer/SVG/SVGRenderer.js"],c["Core/Utilities.js"]],function(c,p,q,l,n){var h=this&&this.__extends||function(){var b=function(a,d){b=Object.setPrototypeOf||{__proto__:[]}instanceof
Array&&function(a,b){a.__proto__=b}||function(a,b){for(var d in b)b.hasOwnProperty(d)&&(a[d]=b[d])};return b(a,d)};return function(a,d){function c(){this.constructor=a}b(a,d);a.prototype=null===d?Object.create(d):(c.prototype=d.prototype,new c)}}(),b=p.getOptions,e=n.addEvent,g=n.merge,t=n.pick;c=function(c){function a(){return null!==c&&c.apply(this,arguments)||this}h(a,c);a.prototype.init=function(a,f){e(this,"afterInit",function(){this.mapView=new q(this,this.options.mapView)});var d=b().credits;
a=g({chart:{panning:{enabled:!0,type:"xy"},type:"map"},credits:{mapText:t(d.mapText,' \u00a9 <a href="{geojson.copyrightUrl}">{geojson.copyrightShort}</a>'),mapTextFull:t(d.mapTextFull,"{geojson.copyright}")},mapView:{},tooltip:{followTouchMove:!1}},a);c.prototype.init.call(this,a,f)};return a}(c);(function(b){b.maps={};b.mapChart=function(a,d,c){return new b(a,d,c)};b.splitPath=function(a){"string"===typeof a&&(a=a.replace(/([A-Za-z])/g," $1 ").replace(/^\s*/,"").replace(/\s*$/,""),a=a.split(/[ ,;]+/).map(function(a){return/[A-za-z]/.test(a)?
a:parseFloat(a)}));return l.prototype.pathToSegments(a)}})(c||(c={}));return c});x(c,"Series/Map/MapPoint.js",[c["Core/Series/SeriesRegistry.js"],c["Core/Utilities.js"]],function(c,p){var q=this&&this.__extends||function(){var c=function(h,b){c=Object.setPrototypeOf||{__proto__:[]}instanceof Array&&function(b,c){b.__proto__=c}||function(b,c){for(var e in c)c.hasOwnProperty(e)&&(b[e]=c[e])};return c(h,b)};return function(h,b){function e(){this.constructor=h}c(h,b);h.prototype=null===b?Object.create(b):
(e.prototype=b.prototype,new e)}}(),l=p.extend;return function(c){function h(){var b=null!==c&&c.apply(this,arguments)||this;b.options=void 0;b.path=void 0;b.series=void 0;return b}q(h,c);h.getProjectedPath=function(b,c){b.projectedPath||(c&&b.geometry?(c.hasCoordinates=!0,b.projectedPath=c.path(b.geometry)):b.projectedPath=b.path);return b.projectedPath||[]};h.prototype.applyOptions=function(b,e){var g=this.series;b=c.prototype.applyOptions.call(this,b,e);e=g.joinBy;g.mapData&&g.mapMap&&(e=c.prototype.getNestedProperty.call(b,
e[1]),(g="undefined"!==typeof e&&g.mapMap[e])?l(b,g):b.value=b.value||null);return b};h.prototype.onMouseOver=function(b){p.clearTimeout(this.colorInterval);if(null!==this.value||this.series.options.nullInteraction)c.prototype.onMouseOver.call(this,b);else this.series.onMouseOut(b)};h.prototype.zoomTo=function(){var b=this.series.chart;b.mapView&&this.bounds&&(b.mapView.fitToBounds(this.bounds,void 0,!1),this.series.isDirty=!0,b.redraw())};return h}(c.seriesTypes.scatter.prototype.pointClass)});x(c,
"Series/Map/MapSeries.js",[c["Core/Animation/AnimationUtilities.js"],c["Series/ColorMapComposition.js"],c["Series/CenteredUtilities.js"],c["Core/Globals.js"],c["Core/Legend/LegendSymbol.js"],c["Core/Chart/MapChart.js"],c["Series/Map/MapPoint.js"],c["Maps/MapView.js"],c["Core/Series/Series.js"],c["Core/Series/SeriesRegistry.js"],c["Core/Renderer/SVG/SVGRenderer.js"],c["Core/Utilities.js"]],function(c,p,q,l,n,h,b,e,g,t,k,a){var d=this&&this.__extends||function(){var a=function(b,c){a=Object.setPrototypeOf||
{__proto__:[]}instanceof Array&&function(a,b){a.__proto__=b}||function(a,b){for(var c in b)b.hasOwnProperty(c)&&(a[c]=b[c])};return a(b,c)};return function(b,c){function d(){this.constructor=b}a(b,c);b.prototype=null===c?Object.create(c):(d.prototype=c.prototype,new d)}}(),f=c.animObject;c=p.colorMapSeriesMixin;var r=l.noop,y=h.maps,z=h.splitPath;h=t.seriesTypes;var u=h.column,v=h.scatter;h=a.extend;var B=a.fireEvent,w=a.getNestedProperty,m=a.isArray,A=a.isNumber,E=a.merge,F=a.objectEach,D=a.pick,
G=a.splat;a=function(a){function c(){var b=null!==a&&a.apply(this,arguments)||this;b.chart=void 0;b.data=void 0;b.group=void 0;b.joinBy=void 0;b.options=void 0;b.points=void 0;b.transformGroup=void 0;return b}d(c,a);c.prototype.animate=function(a){var b=this.chart,c=this.group,d=f(this.options.animation);b.renderer.isSVG&&(a?c.attr({translateX:b.plotLeft+b.plotWidth/2,translateY:b.plotTop+b.plotHeight/2,scaleX:.001,scaleY:.001}):c.animate({translateX:b.plotLeft,translateY:b.plotTop,scaleX:1,scaleY:1},
d))};c.prototype.animateDrilldown=function(a){var b=this.chart,c=this.group;b.renderer.isSVG&&(a?c.attr({translateX:b.plotLeft+b.plotWidth/2,translateY:b.plotTop+b.plotHeight/2,scaleX:.1,scaleY:.1,opacity:.01}):c.animate({translateX:b.plotLeft,translateY:b.plotTop,scaleX:1,scaleY:1,opacity:1}))};c.prototype.animateDrillupFrom=function(){var a=this.chart;a.renderer.isSVG&&this.group.animate({translateX:a.plotLeft+a.plotWidth/2,translateY:a.plotTop+a.plotHeight/2,scaleX:.1,scaleY:.1,opacity:.01})};
c.prototype.animateDrillupTo=function(a){u.prototype.animateDrillupTo.call(this,a)};c.prototype.clearBounds=function(){this.points.forEach(function(a){delete a.bounds;delete a.projectedPath});delete this.bounds};c.prototype.doFullTranslate=function(){return!(!(this.isDirtyData||this.chart.isResizing||this.chart.renderer.isVML)&&this.hasRendered)};c.prototype.drawMapDataLabels=function(){g.prototype.drawDataLabels.call(this);this.dataLabelsGroup&&this.dataLabelsGroup.clip(this.chart.clipRect)};c.prototype.drawPoints=
function(){var a=this,b=this.chart,c=this.group,d=this.svgTransform,f=b.mapView,e=b.renderer;this.transformGroup||(this.transformGroup=e.g().add(c),this.transformGroup.survive=!0);this.doFullTranslate()&&(b.hasRendered&&!b.styledMode&&this.points.forEach(function(b){b.shapeArgs&&(b.shapeArgs.fill=a.pointAttribs(b,b.state).fill)}),this.group=this.transformGroup,u.prototype.drawPoints.apply(this),this.group=c,this.points.forEach(function(c){if(c.graphic){var d="";c.name&&(d+="highcharts-name-"+c.name.replace(/ /g,
"-").toLowerCase());c.properties&&c.properties["hc-key"]&&(d+=" highcharts-key-"+c.properties["hc-key"].toLowerCase());d&&c.graphic.addClass(d);b.styledMode&&c.graphic.css(a.pointAttribs(c,c.selected&&"select"||void 0))}}));if(f&&d){var m=D(this.options[this.pointAttrToOptions&&this.pointAttrToOptions["stroke-width"]||"borderWidth"],1),k=d.scaleX,g=0<d.scaleY?1:-1,r=this.transformGroup;if(e.globalAnimation&&b.hasRendered){var w=Number(r.attr("translateX")),h=Number(r.attr("translateY")),v=Number(r.attr("scaleX"));
r.attr({animator:0}).animate({animator:1},{step:function(a,b){a=v+(k-v)*b.pos;r.attr({translateX:w+(d.translateX-w)*b.pos,translateY:h+(d.translateY-h)*b.pos,scaleX:a,scaleY:a*g});c.element.setAttribute("stroke-width",m/a)}})}else r.attr(d),c.element.setAttribute("stroke-width",m/k)}this.drawMapDataLabels()};c.prototype.getProjectedBounds=function(){if(!this.bounds){var a=Number.MAX_VALUE,c=this.chart.mapView&&this.chart.mapView.projection,d=[];(this.points||[]).forEach(function(f){if(f.path||f.geometry){"string"===
typeof f.path?f.path=z(f.path):m(f.path)&&"M"===f.path[0]&&(f.path=k.prototype.pathToSegments(f.path));if(!f.bounds){var e=b.getProjectedPath(f,c),g=f.properties,r=-a,w=a,h=-a,u=a,v;e.forEach(function(a){var b=a[a.length-2];a=a[a.length-1];"number"===typeof b&&"number"===typeof a&&(w=Math.min(w,b),r=Math.max(r,b),u=Math.min(u,a),h=Math.max(h,a),v=!0)});v&&(e=g&&g["hc-middle-x"],e=w+(r-w)*D(f.middleX,A(e)?e:.5),g=g&&g["hc-middle-y"],g=D(f.middleY,A(g)?g:.5),f.geometry||(g=1-g),f.bounds={midX:e,midY:h-
(h-u)*g,x1:w,y1:u,x2:r,y2:h},f.labelrank=D(f.labelrank,(r-w)*(h-u)))}f.bounds&&d.push(f.bounds)}});this.bounds=e.compositeBounds(d)}return this.bounds};c.prototype.hasData=function(){return!!this.processedXData.length};c.prototype.pointAttribs=function(a,b){var c=a.series.chart,d=c.mapView;b=c.styledMode?this.colorAttribs(a):u.prototype.pointAttribs.call(this,a,b);(a=a.options[this.pointAttrToOptions&&this.pointAttrToOptions["stroke-width"]||"borderWidth"])&&d&&(a/=d.getScale());b.dashstyle&&d&&this.options.borderWidth&&
(a=this.options.borderWidth/d.getScale());b["stroke-width"]=D(a,"inherit");return b};c.prototype.setData=function(a,c,d,f){var e=this.options,k=this.chart.options.chart,r=k&&k.map,h=e.mapData,u=this.joinBy,v=e.keys||this.pointArrayMap,t=[],p={},n=this.chart.mapTransforms;!h&&r&&(h="string"===typeof r?y[r]:r);a&&a.forEach(function(c,d){var f=0;if(A(c))a[d]={value:c};else if(m(c)){a[d]={};!e.keys&&c.length>v.length&&"string"===typeof c[0]&&(a[d]["hc-key"]=c[0],++f);for(var k=0;k<v.length;++k,++f)v[k]&&
"undefined"!==typeof c[f]&&(0<v[k].indexOf(".")?b.prototype.setNestedProperty(a[d],c[f],v[k]):a[d][v[k]]=c[f])}u&&"_i"===u[0]&&(a[d]._i=d)});(this.chart.mapTransforms=n=k.mapTransforms||h&&h["hc-transform"]||n)&&F(n,function(a){a.rotation&&(a.cosAngle=Math.cos(a.rotation),a.sinAngle=Math.sin(a.rotation))});if(h){"FeatureCollection"===h.type&&(this.mapTitle=h.title,h=l.geojson(h,this.type,this));this.mapData=h;this.mapMap={};for(n=0;n<h.length;n++)k=h[n],r=k.properties,k._i=n,u[0]&&r&&r[u[0]]&&(k[u[0]]=
r[u[0]]),p[k[u[0]]]=k;this.mapMap=p;if(a&&u[1]){var q=u[1];a.forEach(function(a){a=w(q,a);p[a]&&t.push(p[a])})}if(e.allAreas){a=a||[];if(u[1]){var B=u[1];a.forEach(function(a){t.push(w(B,a))})}t="|"+t.map(function(a){return a&&a[u[0]]}).join("|")+"|";h.forEach(function(b){u[0]&&-1!==t.indexOf("|"+b[u[0]]+"|")||(a.push(E(b,{value:null})),f=!1)})}}g.prototype.setData.call(this,a,c,d,f);this.processData();this.generatePoints()};c.prototype.setOptions=function(a){a=g.prototype.setOptions.call(this,a);
var b=a.joinBy;null===b&&(b="_i");b=this.joinBy=G(b);b[1]||(b[1]=b[0]);return a};c.prototype.translate=function(){var a=this.doFullTranslate(),c=this.chart.mapView,d=c&&c.projection;!this.chart.hasRendered||!this.isDirtyData&&this.hasRendered||(this.processData(),this.generatePoints(),delete this.bounds,this.getProjectedBounds());var f;if(c){var e=c.getScale(),m=c.projection.forward(c.center);c=c.projection.hasCoordinates?-1:1;this.svgTransform=f={scaleX:e,scaleY:e*c,translateX:this.chart.plotWidth/
2-m[0]*e,translateY:this.chart.plotHeight/2-m[1]*e*c}}this.points.forEach(function(c){f&&c.bounds&&A(c.bounds.midX)&&A(c.bounds.midY)&&(c.plotX=c.bounds.midX*f.scaleX+f.translateX,c.plotY=c.bounds.midY*f.scaleY+f.translateY);a&&(c.shapeType="path",c.shapeArgs={d:b.getProjectedPath(c,d)})});B(this,"afterTranslate")};c.defaultOptions=E(v.defaultOptions,{animation:!1,dataLabels:{crop:!1,formatter:function(){var a=this.series.chart.numberFormatter,b=this.point.value;return A(b)?a(b,-1):""},inside:!0,
overflow:!1,padding:0,verticalAlign:"middle"},marker:null,nullColor:"#f7f7f7",stickyTracking:!1,tooltip:{followPointer:!0,pointFormat:"{point.name}: {point.value}<br/>"},turboThreshold:0,allAreas:!0,borderColor:"#cccccc",borderWidth:1,joinBy:"hc-key",states:{hover:{halo:null,brightness:.2},normal:{animation:!0},select:{color:"#cccccc"},inactive:{opacity:1}}});return c}(v);h(a.prototype,{type:"map",axisTypes:["colorAxis"],colorKey:"value",directTouch:!0,drawDataLabels:r,drawGraph:r,drawLegendSymbol:n.drawRectangle,
forceDL:!0,getCenter:q.getCenter,getExtremesFromAll:!0,getSymbol:r,isCartesian:!1,parallelArrays:c.parallelArrays,pointArrayMap:c.pointArrayMap,pointClass:b,preserveAspectRatio:!0,searchPoint:r,trackerGroups:c.trackerGroups,useMapGeometry:!0});p.compose(a,b);t.registerSeriesType("map",a);"";return a});x(c,"Series/MapLine/MapLineSeries.js",[c["Series/Map/MapSeries.js"],c["Core/Series/SeriesRegistry.js"],c["Core/Utilities.js"]],function(c,p,q){var l=this&&this.__extends||function(){var b=function(c,
e){b=Object.setPrototypeOf||{__proto__:[]}instanceof Array&&function(b,a){b.__proto__=a}||function(b,a){for(var c in a)a.hasOwnProperty(c)&&(b[c]=a[c])};return b(c,e)};return function(c,e){function k(){this.constructor=c}b(c,e);c.prototype=null===e?Object.create(e):(k.prototype=e.prototype,new k)}}(),n=p.series,h=q.extend,b=q.merge;q=function(e){function g(){var b=null!==e&&e.apply(this,arguments)||this;b.data=void 0;b.options=void 0;b.points=void 0;return b}l(g,e);g.prototype.pointAttribs=function(b,
e){b=c.prototype.pointAttribs.call(this,b,e);b.fill=this.options.fillColor;return b};g.defaultOptions=b(c.defaultOptions,{lineWidth:1,fillColor:"none"});return g}(c);h(q.prototype,{type:"mapline",colorProp:"stroke",drawLegendSymbol:n.prototype.drawLegendSymbol,pointAttrToOptions:{stroke:"color","stroke-width":"lineWidth"}});p.registerSeriesType("mapline",q);"";return q});x(c,"Series/MapPoint/MapPointPoint.js",[c["Core/Series/SeriesRegistry.js"],c["Core/Utilities.js"]],function(c,p){var q=this&&this.__extends||
function(){var c=function(b,e){c=Object.setPrototypeOf||{__proto__:[]}instanceof Array&&function(b,c){b.__proto__=c}||function(b,c){for(var e in c)c.hasOwnProperty(e)&&(b[e]=c[e])};return c(b,e)};return function(b,e){function g(){this.constructor=b}c(b,e);b.prototype=null===e?Object.create(e):(g.prototype=e.prototype,new g)}}(),l=p.isNumber,n=p.merge;return function(c){function b(){var b=null!==c&&c.apply(this,arguments)||this;b.options=void 0;b.series=void 0;return b}q(b,c);b.prototype.applyOptions=
function(b,g){b="undefined"!==typeof b.lat&&"undefined"!==typeof b.lon?n(b,this.series.chart.fromLatLonToPoint(b)):b;return c.prototype.applyOptions.call(this,b,g)};b.prototype.isValid=function(){return!!(this.options.geometry||l(this.x)&&l(this.y))};return b}(c.seriesTypes.scatter.prototype.pointClass)});x(c,"Series/MapPoint/MapPointSeries.js",[c["Core/Globals.js"],c["Series/MapPoint/MapPointPoint.js"],c["Core/Series/SeriesRegistry.js"],c["Core/Utilities.js"]],function(c,p,q,l){var n=this&&this.__extends||
function(){var b=function(a,c){b=Object.setPrototypeOf||{__proto__:[]}instanceof Array&&function(a,b){a.__proto__=b}||function(a,b){for(var c in b)b.hasOwnProperty(c)&&(a[c]=b[c])};return b(a,c)};return function(a,c){function d(){this.constructor=a}b(a,c);a.prototype=null===c?Object.create(c):(d.prototype=c.prototype,new d)}}();c=c.noop;var h=q.seriesTypes.scatter,b=l.extend,e=l.fireEvent,g=l.isNumber,t=l.merge;l=function(b){function a(){var a=null!==b&&b.apply(this,arguments)||this;a.data=void 0;
a.options=void 0;a.points=void 0;return a}n(a,b);a.prototype.drawDataLabels=function(){b.prototype.drawDataLabels.call(this);this.dataLabelsGroup&&this.dataLabelsGroup.clip(this.chart.clipRect)};a.prototype.translate=function(){var a=this,b=this.chart.mapView;this.processedXData||this.processData();this.generatePoints();if(b){var c=b.projection,k=c.forward,h=c.hasCoordinates;this.points.forEach(function(c){var d=c.x;d=void 0===d?void 0:d;var f=c.y;f=void 0===f?void 0:f;var e=c.options.geometry;(e=
e&&"Point"===e.type&&e.coordinates)?(f=k(e),d=f[0],f=f[1]):c.bounds&&(d=c.bounds.midX,f=c.bounds.midY);g(d)&&g(f)?(d=b.projectedUnitsToPixels({x:d,y:f}),c.plotX=d.x,c.plotY=h?d.y:a.chart.plotHeight-d.y):(c.plotX=void 0,c.plotY=void 0);c.isInside=a.isPointInside(c);c.zone=a.zones.length?c.getZone():void 0})}e(this,"afterTranslate")};a.defaultOptions=t(h.defaultOptions,{dataLabels:{crop:!1,defer:!1,enabled:!0,formatter:function(){return this.point.name},overflow:!1,style:{color:"#000000"}}});return a}(h);
b(l.prototype,{type:"mappoint",axisTypes:["colorAxis"],forceDL:!0,isCartesian:!1,pointClass:p,searchPoint:c,useMapGeometry:!0});q.registerSeriesType("mappoint",l);"";return l});x(c,"Series/Bubble/BubbleLegendDefaults.js",[],function(){return{borderColor:void 0,borderWidth:2,className:void 0,color:void 0,connectorClassName:void 0,connectorColor:void 0,connectorDistance:60,connectorWidth:1,enabled:!1,labels:{className:void 0,allowOverlap:!1,format:"",formatter:void 0,align:"right",style:{fontSize:"10px",
color:"#000000"},x:0,y:0},maxSize:60,minSize:10,legendIndex:0,ranges:{value:void 0,borderColor:void 0,color:void 0,connectorColor:void 0},sizeBy:"area",sizeByAbsoluteValue:!1,zIndex:1,zThreshold:0}});x(c,"Series/Bubble/BubbleLegendItem.js",[c["Core/Color/Color.js"],c["Core/FormatUtilities.js"],c["Core/Globals.js"],c["Core/Utilities.js"]],function(c,p,q,l){var n=c.parse,h=q.noop,b=l.arrayMax,e=l.arrayMin,g=l.isNumber,t=l.merge,k=l.pick,a=l.stableSort;"";return function(){function c(a,b){this.options=
this.symbols=this.visible=this.selected=this.ranges=this.movementX=this.maxLabel=this.legendSymbol=this.legendItemWidth=this.legendItemHeight=this.legendItem=this.legendGroup=this.legend=this.fontMetrics=this.chart=void 0;this.setState=h;this.init(a,b)}c.prototype.init=function(a,b){this.options=a;this.visible=!0;this.chart=b.chart;this.legend=b};c.prototype.addToLegend=function(a){a.splice(this.options.legendIndex,0,this)};c.prototype.drawLegendSymbol=function(b){var c=this.chart,d=this.options,
f=k(b.options.itemDistance,20),e=d.ranges,h=d.connectorDistance;this.fontMetrics=c.renderer.fontMetrics(d.labels.style.fontSize);e&&e.length&&g(e[0].value)?(a(e,function(a,b){return b.value-a.value}),this.ranges=e,this.setOptions(),this.render(),b=this.getMaxLabelSize(),e=this.ranges[0].radius,c=2*e,h=h-e+b.width,h=0<h?h:0,this.maxLabel=b,this.movementX="left"===d.labels.align?h:0,this.legendItemWidth=c+h+f,this.legendItemHeight=c+this.fontMetrics.h/2):b.options.bubbleLegend.autoRanges=!0};c.prototype.setOptions=
function(){var a=this.ranges,b=this.options,c=this.chart.series[b.seriesIndex],d=this.legend.baseline,e={zIndex:b.zIndex,"stroke-width":b.borderWidth},g={zIndex:b.zIndex,"stroke-width":b.connectorWidth},h={align:this.legend.options.rtl||"left"===b.labels.align?"right":"left",zIndex:b.zIndex},w=c.options.marker.fillOpacity,m=this.chart.styledMode;a.forEach(function(f,r){m||(e.stroke=k(f.borderColor,b.borderColor,c.color),e.fill=k(f.color,b.color,1!==w?n(c.color).setOpacity(w).get("rgba"):c.color),
g.stroke=k(f.connectorColor,b.connectorColor,c.color));a[r].radius=this.getRangeRadius(f.value);a[r]=t(a[r],{center:a[0].radius-a[r].radius+d});m||t(!0,a[r],{bubbleAttribs:t(e),connectorAttribs:t(g),labelAttribs:h})},this)};c.prototype.getRangeRadius=function(a){var b=this.options;return this.chart.series[this.options.seriesIndex].getRadius.call(this,b.ranges[b.ranges.length-1].value,b.ranges[0].value,b.minSize,b.maxSize,a)};c.prototype.render=function(){var a=this.chart.renderer,b=this.options.zThreshold;
this.symbols||(this.symbols={connectors:[],bubbleItems:[],labels:[]});this.legendSymbol=a.g("bubble-legend");this.legendItem=a.g("bubble-legend-item");this.legendSymbol.translateX=0;this.legendSymbol.translateY=0;this.ranges.forEach(function(a){a.value>=b&&this.renderRange(a)},this);this.legendSymbol.add(this.legendItem);this.legendItem.add(this.legendGroup);this.hideOverlappingLabels()};c.prototype.renderRange=function(a){var b=this.options,c=b.labels,d=this.chart,e=d.series[b.seriesIndex],f=d.renderer,
g=this.symbols;d=g.labels;var k=a.center,m=Math.abs(a.radius),h=b.connectorDistance||0,l=c.align,n=b.connectorWidth,p=this.ranges[0].radius||0,t=k-m-b.borderWidth/2+n/2,q=this.fontMetrics;q=q.f/2-(q.h-q.f)/2;var C=f.styledMode;h=this.legend.options.rtl||"left"===l?-h:h;"center"===l&&(h=0,b.connectorDistance=0,a.labelAttribs.align="center");l=t+b.labels.y;var x=p+h+b.labels.x;g.bubbleItems.push(f.circle(p,k+((t%1?1:.5)-(n%2?0:.5)),m).attr(C?{}:a.bubbleAttribs).addClass((C?"highcharts-color-"+e.colorIndex+
" ":"")+"highcharts-bubble-legend-symbol "+(b.className||"")).add(this.legendSymbol));g.connectors.push(f.path(f.crispLine([["M",p,t],["L",p+h,t]],b.connectorWidth)).attr(C?{}:a.connectorAttribs).addClass((C?"highcharts-color-"+this.options.seriesIndex+" ":"")+"highcharts-bubble-legend-connectors "+(b.connectorClassName||"")).add(this.legendSymbol));a=f.text(this.formatLabel(a),x,l+q).attr(C?{}:a.labelAttribs).css(C?{}:c.style).addClass("highcharts-bubble-legend-labels "+(b.labels.className||"")).add(this.legendSymbol);
d.push(a);a.placed=!0;a.alignAttr={x:x,y:l+q}};c.prototype.getMaxLabelSize=function(){var a,b;this.symbols.labels.forEach(function(c){b=c.getBBox(!0);a=a?b.width>a.width?b:a:b});return a||{}};c.prototype.formatLabel=function(a){var b=this.options,c=b.labels.formatter;b=b.labels.format;var d=this.chart.numberFormatter;return b?p.format(b,a):c?c.call(a):d(a.value,1)};c.prototype.hideOverlappingLabels=function(){var a=this.chart,b=this.symbols;!this.options.labels.allowOverlap&&b&&(a.hideOverlappingLabels(b.labels),
b.labels.forEach(function(a,c){a.newOpacity?a.newOpacity!==a.oldOpacity&&b.connectors[c].show():b.connectors[c].hide()}))};c.prototype.getRanges=function(){var a=this.legend.bubbleLegend,c=a.options.ranges,d,h=Number.MAX_VALUE,l=-Number.MAX_VALUE;a.chart.series.forEach(function(a){a.isBubble&&!a.ignoreSeries&&(d=a.zData.filter(g),d.length&&(h=k(a.options.zMin,Math.min(h,Math.max(e(d),!1===a.options.displayNegative?a.options.zThreshold:-Number.MAX_VALUE))),l=k(a.options.zMax,Math.max(l,b(d)))))});
var n=h===l?[{value:l}]:[{value:h},{value:(h+l)/2},{value:l,autoRanges:!0}];c.length&&c[0].radius&&n.reverse();n.forEach(function(a,b){c&&c[b]&&(n[b]=t(c[b],a))});return n};c.prototype.predictBubbleSizes=function(){var a=this.chart,b=this.fontMetrics,c=a.legend.options,d=c.floating,e=(c="horizontal"===c.layout)?a.legend.lastLineHeight:0,g=a.plotSizeX,k=a.plotSizeY,h=a.series[this.options.seriesIndex],m=h.getPxExtremes();a=Math.ceil(m.minPxSize);m=Math.ceil(m.maxPxSize);var l=Math.min(k,g);h=h.options.maxSize;
if(d||!/%$/.test(h))b=m;else if(h=parseFloat(h),b=(l+e-b.h/2)*h/100/(h/100+1),c&&k-b>=g||!c&&g-b>=k)b=m;return[a,Math.ceil(b)]};c.prototype.updateRanges=function(a,b){var c=this.legend.options.bubbleLegend;c.minSize=a;c.maxSize=b;c.ranges=this.getRanges()};c.prototype.correctSizes=function(){var a=this.legend,b=this.chart.series[this.options.seriesIndex].getPxExtremes();1<Math.abs(Math.ceil(b.maxPxSize)-this.options.maxSize)&&(this.updateRanges(this.options.minSize,b.maxPxSize),a.render())};return c}()});
x(c,"Series/Bubble/BubbleLegendComposition.js",[c["Series/Bubble/BubbleLegendDefaults.js"],c["Series/Bubble/BubbleLegendItem.js"],c["Core/DefaultOptions.js"],c["Core/Utilities.js"]],function(c,p,q,l){var n=q.setOptions,h=l.addEvent,b=l.objectEach,e=l.wrap,g;(function(g){function k(c,e,f){var g=this.legend,m=0<=a(this);if(g&&g.options.enabled&&g.bubbleLegend&&g.options.bubbleLegend.autoRanges&&m){var k=g.bubbleLegend.options;m=g.bubbleLegend.predictBubbleSizes();g.bubbleLegend.updateRanges(m[0],m[1]);
k.placed||(g.group.placed=!1,g.allItems.forEach(function(a){a.legendGroup.translateY=null}));g.render();this.getMargins();this.axes.forEach(function(a){a.visible&&a.render();k.placed||(a.setScale(),a.updateNames(),b(a.ticks,function(a){a.isNew=!0;a.isNewLabel=!0}))});k.placed=!0;this.getMargins();c.call(this,e,f);g.bubbleLegend.correctSizes();t(g,d(g))}else c.call(this,e,f),g&&g.options.enabled&&g.bubbleLegend&&(g.render(),t(g,d(g)))}function a(a){a=a.series;for(var b=0;b<a.length;){if(a[b]&&a[b].isBubble&&
a[b].visible&&a[b].zData.length)return b;b++}return-1}function d(a){a=a.allItems;var b=[],c=a.length,d,e=0;for(d=0;d<c;d++)if(a[d].legendItemHeight&&(a[d].itemHeight=a[d].legendItemHeight),a[d]===a[c-1]||a[d+1]&&a[d]._legendItemPos[1]!==a[d+1]._legendItemPos[1]){b.push({height:0});var f=b[b.length-1];for(e;e<=d;e++)a[e].itemHeight>f.height&&(f.height=a[e].itemHeight);f.step=d}return b}function f(b){var c=this.bubbleLegend,d=this.options,e=d.bubbleLegend,f=a(this.chart);c&&c.ranges&&c.ranges.length&&
(e.ranges.length&&(e.autoRanges=!!e.ranges[0].autoRanges),this.destroyItem(c));0<=f&&d.enabled&&e.enabled&&(e.seriesIndex=f,this.bubbleLegend=new p(e,this),this.bubbleLegend.addToLegend(b.allItems))}function l(){var b=this.chart,c=this.visible,d=this.chart.legend;d&&d.bubbleLegend&&(this.visible=!c,this.ignoreSeries=c,b=0<=a(b),d.bubbleLegend.visible!==b&&(d.update({bubbleLegend:{enabled:b}}),d.bubbleLegend.visible=b),this.visible=c)}function t(a,b){var c=a.options.rtl,d,e,f,g=0;a.allItems.forEach(function(a,
m){d=a.legendGroup.translateX;e=a._legendItemPos[1];if((f=a.movementX)||c&&a.ranges)f=c?d-a.options.maxSize/2:d+f,a.legendGroup.attr({translateX:f});m>b[g].step&&g++;a.legendGroup.attr({translateY:Math.round(e+b[g].height/2)});a._legendItemPos[1]=e+b[g].height/2})}var q=[];g.compose=function(a,b,d){-1===q.indexOf(a)&&(q.push(a),n({legend:{bubbleLegend:c}}),e(a.prototype,"drawChartBox",k));-1===q.indexOf(b)&&(q.push(b),h(b,"afterGetAllItems",f));-1===q.indexOf(d)&&(q.push(d),h(d,"legendItemClick",
l))}})(g||(g={}));return g});x(c,"Series/Bubble/BubblePoint.js",[c["Core/Series/Point.js"],c["Core/Series/SeriesRegistry.js"],c["Core/Utilities.js"]],function(c,p,q){var l=this&&this.__extends||function(){var c=function(h,b){c=Object.setPrototypeOf||{__proto__:[]}instanceof Array&&function(b,c){b.__proto__=c}||function(b,c){for(var e in c)c.hasOwnProperty(e)&&(b[e]=c[e])};return c(h,b)};return function(h,b){function e(){this.constructor=h}c(h,b);h.prototype=null===b?Object.create(b):(e.prototype=
b.prototype,new e)}}();q=q.extend;p=function(n){function h(){var b=null!==n&&n.apply(this,arguments)||this;b.options=void 0;b.series=void 0;return b}l(h,n);h.prototype.haloPath=function(b){return c.prototype.haloPath.call(this,0===b?0:(this.marker?this.marker.radius||0:0)+b)};return h}(p.seriesTypes.scatter.prototype.pointClass);q(p.prototype,{ttBelow:!1});return p});x(c,"Series/Bubble/BubbleSeries.js",[c["Core/Axis/Axis.js"],c["Series/Bubble/BubbleLegendComposition.js"],c["Series/Bubble/BubblePoint.js"],
c["Core/Color/Color.js"],c["Core/Globals.js"],c["Core/Series/Series.js"],c["Core/Series/SeriesRegistry.js"],c["Core/Utilities.js"]],function(c,p,q,l,n,h,b,e){var g=this&&this.__extends||function(){var a=function(b,c){a=Object.setPrototypeOf||{__proto__:[]}instanceof Array&&function(a,b){a.__proto__=b}||function(a,b){for(var c in b)b.hasOwnProperty(c)&&(a[c]=b[c])};return a(b,c)};return function(b,c){function d(){this.constructor=b}a(b,c);b.prototype=null===c?Object.create(c):(d.prototype=c.prototype,
new d)}}(),t=l.parse;l=n.noop;var k=b.seriesTypes;n=k.column;var a=k.scatter;k=e.addEvent;var d=e.arrayMax,f=e.arrayMin,r=e.clamp,y=e.extend,z=e.isNumber,u=e.merge,v=e.pick;e=function(b){function c(){var a=null!==b&&b.apply(this,arguments)||this;a.data=void 0;a.maxPxSize=void 0;a.minPxSize=void 0;a.options=void 0;a.points=void 0;a.radii=void 0;a.yData=void 0;a.zData=void 0;return a}g(c,b);c.prototype.animate=function(a){!a&&this.points.length<this.options.animationLimit&&this.points.forEach(function(a){var b=
a.graphic;b&&b.width&&(this.hasRendered||b.attr({x:a.plotX,y:a.plotY,width:1,height:1}),b.animate(this.markerAttribs(a),this.options.animation))},this)};c.prototype.getRadii=function(){var a=this,b=this.zData,c=this.yData,d=[],e=this.chart.bubbleZExtremes;var f=this.getPxExtremes();var g=f.minPxSize,k=f.maxPxSize;if(!e){var h=Number.MAX_VALUE,l=-Number.MAX_VALUE,r;this.chart.series.forEach(function(b){b.bubblePadding&&(b.visible||!a.chart.options.chart.ignoreHiddenSeries)&&(b=b.getZExtremes())&&(h=
Math.min(h||b.zMin,b.zMin),l=Math.max(l||b.zMax,b.zMax),r=!0)});r?(e={zMin:h,zMax:l},this.chart.bubbleZExtremes=e):e={zMin:0,zMax:0}}var n=0;for(f=b.length;n<f;n++){var p=b[n];d.push(this.getRadius(e.zMin,e.zMax,g,k,p,c[n]))}this.radii=d};c.prototype.getRadius=function(a,b,c,d,e,f){var m=this.options,g="width"!==m.sizeBy,k=m.zThreshold,h=b-a,l=.5;if(null===f||null===e)return null;if(z(e)){m.sizeByAbsoluteValue&&(e=Math.abs(e-k),h=Math.max(b-k,Math.abs(a-k)),a=0);if(e<a)return c/2-1;0<h&&(l=(e-a)/
h)}g&&0<=l&&(l=Math.sqrt(l));return Math.ceil(c+l*(d-c))/2};c.prototype.hasData=function(){return!!this.processedXData.length};c.prototype.pointAttribs=function(a,b){var c=this.options.marker.fillOpacity;a=h.prototype.pointAttribs.call(this,a,b);1!==c&&(a.fill=t(a.fill).setOpacity(c).get("rgba"));return a};c.prototype.translate=function(){b.prototype.translate.call(this);this.getRadii();this.translateBubble()};c.prototype.translateBubble=function(){for(var a=this.data,b=this.radii,c=this.getPxExtremes().minPxSize,
d=a.length;d--;){var e=a[d],f=b?b[d]:0;z(f)&&f>=c/2?(e.marker=y(e.marker,{radius:f,width:2*f,height:2*f}),e.dlBox={x:e.plotX-f,y:e.plotY-f,width:2*f,height:2*f}):e.shapeArgs=e.plotY=e.dlBox=void 0}};c.prototype.getPxExtremes=function(){var a=Math.min(this.chart.plotWidth,this.chart.plotHeight),b=function(b){if("string"===typeof b){var c=/%$/.test(b);b=parseInt(b,10)}return c?a*b/100:b},c=b(v(this.options.minSize,8));b=Math.max(b(v(this.options.maxSize,"20%")),c);return{minPxSize:c,maxPxSize:b}};c.prototype.getZExtremes=
function(){var a=this.options,b=(this.zData||[]).filter(z);if(b.length){var c=v(a.zMin,r(f(b),!1===a.displayNegative?a.zThreshold||0:-Number.MAX_VALUE,Number.MAX_VALUE));a=v(a.zMax,d(b));if(z(c)&&z(a))return{zMin:c,zMax:a}}};c.compose=p.compose;c.defaultOptions=u(a.defaultOptions,{dataLabels:{formatter:function(){var a=this.series.chart.numberFormatter,b=this.point.z;return z(b)?a(b,-1):""},inside:!0,verticalAlign:"middle"},animationLimit:250,marker:{lineColor:null,lineWidth:1,fillOpacity:.5,radius:null,
states:{hover:{radiusPlus:0}},symbol:"circle"},minSize:8,maxSize:"20%",softThreshold:!1,states:{hover:{halo:{size:5}}},tooltip:{pointFormat:"({point.x}, {point.y}), Size: {point.z}"},turboThreshold:0,zThreshold:0,zoneAxis:"z"});return c}(a);y(e.prototype,{alignDataLabel:n.prototype.alignDataLabel,applyZones:l,bubblePadding:!0,buildKDTree:l,directTouch:!0,isBubble:!0,pointArrayMap:["y","z"],pointClass:q,parallelArrays:["x","y","z"],trackerGroups:["group","dataLabelsGroup"],specialGroup:"group",zoneAxis:"z"});
k(e,"updatedData",function(a){delete a.target.chart.bubbleZExtremes});c.prototype.beforePadding=function(){var a=this,b=this.len,c=this.chart,d=0,e=b,f=this.isXAxis,g=f?"xData":"yData",k=this.min,h=this.max-k,l=b/h,r;this.series.forEach(function(b){if(b.bubblePadding&&(b.visible||!c.options.chart.ignoreHiddenSeries)){r=a.allowZoomOutside=!0;var m=b[g];f&&b.getRadii(0,0,b);if(0<h)for(var n=m.length;n--;)if(z(m[n])&&a.dataMin<=m[n]&&m[n]<=a.max){var p=b.radii&&b.radii[n]||0;d=Math.min((m[n]-k)*l-p,
d);e=Math.max((m[n]-k)*l+p,e)}}});r&&0<h&&!this.logarithmic&&(e-=b,l*=(b+Math.max(0,d)-Math.min(e,b))/b,[["min","userMin",d],["max","userMax",e]].forEach(function(b){"undefined"===typeof v(a.options[b[0]],a[b[1]])&&(a[b[0]]+=b[2]/l)}))};b.registerSeriesType("bubble",e);"";"";return e});x(c,"Series/MapBubble/MapBubblePoint.js",[c["Core/Series/SeriesRegistry.js"],c["Core/Utilities.js"]],function(c,p){var q=this&&this.__extends||function(){var c=function(b,e){c=Object.setPrototypeOf||{__proto__:[]}instanceof
Array&&function(b,c){b.__proto__=c}||function(b,c){for(var e in c)c.hasOwnProperty(e)&&(b[e]=c[e])};return c(b,e)};return function(b,e){function g(){this.constructor=b}c(b,e);b.prototype=null===e?Object.create(e):(g.prototype=e.prototype,new g)}}();c=c.seriesTypes;var l=c.map,n=p.merge;return function(c){function b(){return null!==c&&c.apply(this,arguments)||this}q(b,c);b.prototype.applyOptions=function(b,g){return b&&"undefined"!==typeof b.lat&&"undefined"!==typeof b.lon?c.prototype.applyOptions.call(this,
n(b,this.series.chart.fromLatLonToPoint(b)),g):l.prototype.pointClass.prototype.applyOptions.call(this,b,g)};b.prototype.isValid=function(){return"number"===typeof this.z};return b}(c.bubble.prototype.pointClass)});x(c,"Series/MapBubble/MapBubbleSeries.js",[c["Series/Bubble/BubbleSeries.js"],c["Series/MapBubble/MapBubblePoint.js"],c["Series/Map/MapSeries.js"],c["Core/Series/SeriesRegistry.js"],c["Core/Utilities.js"]],function(c,p,q,l,n){var h=this&&this.__extends||function(){var b=function(c,a){b=
Object.setPrototypeOf||{__proto__:[]}instanceof Array&&function(a,b){a.__proto__=b}||function(a,b){for(var c in b)b.hasOwnProperty(c)&&(a[c]=b[c])};return b(c,a)};return function(c,a){function d(){this.constructor=c}b(c,a);c.prototype=null===a?Object.create(a):(d.prototype=a.prototype,new d)}}(),b=l.seriesTypes.mappoint,e=n.extend,g=n.merge;n=function(e){function k(){var a=null!==e&&e.apply(this,arguments)||this;a.data=void 0;a.options=void 0;a.points=void 0;return a}h(k,e);k.prototype.translate=
function(){b.prototype.translate.call(this);this.getRadii();this.translateBubble()};k.compose=c.compose;k.defaultOptions=g(c.defaultOptions,{animationLimit:500,tooltip:{pointFormat:"{point.name}: {point.z}"}});return k}(c);e(n.prototype,{type:"mapbubble",axisTypes:["colorAxis"],getProjectedBounds:q.prototype.getProjectedBounds,isCartesian:!1,pointArrayMap:["z"],pointClass:p,setData:q.prototype.setData,setOptions:q.prototype.setOptions,useMapGeometry:!0,xyFromShape:!0});l.registerSeriesType("mapbubble",
n);"";return n});x(c,"Series/Heatmap/HeatmapPoint.js",[c["Core/Series/SeriesRegistry.js"],c["Core/Utilities.js"]],function(c,p){var q=this&&this.__extends||function(){var b=function(c,g){b=Object.setPrototypeOf||{__proto__:[]}instanceof Array&&function(b,c){b.__proto__=c}||function(b,c){for(var a in c)c.hasOwnProperty(a)&&(b[a]=c[a])};return b(c,g)};return function(c,g){function e(){this.constructor=c}b(c,g);c.prototype=null===g?Object.create(g):(e.prototype=g.prototype,new e)}}(),l=p.clamp,n=p.extend,
h=p.pick;c=function(b){function c(){var c=null!==b&&b.apply(this,arguments)||this;c.options=void 0;c.series=void 0;c.value=void 0;c.x=void 0;c.y=void 0;return c}q(c,b);c.prototype.applyOptions=function(c,e){c=b.prototype.applyOptions.call(this,c,e);c.formatPrefix=c.isNull||null===c.value?"null":"point";return c};c.prototype.getCellAttributes=function(){var b=this.series,c=b.options,e=(c.colsize||1)/2,a=(c.rowsize||1)/2,d=b.xAxis,f=b.yAxis,n=this.options.marker||b.options.marker;b=b.pointPlacementToXValue();
var p=h(this.pointPadding,c.pointPadding,0),q={x1:l(Math.round(d.len-(d.translate(this.x-e,!1,!0,!1,!0,-b)||0)),-d.len,2*d.len),x2:l(Math.round(d.len-(d.translate(this.x+e,!1,!0,!1,!0,-b)||0)),-d.len,2*d.len),y1:l(Math.round(f.translate(this.y-a,!1,!0,!1,!0)||0),-f.len,2*f.len),y2:l(Math.round(f.translate(this.y+a,!1,!0,!1,!0)||0),-f.len,2*f.len)};[["width","x"],["height","y"]].forEach(function(a){var b=a[0];a=a[1];var c=a+"1",d=a+"2",e=Math.abs(q[c]-q[d]),f=n&&n.lineWidth||0,g=Math.abs(q[c]+q[d])/
2;n[b]&&n[b]<e&&(q[c]=g-n[b]/2-f/2,q[d]=g+n[b]/2+f/2);p&&("y"===a&&(c=d,d=a+"1"),q[c]+=p,q[d]-=p)});return q};c.prototype.haloPath=function(b){if(!b)return[];var c=this.shapeArgs;return["M",c.x-b,c.y-b,"L",c.x-b,c.y+c.height+b,c.x+c.width+b,c.y+c.height+b,c.x+c.width+b,c.y-b,"Z"]};c.prototype.isValid=function(){return Infinity!==this.value&&-Infinity!==this.value};return c}(c.seriesTypes.scatter.prototype.pointClass);n(c.prototype,{dataLabelOnNull:!0,moveToTopOnHover:!0,ttBelow:!1});return c});x(c,
"Series/Heatmap/HeatmapSeries.js",[c["Core/Color/Color.js"],c["Series/ColorMapComposition.js"],c["Series/Heatmap/HeatmapPoint.js"],c["Core/Legend/LegendSymbol.js"],c["Core/Series/SeriesRegistry.js"],c["Core/Renderer/SVG/SVGRenderer.js"],c["Core/Utilities.js"]],function(c,p,q,l,n,h,b){var e=this&&this.__extends||function(){var a=function(b,c){a=Object.setPrototypeOf||{__proto__:[]}instanceof Array&&function(a,b){a.__proto__=b}||function(a,b){for(var c in b)b.hasOwnProperty(c)&&(a[c]=b[c])};return a(b,
c)};return function(b,c){function d(){this.constructor=b}a(b,c);b.prototype=null===c?Object.create(c):(d.prototype=c.prototype,new d)}}(),g=p.colorMapSeriesMixin,t=n.series,k=n.seriesTypes,a=k.column,d=k.scatter,f=h.prototype.symbols,r=b.extend,y=b.fireEvent,x=b.isNumber,u=b.merge,v=b.pick;h=function(a){function b(){var b=null!==a&&a.apply(this,arguments)||this;b.colorAxis=void 0;b.data=void 0;b.options=void 0;b.points=void 0;b.valueMax=NaN;b.valueMin=NaN;return b}e(b,a);b.prototype.drawPoints=function(){var a=
this;if((this.options.marker||{}).enabled||this._hasPointMarkers)t.prototype.drawPoints.call(this),this.points.forEach(function(b){b.graphic&&(b.graphic[a.chart.styledMode?"css":"animate"](a.colorAttribs(b)),a.options.borderRadius&&b.graphic.attr({r:a.options.borderRadius}),(b.shapeArgs||{}).r=a.options.borderRadius,(b.shapeArgs||{}).d=b.graphic.pathArray,null===b.value&&b.graphic.addClass("highcharts-null-point"))})};b.prototype.getExtremes=function(){var a=t.prototype.getExtremes.call(this,this.valueData),
b=a.dataMin;a=a.dataMax;x(b)&&(this.valueMin=b);x(a)&&(this.valueMax=a);return t.prototype.getExtremes.call(this)};b.prototype.getValidPoints=function(a,b){return t.prototype.getValidPoints.call(this,a,b,!0)};b.prototype.hasData=function(){return!!this.processedXData.length};b.prototype.init=function(){t.prototype.init.apply(this,arguments);var a=this.options;a.pointRange=v(a.pointRange,a.colsize||1);this.yAxis.axisPointRange=a.rowsize||1;f.ellipse=f.circle};b.prototype.markerAttribs=function(a,b){var c=
a.marker||{},d=this.options.marker||{},e=a.shapeArgs||{},f={};if(a.hasImage)return{x:a.plotX,y:a.plotY};if(b){var g=d.states[b]||{};var k=c.states&&c.states[b]||{};[["width","x"],["height","y"]].forEach(function(a){f[a[0]]=(k[a[0]]||g[a[0]]||e[a[0]])+(k[a[0]+"Plus"]||g[a[0]+"Plus"]||0);f[a[1]]=e[a[1]]+(e[a[0]]-f[a[0]])/2})}return b?f:e};b.prototype.pointAttribs=function(a,b){var d=t.prototype.pointAttribs.call(this,a,b),e=this.options||{},f=this.chart.options.plotOptions||{},g=f.series||{},k=f.heatmap||
{};f=a&&a.options.borderColor||e.borderColor||k.borderColor||g.borderColor;g=a&&a.options.borderWidth||e.borderWidth||k.borderWidth||g.borderWidth||d["stroke-width"];d.stroke=a&&a.marker&&a.marker.lineColor||e.marker&&e.marker.lineColor||f||this.color;d["stroke-width"]=g;b&&(a=u(e.states[b],e.marker&&e.marker.states[b],a&&a.options.states&&a.options.states[b]||{}),b=a.brightness,d.fill=a.color||c.parse(d.fill).brighten(b||0).get(),d.stroke=a.lineColor);return d};b.prototype.setClip=function(a){var b=
this.chart;t.prototype.setClip.apply(this,arguments);(!1!==this.options.clip||a)&&this.markerGroup.clip((a||this.clipBox)&&this.sharedClipKey?b.sharedClips[this.sharedClipKey]:b.clipRect)};b.prototype.translate=function(){var a=this.options,b=a.marker&&a.marker.symbol||"rect",c=f[b]?b:"rect",d=-1!==["circle","square"].indexOf(c);this.generatePoints();this.points.forEach(function(a){var e=a.getCellAttributes(),g={};g.x=Math.min(e.x1,e.x2);g.y=Math.min(e.y1,e.y2);g.width=Math.max(Math.abs(e.x2-e.x1),
0);g.height=Math.max(Math.abs(e.y2-e.y1),0);var k=a.hasImage=0===(a.marker&&a.marker.symbol||b||"").indexOf("url");if(d){var h=Math.abs(g.width-g.height);g.x=Math.min(e.x1,e.x2)+(g.width<g.height?0:h/2);g.y=Math.min(e.y1,e.y2)+(g.width<g.height?h/2:0);g.width=g.height=Math.min(g.width,g.height)}h={plotX:(e.x1+e.x2)/2,plotY:(e.y1+e.y2)/2,clientX:(e.x1+e.x2)/2,shapeType:"path",shapeArgs:u(!0,g,{d:f[c](g.x,g.y,g.width,g.height)})};k&&(a.marker={width:g.width,height:g.height});r(a,h)});y(this,"afterTranslate")};
b.defaultOptions=u(d.defaultOptions,{animation:!1,borderRadius:0,borderWidth:0,nullColor:"#f7f7f7",dataLabels:{formatter:function(){var a=this.series.chart.numberFormatter,b=this.point.value;return x(b)?a(b,-1):""},inside:!0,verticalAlign:"middle",crop:!1,overflow:!1,padding:0},marker:{symbol:"rect",radius:0,lineColor:void 0,states:{hover:{lineWidthPlus:0},select:{}}},clip:!0,pointRange:null,tooltip:{pointFormat:"{point.x}, {point.y}: {point.value}<br/>"},states:{hover:{halo:!1,brightness:.2}}});
return b}(d);r(h.prototype,{alignDataLabel:a.prototype.alignDataLabel,axisTypes:g.axisTypes,colorKey:"value",directTouch:!0,drawLegendSymbol:l.drawRectangle,getExtremesFromAll:!0,getSymbol:t.prototype.getSymbol,parallelArrays:g.parallelArrays,pointArrayMap:["y","value"],pointClass:q,trackerGroups:g.trackerGroups});p.compose(h);n.registerSeriesType("heatmap",h);"";"";return h});x(c,"Extensions/GeoJSON.js",[c["Core/Chart/Chart.js"],c["Core/FormatUtilities.js"],c["Core/Globals.js"],c["Core/Utilities.js"]],
function(c,p,q,l){function n(b,a){var c,e=!1,g=b.x,k=b.y;b=0;for(c=a.length-1;b<a.length;c=b++){var h=a[b][1]>k;var l=a[c][1]>k;h!==l&&g<(a[c][0]-a[b][0])*(k-a[b][1])/(a[c][1]-a[b][1])+a[b][0]&&(e=!e)}return e}var h=p.format,b=q.win,e=l.error,g=l.extend,t=l.merge;p=l.wrap;"";c.prototype.transformFromLatLon=function(c,a){var d=this.options.chart.proj4||b.proj4;if(!d)return e(21,!1,this),{x:0,y:null};var f=a.jsonmarginX;f=void 0===f?0:f;var g=a.jsonmarginY;g=void 0===g?0:g;var k=a.jsonres;k=void 0===
k?1:k;var h=a.scale;h=void 0===h?1:h;var l=a.xoffset;l=void 0===l?0:l;var n=a.xpan;n=void 0===n?0:n;var p=a.yoffset;p=void 0===p?0:p;var q=a.ypan;q=void 0===q?0:q;c=d(a.crs,[c.lon,c.lat]);d=a.cosAngle||a.rotation&&Math.cos(a.rotation);var m=a.sinAngle||a.rotation&&Math.sin(a.rotation);a=a.rotation?[c[0]*d+c[1]*m,-c[0]*m+c[1]*d]:c;return{x:((a[0]-l)*h+n)*k+f,y:-(((p-a[1])*h+q)*k-g)}};c.prototype.transformToLatLon=function(c,a){if(!this.options.chart.proj4&&!b.proj4)e(21,!1,this);else if(null!==c.y){var d=
a.jsonmarginX,f=a.jsonmarginY,g=a.jsonres;g=void 0===g?1:g;var h=a.scale;h=void 0===h?1:h;var k=a.xoffset,l=a.xpan,n=a.yoffset,p=a.ypan;c={x:((c.x-(void 0===d?0:d))/g-(void 0===l?0:l))/h+(void 0===k?0:k),y:((c.y-(void 0===f?0:f))/g+(void 0===p?0:p))/h+(void 0===n?0:n)};d=a.cosAngle||a.rotation&&Math.cos(a.rotation);f=a.sinAngle||a.rotation&&Math.sin(a.rotation);a=b.proj4(a.crs,"WGS84",a.rotation?{x:c.x*d+c.y*-f,y:c.x*f+c.y*d}:c);return{lat:a.y,lon:a.x}}};c.prototype.fromPointToLatLon=function(b){var a=
this.mapTransforms;if(a){for(var c in a)if(Object.hasOwnProperty.call(a,c)&&a[c].hitZone&&n(b,a[c].hitZone.coordinates[0]))return this.transformToLatLon(b,a[c]);return this.transformToLatLon(b,a["default"])}e(22,!1,this)};c.prototype.fromLatLonToPoint=function(b){var a=this.mapTransforms,c;if(!a)return e(22,!1,this),{x:0,y:null};for(c in a)if(Object.hasOwnProperty.call(a,c)&&a[c].hitZone){var f=this.transformFromLatLon(b,a[c]);if(n(f,a[c].hitZone.coordinates[0]))return f}return this.transformFromLatLon(b,
a["default"])};q.geojson=function(b,a,c){void 0===a&&(a="map");var d=[];b.features.forEach(function(b){var c=b.geometry||{},e=c.type;c=c.coordinates;b=b.properties;var f;"map"!==a&&"mapbubble"!==a||"Polygon"!==e&&"MultiPolygon"!==e?"mapline"!==a||"LineString"!==e&&"MultiLineString"!==e?"mappoint"===a&&"Point"===e&&c.length&&(f={geometry:{coordinates:c,type:e}}):c.length&&(f={geometry:{coordinates:c,type:e}}):c.length&&(f={geometry:{coordinates:c,type:e}});f&&d.push(g(f,{name:b.name||b.NAME,properties:b}))});
c&&b.copyrightShort&&(c.chart.mapCredits=h(c.chart.options.credits.mapText,{geojson:b}),c.chart.mapCreditsFull=h(c.chart.options.credits.mapTextFull,{geojson:b}));return d};p(c.prototype,"addCredits",function(b,a){a=t(!0,this.options.credits,a);this.mapCredits&&(a.href=null);b.call(this,a);this.credits&&this.mapCreditsFull&&this.credits.attr({title:this.mapCreditsFull})})});x(c,"masters/modules/map.src.js",[c["Core/Globals.js"],c["Core/Axis/Color/ColorAxis.js"],c["Series/MapBubble/MapBubbleSeries.js"],
c["Core/Chart/MapChart.js"]],function(c,p,q,l){c.ColorAxis=p;c.MapChart=l;c.mapChart=c.Map=l.mapChart;c.maps=l.maps;p.compose(c.Chart,c.Fx,c.Legend,c.Series);q.compose(c.Chart,c.Legend,c.Series)})});
//# sourceMappingURL=map.js.map
```
