## Supplementary material for "Exploring natural odour landscapes: A case study with implications for human-biting insects": Figure S4: marker-clusters.js

```
/*
 Highcharts JS v9.3.1 (2021-11-05)

 Marker clusters module for Highcharts

 (c) 2010-2021 Wojciech Chmiel

 License: www.highcharts.com/license
*/
'use strict';(function(m){"object"===typeof module&&module.exports?(m["default"]=m,module.exports=m):"function"===typeof define&&define.amd?define("highcharts/modules/marker-clusters",["highcharts"],function(v){m(v);m.Highcharts=v;return m}):m("undefined"!==typeof Highcharts?Highcharts:void 0)})(function(m){function v(m,v,M,F){m.hasOwnProperty(v)||(m[v]=F.apply(null,M))}m=m?m._modules:{};v(m,"Extensions/MarkerClusters.js",[m["Core/Animation/AnimationUtilities.js"],m["Core/Chart/Chart.js"],m["Core/DefaultOptions.js"],
m["Core/Series/Point.js"],m["Core/Series/Series.js"],m["Core/Series/SeriesRegistry.js"],m["Core/Renderer/SVG/SVGRenderer.js"],m["Core/Utilities.js"],m["Core/Axis/Axis.js"]],function(m,v,M,F,N,O,t,u,da){function J(a){var b=a.length,c=0,e=0,d;for(d=0;d<b;d++)c+=a[d].x,e+=a[d].y;return{x:c/b,y:e/b}}function T(a,b){var c=[];c.length=b;a.clusters.forEach(function(a){a.data.forEach(function(a){c[a.dataIndex]=a})});a.noise.forEach(function(a){c[a.data[0].dataIndex]=a.data[0]});return c}function U(a,b,c,
e,d){a.point&&(e&&a.point.graphic&&(a.point.graphic.show(),a.point.graphic.attr({opacity:b}).animate({opacity:1},c)),d&&a.point.dataLabel&&(a.point.dataLabel.show(),a.point.dataLabel.attr({opacity:b}).animate({opacity:1},c)))}function V(a,b,c){a.point&&(b&&a.point.graphic&&a.point.graphic.hide(),c&&a.point.dataLabel&&a.point.dataLabel.hide())}function ea(a){a&&W(a,function(a){a.point&&a.point.destroy&&a.point.destroy()})}function P(a,b,c,e){U(a,e,c,!0,!0);b.forEach(function(a){a.point&&a.point.destroy&&
a.point.destroy()})}var X=m.animObject;m=M.defaultOptions;O=O.seriesTypes;var G=t.prototype.symbols,z=u.addEvent,A=u.defined,Y=u.error,Z=u.isArray,Q=u.isFunction,R=u.isObject,D=u.isNumber,S=u.merge,W=u.objectEach,aa=u.relativeLength,K=u.syncTimeout;"";t=O.scatter;var ba=N.prototype.generatePoints,L=[],ca=0,B={enabled:!1,allowOverlap:!0,animation:{duration:500},drillToCluster:!0,minimumClusterSize:2,layoutAlgorithm:{gridSize:50,distance:40,kmeansThreshold:100},marker:{symbol:"cluster",radius:15,lineWidth:0,
lineColor:"#ffffff"},dataLabels:{enabled:!0,format:"{point.clusterPointsAmount}",verticalAlign:"middle",align:"center",style:{color:"contrast"},inside:!0}};(m.plotOptions||{}).series=S((m.plotOptions||{}).series,{cluster:B,tooltip:{clusterFormat:"<span>Clustered points: {point.clusterPointsAmount}</span><br/>"}});var I=function(a,b){var c=a.chart,e=a.xAxis;a=a.yAxis;return c.mapView?c.mapView.pixelsToProjectedUnits(b):{x:e?e.toValue(b.x):0,y:a?a.toValue(b.y):0}},H=function(a,b){var c=a.chart,e=a.xAxis;
a=a.yAxis;return c.mapView?c.mapView.projectedUnitsToPixels(b):{x:e?e.toPixels(b.x):0,y:a?a.toPixels(b.y):0}};G.cluster=function(a,b,c,e){c/=2;e/=2;var d=G.arc(a+c,b+e,c-4,e-4,{start:.5*Math.PI,end:2.5*Math.PI,open:!1}),k=G.arc(a+c,b+e,c-3,e-3,{start:.5*Math.PI,end:2.5*Math.PI,innerR:c-2,open:!1});return G.arc(a+c,b+e,c-1,e-1,{start:.5*Math.PI,end:2.5*Math.PI,innerR:c,open:!1}).concat(k,d)};t.prototype.animateClusterPoint=function(a){var b=this.chart,c=b.mapView,e=X((this.options.cluster||{}).animation),
d=e.duration||500,k=(this.markerClusterInfo||{}).pointsState,r=(k||{}).newState,n=(k||{}).oldState,p=[],l=k=0,q=0,h=!1,w=!1;if(n&&r){var g=r[a.stateId];k=H(this,g);l=k.x-(c?0:b.plotLeft);q=k.y-(c?0:b.plotTop);if(1===g.parentsId.length){a=(r||{})[a.stateId].parentsId[0];var f=n[a];g.point&&g.point.graphic&&f&&f.point&&f.point.plotX&&f.point.plotY&&f.point.plotX!==g.point.plotX&&f.point.plotY!==g.point.plotY&&(a=g.point.graphic.getBBox(),k=g.point.graphic&&g.point.graphic.isImg?0:a.width/2,g.point.graphic.attr({x:f.point.plotX-
k,y:f.point.plotY-k}),g.point.graphic.animate({x:l-(g.point.graphic.radius||0),y:q-(g.point.graphic.radius||0)},e,function(){w=!0;f.point&&f.point.destroy&&f.point.destroy()}),g.point.dataLabel&&g.point.dataLabel.alignAttr&&f.point.dataLabel&&f.point.dataLabel.alignAttr&&(g.point.dataLabel.attr({x:f.point.dataLabel.alignAttr.x,y:f.point.dataLabel.alignAttr.y}),g.point.dataLabel.animate({x:g.point.dataLabel.alignAttr.x,y:g.point.dataLabel.alignAttr.y},e)))}else 0===g.parentsId.length?(V(g,!0,!0),K(function(){U(g,
.1,e,!0,!0)},d/2)):(V(g,!0,!0),g.parentsId.forEach(function(a){n&&n[a]&&(f=n[a],p.push(f),f.point&&f.point.graphic&&(h=!0,f.point.graphic.show(),f.point.graphic.animate({x:l-(f.point.graphic.radius||0),y:q-(f.point.graphic.radius||0),opacity:.4},e,function(){w=!0;P(g,p,e,.7)}),f.point.dataLabel&&-9999!==f.point.dataLabel.y&&g.point&&g.point.dataLabel&&g.point.dataLabel.alignAttr&&(f.point.dataLabel.show(),f.point.dataLabel.animate({x:g.point.dataLabel.alignAttr.x,y:g.point.dataLabel.alignAttr.y,opacity:.4},
e))))}),K(function(){w||P(g,p,e,.85)},d),h||K(function(){P(g,p,e,.1)},d/2))}};t.prototype.getGridOffset=function(){var a=this.chart,b=this.xAxis,c=this.yAxis;b=b&&this.dataMinX&&this.dataMaxX?b.reversed?b.toPixels(this.dataMaxX):b.toPixels(this.dataMinX):a.plotLeft;a=c&&this.dataMinY&&this.dataMaxY?c.reversed?c.toPixels(this.dataMinY):c.toPixels(this.dataMaxY):a.plotTop;return{plotLeft:b,plotTop:a}};t.prototype.getScaledGridSize=function(a){var b=this.xAxis,c=this.chart.mapView;a=a.processedGridSize||
B.layoutAlgorithm.gridSize;var e=!0,d=1,k=1;this.gridValueSize||(this.gridValueSize=c?a/c.getScale():Math.abs(b.toValue(a)-b.toValue(0)));b=c?this.gridValueSize*c.getScale():b.toPixels(this.gridValueSize)-b.toPixels(0);for(b=+(a/b).toFixed(14);e&&1!==b;)c=Math.pow(2,d),.75<b&&1.25>b?e=!1:b>=1/c&&b<1/c*2?(e=!1,k=c):b<=c&&b>c/2&&(e=!1,k=1/c),d++;return a/k/b};t.prototype.getRealExtremes=function(){var a=this.chart,b=a.mapView?0:a.plotLeft,c=I(this,{x:b,y:a.mapView?0:a.plotTop}),e=I(this,{x:b+a.plotWidth,
y:b+a.plotHeight});a=c.x;b=e.x;c=c.y;e=e.y;return{minX:Math.min(a,b),maxX:Math.max(a,b),minY:Math.min(c,e),maxY:Math.max(c,e)}};t.prototype.onDrillToCluster=function(a){(a.point||a.target).firePointEvent("drillToCluster",a,function(a){var b=a.point||a.target,e=b.series.xAxis,d=b.series.yAxis,k=b.series.chart,r=k.mapView;if((b.series.options.cluster||{}).drillToCluster&&b.clusteredData){var n=b.clusteredData.map(function(a){return a.x}).sort(function(a,b){return a-b});b=b.clusteredData.map(function(a){return a.y}).sort(function(a,
b){return a-b});var p=n[0],l=n[n.length-1];n=b[0];var q=b[b.length-1],h=Math.abs(.1*(l-p)),w=Math.abs(.1*(q-n));b=Math.min(p,l)-h;p=Math.max(p,l)+h;l=Math.min(n,q)-w;n=Math.max(n,q)+w;r?r.fitToBounds({x1:b,x2:p,y1:l,y2:n}):e&&d&&(k.pointer.zoomX=!0,k.pointer.zoomY=!0,k.zoom({originalEvent:a,xAxis:[{axis:e,min:b,max:p}],yAxis:[{axis:d,min:l,max:n}]}))}})};t.prototype.getClusterDistancesFromPoint=function(a,b,c){for(var e=[],d=0;d<a.length;d++){var k=H(this,{x:b,y:c}),r=H(this,{x:a[d].posX,y:a[d].posY});
e.push({clusterIndex:d,distance:Math.sqrt(Math.pow(k.x-r.x,2)+Math.pow(k.y-r.y,2))})}return e.sort(function(a,b){return a.distance-b.distance})};t.prototype.getPointsState=function(a,b,c){b=b?T(b,c):[];c=T(a,c);var e={},d;L=[];a.clusters.forEach(function(a){e[a.stateId]={x:a.x,y:a.y,id:a.stateId,point:a.point,parentsId:[]}});a.noise.forEach(function(a){e[a.stateId]={x:a.x,y:a.y,id:a.stateId,point:a.point,parentsId:[]}});for(d=0;d<c.length;d++){a=c[d];var k=b[d];a&&k&&a.parentStateId&&k.parentStateId&&
e[a.parentStateId]&&-1===e[a.parentStateId].parentsId.indexOf(k.parentStateId)&&(e[a.parentStateId].parentsId.push(k.parentStateId),-1===L.indexOf(k.parentStateId)&&L.push(k.parentStateId))}return e};t.prototype.markerClusterAlgorithms={grid:function(a,b,c,e){var d={},k=this.getGridOffset(),r=this.getScaledGridSize(e);for(e=0;e<a.length;e++){var n=H(this,{x:a[e],y:b[e]});var p=n.x-k.plotLeft;n=n.y-k.plotTop;p=Math.floor(p/r);n=Math.floor(n/r);p=n+"-"+p;d[p]||(d[p]=[]);d[p].push({dataIndex:c[e],x:a[e],
y:b[e]})}return d},kmeans:function(a,b,c,e){var d=[],k=[],r={},n=e.processedDistance||B.layoutAlgorithm.distance,p=e.iterations,l=0,q=!0,h=0,w=0;var g=[];var f;e.processedGridSize=e.processedDistance;h=this.markerClusterAlgorithms?this.markerClusterAlgorithms.grid.call(this,a,b,c,e):{};for(f in h)1<h[f].length&&(g=J(h[f]),d.push({posX:g.x,posY:g.y,oldX:0,oldY:0,startPointsLen:h[f].length,points:[]}));for(;q;){d.map(function(a){a.points.length=0;return a});for(q=k.length=0;q<a.length;q++)h=a[q],w=
b[q],g=this.getClusterDistancesFromPoint(d,h,w),g.length&&g[0].distance<n?d[g[0].clusterIndex].points.push({x:h,y:w,dataIndex:c[q]}):k.push({x:h,y:w,dataIndex:c[q]});for(f=0;f<d.length;f++)1===d[f].points.length&&(g=this.getClusterDistancesFromPoint(d,d[f].points[0].x,d[f].points[0].y),g[1].distance<n&&(d[g[1].clusterIndex].points.push(d[f].points[0]),d[g[0].clusterIndex].points.length=0));q=!1;for(f=0;f<d.length;f++)if(g=J(d[f].points),d[f].oldX=d[f].posX,d[f].oldY=d[f].posY,d[f].posX=g.x,d[f].posY=
g.y,d[f].posX>d[f].oldX+1||d[f].posX<d[f].oldX-1||d[f].posY>d[f].oldY+1||d[f].posY<d[f].oldY-1)q=!0;p&&(q=l<p-1);l++}d.forEach(function(a,b){r["cluster"+b]=a.points});k.forEach(function(a,b){r["noise"+b]=[a]});return r},optimizedKmeans:function(a,b,c,e){var d=this,k=e.processedDistance||B.layoutAlgorithm.gridSize,r={},n=d.getRealExtremes(),p=(d.options.cluster||{}).marker,l,q,h;!d.markerClusterInfo||d.initMaxX&&d.initMaxX<n.maxX||d.initMinX&&d.initMinX>n.minX||d.initMaxY&&d.initMaxY<n.maxY||d.initMinY&&
d.initMinY>n.minY?(d.initMaxX=n.maxX,d.initMinX=n.minX,d.initMaxY=n.maxY,d.initMinY=n.minY,r=d.markerClusterAlgorithms?d.markerClusterAlgorithms.kmeans.call(d,a,b,c,e):{},d.baseClusters=null):(d.baseClusters||(d.baseClusters={clusters:d.markerClusterInfo.clusters,noise:d.markerClusterInfo.noise}),d.baseClusters.clusters.forEach(function(a){a.pointsOutside=[];a.pointsInside=[];a.data.forEach(function(b){var c=H(d,b),e=H(d,a);q=Math.sqrt(Math.pow(c.x-e.x,2)+Math.pow(c.y-e.y,2));h=a.clusterZone&&a.clusterZone.marker&&
a.clusterZone.marker.radius?a.clusterZone.marker.radius:p&&p.radius?p.radius:B.marker.radius;l=0<=k-h?k-h:h;q>h+l&&A(a.pointsOutside)?a.pointsOutside.push(b):A(a.pointsInside)&&a.pointsInside.push(b)});a.pointsInside.length&&(r[a.id]=a.pointsInside);a.pointsOutside.forEach(function(b,c){r[a.id+"_noise"+c]=[b]})}),d.baseClusters.noise.forEach(function(a){r[a.id]=a.data}));return r}};t.prototype.preventClusterCollisions=function(a){var b=this,c=a.key.split("-").map(parseFloat),e=c[0],d=c[1],k=a.gridSize,
r=a.groupedData,n=a.defaultRadius,p=a.clusterRadius,l=d*k,q=e*k;c=H(b,a);var h=c.x,w=c.y;c=[];var g=0,f=0,m=(b.options.cluster||{}).marker,t=(b.options.cluster||{}).zones,x=b.getGridOffset(),u,v,y,z,D,F,G;h-=x.plotLeft;w-=x.plotTop;for(y=1;5>y;y++){var E=y%2?-1:1;var C=3>y?-1:1;E=Math.floor((h+E*p)/k);C=Math.floor((w+C*p)/k);E=[C+"-"+E,C+"-"+d,e+"-"+E];for(C=0;C<E.length;C++)-1===c.indexOf(E[C])&&E[C]!==a.key&&c.push(E[C])}c.forEach(function(a){if(r[a]){r[a].posX||(F=J(r[a]),r[a].posX=F.x,r[a].posY=
F.y);var c=H(b,{x:r[a].posX||0,y:r[a].posY||0});u=c.x-x.plotLeft;v=c.y-x.plotTop;c=a.split("-").map(parseFloat);D=c[0];z=c[1];if(t)for(g=r[a].length,y=0;y<t.length;y++)g>=t[y].from&&g<=t[y].to&&(f=A((t[y].marker||{}).radius)?t[y].marker.radius||0:m&&m.radius?m.radius:B.marker.radius);1<r[a].length&&0===f&&m&&m.radius?f=m.radius:1===r[a].length&&(f=n);G=p+f;f=0;z!==d&&Math.abs(h-u)<G&&(h=0>z-d?l+p:l+k-p);D!==e&&Math.abs(w-v)<G&&(w=0>D-e?q+p:q+k-p)}});c=I(b,{x:h+x.plotLeft,y:w+x.plotTop});r[a.key].posX=
c.x;r[a.key].posY=c.y;return c};t.prototype.isValidGroupedDataObject=function(a){var b=!1,c;if(!R(a))return!1;W(a,function(a){b=!0;if(Z(a)&&a.length)for(c=0;c<a.length;c++){if(!R(a[c])||!a[c].x||!a[c].y){b=!1;break}}else b=!1});return b};t.prototype.getClusteredData=function(a,b){var c=[],e=[],d=[],k=[],r=[],n=0,p=Math.max(2,b.minimumClusterSize||2),l,q;if(Q(b.layoutAlgorithm.type)&&!this.isValidGroupedDataObject(a))return Y("Highcharts marker-clusters module: The custom algorithm result is not valid!",
!1,this.chart),!1;for(q in a)if(a[q].length>=p){var h=a[q];var m=Math.random().toString(36).substring(2,7)+"-"+ca++;var g=h.length;if(b.zones)for(l=0;l<b.zones.length;l++)if(g>=b.zones[l].from&&g<=b.zones[l].to){var f=b.zones[l];f.zoneIndex=l;var t=b.zones[l].marker;var u=b.zones[l].className}var x=J(h);"grid"!==b.layoutAlgorithm.type||b.allowOverlap?x={x:x.x,y:x.y}:(l=this.options.marker||{},x=this.preventClusterCollisions({x:x.x,y:x.y,key:q,groupedData:a,gridSize:this.getScaledGridSize(b.layoutAlgorithm),
defaultRadius:l.radius||3+(l.lineWidth||0),clusterRadius:t&&t.radius?t.radius:(b.marker||{}).radius||B.marker.radius}));for(l=0;l<g;l++)h[l].parentStateId=m;d.push({x:x.x,y:x.y,id:q,stateId:m,index:n,data:h,clusterZone:f,clusterZoneClassName:u});c.push(x.x);e.push(x.y);r.push({options:{formatPrefix:"cluster",dataLabels:b.dataLabels,marker:S(b.marker,{states:b.states},t||{})}});if(this.options.data&&this.options.data.length)for(l=0;l<g;l++)R(this.options.data[h[l].dataIndex])&&(h[l].options=this.options.data[h[l].dataIndex]);
n++;t=null}else for(l=0;l<a[q].length;l++)h=a[q][l],m=Math.random().toString(36).substring(2,7)+"-"+ca++,g=((this.options||{}).data||[])[h.dataIndex],c.push(h.x),e.push(h.y),h.parentStateId=m,k.push({x:h.x,y:h.y,id:q,stateId:m,index:n,data:a[q]}),m=g&&"object"===typeof g&&!Z(g)?S(g,{x:h.x,y:h.y}):{userOptions:g,x:h.x,y:h.y},r.push({options:m}),n++;return{clusters:d,noise:k,groupedXData:c,groupedYData:e,groupMap:r}};t.prototype.destroyClusteredData=function(){(this.markerClusterSeriesData||[]).forEach(function(a){a&&
a.destroy&&a.destroy()});this.markerClusterSeriesData=null};t.prototype.hideClusteredData=function(){var a=this.markerClusterSeriesData,b=((this.markerClusterInfo||{}).pointsState||{}).oldState||{},c=L.map(function(a){return(b[a].point||{}).id||""});(a||[]).forEach(function(a){a&&-1!==c.indexOf(a.id)?(a.graphic&&a.graphic.hide(),a.dataLabel&&a.dataLabel.hide()):a&&a.destroy&&a.destroy()})};t.prototype.generatePoints=function(){var a=this,b=a.chart,c=a.options.cluster,e=a.getRealExtremes(),d=[],k=
[],m=[],n,p,l;if(c&&c.enabled&&a.xData&&a.xData.length&&a.yData&&a.yData.length&&!b.polar){var q=c.layoutAlgorithm.type;var h=c.layoutAlgorithm;h.processedGridSize=aa(h.gridSize||B.layoutAlgorithm.gridSize,b.plotWidth);h.processedDistance=aa(h.distance||B.layoutAlgorithm.distance,b.plotWidth);b=h.kmeansThreshold||B.layoutAlgorithm.kmeansThreshold;var t=h.processedGridSize/2;var g=I(a,{x:0,y:0});var f=I(a,{x:t,y:t});t=Math.abs(g.x-f.x);g=Math.abs(g.y-f.y);for(f=0;f<a.xData.length;f++){if(!a.dataMaxX)if(A(u)&&
A(n)&&A(v)&&A(p))D(a.yData[f])&&D(v)&&D(p)&&(u=Math.max(a.xData[f],u),n=Math.min(a.xData[f],n),v=Math.max(a.yData[f]||v,v),p=Math.min(a.yData[f]||p,p));else{var u=n=a.xData[f];var v=p=a.yData[f]}a.xData[f]>=e.minX-t&&a.xData[f]<=e.maxX+t&&(a.yData[f]||e.minY)>=e.minY-g&&(a.yData[f]||e.maxY)<=e.maxY+g&&(d.push(a.xData[f]),k.push(a.yData[f]),m.push(f))}A(u)&&A(n)&&D(v)&&D(p)&&(a.dataMaxX=u,a.dataMinX=n,a.dataMaxY=v,a.dataMinY=p);e=Q(q)?q:a.markerClusterAlgorithms?q&&a.markerClusterAlgorithms[q]?a.markerClusterAlgorithms[q]:
d.length<b?a.markerClusterAlgorithms.kmeans:a.markerClusterAlgorithms.grid:function(){return!1};h=(d=e.call(this,d,k,m,h))?a.getClusteredData(d,c):d;c.animation&&a.markerClusterInfo&&a.markerClusterInfo.pointsState&&a.markerClusterInfo.pointsState.oldState?(ea(a.markerClusterInfo.pointsState.oldState),d=a.markerClusterInfo.pointsState.newState):d={};k=a.xData.length;m=a.markerClusterInfo;h&&(a.processedXData=h.groupedXData,a.processedYData=h.groupedYData,a.hasGroupedData=!0,a.markerClusterInfo=h,
a.groupMap=h.groupMap);ba.apply(this);h&&a.markerClusterInfo&&((a.markerClusterInfo.clusters||[]).forEach(function(b){l=a.points[b.index];l.isCluster=!0;l.clusteredData=b.data;l.clusterPointsAmount=b.data.length;b.point=l;z(l,"click",a.onDrillToCluster)}),(a.markerClusterInfo.noise||[]).forEach(function(b){b.point=a.points[b.index]}),c.animation&&a.markerClusterInfo&&(a.markerClusterInfo.pointsState={oldState:d,newState:a.getPointsState(h,m,k)}),c.animation?this.hideClusteredData():this.destroyClusteredData(),
this.markerClusterSeriesData=this.hasGroupedData?this.points:null)}else ba.apply(this)};z(v,"render",function(){(this.series||[]).forEach(function(a){if(a.markerClusterInfo){var b=((a.markerClusterInfo||{}).pointsState||{}).oldState;(a.options.cluster||{}).animation&&a.markerClusterInfo&&0===a.chart.pointer.pinchDown.length&&"pan"!==((a.xAxis||{}).eventArgs||{}).trigger&&b&&Object.keys(b).length&&(a.markerClusterInfo.clusters.forEach(function(b){a.animateClusterPoint(b)}),a.markerClusterInfo.noise.forEach(function(b){a.animateClusterPoint(b)}))}})});
z(F,"update",function(){if(this.dataGroup)return Y("Highcharts marker-clusters module: Running `Point.update` when point belongs to clustered series is not supported.",!1,this.series.chart),!1});z(N,"destroy",t.prototype.destroyClusteredData);z(N,"afterRender",function(){var a=(this.options.cluster||{}).drillToCluster;this.markerClusterInfo&&this.markerClusterInfo.clusters&&this.markerClusterInfo.clusters.forEach(function(b){b.point&&b.point.graphic&&(b.point.graphic.addClass("highcharts-cluster-point"),
a&&b.point&&(b.point.graphic.css({cursor:"pointer"}),b.point.dataLabel&&b.point.dataLabel.css({cursor:"pointer"})),A(b.clusterZone)&&b.point.graphic.addClass(b.clusterZoneClassName||"highcharts-cluster-zone-"+b.clusterZone.zoneIndex))})});z(F,"drillToCluster",function(a){var b=(((a.point||a.target).series.options.cluster||{}).events||{}).drillToCluster;Q(b)&&b.call(this,a)});z(da,"setExtremes",function(){var a=this.chart,b=0,c;a.series.forEach(function(a){a.markerClusterInfo&&(c=X((a.options.cluster||
{}).animation),b=c.duration||0)});K(function(){a.tooltip&&a.tooltip.destroy()},b)})});v(m,"masters/modules/marker-clusters.src.js",[],function(){})});
//# sourceMappingURL=marker-clusters.js.map
```
