## Supplementary material for "Exploring natural odour landscapes: A case study with implications for human-biting insects": Figure S4: networkgraph.js

```
/*
 Highcharts JS v9.3.1 (2021-11-05)

 Force directed graph module

 (c) 2010-2021 Torstein Honsi

 License: www.highcharts.com/license
*/
'use strict';(function(f){"object"===typeof module&&module.exports?(f["default"]=f,module.exports=f):"function"===typeof define&&define.amd?define("highcharts/modules/networkgraph",["highcharts"],function(m){f(m);f.Highcharts=m;return f}):f("undefined"!==typeof Highcharts?Highcharts:void 0)})(function(f){function m(f,a,d,e){f.hasOwnProperty(a)||(f[a]=e.apply(null,d))}f=f?f._modules:{};m(f,"Series/NodesComposition.js",[f["Core/Series/Point.js"],f["Core/Series/Series.js"],f["Core/Utilities.js"]],function(f,
a,d){var e=d.defined,c=d.extend,h=d.find,p=d.pick,n;(function(d){function l(){this.data=[].concat(this.points||[],this.nodes);return a.prototype.destroy.apply(this,arguments)}function b(){this.nodes&&(this.nodes.forEach(function(b){b.destroy()}),this.nodes.length=0);a.prototype.setData.apply(this,arguments)}function t(b){var c=arguments,g=this.isNode?this.linksTo.concat(this.linksFrom):[this.fromNode,this.toNode];"select"!==b&&g.forEach(function(b){b&&b.series&&(f.prototype.setState.apply(b,c),b.isNode||
(b.fromNode.graphic&&f.prototype.setState.apply(b.fromNode,c),b.toNode&&b.toNode.graphic&&f.prototype.setState.apply(b.toNode,c)))});f.prototype.setState.apply(this,c)}var u=[];d.compose=function(c,a){-1===u.indexOf(c)&&(u.push(c),c=c.prototype,c.setNodeState=t,c.setState=t);-1===u.indexOf(a)&&(u.push(a),c=a.prototype,c.destroy=l,c.setData=b);return a};d.createNode=function(b){var a=this.pointClass,g=function(b,g){return h(b,function(b){return b.id===g})},k=g(this.nodes,b);k||(g=this.options.nodes&&
g(this.options.nodes,b),k=(new a).init(this,c({className:"highcharts-node",isNode:!0,id:b,y:1},g)),k.linksTo=[],k.linksFrom=[],k.formatPrefix="node",k.name=k.name||k.options.id||"",k.mass=p(k.options.mass,k.options.marker&&k.options.marker.radius,this.options.marker&&this.options.marker.radius,4),k.getSum=function(){var b=0,g=0;k.linksTo.forEach(function(g){b+=g.weight});k.linksFrom.forEach(function(b){g+=b.weight});return Math.max(b,g)},k.offset=function(b,g){for(var c=0,a=0;a<k[g].length;a++){if(k[g][a]===
b)return c;c+=k[g][a].weight}},k.hasShape=function(){var b=0;k.linksTo.forEach(function(g){g.outgoing&&b++});return!k.linksTo.length||b!==k.linksTo.length},this.nodes.push(k));return k};d.destroy=l;d.generatePoints=function(){var b=this.chart,c={};a.prototype.generatePoints.call(this);this.nodes||(this.nodes=[]);this.colorCounter=0;this.nodes.forEach(function(b){b.linksFrom.length=0;b.linksTo.length=0;b.level=b.options.level});this.points.forEach(function(g){e(g.from)&&(c[g.from]||(c[g.from]=this.createNode(g.from)),
c[g.from].linksFrom.push(g),g.fromNode=c[g.from],b.styledMode?g.colorIndex=p(g.options.colorIndex,c[g.from].colorIndex):g.color=g.options.color||c[g.from].color);e(g.to)&&(c[g.to]||(c[g.to]=this.createNode(g.to)),c[g.to].linksTo.push(g),g.toNode=c[g.to]);g.name=g.name||g.id},this);this.nodeLookup=c};d.setNodeState=t})(n||(n={}));return n});m(f,"Series/Networkgraph/Integrations.js",[f["Core/Globals.js"]],function(f){f.networkgraphIntegrations={verlet:{attractiveForceFunction:function(a,d){return(d-
a)/a},repulsiveForceFunction:function(a,d){return(d-a)/a*(d>a?1:0)},barycenter:function(){var a=this.options.gravitationalConstant,d=this.barycenter.xFactor,e=this.barycenter.yFactor;d=(d-(this.box.left+this.box.width)/2)*a;e=(e-(this.box.top+this.box.height)/2)*a;this.nodes.forEach(function(c){c.fixedPosition||(c.plotX-=d/c.mass/c.degree,c.plotY-=e/c.mass/c.degree)})},repulsive:function(a,d,e){d=d*this.diffTemperature/a.mass/a.degree;a.fixedPosition||(a.plotX+=e.x*d,a.plotY+=e.y*d)},attractive:function(a,
d,e){var c=a.getMass(),h=-e.x*d*this.diffTemperature;d=-e.y*d*this.diffTemperature;a.fromNode.fixedPosition||(a.fromNode.plotX-=h*c.fromNode/a.fromNode.degree,a.fromNode.plotY-=d*c.fromNode/a.fromNode.degree);a.toNode.fixedPosition||(a.toNode.plotX+=h*c.toNode/a.toNode.degree,a.toNode.plotY+=d*c.toNode/a.toNode.degree)},integrate:function(a,d){var e=-a.options.friction,c=a.options.maxSpeed,h=(d.plotX+d.dispX-d.prevX)*e;e*=d.plotY+d.dispY-d.prevY;var f=Math.abs,n=f(h)/(h||1);f=f(e)/(e||1);h=n*Math.min(c,
Math.abs(h));e=f*Math.min(c,Math.abs(e));d.prevX=d.plotX+d.dispX;d.prevY=d.plotY+d.dispY;d.plotX+=h;d.plotY+=e;d.temperature=a.vectorLength({x:h,y:e})},getK:function(a){return Math.pow(a.box.width*a.box.height/a.nodes.length,.5)}},euler:{attractiveForceFunction:function(a,d){return a*a/d},repulsiveForceFunction:function(a,d){return d*d/a},barycenter:function(){var a=this.options.gravitationalConstant,d=this.barycenter.xFactor,e=this.barycenter.yFactor;this.nodes.forEach(function(c){if(!c.fixedPosition){var h=
c.getDegree();h*=1+h/2;c.dispX+=(d-c.plotX)*a*h/c.degree;c.dispY+=(e-c.plotY)*a*h/c.degree}})},repulsive:function(a,d,e,c){a.dispX+=e.x/c*d/a.degree;a.dispY+=e.y/c*d/a.degree},attractive:function(a,d,e,c){var h=a.getMass(),f=e.x/c*d;d*=e.y/c;a.fromNode.fixedPosition||(a.fromNode.dispX-=f*h.fromNode/a.fromNode.degree,a.fromNode.dispY-=d*h.fromNode/a.fromNode.degree);a.toNode.fixedPosition||(a.toNode.dispX+=f*h.toNode/a.toNode.degree,a.toNode.dispY+=d*h.toNode/a.toNode.degree)},integrate:function(a,
d){d.dispX+=d.dispX*a.options.friction;d.dispY+=d.dispY*a.options.friction;var e=d.temperature=a.vectorLength({x:d.dispX,y:d.dispY});0!==e&&(d.plotX+=d.dispX/e*Math.min(Math.abs(d.dispX),a.temperature),d.plotY+=d.dispY/e*Math.min(Math.abs(d.dispY),a.temperature))},getK:function(a){return Math.pow(a.box.width*a.box.height/a.nodes.length,.3)}}}});m(f,"Series/Networkgraph/QuadTree.js",[f["Core/Globals.js"],f["Core/Utilities.js"]],function(f,a){a=a.extend;var d=f.QuadTreeNode=function(a){this.box=a;this.boxSize=
Math.min(a.width,a.height);this.nodes=[];this.body=this.isInternal=!1;this.isEmpty=!0};a(d.prototype,{insert:function(a,c){this.isInternal?this.nodes[this.getBoxPosition(a)].insert(a,c-1):(this.isEmpty=!1,this.body?c?(this.isInternal=!0,this.divideBox(),!0!==this.body&&(this.nodes[this.getBoxPosition(this.body)].insert(this.body,c-1),this.body=!0),this.nodes[this.getBoxPosition(a)].insert(a,c-1)):(c=new d({top:a.plotX,left:a.plotY,width:.1,height:.1}),c.body=a,c.isInternal=!1,this.nodes.push(c)):
(this.isInternal=!1,this.body=a))},updateMassAndCenter:function(){var a=0,c=0,d=0;this.isInternal?(this.nodes.forEach(function(e){e.isEmpty||(a+=e.mass,c+=e.plotX*e.mass,d+=e.plotY*e.mass)}),c/=a,d/=a):this.body&&(a=this.body.mass,c=this.body.plotX,d=this.body.plotY);this.mass=a;this.plotX=c;this.plotY=d},divideBox:function(){var a=this.box.width/2,c=this.box.height/2;this.nodes[0]=new d({left:this.box.left,top:this.box.top,width:a,height:c});this.nodes[1]=new d({left:this.box.left+a,top:this.box.top,
width:a,height:c});this.nodes[2]=new d({left:this.box.left+a,top:this.box.top+c,width:a,height:c});this.nodes[3]=new d({left:this.box.left,top:this.box.top+c,width:a,height:c})},getBoxPosition:function(a){var c=a.plotY<this.box.top+this.box.height/2;return a.plotX<this.box.left+this.box.width/2?c?0:3:c?1:2}});f=f.QuadTree=function(a,c,f,p){this.box={left:a,top:c,width:f,height:p};this.maxDepth=25;this.root=new d(this.box,"0");this.root.isInternal=!0;this.root.isRoot=!0;this.root.divideBox()};a(f.prototype,
{insertNodes:function(a){a.forEach(function(a){this.root.insert(a,this.maxDepth)},this)},visitNodeRecursive:function(a,c,d){var e;a||(a=this.root);a===this.root&&c&&(e=c(a));!1!==e&&(a.nodes.forEach(function(a){if(a.isInternal){c&&(e=c(a));if(!1===e)return;this.visitNodeRecursive(a,c,d)}else a.body&&c&&c(a.body);d&&d(a)},this),a===this.root&&d&&d(a))},calculateMassAndCenter:function(){this.visitNodeRecursive(null,null,function(a){a.updateMassAndCenter()})}})});m(f,"Series/Networkgraph/Layouts.js",
[f["Core/Chart/Chart.js"],f["Core/Animation/AnimationUtilities.js"],f["Core/Globals.js"],f["Core/Utilities.js"]],function(f,a,d,e){var c=a.setAnimation;a=e.addEvent;var h=e.clamp,p=e.defined,n=e.extend,r=e.isFunction,l=e.pick;d.layouts={"reingold-fruchterman":function(){}};n(d.layouts["reingold-fruchterman"].prototype,{init:function(b){this.options=b;this.nodes=[];this.links=[];this.series=[];this.box={x:0,y:0,width:0,height:0};this.setInitialRendering(!0);this.integration=d.networkgraphIntegrations[b.integration];
this.enableSimulation=b.enableSimulation;this.attractiveForce=l(b.attractiveForce,this.integration.attractiveForceFunction);this.repulsiveForce=l(b.repulsiveForce,this.integration.repulsiveForceFunction);this.approximation=b.approximation},updateSimulation:function(b){this.enableSimulation=l(b,this.options.enableSimulation)},start:function(){var b=this.series,a=this.options;this.currentStep=0;this.forces=b[0]&&b[0].forces||[];this.chart=b[0]&&b[0].chart;this.initialRendering&&(this.initPositions(),
b.forEach(function(b){b.finishedAnimating=!0;b.render()}));this.setK();this.resetSimulation(a);this.enableSimulation&&this.step()},step:function(){var b=this,a=this.series;b.currentStep++;"barnes-hut"===b.approximation&&(b.createQuadTree(),b.quadTree.calculateMassAndCenter());b.forces.forEach(function(a){b[a+"Forces"](b.temperature)});b.applyLimits(b.temperature);b.temperature=b.coolDown(b.startTemperature,b.diffTemperature,b.currentStep);b.prevSystemTemperature=b.systemTemperature;b.systemTemperature=
b.getSystemTemperature();b.enableSimulation&&(a.forEach(function(b){b.chart&&b.render()}),b.maxIterations--&&isFinite(b.temperature)&&!b.isStable()?(b.simulation&&d.win.cancelAnimationFrame(b.simulation),b.simulation=d.win.requestAnimationFrame(function(){b.step()})):b.simulation=!1)},stop:function(){this.simulation&&d.win.cancelAnimationFrame(this.simulation)},setArea:function(b,a,c,d){this.box={left:b,top:a,width:c,height:d}},setK:function(){this.k=this.options.linkLength||this.integration.getK(this)},
addElementsToCollection:function(b,a){b.forEach(function(b){-1===a.indexOf(b)&&a.push(b)})},removeElementFromCollection:function(b,a){b=a.indexOf(b);-1!==b&&a.splice(b,1)},clear:function(){this.nodes.length=0;this.links.length=0;this.series.length=0;this.resetSimulation()},resetSimulation:function(){this.forcedStop=!1;this.systemTemperature=0;this.setMaxIterations();this.setTemperature();this.setDiffTemperature()},restartSimulation:function(){this.simulation?this.resetSimulation():(this.setInitialRendering(!1),
this.enableSimulation?this.start():this.setMaxIterations(1),this.chart&&this.chart.redraw(),this.setInitialRendering(!0))},setMaxIterations:function(b){this.maxIterations=l(b,this.options.maxIterations)},setTemperature:function(){this.temperature=this.startTemperature=Math.sqrt(this.nodes.length)},setDiffTemperature:function(){this.diffTemperature=this.startTemperature/(this.options.maxIterations+1)},setInitialRendering:function(b){this.initialRendering=b},createQuadTree:function(){this.quadTree=
new d.QuadTree(this.box.left,this.box.top,this.box.width,this.box.height);this.quadTree.insertNodes(this.nodes)},initPositions:function(){var b=this.options.initialPositions;r(b)?(b.call(this),this.nodes.forEach(function(b){p(b.prevX)||(b.prevX=b.plotX);p(b.prevY)||(b.prevY=b.plotY);b.dispX=0;b.dispY=0})):"circle"===b?this.setCircularPositions():this.setRandomPositions()},setCircularPositions:function(){function b(a){a.linksFrom.forEach(function(a){k[a.toNode.id]||(k[a.toNode.id]=!0,g.push(a.toNode),
b(a.toNode))})}var a=this.box,c=this.nodes,d=2*Math.PI/(c.length+1),e=c.filter(function(b){return 0===b.linksTo.length}),g=[],k={},f=this.options.initialPositionRadius;e.forEach(function(a){g.push(a);b(a)});g.length?c.forEach(function(b){-1===g.indexOf(b)&&g.push(b)}):g=c;g.forEach(function(b,g){b.plotX=b.prevX=l(b.plotX,a.width/2+f*Math.cos(g*d));b.plotY=b.prevY=l(b.plotY,a.height/2+f*Math.sin(g*d));b.dispX=0;b.dispY=0})},setRandomPositions:function(){function b(b){b=b*b/Math.PI;return b-=Math.floor(b)}
var a=this.box,c=this.nodes,d=c.length+1;c.forEach(function(c,g){c.plotX=c.prevX=l(c.plotX,a.width*b(g));c.plotY=c.prevY=l(c.plotY,a.height*b(d+g));c.dispX=0;c.dispY=0})},force:function(b){this.integration[b].apply(this,Array.prototype.slice.call(arguments,1))},barycenterForces:function(){this.getBarycenter();this.force("barycenter")},getBarycenter:function(){var b=0,a=0,c=0;this.nodes.forEach(function(d){a+=d.plotX*d.mass;c+=d.plotY*d.mass;b+=d.mass});return this.barycenter={x:a,y:c,xFactor:a/b,
yFactor:c/b}},barnesHutApproximation:function(b,a){var c=this.getDistXY(b,a),d=this.vectorLength(c);if(b!==a&&0!==d)if(a.isInternal)if(a.boxSize/d<this.options.theta&&0!==d){var e=this.repulsiveForce(d,this.k);this.force("repulsive",b,e*a.mass,c,d);var g=!1}else g=!0;else e=this.repulsiveForce(d,this.k),this.force("repulsive",b,e*a.mass,c,d);return g},repulsiveForces:function(){var b=this;"barnes-hut"===b.approximation?b.nodes.forEach(function(a){b.quadTree.visitNodeRecursive(null,function(c){return b.barnesHutApproximation(a,
c)})}):b.nodes.forEach(function(a){b.nodes.forEach(function(c){if(a!==c&&!a.fixedPosition){var d=b.getDistXY(a,c);var e=b.vectorLength(d);if(0!==e){var g=b.repulsiveForce(e,b.k);b.force("repulsive",a,g*c.mass,d,e)}}})})},attractiveForces:function(){var b=this,a,c,d;b.links.forEach(function(e){e.fromNode&&e.toNode&&(a=b.getDistXY(e.fromNode,e.toNode),c=b.vectorLength(a),0!==c&&(d=b.attractiveForce(c,b.k),b.force("attractive",e,d,a,c)))})},applyLimits:function(){var a=this;a.nodes.forEach(function(b){b.fixedPosition||
(a.integration.integrate(a,b),a.applyLimitBox(b,a.box),b.dispX=0,b.dispY=0)})},applyLimitBox:function(a,c){var b=a.radius;a.plotX=h(a.plotX,c.left+b,c.width-b);a.plotY=h(a.plotY,c.top+b,c.height-b)},coolDown:function(a,c,d){return a-c*d},isStable:function(){return.00001>Math.abs(this.systemTemperature-this.prevSystemTemperature)||0>=this.temperature},getSystemTemperature:function(){return this.nodes.reduce(function(a,c){return a+c.temperature},0)},vectorLength:function(a){return Math.sqrt(a.x*a.x+
a.y*a.y)},getDistR:function(a,c){a=this.getDistXY(a,c);return this.vectorLength(a)},getDistXY:function(a,c){var b=a.plotX-c.plotX;a=a.plotY-c.plotY;return{x:b,y:a,absX:Math.abs(b),absY:Math.abs(a)}}});a(f,"predraw",function(){this.graphLayoutsLookup&&this.graphLayoutsLookup.forEach(function(a){a.stop()})});a(f,"render",function(){function a(a){a.maxIterations--&&isFinite(a.temperature)&&!a.isStable()&&!a.enableSimulation&&(a.beforeStep&&a.beforeStep(),a.step(),e=!1,d=!0)}var d=!1;if(this.graphLayoutsLookup){c(!1,
this);for(this.graphLayoutsLookup.forEach(function(a){a.start()});!e;){var e=!0;this.graphLayoutsLookup.forEach(a)}d&&this.series.forEach(function(a){a&&a.layout&&a.render()})}});a(f,"beforePrint",function(){this.graphLayoutsLookup&&(this.graphLayoutsLookup.forEach(function(a){a.updateSimulation(!1)}),this.redraw())});a(f,"afterPrint",function(){this.graphLayoutsLookup&&this.graphLayoutsLookup.forEach(function(a){a.updateSimulation()});this.redraw()})});m(f,"Series/Networkgraph/DraggableNodes.js",
[f["Core/Chart/Chart.js"],f["Core/Globals.js"],f["Core/Utilities.js"]],function(f,a,d){var e=d.addEvent;a.dragNodesMixin={onMouseDown:function(a,d){d=this.chart.pointer.normalize(d);a.fixedPosition={chartX:d.chartX,chartY:d.chartY,plotX:a.plotX,plotY:a.plotY};a.inDragMode=!0},onMouseMove:function(a,d){if(a.fixedPosition&&a.inDragMode){var c=this.chart,e=c.pointer.normalize(d);d=a.fixedPosition.chartX-e.chartX;e=a.fixedPosition.chartY-e.chartY;var f=void 0,h=void 0,b=c.graphLayoutsLookup;if(5<Math.abs(d)||
5<Math.abs(e))f=a.fixedPosition.plotX-d,h=a.fixedPosition.plotY-e,c.isInsidePlot(f,h)&&(a.plotX=f,a.plotY=h,a.hasDragged=!0,this.redrawHalo(a),b.forEach(function(a){a.restartSimulation()}))}},onMouseUp:function(a,d){a.fixedPosition&&(a.hasDragged&&(this.layout.enableSimulation?this.layout.start():this.chart.redraw()),a.inDragMode=a.hasDragged=!1,this.options.fixedDraggable||delete a.fixedPosition)},redrawHalo:function(a){a&&this.halo&&this.halo.attr({d:a.haloPath(this.options.states.hover.halo.size)})}};
e(f,"load",function(){var a=this,d,f,n;a.container&&(d=e(a.container,"mousedown",function(d){var c=a.hoverPoint;c&&c.series&&c.series.hasDraggableNodes&&c.series.options.draggable&&(c.series.onMouseDown(c,d),f=e(a.container,"mousemove",function(a){return c&&c.series&&c.series.onMouseMove(c,a)}),n=e(a.container.ownerDocument,"mouseup",function(a){f();n();return c&&c.series&&c.series.onMouseUp(c,a)}))}));e(a,"destroy",function(){d()})})});m(f,"Series/Networkgraph/Networkgraph.js",[f["Core/Globals.js"],
f["Series/NodesComposition.js"],f["Core/Series/Point.js"],f["Core/Series/Series.js"],f["Core/Series/SeriesRegistry.js"],f["Core/Utilities.js"]],function(f,a,d,e,c,h){var m=this&&this.__extends||function(){var a=function(b,c){a=Object.setPrototypeOf||{__proto__:[]}instanceof Array&&function(a,b){a.__proto__=b}||function(a,b){for(var c in b)b.hasOwnProperty(c)&&(a[c]=b[c])};return a(b,c)};return function(b,c){function d(){this.constructor=b}a(b,c);b.prototype=null===c?Object.create(c):(d.prototype=
c.prototype,new d)}}(),n=c.seriesTypes,r=h.addEvent,l=h.css,b=h.defined,t=h.extend,u=h.merge,v=h.pick,q=f.dragNodesMixin;"";h=function(a){function b(){var b=null!==a&&a.apply(this,arguments)||this;b.data=void 0;b.nodes=void 0;b.options=void 0;b.points=void 0;return b}m(b,a);b.defaultOptions=u(e.defaultOptions,{stickyTracking:!1,inactiveOtherPoints:!0,marker:{enabled:!0,states:{inactive:{opacity:.3,animation:{duration:50}}}},states:{inactive:{linkOpacity:.3,animation:{duration:50}}},dataLabels:{formatter:function(){return this.key},
linkFormatter:function(){return this.point.fromNode.name+"<br>"+this.point.toNode.name},linkTextPath:{enabled:!0},textPath:{enabled:!1},style:{transition:"opacity 2000ms"}},link:{color:"rgba(100, 100, 100, 0.5)",width:1},draggable:!0,layoutAlgorithm:{initialPositions:"circle",initialPositionRadius:1,enableSimulation:!1,theta:.5,maxSpeed:10,approximation:"none",type:"reingold-fruchterman",integration:"euler",maxIterations:1E3,gravitationalConstant:.0625,friction:-.981},showInLegend:!1});return b}(e);
t(h.prototype,{forces:["barycenter","repulsive","attractive"],hasDraggableNodes:!0,drawGraph:void 0,isCartesian:!1,requireSorting:!1,directTouch:!0,noSharedTooltip:!0,pointArrayMap:["from","to"],trackerGroups:["group","markerGroup","dataLabelsGroup"],drawTracker:n.column.prototype.drawTracker,animate:void 0,buildKDTree:f.noop,createNode:a.createNode,destroy:function(){this.layout&&this.layout.removeElementFromCollection(this,this.layout.series);a.destroy.call(this)},init:function(){var a=this;e.prototype.init.apply(this,
arguments);r(this,"updatedData",function(){a.layout&&a.layout.stop()});r(this,"afterUpdate",function(){a.nodes.forEach(function(a){a&&a.series&&a.resolveColor()})});return this},generatePoints:function(){var b;a.generatePoints.apply(this,arguments);this.options.nodes&&this.options.nodes.forEach(function(a){this.nodeLookup[a.id]||(this.nodeLookup[a.id]=this.createNode(a.id))},this);for(b=this.nodes.length-1;0<=b;b--){var c=this.nodes[b];c.degree=c.getDegree();c.radius=v(c.marker&&c.marker.radius,this.options.marker&&
this.options.marker.radius,0);this.nodeLookup[c.id]||c.remove()}this.data.forEach(function(a){a.formatPrefix="link"});this.indexateNodes()},getPointsCollection:function(){return this.nodes||[]},indexateNodes:function(){this.nodes.forEach(function(a,b){a.index=b})},markerAttribs:function(a,c){c=e.prototype.markerAttribs.call(this,a,c);b(a.plotY)||(c.y=0);c.x=(a.plotX||0)-(c.width||0)/2;return c},translate:function(){this.processedXData||this.processData();this.generatePoints();this.deferLayout();this.nodes.forEach(function(a){a.isInside=
!0;a.linksFrom.forEach(function(a){a.shapeType="path";a.y=1})})},deferLayout:function(){var a=this.options.layoutAlgorithm,c=this.chart.graphLayoutsStorage,d=this.chart.graphLayoutsLookup,e=this.chart.options.chart;if(this.visible){c||(this.chart.graphLayoutsStorage=c={},this.chart.graphLayoutsLookup=d=[]);var h=c[a.type];h||(a.enableSimulation=b(e.forExport)?!e.forExport:a.enableSimulation,c[a.type]=h=new f.layouts[a.type],h.init(a),d.splice(h.index,0,h));this.layout=h;h.setArea(0,0,this.chart.plotWidth,
this.chart.plotHeight);h.addElementsToCollection([this],h.series);h.addElementsToCollection(this.nodes,h.nodes);h.addElementsToCollection(this.points,h.links)}},render:function(){var a=this.points,b=this.chart.hoverPoint,c=[];this.points=this.nodes;n.line.prototype.render.call(this);this.points=a;a.forEach(function(a){a.fromNode&&a.toNode&&(a.renderLink(),a.redrawLink())});b&&b.series===this&&this.redrawHalo(b);this.chart.hasRendered&&!this.options.dataLabels.allowOverlap&&(this.nodes.concat(this.points).forEach(function(a){a.dataLabel&&
c.push(a.dataLabel)}),this.chart.hideOverlappingLabels(c))},drawDataLabels:function(){var a=this.options.dataLabels.textPath;e.prototype.drawDataLabels.apply(this,arguments);this.points=this.data;this.options.dataLabels.textPath=this.options.dataLabels.linkTextPath;e.prototype.drawDataLabels.apply(this,arguments);this.points=this.nodes;this.options.dataLabels.textPath=a},pointAttribs:function(a,b){var c=b||a&&a.state||"normal";b=e.prototype.pointAttribs.call(this,a,c);c=this.options.states[c];a&&
!a.isNode&&(b=a.getLinkAttributes(),c&&(b={stroke:c.linkColor||b.stroke,dashstyle:c.linkDashStyle||b.dashstyle,opacity:v(c.linkOpacity,b.opacity),"stroke-width":c.linkColor||b["stroke-width"]}));return b},redrawHalo:q.redrawHalo,onMouseDown:q.onMouseDown,onMouseMove:q.onMouseMove,onMouseUp:q.onMouseUp,setState:function(a,b){b?(this.points=this.nodes.concat(this.data),e.prototype.setState.apply(this,arguments),this.points=this.data):e.prototype.setState.apply(this,arguments);this.layout.simulation||
a||this.render()}});q=function(a){function b(){var b=null!==a&&a.apply(this,arguments)||this;b.degree=void 0;b.linksFrom=void 0;b.linksTo=void 0;b.options=void 0;b.radius=void 0;b.series=void 0;b.toNode=void 0;return b}m(b,a);return b}(e.prototype.pointClass);t(q.prototype,{setState:a.setNodeState,init:function(){d.prototype.init.apply(this,arguments);this.series.options.draggable&&!this.series.chart.styledMode&&(r(this,"mouseOver",function(){l(this.series.chart.container,{cursor:"move"})}),r(this,
"mouseOut",function(){l(this.series.chart.container,{cursor:"default"})}));return this},getDegree:function(){var a=this.isNode?this.linksFrom.length+this.linksTo.length:0;return 0===a?1:a},getLinkAttributes:function(){var a=this.series.options.link,b=this.options;return{"stroke-width":v(b.width,a.width),stroke:b.color||a.color,dashstyle:b.dashStyle||a.dashStyle,opacity:v(b.opacity,a.opacity,1)}},renderLink:function(){if(!this.graphic&&(this.graphic=this.series.chart.renderer.path(this.getLinkPath()).addClass(this.getClassName(),
!0).add(this.series.group),!this.series.chart.styledMode)){var a=this.series.pointAttribs(this);this.graphic.attr(a);(this.dataLabels||[]).forEach(function(b){b&&b.attr({opacity:a.opacity})})}},redrawLink:function(){var a=this.getLinkPath();if(this.graphic){this.shapeArgs={d:a};if(!this.series.chart.styledMode){var b=this.series.pointAttribs(this);this.graphic.attr(b);(this.dataLabels||[]).forEach(function(a){a&&a.attr({opacity:b.opacity})})}this.graphic.animate(this.shapeArgs);var c=a[0];a=a[1];
"M"===c[0]&&"L"===a[0]&&(this.plotX=(c[1]+a[1])/2,this.plotY=(c[2]+a[2])/2)}},getMass:function(){var a=this.fromNode.mass,b=this.toNode.mass,c=a+b;return{fromNode:1-a/c,toNode:1-b/c}},getLinkPath:function(){var a=this.fromNode,b=this.toNode;a.plotX>b.plotX&&(a=this.toNode,b=this.fromNode);return[["M",a.plotX||0,a.plotY||0],["L",b.plotX||0,b.plotY||0]]},isValid:function(){return!this.isNode||b(this.id)},remove:function(a,b){var c=this.series,d=c.options.nodes||[],f,g=d.length;if(this.isNode){c.points=
[];[].concat(this.linksFrom).concat(this.linksTo).forEach(function(a){f=a.fromNode.linksFrom.indexOf(a);-1<f&&a.fromNode.linksFrom.splice(f,1);f=a.toNode.linksTo.indexOf(a);-1<f&&a.toNode.linksTo.splice(f,1);e.prototype.removePoint.call(c,c.data.indexOf(a),!1,!1)});c.points=c.data.slice();for(c.nodes.splice(c.nodes.indexOf(this),1);g--;)if(d[g].id===this.options.id){c.options.nodes.splice(g,1);break}this&&this.destroy();c.isDirty=!0;c.isDirtyData=!0;a&&c.chart.redraw(a)}else c.removePoint(c.data.indexOf(this),
a,b)},destroy:function(){this.isNode&&this.linksFrom.concat(this.linksTo).forEach(function(a){a.destroyElements&&a.destroyElements()});this.series.layout.removeElementFromCollection(this,this.series.layout[this.isNode?"nodes":"links"]);return d.prototype.destroy.apply(this,arguments)}});h.prototype.pointClass=q;c.registerSeriesType("networkgraph",h);"";return h});m(f,"masters/modules/networkgraph.src.js",[],function(){})});
//# sourceMappingURL=networkgraph.js.map
```
