## Supplementary material for "Exploring natural odour landscapes: A case study with implications for human-biting insects": Figure S4: no-data-to-display.js

```
/*
 Highcharts JS v9.3.1 (2021-11-05)

 Plugin for displaying a message when there is no data visible in chart.

 (c) 2010-2021 Highsoft AS
 Author: Oystein Moseng

 License: www.highcharts.com/license
*/
'use strict';(function(a){"object"===typeof module&&module.exports?(a["default"]=a,module.exports=a):"function"===typeof define&&define.amd?define("highcharts/modules/no-data-to-display",["highcharts"],function(b){a(b);a.Highcharts=b;return a}):a("undefined"!==typeof Highcharts?Highcharts:void 0)})(function(a){function b(a,b,e,d){a.hasOwnProperty(b)||(a[b]=d.apply(null,e))}a=a?a._modules:{};b(a,"Extensions/NoDataToDisplay.js",[a["Core/Renderer/HTML/AST.js"],a["Core/Chart/Chart.js"],a["Core/DefaultOptions.js"],
a["Core/Utilities.js"]],function(a,b,e,d){var f=e.getOptions;e=d.addEvent;var g=d.extend;d=b.prototype;f=f();g(f.lang,{noData:"No data to display"});f.noData={attr:{zIndex:1},position:{x:0,y:0,align:"center",verticalAlign:"middle"},style:{fontWeight:"bold",fontSize:"12px",color:"#666666"}};d.showNoData=function(b){var c=this.options;b=b||c&&c.lang.noData||"";c=c&&(c.noData||{});this.renderer&&(this.noDataLabel||(this.noDataLabel=this.renderer.label(b,0,0,void 0,void 0,void 0,c.useHTML,void 0,"no-data").add()),
this.styledMode||this.noDataLabel.attr(a.filterUserAttributes(c.attr||{})).css(c.style||{}),this.noDataLabel.align(g(this.noDataLabel.getBBox(),c.position||{}),!1,"plotBox"))};d.hideNoData=function(){this.noDataLabel&&(this.noDataLabel=this.noDataLabel.destroy())};d.hasData=function(){for(var a=this.series||[],b=a.length;b--;)if(a[b].hasData()&&!a[b].options.isInternal)return!0;return this.loadingShown};e(b,"render",function(){this.hasData()?this.hideNoData():this.showNoData()})});b(a,"masters/modules/no-data-to-display.src.js",
[],function(){})});
//# sourceMappingURL=no-data-to-display.js.map
```
