## Supplementary material for "Exploring natural odour landscapes: A case study with implications for human-biting insects": Figure S4: offline-exporting.js

```
/*
 Highcharts JS v9.3.1 (2021-11-05)

 Client side exporting module

 (c) 2015-2021 Torstein Honsi / Oystein Moseng

 License: www.highcharts.com/license
*/
'use strict';(function(a){"object"===typeof module&&module.exports?(a["default"]=a,module.exports=a):"function"===typeof define&&define.amd?define("highcharts/modules/offline-exporting",["highcharts","highcharts/modules/exporting"],function(k){a(k);a.Highcharts=k;return a}):a("undefined"!==typeof Highcharts?Highcharts:void 0)})(function(a){function k(a,t,e,r){a.hasOwnProperty(t)||(a[t]=r.apply(null,e))}a=a?a._modules:{};k(a,"Extensions/DownloadURL.js",[a["Core/Globals.js"]],function(a){var t=a.isSafari,
e=a.win,r=e.document,l=e.URL||e.webkitURL||e,k=a.dataURLtoBlob=function(a){if((a=a.replace(/filename=.*;/,"").match(/data:([^;]*)(;base64)?,([0-9A-Za-z+/]+)/))&&3<a.length&&e.atob&&e.ArrayBuffer&&e.Uint8Array&&e.Blob&&l.createObjectURL){var f=e.atob(a[3]),m=new e.ArrayBuffer(f.length);m=new e.Uint8Array(m);for(var b=0;b<m.length;++b)m[b]=f.charCodeAt(b);a=new e.Blob([m],{type:a[1]});return l.createObjectURL(a)}};a=a.downloadURL=function(a,l){var m=e.navigator,b=r.createElement("a");if("string"===
typeof a||a instanceof String||!m.msSaveOrOpenBlob){a=""+a;m=/Edge\/\d+/.test(m.userAgent);if(t&&"string"===typeof a&&0===a.indexOf("data:application/pdf")||m||2E6<a.length)if(a=k(a)||"",!a)throw Error("Failed to convert to blob");if("undefined"!==typeof b.download)b.href=a,b.download=l,r.body.appendChild(b),b.click(),r.body.removeChild(b);else try{var f=e.open(a,"chart");if("undefined"===typeof f||null===f)throw Error("Failed to open window");}catch(L){e.location.href=a}}else m.msSaveOrOpenBlob(a,
l)};return{dataURLtoBlob:k,downloadURL:a}});k(a,"Extensions/OfflineExporting/OfflineExportingDefaults.js",[],function(){return{libURL:"https://code.highcharts.com/9.3.1/lib/",menuItemDefinitions:{downloadPNG:{textKey:"downloadPNG",onclick:function(){this.exportChartLocal()}},downloadJPEG:{textKey:"downloadJPEG",onclick:function(){this.exportChartLocal({type:"image/jpeg"})}},downloadSVG:{textKey:"downloadSVG",onclick:function(){this.exportChartLocal({type:"image/svg+xml"})}},downloadPDF:{textKey:"downloadPDF",
onclick:function(){this.exportChartLocal({type:"application/pdf"})}}}}});k(a,"Extensions/OfflineExporting/OfflineExporting.js",[a["Core/Chart/Chart.js"],a["Core/DefaultOptions.js"],a["Extensions/DownloadURL.js"],a["Extensions/Exporting/Exporting.js"],a["Core/Globals.js"],a["Extensions/OfflineExporting/OfflineExportingDefaults.js"],a["Core/Utilities.js"]],function(a,k,e,r,l,G,f){var t=k.defaultOptions,m=e.downloadURL,b=l.win,w=l.doc,y=f.addEvent,x=f.error,H=f.extend,I=f.fireEvent,B=f.merge,C=[],v;
(function(a){function e(g,d){var h=this,c=B(h.options.exporting,g),b=function(a){!1===c.fallbackToExportServer?c.error?c.error(c,a):x(28,!0):h.exportChart(c)};g=function(){return[].some.call(h.container.getElementsByTagName("image"),function(a){a=a.getAttribute("href");return""!==a&&0!==a.indexOf("data:")})};l.isMS&&h.styledMode&&!r.inlineWhitelist.length&&r.inlineWhitelist.push(/^blockSize/,/^border/,/^caretColor/,/^color/,/^columnRule/,/^columnRuleColor/,/^cssFloat/,/^cursor/,/^fill$/,/^fillOpacity/,
/^font/,/^inlineSize/,/^length/,/^lineHeight/,/^opacity/,/^outline/,/^parentRule/,/^rx$/,/^ry$/,/^stroke/,/^textAlign/,/^textAnchor/,/^textDecoration/,/^transform/,/^vectorEffect/,/^visibility/,/^x$/,/^y$/);l.isMS&&("application/pdf"===c.type||h.container.getElementsByTagName("image").length&&"image/svg+xml"!==c.type)||"application/pdf"===c.type&&g()?b("Image type not supported for this chart/browser."):h.getSVGForLocalExport(c,d||{},b,function(g){-1<g.indexOf("<foreignObject")&&"image/svg+xml"!==
c.type&&(l.isMS||"application/pdf"===c.type)?b("Image type not supportedfor charts with embedded HTML"):a.downloadSVGLocal(g,H({filename:h.getFilename()},c),b,function(){return I(h,"exportChartLocalSuccess")})})}function k(a,d){var g=w.getElementsByTagName("head")[0],c=w.createElement("script");c.type="text/javascript";c.src=a;c.onload=d;c.onerror=function(){x("Error loading script "+a)};g.appendChild(c)}function f(g,d,h,c){var b=this,e=function(){f&&q===l&&c(b.sanitizeSVG(m.innerHTML,p))},k=function(a,
g,c){++q;c.imageElement.setAttributeNS("http://www.w3.org/1999/xlink","href",a);e()},m,p,n=null,f,l=0,q=0;b.unbindGetSVG=y(b,"getSVG",function(a){p=a.chartCopy.options;f=(m=a.chartCopy.container.cloneNode(!0))&&m.getElementsByTagName("image")||[];l=f.length});b.getSVGForExport(g,d);try{if(!f||!f.length){c(b.sanitizeSVG(m.innerHTML,p));return}for(d=0;d<f.length;d++){var u=f[d];(n=u.getAttributeNS("http://www.w3.org/1999/xlink","href"))?a.imageToDataUrl(n,"image/png",{imageElement:u},g.scale,k,h,h,
h):(q++,u.parentNode.removeChild(u),d--,e())}}catch(z){h(z)}b.unbindGetSVG()}function v(g,d,h,c,e,f,m,k,p){var n=new b.Image,l=function(){setTimeout(function(){var a=w.createElement("canvas"),b=a.getContext&&a.getContext("2d");try{if(b){a.height=n.height*c;a.width=n.width*c;b.drawImage(n,0,0,a.width,a.height);try{var f=a.toDataURL(d);e(f,d,h,c)}catch(E){q(g,d,h,c)}}else m(g,d,h,c)}finally{p&&p(g,d,h,c)}},a.loadEventDeferDelay)},D=function(){k(g,d,h,c);p&&p(g,d,h,c)};var q=function(){n=new b.Image;
q=f;n.crossOrigin="Anonymous";n.onload=l;n.onerror=D;n.src=g};n.onload=l;n.onerror=D;n.src=g}function A(g){var d=b.navigator.userAgent;d=-1<d.indexOf("WebKit")&&0>d.indexOf("Chrome");try{if(!d&&-1===g.indexOf("<foreignObject"))return a.domurl.createObjectURL(new b.Blob([g],{type:"image/svg+xml;charset-utf-16"}))}catch(h){}return"data:image/svg+xml;charset=UTF-8,"+encodeURIComponent(g)}function F(a,d){var g=a.width.baseVal.value+2*d;d=a.height.baseVal.value+2*d;g=new b.jsPDF(d>g?"p":"l","pt",[g,d]);
[].forEach.call(a.querySelectorAll('*[visibility="hidden"]'),function(a){a.parentNode.removeChild(a)});d=a.querySelectorAll("linearGradient");for(var c=0;c<d.length;c++)for(var e=d[c].querySelectorAll("stop"),f=0;f<e.length&&"0"===e[f].getAttribute("offset")&&"0"===e[f+1].getAttribute("offset");)e[f].remove(),f++;[].forEach.call(a.querySelectorAll("tspan"),function(a){"\u200b"===a.textContent&&(a.textContent=" ",a.setAttribute("dx",-5))});b.svg2pdf(a,g,{removeInvalid:!0});return g.output("datauristring")}
a.CanVGRenderer={};a.domurl=b.URL||b.webkitURL||b;a.loadEventDeferDelay=l.isMS?150:0;a.compose=function(a){if(-1===C.indexOf(a)){C.push(a);var b=a.prototype;b.getSVGForLocalExport=f;b.exportChartLocal=e;B(!0,t.exporting,G)}return a};a.downloadSVGLocal=function(g,d,e,c){var f=w.createElement("div"),h=d.type||"image/png",l=(d.filename||"chart")+"."+("image/svg+xml"===h?"svg":h.split("/")[1]),r=d.scale||1,p=d.libURL||t.exporting.libURL,n=!0;p="/"!==p.slice(-1)?p+"/":p;var y=function(){f.innerHTML=g;
var a=f.getElementsByTagName("text"),b;[].forEach.call(a,function(a){["font-family","font-size"].forEach(function(b){for(var c=a;c&&c!==f;){if(c.style[b]){a.style[b]=c.style[b];break}c=c.parentNode}});a.style["font-family"]=a.style["font-family"]&&a.style["font-family"].split(" ").splice(-1);b=a.getElementsByTagName("title");[].forEach.call(b,function(b){a.removeChild(b)})});a=F(f.firstChild,0);try{m(a,l),c&&c()}catch(E){e(E)}};if("image/svg+xml"===h)try{if("undefined"!==typeof b.navigator.msSaveOrOpenBlob){var x=
new MSBlobBuilder;x.append(g);var q=x.getBlob("image/svg+xml")}else q=A(g);m(q,l);c&&c()}catch(z){e(z)}else if("application/pdf"===h)b.jsPDF&&b.svg2pdf?y():(n=!0,k(p+"jspdf.js",function(){k(p+"svg2pdf.js",function(){y()})}));else{q=A(g);var u=function(){try{a.domurl.revokeObjectURL(q)}catch(z){}};v(q,h,{},r,function(a){try{m(a,l),c&&c()}catch(J){e(J)}},function(){var a=w.createElement("canvas"),d=a.getContext("2d"),f=g.match(/^<svg[^>]*width\s*=\s*"?(\d+)"?[^>]*>/)[1]*r,q=g.match(/^<svg[^>]*height\s*=\s*"?(\d+)"?[^>]*>/)[1]*
r,t=function(){b.canvg.Canvg.fromString(d,g).start();try{m(b.navigator.msSaveOrOpenBlob?a.msToBlob():a.toDataURL(h),l),c&&c()}catch(K){e(K)}finally{u()}};a.width=f;a.height=q;b.canvg?t():(n=!0,k(p+"canvg.js",function(){t()}))},e,e,function(){n&&u()})}};a.getScript=k;a.imageToDataUrl=v;a.svgToDataUrl=A;a.svgToPdf=F})(v||(v={}));return v});k(a,"masters/modules/offline-exporting.src.js",[a["Core/Globals.js"],a["Extensions/OfflineExporting/OfflineExporting.js"]],function(a,k){a.downloadSVGLocal=k.downloadSVGLocal;
k.compose(a.Chart)})});
//# sourceMappingURL=offline-exporting.js.map
```
