## Supplementary material for "Exploring natural odour landscapes: A case study with implications for human-biting insects": Figure S4: oldie-polyfills.js

```
/*
 Highcharts JS v9.3.1 (2021-11-05)

 Old IE (v6, v7, v8) array polyfills for Highcharts v7+.

 (c) 2010-2021 Highsoft AS
 Author: Torstein Honsi

 License: www.highcharts.com/license
*/
'use strict';(function(e){"object"===typeof module&&module.exports?(e["default"]=e,module.exports=e):"function"===typeof define&&define.amd?define("highcharts/modules/oldie-polyfills",["highcharts"],function(f){e(f);e.Highcharts=f;return e}):e("undefined"!==typeof Highcharts?Highcharts:void 0)})(function(e){function f(c,b,a,d){c.hasOwnProperty(b)||(c[b]=d.apply(null,a))}e=e?e._modules:{};f(e,"Extensions/OldiePolyfills.js",[],function(){String.prototype.trim||(String.prototype.trim=function(){return this.replace(/^[\s\uFEFF\xA0]+|[\s\uFEFF\xA0]+$/g,
"")});Array.prototype.forEach||(Array.prototype.forEach=function(c,b){for(var a=0,d=this.length;a<d;a++)if("undefined"!==typeof this[a]&&!1===c.call(b,this[a],a,this))return a});Array.prototype.map||(Array.prototype.map=function(c){for(var b=[],a=0,d=this.length;a<d;a++)b[a]=c.call(this[a],this[a],a,this);return b});Array.prototype.indexOf||(Array.prototype.indexOf=function(c,b){var a=b||0;if(this)for(b=this.length;a<b;a++)if(this[a]===c)return a;return-1});Array.prototype.filter||(Array.prototype.filter=
function(c){for(var b=[],a=0,d=this.length;a<d;a++)c(this[a],a)&&b.push(this[a]);return b});Array.prototype.some||(Array.prototype.some=function(c,b){for(var a=0,d=this.length;a<d;a++)if(!0===c.call(b,this[a],a,this))return!0;return!1});Array.prototype.reduce||(Array.prototype.reduce=function(c,b){for(var a=1<arguments.length?0:1,d=1<arguments.length?b:this[0],e=this.length;a<e;++a)d=c.call(this,d,this[a],a,this);return d});Function.prototype.bind||(Function.prototype.bind=function(){var c=this,b=
arguments[0],a=Array.prototype.slice.call(arguments,1);if("function"!==typeof c)throw new TypeError("Function.prototype.bind - what is trying to be bound is not callable");return function(){var d=a.concat(Array.prototype.slice.call(arguments));return c.apply(b,d)}});Object.getPrototypeOf||(Object.getPrototypeOf="object"===typeof"test".__proto__?function(c){return c.__proto__}:function(c){var b=c.constructor.prototype;return b===c?{}.constructor.prototype:b});Object.keys||(Object.keys=function(c){var b=
[],a;for(a in c)Object.hasOwnProperty.call(c,a)&&b.push(a);return b});document.getElementsByClassName||(document.getElementsByClassName=function(c){var b=document,a,d=[];if(b.querySelectorAll)return b.querySelectorAll("."+c);if(b.evaluate)for(b=b.evaluate(".//*[contains(concat(' ', @class, ' '), ' "+c+" ')]",b,null,0,null);a=b.iterateNext();)d.push(a);else for(b=b.getElementsByTagName("*"),c=new RegExp("(^|\\s)"+c+"(\\s|$)"),a=0;a<b.length;a++)c.test(b[a].className)&&d.push(b[a]);return d})});f(e,
"masters/modules/oldie-polyfills.src.js",[],function(){})});
//# sourceMappingURL=oldie-polyfills.js.map
```
