## Supplementary material for "Exploring natural odour landscapes: A case study with implications for human-biting insects": Figure S4: oldie.js

```
/*
 Highcharts JS v9.3.1 (2021-11-05)

 Old IE (v6, v7, v8) module for Highcharts v6+.

 (c) 2010-2021 Highsoft AS
 Author: Torstein Honsi

 License: www.highcharts.com/license
*/
'use strict';(function(c){"object"===typeof module&&module.exports?(c["default"]=c,module.exports=c):"function"===typeof define&&define.amd?define("highcharts/modules/oldie",["highcharts"],function(p){c(p);c.Highcharts=p;return c}):c("undefined"!==typeof Highcharts?Highcharts:void 0)})(function(c){function p(c,r,e,D){c.hasOwnProperty(r)||(c[r]=D.apply(null,e))}c=c?c._modules:{};p(c,"Extensions/Oldie/VMLAxis3D.js",[c["Core/Utilities.js"]],function(c){var x=c.addEvent,e=function(){return function(c){this.axis=
c}}();return function(){function c(){}c.compose=function(l){l.keepProps.push("vml");x(l,"destroy",c.onDestroy);x(l,"init",c.onInit);x(l,"render",c.onRender)};c.onDestroy=function(){var c=this.vml;if(c){var e;["backFrame","bottomFrame","sideFrame"].forEach(function(l){(e=c[l])&&(c[l]=e.destroy())},this)}};c.onInit=function(){this.vml||(this.vml=new e(this))};c.onRender=function(){var c=this.vml;c.sideFrame&&(c.sideFrame.css({zIndex:0}),c.sideFrame.front.attr({fill:c.sideFrame.color}));c.bottomFrame&&
(c.bottomFrame.css({zIndex:1}),c.bottomFrame.front.attr({fill:c.bottomFrame.color}));c.backFrame&&(c.backFrame.css({zIndex:0}),c.backFrame.front.attr({fill:c.backFrame.color}))};return c}()});p(c,"Extensions/Oldie/VMLRenderer3D.js",[c["Core/Axis/Axis.js"],c["Core/DefaultOptions.js"],c["Extensions/Oldie/VMLAxis3D.js"]],function(c,r,e){var x=r.setOptions;return function(){function l(){}l.compose=function(g,l){var h=l.prototype;g=g.prototype;x({animate:!1});g.face3d=h.face3d;g.polyhedron=h.polyhedron;
g.elements3d=h.elements3d;g.element3d=h.element3d;g.cuboid=h.cuboid;g.cuboidPath=h.cuboidPath;g.toLinePath=h.toLinePath;g.toLineSegments=h.toLineSegments;g.arc3d=function(c){c=h.arc3d.call(this,c);c.css({zIndex:c.zIndex});return c};g.arc3dPath=h.arc3dPath;e.compose(c)};return l}()});p(c,"Extensions/Oldie/Oldie.js",[c["Core/Chart/Chart.js"],c["Core/Color/Color.js"],c["Core/Globals.js"],c["Core/DefaultOptions.js"],c["Core/Pointer.js"],c["Core/Renderer/RendererRegistry.js"],c["Core/Renderer/SVG/SVGElement.js"],
c["Core/Renderer/SVG/SVGRenderer.js"],c["Core/Utilities.js"],c["Extensions/Oldie/VMLRenderer3D.js"]],function(c,r,e,p,l,g,y,h,m,Q){var x=r.parse,u=e.deg2rad,k=e.doc;r=e.noop;var E=e.svg,v=e.win,M=p.getOptions,D=m.addEvent,F=m.createElement,z=m.css,H=m.defined,I=m.discardElement,N=m.erase,w=m.extend;p=m.extendClass;var R=m.isArray,O=m.isNumber,G=m.isObject,B=m.pick,t=m.pInt,S=m.uniqueKey;M().global.VMLRadialGradientURL="http://code.highcharts.com/9.3.1/gfx/vml-radial-gradient.png";k&&!k.defaultView&&
(e.getStyle=m.getStyle=function q(b,d){var c={width:"clientWidth",height:"clientHeight"}[d];if(b.style[d])return t(b.style[d]);"opacity"===d&&(d="filter");if(c)return b.style.zoom=1,Math.max(b[c]-2*q(b,"padding"),0);b=b.currentStyle[d.replace(/\-(\w)/g,function(b,d){return d.toUpperCase()})];"filter"===d&&(b=b.replace(/alpha\(opacity=([0-9]+)\)/,function(b,d){return d/100}));return""===b?1:t(b)});E||(D(y,"afterInit",function(){"text"===this.element.nodeName&&this.css({position:"absolute"})}),l.prototype.normalize=
function(a,b){a=a||v.event;a.target||(a.target=a.srcElement);b||(this.chartPosition=b=this.getChartPosition());return w(a,{chartX:Math.round(Math.max(a.x,a.clientX-b.left)),chartY:Math.round(a.y)})},c.prototype.ieSanitizeSVG=function(a){return a=a.replace(/<IMG /g,"<image ").replace(/<(\/?)TITLE>/g,"<$1title>").replace(/height=([^" ]+)/g,'height="$1"').replace(/width=([^" ]+)/g,'width="$1"').replace(/hc-svg-href="([^"]+)">/g,'xlink:href="$1"/>').replace(/ id=([^" >]+)/g,' id="$1"').replace(/class=([^" >]+)/g,
'class="$1"').replace(/ transform /g," ").replace(/:(path|rect)/g,"$1").replace(/style="([^"]+)"/g,function(a){return a.toLowerCase()})},c.prototype.isReadyToRender=function(){var a=this;return E||v!=v.top||"complete"===k.readyState?!0:(k.attachEvent("onreadystatechange",function(){k.detachEvent("onreadystatechange",a.firstRender);"complete"===k.readyState&&a.firstRender()}),!1)},k.createElementNS||(k.createElementNS=function(a,b){return k.createElement(b)}),e.addEventListenerPolyfill=function(a,
b){function d(a){a.target=a.srcElement||v;b.call(c,a)}var c=this;c.attachEvent&&(c.hcEventsIE||(c.hcEventsIE={}),b.hcKey||(b.hcKey=S()),c.hcEventsIE[b.hcKey]=d,c.attachEvent("on"+a,d))},e.removeEventListenerPolyfill=function(a,b){this.detachEvent&&(b=this.hcEventsIE[b.hcKey],this.detachEvent("on"+a,b))},c={docMode8:k&&8===k.documentMode,init:function(a,b){var d=["<",b,' filled="f" stroked="f"'],c=["position: ","absolute",";"],f="div"===b;("shape"===b||f)&&c.push("left:0;top:0;width:1px;height:1px;");
c.push("visibility: ",f?"hidden":"visible");d.push(' style="',c.join(""),'"/>');b&&(d=f||"span"===b||"img"===b?d.join(""):a.prepVML(d),this.element=F(d));this.renderer=a},add:function(a){var b=this.renderer,d=this.element,c=b.box,f=a&&a.inverted;c=a?a.element||a:c;a&&(this.parentGroup=a);f&&b.invertChild(d,c);c.appendChild(d);this.added=!0;this.alignOnAdd&&!this.deferUpdateTransform&&this.updateTransform();if(this.onAdd)this.onAdd();this.className&&this.attr("class",this.className);return this},updateTransform:y.prototype.htmlUpdateTransform,
setSpanRotation:function(){var a=this.rotation,b=Math.cos(a*u),d=Math.sin(a*u);z(this.element,{filter:a?["progid:DXImageTransform.Microsoft.Matrix(M11=",b,", M12=",-d,", M21=",d,", M22=",b,", sizingMethod='auto expand')"].join(""):"none"})},getSpanCorrection:function(a,b,d,c,f){var q=c?Math.cos(c*u):1,e=c?Math.sin(c*u):0,J=B(this.elemHeight,this.element.offsetHeight);this.xCorr=0>q&&-a;this.yCorr=0>e&&-J;var A=0>q*e;this.xCorr+=e*b*(A?1-d:d);this.yCorr-=q*b*(c?A?d:1-d:1);f&&"left"!==f&&(this.xCorr-=
a*d*(0>q?-1:1),c&&(this.yCorr-=J*d*(0>e?-1:1)),z(this.element,{textAlign:f}))},pathToVML:function(a){for(var b=a.length,d=[];b--;)O(a[b])?d[b]=Math.round(10*a[b])-5:"Z"===a[b]?d[b]="x":(d[b]=a[b],!a.isArc||"wa"!==a[b]&&"at"!==a[b]||(d[b+5]===d[b+7]&&(d[b+7]+=a[b+7]>a[b+5]?1:-1),d[b+6]===d[b+8]&&(d[b+8]+=a[b+8]>a[b+6]?1:-1)));return d.join(" ")||"x"},clip:function(a){var b=this;if(a){var d=a.members;N(d,b);d.push(b);b.destroyClip=function(){N(d,b)};a=a.getCSS(b)}else b.destroyClip&&b.destroyClip(),
a={clip:b.docMode8?"inherit":"rect(auto)"};return b.css(a)},css:y.prototype.htmlCss,safeRemoveChild:function(a){a.parentNode&&I(a)},destroy:function(){this.destroyClip&&this.destroyClip();return y.prototype.destroy.apply(this)},on:function(a,b){this.element["on"+a]=function(){var a=v.event;a.target=a.srcElement;b(a)};return this},cutOffPath:function(a,b){a=a.split(/[ ,]/);var d=a.length;if(9===d||11===d)a[d-4]=a[d-2]=t(a[d-2])-10*b;return a.join(" ")},shadow:function(a,b,d){var c=[],f,e=this.element,
h=this.renderer,J=e.style,A=e.path;A&&"string"!==typeof A.value&&(A="x");var k=A;if(a){var P=B(a.width,3);var n=(a.opacity||.15)/P;for(f=1;3>=f;f++){var m=2*P+1-2*f;d&&(k=this.cutOffPath(A.value,m+.5));var l=['<shape isShadow="true" strokeweight="',m,'" filled="false" path="',k,'" coordsize="10 10" style="',e.style.cssText,'" />'];var g=F(h.prepVML(l),null,{left:t(J.left)+B(a.offsetX,1)+"px",top:t(J.top)+B(a.offsetY,1)+"px"});d&&(g.cutOff=m+1);l=['<stroke color="',a.color||"#000000",'" opacity="',
n*f,'"/>'];F(h.prepVML(l),null,null,g);b?b.element.appendChild(g):e.parentNode.insertBefore(g,e);c.push(g)}this.shadows=c}return this},updateShadows:r,setAttr:function(a,b){this.docMode8?this.element[a]=b:this.element.setAttribute(a,b)},getAttr:function(a){return this.docMode8?this.element[a]:this.element.getAttribute(a)},classSetter:function(a){(this.added?this.element:this).className=a},dashstyleSetter:function(a,b,d){(d.getElementsByTagName("stroke")[0]||F(this.renderer.prepVML(["<stroke/>"]),
null,null,d))[b]=a||"solid";this[b]=a},dSetter:function(a,b,d){var c=this.shadows;a=a||[];this.d=a.join&&a.join(" ");d.path=a=this.pathToVML(a);if(c)for(d=c.length;d--;)c[d].path=c[d].cutOff?this.cutOffPath(a,c[d].cutOff):a;this.setAttr(b,a)},fillSetter:function(a,b,d){var c=d.nodeName;"SPAN"===c?d.style.color=a:"IMG"!==c&&(d.filled="none"!==a,this.setAttr("fillcolor",this.renderer.color(a,d,b,this)))},"fill-opacitySetter":function(a,b,d){F(this.renderer.prepVML(["<",b.split("-")[0],' opacity="',
a,'"/>']),null,null,d)},opacitySetter:r,rotationSetter:function(a,b,d){d=d.style;this[b]=d[b]=a;d.left=-Math.round(Math.sin(a*u)+1)+"px";d.top=Math.round(Math.cos(a*u))+"px"},strokeSetter:function(a,b,d){this.setAttr("strokecolor",this.renderer.color(a,d,b,this))},"stroke-widthSetter":function(a,b,d){d.stroked=!!a;this[b]=a;O(a)&&(a+="px");this.setAttr("strokeweight",a)},titleSetter:function(a,b){this.setAttr(b,a)},visibilitySetter:function(a,b,d){"inherit"===a&&(a="visible");this.shadows&&this.shadows.forEach(function(d){d.style[b]=
a});"DIV"===d.nodeName&&(a="hidden"===a?"-999em":0,this.docMode8||(d.style[b]=a?"visible":"hidden"),b="top");d.style[b]=a},xSetter:function(a,b,d){this[b]=a;"x"===b?b="left":"y"===b&&(b="top");this.updateClipping?(this[b]=a,this.updateClipping()):d.style[b]=a},zIndexSetter:function(a,b,d){d.style[b]=a},fillGetter:function(){return this.getAttr("fillcolor")||""},strokeGetter:function(){return this.getAttr("strokecolor")||""},classGetter:function(){return this.getAttr("className")||""}},c["stroke-opacitySetter"]=
c["fill-opacitySetter"],e.VMLElement=c=p(y,c),c.prototype.ySetter=c.prototype.widthSetter=c.prototype.heightSetter=c.prototype.xSetter,c={Element:c,isIE8:-1<v.navigator.userAgent.indexOf("MSIE 8.0"),init:function(a,b,d){this.crispPolyLine=h.prototype.crispPolyLine;this.alignedObjects=[];var c=this.createElement("div").css({position:"relative"});var f=c.element;a.appendChild(c.element);this.isVML=!0;this.box=f;this.boxWrapper=c;this.gradients={};this.cache={};this.cacheKeys=[];this.imgCount=0;this.setSize(b,
d,!1);if(!k.namespaces.hcv){k.namespaces.add("hcv","urn:schemas-microsoft-com:vml");try{k.createStyleSheet().cssText="hcv\\:fill, hcv\\:path, hcv\\:shape, hcv\\:stroke{ behavior:url(#default#VML); display: inline-block; } "}catch(T){k.styleSheets[0].cssText+="hcv\\:fill, hcv\\:path, hcv\\:shape, hcv\\:stroke{ behavior:url(#default#VML); display: inline-block; } "}}},isHidden:function(){return!this.box.offsetWidth},clipRect:function(a,b,d,c){var f=this.createElement(),e=G(a);return w(f,{members:[],
count:0,left:(e?a.x:a)+1,top:(e?a.y:b)+1,width:(e?a.width:d)-1,height:(e?a.height:c)-1,getCSS:function(a){var b=a.element,d=b.nodeName,c=a.inverted,f=this.top-("shape"===d?b.offsetTop:0),e=this.left;b=e+this.width;var q=f+this.height;f={clip:"rect("+Math.round(c?e:f)+"px,"+Math.round(c?q:b)+"px,"+Math.round(c?b:q)+"px,"+Math.round(c?f:e)+"px)"};!c&&a.docMode8&&"DIV"===d&&w(f,{width:b+"px",height:q+"px"});return f},updateClipping:function(){f.members.forEach(function(a){a.element&&a.css(f.getCSS(a))})}})},
color:function(a,b,d,c){var f=this,e=/^rgba/,q,h,k="none";a&&a.linearGradient?h="gradient":a&&a.radialGradient&&(h="pattern");if(h){var g,m,n=a.linearGradient||a.radialGradient,l=void 0,p=void 0,r=void 0,t=void 0,u,v,w,y,z="";a=a.stops;p=l=void 0;var B=[],D=function(){q=['<fill colors="'+B.join(",")+'" opacity="',v,'" o:opacity2="',u,'" type="',h,'" ',z,'focus="100%" method="any" />'];F(f.prepVML(q),null,null,b)};l=a[0];p=a[a.length-1];0<l[0]&&a.unshift([0,l[1]]);1>p[0]&&a.push([1,p[1]]);a.forEach(function(a,
b){e.test(a[1])?(K=x(a[1]),g=K.get("rgb"),m=K.get("a")):(g=a[1],m=1);B.push(100*a[0]+"% "+g);b?(v=m,w=g):(u=m,y=g)});if("fill"===d)if("gradient"===h)l=n.x1||n[0]||0,p=n.y1||n[1]||0,r=n.x2||n[2]||0,t=n.y2||n[3]||0,z='angle="'+(90-180*Math.atan((t-p)/(r-l))/Math.PI)+'"',D();else{d=n.r;var E=2*d,G=2*d,H=n.cx,I=n.cy,L=b.radialReference,C;n=function(){L&&(C=c.getBBox(),H+=(L[0]-C.x)/C.width-.5,I+=(L[1]-C.y)/C.height-.5,E*=L[2]/C.width,G*=L[2]/C.height);z='src="'+M().global.VMLRadialGradientURL+'" size="'+
E+","+G+'" origin="0.5,0.5" position="'+H+","+I+'" color2="'+y+'" ';D()};c.added?n():c.onAdd=n;k=w}else k=g}else if(e.test(a)&&"IMG"!==b.tagName){var K=x(a);c[d+"-opacitySetter"](K.get("a"),d,b);k=K.get("rgb")}else n=b.getElementsByTagName(d),n.length&&(n[0].opacity=1,n[0].type="solid"),k=a;return k},prepVML:function(a){var b=this.isIE8;a=a.join("");b?(a=a.replace("/>",' xmlns="urn:schemas-microsoft-com:vml" />'),a=-1===a.indexOf('style="')?a.replace("/>",' style="display:inline-block;behavior:url(#default#VML);" />'):
a.replace('style="','style="display:inline-block;behavior:url(#default#VML);')):a=a.replace("<","<hcv:");return a},text:h.prototype.html,path:function(a){var b={coordsize:"10 10"};R(a)?b.d=a:G(a)&&w(b,a);return this.createElement("shape").attr(b)},circle:function(a,b,d){var c=this.symbol("circle");G(a)&&(d=a.r,b=a.y,a=a.x);c.isCircle=!0;c.r=d;return c.attr({x:a,y:b})},g:function(a){var b;a&&(b={className:"highcharts-"+a,"class":"highcharts-"+a});return this.createElement("div").attr(b)},image:function(a,
b,d,c,f){var e=this.createElement("img").attr({src:a});1<arguments.length&&e.attr({x:b,y:d,width:c,height:f});return e},createElement:function(a){return"rect"===a?this.symbol(a):h.prototype.createElement.call(this,a)},invertChild:function(a,b){var d=this;b=b.style;var c="IMG"===a.tagName&&a.style;z(a,{flip:"x",left:t(b.width)-(c?t(c.top):1)+"px",top:t(b.height)-(c?t(c.left):1)+"px",rotation:-90});[].forEach.call(a.childNodes,function(b){d.invertChild(b,a)})},symbols:{arc:function(a,b,c,e,f){var d=
f.start,h=f.end,g=f.r||c||e;c=f.innerR;e=Math.cos(d);var k=Math.sin(d),l=Math.cos(h),m=Math.sin(h);if(0===h-d)return["x"];d=["wa",a-g,b-g,a+g,b+g,a+g*e,b+g*k,a+g*l,b+g*m];f.open&&!c&&d.push("e","M",a,b);d.push("at",a-c,b-c,a+c,b+c,a+c*l,b+c*m,a+c*e,b+c*k,"x","e");d.isArc=!0;return d},circle:function(a,b,c,e,f){f&&H(f.r)&&(c=e=2*f.r);f&&f.isCircle&&(a-=c/2,b-=e/2);return["wa",a,b,a+c,b+e,a+c,b+e/2,a+c,b+e/2,"e"]},rect:function(a,b,c,e,f){return h.prototype.symbols[H(f)&&f.r?"callout":"square"].call(0,
a,b,c,e,f)}}},e.VMLRenderer=e=function(){this.init.apply(this,arguments)},w(e.prototype,h.prototype),w(e.prototype,c),g.registerRendererType("VMLRenderer",e,!0),Q.compose(e,h));h.prototype.getSpanWidth=function(a,b){var c=a.getBBox(!0).width;!E&&this.forExport&&(c=this.measureSpanWidth(b.firstChild.data,a.styles));return c};h.prototype.measureSpanWidth=function(a,b){var c=k.createElement("span");a=k.createTextNode(a);c.appendChild(a);z(c,b);this.box.appendChild(c);b=c.offsetWidth;I(c);return b}});
p(c,"masters/modules/oldie.src.js",[],function(){})});
//# sourceMappingURL=oldie.js.map
```
