## Supplementary material for "Exploring natural odour landscapes: A case study with implications for human-biting insects": Figure S4: organization.js

```
/*
 Highcharts JS v9.3.1 (2021-11-05)
 Organization chart series type

 (c) 2019-2021 Torstein Honsi

 License: www.highcharts.com/license
*/
'use strict';(function(b){"object"===typeof module&&module.exports?(b["default"]=b,module.exports=b):"function"===typeof define&&define.amd?define("highcharts/modules/organization",["highcharts","highcharts/modules/sankey"],function(h){b(h);b.Highcharts=h;return b}):b("undefined"!==typeof Highcharts?Highcharts:void 0)})(function(b){function h(b,l,a,t){b.hasOwnProperty(l)||(b[l]=t.apply(null,a))}b=b?b._modules:{};h(b,"Series/Organization/OrganizationPoint.js",[b["Core/Series/SeriesRegistry.js"]],function(b){var l=
this&&this.__extends||function(){var b=function(a,d){b=Object.setPrototypeOf||{__proto__:[]}instanceof Array&&function(b,d){b.__proto__=d}||function(b,d){for(var a in d)d.hasOwnProperty(a)&&(b[a]=d[a])};return b(a,d)};return function(a,d){function l(){this.constructor=a}b(a,d);a.prototype=null===d?Object.create(d):(l.prototype=d.prototype,new l)}}();return function(b){function a(){var a=null!==b&&b.apply(this,arguments)||this;a.fromNode=void 0;a.linksFrom=void 0;a.linksTo=void 0;a.options=void 0;
a.series=void 0;a.toNode=void 0;return a}l(a,b);a.prototype.getSum=function(){return 1};return a}(b.seriesTypes.sankey.prototype.pointClass)});h(b,"Series/Organization/OrganizationSeries.js",[b["Series/Organization/OrganizationPoint.js"],b["Core/Series/SeriesRegistry.js"],b["Core/Utilities.js"]],function(b,l,a){var h=this&&this.__extends||function(){var b=function(a,c){b=Object.setPrototypeOf||{__proto__:[]}instanceof Array&&function(c,b){c.__proto__=b}||function(c,b){for(var e in b)b.hasOwnProperty(e)&&
(c[e]=b[e])};return b(a,c)};return function(a,c){function e(){this.constructor=a}b(a,c);a.prototype=null===c?Object.create(c):(e.prototype=c.prototype,new e)}}(),d=l.seriesTypes.sankey,r=a.css,u=a.extend,v=a.merge,x=a.pick,y=a.wrap;a=function(b){function a(){var c=null!==b&&b.apply(this,arguments)||this;c.data=void 0;c.options=void 0;c.points=void 0;return c}h(a,b);a.curvedPath=function(c,b){for(var a=[],e=0;e<c.length;e++){var f=c[e][1],g=c[e][2];if("number"===typeof f&&"number"===typeof g)if(0===
e)a.push(["M",f,g]);else if(e===c.length-1)a.push(["L",f,g]);else if(b){var k=c[e-1],m=c[e+1];if(k&&m){var d=k[1];k=k[2];var n=m[1];m=m[2];if("number"===typeof d&&"number"===typeof n&&"number"===typeof k&&"number"===typeof m&&d!==n&&k!==m){var l=d<n?1:-1,h=k<m?1:-1;a.push(["L",f-l*Math.min(Math.abs(f-d),b),g-h*Math.min(Math.abs(g-k),b)],["C",f,g,f,g,f+l*Math.min(Math.abs(f-n),b),g+h*Math.min(Math.abs(g-m),b)])}}}else a.push(["L",f,g])}return a};a.prototype.alignDataLabel=function(c,a,w){if(w.useHTML){var e=
c.shapeArgs.width,f=c.shapeArgs.height,g=this.options.borderWidth+2*this.options.dataLabels.padding;this.chart.inverted&&(e=f,f=c.shapeArgs.width);f-=g;e-=g;if(g=a.text)r(g.element.parentNode,{width:e+"px",height:f+"px"}),r(g.element,{left:0,top:0,width:"100%",height:"100%",overflow:"hidden"});a.getBBox=function(){return{width:e,height:f}};a.width=e;a.height=f}b.prototype.alignDataLabel.apply(this,arguments)};a.prototype.createNode=function(c){c=b.prototype.createNode.call(this,c);c.getSum=function(){return 1};
return c};a.prototype.createNodeColumn=function(){var c=b.prototype.createNodeColumn.call(this);y(c,"offset",function(c,a,b){c=c.call(this,a,b);return a.hangsFrom?{absoluteTop:a.hangsFrom.nodeY}:c});return c};a.prototype.pointAttribs=function(c,a){var b=this,e=d.prototype.pointAttribs.call(b,c,a),f=b.mapOptionsToLevel[(c.isNode?c.level:c.fromNode.level)||0]||{},g=c.options,k=f.states&&f.states[a]||{};a=["borderRadius","linkColor","linkLineWidth"].reduce(function(a,c){a[c]=x(k[c],g[c],f[c],b.options[c]);
return a},{});c.isNode?a.borderRadius&&(e.r=a.borderRadius):(e.stroke=a.linkColor,e["stroke-width"]=a.linkLineWidth,delete e.fill);return e};a.prototype.translateLink=function(c){var b=c.fromNode,d=c.toNode,p=Math.round(this.options.linkLineWidth)%2/2,f=Math.floor(b.shapeArgs.x+b.shapeArgs.width)+p,g=Math.floor(b.shapeArgs.y+b.shapeArgs.height/2)+p,k=Math.floor(d.shapeArgs.x)+p,m=Math.floor(d.shapeArgs.y+d.shapeArgs.height/2)+p,l=this.options.hangingIndent;var n=d.options.offset;var h=/%$/.test(n)&&
parseInt(n,10),q=this.chart.inverted;q&&(f-=b.shapeArgs.width,k+=d.shapeArgs.width);n=Math.floor(k+(q?1:-1)*(this.colDistance-this.nodeWidth)/2)+p;h&&(50<=h||-50>=h)&&(n=k=Math.floor(k+(q?-.5:.5)*d.shapeArgs.width)+p,m=d.shapeArgs.y,0<h&&(m+=d.shapeArgs.height));d.hangsFrom===b&&(this.chart.inverted?(g=Math.floor(b.shapeArgs.y+b.shapeArgs.height-l/2)+p,m=d.shapeArgs.y+d.shapeArgs.height):g=Math.floor(b.shapeArgs.y+l/2)+p,n=k=Math.floor(d.shapeArgs.x+d.shapeArgs.width/2)+p);c.plotY=1;c.shapeType="path";
c.shapeArgs={d:a.curvedPath([["M",f,g],["L",n,g],["L",n,m],["L",k,m]],this.options.linkRadius)}};a.prototype.translateNode=function(a,b){d.prototype.translateNode.call(this,a,b);a.hangsFrom&&(a.shapeArgs.height-=this.options.hangingIndent,this.chart.inverted||(a.shapeArgs.y+=this.options.hangingIndent));a.nodeHeight=this.chart.inverted?a.shapeArgs.width:a.shapeArgs.height};a.defaultOptions=v(d.defaultOptions,{borderColor:"#666666",borderRadius:3,linkRadius:10,borderWidth:1,dataLabels:{nodeFormatter:function(){function a(a){return Object.keys(a).reduce(function(b,
c){return b+c+":"+a[c]+";"},'style="')+'"'}var b={width:"100%",height:"100%",display:"flex","flex-direction":"row","align-items":"center","justify-content":"center"},d={"max-height":"100%","border-radius":"50%"},h={width:"100%",padding:0,"text-align":"center","white-space":"normal"},f={margin:0},g={margin:0},k={opacity:.75,margin:"5px"};this.point.image&&(d["max-width"]="30%",h.width="70%");this.series.chart.renderer.forExport&&(b.display="block",h.position="absolute",h.left=this.point.image?"30%":
0,h.top=0);b="<div "+a(b)+">";this.point.image&&(b+='<img src="'+this.point.image+'" '+a(d)+">");b+="<div "+a(h)+">";this.point.name&&(b+="<h4 "+a(f)+">"+this.point.name+"</h4>");this.point.title&&(b+="<p "+a(g)+">"+(this.point.title||"")+"</p>");this.point.description&&(b+="<p "+a(k)+">"+this.point.description+"</p>");return b+"</div></div>"},style:{fontWeight:"normal",fontSize:"13px"},useHTML:!0},hangingIndent:20,linkColor:"#666666",linkLineWidth:1,nodeWidth:50,tooltip:{nodeFormat:"{point.name}<br>{point.title}<br>{point.description}"}});
return a}(d);u(a.prototype,{pointClass:b});l.registerSeriesType("organization",a);"";"";return a});h(b,"masters/modules/organization.src.js",[],function(){})});
//# sourceMappingURL=organization.js.map
```
