## Supplementary material for "Exploring natural odour landscapes: A case study with implications for human-biting insects": Figure S4: overlapping-datalabels.js

```
/*
 Highcharts JS v9.3.1 (2021-11-05)

 (c) 2009-2021 Torstein Honsi

 License: www.highcharts.com/license
*/
'use strict';(function(a){"object"===typeof module&&module.exports?(a["default"]=a,module.exports=a):"function"===typeof define&&define.amd?define("highcharts/modules/overlapping-datalabels",["highcharts"],function(b){a(b);a.Highcharts=b;return a}):a("undefined"!==typeof Highcharts?Highcharts:void 0)})(function(a){(function(a,c,d,e){a.hasOwnProperty(c)||(a[c]=e.apply(null,d))})(a?a._modules:{},"masters/modules/overlapping-datalabels.src.js",[],function(){})});
//# sourceMappingURL=overlapping-datalabels.js.map
```
