## Supplementary material for "Exploring natural odour landscapes: A case study with implications for human-biting insects": Figure S4: parallel-coordinates.js

```
/*
 Highcharts JS v9.3.1 (2021-11-05)

 Support for parallel coordinates in Highcharts

 (c) 2010-2021 Pawel Fus

 License: www.highcharts.com/license
*/
'use strict';(function(b){"object"===typeof module&&module.exports?(b["default"]=b,module.exports=b):"function"===typeof define&&define.amd?define("highcharts/modules/parallel-coordinates",["highcharts"],function(g){b(g);b.Highcharts=g;return b}):b("undefined"!==typeof Highcharts?Highcharts:void 0)})(function(b){function g(b,m,g,r){b.hasOwnProperty(m)||(b[m]=r.apply(null,g))}b=b?b._modules:{};g(b,"Extensions/ParallelCoordinates.js",[b["Core/Axis/Axis.js"],b["Core/Chart/Chart.js"],b["Core/FormatUtilities.js"],
b["Core/Globals.js"],b["Core/DefaultOptions.js"],b["Core/Series/Series.js"],b["Core/Utilities.js"]],function(b,g,q,r,y,t,d){function m(a){var c=this.series&&this.series.chart,b=a.apply(this,Array.prototype.slice.call(arguments,1)),h;if(c&&c.hasParallelCoordinates&&!n(b.formattedValue)){var e=c.yAxis[this.x];var f=e.options;c=(h=u(f.tooltipValueFormat,f.labels.format))?z(h,v(this,{value:this.y}),c):e.dateTime?c.time.dateFormat(c.time.resolveDTLFormat(f.dateTimeLabelFormats[e.tickPositions.info.unitName]).main,
this.y):f.categories?f.categories[this.y]:this.y;b.formattedValue=b.point.formattedValue=c}return b}var z=q.format;q=y.setOptions;var k=d.addEvent,A=d.arrayMax,B=d.arrayMin,n=d.defined,C=d.erase,v=d.extend,l=d.merge,u=d.pick,w=d.splat,D=d.wrap;d=g.prototype;var x={lineWidth:0,tickLength:0,opposite:!0,type:"category"};q({chart:{parallelCoordinates:!1,parallelAxes:{lineWidth:1,title:{text:"",reserveSpace:!1},labels:{x:0,y:4,align:"center",reserveSpace:!1},offset:0}}});k(g,"init",function(a){a=a.args[0];
var c=w(a.yAxis||{}),b=[],h=c.length;if(this.hasParallelCoordinates=a.chart&&a.chart.parallelCoordinates){for(this.setParallelInfo(a);h<=this.parallelInfo.counter;h++)b.push({});a.legend||(a.legend={});"undefined"===typeof a.legend.enabled&&(a.legend.enabled=!1);l(!0,a,{boost:{seriesThreshold:Number.MAX_VALUE},plotOptions:{series:{boostThreshold:Number.MAX_VALUE}}});a.yAxis=c.concat(b);a.xAxis=l(x,w(a.xAxis||{})[0])}});k(g,"update",function(a){a=a.options;a.chart&&(n(a.chart.parallelCoordinates)&&
(this.hasParallelCoordinates=a.chart.parallelCoordinates),this.options.chart.parallelAxes=l(this.options.chart.parallelAxes,a.chart.parallelAxes));this.hasParallelCoordinates&&(a.series&&this.setParallelInfo(a),this.yAxis.forEach(function(a){a.update({},!1)}))});v(d,{setParallelInfo:function(a){var c=this;a=a.series;c.parallelInfo={counter:0};a.forEach(function(a){a.data&&(c.parallelInfo.counter=Math.max(c.parallelInfo.counter,a.data.length-1))})}});k(t,"bindAxes",function(a){if(this.chart.hasParallelCoordinates){var c=
this;this.chart.axes.forEach(function(a){c.insert(a.series);a.isDirty=!0});c.xAxis=this.chart.xAxis[0];c.yAxis=this.chart.yAxis[0];a.preventDefault()}});k(t,"afterTranslate",function(){var a=this.chart,c=this.points,b=c&&c.length,h=Number.MAX_VALUE,e;if(this.chart.hasParallelCoordinates){for(e=0;e<b;e++){var f=c[e];if(n(f.y)){f.plotX=a.polar?a.yAxis[e].angleRad||0:a.inverted?a.plotHeight-a.yAxis[e].top+a.plotTop:a.yAxis[e].left-a.plotLeft;f.clientX=f.plotX;f.plotY=a.yAxis[e].translate(f.y,!1,!0,null,
!0);"undefined"!==typeof d&&(h=Math.min(h,Math.abs(f.plotX-d)));var d=f.plotX;f.isInside=a.isInsidePlot(f.plotX,f.plotY,{inverted:a.inverted})}else f.isNull=!0}this.closestPointRangePx=h}},{order:1});k(t,"destroy",function(){this.chart.hasParallelCoordinates&&(this.chart.axes||[]).forEach(function(a){a&&a.series&&(C(a.series,this),a.isDirty=a.forceRedraw=!0)},this)});["line","spline"].forEach(function(a){D(r.seriesTypes[a].prototype.pointClass.prototype,"getLabelConfig",m)});var E=function(){function a(a){this.axis=
a}a.prototype.setPosition=function(a,b){var c=this.axis,e=c.chart,f=((this.position||0)+.5)/(e.parallelInfo.counter+1);e.polar?b.angle=360*f:(b[a[0]]=100*f+"%",c[a[1]]=b[a[1]]=0,c[a[2]]=b[a[2]]=null,c[a[3]]=b[a[3]]=null)};return a}(),p;(function(a){function b(a){var b=this.chart,c=this.parallelCoordinates,e=["left","width","height","top"];if(b.hasParallelCoordinates)if(b.inverted&&(e=e.reverse()),this.isXAxis)this.options=l(this.options,x,a.userOptions);else{var d=b.yAxis.indexOf(this);this.options=
l(this.options,this.chart.options.chart.parallelAxes,a.userOptions);c.position=u(c.position,0<=d?d:b.yAxis.length);c.setPosition(e,this.options)}}function d(a){var b=this.chart,c=this.parallelCoordinates;if(c&&b&&b.hasParallelCoordinates&&!this.isXAxis){var e=c.position,d=[];this.series.forEach(function(a){a.visible&&n(a.yData[e])&&d.push(a.yData[e])});this.dataMin=B(d);this.dataMax=A(d);a.preventDefault()}}function g(){this.parallelCoordinates||(this.parallelCoordinates=new E(this))}a.compose=function(a){a.keepProps.push("parallel");
k(a,"init",g);k(a,"afterSetOptions",b);k(a,"getSeriesExtremes",d)}})(p||(p={}));p.compose(b);return p});g(b,"masters/modules/parallel-coordinates.src.js",[],function(){})});
//# sourceMappingURL=parallel-coordinates.js.map
```
