## Supplementary material for "Exploring natural odour landscapes: A case study with implications for human-biting insects": Figure S4: pareto.js

```
/*
 Highcharts JS v9.3.1 (2021-11-05)

 Pareto series type for Highcharts

 (c) 2010-2021 Sebastian Bochan

 License: www.highcharts.com/license
*/
'use strict';(function(a){"object"===typeof module&&module.exports?(a["default"]=a,module.exports=a):"function"===typeof define&&define.amd?define("highcharts/modules/pareto",["highcharts"],function(b){a(b);a.Highcharts=b;return a}):a("undefined"!==typeof Highcharts?Highcharts:void 0)})(function(a){function b(a,e,d,b){a.hasOwnProperty(e)||(a[e]=b.apply(null,d))}a=a?a._modules:{};b(a,"Series/DerivedComposition.js",[a["Core/Globals.js"],a["Core/Series/Series.js"],a["Core/Utilities.js"]],function(a,
b,d){var e=a.noop,f=d.addEvent,l=d.defined,g;(function(a){function d(){b.prototype.init.apply(this,arguments);this.initialised=!1;this.baseSeries=null;this.eventRemovers=[];this.addEvents()}function m(){var a=this.chart,c=this.options.baseSeries;this.baseSeries=l(c)&&(a.series[c]||a.get(c))||null}function c(){var a=this;this.eventRemovers.push(f(this.chart,"afterLinkSeries",function(){a.setBaseSeries();a.baseSeries&&!a.initialised&&(a.setDerivedData(),a.addBaseSeriesEvents(),a.initialised=!0)}))}
function k(){var a=this;this.eventRemovers.push(f(this.baseSeries,"updatedData",function(){a.setDerivedData()}),f(this.baseSeries,"destroy",function(){a.baseSeries=null;a.initialised=!1}))}function n(){this.eventRemovers.forEach(function(a){a()});b.prototype.destroy.apply(this,arguments)}var p=[];a.hasDerivedData=!0;a.setDerivedData=e;a.compose=function(a){if(-1===p.indexOf(a)){p.push(a);var b=a.prototype;b.addBaseSeriesEvents=k;b.addEvents=c;b.destroy=n;b.init=d;b.setBaseSeries=m}return a};a.init=
d;a.setBaseSeries=m;a.addEvents=c;a.addBaseSeriesEvents=k;a.destroy=n})(g||(g={}));return g});b(a,"Series/ParetoSeries/ParetoSeries.js",[a["Series/DerivedComposition.js"],a["Core/Series/SeriesRegistry.js"],a["Core/Utilities.js"]],function(a,b,d){var e=this&&this.__extends||function(){var a=function(b,c){a=Object.setPrototypeOf||{__proto__:[]}instanceof Array&&function(a,c){a.__proto__=c}||function(a,c){for(var b in c)c.hasOwnProperty(b)&&(a[b]=c[b])};return a(b,c)};return function(b,c){function d(){this.constructor=
b}a(b,c);b.prototype=null===c?Object.create(c):(d.prototype=c.prototype,new d)}}(),f=b.seriesTypes.line,l=d.correctFloat,g=d.merge;d=d.extend;var h=function(a){function b(){var b=null!==a&&a.apply(this,arguments)||this;b.data=void 0;b.points=void 0;b.options=void 0;return b}e(b,a);b.prototype.sumPointsPercents=function(a,b,d,e){var c=0,f=0,g=[],k;a.forEach(function(a,h){null!==a&&(e?c+=a:(k=a/d*100,g.push([b[h],l(f+k)]),f+=k))});return e?c:g};b.prototype.setDerivedData=function(){var a=this.baseSeries.xData,
b=this.baseSeries.yData,d=this.sumPointsPercents(b,a,null,!0);this.setData(this.sumPointsPercents(b,a,d,!1),!1)};b.defaultOptions=g(f.defaultOptions,{zIndex:3});return b}(f);d(h.prototype,{hasDerivedData:a.hasDerivedData});a.compose(h);b.registerSeriesType("pareto",h);"";return h});b(a,"masters/modules/pareto.src.js",[],function(){})});
//# sourceMappingURL=pareto.js.map
```
