## Supplementary material for "Exploring natural odour landscapes: A case study with implications for human-biting insects": Figure S4: pathfinder.js

```
/*
 Highcharts Gantt JS v9.3.1 (2021-11-05)

 Pathfinder

 (c) 2016-2021 ystein Moseng

 License: www.highcharts.com/license
*/
'use strict';(function(a){"object"===typeof module&&module.exports?(a["default"]=a,module.exports=a):"function"===typeof define&&define.amd?define("highcharts/modules/pathfinder",["highcharts"],function(w){a(w);a.Highcharts=w;return a}):a("undefined"!==typeof Highcharts?Highcharts:void 0)})(function(a){function w(a,t,z,A){a.hasOwnProperty(t)||(a[t]=A.apply(null,z))}a=a?a._modules:{};w(a,"Extensions/ArrowSymbols.js",[a["Core/Renderer/SVG/SVGRenderer.js"]],function(a){function t(a,n,p,l){return[["M",
a,n+l/2],["L",a+p,n],["L",a,n+l/2],["L",a+p,n+l]]}function z(a,n,p,l){return[["M",a+p,n],["L",a,n+l/2],["L",a+p,n+l],["Z"]]}function A(a,n,p,l){return z(a,n,p/2,l)}a=a.prototype.symbols;a.arrow=t;a["arrow-filled"]=z;a["arrow-filled-half"]=A;a["arrow-half"]=function(a,n,p,l){return t(a,n,p/2,l)};a["triangle-left"]=z;a["triangle-left-half"]=A;return a});w(a,"Gantt/Connection.js",[a["Core/Globals.js"],a["Core/DefaultOptions.js"],a["Core/Series/Point.js"],a["Core/Utilities.js"]],function(a,t,z,A){function J(b){var c=
b.shapeArgs;return c?{xMin:c.x||0,xMax:(c.x||0)+(c.width||0),yMin:c.y||0,yMax:(c.y||0)+(c.height||0)}:(c=b.graphic&&b.graphic.getBBox())?{xMin:b.plotX-c.width/2,xMax:b.plotX+c.width/2,yMin:b.plotY-c.height/2,yMax:b.plotY+c.height/2}:null}"";var n=A.defined,p=A.error,l=A.extend,x=A.merge,w=A.objectEach,f=a.deg2rad,d=Math.max,h=Math.min;l(t.defaultOptions,{connectors:{type:"straight",lineWidth:1,marker:{enabled:!1,align:"center",verticalAlign:"middle",inside:!1,lineWidth:1},startMarker:{symbol:"diamond"},
endMarker:{symbol:"arrow-filled"}}});t=function(){function b(c,y,k){this.toPoint=this.pathfinder=this.graphics=this.fromPoint=this.chart=void 0;this.init(c,y,k)}b.prototype.init=function(c,y,k){this.fromPoint=c;this.toPoint=y;this.options=k;this.chart=c.series.chart;this.pathfinder=this.chart.pathfinder};b.prototype.renderPath=function(c,y,k){var b=this.chart,a=b.styledMode,f=b.pathfinder,e=!b.options.chart.forExport&&!1!==k,g=this.graphics&&this.graphics.path;f.group||(f.group=b.renderer.g().addClass("highcharts-pathfinder-group").attr({zIndex:-1}).add(b.seriesGroup));
f.group.translate(b.plotLeft,b.plotTop);g&&g.renderer||(g=b.renderer.path().add(f.group),a||g.attr({opacity:0}));g.attr(y);c={d:c};a||(c.opacity=1);g[e?"animate":"attr"](c,k);this.graphics=this.graphics||{};this.graphics.path=g};b.prototype.addMarker=function(c,b,k){var a=this.fromPoint.series.chart,d=a.pathfinder;a=a.renderer;var y="start"===c?this.fromPoint:this.toPoint,e=y.getPathfinderAnchorPoint(b);if(b.enabled&&((k="start"===c?k[1]:k[k.length-2])&&"M"===k[0]||"L"===k[0])){k={x:k[1],y:k[2]};
k=y.getRadiansToVector(k,e);e=y.getMarkerVector(k,b.radius,e);k=-k/f;if(b.width&&b.height){var g=b.width;var u=b.height}else g=u=2*b.radius;this.graphics=this.graphics||{};e={x:e.x-g/2,y:e.y-u/2,width:g,height:u,rotation:k,rotationOriginX:e.x,rotationOriginY:e.y};this.graphics[c]?this.graphics[c].animate(e):(this.graphics[c]=a.symbol(b.symbol).addClass("highcharts-point-connecting-path-"+c+"-marker").attr(e).add(d.group),a.styledMode||this.graphics[c].attr({fill:b.color||this.fromPoint.color,stroke:b.lineColor,
"stroke-width":b.lineWidth,opacity:0}).animate({opacity:1},y.series.options.animation))}};b.prototype.getPath=function(b){var c=this.pathfinder,a=this.chart,f=c.algorithms[b.type],d=c.chartObstacles;if("function"!==typeof f)return p('"'+b.type+'" is not a Pathfinder algorithm.'),{path:[],obstacles:[]};f.requiresObstacles&&!d&&(d=c.chartObstacles=c.getChartObstacles(b),a.options.connectors.algorithmMargin=b.algorithmMargin,c.chartObstacleMetrics=c.getObstacleMetrics(d));return f(this.fromPoint.getPathfinderAnchorPoint(b.startMarker),
this.toPoint.getPathfinderAnchorPoint(b.endMarker),x({chartObstacles:d,lineObstacles:c.lineObstacles||[],obstacleMetrics:c.chartObstacleMetrics,hardBounds:{xMin:0,xMax:a.plotWidth,yMin:0,yMax:a.plotHeight},obstacleOptions:{margin:b.algorithmMargin},startDirectionX:c.getAlgorithmStartDirection(b.startMarker)},b))};b.prototype.render=function(){var b=this.fromPoint,a=b.series,f=a.chart,D=f.pathfinder,q=x(f.options.connectors,a.options.connectors,b.options.connectors,this.options),l={};f.styledMode||
(l.stroke=q.lineColor||b.color,l["stroke-width"]=q.lineWidth,q.dashStyle&&(l.dashstyle=q.dashStyle));l["class"]="highcharts-point-connecting-path highcharts-color-"+b.colorIndex;q=x(l,q);n(q.marker.radius)||(q.marker.radius=h(d(Math.ceil((q.algorithmMargin||8)/2)-1,1),5));b=this.getPath(q);f=b.path;b.obstacles&&(D.lineObstacles=D.lineObstacles||[],D.lineObstacles=D.lineObstacles.concat(b.obstacles));this.renderPath(f,l,a.options.animation);this.addMarker("start",x(q.marker,q.startMarker),f);this.addMarker("end",
x(q.marker,q.endMarker),f)};b.prototype.destroy=function(){this.graphics&&(w(this.graphics,function(b){b.destroy()}),delete this.graphics)};return b}();a.Connection=t;l(z.prototype,{getPathfinderAnchorPoint:function(b){var a=J(this);switch(b.align){case "right":var f="xMax";break;case "left":f="xMin"}switch(b.verticalAlign){case "top":var d="yMin";break;case "bottom":d="yMax"}return{x:f?a[f]:(a.xMin+a.xMax)/2,y:d?a[d]:(a.yMin+a.yMax)/2}},getRadiansToVector:function(b,a){var f;n(a)||(f=J(this))&&(a=
{x:(f.xMin+f.xMax)/2,y:(f.yMin+f.yMax)/2});return Math.atan2(a.y-b.y,b.x-a.x)},getMarkerVector:function(b,a,f){var d=2*Math.PI,c=J(this),h=c.xMax-c.xMin,l=c.yMax-c.yMin,e=Math.atan2(l,h),g=!1;h/=2;var u=l/2,H=c.xMin+h;c=c.yMin+u;for(var F=H,G=c,m=1,v=1;b<-Math.PI;)b+=d;for(;b>Math.PI;)b-=d;d=Math.tan(b);b>-e&&b<=e?(v=-1,g=!0):b>e&&b<=Math.PI-e?v=-1:b>Math.PI-e||b<=-(Math.PI-e)?(m=-1,g=!0):m=-1;g?(F+=m*h,G+=v*h*d):(F+=l/(2*d)*m,G+=v*u);f.x!==H&&(F=f.x);f.y!==c&&(G=f.y);return{x:F+a*Math.cos(b),y:G-
a*Math.sin(b)}}});return t});w(a,"Gantt/PathfinderAlgorithms.js",[a["Core/Utilities.js"]],function(a){function t(a,d,h){h=h||0;var b=a.length-1;d-=1e-7;for(var f,l;h<=b;)if(f=b+h>>1,l=d-a[f].xMin,0<l)h=f+1;else if(0>l)b=f-1;else return f;return 0<h?h-1:0}function z(a,d){for(var f=t(a,d.x+1)+1;f--;){var b;if(b=a[f].xMax>=d.x)b=a[f],b=d.x<=b.xMax&&d.x>=b.xMin&&d.y<=b.yMax&&d.y>=b.yMin;if(b)return f}return-1}function A(a){var f=[];if(a.length){f.push(["M",a[0].start.x,a[0].start.y]);for(var h=0;h<a.length;++h)f.push(["L",
a[h].end.x,a[h].end.y])}return f}function w(a,d){a.yMin=l(a.yMin,d.yMin);a.yMax=p(a.yMax,d.yMax);a.xMin=l(a.xMin,d.xMin);a.xMax=p(a.xMax,d.xMax)}var n=a.pick,p=Math.min,l=Math.max,x=Math.abs;a=function(a,d,h){function b(a,b,e,f,d){a={x:a.x,y:a.y};a[b]=e[f||b]+(d||0);return a}function f(a,e,f){var g=x(e[f]-a[f+"Min"])>x(e[f]-a[f+"Max"]);return b(e,f,a,f+(g?"Max":"Min"),g?1:-1)}var l=[],k=n(h.startDirectionX,x(d.x-a.x)>x(d.y-a.y))?"x":"y",p=h.chartObstacles,q=z(p,a);h=z(p,d);if(-1<h){var t=p[h];h=f(t,
d,k);t={start:h,end:d};var e=h}else e=d;-1<q&&(p=p[q],h=f(p,a,k),l.push({start:a,end:h}),h[k]>=a[k]===h[k]>=e[k]&&(k="y"===k?"x":"y",d=a[k]<d[k],l.push({start:h,end:b(h,k,p,k+(d?"Max":"Min"),d?1:-1)}),k="y"===k?"x":"y"));a=l.length?l[l.length-1].end:a;h=b(a,k,e);l.push({start:a,end:h});k=b(h,"y"===k?"x":"y",e);l.push({start:h,end:k});l.push(t);return{path:A(l),obstacles:l}};a.requiresObstacles=!0;var I=function(a,d,h){function b(a,b,g){var e,f=a.x<b.x?1:-1;if(a.x<b.x){var d=a;var u=b}else d=b,u=a;
if(a.y<b.y){var C=a;var r=b}else C=b,r=a;for(e=0>f?p(t(m,u.x),m.length-1):0;m[e]&&(0<f&&m[e].xMin<=u.x||0>f&&m[e].xMax>=d.x);){if(m[e].xMin<=u.x&&m[e].xMax>=d.x&&m[e].yMin<=r.y&&m[e].yMax>=C.y)return g?{y:a.y,x:a.x<b.x?m[e].xMin-1:m[e].xMax+1,obstacle:m[e]}:{x:a.x,y:a.y<b.y?m[e].yMin-1:m[e].yMax+1,obstacle:m[e]};e+=f}return b}function f(a,e,g,f,d){var u=d.soft,m=d.hard,c=f?"x":"y",h={x:e.x,y:e.y},C={x:e.x,y:e.y};d=a[c+"Max"]>=u[c+"Max"];u=a[c+"Min"]<=u[c+"Min"];var r=a[c+"Max"]>=m[c+"Max"];m=a[c+
"Min"]<=m[c+"Min"];var l=x(a[c+"Min"]-e[c]),k=x(a[c+"Max"]-e[c]);g=10>x(l-k)?e[c]<g[c]:k<l;C[c]=a[c+"Min"];h[c]=a[c+"Max"];a=b(e,C,f)[c]!==C[c];e=b(e,h,f)[c]!==h[c];g=a?e?g:!0:e?!1:g;g=u?d?g:!0:d?!1:g;return m?r?g:!0:r?!1:g}function y(a,g,d){if(a.x===g.x&&a.y===g.y)return[];var c=d?"x":"y",k=h.obstacleOptions.margin;var r={soft:{xMin:u,xMax:H,yMin:F,yMax:G},hard:h.hardBounds};var v=z(m,a);if(-1<v){v=m[v];r=f(v,a,g,d,r);w(v,h.hardBounds);var C=d?{y:a.y,x:v[r?"xMax":"xMin"]+(r?1:-1)}:{x:a.x,y:v[r?"yMax":
"yMin"]+(r?1:-1)};var B=z(m,C);-1<B&&(B=m[B],w(B,h.hardBounds),C[c]=r?l(v[c+"Max"]-k+1,(B[c+"Min"]+v[c+"Max"])/2):p(v[c+"Min"]+k-1,(B[c+"Max"]+v[c+"Min"])/2),a.x===C.x&&a.y===C.y?(e&&(C[c]=r?l(v[c+"Max"],B[c+"Max"])+1:p(v[c+"Min"],B[c+"Min"])-1),e=!e):e=!1);a=[{start:a,end:C}]}else c=b(a,{x:d?g.x:a.x,y:d?a.y:g.y},d),a=[{start:a,end:{x:c.x,y:c.y}}],c[d?"x":"y"]!==g[d?"x":"y"]&&(r=f(c.obstacle,c,g,!d,r),w(c.obstacle,h.hardBounds),r={x:d?c.x:c.obstacle[r?"xMax":"xMin"]+(r?1:-1),y:d?c.obstacle[r?"yMax":
"yMin"]+(r?1:-1):c.y},d=!d,a=a.concat(y({x:c.x,y:c.y},r,d)));return a=a.concat(y(a[a.length-1].end,g,!d))}function k(a,b,e){var g=p(a.xMax-b.x,b.x-a.xMin)<p(a.yMax-b.y,b.y-a.yMin);e=f(a,b,e,g,{soft:h.hardBounds,hard:h.hardBounds});return g?{y:b.y,x:a[e?"xMax":"xMin"]+(e?1:-1)}:{x:b.x,y:a[e?"yMax":"yMin"]+(e?1:-1)}}var D=n(h.startDirectionX,x(d.x-a.x)>x(d.y-a.y)),q=D?"x":"y",E=[],e=!1,g=h.obstacleMetrics,u=p(a.x,d.x)-g.maxWidth-10,H=l(a.x,d.x)+g.maxWidth+10,F=p(a.y,d.y)-g.maxHeight-10,G=l(a.y,d.y)+
g.maxHeight+10,m=h.chartObstacles;var v=t(m,u);g=t(m,H);m=m.slice(v,g+1);if(-1<(g=z(m,d))){var B=k(m[g],d,a);E.push({end:d,start:B});d=B}for(;-1<(g=z(m,d));)v=0>d[q]-a[q],B={x:d.x,y:d.y},B[q]=m[g][v?q+"Max":q+"Min"]+(v?1:-1),E.push({end:d,start:B}),d=B;a=y(a,d,D);a=a.concat(E.reverse());return{path:A(a),obstacles:a}};I.requiresObstacles=!0;return{fastAvoid:I,straight:function(a,d){return{path:[["M",a.x,a.y],["L",d.x,d.y]],obstacles:[{start:a,end:d}]}},simpleConnect:a}});w(a,"Gantt/Pathfinder.js",
[a["Gantt/Connection.js"],a["Core/Chart/Chart.js"],a["Core/Globals.js"],a["Core/DefaultOptions.js"],a["Core/Series/Point.js"],a["Core/Utilities.js"],a["Gantt/PathfinderAlgorithms.js"]],function(a,t,z,A,w,n,p){function l(a){var b=a.shapeArgs;return b?{xMin:b.x||0,xMax:(b.x||0)+(b.width||0),yMin:b.y||0,yMax:(b.y||0)+(b.height||0)}:(b=a.graphic&&a.graphic.getBBox())?{xMin:a.plotX-b.width/2,xMax:a.plotX+b.width/2,yMin:a.plotY-b.height/2,yMax:a.plotY+b.height/2}:null}function x(a){for(var b=a.length,e=
0,c,d,f=[],m=function(a,b,e){e=y(e,10);var c=a.yMax+e>b.yMin-e&&a.yMin-e<b.yMax+e,d=a.xMax+e>b.xMin-e&&a.xMin-e<b.xMax+e,g=c?a.xMin>b.xMax?a.xMin-b.xMax:b.xMin-a.xMax:Infinity,f=d?a.yMin>b.yMax?a.yMin-b.yMax:b.yMin-a.yMax:Infinity;return d&&c?e?m(a,b,Math.floor(e/2)):Infinity:q(g,f)};e<b;++e)for(c=e+1;c<b;++c)d=m(a[e],a[c]),80>d&&f.push(d);f.push(80);return D(Math.floor(f.sort(function(a,b){return a-b})[Math.floor(f.length/10)]/2-1),1)}function I(a){if(a.options.pathfinder||a.series.reduce(function(a,
b){b.options&&c(!0,b.options.connectors=b.options.connectors||{},b.options.pathfinder);return a||b.options&&b.options.pathfinder},!1))c(!0,a.options.connectors=a.options.connectors||{},a.options.pathfinder),h('WARNING: Pathfinder options have been renamed. Use "chart.connectors" or "series.connectors" instead.')}"";var f=n.addEvent,d=n.defined,h=n.error,b=n.extend,c=n.merge,y=n.pick,k=n.splat,D=Math.max,q=Math.min;b(A.defaultOptions,{connectors:{type:"straight",lineWidth:1,marker:{enabled:!1,align:"center",
verticalAlign:"middle",inside:!1,lineWidth:1},startMarker:{symbol:"diamond"},endMarker:{symbol:"arrow-filled"}}});var E=function(){function b(a){this.lineObstacles=this.group=this.connections=this.chartObstacleMetrics=this.chartObstacles=this.chart=void 0;this.init(a)}b.prototype.init=function(a){this.chart=a;this.connections=[];f(a,"redraw",function(){this.pathfinder.update()})};b.prototype.update=function(b){var e=this.chart,c=this,d=c.connections;c.connections=[];e.series.forEach(function(b){b.visible&&
!b.options.isInternal&&b.points.forEach(function(b){var d=b.options;d&&d.dependency&&(d.connect=d.dependency);var g;d=b.options&&b.options.connect&&k(b.options.connect);b.visible&&!1!==b.isInside&&d&&d.forEach(function(d){g=e.get("string"===typeof d?d:d.to);g instanceof w&&g.series.visible&&g.visible&&!1!==g.isInside&&c.connections.push(new a(b,g,"string"===typeof d?{}:d))})})});for(var g=0,f=void 0,h=void 0,l=d.length,p=c.connections.length;g<l;++g){h=!1;for(f=0;f<p;++f)if(d[g].fromPoint===c.connections[f].fromPoint&&
d[g].toPoint===c.connections[f].toPoint){c.connections[f].graphics=d[g].graphics;h=!0;break}h||d[g].destroy()}delete this.chartObstacles;delete this.lineObstacles;c.renderConnections(b)};b.prototype.renderConnections=function(a){a?this.chart.series.forEach(function(a){var b=function(){var b=a.chart.pathfinder;(b&&b.connections||[]).forEach(function(b){b.fromPoint&&b.fromPoint.series===a&&b.render()});a.pathfinderRemoveRenderEvent&&(a.pathfinderRemoveRenderEvent(),delete a.pathfinderRemoveRenderEvent)};
!1===a.options.animation?b():a.pathfinderRemoveRenderEvent=f(a,"afterAnimate",b)}):this.connections.forEach(function(a){a.render()})};b.prototype.getChartObstacles=function(a){for(var b=[],c=this.chart.series,e=y(a.algorithmMargin,0),g,f=0,h=c.length;f<h;++f)if(c[f].visible&&!c[f].options.isInternal){var k=0,p=c[f].points.length,n=void 0;for(n=void 0;k<p;++k)n=c[f].points[k],n.visible&&(n=l(n))&&b.push({xMin:n.xMin-e,xMax:n.xMax+e,yMin:n.yMin-e,yMax:n.yMax+e})}b=b.sort(function(a,b){return a.xMin-
b.xMin});d(a.algorithmMargin)||(g=a.algorithmMargin=x(b),b.forEach(function(a){a.xMin-=g;a.xMax+=g;a.yMin-=g;a.yMax+=g}));return b};b.prototype.getObstacleMetrics=function(a){for(var b=0,c=0,d,e,f=a.length;f--;)d=a[f].xMax-a[f].xMin,e=a[f].yMax-a[f].yMin,b<d&&(b=d),c<e&&(c=e);return{maxHeight:c,maxWidth:b}};b.prototype.getAlgorithmStartDirection=function(a){var b="top"!==a.verticalAlign&&"bottom"!==a.verticalAlign;return"left"!==a.align&&"right"!==a.align?b?void 0:!1:b?!0:void 0};return b}();E.prototype.algorithms=
p;z.Pathfinder=E;b(w.prototype,{getPathfinderAnchorPoint:function(a){var b=l(this);switch(a.align){case "right":var c="xMax";break;case "left":c="xMin"}switch(a.verticalAlign){case "top":var d="yMin";break;case "bottom":d="yMax"}return{x:c?b[c]:(b.xMin+b.xMax)/2,y:d?b[d]:(b.yMin+b.yMax)/2}},getRadiansToVector:function(a,b){var c;d(b)||(c=l(this))&&(b={x:(c.xMin+c.xMax)/2,y:(c.yMin+c.yMax)/2});return Math.atan2(b.y-a.y,a.x-b.x)},getMarkerVector:function(a,b,c){var d=2*Math.PI,e=l(this),f=e.xMax-e.xMin,
g=e.yMax-e.yMin,h=Math.atan2(g,f),k=!1;f/=2;var n=g/2,p=e.xMin+f;e=e.yMin+n;for(var r=p,q=e,t=1,u=1;a<-Math.PI;)a+=d;for(;a>Math.PI;)a-=d;d=Math.tan(a);a>-h&&a<=h?(u=-1,k=!0):a>h&&a<=Math.PI-h?u=-1:a>Math.PI-h||a<=-(Math.PI-h)?(t=-1,k=!0):t=-1;k?(r+=t*f,q+=u*f*d):(r+=g/(2*d)*t,q+=u*n);c.x!==p&&(r=c.x);c.y!==e&&(q=c.y);return{x:r+b*Math.cos(a),y:q-b*Math.sin(a)}}});t.prototype.callbacks.push(function(a){!1!==a.options.connectors.enabled&&(I(a),this.pathfinder=new E(this),this.pathfinder.update(!0))});
return E});w(a,"masters/modules/pathfinder.src.js",[],function(){})});
//# sourceMappingURL=pathfinder.js.map
```
