## Supplementary material for "Exploring natural odour landscapes: A case study with implications for human-biting insects": Figure S4: pattern-fill.js

```
/*
 Highcharts JS v9.3.1 (2021-11-05)

 Module for adding patterns and images as point fills.

 (c) 2010-2021 Highsoft AS
 Author: Torstein Hnsi, ystein Moseng

 License: www.highcharts.com/license
*/
'use strict';(function(b){"object"===typeof module&&module.exports?(b["default"]=b,module.exports=b):"function"===typeof define&&define.amd?define("highcharts/modules/pattern-fill",["highcharts"],function(f){b(f);b.Highcharts=f;return b}):b("undefined"!==typeof Highcharts?Highcharts:void 0)})(function(b){function f(b,f,r,p){b.hasOwnProperty(f)||(b[f]=p.apply(null,r))}b=b?b._modules:{};f(b,"Extensions/PatternFill.js",[b["Core/Animation/AnimationUtilities.js"],b["Core/Chart/Chart.js"],b["Core/Globals.js"],
b["Core/DefaultOptions.js"],b["Core/Series/Point.js"],b["Core/Series/Series.js"],b["Core/Renderer/SVG/SVGRenderer.js"],b["Core/Utilities.js"]],function(b,f,r,p,t,u,v,l){function w(a,c){a=JSON.stringify(a);var b=a.length||0,e=0,d=0;if(c){c=Math.max(Math.floor(b/500),1);for(var n=0;n<b;n+=c)e+=a.charCodeAt(n);e&=e}for(;d<b;++d)c=a.charCodeAt(d),e=(e<<5)-e+c,e&=e;return e.toString(16).replace("-","1")}var z=b.animObject,A=p.getOptions;b=l.addEvent;var B=l.erase,x=l.merge,q=l.pick,C=l.removeEvent;p=l.wrap;
var y=r.patterns=function(){var a=[],c=A().colors;["M 0 0 L 5 5 M 4.5 -0.5 L 5.5 0.5 M -0.5 4.5 L 0.5 5.5","M 0 5 L 5 0 M -0.5 0.5 L 0.5 -0.5 M 4.5 5.5 L 5.5 4.5","M 2 0 L 2 5 M 4 0 L 4 5","M 0 2 L 5 2 M 0 4 L 5 4","M 0 1.5 L 2.5 1.5 L 2.5 0 M 2.5 5 L 2.5 3.5 L 5 3.5"].forEach(function(b,e){a.push({path:b,color:c[e],width:5,height:5,patternTransform:"scale(1.4 1.4)"})});["M 0 0 L 5 10 L 10 0","M 3 3 L 8 3 L 8 8 L 3 8 Z","M 5 5 m -4 0 a 4 4 0 1 1 8 0 a 4 4 0 1 1 -8 0","M 0 0 L 10 10 M 9 -1 L 11 1 M -1 9 L 1 11",
"M 0 10 L 10 0 M -1 1 L 1 -1 M 9 11 L 11 9"].forEach(function(b,e){a.push({path:b,color:c[e+5],width:10,height:10})});return a}();t.prototype.calculatePatternDimensions=function(a){if(!a.width||!a.height){var c=this.graphic&&(this.graphic.getBBox&&this.graphic.getBBox(!0)||this.graphic.element&&this.graphic.element.getBBox())||{},b=this.shapeArgs;b&&(c.width=b.width||c.width,c.height=b.height||c.height,c.x=b.x||c.x,c.y=b.y||c.y);if(a.image){if(!c.width||!c.height){a._width="defer";a._height="defer";
return}a.aspectRatio&&(c.aspectRatio=c.width/c.height,a.aspectRatio>c.aspectRatio?c.aspectWidth=c.height*a.aspectRatio:c.aspectHeight=c.width/a.aspectRatio);a._width=a.width||Math.ceil(c.aspectWidth||c.width);a._height=a.height||Math.ceil(c.aspectHeight||c.height)}a.width||(a._x=a.x||0,a._x+=c.x-Math.round(c.aspectWidth?Math.abs(c.aspectWidth-c.width)/2:0));a.height||(a._y=a.y||0,a._y+=c.y-Math.round(c.aspectHeight?Math.abs(c.aspectHeight-c.height)/2:0))}};v.prototype.addPattern=function(a,c){c=q(c,
!0);var b=z(c),e=a.width||a._width||32,d=a.height||a._height||32,n=a.color||"#343434",g=a.id,f=this,m=function(a){f.rect(0,0,e,d).attr({fill:a}).add(k)};g||(this.idCounter=this.idCounter||0,g="highcharts-pattern-"+this.idCounter+"-"+(this.chartIndex||0),++this.idCounter);this.forExport&&(g+="-export");this.defIds=this.defIds||[];if(!(-1<this.defIds.indexOf(g))){this.defIds.push(g);var h={id:g,patternUnits:"userSpaceOnUse",patternContentUnits:a.patternContentUnits||"userSpaceOnUse",width:e,height:d,
x:a._x||a.x||0,y:a._y||a.y||0};a.patternTransform&&(h.patternTransform=a.patternTransform);var k=this.createElement("pattern").attr(h).add(this.defs);k.id=g;a.path?(h=l.isObject(a.path)?a.path:{d:a.path},a.backgroundColor&&m(a.backgroundColor),m={d:h.d},this.styledMode||(m.stroke=h.stroke||n,m["stroke-width"]=q(h.strokeWidth,2),m.fill=h.fill||"none"),h.transform&&(m.transform=h.transform),this.createElement("path").attr(m).add(k),k.color=n):a.image&&(c?this.image(a.image,0,0,e,d,function(){this.animate({opacity:q(a.opacity,
1)},b);C(this.element,"load")}).attr({opacity:0}).add(k):this.image(a.image,0,0,e,d).add(k));a.image&&c||"undefined"===typeof a.opacity||[].forEach.call(k.element.childNodes,function(c){c.setAttribute("opacity",a.opacity)});this.patternElements=this.patternElements||{};return this.patternElements[g]=k}};p(u.prototype,"getColor",function(a){var c=this.options.color;c&&c.pattern&&!c.pattern.color?(delete this.options.color,a.apply(this,Array.prototype.slice.call(arguments,1)),c.pattern.color=this.color,
this.color=this.options.color=c):a.apply(this,Array.prototype.slice.call(arguments,1))});b(u,"render",function(){var a=this.chart.isResizing;(this.isDirtyData||a||!this.chart.hasRendered)&&(this.points||[]).forEach(function(c){var b=c.options&&c.options.color;b&&b.pattern&&(!a||c.shapeArgs&&c.shapeArgs.width&&c.shapeArgs.height?c.calculatePatternDimensions(b.pattern):(b.pattern._width="defer",b.pattern._height="defer"))})});b(t,"afterInit",function(){var a=this.options.color;a&&a.pattern&&("string"===
typeof a.pattern.path&&(a.pattern.path={d:a.pattern.path}),this.color=this.options.color=x(this.series.options.color,a))});b(v,"complexColor",function(a){var c=a.args[0],b=a.args[1];a=a.args[2];var e=this.chartIndex||0,d=c.pattern,f="#343434";"undefined"!==typeof c.patternIndex&&y&&(d=y[c.patternIndex]);if(!d)return!0;if(d.image||"string"===typeof d.path||d.path&&d.path.d){var g=a.parentNode&&a.parentNode.getAttribute("class");g=g&&-1<g.indexOf("highcharts-legend");"defer"!==d._width&&"defer"!==d._height||
t.prototype.calculatePatternDimensions.call({graphic:{element:a}},d);if(g||!d.id)d=x({},d),d.id="highcharts-pattern-"+e+"-"+w(d)+w(d,!0);this.addPattern(d,!this.forExport&&q(d.animation,this.globalAnimation,{duration:100}));f="url("+this.url+"#"+(d.id+(this.forExport?"-export":""))+")"}else f=d.color||f;a.setAttribute(b,f);c.toString=function(){return f};return!1});b(f,"endResize",function(){(this.renderer&&this.renderer.defIds||[]).filter(function(a){return a&&a.indexOf&&0===a.indexOf("highcharts-pattern-")}).length&&
(this.series.forEach(function(a){a.points.forEach(function(a){(a=a.options&&a.options.color)&&a.pattern&&(a.pattern._width="defer",a.pattern._height="defer")})}),this.redraw(!1))});b(f,"redraw",function(){var a={},c=this.renderer,b=(c.defIds||[]).filter(function(a){return a.indexOf&&0===a.indexOf("highcharts-pattern-")});b.length&&([].forEach.call(this.renderTo.querySelectorAll('[color^="url("], [fill^="url("], [stroke^="url("]'),function(b){if(b=b.getAttribute("fill")||b.getAttribute("color")||b.getAttribute("stroke"))b=
b.replace(c.url,"").replace("url(#","").replace(")",""),a[b]=!0}),b.forEach(function(b){a[b]||(B(c.defIds,b),c.patternElements[b]&&(c.patternElements[b].destroy(),delete c.patternElements[b]))}))});""});f(b,"masters/modules/pattern-fill.src.js",[],function(){})});
//# sourceMappingURL=pattern-fill.js.map
```
