## Supplementary material for "Exploring natural odour landscapes: A case study with implications for human-biting insects": Figure S4: price-indicator.js

```
/*
 Highstock JS v9.3.1 (2021-11-05)

 Advanced Highcharts Stock tools

 (c) 2010-2021 Highsoft AS
 Author: Torstein Honsi

 License: www.highcharts.com/license
*/
'use strict';(function(a){"object"===typeof module&&module.exports?(a["default"]=a,module.exports=a):"function"===typeof define&&define.amd?define("highcharts/modules/price-indicator",["highcharts","highcharts/modules/stock"],function(c){a(c);a.Highcharts=c;return a}):a("undefined"!==typeof Highcharts?Highcharts:void 0)})(function(a){function c(a,d,c,f){a.hasOwnProperty(d)||(a[d]=f.apply(null,c))}a=a?a._modules:{};c(a,"Extensions/PriceIndication.js",[a["Core/Series/Series.js"],a["Core/Utilities.js"]],
function(a,c){var d=c.addEvent,f=c.isArray,m=c.merge;d(a,"afterRender",function(){var a=this.options,c=a.pointRange,d=a.lastVisiblePrice,e=a.lastPrice;if((d||e)&&"highcharts-navigator-series"!==a.id){var n=this.xAxis,b=this.yAxis,p=b.crosshair,q=b.cross,r=b.crossLabel,g=this.points,h=g.length,k=this.xData[this.xData.length-1],l=this.yData[this.yData.length-1];e&&e.enabled&&(b.crosshair=b.options.crosshair=a.lastPrice,!this.chart.styledMode&&b.crosshair&&b.options.crosshair&&a.lastPrice&&(b.crosshair.color=
b.options.crosshair.color=a.lastPrice.color||this.color),b.cross=this.lastPrice,e=f(l)?l[3]:l,b.drawCrosshair(null,{x:k,y:e,plotX:n.toPixels(k,!0),plotY:b.toPixels(e,!0)}),this.yAxis.cross&&(this.lastPrice=this.yAxis.cross,this.lastPrice.addClass("highcharts-color-"+this.colorIndex),this.lastPrice.y=e));d&&d.enabled&&0<h&&(c=g[h-1].x===k||null===c?1:2,b.crosshair=b.options.crosshair=m({color:"transparent"},a.lastVisiblePrice),b.cross=this.lastVisiblePrice,a=g[h-c],this.crossLabel&&this.crossLabel.destroy(),
delete b.crossLabel,b.drawCrosshair(null,a),b.cross&&(this.lastVisiblePrice=b.cross,"number"===typeof a.y&&(this.lastVisiblePrice.y=a.y)),this.crossLabel=b.crossLabel);b.crosshair=b.options.crosshair=p;b.cross=q;b.crossLabel=r}})});c(a,"masters/modules/price-indicator.src.js",[],function(){})});
//# sourceMappingURL=price-indicator.js.map
```
