## Supplementary material for "Exploring natural odour landscapes: A case study with implications for human-biting insects": Figure S4: pyramid3d.js

```
/*
 Highcharts JS v9.3.1 (2021-11-05)

 Highcharts 3D funnel module

 (c) 2010-2021 Kacper Madej

 License: www.highcharts.com/license
*/
'use strict';(function(a){"object"===typeof module&&module.exports?(a["default"]=a,module.exports=a):"function"===typeof define&&define.amd?define("highcharts/modules/pyramid3d",["highcharts","highcharts/highcharts-3d","highcharts/modules/cylinder","highcharts/modules/funnel3d"],function(b){a(b);a.Highcharts=b;return a}):a("undefined"!==typeof Highcharts?Highcharts:void 0)})(function(a){function b(a,d,b,e){a.hasOwnProperty(d)||(a[d]=e.apply(null,b))}a=a?a._modules:{};b(a,"Series/Pyramid3D/Pyramid3DSeries.js",
[a["Core/Series/SeriesRegistry.js"],a["Core/Utilities.js"]],function(a,b){var d=this&&this.__extends||function(){var a=function(b,c){a=Object.setPrototypeOf||{__proto__:[]}instanceof Array&&function(a,b){a.__proto__=b}||function(a,b){for(var c in b)b.hasOwnProperty(c)&&(a[c]=b[c])};return a(b,c)};return function(b,c){function d(){this.constructor=b}a(b,c);b.prototype=null===c?Object.create(c):(d.prototype=c.prototype,new d)}}(),e=a.seriesTypes.funnel3d,f=b.merge;b=function(a){function b(){var b=null!==
a&&a.apply(this,arguments)||this;b.data=void 0;b.options=void 0;b.points=void 0;return b}d(b,a);b.defaultOptions=f(e.defaultOptions,{reversed:!0,neckHeight:0,neckWidth:0,dataLabels:{verticalAlign:"top"}});return b}(e);a.registerSeriesType("pyramid3d",b);"";return b});b(a,"masters/modules/pyramid3d.src.js",[],function(){})});
//# sourceMappingURL=pyramid3d.js.map
```
