## Supplementary material for "Exploring natural odour landscapes: A case study with implications for human-biting insects": Figure S4: sankey.js

```
/*
 Highcharts JS v9.3.1 (2021-11-05)

 Sankey diagram module

 (c) 2010-2021 Torstein Honsi

 License: www.highcharts.com/license
*/
'use strict';(function(c){"object"===typeof module&&module.exports?(c["default"]=c,module.exports=c):"function"===typeof define&&define.amd?define("highcharts/modules/sankey",["highcharts"],function(n){c(n);c.Highcharts=n;return c}):c("undefined"!==typeof Highcharts?Highcharts:void 0)})(function(c){function n(c,e,k,l){c.hasOwnProperty(e)||(c[e]=l.apply(null,k))}c=c?c._modules:{};n(c,"Series/NodesComposition.js",[c["Core/Series/Point.js"],c["Core/Series/Series.js"],c["Core/Utilities.js"]],function(c,
e,k){var l=k.defined,u=k.extend,w=k.find,d=k.pick,a;(function(a){function b(){this.data=[].concat(this.points||[],this.nodes);return e.prototype.destroy.apply(this,arguments)}function q(){this.nodes&&(this.nodes.forEach(function(b){b.destroy()}),this.nodes.length=0);e.prototype.setData.apply(this,arguments)}function m(b){var a=arguments,f=this.isNode?this.linksTo.concat(this.linksFrom):[this.fromNode,this.toNode];"select"!==b&&f.forEach(function(b){b&&b.series&&(c.prototype.setState.apply(b,a),b.isNode||
(b.fromNode.graphic&&c.prototype.setState.apply(b.fromNode,a),b.toNode&&b.toNode.graphic&&c.prototype.setState.apply(b.toNode,a)))});c.prototype.setState.apply(this,a)}var x=[];a.compose=function(a,c){-1===x.indexOf(a)&&(x.push(a),a=a.prototype,a.setNodeState=m,a.setState=m);-1===x.indexOf(c)&&(x.push(c),a=c.prototype,a.destroy=b,a.setData=q);return c};a.createNode=function(b){var a=this.pointClass,f=function(b,a){return w(b,function(b){return b.id===a})},g=f(this.nodes,b);g||(f=this.options.nodes&&
f(this.options.nodes,b),g=(new a).init(this,u({className:"highcharts-node",isNode:!0,id:b,y:1},f)),g.linksTo=[],g.linksFrom=[],g.formatPrefix="node",g.name=g.name||g.options.id||"",g.mass=d(g.options.mass,g.options.marker&&g.options.marker.radius,this.options.marker&&this.options.marker.radius,4),g.getSum=function(){var b=0,a=0;g.linksTo.forEach(function(a){b+=a.weight});g.linksFrom.forEach(function(b){a+=b.weight});return Math.max(b,a)},g.offset=function(b,a){for(var f=0,h=0;h<g[a].length;h++){if(g[a][h]===
b)return f;f+=g[a][h].weight}},g.hasShape=function(){var b=0;g.linksTo.forEach(function(a){a.outgoing&&b++});return!g.linksTo.length||b!==g.linksTo.length},this.nodes.push(g));return g};a.destroy=b;a.generatePoints=function(){var b=this.chart,a={};e.prototype.generatePoints.call(this);this.nodes||(this.nodes=[]);this.colorCounter=0;this.nodes.forEach(function(b){b.linksFrom.length=0;b.linksTo.length=0;b.level=b.options.level});this.points.forEach(function(f){l(f.from)&&(a[f.from]||(a[f.from]=this.createNode(f.from)),
a[f.from].linksFrom.push(f),f.fromNode=a[f.from],b.styledMode?f.colorIndex=d(f.options.colorIndex,a[f.from].colorIndex):f.color=f.options.color||a[f.from].color);l(f.to)&&(a[f.to]||(a[f.to]=this.createNode(f.to)),a[f.to].linksTo.push(f),f.toNode=a[f.to]);f.name=f.name||f.id},this);this.nodeLookup=a};a.setNodeState=m})(a||(a={}));return a});n(c,"Series/Sankey/SankeyPoint.js",[c["Core/Series/Point.js"],c["Core/Series/SeriesRegistry.js"],c["Core/Utilities.js"]],function(c,e,k){var l=this&&this.__extends||
function(){var c=function(d,a){c=Object.setPrototypeOf||{__proto__:[]}instanceof Array&&function(a,b){a.__proto__=b}||function(a,b){for(var c in b)b.hasOwnProperty(c)&&(a[c]=b[c])};return c(d,a)};return function(d,a){function e(){this.constructor=d}c(d,a);d.prototype=null===a?Object.create(a):(e.prototype=a.prototype,new e)}}(),u=k.defined;return function(e){function d(){var a=null!==e&&e.apply(this,arguments)||this;a.className=void 0;a.fromNode=void 0;a.level=void 0;a.linkBase=void 0;a.linksFrom=
void 0;a.linksTo=void 0;a.mass=void 0;a.nodeX=void 0;a.nodeY=void 0;a.options=void 0;a.series=void 0;a.toNode=void 0;return a}l(d,e);d.prototype.applyOptions=function(a,d){c.prototype.applyOptions.call(this,a,d);u(this.options.level)&&(this.options.column=this.column=this.options.level);return this};d.prototype.getClassName=function(){return(this.isNode?"highcharts-node ":"highcharts-link ")+c.prototype.getClassName.call(this)};d.prototype.isValid=function(){return this.isNode||"number"===typeof this.weight};
return d}(e.seriesTypes.column.prototype.pointClass)});n(c,"Series/TreeUtilities.js",[c["Core/Color/Color.js"],c["Core/Utilities.js"]],function(c,e){function k(b,a){var c=a.before,d=a.idRoot,e=a.mapIdToNode[d],q=a.points[b.i],f=q&&q.options||{},g=[],t=0;b.levelDynamic=b.level-(!1!==a.levelIsConstant?0:e.level);b.name=m(q&&q.name,"");b.visible=d===b.id||!0===a.visible;"function"===typeof c&&(b=c(b,a));b.children.forEach(function(c,f){var h=l({},a);l(h,{index:f,siblings:b.children.length,visible:b.visible});
c=k(c,h);g.push(c);c.visible&&(t+=c.val)});c=m(f.value,t);b.visible=0<=c&&(0<t||b.visible);b.children=g;b.childrenTotal=t;b.isLeaf=b.visible&&!t;b.val=c;return b}var l=e.extend,u=e.isArray,w=e.isNumber,d=e.isObject,a=e.merge,m=e.pick;return{getColor:function(b,a){var d=a.index,e=a.mapOptionsToLevel,q=a.parentColor,k=a.parentColorIndex,f=a.series,g=a.colors,t=a.siblings,y=f.points,A=f.chart.options.chart,h;if(b){y=y[b.i];b=e[b.level]||{};if(e=y&&b.colorByPoint){var v=y.index%(g?g.length:A.colorCount);
var G=g&&g[v]}if(!f.chart.styledMode){g=y&&y.options.color;A=b&&b.color;if(h=q)h=(h=b&&b.colorVariation)&&"brightness"===h.key&&d&&t?c.parse(q).brighten(d/t*h.to).get():q;h=m(g,A,G,h,f.color)}var B=m(y&&y.options.colorIndex,b&&b.colorIndex,v,k,a.colorIndex)}return{color:h,colorIndex:B}},getLevelOptions:function(b){var c=null;if(d(b)){c={};var e=w(b.from)?b.from:1;var k=b.levels;var l={};var n=d(b.defaults)?b.defaults:{};u(k)&&(l=k.reduce(function(b,c){if(d(c)&&w(c.level)){var f=a({},c);var g=m(f.levelIsConstant,
n.levelIsConstant);delete f.levelIsConstant;delete f.level;c=c.level+(g?0:e-1);d(b[c])?a(!0,b[c],f):b[c]=f}return b},{}));k=w(b.to)?b.to:1;for(b=0;b<=k;b++)c[b]=a({},n,d(l[b])?l[b]:{})}return c},setTreeValues:k,updateRootId:function(b){if(d(b)){var a=d(b.options)?b.options:{};a=m(b.rootNode,a.rootId,"");d(b.userOptions)&&(b.userOptions.rootId=a);b.rootNode=a}return a}}});n(c,"Series/Sankey/SankeySeries.js",[c["Core/Color/Color.js"],c["Core/Globals.js"],c["Series/NodesComposition.js"],c["Series/Sankey/SankeyPoint.js"],
c["Core/Series/SeriesRegistry.js"],c["Series/TreeUtilities.js"],c["Core/Utilities.js"]],function(c,e,k,l,n,w,d){var a=this&&this.__extends||function(){var b=function(a,h){b=Object.setPrototypeOf||{__proto__:[]}instanceof Array&&function(h,b){h.__proto__=b}||function(h,b){for(var a in b)b.hasOwnProperty(a)&&(h[a]=b[a])};return b(a,h)};return function(a,h){function c(){this.constructor=a}b(a,h);a.prototype=null===h?Object.create(h):(c.prototype=h.prototype,new c)}}(),m=n.series,b=n.seriesTypes.column,
q=w.getLevelOptions,u=d.defined;w=d.extend;var x=d.find,C=d.isObject,D=d.merge,f=d.pick,g=d.relativeLength,t=d.stableSort;d=function(d){function e(){var b=null!==d&&d.apply(this,arguments)||this;b.colDistance=void 0;b.data=void 0;b.group=void 0;b.nodeLookup=void 0;b.nodePadding=void 0;b.nodes=void 0;b.nodeWidth=void 0;b.options=void 0;b.points=void 0;b.translationFactor=void 0;return b}a(e,d);e.getDLOptions=function(b){var a=C(b.optionsPoint)?b.optionsPoint.dataLabels:{};b=C(b.level)?b.level.dataLabels:
{};return D({style:{}},b,a)};e.prototype.createNodeColumn=function(){var b=this,a=this.chart,c=[];c.sum=function(){return this.reduce(function(b,a){return b+a.getSum()},0)};c.offset=function(h,f){for(var v=0,r,d=b.nodePadding,p=0;p<c.length;p++){r=c[p].getSum();var e=Math.max(r*f,b.options.minLinkWidth),B=h.options[a.inverted?"offsetHorizontal":"offsetVertical"],G=h.options.offset||0;r=r?e+d:0;if(c[p]===h)return{relativeTop:v+(u(B)?g(B,e):g(G,r))};v+=r}};c.top=function(h){var c=b.nodePadding,f=this.reduce(function(a,
f){0<a&&(a+=c);f=Math.max(f.getSum()*h,b.options.minLinkWidth);return a+f},0);return(a.plotSizeY-f)/2};return c};e.prototype.createNodeColumns=function(){var b=[];this.nodes.forEach(function(a){var c=-1;if(!u(a.options.column))if(0===a.linksTo.length)a.column=0;else{for(var h=0;h<a.linksTo.length;h++){var f=a.linksTo[h];if(f.fromNode.column>c&&f.fromNode!==a){var d=f.fromNode;c=d.column}}a.column=c+1;if(d&&"hanging"===d.options.layout){a.hangsFrom=d;var v=-1;x(d.linksFrom,function(b,c){(b=b.toNode===
a)&&(v=c);return b});a.column+=v}}b[a.column]||(b[a.column]=this.createNodeColumn());b[a.column].push(a)},this);for(var a=0;a<b.length;a++)"undefined"===typeof b[a]&&(b[a]=this.createNodeColumn());return b};e.prototype.generatePoints=function(){function b(a,c){"undefined"===typeof a.level&&(a.level=c,a.linksFrom.forEach(function(a){a.toNode&&b(a.toNode,c+1)}))}k.generatePoints.apply(this,arguments);this.orderNodes&&(this.nodes.filter(function(b){return 0===b.linksTo.length}).forEach(function(a){b(a,
0)}),t(this.nodes,function(b,a){return b.level-a.level}))};e.prototype.getNodePadding=function(){var b=this.options.nodePadding||0;if(this.nodeColumns){var a=this.nodeColumns.reduce(function(b,a){return Math.max(b,a.length)},0);a*b>this.chart.plotSizeY&&(b=this.chart.plotSizeY/a)}return b};e.prototype.hasData=function(){return!!this.processedXData.length};e.prototype.pointAttribs=function(b,a){if(!b)return{};var h=this,d=h.mapOptionsToLevel[(b.isNode?b.level:b.fromNode.level)||0]||{},e=b.options,
g=d.states&&d.states[a||""]||{};a=["colorByPoint","borderColor","borderWidth","linkOpacity"].reduce(function(b,a){b[a]=f(g[a],e[a],d[a],h.options[a]);return b},{});var r=f(g.color,e.color,a.colorByPoint?b.color:d.color);return b.isNode?{fill:r,stroke:a.borderColor,"stroke-width":a.borderWidth}:{fill:c.parse(r).setOpacity(a.linkOpacity).get()}};e.prototype.render=function(){var a=this.points;this.points=this.points.concat(this.nodes||[]);b.prototype.render.call(this);this.points=a};e.prototype.translate=
function(){var b=this,a=function(a){for(var h=a.slice(),e=b.options.minLinkWidth||0,g,r=0,v,B=f.plotSizeY-d.borderWidth-(a.length-1)*c.nodePadding;a.length;){r=B/a.sum();g=!1;for(v=a.length;v--;)a[v].getSum()*r<e&&(a.splice(v,1),B-=e,g=!0);if(!g)break}a.length=0;h.forEach(function(b){return a.push(b)});return r};this.processedXData||this.processData();this.generatePoints();this.nodeColumns=this.createNodeColumns();this.nodeWidth=g(this.options.nodeWidth,this.chart.plotSizeX);var c=this,f=this.chart,
d=this.options,e=this.nodeWidth,r=this.nodeColumns;this.nodePadding=this.getNodePadding();this.translationFactor=r.reduce(function(b,c){return Math.min(b,a(c))},Infinity);this.colDistance=(f.plotSizeX-e-d.borderWidth)/Math.max(1,r.length-1);c.mapOptionsToLevel=q({from:1,levels:d.levels,to:r.length-1,defaults:{borderColor:d.borderColor,borderRadius:d.borderRadius,borderWidth:d.borderWidth,color:c.color,colorByPoint:d.colorByPoint,levelIsConstant:!0,linkColor:d.linkColor,linkLineWidth:d.linkLineWidth,
linkOpacity:d.linkOpacity,states:d.states}});r.forEach(function(a){a.forEach(function(b){c.translateNode(b,a)})},this);this.nodes.forEach(function(a){a.linksFrom.forEach(function(a){(a.weight||a.isNull)&&a.to&&(c.translateLink(a),a.allowShadow=!1)})})};e.prototype.translateLink=function(a){var b=function(b,c){c=b.offset(a,c)*h;return Math.min(b.nodeY+c,b.nodeY+(b.shapeArgs&&b.shapeArgs.height||0)-e)},c=a.fromNode,d=a.toNode,f=this.chart,h=this.translationFactor,e=Math.max(a.weight*h,this.options.minLinkWidth),
g=(f.inverted?-this.colDistance:this.colDistance)*this.options.curveFactor,p=b(c,"linksFrom");b=b(d,"linksTo");var k=c.nodeX,l=this.nodeWidth;d=d.nodeX;var n=a.outgoing,m=d>k+l;f.inverted&&(p=f.plotSizeY-p,b=(f.plotSizeY||0)-b,l=-l,e=-e,m=k>d);a.shapeType="path";a.linkBase=[p,p+e,b,b+e];if(m&&"number"===typeof b)a.shapeArgs={d:[["M",k+l,p],["C",k+l+g,p,d-g,b,d,b],["L",d+(n?l:0),b+e/2],["L",d,b+e],["C",d-g,b+e,k+l+g,p+e,k+l,p+e],["Z"]]};else if("number"===typeof b){g=d-20-e;n=d-20;m=k+l;var q=m+20,
t=q+e,w=p,u=p+e,y=u+20,x=y+(f.plotHeight-p-e),z=x+20,F=z+e,A=b,E=A+e,C=E+20,D=z+.7*e,H=d-.7*e,I=m+.7*e;a.shapeArgs={d:[["M",m,w],["C",I,w,t,u-.7*e,t,y],["L",t,x],["C",t,D,I,F,m,F],["L",d,F],["C",H,F,g,D,g,x],["L",g,C],["C",g,E-.7*e,H,A,d,A],["L",d,E],["C",n,E,n,E,n,C],["L",n,x],["C",n,z,n,z,d,z],["L",m,z],["C",q,z,q,z,q,x],["L",q,y],["C",q,u,q,u,m,u],["Z"]]}}a.dlBox={x:k+(d-k+l)/2,y:p+(b-p)/2,height:e,width:0};a.tooltipPos=f.inverted?[f.plotSizeY-a.dlBox.y-e/2,f.plotSizeX-a.dlBox.x]:[a.dlBox.x,a.dlBox.y+
e/2];a.y=a.plotY=1;a.color||(a.color=c.color)};e.prototype.translateNode=function(a,b){var c=this.translationFactor,d=this.chart,h=this.options,k=a.getSum(),l=Math.max(Math.round(k*c),this.options.minLinkWidth),n=Math.round(this.nodeWidth),p=Math.round(h.borderWidth)%2/2,m=b.offset(a,c);b=Math.floor(f(m.absoluteTop,b.top(c)+m.relativeTop))+p;p=Math.floor(this.colDistance*a.column+h.borderWidth/2)+g(a.options.offsetHorizontal||0,n)+p;p=d.inverted?d.plotSizeX-p:p;if(a.sum=k){a.shapeType="rect";a.nodeX=
p;a.nodeY=b;k=p;c=b;m=a.options.width||h.width||n;var q=a.options.height||h.height||l;d.inverted&&(k=p-n,c=d.plotSizeY-b-l,m=a.options.height||h.height||n,q=a.options.width||h.width||l);a.dlOptions=e.getDLOptions({level:this.mapOptionsToLevel[a.level],optionsPoint:a.options});a.plotX=1;a.plotY=1;a.tooltipPos=d.inverted?[d.plotSizeY-c-q/2,d.plotSizeX-k-m/2]:[k+m/2,c+q/2];a.shapeArgs={x:k,y:c,width:m,height:q,display:a.hasShape()?"":"none"}}else a.dlOptions={enabled:!1}};e.defaultOptions=D(b.defaultOptions,
{borderWidth:0,colorByPoint:!0,curveFactor:.33,dataLabels:{enabled:!0,backgroundColor:"none",crop:!1,nodeFormat:void 0,nodeFormatter:function(){return this.point.name},format:void 0,formatter:function(){},inside:!0},inactiveOtherPoints:!0,linkOpacity:.5,minLinkWidth:0,nodeWidth:20,nodePadding:10,showInLegend:!1,states:{hover:{linkOpacity:1},inactive:{linkOpacity:.1,opacity:.1,animation:{duration:50}}},tooltip:{followPointer:!0,headerFormat:'<span style="font-size: 10px">{series.name}</span><br/>',
pointFormat:"{point.fromNode.name} \u2192 {point.toNode.name}: <b>{point.weight}</b><br/>",nodeFormat:"{point.name}: <b>{point.sum}</b><br/>"}});return e}(b);k.compose(l,d);w(d.prototype,{animate:m.prototype.animate,createNode:k.createNode,forceDL:!0,invertible:!0,isCartesian:!1,orderNodes:!0,noSharedTooltip:!0,pointArrayMap:["from","to"],pointClass:l,searchPoint:e.noop});n.registerSeriesType("sankey",d);"";"";return d});n(c,"masters/modules/sankey.src.js",[],function(){})});
//# sourceMappingURL=sankey.js.map
```
