## Supplementary material for "Exploring natural odour landscapes: A case study with implications for human-biting insects": Figure S4: series-label.js

```
/*
 Highcharts JS v9.3.1 (2021-11-05)

 (c) 2009-2021 Torstein Honsi

 License: www.highcharts.com/license
*/
'use strict';(function(k){"object"===typeof module&&module.exports?(k["default"]=k,module.exports=k):"function"===typeof define&&define.amd?define("highcharts/modules/series-label",["highcharts"],function(u){k(u);k.Highcharts=u;return k}):k("undefined"!==typeof Highcharts?Highcharts:void 0)})(function(k){function u(k,A,y,u){k.hasOwnProperty(A)||(k[A]=u.apply(null,y))}k=k?k._modules:{};u(k,"Extensions/SeriesLabel.js",[k["Core/Animation/AnimationUtilities.js"],k["Core/Chart/Chart.js"],k["Core/FormatUtilities.js"],
k["Core/DefaultOptions.js"],k["Core/Series/Series.js"],k["Core/Renderer/SVG/SVGRenderer.js"],k["Core/Utilities.js"]],function(k,u,y,K,E,F,x){function C(b,c,a,f,d,e){b=(e-c)*(a-b)-(f-c)*(d-b);return 0<b?!0:!(0>b)}function D(b,c,a,f,d,e,r,h){return C(b,c,d,e,r,h)!==C(a,f,d,e,r,h)&&C(b,c,a,f,d,e)!==C(b,c,a,f,r,h)}function A(b,c,a,f,d,e,r,h){return D(b,c,b+a,c,d,e,r,h)||D(b+a,c,b+a,c+f,d,e,r,h)||D(b,c+f,b+a,c+f,d,e,r,h)||D(b,c,b,c+f,d,e,r,h)}function I(b){if(this.renderer){var c=this,a=G(c.renderer.globalAnimation).duration;
c.labelSeries=[];c.labelSeriesMaxSum=0;x.clearTimeout(c.seriesLabelTimer);c.series.forEach(function(f){var d=f.options.label,e=f.labelBySeries,r=e&&e.closest;d.enabled&&f.visible&&(f.graph||f.area)&&!f.isSeriesBoosting&&(c.labelSeries.push(f),d.minFontSize&&d.maxFontSize&&(f.sum=f.yData.reduce(function(a,b){return(a||0)+(b||0)},0),c.labelSeriesMaxSum=Math.max(c.labelSeriesMaxSum,f.sum)),"load"===b.type&&(a=Math.max(a,G(f.options.animation).duration)),r&&("undefined"!==typeof r[0].plotX?e.animate({x:r[0].plotX+
r[1],y:r[0].plotY+r[2]}):e.attr({opacity:0})))});c.seriesLabelTimer=L(function(){c.series&&c.labelSeries&&c.drawSeriesLabels()},c.renderer.forExport||!a?0:a)}}var G=k.animObject,M=y.format;k=K.setOptions;F=F.prototype.symbols;y=x.addEvent;var J=x.extend,N=x.fireEvent,H=x.isNumber,B=x.pick,L=x.syncTimeout;"";k({plotOptions:{series:{label:{enabled:!0,connectorAllowed:!1,connectorNeighbourDistance:24,format:void 0,formatter:void 0,minFontSize:null,maxFontSize:null,onArea:null,style:{fontWeight:"bold"},
boxesToAvoid:[]}}}});F.connector=function(b,c,a,f,d){var e=d&&d.anchorX;d=d&&d.anchorY;var r=a/2;if(H(e)&&H(d)){var h=[["M",e,d]];var n=c-d;0>n&&(n=-f-n);n<a&&(r=e<b+a/2?n:a-n);d>c+f?h.push(["L",b+r,c+f]):d<c?h.push(["L",b+r,c]):e<b?h.push(["L",b,c+f/2]):e>b+a&&h.push(["L",b+a,c+f/2])}return h||[]};E.prototype.getPointsOnGraph=function(){function b(b){var c=Math.round(b.plotX/8)+","+Math.round(b.plotY/8);p[c]||(p[c]=1,a.push(b))}if(this.xAxis||this.yAxis){var c=this.points,a=[],f=this.graph||this.area,
d=f.element,e=this.chart.inverted,r=this.xAxis,h=this.yAxis,n=e?h.pos:r.pos;e=e?r.pos:h.pos;r=B(this.options.label.onArea,!!this.area);h=h.getThreshold(this.options.threshold);var p={},k;if(this.getPointSpline&&d.getPointAtLength&&!r&&c.length<this.chart.plotSizeX/16){if(f.toD){var g=f.attr("d");f.attr({d:f.toD})}var m=d.getTotalLength();for(k=0;k<m;k+=16){var l=d.getPointAtLength(k);b({chartX:n+l.x,chartY:e+l.y,plotX:l.x,plotY:l.y})}g&&f.attr({d:g});l=c[c.length-1];l.chartX=n+l.plotX;l.chartY=e+
l.plotY;b(l)}else for(m=c.length,k=0;k<m;k+=1){l=c[k];f=c[k-1];l.chartX=n+l.plotX;l.chartY=e+l.plotY;r&&(l.chartCenterY=e+(l.plotY+B(l.yBottom,h))/2);if(0<k&&(d=Math.abs(l.chartX-f.chartX),g=Math.abs(l.chartY-f.chartY),d=Math.max(d,g),16<d))for(d=Math.ceil(d/16),g=1;g<d;g+=1)b({chartX:f.chartX+g/d*(l.chartX-f.chartX),chartY:f.chartY+g/d*(l.chartY-f.chartY),chartCenterY:f.chartCenterY+g/d*(l.chartCenterY-f.chartCenterY),plotX:f.plotX+g/d*(l.plotX-f.plotX),plotY:f.plotY+g/d*(l.plotY-f.plotY)});H(l.plotY)&&
b(l)}return a}};E.prototype.labelFontSize=function(b,c){return b+this.sum/this.chart.labelSeriesMaxSum*(c-b)+"px"};E.prototype.checkClearPoint=function(b,c,a,f){var d=this.chart,e=B(this.options.label.onArea,!!this.area),k=e||this.options.label.connectorAllowed,h=Number.MAX_VALUE,n=Number.MAX_VALUE,p,z;for(z=0;z<d.boxesToAvoid.length;z+=1){var g=d.boxesToAvoid[z];var m=b+a.width;var l=c;var t=c+a.height;if(!(b>g.right||m<g.left||l>g.bottom||t<g.top))return!1}for(z=0;z<d.series.length;z+=1)if(l=d.series[z],
g=l.interpolatedPoints,l.visible&&g){for(m=1;m<g.length;m+=1){if(g[m].chartX>=b-16&&g[m-1].chartX<=b+a.width+16){if(A(b,c,a.width,a.height,g[m-1].chartX,g[m-1].chartY,g[m].chartX,g[m].chartY))return!1;this===l&&!p&&f&&(p=A(b-16,c-16,a.width+32,a.height+32,g[m-1].chartX,g[m-1].chartY,g[m].chartX,g[m].chartY))}if((k||p)&&(this!==l||e)){t=b+a.width/2-g[m].chartX;var u=c+a.height/2-g[m].chartY;h=Math.min(h,t*t+u*u)}}if(!e&&k&&this===l&&(f&&!p||h<Math.pow(this.options.label.connectorNeighbourDistance,
2))){for(m=1;m<g.length;m+=1)if(p=Math.min(Math.pow(b+a.width/2-g[m].chartX,2)+Math.pow(c+a.height/2-g[m].chartY,2),Math.pow(b-g[m].chartX,2)+Math.pow(c-g[m].chartY,2),Math.pow(b+a.width-g[m].chartX,2)+Math.pow(c-g[m].chartY,2),Math.pow(b+a.width-g[m].chartX,2)+Math.pow(c+a.height-g[m].chartY,2),Math.pow(b-g[m].chartX,2)+Math.pow(c+a.height-g[m].chartY,2)),p<n){n=p;var w=g[m]}p=!0}}return!f||p?{x:b,y:c,weight:h-(w?n:0),connectorPoint:w}:!1};u.prototype.drawSeriesLabels=function(){var b=this,c=this.labelSeries;
b.boxesToAvoid=[];c.forEach(function(a){a.interpolatedPoints=a.getPointsOnGraph();(a.options.label.boxesToAvoid||[]).forEach(function(a){b.boxesToAvoid.push(a)})});b.series.forEach(function(a){function c(a,b,c){var d=Math.max(u,B(y,-Infinity)),e=Math.min(u+m,B(A,Infinity));return a>d&&a<=e-c.width&&b>=g&&b<=g+l-c.height}var d=a.options.label;if(d&&(a.xAxis||a.yAxis)){var e="highcharts-color-"+B(a.colorIndex,"none"),k=!a.labelBySeries,h=d.minFontSize,n=d.maxFontSize,p=b.inverted,u=p?a.yAxis.pos:a.xAxis.pos,
g=p?a.xAxis.pos:a.yAxis.pos,m=b.inverted?a.yAxis.len:a.xAxis.len,l=b.inverted?a.xAxis.len:a.yAxis.len,t=a.interpolatedPoints,x=B(d.onArea,!!a.area),w=[],q,v=a.labelBySeries;if(x&&!p){p=[a.xAxis.toPixels(a.xData[0]),a.xAxis.toPixels(a.xData[a.xData.length-1])];var y=Math.min.apply(Math,p);var A=Math.max.apply(Math,p)}if(a.visible&&!a.isSeriesBoosting&&t){v||(v=a.name,"string"===typeof d.format?v=M(d.format,a,b):d.formatter&&(v=d.formatter.call(a)),a.labelBySeries=v=b.renderer.label(v,0,-9999,"connector").addClass("highcharts-series-label highcharts-series-label-"+
a.index+" "+(a.options.className||"")+" "+e),b.renderer.styledMode||(v.css(J({color:x?b.renderer.getContrast(a.color):a.color},d.style||{})),v.attr({opacity:b.renderer.forExport?1:0,stroke:a.color,"stroke-width":1})),h&&n&&v.css({fontSize:a.labelFontSize(h,n)}),v.attr({padding:0,zIndex:3}).add());e=v.getBBox();e.width=Math.round(e.width);for(p=t.length-1;0<p;--p)x?(h=t[p].chartX-e.width/2,n=t[p].chartCenterY-e.height/2,c(h,n,e)&&(q=a.checkClearPoint(h,n,e))):(h=t[p].chartX+3,n=t[p].chartY-e.height-
3,c(h,n,e)&&(q=a.checkClearPoint(h,n,e,!0)),q&&w.push(q),h=t[p].chartX+3,n=t[p].chartY+3,c(h,n,e)&&(q=a.checkClearPoint(h,n,e,!0)),q&&w.push(q),h=t[p].chartX-e.width-3,n=t[p].chartY+3,c(h,n,e)&&(q=a.checkClearPoint(h,n,e,!0)),q&&w.push(q),h=t[p].chartX-e.width-3,n=t[p].chartY-e.height-3,c(h,n,e)&&(q=a.checkClearPoint(h,n,e,!0))),q&&w.push(q);if(d.connectorAllowed&&!w.length&&!x)for(h=u+m-e.width;h>=u;h-=16)for(n=g;n<g+l-e.height;n+=16)(q=a.checkClearPoint(h,n,e,!0))&&w.push(q);if(w.length){if(w.sort(function(a,
b){return b.weight-a.weight}),q=w[0],b.boxesToAvoid.push({left:q.x,right:q.x+e.width,top:q.y,bottom:q.y+e.height}),(t=Math.sqrt(Math.pow(Math.abs(q.x-(v.x||0)),2)+Math.pow(Math.abs(q.y-(v.y||0)),2)))&&a.labelBySeries&&(w={opacity:b.renderer.forExport?1:0,x:q.x,y:q.y},d={opacity:1},10>=t&&(d={x:w.x,y:w.y},w={}),t=void 0,k&&(t=G(a.options.animation),t.duration*=.2),a.labelBySeries.attr(J(w,{anchorX:q.connectorPoint&&q.connectorPoint.plotX+u,anchorY:q.connectorPoint&&q.connectorPoint.plotY+g})).animate(d,
t),a.options.kdNow=!0,a.buildKDTree(),a=a.searchPoint({chartX:q.x,chartY:q.y},!0)))v.closest=[a,q.x-(a.plotX||0),q.y-(a.plotY||0)]}else v&&(a.labelBySeries=v.destroy())}else v&&(a.labelBySeries=v.destroy())}});N(b,"afterDrawSeriesLabels")};y(u,"load",I);y(u,"redraw",I)});u(k,"masters/modules/series-label.src.js",[],function(){})});
//# sourceMappingURL=series-label.js.map
```
