## Supplementary material for "Exploring natural odour landscapes: A case study with implications for human-biting insects": Figure S4: solid-gauge.js

```
/*
 Highcharts JS v9.3.1 (2021-11-05)

 Solid angular gauge module

 (c) 2010-2021 Torstein Honsi

 License: www.highcharts.com/license
*/
'use strict';(function(a){"object"===typeof module&&module.exports?(a["default"]=a,module.exports=a):"function"===typeof define&&define.amd?define("highcharts/modules/solid-gauge",["highcharts","highcharts/highcharts-more"],function(f){a(f);a.Highcharts=f;return a}):a("undefined"!==typeof Highcharts?Highcharts:void 0)})(function(a){function f(a,k,l,c){a.hasOwnProperty(k)||(a[k]=c.apply(null,l))}a=a?a._modules:{};f(a,"Core/Axis/SolidGaugeAxis.js",[a["Core/Color/Color.js"],a["Core/Utilities.js"]],function(a,
k){var l=a.parse,c=k.extend,e=k.merge,m;(function(a){var b={initDataClasses:function(a){var c=this.chart,n,p=0,g=this.options;this.dataClasses=n=[];a.dataClasses.forEach(function(b,d){b=e(b);n.push(b);b.color||("category"===g.dataClassColor?(d=c.options.colors,b.color=d[p++],p===d.length&&(p=0)):b.color=l(g.minColor).tweenTo(l(g.maxColor),d/(a.dataClasses.length-1)))})},initStops:function(a){this.stops=a.stops||[[0,this.options.minColor],[1,this.options.maxColor]];this.stops.forEach(function(a){a.color=
l(a[1])})},toColor:function(a,c){var b=this.stops,l=this.dataClasses,g;if(l)for(g=l.length;g--;){var e=l[g];var d=e.from;b=e.to;if(("undefined"===typeof d||a>=d)&&("undefined"===typeof b||a<=b)){var k=e.color;c&&(c.dataClass=g);break}}else{this.logarithmic&&(a=this.val2lin(a));a=1-(this.max-a)/(this.max-this.min);for(g=b.length;g--&&!(a>b[g][0]););d=b[g]||b[g+1];b=b[g+1]||d;a=1-(b[0]-a)/(b[0]-d[0]||1);k=d.color.tweenTo(b.color,a)}return k}};a.init=function(a){c(a,b)}})(m||(m={}));return m});f(a,"Series/SolidGauge/SolidGaugeComposition.js",
[a["Core/Renderer/SVG/SVGRenderer.js"]],function(a){a=a.prototype;var k=a.symbols.arc;a.symbols.arc=function(a,c,e,m,b){a=k(a,c,e,m,b);b&&b.rounded&&(e=((b.r||e)-(b.innerR||0))/2,c=a[0],b=a[2],"M"===c[0]&&"L"===b[0]&&(c=["A",e,e,0,1,1,c[1],c[2]],a[2]=["A",e,e,0,1,1,b[1],b[2]],a[4]=c));return a}});f(a,"Series/SolidGauge/SolidGaugeSeries.js",[a["Core/Legend/LegendSymbol.js"],a["Core/Series/SeriesRegistry.js"],a["Core/Axis/SolidGaugeAxis.js"],a["Core/Utilities.js"]],function(a,k,l,c){var e=this&&this.__extends||
function(){var a=function(b,h){a=Object.setPrototypeOf||{__proto__:[]}instanceof Array&&function(a,b){a.__proto__=b}||function(a,b){for(var h in b)b.hasOwnProperty(h)&&(a[h]=b[h])};return a(b,h)};return function(b,h){function c(){this.constructor=b}a(b,h);b.prototype=null===h?Object.create(h):(c.prototype=h.prototype,new c)}}(),m=k.seriesTypes,b=m.gauge,f=m.pie.prototype,p=c.clamp,u=c.extend,n=c.isNumber,w=c.merge,g=c.pick,v=c.pInt,d={colorByPoint:!0,dataLabels:{y:0}};c=function(a){function c(){var b=
null!==a&&a.apply(this,arguments)||this;b.data=void 0;b.points=void 0;b.options=void 0;b.axis=void 0;b.yAxis=void 0;b.startAngleRad=void 0;b.thresholdAngleRad=void 0;return b}e(c,a);c.prototype.translate=function(){var a=this.yAxis;l.init(a);!a.dataClasses&&a.options.dataClasses&&a.initDataClasses(a.options);a.initStops(a.options);b.prototype.translate.call(this)};c.prototype.drawPoints=function(){var a=this,b=a.yAxis,c=b.center,e=a.options,k=a.chart.renderer,d=e.overshoot,l=n(d)?d/180*Math.PI:0,
f;n(e.threshold)&&(f=b.startAngleRad+b.translate(e.threshold,null,null,null,!0));this.thresholdAngleRad=g(f,b.startAngleRad);a.points.forEach(function(d){if(!d.isNull){var h=d.graphic,f=b.startAngleRad+b.translate(d.y,null,null,null,!0),m=v(g(d.options.radius,e.radius,100))*c[2]/200,q=v(g(d.options.innerRadius,e.innerRadius,60))*c[2]/200,r=b.toColor(d.y,d),t=Math.min(b.startAngleRad,b.endAngleRad),n=Math.max(b.startAngleRad,b.endAngleRad);"none"===r&&(r=d.color||a.color||"none");"none"!==r&&(d.color=
r);f=p(f,t-l,n+l);!1===e.wrap&&(f=p(f,t,n));t=Math.min(f,a.thresholdAngleRad);f=Math.max(f,a.thresholdAngleRad);f-t>2*Math.PI&&(f=t+2*Math.PI);d.shapeArgs=q={x:c[0],y:c[1],r:m,innerR:q,start:t,end:f,rounded:e.rounded};d.startR=m;h?(m=q.d,h.animate(u({fill:r},q)),m&&(q.d=m)):d.graphic=h=k.arc(q).attr({fill:r,"sweep-flag":0}).add(a.group);a.chart.styledMode||("square"!==e.linecap&&h.attr({"stroke-linecap":"round","stroke-linejoin":"round"}),h.attr({stroke:e.borderColor||"none","stroke-width":e.borderWidth||
0}));h&&h.addClass(d.getClassName(),!0)}})};c.prototype.animate=function(a){a||(this.startAngleRad=this.thresholdAngleRad,f.animate.call(this,a))};c.defaultOptions=w(b.defaultOptions,d);return c}(b);u(c.prototype,{drawLegendSymbol:a.drawRectangle});k.registerSeriesType("solidgauge",c);"";return c});f(a,"masters/modules/solid-gauge.src.js",[],function(){})});
//# sourceMappingURL=solid-gauge.js.map
```
