## Supplementary material for "Exploring natural odour landscapes: A case study with implications for human-biting insects": Figure S4: sonification.js

```
/*
 Highcharts JS v9.3.1 (2021-11-05)

 Sonification module

 (c) 2012-2021 ystein Moseng

 License: www.highcharts.com/license
*/
'use strict';(function(a){"object"===typeof module&&module.exports?(a["default"]=a,module.exports=a):"function"===typeof define&&define.amd?define("highcharts/modules/sonification",["highcharts"],function(l){a(l);a.Highcharts=l;return a}):a("undefined"!==typeof Highcharts?Highcharts:void 0)})(function(a){function l(a,f,h,k){a.hasOwnProperty(f)||(a[f]=k.apply(null,h))}a=a?a._modules:{};l(a,"Extensions/Sonification/MusicalFrequencies.js",[],function(){return[16.351597831287414,17.323914436054505,18.354047994837977,
19.445436482630058,20.601722307054366,21.826764464562746,23.12465141947715,24.499714748859326,25.956543598746574,27.5,29.13523509488062,30.86770632850775,32.70319566257483,34.64782887210901,36.70809598967594,38.890872965260115,41.20344461410875,43.653528929125486,46.2493028389543,48.999429497718666,51.91308719749314,55,58.27047018976124,61.7354126570155,65.40639132514966,69.29565774421802,73.41619197935188,77.78174593052023,82.4068892282175,87.30705785825097,92.4986056779086,97.99885899543733,103.82617439498628,
110,116.54094037952248,123.47082531403103,130.8127826502993,138.59131548843604,146.8323839587038,155.56349186104046,164.81377845643496,174.61411571650194,184.9972113558172,195.99771799087463,207.65234878997256,220,233.08188075904496,246.94165062806206,261.6255653005986,277.1826309768721,293.6647679174076,311.1269837220809,329.6275569128699,349.2282314330039,369.9944227116344,391.99543598174927,415.3046975799451,440,466.1637615180899,493.8833012561241,523.2511306011972,554.3652619537442,587.3295358348151,
622.2539674441618,659.2551138257398,698.4564628660078,739.9888454232688,783.9908719634985,830.6093951598903,880,932.3275230361799,987.7666025122483,1046.5022612023945,1108.7305239074883,1174.6590716696303,1244.5079348883237,1318.5102276514797,1396.9129257320155,1479.9776908465376,1567.981743926997,1661.2187903197805,1760,1864.6550460723597,1975.533205024496,2093.004522404789,2217.4610478149766,2349.31814333926,2489.0158697766474,2637.02045530296,2793.825851464031,2959.955381693075,3135.9634878539946,
3322.437580639561,3520,3729.3100921447194,3951.066410048992,4186.009044809578]});l(a,"Extensions/Sonification/SignalHandler.js",[],function(){return function(){function a(a){this.supportedSignals=this.signals=void 0;this.init(a||[])}a.prototype.init=function(a){this.supportedSignals=a;this.signals={}};a.prototype.registerSignalCallbacks=function(a){var f=this;f.supportedSignals.forEach(function(k){var n=a[k];n&&(f.signals[k]=f.signals[k]||[]).push(n)})};a.prototype.clearSignalCallbacks=function(a){var f=
this;a?a.forEach(function(a){f.signals[a]&&delete f.signals[a]}):f.signals={}};a.prototype.emitSignal=function(a,h){var f;this.signals[a]&&this.signals[a].forEach(function(a){a=a(h);f="undefined"!==typeof a?a:f});return f};return a}()});l(a,"Extensions/Sonification/SonificationUtilities.js",[a["Extensions/Sonification/MusicalFrequencies.js"],a["Extensions/Sonification/SignalHandler.js"],a["Core/Utilities.js"]],function(a,f,h){var k=h.clamp,n=h.merge,e={musicalFrequencies:a,SignalHandler:f,getExtremesForInstrumentProps:function(a,
b,g){var c=a.options.sonification&&a.options.sonification.defaultInstrumentOptions,q=function(a){return{instrumentMapping:a.mapping}},r=(b||[]).slice(0);c&&r.push(q(c));a.series.forEach(function(a){(a=a.options.sonification&&a.options.sonification.instruments)&&(r=r.concat(a.map(q)))});return r.reduce(function(c,b){Object.keys(b.instrumentMapping||{}).forEach(function(q){q=b.instrumentMapping[q];"string"!==typeof q||c[q]||(c[q]=e.calculateDataExtremes(a,q))});return c},n(g))},getMusicalScale:function(c){return a.filter(function(a,
g){var b=g%12+1;return c.some(function(a){return a===b})})},calculateDataExtremes:function(a,b){return a.series.reduce(function(a,c){c.points.forEach(function(c){c="undefined"!==typeof c[b]?c[b]:c.options[b];a.min=Math.min(a.min,c);a.max=Math.max(a.max,c)});return a},{min:Infinity,max:-Infinity})},virtualAxisTranslate:function(a,b,g,e){var c=b.max-b.min;a=g.min+Math.abs(g.max-g.min)*(e?b.max-a:a-b.min)/c;return 0<c?k(a,g.min,g.max):g.min}};return e});l(a,"Extensions/Sonification/Options.js",[],function(){return{sonification:{enabled:!1,
duration:2500,afterSeriesWait:700,masterVolume:1,order:"sequential",defaultInstrumentOptions:{instrument:"sineMusical",minFrequency:392,maxFrequency:1046,mapping:{pointPlayTime:"x",duration:200,frequency:"y"}}}}});l(a,"Extensions/Sonification/Sonification.js",[a["Core/DefaultOptions.js"],a["Core/Utilities.js"],a["Extensions/Sonification/SonificationUtilities.js"],a["Extensions/Sonification/Options.js"]],function(a,f,h,k){f=f.merge;f(!0,a.defaultOptions,k);a={fadeOutDuration:20,utilities:h};"";return a});
l(a,"Extensions/Sonification/Instrument.js",[a["Core/Globals.js"],a["Extensions/Sonification/Sonification.js"],a["Extensions/Sonification/SonificationUtilities.js"],a["Core/Utilities.js"]],function(a,f,h,k){var n=a.win,e=k.error,c=k.merge,b=k.pick,g=k.uniqueKey,t=function(){function a(a){this.playCallbackTimers=this.options=this.masterVolume=this.id=void 0;this.init(a)}a.prototype.init=function(b){if(this.initAudioContext()){this.options=c(a.defaultOptions,b);this.id=this.options.id=b&&b.id||g();
this.masterVolume=this.options.masterVolume||0;b=a.audioContext;var d=this.destinationNode||b.destination;this.gainNode=b.createGain();this.setGain(0);(this.panNode=b.createStereoPanner&&b.createStereoPanner())?(this.setPan(0),this.gainNode.connect(this.panNode),this.panNode.connect(d)):this.gainNode.connect(d);"oscillator"===this.options.type&&this.initOscillator(this.options.oscillator);this.playCallbackTimers=[]}else e(29)};a.prototype.copy=function(b){return new a(c(this.options,{id:null},b))};
a.prototype.initAudioContext=function(){var b=n.AudioContext||n.webkitAudioContext,c=!!a.audioContext;return b?(a.audioContext=a.audioContext||new b,!c&&a.audioContext&&"running"===a.audioContext.state&&a.audioContext.suspend(),!!(a.audioContext&&a.audioContext.createOscillator&&a.audioContext.createGain)):!1};a.prototype.initOscillator=function(b){this.oscillator=a.audioContext.createOscillator();this.oscillator.type=b.waveformShape;this.oscillator.connect(this.gainNode);this.oscillatorStarted=!1};
a.prototype.setPan=function(b){this.panNode&&this.panNode.pan.setValueAtTime(b,a.audioContext.currentTime)};a.prototype.setGain=function(b,c){var d=this.gainNode;b*=this.masterVolume;d&&(1.2<b&&(console.warn("Highcharts sonification warning: Volume of instrument set too high."),b=1.2),c?(d.gain.setValueAtTime(d.gain.value,a.audioContext.currentTime),d.gain.linearRampToValueAtTime(b,a.audioContext.currentTime+c/1E3)):d.gain.setValueAtTime(b,a.audioContext.currentTime))};a.prototype.cancelGainRamp=
function(){this.gainNode&&this.gainNode.gain.cancelScheduledValues(0)};a.prototype.setMasterVolume=function(a){this.masterVolume=a||0};a.prototype.getValidFrequency=function(a,c,m){var d=this.options.allowedFrequencies,g=b(m,Infinity),e=b(c,-Infinity);return d&&d.length?d.reduce(function(b,c){return Math.abs(c-a)<Math.abs(b-a)&&c<g&&c>e?c:b},Infinity):a};a.prototype.clearPlayCallbackTimers=function(){this.playCallbackTimers.forEach(function(a){clearInterval(a)});this.playCallbackTimers=[]};a.prototype.setFrequency=
function(a,b){b=b||{};a=this.getValidFrequency(a,b.min,b.max);"oscillator"===this.options.type&&this.oscillatorPlay(a)};a.prototype.oscillatorPlay=function(b){this.oscillatorStarted||(this.oscillator.start(),this.oscillatorStarted=!0);this.oscillator.frequency.setValueAtTime(b,a.audioContext.currentTime)};a.prototype.preparePlay=function(){this.setGain(.001);"suspended"===a.audioContext.state&&a.audioContext.resume();this.oscillator&&!this.oscillatorStarted&&(this.oscillator.start(),this.oscillatorStarted=
!0)};a.prototype.play=function(c){var d=this,m=c.duration||0,g=function(a,b,g){var m=c.duration,e=d.options.playCallbackInterval,w=0;if("function"===typeof a){var f=setInterval(function(){w++;var c=w*e/m;if(1<=c)d[b](a(1),g),clearInterval(f);else d[b](a(c),g)},e);d.playCallbackTimers.push(f)}else d[b](a,g)};if(d.id)if("suspended"===a.audioContext.state||this.oscillator&&!this.oscillatorStarted)d.preparePlay(),setTimeout(function(){d.play(c)},10);else{d.playCallbackTimers.length&&d.clearPlayCallbackTimers();
d.cancelGainRamp();d.stopOscillatorTimeout&&(clearTimeout(d.stopOscillatorTimeout),delete d.stopOscillatorTimeout);d.stopTimeout&&(clearTimeout(d.stopTimeout),delete d.stopTimeout,d.stopCallback&&(d._play=d.play,d.play=function(){},d.stopCallback("cancelled"),d.play=d._play));var e=m<f.fadeOutDuration+20;d.stopCallback=c.onEnd;var n=function(){delete d.stopTimeout;d.stop(e)};m?(d.stopTimeout=setTimeout(n,e?m:m-f.fadeOutDuration),g(c.frequency,"setFrequency",{minFrequency:c.minFrequency,maxFrequency:c.maxFrequency}),
g(b(c.volume,1),"setGain",4),g(b(c.pan,0),"setPan")):n()}};a.prototype.mute=function(){this.setGain(.0001,.8*f.fadeOutDuration)};a.prototype.stop=function(a,b,c){var d=this,g=function(){d.stopOscillatorTimeout&&delete d.stopOscillatorTimeout;if(d.oscillator&&d.options.oscillator){try{d.oscillator.stop()}catch(u){}d.gainNode&&d.oscillator.disconnect(d.gainNode);d.initOscillator(d.options.oscillator)}b&&b(c);d.stopCallback&&d.stopCallback(c)};d.playCallbackTimers.length&&d.clearPlayCallbackTimers();
d.stopTimeout&&clearTimeout(d.stopTimeout);a?(d.setGain(0),g()):(d.mute(),d.stopOscillatorTimeout=setTimeout(g,f.fadeOutDuration+100))};a.defaultOptions={type:"oscillator",playCallbackInterval:20,masterVolume:1,oscillator:{waveformShape:"sine"}};a.definitions={};return a}();["sine","square","triangle","sawtooth"].forEach(function(a){t.definitions[a]=new t({oscillator:{waveformShape:a}});t.definitions[a+"Musical"]=new t({allowedFrequencies:h.musicalFrequencies,oscillator:{waveformShape:a}});t.definitions[a+
"Major"]=new t({allowedFrequencies:h.getMusicalScale([1,3,5,6,8,10,12]),oscillator:{waveformShape:a}})});"";return t});l(a,"Extensions/Sonification/Earcon.js",[a["Extensions/Sonification/Instrument.js"],a["Core/Utilities.js"]],function(a,f){var h=f.error,k=f.merge,n=f.pick,e=f.uniqueKey;f=function(){function c(a){this.options=this.instrumentsPlaying=this.id=void 0;this.init(a||{})}c.prototype.init=function(a){this.options=a;this.options.id||(this.options.id=this.id=e());this.instrumentsPlaying={}};
c.prototype.sonify=function(c){var b=k(this.options,c),e=n(b.volume,1),f=b.pan,r=this,d=c&&c.onEnd,m=r.options.onEnd;b.instruments.forEach(function(c){var b="string"===typeof c.instrument?a.definitions[c.instrument]:c.instrument,g=k(c.playOptions),q="";if(b&&b.play){if(c.playOptions){g.pan=n(f,g.pan);var t=g.onEnd;g.onEnd=function(){delete r.instrumentsPlaying[q];t&&t.apply(this,arguments);Object.keys(r.instrumentsPlaying).length||(d&&d.apply(this,arguments),m&&m.apply(this,arguments))};c=b.copy();
c.setMasterVolume(e);q=c.id;r.instrumentsPlaying[q]=c;c.play(g)}}else h(30)})};c.prototype.cancelSonify=function(a){var c=this.instrumentsPlaying,b=c&&Object.keys(c);b&&b.length&&(b.forEach(function(b){c[b].stop(!a,null,"cancelled")}),this.instrumentsPlaying={})};return c}();"";return f});l(a,"Extensions/Sonification/Timeline.js",[a["Extensions/Sonification/Sonification.js"],a["Core/Utilities.js"],a["Extensions/Sonification/SonificationUtilities.js"]],function(a,f,h){var k=f.merge,n=f.splat;f=function(){function e(a){this.signalHandler=
this.pathsPlaying=this.paths=this.options=this.cursor=void 0;this.init(a||{})}e.prototype.init=function(a){this.options=a;this.cursor=0;this.paths=a.paths||[];this.pathsPlaying={};this.signalHandler=new h.SignalHandler(["playOnEnd","masterOnEnd","onPathStart","onPathEnd"]);this.signalHandler.registerSignalCallbacks(k(a,{masterOnEnd:a.onEnd}))};e.prototype.play=function(a){this.pause();this.signalHandler.clearSignalCallbacks(["playOnEnd"]);this.signalHandler.registerSignalCallbacks({playOnEnd:a});
this.playPaths(1)};e.prototype.rewind=function(a){this.pause();this.signalHandler.clearSignalCallbacks(["playOnEnd"]);this.signalHandler.registerSignalCallbacks({playOnEnd:a});this.playPaths(-1)};e.prototype.playPaths=function(c){var b=this,g=b.signalHandler;if(b.paths.length){var e=n(this.paths[this.cursor]),f=this.paths[this.cursor+c],k=function(a){g.emitSignal("onPathStart",a);b.pathsPlaying[a.id]=a;a[0<c?"play":"rewind"](function(d){d=d&&d.cancelled;var m={path:a,cancelled:d};delete b.pathsPlaying[a.id];
g.emitSignal("onPathEnd",m);var k=0;k++;k>=e.length&&(f&&!d?(b.cursor+=c,n(f).forEach(function(a){a[0<c?"resetCursor":"resetCursorEnd"]()}),b.playPaths(c)):(g.emitSignal("playOnEnd",m),g.emitSignal("masterOnEnd",m)))})};e.forEach(function(c){c&&(c.timeline=b,setTimeout(function(){k(c)},a.fadeOutDuration))})}else{var d={cancelled:!1};g.emitSignal("playOnEnd",d);g.emitSignal("masterOnEnd",d)}};e.prototype.pause=function(a){var c=this;Object.keys(c.pathsPlaying).forEach(function(b){c.pathsPlaying[b]&&
c.pathsPlaying[b].pause(a)});c.pathsPlaying={}};e.prototype.resetCursor=function(){this.paths.forEach(function(a){n(a).forEach(function(a){a.resetCursor()})});this.cursor=0};e.prototype.resetCursorEnd=function(){this.paths.forEach(function(a){n(a).forEach(function(a){a.resetCursorEnd()})});this.cursor=this.paths.length-1};e.prototype.setCursor=function(a){return this.paths.some(function(c){return n(c).some(function(c){return c.setCursor(a)})})};e.prototype.getCursor=function(){return this.getCurrentPlayingPaths().reduce(function(a,
b){a[b.id]=b.getCursor();return a},{})};e.prototype.atStart=function(){return this.cursor?!1:!n(this.paths[0]).some(function(a){return a.cursor})};e.prototype.getCurrentPlayingPaths=function(){return this.paths.length?n(this.paths[this.cursor]):[]};return e}();"";return f});l(a,"Extensions/Sonification/TimelineEvent.js",[a["Core/Utilities.js"]],function(a){var f=a.merge,h=a.uniqueKey;a=function(){function a(a){this.time=this.options=this.id=void 0;this.init(a)}a.prototype.init=function(a){this.options=
a;this.time=a.time||0;this.id=this.options.id=a.id||h()};a.prototype.play=function(a){var e=this.options.eventObject,c=this.options.onEnd,b=a&&a.onEnd,g=this.options.playOptions&&this.options.playOptions.onEnd;a=f(this.options.playOptions,a);e&&e.sonify?(a.onEnd=c||b||g?function(){var a=arguments;[c,b,g].forEach(function(c){c&&c.apply(this,a)})}:void 0,e.sonify(a)):(b&&b(),c&&c())};a.prototype.cancel=function(a){var e=this.options.eventObject;e&&e.cancelSonify(a)};return a}();"";return a});l(a,"Extensions/Sonification/TimelinePath.js",
[a["Extensions/Sonification/TimelineEvent.js"],a["Extensions/Sonification/SonificationUtilities.js"],a["Core/Utilities.js"]],function(a,f,h){var k=h.merge,n=h.uniqueKey;h=function(){function e(a){this.signalHandler=this.options=this.id=this.eventsPlaying=this.eventIdMap=this.events=this.cursor=void 0;this.init(a)}e.prototype.init=function(c){this.options=c;this.id=this.options.id=c.id||n();this.cursor=0;this.eventsPlaying={};this.events=c.silentWait?[new a({time:0}),new a({time:c.silentWait})]:this.options.events;
this.targetDuration=c.targetDuration||c.silentWait;this.sortEvents();this.updateEventIdMap();this.signalHandler=new f.SignalHandler(["playOnEnd","masterOnEnd","onStart","onEventStart","onEventEnd"]);this.signalHandler.registerSignalCallbacks(k(c,{masterOnEnd:c.onEnd}))};e.prototype.sortEvents=function(){this.events=this.events.sort(function(a,b){return a.time-b.time})};e.prototype.updateEventIdMap=function(){this.eventIdMap=this.events.reduce(function(a,b,e){a[b.id]=e;return a},{})};e.prototype.addTimelineEvents=
function(a){this.events=this.events.concat(a);this.sortEvents();this.updateEventIdMap()};e.prototype.getCursor=function(){return this.events[this.cursor]};e.prototype.setCursor=function(a){a=this.eventIdMap[a];return"undefined"!==typeof a?(this.cursor=a,!0):!1};e.prototype.play=function(a){this.pause();this.signalHandler.emitSignal("onStart");this.signalHandler.clearSignalCallbacks(["playOnEnd"]);this.signalHandler.registerSignalCallbacks({playOnEnd:a});this.playEvents(1)};e.prototype.rewind=function(a){this.pause();
this.signalHandler.emitSignal("onStart");this.signalHandler.clearSignalCallbacks(["playOnEnd"]);this.signalHandler.registerSignalCallbacks({playOnEnd:a});this.playEvents(-1)};e.prototype.resetCursor=function(){this.cursor=0};e.prototype.resetCursorEnd=function(){this.cursor=this.events.length-1};e.prototype.pause=function(a){var b=this;clearTimeout(b.nextScheduledPlay);Object.keys(b.eventsPlaying).forEach(function(c){b.eventsPlaying[c]&&b.eventsPlaying[c].cancel(a)});b.eventsPlaying={}};e.prototype.playEvents=
function(a){var b=this,c=b.events[this.cursor],e=b.events[this.cursor+a],f=function(a){b.signalHandler.emitSignal("masterOnEnd",a);b.signalHandler.emitSignal("playOnEnd",a)};c.timelinePath=b;if(!1===b.signalHandler.emitSignal("onEventStart",c))f({event:c,cancelled:!0});else if(b.eventsPlaying[c.id]=c,c.play({onEnd:function(a){a={event:c,cancelled:!!a};delete b.eventsPlaying[c.id];b.signalHandler.emitSignal("onEventEnd",a);e||f(a)}}),e){var k=Math.abs(e.time-c.time);1>k?(b.cursor+=a,b.playEvents(a)):
this.nextScheduledPlay=setTimeout(function(){b.cursor+=a;b.playEvents(a)},k)}};return e}();"";return h});l(a,"Extensions/Sonification/SeriesSonify.js",[a["Extensions/Sonification/Earcon.js"],a["Extensions/Sonification/Instrument.js"],a["Core/Series/Point.js"],a["Extensions/Sonification/SonificationUtilities.js"],a["Extensions/Sonification/Timeline.js"],a["Extensions/Sonification/TimelineEvent.js"],a["Extensions/Sonification/TimelinePath.js"],a["Core/Utilities.js"]],function(a,f,h,k,n,e,c,b){var g=
k.getExtremesForInstrumentProps,t=k.virtualAxisTranslate,q=b.extend,r=b.find,d=b.isArray,m=b.merge,w=b.objectEach,l=b.pick,u;(function(b){function k(a,b){a.forEach(function(a){a=a.instrument;"string"!==typeof a&&a.setMasterVolume(b)});return a}function y(a,b){var d=b.timeExtremes||G(a,b.pointPlayTime),p=g(a.chart,b.instruments,b.dataExtremes),f=u(a,b.instruments,p),m=l(b.masterVolume,1),A=E(b.instruments),v=k(A,m);A=a.points.reduce(function(a,c){var N=D(c,b.earcons||[]),g=t(x(c,b.pointPlayTime),d,
{min:0,max:Math.max(b.duration-f,10)});return a.concat(new e({eventObject:c,time:g,id:c.id,playOptions:{instruments:v,dataExtremes:p,masterVolume:m}}),N.map(function(a){return new e({eventObject:a,time:g,playOptions:{volume:m}})}))},[]);return new c({events:A,onStart:function(){if(b.onStart)b.onStart(a)},onEventStart:function(a){var c=a.options&&a.options.eventObject;if(c instanceof h){if(!c.series.visible&&!c.series.chart.series.some(function(a){return a.visible}))return a.timelinePath.timeline.pause(),
a.timelinePath.timeline.resetCursor(),!1;if(b.onPointStart)b.onPointStart(a,c)}},onEventEnd:function(a){var c=a.event&&a.event.options&&a.event.options.eventObject;if(c instanceof h&&b.onPointEnd)b.onPointEnd(a.event,c)},onEnd:function(){if(b.onEnd)b.onEnd(a)},targetDuration:b.duration})}function z(a){var b=a.options.sonification||{},c=a.chart.options.sonification||{},d=c.events||{},e=b.events||{};return{onEnd:e.onSeriesEnd||d.onSeriesEnd,onStart:e.onSeriesStart||d.onSeriesStart,onPointEnd:e.onPointEnd||
d.onPointEnd,onPointStart:e.onPointStart||d.onPointStart,pointPlayTime:c.defaultInstrumentOptions&&c.defaultInstrumentOptions.mapping&&c.defaultInstrumentOptions.mapping.pointPlayTime,masterVolume:c.masterVolume,instruments:C(a),earcons:b.earcons||c.earcons}}function u(a,b,c){var d=a.points[a.points.length-1];return b.reduce(function(a,b){b=b.instrumentMapping.duration;b="string"===typeof b?0:"function"===typeof b?b(d,c):b;return Math.max(a,b)},0)}function D(b,c){return c.reduce(function(c,d){var p=
d.earcon;d.condition?(d=d.condition(b),d instanceof a?c.push(d):d&&c.push(p)):d.onPoint&&b.id===d.onPoint&&c.push(p);return c},[])}function x(a,b){return"function"===typeof b?b(a):l(a[b],a.options[b])}function C(a,b){if(b&&b.instruments)return b.instruments;var c=a.chart.options.sonification&&a.chart.options.sonification.defaultInstrumentOptions||{},d=function(a){w(a,function(b,c){null===b&&delete a[c]})};return(a.options.sonification&&a.options.sonification.instruments||[{}]).map(function(a){d(a.mapping||
{});d(a);return{instrument:a.instrument||c.instrument,instrumentOptions:m(c,a,{mapping:void 0,instrument:void 0}),instrumentMapping:m(c.mapping,a.mapping)}})}function G(a,b){return a.points.reduce(function(a,c){c=x(c,b);a.min=Math.min(a.min,c);a.max=Math.max(a.max,c);return a},{min:Infinity,max:-Infinity})}function E(a){return a.map(function(a){var b=a.instrument;b=("string"===typeof b?f.definitions[b]:b).copy();return m(a,{instrument:b})})}function F(a){var b=this.chart.options.sonification,c=this.options.sonification;
a=m({duration:c&&c.duration||b&&b.duration},z(this),a);b=y(this,a);if(c=this.chart.sonification)c.timeline&&c.timeline.pause(),c.duration=a.duration,c.timeline=new n({paths:[b]}),c.timeline.play()}var B=[];b.compose=function(a){-1===B.indexOf(a)&&(B.push(a),q(a.prototype,{sonify:F}));return a};b.buildChartSonifySeriesOptions=function(a,b,c){var e=c.seriesOptions||{},f=a.chart.options.sonification&&a.chart.options.sonification.defaultInstrumentOptions&&a.chart.options.sonification.defaultInstrumentOptions.mapping&&
a.chart.options.sonification.defaultInstrumentOptions.mapping.pointPlayTime||"x",g=z(a);return m(g,{dataExtremes:b,timeExtremes:G(a,f),instruments:c.instruments||g.instruments,onStart:c.onSeriesStart||g.onStart,onEnd:c.onSeriesEnd||g.onEnd,earcons:c.earcons||g.earcons,masterVolume:l(c.masterVolume,g.masterVolume)},d(e)?r(e,function(b){return b.id===l(a.id,a.options.id)})||{}:e,{pointPlayTime:f})};b.buildTimelinePathFromSeries=y})(u||(u={}));return u});l(a,"Extensions/Sonification/ChartSonify.js",
[a["Extensions/Sonification/Earcon.js"],a["Core/Series/Point.js"],a["Extensions/Sonification/SeriesSonify.js"],a["Extensions/Sonification/SonificationUtilities.js"],a["Extensions/Sonification/Timeline.js"],a["Extensions/Sonification/TimelineEvent.js"],a["Extensions/Sonification/TimelinePath.js"],a["Core/Utilities.js"]],function(a,f,h,k,n,e,c,b){function g(b,d,f){if("sequential"===b||"simultaneous"===b){var p=d.series.reduce(function(a,b){b.visible&&!1!==(b.options.sonification&&b.options.sonification.enabled)&&
a.push({series:b,seriesOptions:f(b)});return a},[]);"simultaneous"===b&&(p=[p])}else p=b.reduce(function(b,p){p=v(p).reduce(function(b,p){var g;if("string"===typeof p){var m=d.get(p);m.visible&&(g={series:m,seriesOptions:f(m)})}else p instanceof a&&(g=new c({events:[new e({eventObject:p})]}));p.silentWait&&(g=new c({silentWait:p.silentWait}));g&&b.push(g);return b},[]);p.length&&b.push(p);return b},[]);return p}function l(a,b){return b?a.reduce(function(d,e,p){e=v(e);d.push(e);p<a.length-1&&e.some(function(a){return a.series})&&
d.push(new c({silentWait:b}));return d},[]):a}function q(a){return a.reduce(function(a,b){b=v(b);return a+(1===b.length&&b[0].options&&b[0].options.silentWait||0)},0)}function r(a){var b=a.reduce(function(a,b){(b=b.events)&&b.length&&(a.min=Math.min(b[0].time,a.min),a.max=Math.max(b[b.length-1].time,a.max));return a},{min:Infinity,max:-Infinity});a.forEach(function(a){var c=a.events,d=c&&c.length,p=[];d&&c[0].time<=b.min||p.push(new e({time:b.min}));d&&c[c.length-1].time>=b.max||p.push(new e({time:b.max}));
p.length&&a.addTimelineEvents(p)})}function d(a){return a.reduce(function(a,b){return a+v(b).reduce(function(a,b){return(b=b.series&&b.seriesOptions&&b.seriesOptions.timeExtremes)?Math.max(a,b.max-b.min):a},0)},0)}function m(a,b){var e=Math.max(b-q(a),0),p=d(a);return a.reduce(function(a,b){b=v(b).reduce(function(a,b){b instanceof c?a.push(b):b.series&&(b.seriesOptions.duration=b.seriesOptions.duration||M(b.seriesOptions.timeExtremes.max-b.seriesOptions.timeExtremes.min,{min:0,max:p},{min:0,max:e}),
a.push(h.buildTimelinePathFromSeries(b.series,b.seriesOptions)));return a},[]);a.push(b);return a},[])}function w(a,b){a=a.options.sonification||{};return B({duration:a.duration,afterSeriesWait:a.afterSeriesWait,pointPlayTime:a.defaultInstrumentOptions&&a.defaultInstrumentOptions.mapping&&a.defaultInstrumentOptions.mapping.pointPlayTime,order:a.order,onSeriesStart:a.events&&a.events.onSeriesStart,onSeriesEnd:a.events&&a.events.onSeriesEnd,onEnd:a.events&&a.events.onEnd},b)}function x(a){var b=w(this,
a);this.sonification.timeline&&this.sonification.timeline.pause();this.sonification.duration=b.duration;var c=L(this,b.instruments,b.dataExtremes);a=g(b.order,this,function(a){return h.buildChartSonifySeriesOptions(a,c,b)});a=l(a,b.afterSeriesWait||0);a=m(a,b.duration);a.forEach(function(a){r(a)});this.sonification.timeline=new n({paths:a,onEnd:b.onEnd});this.sonification.timeline.play()}function u(){if(this.sonification.timeline){var a=this.sonification.timeline.getCursor();return Object.keys(a).map(function(b){return a[b].options.eventObject}).filter(function(a){return a instanceof
f})}return[]}function C(a){var b=this.sonification.timeline;b&&v(a).forEach(function(a){b.setCursor(a.id)})}function H(a){this.sonification.timeline?this.sonification.timeline.pause(A(a,!0)):this.sonification.currentlyPlayingPoint&&this.sonification.currentlyPlayingPoint.cancelSonify(a)}function y(a){this.sonification.timeline&&this.sonification.timeline.play(a)}function z(a){this.sonification.timeline&&this.sonification.timeline.rewind(a)}function I(a){this.pauseSonify(a);this.resetSonifyCursor()}
function D(){this.sonification.timeline&&this.sonification.timeline.resetCursor()}function J(){this.sonification.timeline&&this.sonification.timeline.resetCursorEnd()}var L=k.getExtremesForInstrumentProps,M=k.virtualAxisTranslate,E=b.addEvent,F=b.extend,B=b.merge,A=b.pick,v=b.splat,K=[];k={chartSonify:x,compose:function(a){-1===K.indexOf(a)&&(K.push(a),F(a.prototype,{sonify:x,pauseSonify:H,resumeSonify:y,rewindSonify:z,cancelSonify:I,getCurrentSonifyPoints:u,setSonifyCursor:C,resetSonifyCursor:D,
resetSonifyCursorEnd:J}),E(a,"init",function(){this.sonification={}}),E(a,"update",function(a){(a=a.options.sonification)&&B(!0,this.options.sonification,a)}));return a},pause:H,resume:y,rewind:z,cancel:I,getCurrentPoints:u,setCursor:C,resetCursor:D,resetCursorEnd:J};"";return k});l(a,"Extensions/Sonification/PointSonify.js",[a["Extensions/Sonification/Instrument.js"],a["Core/Utilities.js"],a["Extensions/Sonification/SonificationUtilities.js"]],function(a,f,h){var k=f.error,n=f.merge,e=f.pick,c=[],
b={minDuration:20,maxDuration:2E3,minVolume:.1,maxVolume:1,minPan:-1,maxPan:1,minFrequency:220,maxFrequency:2200},g;(function(f){function g(c){var d=this,f=d.series.chart,g=e(c.masterVolume,f.options.sonification&&f.options.sonification.masterVolume),l=c.dataExtremes||{},q=function(a,b,c){if("function"===typeof a)return b?function(b){return a(d,l,b)}:a(d,l);if("string"===typeof a){var f=(b="-"===a.charAt(0))?a.slice(1):a,g=e(d[f],d.options[f]);l[f]=l[f]||h.calculateDataExtremes(d.series.chart,f);
return h.virtualAxisTranslate(g,l[f],c,b)}return a};f.sonification.currentlyPlayingPoint=d;d.sonification=d.sonification||{};d.sonification.instrumentsPlaying=d.sonification.instrumentsPlaying||{};var r=d.sonification.signalHandler=d.sonification.signalHandler||new h.SignalHandler(["onEnd"]);r.clearSignalCallbacks();r.registerSignalCallbacks({onEnd:c.onEnd});!d.isNull&&d.visible&&d.series.visible?c.instruments.forEach(function(c){var e="string"===typeof c.instrument?a.definitions[c.instrument]:c.instrument,
m=c.instrumentMapping||{},h=n(b,c.instrumentOptions),l=e.id,t=function(a){c.onEnd&&c.onEnd.apply(this,arguments);f.sonification&&f.sonification.currentlyPlayingPoint&&delete f.sonification.currentlyPlayingPoint;d.sonification&&d.sonification.instrumentsPlaying&&(delete d.sonification.instrumentsPlaying[l],Object.keys(d.sonification.instrumentsPlaying).length||r.emitSignal("onEnd",a))};e&&e.play?("undefined"!==typeof g&&e.setMasterVolume(g),d.sonification.instrumentsPlaying[e.id]=e,e.play({frequency:q(m.frequency,
!0,{min:h.minFrequency,max:h.maxFrequency}),duration:q(m.duration,!1,{min:h.minDuration,max:h.maxDuration}),pan:q(m.pan,!0,{min:h.minPan,max:h.maxPan}),volume:q(m.volume,!0,{min:h.minVolume,max:h.maxVolume}),onEnd:t,minFrequency:h.minFrequency,maxFrequency:h.maxFrequency})):k(30)}):r.emitSignal("onEnd")}function l(a){var b=this.sonification&&this.sonification.instrumentsPlaying,c=b&&Object.keys(b);c&&c.length&&(c.forEach(function(c){b[c].stop(!a,null,"cancelled")}),this.sonification.instrumentsPlaying=
{},this.sonification.signalHandler.emitSignal("onEnd","cancelled"))}f.compose=function(a){if(-1===c.indexOf(a)){c.push(a);var b=a.prototype;b.sonify=g;b.cancelSonify=l}return a}})(g||(g={}));"";return g});l(a,"masters/modules/sonification.src.js",[a["Core/Globals.js"],a["Extensions/Sonification/ChartSonify.js"],a["Extensions/Sonification/Earcon.js"],a["Extensions/Sonification/Instrument.js"],a["Extensions/Sonification/PointSonify.js"],a["Extensions/Sonification/SeriesSonify.js"],a["Extensions/Sonification/Sonification.js"],
a["Extensions/Sonification/Timeline.js"],a["Extensions/Sonification/TimelineEvent.js"],a["Extensions/Sonification/TimelinePath.js"]],function(a,f,h,k,l,e,c,b,g,t){var n=this&&this.__assign||function(){n=Object.assign||function(a){for(var b,c=1,e=arguments.length;c<e;c++){b=arguments[c];for(var f in b)Object.prototype.hasOwnProperty.call(b,f)&&(a[f]=b[f])}return a};return n.apply(this,arguments)};a.sonification=n(n({},c),{instruments:k.definitions,Earcon:h,Instrument:k,Timeline:b,TimelineEvent:g,TimelinePath:t});
a.Earcon=h;a.Instrument=k;f.compose(a.Chart);e.compose(a.Series);l.compose(a.Point)})});
//# sourceMappingURL=sonification.js.map
```
