## Supplementary material for "Exploring natural odour landscapes: A case study with implications for human-biting insects": Figure S4: static-scale.js

```
/*
 Highcharts Gantt JS v9.3.1 (2021-11-05)

 StaticScale

 (c) 2016-2021 Torstein Honsi, Lars A. V. Cabrera

 License: www.highcharts.com/license
*/
'use strict';(function(a){"object"===typeof module&&module.exports?(a["default"]=a,module.exports=a):"function"===typeof define&&define.amd?define("highcharts/modules/static-scale",["highcharts"],function(b){a(b);a.Highcharts=b;return a}):a("undefined"!==typeof Highcharts?Highcharts:void 0)})(function(a){function b(a,e,c,b){a.hasOwnProperty(e)||(a[e]=b.apply(null,c))}a=a?a._modules:{};b(a,"Extensions/StaticScale.js",[a["Core/Axis/Axis.js"],a["Core/Chart/Chart.js"],a["Core/Utilities.js"]],function(a,
b,c){var e=c.addEvent,f=c.defined,g=c.isNumber,h=c.pick;e(a,"afterSetOptions",function(){var a=this.chart.options.chart;!this.horiz&&g(this.options.staticScale)&&(!a.height||a.scrollablePlotArea&&a.scrollablePlotArea.minHeight)&&(this.staticScale=this.options.staticScale)});b.prototype.adjustHeight=function(){"adjustHeight"!==this.redrawTrigger&&((this.axes||[]).forEach(function(a){var d=a.chart,b=!!d.initiatedScale&&d.options.animation,c=a.options.staticScale;if(a.staticScale&&f(a.min)){var e=h(a.brokenAxis&&
a.brokenAxis.unitLength,a.max+a.tickInterval-a.min)*c;e=Math.max(e,c);c=e-d.plotHeight;!d.scrollablePixelsY&&1<=Math.abs(c)&&(d.plotHeight=e,d.redrawTrigger="adjustHeight",d.setSize(void 0,d.chartHeight+c,b));a.series.forEach(function(a){(a=a.sharedClipKey&&d.sharedClips[a.sharedClipKey])&&a.attr(d.inverted?{width:d.plotHeight}:{height:d.plotHeight})})}}),this.initiatedScale=!0);this.redrawTrigger=null};e(b,"render",b.prototype.adjustHeight)});b(a,"masters/modules/static-scale.src.js",[],function(){})});
//# sourceMappingURL=static-scale.js.map
```
