## Supplementary material for "Exploring natural odour landscapes: A case study with implications for human-biting insects": Figure S4: stock-tools.js

```
/*
 Highstock JS v9.3.1 (2021-11-05)

 Advanced Highcharts Stock tools

 (c) 2010-2021 Highsoft AS
 Author: Torstein Honsi

 License: www.highcharts.com/license
*/
'use strict';(function(a){"object"===typeof module&&module.exports?(a["default"]=a,module.exports=a):"function"===typeof define&&define.amd?define("highcharts/modules/stock-tools",["highcharts","highcharts/modules/stock"],function(w){a(w);a.Highcharts=w;return a}):a("undefined"!==typeof Highcharts?Highcharts:void 0)})(function(a){function w(c,a,r,v){c.hasOwnProperty(a)||(c[a]=v.apply(null,r))}a=a?a._modules:{};w(a,"Extensions/Annotations/Mixins/EventEmitterMixin.js",[a["Core/Globals.js"],a["Core/Utilities.js"]],
function(c,a){var n=a.addEvent,v=a.fireEvent,m=a.objectEach,p=a.pick,b=a.removeEvent;return{addEvents:function(){var b=this,e=function(e){n(e,c.isTouchDevice?"touchstart":"mousedown",function(e){b.onMouseDown(e)},{passive:!1})};e(this.graphic.element);(b.labels||[]).forEach(function(b){b.options.useHTML&&b.graphic.text&&e(b.graphic.text.element)});m(b.options.events,function(e,g){var c=function(c){"click"===g&&b.cancelClick||e.call(b,b.chart.pointer.normalize(c),b.target)};if(-1===(b.nonDOMEvents||
[]).indexOf(g))b.graphic.on(g,c);else n(b,g,c,{passive:!1})});if(b.options.draggable&&(n(b,"drag",b.onDrag),!b.graphic.renderer.styledMode)){var g={cursor:{x:"ew-resize",y:"ns-resize",xy:"move"}[b.options.draggable]};b.graphic.css(g);(b.labels||[]).forEach(function(b){b.options.useHTML&&b.graphic.text&&b.graphic.text.css(g)})}b.isUpdating||v(b,"add")},removeDocEvents:function(){this.removeDrag&&(this.removeDrag=this.removeDrag());this.removeMouseUp&&(this.removeMouseUp=this.removeMouseUp())},onMouseDown:function(b){var e=
this,g=e.chart.pointer;b.preventDefault&&b.preventDefault();if(2!==b.button){b=g.normalize(b);var t=b.chartX;var f=b.chartY;e.cancelClick=!1;e.chart.hasDraggedAnnotation=!0;e.removeDrag=n(c.doc,c.isTouchDevice?"touchmove":"mousemove",function(b){e.hasDragged=!0;b=g.normalize(b);b.prevChartX=t;b.prevChartY=f;v(e,"drag",b);t=b.chartX;f=b.chartY},c.isTouchDevice?{passive:!1}:void 0);e.removeMouseUp=n(c.doc,c.isTouchDevice?"touchend":"mouseup",function(b){var g=p(e.target&&e.target.annotation,e.target);
g&&(g.cancelClick=e.hasDragged);e.cancelClick=e.hasDragged;e.hasDragged=!1;e.chart.hasDraggedAnnotation=!1;v(p(g,e),"afterUpdate");e.onMouseUp(b)},c.isTouchDevice?{passive:!1}:void 0)}},onMouseUp:function(b){var e=this.chart;b=this.target||this;var g=e.options.annotations;e=e.annotations.indexOf(b);this.removeDocEvents();g[e]=b.options},onDrag:function(b){if(this.chart.isInsidePlot(b.chartX-this.chart.plotLeft,b.chartY-this.chart.plotTop,{visiblePlotOnly:!0})){var e=this.mouseMoveToTranslation(b);
"x"===this.options.draggable&&(e.y=0);"y"===this.options.draggable&&(e.x=0);this.points.length?this.translate(e.x,e.y):(this.shapes.forEach(function(b){b.translate(e.x,e.y)}),this.labels.forEach(function(b){b.translate(e.x,e.y)}));this.redraw(!1)}},mouseMoveToRadians:function(b,e,g){var c=b.prevChartY-g,f=b.prevChartX-e;g=b.chartY-g;b=b.chartX-e;this.chart.inverted&&(e=f,f=c,c=e,e=b,b=g,g=e);return Math.atan2(g,b)-Math.atan2(c,f)},mouseMoveToTranslation:function(b){var e=b.chartX-b.prevChartX;b=b.chartY-
b.prevChartY;if(this.chart.inverted){var g=b;b=e;e=g}return{x:e,y:b}},mouseMoveToScale:function(b,e,g){e=(b.chartX-e||1)/(b.prevChartX-e||1);b=(b.chartY-g||1)/(b.prevChartY-g||1);this.chart.inverted&&(g=b,b=e,e=g);return{x:e,y:b}},destroy:function(){this.removeDocEvents();b(this);this.hcEvents=null}}});w(a,"Extensions/Annotations/ControlPoint.js",[a["Core/Utilities.js"],a["Extensions/Annotations/Mixins/EventEmitterMixin.js"]],function(c,a){var n=c.merge,v=c.pick;return function(){function c(c,b,E,
e){this.addEvents=a.addEvents;this.graphic=void 0;this.mouseMoveToRadians=a.mouseMoveToRadians;this.mouseMoveToScale=a.mouseMoveToScale;this.mouseMoveToTranslation=a.mouseMoveToTranslation;this.onDrag=a.onDrag;this.onMouseDown=a.onMouseDown;this.onMouseUp=a.onMouseUp;this.removeDocEvents=a.removeDocEvents;this.nonDOMEvents=["drag"];this.chart=c;this.target=b;this.options=E;this.index=v(E.index,e)}c.prototype.setVisibility=function(c){this.graphic.attr("visibility",c?"visible":"hidden");this.options.visible=
c};c.prototype.render=function(){var c=this.chart,b=this.options;this.graphic=c.renderer.symbol(b.symbol,0,0,b.width,b.height).add(c.controlPointsGroup).css(b.style);this.setVisibility(b.visible);this.addEvents()};c.prototype.redraw=function(c){this.graphic[c?"animate":"attr"](this.options.positioner.call(this,this.target))};c.prototype.destroy=function(){a.destroy.call(this);this.graphic&&(this.graphic=this.graphic.destroy());this.options=this.target=this.chart=null};c.prototype.update=function(c){var b=
this.chart,a=this.target,e=this.index;c=n(!0,this.options,c);this.destroy();this.constructor(b,a,c,e);this.render(b.controlPointsGroup);this.redraw()};return c}()});w(a,"Extensions/Annotations/MockPoint.js",[a["Core/Series/Series.js"],a["Core/Utilities.js"],a["Core/Axis/Axis.js"]],function(c,a,r){var n=a.defined,m=a.fireEvent;return function(){function a(b,a,e){this.y=this.x=this.ttBelow=this.plotY=this.plotX=this.negative=this.isInside=void 0;this.mock=!0;this.series={visible:!0,chart:b,getPlotBox:c.prototype.getPlotBox};
this.target=a||null;this.options=e;this.applyOptions(this.getOptions())}a.fromPoint=function(b){return new a(b.series.chart,null,{x:b.x,y:b.y,xAxis:b.series.xAxis,yAxis:b.series.yAxis})};a.pointToPixels=function(b,c){var e=b.series,g=e.chart,a=b.plotX,f=b.plotY;g.inverted&&(b.mock?(a=b.plotY,f=b.plotX):(a=g.plotWidth-b.plotY,f=g.plotHeight-b.plotX));e&&!c&&(b=e.getPlotBox(),a+=b.translateX,f+=b.translateY);return{x:a,y:f}};a.pointToOptions=function(b){return{x:b.x,y:b.y,xAxis:b.series.xAxis,yAxis:b.series.yAxis}};
a.prototype.hasDynamicOptions=function(){return"function"===typeof this.options};a.prototype.getOptions=function(){return this.hasDynamicOptions()?this.options(this.target):this.options};a.prototype.applyOptions=function(b){this.command=b.command;this.setAxis(b,"x");this.setAxis(b,"y");this.refresh()};a.prototype.setAxis=function(b,c){c+="Axis";b=b[c];var e=this.series.chart;this.series[c]=b instanceof r?b:n(b)?e[c][b]||e.get(b):null};a.prototype.toAnchor=function(){var b=[this.plotX,this.plotY,0,
0];this.series.chart.inverted&&(b[0]=this.plotY,b[1]=this.plotX);return b};a.prototype.getLabelConfig=function(){return{x:this.x,y:this.y,point:this}};a.prototype.isInsidePlot=function(){var b=this.plotX,c=this.plotY,e=this.series.xAxis,g=this.series.yAxis,a={x:b,y:c,isInsidePlot:!0};e&&(a.isInsidePlot=n(b)&&0<=b&&b<=e.len);g&&(a.isInsidePlot=a.isInsidePlot&&n(c)&&0<=c&&c<=g.len);m(this.series.chart,"afterIsInsidePlot",a);return a.isInsidePlot};a.prototype.refresh=function(){var b=this.series,c=b.xAxis;
b=b.yAxis;var e=this.getOptions();c?(this.x=e.x,this.plotX=c.toPixels(e.x,!0)):(this.x=null,this.plotX=e.x);b?(this.y=e.y,this.plotY=b.toPixels(e.y,!0)):(this.y=null,this.plotY=e.y);this.isInside=this.isInsidePlot()};a.prototype.translate=function(b,c,e,g){this.hasDynamicOptions()||(this.plotX+=e,this.plotY+=g,this.refreshOptions())};a.prototype.scale=function(b,c,e,g){if(!this.hasDynamicOptions()){var a=this.plotY*g;this.plotX=(1-e)*b+this.plotX*e;this.plotY=(1-g)*c+a;this.refreshOptions()}};a.prototype.rotate=
function(b,c,e){if(!this.hasDynamicOptions()){var g=Math.cos(e);e=Math.sin(e);var a=this.plotX,f=this.plotY;a-=b;f-=c;this.plotX=a*g-f*e+b;this.plotY=a*e+f*g+c;this.refreshOptions()}};a.prototype.refreshOptions=function(){var b=this.series,c=b.xAxis;b=b.yAxis;this.x=this.options.x=c?this.options.x=c.toValue(this.plotX,!0):this.plotX;this.y=this.options.y=b?b.toValue(this.plotY,!0):this.plotY};return a}()});w(a,"Extensions/Annotations/Mixins/ControllableMixin.js",[a["Extensions/Annotations/ControlPoint.js"],
a["Extensions/Annotations/MockPoint.js"],a["Core/Tooltip.js"],a["Core/Utilities.js"]],function(c,a,r,v){var m=v.isObject,p=v.isString,b=v.merge,E=v.splat;return{init:function(b,c,a){this.annotation=b;this.chart=b.chart;this.options=c;this.points=[];this.controlPoints=[];this.index=a;this.linkPoints();this.addControlPoints()},attr:function(){this.graphic.attr.apply(this.graphic,arguments)},getPointsOptions:function(){var b=this.options;return b.points||b.point&&E(b.point)},attrsFromOptions:function(b){var c=
this.constructor.attrsMap,e={},a,u=this.chart.styledMode;for(a in b){var x=c[a];!x||u&&-1!==["fill","stroke","stroke-width"].indexOf(x)||(e[x]=b[a])}return e},anchor:function(c){var e=c.series.getPlotBox(),a=c.series.chart,f=c.mock?c.toAnchor():r.prototype.getAnchor.call({chart:c.series.chart},c);f={x:f[0]+(this.options.x||0),y:f[1]+(this.options.y||0),height:f[2]||0,width:f[3]||0};return{relativePosition:f,absolutePosition:b(f,{x:f.x+(c.mock?e.translateX:a.plotLeft),y:f.y+(c.mock?e.translateY:a.plotTop)})}},
point:function(b,c){if(b&&b.series)return b;c&&null!==c.series||(m(b)?c=new a(this.chart,this,b):p(b)?c=this.chart.get(b)||null:"function"===typeof b&&(c=b.call(c,this),c=c.series?c:new a(this.chart,this,b)));return c},linkPoints:function(){var b=this.getPointsOptions(),c=this.points,a=b&&b.length||0,f;for(f=0;f<a;f++){var u=this.point(b[f],c[f]);if(!u){c.length=0;return}u.mock&&u.refresh();c[f]=u}return c},addControlPoints:function(){var e=this.options.controlPoints;(e||[]).forEach(function(a,t){a=
b(this.options.controlPointOptions,a);a.index||(a.index=t);e[t]=a;this.controlPoints.push(new c(this.chart,this,a))},this)},shouldBeDrawn:function(){return!!this.points.length},render:function(b){this.controlPoints.forEach(function(b){b.render()})},redraw:function(b){this.controlPoints.forEach(function(c){c.redraw(b)})},transform:function(b,c,a,f,u){if(this.chart.inverted){var e=c;c=a;a=e}this.points.forEach(function(e,d){this.transformPoint(b,c,a,f,u,d)},this)},transformPoint:function(b,c,t,f,u,
x){var e=this.points[x];e.mock||(e=this.points[x]=a.fromPoint(e));e[b](c,t,f,u)},translate:function(b,c){this.transform("translate",null,null,b,c)},translatePoint:function(b,c,a){this.transformPoint("translate",null,null,b,c,a)},translateShape:function(b,c,a){var e=this.annotation.chart,g=this.annotation.userOptions,t=e.annotations.indexOf(this.annotation);e=e.options.annotations[t];this.translatePoint(b,c,0);a&&this.translatePoint(b,c,1);e[this.collection][this.index].point=this.options.point;g[this.collection][this.index].point=
this.options.point},rotate:function(b,c,a){this.transform("rotate",b,c,a)},scale:function(b,c,a,f){this.transform("scale",b,c,a,f)},setControlPointsVisibility:function(b){this.controlPoints.forEach(function(c){c.setVisibility(b)})},destroy:function(){this.graphic&&(this.graphic=this.graphic.destroy());this.tracker&&(this.tracker=this.tracker.destroy());this.controlPoints.forEach(function(b){b.destroy()});this.options=this.controlPoints=this.points=this.chart=null;this.annotation&&(this.annotation=
null)},update:function(c){var a=this.annotation;c=b(!0,this.options,c);var e=this.graphic.parentGroup;this.destroy();this.constructor(a,c,this.index);this.render(e);this.redraw()}}});w(a,"Extensions/Annotations/Mixins/MarkerMixin.js",[a["Core/Chart/Chart.js"],a["Core/Renderer/SVG/SVGRenderer.js"],a["Core/Utilities.js"]],function(c,a,r){function n(b){return function(c){this.attr(b,"url(#"+c+")")}}var m=r.addEvent,p=r.defined,b=r.merge,E=r.uniqueKey,e={arrow:{tagName:"marker",attributes:{id:"arrow",
refY:5,refX:9,markerWidth:10,markerHeight:10},children:[{tagName:"path",attributes:{d:"M 0 0 L 10 5 L 0 10 Z","stroke-width":0}}]},"reverse-arrow":{tagName:"marker",attributes:{id:"reverse-arrow",refY:5,refX:1,markerWidth:10,markerHeight:10},children:[{tagName:"path",attributes:{d:"M 0 5 L 10 0 L 10 10 Z","stroke-width":0}}]}};a.prototype.addMarker=function(c,a){var e={attributes:{id:c}},u={stroke:a.color||"none",fill:a.color||"rgba(0, 0, 0, 0.75)"};e.children=a.children&&a.children.map(function(c){return b(u,
c)});a=b(!0,{attributes:{markerWidth:20,markerHeight:20,refX:0,refY:0,orient:"auto"}},a,e);a=this.definition(a);a.id=c;return a};a={markerEndSetter:n("marker-end"),markerStartSetter:n("marker-start"),setItemMarkers:function(c){var a=c.options,e=c.chart,u=e.options.defs,g=a.fill,k=p(g)&&"none"!==g?g:a.stroke;["markerStart","markerEnd"].forEach(function(d){var h=a[d],l;if(h){for(l in u){var q=u[l];if((h===(q.attributes&&q.attributes.id)||h===q.id)&&"marker"===q.tagName){var f=q;break}}f&&(h=c[d]=e.renderer.addMarker((a.id||
E())+"-"+h,b(f,{color:k})),c.attr(d,h.getAttribute("id")))}})}};m(c,"afterGetContainer",function(){this.options.defs=b(e,this.options.defs||{})});return a});w(a,"Extensions/Annotations/Controllables/ControllablePath.js",[a["Extensions/Annotations/Mixins/ControllableMixin.js"],a["Core/Globals.js"],a["Extensions/Annotations/Mixins/MarkerMixin.js"],a["Core/Utilities.js"]],function(c,a,r,v){var m=v.extend,p="rgba(192,192,192,"+(a.svg?.0001:.002)+")";return function(){function b(b,a,g){this.addControlPoints=
c.addControlPoints;this.anchor=c.anchor;this.attr=c.attr;this.attrsFromOptions=c.attrsFromOptions;this.destroy=c.destroy;this.getPointsOptions=c.getPointsOptions;this.init=c.init;this.linkPoints=c.linkPoints;this.point=c.point;this.rotate=c.rotate;this.scale=c.scale;this.setControlPointsVisibility=c.setControlPointsVisibility;this.setMarkers=r.setItemMarkers;this.transform=c.transform;this.transformPoint=c.transformPoint;this.translate=c.translate;this.translatePoint=c.translatePoint;this.translateShape=
c.translateShape;this.update=c.update;this.type="path";this.init(b,a,g);this.collection="shapes"}b.prototype.toD=function(){var b=this.options.d;if(b)return"function"===typeof b?b.call(this):b;b=this.points;var c=b.length,a=c,p=b[0],f=a&&this.anchor(p).absolutePosition,u=0,x=[];if(f)for(x.push(["M",f.x,f.y]);++u<c&&a;)p=b[u],a=p.command||"L",f=this.anchor(p).absolutePosition,"M"===a?x.push([a,f.x,f.y]):"L"===a?x.push([a,f.x,f.y]):"Z"===a&&x.push([a]),a=p.series.visible;return a?this.chart.renderer.crispLine(x,
this.graphic.strokeWidth()):null};b.prototype.shouldBeDrawn=function(){return c.shouldBeDrawn.call(this)||!!this.options.d};b.prototype.render=function(b){var a=this.options,g=this.attrsFromOptions(a);this.graphic=this.annotation.chart.renderer.path([["M",0,0]]).attr(g).add(b);a.className&&this.graphic.addClass(a.className);this.tracker=this.annotation.chart.renderer.path([["M",0,0]]).addClass("highcharts-tracker-line").attr({zIndex:2}).add(b);this.annotation.chart.styledMode||this.tracker.attr({"stroke-linejoin":"round",
stroke:p,fill:p,"stroke-width":this.graphic.strokeWidth()+2*a.snap});c.render.call(this);m(this.graphic,{markerStartSetter:r.markerStartSetter,markerEndSetter:r.markerEndSetter});this.setMarkers(this)};b.prototype.redraw=function(b){var a=this.toD(),g=b?"animate":"attr";a?(this.graphic[g]({d:a}),this.tracker[g]({d:a})):(this.graphic.attr({d:"M 0 -9000000000"}),this.tracker.attr({d:"M 0 -9000000000"}));this.graphic.placed=this.tracker.placed=!!a;c.redraw.call(this,b)};b.attrsMap={dashStyle:"dashstyle",
strokeWidth:"stroke-width",stroke:"stroke",fill:"fill",zIndex:"zIndex"};return b}()});w(a,"Extensions/Annotations/Controllables/ControllableRect.js",[a["Extensions/Annotations/Mixins/ControllableMixin.js"],a["Extensions/Annotations/Controllables/ControllablePath.js"],a["Core/Utilities.js"]],function(c,a,r){var n=r.merge;return function(){function m(a,b,m){this.addControlPoints=c.addControlPoints;this.anchor=c.anchor;this.attr=c.attr;this.attrsFromOptions=c.attrsFromOptions;this.destroy=c.destroy;
this.getPointsOptions=c.getPointsOptions;this.init=c.init;this.linkPoints=c.linkPoints;this.point=c.point;this.rotate=c.rotate;this.scale=c.scale;this.setControlPointsVisibility=c.setControlPointsVisibility;this.shouldBeDrawn=c.shouldBeDrawn;this.transform=c.transform;this.transformPoint=c.transformPoint;this.translatePoint=c.translatePoint;this.translateShape=c.translateShape;this.update=c.update;this.type="rect";this.translate=c.translateShape;this.init(a,b,m);this.collection="shapes"}m.prototype.render=
function(a){var b=this.attrsFromOptions(this.options);this.graphic=this.annotation.chart.renderer.rect(0,-9E9,0,0).attr(b).add(a);c.render.call(this)};m.prototype.redraw=function(a){var b=this.anchor(this.points[0]).absolutePosition;if(b)this.graphic[a?"animate":"attr"]({x:b.x,y:b.y,width:this.options.width,height:this.options.height});else this.attr({x:0,y:-9E9});this.graphic.placed=!!b;c.redraw.call(this,a)};m.attrsMap=n(a.attrsMap,{width:"width",height:"height"});return m}()});w(a,"Extensions/Annotations/Controllables/ControllableCircle.js",
[a["Extensions/Annotations/Mixins/ControllableMixin.js"],a["Extensions/Annotations/Controllables/ControllablePath.js"],a["Core/Utilities.js"]],function(c,a,r){var n=r.merge;return function(){function m(a,b,m){this.addControlPoints=c.addControlPoints;this.anchor=c.anchor;this.attr=c.attr;this.attrsFromOptions=c.attrsFromOptions;this.destroy=c.destroy;this.getPointsOptions=c.getPointsOptions;this.init=c.init;this.linkPoints=c.linkPoints;this.point=c.point;this.rotate=c.rotate;this.scale=c.scale;this.setControlPointsVisibility=
c.setControlPointsVisibility;this.shouldBeDrawn=c.shouldBeDrawn;this.transform=c.transform;this.transformPoint=c.transformPoint;this.translatePoint=c.translatePoint;this.translateShape=c.translateShape;this.update=c.update;this.type="circle";this.translate=c.translateShape;this.init(a,b,m);this.collection="shapes"}m.prototype.render=function(a){var b=this.attrsFromOptions(this.options);this.graphic=this.annotation.chart.renderer.circle(0,-9E9,0).attr(b).add(a);c.render.call(this)};m.prototype.redraw=
function(a){var b=this.anchor(this.points[0]).absolutePosition;if(b)this.graphic[a?"animate":"attr"]({x:b.x,y:b.y,r:this.options.r});else this.graphic.attr({x:0,y:-9E9});this.graphic.placed=!!b;c.redraw.call(this,a)};m.prototype.setRadius=function(c){this.options.r=c};m.attrsMap=n(a.attrsMap,{r:"r"});return m}()});w(a,"Extensions/Annotations/Controllables/ControllableEllipse.js",[a["Extensions/Annotations/Mixins/ControllableMixin.js"],a["Extensions/Annotations/Controllables/ControllablePath.js"],
a["Core/Utilities.js"]],function(c,a,r){var n=r.merge,m=r.defined;return function(){function p(b,a,e){this.addControlPoints=c.addControlPoints;this.anchor=c.anchor;this.attr=c.attr;this.attrsFromOptions=c.attrsFromOptions;this.destroy=c.destroy;this.getPointsOptions=c.getPointsOptions;this.linkPoints=c.linkPoints;this.point=c.point;this.scale=c.scale;this.setControlPointsVisibility=c.setControlPointsVisibility;this.shouldBeDrawn=c.shouldBeDrawn;this.transform=c.transform;this.translatePoint=c.translatePoint;
this.transformPoint=c.transformPoint;this.type="ellipse";this.init(b,a,e);this.collection="shapes"}p.prototype.init=function(b,a,e){m(a.yAxis)&&a.points.forEach(function(b){b.yAxis=a.yAxis});m(a.xAxis)&&a.points.forEach(function(b){b.xAxis=a.xAxis});c.init.call(this,b,a,e)};p.prototype.render=function(b){this.graphic=this.annotation.chart.renderer.createElement("ellipse").attr(this.attrsFromOptions(this.options)).add(b);c.render.call(this)};p.prototype.translate=function(b,a){c.translateShape.call(this,
b,a,!0)};p.prototype.getDistanceFromLine=function(b,c,a,g){return Math.abs((c.y-b.y)*a-(c.x-b.x)*g+c.x*b.y-c.y*b.x)/Math.sqrt((c.y-b.y)*(c.y-b.y)+(c.x-b.x)*(c.x-b.x))};p.prototype.getAttrs=function(b,c){var a=b.x,g=b.y,m=c.x,f=c.y;c=(a+m)/2;b=(g+f)/2;var u=Math.sqrt((a-m)*(a-m)/4+(g-f)*(g-f)/4);g=180*Math.atan((f-g)/(m-a))/Math.PI;c<a&&(g+=180);a=this.getRY();return{cx:c,cy:b,rx:u,ry:a,angle:g}};p.prototype.getRY=function(){var b=this.getYAxis();return m(b)?Math.abs(b.toPixels(this.options.ry)-b.toPixels(0)):
this.options.ry};p.prototype.getYAxis=function(){return this.chart.yAxis[this.options.yAxis]};p.prototype.getAbsolutePosition=function(b){return this.anchor(b).absolutePosition};p.prototype.redraw=function(b){var a=this.getAbsolutePosition(this.points[0]),e=this.getAbsolutePosition(this.points[1]);e=this.getAttrs(a,e);if(a)this.graphic[b?"animate":"attr"]({cx:e.cx,cy:e.cy,rx:e.rx,ry:e.ry,rotation:e.angle,rotationOriginX:e.cx,rotationOriginY:e.cy});else this.graphic.attr({x:0,y:-9E9});this.graphic.placed=
!!a;c.redraw.call(this,b)};p.prototype.setYRadius=function(b){this.options.ry=b;this.annotation.userOptions.shapes[0].ry=b;this.annotation.options.shapes[0].ry=b};p.attrsMap=n(a.attrsMap,{ry:"ry"});return p}()});w(a,"Extensions/Annotations/Controllables/ControllableLabel.js",[a["Extensions/Annotations/Mixins/ControllableMixin.js"],a["Core/FormatUtilities.js"],a["Extensions/Annotations/MockPoint.js"],a["Core/Renderer/SVG/SVGRenderer.js"],a["Core/Tooltip.js"],a["Core/Utilities.js"]],function(c,a,r,
v,m,p){var b=a.format;a=v.prototype.symbols;var n=p.extend,e=p.isNumber,g=p.pick;p=function(){function a(b,a,e){this.addControlPoints=c.addControlPoints;this.attr=c.attr;this.attrsFromOptions=c.attrsFromOptions;this.destroy=c.destroy;this.getPointsOptions=c.getPointsOptions;this.init=c.init;this.linkPoints=c.linkPoints;this.point=c.point;this.rotate=c.rotate;this.scale=c.scale;this.setControlPointsVisibility=c.setControlPointsVisibility;this.shouldBeDrawn=c.shouldBeDrawn;this.transform=c.transform;
this.transformPoint=c.transformPoint;this.translateShape=c.translateShape;this.update=c.update;this.init(b,a,e);this.collection="labels"}a.alignedPosition=function(b,a){var c=b.align,e=b.verticalAlign,d=(a.x||0)+(b.x||0),h=(a.y||0)+(b.y||0),l,q;"right"===c?l=1:"center"===c&&(l=2);l&&(d+=(a.width-(b.width||0))/l);"bottom"===e?q=1:"middle"===e&&(q=2);q&&(h+=(a.height-(b.height||0))/q);return{x:Math.round(d),y:Math.round(h)}};a.justifiedOptions=function(b,a,c,e){var d=c.align,h=c.verticalAlign,l=a.box?
0:a.padding||0,q=a.getBBox();a={align:d,verticalAlign:h,x:c.x,y:c.y,width:a.width,height:a.height};c=(e.x||0)-b.plotLeft;e=(e.y||0)-b.plotTop;var f=c+l;0>f&&("right"===d?a.align="left":a.x=(a.x||0)-f);f=c+q.width-l;f>b.plotWidth&&("left"===d?a.align="right":a.x=(a.x||0)+b.plotWidth-f);f=e+l;0>f&&("bottom"===h?a.verticalAlign="top":a.y=(a.y||0)-f);f=e+q.height-l;f>b.plotHeight&&("top"===h?a.verticalAlign="bottom":a.y=(a.y||0)+b.plotHeight-f);return a};a.prototype.translatePoint=function(b,a){c.translatePoint.call(this,
b,a,0)};a.prototype.translate=function(b,a){var c=this.annotation.chart,e=this.annotation.userOptions,d=c.annotations.indexOf(this.annotation);d=c.options.annotations[d];c.inverted&&(c=b,b=a,a=c);this.options.x+=b;this.options.y+=a;d[this.collection][this.index].x=this.options.x;d[this.collection][this.index].y=this.options.y;e[this.collection][this.index].x=this.options.x;e[this.collection][this.index].y=this.options.y};a.prototype.render=function(b){var e=this.options,f=this.attrsFromOptions(e),
k=e.style;this.graphic=this.annotation.chart.renderer.label("",0,-9999,e.shape,null,null,e.useHTML,null,"annotation-label").attr(f).add(b);this.annotation.chart.styledMode||("contrast"===k.color&&(k.color=this.annotation.chart.renderer.getContrast(-1<a.shapesWithoutBackground.indexOf(e.shape)?"#FFFFFF":e.backgroundColor)),this.graphic.css(e.style).shadow(e.shadow));e.className&&this.graphic.addClass(e.className);this.graphic.labelrank=e.labelrank;c.render.call(this)};a.prototype.redraw=function(a){var e=
this.options,f=this.text||e.format||e.text,k=this.graphic,d=this.points[0];k.attr({text:f?b(f,d.getLabelConfig(),this.annotation.chart):e.formatter.call(d,this)});e=this.anchor(d);(f=this.position(e))?(k.alignAttr=f,f.anchorX=e.absolutePosition.x,f.anchorY=e.absolutePosition.y,k[a?"animate":"attr"](f)):k.attr({x:0,y:-9999});k.placed=!!f;c.redraw.call(this,a)};a.prototype.anchor=function(b){var a=c.anchor.apply(this,arguments),e=this.options.x||0,f=this.options.y||0;a.absolutePosition.x-=e;a.absolutePosition.y-=
f;a.relativePosition.x-=e;a.relativePosition.y-=f;return a};a.prototype.position=function(b){var c=this.graphic,e=this.annotation.chart,f=this.points[0],d=this.options,h=b.absolutePosition,l=b.relativePosition,q=f.series.visible&&r.prototype.isInsidePlot.call(f);b=c.width;b=void 0===b?0:b;var z=c.height;z=void 0===z?0:z;if(q){if(d.distance)var A=m.prototype.getPosition.call({chart:e,distance:g(d.distance,16)},b,z,{plotX:l.x,plotY:l.y,negative:f.negative,ttBelow:f.ttBelow,h:l.height||l.width});else d.positioner?
A=d.positioner.call(this):(f={x:h.x,y:h.y,width:0,height:0},A=a.alignedPosition(n(d,{width:b,height:z}),f),"justify"===this.options.overflow&&(A=a.alignedPosition(a.justifiedOptions(e,c,d,A),f)));d.crop&&(c=A.x-e.plotLeft,d=A.y-e.plotTop,q=e.isInsidePlot(c,d)&&e.isInsidePlot(c+b,d+z))}return q?A:null};a.attrsMap={backgroundColor:"fill",borderColor:"stroke",borderWidth:"stroke-width",zIndex:"zIndex",borderRadius:"r",padding:"padding"};a.shapesWithoutBackground=["connector"];return a}();a.connector=
function(b,a,c,g,k){var d=k&&k.anchorX;k=k&&k.anchorY;var h=c/2;if(e(d)&&e(k)){var l=[["M",d,k]];var q=a-k;0>q&&(q=-g-q);q<c&&(h=d<b+c/2?q:c-q);k>a+g?l.push(["L",b+h,a+g]):k<a?l.push(["L",b+h,a]):d<b?l.push(["L",b,a+g/2]):d>b+c&&l.push(["L",b+c,a+g/2])}return l||[]};return p});w(a,"Extensions/Annotations/Controllables/ControllableImage.js",[a["Extensions/Annotations/Controllables/ControllableLabel.js"],a["Extensions/Annotations/Mixins/ControllableMixin.js"]],function(a,n){return function(){function c(a,
c,p){this.addControlPoints=n.addControlPoints;this.anchor=n.anchor;this.attr=n.attr;this.attrsFromOptions=n.attrsFromOptions;this.destroy=n.destroy;this.getPointsOptions=n.getPointsOptions;this.init=n.init;this.linkPoints=n.linkPoints;this.point=n.point;this.rotate=n.rotate;this.scale=n.scale;this.setControlPointsVisibility=n.setControlPointsVisibility;this.shouldBeDrawn=n.shouldBeDrawn;this.transform=n.transform;this.transformPoint=n.transformPoint;this.translatePoint=n.translatePoint;this.translateShape=
n.translateShape;this.update=n.update;this.type="image";this.translate=n.translateShape;this.init(a,c,p);this.collection="shapes"}c.prototype.render=function(a){var c=this.attrsFromOptions(this.options),p=this.options;this.graphic=this.annotation.chart.renderer.image(p.src,0,-9E9,p.width,p.height).attr(c).add(a);this.graphic.width=p.width;this.graphic.height=p.height;n.render.call(this)};c.prototype.redraw=function(c){var m=this.anchor(this.points[0]);if(m=a.prototype.position.call(this,m))this.graphic[c?
"animate":"attr"]({x:m.x,y:m.y});else this.graphic.attr({x:0,y:-9E9});this.graphic.placed=!!m;n.redraw.call(this,c)};c.attrsMap={width:"width",height:"height",zIndex:"zIndex"};return c}()});w(a,"Extensions/Annotations/Annotations.js",[a["Core/Animation/AnimationUtilities.js"],a["Core/Chart/Chart.js"],a["Extensions/Annotations/Mixins/ControllableMixin.js"],a["Extensions/Annotations/Controllables/ControllableRect.js"],a["Extensions/Annotations/Controllables/ControllableCircle.js"],a["Extensions/Annotations/Controllables/ControllableEllipse.js"],
a["Extensions/Annotations/Controllables/ControllablePath.js"],a["Extensions/Annotations/Controllables/ControllableImage.js"],a["Extensions/Annotations/Controllables/ControllableLabel.js"],a["Extensions/Annotations/ControlPoint.js"],a["Extensions/Annotations/Mixins/EventEmitterMixin.js"],a["Core/Globals.js"],a["Extensions/Annotations/MockPoint.js"],a["Core/Pointer.js"],a["Core/Utilities.js"]],function(a,n,r,v,m,p,b,w,e,g,t,f,u,x,k){var d=a.getDeferredAnimation;a=n.prototype;var c=k.addEvent,l=k.defined,
q=k.destroyObjectProperties,z=k.erase,A=k.extend,F=k.find,D=k.fireEvent,B=k.merge,y=k.pick,G=k.splat;k=k.wrap;var C=function(){function a(a,d){this.annotation=void 0;this.coll="annotations";this.shapesGroup=this.labelsGroup=this.labelCollector=this.group=this.graphic=this.animationConfig=this.collection=void 0;this.chart=a;this.points=[];this.controlPoints=[];this.coll="annotations";this.labels=[];this.shapes=[];this.options=B(this.defaultOptions,d);this.userOptions=d;d=this.getLabelsAndShapesOptions(this.options,
d);this.options.labels=d.labels;this.options.shapes=d.shapes;this.init(a,this.options)}a.prototype.init=function(){var a=this.chart,b=this.options.animation;this.linkPoints();this.addControlPoints();this.addShapes();this.addLabels();this.setLabelCollector();this.animationConfig=d(a,b)};a.prototype.getLabelsAndShapesOptions=function(d,a){var b={};["labels","shapes"].forEach(function(c){d[c]&&(b[c]=a[c]?G(a[c]).map(function(a,b){return B(d[c][b],a)}):d[c])});return b};a.prototype.addShapes=function(){(this.options.shapes||
[]).forEach(function(a,d){a=this.initShape(a,d);B(!0,this.options.shapes[d],a.options)},this)};a.prototype.addLabels=function(){(this.options.labels||[]).forEach(function(a,d){a=this.initLabel(a,d);B(!0,this.options.labels[d],a.options)},this)};a.prototype.addClipPaths=function(){this.setClipAxes();this.clipXAxis&&this.clipYAxis&&this.options.crop&&(this.clipRect=this.chart.renderer.clipRect(this.getClipBox()))};a.prototype.setClipAxes=function(){var a=this.chart.xAxis,d=this.chart.yAxis,b=(this.options.labels||
[]).concat(this.options.shapes||[]).reduce(function(b,c){c=c&&(c.point||c.points&&c.points[0]);return[a[c&&c.xAxis]||b[0],d[c&&c.yAxis]||b[1]]},[]);this.clipXAxis=b[0];this.clipYAxis=b[1]};a.prototype.getClipBox=function(){if(this.clipXAxis&&this.clipYAxis)return{x:this.clipXAxis.left,y:this.clipYAxis.top,width:this.clipXAxis.width,height:this.clipYAxis.height}};a.prototype.setLabelCollector=function(){var a=this;a.labelCollector=function(){return a.labels.reduce(function(a,d){d.options.allowOverlap||
a.push(d.graphic);return a},[])};a.chart.labelCollectors.push(a.labelCollector)};a.prototype.setOptions=function(a){this.options=B(this.defaultOptions,a)};a.prototype.redraw=function(a){this.linkPoints();this.graphic||this.render();this.clipRect&&this.clipRect.animate(this.getClipBox());this.redrawItems(this.shapes,a);this.redrawItems(this.labels,a);r.redraw.call(this,a)};a.prototype.redrawItems=function(a,d){for(var b=a.length;b--;)this.redrawItem(a[b],d)};a.prototype.renderItems=function(a){for(var d=
a.length;d--;)this.renderItem(a[d])};a.prototype.render=function(){var a=this.chart.renderer;this.graphic=a.g("annotation").attr({opacity:0,zIndex:this.options.zIndex,visibility:this.options.visible?"visible":"hidden"}).add();this.shapesGroup=a.g("annotation-shapes").add(this.graphic);this.options.crop&&this.shapesGroup.clip(this.chart.plotBoxClip);this.labelsGroup=a.g("annotation-labels").attr({translateX:0,translateY:0}).add(this.graphic);this.addClipPaths();this.clipRect&&this.graphic.clip(this.clipRect);
this.renderItems(this.shapes);this.renderItems(this.labels);this.addEvents();r.render.call(this)};a.prototype.setVisibility=function(a){var d=this.options,b=this.chart.navigationBindings;a=y(a,!d.visible);this.graphic.attr("visibility",a?"visible":"hidden");a||(this.setControlPointsVisibility(!1),b.activeAnnotation===this&&b.popup&&"annotation-toolbar"===b.popup.formType&&D(b,"closePopup"));d.visible=a};a.prototype.setControlPointsVisibility=function(a){var d=function(d){d.setControlPointsVisibility(a)};
r.setControlPointsVisibility.call(this,a);this.shapes.forEach(d);this.labels.forEach(d)};a.prototype.destroy=function(){var a=this.chart,d=function(a){a.destroy()};this.labels.forEach(d);this.shapes.forEach(d);this.clipYAxis=this.clipXAxis=null;z(a.labelCollectors,this.labelCollector);t.destroy.call(this);r.destroy.call(this);q(this,a)};a.prototype.remove=function(){return this.chart.removeAnnotation(this)};a.prototype.update=function(a,d){var b=this.chart,c=this.getLabelsAndShapesOptions(this.userOptions,
a),h=b.annotations.indexOf(this);a=B(!0,this.userOptions,a);a.labels=c.labels;a.shapes=c.shapes;this.destroy();this.constructor(b,a);b.options.annotations[h]=a;this.isUpdating=!0;y(d,!0)&&b.redraw();D(this,"afterUpdate");this.isUpdating=!1};a.prototype.initShape=function(d,b){d=B(this.options.shapeOptions,{controlPointOptions:this.options.controlPointOptions},d);b=new a.shapesMap[d.type](this,d,b);b.itemType="shape";this.shapes.push(b);return b};a.prototype.initLabel=function(a,d){a=B(this.options.labelOptions,
{controlPointOptions:this.options.controlPointOptions},a);d=new e(this,a,d);d.itemType="label";this.labels.push(d);return d};a.prototype.redrawItem=function(a,d){a.linkPoints();a.shouldBeDrawn()?(a.graphic||this.renderItem(a),a.redraw(y(d,!0)&&a.graphic.placed),a.points.length&&this.adjustVisibility(a)):this.destroyItem(a)};a.prototype.adjustVisibility=function(a){var d=!1,b=a.graphic;a.points.forEach(function(a){!1!==a.series.visible&&!1!==a.visible&&(d=!0)});d?"hidden"===b.visibility&&b.show():
b.hide()};a.prototype.destroyItem=function(a){z(this[a.itemType+"s"],a);a.destroy()};a.prototype.renderItem=function(a){a.render("label"===a.itemType?this.labelsGroup:this.shapesGroup)};a.ControlPoint=g;a.MockPoint=u;a.shapesMap={rect:v,circle:m,ellipse:p,path:b,image:w};a.types={};return a}();B(!0,C.prototype,r,t,B(C.prototype,{nonDOMEvents:["add","afterUpdate","drag","remove"],defaultOptions:{visible:!0,animation:{},crop:!0,draggable:"xy",labelOptions:{align:"center",allowOverlap:!1,backgroundColor:"rgba(0, 0, 0, 0.75)",
borderColor:"#000000",borderRadius:3,borderWidth:1,className:"highcharts-no-tooltip",crop:!1,formatter:function(){return l(this.y)?this.y:"Annotation label"},includeInDataExport:!0,overflow:"justify",padding:5,shadow:!1,shape:"callout",style:{fontSize:"11px",fontWeight:"normal",color:"contrast"},useHTML:!1,verticalAlign:"bottom",x:0,y:-16},shapeOptions:{stroke:"rgba(0, 0, 0, 0.75)",strokeWidth:1,fill:"rgba(0, 0, 0, 0.75)",r:0,snap:2},controlPointOptions:{symbol:"circle",width:10,height:10,style:{cursor:"pointer",
fill:"#ffffff",stroke:"#000000","stroke-width":2},visible:!1,events:{}},events:{},zIndex:6}}));f.extendAnnotation=function(a,d,b,c){d=d||C;A(a.prototype,B(d.prototype,b));a.prototype.defaultOptions=B(a.prototype.defaultOptions,c||{})};A(a,{initAnnotation:function(a){a=new (C.types[a.type]||C)(this,a);this.annotations.push(a);return a},addAnnotation:function(a,d){a=this.initAnnotation(a);this.options.annotations.push(a.options);y(d,!0)&&(a.redraw(),a.graphic.attr({opacity:1}));return a},removeAnnotation:function(a){var d=
this.annotations,b="annotations"===a.coll?a:F(d,function(d){return d.options.id===a});b&&(D(b,"remove"),z(this.options.annotations,b.options),z(d,b),b.destroy())},drawAnnotations:function(){this.plotBoxClip.attr(this.plotBox);this.annotations.forEach(function(a){a.redraw();a.graphic.animate({opacity:1},a.animationConfig)})}});a.collectionsWithUpdate.push("annotations");a.collectionsWithInit.annotations=[a.addAnnotation];c(n,"afterInit",function(){this.annotations=[];this.options.annotations||(this.options.annotations=
[])});a.callbacks.push(function(a){a.plotBoxClip=this.renderer.clipRect(this.plotBox);a.controlPointsGroup=a.renderer.g("control-points").attr({zIndex:99}).clip(a.plotBoxClip).add();a.options.annotations.forEach(function(d,b){if(!a.annotations.some(function(a){return a.options===d})){var c=a.initAnnotation(d);a.options.annotations[b]=c.options}});a.drawAnnotations();c(a,"redraw",a.drawAnnotations);c(a,"destroy",function(){a.plotBoxClip.destroy();a.controlPointsGroup.destroy()});c(a,"exportData",function(d){var b=
(this.options.exporting&&this.options.exporting.csv||{}).columnHeaderFormatter,c=!d.dataRows[1].xValues,h=a.options.lang&&a.options.lang.exportData&&a.options.lang.exportData.annotationHeader,l=function(a){if(b){var d=b(a);if(!1!==d)return d}d=h+" "+a;return c?{columnTitle:d,topLevelColumnTitle:d}:d},e=d.dataRows[0].length,q=a.options.exporting&&a.options.exporting.csv&&a.options.exporting.csv.annotations&&a.options.exporting.csv.annotations.itemDelimiter,B=a.options.exporting&&a.options.exporting.csv&&
a.options.exporting.csv.annotations&&a.options.exporting.csv.annotations.join;a.annotations.forEach(function(a){a.options.labelOptions.includeInDataExport&&a.labels.forEach(function(a){if(a.options.text){var b=a.options.text;a.points.forEach(function(a){var c=a.x,h=a.series.xAxis?a.series.xAxis.options.index:-1,l=!1;if(-1===h){a=d.dataRows[0].length;for(var y=Array(a),f=0;f<a;++f)y[f]="";y.push(b);y.xValues=[];y.xValues[h]=c;d.dataRows.push(y);l=!0}l||d.dataRows.forEach(function(a,d){!l&&a.xValues&&
void 0!==h&&c===a.xValues[h]&&(B&&a.length>e?a[a.length-1]+=q+b:a.push(b),l=!0)});if(!l){a=d.dataRows[0].length;y=Array(a);for(f=0;f<a;++f)y[f]="";y[0]=c;y.push(b);y.xValues=[];void 0!==h&&(y.xValues[h]=c);d.dataRows.push(y)}})}})});var y=0;d.dataRows.forEach(function(a){y=Math.max(y,a.length)});for(var G=y-d.dataRows[0].length,f=0;f<G;f++){var C=l(f+1);c?(d.dataRows[0].push(C.topLevelColumnTitle),d.dataRows[1].push(C.columnTitle)):d.dataRows[0].push(C)}})});k(x.prototype,"onContainerMouseDown",function(a){this.chart.hasDraggedAnnotation||
a.apply(this,Array.prototype.slice.call(arguments,1))});f.Annotation=C;"";return C});w(a,"Core/Chart/ChartNavigationComposition.js",[],function(){var a;(function(a){a.compose=function(a){a.navigation||(a.navigation=new c(a));return a};var c=function(){function a(a){this.updates=[];this.chart=a}a.prototype.addUpdate=function(a){this.chart.navigation.updates.push(a)};a.prototype.update=function(a,c){var b=this;this.updates.forEach(function(m){m.call(b.chart,a,c)})};return a}();a.Additions=c})(a||(a=
{}));return a});w(a,"Extensions/Annotations/NavigationBindings.js",[a["Extensions/Annotations/Annotations.js"],a["Core/Chart/Chart.js"],a["Core/Chart/ChartNavigationComposition.js"],a["Core/FormatUtilities.js"],a["Core/Globals.js"],a["Core/DefaultOptions.js"],a["Core/Utilities.js"]],function(a,n,r,v,m,p,b){function c(a,d){var b=F.Element.prototype,c=b.matches||b.msMatchesSelector||b.webkitMatchesSelector,h=null;if(b.closest)h=b.closest.call(a,d);else{do{if(c.call(a,d))return a;a=a.parentElement||
a.parentNode}while(null!==a&&1===a.nodeType)}return h}function e(a){var d=a.prototype.defaultOptions.events&&a.prototype.defaultOptions.events.click;l(!0,a.prototype.defaultOptions.events,{click:function(a){var b=this,c=b.chart.navigationBindings,h=c.activeAnnotation;d&&d.call(b,a);h!==b?(c.deselectAnnotation(),c.activeAnnotation=b,b.setControlPointsVisibility(!0),u(c,"showPopup",{annotation:b,formType:"annotation-toolbar",options:c.annotationToFields(b),onSubmit:function(a){var d={};"remove"===a.actionType?
(c.activeAnnotation=!1,c.chart.removeAnnotation(b)):(c.fieldsToOptions(a.fields,d),c.deselectAnnotation(),a=d.typeOptions,"measure"===b.options.type&&(a.crosshairY.enabled=0!==a.crosshairY.strokeWidth,a.crosshairX.enabled=0!==a.crosshairX.strokeWidth),b.update(d))}})):u(c,"closePopup");a.activeAnnotation=!0}})}var g=v.format;v=p.setOptions;var t=b.addEvent,f=b.attr,u=b.fireEvent,x=b.isArray,k=b.isFunction,d=b.isNumber,h=b.isObject,l=b.merge,q=b.objectEach,z=b.pick,A=m.doc,F=m.win,D=function(){function a(a,
d){this.selectedButton=this.boundClassNames=void 0;this.chart=a;this.options=d;this.eventsToUnbind=[];this.container=A.getElementsByClassName(this.options.bindingsClassName||"")}a.prototype.initEvents=function(){var a=this,d=a.chart,b=a.container,c=a.options;a.boundClassNames={};q(c.bindings||{},function(d){a.boundClassNames[d.className]=d});[].forEach.call(b,function(d){a.eventsToUnbind.push(t(d,"click",function(b){var c=a.getButtonEvents(d,b);c&&-1===c.button.className.indexOf("highcharts-disabled-btn")&&
a.bindingsButtonClick(c.button,c.events,b)}))});q(c.events||{},function(d,b){k(d)&&a.eventsToUnbind.push(t(a,b,d,{passive:!1}))});a.eventsToUnbind.push(t(d.container,"click",function(b){!d.cancelClick&&d.isInsidePlot(b.chartX-d.plotLeft,b.chartY-d.plotTop,{visiblePlotOnly:!0})&&a.bindingsChartClick(this,b)}));a.eventsToUnbind.push(t(d.container,m.isTouchDevice?"touchmove":"mousemove",function(d){a.bindingsContainerMouseMove(this,d)},m.isTouchDevice?{passive:!1}:void 0))};a.prototype.initUpdate=function(){var a=
this;r.compose(this.chart).navigation.addUpdate(function(d){a.update(d)})};a.prototype.bindingsButtonClick=function(a,d,b){var c=this.chart,h=c.renderer.boxWrapper,l=!0;this.selectedButtonElement&&(this.selectedButtonElement.classList===a.classList&&(l=!1),u(this,"deselectButton",{button:this.selectedButtonElement}),this.nextEvent&&(this.currentUserDetails&&"annotations"===this.currentUserDetails.coll&&c.removeAnnotation(this.currentUserDetails),this.mouseMoveEvent=this.nextEvent=!1));l?(this.selectedButton=
d,this.selectedButtonElement=a,u(this,"selectButton",{button:a}),d.init&&d.init.call(this,a,b),(d.start||d.steps)&&c.renderer.boxWrapper.addClass("highcharts-draw-mode")):(c.stockTools&&c.stockTools.toggleButtonAciveClass(a),h.removeClass("highcharts-draw-mode"),this.mouseMoveEvent=this.nextEvent=!1,this.selectedButton=null)};a.prototype.bindingsChartClick=function(a,d){a=this.chart;var b=this.activeAnnotation,h=this.selectedButton;a=a.renderer.boxWrapper;b&&(b.cancelClick||d.activeAnnotation||!d.target.parentNode||
c(d.target,".highcharts-popup")?b.cancelClick&&setTimeout(function(){b.cancelClick=!1},0):u(this,"closePopup"));h&&h.start&&(this.nextEvent?(this.nextEvent(d,this.currentUserDetails),this.steps&&(this.stepIndex++,h.steps[this.stepIndex]?this.mouseMoveEvent=this.nextEvent=h.steps[this.stepIndex]:(u(this,"deselectButton",{button:this.selectedButtonElement}),a.removeClass("highcharts-draw-mode"),h.end&&h.end.call(this,d,this.currentUserDetails),this.mouseMoveEvent=this.nextEvent=!1,this.selectedButton=
null))):(this.currentUserDetails=h.start.call(this,d))&&h.steps?(this.stepIndex=0,this.steps=!0,this.mouseMoveEvent=this.nextEvent=h.steps[this.stepIndex]):(u(this,"deselectButton",{button:this.selectedButtonElement}),a.removeClass("highcharts-draw-mode"),this.steps=!1,this.selectedButton=null,h.end&&h.end.call(this,d,this.currentUserDetails)))};a.prototype.bindingsContainerMouseMove=function(a,d){this.mouseMoveEvent&&this.mouseMoveEvent(d,this.currentUserDetails)};a.prototype.fieldsToOptions=function(a,
b){q(a,function(a,c){var h=parseFloat(a),l=c.split("."),e=b,q=l.length-1;!d(h)||a.match(/px/g)||c.match(/format/g)||(a=h);""!==a&&"undefined"!==a&&l.forEach(function(d,b){var c=z(l[b+1],"");q===b?e[d]=a:(e[d]||(e[d]=c.match(/\d/g)?[]:{}),e=e[d])})});return b};a.prototype.deselectAnnotation=function(){this.activeAnnotation&&(this.activeAnnotation.setControlPointsVisibility(!1),this.activeAnnotation=!1)};a.prototype.annotationToFields=function(d){function b(a,c,l,f){if(l&&a&&-1===y.indexOf(c)&&(0<=
(l.indexOf&&l.indexOf(c))||l[c]||!0===l))if(x(a))f[c]=[],a.forEach(function(a,d){h(a)?(f[c][d]={},q(a,function(a,h){b(a,h,e[c],f[c][d])})):b(a,0,e[c],f[c])});else if(h(a)){var z={};x(f)?(f.push(z),z[c]={},z=z[c]):f[c]=z;q(a,function(a,d){b(a,d,0===c?l:e[c],z)})}else"format"===c?f[c]=[g(a,d.labels[0].points[0]).toString(),"text"]:x(f)?f.push([a,B(a)]):f[c]=[a,B(a)]}var c=d.options,l=a.annotationsEditable,e=l.nestedOptions,B=this.utils.getFieldType,f=z(c.type,c.shapes&&c.shapes[0]&&c.shapes[0].type,
c.labels&&c.labels[0]&&c.labels[0].itemType,"label"),y=a.annotationsNonEditable[c.langKey]||[],k={langKey:c.langKey,type:f};q(c,function(a,d){"typeOptions"===d?(k[d]={},q(c[d],function(a,c){b(a,c,e,k[d],!0)})):b(a,d,l[f],k)});return k};a.prototype.getClickedClassNames=function(a,d){var b=d.target;d=[];for(var c;b&&((c=f(b,"class"))&&(d=d.concat(c.split(" ").map(function(a){return[a,b]}))),b=b.parentNode,b!==a););return d};a.prototype.getButtonEvents=function(a,d){var b=this,c;this.getClickedClassNames(a,
d).forEach(function(a){b.boundClassNames[a[0]]&&!c&&(c={events:b.boundClassNames[a[0]],button:a[1]})});return c};a.prototype.update=function(a){this.options=l(!0,this.options,a);this.removeEvents();this.initEvents()};a.prototype.removeEvents=function(){this.eventsToUnbind.forEach(function(a){a()})};a.prototype.destroy=function(){this.removeEvents()};a.annotationsEditable={nestedOptions:{labelOptions:["style","format","backgroundColor"],labels:["style"],label:["style"],style:["fontSize","color"],background:["fill",
"strokeWidth","stroke"],innerBackground:["fill","strokeWidth","stroke"],outerBackground:["fill","strokeWidth","stroke"],shapeOptions:["fill","strokeWidth","stroke"],shapes:["fill","strokeWidth","stroke"],line:["strokeWidth","stroke"],backgroundColors:[!0],connector:["fill","strokeWidth","stroke"],crosshairX:["strokeWidth","stroke"],crosshairY:["strokeWidth","stroke"]},circle:["shapes"],ellipse:["shapes"],verticalLine:[],label:["labelOptions"],measure:["background","crosshairY","crosshairX"],fibonacci:[],
tunnel:["background","line","height"],pitchfork:["innerBackground","outerBackground"],rect:["shapes"],crookedLine:[],basicAnnotation:["shapes","labelOptions"]};a.annotationsNonEditable={rectangle:["crosshairX","crosshairY","labelOptions"],ellipse:["labelOptions"],circle:["labelOptions"]};return a}();D.prototype.utils={getFieldType:function(a){return{string:"text",number:"number","boolean":"checkbox"}[typeof a]},updateRectSize:function(a,b){var c=b.chart,h=b.options.typeOptions,l=d(h.xAxis)&&c.xAxis[h.xAxis],
e=d(h.yAxis)&&c.yAxis[h.yAxis];l&&e&&(l=l.toValue(a[l.horiz?"chartX":"chartY"]),a=e.toValue(a[e.horiz?"chartX":"chartY"]),e=l-h.point.x,h=h.point.y-a,b.update({typeOptions:{background:{width:c.inverted?h:e,height:c.inverted?e:h}}}))},getAssignedAxis:function(a){return a.filter(function(a){var b=a.axis.getExtremes(),c=b.min;b=b.max;var h=z(a.axis.minPointOffset,0);return d(c)&&d(b)&&a.value>=c-h&&a.value<=b+h&&!a.axis.options.isInternal})[0]}};n.prototype.initNavigationBindings=function(){var a=this.options;
a&&a.navigation&&a.navigation.bindings&&(this.navigationBindings=new D(this,a.navigation),this.navigationBindings.initEvents(),this.navigationBindings.initUpdate())};t(n,"load",function(){this.initNavigationBindings()});t(n,"destroy",function(){this.navigationBindings&&this.navigationBindings.destroy()});t(D,"deselectButton",function(){this.selectedButtonElement=null});t(a,"remove",function(){this.chart.navigationBindings&&this.chart.navigationBindings.deselectAnnotation()});m.Annotation&&(e(a),q(a.types,
function(a){e(a)}));v({lang:{navigation:{popup:{simpleShapes:"Simple shapes",lines:"Lines",circle:"Circle",ellipse:"Ellipse",rectangle:"Rectangle",label:"Label",shapeOptions:"Shape options",typeOptions:"Details",fill:"Fill",format:"Text",strokeWidth:"Line width",stroke:"Line color",title:"Title",name:"Name",labelOptions:"Label options",labels:"Labels",backgroundColor:"Background color",backgroundColors:"Background colors",borderColor:"Border color",borderRadius:"Border radius",borderWidth:"Border width",
style:"Style",padding:"Padding",fontSize:"Font size",color:"Color",height:"Height",shapes:"Shape options"}}},navigation:{bindingsClassName:"highcharts-bindings-container",bindings:{circleAnnotation:{className:"highcharts-circle-annotation",start:function(a){var d=this.chart.pointer.getCoordinates(a);a=this.utils.getAssignedAxis(d.xAxis);d=this.utils.getAssignedAxis(d.yAxis);var b=this.chart.options.navigation;if(a&&d)return this.chart.addAnnotation(l({langKey:"circle",type:"basicAnnotation",shapes:[{type:"circle",
point:{x:a.value,y:d.value,xAxis:a.axis.options.index,yAxis:d.axis.options.index},r:5}]},b.annotationsOptions,b.bindings.circleAnnotation.annotationsOptions))},steps:[function(a,b){var c=b.options.shapes[0].point;if(d(c.xAxis)&&d(c.yAxis)){var h=this.chart.inverted;var l=this.chart.xAxis[c.xAxis].toPixels(c.x);c=this.chart.yAxis[c.yAxis].toPixels(c.y);h=Math.max(Math.sqrt(Math.pow(h?c-a.chartX:l-a.chartX,2)+Math.pow(h?l-a.chartY:c-a.chartY,2)),5)}b.update({shapes:[{r:h}]})}]},ellipseAnnotation:{className:"highcharts-ellipse-annotation",
start:function(a){var d=this.chart.pointer.getCoordinates(a);a=this.utils.getAssignedAxis(d.xAxis);d=this.utils.getAssignedAxis(d.yAxis);var b=this.chart.options.navigation;if(a&&d)return this.chart.addAnnotation(l({langKey:"ellipse",type:"basicAnnotation",shapes:[{type:"ellipse",xAxis:a.axis.options.index,yAxis:d.axis.options.index,points:[{x:a.value,y:d.value},{x:a.value,y:d.value}],ry:1}]},b.annotationsOptions,b.bindings.ellipseAnnotation.annotationOptions))},steps:[function(a,d){d=d.shapes[0];
var b=d.getAbsolutePosition(d.points[1]);d.translatePoint(a.chartX-b.x,a.chartY-b.y,1);d.redraw(!1)},function(a,d){d=d.shapes[0];var b=d.getAbsolutePosition(d.points[0]),c=d.getAbsolutePosition(d.points[1]);a=d.getDistanceFromLine(b,c,a.chartX,a.chartY);b=d.getYAxis();a=Math.abs(b.toValue(0)-b.toValue(a));d.setYRadius(a);d.redraw(!1)}]},rectangleAnnotation:{className:"highcharts-rectangle-annotation",start:function(a){a=this.chart.pointer.getCoordinates(a);var d=this.utils.getAssignedAxis(a.xAxis),
b=this.utils.getAssignedAxis(a.yAxis);if(d&&b){a=d.value;var c=b.value;d=d.axis.options.index;b=b.axis.options.index;var h=this.chart.options.navigation;return this.chart.addAnnotation(l({langKey:"rectangle",type:"basicAnnotation",shapes:[{type:"path",points:[{xAxis:d,yAxis:b,x:a,y:c},{xAxis:d,yAxis:b,x:a,y:c},{xAxis:d,yAxis:b,x:a,y:c},{xAxis:d,yAxis:b,x:a,y:c},{command:"Z"}]}]},h.annotationsOptions,h.bindings.rectangleAnnotation.annotationsOptions))}},steps:[function(a,d){var b=d.options.shapes[0].points,
c=this.chart.pointer.getCoordinates(a);a=this.utils.getAssignedAxis(c.xAxis);c=this.utils.getAssignedAxis(c.yAxis);a&&c&&(a=a.value,c=c.value,b[1].x=a,b[2].x=a,b[2].y=c,b[3].y=c,d.update({shapes:[{points:b}]}))}]},labelAnnotation:{className:"highcharts-label-annotation",start:function(a){var d=this.chart.pointer.getCoordinates(a);a=this.utils.getAssignedAxis(d.xAxis);d=this.utils.getAssignedAxis(d.yAxis);var b=this.chart.options.navigation;if(a&&d)return this.chart.addAnnotation(l({langKey:"label",
type:"basicAnnotation",labelOptions:{format:"{y:.2f}"},labels:[{point:{xAxis:a.axis.options.index,yAxis:d.axis.options.index,x:a.value,y:d.value},overflow:"none",crop:!0}]},b.annotationsOptions,b.bindings.labelAnnotation.annotationsOptions))}}},events:{},annotationsOptions:{animation:{defer:0}}}});t(n,"render",function(){var a=this,d=a.navigationBindings;if(a&&d){var b=!1;a.series.forEach(function(a){!a.options.isInternal&&a.visible&&(b=!0)});q(d.boundClassNames,function(d,c){if(a.navigationBindings&&
a.navigationBindings.container&&a.navigationBindings.container[0]&&(c=a.navigationBindings.container[0].querySelectorAll("."+c)))for(var h=0;h<c.length;h++){var l=c[h];"normal"===d.noDataState?-1!==l.className.indexOf("highcharts-disabled-btn")&&l.classList.remove("highcharts-disabled-btn"):b?-1!==l.className.indexOf("highcharts-disabled-btn")&&l.classList.remove("highcharts-disabled-btn"):-1===l.className.indexOf("highcharts-disabled-btn")&&(l.className+=" highcharts-disabled-btn")}})}});t(D,"closePopup",
function(){this.deselectAnnotation()});return D});w(a,"Stock/StockToolsBindings.js",[a["Core/DefaultOptions.js"],a["Core/Globals.js"],a["Extensions/Annotations/NavigationBindings.js"],a["Core/Series/Series.js"],a["Core/Utilities.js"]],function(a,n,r,v,m){var c=a.getOptions;a=a.setOptions;var b=m.correctFloat,w=m.defined,e=m.extend,g=m.fireEvent,t=m.isNumber,f=m.merge,u=m.pick,x=m.uniqueKey,k=r.prototype.utils;k.addFlagFromForm=function(a){return function(d){var b=this,c=b.chart,h=c.stockTools,e=k.getFieldType;
d=k.attractToPoint(d,c);if(d){var f={type:"flags",onSeries:d.series.id,shape:a,data:[{x:d.x,y:d.y}],xAxis:d.xAxis,yAxis:d.yAxis,point:{events:{click:function(){var a=this,d=a.options;g(b,"showPopup",{point:a,formType:"annotation-toolbar",options:{langKey:"flags",type:"flags",title:[d.title,e(d.title)],name:[d.name,e(d.name)]},onSubmit:function(d){"remove"===d.actionType?a.remove():a.update(b.fieldsToOptions(d.fields,{}))}})}}}};h&&h.guiEnabled||c.addSeries(f);g(b,"showPopup",{formType:"flag",options:{langKey:"flags",
type:"flags",title:["A",e("A")],name:["Flag A",e("Flag A")]},onSubmit:function(a){b.fieldsToOptions(a.fields,f.data[0]);c.addSeries(f)}})}}};k.indicatorsWithAxes="apo ad aroon aroonoscillator atr ao cci chaikin cmf cmo disparityindex dmi dpo linearRegressionAngle linearRegressionIntercept linearRegressionSlope klinger macd mfi momentum natr obv ppo roc rsi slowstochastic stochastic trix williamsr".split(" ");k.manageIndicators=function(a){var d=this.chart,b={linkedTo:a.linkedTo,type:a.type},e="ad cmf klinger mfi obv vbp vwap".split(" "),
f=k.indicatorsWithAxes;if("edit"===a.actionType)this.fieldsToOptions(a.fields,b),(a=d.get(a.seriesId))&&a.update(b,!1);else if("remove"===a.actionType){if(a=d.get(a.seriesId)){var A=a.yAxis;a.linkedSeries&&a.linkedSeries.forEach(function(a){a.remove(!1)});a.remove(!1);0<=f.indexOf(a.type)&&(b=A.options.height,A.remove(!1),this.resizeYAxes(b))}}else{b.id=x();this.fieldsToOptions(a.fields,b);A=d.get(b.linkedTo);var m=c().plotOptions;"undefined"!==typeof A&&A instanceof v&&"sum"===A.getDGApproximation()&&
!w(m&&m[b.type]&&m.dataGrouping&&m.dataGrouping.approximation)&&(b.dataGrouping={approximation:"sum"});0<=f.indexOf(a.type)?(A=d.addAxis({id:x(),offset:0,opposite:!0,title:{text:""},tickPixelInterval:40,showLastLabel:!1,labels:{align:"left",y:-2}},!1,!1),b.yAxis=A.options.id,this.resizeYAxes()):b.yAxis=d.get(a.linkedTo).options.yAxis;0<=e.indexOf(a.type)&&(b.params.volumeSeriesID=d.series.filter(function(a){return"column"===a.options.type})[0].options.id);d.addSeries(b,!1)}g(this,"deselectButton",
{button:this.selectedButtonElement});d.redraw()};k.updateHeight=function(a,b){var d=b.options.typeOptions,c=t(d.yAxis)&&this.chart.yAxis[d.yAxis];c&&d.points&&b.update({typeOptions:{height:c.toValue(a[c.horiz?"chartX":"chartY"])-(d.points[1].y||0)}})};k.attractToPoint=function(a,b){a=b.pointer.getCoordinates(a);var d=Number.MAX_VALUE,c;if(b.navigationBindings){var h=b.navigationBindings.utils.getAssignedAxis(a.xAxis);var e=b.navigationBindings.utils.getAssignedAxis(a.yAxis)}if(h&&e){var f=h.value;
b=e.value;e.axis.series.forEach(function(a){a.points&&a.points.forEach(function(a){a&&d>Math.abs(a.x-f)&&(d=Math.abs(a.x-f),c=a)})});if(c&&c.x&&c.y)return{x:c.x,y:c.y,below:b<c.y,series:c.series,xAxis:c.series.xAxis.options.index||0,yAxis:c.series.yAxis.options.index||0}}};k.isNotNavigatorYAxis=function(a){return"highcharts-navigator-yaxis"!==a.userOptions.className};k.isPriceIndicatorEnabled=function(a){return a.some(function(a){return a.lastVisiblePrice||a.lastPrice})};k.updateNthPoint=function(a){return function(d,
b){var c=b.options.typeOptions,h=t(c.xAxis)&&this.chart.xAxis[c.xAxis],l=t(c.yAxis)&&this.chart.yAxis[c.yAxis];h&&l&&(c.points.forEach(function(b,c){c>=a&&(b.x=h.toValue(d[h.horiz?"chartX":"chartY"]),b.y=l.toValue(d[l.horiz?"chartX":"chartY"]))}),b.update({typeOptions:{points:c.points}}))}};e(r.prototype,{getYAxisPositions:function(a,c,l,e){function d(a){return w(a)&&!t(a)&&a.match("%")}var h=0,f,q;e&&(q=b(parseFloat(e)/100));return{positions:a.map(function(e,g){var k=b(d(e.options.height)?parseFloat(e.options.height)/
100:e.height/c);e=b(d(e.options.top)?parseFloat(e.options.top)/100:(e.top-e.chart.plotTop)/c);q?e<=h?h=b(Math.max(h,(e||0)+(k||0))):(e=b(e-q),h=b(h+k)):(t(k)||(k=a[g-1].series.every(function(a){return a.is("sma")})?f:l/100),t(e)||(e=h),f=k,h=b(Math.max(h,(e||0)+(k||0))));return{height:100*k,top:100*e}}),allAxesHeight:h}},getYAxisResizers:function(a){var d=[];a.forEach(function(b,c){b=a[c+1];d[c]=b?{enabled:!0,controlledAxis:{next:[u(b.options.id,b.options.index)]}}:{enabled:!1}});return d},resizeYAxes:function(a){var d=
this.chart,c=d.yAxis.filter(k.isNotNavigatorYAxis);d=this.getYAxisPositions(c,d.plotHeight,20,a);var e=d.positions,f=d.allAxesHeight,g=this.getYAxisResizers(c);!a&&f<=b(1)?e[e.length-1]={height:20,top:b(100*f-20)}:e.forEach(function(a){a.height=a.height/(100*f)*100;a.top=a.top/(100*f)*100});e.forEach(function(a,d){c[d].update({height:a.height+"%",top:a.top+"%",resize:g[d],offset:0},!1)})},recalculateYAxisPositions:function(a,c,e,f){a.forEach(function(d,h){h=a[h-1];d.top=h?b(h.height+h.top):0;e&&(d.height=
b(d.height+f*c))});return a}});m={segment:{className:"highcharts-segment",start:function(a){var d=this.chart.pointer.getCoordinates(a);a=this.utils.getAssignedAxis(d.xAxis);d=this.utils.getAssignedAxis(d.yAxis);var b=this.chart.options.navigation;if(a&&d)return a=f({langKey:"segment",type:"crookedLine",typeOptions:{xAxis:a.axis.options.index,yAxis:d.axis.options.index,points:[{x:a.value,y:d.value},{x:a.value,y:d.value}]}},b.annotationsOptions,b.bindings.segment.annotationsOptions),this.chart.addAnnotation(a)},
steps:[k.updateNthPoint(1)]},arrowSegment:{className:"highcharts-arrow-segment",start:function(a){var d=this.chart.pointer.getCoordinates(a);a=this.utils.getAssignedAxis(d.xAxis);d=this.utils.getAssignedAxis(d.yAxis);var b=this.chart.options.navigation;if(a&&d)return a=f({langKey:"arrowSegment",type:"crookedLine",typeOptions:{line:{markerEnd:"arrow"},xAxis:a.axis.options.index,yAxis:d.axis.options.index,points:[{x:a.value,y:d.value},{x:a.value,y:d.value}]}},b.annotationsOptions,b.bindings.arrowSegment.annotationsOptions),
this.chart.addAnnotation(a)},steps:[k.updateNthPoint(1)]},ray:{className:"highcharts-ray",start:function(a){var d=this.chart.pointer.getCoordinates(a);a=this.utils.getAssignedAxis(d.xAxis);d=this.utils.getAssignedAxis(d.yAxis);var b=this.chart.options.navigation;if(a&&d)return a=f({langKey:"ray",type:"infinityLine",typeOptions:{type:"ray",xAxis:a.axis.options.index,yAxis:d.axis.options.index,points:[{x:a.value,y:d.value},{x:a.value,y:d.value}]}},b.annotationsOptions,b.bindings.ray.annotationsOptions),
this.chart.addAnnotation(a)},steps:[k.updateNthPoint(1)]},arrowRay:{className:"highcharts-arrow-ray",start:function(a){var d=this.chart.pointer.getCoordinates(a);a=this.utils.getAssignedAxis(d.xAxis);d=this.utils.getAssignedAxis(d.yAxis);var b=this.chart.options.navigation;if(a&&d)return a=f({langKey:"arrowRay",type:"infinityLine",typeOptions:{type:"ray",line:{markerEnd:"arrow"},xAxis:a.axis.options.index,yAxis:d.axis.options.index,points:[{x:a.value,y:d.value},{x:a.value,y:d.value}]}},b.annotationsOptions,
b.bindings.arrowRay.annotationsOptions),this.chart.addAnnotation(a)},steps:[k.updateNthPoint(1)]},infinityLine:{className:"highcharts-infinity-line",start:function(a){var d=this.chart.pointer.getCoordinates(a);a=this.utils.getAssignedAxis(d.xAxis);d=this.utils.getAssignedAxis(d.yAxis);var b=this.chart.options.navigation;if(a&&d)return a=f({langKey:"infinityLine",type:"infinityLine",typeOptions:{type:"line",xAxis:a.axis.options.index,yAxis:d.axis.options.index,points:[{x:a.value,y:d.value},{x:a.value,
y:d.value}]}},b.annotationsOptions,b.bindings.infinityLine.annotationsOptions),this.chart.addAnnotation(a)},steps:[k.updateNthPoint(1)]},arrowInfinityLine:{className:"highcharts-arrow-infinity-line",start:function(a){var d=this.chart.pointer.getCoordinates(a);a=this.utils.getAssignedAxis(d.xAxis);d=this.utils.getAssignedAxis(d.yAxis);var b=this.chart.options.navigation;if(a&&d)return a=f({langKey:"arrowInfinityLine",type:"infinityLine",typeOptions:{type:"line",line:{markerEnd:"arrow"},xAxis:a.axis.options.index,
yAxis:d.axis.options.index,points:[{x:a.value,y:d.value},{x:a.value,y:d.value}]}},b.annotationsOptions,b.bindings.arrowInfinityLine.annotationsOptions),this.chart.addAnnotation(a)},steps:[k.updateNthPoint(1)]},horizontalLine:{className:"highcharts-horizontal-line",start:function(a){var d=this.chart.pointer.getCoordinates(a);a=this.utils.getAssignedAxis(d.xAxis);d=this.utils.getAssignedAxis(d.yAxis);var b=this.chart.options.navigation;a&&d&&(a=f({langKey:"horizontalLine",type:"infinityLine",draggable:"y",
typeOptions:{type:"horizontalLine",xAxis:a.axis.options.index,yAxis:d.axis.options.index,points:[{x:a.value,y:d.value}]}},b.annotationsOptions,b.bindings.horizontalLine.annotationsOptions),this.chart.addAnnotation(a))}},verticalLine:{className:"highcharts-vertical-line",start:function(a){var d=this.chart.pointer.getCoordinates(a);a=this.utils.getAssignedAxis(d.xAxis);d=this.utils.getAssignedAxis(d.yAxis);var b=this.chart.options.navigation;a&&d&&(a=f({langKey:"verticalLine",type:"infinityLine",draggable:"x",
typeOptions:{type:"verticalLine",xAxis:a.axis.options.index,yAxis:d.axis.options.index,points:[{x:a.value,y:d.value}]}},b.annotationsOptions,b.bindings.verticalLine.annotationsOptions),this.chart.addAnnotation(a))}},crooked3:{className:"highcharts-crooked3",start:function(a){var d=this.chart.pointer.getCoordinates(a);a=this.utils.getAssignedAxis(d.xAxis);d=this.utils.getAssignedAxis(d.yAxis);if(a&&d){var b=a.value,c=d.value,e=this.chart.options.navigation;a=f({langKey:"crooked3",type:"crookedLine",
typeOptions:{xAxis:a.axis.options.index,yAxis:d.axis.options.index,points:[{x:b,y:c},{x:b,y:c},{x:b,y:c}]}},e.annotationsOptions,e.bindings.crooked3.annotationsOptions);return this.chart.addAnnotation(a)}},steps:[k.updateNthPoint(1),k.updateNthPoint(2)]},crooked5:{className:"highcharts-crooked5",start:function(a){var d=this.chart.pointer.getCoordinates(a);a=this.utils.getAssignedAxis(d.xAxis);d=this.utils.getAssignedAxis(d.yAxis);if(a&&d){var b=a.value,c=d.value,e=this.chart.options.navigation;a=
f({langKey:"crooked5",type:"crookedLine",typeOptions:{xAxis:a.axis.options.index,yAxis:d.axis.options.index,points:[{x:b,y:c},{x:b,y:c},{x:b,y:c},{x:b,y:c},{x:b,y:c}]}},e.annotationsOptions,e.bindings.crooked5.annotationsOptions);return this.chart.addAnnotation(a)}},steps:[k.updateNthPoint(1),k.updateNthPoint(2),k.updateNthPoint(3),k.updateNthPoint(4)]},elliott3:{className:"highcharts-elliott3",start:function(a){var b=this.chart.pointer.getCoordinates(a);a=this.utils.getAssignedAxis(b.xAxis);b=this.utils.getAssignedAxis(b.yAxis);
if(a&&b){var d=a.value,c=b.value,e=this.chart.options.navigation;a=f({langKey:"elliott3",type:"elliottWave",typeOptions:{xAxis:a.axis.options.index,yAxis:b.axis.options.index,points:[{x:d,y:c},{x:d,y:c},{x:d,y:c},{x:d,y:c}]},labelOptions:{style:{color:"#666666"}}},e.annotationsOptions,e.bindings.elliott3.annotationsOptions);return this.chart.addAnnotation(a)}},steps:[k.updateNthPoint(1),k.updateNthPoint(2),k.updateNthPoint(3)]},elliott5:{className:"highcharts-elliott5",start:function(a){var b=this.chart.pointer.getCoordinates(a);
a=this.utils.getAssignedAxis(b.xAxis);b=this.utils.getAssignedAxis(b.yAxis);if(a&&b){var d=a.value,c=b.value,e=this.chart.options.navigation;a=f({langKey:"elliott5",type:"elliottWave",typeOptions:{xAxis:a.axis.options.index,yAxis:b.axis.options.index,points:[{x:d,y:c},{x:d,y:c},{x:d,y:c},{x:d,y:c},{x:d,y:c},{x:d,y:c}]},labelOptions:{style:{color:"#666666"}}},e.annotationsOptions,e.bindings.elliott5.annotationsOptions);return this.chart.addAnnotation(a)}},steps:[k.updateNthPoint(1),k.updateNthPoint(2),
k.updateNthPoint(3),k.updateNthPoint(4),k.updateNthPoint(5)]},measureX:{className:"highcharts-measure-x",start:function(a){var b=this.chart.pointer.getCoordinates(a);a=this.utils.getAssignedAxis(b.xAxis);b=this.utils.getAssignedAxis(b.yAxis);if(a&&b){var d=this.chart.options.navigation;a=f({langKey:"measure",type:"measure",typeOptions:{selectType:"x",xAxis:a.axis.options.index,yAxis:b.axis.options.index,point:{x:a.value,y:b.value},crosshairX:{strokeWidth:1,stroke:"#000000"},crosshairY:{enabled:!1,
strokeWidth:0,stroke:"#000000"},background:{width:0,height:0,strokeWidth:0,stroke:"#ffffff"}},labelOptions:{style:{color:"#666666"}}},d.annotationsOptions,d.bindings.measureX.annotationsOptions);return this.chart.addAnnotation(a)}},steps:[k.updateRectSize]},measureY:{className:"highcharts-measure-y",start:function(a){var b=this.chart.pointer.getCoordinates(a);a=this.utils.getAssignedAxis(b.xAxis);b=this.utils.getAssignedAxis(b.yAxis);if(a&&b){var d=this.chart.options.navigation;a=f({langKey:"measure",
type:"measure",typeOptions:{selectType:"y",xAxis:a.axis.options.index,yAxis:b.axis.options.index,point:{x:a.value,y:b.value},crosshairX:{enabled:!1,strokeWidth:0,stroke:"#000000"},crosshairY:{strokeWidth:1,stroke:"#000000"},background:{width:0,height:0,strokeWidth:0,stroke:"#ffffff"}},labelOptions:{style:{color:"#666666"}}},d.annotationsOptions,d.bindings.measureY.annotationsOptions);return this.chart.addAnnotation(a)}},steps:[k.updateRectSize]},measureXY:{className:"highcharts-measure-xy",start:function(a){var b=
this.chart.pointer.getCoordinates(a);a=this.utils.getAssignedAxis(b.xAxis);b=this.utils.getAssignedAxis(b.yAxis);if(a&&b){var d=this.chart.options.navigation;a=f({langKey:"measure",type:"measure",typeOptions:{selectType:"xy",xAxis:a.axis.options.index,yAxis:b.axis.options.index,point:{x:a.value,y:b.value},background:{width:0,height:0,strokeWidth:10},crosshairX:{strokeWidth:1,stroke:"#000000"},crosshairY:{strokeWidth:1,stroke:"#000000"}},labelOptions:{style:{color:"#666666"}}},d.annotationsOptions,
d.bindings.measureXY.annotationsOptions);return this.chart.addAnnotation(a)}},steps:[k.updateRectSize]},fibonacci:{className:"highcharts-fibonacci",start:function(a){var b=this.chart.pointer.getCoordinates(a);a=this.utils.getAssignedAxis(b.xAxis);b=this.utils.getAssignedAxis(b.yAxis);if(a&&b){var d=a.value,c=b.value,e=this.chart.options.navigation;a=f({langKey:"fibonacci",type:"fibonacci",typeOptions:{xAxis:a.axis.options.index,yAxis:b.axis.options.index,points:[{x:d,y:c},{x:d,y:c}]},labelOptions:{style:{color:"#666666"}}},
e.annotationsOptions,e.bindings.fibonacci.annotationsOptions);return this.chart.addAnnotation(a)}},steps:[k.updateNthPoint(1),k.updateHeight]},parallelChannel:{className:"highcharts-parallel-channel",start:function(a){var b=this.chart.pointer.getCoordinates(a);a=this.utils.getAssignedAxis(b.xAxis);b=this.utils.getAssignedAxis(b.yAxis);if(a&&b){var d=a.value,c=b.value,e=this.chart.options.navigation;a=f({langKey:"parallelChannel",type:"tunnel",typeOptions:{xAxis:a.axis.options.index,yAxis:b.axis.options.index,
points:[{x:d,y:c},{x:d,y:c}]}},e.annotationsOptions,e.bindings.parallelChannel.annotationsOptions);return this.chart.addAnnotation(a)}},steps:[k.updateNthPoint(1),k.updateHeight]},pitchfork:{className:"highcharts-pitchfork",start:function(a){var b=this.chart.pointer.getCoordinates(a);a=this.utils.getAssignedAxis(b.xAxis);b=this.utils.getAssignedAxis(b.yAxis);if(a&&b){var d=a.value,c=b.value,e=this.chart.options.navigation;a=f({langKey:"pitchfork",type:"pitchfork",typeOptions:{xAxis:a.axis.options.index,
yAxis:b.axis.options.index,points:[{x:a.value,y:b.value,controlPoint:{style:{fill:"#f21313"}}},{x:d,y:c},{x:d,y:c}],innerBackground:{fill:"rgba(100, 170, 255, 0.8)"}},shapeOptions:{strokeWidth:2}},e.annotationsOptions,e.bindings.pitchfork.annotationsOptions);return this.chart.addAnnotation(a)}},steps:[k.updateNthPoint(1),k.updateNthPoint(2)]},verticalCounter:{className:"highcharts-vertical-counter",start:function(a){a=k.attractToPoint(a,this.chart);var b=this.chart.options.navigation;a&&(this.verticalCounter=
this.verticalCounter||0,a=f({langKey:"verticalCounter",type:"verticalLine",typeOptions:{point:{x:a.x,y:a.y,xAxis:a.xAxis,yAxis:a.yAxis},label:{offset:a.below?40:-40,text:this.verticalCounter.toString()}},labelOptions:{style:{color:"#666666",fontSize:"11px"}},shapeOptions:{stroke:"rgba(0, 0, 0, 0.75)",strokeWidth:1}},b.annotationsOptions,b.bindings.verticalCounter.annotationsOptions),a=this.chart.addAnnotation(a),this.verticalCounter++,a.options.events.click.call(a,{}))}},timeCycles:{className:"highcharts-time-cycles",
start:function(a){a=k.attractToPoint(a,this.chart);var b=this.chart.options.navigation;if(a)return a=f({langKey:"timeCycles",type:"timeCycles",typeOptions:{xAxis:a.xAxis,yAxis:a.yAxis,points:[{x:a.x},{x:a.x}],line:{stroke:"rgba(0, 0, 0, 0.75)",fill:"transparent",strokeWidth:2}}},b.annotationsOptions,b.bindings.timeCycles.annotationsOptions),a=this.chart.addAnnotation(a),a.options.events.click.call(a,{}),a},steps:[k.updateNthPoint(1)]},verticalLabel:{className:"highcharts-vertical-label",start:function(a){a=
k.attractToPoint(a,this.chart);var b=this.chart.options.navigation;a&&(a=f({langKey:"verticalLabel",type:"verticalLine",typeOptions:{point:{x:a.x,y:a.y,xAxis:a.xAxis,yAxis:a.yAxis},label:{offset:a.below?40:-40}},labelOptions:{style:{color:"#666666",fontSize:"11px"}},shapeOptions:{stroke:"rgba(0, 0, 0, 0.75)",strokeWidth:1}},b.annotationsOptions,b.bindings.verticalLabel.annotationsOptions),a=this.chart.addAnnotation(a),a.options.events.click.call(a,{}))}},verticalArrow:{className:"highcharts-vertical-arrow",
start:function(a){a=k.attractToPoint(a,this.chart);var b=this.chart.options.navigation;a&&(a=f({langKey:"verticalArrow",type:"verticalLine",typeOptions:{point:{x:a.x,y:a.y,xAxis:a.xAxis,yAxis:a.yAxis},label:{offset:a.below?40:-40,format:" "},connector:{fill:"none",stroke:a.below?"#f21313":"#06b535"}},shapeOptions:{stroke:"rgba(0, 0, 0, 0.75)",strokeWidth:1}},b.annotationsOptions,b.bindings.verticalArrow.annotationsOptions),a=this.chart.addAnnotation(a),a.options.events.click.call(a,{}))}},fibonacciTimeZones:{className:"highcharts-fibonacci-time-zones",
start:function(a){var b=this.chart.pointer.getCoordinates(a);a=this.utils.getAssignedAxis(b.xAxis);b=this.utils.getAssignedAxis(b.yAxis);if(a&&b){var c=this.chart.options.navigation;a=f({type:"fibonacciTimeZones",langKey:"fibonacciTimeZones",typeOptions:{xAxis:a.axis.options.index,yAxis:b.axis.options.index,points:[{x:a.value}]}},c.annotationsOptions,c.bindings.fibonacciTimeZones.annotationsOptions);return this.chart.addAnnotation(a)}},steps:[function(a,b){var c=b.options.typeOptions.points;c=c&&
c[0].x;var d=this.chart.pointer.getCoordinates(a);a=this.utils.getAssignedAxis(d.xAxis);d=this.utils.getAssignedAxis(d.yAxis);b.update({typeOptions:{xAxis:a.axis.options.index,yAxis:d.axis.options.index,points:[{x:c},{x:a.value}]}})}]},flagCirclepin:{className:"highcharts-flag-circlepin",start:k.addFlagFromForm("circlepin")},flagDiamondpin:{className:"highcharts-flag-diamondpin",start:k.addFlagFromForm("flag")},flagSquarepin:{className:"highcharts-flag-squarepin",start:k.addFlagFromForm("squarepin")},
flagSimplepin:{className:"highcharts-flag-simplepin",start:k.addFlagFromForm("nopin")},zoomX:{className:"highcharts-zoom-x",init:function(a){this.chart.update({chart:{zoomType:"x"}});g(this,"deselectButton",{button:a})}},zoomY:{className:"highcharts-zoom-y",init:function(a){this.chart.update({chart:{zoomType:"y"}});g(this,"deselectButton",{button:a})}},zoomXY:{className:"highcharts-zoom-xy",init:function(a){this.chart.update({chart:{zoomType:"xy"}});g(this,"deselectButton",{button:a})}},seriesTypeLine:{className:"highcharts-series-type-line",
init:function(a){this.chart.series[0].update({type:"line",useOhlcData:!0});g(this,"deselectButton",{button:a})}},seriesTypeOhlc:{className:"highcharts-series-type-ohlc",init:function(a){this.chart.series[0].update({type:"ohlc"});g(this,"deselectButton",{button:a})}},seriesTypeCandlestick:{className:"highcharts-series-type-candlestick",init:function(a){this.chart.series[0].update({type:"candlestick"});g(this,"deselectButton",{button:a})}},seriesTypeHeikinAshi:{className:"highcharts-series-type-heikinashi",
init:function(a){this.chart.series[0].update({type:"heikinashi"});g(this,"deselectButton",{button:a})}},seriesTypeHLC:{className:"highcharts-series-type-hlc",init:function(a){this.chart.series[0].update({type:"hlc",useOhlcData:!0});g(this,"deselectButton",{button:a})}},seriesTypeHollowCandlestick:{className:"highcharts-series-type-hollowcandlestick",init:function(a){this.chart.series[0].update({type:"hollowcandlestick"});g(this,"deselectButton",{button:a})}},fullScreen:{className:"highcharts-full-screen",
noDataState:"normal",init:function(a){this.chart.fullscreen.toggle();g(this,"deselectButton",{button:a})}},currentPriceIndicator:{className:"highcharts-current-price-indicator",init:function(a){var b=this.chart,c=b.series,d=b.stockTools,e=k.isPriceIndicatorEnabled(b.series);d&&d.guiEnabled&&(c.forEach(function(a){a.update({lastPrice:{enabled:!e},lastVisiblePrice:{enabled:!e,label:{enabled:!0}}},!1)}),b.redraw());g(this,"deselectButton",{button:a})}},indicators:{className:"highcharts-indicators",init:function(){var a=
this;g(a,"showPopup",{formType:"indicators",options:{},onSubmit:function(b){a.utils.manageIndicators.call(a,b)}})}},toggleAnnotations:{className:"highcharts-toggle-annotations",init:function(a){var b=this.chart,c=b.stockTools,d=c.getIconsURL();this.toggledAnnotations=!this.toggledAnnotations;(b.annotations||[]).forEach(function(a){a.setVisibility(!this.toggledAnnotations)},this);c&&c.guiEnabled&&(a.firstChild.style["background-image"]=this.toggledAnnotations?'url("'+d+'annotations-hidden.svg")':'url("'+
d+'annotations-visible.svg")');g(this,"deselectButton",{button:a})}},saveChart:{className:"highcharts-save-chart",noDataState:"normal",init:function(a){var b=this.chart,c=[],d=[],e=[],f=[];b.annotations.forEach(function(a,b){c[b]=a.userOptions});b.series.forEach(function(a){a.is("sma")?d.push(a.userOptions):"flags"===a.type&&e.push(a.userOptions)});b.yAxis.forEach(function(a){k.isNotNavigatorYAxis(a)&&f.push(a.options)});n.win.localStorage.setItem("highcharts-chart",JSON.stringify({annotations:c,
indicators:d,flags:e,yAxes:f}));g(this,"deselectButton",{button:a})}}};a({navigation:{bindings:m}});r.prototype.utils=f(k,r.prototype.utils)});w(a,"Stock/StockToolsGui.js",[a["Core/Chart/Chart.js"],a["Core/Globals.js"],a["Extensions/Annotations/NavigationBindings.js"],a["Core/DefaultOptions.js"],a["Core/Utilities.js"]],function(a,n,r,v,m){v=v.setOptions;var c=m.addEvent,b=m.createElement,w=m.css,e=m.extend,g=m.fireEvent,t=m.getStyle,f=m.isArray,u=m.merge,x=m.pick;v({lang:{stockTools:{gui:{simpleShapes:"Simple shapes",
lines:"Lines",crookedLines:"Crooked lines",measure:"Measure",advanced:"Advanced",toggleAnnotations:"Toggle annotations",verticalLabels:"Vertical labels",flags:"Flags",zoomChange:"Zoom change",typeChange:"Type change",saveChart:"Save chart",indicators:"Indicators",currentPriceIndicator:"Current Price Indicators",zoomX:"Zoom X",zoomY:"Zoom Y",zoomXY:"Zooom XY",fullScreen:"Fullscreen",typeOHLC:"OHLC",typeLine:"Line",typeCandlestick:"Candlestick",typeHLC:"HLC",typeHollowCandlestick:"Hollow Candlestick",
typeHeikinAshi:"Heikin Ashi",circle:"Circle",ellipse:"Ellipse",label:"Label",rectangle:"Rectangle",flagCirclepin:"Flag circle",flagDiamondpin:"Flag diamond",flagSquarepin:"Flag square",flagSimplepin:"Flag simple",measureXY:"Measure XY",measureX:"Measure X",measureY:"Measure Y",segment:"Segment",arrowSegment:"Arrow segment",ray:"Ray",arrowRay:"Arrow ray",line:"Line",arrowInfinityLine:"Arrow line",horizontalLine:"Horizontal line",verticalLine:"Vertical line",infinityLine:"Infinity line",crooked3:"Crooked 3 line",
crooked5:"Crooked 5 line",elliott3:"Elliott 3 line",elliott5:"Elliott 5 line",verticalCounter:"Vertical counter",verticalLabel:"Vertical label",verticalArrow:"Vertical arrow",fibonacci:"Fibonacci",fibonacciTimeZones:"Fibonacci Time Zones",pitchfork:"Pitchfork",parallelChannel:"Parallel channel",timeCycles:"Time Cycles"}},navigation:{popup:{circle:"Circle",ellipse:"Ellipse",rectangle:"Rectangle",label:"Label",segment:"Segment",arrowSegment:"Arrow segment",ray:"Ray",arrowRay:"Arrow ray",line:"Line",
arrowInfinityLine:"Arrow line",horizontalLine:"Horizontal line",verticalLine:"Vertical line",crooked3:"Crooked 3 line",crooked5:"Crooked 5 line",elliott3:"Elliott 3 line",elliott5:"Elliott 5 line",verticalCounter:"Vertical counter",verticalLabel:"Vertical label",verticalArrow:"Vertical arrow",fibonacci:"Fibonacci",fibonacciTimeZones:"Fibonacci Time Zones",pitchfork:"Pitchfork",parallelChannel:"Parallel channel",infinityLine:"Infinity line",measure:"Measure",measureXY:"Measure XY",measureX:"Measure X",
measureY:"Measure Y",timeCycles:"Time Cycles",flags:"Flags",addButton:"add",saveButton:"save",editButton:"edit",removeButton:"remove",series:"Series",volume:"Volume",connector:"Connector",innerBackground:"Inner background",outerBackground:"Outer background",crosshairX:"Crosshair X",crosshairY:"Crosshair Y",tunnel:"Tunnel",background:"Background",noFilterMatch:"No match",searchIndicators:"Search Indicators",clearFilter:"\u2715 clear filter",index:"Index",period:"Period",periods:"Periods",standardDeviation:"Standard deviation",
periodTenkan:"Tenkan period",periodSenkouSpanB:"Senkou Span B period",periodATR:"ATR period",multiplierATR:"ATR multiplier",shortPeriod:"Short period",longPeriod:"Long period",signalPeriod:"Signal period",decimals:"Decimals",algorithm:"Algorithm",topBand:"Top band",bottomBand:"Bottom band",initialAccelerationFactor:"Initial acceleration factor",maxAccelerationFactor:"Max acceleration factor",increment:"Increment",multiplier:"Multiplier",ranges:"Ranges",highIndex:"High index",lowIndex:"Low index",
deviation:"Deviation",xAxisUnit:"x-axis unit",factor:"Factor",fastAvgPeriod:"Fast average period",slowAvgPeriod:"Slow average period",average:"Average",indicatorAliases:{abands:["Acceleration Bands"],bb:["Bollinger Bands"],dema:["Double Exponential Moving Average"],ema:["Exponential Moving Average"],ikh:["Ichimoku Kinko Hyo"],keltnerchannels:["Keltner Channels"],linearRegression:["Linear Regression"],pivotpoints:["Pivot Points"],pc:["Price Channel"],priceenvelopes:["Price Envelopes"],psar:["Parabolic SAR"],
sma:["Simple Moving Average"],supertrend:["Super Trend"],tema:["Triple Exponential Moving Average"],vbp:["Volume by Price"],vwap:["Volume Weighted Moving Average"],wma:["Weighted Moving Average"],zigzag:["Zig Zag"],apo:["Absolute price indicator"],ad:["Accumulation/Distribution"],aroon:["Aroon"],aroonoscillator:["Aroon oscillator"],atr:["Average True Range"],ao:["Awesome oscillator"],cci:["Commodity Channel Index"],chaikin:["Chaikin"],cmf:["Chaikin Money Flow"],cmo:["Chande Momentum Oscillator"],
disparityindex:["Disparity Index"],dmi:["Directional Movement Index"],dpo:["Detrended price oscillator"],klinger:["Klinger Oscillator"],linearRegressionAngle:["Linear Regression Angle"],linearRegressionIntercept:["Linear Regression Intercept"],linearRegressionSlope:["Linear Regression Slope"],macd:["Moving Average Convergence Divergence"],mfi:["Money Flow Index"],momentum:["Momentum"],natr:["Normalized Average True Range"],obv:["On-Balance Volume"],ppo:["Percentage Price oscillator"],roc:["Rate of Change"],
rsi:["Relative Strength Index"],slowstochastic:["Slow Stochastic"],stochastic:["Stochastic"],trix:["TRIX"],williamsr:["Williams %R"]}}}},stockTools:{gui:{enabled:!0,className:"highcharts-bindings-wrapper",toolbarClassName:"stocktools-toolbar",buttons:"indicators separator simpleShapes lines crookedLines measure advanced toggleAnnotations separator verticalLabels flags separator zoomChange fullScreen typeChange separator currentPriceIndicator saveChart".split(" "),definitions:{separator:{symbol:"separator.svg"},
simpleShapes:{items:["label","circle","ellipse","rectangle"],circle:{symbol:"circle.svg"},ellipse:{symbol:"ellipse.svg"},rectangle:{symbol:"rectangle.svg"},label:{symbol:"label.svg"}},flags:{items:["flagCirclepin","flagDiamondpin","flagSquarepin","flagSimplepin"],flagSimplepin:{symbol:"flag-basic.svg"},flagDiamondpin:{symbol:"flag-diamond.svg"},flagSquarepin:{symbol:"flag-trapeze.svg"},flagCirclepin:{symbol:"flag-elipse.svg"}},lines:{items:"segment arrowSegment ray arrowRay line arrowInfinityLine horizontalLine verticalLine".split(" "),
segment:{symbol:"segment.svg"},arrowSegment:{symbol:"arrow-segment.svg"},ray:{symbol:"ray.svg"},arrowRay:{symbol:"arrow-ray.svg"},line:{symbol:"line.svg"},arrowInfinityLine:{symbol:"arrow-line.svg"},verticalLine:{symbol:"vertical-line.svg"},horizontalLine:{symbol:"horizontal-line.svg"}},crookedLines:{items:["elliott3","elliott5","crooked3","crooked5"],crooked3:{symbol:"crooked-3.svg"},crooked5:{symbol:"crooked-5.svg"},elliott3:{symbol:"elliott-3.svg"},elliott5:{symbol:"elliott-5.svg"}},verticalLabels:{items:["verticalCounter",
"verticalLabel","verticalArrow"],verticalCounter:{symbol:"vertical-counter.svg"},verticalLabel:{symbol:"vertical-label.svg"},verticalArrow:{symbol:"vertical-arrow.svg"}},advanced:{items:["fibonacci","fibonacciTimeZones","pitchfork","parallelChannel","timeCycles"],pitchfork:{symbol:"pitchfork.svg"},fibonacci:{symbol:"fibonacci.svg"},fibonacciTimeZones:{symbol:"fibonacci-timezone.svg"},parallelChannel:{symbol:"parallel-channel.svg"},timeCycles:{symbol:"time-cycles.svg"}},measure:{items:["measureXY",
"measureX","measureY"],measureX:{symbol:"measure-x.svg"},measureY:{symbol:"measure-y.svg"},measureXY:{symbol:"measure-xy.svg"}},toggleAnnotations:{symbol:"annotations-visible.svg"},currentPriceIndicator:{symbol:"current-price-show.svg"},indicators:{symbol:"indicators.svg"},zoomChange:{items:["zoomX","zoomY","zoomXY"],zoomX:{symbol:"zoom-x.svg"},zoomY:{symbol:"zoom-y.svg"},zoomXY:{symbol:"zoom-xy.svg"}},typeChange:{items:"typeOHLC typeLine typeCandlestick typeHollowCandlestick typeHLC typeHeikinAshi".split(" "),
typeOHLC:{symbol:"series-ohlc.svg"},typeLine:{symbol:"series-line.svg"},typeCandlestick:{symbol:"series-candlestick.svg"},typeHLC:{symbol:"series-hlc.svg"},typeHeikinAshi:{symbol:"series-heikin-ashi.svg"},typeHollowCandlestick:{symbol:"series-hollow-candlestick.svg"}},fullScreen:{symbol:"fullscreen.svg"},saveChart:{symbol:"save-chart.svg"}}}}});c(a,"afterGetContainer",function(){this.setStockTools()});c(a,"getMargins",function(){var a=this.stockTools&&this.stockTools.listWrapper;(a=a&&(a.startWidth+
t(a,"padding-left")+t(a,"padding-right")||a.offsetWidth))&&a<this.plotWidth&&(this.plotLeft+=a,this.spacing[3]+=a)},{order:0});["beforeRender","beforeRedraw"].forEach(function(b){c(a,b,function(){if(this.stockTools){var a=this.options.chart,b=this.stockTools.listWrapper;b=b&&(b.startWidth+t(b,"padding-left")+t(b,"padding-right")||b.offsetWidth);var c=!1;b&&b<this.plotWidth?(a=x(a.spacingLeft,a.spacing&&a.spacing[3],0)+b,c=a-this.spacingBox.x,this.spacingBox.x=a,this.spacingBox.width-=c,c=!0):0===
b&&(c=!0);b!==this.stockTools.prevOffsetWidth&&(this.stockTools.prevOffsetWidth=b,c&&(this.isDirtyLegend=!0))}})});c(a,"destroy",function(){this.stockTools&&this.stockTools.destroy()});c(a,"redraw",function(){this.stockTools&&this.stockTools.guiEnabled&&this.stockTools.redraw()});var k=function(){function a(a,b,c){this.wrapper=this.toolbar=this.submenu=this.showhideBtn=this.listWrapper=this.arrowWrapper=this.arrowUp=this.arrowDown=void 0;this.chart=c;this.options=a;this.lang=b;this.iconsURL=this.getIconsURL();
this.guiEnabled=a.enabled;this.visible=x(a.visible,!0);this.placed=x(a.placed,!1);this.eventsToUnbind=[];this.guiEnabled&&(this.createHTML(),this.init(),this.showHideNavigatorion());g(this,"afterInit")}a.prototype.init=function(){var a=this,b=this.lang,d=this.options,e=this.toolbar,k=a.addSubmenu,g=d.definitions,m=e.childNodes,n;d.buttons.forEach(function(d){n=a.addButton(e,g,d,b);a.eventsToUnbind.push(c(n.buttonWrapper,"click",function(){a.eraseActiveButtons(m,n.buttonWrapper)}));f(g[d].items)&&
k.call(a,n,g[d])})};a.prototype.addSubmenu=function(a,d){var e=this,f=a.submenuArrow,h=a.buttonWrapper,l=t(h,"width"),k=this.wrapper,g=this.listWrapper,m=this.toolbar.childNodes,n=0,p;this.submenu=p=b("ul",{className:"highcharts-submenu-wrapper"},void 0,h);this.addSubmenuItems(h,d);e.eventsToUnbind.push(c(f,"click",function(a){a.stopPropagation();e.eraseActiveButtons(m,h);0<=h.className.indexOf("highcharts-current")?(g.style.width=g.startWidth+"px",h.classList.remove("highcharts-current"),p.style.display=
"none"):(p.style.display="block",n=p.offsetHeight-h.offsetHeight-3,p.offsetHeight+h.offsetTop>k.offsetHeight&&h.offsetTop>n||(n=0),w(p,{top:-n+"px",left:l+3+"px"}),h.className+=" highcharts-current",g.startWidth=k.offsetWidth,g.style.width=g.startWidth+t(g,"padding-left")+p.offsetWidth+3+"px")}))};a.prototype.addSubmenuItems=function(a,b){var d=this,e=this.submenu,h=this.lang,f=this.listWrapper,l;b.items.forEach(function(g){l=d.addButton(e,b,g,h);d.eventsToUnbind.push(c(l.mainButton,"click",function(){d.switchSymbol(this,
a,!0);f.style.width=f.startWidth+"px";e.style.display="none"}))});var g=e.querySelectorAll("li > .highcharts-menu-item-btn")[0];d.switchSymbol(g,!1)};a.prototype.eraseActiveButtons=function(a,b,c){[].forEach.call(a,function(a){a!==b&&(a.classList.remove("highcharts-current"),a.classList.remove("highcharts-active"),c=a.querySelectorAll(".highcharts-submenu-wrapper"),0<c.length&&(c[0].style.display="none"))})};a.prototype.addButton=function(c,d,e,f){void 0===f&&(f={});d=d[e];var h=d.items,g=d.className||
"";e=b("li",{className:x(a.prototype.classMapping[e],"")+" "+g,title:f[e]||e},void 0,c);c=b("span",{className:"highcharts-menu-item-btn"},void 0,e);if(h&&h.length){var l=b("span",{className:"highcharts-submenu-item-arrow highcharts-arrow-right"},void 0,e);l.style.backgroundImage="url("+this.iconsURL+"arrow-bottom.svg)"}else c.style.backgroundImage="url("+this.iconsURL+d.symbol+")";return{buttonWrapper:e,mainButton:c,submenuArrow:l}};a.prototype.addNavigation=function(){var a=this.wrapper;this.arrowWrapper=
b("div",{className:"highcharts-arrow-wrapper"});this.arrowUp=b("div",{className:"highcharts-arrow-up"},void 0,this.arrowWrapper);this.arrowUp.style.backgroundImage="url("+this.iconsURL+"arrow-right.svg)";this.arrowDown=b("div",{className:"highcharts-arrow-down"},void 0,this.arrowWrapper);this.arrowDown.style.backgroundImage="url("+this.iconsURL+"arrow-right.svg)";a.insertBefore(this.arrowWrapper,a.childNodes[0]);this.scrollButtons()};a.prototype.scrollButtons=function(){var a=0,b=this.wrapper,d=this.toolbar,
e=.1*b.offsetHeight;this.eventsToUnbind.push(c(this.arrowUp,"click",function(){0<a&&(a-=e,d.style.marginTop=-a+"px")}));this.eventsToUnbind.push(c(this.arrowDown,"click",function(){b.offsetHeight+a<=d.offsetHeight+e&&(a+=e,d.style.marginTop=-a+"px")}))};a.prototype.createHTML=function(){var a=this.chart,d=this.options,e=a.container,f=a.options.navigation,g=this.wrapper=b("div",{className:"highcharts-stocktools-wrapper "+d.className+" "+(f&&f.bindingsClassName)});e.appendChild(g);["mousedown","mousemove",
"click","touchstart"].forEach(function(a){c(g,a,function(a){return a.stopPropagation()})});c(g,"mouseover",function(b){return a.pointer.onContainerMouseLeave(b)});this.toolbar=e=b("ul",{className:"highcharts-stocktools-toolbar "+d.toolbarClassName});this.listWrapper=d=b("div",{className:"highcharts-menu-wrapper"});g.insertBefore(d,g.childNodes[0]);d.insertBefore(e,d.childNodes[0]);this.showHideToolbar();this.addNavigation()};a.prototype.showHideNavigatorion=function(){this.visible&&this.toolbar.offsetHeight>
this.wrapper.offsetHeight-50?this.arrowWrapper.style.display="block":(this.toolbar.style.marginTop="0px",this.arrowWrapper.style.display="none")};a.prototype.showHideToolbar=function(){var a=this.chart,d=this.wrapper,e=this.listWrapper,f=this.submenu,g=this.visible,k;this.showhideBtn=k=b("div",{className:"highcharts-toggle-toolbar highcharts-arrow-left"},void 0,d);k.style.backgroundImage="url("+this.iconsURL+"arrow-right.svg)";g?(d.style.height="100%",k.style.top=t(e,"padding-top")+"px",k.style.left=
d.offsetWidth+t(e,"padding-left")+"px"):(f&&(f.style.display="none"),k.style.left="0px",this.visible=g=!1,e.classList.add("highcharts-hide"),k.classList.toggle("highcharts-arrow-right"),d.style.height=k.offsetHeight+"px");this.eventsToUnbind.push(c(k,"click",function(){a.update({stockTools:{gui:{visible:!g,placed:!0}}})}))};a.prototype.switchSymbol=function(a,b){var c=a.parentNode,d=c.className;c=c.parentNode.parentNode;-1<d.indexOf("highcharts-disabled-btn")||(c.className="",d&&c.classList.add(d.trim()),
c.querySelectorAll(".highcharts-menu-item-btn")[0].style.backgroundImage=a.style.backgroundImage,b&&this.toggleButtonAciveClass(c))};a.prototype.toggleButtonAciveClass=function(a){0<=a.className.indexOf("highcharts-active")?a.classList.remove("highcharts-active"):a.classList.add("highcharts-active")};a.prototype.unselectAllButtons=function(a){var b=a.parentNode.querySelectorAll(".highcharts-active");[].forEach.call(b,function(b){b!==a&&b.classList.remove("highcharts-active")})};a.prototype.update=
function(a,b){u(!0,this.chart.options.stockTools,a);this.destroy();this.chart.setStockTools(a);this.chart.navigationBindings&&this.chart.navigationBindings.update();this.chart.isDirtyBox=!0;x(b,!0)&&this.chart.redraw()};a.prototype.destroy=function(){var a=this.wrapper,b=a&&a.parentNode;this.eventsToUnbind.forEach(function(a){a()});b&&b.removeChild(a)};a.prototype.redraw=function(){this.showHideNavigatorion()};a.prototype.getIconsURL=function(){return this.chart.options.navigation.iconsURL||this.options.iconsURL||
"https://code.highcharts.com/9.3.1/gfx/stock-icons/"};return a}();k.prototype.classMapping={circle:"highcharts-circle-annotation",ellipse:"highcharts-ellipse-annotation",rectangle:"highcharts-rectangle-annotation",label:"highcharts-label-annotation",segment:"highcharts-segment",arrowSegment:"highcharts-arrow-segment",ray:"highcharts-ray",arrowRay:"highcharts-arrow-ray",line:"highcharts-infinity-line",arrowInfinityLine:"highcharts-arrow-infinity-line",verticalLine:"highcharts-vertical-line",horizontalLine:"highcharts-horizontal-line",
crooked3:"highcharts-crooked3",crooked5:"highcharts-crooked5",elliott3:"highcharts-elliott3",elliott5:"highcharts-elliott5",pitchfork:"highcharts-pitchfork",fibonacci:"highcharts-fibonacci",fibonacciTimeZones:"highcharts-fibonacci-time-zones",parallelChannel:"highcharts-parallel-channel",measureX:"highcharts-measure-x",measureY:"highcharts-measure-y",measureXY:"highcharts-measure-xy",timeCycles:"highcharts-time-cycles",verticalCounter:"highcharts-vertical-counter",verticalLabel:"highcharts-vertical-label",
verticalArrow:"highcharts-vertical-arrow",currentPriceIndicator:"highcharts-current-price-indicator",indicators:"highcharts-indicators",flagCirclepin:"highcharts-flag-circlepin",flagDiamondpin:"highcharts-flag-diamondpin",flagSquarepin:"highcharts-flag-squarepin",flagSimplepin:"highcharts-flag-simplepin",zoomX:"highcharts-zoom-x",zoomY:"highcharts-zoom-y",zoomXY:"highcharts-zoom-xy",typeLine:"highcharts-series-type-line",typeOHLC:"highcharts-series-type-ohlc",typeHLC:"highcharts-series-type-hlc",
typeCandlestick:"highcharts-series-type-candlestick",typeHollowCandlestick:"highcharts-series-type-hollowcandlestick",typeHeikinAshi:"highcharts-series-type-heikinashi",fullScreen:"highcharts-full-screen",toggleAnnotations:"highcharts-toggle-annotations",saveChart:"highcharts-save-chart",separator:"highcharts-separator"};e(a.prototype,{setStockTools:function(a){var b=this.options,c=b.lang;a=u(b.stockTools&&b.stockTools.gui,a&&a.gui);this.stockTools=new k(a,c&&c.stockTools&&c.stockTools.gui,this);
this.stockTools.guiEnabled&&(this.isDirtyBox=!0)}});c(r,"selectButton",function(a){var b=a.button,c=this.chart.stockTools;c&&c.guiEnabled&&(c.unselectAllButtons(a.button),0<=b.parentNode.className.indexOf("highcharts-submenu-wrapper")&&(b=b.parentNode.parentNode),c.toggleButtonAciveClass(b))});c(r,"deselectButton",function(a){a=a.button;var b=this.chart.stockTools;b&&b.guiEnabled&&(0<=a.parentNode.className.indexOf("highcharts-submenu-wrapper")&&(a=a.parentNode.parentNode),b.toggleButtonAciveClass(a))});
c(a,"render",function(){var a=this.stockTools,b=a&&a.toolbar&&a.toolbar.querySelector(".highcharts-current-price-indicator");a&&this.navigationBindings&&this.options.series&&b&&(this.navigationBindings.constructor.prototype.utils.isPriceIndicatorEnabled(this.series)?b.firstChild.style["background-image"]='url("'+a.getIconsURL()+'current-price-hide.svg")':b.firstChild.style["background-image"]='url("'+a.getIconsURL()+'current-price-show.svg")')});n.Toolbar=k;return n.Toolbar});w(a,"masters/modules/stock-tools.src.js",
[],function(){})});
//# sourceMappingURL=stock-tools.js.map
```
