## Supplementary material for "Exploring natural odour landscapes: A case study with implications for human-biting insects": Figure S4: stock.js

```
/*
 Highstock JS v9.3.1 (2021-11-05)

 Highcharts Stock as a plugin for Highcharts

 (c) 2010-2021 Torstein Honsi

 License: www.highcharts.com/license
*/
'use strict';(function(a){"object"===typeof module&&module.exports?(a["default"]=a,module.exports=a):"function"===typeof define&&define.amd?define("highcharts/modules/stock",["highcharts"],function(L){a(L);a.Highcharts=L;return a}):a("undefined"!==typeof Highcharts?Highcharts:void 0)})(function(a){function L(a,r,y,t){a.hasOwnProperty(r)||(a[r]=t.apply(null,y))}a=a?a._modules:{};L(a,"Core/Axis/OrdinalAxis.js",[a["Core/Axis/Axis.js"],a["Core/Globals.js"],a["Core/Series/Series.js"],a["Core/Utilities.js"]],
function(a,r,y,t){var l=t.addEvent,D=t.correctFloat,E=t.css,x=t.defined,b=t.error,m=t.pick,d=t.timeUnits,z=[],h;(function(h){function G(f,e,c,n,p,g,v){void 0===p&&(p=[]);void 0===g&&(g=0);var w={},Q=this.options.tickPixelInterval,O=this.chart.time,B=[],H,A,k=0,u=[],h=-Number.MAX_VALUE;if(!this.options.ordinal&&!this.options.breaks||!p||3>p.length||"undefined"===typeof e)return O.getTimeTicks.apply(O,arguments);var m=p.length;for(H=0;H<m;H++){var F=H&&p[H-1]>c;p[H]<e&&(k=H);if(H===m-1||p[H+1]-p[H]>
5*g||F){if(p[H]>h){for(A=O.getTimeTicks(f,p[k],p[H],n);A.length&&A[0]<=h;)A.shift();A.length&&(h=A[A.length-1]);B.push(u.length);u=u.concat(A)}k=H+1}if(F)break}if(A){A=A.info;if(v&&A.unitRange<=d.hour){H=u.length-1;for(k=1;k<H;k++)if(O.dateFormat("%d",u[k])!==O.dateFormat("%d",u[k-1])){w[u[k]]="day";var q=!0}q&&(w[u[0]]="day");A.higherRanks=w}A.segmentStarts=B;u.info=A}else b(12,!1,this.chart);if(v&&x(Q)){A=u.length;O=[];k=[];H=void 0;for(q=A;q--;)B=this.translate(u[q]),H&&(k[q]=H-B),O[q]=H=B;k.sort();
k=k[Math.floor(k.length/2)];k<.6*Q&&(k=null);q=u[A-1]>c?A-1:A;for(H=void 0;q--;)B=O[q],A=Math.abs(H-B),H&&A<.8*Q&&(null===k||A<.8*k)?(w[u[q]]&&!w[u[q+1]]?(A=q+1,H=B):A=q,u.splice(A,1)):H=B}return u}function I(f){var e=this.ordinal.positions;if(!e)return f;var c=e.length-1;if(0>f)f=e[0];else if(f>c)f=e[c];else{c=Math.floor(f);var n=f-c}return"undefined"!==typeof n&&"undefined"!==typeof e[c]?e[c]+(n?n*(e[c+1]-e[c]):0):f}function C(f){var e=this.ordinal,c=e.positions;if(!c)return f;var n=(f-(this.old?
this.old.min:this.min))*(this.old?this.old.transA:this.transA)+this.minPixelPadding;0<n&&n<this.left+this.len||(e.extendedOrdinalPositions||(e.extendedOrdinalPositions=e.getExtendedPositions()),c=e.extendedOrdinalPositions);if(c&&c.length){f=e.getIndexOfPoint(n,c);e=D(f%1);if(0<=f&&f<c.length-1)return c[Math.floor(f)]+e*(c[Math.ceil(f)]-c[Math.floor(f)]);e=c.length;n=c[0];c=c[e-1];var p=(c-n)/(e-1);return 0>f?n+p*f:c+p*(f-e)}return f}function q(f,e){var c=h.Additions.findIndexOf(f,e,!0);return f[c]===
e?c:c+(e-f[c])/(f[c+1]-f[c])}function K(){this.ordinal||(this.ordinal=new h.Additions(this))}function a(){this.isXAxis&&x(this.options.overscroll)&&this.max===this.dataMax&&(!this.chart.mouseIsDown||this.isInternal)&&(!this.eventArgs||this.eventArgs&&"navigator"!==this.eventArgs.trigger)&&(this.max+=this.options.overscroll,!this.isInternal&&x(this.userMin)&&(this.min+=this.options.overscroll))}function F(){this.horiz&&!this.isDirty&&(this.isDirty=this.isOrdinal&&this.chart.navigator&&!this.chart.navigator.adaptToUpdatedData)}
function N(){this.ordinal&&(this.ordinal.beforeSetTickPositions(),this.tickInterval=this.ordinal.postProcessTickInterval(this.tickInterval))}function u(f){var e=this.xAxis[0],c=e.options.overscroll,n=f.originalEvent.chartX,p=this.options.chart.panning,g=!1;if(p&&"y"!==p.type&&e.options.ordinal&&e.series.length){var v=this.mouseDownX,w=e.getExtremes(),Q=w.dataMax,O=w.min,B=w.max,H=this.hoverPoints,A=e.closestPointRange||e.ordinal&&e.ordinal.overscrollPointsRange;v=(v-n)/(e.translationSlope*(e.ordinal.slope||
A));A={ordinal:{positions:e.ordinal.getExtendedPositions()}};var k=e.index2val,u=e.val2lin,d=void 0,h=d=void 0,q=void 0;A.ordinal.positions?1<Math.abs(v)&&(H&&H.forEach(function(g){g.setState()}),0>v?(h=A,q=e.ordinal.positions?e:A):(h=e.ordinal.positions?e:A,q=A),d=q.ordinal.positions,Q>d[d.length-1]&&d.push(Q),this.fixedRange=B-O,d=e.navigatorAxis.toFixedRange(null,null,k.apply(h,[u.apply(h,[O,!0])+v]),k.apply(q,[u.apply(q,[B,!0])+v])),d.min>=Math.min(w.dataMin,O)&&d.max<=Math.max(Q,B)+c&&e.setExtremes(d.min,
d.max,!0,!1,{trigger:"pan"}),this.mouseDownX=n,E(this.container,{cursor:"move"})):g=!0}else g=!0;g||p&&/y/.test(p.type)?c&&(e.max=e.dataMax+c):f.preventDefault()}function M(){var f=this.xAxis;f&&f.options.ordinal&&(delete f.ordinal.index,delete f.ordinal.extendedOrdinalPositions)}function k(f,e){var c=this.ordinal,n=c.positions,p=c.slope,g=c.extendedOrdinalPositions;if(!n)return f;var v=n.length;if(n[0]<=f&&n[v-1]>=f)f=q(n,f);else{g||(g=c.getExtendedPositions&&c.getExtendedPositions(),c.extendedOrdinalPositions=
g);if(!g||!g.length)return f;v=g.length;p||(p=(g[v-1]-g[0])/v);n=q(g,n[0]);f>=g[0]&&f<=g[v-1]?f=q(g,f)-n:f<g[0]?(f=g[0]-f,f=-n-f/p):(f-=g[v-1],f=f/p+v-n)}return e?f:p*(f||0)+c.offset}h.compose=function(f,e,c){if(-1===z.indexOf(f)){z.push(f);var n=f.prototype;n.getTimeTicks=G;n.index2val=I;n.lin2val=C;n.val2lin=k;n.ordinal2lin=n.val2lin;l(f,"afterInit",K);l(f,"foundExtremes",a);l(f,"afterSetScale",F);l(f,"initialAxisTranslation",N)}-1===z.indexOf(c)&&(z.push(c),l(c,"pan",u));-1===z.indexOf(e)&&(z.push(e),
l(e,"updatedData",M));return f};var J=function(){function f(e){this.index={};this.axis=e}f.prototype.beforeSetTickPositions=function(){var e=this.axis,c=e.ordinal,n=e.getExtremes(),p=n.min,g=n.max,v=e.isXAxis&&!!e.options.breaks;n=e.options.ordinal;var w=e.chart.options.chart.ignoreHiddenSeries,f,O,B=[],H=Number.MAX_VALUE,A=!1;if(n||v){e.series.forEach(function(g,c){f=[];if(!(w&&!1===g.visible||!1===g.takeOrdinalPosition&&!v)&&(B=B.concat(g.processedXData),k=B.length,B.sort(function(g,c){return g-
c}),H=Math.min(H,m(g.closestPointRange,H)),k)){for(c=0;c<k-1;)B[c]!==B[c+1]&&f.push(B[c+1]),c++;f[0]!==B[0]&&f.unshift(B[0]);B=f}});var k=B.length;if(2<k){var d=B[1]-B[0];for(O=k-1;O--&&!A;)B[O+1]-B[O]!==d&&(A=!0);!e.options.keepOrdinalPadding&&(B[0]-p>d||g-B[B.length-1]>d)&&(A=!0)}else e.options.overscroll&&(2===k?H=B[1]-B[0]:1===k?(H=e.options.overscroll,B=[B[0],B[0]+H]):H=c.overscrollPointsRange);A||e.forceOrdinal?(e.options.overscroll&&(c.overscrollPointsRange=H,B=B.concat(c.getOverscrollPositions())),
c.positions=B,d=e.ordinal2lin(Math.max(p,B[0]),!0),O=Math.max(e.ordinal2lin(Math.min(g,B[B.length-1]),!0),1),c.slope=g=(g-p)/(O-d),c.offset=p-d*g):(c.overscrollPointsRange=m(e.closestPointRange,c.overscrollPointsRange),c.positions=e.ordinal.slope=c.offset=void 0)}e.isOrdinal=n&&A;c.groupIntervalFactor=null};f.findIndexOf=function(e,c,n){for(var p=0,g=e.length-1,v;p<g;)v=Math.ceil((p+g)/2),e[v]<=c?p=v:g=v-1;return e[p]===c?p:n?p:-1};f.prototype.getExtendedPositions=function(){var e=this,c=e.axis,n=
c.constructor.prototype,p=c.chart,g=c.series[0].currentDataGrouping,v=g?g.count+g.unitName:"raw",w=c.options.overscroll,f=c.getExtremes(),k=void 0,B=e.index;B||(B=e.index={});if(!B[v]){var H={series:[],chart:p,forceOrdinal:!1,getExtremes:function(){return{min:f.dataMin,max:f.dataMax+w}},getGroupPixelWidth:n.getGroupPixelWidth,getTimeTicks:n.getTimeTicks,options:{ordinal:!0},ordinal:{getGroupIntervalFactor:this.getGroupIntervalFactor},ordinal2lin:n.ordinal2lin,getIndexOfPoint:n.getIndexOfPoint,val2lin:n.val2lin};
H.ordinal.axis=H;c.series.forEach(function(c){k={xAxis:H,xData:c.xData.slice(),chart:p,destroyGroupedData:r.noop,getProcessedData:y.prototype.getProcessedData,applyGrouping:y.prototype.applyGrouping};k.xData=k.xData.concat(e.getOverscrollPositions());k.options={dataGrouping:g?{firstAnchor:"firstPoint",anchor:"middle",lastAnchor:"lastPoint",enabled:!0,forced:!0,approximation:"open",units:[[g.unitName,[g.count]]]}:{enabled:!1}};H.series.push(k);c.processData.apply(k)});k.closestPointRange!==k.basePointRange&&
k.currentDataGrouping&&(H.forceOrdinal=!0);c.ordinal.beforeSetTickPositions.apply({axis:H});B[v]=H.ordinal.positions}return B[v]};f.prototype.getGroupIntervalFactor=function(e,c,n){n=n.processedXData;var p=n.length,g=[];var v=this.groupIntervalFactor;if(!v){for(v=0;v<p-1;v++)g[v]=n[v+1]-n[v];g.sort(function(g,c){return g-c});g=g[Math.floor(p/2)];e=Math.max(e,n[0]);c=Math.min(c,n[p-1]);this.groupIntervalFactor=v=p*g/(c-e)}return v};f.prototype.getIndexOfPoint=function(e,c){var n=this.axis,p=this.positions?
this.positions[0]:0,g=n.series[0].points&&n.series[0].points[0]&&n.series[0].points[0].plotX||n.minPixelPadding;1<n.series.length&&n.series.forEach(function(c){c.points&&x(c.points[0])&&x(c.points[0].plotX)&&c.points[0].plotX<g&&(g=c.points[0].plotX)});e=(e-g)/(n.translationSlope*(this.slope||n.closestPointRange||this.overscrollPointsRange));return f.findIndexOf(c,p)+e};f.prototype.getOverscrollPositions=function(){var e=this.axis,c=e.options.overscroll,n=this.overscrollPointsRange,p=[],g=e.dataMax;
if(x(n))for(;g<=e.dataMax+c;)g+=n,p.push(g);return p};f.prototype.postProcessTickInterval=function(e){var c=this.axis,n=this.slope;return n?c.options.breaks?c.closestPointRange||e:e/(n/c.closestPointRange):e};return f}();h.Additions=J})(h||(h={}));return h});L(a,"Series/DataModifyComposition.js",[a["Core/Axis/Axis.js"],a["Core/Series/Point.js"],a["Core/Series/Series.js"],a["Core/Utilities.js"]],function(a,r,y,t){var l=r.prototype.tooltipFormatter,D=t.addEvent,E=t.arrayMax,x=t.arrayMin,b=t.correctFloat,
m=t.defined,d=t.isArray,z=t.isNumber,h=t.isString,K=t.pick,G;(function(I){function C(e,c,n){this.isXAxis||(this.series.forEach(function(p){"compare"===e&&"boolean"!==typeof c?p.setCompare(c,!1):"cumulative"!==e||h(c)||p.setCumulative(c,!1)}),K(n,!0)&&this.chart.redraw())}function q(e){var c=this,n=c.series.chart.numberFormatter,p=function(g){e=e.replace("{point."+g+"}",(0<c[g]&&"change"===g?"+":"")+n(c[g],K(c.series.tooltipOptions.changeDecimals,2)))};m(c.change)&&p("change");m(c.cumulativeSum)&&
p("cumulativeSum");return l.apply(this,[e])}function a(){var e=this.options.compare;if("percent"===e||"value"===e||this.options.cumulative){var c=new f(this);"percent"===e||"value"===e?c.initCompare(e):c.initCumulative()}this.dataModify=c}function G(e){e=e.dataExtremes;var c=e.activeYData;if(this.dataModify&&e){var n=void 0;this.options.compare?n=[this.dataModify.modifyValue(e.dataMin),this.dataModify.modifyValue(e.dataMax)]:this.options.cumulative&&d(c)&&2<=c.length&&(n=f.getCumulativeExtremes(c));
n&&(e.dataMin=x(n),e.dataMax=E(n))}}function F(e,c){this.options.compare=this.userOptions.compare=e;this.update({},K(c,!0));!this.dataModify||"value"!==e&&"percent"!==e?this.points.forEach(function(c){delete c.change}):this.dataModify.initCompare(e)}function N(){if(this.xAxis&&this.processedYData&&this.dataModify){var e=this.processedXData,c=this.processedYData,n=c.length,p=!0===this.options.compareStart?0:1,g=-1,v;this.pointArrayMap&&(g=this.pointArrayMap.indexOf(this.options.pointValKey||this.pointValKey||
"y"));for(v=0;v<n-p;v++){var w=c[v]&&-1<g?c[v][g]:c[v];if(z(w)&&0!==w&&e[v+p]>=(this.xAxis.min||0)){this.dataModify.compareValue=w;break}}}}function u(e,c){this.setModifier("compare",e,c)}function M(e,c){e=K(e,!1);this.options.cumulative=this.userOptions.cumulative=e;this.update({},K(c,!0));this.dataModify?this.dataModify.initCumulative():this.points.forEach(function(c){delete c.cumulativeSum})}function k(e,c){this.setModifier("cumulative",e,c)}var J=[];I.compose=function(e,c,n){if(-1===J.indexOf(e)){J.push(e);
var p=e.prototype;p.setCompare=F;p.setCumulative=M;D(e,"afterInit",a);D(e,"afterGetExtremes",G);D(e,"afterProcessData",N)}-1===J.indexOf(c)&&(J.push(c),c=c.prototype,c.setCompare=u,c.setModifier=C,c.setCumulative=k);-1===J.indexOf(n)&&(J.push(n),n.prototype.tooltipFormatter=q);return e};var f=function(){function e(c){this.series=c}e.prototype.modifyValue=function(){return 0};e.getCumulativeExtremes=function(c){var e=Infinity,p=-Infinity;c.reduce(function(g,c){c=g+c;e=Math.min(e,c,g);p=Math.max(p,
c,g);return c});return[e,p]};e.prototype.initCompare=function(c){this.modifyValue=function(e,p){null===e&&(e=0);var g=this.compareValue;return"undefined"!==typeof e&&"undefined"!==typeof g?(e="value"===c?e-g:e/g*100-(100===this.series.options.compareBase?0:100),"undefined"!==typeof p&&(p=this.series.points[p])&&(p.change=e),e):0}};e.prototype.initCumulative=function(){this.modifyValue=function(c,e){null===c&&(c=0);if(void 0!==c&&void 0!==e){var p=0<e?this.series.points[e-1]:null;p&&p.cumulativeSum&&
(c=b(p.cumulativeSum+c));if(e=this.series.points[e])e.cumulativeSum=c;return c}return 0}};return e}();I.Additions=f})(G||(G={}));"";return G});L(a,"Core/Axis/BrokenAxis.js",[a["Extensions/Stacking.js"],a["Core/Utilities.js"]],function(a,r){var y=r.addEvent,t=r.find,l=r.fireEvent,D=r.isArray,E=r.isNumber,x=r.pick,b;(function(m){function d(){"undefined"!==typeof this.brokenAxis&&this.brokenAxis.setBreaks(this.options.breaks,!1)}function b(){this.brokenAxis&&this.brokenAxis.hasBreaks&&(this.options.ordinal=
!1)}function h(){var d=this.brokenAxis;if(d&&d.hasBreaks){for(var q=this.tickPositions,u=this.tickPositions.info,h=[],k=0;k<q.length;k++)d.isInAnyBreak(q[k])||h.push(q[k]);this.tickPositions=h;this.tickPositions.info=u}}function K(){this.brokenAxis||(this.brokenAxis=new S(this))}function G(){var d=this.options.connectNulls,q=this.points,u=this.xAxis,h=this.yAxis;if(this.isDirty)for(var k=q.length;k--;){var b=q[k],f=!(null===b.y&&!1===d)&&(u&&u.brokenAxis&&u.brokenAxis.isInAnyBreak(b.x,!0)||h&&h.brokenAxis&&
h.brokenAxis.isInAnyBreak(b.y,!0));b.visible=f?!1:!1!==b.options.visible}}function I(){this.drawBreaks(this.xAxis,["x"]);this.drawBreaks(this.yAxis,x(this.pointArrayMap,["y"]))}function C(d,q){var u=this,h=u.points,k,b,f,e;if(d&&d.brokenAxis&&d.brokenAxis.hasBreaks){var c=d.brokenAxis;q.forEach(function(n){k=c&&c.breakArray||[];b=d.isXAxis?d.min:x(u.options.threshold,d.min);h.forEach(function(c){e=x(c["stack"+n.toUpperCase()],c[n]);k.forEach(function(g){if(E(b)&&E(e)){f=!1;if(b<g.from&&e>g.to||b>
g.from&&e<g.from)f="pointBreak";else if(b<g.from&&e>g.from&&e<g.to||b>g.from&&e>g.to&&e<g.from)f="pointInBreak";f&&l(d,f,{point:c,brk:g})}})})})}}function q(){var d=this.currentDataGrouping,q=d&&d.gapSize;d=this.points.slice();var u=this.yAxis,h=this.options.gapSize,k=d.length-1,b;if(h&&0<k)for("value"!==this.options.gapUnit&&(h*=this.basePointRange),q&&q>h&&q>=this.basePointRange&&(h=q),b=void 0;k--;)b&&!1!==b.visible||(b=d[k+1]),q=d[k],!1!==b.visible&&!1!==q.visible&&(b.x-q.x>h&&(b=(q.x+b.x)/2,
d.splice(k+1,0,{isNull:!0,x:b}),u.stacking&&this.options.stacking&&(b=u.stacking.stacks[this.stackKey][b]=new a(u,u.options.stackLabels,!1,b,this.stack),b.total=0)),b=q);return this.getGraphPath(d)}var R=[];m.compose=function(m,a){-1===R.indexOf(m)&&(R.push(m),m.keepProps.push("brokenAxis"),y(m,"init",K),y(m,"afterInit",d),y(m,"afterSetTickPositions",h),y(m,"afterSetOptions",b));if(-1===R.indexOf(a)){R.push(a);var u=a.prototype;u.drawBreaks=C;u.gappedPath=q;y(a,"afterGeneratePoints",G);y(a,"afterRender",
I)}return m};var S=function(){function d(d){this.hasBreaks=!1;this.axis=d}d.isInBreak=function(d,b){var q=d.repeat||Infinity,k=d.from,h=d.to-d.from;b=b>=k?(b-k)%q:q-(k-b)%q;return d.inclusive?b<=h:b<h&&0!==b};d.lin2Val=function(b){var q=this.brokenAxis;q=q&&q.breakArray;if(!q||!E(b))return b;var h;for(h=0;h<q.length;h++){var k=q[h];if(k.from>=b)break;else k.to<b?b+=k.len:d.isInBreak(k,b)&&(b+=k.len)}return b};d.val2Lin=function(b){var q=this.brokenAxis;q=q&&q.breakArray;if(!q||!E(b))return b;var h=
b,k;for(k=0;k<q.length;k++){var m=q[k];if(m.to<=b)h-=m.len;else if(m.from>=b)break;else if(d.isInBreak(m,b)){h-=b-m.from;break}}return h};d.prototype.findBreakAt=function(d,b){return t(b,function(b){return b.from<d&&d<b.to})};d.prototype.isInAnyBreak=function(b,q){var h=this.axis,k=h.options.breaks||[],m=k.length,f;if(m&&E(b)){for(;m--;)if(d.isInBreak(k[m],b)){var e=!0;f||(f=x(k[m].showPoints,!h.isXAxis))}var c=e&&q?e&&!f:e}return c};d.prototype.setBreaks=function(b,q){var h=this,k=h.axis,m=D(b)&&
!!b.length;k.isDirty=h.hasBreaks!==m;h.hasBreaks=m;k.options.breaks=k.userOptions.breaks=b;k.forceRedraw=!0;k.series.forEach(function(f){f.isDirty=!0});m||k.val2lin!==d.val2Lin||(delete k.val2lin,delete k.lin2val);m&&(k.userOptions.ordinal=!1,k.lin2val=d.lin2Val,k.val2lin=d.val2Lin,k.setExtremes=function(f,e,c,n,p){if(h.hasBreaks){for(var g=this.options.breaks||[],v;v=h.findBreakAt(f,g);)f=v.to;for(;v=h.findBreakAt(e,g);)e=v.from;e<f&&(e=f)}k.constructor.prototype.setExtremes.call(this,f,e,c,n,p)},
k.setAxisTranslation=function(){k.constructor.prototype.setAxisTranslation.call(this);h.unitLength=void 0;if(h.hasBreaks){var f=k.options.breaks||[],e=[],c=[],n=x(k.pointRangePadding,0),p=0,g,v=k.userMin||k.min,w=k.userMax||k.max,b;f.forEach(function(c){g=c.repeat||Infinity;E(v)&&E(w)&&(d.isInBreak(c,v)&&(v+=c.to%g-v%g),d.isInBreak(c,w)&&(w-=w%g-c.from%g))});f.forEach(function(c){B=c.from;g=c.repeat||Infinity;if(E(v)&&E(w)){for(;B-g>v;)B-=g;for(;B<v;)B+=g;for(b=B;b<w;b+=g)e.push({value:b,move:"in"}),
e.push({value:b+c.to-c.from,move:"out",size:c.breakSize})}});e.sort(function(g,c){return g.value===c.value?("in"===g.move?0:1)-("in"===c.move?0:1):g.value-c.value});var q=0;var B=v;e.forEach(function(g){q+="in"===g.move?1:-1;1===q&&"in"===g.move&&(B=g.value);0===q&&E(B)&&(c.push({from:B,to:g.value,len:g.value-B-(g.size||0)}),p+=g.value-B-(g.size||0))});h.breakArray=c;E(v)&&E(w)&&E(k.min)&&(h.unitLength=w-v-p+n,l(k,"afterBreaks"),k.staticScale?k.transA=k.staticScale:h.unitLength&&(k.transA*=(w-k.min+
n)/h.unitLength),n&&(k.minPixelPadding=k.transA*(k.minPointOffset||0)),k.min=v,k.max=w)}});x(q,!0)&&k.chart.redraw()};return d}();m.Additions=S})(b||(b={}));return b});L(a,"masters/modules/broken-axis.src.js",[a["Core/Globals.js"],a["Core/Axis/BrokenAxis.js"]],function(a,r){r.compose(a.Axis,a.Series)});L(a,"Extensions/DataGrouping.js",[a["Core/Axis/Axis.js"],a["Core/Axis/DateTimeAxis.js"],a["Core/FormatUtilities.js"],a["Core/Globals.js"],a["Core/Series/Point.js"],a["Core/Series/Series.js"],a["Core/Tooltip.js"],
a["Core/DefaultOptions.js"],a["Core/Utilities.js"]],function(a,r,y,t,l,D,E,x,b){var m=y.format,d=D.prototype;y=b.addEvent;var z=b.arrayMax,h=b.arrayMin,K=b.correctFloat,G=b.defined,I=b.error,C=b.extend,q=b.isNumber,R=b.merge,S=b.pick;"";var F=t.approximations={sum:function(f){var e=f.length;if(!e&&f.hasNulls)var c=null;else if(e)for(c=0;e--;)c+=f[e];return c},average:function(f){var e=f.length;f=F.sum(f);q(f)&&e&&(f=K(f/e));return f},averages:function(){var f=[];[].forEach.call(arguments,function(e){f.push(F.average(e))});
return"undefined"===typeof f[0]?void 0:f},open:function(f){return f.length?f[0]:f.hasNulls?null:void 0},high:function(f){return f.length?z(f):f.hasNulls?null:void 0},low:function(f){return f.length?h(f):f.hasNulls?null:void 0},close:function(f){return f.length?f[f.length-1]:f.hasNulls?null:void 0},hlc:function(f,e,c){f=F.high(f);e=F.low(e);c=F.close(c);if(q(f)||q(e)||q(c))return[f,e,c]},ohlc:function(f,e,c,n){f=F.open(f);e=F.high(e);c=F.low(c);n=F.close(n);if(q(f)||q(e)||q(c)||q(n))return[f,e,c,n]},
range:function(f,e){f=F.low(f);e=F.high(e);if(q(f)||q(e))return[f,e];if(null===f&&null===e)return null}};b=function(f,e,c,n){var p=this,g=p.data,v=p.options&&p.options.data,w=[],d=[],b=[],B=f.length,H=!!e,A=[],h=p.pointArrayMap,k=h&&h.length,m=["x"].concat(h||["y"]),u=this.options.dataGrouping&&this.options.dataGrouping.groupAll,a=0,I=0,C;n="function"===typeof n?n:F[n]?F[n]:F[p.getDGApproximation&&p.getDGApproximation()||"average"];k?h.forEach(function(){A.push([])}):A.push([]);var K=k||1;for(C=0;C<=
B&&!(f[C]>=c[0]);C++);for(C;C<=B;C++){for(;"undefined"!==typeof c[a+1]&&f[C]>=c[a+1]||C===B;){var z=c[a];p.dataGroupInfo={start:u?I:p.cropStart+I,length:A[0].length};var l=n.apply(p,A);p.pointClass&&!G(p.dataGroupInfo.options)&&(p.dataGroupInfo.options=R(p.pointClass.prototype.optionsToObject.call({series:p},p.options.data[p.cropStart+I])),m.forEach(function(g){delete p.dataGroupInfo.options[g]}));"undefined"!==typeof l&&(w.push(z),d.push(l),b.push(p.dataGroupInfo));I=C;for(z=0;z<K;z++)A[z].length=
0,A[z].hasNulls=!1;a+=1;if(C===B)break}if(C===B)break;if(h){z=p.options.dataGrouping&&p.options.dataGrouping.groupAll?C:p.cropStart+C;l=g&&g[z]||p.pointClass.prototype.applyOptions.apply({series:p},[v[z]]);var x=void 0;for(z=0;z<k;z++)x=l[h[z]],q(x)?A[z].push(x):null===x&&(A[z].hasNulls=!0)}else z=H?e[C]:null,q(z)?A[0].push(z):null===z&&(A[0].hasNulls=!0)}return{groupedXData:w,groupedYData:d,groupMap:b}};var P={approximations:F,groupData:b},u=d.generatePoints,M={groupPixelWidth:2,dateTimeLabelFormats:{millisecond:["%A, %b %e, %H:%M:%S.%L",
"%A, %b %e, %H:%M:%S.%L","-%H:%M:%S.%L"],second:["%A, %b %e, %H:%M:%S","%A, %b %e, %H:%M:%S","-%H:%M:%S"],minute:["%A, %b %e, %H:%M","%A, %b %e, %H:%M","-%H:%M"],hour:["%A, %b %e, %H:%M","%A, %b %e, %H:%M","-%H:%M"],day:["%A, %b %e, %Y","%A, %b %e","-%A, %b %e, %Y"],week:["Week from %A, %b %e, %Y","%A, %b %e","-%A, %b %e, %Y"],month:["%B %Y","%B","-%B %Y"],year:["%Y","%Y","-%Y"]}},k={line:{},spline:{},area:{},areaspline:{},arearange:{},column:{groupPixelWidth:10},columnrange:{groupPixelWidth:10},
candlestick:{groupPixelWidth:10},ohlc:{groupPixelWidth:5},hlc:{groupPixelWidth:5},heikinashi:{groupPixelWidth:10}},J=t.defaultDataGroupingUnits=[["millisecond",[1,2,5,10,20,25,50,100,200,500]],["second",[1,2,5,10,15,30]],["minute",[1,2,5,10,15,30]],["hour",[1,2,3,4,6,8,12]],["day",[1]],["week",[1]],["month",[1,3,6]],["year",null]];d.getDGApproximation=function(){return this.is("arearange")?"range":this.is("ohlc")?"ohlc":this.is("hlc")?"hlc":this.is("column")?"sum":"average"};d.groupData=b;d.applyGrouping=
function(f){var e=this.chart,c=this.options.dataGrouping,n=!1!==this.allowDG&&c&&S(c.enabled,e.options.isStock),p=this.visible||!e.options.chart.ignoreHiddenSeries,g,v=this.currentDataGrouping,w=!1;n&&!this.requireSorting&&(this.requireSorting=w=!0);f=!1===!(this.isCartesian&&!this.isDirty&&!this.xAxis.isDirty&&!this.yAxis.isDirty&&!f)||!n;w&&(this.requireSorting=!1);if(!f){this.destroyGroupedData();n=c.groupAll?this.xData:this.processedXData;var b=c.groupAll?this.yData:this.processedYData;f=e.plotSizeX;
w=this.xAxis;var h=w.options.ordinal,B=this.groupPixelWidth;if(B&&n&&n.length){this.isDirty=g=!0;this.points=null;var k=w.getExtremes();var A=k.min;k=k.max;h=h&&w.ordinal&&w.ordinal.getGroupIntervalFactor(A,k,this)||1;f=w.getTimeTicks(r.Additions.prototype.normalizeTimeTickInterval(B*(k-A)/f*h,c.units||J),Math.min(A,n[0]),Math.max(k,n[n.length-1]),w.options.startOfWeek,n,this.closestPointRange);B=d.groupData.apply(this,[n,b,f,c.approximation]);n=B.groupedXData;b=B.groupedYData;h=0;c&&c.smoothed&&
n.length&&(c.firstAnchor="firstPoint",c.anchor="middle",c.lastAnchor="lastPoint",I(32,!1,e,{"dataGrouping.smoothed":"use dataGrouping.anchor"}));e=n;var m=this.options.dataGrouping;A=this.currentDataGrouping&&this.currentDataGrouping.gapSize;if(m&&this.xData&&A&&this.groupMap){var u=e.length-1;var a=m.anchor;var C=S(m.firstAnchor,a);m=S(m.lastAnchor,a);if(a&&"start"!==a){var z=A*{middle:.5,end:1}[a];for(a=e.length-1;a--&&0<a;)e[a]+=z}if(C&&"start"!==C&&this.xData[0]>=e[0]){a=this.groupMap[0].start;
z=this.groupMap[0].length;var K=void 0;q(a)&&q(z)&&(K=a+(z-1));e[0]={middle:e[0]+.5*A,end:e[0]+A,firstPoint:this.xData[0],lastPoint:K&&this.xData[K]}[C]}m&&"start"!==m&&A&&e[u]>=k-A&&(k=this.groupMap[this.groupMap.length-1].start,e[u]={middle:e[u]+.5*A,end:e[u]+A,firstPoint:k&&this.xData[k],lastPoint:this.xData[this.xData.length-1]}[m])}for(k=1;k<f.length;k++)f.info.segmentStarts&&-1!==f.info.segmentStarts.indexOf(k)||(h=Math.max(f[k]-f[k-1],h));k=f.info;k.gapSize=h;this.closestPointRange=f.info.totalRange;
this.groupMap=B.groupMap;if(p){p=n;if(G(p[0])&&q(w.min)&&q(w.dataMin)&&p[0]<w.min){if(!G(w.options.min)&&w.min<=w.dataMin||w.min===w.dataMin)w.min=Math.min(p[0],w.min);w.dataMin=Math.min(p[0],w.dataMin)}if(G(p[p.length-1])&&q(w.max)&&q(w.dataMax)&&p[p.length-1]>w.max){if(!G(w.options.max)&&q(w.dataMax)&&w.max>=w.dataMax||w.max===w.dataMax)w.max=Math.max(p[p.length-1],w.max);w.dataMax=Math.max(p[p.length-1],w.dataMax)}}c.groupAll&&(this.allGroupedData=b,c=this.cropData(n,b,w.min,w.max,1),n=c.xData,
b=c.yData,this.cropStart=c.start);this.processedXData=n;this.processedYData=b}else this.groupMap=null;this.hasGroupedData=g;this.currentDataGrouping=k;this.preventGraphAnimation=(v&&v.totalRange)!==(k&&k.totalRange)}};d.destroyGroupedData=function(){this.groupedData&&(this.groupedData.forEach(function(f,e){f&&(this.groupedData[e]=f.destroy?f.destroy():null)},this),this.groupedData.length=0)};d.generatePoints=function(){u.apply(this);this.destroyGroupedData();this.groupedData=this.hasGroupedData?this.points:
null};a.prototype.applyGrouping=function(f){var e=this;e.series.forEach(function(c){c.groupPixelWidth=void 0;c.groupPixelWidth=e.getGroupPixelWidth&&e.getGroupPixelWidth();c.groupPixelWidth&&(c.hasProcessed=!0);c.applyGrouping(!!f.hasExtemesChanged)})};a.prototype.getGroupPixelWidth=function(){var f=this.series,e=f.length,c,n=0,p=!1,g;for(c=e;c--;)(g=f[c].options.dataGrouping)&&(n=Math.max(n,S(g.groupPixelWidth,M.groupPixelWidth)));for(c=e;c--;)if(g=f[c].options.dataGrouping)if(e=(f[c].processedXData||
f[c].data).length,f[c].groupPixelWidth||e>this.chart.plotSizeX/n||e&&g.forced)p=!0;return p?n:0};a.prototype.setDataGrouping=function(f,e){var c;e=S(e,!0);f||(f={forced:!1,units:null});if(this instanceof a)for(c=this.series.length;c--;)this.series[c].update({dataGrouping:f},!1);else this.chart.options.series.forEach(function(c){c.dataGrouping=f},!1);this.ordinal&&(this.ordinal.slope=void 0);e&&this.chart.redraw()};y(a,"postProcessData",a.prototype.applyGrouping);y(l,"update",function(){if(this.dataGroup)return I(24,
!1,this.series.chart),!1});y(E,"headerFormatter",function(f){var e=this.chart,c=e.time,n=f.labelConfig,p=n.series,g=p.tooltipOptions,v=p.options.dataGrouping,w=g.xDateFormat,d=p.xAxis,b=g[f.isFooter?"footerFormat":"headerFormat"];if(d&&"datetime"===d.options.type&&v&&q(n.key)){var k=p.currentDataGrouping;v=v.dateTimeLabelFormats||M.dateTimeLabelFormats;if(k)if(g=v[k.unitName],1===k.count)w=g[0];else{w=g[1];var h=g[2]}else!w&&v&&d.dateTime&&(w=d.dateTime.getXDateFormat(n.x,g.dateTimeLabelFormats));
w=c.dateFormat(w,n.key);h&&(w+=c.dateFormat(h,n.key+k.totalRange-1));p.chart.styledMode&&(b=this.styledModeFormat(b));f.text=m(b,{point:C(n.point,{key:w}),series:p},e);f.preventDefault()}});y(D,"destroy",d.destroyGroupedData);y(D,"afterSetOptions",function(f){f=f.options;var e=this.type,c=this.chart.options.plotOptions,n=x.defaultOptions.plotOptions[e].dataGrouping,p=this.useCommonDataGrouping&&M;if(k[e]||p){n||(n=R(M,k[e]));var g=this.chart.rangeSelector;f.dataGrouping=R(p,n,c.series&&c.series.dataGrouping,
c[e].dataGrouping,this.userOptions.dataGrouping,!f.isInternal&&g&&q(g.selected)&&g.buttonOptions[g.selected].dataGrouping)}});y(a,"afterSetScale",function(){this.series.forEach(function(f){f.hasProcessed=!1})});t.dataGrouping=P;"";return P});L(a,"Series/HLC/HLCPoint.js",[a["Core/Series/SeriesRegistry.js"]],function(a){var r=this&&this.__extends||function(){var a=function(t,l){a=Object.setPrototypeOf||{__proto__:[]}instanceof Array&&function(a,l){a.__proto__=l}||function(a,l){for(var x in l)l.hasOwnProperty(x)&&
(a[x]=l[x])};return a(t,l)};return function(t,l){function D(){this.constructor=t}a(t,l);t.prototype=null===l?Object.create(l):(D.prototype=l.prototype,new D)}}();return function(a){function t(){var l=null!==a&&a.apply(this,arguments)||this;l.close=void 0;l.high=void 0;l.low=void 0;l.options=void 0;l.plotClose=void 0;l.series=void 0;return l}r(t,a);return t}(a.seriesTypes.column.prototype.pointClass)});L(a,"Series/HLC/HLCSeries.js",[a["Series/HLC/HLCPoint.js"],a["Core/Series/SeriesRegistry.js"],a["Core/Utilities.js"]],
function(a,r,y){var t=this&&this.__extends||function(){var a=function(b,m){a=Object.setPrototypeOf||{__proto__:[]}instanceof Array&&function(d,b){d.__proto__=b}||function(d,b){for(var h in b)b.hasOwnProperty(h)&&(d[h]=b[h])};return a(b,m)};return function(b,m){function d(){this.constructor=b}a(b,m);b.prototype=null===m?Object.create(m):(d.prototype=m.prototype,new d)}}(),l=r.seriesTypes.column,D=y.extend,E=y.merge;y=function(a){function b(){var b=null!==a&&a.apply(this,arguments)||this;b.data=void 0;
b.options=void 0;b.points=void 0;b.yData=void 0;return b}t(b,a);b.prototype.extendStem=function(b,d,a){var h=b[0];b=b[1];"number"===typeof h[2]&&(h[2]=Math.max(a+d,h[2]));"number"===typeof b[2]&&(b[2]=Math.min(a-d,b[2]))};b.prototype.getPointPath=function(b,d){d=d.strokeWidth();var a=b.series,h=d%2/2,m=Math.round(b.plotX)-h,G=Math.round(b.shapeArgs.width/2);var I=[["M",m,Math.round(b.yBottom)],["L",m,Math.round(b.plotHigh)]];null!==b.close&&(b=Math.round(b.plotClose)+h,I.push(["M",m,b],["L",m+G,b]),
a.extendStem(I,d/2,b));return I};b.prototype.drawSinglePoint=function(b){var d=b.series,a=d.chart,h=b.graphic,m=!h;"undefined"!==typeof b.plotY&&(h||(b.graphic=h=a.renderer.path().add(d.group)),a.styledMode||h.attr(d.pointAttribs(b,b.selected&&"select")),d=d.getPointPath(b,h),h[m?"attr":"animate"]({d:d}).addClass(b.getClassName(),!0))};b.prototype.drawPoints=function(){this.points.forEach(this.drawSinglePoint)};b.prototype.init=function(){a.prototype.init.apply(this,arguments);this.options.stacking=
void 0};b.prototype.pointAttribs=function(b,d){b=a.prototype.pointAttribs.call(this,b,d);delete b.fill;return b};b.prototype.toYData=function(b){return[b.high,b.low,b.close]};b.prototype.translate=function(){var b=this,d=b.yAxis,z=this.pointArrayMap&&this.pointArrayMap.slice()||[],h=z.map(function(b){return"plot"+(b.charAt(0).toUpperCase()+b.slice(1))});h.push("yBottom");z.push("low");a.prototype.translate.apply(b);b.points.forEach(function(a){z.forEach(function(m,I){m=a[m];null!==m&&(b.dataModify&&
(m=b.dataModify.modifyValue(m)),a[h[I]]=d.toPixels(m,!0))});a.tooltipPos[1]=a.plotHigh+d.pos-b.chart.plotTop})};b.defaultOptions=E(l.defaultOptions,{lineWidth:1,tooltip:{pointFormat:'<span style="color:{point.color}">\u25cf</span> <b> {series.name}</b><br/>High: {point.high}<br/>Low: {point.low}<br/>Close: {point.close}<br/>'},threshold:null,states:{hover:{lineWidth:3}},stickyTracking:!0});return b}(l);D(y.prototype,{animate:null,directTouch:!1,pointArrayMap:["high","low","close"],pointAttrToOptions:{stroke:"color",
"stroke-width":"lineWidth"},pointValKey:"close"});y.prototype.pointClass=a;r.registerSeriesType("hlc",y);"";return y});L(a,"Series/OHLC/OHLCPoint.js",[a["Core/Series/SeriesRegistry.js"]],function(a){var r=this&&this.__extends||function(){var a=function(t,l){a=Object.setPrototypeOf||{__proto__:[]}instanceof Array&&function(a,l){a.__proto__=l}||function(a,l){for(var t in l)l.hasOwnProperty(t)&&(a[t]=l[t])};return a(t,l)};return function(t,l){function D(){this.constructor=t}a(t,l);t.prototype=null===
l?Object.create(l):(D.prototype=l.prototype,new D)}}();return function(a){function t(){var l=null!==a&&a.apply(this,arguments)||this;l.open=void 0;l.options=void 0;l.plotOpen=void 0;l.series=void 0;return l}r(t,a);t.prototype.getClassName=function(){return a.prototype.getClassName.call(this)+(this.open<this.close?" highcharts-point-up":" highcharts-point-down")};t.prototype.resolveUpColor=function(){this.open<this.close&&!this.options.color&&this.series.options.upColor&&(this.color=this.series.options.upColor)};
t.prototype.resolveColor=function(){a.prototype.resolveColor.call(this);this.resolveUpColor()};t.prototype.getZone=function(){var l=a.prototype.getZone.call(this);this.resolveUpColor();return l};t.prototype.applyOptions=function(){a.prototype.applyOptions.apply(this,arguments);this.resolveColor&&this.resolveColor();return this};return t}(a.seriesTypes.hlc.prototype.pointClass)});L(a,"Series/OHLC/OHLCSeries.js",[a["Series/OHLC/OHLCPoint.js"],a["Core/Series/SeriesRegistry.js"],a["Core/Utilities.js"]],
function(a,r,y){var t=this&&this.__extends||function(){var b=function(d,a){b=Object.setPrototypeOf||{__proto__:[]}instanceof Array&&function(b,d){b.__proto__=d}||function(b,d){for(var a in d)d.hasOwnProperty(a)&&(b[a]=d[a])};return b(d,a)};return function(d,a){function h(){this.constructor=d}b(d,a);d.prototype=null===a?Object.create(a):(h.prototype=a.prototype,new h)}}(),l=r.series,D=r.seriesTypes.hlc,E=y.addEvent,x=y.extend,b=y.merge,m=function(d){function a(){var b=null!==d&&d.apply(this,arguments)||
this;b.data=void 0;b.options=void 0;b.points=void 0;return b}t(a,d);a.prototype.getPointPath=function(b,a){var h=d.prototype.getPointPath.call(this,b,a);a=a.strokeWidth();var m=a%2/2,C=Math.round(b.plotX)-m,q=Math.round(b.shapeArgs.width/2);null!==b.open&&(b=Math.round(b.plotOpen)+m,h.push(["M",C,b],["L",C-q,b]),d.prototype.extendStem.call(this,h,a/2,b));return h};a.prototype.pointAttribs=function(b,a){a=d.prototype.pointAttribs.call(this,b,a);var h=this.options;delete a.fill;!b.options.color&&h.upColor&&
b.open<b.close&&(a.stroke=h.upColor);return a};a.prototype.toYData=function(b){return[b.open,b.high,b.low,b.close]};a.defaultOptions=b(D.defaultOptions,{tooltip:{pointFormat:'<span style="color:{point.color}">\u25cf</span> <b> {series.name}</b><br/>Open: {point.open}<br/>High: {point.high}<br/>Low: {point.low}<br/>Close: {point.close}<br/>'}});return a}(D);x(m.prototype,{pointArrayMap:["open","high","low","close"]});m.prototype.pointClass=a;r.registerSeriesType("ohlc",m);E(l,"init",function(b){b=
b.options;b.useOhlcData&&"highcharts-navigator-series"!==b.id&&x(this,{pointValKey:m.prototype.pointValKey,pointArrayMap:m.prototype.pointArrayMap,toYData:m.prototype.toYData})});E(l,"afterSetOptions",function(b){b=b.options;var a=b.dataGrouping;a&&b.useOhlcData&&"highcharts-navigator-series"!==b.id&&(a.approximation="ohlc")});"";return m});L(a,"Series/Candlestick/CandlestickSeries.js",[a["Core/DefaultOptions.js"],a["Core/Series/SeriesRegistry.js"],a["Core/Utilities.js"]],function(a,r,y){var t=this&&
this.__extends||function(){var b=function(a,d){b=Object.setPrototypeOf||{__proto__:[]}instanceof Array&&function(b,a){b.__proto__=a}||function(b,a){for(var d in a)a.hasOwnProperty(d)&&(b[d]=a[d])};return b(a,d)};return function(a,d){function m(){this.constructor=a}b(a,d);a.prototype=null===d?Object.create(d):(m.prototype=d.prototype,new m)}}(),l=a.defaultOptions;a=r.seriesTypes;var D=a.column,E=a.ohlc,x=y.merge;y=function(b){function a(){var a=null!==b&&b.apply(this,arguments)||this;a.data=void 0;
a.options=void 0;a.points=void 0;return a}t(a,b);a.prototype.pointAttribs=function(b,a){var d=D.prototype.pointAttribs.call(this,b,a),m=this.options,l=b.open<b.close,I=m.lineColor||this.color,C=b.color||this.color;d["stroke-width"]=m.lineWidth;d.fill=b.options.color||(l?m.upColor||C:C);d.stroke=b.options.lineColor||(l?m.upLineColor||I:I);a&&(b=m.states[a],d.fill=b.color||d.fill,d.stroke=b.lineColor||d.stroke,d["stroke-width"]=b.lineWidth||d["stroke-width"]);return d};a.prototype.drawPoints=function(){var b=
this,a=b.chart,h=b.yAxis.reversed;b.points.forEach(function(d){var m=d.graphic,I=!m;if("undefined"!==typeof d.plotY){m||(d.graphic=m=a.renderer.path().add(b.group));b.chart.styledMode||m.attr(b.pointAttribs(d,d.selected&&"select")).shadow(b.options.shadow);var C=m.strokeWidth()%2/2;var q=Math.round(d.plotX)-C;var l=d.plotOpen;var t=d.plotClose;var F=Math.min(l,t);l=Math.max(l,t);var z=Math.round(d.shapeArgs.width/2);t=h?l!==d.yBottom:Math.round(F)!==Math.round(d.plotHigh);var u=h?Math.round(F)!==
Math.round(d.plotHigh):l!==d.yBottom;F=Math.round(F)+C;l=Math.round(l)+C;C=[];C.push(["M",q-z,l],["L",q-z,F],["L",q+z,F],["L",q+z,l],["Z"],["M",q,F],["L",q,t?Math.round(h?d.yBottom:d.plotHigh):F],["M",q,l],["L",q,u?Math.round(h?d.plotHigh:d.yBottom):l]);m[I?"attr":"animate"]({d:C}).addClass(d.getClassName(),!0)}})};a.defaultOptions=x(E.defaultOptions,l.plotOptions,{states:{hover:{lineWidth:2}},tooltip:l.plotOptions.ohlc.tooltip,threshold:null,lineColor:"#000000",lineWidth:1,upColor:"#ffffff",stickyTracking:!0});
return a}(E);r.registerSeriesType("candlestick",y);"";return y});L(a,"Series/Flags/FlagsPoint.js",[a["Core/Series/SeriesRegistry.js"],a["Core/Utilities.js"]],function(a,r){var y=this&&this.__extends||function(){var a=function(l,t){a=Object.setPrototypeOf||{__proto__:[]}instanceof Array&&function(a,b){a.__proto__=b}||function(a,b){for(var m in b)b.hasOwnProperty(m)&&(a[m]=b[m])};return a(l,t)};return function(l,t){function x(){this.constructor=l}a(l,t);l.prototype=null===t?Object.create(t):(x.prototype=
t.prototype,new x)}}(),t=r.isNumber;return function(a){function l(){var l=null!==a&&a.apply(this,arguments)||this;l.options=void 0;l.series=void 0;return l}y(l,a);l.prototype.isValid=function(){return t(this.y)||"undefined"===typeof this.y};l.prototype.hasNewShapeType=function(){var a=this.options.shape||this.series.options.shape;return this.graphic&&a&&a!==this.graphic.symbolKey};return l}(a.seriesTypes.column.prototype.pointClass)});L(a,"Series/OnSeriesComposition.js",[a["Series/Column/ColumnSeries.js"],
a["Core/Series/Series.js"],a["Core/Utilities.js"]],function(a,r,y){var t=a.prototype,l=r.prototype,D=y.defined,E=y.stableSort,x;(function(b){function a(){return l.getPlotBox.call(this.options.onSeries&&this.chart.get(this.options.onSeries)||this)}function d(){t.translate.apply(this);var b=this,a=b.options,d=b.chart,m=b.points,C=a.onSeries,q=(C=C&&d.get(C))&&C.options.step,l=C&&C.points,z=d.inverted,F=b.xAxis,x=b.yAxis;d=m.length-1;var u;a=a.onKey||"y";var M=l&&l.length,k=0,r;if(C&&C.visible&&M){k=
(C.pointXOffset||0)+(C.barW||0)/2;var f=C.currentDataGrouping;var e=l[M-1].x+(f?f.totalRange:0);E(m,function(c,g){return c.x-g.x});for(a="plot"+a[0].toUpperCase()+a.substr(1);M--&&m[d];){var c=l[M];f=m[d];f.y=c.y;if(c.x<=f.x&&"undefined"!==typeof c[a]){if(f.x<=e&&(f.plotY=c[a],c.x<f.x&&!q&&(r=l[M+1])&&"undefined"!==typeof r[a])){var n=(f.x-c.x)/(r.x-c.x);f.plotY+=n*(r[a]-c[a]);f.y+=n*(r.y-c.y)}d--;M++;if(0>d)break}}}m.forEach(function(c,g){c.plotX+=k;if("undefined"===typeof c.plotY||z)0<=c.plotX&&
c.plotX<=F.len?z?(c.plotY=F.translate(c.x,0,1,0,1),c.plotX=D(c.y)?x.translate(c.y,0,0,0,1):0):c.plotY=(F.opposite?0:b.yAxis.len)+F.offset:c.shapeArgs={};if((u=m[g-1])&&u.plotX===c.plotX){"undefined"===typeof u.stackIndex&&(u.stackIndex=0);var v=u.stackIndex+1}c.stackIndex=v});this.onSeries=C}var z=[];b.compose=function(b){if(-1===z.indexOf(b)){z.push(b);var h=b.prototype;h.getPlotBox=a;h.translate=d}return b};b.getPlotBox=a;b.translate=d})(x||(x={}));return x});L(a,"Series/Flags/FlagsSymbols.js",
[a["Core/Renderer/RendererRegistry.js"],a["Core/Renderer/SVG/SVGRenderer.js"]],function(a,r){function y(a){t[a+"pin"]=function(l,r,x,b,m){var d=m&&m.anchorX;m=m&&m.anchorY;"circle"===a&&b>x&&(l-=Math.round((b-x)/2),x=b);var z=t[a](l,r,x,b);if(d&&m){var h=d;"circle"===a?h=l+x/2:(l=z[0],x=z[1],"M"===l[0]&&"L"===x[0]&&(h=(l[1]+x[1])/2));z.push(["M",h,r>m?r:r+b],["L",d,m]);z=z.concat(t.circle(d-1,m-1,2,2))}return z}}var t=r.prototype.symbols;t.flag=function(a,r,y,x,b){var m=b&&b.anchorX||a;b=b&&b.anchorY||
r;var d=t.circle(m-1,b-1,2,2);d.push(["M",m,b],["L",a,r+x],["L",a,r],["L",a+y,r],["L",a+y,r+x],["L",a,r+x],["Z"]);return d};y("circle");y("square");a=a.getRendererType();a!==r&&(a.prototype.symbols.circlepin=t.circlepin,a.prototype.symbols.flag=t.flag,a.prototype.symbols.squarepin=t.squarepin);return t});L(a,"Series/Flags/FlagsSeries.js",[a["Series/Flags/FlagsPoint.js"],a["Core/Globals.js"],a["Series/OnSeriesComposition.js"],a["Core/Renderer/RendererUtilities.js"],a["Core/Series/SeriesRegistry.js"],
a["Core/Renderer/SVG/SVGElement.js"],a["Core/Utilities.js"]],function(a,r,y,t,l,D,E){var x=this&&this.__extends||function(){var b=function(a,d){b=Object.setPrototypeOf||{__proto__:[]}instanceof Array&&function(b,a){b.__proto__=a}||function(b,a){for(var d in a)a.hasOwnProperty(d)&&(b[d]=a[d])};return b(a,d)};return function(a,d){function q(){this.constructor=a}b(a,d);a.prototype=null===d?Object.create(d):(q.prototype=d.prototype,new q)}}();r=r.noop;var b=t.distribute,m=l.series,d=l.seriesTypes.column,
z=E.addEvent,h=E.defined;t=E.extend;var K=E.merge,G=E.objectEach,I=E.wrap;E=function(a){function q(){var b=null!==a&&a.apply(this,arguments)||this;b.data=void 0;b.options=void 0;b.points=void 0;return b}x(q,a);q.prototype.animate=function(b){b&&this.setClip()};q.prototype.drawPoints=function(){var a=this.points,d=this.chart,q=d.renderer,m=d.inverted,u=this.options,l=u.y,k,C=this.yAxis,f={},e=[];for(k=a.length;k--;){var c=a[k];var n=(m?c.plotY:c.plotX)>this.xAxis.len;var p=c.plotX;var g=c.stackIndex;
var v=c.options.shape||u.shape;var w=c.plotY;"undefined"!==typeof w&&(w=c.plotY+l-("undefined"!==typeof g&&g*u.stackDistance));c.anchorX=g?void 0:c.plotX;var Q=g?void 0:c.plotY;var O="flag"!==v;g=c.graphic;"undefined"!==typeof w&&0<=p&&!n?(g&&c.hasNewShapeType()&&(g=g.destroy()),g||(g=c.graphic=q.label("",null,null,v,null,null,u.useHTML).addClass("highcharts-point").add(this.markerGroup),c.graphic.div&&(c.graphic.div.point=c),g.isNew=!0),g.attr({align:O?"center":"left",width:u.width,height:u.height,
"text-align":u.textAlign}),d.styledMode||g.attr(this.pointAttribs(c)).css(K(u.style,c.style)).shadow(u.shadow),0<p&&(p-=g.strokeWidth()%2),v={y:w,anchorY:Q},u.allowOverlapX&&(v.x=p,v.anchorX=c.anchorX),g.attr({text:c.options.title||u.title||"A"})[g.isNew?"attr":"animate"](v),u.allowOverlapX||(f[c.plotX]?f[c.plotX].size=Math.max(f[c.plotX].size,g.width):f[c.plotX]={align:O?.5:0,size:g.width,target:p,anchorX:p}),c.tooltipPos=[p,w+C.pos-d.plotTop]):g&&(c.graphic=g.destroy())}if(!u.allowOverlapX){var B=
100;G(f,function(g){g.plotX=g.anchorX;e.push(g);B=Math.max(g.size,B)});b(e,m?C.len:this.xAxis.len,B);a.forEach(function(g){var c=g.graphic&&f[g.plotX];c&&(g.graphic[g.graphic.isNew?"attr":"animate"]({x:c.pos+c.align*c.size,anchorX:g.anchorX}),h(c.pos)?g.graphic.isNew=!1:(g.graphic.attr({x:-9999,anchorX:-9999}),g.graphic.isNew=!0))})}u.useHTML&&I(this.markerGroup,"on",function(g){return D.prototype.on.apply(g.apply(this,[].slice.call(arguments,1)),[].slice.call(arguments,1))})};q.prototype.drawTracker=
function(){var b=this.points;a.prototype.drawTracker.call(this);b.forEach(function(a){var d=a.graphic;d&&(a.unbindMouseOver&&a.unbindMouseOver(),a.unbindMouseOver=z(d.element,"mouseover",function(){0<a.stackIndex&&!a.raised&&(a._y=d.y,d.attr({y:a._y-8}),a.raised=!0);b.forEach(function(b){b!==a&&b.raised&&b.graphic&&(b.graphic.attr({y:b._y}),b.raised=!1)})}))})};q.prototype.pointAttribs=function(b,a){var d=this.options,q=b&&b.color||this.color,h=d.lineColor,m=b&&b.lineWidth;b=b&&b.fillColor||d.fillColor;
a&&(b=d.states[a].fillColor,h=d.states[a].lineColor,m=d.states[a].lineWidth);return{fill:b||q,stroke:h||q,"stroke-width":m||d.lineWidth||0}};q.prototype.setClip=function(){m.prototype.setClip.apply(this,arguments);!1!==this.options.clip&&this.sharedClipKey&&this.markerGroup&&this.markerGroup.clip(this.chart.sharedClips[this.sharedClipKey])};q.defaultOptions=K(d.defaultOptions,{pointRange:0,allowOverlapX:!1,shape:"flag",stackDistance:12,textAlign:"center",tooltip:{pointFormat:"{point.text}"},threshold:null,
y:-30,fillColor:"#ffffff",lineWidth:1,states:{hover:{lineColor:"#000000",fillColor:"#ccd6eb"}},style:{fontSize:"11px",fontWeight:"bold"}});return q}(d);y.compose(E);t(E.prototype,{allowDG:!1,forceCrop:!0,invertible:!1,noSharedTooltip:!0,pointClass:a,sorted:!1,takeOrdinalPosition:!1,trackerGroups:["markerGroup"],buildKDTree:r,init:m.prototype.init,invertGroups:r});l.registerSeriesType("flags",E);"";"";return E});L(a,"Core/Axis/ScrollbarAxis.js",[a["Core/Utilities.js"]],function(a){var r=a.addEvent,
y=a.defined,t=a.pick;return function(){function a(){}a.compose=function(l,E){if(-1===a.composed.indexOf(l))a.composed.push(l);else return l;var x=function(b){var a=t(b.options&&b.options.min,b.min),d=t(b.options&&b.options.max,b.max);return{axisMin:a,axisMax:d,scrollMin:y(b.dataMin)?Math.min(a,b.min,b.dataMin,t(b.threshold,Infinity)):a,scrollMax:y(b.dataMax)?Math.max(d,b.max,b.dataMax,t(b.threshold,-Infinity)):d}};r(l,"afterInit",function(){var b=this;b.options&&b.options.scrollbar&&b.options.scrollbar.enabled&&
(b.options.scrollbar.vertical=!b.horiz,b.options.startOnTick=b.options.endOnTick=!1,b.scrollbar=new E(b.chart.renderer,b.options.scrollbar,b.chart),r(b.scrollbar,"changed",function(a){var d=x(b),m=d.axisMax,h=d.scrollMin,l=d.scrollMax-h;y(d.axisMin)&&y(m)&&(b.horiz&&!b.reversed||!b.horiz&&b.reversed?(d=h+l*this.to,h+=l*this.from):(d=h+l*(1-this.from),h+=l*(1-this.to)),this.shouldUpdateExtremes(a.DOMType)?b.setExtremes(h,d,!0,"mousemove"!==a.DOMType&&"touchmove"!==a.DOMType,a):this.setRange(this.from,
this.to))}))});r(l,"afterRender",function(){var b=x(this),a=b.scrollMin,d=b.scrollMax;b=this.scrollbar;var l=this.axisTitleMargin+(this.titleOffset||0),h=this.chart.scrollbarsOffsets,t=this.options.margin||0;b&&(this.horiz?(this.opposite||(h[1]+=l),b.position(this.left,this.top+this.height+2+h[1]-(this.opposite?t:0),this.width,this.height),this.opposite||(h[1]+=t),l=1):(this.opposite&&(h[0]+=l),b.position(b.options.opposite?this.left+this.width+2+h[0]-(this.opposite?0:t):this.opposite?0:t,this.top,
this.width,this.height),this.opposite&&(h[0]+=t),l=0),h[l]+=b.size+b.options.margin,isNaN(a)||isNaN(d)||!y(this.min)||!y(this.max)||this.min===this.max?b.setRange(0,1):(h=(this.min-a)/(d-a),a=(this.max-a)/(d-a),this.horiz&&!this.reversed||!this.horiz&&this.reversed?b.setRange(h,a):b.setRange(1-a,1-h)))});r(l,"afterGetOffset",function(){var b=this.scrollbar&&!this.scrollbar.options.opposite;b=this.horiz?2:b?3:1;var a=this.scrollbar;a&&(this.chart.scrollbarsOffsets=[0,0],this.chart.axisOffset[b]+=a.size+
a.options.margin)});return l};a.composed=[];return a}()});L(a,"Core/ScrollbarDefaults.js",[a["Core/Globals.js"]],function(a){return{height:a.isTouchDevice?20:14,barBorderRadius:0,buttonBorderRadius:0,liveRedraw:void 0,margin:10,minWidth:6,opposite:!0,step:.2,zIndex:3,barBackgroundColor:"#cccccc",barBorderWidth:1,barBorderColor:"#cccccc",buttonArrowColor:"#333333",buttonBackgroundColor:"#e6e6e6",buttonBorderColor:"#cccccc",buttonBorderWidth:1,rifleColor:"#333333",trackBackgroundColor:"#f2f2f2",trackBorderColor:"#f2f2f2",
trackBorderWidth:1}});L(a,"Core/Scrollbar.js",[a["Core/DefaultOptions.js"],a["Core/Globals.js"],a["Core/Axis/ScrollbarAxis.js"],a["Core/ScrollbarDefaults.js"],a["Core/Utilities.js"]],function(a,r,y,t,l){var D=a.defaultOptions,E=l.addEvent,x=l.correctFloat,b=l.defined,m=l.destroyObjectProperties,d=l.fireEvent,z=l.merge,h=l.pick,K=l.removeEvent;a=function(){function a(a,b,d){this._events=[];this.chart=void 0;this.from=this.chartY=this.chartX=0;this.scrollbar=this.renderer=this.options=this.group=void 0;
this.scrollbarButtons=[];this.scrollbarGroup=void 0;this.scrollbarLeft=0;this.scrollbarRifles=void 0;this.scrollbarStrokeWidth=1;this.to=this.size=this.scrollbarTop=0;this.track=void 0;this.trackBorderWidth=1;this.userOptions=void 0;this.y=this.x=0;this.init(a,b,d)}a.compose=function(b){y.compose(b,a)};a.swapXY=function(a,b){b&&a.forEach(function(a){for(var b=a.length,d,q=0;q<b;q+=2)d=a[q+1],"number"===typeof d&&(a[q+1]=a[q+2],a[q+2]=d)});return a};a.prototype.addEvents=function(){var a=this.options.inverted?
[1,0]:[0,1],b=this.scrollbarButtons,d=this.scrollbarGroup.element,h=this.track.element,l=this.mouseDownHandler.bind(this),m=this.mouseMoveHandler.bind(this),t=this.mouseUpHandler.bind(this);a=[[b[a[0]].element,"click",this.buttonToMinClick.bind(this)],[b[a[1]].element,"click",this.buttonToMaxClick.bind(this)],[h,"click",this.trackClick.bind(this)],[d,"mousedown",l],[d.ownerDocument,"mousemove",m],[d.ownerDocument,"mouseup",t]];r.hasTouch&&a.push([d,"touchstart",l],[d.ownerDocument,"touchmove",m],
[d.ownerDocument,"touchend",t]);a.forEach(function(a){E.apply(null,a)});this._events=a};a.prototype.buttonToMaxClick=function(a){var b=(this.to-this.from)*h(this.options.step,.2);this.updatePosition(this.from+b,this.to+b);d(this,"changed",{from:this.from,to:this.to,trigger:"scrollbar",DOMEvent:a})};a.prototype.buttonToMinClick=function(a){var b=x(this.to-this.from)*h(this.options.step,.2);this.updatePosition(x(this.from-b),x(this.to-b));d(this,"changed",{from:this.from,to:this.to,trigger:"scrollbar",
DOMEvent:a})};a.prototype.cursorToScrollbarPosition=function(a){var b=this.options;b=b.minWidth>this.calculatedWidth?b.minWidth:0;return{chartX:(a.chartX-this.x-this.xOffset)/(this.barWidth-b),chartY:(a.chartY-this.y-this.yOffset)/(this.barWidth-b)}};a.prototype.destroy=function(){var a=this,b=a.chart.scroller;a.removeEvents();["track","scrollbarRifles","scrollbar","scrollbarGroup","group"].forEach(function(b){a[b]&&a[b].destroy&&(a[b]=a[b].destroy())});b&&a===b.scrollbar&&(b.scrollbar=null,m(b.scrollbarButtons))};
a.prototype.drawScrollbarButton=function(b){var d=this.renderer,q=this.scrollbarButtons,h=this.options,l=this.size,m=d.g().add(this.group);q.push(m);m=d.rect().addClass("highcharts-scrollbar-button").add(m);this.chart.styledMode||m.attr({stroke:h.buttonBorderColor,"stroke-width":h.buttonBorderWidth,fill:h.buttonBackgroundColor});m.attr(m.crisp({x:-.5,y:-.5,width:l+1,height:l+1,r:h.buttonBorderRadius},m.strokeWidth()));m=d.path(a.swapXY([["M",l/2+(b?-1:1),l/2-3],["L",l/2+(b?-1:1),l/2+3],["L",l/2+(b?
2:-2),l/2]],h.vertical)).addClass("highcharts-scrollbar-arrow").add(q[b]);this.chart.styledMode||m.attr({fill:h.buttonArrowColor})};a.prototype.init=function(a,b,d){this.scrollbarButtons=[];this.renderer=a;this.userOptions=b;this.options=z(t,D.scrollbar,b);this.chart=d;this.size=h(this.options.size,this.options.height);b.enabled&&(this.render(),this.addEvents())};a.prototype.mouseDownHandler=function(a){a=this.chart.pointer.normalize(a);a=this.cursorToScrollbarPosition(a);this.chartX=a.chartX;this.chartY=
a.chartY;this.initPositions=[this.from,this.to];this.grabbedCenter=!0};a.prototype.mouseMoveHandler=function(a){var b=this.chart.pointer.normalize(a),q=this.options.vertical?"chartY":"chartX",h=this.initPositions||[];!this.grabbedCenter||a.touches&&0===a.touches[0][q]||(b=this.cursorToScrollbarPosition(b)[q],q=this[q],q=b-q,this.hasDragged=!0,this.updatePosition(h[0]+q,h[1]+q),this.hasDragged&&d(this,"changed",{from:this.from,to:this.to,trigger:"scrollbar",DOMType:a.type,DOMEvent:a}))};a.prototype.mouseUpHandler=
function(a){this.hasDragged&&d(this,"changed",{from:this.from,to:this.to,trigger:"scrollbar",DOMType:a.type,DOMEvent:a});this.grabbedCenter=this.hasDragged=this.chartX=this.chartY=null};a.prototype.position=function(a,b,d,h){var q=this.options.vertical,l=this.rendered?"animate":"attr",m=0;this.x=a;this.y=b+this.trackBorderWidth;this.width=d;this.xOffset=this.height=h;this.yOffset=m;q?(this.width=this.yOffset=d=m=this.size,this.xOffset=b=0,this.barWidth=h-2*d,this.x=a+=this.options.margin):(this.height=
this.xOffset=h=b=this.size,this.barWidth=d-2*h,this.y+=this.options.margin);this.group[l]({translateX:a,translateY:this.y});this.track[l]({width:d,height:h});this.scrollbarButtons[1][l]({translateX:q?0:d-b,translateY:q?h-m:0})};a.prototype.removeEvents=function(){this._events.forEach(function(a){K.apply(null,a)});this._events.length=0};a.prototype.render=function(){var b=this.renderer,d=this.options,h=this.size,l=this.chart.styledMode,m=b.g("scrollbar").attr({zIndex:d.zIndex,translateY:-99999}).add();
this.group=m;this.track=b.rect().addClass("highcharts-scrollbar-track").attr({x:0,r:d.trackBorderRadius||0,height:h,width:h}).add(m);l||this.track.attr({fill:d.trackBackgroundColor,stroke:d.trackBorderColor,"stroke-width":d.trackBorderWidth});this.trackBorderWidth=this.track.strokeWidth();this.track.attr({y:-this.trackBorderWidth%2/2});this.scrollbarGroup=b.g().add(m);this.scrollbar=b.rect().addClass("highcharts-scrollbar-thumb").attr({height:h,width:h,r:d.barBorderRadius||0}).add(this.scrollbarGroup);
this.scrollbarRifles=b.path(a.swapXY([["M",-3,h/4],["L",-3,2*h/3],["M",0,h/4],["L",0,2*h/3],["M",3,h/4],["L",3,2*h/3]],d.vertical)).addClass("highcharts-scrollbar-rifles").add(this.scrollbarGroup);l||(this.scrollbar.attr({fill:d.barBackgroundColor,stroke:d.barBorderColor,"stroke-width":d.barBorderWidth}),this.scrollbarRifles.attr({stroke:d.rifleColor,"stroke-width":1}));this.scrollbarStrokeWidth=this.scrollbar.strokeWidth();this.scrollbarGroup.translate(-this.scrollbarStrokeWidth%2/2,-this.scrollbarStrokeWidth%
2/2);this.drawScrollbarButton(0);this.drawScrollbarButton(1)};a.prototype.setRange=function(a,d){var h=this.options,l=h.vertical,m=h.minWidth,t=this.barWidth,r=!this.rendered||this.hasDragged||this.chart.navigator&&this.chart.navigator.hasDragged?"attr":"animate";if(b(t)){var u=t*Math.min(d,1);a=Math.max(a,0);var z=Math.ceil(t*a);this.calculatedWidth=u=x(u-z);u<m&&(z=(t-m+u)*a,u=m);m=Math.floor(z+this.xOffset+this.yOffset);t=u/2-.5;this.from=a;this.to=d;l?(this.scrollbarGroup[r]({translateY:m}),this.scrollbar[r]({height:u}),
this.scrollbarRifles[r]({translateY:t}),this.scrollbarTop=m,this.scrollbarLeft=0):(this.scrollbarGroup[r]({translateX:m}),this.scrollbar[r]({width:u}),this.scrollbarRifles[r]({translateX:t}),this.scrollbarLeft=m,this.scrollbarTop=0);12>=u?this.scrollbarRifles.hide():this.scrollbarRifles.show(!0);!1===h.showFull&&(0>=a&&1<=d?this.group.hide():this.group.show());this.rendered=!0}};a.prototype.shouldUpdateExtremes=function(a){return h(this.options.liveRedraw,r.svg&&!r.isTouchDevice&&!this.chart.isBoosting)||
"mouseup"===a||"touchend"===a||!b(a)};a.prototype.trackClick=function(a){var b=this.chart.pointer.normalize(a),h=this.to-this.from,m=this.y+this.scrollbarTop,l=this.x+this.scrollbarLeft;this.options.vertical&&b.chartY>m||!this.options.vertical&&b.chartX>l?this.updatePosition(this.from+h,this.to+h):this.updatePosition(this.from-h,this.to-h);d(this,"changed",{from:this.from,to:this.to,trigger:"scrollbar",DOMEvent:a})};a.prototype.update=function(a){this.destroy();this.init(this.chart.renderer,z(!0,
this.options,a),this.chart)};a.prototype.updatePosition=function(a,b){1<b&&(a=x(1-x(b-a)),b=1);0>a&&(b=x(b-a),a=0);this.from=a;this.to=b};a.defaultOptions=t;return a}();D.scrollbar=z(!0,a.defaultOptions,D.scrollbar);return a});L(a,"Core/Axis/NavigatorAxis.js",[a["Core/Globals.js"],a["Core/Utilities.js"]],function(a,r){var y=a.isTouchDevice,t=r.addEvent,l=r.correctFloat,D=r.defined,E=r.isNumber,x=r.pick,b=function(){function a(a){this.axis=a}a.prototype.destroy=function(){this.axis=void 0};a.prototype.toFixedRange=
function(a,b,h,m){var d=this.axis,t=d.chart;t=t&&t.fixedRange;var r=(d.pointRange||0)/2;a=x(h,d.translate(a,!0,!d.horiz));b=x(m,d.translate(b,!0,!d.horiz));d=t&&(b-a)/t;D(h)||(a=l(a+r));D(m)||(b=l(b-r));.7<d&&1.3>d&&(m?a=b-t:b=a+t);E(a)&&E(b)||(a=b=void 0);return{min:a,max:b}};return a}();return function(){function a(){}a.compose=function(a){a.keepProps.push("navigatorAxis");t(a,"init",function(){this.navigatorAxis||(this.navigatorAxis=new b(this))});t(a,"zoom",function(a){var b=this.chart.options,
d=b.navigator,l=this.navigatorAxis,m=b.chart.pinchType,t=b.rangeSelector;b=b.chart.zoomType;this.isXAxis&&(d&&d.enabled||t&&t.enabled)&&("y"===b?a.zoomed=!1:(!y&&"xy"===b||y&&"xy"===m)&&this.options.range&&(d=l.previousZoom,D(a.newMin)?l.previousZoom=[this.min,this.max]:d&&(a.newMin=d[0],a.newMax=d[1],l.previousZoom=void 0)));"undefined"!==typeof a.zoomed&&a.preventDefault()})};a.AdditionsClass=b;return a}()});L(a,"Core/Navigator.js",[a["Core/Axis/Axis.js"],a["Core/Chart/Chart.js"],a["Core/Color/Color.js"],
a["Core/Globals.js"],a["Core/Axis/NavigatorAxis.js"],a["Core/DefaultOptions.js"],a["Core/Renderer/RendererRegistry.js"],a["Core/Scrollbar.js"],a["Core/Series/Series.js"],a["Core/Series/SeriesRegistry.js"],a["Core/Utilities.js"]],function(a,r,y,t,l,D,E,x,b,m,d){y=y.parse;var z=t.hasTouch,h=t.isTouchDevice,K=D.defaultOptions,G=d.addEvent,I=d.clamp,C=d.correctFloat,q=d.defined,L=d.destroyObjectProperties,P=d.erase,F=d.extend,N=d.find,u=d.isArray,M=d.isNumber,k=d.merge,J=d.pick,f=d.removeEvent,e=d.splat,
c=function(a){for(var g=[],c=1;c<arguments.length;c++)g[c-1]=arguments[c];g=[].filter.call(g,M);if(g.length)return Math[a].apply(0,g)};D="undefined"===typeof m.seriesTypes.areaspline?"line":"areaspline";F(K,{navigator:{height:40,margin:25,maskInside:!0,handles:{width:7,height:15,symbols:["navigator-handle","navigator-handle"],enabled:!0,lineWidth:1,backgroundColor:"#f2f2f2",borderColor:"#999999"},maskFill:y("#6685c2").setOpacity(.3).get(),outlineColor:"#cccccc",outlineWidth:1,series:{type:D,fillOpacity:.05,
lineWidth:1,compare:null,dataGrouping:{approximation:"average",enabled:!0,groupPixelWidth:2,firstAnchor:"firstPoint",anchor:"middle",lastAnchor:"lastPoint",units:[["millisecond",[1,2,5,10,20,25,50,100,200,500]],["second",[1,2,5,10,15,30]],["minute",[1,2,5,10,15,30]],["hour",[1,2,3,4,6,8,12]],["day",[1,2,3,4]],["week",[1,2,3]],["month",[1,3,6]],["year",null]]},dataLabels:{enabled:!1,zIndex:2},id:"highcharts-navigator-series",className:"highcharts-navigator-series",lineColor:null,marker:{enabled:!1},
threshold:null},xAxis:{overscroll:0,className:"highcharts-navigator-xaxis",tickLength:0,lineWidth:0,gridLineColor:"#e6e6e6",gridLineWidth:1,tickPixelInterval:200,labels:{align:"left",style:{color:"#999999"},x:3,y:-4},crosshair:!1},yAxis:{className:"highcharts-navigator-yaxis",gridLineWidth:0,startOnTick:!1,endOnTick:!1,minPadding:.1,maxPadding:.1,labels:{enabled:!1},crosshair:!1,title:{text:null},tickLength:0,tickWidth:0}}});E.getRendererType().prototype.symbols["navigator-handle"]=function(a,g,c,
b,e){a=(e&&e.width||0)/2;g=Math.round(a/3)+.5;e=e&&e.height||0;return[["M",-a-1,.5],["L",a,.5],["L",a,e+.5],["L",-a-1,e+.5],["L",-a-1,.5],["M",-g,4],["L",-g,e-3],["M",g-1,4],["L",g-1,e-3]]};var n=function(){function b(g){this.zoomedMin=this.zoomedMax=this.yAxis=this.xAxis=this.top=this.size=this.shades=this.rendered=this.range=this.outlineHeight=this.outline=this.opposite=this.navigatorSize=this.navigatorSeries=this.navigatorOptions=this.navigatorGroup=this.navigatorEnabled=this.left=this.height=
this.handles=this.chart=this.baseSeries=void 0;this.init(g)}b.prototype.drawHandle=function(g,a,c,b){var e=this.navigatorOptions.handles.height;this.handles[a][b](c?{translateX:Math.round(this.left+this.height/2),translateY:Math.round(this.top+parseInt(g,10)+.5-e)}:{translateX:Math.round(this.left+parseInt(g,10)),translateY:Math.round(this.top+this.height/2-e/2-1)})};b.prototype.drawOutline=function(g,a,c,b){var e=this.navigatorOptions.maskInside,v=this.outline.strokeWidth(),d=v/2,w=v%2/2;v=this.outlineHeight;
var p=this.scrollbarHeight||0,n=this.size,f=this.left-p,k=this.top;c?(f-=d,c=k+a+w,a=k+g+w,w=[["M",f+v,k-p-w],["L",f+v,c],["L",f,c],["L",f,a],["L",f+v,a],["L",f+v,k+n+p]],e&&w.push(["M",f+v,c-d],["L",f+v,a+d])):(g+=f+p-w,a+=f+p-w,k+=d,w=[["M",f,k],["L",g,k],["L",g,k+v],["L",a,k+v],["L",a,k],["L",f+n+2*p,k]],e&&w.push(["M",g-d,k],["L",a+d,k]));this.outline[b]({d:w})};b.prototype.drawMasks=function(a,c,b,e){var g=this.left,v=this.top,d=this.height;if(b){var w=[g,g,g];var p=[v,v+a,v+c];var n=[d,d,d];
var f=[a,c-a,this.size-c]}else w=[g,g+a,g+c],p=[v,v,v],n=[a,c-a,this.size-c],f=[d,d,d];this.shades.forEach(function(a,g){a[e]({x:w[g],y:p[g],width:n[g],height:f[g]})})};b.prototype.renderElements=function(){var a=this,c=a.navigatorOptions,b=c.maskInside,e=a.chart,d=e.renderer,p,n={cursor:e.inverted?"ns-resize":"ew-resize"};a.navigatorGroup=p=d.g("navigator").attr({zIndex:8,visibility:"hidden"}).add();[!b,b,!b].forEach(function(g,b){a.shades[b]=d.rect().addClass("highcharts-navigator-mask"+(1===b?
"-inside":"-outside")).add(p);e.styledMode||a.shades[b].attr({fill:g?c.maskFill:"rgba(0,0,0,0)"}).css(1===b&&n)});a.outline=d.path().addClass("highcharts-navigator-outline").add(p);e.styledMode||a.outline.attr({"stroke-width":c.outlineWidth,stroke:c.outlineColor});c.handles.enabled&&[0,1].forEach(function(g){c.handles.inverted=e.inverted;a.handles[g]=d.symbol(c.handles.symbols[g],-c.handles.width/2-1,0,c.handles.width,c.handles.height,c.handles);a.handles[g].attr({zIndex:7-g}).addClass("highcharts-navigator-handle highcharts-navigator-handle-"+
["left","right"][g]).add(p);if(!e.styledMode){var b=c.handles;a.handles[g].attr({fill:b.backgroundColor,stroke:b.borderColor,"stroke-width":b.lineWidth}).css(n)}})};b.prototype.update=function(a){(this.series||[]).forEach(function(a){a.baseSeries&&delete a.baseSeries.navigatorSeries});this.destroy();k(!0,this.chart.options.navigator,this.options,a);this.init(this.chart)};b.prototype.render=function(a,c,b,e){var g=this.chart,v=this.scrollbarHeight,d,p=this.xAxis,n=p.pointRange||0;var w=p.navigatorAxis.fake?
g.xAxis[0]:p;var f=this.navigatorEnabled,k,h=this.rendered;var l=g.inverted;var m=g.xAxis[0].minRange,u=g.xAxis[0].options.maxRange;if(!this.hasDragged||q(b)){a=C(a-n/2);c=C(c+n/2);if(!M(a)||!M(c))if(h)b=0,e=J(p.width,w.width);else return;this.left=J(p.left,g.plotLeft+v+(l?g.plotWidth:0));this.size=k=d=J(p.len,(l?g.plotHeight:g.plotWidth)-2*v);g=l?v:d+2*v;b=J(b,p.toPixels(a,!0));e=J(e,p.toPixels(c,!0));M(b)&&Infinity!==Math.abs(b)||(b=0,e=g);a=p.toValue(b,!0);c=p.toValue(e,!0);var Q=Math.abs(C(c-
a));Q<m?this.grabbedLeft?b=p.toPixels(c-m-n,!0):this.grabbedRight&&(e=p.toPixels(a+m+n,!0)):q(u)&&C(Q-n)>u&&(this.grabbedLeft?b=p.toPixels(c-u-n,!0):this.grabbedRight&&(e=p.toPixels(a+u+n,!0)));this.zoomedMax=I(Math.max(b,e),0,k);this.zoomedMin=I(this.fixedWidth?this.zoomedMax-this.fixedWidth:Math.min(b,e),0,k);this.range=this.zoomedMax-this.zoomedMin;k=Math.round(this.zoomedMax);b=Math.round(this.zoomedMin);f&&(this.navigatorGroup.attr({visibility:"visible"}),h=h&&!this.hasDragged?"animate":"attr",
this.drawMasks(b,k,l,h),this.drawOutline(b,k,l,h),this.navigatorOptions.handles.enabled&&(this.drawHandle(b,0,l,h),this.drawHandle(k,1,l,h)));this.scrollbar&&(l?(l=this.top-v,w=this.left-v+(f||!w.opposite?0:(w.titleOffset||0)+w.axisTitleMargin),v=d+2*v):(l=this.top+(f?this.height:-v),w=this.left-v),this.scrollbar.position(w,l,g,v),this.scrollbar.setRange(this.zoomedMin/(d||1),this.zoomedMax/(d||1)));this.rendered=!0}};b.prototype.addMouseEvents=function(){var a=this,c=a.chart,b=c.container,e=[],d,
p;a.mouseMoveHandler=d=function(c){a.onMouseMove(c)};a.mouseUpHandler=p=function(c){a.onMouseUp(c)};e=a.getPartsEvents("mousedown");e.push(G(c.renderTo,"mousemove",d),G(b.ownerDocument,"mouseup",p));z&&(e.push(G(c.renderTo,"touchmove",d),G(b.ownerDocument,"touchend",p)),e.concat(a.getPartsEvents("touchstart")));a.eventsToUnbind=e;a.series&&a.series[0]&&e.push(G(a.series[0].xAxis,"foundExtremes",function(){c.navigator.modifyNavigatorAxisExtremes()}))};b.prototype.getPartsEvents=function(a){var c=this,
g=[];["shades","handles"].forEach(function(b){c[b].forEach(function(e,v){g.push(G(e.element,a,function(a){c[b+"Mousedown"](a,v)}))})});return g};b.prototype.shadesMousedown=function(a,c){a=this.chart.pointer.normalize(a);var b=this.chart,g=this.xAxis,e=this.zoomedMin,v=this.left,d=this.size,p=this.range,n=a.chartX;b.inverted&&(n=a.chartY,v=this.top);if(1===c)this.grabbedCenter=n,this.fixedWidth=p,this.dragOffset=n-e;else{a=n-v-p/2;if(0===c)a=Math.max(0,a);else if(2===c&&a+p>=d)if(a=d-p,this.reversedExtremes){a-=
p;var f=this.getUnionExtremes().dataMin}else var k=this.getUnionExtremes().dataMax;a!==e&&(this.fixedWidth=p,c=g.navigatorAxis.toFixedRange(a,a+p,f,k),q(c.min)&&b.xAxis[0].setExtremes(Math.min(c.min,c.max),Math.max(c.min,c.max),!0,null,{trigger:"navigator"}))}};b.prototype.handlesMousedown=function(a,c){this.chart.pointer.normalize(a);a=this.chart;var b=a.xAxis[0],g=this.reversedExtremes;0===c?(this.grabbedLeft=!0,this.otherHandlePos=this.zoomedMax,this.fixedExtreme=g?b.min:b.max):(this.grabbedRight=
!0,this.otherHandlePos=this.zoomedMin,this.fixedExtreme=g?b.max:b.min);a.fixedRange=null};b.prototype.onMouseMove=function(a){var c=this,b=c.chart,g=c.left,e=c.navigatorSize,d=c.range,p=c.dragOffset,n=b.inverted;a.touches&&0===a.touches[0].pageX||(a=b.pointer.normalize(a),b=a.chartX,n&&(g=c.top,b=a.chartY),c.grabbedLeft?(c.hasDragged=!0,c.render(0,0,b-g,c.otherHandlePos)):c.grabbedRight?(c.hasDragged=!0,c.render(0,0,c.otherHandlePos,b-g)):c.grabbedCenter&&(c.hasDragged=!0,b<p?b=p:b>e+p-d&&(b=e+p-
d),c.render(0,0,b-p,b-p+d)),c.hasDragged&&c.scrollbar&&J(c.scrollbar.options.liveRedraw,t.svg&&!h&&!this.chart.isBoosting)&&(a.DOMType=a.type,setTimeout(function(){c.onMouseUp(a)},0)))};b.prototype.onMouseUp=function(a){var c=this.chart,b=this.xAxis,g=this.scrollbar,e=a.DOMEvent||a,d=c.inverted,p=this.rendered&&!this.hasDragged?"animate":"attr";if(this.hasDragged&&(!g||!g.hasDragged)||"scrollbar"===a.trigger){g=this.getUnionExtremes();if(this.zoomedMin===this.otherHandlePos)var n=this.fixedExtreme;
else if(this.zoomedMax===this.otherHandlePos)var f=this.fixedExtreme;this.zoomedMax===this.size&&(f=this.reversedExtremes?g.dataMin:g.dataMax);0===this.zoomedMin&&(n=this.reversedExtremes?g.dataMax:g.dataMin);b=b.navigatorAxis.toFixedRange(this.zoomedMin,this.zoomedMax,n,f);q(b.min)&&c.xAxis[0].setExtremes(Math.min(b.min,b.max),Math.max(b.min,b.max),!0,this.hasDragged?!1:null,{trigger:"navigator",triggerOp:"navigator-drag",DOMEvent:e})}"mousemove"!==a.DOMType&&"touchmove"!==a.DOMType&&(this.grabbedLeft=
this.grabbedRight=this.grabbedCenter=this.fixedWidth=this.fixedExtreme=this.otherHandlePos=this.hasDragged=this.dragOffset=null);this.navigatorEnabled&&M(this.zoomedMin)&&M(this.zoomedMax)&&(c=Math.round(this.zoomedMin),a=Math.round(this.zoomedMax),this.shades&&this.drawMasks(c,a,d,p),this.outline&&this.drawOutline(c,a,d,p),this.navigatorOptions.handles.enabled&&Object.keys(this.handles).length===this.handles.length&&(this.drawHandle(c,0,d,p),this.drawHandle(a,1,d,p)))};b.prototype.removeEvents=function(){this.eventsToUnbind&&
(this.eventsToUnbind.forEach(function(a){a()}),this.eventsToUnbind=void 0);this.removeBaseSeriesEvents()};b.prototype.removeBaseSeriesEvents=function(){var a=this.baseSeries||[];this.navigatorEnabled&&a[0]&&(!1!==this.navigatorOptions.adaptToUpdatedData&&a.forEach(function(a){f(a,"updatedData",this.updatedDataHandler)},this),a[0].xAxis&&f(a[0].xAxis,"foundExtremes",this.modifyBaseAxisExtremes))};b.prototype.init=function(b){var g=b.options,e=g.navigator,d=e.enabled,p=g.scrollbar,n=p.enabled;g=d?e.height:
0;var f=n?p.height:0;this.handles=[];this.shades=[];this.chart=b;this.setBaseSeries();this.height=g;this.scrollbarHeight=f;this.scrollbarEnabled=n;this.navigatorEnabled=d;this.navigatorOptions=e;this.scrollbarOptions=p;this.outlineHeight=g+f;this.opposite=J(e.opposite,!(d||!b.inverted));var h=this;d=h.baseSeries;p=b.xAxis.length;n=b.yAxis.length;var m=d&&d[0]&&d[0].xAxis||b.xAxis[0]||{options:{}};b.isDirtyBox=!0;h.navigatorEnabled?(h.xAxis=new a(b,k({breaks:m.options.breaks,ordinal:m.options.ordinal},
e.xAxis,{id:"navigator-x-axis",yAxis:"navigator-y-axis",isX:!0,type:"datetime",index:p,isInternal:!0,offset:0,keepOrdinalPadding:!0,startOnTick:!1,endOnTick:!1,minPadding:0,maxPadding:0,zoomEnabled:!1},b.inverted?{offsets:[f,0,-f,0],width:g}:{offsets:[0,-f,0,f],height:g})),h.yAxis=new a(b,k(e.yAxis,{id:"navigator-y-axis",alignTicks:!1,offset:0,index:n,isInternal:!0,reversed:J(e.yAxis&&e.yAxis.reversed,b.yAxis[0]&&b.yAxis[0].reversed,!1),zoomEnabled:!1},b.inverted?{width:g}:{height:g})),d||e.series.data?
h.updateNavigatorSeries(!1):0===b.series.length&&(h.unbindRedraw=G(b,"beforeRedraw",function(){0<b.series.length&&!h.series&&(h.setBaseSeries(),h.unbindRedraw())})),h.reversedExtremes=b.inverted&&!h.xAxis.reversed||!b.inverted&&h.xAxis.reversed,h.renderElements(),h.addMouseEvents()):(h.xAxis={chart:b,navigatorAxis:{fake:!0},translate:function(a,g){var e=b.xAxis[0],d=e.getExtremes(),p=e.len-2*f,n=c("min",e.options.min,d.dataMin);e=c("max",e.options.max,d.dataMax)-n;return g?a*e/p+n:p*(a-n)/e},toPixels:function(a){return this.translate(a)},
toValue:function(a){return this.translate(a,!0)}},h.xAxis.navigatorAxis.axis=h.xAxis,h.xAxis.navigatorAxis.toFixedRange=l.AdditionsClass.prototype.toFixedRange.bind(h.xAxis.navigatorAxis));b.options.scrollbar.enabled&&(b.scrollbar=h.scrollbar=new x(b.renderer,k(b.options.scrollbar,{margin:h.navigatorEnabled?0:10,vertical:b.inverted}),b),G(h.scrollbar,"changed",function(a){var c=h.size,b=c*this.to;c*=this.from;h.hasDragged=h.scrollbar.hasDragged;h.render(0,0,c,b);this.shouldUpdateExtremes(a.DOMType)&&
setTimeout(function(){h.onMouseUp(a)})}));h.addBaseSeriesEvents();h.addChartEvents()};b.prototype.getUnionExtremes=function(a){var b=this.chart.xAxis[0],g=this.xAxis,e=g.options,d=b.options,p;a&&null===b.dataMin||(p={dataMin:J(e&&e.min,c("min",d.min,b.dataMin,g.dataMin,g.min)),dataMax:J(e&&e.max,c("max",d.max,b.dataMax,g.dataMax,g.max))});return p};b.prototype.setBaseSeries=function(a,c){var b=this.chart,g=this.baseSeries=[];a=a||b.options&&b.options.navigator.baseSeries||(b.series.length?N(b.series,
function(a){return!a.options.isInternal}).index:0);(b.series||[]).forEach(function(c,b){c.options.isInternal||!c.options.showInNavigator&&(b!==a&&c.options.id!==a||!1===c.options.showInNavigator)||g.push(c)});this.xAxis&&!this.xAxis.navigatorAxis.fake&&this.updateNavigatorSeries(!0,c)};b.prototype.updateNavigatorSeries=function(a,c){var b=this,g=b.chart,d=b.baseSeries,p,n,v=b.navigatorOptions.series,h,l={enableMouseTracking:!1,index:null,linkedTo:null,group:"nav",padXAxis:!1,xAxis:"navigator-x-axis",
yAxis:"navigator-y-axis",showInLegend:!1,stacking:void 0,isInternal:!0,states:{inactive:{opacity:1}}},m=b.series=(b.series||[]).filter(function(a){var c=a.baseSeries;return 0>d.indexOf(c)?(c&&(f(c,"updatedData",b.updatedDataHandler),delete c.navigatorSeries),a.chart&&a.destroy(),!1):!0});d&&d.length&&d.forEach(function(a){var f=a.navigatorSeries,w=F({color:a.color,visible:a.visible},u(v)?K.navigator.series:v);f&&!1===b.navigatorOptions.adaptToUpdatedData||(l.name="Navigator "+d.length,p=a.options||
{},h=p.navigatorOptions||{},w.dataLabels=e(w.dataLabels),n=k(p,l,w,h),n.pointRange=J(w.pointRange,h.pointRange,K.plotOptions[n.type||"line"].pointRange),w=h.data||w.data,b.hasNavigatorData=b.hasNavigatorData||!!w,n.data=w||p.data&&p.data.slice(0),f&&f.options?f.update(n,c):(a.navigatorSeries=g.initSeries(n),a.navigatorSeries.baseSeries=a,m.push(a.navigatorSeries)))});if(v.data&&(!d||!d.length)||u(v))b.hasNavigatorData=!1,v=e(v),v.forEach(function(a,c){l.name="Navigator "+(m.length+1);n=k(K.navigator.series,
{color:g.series[c]&&!g.series[c].options.isInternal&&g.series[c].color||g.options.colors[c]||g.options.colors[0]},l,a);n.data=a.data;n.data&&(b.hasNavigatorData=!0,m.push(g.initSeries(n)))});a&&this.addBaseSeriesEvents()};b.prototype.addBaseSeriesEvents=function(){var a=this,c=a.baseSeries||[];c[0]&&c[0].xAxis&&c[0].eventsToUnbind.push(G(c[0].xAxis,"foundExtremes",this.modifyBaseAxisExtremes));c.forEach(function(c){c.eventsToUnbind.push(G(c,"show",function(){this.navigatorSeries&&this.navigatorSeries.setVisible(!0,
!1)}));c.eventsToUnbind.push(G(c,"hide",function(){this.navigatorSeries&&this.navigatorSeries.setVisible(!1,!1)}));!1!==this.navigatorOptions.adaptToUpdatedData&&c.xAxis&&c.eventsToUnbind.push(G(c,"updatedData",this.updatedDataHandler));c.eventsToUnbind.push(G(c,"remove",function(){this.navigatorSeries&&(P(a.series,this.navigatorSeries),q(this.navigatorSeries.options)&&this.navigatorSeries.remove(!1),delete this.navigatorSeries)}))},this)};b.prototype.getBaseSeriesMin=function(a){return this.baseSeries.reduce(function(a,
c){return Math.min(a,c.xData?c.xData[0]:a)},a)};b.prototype.modifyNavigatorAxisExtremes=function(){var a=this.xAxis,c;"undefined"!==typeof a.getExtremes&&(!(c=this.getUnionExtremes(!0))||c.dataMin===a.min&&c.dataMax===a.max||(a.min=c.dataMin,a.max=c.dataMax))};b.prototype.modifyBaseAxisExtremes=function(){var a=this.chart.navigator,c=this.getExtremes(),b=c.dataMin,e=c.dataMax;c=c.max-c.min;var d=a.stickToMin,p=a.stickToMax,n=J(this.options.overscroll,0),f=a.series&&a.series[0],k=!!this.setExtremes;
if(!this.eventArgs||"rangeSelectorButton"!==this.eventArgs.trigger){if(d){var h=b;var l=h+c}p&&(l=e+n,d||(h=Math.max(b,l-c,a.getBaseSeriesMin(f&&f.xData?f.xData[0]:-Number.MAX_VALUE))));k&&(d||p)&&M(h)&&(this.min=this.userMin=h,this.max=this.userMax=l)}a.stickToMin=a.stickToMax=null};b.prototype.updatedDataHandler=function(){var a=this.chart.navigator,c=this.navigatorSeries;a.stickToMax=a.reversedExtremes?0===Math.round(a.zoomedMin):Math.round(a.zoomedMax)>=Math.round(a.size);a.stickToMin=a.shouldStickToMin(this,
a);c&&!a.hasNavigatorData&&(c.options.pointStart=this.xData[0],c.setData(this.options.data,!1,null,!1))};b.prototype.shouldStickToMin=function(a,c){c=c.getBaseSeriesMin(a.xData[0]);var b=a.xAxis;a=b.max;var g=b.min;b=b.options.range;return M(a)&&M(g)?b&&0<a-c?a-c<b:g<=c:!1};b.prototype.addChartEvents=function(){this.eventsToUnbind||(this.eventsToUnbind=[]);this.eventsToUnbind.push(G(this.chart,"redraw",function(){var a=this.navigator,c=a&&(a.baseSeries&&a.baseSeries[0]&&a.baseSeries[0].xAxis||this.xAxis[0]);
c&&a.render(c.min,c.max)}),G(this.chart,"getMargins",function(){var a=this.navigator,c=a.opposite?"plotTop":"marginBottom";this.inverted&&(c=a.opposite?"marginRight":"plotLeft");this[c]=(this[c]||0)+(a.navigatorEnabled||!this.inverted?a.outlineHeight:0)+a.navigatorOptions.margin}))};b.prototype.destroy=function(){this.removeEvents();this.xAxis&&(P(this.chart.xAxis,this.xAxis),P(this.chart.axes,this.xAxis));this.yAxis&&(P(this.chart.yAxis,this.yAxis),P(this.chart.axes,this.yAxis));(this.series||[]).forEach(function(a){a.destroy&&
a.destroy()});"series xAxis yAxis shades outline scrollbarTrack scrollbarRifles scrollbarGroup scrollbar navigatorGroup rendered".split(" ").forEach(function(a){this[a]&&this[a].destroy&&this[a].destroy();this[a]=null},this);[this.handles].forEach(function(a){L(a)},this)};return b}();t.Navigator||(t.Navigator=n,l.compose(a),G(r,"beforeShowResetZoom",function(){var a=this.options,c=a.navigator,b=a.rangeSelector;if((c&&c.enabled||b&&b.enabled)&&(!h&&"x"===a.chart.zoomType||h&&"x"===a.chart.pinchType))return!1}),
G(r,"beforeRender",function(){var a=this.options;if(a.navigator.enabled||a.scrollbar.enabled)this.scroller=this.navigator=new n(this)}),G(r,"afterSetChartSize",function(){var a=this.legend,c=this.navigator;if(c){var b=a&&a.options;var e=c.xAxis;var d=c.yAxis;var n=c.scrollbarHeight;this.inverted?(c.left=c.opposite?this.chartWidth-n-c.height:this.spacing[3]+n,c.top=this.plotTop+n):(c.left=J(e.left,this.plotLeft+n),c.top=c.navigatorOptions.top||this.chartHeight-c.height-n-this.spacing[2]-(this.rangeSelector&&
this.extraBottomMargin?this.rangeSelector.getHeight():0)-(b&&"bottom"===b.verticalAlign&&"proximate"!==b.layout&&b.enabled&&!b.floating?a.legendHeight+J(b.margin,10):0)-(this.titleOffset?this.titleOffset[2]:0));e&&d&&(this.inverted?e.options.left=d.options.left=c.left:e.options.top=d.options.top=c.top,e.setAxisSize(),d.setAxisSize())}}),G(r,"update",function(a){var c=a.options.navigator||{},b=a.options.scrollbar||{};this.navigator||this.scroller||!c.enabled&&!b.enabled||(k(!0,this.options.navigator,
c),k(!0,this.options.scrollbar,b),delete a.options.navigator,delete a.options.scrollbar)}),G(r,"afterUpdate",function(a){this.navigator||this.scroller||!this.options.navigator.enabled&&!this.options.scrollbar.enabled||(this.scroller=this.navigator=new n(this),J(a.redraw,!0)&&this.redraw(a.animation))}),G(r,"afterAddSeries",function(){this.navigator&&this.navigator.setBaseSeries(null,!1)}),G(b,"afterUpdate",function(){this.chart.navigator&&!this.options.isInternal&&this.chart.navigator.setBaseSeries(null,
!1)}),r.prototype.callbacks.push(function(a){var c=a.navigator;c&&a.xAxis[0]&&(a=a.xAxis[0].getExtremes(),c.render(a.min,a.max))}));t.Navigator=n;return t.Navigator});L(a,"Extensions/RangeSelector.js",[a["Core/Axis/Axis.js"],a["Core/Chart/Chart.js"],a["Core/Globals.js"],a["Core/DefaultOptions.js"],a["Core/Renderer/SVG/SVGElement.js"],a["Core/Utilities.js"]],function(a,r,y,t,l,D){function E(a){if(-1!==a.indexOf("%L"))return"text";var c="aAdewbBmoyY".split("").some(function(c){return-1!==a.indexOf("%"+
c)}),b="HkIlMS".split("").some(function(c){return-1!==a.indexOf("%"+c)});return c&&b?"datetime-local":c?"date":b?"time":"text"}var x=t.defaultOptions,b=D.addEvent,m=D.createElement,d=D.css,z=D.defined,h=D.destroyObjectProperties,K=D.discardElement,G=D.extend,I=D.find,C=D.fireEvent,q=D.isNumber,P=D.merge,L=D.objectEach,F=D.pad,N=D.pick,u=D.pInt,M=D.splat;G(x,{rangeSelector:{allButtonsEnabled:!1,buttons:void 0,buttonSpacing:5,dropdown:"responsive",enabled:void 0,verticalAlign:"top",buttonTheme:{width:28,
height:18,padding:2,zIndex:7},floating:!1,x:0,y:0,height:void 0,inputBoxBorderColor:"none",inputBoxHeight:17,inputBoxWidth:void 0,inputDateFormat:"%b %e, %Y",inputDateParser:void 0,inputEditDateFormat:"%Y-%m-%d",inputEnabled:!0,inputPosition:{align:"right",x:0,y:0},inputSpacing:5,selected:void 0,buttonPosition:{align:"left",x:0,y:0},inputStyle:{color:"#335cad",cursor:"pointer"},labelStyle:{color:"#666666"}}});G(x.lang,{rangeSelectorZoom:"Zoom",rangeSelectorFrom:"",rangeSelectorTo:"\u2192"});var k=
function(){function e(a){this.buttons=void 0;this.buttonOptions=e.prototype.defaultButtons;this.initialButtonGroupWidth=0;this.options=void 0;this.chart=a;this.init(a)}e.prototype.clickButton=function(c,e){var d=this.chart,g=this.buttonOptions[c],n=d.xAxis[0],f=d.scroller&&d.scroller.getUnionExtremes()||n||{},k=f.dataMin,h=f.dataMax,l=n&&Math.round(Math.min(n.max,N(h,n.max))),m=g.type;f=g._range;var u,t=g.dataGrouping;if(null!==k&&null!==h){d.fixedRange=f;this.setSelected(c);t&&(this.forcedDataGrouping=
!0,a.prototype.setDataGrouping.call(n||{chart:this.chart},t,!1),this.frozenStates=g.preserveDataGrouping);if("month"===m||"year"===m)if(n){m={range:g,max:l,chart:d,dataMin:k,dataMax:h};var r=n.minFromRange.call(m);q(m.newMax)&&(l=m.newMax)}else f=g;else if(f)r=Math.max(l-f,k),l=Math.min(r+f,h);else if("ytd"===m)if(n)"undefined"===typeof h&&(k=Number.MAX_VALUE,h=Number.MIN_VALUE,d.series.forEach(function(a){a=a.xData;k=Math.min(a[0],k);h=Math.max(a[a.length-1],h)}),e=!1),l=this.getYTDExtremes(h,k,
d.time.useUTC),r=u=l.min,l=l.max;else{this.deferredYTDClick=c;return}else"all"===m&&n&&(d.navigator&&d.navigator.baseSeries[0]&&(d.navigator.baseSeries[0].xAxis.options.range=void 0),r=k,l=h);z(r)&&(r+=g._offsetMin);z(l)&&(l+=g._offsetMax);this.dropdown&&(this.dropdown.selectedIndex=c+1);if(n)n.setExtremes(r,l,N(e,!0),void 0,{trigger:"rangeSelectorButton",rangeSelectorButton:g});else{var x=M(d.options.xAxis)[0];var y=x.range;x.range=f;var J=x.min;x.min=u;b(d,"load",function(){x.range=y;x.min=J})}C(this,
"afterBtnClick")}};e.prototype.setSelected=function(a){this.selected=this.options.selected=a};e.prototype.init=function(a){var c=this,e=a.options.rangeSelector,g=e.buttons||c.defaultButtons.slice(),d=e.selected,f=function(){var a=c.minInput,b=c.maxInput;a&&a.blur&&C(a,"blur");b&&b.blur&&C(b,"blur")};c.chart=a;c.options=e;c.buttons=[];c.buttonOptions=g;this.eventsToUnbind=[];this.eventsToUnbind.push(b(a.container,"mousedown",f));this.eventsToUnbind.push(b(a,"resize",f));g.forEach(c.computeButtonRange);
"undefined"!==typeof d&&g[d]&&this.clickButton(d,!1);this.eventsToUnbind.push(b(a,"load",function(){a.xAxis&&a.xAxis[0]&&b(a.xAxis[0],"setExtremes",function(b){this.max-this.min!==a.fixedRange&&"rangeSelectorButton"!==b.trigger&&"updatedData"!==b.trigger&&c.forcedDataGrouping&&!c.frozenStates&&this.setDataGrouping(!1,!1)})}))};e.prototype.updateButtonStates=function(){var a=this,b=this.chart,e=this.dropdown,g=b.xAxis[0],d=Math.round(g.max-g.min),f=!g.hasVisibleSeries,k=b.scroller&&b.scroller.getUnionExtremes()||
g,h=k.dataMin,l=k.dataMax;b=a.getYTDExtremes(l,h,b.time.useUTC);var m=b.min,u=b.max,t=a.selected,r=q(t),x=a.options.allButtonsEnabled,y=a.buttons;a.buttonOptions.forEach(function(c,b){var n=c._range,p=c.type,k=c.count||1,v=y[b],w=0,q=c._offsetMax-c._offsetMin;c=b===t;var B=n>l-h,A=n<g.minRange,H=!1,z=!1;n=n===d;("month"===p||"year"===p)&&d+36E5>=864E5*{month:28,year:365}[p]*k-q&&d-36E5<=864E5*{month:31,year:366}[p]*k+q?n=!0:"ytd"===p?(n=u-m+q===d,H=!c):"all"===p&&(n=g.max-g.min>=l-h,z=!c&&r&&n);p=
!x&&(B||A||z||f);k=c&&n||n&&!r&&!H||c&&a.frozenStates;p?w=3:k&&(r=!0,w=2);v.state!==w&&(v.setState(w),e&&(e.options[b+1].disabled=p,2===w&&(e.selectedIndex=b+1)),0===w&&t===b&&a.setSelected())})};e.prototype.computeButtonRange=function(a){var c=a.type,b=a.count||1,e={millisecond:1,second:1E3,minute:6E4,hour:36E5,day:864E5,week:6048E5};if(e[c])a._range=e[c]*b;else if("month"===c||"year"===c)a._range=864E5*{month:30,year:365}[c]*b;a._offsetMin=N(a.offsetMin,0);a._offsetMax=N(a.offsetMax,0);a._range+=
a._offsetMax-a._offsetMin};e.prototype.getInputValue=function(a){a="min"===a?this.minInput:this.maxInput;var c=this.chart.options.rangeSelector,b=this.chart.time;return a?("text"===a.type&&c.inputDateParser||this.defaultInputDateParser)(a.value,b.useUTC,b):0};e.prototype.setInputValue=function(a,b){var c=this.options,e=this.chart.time,d="min"===a?this.minInput:this.maxInput;a="min"===a?this.minDateBox:this.maxDateBox;if(d){var n=d.getAttribute("data-hc-time");n=z(n)?Number(n):void 0;z(b)&&(z(n)&&
d.setAttribute("data-hc-time-previous",n),d.setAttribute("data-hc-time",b),n=b);d.value=e.dateFormat(this.inputTypeFormats[d.type]||c.inputEditDateFormat,n);a&&a.attr({text:e.dateFormat(c.inputDateFormat,n)})}};e.prototype.setInputExtremes=function(a,b,e){if(a="min"===a?this.minInput:this.maxInput){var c=this.inputTypeFormats[a.type],d=this.chart.time;c&&(b=d.dateFormat(c,b),a.min!==b&&(a.min=b),e=d.dateFormat(c,e),a.max!==e&&(a.max=e))}};e.prototype.showInput=function(a){var b="min"===a?this.minDateBox:
this.maxDateBox;if((a="min"===a?this.minInput:this.maxInput)&&b&&this.inputGroup){var c="text"===a.type,e=this.inputGroup,f=e.translateX;e=e.translateY;var k=this.options.inputBoxWidth;d(a,{width:c?b.width+(k?-2:20)+"px":"auto",height:c?b.height-2+"px":"auto",border:"2px solid silver"});c&&k?d(a,{left:f+b.x+"px",top:e+"px"}):d(a,{left:Math.min(Math.round(b.x+f-(a.offsetWidth-b.width)/2),this.chart.chartWidth-a.offsetWidth)+"px",top:e-(a.offsetHeight-b.height)/2+"px"})}};e.prototype.hideInput=function(a){(a=
"min"===a?this.minInput:this.maxInput)&&d(a,{top:"-9999em",border:0,width:"1px",height:"1px"})};e.prototype.defaultInputDateParser=function(a,b,e){var c=a.split("/").join("-").split(" ").join("T");-1===c.indexOf("T")&&(c+="T00:00");if(b)c+="Z";else{var d;if(d=y.isSafari)d=c,d=!(6<d.length&&(d.lastIndexOf("-")===d.length-6||d.lastIndexOf("+")===d.length-6));d&&(d=(new Date(c)).getTimezoneOffset()/60,c+=0>=d?"+"+F(-d)+":00":"-"+F(d)+":00")}c=Date.parse(c);q(c)||(a=a.split("-"),c=Date.UTC(u(a[0]),u(a[1])-
1,u(a[2])));e&&b&&q(c)&&(c+=e.getTimezoneOffset(c));return c};e.prototype.drawInput=function(a){function b(){var b=k.getInputValue(a),e=c.xAxis[0],d=c.scroller&&c.scroller.xAxis?c.scroller.xAxis:e,g=d.dataMin;d=d.dataMax;var f=k.maxInput,n=k.minInput;b!==Number(r.getAttribute("data-hc-time-previous"))&&q(b)&&(r.setAttribute("data-hc-time-previous",b),t&&f&&q(g)?b>Number(f.getAttribute("data-hc-time"))?b=void 0:b<g&&(b=g):n&&q(d)&&(b<Number(n.getAttribute("data-hc-time"))?b=void 0:b>d&&(b=d)),"undefined"!==
typeof b&&e.setExtremes(t?b:e.min,t?e.max:b,void 0,void 0,{trigger:"rangeSelectorInput"}))}var c=this.chart,e=this.div,f=this.inputGroup,k=this,h=c.renderer.style||{},l=c.renderer,u=c.options.rangeSelector,t="min"===a,A=x.lang[t?"rangeSelectorFrom":"rangeSelectorTo"]||"";A=l.label(A,0).addClass("highcharts-range-label").attr({padding:A?2:0,height:A?u.inputBoxHeight:0}).add(f);l=l.label("",0).addClass("highcharts-range-input").attr({padding:2,width:u.inputBoxWidth,height:u.inputBoxHeight,"text-align":"center"}).on("click",
function(){k.showInput(a);k[a+"Input"].focus()});c.styledMode||l.attr({stroke:u.inputBoxBorderColor,"stroke-width":1});l.add(f);var r=m("input",{name:a,className:"highcharts-range-selector"},void 0,e);r.setAttribute("type",E(u.inputDateFormat||"%b %e, %Y"));c.styledMode||(A.css(P(h,u.labelStyle)),l.css(P({color:"#333333"},h,u.inputStyle)),d(r,G({position:"absolute",border:0,boxShadow:"0 0 15px rgba(0,0,0,0.3)",width:"1px",height:"1px",padding:0,textAlign:"center",fontSize:h.fontSize,fontFamily:h.fontFamily,
top:"-9999em"},u.inputStyle)));r.onfocus=function(){k.showInput(a)};r.onblur=function(){r===y.doc.activeElement&&b();k.hideInput(a);k.setInputValue(a);r.blur()};var z=!1;r.onchange=function(){z||(b(),k.hideInput(a),r.blur())};r.onkeypress=function(a){13===a.keyCode&&b()};r.onkeydown=function(a){z=!0;38!==a.keyCode&&40!==a.keyCode||b()};r.onkeyup=function(){z=!1};return{dateBox:l,input:r,label:A}};e.prototype.getPosition=function(){var a=this.chart,b=a.options.rangeSelector;a="top"===b.verticalAlign?
a.plotTop-a.axisOffset[0]:0;return{buttonTop:a+b.buttonPosition.y,inputTop:a+b.inputPosition.y-10}};e.prototype.getYTDExtremes=function(a,b,e){var c=this.chart.time,d=new c.Date(a),f=c.get("FullYear",d);e=e?c.Date.UTC(f,0,1):+new c.Date(f,0,1);b=Math.max(b,e);d=d.getTime();return{max:Math.min(a||d,d),min:b}};e.prototype.render=function(a,b){var c=this.chart,e=c.renderer,d=c.container,f=c.options,n=f.rangeSelector,k=N(f.chart.style&&f.chart.style.zIndex,0)+1;f=n.inputEnabled;if(!1!==n.enabled){this.rendered||
(this.group=e.g("range-selector-group").attr({zIndex:7}).add(),this.div=m("div",void 0,{position:"relative",height:0,zIndex:k}),this.buttonOptions.length&&this.renderButtons(),d.parentNode&&d.parentNode.insertBefore(this.div,d),f&&(this.inputGroup=e.g("input-group").add(this.group),e=this.drawInput("min"),this.minDateBox=e.dateBox,this.minLabel=e.label,this.minInput=e.input,e=this.drawInput("max"),this.maxDateBox=e.dateBox,this.maxLabel=e.label,this.maxInput=e.input));if(f&&(this.setInputValue("min",
a),this.setInputValue("max",b),a=c.scroller&&c.scroller.getUnionExtremes()||c.xAxis[0]||{},z(a.dataMin)&&z(a.dataMax)&&(c=c.xAxis[0].minRange||0,this.setInputExtremes("min",a.dataMin,Math.min(a.dataMax,this.getInputValue("max"))-c),this.setInputExtremes("max",Math.max(a.dataMin,this.getInputValue("min"))+c,a.dataMax)),this.inputGroup)){var h=0;[this.minLabel,this.minDateBox,this.maxLabel,this.maxDateBox].forEach(function(a){if(a){var b=a.getBBox().width;b&&(a.attr({x:h}),h+=b+n.inputSpacing)}})}this.alignElements();
this.rendered=!0}};e.prototype.renderButtons=function(){var a=this,e=this.buttons,d=this.options,g=x.lang,f=this.chart.renderer,k=P(d.buttonTheme),h=k&&k.states,l=k.width||28;delete k.width;delete k.states;this.buttonGroup=f.g("range-selector-buttons").add(this.group);var q=this.dropdown=m("select",void 0,{position:"absolute",width:"1px",height:"1px",padding:0,border:0,top:"-9999em",cursor:"pointer",opacity:.0001},this.div);b(q,"touchstart",function(){q.style.fontSize="16px"});[[y.isMS?"mouseover":
"mouseenter"],[y.isMS?"mouseout":"mouseleave"],["change","click"]].forEach(function(c){var d=c[0],g=c[1];b(q,d,function(){var b=e[a.currentButtonIndex()];b&&C(b.element,g||d)})});this.zoomText=f.label(g&&g.rangeSelectorZoom||"",0).attr({padding:d.buttonTheme.padding,height:d.buttonTheme.height,paddingLeft:0,paddingRight:0}).add(this.buttonGroup);this.chart.styledMode||(this.zoomText.css(d.labelStyle),k["stroke-width"]=N(k["stroke-width"],0));m("option",{textContent:this.zoomText.textStr,disabled:!0},
void 0,q);this.buttonOptions.forEach(function(b,c){m("option",{textContent:b.title||b.text},void 0,q);e[c]=f.button(b.text,0,0,function(e){var d=b.events&&b.events.click,g;d&&(g=d.call(b,e));!1!==g&&a.clickButton(c);a.isActive=!0},k,h&&h.hover,h&&h.select,h&&h.disabled).attr({"text-align":"center",width:l}).add(a.buttonGroup);b.title&&e[c].attr("title",b.title)})};e.prototype.alignElements=function(){var a=this,b=this.buttonGroup,e=this.buttons,d=this.chart,f=this.group,k=this.inputGroup,h=this.options,
l=this.zoomText,m=d.options,q=m.exporting&&!1!==m.exporting.enabled&&m.navigation&&m.navigation.buttonOptions;m=h.buttonPosition;var u=h.inputPosition,t=h.verticalAlign,r=function(b,c){return q&&a.titleCollision(d)&&"top"===t&&"right"===c.align&&c.y-b.getBBox().height-12<(q.y||0)+(q.height||0)+d.spacing[0]?-40:0},x=d.plotLeft;if(f&&m&&u){var z=m.x-d.spacing[3];if(b){this.positionButtons();if(!this.initialButtonGroupWidth){var y=0;l&&(y+=l.getBBox().width+5);e.forEach(function(a,b){y+=a.width;b!==
e.length-1&&(y+=h.buttonSpacing)});this.initialButtonGroupWidth=y}x-=d.spacing[3];this.updateButtonStates();l=r(b,m);this.alignButtonGroup(l);f.placed=b.placed=d.hasLoaded}b=0;k&&(b=r(k,u),"left"===u.align?z=x:"right"===u.align&&(z=-Math.max(d.axisOffset[1],-b)),k.align({y:u.y,width:k.getBBox().width,align:u.align,x:u.x+z-2},!0,d.spacingBox),k.placed=d.hasLoaded);this.handleCollision(b);f.align({verticalAlign:t},!0,d.spacingBox);k=f.alignAttr.translateY;b=f.getBBox().height+20;r=0;"bottom"===t&&(r=
(r=d.legend&&d.legend.options)&&"bottom"===r.verticalAlign&&r.enabled&&!r.floating?d.legend.legendHeight+N(r.margin,10):0,b=b+r-20,r=k-b-(h.floating?0:h.y)-(d.titleOffset?d.titleOffset[2]:0)-10);if("top"===t)h.floating&&(r=0),d.titleOffset&&d.titleOffset[0]&&(r=d.titleOffset[0]),r+=d.margin[0]-d.spacing[0]||0;else if("middle"===t)if(u.y===m.y)r=k;else if(u.y||m.y)r=0>u.y||0>m.y?r-Math.min(u.y,m.y):k-b;f.translate(h.x,h.y+Math.floor(r));m=this.minInput;u=this.maxInput;k=this.dropdown;h.inputEnabled&&
m&&u&&(m.style.marginTop=f.translateY+"px",u.style.marginTop=f.translateY+"px");k&&(k.style.marginTop=f.translateY+"px")}};e.prototype.alignButtonGroup=function(a,b){var c=this.chart,e=this.buttonGroup,d=this.options.buttonPosition,f=c.plotLeft-c.spacing[3],k=d.x-c.spacing[3];"right"===d.align?k+=a-f:"center"===d.align&&(k-=f/2);e&&e.align({y:d.y,width:N(b,this.initialButtonGroupWidth),align:d.align,x:k},!0,c.spacingBox)};e.prototype.positionButtons=function(){var a=this.buttons,b=this.chart,e=this.options,
d=this.zoomText,f=b.hasLoaded?"animate":"attr",k=e.buttonPosition,h=b.plotLeft,l=h;d&&"hidden"!==d.visibility&&(d[f]({x:N(h+k.x,h)}),l+=k.x+d.getBBox().width+5);this.buttonOptions.forEach(function(b,c){if("hidden"!==a[c].visibility)a[c][f]({x:l}),l+=a[c].width+e.buttonSpacing;else a[c][f]({x:h})})};e.prototype.handleCollision=function(a){var b=this,c=this.chart,e=this.buttonGroup,d=this.inputGroup,f=this.options,k=f.buttonPosition,h=f.dropdown,l=f.inputPosition;f=function(){var a=0;b.buttons.forEach(function(b){b=
b.getBBox();b.width>a&&(a=b.width)});return a};var m=function(b){if(d&&e){var c=d.alignAttr.translateX+d.alignOptions.x-a+d.getBBox().x+2,f=d.alignOptions.width,g=e.alignAttr.translateX+e.getBBox().x;return g+b>c&&c+f>g&&k.y<l.y+d.getBBox().height}return!1},u=function(){d&&e&&d.attr({translateX:d.alignAttr.translateX+(c.axisOffset[1]>=-a?0:-a),translateY:d.alignAttr.translateY+e.getBBox().height+10})};if(e){if("always"===h){this.collapseButtons(a);m(f())&&u();return}"never"===h&&this.expandButtons()}d&&
e?l.align===k.align||m(this.initialButtonGroupWidth+20)?"responsive"===h?(this.collapseButtons(a),m(f())&&u()):u():"responsive"===h&&this.expandButtons():e&&"responsive"===h&&(this.initialButtonGroupWidth>c.plotWidth?this.collapseButtons(a):this.expandButtons())};e.prototype.collapseButtons=function(a){var b=this.buttons,c=this.buttonOptions,e=this.chart,d=this.dropdown,f=this.options,k=this.zoomText,h=e.userOptions.rangeSelector&&e.userOptions.rangeSelector.buttonTheme||{},l=function(a){return{text:a?
a+" \u25be":"\u25be",width:"auto",paddingLeft:N(f.buttonTheme.paddingLeft,h.padding,8),paddingRight:N(f.buttonTheme.paddingRight,h.padding,8)}};k&&k.hide();var m=!1;c.forEach(function(a,c){c=b[c];2!==c.state?c.hide():(c.show(),c.attr(l(a.text)),m=!0)});m||(d&&(d.selectedIndex=0),b[0].show(),b[0].attr(l(this.zoomText&&this.zoomText.textStr)));c=f.buttonPosition.align;this.positionButtons();"right"!==c&&"center"!==c||this.alignButtonGroup(a,b[this.currentButtonIndex()].getBBox().width);this.showDropdown()};
e.prototype.expandButtons=function(){var a=this.buttons,b=this.buttonOptions,e=this.options,d=this.zoomText;this.hideDropdown();d&&d.show();b.forEach(function(b,c){c=a[c];c.show();c.attr({text:b.text,width:e.buttonTheme.width||28,paddingLeft:N(e.buttonTheme.paddingLeft,"unset"),paddingRight:N(e.buttonTheme.paddingRight,"unset")});2>c.state&&c.setState(0)});this.positionButtons()};e.prototype.currentButtonIndex=function(){var a=this.dropdown;return a&&0<a.selectedIndex?a.selectedIndex-1:0};e.prototype.showDropdown=
function(){var a=this.buttonGroup,b=this.buttons,e=this.chart,f=this.dropdown;if(a&&f){var k=a.translateX;a=a.translateY;b=b[this.currentButtonIndex()].getBBox();d(f,{left:e.plotLeft+k+"px",top:a+.5+"px",width:b.width+"px",height:b.height+"px"});this.hasVisibleDropdown=!0}};e.prototype.hideDropdown=function(){var a=this.dropdown;a&&(d(a,{top:"-9999em",width:"1px",height:"1px"}),this.hasVisibleDropdown=!1)};e.prototype.getHeight=function(){var a=this.options,b=this.group,e=a.y,d=a.buttonPosition.y,
f=a.inputPosition.y;if(a.height)return a.height;this.alignElements();a=b?b.getBBox(!0).height+13+e:0;b=Math.min(f,d);if(0>f&&0>d||0<f&&0<d)a+=Math.abs(b);return a};e.prototype.titleCollision=function(a){return!(a.options.title.text||a.options.subtitle.text)};e.prototype.update=function(a){var b=this.chart;P(!0,b.options.rangeSelector,a);this.destroy();this.init(b);this.render()};e.prototype.destroy=function(){var a=this,b=a.minInput,d=a.maxInput;a.eventsToUnbind&&(a.eventsToUnbind.forEach(function(a){return a()}),
a.eventsToUnbind=void 0);h(a.buttons);b&&(b.onfocus=b.onblur=b.onchange=null);d&&(d.onfocus=d.onblur=d.onchange=null);L(a,function(b,c){b&&"chart"!==c&&(b instanceof l?b.destroy():b instanceof window.HTMLElement&&K(b));b!==e.prototype[c]&&(a[c]=null)},this)};return e}();k.prototype.defaultButtons=[{type:"month",count:1,text:"1m",title:"View 1 month"},{type:"month",count:3,text:"3m",title:"View 3 months"},{type:"month",count:6,text:"6m",title:"View 6 months"},{type:"ytd",text:"YTD",title:"View year to date"},
{type:"year",count:1,text:"1y",title:"View 1 year"},{type:"all",text:"All",title:"View all"}];k.prototype.inputTypeFormats={"datetime-local":"%Y-%m-%dT%H:%M:%S",date:"%Y-%m-%d",time:"%H:%M:%S"};a.prototype.minFromRange=function(){var a=this.range,b=a.type,d=this.max,f=this.chart.time,g=function(a,c){var e="year"===b?"FullYear":"Month",d=new f.Date(a),g=f.get(e,d);f.set(e,d,g+c);g===f.get(e,d)&&f.set("Date",d,0);return d.getTime()-a};if(q(a)){var k=d-a;var h=a}else k=d+g(d,-a.count),this.chart&&(this.chart.fixedRange=
d-k);var l=N(this.dataMin,Number.MIN_VALUE);q(k)||(k=l);k<=l&&(k=l,"undefined"===typeof h&&(h=g(k,a.count)),this.newMax=Math.min(k+h,this.dataMax));q(d)||(k=void 0);return k};if(!y.RangeSelector){var J=[],f=function(a){function c(){e&&(d=a.xAxis[0].getExtremes(),f=a.legend,h=e&&e.options.verticalAlign,q(d.min)&&e.render(d.min,d.max),f.display&&"top"===h&&h===f.options.verticalAlign&&(k=P(a.spacingBox),k.y="vertical"===f.options.layout?a.plotTop:k.y+e.getHeight(),f.group.placed=!1,f.align(k)))}var d,
e=a.rangeSelector,f,k,h;e&&(I(J,function(b){return b[0]===a})||J.push([a,[b(a.xAxis[0],"afterSetExtremes",function(a){e&&e.render(a.min,a.max)}),b(a,"redraw",c)]]),c())};b(r,"afterGetContainer",function(){this.options.rangeSelector&&this.options.rangeSelector.enabled&&(this.rangeSelector=new k(this))});b(r,"beforeRender",function(){var a=this.axes,b=this.rangeSelector;b&&(q(b.deferredYTDClick)&&(b.clickButton(b.deferredYTDClick),delete b.deferredYTDClick),a.forEach(function(a){a.updateNames();a.setScale()}),
this.getAxisMargins(),b.render(),a=b.options.verticalAlign,b.options.floating||("bottom"===a?this.extraBottomMargin=!0:"middle"!==a&&(this.extraTopMargin=!0)))});b(r,"update",function(a){var b=a.options.rangeSelector;a=this.rangeSelector;var d=this.extraBottomMargin,e=this.extraTopMargin;b&&b.enabled&&!z(a)&&this.options.rangeSelector&&(this.options.rangeSelector.enabled=!0,this.rangeSelector=a=new k(this));this.extraTopMargin=this.extraBottomMargin=!1;a&&(f(this),b=b&&b.verticalAlign||a.options&&
a.options.verticalAlign,a.options.floating||("bottom"===b?this.extraBottomMargin=!0:"middle"!==b&&(this.extraTopMargin=!0)),this.extraBottomMargin!==d||this.extraTopMargin!==e)&&(this.isDirtyBox=!0)});b(r,"render",function(){var a=this.rangeSelector;a&&!a.options.floating&&(a.render(),a=a.options.verticalAlign,"bottom"===a?this.extraBottomMargin=!0:"middle"!==a&&(this.extraTopMargin=!0))});b(r,"getMargins",function(){var a=this.rangeSelector;a&&(a=a.getHeight(),this.extraTopMargin&&(this.plotTop+=
a),this.extraBottomMargin&&(this.marginBottom+=a))});r.prototype.callbacks.push(f);b(r,"destroy",function(){for(var a=0;a<J.length;a++){var b=J[a];if(b[0]===this){b[1].forEach(function(a){return a()});J.splice(a,1);break}}});y.RangeSelector=k}return k});L(a,"Core/Chart/StockChart.js",[a["Core/Animation/AnimationUtilities.js"],a["Core/Axis/Axis.js"],a["Core/Chart/Chart.js"],a["Core/FormatUtilities.js"],a["Core/DefaultOptions.js"],a["Core/Series/Series.js"],a["Core/Renderer/SVG/SVGRenderer.js"],a["Core/Utilities.js"]],
function(a,r,y,t,l,D,E,x){function b(a,b){return"xAxis"===a?{minPadding:0,maxPadding:0,overscroll:0,ordinal:!0,title:{text:null},labels:{overflow:"justify"},showLastLabel:!0}:"yAxis"===a?{labels:{y:-2},opposite:F(b.opposite,!0),showLastLabel:!(!b.categories&&"category"!==b.type),title:{text:null}}:{}}function m(a,b){if("xAxis"===a){a=h();var d={type:"datetime",categories:void 0};F(b.navigator&&b.navigator.enabled,a.navigator.enabled,!0)&&(d.startOnTick=!1,d.endOnTick=!1);return d}return{}}var d=this&&
this.__extends||function(){var a=function(b,d){a=Object.setPrototypeOf||{__proto__:[]}instanceof Array&&function(a,b){a.__proto__=b}||function(a,b){for(var d in b)b.hasOwnProperty(d)&&(a[d]=b[d])};return a(b,d)};return function(b,d){function k(){this.constructor=b}a(b,d);b.prototype=null===d?Object.create(d):(k.prototype=d.prototype,new k)}}(),z=t.format,h=l.getOptions;a=x.addEvent;var K=x.clamp,G=x.defined,I=x.extend,C=x.find,q=x.isNumber,L=x.isString,P=x.merge,F=x.pick,N=x.splat;x=function(a){function l(){return null!==
a&&a.apply(this,arguments)||this}d(l,a);l.prototype.init=function(d,l){var f=h(),e=d.xAxis,c=d.yAxis,k=F(d.navigator&&d.navigator.enabled,f.navigator.enabled,!0);d.xAxis=d.yAxis=void 0;k=P({chart:{panning:{enabled:!0,type:"x"},pinchType:"x"},navigator:{enabled:k},scrollbar:{enabled:F(f.scrollbar&&f.scrollbar.enabled,!0)},rangeSelector:{enabled:F(f.rangeSelector.enabled,!0)},title:{text:null},tooltip:{split:F(f.tooltip.split,!0),crosshairs:!0},legend:{enabled:!1}},d,{isStock:!0});d.xAxis=e;d.yAxis=
c;k.xAxis=N(d.xAxis||{}).map(function(a,c){return P(b("xAxis",a),f.xAxis,f.xAxis&&f.xAxis[c],a,m("xAxis",d))});k.yAxis=N(d.yAxis||{}).map(function(a,c){return P(b("yAxis",a),f.yAxis,f.yAxis&&f.yAxis[c],a)});a.prototype.init.call(this,k,l)};l.prototype.createAxis=function(d,h){h.axis=P(b(d,h.axis),h.axis,m(d,this.userOptions));return a.prototype.createAxis.call(this,d,h)};return l}(y);(function(a){a.stockChart=function(b,d,h){return new a(b,d,h)}})(x||(x={}));a(D,"setOptions",function(a){var b;this.chart.options.isStock&&
(this.is("column")||this.is("columnrange")?b={borderWidth:0,shadow:!1}:this.is("scatter")||this.is("sma")||(b={marker:{enabled:!1,radius:2}}),b&&(a.plotOptions[this.type]=P(a.plotOptions[this.type],b)))});a(r,"autoLabelAlign",function(a){var b=this.chart,d=this.options;b=b._labelPanes=b._labelPanes||{};var h=this.options.labels;this.chart.options.isStock&&"yAxis"===this.coll&&(d=d.top+","+d.height,!b[d]&&h.enabled&&(15===h.x&&(h.x=0),"undefined"===typeof h.align&&(h.align="right"),b[d]=this,a.align=
"right",a.preventDefault()))});a(r,"destroy",function(){var a=this.chart,b=this.options&&this.options.top+","+this.options.height;b&&a._labelPanes&&a._labelPanes[b]===this&&delete a._labelPanes[b]});a(r,"getPlotLinePath",function(a){function b(a){var b="xAxis"===a?"yAxis":"xAxis";a=d.options[b];return q(a)?[f[b][a]]:L(a)?[f.get(a)]:h.map(function(a){return a[b]})}var d=this,h=this.isLinked&&!this.series?this.linkedParent.series:this.series,f=d.chart,e=f.renderer,c=d.left,l=d.top,p,g,m,t,r=[],u=[],
x=a.translatedValue,z=a.value,y=a.force;if(f.options.isStock&&!1!==a.acrossPanes&&"xAxis"===d.coll||"yAxis"===d.coll){a.preventDefault();u=b(d.coll);var D=d.isXAxis?f.yAxis:f.xAxis;D.forEach(function(a){if(G(a.options.id)?-1===a.options.id.indexOf("navigator"):1){var b=a.isXAxis?"yAxis":"xAxis";b=G(a.options[b])?f[b][a.options[b]]:f[b][0];d===b&&u.push(a)}});var E=u.length?[]:[d.isXAxis?f.yAxis[0]:f.xAxis[0]];u.forEach(function(a){-1!==E.indexOf(a)||C(E,function(b){return b.pos===a.pos&&b.len===a.len})||
E.push(a)});var I=F(x,d.translate(z,null,null,a.old));q(I)&&(d.horiz?E.forEach(function(a){var b;g=a.pos;t=g+a.len;p=m=Math.round(I+d.transB);"pass"!==y&&(p<c||p>c+d.width)&&(y?p=m=K(p,c,c+d.width):b=!0);b||r.push(["M",p,g],["L",m,t])}):E.forEach(function(a){var b;p=a.pos;m=p+a.len;g=t=Math.round(l+d.height-I);"pass"!==y&&(g<l||g>l+d.height)&&(y?g=t=K(g,l,l+d.height):b=!0);b||r.push(["M",p,g],["L",m,t])}));a.path=0<r.length?e.crispPolyLine(r,a.lineWidth||1):null}});E.prototype.crispPolyLine=function(a,
b){for(var d=0;d<a.length;d+=2){var h=a[d],f=a[d+1];h[1]===f[1]&&(h[1]=f[1]=Math.round(h[1])-b%2/2);h[2]===f[2]&&(h[2]=f[2]=Math.round(h[2])+b%2/2)}return a};a(r,"afterHideCrosshair",function(){this.crossLabel&&(this.crossLabel=this.crossLabel.hide())});a(r,"afterDrawCrosshair",function(a){var b,d;if(this.crosshair&&this.crosshair.label&&this.crosshair.label.enabled&&this.cross&&q(this.min)&&q(this.max)){var h=this.chart,f=this.logarithmic,e=this.crosshair.label,c=this.horiz,l=this.opposite,p=this.left,
g=this.top,m=this.crossLabel,t=e.format,r="",u="inside"===this.options.tickPosition,x=!1!==this.crosshair.snap,y=0,A=a.e||this.cross&&this.cross.e;a=a.point;var C=this.min,D=this.max;f&&(C=f.lin2log(C),D=f.lin2log(D));f=c?"center":l?"right"===this.labelAlign?"right":"left":"left"===this.labelAlign?"left":"center";m||(m=this.crossLabel=h.renderer.label("",0,void 0,e.shape||"callout").addClass("highcharts-crosshair-label highcharts-color-"+(a?a.series.colorIndex:this.series[0]&&this.series[0].colorIndex)).attr({align:e.align||
f,padding:F(e.padding,8),r:F(e.borderRadius,3),zIndex:2}).add(this.labelGroup),h.styledMode||m.attr({fill:e.backgroundColor||a&&a.series&&a.series.color||"#666666",stroke:e.borderColor||"","stroke-width":e.borderWidth||0}).css(I({color:"#ffffff",fontWeight:"normal",fontSize:"11px",textAlign:"center"},e.style||{})));c?(f=x?(a.plotX||0)+p:A.chartX,g+=l?0:this.height):(f=l?this.width+p:0,g=x?(a.plotY||0)+g:A.chartY);t||e.formatter||(this.dateTime&&(r="%b %d, %Y"),t="{value"+(r?":"+r:"")+"}");r=x?this.isXAxis?
a.x:a.y:this.toValue(c?A.chartX:A.chartY);x=a?a.series.isPointInside(a):q(r)&&r>C&&r<D;A="";t?A=z(t,{value:r},h):e.formatter&&q(r)&&(A=e.formatter.call(this,r));m.attr({text:A,x:f,y:g,visibility:x?"visible":"hidden"});e=m.getBBox();if(q(m.y))if(c){if(u&&!l||!u&&l)g=m.y-e.height}else g=m.y-e.height/2;c?(b=p-e.x,d=p+this.width-e.x):(b="left"===this.labelAlign?p:0,d="right"===this.labelAlign?p+this.width:h.chartWidth);m.translateX<b&&(y=b-m.translateX);m.translateX+e.width>=d&&(y=-(m.translateX+e.width-
d));m.attr({x:f+y,y:g,anchorX:c?f:this.opposite?0:h.chartWidth,anchorY:c?this.opposite?h.chartHeight:0:g+e.height/2})}});D.prototype.forceCropping=function(){var a=this.chart,b=this.options.dataGrouping;return!1!==this.allowDG&&b&&F(b.enabled,a.options.isStock)};a(y,"update",function(a){a=a.options;"scrollbar"in a&&this.navigator&&(P(!0,this.options.scrollbar,a.scrollbar),this.navigator.update({},!1),delete a.scrollbar)});return x});L(a,"masters/modules/stock.src.js",[a["Core/Globals.js"],a["Core/Axis/OrdinalAxis.js"],
a["Series/DataModifyComposition.js"],a["Core/Scrollbar.js"],a["Core/Chart/StockChart.js"]],function(a,r,y,t,l){a.Scrollbar=t;a.StockChart=a.stockChart=l.stockChart;t.compose(a.Axis);r.compose(a.Axis,a.Series,a.Chart);y.compose(a.Series,a.Axis,a.Point)})});
//# sourceMappingURL=stock.js.map
```
