## Supplementary material for "Exploring natural odour landscapes: A case study with implications for human-biting insects": Figure S4: streamgraph.js

```
/*
 Highcharts JS v9.3.1 (2021-11-05)

 Streamgraph module

 (c) 2010-2021 Torstein Honsi

 License: www.highcharts.com/license
*/
'use strict';(function(a){"object"===typeof module&&module.exports?(a["default"]=a,module.exports=a):"function"===typeof define&&define.amd?define("highcharts/modules/streamgraph",["highcharts"],function(b){a(b);a.Highcharts=b;return a}):a("undefined"!==typeof Highcharts?Highcharts:void 0)})(function(a){function b(a,d,b,e){a.hasOwnProperty(d)||(a[d]=e.apply(null,b))}a=a?a._modules:{};b(a,"Series/Streamgraph/StreamgraphSeries.js",[a["Core/Series/SeriesRegistry.js"],a["Core/Utilities.js"]],function(a,
b){var d=this&&this.__extends||function(){var a=function(b,c){a=Object.setPrototypeOf||{__proto__:[]}instanceof Array&&function(a,c){a.__proto__=c}||function(a,c){for(var b in c)c.hasOwnProperty(b)&&(a[b]=c[b])};return a(b,c)};return function(b,c){function g(){this.constructor=b}a(b,c);b.prototype=null===c?Object.create(c):(g.prototype=c.prototype,new g)}}(),e=a.seriesTypes.areaspline,h=b.merge;b=b.extend;var f=function(a){function b(){var b=null!==a&&a.apply(this,arguments)||this;b.data=void 0;b.points=
void 0;b.options=void 0;return b}d(b,a);b.prototype.streamStacker=function(a,b,d){a[0]-=b.total/2;a[1]-=b.total/2;this.stackedYData[d]=a};b.defaultOptions=h(e.defaultOptions,{fillOpacity:1,lineWidth:0,marker:{enabled:!1},stacking:"stream"});return b}(e);b(f.prototype,{negStacks:!1});a.registerSeriesType("streamgraph",f);"";return f});b(a,"masters/modules/streamgraph.src.js",[],function(){})});
//# sourceMappingURL=streamgraph.js.map
```
