## Supplementary material for "Exploring natural odour landscapes: A case study with implications for human-biting insects": Figure S4: sunburst.js

```
/*
 Highcharts JS v9.3.1 (2021-11-05)

 (c) 2016-2021 Highsoft AS
 Authors: Jon Arild Nygard

 License: www.highcharts.com/license
*/
'use strict';(function(a){"object"===typeof module&&module.exports?(a["default"]=a,module.exports=a):"function"===typeof define&&define.amd?define("highcharts/modules/sunburst",["highcharts"],function(u){a(u);a.Highcharts=u;return a}):a("undefined"!==typeof Highcharts?Highcharts:void 0)})(function(a){function u(a,h,g,r){a.hasOwnProperty(h)||(a[h]=r.apply(null,g))}a=a?a._modules:{};u(a,"Series/ColorMapComposition.js",[a["Core/Utilities.js"]],function(a){var h=a.defined,g=a.wrap,r={pointArrayMap:["value"],
axisTypes:["xAxis","yAxis","colorAxis"],trackerGroups:["group","markerGroup","dataLabelsGroup"],parallelArrays:["x","y","value"],colorKey:"value"},q;(function(a){function k(){return null!==this.value&&Infinity!==this.value&&-Infinity!==this.value}function n(b){var d={};!h(b.color)||b.state&&"normal"!==b.state||(d[this.colorProp||"fill"]=b.color);return d}function p(b,t,a){b=b.call(this,t,a);t.moveToTopOnHover&&(b.zIndex="hover"===a?1:0);return b}a.colorMapSeriesMixin=r;var b=[];a.compose=function(d,
t){t&&-1===b.indexOf(t)&&(b.push(t),t=t.prototype,t.dataLabelOnNull=!0,t.moveToTopOnHover=!0,t.isValid=k);-1===b.indexOf(d)&&(b.push(d),t=d.prototype,t.colorAttribs=n,g(t,"pointAttribs",p));return d};a.seriesColorAttribs=n})(q||(q={}));return q});u(a,"Series/Treemap/TreemapAlgorithmGroup.js",[],function(){return function(){function a(a,g,r,q){this.height=a;this.width=g;this.plot=q;this.startDirection=this.direction=r;this.lH=this.nH=this.lW=this.nW=this.total=0;this.elArr=[];this.lP={total:0,lH:0,
nH:0,lW:0,nW:0,nR:0,lR:0,aspectRatio:function(a,c){return Math.max(a/c,c/a)}}}a.prototype.addElement=function(a){this.lP.total=this.elArr[this.elArr.length-1];this.total+=a;0===this.direction?(this.lW=this.nW,this.lP.lH=this.lP.total/this.lW,this.lP.lR=this.lP.aspectRatio(this.lW,this.lP.lH),this.nW=this.total/this.height,this.lP.nH=this.lP.total/this.nW,this.lP.nR=this.lP.aspectRatio(this.nW,this.lP.nH)):(this.lH=this.nH,this.lP.lW=this.lP.total/this.lH,this.lP.lR=this.lP.aspectRatio(this.lP.lW,
this.lH),this.nH=this.total/this.width,this.lP.nW=this.lP.total/this.nH,this.lP.nR=this.lP.aspectRatio(this.lP.nW,this.nH));this.elArr.push(a)};a.prototype.reset=function(){this.lW=this.nW=0;this.elArr=[];this.total=0};return a}()});u(a,"Series/DrawPointComposition.js",[],function(){var a;(function(a){function g(a){var c=this,n=a.animatableAttribs,p=a.onComplete,b=a.css,d=a.renderer,t=this.series&&this.series.chart.hasRendered?void 0:this.series&&this.series.options.animation,x=this.graphic;a.attribs=
a.attribs||{};a.attribs["class"]=this.getClassName();if(this.shouldDraw())x||(this.graphic=x=d[a.shapeType](a.shapeArgs).add(a.group)),x.css(b).attr(a.attribs).animate(n,a.isNew?!1:t,p);else if(x){var B=function(){c.graphic=x=x&&x.destroy();"function"===typeof p&&p()};Object.keys(n).length?x.animate(n,void 0,function(){B()}):B()}}function h(){return!this.isNull}var q=[];a.compose=function(a){if(-1===q.indexOf(a)){q.push(a);var c=a.prototype;c.draw=g;c.shouldDraw||(c.shouldDraw=h)}return a}})(a||(a=
{}));return a});u(a,"Series/Treemap/TreemapPoint.js",[a["Series/DrawPointComposition.js"],a["Core/Series/SeriesRegistry.js"],a["Core/Utilities.js"]],function(a,h,g){var r=this&&this.__extends||function(){var b=function(a,t){b=Object.setPrototypeOf||{__proto__:[]}instanceof Array&&function(b,a){b.__proto__=a}||function(b,a){for(var d in a)a.hasOwnProperty(d)&&(b[d]=a[d])};return b(a,t)};return function(a,t){function d(){this.constructor=a}b(a,t);a.prototype=null===t?Object.create(t):(d.prototype=t.prototype,
new d)}}(),q=h.series.prototype.pointClass,k=h.seriesTypes;h=k.pie.prototype.pointClass;var c=g.extend,n=g.isNumber,p=g.pick;g=function(b){function a(){var a=null!==b&&b.apply(this,arguments)||this;a.name=void 0;a.node=void 0;a.options=void 0;a.series=void 0;a.value=void 0;return a}r(a,b);a.prototype.getClassName=function(){var a=q.prototype.getClassName.call(this),b=this.series,d=b.options;this.node.level<=b.nodeMap[b.rootNode].level?a+=" highcharts-above-level":this.node.isLeaf||p(d.interactByLeaf,
!d.allowTraversingTree)?this.node.isLeaf||(a+=" highcharts-internal-node"):a+=" highcharts-internal-node-interactive";return a};a.prototype.isValid=function(){return!(!this.id&&!n(this.value))};a.prototype.setState=function(a){q.prototype.setState.call(this,a);this.graphic&&this.graphic.attr({zIndex:"hover"===a?1:0})};a.prototype.shouldDraw=function(){return n(this.plotY)&&null!==this.y};return a}(k.scatter.prototype.pointClass);c(g.prototype,{setVisible:h.prototype.setVisible});a.compose(g);return g});
u(a,"Series/Treemap/TreemapUtilities.js",[a["Core/Utilities.js"]],function(a){var h=a.objectEach,g;(function(a){function g(a,c,n){void 0===n&&(n=this);a=c.call(n,a);!1!==a&&g(a,c,n)}a.AXIS_MAX=100;a.isBoolean=function(a){return"boolean"===typeof a};a.eachObject=function(a,c,n){n=n||this;h(a,function(p,b){c.call(n,p,b,a)})};a.recursive=g})(g||(g={}));return g});u(a,"Series/TreeUtilities.js",[a["Core/Color/Color.js"],a["Core/Utilities.js"]],function(a,h){function g(a,d){var b=d.before,n=d.idRoot,c=
d.mapIdToNode[n],h=d.points[a.i],q=h&&h.options||{},k=[],w=0;a.levelDynamic=a.level-(!1!==d.levelIsConstant?0:c.level);a.name=p(h&&h.name,"");a.visible=n===a.id||!0===d.visible;"function"===typeof b&&(a=b(a,d));a.children.forEach(function(b,n){var c=r({},d);r(c,{index:n,siblings:a.children.length,visible:a.visible});b=g(b,c);k.push(b);b.visible&&(w+=b.val)});b=p(q.value,w);a.visible=0<=b&&(0<w||a.visible);a.children=k;a.childrenTotal=w;a.isLeaf=a.visible&&!w;a.val=b;return a}var r=h.extend,q=h.isArray,
k=h.isNumber,c=h.isObject,n=h.merge,p=h.pick;return{getColor:function(b,d){var n=d.index,c=d.mapOptionsToLevel,h=d.parentColor,g=d.parentColorIndex,q=d.series,k=d.colors,w=d.siblings,r=q.points,u=q.chart.options.chart,z;if(b){r=r[b.i];b=c[b.level]||{};if(c=r&&b.colorByPoint){var D=r.index%(k?k.length:u.colorCount);var F=k&&k[D]}if(!q.chart.styledMode){k=r&&r.options.color;u=b&&b.color;if(z=h)z=(z=b&&b.colorVariation)&&"brightness"===z.key&&n&&w?a.parse(h).brighten(n/w*z.to).get():h;z=p(k,u,F,z,q.color)}var G=
p(r&&r.options.colorIndex,b&&b.colorIndex,D,g,d.colorIndex)}return{color:z,colorIndex:G}},getLevelOptions:function(a){var b=null;if(c(a)){b={};var h=k(a.from)?a.from:1;var g=a.levels;var r={};var z=c(a.defaults)?a.defaults:{};q(g)&&(r=g.reduce(function(a,b){if(c(b)&&k(b.level)){var d=n({},b);var g=p(d.levelIsConstant,z.levelIsConstant);delete d.levelIsConstant;delete d.level;b=b.level+(g?0:h-1);c(a[b])?n(!0,a[b],d):a[b]=d}return a},{}));g=k(a.to)?a.to:1;for(a=0;a<=g;a++)b[a]=n({},z,c(r[a])?r[a]:{})}return b},
setTreeValues:g,updateRootId:function(a){if(c(a)){var b=c(a.options)?a.options:{};b=p(a.rootNode,b.rootId,"");c(a.userOptions)&&(a.userOptions.rootId=b);a.rootNode=b}return b}}});u(a,"Series/Treemap/TreemapComposition.js",[a["Core/Series/SeriesRegistry.js"],a["Series/Treemap/TreemapUtilities.js"],a["Core/Utilities.js"]],function(a,h,g){var r=g.addEvent,q=g.extend,k=!1;r(a.series,"afterBindAxes",function(){var a=this.xAxis,n=this.yAxis;if(a&&n)if(this.is("treemap")){var g={endOnTick:!1,gridLineWidth:0,
lineWidth:0,min:0,minPadding:0,max:h.AXIS_MAX,maxPadding:0,startOnTick:!1,title:void 0,tickPositions:[]};q(n.options,g);q(a.options,g);k=!0}else k&&(n.setOptions(n.userOptions),a.setOptions(a.userOptions),k=!1)})});u(a,"Series/Treemap/TreemapSeries.js",[a["Core/Color/Color.js"],a["Series/ColorMapComposition.js"],a["Core/Globals.js"],a["Core/Legend/LegendSymbol.js"],a["Core/Series/SeriesRegistry.js"],a["Series/Treemap/TreemapAlgorithmGroup.js"],a["Series/Treemap/TreemapPoint.js"],a["Series/Treemap/TreemapUtilities.js"],
a["Series/TreeUtilities.js"],a["Core/Utilities.js"]],function(a,h,g,r,q,k,c,n,p,b){var d=this&&this.__extends||function(){var a=function(b,e){a=Object.setPrototypeOf||{__proto__:[]}instanceof Array&&function(e,a){e.__proto__=a}||function(e,a){for(var m in a)a.hasOwnProperty(m)&&(e[m]=a[m])};return a(b,e)};return function(b,e){function m(){this.constructor=b}a(b,e);b.prototype=null===e?Object.create(e):(m.prototype=e.prototype,new m)}}(),t=a.parse,x=h.colorMapSeriesMixin;a=g.noop;var B=q.series;g=
q.seriesTypes;var z=g.column,u=g.heatmap,L=g.scatter,w=p.getColor,N=p.getLevelOptions,K=p.updateRootId,I=b.addEvent,D=b.correctFloat,F=b.defined,G=b.error,J=b.extend,O=b.fireEvent,P=b.isArray,T=b.isObject,f=b.isString,C=b.merge,v=b.pick,U=b.stableSort;p=function(a){function b(){var e=null!==a&&a.apply(this,arguments)||this;e.axisRatio=void 0;e.data=void 0;e.mapOptionsToLevel=void 0;e.nodeMap=void 0;e.options=void 0;e.points=void 0;e.rootNode=void 0;e.tree=void 0;return e}d(b,a);b.prototype.algorithmCalcPoints=
function(a,b,l,f){var e,m,A,d,n=l.lW,g=l.lH,c=l.plot,h=0,t=l.elArr.length-1;if(b)n=l.nW,g=l.nH;else var H=l.elArr[l.elArr.length-1];l.elArr.forEach(function(a){if(b||h<t)0===l.direction?(e=c.x,m=c.y,A=n,d=a/A):(e=c.x,m=c.y,d=g,A=a/d),f.push({x:e,y:m,width:A,height:D(d)}),0===l.direction?c.y+=d:c.x+=A;h+=1});l.reset();0===l.direction?l.width-=n:l.height-=g;c.y=c.parent.y+(c.parent.height-l.height);c.x=c.parent.x+(c.parent.width-l.width);a&&(l.direction=1-l.direction);b||l.addElement(H)};b.prototype.algorithmFill=
function(a,b,l){var e=[],m,f=b.direction,d=b.x,c=b.y,n=b.width,g=b.height,h,t,p,H;l.forEach(function(l){m=l.val/b.val*b.height*b.width;h=d;t=c;0===f?(H=g,p=m/H,n-=p,d+=p):(p=n,H=m/p,g-=H,c+=H);e.push({x:h,y:t,width:p,height:H});a&&(f=1-f)});return e};b.prototype.algorithmLowAspectRatio=function(a,b,l){var e=[],m=this,f,d={x:b.x,y:b.y,parent:b},c=0,n=l.length-1,g=new k(b.height,b.width,b.direction,d);l.forEach(function(l){f=l.val/b.val*b.height*b.width;g.addElement(f);g.lP.nR>g.lP.lR&&m.algorithmCalcPoints(a,
!1,g,e,d);c===n&&m.algorithmCalcPoints(a,!0,g,e,d);c+=1});return e};b.prototype.alignDataLabel=function(a,b,l){var e=l.style;e&&!F(e.textOverflow)&&b.text&&b.getBBox().width>b.text.textWidth&&b.css({textOverflow:"ellipsis",width:e.width+="px"});z.prototype.alignDataLabel.apply(this,arguments);a.dataLabel&&a.dataLabel.attr({zIndex:(a.node.zIndex||0)+1})};b.prototype.buildNode=function(a,b,l,f,d){var e=this,m=[],E=e.points[b],c=0,A;(f[a]||[]).forEach(function(b){A=e.buildNode(e.points[b].id,b,l+1,f,
a);c=Math.max(A.height+1,c);m.push(A)});b={id:a,i:b,children:m,height:c,level:l,parent:d,visible:!1};e.nodeMap[b.id]=b;E&&(E.node=b);return b};b.prototype.calculateChildrenAreas=function(a,b){var e=this,m=e.options,f=e.mapOptionsToLevel[a.level+1],d=v(e[f&&f.layoutAlgorithm]&&f.layoutAlgorithm,m.layoutAlgorithm),c=m.alternateStartingDirection,g=[];a=a.children.filter(function(a){return!a.ignore});f&&f.layoutStartingDirection&&(b.direction="vertical"===f.layoutStartingDirection?0:1);g=e[d](b,a);a.forEach(function(a,
m){m=g[m];a.values=C(m,{val:a.childrenTotal,direction:c?1-b.direction:b.direction});a.pointValues=C(m,{x:m.x/e.axisRatio,y:n.AXIS_MAX-m.y-m.height,width:m.width/e.axisRatio});a.children.length&&e.calculateChildrenAreas(a,a.values)})};b.prototype.drawDataLabels=function(){var a=this,b=a.mapOptionsToLevel,f,d;a.points.filter(function(a){return a.node.visible}).forEach(function(e){d=b[e.node.level];f={style:{}};e.node.isLeaf||(f.enabled=!1);d&&d.dataLabels&&(f=C(f,d.dataLabels),a._hasPointLabels=!0);
e.shapeArgs&&(f.style.width=e.shapeArgs.width,e.dataLabel&&e.dataLabel.css({width:e.shapeArgs.width+"px"}));e.dlOptions=C(f,e.options.dataLabels)});B.prototype.drawDataLabels.call(this)};b.prototype.drawPoints=function(){var a=this,b=a.chart,f=b.renderer,d=b.styledMode,c=a.options,g=d?{}:c.shadow,n=c.borderRadius,h=b.pointCount<c.animationLimit,t=c.allowTraversingTree;a.points.forEach(function(e){var b=e.node.levelDynamic,m={},l={},A={},E="level-group-"+e.node.level,p=!!e.graphic,V=h&&p,Q=e.shapeArgs;
e.shouldDraw()&&(e.isInside=!0,n&&(l.r=n),C(!0,V?m:l,p?Q:{},d?{}:a.pointAttribs(e,e.selected?"select":void 0)),a.colorAttribs&&d&&J(A,a.colorAttribs(e)),a[E]||(a[E]=f.g(E).attr({zIndex:1E3-(b||0)}).add(a.group),a[E].survive=!0));e.draw({animatableAttribs:m,attribs:l,css:A,group:a[E],renderer:f,shadow:g,shapeArgs:Q,shapeType:"rect"});t&&e.graphic&&(e.drillId=c.interactByLeaf?a.drillToByLeaf(e):a.drillToByGroup(e))})};b.prototype.drillToByGroup=function(a){var e=!1;1!==a.node.level-this.nodeMap[this.rootNode].level||
a.node.isLeaf||(e=a.id);return e};b.prototype.drillToByLeaf=function(a){var e=!1;if(a.node.parent!==this.rootNode&&a.node.isLeaf)for(a=a.node;!e;)a=this.nodeMap[a.parent],a.parent===this.rootNode&&(e=a.id);return e};b.prototype.drillToNode=function(a,b){G(32,!1,void 0,{"treemap.drillToNode":"use treemap.setRootNode"});this.setRootNode(a,b)};b.prototype.drillUp=function(){var a=this.nodeMap[this.rootNode];a&&f(a.parent)&&this.setRootNode(a.parent,!0,{trigger:"traverseUpButton"})};b.prototype.getExtremes=
function(){var a=B.prototype.getExtremes.call(this,this.colorValueData),b=a.dataMax;this.valueMin=a.dataMin;this.valueMax=b;return B.prototype.getExtremes.call(this)};b.prototype.getListOfParents=function(a,b){a=P(a)?a:[];var e=P(b)?b:[];b=a.reduce(function(a,e,b){e=v(e.parent,"");"undefined"===typeof a[e]&&(a[e]=[]);a[e].push(b);return a},{"":[]});n.eachObject(b,function(a,b,f){""!==b&&-1===e.indexOf(b)&&(a.forEach(function(a){f[""].push(a)}),delete f[b])});return b};b.prototype.getTree=function(){var a=
this.data.map(function(a){return a.id});a=this.getListOfParents(this.data,a);this.nodeMap={};return this.buildNode("",-1,0,a)};b.prototype.hasData=function(){return!!this.processedXData.length};b.prototype.init=function(a,b){x&&(this.colorAttribs=h.seriesColorAttribs);var e=I(this,"setOptions",function(a){a=a.userOptions;F(a.allowDrillToNode)&&!F(a.allowTraversingTree)&&(a.allowTraversingTree=a.allowDrillToNode,delete a.allowDrillToNode);F(a.drillUpButton)&&!F(a.traverseUpButton)&&(a.traverseUpButton=
a.drillUpButton,delete a.drillUpButton)});B.prototype.init.call(this,a,b);delete this.opacity;this.eventsToUnbind.push(e);this.options.allowTraversingTree&&this.eventsToUnbind.push(I(this,"click",this.onClickDrillToNode))};b.prototype.onClickDrillToNode=function(a){var b=(a=a.point)&&a.drillId;f(b)&&(a.setState(""),this.setRootNode(b,!0,{trigger:"click"}))};b.prototype.pointAttribs=function(a,b){var e=T(this.mapOptionsToLevel)?this.mapOptionsToLevel:{},f=a&&e[a.node.level]||{};e=this.options;var d=
b&&e.states[b]||{},m=a&&a.getClassName()||"";a={stroke:a&&a.borderColor||f.borderColor||d.borderColor||e.borderColor,"stroke-width":v(a&&a.borderWidth,f.borderWidth,d.borderWidth,e.borderWidth),dashstyle:a&&a.borderDashStyle||f.borderDashStyle||d.borderDashStyle||e.borderDashStyle,fill:a&&a.color||this.color};-1!==m.indexOf("highcharts-above-level")?(a.fill="none",a["stroke-width"]=0):-1!==m.indexOf("highcharts-internal-node-interactive")?(b=v(d.opacity,e.opacity),a.fill=t(a.fill).setOpacity(b).get(),
a.cursor="pointer"):-1!==m.indexOf("highcharts-internal-node")?a.fill="none":b&&(a.fill=t(a.fill).brighten(d.brightness).get());return a};b.prototype.renderTraverseUpButton=function(a){var b=this,e=b.options.traverseUpButton,f=v(e.text,b.nodeMap[a].name,"\u25c1 Back");if(""===a||b.is("sunburst")&&1===b.tree.children.length&&a===b.tree.children[0].id)b.drillUpButton&&(b.drillUpButton=b.drillUpButton.destroy());else if(this.drillUpButton)this.drillUpButton.placed=!1,this.drillUpButton.attr({text:f}).align();
else{var d=(a=e.theme)&&a.states;this.drillUpButton=this.chart.renderer.button(f,0,0,function(){b.drillUp()},a,d&&d.hover,d&&d.select).addClass("highcharts-drillup-button").attr({align:e.position.align,zIndex:7}).add().align(e.position,!1,e.relativeTo||"plotBox")}};b.prototype.setColorRecursive=function(a,b,f,d,c){var e=this,m=e&&e.chart;m=m&&m.options&&m.options.colors;if(a){var l=w(a,{colors:m,index:d,mapOptionsToLevel:e.mapOptionsToLevel,parentColor:b,parentColorIndex:f,series:e,siblings:c});if(b=
e.points[a.i])b.color=l.color,b.colorIndex=l.colorIndex;(a.children||[]).forEach(function(b,f){e.setColorRecursive(b,l.color,l.colorIndex,f,a.children.length)})}};b.prototype.setPointValues=function(){var a=this,b=a.xAxis,f=a.yAxis,d=a.chart.styledMode;a.points.forEach(function(e){var m=e.node,c=m.pointValues;m=m.visible;if(c&&m){m=c.height;var l=c.width,g=c.x,n=c.y,h=d?0:(a.pointAttribs(e)["stroke-width"]||0)%2/2;c=Math.round(b.toPixels(g,!0))-h;l=Math.round(b.toPixels(g+l,!0))-h;g=Math.round(f.toPixels(n,
!0))-h;m=Math.round(f.toPixels(n+m,!0))-h;m={x:Math.min(c,l),y:Math.min(g,m),width:Math.abs(l-c),height:Math.abs(m-g)};e.plotX=m.x+m.width/2;e.plotY=m.y+m.height/2;e.shapeArgs=m}else delete e.plotX,delete e.plotY})};b.prototype.setRootNode=function(a,b,f){a=J({newRootId:a,previousRootId:this.rootNode,redraw:v(b,!0),series:this},f);O(this,"setRootNode",a,function(a){var b=a.series;b.idPreviousRoot=a.previousRootId;b.rootNode=a.newRootId;b.isDirty=!0;a.redraw&&b.chart.redraw()})};b.prototype.setState=
function(a){this.options.inactiveOtherPoints=!0;B.prototype.setState.call(this,a,!1);this.options.inactiveOtherPoints=!1};b.prototype.setTreeValues=function(a){var b=this,e=b.options,f=b.nodeMap[b.rootNode];e=n.isBoolean(e.levelIsConstant)?e.levelIsConstant:!0;var d=0,c=[],g=b.points[a.i];a.children.forEach(function(a){a=b.setTreeValues(a);c.push(a);a.ignore||(d+=a.val)});U(c,function(a,b){return(a.sortIndex||0)-(b.sortIndex||0)});var h=v(g&&g.options.value,d);g&&(g.value=h);J(a,{children:c,childrenTotal:d,
ignore:!(v(g&&g.visible,!0)&&0<h),isLeaf:a.visible&&!d,levelDynamic:a.level-(e?0:f.level),name:v(g&&g.name,""),sortIndex:v(g&&g.sortIndex,-h),val:h});return a};b.prototype.sliceAndDice=function(a,b){return this.algorithmFill(!0,a,b)};b.prototype.squarified=function(a,b){return this.algorithmLowAspectRatio(!0,a,b)};b.prototype.strip=function(a,b){return this.algorithmLowAspectRatio(!1,a,b)};b.prototype.stripes=function(a,b){return this.algorithmFill(!1,a,b)};b.prototype.translate=function(){var a=
this,b=a.options,f=K(a);B.prototype.translate.call(a);var d=a.tree=a.getTree();var c=a.nodeMap[f];""===f||c&&c.children.length||(a.setRootNode("",!1),f=a.rootNode,c=a.nodeMap[f]);a.renderTraverseUpButton(f);a.mapOptionsToLevel=N({from:c.level+1,levels:b.levels,to:d.height,defaults:{levelIsConstant:a.options.levelIsConstant,colorByPoint:b.colorByPoint}});n.recursive(a.nodeMap[a.rootNode],function(b){var e=!1,f=b.parent;b.visible=!0;if(f||""===f)e=a.nodeMap[f];return e});n.recursive(a.nodeMap[a.rootNode].children,
function(a){var b=!1;a.forEach(function(a){a.visible=!0;a.children.length&&(b=(b||[]).concat(a.children))});return b});a.setTreeValues(d);a.axisRatio=a.xAxis.len/a.yAxis.len;a.nodeMap[""].pointValues=f={x:0,y:0,width:n.AXIS_MAX,height:n.AXIS_MAX};a.nodeMap[""].values=f=C(f,{width:f.width*a.axisRatio,direction:"vertical"===b.layoutStartingDirection?0:1,val:d.val});a.calculateChildrenAreas(d,f);a.colorAxis||b.colorByPoint||a.setColorRecursive(a.tree);b.allowTraversingTree&&(b=c.pointValues,a.xAxis.setExtremes(b.x,
b.x+b.width,!1),a.yAxis.setExtremes(b.y,b.y+b.height,!1),a.xAxis.setScale(),a.yAxis.setScale());a.setPointValues()};b.defaultOptions=C(L.defaultOptions,{allowTraversingTree:!1,animationLimit:250,borderRadius:0,showInLegend:!1,marker:void 0,colorByPoint:!1,dataLabels:{defer:!1,enabled:!0,formatter:function(){var a=this&&this.point?this.point:{};return f(a.name)?a.name:""},inside:!0,verticalAlign:"middle"},tooltip:{headerFormat:"",pointFormat:"<b>{point.name}</b>: {point.value}<br/>"},ignoreHiddenPoint:!0,
layoutAlgorithm:"sliceAndDice",layoutStartingDirection:"vertical",alternateStartingDirection:!1,levelIsConstant:!0,drillUpButton:{position:{align:"right",x:-10,y:10}},traverseUpButton:{position:{align:"right",x:-10,y:10}},borderColor:"#e6e6e6",borderWidth:1,colorKey:"colorValue",opacity:.15,states:{hover:{borderColor:"#999999",brightness:u?0:.1,halo:!1,opacity:.75,shadow:!1}}});return b}(L);J(p.prototype,{buildKDTree:a,colorKey:"colorValue",directTouch:!0,drawLegendSymbol:r.drawRectangle,getExtremesFromAll:!0,
getSymbol:a,optionalAxis:"colorAxis",parallelArrays:["x","y","value","colorValue"],pointArrayMap:["value"],pointClass:c,trackerGroups:["group","dataLabelsGroup"],utils:{recursive:n.recursive}});h.compose(p);q.registerSeriesType("treemap",p);"";return p});u(a,"Series/Sunburst/SunburstPoint.js",[a["Series/DrawPointComposition.js"],a["Core/Series/SeriesRegistry.js"],a["Core/Utilities.js"]],function(a,h,g){var r=this&&this.__extends||function(){var a=function(c,g){a=Object.setPrototypeOf||{__proto__:[]}instanceof
Array&&function(a,d){a.__proto__=d}||function(a,d){for(var b in d)d.hasOwnProperty(b)&&(a[b]=d[b])};return a(c,g)};return function(c,g){function b(){this.constructor=c}a(c,g);c.prototype=null===g?Object.create(g):(b.prototype=g.prototype,new b)}}(),q=h.series.prototype.pointClass,k=g.correctFloat;g=g.extend;h=function(a){function c(){var c=null!==a&&a.apply(this,arguments)||this;c.node=void 0;c.options=void 0;c.series=void 0;c.shapeExisting=void 0;return c}r(c,a);c.prototype.getDataLabelPath=function(a){var b=
this.series.chart.renderer,d=this.shapeExisting,c=d.start,g=d.end,h=c+(g-c)/2;h=0>h&&h>-Math.PI||h>Math.PI;var n=d.r+(a.options.distance||0);c===-Math.PI/2&&k(g)===k(1.5*Math.PI)&&(c=-Math.PI+Math.PI/360,g=-Math.PI/360,h=!0);if(g-c>Math.PI){h=!1;var p=!0}this.dataLabelPath&&(this.dataLabelPath=this.dataLabelPath.destroy());this.dataLabelPath=b.arc({open:!0,longArc:p?1:0}).add(a);this.dataLabelPath.attr({start:h?c:g,end:h?g:c,clockwise:+h,x:d.x,y:d.y,r:(n+d.innerR)/2});return this.dataLabelPath};c.prototype.isValid=
function(){return!0};return c}(h.seriesTypes.treemap.prototype.pointClass);g(h.prototype,{getClassName:q.prototype.getClassName,haloPath:q.prototype.haloPath,setState:q.prototype.setState});a.compose(h);return h});u(a,"Series/Sunburst/SunburstUtilities.js",[a["Core/Series/SeriesRegistry.js"],a["Core/Utilities.js"]],function(a,h){var g=a.seriesTypes.treemap,r=h.isNumber,q=h.isObject,k=h.merge,c;(function(a){function c(a,d){var b=[];if(r(a)&&r(d)&&a<=d)for(;a<=d;a++)b.push(a);return b}a.recursive=g.prototype.utils.recursive;
a.calculateLevelSizes=function(a,d){d=q(d)?d:{};var b=0,g;if(q(a)){var h=k({},a);a=r(d.from)?d.from:0;var n=r(d.to)?d.to:0;var p=c(a,n);a=Object.keys(h).filter(function(a){return-1===p.indexOf(+a)});var u=g=r(d.diffRadius)?d.diffRadius:0;p.forEach(function(a){a=h[a];var d=a.levelSize.unit,c=a.levelSize.value;"weight"===d?b+=c:"percentage"===d?(a.levelSize={unit:"pixels",value:c/100*u},g-=a.levelSize.value):"pixels"===d&&(g-=c)});p.forEach(function(a){var d=h[a];"weight"===d.levelSize.unit&&(d=d.levelSize.value,
h[a].levelSize={unit:"pixels",value:d/b*g})});a.forEach(function(a){h[a].levelSize={value:0,unit:"pixels"}})}return h};a.getLevelFromAndTo=function(a){var b=a.level;return{from:0<b?b:1,to:b+a.height}};a.range=c})(c||(c={}));return c});u(a,"Series/Sunburst/SunburstSeries.js",[a["Series/CenteredUtilities.js"],a["Core/Globals.js"],a["Core/Series/SeriesRegistry.js"],a["Series/Sunburst/SunburstPoint.js"],a["Series/Sunburst/SunburstUtilities.js"],a["Series/TreeUtilities.js"],a["Core/Utilities.js"]],function(a,
h,g,r,q,k,c){function n(a,b){var f=b.mapIdToNode[a.parent],d=b.series,c=d.chart,g=d.points[a.i];f=R(a,{colors:d.options.colors||c&&c.options.colors,colorIndex:d.colorIndex,index:b.index,mapOptionsToLevel:b.mapOptionsToLevel,parentColor:f&&f.color,parentColorIndex:f&&f.colorIndex,series:b.series,siblings:b.siblings});a.color=f.color;a.colorIndex=f.colorIndex;g&&(g.color=a.color,g.colorIndex=a.colorIndex,a.sliced=a.id!==b.idRoot?g.sliced:!1);return a}var p=this&&this.__extends||function(){var a=function(b,
f){a=Object.setPrototypeOf||{__proto__:[]}instanceof Array&&function(a,b){a.__proto__=b}||function(a,b){for(var f in b)b.hasOwnProperty(f)&&(a[f]=b[f])};return a(b,f)};return function(b,f){function d(){this.constructor=b}a(b,f);b.prototype=null===f?Object.create(f):(d.prototype=f.prototype,new d)}}(),b=a.getCenter,d=a.getStartAndEndRadians;a=h.noop;var t=g.series,u=g.seriesTypes;h=u.column;var z=u.treemap,R=k.getColor,S=k.getLevelOptions,L=k.setTreeValues,w=k.updateRootId,N=c.error,K=c.extend,I=c.isNumber,
D=c.isObject,F=c.isString,G=c.merge,J=c.splat,O=180/Math.PI;k=function(a){function c(){var b=null!==a&&a.apply(this,arguments)||this;b.center=void 0;b.data=void 0;b.mapOptionsToLevel=void 0;b.nodeMap=void 0;b.options=void 0;b.points=void 0;b.shapeRoot=void 0;b.startAndEndRadians=void 0;b.tree=void 0;return b}p(c,a);c.prototype.alignDataLabel=function(b,c,d){if(!d.textPath||!d.textPath.enabled)return a.prototype.alignDataLabel.apply(this,arguments)};c.prototype.animate=function(a){var b=this.chart,
c=[b.plotWidth/2,b.plotHeight/2],d=b.plotLeft,f=b.plotTop;b=this.group;a?(a={translateX:c[0]+d,translateY:c[1]+f,scaleX:.001,scaleY:.001,rotation:10,opacity:.01},b.attr(a)):(a={translateX:d,translateY:f,scaleX:1,scaleY:1,rotation:0,opacity:1},b.animate(a,this.options.animation))};c.prototype.drawPoints=function(){var a=this,b=a.mapOptionsToLevel,c=a.shapeRoot,d=a.group,g=a.hasRendered,h=a.rootNode,e=a.idPreviousRoot,m=a.nodeMap,l=m[e],n=l&&l.shapeArgs;l=a.points;var p=a.startAndEndRadians,q=a.chart,
k=q&&q.options&&q.options.chart||{},r="boolean"===typeof k.animation?k.animation:!0,u=a.center[3]/2,z=a.chart.renderer,w=!1,x=!1;if(k=!!(r&&g&&h!==e&&a.dataLabelsGroup)){a.dataLabelsGroup.attr({opacity:0});var B=function(){w=!0;a.dataLabelsGroup&&a.dataLabelsGroup.animate({opacity:1,visibility:"visible"})}}l.forEach(function(f){var l=f.node,k=b[l.level];var t=f.shapeExisting||{};var v=l.shapeArgs||{},A=!(!l.visible||!l.shapeArgs);if(g&&r){var C={};var M={end:v.end,start:v.start,innerR:v.innerR,r:v.r,
x:v.x,y:v.y};A?!f.graphic&&n&&(C=h===f.id?{start:p.start,end:p.end}:n.end<=v.start?{start:p.end,end:p.end}:{start:p.start,end:p.start},C.innerR=C.r=u):f.graphic&&(e===f.id?M={innerR:u,r:u}:c&&(M=c.end<=t.start?{innerR:u,r:u,start:p.end,end:p.end}:{innerR:u,r:u,start:p.start,end:p.start}));t=C}else M=v,t={};C=[v.plotX,v.plotY];if(!f.node.isLeaf)if(h===f.id){var y=m[h];y=y.parent}else y=f.id;K(f,{shapeExisting:v,tooltipPos:C,drillId:y,name:""+(f.name||f.id||f.index),plotX:v.plotX,plotY:v.plotY,value:l.val,
isInside:A,isNull:!A});y=f.options;l=D(v)?v:{};y=D(y)?y.dataLabels:{};k=J(D(k)?k.dataLabels:{})[0];k=G({style:{}},k,y);y=k.rotationMode;if(!I(k.rotation)){if("auto"===y||"circular"===y)if(1>f.innerArcLength&&f.outerArcLength>l.radius){var w=0;f.dataLabelPath&&"circular"===y&&(k.textPath={enabled:!0})}else 1<f.innerArcLength&&f.outerArcLength>1.5*l.radius?"circular"===y?k.textPath={enabled:!0,attributes:{dy:5}}:y="parallel":(f.dataLabel&&f.dataLabel.textPathWrapper&&"circular"===y&&(k.textPath={enabled:!1}),
y="perpendicular");"auto"!==y&&"circular"!==y&&(w=l.end-(l.end-l.start)/2);k.style.width="parallel"===y?Math.min(2.5*l.radius,(f.outerArcLength+f.innerArcLength)/2):l.radius;"perpendicular"===y&&f.series.chart.renderer.fontMetrics(k.style.fontSize).h>f.outerArcLength&&(k.style.width=1);k.style.width=Math.max(k.style.width-2*(k.padding||0),1);w=w*O%180;"parallel"===y&&(w-=90);90<w?w-=180:-90>w&&(w+=180);k.rotation=w}k.textPath&&(0===f.shapeExisting.innerR&&k.textPath.enabled?(k.rotation=0,k.textPath.enabled=
!1,k.style.width=Math.max(2*f.shapeExisting.r-2*(k.padding||0),1)):f.dlOptions&&f.dlOptions.textPath&&!f.dlOptions.textPath.enabled&&"circular"===y&&(k.textPath.enabled=!0),k.textPath.enabled&&(k.rotation=0,k.style.width=Math.max((f.outerArcLength+f.innerArcLength)/2-2*(k.padding||0),1)));0===k.rotation&&(k.rotation=.001);f.dlOptions=k;if(!x&&A){x=!0;var E=B}f.draw({animatableAttribs:M,attribs:K(t,!q.styledMode&&a.pointAttribs(f,f.selected&&"select")),onComplete:E,group:d,renderer:z,shapeType:"arc",
shapeArgs:v})});k&&x?(a.hasRendered=!1,a.options.dataLabels.defer=!0,t.prototype.drawDataLabels.call(a),a.hasRendered=!0,w&&B()):t.prototype.drawDataLabels.call(a)};c.prototype.layoutAlgorithm=function(a,b,c){var f=a.start,d=a.end-f,g=a.val,e=a.x,h=a.y,l=c&&D(c.levelSize)&&I(c.levelSize.value)?c.levelSize.value:0,k=a.r,n=k+l,p=c&&I(c.slicedOffset)?c.slicedOffset:0;return(b||[]).reduce(function(a,b){var c=1/g*b.val*d,m=f+c/2,q=e+Math.cos(m)*p;m=h+Math.sin(m)*p;b={x:b.sliced?q:e,y:b.sliced?m:h,innerR:k,
r:n,radius:l,start:f,end:f+c};a.push(b);f=b.end;return a},[])};c.prototype.setShapeArgs=function(a,b,c){var d=[],f=c[a.level+1];a=a.children.filter(function(a){return a.visible});d=this.layoutAlgorithm(b,a,f);a.forEach(function(a,b){b=d[b];var f=b.start+(b.end-b.start)/2,e=b.innerR+(b.r-b.innerR)/2,g=b.end-b.start;e=0===b.innerR&&6.28<g?{x:b.x,y:b.y}:{x:b.x+Math.cos(f)*e,y:b.y+Math.sin(f)*e};var h=a.val?a.childrenTotal>a.val?a.childrenTotal:a.val:a.childrenTotal;this.points[a.i]&&(this.points[a.i].innerArcLength=
g*b.innerR,this.points[a.i].outerArcLength=g*b.r);a.shapeArgs=G(b,{plotX:e.x,plotY:e.y+4*Math.abs(Math.cos(f))});a.values=G(b,{val:h});a.children.length&&this.setShapeArgs(a,a.values,c)},this)};c.prototype.translate=function(){var a=this,c=a.options,g=a.center=b.call(a),h=a.startAndEndRadians=d(c.startAngle,c.endAngle),k=g[3]/2,p=g[2]/2-k,e=w(a),m=a.nodeMap,l=m&&m[e],r={};a.shapeRoot=l&&l.shapeArgs;t.prototype.translate.call(a);var u=a.tree=a.getTree();a.renderTraverseUpButton(e);m=a.nodeMap;l=m[e];
var z=F(l.parent)?l.parent:"";z=m[z];var x=q.getLevelFromAndTo(l);var B=x.from,D=x.to;x=S({from:B,levels:a.options.levels,to:D,defaults:{colorByPoint:c.colorByPoint,dataLabels:c.dataLabels,levelIsConstant:c.levelIsConstant,levelSize:c.levelSize,slicedOffset:c.slicedOffset}});x=q.calculateLevelSizes(x,{diffRadius:p,from:B,to:D});L(u,{before:n,idRoot:e,levelIsConstant:c.levelIsConstant,mapOptionsToLevel:x,mapIdToNode:m,points:a.points,series:a});c=m[""].shapeArgs={end:h.end,r:k,start:h.start,val:l.val,
x:g[0],y:g[1]};this.setShapeArgs(z,c,x);a.mapOptionsToLevel=x;a.data.forEach(function(b){r[b.id]&&N(31,!1,a.chart);r[b.id]=!0});r={}};c.defaultOptions=G(z.defaultOptions,{center:["50%","50%"],colorByPoint:!1,opacity:1,dataLabels:{allowOverlap:!0,defer:!0,rotationMode:"auto",style:{textOverflow:"ellipsis"}},rootId:void 0,levelIsConstant:!0,levelSize:{value:1,unit:"weight"},slicedOffset:10});return c}(z);K(k.prototype,{drawDataLabels:a,pointAttribs:h.prototype.pointAttribs,pointClass:r,utils:q});g.registerSeriesType("sunburst",
k);"";return k});u(a,"masters/modules/sunburst.src.js",[],function(){})});
//# sourceMappingURL=sunburst.js.map
```
