## Supplementary material for "Exploring natural odour landscapes: A case study with implications for human-biting insects": Figure S4: tilemap.js

```
/*
 Highmaps JS v9.3.1 (2021-11-05)

 Tilemap module

 (c) 2010-2021 Highsoft AS

 License: www.highcharts.com/license
*/
'use strict';(function(b){"object"===typeof module&&module.exports?(b["default"]=b,module.exports=b):"function"===typeof define&&define.amd?define("highcharts/modules/tilemap",["highcharts","highcharts/modules/map"],function(l){b(l);b.Highcharts=l;return b}):b("undefined"!==typeof Highcharts?Highcharts:void 0)})(function(b){function l(b,e,k,f){b.hasOwnProperty(e)||(b[e]=f.apply(null,k))}b=b?b._modules:{};l(b,"Series/Tilemap/TilemapPoint.js",[b["Core/Axis/Color/ColorAxisComposition.js"],b["Core/Series/SeriesRegistry.js"],
b["Core/Utilities.js"]],function(b,e,k){var f=this&&this.__extends||function(){var b=function(d,k){b=Object.setPrototypeOf||{__proto__:[]}instanceof Array&&function(a,c){a.__proto__=c}||function(a,c){for(var h in c)c.hasOwnProperty(h)&&(a[h]=c[h])};return b(d,k)};return function(d,k){function a(){this.constructor=d}b(d,k);d.prototype=null===k?Object.create(k):(a.prototype=k.prototype,new a)}}(),g=e.series.prototype.pointClass;k=k.extend;e=function(b){function d(){var d=null!==b&&b.apply(this,arguments)||
this;d.options=void 0;d.radius=void 0;d.series=void 0;d.tileEdges=void 0;return d}f(d,b);d.prototype.haloPath=function(){return this.series.tileShape.haloPath.apply(this,arguments)};return d}(e.seriesTypes.heatmap.prototype.pointClass);k(e.prototype,{setState:g.prototype.setState,setVisible:b.pointSetVisible});return e});l(b,"Series/Tilemap/TilemapShapes.js",[b["Core/Globals.js"],b["Core/Series/SeriesRegistry.js"],b["Core/Utilities.js"]],function(b,e,k){function f(a,c,b){a=a.options;return{xPad:(a.colsize||
1)/-c,yPad:(a.rowsize||1)/-b}}e=e.seriesTypes;var g=e.heatmap,t=e.scatter,d=k.clamp,l=k.pick;return{hexagon:{alignDataLabel:t.prototype.alignDataLabel,getSeriesPadding:function(a){return f(a,3,2)},haloPath:function(a){if(!a)return[];var c=this.tileEdges;return[["M",c.x2-a,c.y1+a],["L",c.x3+a,c.y1+a],["L",c.x4+1.5*a,c.y2],["L",c.x3+a,c.y3-a],["L",c.x2-a,c.y3-a],["L",c.x1-1.5*a,c.y2],["Z"]]},translate:function(){var a=this.options,c=this.xAxis,b=this.yAxis,k=a.pointPadding||0,p=(a.colsize||1)/3,v=(a.rowsize||
1)/2,n;this.generatePoints();this.points.forEach(function(a){var r=d(Math.floor(c.len-c.translate(a.x-2*p,0,1,0,1)),-c.len,2*c.len),h=d(Math.floor(c.len-c.translate(a.x-p,0,1,0,1)),-c.len,2*c.len),e=d(Math.floor(c.len-c.translate(a.x+p,0,1,0,1)),-c.len,2*c.len),u=d(Math.floor(c.len-c.translate(a.x+2*p,0,1,0,1)),-c.len,2*c.len),w=d(Math.floor(b.translate(a.y-v,0,1,0,1)),-b.len,2*b.len),x=d(Math.floor(b.translate(a.y,0,1,0,1)),-b.len,2*b.len),q=d(Math.floor(b.translate(a.y+v,0,1,0,1)),-b.len,2*b.len),
m=l(a.pointPadding,k),g=m*Math.abs(h-r)/Math.abs(q-x);g=c.reversed?-g:g;var f=c.reversed?-m:m;m=b.reversed?-m:m;a.x%2&&(n=n||Math.round(Math.abs(q-w)/2)*(b.reversed?-1:1),w+=n,x+=n,q+=n);a.plotX=a.clientX=(h+e)/2;a.plotY=x;r+=g+f;h+=f;e-=f;u-=g+f;w-=m;q+=m;a.tileEdges={x1:r,x2:h,x3:e,x4:u,y1:w,y2:x,y3:q};a.shapeType="path";a.shapeArgs={d:[["M",h,w],["L",e,w],["L",u,x],["L",e,q],["L",h,q],["L",r,x],["Z"]]}});this.translateColors()}},diamond:{alignDataLabel:t.prototype.alignDataLabel,getSeriesPadding:function(a){return f(a,
2,2)},haloPath:function(a){if(!a)return[];var c=this.tileEdges;return[["M",c.x2,c.y1+a],["L",c.x3+a,c.y2],["L",c.x2,c.y3-a],["L",c.x1-a,c.y2],["Z"]]},translate:function(){var a=this.options,c=this.xAxis,b=this.yAxis,k=a.pointPadding||0,e=a.colsize||1,v=(a.rowsize||1)/2,n;this.generatePoints();this.points.forEach(function(a){var g=d(Math.round(c.len-c.translate(a.x-e,0,1,0,0)),-c.len,2*c.len),p=d(Math.round(c.len-c.translate(a.x,0,1,0,0)),-c.len,2*c.len),r=d(Math.round(c.len-c.translate(a.x+e,0,1,
0,0)),-c.len,2*c.len),u=d(Math.round(b.translate(a.y-v,0,1,0,0)),-b.len,2*b.len),h=d(Math.round(b.translate(a.y,0,1,0,0)),-b.len,2*b.len),f=d(Math.round(b.translate(a.y+v,0,1,0,0)),-b.len,2*b.len),q=l(a.pointPadding,k),m=q*Math.abs(p-g)/Math.abs(f-h);m=c.reversed?-m:m;q=b.reversed?-q:q;a.x%2&&(n=Math.abs(f-u)/2*(b.reversed?-1:1),u+=n,h+=n,f+=n);a.plotX=a.clientX=p;a.plotY=h;g+=m;r-=m;u-=q;f+=q;a.tileEdges={x1:g,x2:p,x3:r,y1:u,y2:h,y3:f};a.shapeType="path";a.shapeArgs={d:[["M",p,u],["L",r,h],["L",
p,f],["L",g,h],["Z"]]}});this.translateColors()}},circle:{alignDataLabel:t.prototype.alignDataLabel,getSeriesPadding:function(a){return f(a,2,2)},haloPath:function(a){return t.prototype.pointClass.prototype.haloPath.call(this,a+(a&&this.radius))},translate:function(){var a=this.options,c=this.xAxis,b=this.yAxis,k=a.pointPadding||0,e=(a.rowsize||1)/2,v=a.colsize||1,n,f,g,t,l=!1;this.generatePoints();this.points.forEach(function(a){var p=d(Math.round(c.len-c.translate(a.x,0,1,0,0)),-c.len,2*c.len),
h=d(Math.round(b.translate(a.y,0,1,0,0)),-b.len,2*b.len),r=k,m=!1;"undefined"!==typeof a.pointPadding&&(r=a.pointPadding,l=m=!0);if(!t||l)n=Math.abs(d(Math.floor(c.len-c.translate(a.x+v,0,1,0,0)),-c.len,2*c.len)-p),f=Math.abs(d(Math.floor(b.translate(a.y+e,0,1,0,0)),-b.len,2*b.len)-h),g=Math.floor(Math.sqrt(n*n+f*f)/2),t=Math.min(n,g,f)-r,l&&!m&&(l=!1);a.x%2&&(h+=f*(b.reversed?-1:1));a.plotX=a.clientX=p;a.plotY=h;a.radius=t;a.shapeType="circle";a.shapeArgs={x:p,y:h,r:t}});this.translateColors()}},
square:{alignDataLabel:g.prototype.alignDataLabel,translate:g.prototype.translate,getSeriesPadding:b.noop,haloPath:g.prototype.pointClass.prototype.haloPath}}});l(b,"Series/Tilemap/TilemapComposition.js",[b["Core/Axis/Axis.js"],b["Core/Utilities.js"]],function(b,e){e=e.addEvent;e(b,"afterSetAxisTranslation",function(){if(!this.recomputingForTilemap&&"colorAxis"!==this.coll){var b=this,f=b.series.map(function(e){return e.getSeriesPixelPadding&&e.getSeriesPixelPadding(b)}).reduce(function(b,d){return(b&&
b.padding)>(d&&d.padding)?b:d},void 0)||{padding:0,axisLengthFactor:1},e=Math.round(f.padding*f.axisLengthFactor);f.padding&&(b.len-=e,b.recomputingForTilemap=!0,b.setAxisTranslation(),delete b.recomputingForTilemap,b.minPixelPadding+=f.padding,b.len+=e)}})});l(b,"Series/Tilemap/TilemapSeries.js",[b["Core/Globals.js"],b["Core/Series/SeriesRegistry.js"],b["Series/Tilemap/TilemapPoint.js"],b["Series/Tilemap/TilemapShapes.js"],b["Core/Utilities.js"]],function(b,e,k,f,g){var l=this&&this.__extends||function(){var b=
function(a,c){b=Object.setPrototypeOf||{__proto__:[]}instanceof Array&&function(b,a){b.__proto__=a}||function(b,a){for(var c in a)a.hasOwnProperty(c)&&(b[c]=a[c])};return b(a,c)};return function(a,c){function d(){this.constructor=a}b(a,c);a.prototype=null===c?Object.create(c):(d.prototype=c.prototype,new d)}}();b=b.noop;var d=e.seriesTypes,y=d.column,a=d.heatmap;d=d.scatter;var c=g.extend,h=g.merge;g=function(b){function c(){var a=null!==b&&b.apply(this,arguments)||this;a.data=void 0;a.options=void 0;
a.points=void 0;a.tileShape=void 0;return a}l(c,b);c.prototype.alignDataLabel=function(){return this.tileShape.alignDataLabel.apply(this,Array.prototype.slice.call(arguments))};c.prototype.drawPoints=function(){var a=this;y.prototype.drawPoints.call(this);this.points.forEach(function(b){b.graphic&&b.graphic[a.chart.styledMode?"css":"animate"](a.colorAttribs(b))})};c.prototype.getSeriesPixelPadding=function(a){var b=a.isXAxis,c=this.tileShape.getSeriesPadding(this);if(!c)return{padding:0,axisLengthFactor:1};
var d=Math.round(a.translate(b?2*c.xPad:c.yPad,0,1,0,1));a=Math.round(a.translate(b?c.xPad:0,0,1,0,1));return{padding:Math.abs(d-a)||0,axisLengthFactor:b?2:1.1}};c.prototype.setOptions=function(){var a=b.prototype.setOptions.apply(this,Array.prototype.slice.call(arguments));this.tileShape=f[a.tileShape];return a};c.prototype.translate=function(){return this.tileShape.translate.apply(this,Array.prototype.slice.call(arguments))};c.defaultOptions=h(a.defaultOptions,{marker:null,states:{hover:{halo:{enabled:!0,
size:2,opacity:.5,attributes:{zIndex:3}}}},pointPadding:2,tileShape:"hexagon"});return c}(a);c(g.prototype,{getSymbol:b,markerAttribs:d.prototype.markerAttribs,pointAttribs:y.prototype.pointAttribs,pointClass:k});e.registerSeriesType("tilemap",g);"";"";return g});l(b,"masters/modules/tilemap.src.js",[],function(){})});
//# sourceMappingURL=tilemap.js.map
```
