## Supplementary material for "Exploring natural odour landscapes: A case study with implications for human-biting insects": Figure S4: timeline.js

```
/*
 Highcharts JS v9.3.1 (2021-11-05)

 Timeline series

 (c) 2010-2021 Highsoft AS
 Author: Daniel Studencki

 License: www.highcharts.com/license
*/
'use strict';(function(a){"object"===typeof module&&module.exports?(a["default"]=a,module.exports=a):"function"===typeof define&&define.amd?define("highcharts/modules/timeline",["highcharts"],function(c){a(c);a.Highcharts=c;return a}):a("undefined"!==typeof Highcharts?Highcharts:void 0)})(function(a){function c(a,p,m,c){a.hasOwnProperty(p)||(a[p]=c.apply(null,m))}a=a?a._modules:{};c(a,"Series/Timeline/TimelinePoint.js",[a["Core/Series/Point.js"],a["Core/Series/SeriesRegistry.js"],a["Core/Utilities.js"]],
function(a,c,m){var p=this&&this.__extends||function(){var a=function(d,f){a=Object.setPrototypeOf||{__proto__:[]}instanceof Array&&function(f,a){f.__proto__=a}||function(f,a){for(var h in a)a.hasOwnProperty(h)&&(f[h]=a[h])};return a(d,f)};return function(d,f){function r(){this.constructor=d}a(d,f);d.prototype=null===f?Object.create(f):(r.prototype=f.prototype,new r)}}(),k=c.seriesTypes.pie.prototype.pointClass,z=m.defined,q=m.isNumber,u=m.merge,v=m.objectEach,n=m.pick;return function(c){function d(){var f=
null!==c&&c.apply(this,arguments)||this;f.options=void 0;f.series=void 0;return f}p(d,c);d.prototype.alignConnector=function(){var f=this.series,a=this.connector,c=this.dataLabel,h=this.dataLabel.options=u(f.options.dataLabels,this.options.dataLabels),g=this.series.chart,b=a.getBBox(),e=b.x+c.translateX;b=b.y+c.translateY;g.inverted?b-=c.options.connectorWidth/2:e+=c.options.connectorWidth/2;g=g.isInsidePlot(e,b);a[g?"animate":"attr"]({d:this.getConnectorPath()});f.chart.styledMode||a.attr({stroke:h.connectorColor||
this.color,"stroke-width":h.connectorWidth,opacity:c[z(c.newOpacity)?"newOpacity":"opacity"]})};d.prototype.drawConnector=function(){var a=this.series;this.connector||(this.connector=a.chart.renderer.path(this.getConnectorPath()).attr({zIndex:-1}).add(this.dataLabel));this.series.chart.isInsidePlot(this.dataLabel.x,this.dataLabel.y)&&this.alignConnector()};d.prototype.getConnectorPath=function(){var a=this.series.chart,c=this.series.xAxis.len,d=a.inverted,h=d?"x2":"y2",g=this.dataLabel,b=g.targetPosition,
e={x1:this.plotX,y1:this.plotY,x2:this.plotX,y2:q(b.y)?b.y:g.y},l=(g.alignAttr||g)[h[0]]<this.series.yAxis.len/2;d&&(e={x1:this.plotY,y1:c-this.plotX,x2:b.x||g.x,y2:c-this.plotX});l&&(e[h]+=g[d?"width":"height"]);v(e,function(b,a){e[a]-=(g.alignAttr||g)[a[0]]});return a.renderer.crispLine([["M",e.x1,e.y1],["L",e.x2,e.y2]],g.options.connectorWidth)};d.prototype.init=function(){var a=c.prototype.init.apply(this,arguments);a.name=n(a.name,"Event");a.y=1;return a};d.prototype.isValid=function(){return null!==
this.options.y};d.prototype.setState=function(){var a=c.prototype.setState;this.isNull||a.apply(this,arguments)};d.prototype.setVisible=function(a,c){var f=this.series;c=n(c,f.options.ignoreHiddenPoint);k.prototype.setVisible.call(this,a,!1);f.processData();c&&f.chart.redraw()};d.prototype.applyOptions=function(f,d){f=a.prototype.optionsToObject.call(this,f);this.userDLOptions=u(this.userDLOptions,f.dataLabels);return c.prototype.applyOptions.call(this,f,d)};return d}(c.series.prototype.pointClass)});
c(a,"Series/Timeline/TimelineSeries.js",[a["Core/Legend/LegendSymbol.js"],a["Core/Series/SeriesRegistry.js"],a["Core/Renderer/SVG/SVGElement.js"],a["Series/Timeline/TimelinePoint.js"],a["Core/Utilities.js"]],function(a,c,m,x,k){var p=this&&this.__extends||function(){var a=function(c,b){a=Object.setPrototypeOf||{__proto__:[]}instanceof Array&&function(b,a){b.__proto__=a}||function(b,a){for(var e in a)a.hasOwnProperty(e)&&(b[e]=a[e])};return a(c,b)};return function(c,b){function e(){this.constructor=
c}a(c,b);c.prototype=null===b?Object.create(b):(e.prototype=b.prototype,new e)}}(),q=c.seriesTypes,u=q.column,v=q.line,n=k.addEvent,y=k.arrayMax,d=k.arrayMin,f=k.defined;q=k.extend;var r=k.merge,w=k.pick;k=function(a){function c(){var b=null!==a&&a.apply(this,arguments)||this;b.data=void 0;b.options=void 0;b.points=void 0;b.userOptions=void 0;b.visibilityMap=void 0;return b}p(c,a);c.prototype.alignDataLabel=function(b,e,c,f){var l=this.chart.inverted,d=this.visibilityMap.filter(function(b){return b}),
t=this.visiblePointsCount,g=d.indexOf(b);d=this.options.dataLabels;var k=b.userDLOptions||{};g=d.alternate?g&&g!==t-1?2:1.5:1;t=Math.floor(this.xAxis.len/t);var h=e.padding;if(b.visible){var m=Math.abs(k.x||b.options.dataLabels.x);l?(l=2*(m-h)-b.itemHeight/2,l={width:l+"px",textOverflow:e.width/l*e.height/2>t*g?"ellipsis":"none"}):l={width:(k.width||d.width||t*g-2*h)+"px"};e.css(l);this.chart.styledMode||e.shadow(d.shadow)}a.prototype.alignDataLabel.apply(this,arguments)};c.prototype.bindAxes=function(){var b=
this;a.prototype.bindAxes.call(b);["xAxis","yAxis"].forEach(function(a){"xAxis"!==a||b[a].userOptions.type||(b[a].categories=b[a].hasNames=!0)})};c.prototype.distributeDL=function(){var a=this,e=a.options.dataLabels,c=1;if(e){var d=e.distance||0;a.points.forEach(function(b){var l;b.options.dataLabels=r((l={},l[a.chart.inverted?"x":"y"]=e.alternate&&c%2?-d:d,l),b.userDLOptions);c++})}};c.prototype.generatePoints=function(){var b=this;a.prototype.generatePoints.apply(b);b.points.forEach(function(a,
c){a.applyOptions({x:b.xData[c]},b.xData[c])})};c.prototype.getVisibilityMap=function(){return(this.data.length?this.data:this.userOptions.data).map(function(a){return a&&!1!==a.visible&&!a.isNull?a:!1})};c.prototype.getXExtremes=function(a){var b=this;a=a.filter(function(a,c){return b.points[c].isValid()&&b.points[c].visible});return{min:d(a),max:y(a)}};c.prototype.init=function(){var b=this;a.prototype.init.apply(b,arguments);b.eventsToUnbind.push(n(b,"afterTranslate",function(){var a,c=Number.MAX_VALUE;
b.points.forEach(function(b){b.isInside=b.isInside&&b.visible;b.visible&&!b.isNull&&(f(a)&&(c=Math.min(c,Math.abs(b.plotX-a))),a=b.plotX)});b.closestPointRangePx=c}));b.eventsToUnbind.push(n(b,"drawDataLabels",function(){b.distributeDL()}));b.eventsToUnbind.push(n(b,"afterDrawDataLabels",function(){var a;b.points.forEach(function(b){if(a=b.dataLabel)return a.animate=function(a){this.targetPosition&&(this.targetPosition=a);return m.prototype.animate.apply(this,arguments)},a.targetPosition||(a.targetPosition=
{}),b.drawConnector()})}));b.eventsToUnbind.push(n(b.chart,"afterHideOverlappingLabel",function(){b.points.forEach(function(a){a.connector&&a.dataLabel&&a.dataLabel.oldOpacity!==a.dataLabel.newOpacity&&a.alignConnector()})}))};c.prototype.markerAttribs=function(b,c){var e=this.options.marker,d=b.marker||{},f=d.symbol||e.symbol,g=w(d.width,e.width,this.closestPointRangePx),k=w(d.height,e.height),h=0;if(this.xAxis.dateTime)return a.prototype.markerAttribs.call(this,b,c);c&&(e=e.states[c]||{},c=d.states&&
d.states[c]||{},h=w(c.radius,e.radius,h+(e.radiusPlus||0)));b.hasImage=f&&0===f.indexOf("url");return{x:Math.floor(b.plotX)-g/2-h/2,y:b.plotY-k/2-h/2,width:g+h,height:k+h}};c.prototype.processData=function(){var b=0,c;this.visibilityMap=this.getVisibilityMap();this.visibilityMap.forEach(function(a){a&&b++});this.visiblePointsCount=b;for(c=0;c<this.xData.length;c++)this.yData[c]=1;a.prototype.processData.call(this,arguments)};c.defaultOptions=r(v.defaultOptions,{colorByPoint:!0,stickyTracking:!1,ignoreHiddenPoint:!0,
legendType:"point",lineWidth:4,tooltip:{headerFormat:'<span style="color:{point.color}">\u25cf</span> <span style="font-size: 10px"> {point.key}</span><br/>',pointFormat:"{point.description}"},states:{hover:{lineWidthPlus:0}},dataLabels:{enabled:!0,allowOverlap:!0,alternate:!0,backgroundColor:"#ffffff",borderWidth:1,borderColor:"#999999",borderRadius:3,color:"#333333",connectorWidth:1,distance:100,formatter:function(){var a=this.series.chart.styledMode?"<span>\u25cf </span>":'<span style="color:'+
this.point.color+'">\u25cf </span>';return a+='<span class="highcharts-strong">'+(this.key||"")+"</span><br/>"+(this.point.label||"")},style:{textOutline:"none",fontWeight:"normal",fontSize:"12px"},shadow:!1,verticalAlign:"middle"},marker:{enabledThreshold:0,symbol:"square",radius:6,lineWidth:2,height:15},showInLegend:!1,colorKey:"x"});return c}(v);q(k.prototype,{drawLegendSymbol:a.drawRectangle,drawTracker:u.prototype.drawTracker,pointClass:x,trackerGroups:["markerGroup","dataLabelsGroup"]});c.registerSeriesType("timeline",
k);"";"";return k});c(a,"masters/modules/timeline.src.js",[],function(){})});
//# sourceMappingURL=timeline.js.map
```
