## Supplementary material for "Exploring natural odour landscapes: A case study with implications for human-biting insects": Figure S4: treegrid.js

```
/*
 Highcharts Gantt JS v9.3.1 (2021-11-05)

 Tree Grid

 (c) 2016-2021 Jon Arild Nygard

 License: www.highcharts.com/license
*/
'use strict';(function(h){"object"===typeof module&&module.exports?(h["default"]=h,module.exports=h):"function"===typeof define&&define.amd?define("highcharts/modules/treegrid",["highcharts"],function(B){h(B);h.Highcharts=B;return h}):h("undefined"!==typeof Highcharts?Highcharts:void 0)})(function(h){function B(h,u,z,r){h.hasOwnProperty(u)||(h[u]=r.apply(null,z))}h=h?h._modules:{};B(h,"Core/Axis/BrokenAxis.js",[h["Extensions/Stacking.js"],h["Core/Utilities.js"]],function(h,u){var z=u.addEvent,r=u.find,
v=u.fireEvent,n=u.isArray,k=u.isNumber,x=u.pick,C;(function(a){function g(){"undefined"!==typeof this.brokenAxis&&this.brokenAxis.setBreaks(this.options.breaks,!1)}function q(){this.brokenAxis&&this.brokenAxis.hasBreaks&&(this.options.ordinal=!1)}function u(){var a=this.brokenAxis;if(a&&a.hasBreaks){for(var b=this.tickPositions,e=this.tickPositions.info,G=[],l=0;l<b.length;l++)a.isInAnyBreak(b[l])||G.push(b[l]);this.tickPositions=G;this.tickPositions.info=e}}function C(){this.brokenAxis||(this.brokenAxis=
new y(this))}function f(){var a=this.options.connectNulls,b=this.points,e=this.xAxis,G=this.yAxis;if(this.isDirty)for(var l=b.length;l--;){var H=b[l],c=!(null===H.y&&!1===a)&&(e&&e.brokenAxis&&e.brokenAxis.isInAnyBreak(H.x,!0)||G&&G.brokenAxis&&G.brokenAxis.isInAnyBreak(H.y,!0));H.visible=c?!1:!1!==H.options.visible}}function d(){this.drawBreaks(this.xAxis,["x"]);this.drawBreaks(this.yAxis,x(this.pointArrayMap,["y"]))}function e(a,b){var e=this,I=e.points,l,F,c,m;if(a&&a.brokenAxis&&a.brokenAxis.hasBreaks){var d=
a.brokenAxis;b.forEach(function(b){l=d&&d.breakArray||[];F=a.isXAxis?a.min:x(e.options.threshold,a.min);I.forEach(function(e){m=x(e["stack"+b.toUpperCase()],e[b]);l.forEach(function(b){if(k(F)&&k(m)){c=!1;if(F<b.from&&m>b.to||F>b.from&&m<b.from)c="pointBreak";else if(F<b.from&&m>b.from&&m<b.to||F>b.from&&m>b.to&&m<b.from)c="pointInBreak";c&&v(a,c,{point:e,brk:b})}})})})}}function c(){var b=this.currentDataGrouping,a=b&&b.gapSize;b=this.points.slice();var e=this.yAxis,c=this.options.gapSize,l=b.length-
1,d;if(c&&0<l)for("value"!==this.options.gapUnit&&(c*=this.basePointRange),a&&a>c&&a>=this.basePointRange&&(c=a),d=void 0;l--;)d&&!1!==d.visible||(d=b[l+1]),a=b[l],!1!==d.visible&&!1!==a.visible&&(d.x-a.x>c&&(d=(a.x+d.x)/2,b.splice(l+1,0,{isNull:!0,x:d}),e.stacking&&this.options.stacking&&(d=e.stacking.stacks[this.stackKey][d]=new h(e,e.options.stackLabels,!1,d,this.stack),d.total=0)),d=a);return this.getGraphPath(b)}var b=[];a.compose=function(a,I){-1===b.indexOf(a)&&(b.push(a),a.keepProps.push("brokenAxis"),
z(a,"init",C),z(a,"afterInit",g),z(a,"afterSetTickPositions",u),z(a,"afterSetOptions",q));if(-1===b.indexOf(I)){b.push(I);var F=I.prototype;F.drawBreaks=e;F.gappedPath=c;z(I,"afterGeneratePoints",f);z(I,"afterRender",d)}return a};var y=function(){function a(a){this.hasBreaks=!1;this.axis=a}a.isInBreak=function(a,b){var e=a.repeat||Infinity,l=a.from,d=a.to-a.from;b=b>=l?(b-l)%e:e-(l-b)%e;return a.inclusive?b<=d:b<d&&0!==b};a.lin2Val=function(b){var e=this.brokenAxis;e=e&&e.breakArray;if(!e||!k(b))return b;
var d;for(d=0;d<e.length;d++){var l=e[d];if(l.from>=b)break;else l.to<b?b+=l.len:a.isInBreak(l,b)&&(b+=l.len)}return b};a.val2Lin=function(b){var e=this.brokenAxis;e=e&&e.breakArray;if(!e||!k(b))return b;var d=b,l;for(l=0;l<e.length;l++){var c=e[l];if(c.to<=b)d-=c.len;else if(c.from>=b)break;else if(a.isInBreak(c,b)){d-=b-c.from;break}}return d};a.prototype.findBreakAt=function(a,b){return r(b,function(b){return b.from<a&&a<b.to})};a.prototype.isInAnyBreak=function(b,e){var d=this.axis,l=d.options.breaks||
[],c=l.length,J;if(c&&k(b)){for(;c--;)if(a.isInBreak(l[c],b)){var m=!0;J||(J=x(l[c].showPoints,!d.isXAxis))}var w=m&&e?m&&!J:m}return w};a.prototype.setBreaks=function(b,e){var d=this,c=d.axis,g=n(b)&&!!b.length;c.isDirty=d.hasBreaks!==g;d.hasBreaks=g;c.options.breaks=c.userOptions.breaks=b;c.forceRedraw=!0;c.series.forEach(function(b){b.isDirty=!0});g||c.val2lin!==a.val2Lin||(delete c.val2lin,delete c.lin2val);g&&(c.userOptions.ordinal=!1,c.lin2val=a.lin2Val,c.val2lin=a.val2Lin,c.setExtremes=function(b,
a,e,E,g){if(d.hasBreaks){for(var m=this.options.breaks||[],w;w=d.findBreakAt(b,m);)b=w.to;for(;w=d.findBreakAt(a,m);)a=w.from;a<b&&(a=b)}c.constructor.prototype.setExtremes.call(this,b,a,e,E,g)},c.setAxisTranslation=function(){c.constructor.prototype.setAxisTranslation.call(this);d.unitLength=void 0;if(d.hasBreaks){var b=c.options.breaks||[],e=[],w=[],g=x(c.pointRangePadding,0),l=0,A,f=c.userMin||c.min,y=c.userMax||c.max,q;b.forEach(function(b){A=b.repeat||Infinity;k(f)&&k(y)&&(a.isInBreak(b,f)&&
(f+=b.to%A-f%A),a.isInBreak(b,y)&&(y-=y%A-b.from%A))});b.forEach(function(b){p=b.from;A=b.repeat||Infinity;if(k(f)&&k(y)){for(;p-A>f;)p-=A;for(;p<f;)p+=A;for(q=p;q<y;q+=A)e.push({value:q,move:"in"}),e.push({value:q+b.to-b.from,move:"out",size:b.breakSize})}});e.sort(function(b,p){return b.value===p.value?("in"===b.move?0:1)-("in"===p.move?0:1):b.value-p.value});var t=0;var p=f;e.forEach(function(b){t+="in"===b.move?1:-1;1===t&&"in"===b.move&&(p=b.value);0===t&&k(p)&&(w.push({from:p,to:b.value,len:b.value-
p-(b.size||0)}),l+=b.value-p-(b.size||0))});d.breakArray=w;k(f)&&k(y)&&k(c.min)&&(d.unitLength=y-f-l+g,v(c,"afterBreaks"),c.staticScale?c.transA=c.staticScale:d.unitLength&&(c.transA*=(y-c.min+g)/d.unitLength),g&&(c.minPixelPadding=c.transA*(c.minPointOffset||0)),c.min=f,c.max=y)}});x(e,!0)&&c.chart.redraw()};return a}();a.Additions=y})(C||(C={}));return C});B(h,"Core/Axis/GridAxis.js",[h["Core/Axis/Axis.js"],h["Core/Axis/AxisDefaults.js"],h["Core/Globals.js"],h["Core/Utilities.js"]],function(h,u,
z,r){var v=z.dateFormats,n=r.addEvent,k=r.defined,x=r.erase,C=r.find,a=r.isArray,g=r.isNumber,q=r.merge,D=r.pick,K=r.timeUnits,f=r.wrap,d;(function(e){function c(b,p){var a={width:0,height:0};p.forEach(function(p){p=b[p];if(r.isObject(p,!0)){var t=r.isObject(p.label,!0)?p.label:{};p=t.getBBox?t.getBBox().height:0;t.textStr&&!g(t.textPxLength)&&(t.textPxLength=t.getBBox().width);var c=g(t.textPxLength)?Math.round(t.textPxLength):0;t.textStr&&(c=Math.round(t.getBBox().width));a.height=Math.max(p,a.height);
a.width=Math.max(c,a.width)}});"treegrid"===this.options.type&&this.treeGrid&&this.treeGrid.mapOfPosToGridNode&&(a.width+=this.options.labels.indentation*((this.treeGrid.mapOfPosToGridNode[-1].height||0)-1));return a}function b(){var b=this.grid;(b&&b.columns||[]).forEach(function(b){b.getOffset()})}function d(b){if(!0===(this.options.grid||{}).enabled){var p=this.axisTitle,a=this.height,t=this.horiz,c=this.left,d=this.offset,m=this.opposite,g=this.options,f=this.top,w=this.width,l=this.tickSize(),
y=p&&p.getBBox().width,E=g.title.x,k=g.title.y,A=D(g.title.margin,t?5:10);p=this.chart.renderer.fontMetrics(g.title.style.fontSize,p).f;l=(t?f+a:c)+(t?1:-1)*(m?-1:1)*(l?l[0]/2:0)+(this.side===e.Side.bottom?p:0);b.titlePosition.x=t?c-(y||0)/2-A+E:l+(m?w:0)+d+E;b.titlePosition.y=t?l-(m?a:0)+(m?p:-p)/2+d+k:f-A+k}}function F(){var b=this.chart,p=this.options.grid;p=void 0===p?{}:p;var a=this.userOptions;if(p.enabled){var c=this.options;c.labels.align=D(c.labels.align,"center");this.categories||(c.showLastLabel=
!1);this.labelRotation=0;c.labels.rotation=0}if(p.columns){c=this.grid.columns=[];for(var e=this.grid.columnIndex=0;++e<p.columns.length;){var d=q(a,p.columns[p.columns.length-e-1],{linkedTo:0,type:"category",scrollbar:{enabled:!1}});delete d.grid.columns;d=new h(this.chart,d);d.grid.isColumn=!0;d.grid.columnIndex=e;x(b.axes,d);x(b[this.coll],d);c.push(d)}}}function I(){var b=this.grid,a=this.options;if(!0===(a.grid||{}).enabled){var c=this.min||0,d=this.max||0;this.maxLabelDimensions=this.getMaxLabelDimensions(this.ticks,
this.tickPositions);this.rightWall&&this.rightWall.destroy();if(this.grid&&this.grid.isOuterAxis()&&this.axisLine){var m=a.lineWidth;if(m){m=this.getLinePath(m);var g=m[0],f=m[1],l=((this.tickSize("tick")||[1])[0]-1)*(this.side===e.Side.top||this.side===e.Side.left?-1:1);"M"===g[0]&&"L"===f[0]&&(this.horiz?(g[2]+=l,f[2]+=l):(g[1]+=l,f[1]+=l));!this.horiz&&this.chart.marginRight&&(g=[g,["L",this.left,g[2]||0]],l=["L",this.chart.chartWidth-this.chart.marginRight,this.toPixels(d+this.tickmarkOffset)],
f=[["M",f[1]||0,this.toPixels(d+this.tickmarkOffset)],l],this.grid.upperBorder||0===c%1||(this.grid.upperBorder=this.grid.renderBorder(g)),this.grid.upperBorder&&(this.grid.upperBorder.attr({stroke:a.lineColor,"stroke-width":a.lineWidth}),this.grid.upperBorder.animate({d:g})),this.grid.lowerBorder||0===d%1||(this.grid.lowerBorder=this.grid.renderBorder(f)),this.grid.lowerBorder&&(this.grid.lowerBorder.attr({stroke:a.lineColor,"stroke-width":a.lineWidth}),this.grid.lowerBorder.animate({d:f})));this.grid.axisLineExtra?
(this.grid.axisLineExtra.attr({stroke:a.lineColor,"stroke-width":a.lineWidth}),this.grid.axisLineExtra.animate({d:m})):this.grid.axisLineExtra=this.grid.renderBorder(m);this.axisLine[this.showAxis?"show":"hide"](!0)}}(b&&b.columns||[]).forEach(function(b){b.render()});if(!this.horiz&&this.chart.hasRendered&&(this.scrollbar||this.linkedParent&&this.linkedParent.scrollbar)){b=this.tickmarkOffset;a=this.tickPositions[this.tickPositions.length-1];m=this.tickPositions[0];for(f=void 0;(f=this.hiddenLabels.pop())&&
f.element;)f.show();(f=this.ticks[m].label)&&(c-m>b?this.hiddenLabels.push(f.hide()):f.show());(f=this.ticks[a].label)&&(a-d>b?this.hiddenLabels.push(f.hide()):f.show());(c=this.ticks[a].mark)&&(a-d<b&&0<a-d&&this.ticks[a].isLast?c.hide():this.ticks[a-1]&&c.show())}}}function v(){var b=this.tickPositions&&this.tickPositions.info,a=this.options,c=this.userOptions.labels||{};(a.grid||{}).enabled&&(this.horiz?(this.series.forEach(function(b){b.options.pointRange=0}),b&&a.dateTimeLabelFormats&&a.labels&&
!k(c.align)&&(!1===a.dateTimeLabelFormats[b.unitName].range||1<b.count)&&(a.labels.align="left",k(c.x)||(a.labels.x=3))):"treegrid"!==this.options.type&&this.grid&&this.grid.columns&&(this.minPointOffset=this.tickInterval))}function G(b){var a=this.options;b=b.userOptions;var c=a&&r.isObject(a.grid,!0)?a.grid:{};if(!0===c.enabled){var d=q(!0,{className:"highcharts-grid-axis "+(b.className||""),dateTimeLabelFormats:{hour:{list:["%H:%M","%H"]},day:{list:["%A, %e. %B","%a, %e. %b","%E"]},week:{list:["Week %W",
"W%W"]},month:{list:["%B","%b","%o"]}},grid:{borderWidth:1},labels:{padding:2,style:{fontSize:"13px"}},margin:0,title:{text:null,reserveSpace:!1,rotation:0},units:[["millisecond",[1,10,100]],["second",[1,10]],["minute",[1,5,15]],["hour",[1,6]],["day",[1]],["week",[1]],["month",[1]],["year",null]]},b);"xAxis"===this.coll&&(k(b.linkedTo)&&!k(b.tickPixelInterval)&&(d.tickPixelInterval=350),k(b.tickPixelInterval)||!k(b.linkedTo)||k(b.tickPositioner)||k(b.tickInterval)||(d.tickPositioner=function(b,a){var c=
this.linkedParent&&this.linkedParent.tickPositions&&this.linkedParent.tickPositions.info;if(c){for(var t=d.units||[],p=void 0,e=void 0,m=void 0,f=0;f<t.length;f++)if(t[f][0]===c.unitName){p=f;break}t[p+1]?(m=t[p+1][0],e=(t[p+1][1]||[1])[0]):"year"===c.unitName&&(m="year",e=10*c.count);c=K[m];this.tickInterval=c*e;return this.getTimeTicks({unitRange:c,count:e,unitName:m},b,a,this.options.startOfWeek)}}));q(!0,this.options,d);this.horiz&&(a.minPadding=D(b.minPadding,0),a.maxPadding=D(b.maxPadding,0));
g(a.grid.borderWidth)&&(a.tickWidth=a.lineWidth=c.borderWidth)}}function l(b){b=(b=b.userOptions)&&b.grid||{};var a=b.columns;b.enabled&&a&&q(!0,this.options,a[a.length-1])}function H(){(this.grid.columns||[]).forEach(function(b){b.setScale()})}function J(b){var c=u.defaultLeftAxisOptions,d=this.horiz,e=this.maxLabelDimensions,t=this.options.grid;t=void 0===t?{}:t;t.enabled&&e&&(c=2*Math.abs(c.labels.x),d=d?t.cellHeight||c+e.height:c+e.width,a(b.tickSize)?b.tickSize[0]=d:b.tickSize=[d,0])}function m(){this.axes.forEach(function(b){(b.grid&&
b.grid.columns||[]).forEach(function(b){b.setAxisSize();b.setAxisTranslation()})})}function w(b){var a=this.grid;(a.columns||[]).forEach(function(a){a.destroy(b.keepEvents)});a.columns=void 0}function E(b){b=b.userOptions||{};var a=b.grid||{};a.enabled&&k(a.borderColor)&&(b.tickColor=b.lineColor=a.borderColor);this.grid||(this.grid=new L(this));this.hiddenLabels=[]}function M(b){var a=this.label,c=this.axis,d=c.reversed,m=c.chart,t=c.options.grid||{},f=c.options.labels,l=f.align,w=e.Side[c.side],
y=b.tickmarkOffset,E=c.tickPositions,k=this.pos-y;E=g(E[b.index+1])?E[b.index+1]-y:(c.max||0)+y;var A=c.tickSize("tick");y=A?A[0]:0;A=A?A[1]/2:0;if(!0===t.enabled){if("top"===w){t=c.top+c.offset;var q=t-y}else"bottom"===w?(q=m.chartHeight-c.bottom+c.offset,t=q+y):(t=c.top+c.len-(c.translate(d?E:k)||0),q=c.top+c.len-(c.translate(d?k:E)||0));"right"===w?(w=m.chartWidth-c.right+c.offset,d=w+y):"left"===w?(d=c.left+c.offset,w=d-y):(w=Math.round(c.left+(c.translate(d?E:k)||0))-A,d=Math.min(Math.round(c.left+
(c.translate(d?k:E)||0))-A,c.left+c.len));this.slotWidth=d-w;b.pos.x="left"===l?w:"right"===l?d:w+(d-w)/2;b.pos.y=q+(t-q)/2;m=m.renderer.fontMetrics(f.style.fontSize,a&&a.element);a=a?a.getBBox().height:0;f.useHTML?b.pos.y+=m.b+-(a/2):(a=Math.round(a/m.h),b.pos.y+=(m.b-(m.h-m.f))/2+-((a-1)*m.h/2));b.pos.x+=c.horiz&&f.x||0}}function A(b){var a=b.axis,c=b.value;if(a.options.grid&&a.options.grid.enabled){var d=a.tickPositions,e=(a.linkedParent||a).series[0],m=c===d[0];d=c===d[d.length-1];var t=e&&C(e.options.data,
function(b){return b[a.isXAxis?"x":"y"]===c}),f=void 0;t&&e.is("gantt")&&(f=q(t),z.seriesTypes.gantt.prototype.pointClass.setGanttPointAliases(f));b.isFirst=m;b.isLast=d;b.point=f}}function N(){var b=this.options,a=this.categories,c=this.tickPositions,d=c[0],e=c[c.length-1],m=this.linkedParent&&this.linkedParent.min||this.min,f=this.linkedParent&&this.linkedParent.max||this.max,g=this.tickInterval;!0!==(b.grid||{}).enabled||a||!this.horiz&&!this.isLinked||(d<m&&d+g>m&&!b.startOnTick&&(c[0]=m),e>f&&
e-g<f&&!b.endOnTick&&(c[c.length-1]=f))}function O(b){var a=this.options.grid;return!0===(void 0===a?{}:a).enabled&&this.categories?this.tickInterval:b.apply(this,Array.prototype.slice.call(arguments,1))}(function(b){b[b.top=0]="top";b[b.right=1]="right";b[b.bottom=2]="bottom";b[b.left=3]="left"})(e.Side||(e.Side={}));e.compose=function(a,e,g){-1===a.keepProps.indexOf("grid")&&(a.keepProps.push("grid"),a.prototype.getMaxLabelDimensions=c,f(a.prototype,"unsquish",O),n(a,"init",E),n(a,"afterGetOffset",
b),n(a,"afterGetTitlePosition",d),n(a,"afterInit",F),n(a,"afterRender",I),n(a,"afterSetAxisTranslation",v),n(a,"afterSetOptions",G),n(a,"afterSetOptions",l),n(a,"afterSetScale",H),n(a,"afterTickSize",J),n(a,"trimTicks",N),n(a,"destroy",w));n(e,"afterSetChartSize",m);n(g,"afterGetLabelPosition",M);n(g,"labelFormat",A);return a};var L=function(){function b(b){this.axis=b}b.prototype.isOuterAxis=function(){var b=this.axis,a=b.grid.columnIndex,c=b.linkedParent&&b.linkedParent.grid.columns||b.grid.columns,
d=a?b.linkedParent:b,e=-1,m=0;b.chart[b.coll].forEach(function(a,c){a.side!==b.side||a.options.isInternal||(m=c,a===d&&(e=c))});return m===e&&(g(a)?c.length===a:!0)};b.prototype.renderBorder=function(b){var a=this.axis,c=a.chart.renderer,d=a.options;b=c.path(b).addClass("highcharts-axis-line").add(a.axisBorder);c.styledMode||b.attr({stroke:d.lineColor,"stroke-width":d.lineWidth,zIndex:7});return b};return b}();e.Additions=L})(d||(d={}));v.E=function(a){return this.dateFormat("%a",a,!0).charAt(0)};
v.W=function(a){a=new this.Date(a);var c=(this.get("Day",a)+6)%7,b=new this.Date(a.valueOf());this.set("Date",b,this.get("Date",a)-c+3);c=new this.Date(this.get("FullYear",b),0,1);4!==this.get("Day",c)&&(this.set("Month",a,0),this.set("Date",a,1+(11-this.get("Day",c))%7));return(1+Math.floor((b.valueOf()-c.valueOf())/6048E5)).toString()};"";return d});B(h,"Gantt/Tree.js",[h["Core/Utilities.js"]],function(h){var u=h.extend,z=h.isNumber,r=h.pick,v=function(k,h){var n=k.reduce(function(a,g){var k=r(g.parent,
"");"undefined"===typeof a[k]&&(a[k]=[]);a[k].push(g);return a},{});Object.keys(n).forEach(function(a,g){var k=n[a];""!==a&&-1===h.indexOf(a)&&(k.forEach(function(a){g[""].push(a)}),delete g[a])});return n},n=function(k,h,v,a,g,q){var D=0,x=0,f=q&&q.after,d=q&&q.before;h={data:a,depth:v-1,id:k,level:v,parent:h};var e,c;"function"===typeof d&&d(h,q);d=(g[k]||[]).map(function(b){var a=n(b.id,k,v+1,b,g,q),d=b.start;b=!0===b.milestone?d:b.end;e=!z(e)||d<e?d:e;c=!z(c)||b>c?b:c;D=D+1+a.descendants;x=Math.max(a.height+
1,x);return a});a&&(a.start=r(a.start,e),a.end=r(a.end,c));u(h,{children:d,descendants:D,height:x});"function"===typeof f&&f(h,q);return h};return{getListOfParents:v,getNode:n,getTree:function(k,h){var u=k.map(function(a){return a.id});k=v(k,u);return n("",null,1,null,k,h)}}});B(h,"Core/Axis/TreeGridTick.js",[h["Core/Utilities.js"]],function(h){var u=h.addEvent,z=h.isObject,r=h.isNumber,v=h.pick,n=h.wrap,k;(function(h){function k(){this.treeGrid||(this.treeGrid=new x(this))}function a(a,d){a=a.treeGrid;
var e=!a.labelIcon,c=d.renderer,b=d.xy,f=d.options,g=f.width||0,h=f.height||0,k=b.x-g/2-(f.padding||0);b=b.y-h/2;var q=d.collapsed?90:180,l=d.show&&r(b),n=a.labelIcon;n||(a.labelIcon=n=c.path(c.symbols[f.type](f.x||0,f.y||0,g,h)).addClass("highcharts-label-icon").add(d.group));n.attr({y:l?0:-9999});c.styledMode||n.attr({cursor:"pointer",fill:v(d.color,"#666666"),"stroke-width":1,stroke:f.lineColor,strokeWidth:f.lineWidth||0});n[e?"attr":"animate"]({translateX:k,translateY:b,rotation:q})}function g(a,
d,e,c,b,g,h,k,q){var f=v(this.options&&this.options.labels,g);g=this.pos;var l=this.axis,n="treegrid"===l.options.type;a=a.apply(this,[d,e,c,b,f,h,k,q]);n&&(d=f&&z(f.symbol,!0)?f.symbol:{},f=f&&r(f.indentation)?f.indentation:0,g=(g=(l=l.treeGrid.mapOfPosToGridNode)&&l[g])&&g.depth||1,a.x+=(d.width||0)+2*(d.padding||0)+(g-1)*f);return a}function q(f){var d=this,e=d.pos,c=d.axis,b=d.label,g=c.treeGrid.mapOfPosToGridNode,h=c.options,k=v(d.options&&d.options.labels,h&&h.labels),q=k&&z(k.symbol,!0)?k.symbol:
{},n=(g=g&&g[e])&&g.depth;h="treegrid"===h.type;var l=-1<c.tickPositions.indexOf(e);e=c.chart.styledMode;h&&g&&b&&b.element&&b.addClass("highcharts-treegrid-node-level-"+n);f.apply(d,Array.prototype.slice.call(arguments,1));h&&b&&b.element&&g&&g.descendants&&0<g.descendants&&(c=c.treeGrid.isCollapsed(g),a(d,{color:!e&&b.styles&&b.styles.color||"",collapsed:c,group:b.parentGroup,options:q,renderer:b.renderer,show:l,xy:b.xy}),q="highcharts-treegrid-node-"+(c?"expanded":"collapsed"),b.addClass("highcharts-treegrid-node-"+
(c?"collapsed":"expanded")).removeClass(q),e||b.css({cursor:"pointer"}),[b,d.treeGrid.labelIcon].forEach(function(a){a&&!a.attachedTreeGridEvents&&(u(a.element,"mouseover",function(){b.addClass("highcharts-treegrid-node-active");b.renderer.styledMode||b.css({textDecoration:"underline"})}),u(a.element,"mouseout",function(){var a=z(k.style)?k.style:{};b.removeClass("highcharts-treegrid-node-active");b.renderer.styledMode||b.css({textDecoration:a.textDecoration})}),u(a.element,"click",function(){d.treeGrid.toggleCollapse()}),
a.attachedTreeGridEvents=!0)}))}var D=!1;h.compose=function(a){D||(u(a,"init",k),n(a.prototype,"getLabelPosition",g),n(a.prototype,"renderLabel",q),a.prototype.collapse=function(a){this.treeGrid.collapse(a)},a.prototype.expand=function(a){this.treeGrid.expand(a)},a.prototype.toggleCollapse=function(a){this.treeGrid.toggleCollapse(a)},D=!0)};var x=function(){function a(a){this.tick=a}a.prototype.collapse=function(a){var d=this.tick,c=d.axis,b=c.brokenAxis;b&&c.treeGrid.mapOfPosToGridNode&&(d=c.treeGrid.collapse(c.treeGrid.mapOfPosToGridNode[d.pos]),
b.setBreaks(d,v(a,!0)))};a.prototype.expand=function(a){var d=this.tick,c=d.axis,b=c.brokenAxis;b&&c.treeGrid.mapOfPosToGridNode&&(d=c.treeGrid.expand(c.treeGrid.mapOfPosToGridNode[d.pos]),b.setBreaks(d,v(a,!0)))};a.prototype.toggleCollapse=function(a){var d=this.tick,c=d.axis,b=c.brokenAxis;b&&c.treeGrid.mapOfPosToGridNode&&(d=c.treeGrid.toggleCollapse(c.treeGrid.mapOfPosToGridNode[d.pos]),b.setBreaks(d,v(a,!0)))};return a}();h.Additions=x})(k||(k={}));return k});B(h,"Series/TreeUtilities.js",[h["Core/Color/Color.js"],
h["Core/Utilities.js"]],function(h,u){function v(a,g){var h=g.before,k=g.idRoot,n=g.mapIdToNode[k],f=g.points[a.i],d=f&&f.options||{},e=[],c=0;a.levelDynamic=a.level-(!1!==g.levelIsConstant?0:n.level);a.name=C(f&&f.name,"");a.visible=k===a.id||!0===g.visible;"function"===typeof h&&(a=h(a,g));a.children.forEach(function(b,d){var f=r({},g);r(f,{index:d,siblings:a.children.length,visible:a.visible});b=v(b,f);e.push(b);b.visible&&(c+=b.val)});h=C(d.value,c);a.visible=0<=h&&(0<c||a.visible);a.children=
e;a.childrenTotal=c;a.isLeaf=a.visible&&!c;a.val=h;return a}var r=u.extend,B=u.isArray,n=u.isNumber,k=u.isObject,x=u.merge,C=u.pick;return{getColor:function(a,g){var k=g.index,n=g.mapOptionsToLevel,u=g.parentColor,f=g.parentColorIndex,d=g.series,e=g.colors,c=g.siblings,b=d.points,v=d.chart.options.chart,r;if(a){b=b[a.i];a=n[a.level]||{};if(n=b&&a.colorByPoint){var x=b.index%(e?e.length:v.colorCount);var z=e&&e[x]}if(!d.chart.styledMode){e=b&&b.options.color;v=a&&a.color;if(r=u)r=(r=a&&a.colorVariation)&&
"brightness"===r.key&&k&&c?h.parse(u).brighten(k/c*r.to).get():u;r=C(e,v,z,r,d.color)}var B=C(b&&b.options.colorIndex,a&&a.colorIndex,x,f,g.colorIndex)}return{color:r,colorIndex:B}},getLevelOptions:function(a){var g=null;if(k(a)){g={};var h=n(a.from)?a.from:1;var r=a.levels;var u={};var f=k(a.defaults)?a.defaults:{};B(r)&&(u=r.reduce(function(a,e){if(k(e)&&n(e.level)){var c=x({},e);var b=C(c.levelIsConstant,f.levelIsConstant);delete c.levelIsConstant;delete c.level;e=e.level+(b?0:h-1);k(a[e])?x(!0,
a[e],c):a[e]=c}return a},{}));r=n(a.to)?a.to:1;for(a=0;a<=r;a++)g[a]=x({},f,k(u[a])?u[a]:{})}return g},setTreeValues:v,updateRootId:function(a){if(k(a)){var g=k(a.options)?a.options:{};g=C(a.rootNode,g.rootId,"");k(a.userOptions)&&(a.userOptions.rootId=g);a.rootNode=g}return g}}});B(h,"Core/Axis/TreeGridAxis.js",[h["Core/Axis/BrokenAxis.js"],h["Core/Axis/GridAxis.js"],h["Gantt/Tree.js"],h["Core/Axis/TreeGridTick.js"],h["Series/TreeUtilities.js"],h["Core/Utilities.js"]],function(h,u,z,r,B,n){var k=
B.getLevelOptions,v=n.addEvent,C=n.find,a=n.fireEvent,g=n.isArray,q=n.isObject,D=n.isString,K=n.merge,f=n.pick,d=n.wrap,e;(function(c){function b(a,b){var c=a.collapseEnd||0;a=a.collapseStart||0;c>=b&&(a-=.5);return{from:a,to:c,showPoints:!1}}function e(a,b,c){var d=[],e=[],m={},g="boolean"===typeof b?b:!1,f={},h=-1;a=z.getTree(a,{after:function(a){a=f[a.pos];var b=0,c=0;a.children.forEach(function(a){c+=(a.descendants||0)+1;b=Math.max((a.height||0)+1,b)});a.descendants=c;a.height=b;a.collapsed&&
e.push(a)},before:function(a){var b=q(a.data,!0)?a.data:{},c=D(b.name)?b.name:"",e=m[a.parent];e=q(e,!0)?f[e.pos]:null;var k=function(a){return a.name===c},l;g&&q(e,!0)&&(l=C(e.children,k))?(k=l.pos,l.nodes.push(a)):k=h++;f[k]||(f[k]=l={depth:e?e.depth+1:0,name:c,id:b.id,nodes:[a],children:[],pos:k},-1!==k&&d.push(c),q(e,!0)&&e.children.push(l));D(a.id)&&(m[a.id]=a);l&&!0===b.collapsed&&(l.collapsed=!0);a.pos=k}});f=function(a,b){var c=function(a,d,e){var m=d+(-1===d?0:b-1),g=(m-d)/2,f=d+g;a.nodes.forEach(function(a){var b=
a.data;q(b,!0)&&(b.y=d+(b.seriesIndex||0),delete b.seriesIndex);a.pos=f});e[f]=a;a.pos=f;a.tickmarkOffset=g+.5;a.collapseStart=m+.5;a.children.forEach(function(a){c(a,m+1,e);m=(a.collapseEnd||0)-.5});a.collapseEnd=m+.5;return e};return c(a["-1"],-1,{})}(f,c);return{categories:d,mapOfIdToNode:m,mapOfPosToGridNode:f,collapsedNodes:e,tree:a}}function n(a){a.target.axes.filter(function(a){return"treegrid"===a.options.type}).forEach(function(b){var c=b.options||{},d=c.labels,m=c.uniqueNames;c=c.max;var f=
0;if(!b.treeGrid.mapOfPosToGridNode||b.series.some(function(a){return!a.hasRendered||a.isDirtyData||a.isDirty})){var h=b.series.reduce(function(a,b){b.visible&&((b.options.data||[]).forEach(function(c){b.options.keys&&b.options.keys.length&&(c=b.pointClass.prototype.optionsToObject.call({series:b},c),b.pointClass.setGanttPointAliases(c));q(c,!0)&&(c.seriesIndex=f,a.push(c))}),!0===m&&f++);return a},[]);if(c&&h.length<c)for(var l=h.length;l<=c;l++)h.push({name:l+"\u200b"});c=e(h,m||!1,!0===m?f:1);
b.categories=c.categories;b.treeGrid.mapOfPosToGridNode=c.mapOfPosToGridNode;b.hasNames=!0;b.treeGrid.tree=c.tree;b.series.forEach(function(a){var b=(a.options.data||[]).map(function(b){g(b)&&a.options.keys&&a.options.keys.length&&h.forEach(function(a){0<=b.indexOf(a.x)&&0<=b.indexOf(a.x2)&&(b=a)});return q(b,!0)?K(b):b});a.visible&&a.setData(b,!1)});b.treeGrid.mapOptionsToLevel=k({defaults:d,from:1,levels:d&&d.levels,to:b.treeGrid.tree&&b.treeGrid.tree.height});"beforeRender"===a.type&&(b.treeGrid.collapsedNodes=
c.collapsedNodes)}})}function x(a,b){var c=this.treeGrid.mapOptionsToLevel||{},d=this.ticks,e=d[b],m;if("treegrid"===this.options.type&&this.treeGrid.mapOfPosToGridNode){var f=this.treeGrid.mapOfPosToGridNode[b];(c=c[f.depth])&&(m={labels:c});!e&&l?d[b]=new l(this,b,void 0,void 0,{category:f.name,tickmarkOffset:f.tickmarkOffset,options:m}):(e.parameters.category=f.name,e.options=m,e.addLabel())}else a.apply(this,Array.prototype.slice.call(arguments,1))}function B(a,b,c){var d=this,f="treegrid"===
c.type;d.treeGrid||(d.treeGrid=new H(d));f&&(v(b,"beforeRender",n),v(b,"beforeRedraw",n),v(b,"addSeries",function(a){a.options.data&&(a=e(a.options.data,c.uniqueNames||!1,1),d.treeGrid.collapsedNodes=(d.treeGrid.collapsedNodes||[]).concat(a.collapsedNodes))}),v(d,"foundExtremes",function(){d.treeGrid.collapsedNodes&&d.treeGrid.collapsedNodes.forEach(function(a){var b=d.treeGrid.collapse(a);d.brokenAxis&&(d.brokenAxis.setBreaks(b,!1),d.treeGrid.collapsedNodes&&(d.treeGrid.collapsedNodes=d.treeGrid.collapsedNodes.filter(function(b){return a.collapseStart!==
b.collapseStart||a.collapseEnd!==b.collapseEnd})))})}),v(d,"afterBreaks",function(){"yAxis"===d.coll&&!d.staticScale&&d.chart.options.chart.height&&(d.isDirty=!0)}),c=K({grid:{enabled:!0},labels:{align:"left",levels:[{level:void 0},{level:1,style:{fontWeight:"bold"}}],symbol:{type:"triangle",x:-5,y:-5,height:10,width:10,padding:5}},uniqueNames:!1},c,{reversed:!0,grid:{columns:void 0}}));a.apply(d,[b,c]);f&&(d.hasNames=!0,d.options.showLastLabel=!0)}function G(b){var c=this.options;"treegrid"===c.type?
(this.min=f(this.userMin,c.min,this.dataMin),this.max=f(this.userMax,c.max,this.dataMax),a(this,"foundExtremes"),this.setAxisTranslation(),this.tickmarkOffset=.5,this.tickInterval=1,this.tickPositions=this.treeGrid.mapOfPosToGridNode?this.treeGrid.getTickPositions():[]):b.apply(this,Array.prototype.slice.call(arguments,1))}var l;c.compose=function(a,b,c,e){-1===a.keepProps.indexOf("treeGrid")&&(a.keepProps.push("treeGrid"),l=e,d(a.prototype,"generateTick",x),d(a.prototype,"init",B),d(a.prototype,
"setTickInterval",G),a.prototype.utils={getNode:z.getNode},u.compose(a,b,e),h.compose(a,c),r.compose(e));return a};var H=function(){function a(a){this.axis=a}a.prototype.setCollapsedStatus=function(a){var b=this.axis,c=b.chart;b.series.forEach(function(b){var d=b.options.data;if(a.id&&d){var e=c.get(a.id);b=d[b.data.indexOf(e)];e&&b&&(e.collapsed=a.collapsed,b.collapsed=a.collapsed)}})};a.prototype.collapse=function(a){var c=this.axis,d=c.options.breaks||[],e=b(a,c.max);d.push(e);a.collapsed=!0;c.treeGrid.setCollapsedStatus(a);
return d};a.prototype.expand=function(a){var c=this.axis,d=c.options.breaks||[],e=b(a,c.max);a.collapsed=!1;c.treeGrid.setCollapsedStatus(a);return d.reduce(function(a,b){b.to===e.to&&b.from===e.from||a.push(b);return a},[])};a.prototype.getTickPositions=function(){var a=this.axis,b=Math.floor(a.min/a.tickInterval)*a.tickInterval,c=Math.ceil(a.max/a.tickInterval)*a.tickInterval;return Object.keys(a.treeGrid.mapOfPosToGridNode||{}).reduce(function(d,e){e=+e;!(e>=b&&e<=c)||a.brokenAxis&&a.brokenAxis.isInAnyBreak(e)||
d.push(e);return d},[])};a.prototype.isCollapsed=function(a){var c=this.axis,d=c.options.breaks||[],e=b(a,c.max);return d.some(function(a){return a.from===e.from&&a.to===e.to})};a.prototype.toggleCollapse=function(a){return this.isCollapsed(a)?this.expand(a):this.collapse(a)};return a}();c.Additions=H})(e||(e={}));return e});B(h,"masters/modules/treegrid.src.js",[h["Core/Globals.js"],h["Core/Axis/TreeGridAxis.js"]],function(h,u){u.compose(h.Axis,h.Chart,h.Series,h.Tick)})});
//# sourceMappingURL=treegrid.js.map
```
