## Supplementary material for "Exploring natural odour landscapes: A case study with implications for human-biting insects": Figure S4: treemap.js

```
/*
 Highcharts JS v9.3.1 (2021-11-05)

 (c) 2014-2021 Highsoft AS
 Authors: Jon Arild Nygard / Oystein Moseng

 License: www.highcharts.com/license
*/
'use strict';(function(a){"object"===typeof module&&module.exports?(a["default"]=a,module.exports=a):"function"===typeof define&&define.amd?define("highcharts/modules/treemap",["highcharts"],function(l){a(l);a.Highcharts=l;return a}):a("undefined"!==typeof Highcharts?Highcharts:void 0)})(function(a){function l(a,e,d,t){a.hasOwnProperty(e)||(a[e]=t.apply(null,d))}a=a?a._modules:{};l(a,"Series/ColorMapComposition.js",[a["Core/Utilities.js"]],function(a){var e=a.defined,d=a.wrap,t={pointArrayMap:["value"],
axisTypes:["xAxis","yAxis","colorAxis"],trackerGroups:["group","markerGroup","dataLabelsGroup"],parallelArrays:["x","y","value"],colorKey:"value"},p;(function(n){function a(){return null!==this.value&&Infinity!==this.value&&-Infinity!==this.value}function f(b){var g={};!e(b.color)||b.state&&"normal"!==b.state||(g[this.colorProp||"fill"]=b.color);return g}function r(b,g,a){b=b.call(this,g,a);g.moveToTopOnHover&&(b.zIndex="hover"===a?1:0);return b}n.colorMapSeriesMixin=t;var b=[];n.compose=function(m,
g){g&&-1===b.indexOf(g)&&(b.push(g),g=g.prototype,g.dataLabelOnNull=!0,g.moveToTopOnHover=!0,g.isValid=a);-1===b.indexOf(m)&&(b.push(m),g=m.prototype,g.colorAttribs=f,d(g,"pointAttribs",r));return m};n.seriesColorAttribs=f})(p||(p={}));return p});l(a,"Series/Treemap/TreemapAlgorithmGroup.js",[],function(){return function(){function a(a,d,t,p){this.height=a;this.width=d;this.plot=p;this.startDirection=this.direction=t;this.lH=this.nH=this.lW=this.nW=this.total=0;this.elArr=[];this.lP={total:0,lH:0,
nH:0,lW:0,nW:0,nR:0,lR:0,aspectRatio:function(a,d){return Math.max(a/d,d/a)}}}a.prototype.addElement=function(a){this.lP.total=this.elArr[this.elArr.length-1];this.total+=a;0===this.direction?(this.lW=this.nW,this.lP.lH=this.lP.total/this.lW,this.lP.lR=this.lP.aspectRatio(this.lW,this.lP.lH),this.nW=this.total/this.height,this.lP.nH=this.lP.total/this.nW,this.lP.nR=this.lP.aspectRatio(this.nW,this.lP.nH)):(this.lH=this.nH,this.lP.lW=this.lP.total/this.lH,this.lP.lR=this.lP.aspectRatio(this.lP.lW,
this.lH),this.nH=this.total/this.width,this.lP.nW=this.lP.total/this.nH,this.lP.nR=this.lP.aspectRatio(this.lP.nW,this.nH));this.elArr.push(a)};a.prototype.reset=function(){this.lW=this.nW=0;this.elArr=[];this.total=0};return a}()});l(a,"Series/DrawPointComposition.js",[],function(){var a;(function(a){function d(a){var d=this,f=a.animatableAttribs,n=a.onComplete,b=a.css,m=a.renderer,g=this.series&&this.series.chart.hasRendered?void 0:this.series&&this.series.options.animation,A=this.graphic;a.attribs=
a.attribs||{};a.attribs["class"]=this.getClassName();if(this.shouldDraw())A||(this.graphic=A=m[a.shapeType](a.shapeArgs).add(a.group)),A.css(b).attr(a.attribs).animate(f,a.isNew?!1:g,n);else if(A){var z=function(){d.graphic=A=A&&A.destroy();"function"===typeof n&&n()};Object.keys(f).length?A.animate(f,void 0,function(){z()}):z()}}function e(){return!this.isNull}var p=[];a.compose=function(a){if(-1===p.indexOf(a)){p.push(a);var n=a.prototype;n.draw=d;n.shouldDraw||(n.shouldDraw=e)}return a}})(a||(a=
{}));return a});l(a,"Series/Treemap/TreemapPoint.js",[a["Series/DrawPointComposition.js"],a["Core/Series/SeriesRegistry.js"],a["Core/Utilities.js"]],function(a,e,d){var t=this&&this.__extends||function(){var b=function(a,g){b=Object.setPrototypeOf||{__proto__:[]}instanceof Array&&function(b,a){b.__proto__=a}||function(b,a){for(var g in a)a.hasOwnProperty(g)&&(b[g]=a[g])};return b(a,g)};return function(a,g){function m(){this.constructor=a}b(a,g);a.prototype=null===g?Object.create(g):(m.prototype=g.prototype,
new m)}}(),p=e.series.prototype.pointClass,n=e.seriesTypes;e=n.pie.prototype.pointClass;var q=d.extend,f=d.isNumber,r=d.pick;d=function(a){function b(){var b=null!==a&&a.apply(this,arguments)||this;b.name=void 0;b.node=void 0;b.options=void 0;b.series=void 0;b.value=void 0;return b}t(b,a);b.prototype.getClassName=function(){var b=p.prototype.getClassName.call(this),a=this.series,m=a.options;this.node.level<=a.nodeMap[a.rootNode].level?b+=" highcharts-above-level":this.node.isLeaf||r(m.interactByLeaf,
!m.allowTraversingTree)?this.node.isLeaf||(b+=" highcharts-internal-node"):b+=" highcharts-internal-node-interactive";return b};b.prototype.isValid=function(){return!(!this.id&&!f(this.value))};b.prototype.setState=function(b){p.prototype.setState.call(this,b);this.graphic&&this.graphic.attr({zIndex:"hover"===b?1:0})};b.prototype.shouldDraw=function(){return f(this.plotY)&&null!==this.y};return b}(n.scatter.prototype.pointClass);q(d.prototype,{setVisible:e.prototype.setVisible});a.compose(d);return d});
l(a,"Series/Treemap/TreemapUtilities.js",[a["Core/Utilities.js"]],function(a){var e=a.objectEach,d;(function(a){function d(a,e,f){void 0===f&&(f=this);a=e.call(f,a);!1!==a&&d(a,e,f)}a.AXIS_MAX=100;a.isBoolean=function(a){return"boolean"===typeof a};a.eachObject=function(a,d,f){f=f||this;e(a,function(e,b){d.call(f,e,b,a)})};a.recursive=d})(d||(d={}));return d});l(a,"Series/TreeUtilities.js",[a["Core/Color/Color.js"],a["Core/Utilities.js"]],function(a,e){function d(a,m){var b=m.before,f=m.idRoot,e=
m.mapIdToNode[f],n=m.points[a.i],p=n&&n.options||{},F=[],q=0;a.levelDynamic=a.level-(!1!==m.levelIsConstant?0:e.level);a.name=r(n&&n.name,"");a.visible=f===a.id||!0===m.visible;"function"===typeof b&&(a=b(a,m));a.children.forEach(function(b,f){var g=t({},m);t(g,{index:f,siblings:a.children.length,visible:a.visible});b=d(b,g);F.push(b);b.visible&&(q+=b.val)});b=r(p.value,q);a.visible=0<=b&&(0<q||a.visible);a.children=F;a.childrenTotal=q;a.isLeaf=a.visible&&!q;a.val=b;return a}var t=e.extend,p=e.isArray,
n=e.isNumber,q=e.isObject,f=e.merge,r=e.pick;return{getColor:function(b,m){var d=m.index,f=m.mapOptionsToLevel,e=m.parentColor,n=m.parentColorIndex,q=m.series,p=m.colors,t=m.siblings,l=q.points,E=q.chart.options.chart,x;if(b){l=l[b.i];b=f[b.level]||{};if(f=l&&b.colorByPoint){var G=l.index%(p?p.length:E.colorCount);var C=p&&p[G]}if(!q.chart.styledMode){p=l&&l.options.color;E=b&&b.color;if(x=e)x=(x=b&&b.colorVariation)&&"brightness"===x.key&&d&&t?a.parse(e).brighten(d/t*x.to).get():e;x=r(p,E,C,x,q.color)}var I=
r(l&&l.options.colorIndex,b&&b.colorIndex,G,n,m.colorIndex)}return{color:x,colorIndex:I}},getLevelOptions:function(a){var b=null;if(q(a)){b={};var d=n(a.from)?a.from:1;var e=a.levels;var l={};var t=q(a.defaults)?a.defaults:{};p(e)&&(l=e.reduce(function(a,b){if(q(b)&&n(b.level)){var e=f({},b);var m=r(e.levelIsConstant,t.levelIsConstant);delete e.levelIsConstant;delete e.level;b=b.level+(m?0:d-1);q(a[b])?f(!0,a[b],e):a[b]=e}return a},{}));e=n(a.to)?a.to:1;for(a=0;a<=e;a++)b[a]=f({},t,q(l[a])?l[a]:{})}return b},
setTreeValues:d,updateRootId:function(a){if(q(a)){var b=q(a.options)?a.options:{};b=r(a.rootNode,b.rootId,"");q(a.userOptions)&&(a.userOptions.rootId=b);a.rootNode=b}return b}}});l(a,"Series/Treemap/TreemapComposition.js",[a["Core/Series/SeriesRegistry.js"],a["Series/Treemap/TreemapUtilities.js"],a["Core/Utilities.js"]],function(a,e,d){var l=d.addEvent,p=d.extend,n=!1;l(a.series,"afterBindAxes",function(){var a=this.xAxis,d=this.yAxis;if(a&&d)if(this.is("treemap")){var l={endOnTick:!1,gridLineWidth:0,
lineWidth:0,min:0,minPadding:0,max:e.AXIS_MAX,maxPadding:0,startOnTick:!1,title:void 0,tickPositions:[]};p(d.options,l);p(a.options,l);n=!0}else n&&(d.setOptions(d.userOptions),a.setOptions(a.userOptions),n=!1)})});l(a,"Series/Treemap/TreemapSeries.js",[a["Core/Color/Color.js"],a["Series/ColorMapComposition.js"],a["Core/Globals.js"],a["Core/Legend/LegendSymbol.js"],a["Core/Series/SeriesRegistry.js"],a["Series/Treemap/TreemapAlgorithmGroup.js"],a["Series/Treemap/TreemapPoint.js"],a["Series/Treemap/TreemapUtilities.js"],
a["Series/TreeUtilities.js"],a["Core/Utilities.js"]],function(a,e,d,l,p,n,q,f,r,b){var m=this&&this.__extends||function(){var a=function(b,c){a=Object.setPrototypeOf||{__proto__:[]}instanceof Array&&function(c,a){c.__proto__=a}||function(c,a){for(var h in a)a.hasOwnProperty(h)&&(c[h]=a[h])};return a(b,c)};return function(b,c){function h(){this.constructor=b}a(b,c);b.prototype=null===c?Object.create(c):(h.prototype=c.prototype,new h)}}(),g=a.parse,t=e.colorMapSeriesMixin;a=d.noop;var z=p.series;d=
p.seriesTypes;var E=d.column,M=d.heatmap,F=d.scatter,N=r.getColor,O=r.getLevelOptions,P=r.updateRootId,x=b.addEvent,G=b.correctFloat,C=b.defined,I=b.error,H=b.extend,Q=b.fireEvent,K=b.isArray,R=b.isObject,J=b.isString,D=b.merge,y=b.pick,S=b.stableSort;r=function(a){function b(){var c=null!==a&&a.apply(this,arguments)||this;c.axisRatio=void 0;c.data=void 0;c.mapOptionsToLevel=void 0;c.nodeMap=void 0;c.options=void 0;c.points=void 0;c.rootNode=void 0;c.tree=void 0;return c}m(b,a);b.prototype.algorithmCalcPoints=
function(a,b,k,B){var c,h,u,d,e=k.lW,w=k.lH,f=k.plot,g=0,m=k.elArr.length-1;if(b)e=k.nW,w=k.nH;else var l=k.elArr[k.elArr.length-1];k.elArr.forEach(function(a){if(b||g<m)0===k.direction?(c=f.x,h=f.y,u=e,d=a/u):(c=f.x,h=f.y,d=w,u=a/d),B.push({x:c,y:h,width:u,height:G(d)}),0===k.direction?f.y+=d:f.x+=u;g+=1});k.reset();0===k.direction?k.width-=e:k.height-=w;f.y=f.parent.y+(f.parent.height-k.height);f.x=f.parent.x+(f.parent.width-k.width);a&&(k.direction=1-k.direction);b||k.addElement(l)};b.prototype.algorithmFill=
function(a,b,k){var c=[],h,d=b.direction,u=b.x,f=b.y,e=b.width,w=b.height,g,m,l,n;k.forEach(function(k){h=k.val/b.val*b.height*b.width;g=u;m=f;0===d?(n=w,l=h/n,e-=l,u+=l):(l=e,n=h/l,w-=n,f+=n);c.push({x:g,y:m,width:l,height:n});a&&(d=1-d)});return c};b.prototype.algorithmLowAspectRatio=function(a,b,k){var c=[],h=this,d,f={x:b.x,y:b.y,parent:b},e=0,g=k.length-1,w=new n(b.height,b.width,b.direction,f);k.forEach(function(k){d=k.val/b.val*b.height*b.width;w.addElement(d);w.lP.nR>w.lP.lR&&h.algorithmCalcPoints(a,
!1,w,c,f);e===g&&h.algorithmCalcPoints(a,!0,w,c,f);e+=1});return c};b.prototype.alignDataLabel=function(a,b,k){var c=k.style;c&&!C(c.textOverflow)&&b.text&&b.getBBox().width>b.text.textWidth&&b.css({textOverflow:"ellipsis",width:c.width+="px"});E.prototype.alignDataLabel.apply(this,arguments);a.dataLabel&&a.dataLabel.attr({zIndex:(a.node.zIndex||0)+1})};b.prototype.buildNode=function(a,b,k,d,v){var c=this,h=[],f=c.points[b],e=0,B;(d[a]||[]).forEach(function(b){B=c.buildNode(c.points[b].id,b,k+1,d,
a);e=Math.max(B.height+1,e);h.push(B)});b={id:a,i:b,children:h,height:e,level:k,parent:v,visible:!1};c.nodeMap[b.id]=b;f&&(f.node=b);return b};b.prototype.calculateChildrenAreas=function(a,b){var c=this,h=c.options,v=c.mapOptionsToLevel[a.level+1],d=y(c[v&&v.layoutAlgorithm]&&v.layoutAlgorithm,h.layoutAlgorithm),e=h.alternateStartingDirection,g=[];a=a.children.filter(function(a){return!a.ignore});v&&v.layoutStartingDirection&&(b.direction="vertical"===v.layoutStartingDirection?0:1);g=c[d](b,a);a.forEach(function(a,
h){h=g[h];a.values=D(h,{val:a.childrenTotal,direction:e?1-b.direction:b.direction});a.pointValues=D(h,{x:h.x/c.axisRatio,y:f.AXIS_MAX-h.y-h.height,width:h.width/c.axisRatio});a.children.length&&c.calculateChildrenAreas(a,a.values)})};b.prototype.drawDataLabels=function(){var a=this,b=a.mapOptionsToLevel,k,d;a.points.filter(function(a){return a.node.visible}).forEach(function(c){d=b[c.node.level];k={style:{}};c.node.isLeaf||(k.enabled=!1);d&&d.dataLabels&&(k=D(k,d.dataLabels),a._hasPointLabels=!0);
c.shapeArgs&&(k.style.width=c.shapeArgs.width,c.dataLabel&&c.dataLabel.css({width:c.shapeArgs.width+"px"}));c.dlOptions=D(k,c.options.dataLabels)});z.prototype.drawDataLabels.call(this)};b.prototype.drawPoints=function(){var a=this,b=a.chart,k=b.renderer,d=b.styledMode,e=a.options,f=d?{}:e.shadow,g=e.borderRadius,l=b.pointCount<e.animationLimit,m=e.allowTraversingTree;a.points.forEach(function(c){var b=c.node.levelDynamic,h={},v={},B={},u="level-group-"+c.node.level,L=!!c.graphic,n=l&&L,p=c.shapeArgs;
c.shouldDraw()&&(c.isInside=!0,g&&(v.r=g),D(!0,n?h:v,L?p:{},d?{}:a.pointAttribs(c,c.selected?"select":void 0)),a.colorAttribs&&d&&H(B,a.colorAttribs(c)),a[u]||(a[u]=k.g(u).attr({zIndex:1E3-(b||0)}).add(a.group),a[u].survive=!0));c.draw({animatableAttribs:h,attribs:v,css:B,group:a[u],renderer:k,shadow:f,shapeArgs:p,shapeType:"rect"});m&&c.graphic&&(c.drillId=e.interactByLeaf?a.drillToByLeaf(c):a.drillToByGroup(c))})};b.prototype.drillToByGroup=function(a){var c=!1;1!==a.node.level-this.nodeMap[this.rootNode].level||
a.node.isLeaf||(c=a.id);return c};b.prototype.drillToByLeaf=function(a){var c=!1;if(a.node.parent!==this.rootNode&&a.node.isLeaf)for(a=a.node;!c;)a=this.nodeMap[a.parent],a.parent===this.rootNode&&(c=a.id);return c};b.prototype.drillToNode=function(a,b){I(32,!1,void 0,{"treemap.drillToNode":"use treemap.setRootNode"});this.setRootNode(a,b)};b.prototype.drillUp=function(){var a=this.nodeMap[this.rootNode];a&&J(a.parent)&&this.setRootNode(a.parent,!0,{trigger:"traverseUpButton"})};b.prototype.getExtremes=
function(){var a=z.prototype.getExtremes.call(this,this.colorValueData),b=a.dataMax;this.valueMin=a.dataMin;this.valueMax=b;return z.prototype.getExtremes.call(this)};b.prototype.getListOfParents=function(a,b){a=K(a)?a:[];var c=K(b)?b:[];b=a.reduce(function(a,b,c){b=y(b.parent,"");"undefined"===typeof a[b]&&(a[b]=[]);a[b].push(c);return a},{"":[]});f.eachObject(b,function(a,b,h){""!==b&&-1===c.indexOf(b)&&(a.forEach(function(a){h[""].push(a)}),delete h[b])});return b};b.prototype.getTree=function(){var a=
this.data.map(function(a){return a.id});a=this.getListOfParents(this.data,a);this.nodeMap={};return this.buildNode("",-1,0,a)};b.prototype.hasData=function(){return!!this.processedXData.length};b.prototype.init=function(a,b){t&&(this.colorAttribs=e.seriesColorAttribs);var c=x(this,"setOptions",function(a){a=a.userOptions;C(a.allowDrillToNode)&&!C(a.allowTraversingTree)&&(a.allowTraversingTree=a.allowDrillToNode,delete a.allowDrillToNode);C(a.drillUpButton)&&!C(a.traverseUpButton)&&(a.traverseUpButton=
a.drillUpButton,delete a.drillUpButton)});z.prototype.init.call(this,a,b);delete this.opacity;this.eventsToUnbind.push(c);this.options.allowTraversingTree&&this.eventsToUnbind.push(x(this,"click",this.onClickDrillToNode))};b.prototype.onClickDrillToNode=function(a){var b=(a=a.point)&&a.drillId;J(b)&&(a.setState(""),this.setRootNode(b,!0,{trigger:"click"}))};b.prototype.pointAttribs=function(a,b){var c=R(this.mapOptionsToLevel)?this.mapOptionsToLevel:{},h=a&&c[a.node.level]||{};c=this.options;var d=
b&&c.states[b]||{},e=a&&a.getClassName()||"";a={stroke:a&&a.borderColor||h.borderColor||d.borderColor||c.borderColor,"stroke-width":y(a&&a.borderWidth,h.borderWidth,d.borderWidth,c.borderWidth),dashstyle:a&&a.borderDashStyle||h.borderDashStyle||d.borderDashStyle||c.borderDashStyle,fill:a&&a.color||this.color};-1!==e.indexOf("highcharts-above-level")?(a.fill="none",a["stroke-width"]=0):-1!==e.indexOf("highcharts-internal-node-interactive")?(b=y(d.opacity,c.opacity),a.fill=g(a.fill).setOpacity(b).get(),
a.cursor="pointer"):-1!==e.indexOf("highcharts-internal-node")?a.fill="none":b&&(a.fill=g(a.fill).brighten(d.brightness).get());return a};b.prototype.renderTraverseUpButton=function(a){var b=this,c=b.options.traverseUpButton,d=y(c.text,b.nodeMap[a].name,"\u25c1 Back");if(""===a||b.is("sunburst")&&1===b.tree.children.length&&a===b.tree.children[0].id)b.drillUpButton&&(b.drillUpButton=b.drillUpButton.destroy());else if(this.drillUpButton)this.drillUpButton.placed=!1,this.drillUpButton.attr({text:d}).align();
else{var e=(a=c.theme)&&a.states;this.drillUpButton=this.chart.renderer.button(d,0,0,function(){b.drillUp()},a,e&&e.hover,e&&e.select).addClass("highcharts-drillup-button").attr({align:c.position.align,zIndex:7}).add().align(c.position,!1,c.relativeTo||"plotBox")}};b.prototype.setColorRecursive=function(a,b,d,e,f){var c=this,h=c&&c.chart;h=h&&h.options&&h.options.colors;if(a){var k=N(a,{colors:h,index:e,mapOptionsToLevel:c.mapOptionsToLevel,parentColor:b,parentColorIndex:d,series:c,siblings:f});if(b=
c.points[a.i])b.color=k.color,b.colorIndex=k.colorIndex;(a.children||[]).forEach(function(b,d){c.setColorRecursive(b,k.color,k.colorIndex,d,a.children.length)})}};b.prototype.setPointValues=function(){var a=this,b=a.xAxis,d=a.yAxis,e=a.chart.styledMode;a.points.forEach(function(c){var h=c.node,k=h.pointValues;h=h.visible;if(k&&h){h=k.height;var f=k.width,g=k.x,l=k.y,m=e?0:(a.pointAttribs(c)["stroke-width"]||0)%2/2;k=Math.round(b.toPixels(g,!0))-m;f=Math.round(b.toPixels(g+f,!0))-m;g=Math.round(d.toPixels(l,
!0))-m;h=Math.round(d.toPixels(l+h,!0))-m;h={x:Math.min(k,f),y:Math.min(g,h),width:Math.abs(f-k),height:Math.abs(h-g)};c.plotX=h.x+h.width/2;c.plotY=h.y+h.height/2;c.shapeArgs=h}else delete c.plotX,delete c.plotY})};b.prototype.setRootNode=function(a,b,d){a=H({newRootId:a,previousRootId:this.rootNode,redraw:y(b,!0),series:this},d);Q(this,"setRootNode",a,function(a){var b=a.series;b.idPreviousRoot=a.previousRootId;b.rootNode=a.newRootId;b.isDirty=!0;a.redraw&&b.chart.redraw()})};b.prototype.setState=
function(a){this.options.inactiveOtherPoints=!0;z.prototype.setState.call(this,a,!1);this.options.inactiveOtherPoints=!1};b.prototype.setTreeValues=function(a){var b=this,c=b.options,d=b.nodeMap[b.rootNode];c=f.isBoolean(c.levelIsConstant)?c.levelIsConstant:!0;var e=0,g=[],l=b.points[a.i];a.children.forEach(function(a){a=b.setTreeValues(a);g.push(a);a.ignore||(e+=a.val)});S(g,function(a,b){return(a.sortIndex||0)-(b.sortIndex||0)});var m=y(l&&l.options.value,e);l&&(l.value=m);H(a,{children:g,childrenTotal:e,
ignore:!(y(l&&l.visible,!0)&&0<m),isLeaf:a.visible&&!e,levelDynamic:a.level-(c?0:d.level),name:y(l&&l.name,""),sortIndex:y(l&&l.sortIndex,-m),val:m});return a};b.prototype.sliceAndDice=function(a,b){return this.algorithmFill(!0,a,b)};b.prototype.squarified=function(a,b){return this.algorithmLowAspectRatio(!0,a,b)};b.prototype.strip=function(a,b){return this.algorithmLowAspectRatio(!1,a,b)};b.prototype.stripes=function(a,b){return this.algorithmFill(!1,a,b)};b.prototype.translate=function(){var a=
this,b=a.options,d=P(a);z.prototype.translate.call(a);var e=a.tree=a.getTree();var g=a.nodeMap[d];""===d||g&&g.children.length||(a.setRootNode("",!1),d=a.rootNode,g=a.nodeMap[d]);a.renderTraverseUpButton(d);a.mapOptionsToLevel=O({from:g.level+1,levels:b.levels,to:e.height,defaults:{levelIsConstant:a.options.levelIsConstant,colorByPoint:b.colorByPoint}});f.recursive(a.nodeMap[a.rootNode],function(b){var c=!1,d=b.parent;b.visible=!0;if(d||""===d)c=a.nodeMap[d];return c});f.recursive(a.nodeMap[a.rootNode].children,
function(a){var b=!1;a.forEach(function(a){a.visible=!0;a.children.length&&(b=(b||[]).concat(a.children))});return b});a.setTreeValues(e);a.axisRatio=a.xAxis.len/a.yAxis.len;a.nodeMap[""].pointValues=d={x:0,y:0,width:f.AXIS_MAX,height:f.AXIS_MAX};a.nodeMap[""].values=d=D(d,{width:d.width*a.axisRatio,direction:"vertical"===b.layoutStartingDirection?0:1,val:e.val});a.calculateChildrenAreas(e,d);a.colorAxis||b.colorByPoint||a.setColorRecursive(a.tree);b.allowTraversingTree&&(b=g.pointValues,a.xAxis.setExtremes(b.x,
b.x+b.width,!1),a.yAxis.setExtremes(b.y,b.y+b.height,!1),a.xAxis.setScale(),a.yAxis.setScale());a.setPointValues()};b.defaultOptions=D(F.defaultOptions,{allowTraversingTree:!1,animationLimit:250,borderRadius:0,showInLegend:!1,marker:void 0,colorByPoint:!1,dataLabels:{defer:!1,enabled:!0,formatter:function(){var a=this&&this.point?this.point:{};return J(a.name)?a.name:""},inside:!0,verticalAlign:"middle"},tooltip:{headerFormat:"",pointFormat:"<b>{point.name}</b>: {point.value}<br/>"},ignoreHiddenPoint:!0,
layoutAlgorithm:"sliceAndDice",layoutStartingDirection:"vertical",alternateStartingDirection:!1,levelIsConstant:!0,drillUpButton:{position:{align:"right",x:-10,y:10}},traverseUpButton:{position:{align:"right",x:-10,y:10}},borderColor:"#e6e6e6",borderWidth:1,colorKey:"colorValue",opacity:.15,states:{hover:{borderColor:"#999999",brightness:M?0:.1,halo:!1,opacity:.75,shadow:!1}}});return b}(F);H(r.prototype,{buildKDTree:a,colorKey:"colorValue",directTouch:!0,drawLegendSymbol:l.drawRectangle,getExtremesFromAll:!0,
getSymbol:a,optionalAxis:"colorAxis",parallelArrays:["x","y","value","colorValue"],pointArrayMap:["value"],pointClass:q,trackerGroups:["group","dataLabelsGroup"],utils:{recursive:f.recursive}});e.compose(r);p.registerSeriesType("treemap",r);"";return r});l(a,"masters/modules/treemap.src.js",[],function(){})});
//# sourceMappingURL=treemap.js.map
```
