## Supplementary material for "Exploring natural odour landscapes: A case study with implications for human-biting insects": Figure S4: variable-pie.js

```
/*
 Highcharts JS v9.3.1 (2021-11-05)

 Variable Pie module for Highcharts

 (c) 2010-2021 Grzegorz Blachliski

 License: www.highcharts.com/license
*/
'use strict';(function(a){"object"===typeof module&&module.exports?(a["default"]=a,module.exports=a):"function"===typeof define&&define.amd?define("highcharts/modules/variable-pie",["highcharts"],function(e){a(e);a.Highcharts=e;return a}):a("undefined"!==typeof Highcharts?Highcharts:void 0)})(function(a){function e(a,b,e,m){a.hasOwnProperty(b)||(a[b]=m.apply(null,e))}a=a?a._modules:{};e(a,"Series/VariablePie/VariablePieSeries.js",[a["Core/Series/SeriesRegistry.js"],a["Core/Utilities.js"]],function(a,
b){var e=this&&this.__extends||function(){var a=function(b,c){a=Object.setPrototypeOf||{__proto__:[]}instanceof Array&&function(c,a){c.__proto__=a}||function(c,a){for(var h in a)a.hasOwnProperty(h)&&(c[h]=a[h])};return a(b,c)};return function(b,c){function r(){this.constructor=b}a(b,c);b.prototype=null===c?Object.create(c):(r.prototype=c.prototype,new r)}}(),m=a.seriesTypes.pie,w=b.arrayMax,x=b.arrayMin,z=b.clamp,A=b.extend,B=b.fireEvent,C=b.merge,n=b.pick;b=function(a){function b(){var c=null!==
a&&a.apply(this,arguments)||this;c.data=void 0;c.options=void 0;c.points=void 0;c.radii=void 0;return c}e(b,a);b.prototype.calculateExtremes=function(){var c=this.chart,a=this.options;var b=this.zData;var e=Math.min(c.plotWidth,c.plotHeight)-2*(a.slicedOffset||0),t={};c=this.center||this.getCenter();["minPointSize","maxPointSize"].forEach(function(c){var b=a[c],h=/%$/.test(b);b=parseInt(b,10);t[c]=h?e*b/100:2*b});this.minPxSize=c[3]+t.minPointSize;this.maxPxSize=z(c[2],c[3]+t.minPointSize,t.maxPointSize);
b.length&&(c=n(a.zMin,x(b.filter(this.zValEval))),b=n(a.zMax,w(b.filter(this.zValEval))),this.getRadii(c,b,this.minPxSize,this.maxPxSize))};b.prototype.getRadii=function(c,a,b,e){var h=0,l=this.zData,r=l.length,k=[],m="radius"!==this.options.sizeBy,n=a-c;for(h;h<r;h++){var g=this.zValEval(l[h])?l[h]:c;g<=c?g=b/2:g>=a?g=e/2:(g=0<n?(g-c)/n:.5,m&&(g=Math.sqrt(g)),g=Math.ceil(b+g*(e-b))/2);k.push(g)}this.radii=k};b.prototype.redraw=function(){this.center=null;a.prototype.redraw.apply(this,arguments)};
b.prototype.translate=function(c){this.generatePoints();var b=0,a=this.options,e=a.slicedOffset,m=e+(a.borderWidth||0),l=a.startAngle||0,u=Math.PI/180*(l-90),k=Math.PI/180*(n(a.endAngle,l+360)-90);l=k-u;var y=this.points,w=a.dataLabels.distance;a=a.ignoreHiddenPoint;var g=y.length;this.startAngleRad=u;this.endAngleRad=k;this.calculateExtremes();c||(this.center=c=this.getCenter());for(k=0;k<g;k++){var f=y[k];var p=this.radii[k];f.labelDistance=n(f.options.dataLabels&&f.options.dataLabels.distance,
w);this.maxLabelDistance=Math.max(this.maxLabelDistance||0,f.labelDistance);var d=u+b*l;if(!a||f.visible)b+=f.percentage/100;var q=u+b*l;f.shapeType="arc";f.shapeArgs={x:c[0],y:c[1],r:p,innerR:c[3]/2,start:Math.round(1E3*d)/1E3,end:Math.round(1E3*q)/1E3};d=(q+d)/2;d>1.5*Math.PI?d-=2*Math.PI:d<-Math.PI/2&&(d+=2*Math.PI);f.slicedTranslation={translateX:Math.round(Math.cos(d)*e),translateY:Math.round(Math.sin(d)*e)};var v=Math.cos(d)*c[2]/2;var x=Math.sin(d)*c[2]/2;q=Math.cos(d)*p;p*=Math.sin(d);f.tooltipPos=
[c[0]+.7*v,c[1]+.7*x];f.half=d<-Math.PI/2||d>Math.PI/2?1:0;f.angle=d;v=Math.min(m,f.labelDistance/5);f.labelPosition={natural:{x:c[0]+q+Math.cos(d)*f.labelDistance,y:c[1]+p+Math.sin(d)*f.labelDistance},"final":{},alignment:f.half?"right":"left",connectorPosition:{breakAt:{x:c[0]+q+Math.cos(d)*v,y:c[1]+p+Math.sin(d)*v},touchingSliceAt:{x:c[0]+q,y:c[1]+p}}}}B(this,"afterTranslate")};b.prototype.zValEval=function(a){return"number"!==typeof a||isNaN(a)?null:!0};b.defaultOptions=C(m.defaultOptions,{minPointSize:"10%",
maxPointSize:"100%",zMin:void 0,zMax:void 0,sizeBy:"area",tooltip:{pointFormat:'<span style="color:{point.color}">\u25cf</span> {series.name}<br/>Value: {point.y}<br/>Size: {point.z}<br/>'}});return b}(m);A(b.prototype,{pointArrayMap:["y","z"],parallelArrays:["x","y","z"]});a.registerSeriesType("variablepie",b);"";"";return b});e(a,"masters/modules/variable-pie.src.js",[],function(){})});
//# sourceMappingURL=variable-pie.js.map
```
