## Supplementary material for "Exploring natural odour landscapes: A case study with implications for human-biting insects": Figure S4: variwide.js

```
/*
 Highcharts JS v9.3.1 (2021-11-05)

 Highcharts variwide module

 (c) 2010-2021 Torstein Honsi

 License: www.highcharts.com/license
*/
'use strict';(function(a){"object"===typeof module&&module.exports?(a["default"]=a,module.exports=a):"function"===typeof define&&define.amd?define("highcharts/modules/variwide",["highcharts"],function(g){a(g);a.Highcharts=g;return a}):a("undefined"!==typeof Highcharts?Highcharts:void 0)})(function(a){function g(a,h,c,f){a.hasOwnProperty(h)||(a[h]=f.apply(null,c))}a=a?a._modules:{};g(a,"Series/Variwide/VariwidePoint.js",[a["Core/Series/SeriesRegistry.js"],a["Core/Utilities.js"]],function(a,h){var c=
this&&this.__extends||function(){var a=function(b,d){a=Object.setPrototypeOf||{__proto__:[]}instanceof Array&&function(a,b){a.__proto__=b}||function(a,b){for(var d in b)b.hasOwnProperty(d)&&(a[d]=b[d])};return a(b,d)};return function(b,d){function c(){this.constructor=b}a(b,d);b.prototype=null===d?Object.create(d):(c.prototype=d.prototype,new c)}}(),f=h.isNumber;return function(a){function b(){var b=null!==a&&a.apply(this,arguments)||this;b.crosshairWidth=void 0;b.options=void 0;b.series=void 0;return b}
c(b,a);b.prototype.isValid=function(){return f(this.y)&&f(this.z)};return b}(a.seriesTypes.column.prototype.pointClass)});g(a,"Series/Variwide/VariwideComposition.js",[a["Core/Axis/Tick.js"],a["Core/Axis/Axis.js"],a["Core/Utilities.js"]],function(a,h,c){var f=c.addEvent;c=c.wrap;a.prototype.postTranslate=function(a,b,c){var d=this.axis,e=a[b]-d.pos;d.horiz||(e=d.len-e);e=d.series[0].postTranslate(c,e);d.horiz||(e=d.len-e);a[b]=d.pos+e};f(h,"afterDrawCrosshair",function(a){this.variwide&&this.cross&&
this.cross.attr("stroke-width",a.point&&a.point.crosshairWidth)});f(h,"afterRender",function(){var a=this;!this.horiz&&this.variwide&&this.chart.labelCollectors.push(function(){return a.tickPositions.filter(function(b){return a.ticks[b].label}).map(function(b,d){b=a.ticks[b].label;b.labelrank=a.zData[d];return b})})});f(a,"afterGetPosition",function(a){var b=this.axis,d=b.horiz?"x":"y";b.variwide&&(this[d+"Orig"]=a.pos[d],this.postTranslate(a.pos,d,this.pos))});c(a.prototype,"getLabelPosition",function(a,
b,c,f,e,r,k,t){var d=Array.prototype.slice.call(arguments,1),p=e?"x":"y";this.axis.variwide&&"number"===typeof this[p+"Orig"]&&(d[e?0:1]=this[p+"Orig"]);d=a.apply(this,d);this.axis.variwide&&this.axis.categories&&this.postTranslate(d,p,this.pos);return d})});g(a,"Series/Variwide/VariwideSeries.js",[a["Core/Series/SeriesRegistry.js"],a["Series/Variwide/VariwidePoint.js"],a["Core/Utilities.js"]],function(a,h,c){var f=this&&this.__extends||function(){var a=function(b,d){a=Object.setPrototypeOf||{__proto__:[]}instanceof
Array&&function(a,b){a.__proto__=b}||function(a,b){for(var d in b)b.hasOwnProperty(d)&&(a[d]=b[d])};return a(b,d)};return function(b,d){function k(){this.constructor=b}a(b,d);b.prototype=null===d?Object.create(d):(k.prototype=d.prototype,new k)}}(),d=a.seriesTypes.column,b=c.extend,g=c.merge,q=c.pick;c=function(b){function c(){var a=null!==b&&b.apply(this,arguments)||this;a.data=void 0;a.options=void 0;a.points=void 0;a.relZ=void 0;a.totalZ=void 0;a.zData=void 0;return a}f(c,b);c.prototype.processData=
function(b){this.totalZ=0;this.relZ=[];a.seriesTypes.column.prototype.processData.call(this,b);(this.xAxis.reversed?this.zData.slice().reverse():this.zData).forEach(function(a,b){this.relZ[b]=this.totalZ;this.totalZ+=a},this);this.xAxis.categories&&(this.xAxis.variwide=!0,this.xAxis.zData=this.zData)};c.prototype.postTranslate=function(a,b,d){var c=this.xAxis,l=this.relZ;a=c.reversed?l.length-a:a;var n=c.reversed?-1:1,m=c.toPixels(c.reversed?(c.dataMax||0)+c.pointRange:c.dataMin||0),f=c.toPixels(c.reversed?
c.dataMin||0:(c.dataMax||0)+c.pointRange),k=Math.abs(f-m),e=this.totalZ;c=this.chart.inverted?f-(this.chart.plotTop-n*c.minPixelPadding):m-this.chart.plotLeft-n*c.minPixelPadding;m=a/l.length*k;f=(a+n)/l.length*k;var g=q(l[a],e)/e*k;l=q(l[a+n],e)/e*k;d&&(d.crosshairWidth=l-g);return c+g+(b-(c+m))*(l-g)/(f-m)};c.prototype.translate=function(){var b=this.options.crisp,c=this.xAxis;this.options.crisp=!1;a.seriesTypes.column.prototype.translate.call(this);this.options.crisp=b;var d=this.chart.inverted,
f=this.borderWidth%2/2;this.points.forEach(function(a,b){if(c.variwide){var e=this.postTranslate(b,a.shapeArgs.x,a);b=this.postTranslate(b,a.shapeArgs.x+a.shapeArgs.width)}else e=a.plotX,b=c.translate(a.x+a.z,0,0,0,1);this.options.crisp&&(e=Math.round(e)-f,b=Math.round(b)-f);a.shapeArgs.x=e;a.shapeArgs.width=Math.max(b-e,1);a.plotX=(e+b)/2;d?a.tooltipPos[1]=c.len-a.shapeArgs.x-a.shapeArgs.width/2:a.tooltipPos[0]=a.shapeArgs.x+a.shapeArgs.width/2},this);this.options.stacking&&this.correctStackLabels()};
c.prototype.correctStackLabels=function(){var a=this,b=a.options,c=a.yAxis,d,e,f,g;a.points.forEach(function(h){g=h.x;e=h.shapeArgs.width;(f=c.stacking.stacks[(a.negStacks&&h.y<(b.startFromThreshold?0:b.threshold)?"-":"")+a.stackKey])&&(d=f[g])&&!h.isNull&&d.setOffset(-(e/2)||0,e||0,void 0,void 0,h.plotX)})};c.defaultOptions=g(d.defaultOptions,{pointPadding:0,groupPadding:0});return c}(d);b(c.prototype,{irregularWidths:!0,pointArrayMap:["y","z"],parallelArrays:["x","y","z"],pointClass:h});a.registerSeriesType("variwide",
c);"";return c});g(a,"masters/modules/variwide.src.js",[],function(){})});
//# sourceMappingURL=variwide.js.map
```
