## Supplementary material for "Exploring natural odour landscapes: A case study with implications for human-biting insects": Figure S4: vector.js

```
/*
 Highcharts JS v9.3.1 (2021-11-05)

 Vector plot series module

 (c) 2010-2021 Torstein Honsi

 License: www.highcharts.com/license
*/
'use strict';(function(a){"object"===typeof module&&module.exports?(a["default"]=a,module.exports=a):"function"===typeof define&&define.amd?define("highcharts/modules/vector",["highcharts"],function(d){a(d);a.Highcharts=d;return a}):a("undefined"!==typeof Highcharts?Highcharts:void 0)})(function(a){function d(a,g,d,e){a.hasOwnProperty(g)||(a[g]=e.apply(null,d))}a=a?a._modules:{};d(a,"Series/Vector/VectorSeries.js",[a["Core/Animation/AnimationUtilities.js"],a["Core/Globals.js"],a["Core/Series/SeriesRegistry.js"],
a["Core/Utilities.js"]],function(a,d,h,e){var g=this&&this.__extends||function(){var a=function(c,b){a=Object.setPrototypeOf||{__proto__:[]}instanceof Array&&function(b,a){b.__proto__=a}||function(b,a){for(var f in a)a.hasOwnProperty(f)&&(b[f]=a[f])};return a(c,b)};return function(c,b){function f(){this.constructor=c}a(c,b);c.prototype=null===b?Object.create(b):(f.prototype=b.prototype,new f)}}(),l=a.animObject,m=h.series,k=h.seriesTypes.scatter,n=e.arrayMax;a=e.extend;var p=e.merge,q=e.pick;e=function(a){function c(){var b=
null!==a&&a.apply(this,arguments)||this;b.data=void 0;b.lengthMax=void 0;b.options=void 0;b.points=void 0;return b}g(c,a);c.prototype.animate=function(b){b?this.markerGroup.attr({opacity:.01}):this.markerGroup.animate({opacity:1},l(this.options.animation))};c.prototype.arrow=function(b){b=b.length/this.lengthMax*this.options.vectorLength/20;var a={start:10*b,center:0,end:-10*b}[this.options.rotationOrigin]||0;return[["M",0,7*b+a],["L",-1.5*b,7*b+a],["L",0,10*b+a],["L",1.5*b,7*b+a],["L",0,7*b+a],["L",
0,-10*b+a]]};c.prototype.drawPoints=function(){var a=this.chart;this.points.forEach(function(b){var c=b.plotX,d=b.plotY;!1===this.options.clip||a.isInsidePlot(c,d,{inverted:a.inverted})?(b.graphic||(b.graphic=this.chart.renderer.path().add(this.markerGroup).addClass("highcharts-point highcharts-color-"+q(b.colorIndex,b.series.colorIndex))),b.graphic.attr({d:this.arrow(b),translateX:c,translateY:d,rotation:b.direction}),this.chart.styledMode||b.graphic.attr(this.pointAttribs(b))):b.graphic&&(b.graphic=
b.graphic.destroy())},this)};c.prototype.pointAttribs=function(b,a){var c=this.options;b=b.color||this.color;var d=this.options.lineWidth;a&&(b=c.states[a].color||b,d=(c.states[a].lineWidth||d)+(c.states[a].lineWidthPlus||0));return{stroke:b,"stroke-width":d}};c.prototype.translate=function(){m.prototype.translate.call(this);this.lengthMax=n(this.lengthData)};c.defaultOptions=p(k.defaultOptions,{lineWidth:2,marker:null,rotationOrigin:"center",states:{hover:{lineWidthPlus:1}},tooltip:{pointFormat:"<b>[{point.x}, {point.y}]</b><br/>Length: <b>{point.length}</b><br/>Direction: <b>{point.direction}\u00b0</b><br/>"},
vectorLength:20});return c}(k);a(e.prototype,{drawGraph:d.noop,getSymbol:d.noop,markerAttribs:d.noop,parallelArrays:["x","y","length","direction"],pointArrayMap:["y","length","direction"]});h.registerSeriesType("vector",e);"";return e});d(a,"masters/modules/vector.src.js",[],function(){})});
//# sourceMappingURL=vector.js.map
```
