## Supplementary material for "Exploring natural odour landscapes: A case study with implications for human-biting insects": Figure S4: venn.js

```
/*
 Highcharts JS v9.3.1 (2021-11-05)

 (c) 2017-2021 Highsoft AS
 Authors: Jon Arild Nygard

 License: www.highcharts.com/license
*/
'use strict';(function(a){"object"===typeof module&&module.exports?(a["default"]=a,module.exports=a):"function"===typeof define&&define.amd?define("highcharts/modules/venn",["highcharts"],function(n){a(n);a.Highcharts=n;return a}):a("undefined"!==typeof Highcharts?Highcharts:void 0)})(function(a){function n(a,r,m,y){a.hasOwnProperty(r)||(a[r]=y.apply(null,m))}a=a?a._modules:{};n(a,"Core/Geometry/GeometryUtilities.js",[],function(){var a;(function(a){a.getCenterOfPoints=function(a){var m=a.reduce(function(a,
h){a.x+=h.x;a.y+=h.y;return a},{x:0,y:0});return{x:m.x/a.length,y:m.y/a.length}};a.getDistanceBetweenPoints=function(a,g){return Math.sqrt(Math.pow(g.x-a.x,2)+Math.pow(g.y-a.y,2))};a.getAngleBetweenPoints=function(a,g){return Math.atan2(g.x-a.x,g.y-a.y)}})(a||(a={}));return a});n(a,"Core/Geometry/CircleUtilities.js",[a["Core/Geometry/GeometryUtilities.js"]],function(a){var g=a.getAngleBetweenPoints,m=a.getCenterOfPoints,y=a.getDistanceBetweenPoints,q;(function(a){function k(a,d){d=Math.pow(10,d);
return Math.round(a*d)/d}function h(a){if(0>=a)throw Error("radius of circle must be a positive number.");return Math.PI*a*a}function x(a,d){return a*a*Math.acos(1-d/a)-(a-d)*Math.sqrt(d*(2*a-d))}function B(a,d){var h=y(a,d),v=a.r,t=d.r,f=[];if(h<v+t&&h>Math.abs(v-t)){v*=v;var u=(v-t*t+h*h)/(2*h);t=Math.sqrt(v-u*u);v=a.x;f=d.x;a=a.y;var g=d.y;d=v+u*(f-v)/h;u=a+u*(g-a)/h;a=t/h*-(g-a);h=t/h*-(f-v);f=[{x:k(d+a,14),y:k(u-h,14)},{x:k(d-a,14),y:k(u+h,14)}]}return f}function r(a){return a.reduce(function(a,
h,k,t){t=t.slice(k+1).reduce(function(a,d,t,f){var g=[k,t+k+1];return a.concat(B(h,d).map(function(a){a.indexes=g;return a}))},[]);return a.concat(t)},[])}function q(a,d){return y(a,d)<=d.r+1e-10}function w(a,d){return!d.some(function(d){return!q(a,d)})}function n(a){return r(a).filter(function(d){return w(d,a)})}a.round=k;a.getAreaOfCircle=h;a.getCircularSegmentArea=x;a.getOverlapBetweenCircles=function(a,d,f){var g=0;f<a+d&&(f<=Math.abs(d-a)?g=h(a<d?a:d):(g=(a*a-d*d+f*f)/(2*f),f-=g,g=x(a,a-g)+x(d,
d-f)),g=k(g,14));return g};a.getCircleCircleIntersection=B;a.getCirclesIntersectionPoints=r;a.isCircle1CompletelyOverlappingCircle2=function(a,d){return y(a,d)+d.r<a.r+1e-10};a.isPointInsideCircle=q;a.isPointInsideAllCircles=w;a.isPointOutsideAllCircles=function(a,d){return!d.some(function(d){return q(a,d)})};a.getCirclesIntersectionPolygon=n;a.getAreaOfIntersectionBetweenCircles=function(a){var d=n(a);if(1<d.length){var h=m(d);d=d.map(function(a){a.angle=g(h,a);return a}).sort(function(a,d){return d.angle-
a.angle});var f=d[d.length-1];d=d.reduce(function(d,h){var f=d.startPoint,k=m([f,h]),x=h.indexes.filter(function(b){return-1<f.indexes.indexOf(b)}).reduce(function(b,c){c=a[c];var e=g(c,h),l=g(c,f);e=l-(l-e+(l<e?2*Math.PI:0))/2;e=y(k,{x:c.x+c.r*Math.sin(e),y:c.y+c.r*Math.cos(e)});c=c.r;e>2*c&&(e=2*c);if(!b||b.width>e)b={r:c,largeArc:e>c?1:0,width:e,x:h.x,y:h.y};return b},null);if(x){var p=x.r;d.arcs.push(["A",p,p,0,x.largeArc,1,x.x,x.y]);d.startPoint=h}return d},{startPoint:f,arcs:[]}).arcs;if(0!==
d.length&&1!==d.length){d.unshift(["M",f.x,f.y]);var k={center:h,d:d}}}return k}})(q||(q={}));return q});n(a,"Series/DrawPointComposition.js",[],function(){var a;(function(a){function g(a){var h=this,f=a.animatableAttribs,g=a.onComplete,m=a.css,q=a.renderer,r=this.series&&this.series.chart.hasRendered?void 0:this.series&&this.series.options.animation,w=this.graphic;a.attribs=a.attribs||{};a.attribs["class"]=this.getClassName();if(this.shouldDraw())w||(this.graphic=w=q[a.shapeType](a.shapeArgs).add(a.group)),
w.css(m).attr(a.attribs).animate(f,a.isNew?!1:r,g);else if(w){var n=function(){h.graphic=w=w&&w.destroy();"function"===typeof g&&g()};Object.keys(f).length?w.animate(f,void 0,function(){n()}):n()}}function r(){return!this.isNull}var q=[];a.compose=function(a){if(-1===q.indexOf(a)){q.push(a);var h=a.prototype;h.draw=g;h.shouldDraw||(h.shouldDraw=r)}return a}})(a||(a={}));return a});n(a,"Series/Venn/VennPoint.js",[a["Series/DrawPointComposition.js"],a["Core/Series/SeriesRegistry.js"],a["Core/Utilities.js"]],
function(a,r,m){var g=this&&this.__extends||function(){var a=function(h,f){a=Object.setPrototypeOf||{__proto__:[]}instanceof Array&&function(a,f){a.__proto__=f}||function(a,f){for(var h in f)f.hasOwnProperty(h)&&(a[h]=f[h])};return a(h,f)};return function(h,f){function g(){this.constructor=h}a(h,f);h.prototype=null===f?Object.create(f):(g.prototype=f.prototype,new g)}}(),q=m.isNumber;r=function(a){function h(){var f=null!==a&&a.apply(this,arguments)||this;f.options=void 0;f.series=void 0;return f}
g(h,a);h.prototype.isValid=function(){return q(this.value)};h.prototype.shouldDraw=function(){return!!this.shapeArgs};return h}(r.seriesTypes.scatter.prototype.pointClass);a.compose(r);return r});n(a,"Series/Venn/VennUtils.js",[a["Core/Geometry/CircleUtilities.js"],a["Core/Geometry/GeometryUtilities.js"],a["Core/Utilities.js"]],function(a,r,m){function g(a){var b=a.filter(function(a){return 2===a.sets.length}).reduce(function(a,b){b.sets.forEach(function(e,c,p){A(a[e])||(a[e]={overlapping:{},totalOverlap:0});
a[e].totalOverlap+=b.value;a[e].overlapping[p[1-c]]=b.value});return a},{});a.filter(n).forEach(function(a){C(a,b[a.sets[0]])});return a}function q(a,b,c,e,l){var p=a(b),d=a(c);l=l||100;e=e||1e-10;var f=c-b,h=1;if(b>=c)throw Error("a must be smaller than b.");if(0<p*d)throw Error("f(a) and f(b) must have opposite signs.");if(0===p)var g=b;else if(0===d)g=c;else for(;h++<=l&&0!==k&&f>e;){f=(c-b)/2;g=b+f;var k=a(g);0<p*k?b=g:c=g}return g}function h(a){a=a.slice(0,-1);for(var b=a.length,c=[],e=function(a,
b){a.sum+=b[a.i];return a},l=0;l<b;l++)c[l]=a.reduce(e,{sum:0,i:l}).sum/b;return c}function k(a,b,c){var e=a+b;return 0>=c?e:w(a<b?a:b)<=c?0:q(function(e){e=t(a,b,e);return c-e},0,e)}function f(a){var b=0;2===a.length&&(b=a[0],a=a[1],b=t(b.r,a.r,z(b,a)));return b}function n(a){return u(a.sets)&&1===a.sets.length}function B(a){var b={};return A(a)&&G(a.value)&&-1<a.value&&u(a.sets)&&0<a.sets.length&&!a.sets.some(function(a){var e=!1;!b[a]&&K(a)?b[a]=!0:e=!0;return e})}function D(a,b){return b.reduce(function(b,
e){var c=0;1<e.sets.length&&(c=e.value,e=f(e.sets.map(function(b){return a[b]})),e=c-e,c=Math.round(e*e*1E11)/1E11);return b+c},0)}function E(a,b){return b.totalOverlap-a.totalOverlap}var w=a.getAreaOfCircle,H=a.getCircleCircleIntersection,t=a.getOverlapBetweenCircles,d=a.isPointInsideAllCircles,F=a.isPointInsideCircle,v=a.isPointOutsideAllCircles,z=r.getDistanceBetweenPoints,C=m.extend,u=m.isArray,G=m.isNumber,A=m.isObject,K=m.isString;return{geometry:r,geometryCircles:a,addOverlapToSets:g,getCentroid:h,
getDistanceBetweenCirclesByOverlap:k,getLabelWidth:function(a,b,c){var e=b.reduce(function(a,b){return Math.min(b.r,a)},Infinity),l=c.filter(function(b){return!F(a,b)});c=function(e,c){return q(function(f){var h={x:a.x+c*f,y:a.y};h=d(h,b)&&v(h,l);return-(e-f)+(h?0:Number.MAX_VALUE)},0,e)};return 2*Math.min(c(e,-1),c(e,1))},getMarginFromCircles:function(a,b,c){b=b.reduce(function(b,c){c=c.r-z(a,c);return c<=b?c:b},Number.MAX_VALUE);return b=c.reduce(function(b,c){c=z(a,c)-c.r;return c<=b?c:b},b)},
isSet:n,layoutGreedyVenn:function(a){var b=[],c={};a.filter(function(a){return 1===a.sets.length}).forEach(function(a){c[a.sets[0]]=a.circle={x:Number.MAX_VALUE,y:Number.MAX_VALUE,r:Math.sqrt(a.value/Math.PI)}});var e=function(a,e){var c=a.circle;c.x=e.x;c.y=e.y;b.push(a)};g(a);var l=a.filter(n).sort(E);e(l.shift(),{x:0,y:0});var d=a.filter(function(a){return 2===a.sets.length});l.forEach(function(a){var l=a.circle,f=l.r,h=a.overlapping,I=b.reduce(function(a,e,I){var g=e.circle,p=k(f,g.r,h[e.sets[0]]),
J=[{x:g.x+p,y:g.y},{x:g.x-p,y:g.y},{x:g.x,y:g.y+p},{x:g.x,y:g.y-p}];b.slice(I+1).forEach(function(a){var b=a.circle;a=k(f,b.r,h[a.sets[0]]);J=J.concat(H({x:g.x,y:g.y,r:p},{x:b.x,y:b.y,r:a}))});J.forEach(function(b){l.x=b.x;l.y=b.y;var e=D(c,d);e<a.loss&&(a.loss=e,a.coordinates=b)});return a},{loss:Number.MAX_VALUE,coordinates:void 0});e(a,I.coordinates)});return c},loss:D,nelderMead:function(a,b){var c=function(a,b){return a.fx-b.fx},e=function(a,b,e,c){return b.map(function(b,l){return a*b+e*c[l]})},
l=function(b,e){e.fx=a(e);b[b.length-1]=e;return b},d=function(b){var c=b[0];return b.map(function(b){b=e(.5,c,.5,b);b.fx=a(b);return b})},f=function(b,c,l,d){b=e(l,b,d,c);b.fx=a(b);return b};b=function(b){var e=b.length,c=Array(e+1);c[0]=b;c[0].fx=a(b);for(var l=0;l<e;++l){var d=b.slice();d[l]=d[l]?1.05*d[l]:.001;d.fx=a(d);c[l+1]=d}return c}(b);for(var g=0;100>g;g++){b.sort(c);var p=b[b.length-1],k=h(b),m=f(k,p,2,-1);if(m.fx<b[0].fx)p=f(k,p,3,-2),b=l(b,p.fx<m.fx?p:m);else if(m.fx>=b[b.length-2].fx){var n=
void 0;m.fx>p.fx?(n=f(k,p,.5,.5),b=n.fx<p.fx?l(b,n):d(b)):(n=f(k,p,1.5,-.5),b=n.fx<m.fx?l(b,n):d(b))}else b=l(b,m)}return b[0]},processVennData:function(a){a=u(a)?a:[];var b=a.reduce(function(a,b){B(b)&&n(b)&&0<b.value&&-1===a.indexOf(b.sets[0])&&a.push(b.sets[0]);return a},[]).sort(),c=a.reduce(function(a,c){B(c)&&!c.sets.some(function(a){return-1===b.indexOf(a)})&&(a[c.sets.sort().join()]=c);return a},{});b.reduce(function(a,b,c,d){d.slice(c+1).forEach(function(c){a.push(b+","+c)});return a},[]).forEach(function(a){if(!c[a]){var b=
{sets:a.split(","),value:0};c[a]=b}});return Object.keys(c).map(function(a){return c[a]})},sortByTotalOverlap:E}});n(a,"Series/Venn/VennSeries.js",[a["Core/Animation/AnimationUtilities.js"],a["Core/Color/Color.js"],a["Core/Geometry/CircleUtilities.js"],a["Core/Geometry/GeometryUtilities.js"],a["Core/Series/SeriesRegistry.js"],a["Series/Venn/VennPoint.js"],a["Series/Venn/VennUtils.js"],a["Core/Utilities.js"]],function(a,n,m,y,q,h,k,f){var g=this&&this.__extends||function(){var a=function(d,b){a=Object.setPrototypeOf||
{__proto__:[]}instanceof Array&&function(a,b){a.__proto__=b}||function(a,b){for(var c in b)b.hasOwnProperty(c)&&(a[c]=b[c])};return a(d,b)};return function(d,b){function c(){this.constructor=d}a(d,b);d.prototype=null===b?Object.create(b):(c.prototype=b.prototype,new c)}}(),r=a.animObject,D=n.parse,E=m.getAreaOfIntersectionBetweenCircles,w=m.getCirclesIntersectionPolygon,H=m.isCircle1CompletelyOverlappingCircle2,t=m.isPointInsideAllCircles,d=m.isPointOutsideAllCircles,F=y.getCenterOfPoints,v=q.seriesTypes.scatter;
a=f.addEvent;var z=f.extend,C=f.isArray,u=f.isNumber,G=f.isObject,A=f.merge;f=function(a){function f(){var b=null!==a&&a.apply(this,arguments)||this;b.data=void 0;b.mapOfIdToRelation=void 0;b.options=void 0;b.points=void 0;return b}g(f,a);f.getLabelPosition=function(a,c){var b=a.reduce(function(b,e){var d=e.r/2;return[{x:e.x,y:e.y},{x:e.x+d,y:e.y},{x:e.x-d,y:e.y},{x:e.x,y:e.y+d},{x:e.x,y:e.y-d}].reduce(function(b,e){var d=k.getMarginFromCircles(e,a,c);b.margin<d&&(b.point=e,b.margin=d);return b},
b)},{point:void 0,margin:-Number.MAX_VALUE}).point;b=k.nelderMead(function(b){return-k.getMarginFromCircles({x:b[0],y:b[1]},a,c)},[b.x,b.y]);b={x:b[0],y:b[1]};t(b,a)&&d(b,c)||(b=1<a.length?F(w(a)):{x:a[0].x,y:a[0].y});return b};f.getLabelValues=function(a,c){var b=a.sets,d=c.reduce(function(a,c){var e=-1<b.indexOf(c.sets[0]);a[e?"internal":"external"].push(c.circle);return a},{internal:[],external:[]});d.external=d.external.filter(function(a){return d.internal.some(function(b){return!H(a,b)})});a=
f.getLabelPosition(d.internal,d.external);c=k.getLabelWidth(a,d.internal,d.external);return{position:a,width:c}};f.layout=function(a){var b={},e={};if(0<a.length){var d=k.layoutGreedyVenn(a),h=a.filter(k.isSet);a.forEach(function(a){var c=a.sets,g=c.join();if(c=k.isSet(a)?d[g]:E(c.map(function(a){return d[a]})))b[g]=c,e[g]=f.getLabelValues(a,h)})}return{mapOfIdToShape:b,mapOfIdToLabelValues:e}};f.getScale=function(a,c,e){var b=e.bottom-e.top,d=e.right-e.left;b=Math.min(0<d?1/d*a:1,0<b?1/b*c:1);return{scale:b,
centerX:a/2-(e.right+e.left)/2*b,centerY:c/2-(e.top+e.bottom)/2*b}};f.updateFieldBoundaries=function(a,c){var b=c.x-c.r,d=c.x+c.r,f=c.y+c.r;c=c.y-c.r;if(!u(a.left)||a.left>b)a.left=b;if(!u(a.right)||a.right<d)a.right=d;if(!u(a.top)||a.top>c)a.top=c;if(!u(a.bottom)||a.bottom<f)a.bottom=f;return a};f.prototype.animate=function(a){if(!a){var b=r(this.options.animation);this.points.forEach(function(a){var c=a.shapeArgs;if(a.graphic&&c){var d={},e={};c.d?d.opacity=.001:(d.r=0,e.r=c.r);a.graphic.attr(d).animate(e,
b);c.d&&setTimeout(function(){a&&a.graphic&&a.graphic.animate({opacity:1})},b.duration)}},this)}};f.prototype.drawPoints=function(){var a=this,c=a.chart,d=a.group,f=c.renderer;(a.points||[]).forEach(function(b){var e={zIndex:C(b.sets)?b.sets.length:0},h=b.shapeArgs;c.styledMode||z(e,a.pointAttribs(b,b.state));b.draw({isNew:!b.graphic,animatableAttribs:h,attribs:e,group:d,renderer:f,shapeType:h&&h.d?"path":"circle"})})};f.prototype.init=function(){v.prototype.init.apply(this,arguments);delete this.opacity};
f.prototype.pointAttribs=function(a,c){var b=this.options||{};a=A(b,{color:a&&a.color},a&&a.options||{},c&&b.states[c]||{});return{fill:D(a.color).brighten(a.brightness).get(),opacity:a.opacity,stroke:a.borderColor,"stroke-width":a.borderWidth,dashstyle:a.borderDashStyle}};f.prototype.translate=function(){var a=this.chart;this.processedXData=this.xData;this.generatePoints();var c=k.processVennData(this.options.data);c=f.layout(c);var d=c.mapOfIdToShape,h=c.mapOfIdToLabelValues;c=Object.keys(d).filter(function(a){return(a=
d[a])&&u(a.r)}).reduce(function(a,b){return f.updateFieldBoundaries(a,d[b])},{top:0,bottom:0,left:0,right:0});a=f.getScale(a.plotWidth,a.plotHeight,c);var g=a.scale,m=a.centerX,n=a.centerY;this.points.forEach(function(a){var b=C(a.sets)?a.sets:[],c=b.join(),e=d[c],f=h[c]||{};c=f.width;f=f.position;var l=a.options&&a.options.dataLabels;if(e){if(e.r)var k={x:m+e.x*g,y:n+e.y*g,r:e.r*g};else e.d&&(e=e.d,e.forEach(function(a){"M"===a[0]?(a[1]=m+a[1]*g,a[2]=n+a[2]*g):"A"===a[0]&&(a[1]*=g,a[2]*=g,a[6]=m+
a[6]*g,a[7]=n+a[7]*g)}),k={d:e});f?(f.x=m+f.x*g,f.y=n+f.y*g):f={};u(c)&&(c=Math.round(c*g))}a.shapeArgs=k;f&&k&&(a.plotX=f.x,a.plotY=f.y);c&&k&&(a.dlOptions=A(!0,{style:{width:c}},G(l,!0)?l:void 0));a.name=a.options.name||b.join("\u2229")})};f.defaultOptions=A(v.defaultOptions,{borderColor:"#cccccc",borderDashStyle:"solid",borderWidth:1,brighten:0,clip:!1,colorByPoint:!0,dataLabels:{enabled:!0,verticalAlign:"middle",formatter:function(){return this.point.name}},inactiveOtherPoints:!0,marker:!1,opacity:.75,
showInLegend:!1,states:{hover:{opacity:1,borderColor:"#333333"},select:{color:"#cccccc",borderColor:"#000000",animation:!1},inactive:{opacity:.075}},tooltip:{pointFormat:"{point.name}: {point.value}"}});return f}(v);z(f.prototype,{axisTypes:[],directTouch:!0,isCartesian:!1,pointArrayMap:["value"],pointClass:h,utils:k});q.registerSeriesType("venn",f);"";a(f,"afterSetOptions",function(a){var d=a.options.states;this.is("venn")&&Object.keys(d).forEach(function(a){d[a].halo=!1})});return f});n(a,"masters/modules/venn.src.js",
[],function(){})});
//# sourceMappingURL=venn.js.map
```
