## Supplementary material for "Exploring natural odour landscapes: A case study with implications for human-biting insects": Figure S4: windbarb.js

```
/*
 Highcharts JS v9.3.1 (2021-11-05)

 Wind barb series module

 (c) 2010-2021 Torstein Honsi

 License: www.highcharts.com/license
*/
'use strict';(function(a){"object"===typeof module&&module.exports?(a["default"]=a,module.exports=a):"function"===typeof define&&define.amd?define("highcharts/modules/windbarb",["highcharts"],function(h){a(h);a.Highcharts=h;return a}):a("undefined"!==typeof Highcharts?Highcharts:void 0)})(function(a){function h(a,b,f,t){a.hasOwnProperty(b)||(a[b]=t.apply(null,f))}a=a?a._modules:{};h(a,"Series/OnSeriesComposition.js",[a["Series/Column/ColumnSeries.js"],a["Core/Series/Series.js"],a["Core/Utilities.js"]],
function(a,b,f){var t=a.prototype,d=b.prototype,w=f.defined,y=f.stableSort,k;(function(a){function b(){return d.getPlotBox.call(this.options.onSeries&&this.chart.get(this.options.onSeries)||this)}function k(){t.translate.apply(this);var a=this,g=a.options,q=a.chart,e=a.points,c=g.onSeries,x=(c=c&&q.get(c))&&c.options.step,r=c&&c.points,m=q.inverted,d=a.xAxis,b=a.yAxis;q=e.length-1;var k;g=g.onKey||"y";var f=r&&r.length,h=0,p;if(c&&c.visible&&f){h=(c.pointXOffset||0)+(c.barW||0)/2;var l=c.currentDataGrouping;
var v=r[f-1].x+(l?l.totalRange:0);y(e,function(e,c){return e.x-c.x});for(g="plot"+g[0].toUpperCase()+g.substr(1);f--&&e[q];){var n=r[f];l=e[q];l.y=n.y;if(n.x<=l.x&&"undefined"!==typeof n[g]){if(l.x<=v&&(l.plotY=n[g],n.x<l.x&&!x&&(p=r[f+1])&&"undefined"!==typeof p[g])){var u=(l.x-n.x)/(p.x-n.x);l.plotY+=u*(p[g]-n[g]);l.y+=u*(p.y-n.y)}q--;f++;if(0>q)break}}}e.forEach(function(c,x){c.plotX+=h;if("undefined"===typeof c.plotY||m)0<=c.plotX&&c.plotX<=d.len?m?(c.plotY=d.translate(c.x,0,1,0,1),c.plotX=w(c.y)?
b.translate(c.y,0,0,0,1):0):c.plotY=(d.opposite?0:a.yAxis.len)+d.offset:c.shapeArgs={};if((k=e[x-1])&&k.plotX===c.plotX){"undefined"===typeof k.stackIndex&&(k.stackIndex=0);var r=k.stackIndex+1}c.stackIndex=r});this.onSeries=c}var f=[];a.compose=function(a){if(-1===f.indexOf(a)){f.push(a);var g=a.prototype;g.getPlotBox=b;g.translate=k}return a};a.getPlotBox=b;a.translate=k})(k||(k={}));return k});h(a,"Series/Windbarb/WindbarbPoint.js",[a["Core/Utilities.js"],a["Series/Column/ColumnSeries.js"]],function(a,
b){var f=this&&this.__extends||function(){var a=function(d,b){a=Object.setPrototypeOf||{__proto__:[]}instanceof Array&&function(a,d){a.__proto__=d}||function(a,d){for(var b in d)d.hasOwnProperty(b)&&(a[b]=d[b])};return a(d,b)};return function(d,b){function f(){this.constructor=d}a(d,b);d.prototype=null===b?Object.create(b):(f.prototype=b.prototype,new f)}}(),h=a.isNumber;return function(a){function b(){var b=null!==a&&a.apply(this,arguments)||this;b.beaufort=void 0;b.beaufortLevel=void 0;b.direction=
void 0;b.options=void 0;b.series=void 0;return b}f(b,a);b.prototype.isValid=function(){return h(this.value)&&0<=this.value};return b}(b.prototype.pointClass)});h(a,"Series/Windbarb/WindbarbSeries.js",[a["Core/Animation/AnimationUtilities.js"],a["Core/Globals.js"],a["Series/OnSeriesComposition.js"],a["Core/Series/SeriesRegistry.js"],a["Core/Utilities.js"],a["Series/Windbarb/WindbarbPoint.js"]],function(a,b,f,h,d,w){var t=this&&this.__extends||function(){var a=function(b,e){a=Object.setPrototypeOf||
{__proto__:[]}instanceof Array&&function(a,e){a.__proto__=e}||function(a,e){for(var c in e)e.hasOwnProperty(c)&&(a[c]=e[c])};return a(b,e)};return function(b,e){function c(){this.constructor=b}a(b,e);b.prototype=null===e?Object.create(e):(c.prototype=e.prototype,new c)}}(),k=a.animObject;a=b.noop;var p=h.series,v=h.seriesTypes.column,u=d.extend,A=d.merge,z=d.pick;d=function(a){function d(){var e=null!==a&&a.apply(this,arguments)||this;e.data=void 0;e.options=void 0;e.points=void 0;return e}t(d,a);
d.registerApproximation=function(){b.approximations&&!b.approximations.windbarb&&(b.approximations.windbarb=function(a,c){var e=0,d=0,m,f=a.length;for(m=0;m<f;m++)e+=a[m]*Math.cos(c[m]*b.deg2rad),d+=a[m]*Math.sin(c[m]*b.deg2rad);return[a.reduce(function(a,c){return a+c},0)/a.length,Math.atan2(d,e)/b.deg2rad]})};d.prototype.init=function(a,c){d.registerApproximation();p.prototype.init.call(this,a,c)};d.prototype.pointAttribs=function(a,c){var e=this.options;a=a.color||this.color;var b=this.options.lineWidth;
c&&(a=e.states[c].color||a,b=(e.states[c].lineWidth||b)+(e.states[c].lineWidthPlus||0));return{stroke:a,"stroke-width":b}};d.prototype.windArrow=function(a){var c=1.943844*a.value,b=this.options.vectorLength/20,e=-10;if(a.isNull)return[];if(0===a.beaufortLevel)return this.chart.renderer.symbols.circle(-10*b,-10*b,20*b,20*b);a=[["M",0,7*b],["L",-1.5*b,7*b],["L",0,10*b],["L",1.5*b,7*b],["L",0,7*b],["L",0,-10*b]];var d=(c-c%50)/50;if(0<d)for(;d--;)a.push(-10===e?["L",0,e*b]:["M",0,e*b],["L",5*b,e*b+
2],["L",0,e*b+4]),c-=50,e+=7;d=(c-c%10)/10;if(0<d)for(;d--;)a.push(-10===e?["L",0,e*b]:["M",0,e*b],["L",7*b,e*b]),c-=10,e+=3;d=(c-c%5)/5;if(0<d)for(;d--;)a.push(-10===e?["L",0,e*b]:["M",0,e*b],["L",4*b,e*b]),c-=5,e+=3;return a};d.prototype.drawPoints=function(){var a=this.chart,b=this.yAxis,d=a.inverted,f=this.options.vectorLength/2;this.points.forEach(function(c){var e=c.plotX,g=c.plotY;!1===this.options.clip||a.isInsidePlot(e,0)?(c.graphic||(c.graphic=this.chart.renderer.path().add(this.markerGroup).addClass("highcharts-point highcharts-color-"+
z(c.colorIndex,c.series.colorIndex))),c.graphic.attr({d:this.windArrow(c),translateX:e+this.options.xOffset,translateY:g+this.options.yOffset,rotation:c.direction}),this.chart.styledMode||c.graphic.attr(this.pointAttribs(c))):c.graphic&&(c.graphic=c.graphic.destroy());c.tooltipPos=[e+this.options.xOffset+(d&&!this.onSeries?f:0),g+this.options.yOffset-(d?0:f+b.pos-a.plotTop)]},this)};d.prototype.animate=function(a){a?this.markerGroup.attr({opacity:.01}):this.markerGroup.animate({opacity:1},k(this.options.animation))};
d.prototype.markerAttribs=function(a,c){return{}};d.prototype.getExtremes=function(){return{}};d.prototype.shouldShowTooltip=function(b,c,d){void 0===d&&(d={});d.ignoreX=this.chart.inverted;d.ignoreY=!d.ignoreX;return a.prototype.shouldShowTooltip.call(this,b,c,d)};d.defaultOptions=A(v.defaultOptions,{dataGrouping:{enabled:!0,approximation:"windbarb",groupPixelWidth:30},lineWidth:2,onSeries:null,states:{hover:{lineWidthPlus:0}},tooltip:{pointFormat:'<span style="color:{point.color}">\u25cf</span> {series.name}: <b>{point.value}</b> ({point.beaufort})<br/>'},
vectorLength:20,colorKey:"value",yOffset:-20,xOffset:0});return d}(v);f.compose(d);u(d.prototype,{beaufortFloor:[0,.3,1.6,3.4,5.5,8,10.8,13.9,17.2,20.8,24.5,28.5,32.7],beaufortName:"Calm;Light air;Light breeze;Gentle breeze;Moderate breeze;Fresh breeze;Strong breeze;Near gale;Gale;Strong gale;Storm;Violent storm;Hurricane".split(";"),parallelArrays:["x","value","direction"],pointArrayMap:["value","direction"],pointClass:w,trackerGroups:["markerGroup"],invertGroups:a,translate:function(){var a=this.beaufortFloor,
b=this.beaufortName;f.translate.call(this);this.points.forEach(function(d){for(var c=0;c<a.length&&!(a[c]>d.value);c++);d.beaufortLevel=c-1;d.beaufort=b[c-1]})}});d.registerApproximation();h.registerSeriesType("windbarb",d);"";return d});h(a,"masters/modules/windbarb.src.js",[],function(){})});
//# sourceMappingURL=windbarb.js.map
```
