## Supplementary material for "Exploring natural odour landscapes: A case study with implications for human-biting insects": Figure S4: wordcloud.js

```
/*
 Highcharts JS v9.3.1 (2021-11-05)

 (c) 2016-2021 Highsoft AS
 Authors: Jon Arild Nygard

 License: www.highcharts.com/license
*/
'use strict';(function(c){"object"===typeof module&&module.exports?(c["default"]=c,module.exports=c):"function"===typeof define&&define.amd?define("highcharts/modules/wordcloud",["highcharts"],function(h){c(h);c.Highcharts=h;return c}):c("undefined"!==typeof Highcharts?Highcharts:void 0)})(function(c){function h(c,f,k,g){c.hasOwnProperty(f)||(c[f]=g.apply(null,k))}c=c?c._modules:{};h(c,"Series/DrawPointComposition.js",[],function(){var c;(function(c){function f(e){var c=this,f=e.animatableAttribs,
g=e.onComplete,m=e.css,k=e.renderer,t=this.series&&this.series.chart.hasRendered?void 0:this.series&&this.series.options.animation,l=this.graphic;e.attribs=e.attribs||{};e.attribs["class"]=this.getClassName();if(this.shouldDraw())l||(this.graphic=l=k[e.shapeType](e.shapeArgs).add(e.group)),l.css(m).attr(e.attribs).animate(f,e.isNew?!1:t,g);else if(l){var h=function(){c.graphic=l=l&&l.destroy();"function"===typeof g&&g()};Object.keys(f).length?l.animate(f,void 0,function(){h()}):h()}}function g(){return!this.isNull}
var m=[];c.compose=function(e){if(-1===m.indexOf(e)){m.push(e);var c=e.prototype;c.draw=f;c.shouldDraw||(c.shouldDraw=g)}return e}})(c||(c={}));return c});h(c,"Series/Wordcloud/WordcloudPoint.js",[c["Series/DrawPointComposition.js"],c["Core/Series/SeriesRegistry.js"],c["Core/Utilities.js"]],function(c,f,k){var g=this&&this.__extends||function(){var c=function(e,f){c=Object.setPrototypeOf||{__proto__:[]}instanceof Array&&function(e,c){e.__proto__=c}||function(e,c){for(var f in c)c.hasOwnProperty(f)&&
(e[f]=c[f])};return c(e,f)};return function(e,f){function g(){this.constructor=e}c(e,f);e.prototype=null===f?Object.create(f):(g.prototype=f.prototype,new g)}}();k=k.extend;f=function(c){function e(){var e=null!==c&&c.apply(this,arguments)||this;e.dimensions=void 0;e.options=void 0;e.polygon=void 0;e.rect=void 0;e.series=void 0;return e}g(e,c);e.prototype.isValid=function(){return!0};return e}(f.seriesTypes.column.prototype.pointClass);k(f.prototype,{weight:1});c.compose(f);return f});h(c,"Series/Wordcloud/WordcloudUtils.js",
[c["Core/Globals.js"],c["Core/Utilities.js"]],function(c,f){function k(b,a){return!(a.left>b.right||a.right<b.left||a.top>b.bottom||a.bottom<b.top)}function g(b,a){var d=a[0]-b[0];b=a[1]-b[1];return[[-b,d],[b,-d]]}function h(b){var a=b.axes||[];if(!a.length){a=[];var d=d=b.concat([b[0]]);d.reduce(function(b,d){var e=g(b,d)[0];D(a,function(a){return a[0]===e[0]&&a[1]===e[1]})||a.push(e);return d});b.axes=a}return a}function e(b,a){b=b.map(function(b){return b[0]*a[0]+b[1]*a[1]});return{min:Math.min.apply(this,
b),max:Math.max.apply(this,b)}}function C(b,a){var d=h(b),c=h(a);d=d.concat(c);return!D(d,function(d){var c=e(b,d);d=e(a,d);return!!(d.min>c.max||d.max<c.min)})}function r(b,a){var d=!1,e=b.rect,c=b.polygon,f=b.lastCollidedWith,g=function(a){var d=k(e,a.rect);d&&(b.rotation%90||a.rotation%90)&&(d=C(c,a.polygon));return d};f&&((d=g(f))||delete b.lastCollidedWith);d||(d=!!D(a,function(a){var d=g(a);d&&(b.lastCollidedWith=a);return d}));return d}function x(b,a){a=4*b;var d=Math.ceil((Math.sqrt(a)-1)/
2),e=2*d+1,c=Math.pow(e,2),f=!1;--e;1E4>=b&&("boolean"===typeof f&&a>=c-e&&(f={x:d-(c-a),y:-d}),c-=e,"boolean"===typeof f&&a>=c-e&&(f={x:-d,y:-d+(c-a)}),c-=e,"boolean"===typeof f&&(f=a>=c-e?{x:-d+(c-a),y:d}:{x:d,y:d-(c-a-e)}),f.x*=5,f.y*=5);return f}function w(b,a){var d=a.width/2,c=-(a.height/2),e=a.height/2;return!(-(a.width/2)<b.left&&d>b.right&&c<b.top&&e>b.bottom)}function A(b,a,d){return d.map(function(d){return[d[0]+b,d[1]+a]})}function t(b,a){a=p(a)?a:14;a=Math.pow(10,a);return Math.round(b*
a)/a}function l(b,a){var d=b[0];b=b[1];var c=E*-a;a=Math.cos(c);c=Math.sin(c);return[t(d*a-b*c),t(d*c+b*a)]}function B(b,a,d){b=l([b[0]-a[0],b[1]-a[1]],d);return[b[0]+a[0],b[1]+a[1]]}var E=c.deg2rad,G=f.extend,D=f.find,p=f.isNumber,q=f.isObject,y=f.merge;return{archimedeanSpiral:function(b,a){var d=a.field;a=!1;d=d.width*d.width+d.height*d.height;var c=.8*b;1E4>=b&&(a={x:c*Math.cos(c),y:c*Math.sin(c)},Math.min(Math.abs(a.x),Math.abs(a.y))<d||(a=!1));return a},extendPlayingField:function(b,a){if(q(b)&&
q(a)){var d=a.bottom-a.top;var c=a.right-a.left;a=b.ratioX;var e=b.ratioY;d=c*a>d*e?c:d;b=y(b,{width:b.width+d*a*2,height:b.height+d*e*2})}return b},getBoundingBoxFromPolygon:function(b){return b.reduce(function(a,b){var d=b[0];b=b[1];a.left=Math.min(d,a.left);a.right=Math.max(d,a.right);a.bottom=Math.max(b,a.bottom);a.top=Math.min(b,a.top);return a},{left:Number.MAX_VALUE,right:-Number.MAX_VALUE,bottom:-Number.MAX_VALUE,top:Number.MAX_VALUE})},getPlayingField:function(b,a,d){d=d.reduce(function(a,
b){b=b.dimensions;var d=Math.max(b.width,b.height);a.maxHeight=Math.max(a.maxHeight,b.height);a.maxWidth=Math.max(a.maxWidth,b.width);a.area+=d*d;return a},{maxHeight:0,maxWidth:0,area:0});d=Math.max(d.maxHeight,d.maxWidth,.85*Math.sqrt(d.area));var c=b>a?b/a:1;b=a>b?a/b:1;return{width:d*c,height:d*b,ratioX:c,ratioY:b}},getPolygon:function(b,a,d,c,e){var f=[b,a],g=b-d/2;b+=d/2;d=a-c/2;a+=c/2;return[[g,d],[b,d],[b,a],[g,a]].map(function(a){return B(a,f,-e)})},getRandomPosition:function(b){return Math.round(b*
(Math.random()+.5)/2)},getRotation:function(b,a,d,c){var e=!1;p(b)&&p(a)&&p(d)&&p(c)&&0<b&&-1<a&&c>d&&(e=d+a%b*((c-d)/(b-1||1)));return e},getScale:function(b,a,d){var c=2*Math.max(Math.abs(d.top),Math.abs(d.bottom));d=2*Math.max(Math.abs(d.left),Math.abs(d.right));return Math.min(0<d?1/d*b:1,0<c?1/c*a:1)},getSpiral:function(b,a){var d,c=[];for(d=1;1E4>d;d++)c.push(b(d,a));return function(a){return 1E4>=a?c[a-1]:!1}},intersectionTesting:function(b,a){var c=a.placed,e=a.field,f=a.rectangle,g=a.polygon,
h=a.spiral,l=1,k={x:0,y:0},z=b.rect=G({},f);b.polygon=g;for(b.rotation=a.rotation;!1!==k&&(r(b,c)||w(z,e));)k=h(l),q(k)&&(z.left=f.left+k.x,z.right=f.right+k.x,z.top=f.top+k.y,z.bottom=f.bottom+k.y,b.polygon=A(k.x,k.y,g)),l++;return k},isPolygonsColliding:C,isRectanglesIntersecting:k,rectangularSpiral:function(b,a){b=x(b,a);a=a.field;b&&(b.x*=a.ratioX,b.y*=a.ratioY);return b},rotate2DToOrigin:l,rotate2DToPoint:B,squareSpiral:x,updateFieldBoundaries:function(b,a){if(!p(b.left)||b.left>a.left)b.left=
a.left;if(!p(b.right)||b.right<a.right)b.right=a.right;if(!p(b.top)||b.top>a.top)b.top=a.top;if(!p(b.bottom)||b.bottom<a.bottom)b.bottom=a.bottom;return b}}});h(c,"Series/Wordcloud/WordcloudSeries.js",[c["Core/Globals.js"],c["Core/Series/Series.js"],c["Core/Series/SeriesRegistry.js"],c["Core/Utilities.js"],c["Series/Wordcloud/WordcloudPoint.js"],c["Series/Wordcloud/WordcloudUtils.js"]],function(c,f,k,g,h,e){var m=this&&this.__extends||function(){var a=function(b,c){a=Object.setPrototypeOf||{__proto__:[]}instanceof
Array&&function(a,b){a.__proto__=b}||function(a,b){for(var c in b)b.hasOwnProperty(c)&&(a[c]=b[c])};return a(b,c)};return function(b,c){function d(){this.constructor=b}a(b,c);b.prototype=null===c?Object.create(c):(d.prototype=c.prototype,new d)}}(),r=c.noop,x=k.seriesTypes.column,w=g.extend,A=g.isArray,t=g.isNumber,l=g.isObject,B=g.merge;g=e.archimedeanSpiral;var E=e.extendPlayingField,G=e.getBoundingBoxFromPolygon,D=e.getPlayingField,p=e.getPolygon,q=e.getRandomPosition,y=e.getRotation,b=e.getScale,
a=e.getSpiral,d=e.intersectionTesting,H=e.isPolygonsColliding,I=e.rectangularSpiral,J=e.rotate2DToOrigin,K=e.rotate2DToPoint,L=e.squareSpiral,M=e.updateFieldBoundaries;e=function(e){function g(){var a=null!==e&&e.apply(this,arguments)||this;a.data=void 0;a.options=void 0;a.points=void 0;return a}m(g,e);g.prototype.bindAxes=function(){var a={endOnTick:!1,gridLineWidth:0,lineWidth:0,maxPadding:0,startOnTick:!1,title:void 0,tickPositions:[]};f.prototype.bindAxes.call(this);w(this.yAxis.options,a);w(this.xAxis.options,
a)};g.prototype.pointAttribs=function(a,b){a=c.seriesTypes.column.prototype.pointAttribs.call(this,a,b);delete a.stroke;delete a["stroke-width"];return a};g.prototype.deriveFontSize=function(a,b,c){a=t(a)?a:0;b=t(b)?b:1;c=t(c)?c:1;return Math.floor(Math.max(c,a*b))};g.prototype.drawPoints=function(){var c=this,e=c.hasRendered,f=c.xAxis,g=c.yAxis,k=c.group,h=c.options,N=h.animation,z=h.allowExtendPlayingField,r=c.chart.renderer,m=r.text().add(k),q=[],x=c.placementStrategy[h.placementStrategy],y=h.rotation,
B=c.points.map(function(a){return a.weight}),A=Math.max.apply(null,B),F=c.points.concat().sort(function(a,b){return b.weight-a.weight});c.group.attr({scaleX:1,scaleY:1});F.forEach(function(a){var b=c.deriveFontSize(1/A*a.weight,h.maxFontSize,h.minFontSize);b=w({fontSize:b+"px"},h.style);m.css(b).attr({x:0,y:0,text:a.name});b=m.getBBox(!0);a.dimensions={height:b.height,width:b.width}});var v=D(f.len,g.len,F);var C=a(c.spirals[h.spiral],{field:v});F.forEach(function(a){var b=c.deriveFontSize(1/A*a.weight,
h.maxFontSize,h.minFontSize);b=w({fontSize:b+"px"},h.style);var f=x(a,{data:F,field:v,placed:q,rotation:y}),g=w(c.pointAttribs(a,a.selected&&"select"),{align:"center","alignment-baseline":"middle","dominant-baseline":"middle",x:f.x,y:f.y,text:a.name,rotation:t(f.rotation)?f.rotation:void 0}),m=p(f.x,f.y,a.dimensions.width,a.dimensions.height,f.rotation),n=G(m),u=d(a,{rectangle:n,polygon:m,field:v,placed:q,spiral:C,rotation:f.rotation});!u&&z&&(v=E(v,n),u=d(a,{rectangle:n,polygon:m,field:v,placed:q,
spiral:C,rotation:f.rotation}));l(u)?(g.x=(g.x||0)+u.x,g.y=(g.y||0)+u.y,n.left+=u.x,n.right+=u.x,n.top+=u.y,n.bottom+=u.y,v=M(v,n),q.push(a),a.isNull=!1,a.isInside=!0):a.isNull=!0;if(N){var O={x:g.x,y:g.y};e?(delete g.x,delete g.y):(g.x=0,g.y=0)}a.draw({animatableAttribs:O,attribs:g,css:b,group:k,renderer:r,shapeArgs:void 0,shapeType:"text"})});m=m.destroy();f=b(f.len,g.len,v);c.group.attr({scaleX:f,scaleY:f})};g.prototype.hasData=function(){return l(this)&&!0===this.visible&&A(this.points)&&0<this.points.length};
g.prototype.getPlotBox=function(){var a=this.chart,b=a.inverted,c=this[b?"yAxis":"xAxis"];b=this[b?"xAxis":"yAxis"];return{translateX:(c?c.left:a.plotLeft)+(c?c.len:a.plotWidth)/2,translateY:(b?b.top:a.plotTop)+(b?b.len:a.plotHeight)/2,scaleX:1,scaleY:1}};g.defaultOptions=B(x.defaultOptions,{allowExtendPlayingField:!0,animation:{duration:500},borderWidth:0,clip:!1,colorByPoint:!0,minFontSize:1,maxFontSize:25,placementStrategy:"center",rotation:{from:0,orientations:2,to:90},showInLegend:!1,spiral:"rectangular",
style:{fontFamily:"sans-serif",fontWeight:"900",whiteSpace:"nowrap"},tooltip:{followPointer:!0,pointFormat:'<span style="color:{point.color}">\u25cf</span> {series.name}: <b>{point.weight}</b><br/>'}});return g}(x);w(e.prototype,{animate:r,animateDrilldown:r,animateDrillupFrom:r,pointClass:h,setClip:r,placementStrategy:{random:function(a,b){var c=b.field;b=b.rotation;return{x:q(c.width)-c.width/2,y:q(c.height)-c.height/2,rotation:y(b.orientations,a.index,b.from,b.to)}},center:function(a,b){b=b.rotation;
return{x:0,y:0,rotation:y(b.orientations,a.index,b.from,b.to)}}},pointArrayMap:["weight"],spirals:{archimedean:g,rectangular:I,square:L},utils:{extendPlayingField:E,getRotation:y,isPolygonsColliding:H,rotate2DToOrigin:J,rotate2DToPoint:K}});k.registerSeriesType("wordcloud",e);"";return e});h(c,"masters/modules/wordcloud.src.js",[],function(){})});
//# sourceMappingURL=wordcloud.js.map
```
