## Supplementary material for "Exploring natural odour landscapes: A case study with implications for human-biting insects": Figure S4: xrange.js

```
/*
 Highcharts JS v9.3.1 (2021-11-05)

 X-range series

 (c) 2010-2021 Torstein Honsi, Lars A. V. Cabrera

 License: www.highcharts.com/license
*/
'use strict';(function(a){"object"===typeof module&&module.exports?(a["default"]=a,module.exports=a):"function"===typeof define&&define.amd?define("highcharts/modules/xrange",["highcharts"],function(h){a(h);a.Highcharts=h;return a}):a("undefined"!==typeof Highcharts?Highcharts:void 0)})(function(a){function h(a,f,g,b){a.hasOwnProperty(f)||(a[f]=b.apply(null,g))}a=a?a._modules:{};h(a,"Series/XRange/XRangePoint.js",[a["Core/Series/Point.js"],a["Core/Series/SeriesRegistry.js"],a["Core/Utilities.js"]],
function(a,f,g){var b=this&&this.__extends||function(){var a=function(c,d){a=Object.setPrototypeOf||{__proto__:[]}instanceof Array&&function(a,d){a.__proto__=d}||function(a,d){for(var c in d)d.hasOwnProperty(c)&&(a[c]=d[c])};return a(c,d)};return function(c,d){function b(){this.constructor=c}a(c,d);c.prototype=null===d?Object.create(d):(b.prototype=d.prototype,new b)}}();g=g.extend;f=function(f){function c(){var a=null!==f&&f.apply(this,arguments)||this;a.options=void 0;a.series=void 0;return a}b(c,
f);c.getColorByCategory=function(a,c){var d=a.options.colors||a.chart.options.colors;a=c.y%(d?d.length:a.chart.options.chart.colorCount);return{colorIndex:a,color:d&&d[a]}};c.prototype.resolveColor=function(){var a=this.series;if(a.options.colorByPoint&&!this.options.color){var b=c.getColorByCategory(a,this);a.chart.styledMode||(this.color=b.color);this.options.colorIndex||(this.colorIndex=b.colorIndex)}else this.color||(this.color=a.color)};c.prototype.init=function(){a.prototype.init.apply(this,
arguments);this.y||(this.y=0);return this};c.prototype.setState=function(){a.prototype.setState.apply(this,arguments);this.series.drawPoint(this,this.series.getAnimationVerb())};c.prototype.getLabelConfig=function(){var d=a.prototype.getLabelConfig.call(this),c=this.series.yAxis.categories;d.x2=this.x2;d.yCategory=this.yCategory=c&&c[this.y];return d};c.prototype.isValid=function(){return"number"===typeof this.x&&"number"===typeof this.x2};return c}(f.seriesTypes.column.prototype.pointClass);g(f.prototype,
{ttBelow:!1,tooltipDateKeys:["x","x2"]});return f});h(a,"Series/XRange/XRangeComposition.js",[a["Core/Axis/Axis.js"],a["Core/Utilities.js"]],function(a,f){var g=f.addEvent,b=f.pick;g(a,"afterGetSeriesExtremes",function(){var a=this.series,c;if(this.isXAxis){var d=b(this.dataMax,-Number.MAX_VALUE);a.forEach(function(a){a.x2Data&&a.x2Data.forEach(function(a){a>d&&(d=a,c=!0)})});c&&(this.dataMax=d)}})});h(a,"Series/XRange/XRangeSeries.js",[a["Core/Globals.js"],a["Core/Color/Color.js"],a["Core/Series/SeriesRegistry.js"],
a["Core/Utilities.js"],a["Series/XRange/XRangePoint.js"]],function(a,f,g,b,h){var c=this&&this.__extends||function(){var a=function(c,e){a=Object.setPrototypeOf||{__proto__:[]}instanceof Array&&function(e,a){e.__proto__=a}||function(e,a){for(var q in a)a.hasOwnProperty(q)&&(e[q]=a[q])};return a(c,e)};return function(c,e){function q(){this.constructor=c}a(c,e);c.prototype=null===e?Object.create(e):(q.prototype=e.prototype,new q)}}(),d=f.parse,z=g.series,r=g.seriesTypes.column,A=r.prototype,v=b.clamp,
D=b.correctFloat,E=b.defined;f=b.extend;var B=b.find,u=b.isNumber,w=b.isObject,t=b.merge,x=b.pick;b=function(a){function b(){var e=null!==a&&a.apply(this,arguments)||this;e.data=void 0;e.options=void 0;e.points=void 0;return e}c(b,a);b.prototype.init=function(){r.prototype.init.apply(this,arguments);this.options.stacking=void 0};b.prototype.getColumnMetrics=function(){function e(){a.series.forEach(function(a){var e=a.xAxis;a.xAxis=a.yAxis;a.yAxis=e})}var a=this.chart;e();var b=A.getColumnMetrics.call(this);
e();return b};b.prototype.cropData=function(a,b,c,d){b=z.prototype.cropData.call(this,this.x2Data,b,c,d);b.xData=a.slice(b.start,b.end);return b};b.prototype.findPointIndex=function(a){var e=this.cropped,b=this.cropStart,c=this.points,C=a.id;if(C)var d=(d=B(c,function(a){return a.id===C}))?d.index:void 0;"undefined"===typeof d&&(d=(d=B(c,function(e){return e.x===a.x&&e.x2===a.x2&&!e.touched}))?d.index:void 0);e&&u(d)&&u(b)&&d>=b&&(d-=b);return d};b.prototype.translatePoint=function(a){var e=this.xAxis,
b=this.yAxis,d=this.columnMetrics,c=this.options,f=c.minPointLength||0,k=(a.shapeArgs&&a.shapeArgs.width||0)/2,l=this.pointXOffset=d.offset,m=a.plotX,g=x(a.x2,a.x+(a.len||0)),n=e.translate(g,0,0,0,1);g=Math.abs(n-m);var p=this.chart.inverted,h=x(c.borderWidth,1)%2/2,r=d.offset,y=Math.round(d.width);f&&(f-=g,0>f&&(f=0),m-=f/2,n+=f/2);m=Math.max(m,-10);n=v(n,-10,e.len+10);E(a.options.pointWidth)&&(r-=(Math.ceil(a.options.pointWidth)-y)/2,y=Math.ceil(a.options.pointWidth));c.pointPlacement&&u(a.plotY)&&
b.categories&&(a.plotY=b.translate(a.y,0,1,0,1,c.pointPlacement));c=Math.floor(Math.min(m,n))+h;c={x:c,y:Math.floor(a.plotY+r)+h,width:Math.floor(Math.max(m,n))+h-c,height:y,r:this.options.borderRadius};a.shapeArgs=c;p?a.tooltipPos[1]+=l+k:a.tooltipPos[0]-=k+l-c.width/2;k=c.x;l=k+c.width;0>k||l>e.len?(k=v(k,0,e.len),l=v(l,0,e.len),n=l-k,a.dlBox=t(c,{x:k,width:l-k,centerX:n?n/2:null})):a.dlBox=null;k=a.tooltipPos;l=p?1:0;n=p?0:1;d=this.columnMetrics?this.columnMetrics.offset:-d.width/2;k[l]=p?k[l]+
c.width/2:k[l]+(e.reversed?-1:0)*c.width;k[n]=v(k[n]+(p?-1:1)*d,0,b.len-1);if(b=a.partialFill)w(b)&&(b=b.amount),u(b)||(b=0),a.partShapeArgs=t(c,{r:this.options.borderRadius}),m=Math.max(Math.round(g*b+a.plotX-m),0),a.clipRectArgs={x:e.reversed?c.x+g-m:c.x,y:c.y,width:m,height:c.height}};b.prototype.translate=function(){A.translate.apply(this,arguments);this.points.forEach(function(a){this.translatePoint(a)},this)};b.prototype.drawPoint=function(a,b){var c=this.options,e=this.chart.renderer,f=a.graphic,
g=a.shapeType,k=a.shapeArgs,l=a.partShapeArgs,m=a.clipRectArgs,h=a.partialFill,n=c.stacking&&!c.borderRadius,p=a.state,q=c.states[p||"normal"]||{},r="undefined"===typeof p?"attr":b;p=this.pointAttribs(a,p);q=x(this.chart.options.chart.animation,q.animation);if(a.isNull||!1===a.visible)f&&(a.graphic=f.destroy());else{if(f)f.rect[b](k);else a.graphic=f=e.g("point").addClass(a.getClassName()).add(a.group||this.group),f.rect=e[g](t(k)).addClass(a.getClassName()).addClass("highcharts-partfill-original").add(f);
l&&(f.partRect?(f.partRect[b](t(l)),f.partialClipRect[b](t(m))):(f.partialClipRect=e.clipRect(m.x,m.y,m.width,m.height),f.partRect=e[g](l).addClass("highcharts-partfill-overlay").add(f).clip(f.partialClipRect)));this.chart.styledMode||(f.rect[b](p,q).shadow(c.shadow,null,n),l&&(w(h)||(h={}),w(c.partialFill)&&(h=t(c.partialFill,h)),a=h.fill||d(p.fill).brighten(-.3).get()||d(a.color||this.color).brighten(-.3).get(),p.fill=a,f.partRect[r](p,q).shadow(c.shadow,null,n)))}};b.prototype.drawPoints=function(){var a=
this,b=a.getAnimationVerb();a.points.forEach(function(c){a.drawPoint(c,b)})};b.prototype.getAnimationVerb=function(){return this.chart.pointCount<(this.options.animationLimit||250)?"animate":"attr"};b.prototype.isPointInside=function(b){var c=b.shapeArgs,d=b.plotX,e=b.plotY;return c?"undefined"!==typeof d&&"undefined"!==typeof e&&0<=e&&e<=this.yAxis.len&&0<=(c.x||0)+(c.width||0)&&d<=this.xAxis.len:a.prototype.isPointInside.apply(this,arguments)};b.defaultOptions=t(r.defaultOptions,{colorByPoint:!0,
dataLabels:{formatter:function(){var a=this.point.partialFill;w(a)&&(a=a.amount);if(u(a)&&0<a)return D(100*a)+"%"},inside:!0,verticalAlign:"middle"},tooltip:{headerFormat:'<span style="font-size: 10px">{point.x} - {point.x2}</span><br/>',pointFormat:'<span style="color:{point.color}">\u25cf</span> {series.name}: <b>{point.yCategory}</b><br/>'},borderRadius:3,pointRange:0});return b}(r);f(b.prototype,{type:"xrange",parallelArrays:["x","x2","y"],requireSorting:!1,animate:z.prototype.animate,cropShoulder:1,
getExtremesFromAll:!0,autoIncrement:a.noop,buildKDTree:a.noop,pointClass:h});g.registerSeriesType("xrange",b);"";return b});h(a,"masters/modules/xrange.src.js",[],function(){})});
//# sourceMappingURL=xrange.js.map
```
