## Supplementary material for "Exploring natural odour landscapes: A case study with implications for human-biting insects": Figure S4: draggable-legend.js

```
/**
 * Highcharts plugin for dragging a legend by its title
 *
 * Author: Torstein Hønsi
 * License: MIT License
 * Version: 1.3.9
 * Requires: Highcharts 3.0+
 *
 * Usage: Set draggable:true and floating:true in the legend options. The legend
 * preserves is alignment after dragging. For example if it is aligned to the right,
 * if will keep the same distance to the right edge even after chart resize or
 * when exporting to a different size.
 */
(function (H) {
    var addEvent = H.addEvent;

    H.wrap(H.Chart.prototype, 'init', function (proceed) {
        proceed.apply(this, Array.prototype.slice.call(arguments, 1));

        var chart = this, 
            legend = chart.legend,
            title = legend.title,
            options = legend.options,
            isDragging,
            downX,
            downY,
            optionsX,
            optionsY,
            currentX,
            currentY;
        

        function pointerDown(e) {
            e = chart.pointer.normalize(e);
            downX = e.chartX;
            downY = e.chartY;
            optionsX = options.x;
            optionsY = options.y;
            currentX = legend.group.attr('translateX');
            currentY = legend.group.attr('translateY');
            isDragging = true;
        }
        
        function pointerMove(e) {
            if (isDragging) {
                e = chart.pointer.normalize(e);
                var draggedX = e.chartX - downX,
                    draggedY = e.chartY - downY;

                // Stop touch-panning the page
                e.preventDefault();

                // Do the move is we're inside the chart
                if (
                    currentX + draggedX > 0 &&
                    currentX + draggedX + legend.legendWidth < chart.chartWidth &&
                    currentY + draggedY > 0 &&
                    currentY + draggedY + legend.legendHeight < chart.chartHeight
                ) {

                    options.x = optionsX + draggedX;
                    options.y = optionsY + draggedY;
                    legend.group.placed = false; // prevent animation
                    legend.group.align(H.extend({
                        width: legend.legendWidth,
                        height: legend.legendHeight
                    }, options), true, 'spacingBox');
                    legend.positionCheckboxes();
                }
                if (chart.pointer.selectionMarker) {
                    chart.pointer.selectionMarker = chart.pointer.selectionMarker.destroy();
                }

            }
        }
        
        function pointerUp() {
            isDragging = false;
        }

        if (options.draggable && title) {

            title.css({ cursor: 'move' });

            // Mouse events
            addEvent(title.element, 'mousedown', pointerDown);
            addEvent(chart.container, 'mousemove', pointerMove);
            addEvent(document, 'mouseup', pointerUp);

            // Touch events
            addEvent(title.element, 'touchstart', pointerDown);
            addEvent(chart.container, 'touchmove', pointerMove);
            addEvent(document, 'touchend', pointerUp);

        }
    });
}(Highcharts));
```
