## Supplementary material for "Exploring natural odour landscapes: A case study with implications for human-biting insects": Figure S4: grouped-categories.js

```
/* global Highcharts module */
(function (factory) {
	if (typeof module === 'object' && module.exports) {
		module.exports = factory;
	} else {
		factory(Highcharts);
	}
}(function (HC) {
	'use strict';
	/**
	 * Grouped Categories v1.2.0 (2021-05-10)
	 *
	 * (c) 2012-2021 Black Label
	 *
	 * License: Creative Commons Attribution (CC)
	 */

	/* jshint expr:true, boss:true */
	var UNDEFINED = void 0,
		mathRound = Math.round,
		mathMin = Math.min,
		mathMax = Math.max,
		merge = HC.merge,
		pick = HC.pick,
		each = HC.each,

		// cache prototypes
		axisProto = HC.Axis.prototype,
		tickProto = HC.Tick.prototype,

		// cache original methods
		protoAxisInit = axisProto.init,
		protoAxisRender = axisProto.render,
		protoAxisSetCategories = axisProto.setCategories,
		protoTickGetLabelSize = tickProto.getLabelSize,
		protoTickAddLabel = tickProto.addLabel,
		protoTickDestroy = tickProto.destroy,
		protoTickRender = tickProto.render;

	function deepClone(thing) {
		return JSON.parse(JSON.stringify(thing));
	}

	function Category(obj, parent) {
		this.userOptions = deepClone(obj);
		this.name = obj.name || obj;
		this.parent = parent;

		return this;
	}

	Category.prototype.toString = function () {
		var parts = [],
			cat = this;

		while (cat) {
			parts.push(cat.name);
			cat = cat.parent;
		}

		return parts.join(', ');
	};

	// returns sum of an array
	function sum(arr) {
		var l = arr.length,
			x = 0;

		while (l--) {
			x += arr[l];
		}

		return x;
	}

	// Adds category leaf to array
	function addLeaf(out, cat, parent) {
		out.unshift(new Category(cat, parent));

		while (parent) {
			parent.leaves = parent.leaves ? (parent.leaves + 1) : 1;
			parent = parent.parent;
		}
	}

	// Builds reverse category tree
	function buildTree(cats, out, options, parent, depth) {
		var len = cats.length,
			cat;

		depth = depth ? depth : 0;
		options.depth = options.depth ? options.depth : 0;

		while (len--) {
			cat = cats[len];
			
			if (cat.categories) {
				if (parent) {
					cat.parent = parent;
				}
				buildTree(cat.categories, out, options, cat, depth + 1);
			} else {
				addLeaf(out, cat, parent);
			}
		}
		options.depth = mathMax(options.depth, depth);
	}

	// Pushes part of grid to path
	function addGridPart(path, d, width) {
		// Based on crispLine from HC (#65)
		if (d[0] === d[2]) {
			d[0] = d[2] = mathRound(d[0]) - (width % 2 / 2);
		}
		if (d[1] === d[3]) {
			d[1] = d[3] = mathRound(d[1]) + (width % 2 / 2);
		}

		path.push(
			'M',
			d[0], d[1],
			'L',
			d[2], d[3]
		);
	}

	// Returns tick position
	function tickPosition(tick, pos) {
		return tick.getPosition(tick.axis.horiz, pos, tick.axis.tickmarkOffset);
	}

	function walk(arr, key, fn) {
		var l = arr.length,
			children;

		while (l--) {
			children = arr[l][key];

			if (children) {
				walk(children, key, fn);
			}
			fn(arr[l]);
		}
	}

	//
	// Axis prototype
	//

	axisProto.init = function (chart, options) {
		// default behaviour
		protoAxisInit.call(this, chart, options);

		if (typeof options === 'object' && options.categories) {
			this.setupGroups(options);
		}
	};

	// setup required axis options
	axisProto.setupGroups = function (options) {
		var categories = deepClone(options.categories),
			reverseTree = [],
			stats = {},
			labelOptions = this.options.labels,
			userAttr = labelOptions.groupedOptions,
			css = labelOptions.style;

		// build categories tree
		buildTree(categories, reverseTree, stats);

		// set axis properties
		this.categoriesTree = categories;
		this.categories = reverseTree;
		this.isGrouped = stats.depth !== 0;
		this.labelsDepth = stats.depth;
		this.labelsSizes = [];
		this.labelsGridPath = [];
		this.tickLength = options.tickLength || this.tickLength || null;
		// #66: tickWidth for x axis defaults to 1, for y to 0
		this.tickWidth = pick(options.tickWidth, this.isXAxis ? 1 : 0);
		this.directionFactor = [-1, 1, 1, -1][this.side];
		this.options.lineWidth = pick(options.lineWidth, 1);
		// #85: align labels vertically
		this.groupFontHeights = [];
		for (var i = 0; i <= stats.depth; i++) {
			var hasOptions = userAttr && userAttr[i - 1],
				mergedCSS = hasOptions && userAttr[i - 1].style ? merge(css, userAttr[i - 1].style) : css;
			this.groupFontHeights[i] = Math.round(this.chart.renderer.fontMetrics(mergedCSS ? mergedCSS.fontSize : 0).b * 0.3);
		}
	};


	axisProto.render = function () {
		// clear grid path
		if (this.isGrouped) {
			this.labelsGridPath = [];
		}

		// cache original tick length
		if (this.originalTickLength === UNDEFINED) {
			this.originalTickLength = this.options.tickLength;
		}

		// use default tickLength for not-grouped axis
		// and generate grid on grouped axes,
		// use tiny number to force highcharts to hide tick
		this.options.tickLength = this.isGrouped ? 0.001 : this.originalTickLength;

		protoAxisRender.call(this);

		if (!this.isGrouped) {
			if (this.labelsGrid) {
				this.labelsGrid.attr({
					visibility: 'hidden'
				});
			}
			return false;
		}

		var axis = this,
			options = axis.options,
			top = axis.top,
			left = axis.left,
			right = left + axis.width,
			bottom = top + axis.height,
			visible = axis.hasVisibleSeries || axis.hasData,
			depth = axis.labelsDepth,
			grid = axis.labelsGrid,
			horiz = axis.horiz,
			d = axis.labelsGridPath,
			i = options.drawHorizontalBorders === false ? (depth + 1) : 0,
			offset = axis.opposite ? (horiz ? top : right) : (horiz ? bottom : left),
			tickWidth = axis.tickWidth,
			part;

		if (axis.userTickLength) {
			depth -= 1;
		}

		// render grid path for the first time
		if (!grid) {
			grid = axis.labelsGrid = axis.chart.renderer.path()
			.attr({
				// #58: use tickWidth/tickColor instead of lineWidth/lineColor:
				strokeWidth: tickWidth, // < 4.0.3
				'stroke-width': tickWidth, // 4.0.3+ #30
				stroke: options.tickColor || '' // for styled mode (tickColor === undefined)
			})
			.add(axis.axisGroup);
			// for styled mode - add class
			if (!options.tickColor) {
				grid.addClass('highcharts-tick');
			}
		}

		// go through every level and draw horizontal grid line
		while (i <= depth) {
			offset += axis.groupSize(i);

			part = horiz ?
				[left, offset, right, offset] :
				[offset, top, offset, bottom];

			addGridPart(d, part, tickWidth);
			i++;
		}

		// draw grid path
		grid.attr({
			d: d,
			visibility: visible ? 'visible' : 'hidden'
		});

		axis.labelGroup.attr({
			visibility: visible ? 'visible' : 'hidden'
		});


		walk(axis.categoriesTree, 'categories', function (group) {
			var tick = group.tick;

			if (!tick) {
				return false;
			}
			if (tick.startAt + tick.leaves - 1 < axis.min || tick.startAt > axis.max) {
				tick.label.hide();
				tick.destroyed = 0;
			} else {
				tick.label.attr({
					visibility: visible ? 'visible' : 'hidden'
				});
			}
			return true;
		});
		return true;
	};

	axisProto.setCategories = function (newCategories, doRedraw) {
		if (this.categories) {
			this.cleanGroups();
		}
		this.setupGroups({
			categories: newCategories
		});
		this.categories = this.userOptions.categories = newCategories;
		protoAxisSetCategories.call(this, this.categories, doRedraw);
	};

	// cleans old categories
	axisProto.cleanGroups = function () {
		var ticks = this.ticks,
			n;

		for (n in ticks) {
			if (ticks[n].parent) {
				delete ticks[n].parent;
			}
		}
		walk(this.categoriesTree, 'categories', function (group) {
			var tick = group.tick;
			
			if (!tick) {
				return false;
			}
			tick.label.destroy();
			
			each(tick, function (v, i) {
				delete tick[i];
			});
			delete group.tick;
			
			return true;
		});
		this.labelsGrid = null;
	};

	// keeps size of each categories level
	axisProto.groupSize = function (level, position) {
		var positions = this.labelsSizes,
			direction = this.directionFactor,
			groupedOptions = this.options.labels.groupedOptions ? this.options.labels.groupedOptions[level - 1] : false,
			userXY = 0;

		if (groupedOptions) {
			if (direction === -1) {
				userXY = groupedOptions.x ? groupedOptions.x : 0;
			} else {
				userXY = groupedOptions.y ? groupedOptions.y : 0;
			}
		}

		if (position !== UNDEFINED) {
			positions[level] = mathMax(positions[level] || 0, position + 10 + Math.abs(userXY));
		}

		if (level === true) {
			return sum(positions) * direction;
		} else if (positions[level]) {
			return positions[level] * direction;
		}

		return 0;
	};

	//
	// Tick prototype
	//

	// Override methods prototypes
	tickProto.addLabel = function () {
		var tick = this,
			axis = tick.axis,
			labelOptions = pick(
				tick.options && tick.options.labels,
				axis.options.labels
			),
			category,
			formatter;
		
		protoTickAddLabel.call(tick);
		
		if (!axis.categories || !(category = axis.categories[tick.pos])) {
			return false;
		}
		
		// set label text - but applied after formatter #46
		if (tick.label) {
			formatter = function (ctx) {
				if (labelOptions.formatter) {
					return labelOptions.formatter.call(ctx, ctx);
				}
				if (labelOptions.format) {
					ctx.text = axis.defaultLabelFormatter.call(ctx);
					return HC.format(labelOptions.format, ctx, axis.chart);
				}
				return axis.defaultLabelFormatter.call(ctx, ctx);
			};

			tick.label.attr('text', formatter({
				axis: axis,
				chart: axis.chart,
				isFirst: tick.isFirst,
				isLast: tick.isLast,
				value: category.name,
				pos: tick.pos
			}));

			// update with new text length, since textSetter removes the size caches when text changes. #137
			tick.label.textPxLength = tick.label.getBBox().width;
		}
		
		// create elements for parent categories
		if (axis.isGrouped && axis.options.labels.enabled) {
			tick.addGroupedLabels(category);
		}
		return true;
	};

	// render ancestor label
	tickProto.addGroupedLabels = function (category) {
		var tick = this,
			axis = this.axis,
			chart = axis.chart,
			options = axis.options.labels,
			useHTML = options.useHTML,
			css = options.style,
			userAttr = options.groupedOptions,
			attr = {
				align: 'center',
				rotation: options.rotation,
				x: 0,
				y: 0
			},
			size = axis.horiz ? 'height' : 'width',
			depth = 0,
			label;


		while (tick) {
			if (depth > 0 && !category.tick) {
				// render label element
				this.value = category.name;
				var name = options.formatter ? options.formatter.call(this, category) : category.name,
					hasOptions = userAttr && userAttr[depth - 1],
					mergedAttrs = hasOptions ? merge(attr, userAttr[depth - 1]) : attr,
					mergedCSS = hasOptions && userAttr[depth - 1].style ? merge(css, userAttr[depth - 1].style) : css;

				// #63: style is passed in CSS and not as an attribute
				delete mergedAttrs.style;

				label = chart.renderer.text(name, 0, 0, useHTML)
					.attr(mergedAttrs)
					.add(axis.labelGroup);

				// css should only be set for non styledMode configuration. #167
				if (label && !chart.styledMode) {
					label.css(mergedCSS);
				}

				// tick properties
				tick.startAt = this.pos;
				tick.childCount = category.categories.length;
				tick.leaves = category.leaves;
				tick.visible = this.childCount;
				tick.label = label;
				tick.labelOffsets = {
					x: mergedAttrs.x,
					y: mergedAttrs.y
				};

				// link tick with category
				category.tick = tick;
			}

			// set level size, #93
			if (tick && tick.label) {
				axis.groupSize(depth, tick.label.getBBox()[size]);
			}

			// go up to the parent category
			category = category.parent;

			if (category) {
				tick = tick.parent = category.tick || {};
			} else {
				tick = null;
			}

			depth++;
		}
	};

	// set labels position & render categories grid
	tickProto.render = function (index, old, opacity) {
		protoTickRender.call(this, index, old, opacity);

		var treeCat = this.axis.categories[this.pos];

		if (!this.axis.isGrouped || !treeCat || this.pos > this.axis.max) {
			return;
		}

		var tick = this,
			group = tick,
			axis = tick.axis,
			tickPos = tick.pos,
			isFirst = tick.isFirst,
			max = axis.max,
			min = axis.min,
			horiz = axis.horiz,
			grid = axis.labelsGridPath,
			size = axis.groupSize(0),
			tickWidth = axis.tickWidth,
			xy = tickPosition(tick, tickPos),
			start = horiz ? xy.y : xy.x,
			baseLine = axis.chart.renderer.fontMetrics(axis.options.labels.style ? axis.options.labels.style.fontSize : 0).b,
			depth = 1,
			reverseCrisp = ((horiz && xy.x === axis.pos + axis.len) || (!horiz && xy.y === axis.pos)) ? -1 : 0, // adjust grid lines for edges
			gridAttrs,
			lvlSize,
			minPos,
			maxPos,
			attrs,
			bBox;

		// render grid for "normal" categories (first-level), render left grid line only for the first category
		if (isFirst) {
			gridAttrs = horiz ?
				[axis.left, xy.y, axis.left, xy.y + axis.groupSize(true)] : axis.isXAxis ?
					[xy.x, axis.top, xy.x + axis.groupSize(true), axis.top] : [xy.x, axis.top + axis.len, xy.x + axis.groupSize(true), axis.top + axis.len];

			addGridPart(grid, gridAttrs, tickWidth);
		}

		if (horiz && axis.left < xy.x) {
			addGridPart(grid, [xy.x - reverseCrisp, xy.y, xy.x - reverseCrisp, xy.y + size], tickWidth);
		} else if (!horiz && axis.top <= xy.y) {
			addGridPart(grid, [xy.x, xy.y + reverseCrisp, xy.x + size, xy.y + reverseCrisp], tickWidth);
		}

		size = start + size;

		function fixOffset(tCat) {
			var ret = 0;
			if (isFirst) {
				ret = tCat.parent.categories.indexOf(tCat.name);
				ret = ret < 0 ? 0 : ret;
				return ret;
			}
			return ret;
		}


		while (group.parent) {
			group = group.parent;
			
			var fix = fixOffset(treeCat),
				userX = group.labelOffsets.x,
				userY = group.labelOffsets.y;
			
			minPos = tickPosition(tick, mathMax(group.startAt - 1, min - 1));
			maxPos = tickPosition(tick, mathMin(group.startAt + group.leaves - 1 - fix, max));
			bBox = group.label.getBBox(true);
			lvlSize = axis.groupSize(depth);
			// check if on the edge to adjust
			reverseCrisp = ((horiz && maxPos.x === axis.pos + axis.len) || (!horiz && maxPos.y === axis.pos)) ? -1 : 0;
			
			attrs = horiz ? {
				x: (minPos.x + maxPos.x) / 2 + userX,
				y: size + axis.groupFontHeights[depth] + lvlSize / 2 + userY / 2
			} : {
				x: size + lvlSize / 2 + userX,
				y: (minPos.y + maxPos.y - bBox.height) / 2 + baseLine + userY
			};
			
			if (!isNaN(attrs.x) && !isNaN(attrs.y)) {
				group.label.attr(attrs);

				if (grid) {
					if (horiz && axis.left < maxPos.x) {
						addGridPart(grid, [maxPos.x - reverseCrisp, size, maxPos.x - reverseCrisp, size + lvlSize], tickWidth);
					} else if (!horiz && axis.top <= maxPos.y) {
						addGridPart(grid, [size, maxPos.y + reverseCrisp, size + lvlSize, maxPos.y + reverseCrisp], tickWidth);
					}
				}
			}

			size += lvlSize;
			depth++;
		}
	};

	tickProto.destroy = function () {
		var group = this.parent;

		while (group) {
			group.destroyed = group.destroyed ? (group.destroyed + 1) : 1;
			group = group.parent;
		}

		protoTickDestroy.call(this);
	};

	// return size of the label (height for horizontal, width for vertical axes)
	tickProto.getLabelSize = function () {
		if (this.axis.isGrouped === true) {
			// #72, getBBox might need recalculating when chart is tall
			var size = protoTickGetLabelSize.call(this) + 10,
				topLabelSize = this.axis.labelsSizes[0];
			if (topLabelSize < size) {
				this.axis.labelsSizes[0] = size;
			}
			return sum(this.axis.labelsSizes);
		}
		return protoTickGetLabelSize.call(this);
	};
	
	// Since datasorting is not supported by the plugin,
	// override replaceMovedLabel method, #146.
	HC.wrap(HC.Tick.prototype, 'replaceMovedLabel', function (proceed) {
		if (!this.axis.isGrouped) {
			proceed.apply(this, Array.prototype.slice.call(arguments, 1));
		}
	});

}));
```
