## Supplementary material for "Exploring natural odour landscapes: A case study with implications for human-biting insects": Figure S4: highcharts-regression.js

```
(function(factory) {
	"use strict";

	if (typeof module === "object" && module.exports) {
        module.exports = factory;
    } else {
		if(typeof define === "function" && define.amd){
			define(function() {
				return factory
			});
		}
		else{
			if(typeof Highcharts !== "undefined"){
				factory(Highcharts);
			}
			else{
				void 0;
			}
		}
    }
}(function (H) {
    var processSerie = function (s, method, chart) {
        if (s.regression && !s.rendered) {
            s.regressionSettings = s.regressionSettings || {};
            s.regressionSettings.tooltip = s.regressionSettings.tooltip || {};
            s.regressionSettings.dashStyle = s.regressionSettings.dashStyle || 'solid';
            s.regressionSettings.decimalPlaces = s.regressionSettings.decimalPlaces || 2;
            s.regressionSettings.useAllSeries = s.regressionSettings.useAllSeries || false;
            s.regressionSettings.regressionSeriesOptions = s.regressionSettings.regressionSeriesOptions || {};

            var regressionType = s.regressionSettings.type || "linear";
            var regression;
            var extraSerie = s.regressionSettings.regressionSeriesOptions;
            
            // Set default values
            extraSerie.data = [];
            extraSerie.isRegressionLine = true;
            if (extraSerie.color === undefined)         extraSerie.color = s.regressionSettings.color || '';
            if (extraSerie.yAxis === undefined)         extraSerie.yAxis = s.yAxis;
            if (extraSerie.lineWidth === undefined)     extraSerie.lineWidth = s.regressionSettings.lineWidth || 2;
            if (extraSerie.marker === undefined)        extraSerie.marker = { enabled: false };
            if (extraSerie.visible === undefined)       extraSerie.visible = s.regressionSettings.visible;
            if (extraSerie.type === undefined)          extraSerie.type = s.regressionSettings.linetype || 'spline';
            if (extraSerie.name === undefined)          extraSerie.name = s.regressionSettings.name || "Equation = %eq";
            if (extraSerie.id === undefined)            extraSerie.id = s.regressionSettings.id;
            if (extraSerie.dashStyle === undefined)     extraSerie.dashStyle = s.regressionSettings.dashStyle || 'solid';
            if (extraSerie.showInLegend === undefined)  extraSerie.showInLegend = !s.regressionSettings.hideInLegend;
            if (extraSerie.tooltip === undefined)       extraSerie.tooltip = s.regressionSettings.tooltip;

            if (s.regressionSettings.tooltip.valueSuffix === undefined) {
                extraSerie.tooltip.valueSuffix = ' ';
            }

            if (typeof s.regressionSettings.index !== 'undefined') {
                extraSerie.index = s.regressionSettings.index;
            }
            if (typeof s.regressionSettings.legendIndex !== 'undefined') {
                extraSerie.legendIndex = s.regressionSettings.legendIndex;
            }

            var mergedData = s.data;
            if (s.regressionSettings.useAllSeries) {
                mergedData = [];
                for (di = 0; di < series.length; di++) {
                    var seriesToMerge = series[di];
                    mergedData = mergedData.concat(seriesToMerge.data);
                }
            }

            if (regressionType == "linear") {
                var extrapolate = s.regressionSettings.extrapolate || 0;
                regression = _linear(mergedData, s.regressionSettings.decimalPlaces, extrapolate);
                extraSerie.type = "line";
            } else if (regressionType == "exponential") {
                var extrapolate = s.regressionSettings.extrapolate || 0;
                regression = _exponential(mergedData, extrapolate);
            }
            else if (regressionType == "polynomial") {
                var order = s.regressionSettings.order || 2;
                var extrapolate = s.regressionSettings.extrapolate || 0;
                regression = _polynomial(mergedData, order, extrapolate);
            } else if (regressionType == "power") {
                var extrapolate = s.regressionSettings.extrapolate || 0;
                regression = _power(mergedData, extrapolate);
            } else if (regressionType == "logarithmic") {
                var extrapolate = s.regressionSettings.extrapolate || 0;
                regression = _logarithmic(mergedData, extrapolate);
            } else if (regressionType == "loess") {
                var loessSmooth = s.regressionSettings.loessSmooth || 25;
                regression = _loess(mergedData, loessSmooth / 100);
            } else {
                console.error("Invalid regression type: ", regressionType);
                return;
            }

            regression.rSquared = coefficientOfDetermination(mergedData, regression.points);
            regression.rValue = _round(Math.sqrt(regression.rSquared), s.regressionSettings.decimalPlaces);
            regression.rSquared = _round(regression.rSquared, s.regressionSettings.decimalPlaces);
            regression.standardError = _round(standardError(mergedData, regression.points), s.regressionSettings.decimalPlaces);
            extraSerie.data = regression.points;
            extraSerie.name = replaceRegressionPlaceholders(extraSerie.name, regression);

            if (s.regressionSettings.dataLabels !== undefined) {
                for (let i = 0; i < s.regressionSettings.dataLabels.length; i++) {
                    var dataLabel = s.regressionSettings.dataLabels[i]
                    var dataLabelPoint = extraSerie.data[dataLabel.pointIndex];
                    var dataLabelFormat = replaceRegressionPlaceholders(dataLabel.format, regression);

                    if (dataLabelPoint !== undefined) {
                        extraSerie.data[dataLabel.pointIndex] = {
                            x: dataLabelPoint[0],
                            y: dataLabelPoint[1],
                            dataLabels: {
                                enabled: true,
                                format: dataLabelFormat
                            }
                        };
                    }
                }
            }

            extraSerie.regressionOutputs = regression;
            return extraSerie;
        }
    }

    H.wrap(H.Chart.prototype, 'init', function (proceed) {
        var series = arguments[1].series;
        var extraSeries = [];
        var i = 0;
        if (series) {
            for (i = 0; i < series.length; i++) {
                var s = series[i];
                if (s.regression) {
                    var extraSerie = processSerie(s, 'init', this);
                    extraSeries.push(extraSerie);
                    arguments[1].series[i].rendered = true;
                }
            }
        }

        if (extraSerie) {
            arguments[1].series = series.concat(extraSeries);
        }

        proceed.apply(this, Array.prototype.slice.call(arguments, 1));


    });

    H.wrap(H.Chart.prototype, 'addSeries', function (proceed) {
        var s = arguments[1];
        var extraSerie = processSerie(s, 'addSeries', this);
        arguments[1].rendered = true;
        if (extraSerie) {
            this.addSeries(extraSerie);
        }
        return proceed.apply(this, Array.prototype.slice.call(arguments, 1));
    });


    /**
     * Code extracted from https://github.com/Tom-Alexander/regression-js/
     */
    function _exponential(data, extrapolate) {
        var sum = [0, 0, 0, 0, 0, 0], n = 0, results = [];

        for (len = data.length; n < len; n++) {
            if (data[n].x != null) {
                data[n][0] = data[n].x;
                data[n][1] = data[n].y;
            }
            if (data[n][1] != null) {
                sum[0] += data[n][0]; // X
                sum[1] += data[n][1]; // Y
                sum[2] += data[n][0] * data[n][0] * data[n][1]; // XXY
                sum[3] += data[n][1] * Math.log(data[n][1]); // Y Log Y
                sum[4] += data[n][0] * data[n][1] * Math.log(data[n][1]); //YY Log Y
                sum[5] += data[n][0] * data[n][1]; //XY
            }
        }

        var denominator = (sum[1] * sum[2] - sum[5] * sum[5]);
        var A = Math.pow(Math.E, (sum[2] * sum[3] - sum[5] * sum[4]) / denominator);
        var B = (sum[1] * sum[4] - sum[5] * sum[3]) / denominator;

        var resultLength = data.length + extrapolate;
        var step = data[data.length - 1][0] - data[data.length - 2][0];

        for (var i = 0, len = resultLength; i < len; i++) {
            var answer = 0;
            if(typeof data[i] !== 'undefined') {
                var x = data[i][0];
            } else {
                var x = data[data.length - 1][0] + (i - data.length) * step;
            }

            var coordinate = [x, A * Math.pow(Math.E, B * x)];
            results.push(coordinate);
        }

        results.sort(function (a, b) {
            if (a[0] > b[0]) {
                return 1;
            }
            if (a[0] < b[0]) {
                return -1;
            }
            return 0;
        });

        var string = 'y = ' + Math.round(A * 100) / 100 + 'e^(' + Math.round(B * 100) / 100 + 'x)';

        return {equation: [A, B], points: results, string: string};
    }


    /**
     * Code extracted from https://github.com/Tom-Alexander/regression-js/
     * Human readable formulas:
     *
     *              N * Σ(XY) - Σ(X)
     * intercept = ---------------------
     *              N * Σ(X^2) - Σ(X)^2
     *
     * correlation = N * Σ(XY) - Σ(X) * Σ (Y) / √ (  N * Σ(X^2) - Σ(X) ) * ( N * Σ(Y^2) - Σ(Y)^2 ) ) )
     *
     */
    function _linear(data, decimalPlaces, extrapolate) {
        var sum = [0, 0, 0, 0, 0], n = 0, results = [], N = data.length;

        for (; n < data.length; n++) {
            if (data[n]['x'] != null) {
                data[n][0] = data[n].x;
                data[n][1] = data[n].y;
            }
            if (data[n][1] != null) {
                sum[0] += data[n][0]; //Σ(X)
                sum[1] += data[n][1]; //Σ(Y)
                sum[2] += data[n][0] * data[n][0]; //Σ(X^2)
                sum[3] += data[n][0] * data[n][1]; //Σ(XY)
                sum[4] += data[n][1] * data[n][1]; //Σ(Y^2)
            } else {
                N -= 1;
            }
        }

        var gradient = (N * sum[3] - sum[0] * sum[1]) / (N * sum[2] - sum[0] * sum[0]);
        var intercept = (sum[1] / N) - (gradient * sum[0]) / N;
        // var correlation = (N * sum[3] - sum[0] * sum[1]) / Math.sqrt((N * sum[2] - sum[0] * sum[0]) * (N * sum[4] - sum[1] * sum[1]));

        var resultLength = data.length + extrapolate;
        var step = data[data.length - 1][0] - data[data.length - 2][0];

        for (var i = 0, len = resultLength; i < len; i++) {
            var answer = 0;
            if(typeof data[i] !== 'undefined') {
                var x = data[i][0];
            } else {
                var x = data[data.length - 1][0] + (i - data.length) * step;
            }

            var coorY = x * gradient + intercept;
            if (decimalPlaces)
                coorY = parseFloat(coorY.toFixed(decimalPlaces));
            var coordinate = [x, coorY];
            results.push(coordinate);
        }

        results.sort(function (a, b) {
            if (a[0] > b[0]) {
                return 1;
            }
            if (a[0] < b[0]) {
                return -1;
            }
            return 0;
        });

        var string = 'y = ' + Math.round(gradient * 100) / 100 + 'x + ' + Math.round(intercept * 100) / 100;
        return {equation: [gradient, intercept], points: results, string: string};
    }

    /**
     *  Code extracted from https://github.com/Tom-Alexander/regression-js/
     */
    function _logarithmic(data, extrapolate) {
        var sum = [0, 0, 0, 0], n = 0, results = [], mean = 0;


        for (len = data.length; n < len; n++) {
            if (data[n].x != null) {
                data[n][0] = data[n].x;
                data[n][1] = data[n].y;
            }
            if (data[n][1] != null) {
                sum[0] += Math.log(data[n][0]);
                sum[1] += data[n][1] * Math.log(data[n][0]);
                sum[2] += data[n][1];
                sum[3] += Math.pow(Math.log(data[n][0]), 2);
            }
        }

        var B = (n * sum[1] - sum[2] * sum[0]) / (n * sum[3] - sum[0] * sum[0]);
        var A = (sum[2] - B * sum[0]) / n;

        var resultLength = data.length + extrapolate;
        var step = data[data.length - 1][0] - data[data.length - 2][0];

        for (var i = 0, len = resultLength; i < len; i++) {
            var answer = 0;
            if(typeof data[i] !== 'undefined') {
                var x = data[i][0];
            } else {
                var x = data[data.length - 1][0] + (i - data.length) * step;
            }

            var coordinate = [x, A + B * Math.log(x)];
            results.push(coordinate);
        }

        results.sort(function (a, b) {
            if (a[0] > b[0]) {
                return 1;
            }
            if (a[0] < b[0]) {
                return -1;
            }
            return 0;
        });

        var string = 'y = ' + Math.round(A * 100) / 100 + ' + ' + Math.round(B * 100) / 100 + ' ln(x)';

        return {equation: [A, B], points: results, string: string};
    }

    /**
     * Code extracted from https://github.com/Tom-Alexander/regression-js/
     */
    function _power(data, extrapolate) {
        var sum = [0, 0, 0, 0], n = 0, results = [];

        for (len = data.length; n < len; n++) {
            if (data[n].x != null) {
                data[n][0] = data[n].x;
                data[n][1] = data[n].y;
            }
            if (data[n][1] != null) {
                sum[0] += Math.log(data[n][0]);
                sum[1] += Math.log(data[n][1]) * Math.log(data[n][0]);
                sum[2] += Math.log(data[n][1]);
                sum[3] += Math.pow(Math.log(data[n][0]), 2);
            }
        }

        var B = (n * sum[1] - sum[2] * sum[0]) / (n * sum[3] - sum[0] * sum[0]);
        var A = Math.pow(Math.E, (sum[2] - B * sum[0]) / n);

        var resultLength = data.length + extrapolate;
        var step = data[data.length - 1][0] - data[data.length - 2][0];

        for (var i = 0, len = resultLength; i < len; i++) {
            var answer = 0;
            if(typeof data[i] !== 'undefined') {
                var x = data[i][0];
            } else {
                var x = data[data.length - 1][0] + (i - data.length) * step;
            }

            var coordinate = [x, A * Math.pow(x, B)];
            results.push(coordinate);
        }

        results.sort(function (a, b) {
            if (a[0] > b[0]) {
                return 1;
            }
            if (a[0] < b[0]) {
                return -1;
            }
            return 0;
        });

        var string = 'y = ' + Math.round(A * 100) / 100 + 'x^' + Math.round(B * 100) / 100;

        return {equation: [A, B], points: results, string: string};
    }

    /**
     * Code extracted from https://github.com/Tom-Alexander/regression-js/
     */
    function _polynomial(data, order, extrapolate) {
        if (typeof order == 'undefined') {
            order = 2;
        }
        var lhs = [], rhs = [], results = [], a = 0, b = 0, i = 0, k = order + 1;

        for (; i < k; i++) {
            for (var l = 0, len = data.length; l < len; l++) {
                if (data[l].x != null) {
                    data[l][0] = data[l].x;
                    data[l][1] = data[l].y;
                }
                if (data[l][1] != null) {
                    a += Math.pow(data[l][0], i) * data[l][1];
                }
            }
            lhs.push(a);
            a = 0;
            var c = [];
            for (var j = 0; j < k; j++) {
                for (var l = 0, len = data.length; l < len; l++) {
                    if (data[l][1]) {
                        b += Math.pow(data[l][0], i + j);
                    }
                }
                c.push(b);
                b = 0;
            }
            rhs.push(c);
        }
        rhs.push(lhs);

        var equation = gaussianElimination(rhs, k);

        var resultLength = data.length + extrapolate;
        var step = data[data.length - 1][0] - data[data.length - 2][0];
        for (var i = 0, len = resultLength; i < len; i++) {
            var answer = 0;
            var x = 0;
            if (typeof data[i] !== 'undefined') {
                x = data[i][0];
            } else {
                x = data[data.length - 1][0] + (i - data.length) * step;
            }

            for (var w = 0; w < equation.length; w++) {
                answer += equation[w] * Math.pow(x, w);
            }
            results.push([x, answer]);
        }

        results.sort(function (a, b) {
            if (a[0] > b[0]) {
                return 1;
            }
            if (a[0] < b[0]) {
                return -1;
            }
            return 0;
        });

        var string = 'y = ';

        for (var i = equation.length - 1; i >= 0; i--) {
            if (i > 1) string += Math.round(equation[i] * 100) / 100 + 'x^' + i + ' + ';
            else if (i == 1) string += Math.round(equation[i] * 100) / 100 + 'x' + ' + ';
            else string += Math.round(equation[i] * 100) / 100;
        }

        return {equation: equation, points: results, string: string};
    }

    /**
     * @author: Ignacio Vazquez
     * Based on
     * - http://commons.apache.org/proper/commons-math/download_math.cgi LoesInterpolator.java
     * - https://gist.github.com/avibryant/1151823
     */
    function _loess(data, bandwidth) {
        bandwidth = bandwidth || 0.25;

        var xval = data.map(function (pair) {
            return pair.x || pair[0];
        });
        var distinctX = array_unique(xval);
        if (2 / distinctX.length > bandwidth) {
            bandwidth = Math.min(2 / distinctX.length, 1);
            console.warn("updated bandwith to " + bandwidth);
        }

        var yval = data.map(function (pair) {
            return pair.y || pair[1];
        });

        function array_unique(values) {
            var o = {}, i, l = values.length, r = [];
            for (i = 0; i < l; i += 1) o[values[i]] = values[i];
            for (i in o) r.push(o[i]);
            return r;
        }

        function tricube(x) {
            var tmp = 1 - x * x * x;
            return tmp * tmp * tmp;
        }

        var res = [];

        var left = 0;
        var right = Math.floor(bandwidth * xval.length) - 1;

        for (var i in xval) {
            var x = xval[i];

            if (i > 0) {
                if (right < xval.length - 1 &&
                    xval[right + 1] - xval[i] < xval[i] - xval[left]) {
                    left++;
                    right++;
                }
            }
            //console.debug("left: "+left  + " right: " + right );
            var edge;
            if (xval[i] - xval[left] > xval[right] - xval[i])
                edge = left;
            else
                edge = right;
            var denom = Math.abs(1.0 / (xval[edge] - x));
            var sumWeights = 0;
            var sumX = 0, sumXSquared = 0, sumY = 0, sumXY = 0;

            var k = left;
            while (k <= right) {
                var xk = xval[k];
                var yk = yval[k];
                var dist;
                if (k < i) {
                    dist = (x - xk);
                } else {
                    dist = (xk - x);
                }
                var w = tricube(dist * denom);
                var xkw = xk * w;
                sumWeights += w;
                sumX += xkw;
                sumXSquared += xk * xkw;
                sumY += yk * w;
                sumXY += yk * xkw;
                k++;
            }

            var meanX = sumX / sumWeights;
            //console.debug(meanX);
            var meanY = sumY / sumWeights;
            var meanXY = sumXY / sumWeights;
            var meanXSquared = sumXSquared / sumWeights;

            var beta;
            if (meanXSquared == meanX * meanX)
                beta = 0;
            else
                beta = (meanXY - meanX * meanY) / (meanXSquared - meanX * meanX);

            var alpha = meanY - beta * meanX;
            res[i] = beta * x + alpha;
        }
        //console.debug(res);
        return {
            equation: "",
            points: xval.map(function (x, i) {
                return [x, res[i]];
            }),
            string: ""
        };
    }


    /**
     * Code extracted from https://github.com/Tom-Alexander/regression-js/
     */
    function gaussianElimination(a, o) {
        var i = 0, j = 0, k = 0, maxrow = 0, tmp = 0, n = a.length - 1, x = new Array(o);
        for (i = 0; i < n; i++) {
            maxrow = i;
            for (j = i + 1; j < n; j++) {
                if (Math.abs(a[i][j]) > Math.abs(a[i][maxrow]))
                    maxrow = j;
            }
            for (k = i; k < n + 1; k++) {
                tmp = a[k][i];
                a[k][i] = a[k][maxrow];
                a[k][maxrow] = tmp;
            }
            for (j = i + 1; j < n; j++) {
                for (k = n; k >= i; k--) {
                    a[k][j] -= a[k][i] * a[i][j] / a[i][i];
                }
            }
        }
        for (j = n - 1; j >= 0; j--) {
            tmp = 0;
            for (k = j + 1; k < n; k++)
                tmp += a[k][j] * x[k];
            x[j] = (a[n][j] - tmp) / a[j][j];
        }
        return (x);
    }

    /**
     * @author Ignacio Vazquez
     * See http://en.wikipedia.org/wiki/Coefficient_of_determination for theaorical details
     */
    function coefficientOfDetermination(data, pred) {
        // Sort the initial data { pred array (model's predictions) is sorted  }
        // The initial data must be sorted in the same way in order to calculate the coefficients
        data.sort(function (a, b) {
            if (a[0] > b[0]) {
                return 1;
            }
            if (a[0] < b[0]) {
                return -1;
            }
            return 0;
        });

        // Calc the mean
        var mean = 0;
        var N = data.length;
        for (var i = 0; i < data.length; i++) {
            if (data[i][1] != null) {
                mean += data[i][1];
            } else {
                N--;
            }
        }
        mean /= N;

        // Calc the coefficent of determination
        var SSE = 0;
        var SSYY = 0;
        for (var i = 0; i < data.length; i++) {
            if (data[i][1] != null) {
                SSYY += Math.pow(data[i][1] - pred[i][1], 2);
                SSE += Math.pow(data[i][1] - mean, 2);
            }
        }
        return 1 - ( SSYY / SSE);
    }

    function standardError(data, pred) {
        var SE = 0, N = data.length;

        for (var i = 0; i < data.length; i++) {
            if (data[i][1] != null) {
                SE += Math.pow(data[i][1] - pred[i][1], 2);
            } else {
                N--;
            }
        }
        SE = Math.sqrt(SE / (N - 2));

        return SE;
    }

    function _round(number, decimalPlaces) {
        var decimalFactor = Math.pow(10, decimalPlaces);
        return Math.round(number * decimalFactor) / decimalFactor;
    }
    function replaceRegressionPlaceholders(text, regression) {
        return text
            .replace("%r2", regression.rSquared)
            .replace("%r", regression.rValue)
            .replace("%eq", regression.string)
            .replace("%se", regression.standardError);
    }
}));
```
