## Supplementary material for "Exploring natural odour landscapes: A case study with implications for human-biting insects": Figure S4: motion.js

```
/**
 * @license http://creativecommons.org/licenses/by-sa/4.0/ Creative Commons Attribution-ShareAlike 4.0 International (CC BY-SA 4.0)
 * @author  Lars Cabrera
 * @version 1.0.9
 */

// JSLint options:
/*global Highcharts, window*/

(function (H) {
    // Check if object is array
    function isArray(obj) {
        return Object.prototype.toString.call(obj) === '[object Array]';
    }


    // Sets up motion ready to use
    function Motion(chart) {
        var motion = this;

        this.chart = chart;
        this.paused = true;
        this.options = H.merge(this.defaultOptions, this.chart.options.motion);
        this.dataSeries = [];
        this.dataLength = 0;
        motion.options.series = H.splat(motion.options.series);
        Highcharts.each(this.chart.series, function (series, index) {
            if (motion.options.series.indexOf(index) >= 0) {
                motion.dataSeries[index] = series;
                for (var i = 0; i < series.data.length; i++) {
                    if (series.data[i].sequence) {
                        motion.dataLength = Math.max(motion.dataLength, series.data[i].sequence.length);
                    }
                }
            }
        });

        // Play-controls HTML-div
        this.playControls = H.createElement('div', {
            id: 'play-controls'
        }, null, this.chart.renderTo, null);

        // Play/pause HTML-button
        this.playPauseBtn = H.createElement('button', {
            id: 'play-pause-button',
            title: 'play'
        }, null, this.playControls, null);
        this.playPauseBtn.className = this.options.playIcon;

        // Play-range HTML-input
        this.playRange = H.createElement('input', {
            id: 'play-range',
            type: 'range',
            min: 0,
            max: this.dataLength - 1,
            step: this.options.magnet.step
        }, null, this.playControls, null);
        // Important: max must be set before value in order to allow for
        // higher numbers than 100.
        this.playRange.value = H.pick(this.options.startIndex, this.dataLength - 1);

        // Play-range HTML-output
        this.playOutput = H.createElement('label', {
            id: 'play-output',
            name: this.options.axisLabel
        }, null, this.playControls, null);
        if (isArray(this.options.labels)) {
            this.playOutput.innerHTML = this.options.labels[this.dataLength - 1] || '';
        } else {
            this.playOutput.innerHTML = this.dataLength - 1;
        }

        // Common key event handler function
        function handleKeyEvents(e) {
            e = e || window.event;
            switch (e.which) {
            case 32: // Space
                motion.togglePlayPause();
                break;
            case 37: // Left
                motion.playRange.value = motion.round(parseFloat(motion.playRange.value) - 1);
                motion.updateChart(motion.playRange.value);
                break;
            case 39: // Right
                motion.playRange.value = motion.round(parseFloat(motion.playRange.value) + 1);
                motion.updateChart(motion.playRange.value);
                break;
            default:
                return;
            }
            e.preventDefault();
        }

        // Bind controls to events
        Highcharts.addEvent(this.playPauseBtn, 'click', function () {
            motion.togglePlayPause();
        });
        Highcharts.addEvent(this.playRange, 'mouseup', function () {
            motion.attractToStep();
        });
        Highcharts.addEvent(this.playRange, 'input', function () {
            motion.updateChart(this.value);
        });

        // Request focus to the controls when clicking on controls div
        Highcharts.addEvent(this.playControls, 'click', function () {
            motion.playRange.focus();
        });
        // Bind keys to events
        Highcharts.addEvent(this.playPauseBtn, 'keydown', handleKeyEvents);
        Highcharts.addEvent(this.playRange, 'keydown', handleKeyEvents);

        // Initial value
        this.inputValue = parseFloat(this.playRange.value);

        // Initial update
        this.updateChart(this.inputValue);

        // Auto-play
        if (this.options.autoPlay) {
            this.play();
        }
    }

    // Default options for Motion
    Motion.prototype.defaultOptions = {
        enabled: true,
        axisLabel: 'year',
        autoPlay : false,
        loop: false,
        series: 0,
        updateInterval: 10,
        magnet: {
            round: 'round',
            step: 0.01
        },
        playIcon: "fa fa-play",
        pauseIcon: "fa fa-pause"
    };

    // Toggles between Play and Pause states, and makes calls to changeButtonType()
    // From http://www.creativebloq.com/html5/build-custom-html5-video-player-9134473
    Motion.prototype.togglePlayPause = function () {
        this[this.paused ? 'play' : 'pause']();
    };

    // Plays the motion, continuously updating the chart
    Motion.prototype.play = function () {
        var motion = this;
        if (this.paused && parseFloat(this.playRange.value) === parseFloat(this.playRange.max)) {
            this.reset();
        }
        this.changeButtonType('pause');
        this.paused = false;
        this.timer = setInterval(function () {
            motion.playUpdate();
        }, this.options.updateInterval);
    };

    // Pauses the motion, which stops updating the chart
    Motion.prototype.pause = function () {
        this.changeButtonType('play');
        this.paused = true;
        window.clearInterval(this.timer);
        this.attractToStep();
    };

    // Resets the motion and updates the chart. Does not pause
    Motion.prototype.reset = function () {
        this.playRange.value = this.playRange.min;
        this.updateChart(this.playRange.value);
    };

    // Updates a button's title, innerHTML and CSS class to a certain value
    Motion.prototype.changeButtonType = function (value) {
        this.playPauseBtn.title = value;
        this.playPauseBtn.className = value + " ";
        if (value == 'play') {
            this.playPauseBtn.className += this.options.playIcon;
        } else if (value == 'pause') {
            this.playPauseBtn.className += this.options.pauseIcon;;
        }
    };

    // Called continuously while playing
    Motion.prototype.playUpdate = function () {
        if (!this.paused) {
            this.inputValue = parseFloat(this.playRange.value);
            this.playRange.value = this.inputValue + this.options.magnet.step;
            this.attractToStep();
            this.updateChart(this.playRange.value); // Use playRange.value to get updated value
            if (this.playRange.value >= parseFloat(this.playRange.max)) { // Auto-pause
                if (this.options.loop) {
                    this.reset();
                } else {
                    this.pause();
                }
            }
        }
    };

    // Updates chart data and redraws the chart
    Motion.prototype.updateChart = function (inputValue) {
        var seriesKey,
            series,
            point,
            roundedInput = this.round(inputValue),
            i;
        if (this.currentAxisValue !== roundedInput) {
            this.currentAxisValue = roundedInput;
            this.chart.options.motion.startIndex = roundedInput;
            for (seriesKey in this.dataSeries) {
                if (this.dataSeries.hasOwnProperty(seriesKey)) {
                    series = this.dataSeries[seriesKey];
                    for (i = 0; i < series.data.length; i++) {
                        point = series.data[i];
                        try {
                            if (point.sequence) {
                                point.update(point.sequence[roundedInput], false, false);
                            }
                        } catch (e) {
                            console.error('Error:', e, ' \nat point:', point, ' \nwith new value:', point.sequence[roundedInput]);
                        }
                    }
                }
            }
            this.chart.redraw();
            this.attractToStep();
        }
    };

    // Moves output value to data point
    Motion.prototype.attractToStep = function () {
        if (isArray(this.options.labels)) {
            this.playOutput.innerHTML = this.options.labels[this.round(this.playRange.value)] || '';
        } else {
            this.playOutput.innerHTML = this.round(this.playRange.value);
        }
    };

    // Returns an integer rounded up, down or even depending on
    // motion.magnet.round options.
    Motion.prototype.round = function (number) {
        return Math[this.options.magnet.round](number);
    };

    // Initiates motion automatically if motion options object exists and
    // is not disabled
    H.Chart.prototype.callbacks.push(function (chart) {
        if (chart.options.motion && chart.options.motion.enabled) {
            chart.motion = new Motion(chart);
        }
    });

    H.Motion = Motion;
}(Highcharts));
```
