## Supplementary material for "Exploring natural odour landscapes: A case study with implications for human-biting insects": Figure S4: multicolor_series.js

```
/**
* Multicolor Series v2.2.7(2020-06-23)
*
* (c) 2012-2016 Black Label
*
* License: Creative Commons Attribution (CC)
*/
/* global Highcharts window document module:true */
/**
 * @fileoverview
 * @suppress {checkTypes}
 */
(function (factory) {
	if (typeof module === 'object' && module.exports) {
		module.exports = factory;
	} else {
		factory(Highcharts);
	}
}(function (H) {
	var each = H.each,
		seriesTypes = H.seriesTypes,
		pick = H.pick,
		UNDEFINED,
		NORMAL_STATE = '',
		VISIBLE = 'visible',
		HIDDEN = 'hidden',
		PREFIX = 'highcharts-',
		NONE = 'none',
		hasTouch = document.documentElement.ontouchstart !== UNDEFINED,
		TRACKER_FILL = 'rgba(192,192,192,' + (H.hasSVG ? 0.0001 : 0.002) + ')', // invisible but clickable
		M = 'M',
		L = 'L';
	
	// handle unsorted data, throw error anyway
	function error(code, stop) {
		var msg = 'Highcharts error #' + code + ': www.highcharts.com/errors/' + code;
		if (stop) {
			throw msg;
		} else if (window.console) {
			console.log(msg); // eslint-disable-line
		}
	}
	
	/**
	If replacing L and M in tracker will be necessary use that getPath():
	
	function getPath(arr){
	var ret = [];
	each(arr, function(el, ind) {
	var len = el[0].length;
	for(var i = 0; i < len; i++){
	var p = el[0][i];
	if(p == M && ind != 0 && i == 0) {
	p = L;
	}
	ret.push(p);
	}
	});
	return ret;
	}
	**/
	
	
	function getPath(arr) {
		var ret = [];
		each(arr, function (el) {
			ret = ret.concat(el[0]);
		});
		return ret;
	}

	/**
	 * Return the graph path of a segment - compatibility with 4.2.3+
	 * @param {Object} segment of the path
	 * @returns {Array} Path (SVG)
	 */
	H.Series.prototype.getSegmentPath = function (segment) {
		var series = this,
			segmentPath = [],
			step = series.options.step;

		// build the segment line
		each(segment, function (point, i) {
			var plotX = point.plotX,
				plotY = point.plotY,
				lastPoint;

			if (series.getPointSpline) {
				// generate the spline as defined in the SplineSeries object
				segmentPath.push.apply(segmentPath, series.getPointSpline(segment, point, i));
			} else {
				// moveTo or lineTo
				segmentPath.push(i ? L : M);

				// step line?
				if (step && i) {
					lastPoint = segment[i - 1];
					if (step === 'right') {
						segmentPath.push(
							lastPoint.plotX,
							plotY,
							L
						);
					} else if (step === 'center') {
						segmentPath.push(
							(lastPoint.plotX + plotX) / 2,
							lastPoint.plotY,
							L,
							(lastPoint.plotX + plotX) / 2,
							plotY,
							L
						);
					} else {
						segmentPath.push(
							plotX,
							lastPoint.plotY,
							L
						);
					}
				}

				// normal line to next point
				segmentPath.push(
					point.plotX,
					point.plotY
				);
			}
		});

		return segmentPath;
	};

	/**
	*
	*   ColoredLine series type
	*
	**/
	
	seriesTypes.coloredline = H.extendClass(seriesTypes.line);
	
	H.seriesTypes.coloredline.prototype.processData = function (force) {
		var series = this,
			processedXData = series.xData, // copied during slice operation below
			processedYData = series.yData,
			cropStart = 0,
			cropped,
			distance,
			closestPointRange,
			xAxis = series.xAxis,
			i, // loop variable
			options = series.options,
			isCartesian = series.isCartesian;
		
		// If the series data or axes haven't changed, don't go through this. Return false to pass
		// the message on to override methods like in data grouping.
		if (isCartesian && !series.isDirty && !xAxis.isDirty && !series.yAxis.isDirty && !force) {
			return false;
		}
		
		// Find the closest distance between processed points
		for (i = processedXData.length - 1; i >= 0; i--) {
			distance = processedXData[i] - processedXData[i - 1];
			if (distance > 0 && (closestPointRange === UNDEFINED || distance < closestPointRange)) {
				closestPointRange = distance;
				
				// Unsorted data is not supported by the line tooltip, as well as data grouping and
				// navigation in Stock charts (#725) and width calculation of columns (#1900)
			} else if (distance < 0 && series.requireSorting) {
				error(15);
			}
		}
		
		// Record the properties
		series.cropped = cropped; // undefined or true
		series.cropStart = cropStart;
		series.processedXData = processedXData;
		series.processedYData = processedYData;
		
		if (options.pointRange === null) { // null means auto, as for columns, candlesticks and OHLC
			series.pointRange = closestPointRange || 1;
		}
		series.closestPointRange = closestPointRange;
		return true;
	};
	
	H.seriesTypes.coloredline.prototype.drawTracker = function () {
		var series = this,
			options = series.options,
			trackByArea = options.trackByArea,
			trackerPath = [].concat(trackByArea ? series.areaPath : getPath(series.graphPath)),
			trackerPathLength = trackerPath.length,
			chart = series.chart,
			pointer = chart.pointer,
			renderer = chart.renderer,
			snap = chart.options.tooltip.snap,
			tracker = series.tracker,
			cursor = options.cursor,
			css = cursor && { cursor: cursor },
			singlePoints = series.singlePoints,
			singlePoint,
			i,
			onMouseOver;

		onMouseOver = function () {
			if (chart.hoverSeries !== series) {
				series.onMouseOver();
			}
		};
		// Extend end points. A better way would be to use round linecaps,
		// but those are not clickable in VML.
		if (trackerPathLength && !trackByArea) {
			i = trackerPathLength + 1;
			while (i--) {
				if (trackerPath[i] === M) { // extend left side
					trackerPath.splice(i + 1, 0, trackerPath[i + 1] - snap, trackerPath[i + 2], L);
				}
				if ((i && trackerPath[i] === M) || i === trackerPathLength) { // extend right side
					trackerPath.splice(i, 0, L, trackerPath[i - 2] + snap, trackerPath[i - 1]);
				}
			}
		}
		
		// handle single points
		for (i = 0; i < singlePoints.length; i++) {
			singlePoint = singlePoints[i];
			if (singlePoint.plotX && singlePoint.plotY) {
				trackerPath.push(M, singlePoint.plotX - snap, singlePoint.plotY,
					L, singlePoint.plotX + snap, singlePoint.plotY);
			}
		}
		
		// draw the tracker
		if (tracker) {
			tracker.attr({ d: trackerPath });
		} else { // create
			series.tracker = renderer.path(trackerPath)
			.attr({
				'stroke-linejoin': 'round', // #1225
				visibility: series.visible ? VISIBLE : HIDDEN,
				stroke: TRACKER_FILL,
				fill: trackByArea ? TRACKER_FILL : NONE,
				'stroke-width': options.lineWidth + (trackByArea ? 0 : 2 * snap),
				zIndex: 2
			})
			.add(series.group);
			
			// The tracker is added to the series group, which is clipped, but is covered
			// by the marker group. So the marker group also needs to capture events.
			each([series.tracker, series.markerGroup], function (track) {
				track.addClass(PREFIX + 'tracker')
				.on('mouseover', onMouseOver)
				.on('mouseout', function (e) { pointer.onTrackerMouseOut(e); })
				.css(css);
				
				if (hasTouch) {
					track.on('touchstart', onMouseOver);
				}
			});
		}
		
	};
	
	H.seriesTypes.coloredline.prototype.setState = function (state) {
		var series = this,
			options = series.options,
			graph = series.graph,
			stateOptions = options.states,
			lineWidth = options.lineWidth,
			attribs;
		
		state = state || NORMAL_STATE;
		
		if (series.state !== state) {
			series.state = state;
			
			if (stateOptions[state] && stateOptions[state].enabled === false) {
				return;
			}
			
			if (state) {
				lineWidth = stateOptions[state].lineWidth || lineWidth + 1;
			}
			
			if (graph && !graph.dashstyle) { // hover is turned off for dashed lines in VML
				attribs = {
					'stroke-width': lineWidth
				};
				// use attr because animate will cause any other animation on the graph to stop
				each(graph, function (seg) {
					seg.attr(attribs);
				});
			}
		}
	};
	
	/**
	* The main change to get multi color isFinite changes segments array.
	* From array of points to object with color and array of points.
	* @returns {undefined}
	**/
	H.seriesTypes.coloredline.prototype.getSegments = function () {
		var series = this,
			lastColor = 0,
			segments = [],
			i,
			points = series.points,
			pointsLength = points.length;
		
		if (pointsLength) { // no action required for []
			
			// if connect nulls, just remove null points
			if (series.options.connectNulls) {
				// iterate backwars for secure point removal
				for (i = pointsLength - 1; i >= 0; --i) {
					if (points[i].y === null) {
						points.splice(i, 1);
					}
				}
				pointsLength = points.length;
				
				each(points, function (point, j) {
					if (j > 0 && points[j].segmentColor !== points[j - 1].segmentColor) {
						segments.push({
							points: points.slice(lastColor, j + 1),
							color: points[j - 1].segmentColor
						});
						lastColor = j;
					}
				});

				if (pointsLength) {
					// add the last segment (only single-point last segement is added)
					if (lastColor !== pointsLength - 1) {
						segments.push({
							points: points.slice(lastColor, pointsLength),
							color: points[pointsLength - 1].segmentColor
						});
					}
				}
				
				if (points.length && segments.length === 0) {
					segments = [points];
				}
				
				// else, split on null points or different colors
			} else {
				var previousColor = null;
				each(points, function (point, j) {
					var colorChanged = j > 0 && (point.y === null || points[j - 1].y === null || (point.segmentColor !== points[j - 1].segmentColor && points[j].segmentColor !== previousColor)),
						colorExists = points[j - 1] && points[j - 1].segmentColor && points[j - 1].y !== null ? true : false;
					
					if (colorChanged) {
						var p = points.slice(lastColor, j + 1);
						if (p.length > 0) {
							// do not create segments with null ponits
							each(p, function (pointObject, k) {
								if (pointObject.y === null) {
									// remove null points (might be on edges)
									p.splice(k, 1);
								}
							});
							
							segments.push({
								points: p,
								color: colorExists ? points[j - 1].segmentColor : previousColor
							});
							lastColor = j;
						}
					} else if (j === pointsLength - 1) {
						var next = j + 1;
						if (point.y === null) {
							next--;
						}
						p = points.slice(lastColor, next);
						if (p.length > 0) {
							// do not create segments with null ponits
							each(p, function (pointObject, k) {
								if (pointObject.y === null) {
									// remove null points (might be on edges)
									p.splice(k, 1);
								}
							});
							segments.push({
								points: p,
								color: colorExists ? points[j - 1].segmentColor : previousColor
							});
							lastColor = j;
						}
						
					}
					
					// store previous color
					if (point) {
						previousColor = point.segmentColor;
					}
				});
			}
		}
		// register it
		series.segments = segments;
	};
	
	H.seriesTypes.coloredline.prototype.getGraphPath = function () {
		// var ret = f.apply(this, Array.prototype.slice.call(arguments, 1));
		var series = this,
			graphPath = [],
			segmentPath,
			singlePoints = []; // used in drawTracker
		// Divide into segments and build graph and area paths
		each(series.segments, function (segment) {
			segmentPath = series.getSegmentPath(segment.points);
			// add the segment to the graph, or a single point for tracking
			if (segment.points.length > 1) {
				graphPath.push([segmentPath, segment.color]);
			} else {
				singlePoints.push(segment.points);
			}
		});
		
		// Record it for use in drawGraph and drawTracker, and return graphPath
		series.singlePoints = singlePoints;
		series.graphPath = graphPath;
		
		return graphPath;
	};
	
	H.seriesTypes.coloredline.prototype.drawGraph = function () {
		var series = this,
			options = series.options,
			props = [['graph', options.lineColor || series.color]],
			lineWidth = options.lineWidth,
			dashStyle = options.dashStyle,
			roundCap = options.linecap !== 'square',
			graphPath = series.getGraphPath(),
			graphPathLength = graphPath.length,
			graphSegmentsLength = 0;
		
		function getSegment(segment, prop, i) {
			var attribs = {
					stroke: prop[1],
					'stroke-width': lineWidth,
					fill: 'none',
					zIndex: 1 // #1069
				},
				item;
			if (dashStyle) {
				attribs.dashstyle = dashStyle;
			} else if (roundCap) {
				attribs['stroke-linecap'] = attribs['stroke-linejoin'] = 'round';
			}
			if (segment[1]) {
				attribs.stroke = segment[1];
			}
			
			item = series.chart.renderer.path(segment[0])
			.attr(attribs)
			.add(series.group);


			if (item.shadow) {
				item.shadow(!i && options.shadow);
			}
			
			return item;
		}

		// draw the graph
		each(props, function (prop, i) {
			var graphKey = prop[0],
				graph = series[graphKey],
				g;
			
			if (graph) { // cancel running animations, #459
				// do we have animation
				each(graphPath, function (segment, j) {
					// update color and path
					
					if (series[graphKey][j]) {
						series[graphKey][j].attr({ d: segment[0], stroke: segment[1] });
					} else {
						series[graphKey][j] = getSegment(segment, prop, i);
					}
				});
				
			} else if (graphPath.length) { // #1487
				graph = [];
				each(graphPath, function (segment, j) {
					graph[j] = getSegment(segment, prop, i);
				});
				series[graphKey] = graph;
				// add destroying elements
				series[graphKey].destroy = function () {
					for (g in series[graphKey]) {  // eslint-disable-line
						var el = series[graphKey][g];
						if (el && el.destroy) {
							el.destroy();
						}
					}
				};
			}
			// Checks if series.graph exists. #3
			graphSegmentsLength = (series.graph && series.graph.length) || -1;
			
			for (var j = graphSegmentsLength; j >= graphPathLength; j--) {
				if (series[graphKey][j]) {
					series[graphKey][j].destroy();
					series[graphKey].splice(j, 1);
				}
			}
		});
	};

	H.wrap(seriesTypes.coloredline.prototype, 'translate', function (proceed) {
		proceed.apply(this, [].slice.call(arguments, 1));
		if (this.getSegments) {
			this.getSegments();
		}
	});
	
	H.wrap(H.Series.prototype, 'applyZones', function (proceed) {
		var series = this,
			parts = ['area', 'graph'];

		parts.forEach(function (part) {
			var shape = series[part];

			if (shape && H.isArray(shape)) {
				shape.show = function () {
					shape.forEach(function (subGraph) {
						subGraph.show(true);
					});
				};
				shape.hide = function () {
					shape.forEach(function (subGraph) {
						subGraph.hide();
					});
				};
			}
		});
	
		return proceed.apply(this, Array.prototype.slice.call(arguments, 1));
	});

	H.wrap(seriesTypes.coloredline.prototype, 'destroy', function (proceed) {
		// destroy all parts
		this.graph.destroy();
		proceed.apply(this, Array.prototype.slice.call(arguments, 1));
	});
	
	/**
	*
	*   ColoredArea series type
	*
	**/
	seriesTypes.coloredarea = H.extendClass(seriesTypes.coloredline);
	
	H.seriesTypes.coloredarea.prototype.init = function (chart, options) {
		options.threshold = options.threshold || null;
		H.Series.prototype.init.call(this, chart, options);
	};
	
	H.seriesTypes.coloredarea.prototype.closeSegment = function (path, segment, translatedThreshold) {
		path.push(
			L,
			segment[segment.length - 1].plotX,
			translatedThreshold,
			L,
			segment[0].plotX,
			translatedThreshold
		);
	};
	
	H.seriesTypes.coloredarea.prototype.drawGraph = function (f) {
		H.seriesTypes.coloredline.prototype.drawGraph.call(this, f);
		var series = this,
			options = this.options,
			props = [['graph', options.lineColor || series.color]];

		each(props, function (prop) {
			var graphKey = prop[0],
				graph = series[graphKey];
			
			if (graph) { // cancel running animations, #459
				// do we have animation
				each(series.graphPath, function (segment, j) {
					// update color and path
					
					if (series[graphKey][j]) {
						series[graphKey][j].attr({ fill: segment[1] });
					}
				});
				
			}
		});
	};

	/**
	* Extend the base Series getSegmentPath method by adding the path for the area.
	* This path is pushed to the series.areaPath property.
	* @param {Object} segment of the path
	* @returns {Array} Path (SVG)
	**/
	H.seriesTypes.coloredarea.prototype.getSegmentPath = function (segment) {
		var segmentPath = H.Series.prototype.getSegmentPath.call(this, segment), // call base method
			areaSegmentPath = [].concat(segmentPath), // work on a copy for the area path
			i,
			options = this.options,
			segLength = segmentPath.length,
			translatedThreshold = this.yAxis.getThreshold(options.threshold), // #2181
			yBottom;

		if (segLength === 3) { // for animation from 1 to two points
			areaSegmentPath.push(L, segmentPath[1], segmentPath[2]);
		}
		if (options.stacking && !this.closedStacks) {
			for (i = segment.length - 1; i >= 0; i--) {

				yBottom = pick(segment[i].yBottom, translatedThreshold);

				// step line?
				if (i < segment.length - 1 && options.step) {
					areaSegmentPath.push(segment[i + 1].plotX, yBottom);
				}
				areaSegmentPath.push(segment[i].plotX, yBottom);
			}
		} else { // follow zero line back
			this.closeSegment(areaSegmentPath, segment, translatedThreshold);
		}
		return areaSegmentPath;
	};
	
	H.seriesTypes.coloredarea.prototype.getGraphPath = function () {
		var series = this,
			graphPath = [],
			segmentPath,
			singlePoints = []; // used in drawTracker
		// Divide into segments and build graph and area paths
		
		this.areaPath = [];
		each(series.segments, function (segment) {
			segmentPath = series.getSegmentPath(segment.points);
			// add the segment to the graph, or a single point for tracking
			if (segment.points.length > 1) {
				graphPath.push([segmentPath, segment.color]);
			} else {
				singlePoints.push(segment.points);
			}
		});
		
		// Record it for use in drawGraph and drawTracker, and return graphPath
		series.singlePoints = singlePoints;
		series.graphPath = graphPath;
		return graphPath;
	
	};
	
	H.seriesTypes.coloredarea.prototype.drawLegendSymbol = H.LegendSymbolMixin.drawRectangle;
}));
```
