## Supplementary material for "Exploring natural odour landscapes: A case study with implications for human-biting insects": Figure S4: proj4.js

```
!function(a){if("object"==typeof exports&&"undefined"!=typeof module)module.exports=a();else if("function"==typeof define&&define.amd)define([],a);else{var b;b="undefined"!=typeof window?window:"undefined"!=typeof global?global:"undefined"!=typeof self?self:this,b.proj4=a()}}(function(){return function a(b,c,d){function e(g,h){if(!c[g]){if(!b[g]){var i="function"==typeof require&&require;if(!h&&i)return i(g,!0);if(f)return f(g,!0);var j=new Error("Cannot find module '"+g+"'");throw j.code="MODULE_NOT_FOUND",j}var k=c[g]={exports:{}};b[g][0].call(k.exports,function(a){var c=b[g][1][a];return e(c?c:a)},k,k.exports,a,b,c,d)}return c[g].exports}for(var f="function"==typeof require&&require,g=0;g<d.length;g++)e(d[g]);return e}({"./includedProjections":[function(a,b,c){var d=[a("./lib/projections/tmerc"),a("./lib/projections/utm"),a("./lib/projections/sterea"),a("./lib/projections/stere"),a("./lib/projections/somerc"),a("./lib/projections/omerc"),a("./lib/projections/lcc"),a("./lib/projections/krovak"),a("./lib/projections/cass"),a("./lib/projections/laea"),a("./lib/projections/aea"),a("./lib/projections/gnom"),a("./lib/projections/cea"),a("./lib/projections/eqc"),a("./lib/projections/poly"),a("./lib/projections/nzmg"),a("./lib/projections/mill"),a("./lib/projections/sinu"),a("./lib/projections/moll"),a("./lib/projections/eqdc"),a("./lib/projections/vandg"),a("./lib/projections/aeqd"),a("./lib/projections/ortho")];b.exports=function(proj4){d.forEach(function(a){proj4.Proj.projections.add(a)})}},{"./lib/projections/aea":40,"./lib/projections/aeqd":41,"./lib/projections/cass":42,"./lib/projections/cea":43,"./lib/projections/eqc":44,"./lib/projections/eqdc":45,"./lib/projections/gnom":47,"./lib/projections/krovak":48,"./lib/projections/laea":49,"./lib/projections/lcc":50,"./lib/projections/mill":53,"./lib/projections/moll":54,"./lib/projections/nzmg":55,"./lib/projections/omerc":56,"./lib/projections/ortho":57,"./lib/projections/poly":58,"./lib/projections/sinu":59,"./lib/projections/somerc":60,"./lib/projections/stere":61,"./lib/projections/sterea":62,"./lib/projections/tmerc":63,"./lib/projections/utm":64,"./lib/projections/vandg":65}],1:[function(a,b,c){function Point(a,b,c){if(!(this instanceof Point))return new Point(a,b,c);if(Array.isArray(a))this.x=a[0],this.y=a[1],this.z=a[2]||0;else if("object"==typeof a)this.x=a.x,this.y=a.y,this.z=a.z||0;else if("string"==typeof a&&"undefined"==typeof b){var d=a.split(",");this.x=parseFloat(d[0],10),this.y=parseFloat(d[1],10),this.z=parseFloat(d[2],10)||0}else this.x=a,this.y=b,this.z=c||0;console.warn("proj4.Point will be removed in version 3, use proj4.toPoint")}var d=a("mgrs");Point.fromMGRS=function(a){return new Point(d.toPoint(a))},Point.prototype.toMGRS=function(a){return d.forward([this.x,this.y],a)},b.exports=Point},{mgrs:68}],2:[function(a,b,c){function Projection(a,b){if(!(this instanceof Projection))return new Projection(a);b=b||function(a){if(a)throw a};var c=d(a);if("object"!=typeof c)return void b(a);var f=g(c),h=Projection.projections.get(f.projName);h?(e(this,f),e(this,h),this.init(),b(null,this)):b(a)}var d=a("./parseCode"),e=a("./extend"),f=a("./projections"),g=a("./deriveConstants");Projection.projections=f,Projection.projections.start(),b.exports=Projection},{"./deriveConstants":33,"./extend":34,"./parseCode":37,"./projections":39}],3:[function(a,b,c){b.exports=function(a,b,c){var d,e,f,g=c.x,h=c.y,i=c.z||0;for(f=0;3>f;f++)if(!b||2!==f||void 0!==c.z)switch(0===f?(d=g,e="x"):1===f?(d=h,e="y"):(d=i,e="z"),a.axis[f]){case"e":c[e]=d;break;case"w":c[e]=-d;break;case"n":c[e]=d;break;case"s":c[e]=-d;break;case"u":void 0!==c[e]&&(c.z=d);break;case"d":void 0!==c[e]&&(c.z=-d);break;default:return null}return c}},{}],4:[function(a,b,c){var d=Math.PI/2,e=a("./sign");b.exports=function(a){return Math.abs(a)<d?a:a-e(a)*Math.PI}},{"./sign":21}],5:[function(a,b,c){var d=2*Math.PI,e=3.14159265359,f=a("./sign");b.exports=function(a){return Math.abs(a)<=e?a:a-f(a)*d}},{"./sign":21}],6:[function(a,b,c){b.exports=function(a){return Math.abs(a)>1&&(a=a>1?1:-1),Math.asin(a)}},{}],7:[function(a,b,c){b.exports=function(a){return 1-.25*a*(1+a/16*(3+1.25*a))}},{}],8:[function(a,b,c){b.exports=function(a){return.375*a*(1+.25*a*(1+.46875*a))}},{}],9:[function(a,b,c){b.exports=function(a){return.05859375*a*a*(1+.75*a)}},{}],10:[function(a,b,c){b.exports=function(a){return a*a*a*(35/3072)}},{}],11:[function(a,b,c){b.exports=function(a,b,c){var d=b*c;return a/Math.sqrt(1-d*d)}},{}],12:[function(a,b,c){b.exports=function(a,b,c,d,e){var f,g;f=a/b;for(var h=0;15>h;h++)if(g=(a-(b*f-c*Math.sin(2*f)+d*Math.sin(4*f)-e*Math.sin(6*f)))/(b-2*c*Math.cos(2*f)+4*d*Math.cos(4*f)-6*e*Math.cos(6*f)),f+=g,Math.abs(g)<=1e-10)return f;return NaN}},{}],13:[function(a,b,c){var d=Math.PI/2;b.exports=function(a,b){var c=1-(1-a*a)/(2*a)*Math.log((1-a)/(1+a));if(Math.abs(Math.abs(b)-c)<1e-6)return 0>b?-1*d:d;for(var e,f,g,h,i=Math.asin(.5*b),j=0;30>j;j++)if(f=Math.sin(i),g=Math.cos(i),h=a*f,e=Math.pow(1-h*h,2)/(2*g)*(b/(1-a*a)-f/(1-h*h)+.5/a*Math.log((1-h)/(1+h))),i+=e,Math.abs(e)<=1e-10)return i;return NaN}},{}],14:[function(a,b,c){b.exports=function(a,b,c,d,e){return a*e-b*Math.sin(2*e)+c*Math.sin(4*e)-d*Math.sin(6*e)}},{}],15:[function(a,b,c){b.exports=function(a,b,c){var d=a*b;return c/Math.sqrt(1-d*d)}},{}],16:[function(a,b,c){var d=Math.PI/2;b.exports=function(a,b){for(var c,e,f=.5*a,g=d-2*Math.atan(b),h=0;15>=h;h++)if(c=a*Math.sin(g),e=d-2*Math.atan(b*Math.pow((1-c)/(1+c),f))-g,g+=e,Math.abs(e)<=1e-10)return g;return-9999}},{}],17:[function(a,b,c){var d=1,e=.25,f=.046875,g=.01953125,h=.01068115234375,i=.75,j=.46875,k=.013020833333333334,l=.007120768229166667,m=.3645833333333333,n=.005696614583333333,o=.3076171875;b.exports=function(a){var b=[];b[0]=d-a*(e+a*(f+a*(g+a*h))),b[1]=a*(i-a*(f+a*(g+a*h)));var c=a*a;return b[2]=c*(j-a*(k+a*l)),c*=a,b[3]=c*(m-a*n),b[4]=c*a*o,b}},{}],18:[function(a,b,c){var d=a("./pj_mlfn"),e=1e-10,f=20;b.exports=function(a,b,c){for(var g=1/(1-b),h=a,i=f;i;--i){var j=Math.sin(h),k=1-b*j*j;if(k=(d(h,j,Math.cos(h),c)-a)*(k*Math.sqrt(k))*g,h-=k,Math.abs(k)<e)return h}return h}},{"./pj_mlfn":19}],19:[function(a,b,c){b.exports=function(a,b,c,d){return c*=b,b*=b,d[0]*a-c*(d[1]+b*(d[2]+b*(d[3]+b*d[4])))}},{}],20:[function(a,b,c){b.exports=function(a,b){var c;return a>1e-7?(c=a*b,(1-a*a)*(b/(1-c*c)-.5/a*Math.log((1-c)/(1+c)))):2*b}},{}],21:[function(a,b,c){b.exports=function(a){return 0>a?-1:1}},{}],22:[function(a,b,c){b.exports=function(a,b){return Math.pow((1-a)/(1+a),b)}},{}],23:[function(a,b,c){b.exports=function(a){var b={x:a[0],y:a[1]};return a.length>2&&(b.z=a[2]),a.length>3&&(b.m=a[3]),b}},{}],24:[function(a,b,c){var d=Math.PI/2;b.exports=function(a,b,c){var e=a*c,f=.5*a;return e=Math.pow((1-e)/(1+e),f),Math.tan(.5*(d-b))/e}},{}],25:[function(a,b,c){c.wgs84={towgs84:"0,0,0",ellipse:"WGS84",datumName:"WGS84"},c.ch1903={towgs84:"674.374,15.056,405.346",ellipse:"bessel",datumName:"swiss"},c.ggrs87={towgs84:"-199.87,74.79,246.62",ellipse:"GRS80",datumName:"Greek_Geodetic_Reference_System_1987"},c.nad83={towgs84:"0,0,0",ellipse:"GRS80",datumName:"North_American_Datum_1983"},c.nad27={nadgrids:"@conus,@alaska,@ntv2_0.gsb,@ntv1_can.dat",ellipse:"clrk66",datumName:"North_American_Datum_1927"},c.potsdam={towgs84:"606.0,23.0,413.0",ellipse:"bessel",datumName:"Potsdam Rauenberg 1950 DHDN"},c.carthage={towgs84:"-263.0,6.0,431.0",ellipse:"clark80",datumName:"Carthage 1934 Tunisia"},c.hermannskogel={towgs84:"653.0,-212.0,449.0",ellipse:"bessel",datumName:"Hermannskogel"},c.ire65={towgs84:"482.530,-130.596,564.557,-1.042,-0.214,-0.631,8.15",ellipse:"mod_airy",datumName:"Ireland 1965"},c.rassadiran={towgs84:"-133.63,-157.5,-158.62",ellipse:"intl",datumName:"Rassadiran"},c.nzgd49={towgs84:"59.47,-5.04,187.44,0.47,-0.1,1.024,-4.5993",ellipse:"intl",datumName:"New Zealand Geodetic Datum 1949"},c.osgb36={towgs84:"446.448,-125.157,542.060,0.1502,0.2470,0.8421,-20.4894",ellipse:"airy",datumName:"Airy 1830"},c.s_jtsk={towgs84:"589,76,480",ellipse:"bessel",datumName:"S-JTSK (Ferro)"},c.beduaram={towgs84:"-106,-87,188",ellipse:"clrk80",datumName:"Beduaram"},c.gunung_segara={towgs84:"-403,684,41",ellipse:"bessel",datumName:"Gunung Segara Jakarta"},c.rnb72={towgs84:"106.869,-52.2978,103.724,-0.33657,0.456955,-1.84218,1",ellipse:"intl",datumName:"Reseau National Belge 1972"}},{}],26:[function(a,b,c){c.MERIT={a:6378137,rf:298.257,ellipseName:"MERIT 1983"},c.SGS85={a:6378136,rf:298.257,ellipseName:"Soviet Geodetic System 85"},c.GRS80={a:6378137,rf:298.257222101,ellipseName:"GRS 1980(IUGG, 1980)"},c.IAU76={a:6378140,rf:298.257,ellipseName:"IAU 1976"},c.airy={a:6377563.396,b:6356256.91,ellipseName:"Airy 1830"},c.APL4={a:6378137,rf:298.25,ellipseName:"Appl. Physics. 1965"},c.NWL9D={a:6378145,rf:298.25,ellipseName:"Naval Weapons Lab., 1965"},c.mod_airy={a:6377340.189,b:6356034.446,ellipseName:"Modified Airy"},c.andrae={a:6377104.43,rf:300,ellipseName:"Andrae 1876 (Den., Iclnd.)"},c.aust_SA={a:6378160,rf:298.25,ellipseName:"Australian Natl & S. Amer. 1969"},c.GRS67={a:6378160,rf:298.247167427,ellipseName:"GRS 67(IUGG 1967)"},c.bessel={a:6377397.155,rf:299.1528128,ellipseName:"Bessel 1841"},c.bess_nam={a:6377483.865,rf:299.1528128,ellipseName:"Bessel 1841 (Namibia)"},c.clrk66={a:6378206.4,b:6356583.8,ellipseName:"Clarke 1866"},c.clrk80={a:6378249.145,rf:293.4663,ellipseName:"Clarke 1880 mod."},c.clrk58={a:6378293.645208759,rf:294.2606763692654,ellipseName:"Clarke 1858"},c.CPM={a:6375738.7,rf:334.29,ellipseName:"Comm. des Poids et Mesures 1799"},c.delmbr={a:6376428,rf:311.5,ellipseName:"Delambre 1810 (Belgium)"},c.engelis={a:6378136.05,rf:298.2566,ellipseName:"Engelis 1985"},c.evrst30={a:6377276.345,rf:300.8017,ellipseName:"Everest 1830"},c.evrst48={a:6377304.063,rf:300.8017,ellipseName:"Everest 1948"},c.evrst56={a:6377301.243,rf:300.8017,ellipseName:"Everest 1956"},c.evrst69={a:6377295.664,rf:300.8017,ellipseName:"Everest 1969"},c.evrstSS={a:6377298.556,rf:300.8017,ellipseName:"Everest (Sabah & Sarawak)"},c.fschr60={a:6378166,rf:298.3,ellipseName:"Fischer (Mercury Datum) 1960"},c.fschr60m={a:6378155,rf:298.3,ellipseName:"Fischer 1960"},c.fschr68={a:6378150,rf:298.3,ellipseName:"Fischer 1968"},c.helmert={a:6378200,rf:298.3,ellipseName:"Helmert 1906"},c.hough={a:6378270,rf:297,ellipseName:"Hough"},c.intl={a:6378388,rf:297,ellipseName:"International 1909 (Hayford)"},c.kaula={a:6378163,rf:298.24,ellipseName:"Kaula 1961"},c.lerch={a:6378139,rf:298.257,ellipseName:"Lerch 1979"},c.mprts={a:6397300,rf:191,ellipseName:"Maupertius 1738"},c.new_intl={a:6378157.5,b:6356772.2,ellipseName:"New International 1967"},c.plessis={a:6376523,rf:6355863,ellipseName:"Plessis 1817 (France)"},c.krass={a:6378245,rf:298.3,ellipseName:"Krassovsky, 1942"},c.SEasia={a:6378155,b:6356773.3205,ellipseName:"Southeast Asia"},c.walbeck={a:6376896,b:6355834.8467,ellipseName:"Walbeck"},c.WGS60={a:6378165,rf:298.3,ellipseName:"WGS 60"},c.WGS66={a:6378145,rf:298.25,ellipseName:"WGS 66"},c.WGS7={a:6378135,rf:298.26,ellipseName:"WGS 72"},c.WGS84={a:6378137,rf:298.257223563,ellipseName:"WGS 84"},c.sphere={a:6370997,b:6370997,ellipseName:"Normal Sphere (r=6370997)"}},{}],27:[function(a,b,c){c.greenwich=0,c.lisbon=-9.131906111111,c.paris=2.337229166667,c.bogota=-74.080916666667,c.madrid=-3.687938888889,c.rome=12.452333333333,c.bern=7.439583333333,c.jakarta=106.807719444444,c.ferro=-17.666666666667,c.brussels=4.367975,c.stockholm=18.058277777778,c.athens=23.7163375,c.oslo=10.722916666667},{}],28:[function(a,b,c){c.ft={to_meter:.3048},c["us-ft"]={to_meter:1200/3937}},{}],29:[function(a,b,c){function d(a,b,c){var d;return Array.isArray(c)?(d=g(a,b,c),3===c.length?[d.x,d.y,d.z]:[d.x,d.y]):g(a,b,c)}function e(a){return a instanceof f?a:a.oProj?a.oProj:f(a)}function proj4(a,b,c){a=e(a);var f,g=!1;return"undefined"==typeof b?(b=a,a=h,g=!0):("undefined"!=typeof b.x||Array.isArray(b))&&(c=b,b=a,a=h,g=!0),b=e(b),c?d(a,b,c):(f={forward:function(c){return d(a,b,c)},inverse:function(c){return d(b,a,c)}},g&&(f.oProj=b),f)}var f=a("./Proj"),g=a("./transform"),h=f("WGS84");b.exports=proj4},{"./Proj":2,"./transform":66}],30:[function(a,b,c){var d=Math.PI/2,e=1,f=2,g=3,h=4,i=5,j=484813681109536e-20,k=1.0026,l=.3826834323650898,m=function(a){return this instanceof m?(this.datum_type=h,void(a&&(a.datumCode&&"none"===a.datumCode&&(this.datum_type=i),a.datum_params&&(this.datum_params=a.datum_params.map(parseFloat),0===this.datum_params[0]&&0===this.datum_params[1]&&0===this.datum_params[2]||(this.datum_type=e),this.datum_params.length>3&&(0===this.datum_params[3]&&0===this.datum_params[4]&&0===this.datum_params[5]&&0===this.datum_params[6]||(this.datum_type=f,this.datum_params[3]*=j,this.datum_params[4]*=j,this.datum_params[5]*=j,this.datum_params[6]=this.datum_params[6]/1e6+1))),this.datum_type=a.grids?g:this.datum_type,this.a=a.a,this.b=a.b,this.es=a.es,this.ep2=a.ep2,this.datum_type===g&&(this.grids=a.grids)))):new m(a)};m.prototype={compare_datums:function(a){return this.datum_type!==a.datum_type?!1:this.a!==a.a||Math.abs(this.es-a.es)>5e-11?!1:this.datum_type===e?this.datum_params[0]===a.datum_params[0]&&this.datum_params[1]===a.datum_params[1]&&this.datum_params[2]===a.datum_params[2]:this.datum_type===f?this.datum_params[0]===a.datum_params[0]&&this.datum_params[1]===a.datum_params[1]&&this.datum_params[2]===a.datum_params[2]&&this.datum_params[3]===a.datum_params[3]&&this.datum_params[4]===a.datum_params[4]&&this.datum_params[5]===a.datum_params[5]&&this.datum_params[6]===a.datum_params[6]:this.datum_type===g||a.datum_type===g?this.nadgrids===a.nadgrids:!0},geodetic_to_geocentric:function(a){var b,c,e,f,g,h,i,j=a.x,k=a.y,l=a.z?a.z:0,m=0;if(-d>k&&k>-1.001*d)k=-d;else if(k>d&&1.001*d>k)k=d;else if(-d>k||k>d)return null;return j>Math.PI&&(j-=2*Math.PI),g=Math.sin(k),i=Math.cos(k),h=g*g,f=this.a/Math.sqrt(1-this.es*h),b=(f+l)*i*Math.cos(j),c=(f+l)*i*Math.sin(j),e=(f*(1-this.es)+l)*g,a.x=b,a.y=c,a.z=e,m},geocentric_to_geodetic:function(a){var b,c,e,f,g,h,i,j,k,l,m,n,o,p,q,r,s,t=1e-12,u=t*t,v=30,w=a.x,x=a.y,y=a.z?a.z:0;if(o=!1,b=Math.sqrt(w*w+x*x),c=Math.sqrt(w*w+x*x+y*y),b/this.a<t){if(o=!0,q=0,c/this.a<t)return r=d,void(s=-this.b)}else q=Math.atan2(x,w);e=y/c,f=b/c,g=1/Math.sqrt(1-this.es*(2-this.es)*f*f),j=f*(1-this.es)*g,k=e*g,p=0;do p++,i=this.a/Math.sqrt(1-this.es*k*k),s=b*j+y*k-i*(1-this.es*k*k),h=this.es*i/(i+s),g=1/Math.sqrt(1-h*(2-h)*f*f),l=f*(1-h)*g,m=e*g,n=m*j-l*k,j=l,k=m;while(n*n>u&&v>p);return r=Math.atan(m/Math.abs(l)),a.x=q,a.y=r,a.z=s,a},geocentric_to_geodetic_noniter:function(a){var b,c,e,f,g,h,i,j,m,n,o,p,q,r,s,t,u,v=a.x,w=a.y,x=a.z?a.z:0;if(v=parseFloat(v),w=parseFloat(w),x=parseFloat(x),u=!1,0!==v)b=Math.atan2(w,v);else if(w>0)b=d;else if(0>w)b=-d;else if(u=!0,b=0,x>0)c=d;else{if(!(0>x))return c=d,void(e=-this.b);c=-d}return g=v*v+w*w,f=Math.sqrt(g),h=x*k,j=Math.sqrt(h*h+g),n=h/j,p=f/j,o=n*n*n,i=x+this.b*this.ep2*o,t=f-this.a*this.es*p*p*p,m=Math.sqrt(i*i+t*t),q=i/m,r=t/m,s=this.a/Math.sqrt(1-this.es*q*q),e=r>=l?f/r-s:-l>=r?f/-r-s:x/q+s*(this.es-1),u===!1&&(c=Math.atan(q/r)),a.x=b,a.y=c,a.z=e,a},geocentric_to_wgs84:function(a){if(this.datum_type===e)a.x+=this.datum_params[0],a.y+=this.datum_params[1],a.z+=this.datum_params[2];else if(this.datum_type===f){var b=this.datum_params[0],c=this.datum_params[1],d=this.datum_params[2],g=this.datum_params[3],h=this.datum_params[4],i=this.datum_params[5],j=this.datum_params[6],k=j*(a.x-i*a.y+h*a.z)+b,l=j*(i*a.x+a.y-g*a.z)+c,m=j*(-h*a.x+g*a.y+a.z)+d;a.x=k,a.y=l,a.z=m}},geocentric_from_wgs84:function(a){if(this.datum_type===e)a.x-=this.datum_params[0],a.y-=this.datum_params[1],a.z-=this.datum_params[2];else if(this.datum_type===f){var b=this.datum_params[0],c=this.datum_params[1],d=this.datum_params[2],g=this.datum_params[3],h=this.datum_params[4],i=this.datum_params[5],j=this.datum_params[6],k=(a.x-b)/j,l=(a.y-c)/j,m=(a.z-d)/j;a.x=k+i*l-h*m,a.y=-i*k+l+g*m,a.z=h*k-g*l+m}}},b.exports=m},{}],31:[function(a,b,c){var d=1,e=2,f=3,g=5,h=6378137,i=.006694379990141316;b.exports=function(a,b,c){function j(a){return a===d||a===e}var k,l,m;if(a.compare_datums(b))return c;if(a.datum_type===g||b.datum_type===g)return c;var n=a.a,o=a.es,p=b.a,q=b.es,r=a.datum_type;if(r===f)if(0===this.apply_gridshift(a,0,c))a.a=h,a.es=i;else{if(!a.datum_params)return a.a=n,a.es=a.es,c;for(k=1,l=0,m=a.datum_params.length;m>l;l++)k*=a.datum_params[l];if(0===k)return a.a=n,a.es=a.es,c;r=a.datum_params.length>3?e:d}return b.datum_type===f&&(b.a=h,b.es=i),(a.es!==b.es||a.a!==b.a||j(r)||j(b.datum_type))&&(a.geodetic_to_geocentric(c),j(a.datum_type)&&a.geocentric_to_wgs84(c),j(b.datum_type)&&b.geocentric_from_wgs84(c),b.geocentric_to_geodetic(c)),b.datum_type===f&&this.apply_gridshift(b,1,c),a.a=n,a.es=o,b.a=p,b.es=q,c}},{}],32:[function(a,b,c){function d(a){var b=this;if(2===arguments.length){var c=arguments[1];"string"==typeof c?"+"===c.charAt(0)?d[a]=f(arguments[1]):d[a]=g(arguments[1]):d[a]=c}else if(1===arguments.length){if(Array.isArray(a))return a.map(function(a){Array.isArray(a)?d.apply(b,a):d(a)});if("string"==typeof a){if(a in d)return d[a]}else"EPSG"in a?d["EPSG:"+a.EPSG]=a:"ESRI"in a?d["ESRI:"+a.ESRI]=a:"IAU2000"in a?d["IAU2000:"+a.IAU2000]=a:console.log(a);return}}var e=a("./global"),f=a("./projString"),g=a("./wkt");e(d),b.exports=d},{"./global":35,"./projString":38,"./wkt":67}],33:[function(a,b,c){var d=a("./constants/Datum"),e=a("./constants/Ellipsoid"),f=a("./extend"),g=a("./datum"),h=1e-10,i=.16666666666666666,j=.04722222222222222,k=.022156084656084655;b.exports=function(a){if(a.datumCode&&"none"!==a.datumCode){var b=d[a.datumCode];b&&(a.datum_params=b.towgs84?b.towgs84.split(","):null,a.ellps=b.ellipse,a.datumName=b.datumName?b.datumName:a.datumCode)}if(!a.a){var c=e[a.ellps]?e[a.ellps]:e.WGS84;f(a,c)}return a.rf&&!a.b&&(a.b=(1-1/a.rf)*a.a),(0===a.rf||Math.abs(a.a-a.b)<h)&&(a.sphere=!0,a.b=a.a),a.a2=a.a*a.a,a.b2=a.b*a.b,a.es=(a.a2-a.b2)/a.a2,a.e=Math.sqrt(a.es),a.R_A&&(a.a*=1-a.es*(i+a.es*(j+a.es*k)),a.a2=a.a*a.a,a.b2=a.b*a.b,a.es=0),a.ep2=(a.a2-a.b2)/a.b2,a.k0||(a.k0=1),a.axis||(a.axis="enu"),a.datum||(a.datum=g(a)),a}},{"./constants/Datum":25,"./constants/Ellipsoid":26,"./datum":30,"./extend":34}],34:[function(a,b,c){b.exports=function(a,b){a=a||{};var c,d;if(!b)return a;for(d in b)c=b[d],void 0!==c&&(a[d]=c);return a}},{}],35:[function(a,b,c){b.exports=function(a){a("EPSG:4326","+title=WGS 84 (long/lat) +proj=longlat +ellps=WGS84 +datum=WGS84 +units=degrees"),a("EPSG:4269","+title=NAD83 (long/lat) +proj=longlat +a=6378137.0 +b=6356752.31414036 +ellps=GRS80 +datum=NAD83 +units=degrees"),a("EPSG:3857","+title=WGS 84 / Pseudo-Mercator +proj=merc +a=6378137 +b=6378137 +lat_ts=0.0 +lon_0=0.0 +x_0=0.0 +y_0=0 +k=1.0 +units=m +nadgrids=@null +no_defs"),a.WGS84=a["EPSG:4326"],a["EPSG:3785"]=a["EPSG:3857"],a.GOOGLE=a["EPSG:3857"],a["EPSG:900913"]=a["EPSG:3857"],a["EPSG:102113"]=a["EPSG:3857"]}},{}],36:[function(a,b,c){var proj4=a("./core");proj4.defaultDatum="WGS84",proj4.Proj=a("./Proj"),proj4.WGS84=new proj4.Proj("WGS84"),proj4.Point=a("./Point"),proj4.toPoint=a("./common/toPoint"),proj4.defs=a("./defs"),proj4.transform=a("./transform"),proj4.mgrs=a("mgrs"),proj4.version=a("../package.json").version,a("./includedProjections")(proj4),b.exports=proj4},{"../package.json":69,"./Point":1,"./Proj":2,"./common/toPoint":23,"./core":29,"./defs":32,"./includedProjections":"./includedProjections","./transform":66,mgrs:68}],37:[function(a,b,c){function d(a){return"string"==typeof a}function e(a){return a in i}function f(a){var b=["GEOGCS","GEOCCS","PROJCS","LOCAL_CS"];return b.reduce(function(b,c){return b+1+a.indexOf(c)},0)}function g(a){return"+"===a[0]}function h(a){return d(a)?e(a)?i[a]:f(a)?j(a):g(a)?k(a):void 0:a}var i=a("./defs"),j=a("./wkt"),k=a("./projString");b.exports=h},{"./defs":32,"./projString":38,"./wkt":67}],38:[function(a,b,c){var d=.017453292519943295,e=a("./constants/PrimeMeridian"),f=a("./constants/units");b.exports=function(a){var b={},c={};a.split("+").map(function(a){return a.trim()}).filter(function(a){return a}).forEach(function(a){var b=a.split("=");b.push(!0),c[b[0].toLowerCase()]=b[1]});var g,h,i,j={proj:"projName",datum:"datumCode",rf:function(a){b.rf=parseFloat(a)},lat_0:function(a){b.lat0=a*d},lat_1:function(a){b.lat1=a*d},lat_2:function(a){b.lat2=a*d},lat_ts:function(a){b.lat_ts=a*d},lon_0:function(a){b.long0=a*d},lon_1:function(a){b.long1=a*d},lon_2:function(a){b.long2=a*d},alpha:function(a){b.alpha=parseFloat(a)*d},lonc:function(a){b.longc=a*d},x_0:function(a){b.x0=parseFloat(a)},y_0:function(a){b.y0=parseFloat(a)},k_0:function(a){b.k0=parseFloat(a)},k:function(a){b.k0=parseFloat(a)},a:function(a){b.a=parseFloat(a)},b:function(a){b.b=parseFloat(a)},r_a:function(){b.R_A=!0},zone:function(a){b.zone=parseInt(a,10)},south:function(){b.utmSouth=!0},towgs84:function(a){b.datum_params=a.split(",").map(function(a){return parseFloat(a)})},to_meter:function(a){b.to_meter=parseFloat(a)},units:function(a){b.units=a,f[a]&&(b.to_meter=f[a].to_meter)},from_greenwich:function(a){b.from_greenwich=a*d},pm:function(a){b.from_greenwich=(e[a]?e[a]:parseFloat(a))*d},nadgrids:function(a){"@null"===a?b.datumCode="none":b.nadgrids=a},axis:function(a){var c="ewnsud";3===a.length&&-1!==c.indexOf(a.substr(0,1))&&-1!==c.indexOf(a.substr(1,1))&&-1!==c.indexOf(a.substr(2,1))&&(b.axis=a)}};for(g in c)h=c[g],g in j?(i=j[g],"function"==typeof i?i(h):b[i]=h):b[g]=h;return"string"==typeof b.datumCode&&"WGS84"!==b.datumCode&&(b.datumCode=b.datumCode.toLowerCase()),b}},{"./constants/PrimeMeridian":27,"./constants/units":28}],39:[function(a,b,c){function d(a,b){var c=g.length;return a.names?(g[c]=a,a.names.forEach(function(a){f[a.toLowerCase()]=c}),this):(console.log(b),!0)}var e=[a("./projections/merc"),a("./projections/longlat")],f={},g=[];c.add=d,c.get=function(a){if(!a)return!1;var b=a.toLowerCase();return"undefined"!=typeof f[b]&&g[f[b]]?g[f[b]]:void 0},c.start=function(){e.forEach(d)}},{"./projections/longlat":51,"./projections/merc":52}],40:[function(a,b,c){var d=1e-10,e=a("../common/msfnz"),f=a("../common/qsfnz"),g=a("../common/adjust_lon"),h=a("../common/asinz");c.init=function(){Math.abs(this.lat1+this.lat2)<d||(this.temp=this.b/this.a,this.es=1-Math.pow(this.temp,2),this.e3=Math.sqrt(this.es),this.sin_po=Math.sin(this.lat1),this.cos_po=Math.cos(this.lat1),this.t1=this.sin_po,this.con=this.sin_po,this.ms1=e(this.e3,this.sin_po,this.cos_po),this.qs1=f(this.e3,this.sin_po,this.cos_po),this.sin_po=Math.sin(this.lat2),this.cos_po=Math.cos(this.lat2),this.t2=this.sin_po,this.ms2=e(this.e3,this.sin_po,this.cos_po),this.qs2=f(this.e3,this.sin_po,this.cos_po),this.sin_po=Math.sin(this.lat0),this.cos_po=Math.cos(this.lat0),this.t3=this.sin_po,this.qs0=f(this.e3,this.sin_po,this.cos_po),Math.abs(this.lat1-this.lat2)>d?this.ns0=(this.ms1*this.ms1-this.ms2*this.ms2)/(this.qs2-this.qs1):this.ns0=this.con,this.c=this.ms1*this.ms1+this.ns0*this.qs1,this.rh=this.a*Math.sqrt(this.c-this.ns0*this.qs0)/this.ns0)},c.forward=function(a){var b=a.x,c=a.y;this.sin_phi=Math.sin(c),this.cos_phi=Math.cos(c);var d=f(this.e3,this.sin_phi,this.cos_phi),e=this.a*Math.sqrt(this.c-this.ns0*d)/this.ns0,h=this.ns0*g(b-this.long0),i=e*Math.sin(h)+this.x0,j=this.rh-e*Math.cos(h)+this.y0;return a.x=i,a.y=j,a},c.inverse=function(a){var b,c,d,e,f,h;return a.x-=this.x0,a.y=this.rh-a.y+this.y0,this.ns0>=0?(b=Math.sqrt(a.x*a.x+a.y*a.y),d=1):(b=-Math.sqrt(a.x*a.x+a.y*a.y),d=-1),e=0,0!==b&&(e=Math.atan2(d*a.x,d*a.y)),d=b*this.ns0/this.a,this.sphere?h=Math.asin((this.c-d*d)/(2*this.ns0)):(c=(this.c-d*d)/this.ns0,h=this.phi1z(this.e3,c)),f=g(e/this.ns0+this.long0),a.x=f,a.y=h,a},c.phi1z=function(a,b){var c,e,f,g,i,j=h(.5*b);if(d>a)return j;for(var k=a*a,l=1;25>=l;l++)if(c=Math.sin(j),e=Math.cos(j),f=a*c,g=1-f*f,i=.5*g*g/e*(b/(1-k)-c/g+.5/a*Math.log((1-f)/(1+f))),j+=i,Math.abs(i)<=1e-7)return j;return null},c.names=["Albers_Conic_Equal_Area","Albers","aea"]},{"../common/adjust_lon":5,"../common/asinz":6,"../common/msfnz":15,"../common/qsfnz":20}],41:[function(a,b,c){var d=a("../common/adjust_lon"),e=Math.PI/2,f=1e-10,g=a("../common/mlfn"),h=a("../common/e0fn"),i=a("../common/e1fn"),j=a("../common/e2fn"),k=a("../common/e3fn"),l=a("../common/gN"),m=a("../common/asinz"),n=a("../common/imlfn");c.init=function(){this.sin_p12=Math.sin(this.lat0),this.cos_p12=Math.cos(this.lat0)},c.forward=function(a){var b,c,m,n,o,p,q,r,s,t,u,v,w,x,y,z,A,B,C,D,E,F,G,H=a.x,I=a.y,J=Math.sin(a.y),K=Math.cos(a.y),L=d(H-this.long0);return this.sphere?Math.abs(this.sin_p12-1)<=f?(a.x=this.x0+this.a*(e-I)*Math.sin(L),a.y=this.y0-this.a*(e-I)*Math.cos(L),a):Math.abs(this.sin_p12+1)<=f?(a.x=this.x0+this.a*(e+I)*Math.sin(L),a.y=this.y0+this.a*(e+I)*Math.cos(L),a):(B=this.sin_p12*J+this.cos_p12*K*Math.cos(L),z=Math.acos(B),A=z/Math.sin(z),a.x=this.x0+this.a*A*K*Math.sin(L),a.y=this.y0+this.a*A*(this.cos_p12*J-this.sin_p12*K*Math.cos(L)),a):(b=h(this.es),c=i(this.es),m=j(this.es),n=k(this.es),Math.abs(this.sin_p12-1)<=f?(o=this.a*g(b,c,m,n,e),p=this.a*g(b,c,m,n,I),a.x=this.x0+(o-p)*Math.sin(L),a.y=this.y0-(o-p)*Math.cos(L),a):Math.abs(this.sin_p12+1)<=f?(o=this.a*g(b,c,m,n,e),p=this.a*g(b,c,m,n,I),a.x=this.x0+(o+p)*Math.sin(L),a.y=this.y0+(o+p)*Math.cos(L),a):(q=J/K,r=l(this.a,this.e,this.sin_p12),s=l(this.a,this.e,J),t=Math.atan((1-this.es)*q+this.es*r*this.sin_p12/(s*K)),u=Math.atan2(Math.sin(L),this.cos_p12*Math.tan(t)-this.sin_p12*Math.cos(L)),C=0===u?Math.asin(this.cos_p12*Math.sin(t)-this.sin_p12*Math.cos(t)):Math.abs(Math.abs(u)-Math.PI)<=f?-Math.asin(this.cos_p12*Math.sin(t)-this.sin_p12*Math.cos(t)):Math.asin(Math.sin(L)*Math.cos(t)/Math.sin(u)),v=this.e*this.sin_p12/Math.sqrt(1-this.es),w=this.e*this.cos_p12*Math.cos(u)/Math.sqrt(1-this.es),x=v*w,y=w*w,D=C*C,E=D*C,F=E*C,G=F*C,z=r*C*(1-D*y*(1-y)/6+E/8*x*(1-2*y)+F/120*(y*(4-7*y)-3*v*v*(1-7*y))-G/48*x),a.x=this.x0+z*Math.sin(u),a.y=this.y0+z*Math.cos(u),a))},c.inverse=function(a){a.x-=this.x0,a.y-=this.y0;var b,c,o,p,q,r,s,t,u,v,w,x,y,z,A,B,C,D,E,F,G,H,I;if(this.sphere){if(b=Math.sqrt(a.x*a.x+a.y*a.y),b>2*e*this.a)return;return c=b/this.a,o=Math.sin(c),p=Math.cos(c),q=this.long0,Math.abs(b)<=f?r=this.lat0:(r=m(p*this.sin_p12+a.y*o*this.cos_p12/b),s=Math.abs(this.lat0)-e,q=d(Math.abs(s)<=f?this.lat0>=0?this.long0+Math.atan2(a.x,-a.y):this.long0-Math.atan2(-a.x,a.y):this.long0+Math.atan2(a.x*o,b*this.cos_p12*p-a.y*this.sin_p12*o))),a.x=q,a.y=r,a}return t=h(this.es),u=i(this.es),v=j(this.es),w=k(this.es),Math.abs(this.sin_p12-1)<=f?(x=this.a*g(t,u,v,w,e),b=Math.sqrt(a.x*a.x+a.y*a.y),y=x-b,r=n(y/this.a,t,u,v,w),q=d(this.long0+Math.atan2(a.x,-1*a.y)),a.x=q,a.y=r,a):Math.abs(this.sin_p12+1)<=f?(x=this.a*g(t,u,v,w,e),b=Math.sqrt(a.x*a.x+a.y*a.y),y=b-x,r=n(y/this.a,t,u,v,w),q=d(this.long0+Math.atan2(a.x,a.y)),a.x=q,a.y=r,a):(b=Math.sqrt(a.x*a.x+a.y*a.y),B=Math.atan2(a.x,a.y),z=l(this.a,this.e,this.sin_p12),C=Math.cos(B),D=this.e*this.cos_p12*C,E=-D*D/(1-this.es),F=3*this.es*(1-E)*this.sin_p12*this.cos_p12*C/(1-this.es),G=b/z,H=G-E*(1+E)*Math.pow(G,3)/6-F*(1+3*E)*Math.pow(G,4)/24,I=1-E*H*H/2-G*H*H*H/6,A=Math.asin(this.sin_p12*Math.cos(H)+this.cos_p12*Math.sin(H)*C),q=d(this.long0+Math.asin(Math.sin(B)*Math.sin(H)/Math.cos(A))),r=Math.atan((1-this.es*I*this.sin_p12/Math.sin(A))*Math.tan(A)/(1-this.es)),a.x=q,a.y=r,a)},c.names=["Azimuthal_Equidistant","aeqd"]},{"../common/adjust_lon":5,"../common/asinz":6,"../common/e0fn":7,"../common/e1fn":8,"../common/e2fn":9,"../common/e3fn":10,"../common/gN":11,"../common/imlfn":12,"../common/mlfn":14}],42:[function(a,b,c){var d=a("../common/mlfn"),e=a("../common/e0fn"),f=a("../common/e1fn"),g=a("../common/e2fn"),h=a("../common/e3fn"),i=a("../common/gN"),j=a("../common/adjust_lon"),k=a("../common/adjust_lat"),l=a("../common/imlfn"),m=Math.PI/2,n=1e-10;c.init=function(){this.sphere||(this.e0=e(this.es),this.e1=f(this.es),this.e2=g(this.es),this.e3=h(this.es),this.ml0=this.a*d(this.e0,this.e1,this.e2,this.e3,this.lat0))},c.forward=function(a){var b,c,e=a.x,f=a.y;if(e=j(e-this.long0),this.sphere)b=this.a*Math.asin(Math.cos(f)*Math.sin(e)),c=this.a*(Math.atan2(Math.tan(f),Math.cos(e))-this.lat0);else{var g=Math.sin(f),h=Math.cos(f),k=i(this.a,this.e,g),l=Math.tan(f)*Math.tan(f),m=e*Math.cos(f),n=m*m,o=this.es*h*h/(1-this.es),p=this.a*d(this.e0,this.e1,this.e2,this.e3,f);b=k*m*(1-n*l*(1/6-(8-l+8*o)*n/120)),c=p-this.ml0+k*g/h*n*(.5+(5-l+6*o)*n/24)}return a.x=b+this.x0,a.y=c+this.y0,a},c.inverse=function(a){a.x-=this.x0,a.y-=this.y0;var b,c,d=a.x/this.a,e=a.y/this.a;if(this.sphere){var f=e+this.lat0;b=Math.asin(Math.sin(f)*Math.cos(d)),c=Math.atan2(Math.tan(d),Math.cos(f))}else{var g=this.ml0/this.a+e,h=l(g,this.e0,this.e1,this.e2,this.e3);if(Math.abs(Math.abs(h)-m)<=n)return a.x=this.long0,a.y=m,0>e&&(a.y*=-1),a;var o=i(this.a,this.e,Math.sin(h)),p=o*o*o/this.a/this.a*(1-this.es),q=Math.pow(Math.tan(h),2),r=d*this.a/o,s=r*r;b=h-o*Math.tan(h)/p*r*r*(.5-(1+3*q)*r*r/24),c=r*(1-s*(q/3+(1+3*q)*q*s/15))/Math.cos(h)}return a.x=j(c+this.long0),a.y=k(b),a},c.names=["Cassini","Cassini_Soldner","cass"]},{"../common/adjust_lat":4,"../common/adjust_lon":5,"../common/e0fn":7,"../common/e1fn":8,"../common/e2fn":9,"../common/e3fn":10,"../common/gN":11,"../common/imlfn":12,"../common/mlfn":14}],43:[function(a,b,c){var d=a("../common/adjust_lon"),e=a("../common/qsfnz"),f=a("../common/msfnz"),g=a("../common/iqsfnz");c.init=function(){this.sphere||(this.k0=f(this.e,Math.sin(this.lat_ts),Math.cos(this.lat_ts)))},c.forward=function(a){var b,c,f=a.x,g=a.y,h=d(f-this.long0);if(this.sphere)b=this.x0+this.a*h*Math.cos(this.lat_ts),c=this.y0+this.a*Math.sin(g)/Math.cos(this.lat_ts);else{var i=e(this.e,Math.sin(g));b=this.x0+this.a*this.k0*h,c=this.y0+this.a*i*.5/this.k0}return a.x=b,a.y=c,a},c.inverse=function(a){a.x-=this.x0,a.y-=this.y0;var b,c;return this.sphere?(b=d(this.long0+a.x/this.a/Math.cos(this.lat_ts)),c=Math.asin(a.y/this.a*Math.cos(this.lat_ts))):(c=g(this.e,2*a.y*this.k0/this.a),b=d(this.long0+a.x/(this.a*this.k0))),a.x=b,a.y=c,a},c.names=["cea"]},{"../common/adjust_lon":5,"../common/iqsfnz":13,"../common/msfnz":15,"../common/qsfnz":20}],44:[function(a,b,c){var d=a("../common/adjust_lon"),e=a("../common/adjust_lat");c.init=function(){this.x0=this.x0||0,this.y0=this.y0||0,this.lat0=this.lat0||0,this.long0=this.long0||0,this.lat_ts=this.lat_ts||0,this.title=this.title||"Equidistant Cylindrical (Plate Carre)",this.rc=Math.cos(this.lat_ts)},c.forward=function(a){var b=a.x,c=a.y,f=d(b-this.long0),g=e(c-this.lat0);return a.x=this.x0+this.a*f*this.rc,a.y=this.y0+this.a*g,a},c.inverse=function(a){var b=a.x,c=a.y;return a.x=d(this.long0+(b-this.x0)/(this.a*this.rc)),a.y=e(this.lat0+(c-this.y0)/this.a),a},c.names=["Equirectangular","Equidistant_Cylindrical","eqc"]},{"../common/adjust_lat":4,"../common/adjust_lon":5}],45:[function(a,b,c){var d=a("../common/e0fn"),e=a("../common/e1fn"),f=a("../common/e2fn"),g=a("../common/e3fn"),h=a("../common/msfnz"),i=a("../common/mlfn"),j=a("../common/adjust_lon"),k=a("../common/adjust_lat"),l=a("../common/imlfn"),m=1e-10;c.init=function(){Math.abs(this.lat1+this.lat2)<m||(this.lat2=this.lat2||this.lat1,this.temp=this.b/this.a,this.es=1-Math.pow(this.temp,2),this.e=Math.sqrt(this.es),this.e0=d(this.es),this.e1=e(this.es),this.e2=f(this.es),this.e3=g(this.es),this.sinphi=Math.sin(this.lat1),this.cosphi=Math.cos(this.lat1),this.ms1=h(this.e,this.sinphi,this.cosphi),this.ml1=i(this.e0,this.e1,this.e2,this.e3,this.lat1),Math.abs(this.lat1-this.lat2)<m?this.ns=this.sinphi:(this.sinphi=Math.sin(this.lat2),this.cosphi=Math.cos(this.lat2),this.ms2=h(this.e,this.sinphi,this.cosphi),this.ml2=i(this.e0,this.e1,this.e2,this.e3,this.lat2),this.ns=(this.ms1-this.ms2)/(this.ml2-this.ml1)),this.g=this.ml1+this.ms1/this.ns,this.ml0=i(this.e0,this.e1,this.e2,this.e3,this.lat0),this.rh=this.a*(this.g-this.ml0))},c.forward=function(a){var b,c=a.x,d=a.y;if(this.sphere)b=this.a*(this.g-d);else{var e=i(this.e0,this.e1,this.e2,this.e3,d);b=this.a*(this.g-e)}var f=this.ns*j(c-this.long0),g=this.x0+b*Math.sin(f),h=this.y0+this.rh-b*Math.cos(f);return a.x=g,a.y=h,a},c.inverse=function(a){a.x-=this.x0,a.y=this.rh-a.y+this.y0;var b,c,d,e;this.ns>=0?(c=Math.sqrt(a.x*a.x+a.y*a.y),
b=1):(c=-Math.sqrt(a.x*a.x+a.y*a.y),b=-1);var f=0;if(0!==c&&(f=Math.atan2(b*a.x,b*a.y)),this.sphere)return e=j(this.long0+f/this.ns),d=k(this.g-c/this.a),a.x=e,a.y=d,a;var g=this.g-c/this.a;return d=l(g,this.e0,this.e1,this.e2,this.e3),e=j(this.long0+f/this.ns),a.x=e,a.y=d,a},c.names=["Equidistant_Conic","eqdc"]},{"../common/adjust_lat":4,"../common/adjust_lon":5,"../common/e0fn":7,"../common/e1fn":8,"../common/e2fn":9,"../common/e3fn":10,"../common/imlfn":12,"../common/mlfn":14,"../common/msfnz":15}],46:[function(a,b,c){var d=Math.PI/4,e=a("../common/srat"),f=Math.PI/2,g=20;c.init=function(){var a=Math.sin(this.lat0),b=Math.cos(this.lat0);b*=b,this.rc=Math.sqrt(1-this.es)/(1-this.es*a*a),this.C=Math.sqrt(1+this.es*b*b/(1-this.es)),this.phic0=Math.asin(a/this.C),this.ratexp=.5*this.C*this.e,this.K=Math.tan(.5*this.phic0+d)/(Math.pow(Math.tan(.5*this.lat0+d),this.C)*e(this.e*a,this.ratexp))},c.forward=function(a){var b=a.x,c=a.y;return a.y=2*Math.atan(this.K*Math.pow(Math.tan(.5*c+d),this.C)*e(this.e*Math.sin(c),this.ratexp))-f,a.x=this.C*b,a},c.inverse=function(a){for(var b=1e-14,c=a.x/this.C,h=a.y,i=Math.pow(Math.tan(.5*h+d)/this.K,1/this.C),j=g;j>0&&(h=2*Math.atan(i*e(this.e*Math.sin(a.y),-.5*this.e))-f,!(Math.abs(h-a.y)<b));--j)a.y=h;return j?(a.x=c,a.y=h,a):null},c.names=["gauss"]},{"../common/srat":22}],47:[function(a,b,c){var d=a("../common/adjust_lon"),e=1e-10,f=a("../common/asinz");c.init=function(){this.sin_p14=Math.sin(this.lat0),this.cos_p14=Math.cos(this.lat0),this.infinity_dist=1e3*this.a,this.rc=1},c.forward=function(a){var b,c,f,g,h,i,j,k,l=a.x,m=a.y;return f=d(l-this.long0),b=Math.sin(m),c=Math.cos(m),g=Math.cos(f),i=this.sin_p14*b+this.cos_p14*c*g,h=1,i>0||Math.abs(i)<=e?(j=this.x0+this.a*h*c*Math.sin(f)/i,k=this.y0+this.a*h*(this.cos_p14*b-this.sin_p14*c*g)/i):(j=this.x0+this.infinity_dist*c*Math.sin(f),k=this.y0+this.infinity_dist*(this.cos_p14*b-this.sin_p14*c*g)),a.x=j,a.y=k,a},c.inverse=function(a){var b,c,e,g,h,i;return a.x=(a.x-this.x0)/this.a,a.y=(a.y-this.y0)/this.a,a.x/=this.k0,a.y/=this.k0,(b=Math.sqrt(a.x*a.x+a.y*a.y))?(g=Math.atan2(b,this.rc),c=Math.sin(g),e=Math.cos(g),i=f(e*this.sin_p14+a.y*c*this.cos_p14/b),h=Math.atan2(a.x*c,b*this.cos_p14*e-a.y*this.sin_p14*c),h=d(this.long0+h)):(i=this.phic0,h=0),a.x=h,a.y=i,a},c.names=["gnom"]},{"../common/adjust_lon":5,"../common/asinz":6}],48:[function(a,b,c){var d=a("../common/adjust_lon");c.init=function(){this.a=6377397.155,this.es=.006674372230614,this.e=Math.sqrt(this.es),this.lat0||(this.lat0=.863937979737193),this.long0||(this.long0=.4334234309119251),this.k0||(this.k0=.9999),this.s45=.785398163397448,this.s90=2*this.s45,this.fi0=this.lat0,this.e2=this.es,this.e=Math.sqrt(this.e2),this.alfa=Math.sqrt(1+this.e2*Math.pow(Math.cos(this.fi0),4)/(1-this.e2)),this.uq=1.04216856380474,this.u0=Math.asin(Math.sin(this.fi0)/this.alfa),this.g=Math.pow((1+this.e*Math.sin(this.fi0))/(1-this.e*Math.sin(this.fi0)),this.alfa*this.e/2),this.k=Math.tan(this.u0/2+this.s45)/Math.pow(Math.tan(this.fi0/2+this.s45),this.alfa)*this.g,this.k1=this.k0,this.n0=this.a*Math.sqrt(1-this.e2)/(1-this.e2*Math.pow(Math.sin(this.fi0),2)),this.s0=1.37008346281555,this.n=Math.sin(this.s0),this.ro0=this.k1*this.n0/Math.tan(this.s0),this.ad=this.s90-this.uq},c.forward=function(a){var b,c,e,f,g,h,i,j=a.x,k=a.y,l=d(j-this.long0);return b=Math.pow((1+this.e*Math.sin(k))/(1-this.e*Math.sin(k)),this.alfa*this.e/2),c=2*(Math.atan(this.k*Math.pow(Math.tan(k/2+this.s45),this.alfa)/b)-this.s45),e=-l*this.alfa,f=Math.asin(Math.cos(this.ad)*Math.sin(c)+Math.sin(this.ad)*Math.cos(c)*Math.cos(e)),g=Math.asin(Math.cos(c)*Math.sin(e)/Math.cos(f)),h=this.n*g,i=this.ro0*Math.pow(Math.tan(this.s0/2+this.s45),this.n)/Math.pow(Math.tan(f/2+this.s45),this.n),a.y=i*Math.cos(h)/1,a.x=i*Math.sin(h)/1,this.czech||(a.y*=-1,a.x*=-1),a},c.inverse=function(a){var b,c,d,e,f,g,h,i,j=a.x;a.x=a.y,a.y=j,this.czech||(a.y*=-1,a.x*=-1),g=Math.sqrt(a.x*a.x+a.y*a.y),f=Math.atan2(a.y,a.x),e=f/Math.sin(this.s0),d=2*(Math.atan(Math.pow(this.ro0/g,1/this.n)*Math.tan(this.s0/2+this.s45))-this.s45),b=Math.asin(Math.cos(this.ad)*Math.sin(d)-Math.sin(this.ad)*Math.cos(d)*Math.cos(e)),c=Math.asin(Math.cos(d)*Math.sin(e)/Math.cos(b)),a.x=this.long0-c/this.alfa,h=b,i=0;var k=0;do a.y=2*(Math.atan(Math.pow(this.k,-1/this.alfa)*Math.pow(Math.tan(b/2+this.s45),1/this.alfa)*Math.pow((1+this.e*Math.sin(h))/(1-this.e*Math.sin(h)),this.e/2))-this.s45),Math.abs(h-a.y)<1e-10&&(i=1),h=a.y,k+=1;while(0===i&&15>k);return k>=15?null:a},c.names=["Krovak","krovak"]},{"../common/adjust_lon":5}],49:[function(a,b,c){var d=Math.PI/2,e=Math.PI/4,f=1e-10,g=a("../common/qsfnz"),h=a("../common/adjust_lon");c.S_POLE=1,c.N_POLE=2,c.EQUIT=3,c.OBLIQ=4,c.init=function(){var a=Math.abs(this.lat0);if(Math.abs(a-d)<f?this.mode=this.lat0<0?this.S_POLE:this.N_POLE:Math.abs(a)<f?this.mode=this.EQUIT:this.mode=this.OBLIQ,this.es>0){var b;switch(this.qp=g(this.e,1),this.mmf=.5/(1-this.es),this.apa=this.authset(this.es),this.mode){case this.N_POLE:this.dd=1;break;case this.S_POLE:this.dd=1;break;case this.EQUIT:this.rq=Math.sqrt(.5*this.qp),this.dd=1/this.rq,this.xmf=1,this.ymf=.5*this.qp;break;case this.OBLIQ:this.rq=Math.sqrt(.5*this.qp),b=Math.sin(this.lat0),this.sinb1=g(this.e,b)/this.qp,this.cosb1=Math.sqrt(1-this.sinb1*this.sinb1),this.dd=Math.cos(this.lat0)/(Math.sqrt(1-this.es*b*b)*this.rq*this.cosb1),this.ymf=(this.xmf=this.rq)/this.dd,this.xmf*=this.dd}}else this.mode===this.OBLIQ&&(this.sinph0=Math.sin(this.lat0),this.cosph0=Math.cos(this.lat0))},c.forward=function(a){var b,c,i,j,k,l,m,n,o,p,q=a.x,r=a.y;if(q=h(q-this.long0),this.sphere){if(k=Math.sin(r),p=Math.cos(r),i=Math.cos(q),this.mode===this.OBLIQ||this.mode===this.EQUIT){if(c=this.mode===this.EQUIT?1+p*i:1+this.sinph0*k+this.cosph0*p*i,f>=c)return null;c=Math.sqrt(2/c),b=c*p*Math.sin(q),c*=this.mode===this.EQUIT?k:this.cosph0*k-this.sinph0*p*i}else if(this.mode===this.N_POLE||this.mode===this.S_POLE){if(this.mode===this.N_POLE&&(i=-i),Math.abs(r+this.phi0)<f)return null;c=e-.5*r,c=2*(this.mode===this.S_POLE?Math.cos(c):Math.sin(c)),b=c*Math.sin(q),c*=i}}else{switch(m=0,n=0,o=0,i=Math.cos(q),j=Math.sin(q),k=Math.sin(r),l=g(this.e,k),this.mode!==this.OBLIQ&&this.mode!==this.EQUIT||(m=l/this.qp,n=Math.sqrt(1-m*m)),this.mode){case this.OBLIQ:o=1+this.sinb1*m+this.cosb1*n*i;break;case this.EQUIT:o=1+n*i;break;case this.N_POLE:o=d+r,l=this.qp-l;break;case this.S_POLE:o=r-d,l=this.qp+l}if(Math.abs(o)<f)return null;switch(this.mode){case this.OBLIQ:case this.EQUIT:o=Math.sqrt(2/o),c=this.mode===this.OBLIQ?this.ymf*o*(this.cosb1*m-this.sinb1*n*i):(o=Math.sqrt(2/(1+n*i)))*m*this.ymf,b=this.xmf*o*n*j;break;case this.N_POLE:case this.S_POLE:l>=0?(b=(o=Math.sqrt(l))*j,c=i*(this.mode===this.S_POLE?o:-o)):b=c=0}}return a.x=this.a*b+this.x0,a.y=this.a*c+this.y0,a},c.inverse=function(a){a.x-=this.x0,a.y-=this.y0;var b,c,e,g,i,j,k,l=a.x/this.a,m=a.y/this.a;if(this.sphere){var n,o=0,p=0;if(n=Math.sqrt(l*l+m*m),c=.5*n,c>1)return null;switch(c=2*Math.asin(c),this.mode!==this.OBLIQ&&this.mode!==this.EQUIT||(p=Math.sin(c),o=Math.cos(c)),this.mode){case this.EQUIT:c=Math.abs(n)<=f?0:Math.asin(m*p/n),l*=p,m=o*n;break;case this.OBLIQ:c=Math.abs(n)<=f?this.phi0:Math.asin(o*this.sinph0+m*p*this.cosph0/n),l*=p*this.cosph0,m=(o-Math.sin(c)*this.sinph0)*n;break;case this.N_POLE:m=-m,c=d-c;break;case this.S_POLE:c-=d}b=0!==m||this.mode!==this.EQUIT&&this.mode!==this.OBLIQ?Math.atan2(l,m):0}else{if(k=0,this.mode===this.OBLIQ||this.mode===this.EQUIT){if(l/=this.dd,m*=this.dd,j=Math.sqrt(l*l+m*m),f>j)return a.x=0,a.y=this.phi0,a;g=2*Math.asin(.5*j/this.rq),e=Math.cos(g),l*=g=Math.sin(g),this.mode===this.OBLIQ?(k=e*this.sinb1+m*g*this.cosb1/j,i=this.qp*k,m=j*this.cosb1*e-m*this.sinb1*g):(k=m*g/j,i=this.qp*k,m=j*e)}else if(this.mode===this.N_POLE||this.mode===this.S_POLE){if(this.mode===this.N_POLE&&(m=-m),i=l*l+m*m,!i)return a.x=0,a.y=this.phi0,a;k=1-i/this.qp,this.mode===this.S_POLE&&(k=-k)}b=Math.atan2(l,m),c=this.authlat(Math.asin(k),this.apa)}return a.x=h(this.long0+b),a.y=c,a},c.P00=.3333333333333333,c.P01=.17222222222222222,c.P02=.10257936507936508,c.P10=.06388888888888888,c.P11=.0664021164021164,c.P20=.016415012942191543,c.authset=function(a){var b,c=[];return c[0]=a*this.P00,b=a*a,c[0]+=b*this.P01,c[1]=b*this.P10,b*=a,c[0]+=b*this.P02,c[1]+=b*this.P11,c[2]=b*this.P20,c},c.authlat=function(a,b){var c=a+a;return a+b[0]*Math.sin(c)+b[1]*Math.sin(c+c)+b[2]*Math.sin(c+c+c)},c.names=["Lambert Azimuthal Equal Area","Lambert_Azimuthal_Equal_Area","laea"]},{"../common/adjust_lon":5,"../common/qsfnz":20}],50:[function(a,b,c){var d=1e-10,e=a("../common/msfnz"),f=a("../common/tsfnz"),g=Math.PI/2,h=a("../common/sign"),i=a("../common/adjust_lon"),j=a("../common/phi2z");c.init=function(){if(this.lat2||(this.lat2=this.lat1),this.k0||(this.k0=1),this.x0=this.x0||0,this.y0=this.y0||0,!(Math.abs(this.lat1+this.lat2)<d)){var a=this.b/this.a;this.e=Math.sqrt(1-a*a);var b=Math.sin(this.lat1),c=Math.cos(this.lat1),g=e(this.e,b,c),h=f(this.e,this.lat1,b),i=Math.sin(this.lat2),j=Math.cos(this.lat2),k=e(this.e,i,j),l=f(this.e,this.lat2,i),m=f(this.e,this.lat0,Math.sin(this.lat0));Math.abs(this.lat1-this.lat2)>d?this.ns=Math.log(g/k)/Math.log(h/l):this.ns=b,isNaN(this.ns)&&(this.ns=b),this.f0=g/(this.ns*Math.pow(h,this.ns)),this.rh=this.a*this.f0*Math.pow(m,this.ns),this.title||(this.title="Lambert Conformal Conic")}},c.forward=function(a){var b=a.x,c=a.y;Math.abs(2*Math.abs(c)-Math.PI)<=d&&(c=h(c)*(g-2*d));var e,j,k=Math.abs(Math.abs(c)-g);if(k>d)e=f(this.e,c,Math.sin(c)),j=this.a*this.f0*Math.pow(e,this.ns);else{if(k=c*this.ns,0>=k)return null;j=0}var l=this.ns*i(b-this.long0);return a.x=this.k0*(j*Math.sin(l))+this.x0,a.y=this.k0*(this.rh-j*Math.cos(l))+this.y0,a},c.inverse=function(a){var b,c,d,e,f,h=(a.x-this.x0)/this.k0,k=this.rh-(a.y-this.y0)/this.k0;this.ns>0?(b=Math.sqrt(h*h+k*k),c=1):(b=-Math.sqrt(h*h+k*k),c=-1);var l=0;if(0!==b&&(l=Math.atan2(c*h,c*k)),0!==b||this.ns>0){if(c=1/this.ns,d=Math.pow(b/(this.a*this.f0),c),e=j(this.e,d),-9999===e)return null}else e=-g;return f=i(l/this.ns+this.long0),a.x=f,a.y=e,a},c.names=["Lambert Tangential Conformal Conic Projection","Lambert_Conformal_Conic","Lambert_Conformal_Conic_2SP","lcc"]},{"../common/adjust_lon":5,"../common/msfnz":15,"../common/phi2z":16,"../common/sign":21,"../common/tsfnz":24}],51:[function(a,b,c){function d(a){return a}c.init=function(){},c.forward=d,c.inverse=d,c.names=["longlat","identity"]},{}],52:[function(a,b,c){var d=a("../common/msfnz"),e=Math.PI/2,f=1e-10,g=57.29577951308232,h=a("../common/adjust_lon"),i=Math.PI/4,j=a("../common/tsfnz"),k=a("../common/phi2z");c.init=function(){var a=this.b/this.a;this.es=1-a*a,"x0"in this||(this.x0=0),"y0"in this||(this.y0=0),this.e=Math.sqrt(this.es),this.lat_ts?this.sphere?this.k0=Math.cos(this.lat_ts):this.k0=d(this.e,Math.sin(this.lat_ts),Math.cos(this.lat_ts)):this.k0||(this.k?this.k0=this.k:this.k0=1)},c.forward=function(a){var b=a.x,c=a.y;if(c*g>90&&-90>c*g&&b*g>180&&-180>b*g)return null;var d,k;if(Math.abs(Math.abs(c)-e)<=f)return null;if(this.sphere)d=this.x0+this.a*this.k0*h(b-this.long0),k=this.y0+this.a*this.k0*Math.log(Math.tan(i+.5*c));else{var l=Math.sin(c),m=j(this.e,c,l);d=this.x0+this.a*this.k0*h(b-this.long0),k=this.y0-this.a*this.k0*Math.log(m)}return a.x=d,a.y=k,a},c.inverse=function(a){var b,c,d=a.x-this.x0,f=a.y-this.y0;if(this.sphere)c=e-2*Math.atan(Math.exp(-f/(this.a*this.k0)));else{var g=Math.exp(-f/(this.a*this.k0));if(c=k(this.e,g),-9999===c)return null}return b=h(this.long0+d/(this.a*this.k0)),a.x=b,a.y=c,a},c.names=["Mercator","Popular Visualisation Pseudo Mercator","Mercator_1SP","Mercator_Auxiliary_Sphere","merc"]},{"../common/adjust_lon":5,"../common/msfnz":15,"../common/phi2z":16,"../common/tsfnz":24}],53:[function(a,b,c){var d=a("../common/adjust_lon");c.init=function(){},c.forward=function(a){var b=a.x,c=a.y,e=d(b-this.long0),f=this.x0+this.a*e,g=this.y0+this.a*Math.log(Math.tan(Math.PI/4+c/2.5))*1.25;return a.x=f,a.y=g,a},c.inverse=function(a){a.x-=this.x0,a.y-=this.y0;var b=d(this.long0+a.x/this.a),c=2.5*(Math.atan(Math.exp(.8*a.y/this.a))-Math.PI/4);return a.x=b,a.y=c,a},c.names=["Miller_Cylindrical","mill"]},{"../common/adjust_lon":5}],54:[function(a,b,c){var d=a("../common/adjust_lon"),e=1e-10;c.init=function(){},c.forward=function(a){for(var b=a.x,c=a.y,f=d(b-this.long0),g=c,h=Math.PI*Math.sin(c),i=0;!0;i++){var j=-(g+Math.sin(g)-h)/(1+Math.cos(g));if(g+=j,Math.abs(j)<e)break}g/=2,Math.PI/2-Math.abs(c)<e&&(f=0);var k=.900316316158*this.a*f*Math.cos(g)+this.x0,l=1.4142135623731*this.a*Math.sin(g)+this.y0;return a.x=k,a.y=l,a},c.inverse=function(a){var b,c;a.x-=this.x0,a.y-=this.y0,c=a.y/(1.4142135623731*this.a),Math.abs(c)>.999999999999&&(c=.999999999999),b=Math.asin(c);var e=d(this.long0+a.x/(.900316316158*this.a*Math.cos(b)));e<-Math.PI&&(e=-Math.PI),e>Math.PI&&(e=Math.PI),c=(2*b+Math.sin(2*b))/Math.PI,Math.abs(c)>1&&(c=1);var f=Math.asin(c);return a.x=e,a.y=f,a},c.names=["Mollweide","moll"]},{"../common/adjust_lon":5}],55:[function(a,b,c){var d=484813681109536e-20;c.iterations=1,c.init=function(){this.A=[],this.A[1]=.6399175073,this.A[2]=-.1358797613,this.A[3]=.063294409,this.A[4]=-.02526853,this.A[5]=.0117879,this.A[6]=-.0055161,this.A[7]=.0026906,this.A[8]=-.001333,this.A[9]=67e-5,this.A[10]=-34e-5,this.B_re=[],this.B_im=[],this.B_re[1]=.7557853228,this.B_im[1]=0,this.B_re[2]=.249204646,this.B_im[2]=.003371507,this.B_re[3]=-.001541739,this.B_im[3]=.04105856,this.B_re[4]=-.10162907,this.B_im[4]=.01727609,this.B_re[5]=-.26623489,this.B_im[5]=-.36249218,this.B_re[6]=-.6870983,this.B_im[6]=-1.1651967,this.C_re=[],this.C_im=[],this.C_re[1]=1.3231270439,this.C_im[1]=0,this.C_re[2]=-.577245789,this.C_im[2]=-.007809598,this.C_re[3]=.508307513,this.C_im[3]=-.112208952,this.C_re[4]=-.15094762,this.C_im[4]=.18200602,this.C_re[5]=1.01418179,this.C_im[5]=1.64497696,this.C_re[6]=1.9660549,this.C_im[6]=2.5127645,this.D=[],this.D[1]=1.5627014243,this.D[2]=.5185406398,this.D[3]=-.03333098,this.D[4]=-.1052906,this.D[5]=-.0368594,this.D[6]=.007317,this.D[7]=.0122,this.D[8]=.00394,this.D[9]=-.0013},c.forward=function(a){var b,c=a.x,e=a.y,f=e-this.lat0,g=c-this.long0,h=f/d*1e-5,i=g,j=1,k=0;for(b=1;10>=b;b++)j*=h,k+=this.A[b]*j;var l,m,n=k,o=i,p=1,q=0,r=0,s=0;for(b=1;6>=b;b++)l=p*n-q*o,m=q*n+p*o,p=l,q=m,r=r+this.B_re[b]*p-this.B_im[b]*q,s=s+this.B_im[b]*p+this.B_re[b]*q;return a.x=s*this.a+this.x0,a.y=r*this.a+this.y0,a},c.inverse=function(a){var b,c,e,f=a.x,g=a.y,h=f-this.x0,i=g-this.y0,j=i/this.a,k=h/this.a,l=1,m=0,n=0,o=0;for(b=1;6>=b;b++)c=l*j-m*k,e=m*j+l*k,l=c,m=e,n=n+this.C_re[b]*l-this.C_im[b]*m,o=o+this.C_im[b]*l+this.C_re[b]*m;for(var p=0;p<this.iterations;p++){var q,r,s=n,t=o,u=j,v=k;for(b=2;6>=b;b++)q=s*n-t*o,r=t*n+s*o,s=q,t=r,u+=(b-1)*(this.B_re[b]*s-this.B_im[b]*t),v+=(b-1)*(this.B_im[b]*s+this.B_re[b]*t);s=1,t=0;var w=this.B_re[1],x=this.B_im[1];for(b=2;6>=b;b++)q=s*n-t*o,r=t*n+s*o,s=q,t=r,w+=b*(this.B_re[b]*s-this.B_im[b]*t),x+=b*(this.B_im[b]*s+this.B_re[b]*t);var y=w*w+x*x;n=(u*w+v*x)/y,o=(v*w-u*x)/y}var z=n,A=o,B=1,C=0;for(b=1;9>=b;b++)B*=z,C+=this.D[b]*B;var D=this.lat0+C*d*1e5,E=this.long0+A;return a.x=E,a.y=D,a},c.names=["New_Zealand_Map_Grid","nzmg"]},{}],56:[function(a,b,c){var d=a("../common/tsfnz"),e=a("../common/adjust_lon"),f=a("../common/phi2z"),g=Math.PI/2,h=Math.PI/4,i=1e-10;c.init=function(){this.no_off=this.no_off||!1,this.no_rot=this.no_rot||!1,isNaN(this.k0)&&(this.k0=1);var a=Math.sin(this.lat0),b=Math.cos(this.lat0),c=this.e*a;this.bl=Math.sqrt(1+this.es/(1-this.es)*Math.pow(b,4)),this.al=this.a*this.bl*this.k0*Math.sqrt(1-this.es)/(1-c*c);var f=d(this.e,this.lat0,a),g=this.bl/b*Math.sqrt((1-this.es)/(1-c*c));1>g*g&&(g=1);var h,i;if(isNaN(this.longc)){var j=d(this.e,this.lat1,Math.sin(this.lat1)),k=d(this.e,this.lat2,Math.sin(this.lat2));this.lat0>=0?this.el=(g+Math.sqrt(g*g-1))*Math.pow(f,this.bl):this.el=(g-Math.sqrt(g*g-1))*Math.pow(f,this.bl);var l=Math.pow(j,this.bl),m=Math.pow(k,this.bl);h=this.el/l,i=.5*(h-1/h);var n=(this.el*this.el-m*l)/(this.el*this.el+m*l),o=(m-l)/(m+l),p=e(this.long1-this.long2);this.long0=.5*(this.long1+this.long2)-Math.atan(n*Math.tan(.5*this.bl*p)/o)/this.bl,this.long0=e(this.long0);var q=e(this.long1-this.long0);this.gamma0=Math.atan(Math.sin(this.bl*q)/i),this.alpha=Math.asin(g*Math.sin(this.gamma0))}else h=this.lat0>=0?g+Math.sqrt(g*g-1):g-Math.sqrt(g*g-1),this.el=h*Math.pow(f,this.bl),i=.5*(h-1/h),this.gamma0=Math.asin(Math.sin(this.alpha)/g),this.long0=this.longc-Math.asin(i*Math.tan(this.gamma0))/this.bl;this.no_off?this.uc=0:this.lat0>=0?this.uc=this.al/this.bl*Math.atan2(Math.sqrt(g*g-1),Math.cos(this.alpha)):this.uc=-1*this.al/this.bl*Math.atan2(Math.sqrt(g*g-1),Math.cos(this.alpha))},c.forward=function(a){var b,c,f,j=a.x,k=a.y,l=e(j-this.long0);if(Math.abs(Math.abs(k)-g)<=i)f=k>0?-1:1,c=this.al/this.bl*Math.log(Math.tan(h+f*this.gamma0*.5)),b=-1*f*g*this.al/this.bl;else{var m=d(this.e,k,Math.sin(k)),n=this.el/Math.pow(m,this.bl),o=.5*(n-1/n),p=.5*(n+1/n),q=Math.sin(this.bl*l),r=(o*Math.sin(this.gamma0)-q*Math.cos(this.gamma0))/p;c=Math.abs(Math.abs(r)-1)<=i?Number.POSITIVE_INFINITY:.5*this.al*Math.log((1-r)/(1+r))/this.bl,b=Math.abs(Math.cos(this.bl*l))<=i?this.al*this.bl*l:this.al*Math.atan2(o*Math.cos(this.gamma0)+q*Math.sin(this.gamma0),Math.cos(this.bl*l))/this.bl}return this.no_rot?(a.x=this.x0+b,a.y=this.y0+c):(b-=this.uc,a.x=this.x0+c*Math.cos(this.alpha)+b*Math.sin(this.alpha),a.y=this.y0+b*Math.cos(this.alpha)-c*Math.sin(this.alpha)),a},c.inverse=function(a){var b,c;this.no_rot?(c=a.y-this.y0,b=a.x-this.x0):(c=(a.x-this.x0)*Math.cos(this.alpha)-(a.y-this.y0)*Math.sin(this.alpha),b=(a.y-this.y0)*Math.cos(this.alpha)+(a.x-this.x0)*Math.sin(this.alpha),b+=this.uc);var d=Math.exp(-1*this.bl*c/this.al),h=.5*(d-1/d),j=.5*(d+1/d),k=Math.sin(this.bl*b/this.al),l=(k*Math.cos(this.gamma0)+h*Math.sin(this.gamma0))/j,m=Math.pow(this.el/Math.sqrt((1+l)/(1-l)),1/this.bl);return Math.abs(l-1)<i?(a.x=this.long0,a.y=g):Math.abs(l+1)<i?(a.x=this.long0,a.y=-1*g):(a.y=f(this.e,m),a.x=e(this.long0-Math.atan2(h*Math.cos(this.gamma0)-k*Math.sin(this.gamma0),Math.cos(this.bl*b/this.al))/this.bl)),a},c.names=["Hotine_Oblique_Mercator","Hotine Oblique Mercator","Hotine_Oblique_Mercator_Azimuth_Natural_Origin","Hotine_Oblique_Mercator_Azimuth_Center","omerc"]},{"../common/adjust_lon":5,"../common/phi2z":16,"../common/tsfnz":24}],57:[function(a,b,c){var d=a("../common/adjust_lon"),e=1e-10,f=a("../common/asinz"),g=Math.PI/2;c.init=function(){this.sin_p14=Math.sin(this.lat0),this.cos_p14=Math.cos(this.lat0)},c.forward=function(a){var b,c,f,g,h,i,j,k,l=a.x,m=a.y;return f=d(l-this.long0),b=Math.sin(m),c=Math.cos(m),g=Math.cos(f),i=this.sin_p14*b+this.cos_p14*c*g,h=1,(i>0||Math.abs(i)<=e)&&(j=this.a*h*c*Math.sin(f),k=this.y0+this.a*h*(this.cos_p14*b-this.sin_p14*c*g)),a.x=j,a.y=k,a},c.inverse=function(a){var b,c,h,i,j,k,l;return a.x-=this.x0,a.y-=this.y0,b=Math.sqrt(a.x*a.x+a.y*a.y),c=f(b/this.a),h=Math.sin(c),i=Math.cos(c),k=this.long0,Math.abs(b)<=e?(l=this.lat0,a.x=k,a.y=l,a):(l=f(i*this.sin_p14+a.y*h*this.cos_p14/b),j=Math.abs(this.lat0)-g,Math.abs(j)<=e?(k=d(this.lat0>=0?this.long0+Math.atan2(a.x,-a.y):this.long0-Math.atan2(-a.x,a.y)),a.x=k,a.y=l,a):(k=d(this.long0+Math.atan2(a.x*h,b*this.cos_p14*i-a.y*this.sin_p14*h)),a.x=k,a.y=l,a))},c.names=["ortho"]},{"../common/adjust_lon":5,"../common/asinz":6}],58:[function(a,b,c){var d=a("../common/e0fn"),e=a("../common/e1fn"),f=a("../common/e2fn"),g=a("../common/e3fn"),h=a("../common/adjust_lon"),i=a("../common/adjust_lat"),j=a("../common/mlfn"),k=1e-10,l=a("../common/gN"),m=20;c.init=function(){this.temp=this.b/this.a,this.es=1-Math.pow(this.temp,2),this.e=Math.sqrt(this.es),this.e0=d(this.es),this.e1=e(this.es),this.e2=f(this.es),this.e3=g(this.es),this.ml0=this.a*j(this.e0,this.e1,this.e2,this.e3,this.lat0)},c.forward=function(a){var b,c,d,e=a.x,f=a.y,g=h(e-this.long0);if(d=g*Math.sin(f),this.sphere)Math.abs(f)<=k?(b=this.a*g,c=-1*this.a*this.lat0):(b=this.a*Math.sin(d)/Math.tan(f),c=this.a*(i(f-this.lat0)+(1-Math.cos(d))/Math.tan(f)));else if(Math.abs(f)<=k)b=this.a*g,c=-1*this.ml0;else{var m=l(this.a,this.e,Math.sin(f))/Math.tan(f);b=m*Math.sin(d),c=this.a*j(this.e0,this.e1,this.e2,this.e3,f)-this.ml0+m*(1-Math.cos(d))}return a.x=b+this.x0,a.y=c+this.y0,a},c.inverse=function(a){var b,c,d,e,f,g,i,l,n;if(d=a.x-this.x0,e=a.y-this.y0,this.sphere)if(Math.abs(e+this.a*this.lat0)<=k)b=h(d/this.a+this.long0),c=0;else{g=this.lat0+e/this.a,i=d*d/this.a/this.a+g*g,l=g;var o;for(f=m;f;--f)if(o=Math.tan(l),n=-1*(g*(l*o+1)-l-.5*(l*l+i)*o)/((l-g)/o-1),l+=n,Math.abs(n)<=k){c=l;break}b=h(this.long0+Math.asin(d*Math.tan(l)/this.a)/Math.sin(c))}else if(Math.abs(e+this.ml0)<=k)c=0,b=h(this.long0+d/this.a);else{g=(this.ml0+e)/this.a,i=d*d/this.a/this.a+g*g,l=g;var p,q,r,s,t;for(f=m;f;--f)if(t=this.e*Math.sin(l),p=Math.sqrt(1-t*t)*Math.tan(l),q=this.a*j(this.e0,this.e1,this.e2,this.e3,l),r=this.e0-2*this.e1*Math.cos(2*l)+4*this.e2*Math.cos(4*l)-6*this.e3*Math.cos(6*l),s=q/this.a,n=(g*(p*s+1)-s-.5*p*(s*s+i))/(this.es*Math.sin(2*l)*(s*s+i-2*g*s)/(4*p)+(g-s)*(p*r-2/Math.sin(2*l))-r),l-=n,Math.abs(n)<=k){c=l;break}p=Math.sqrt(1-this.es*Math.pow(Math.sin(c),2))*Math.tan(c),b=h(this.long0+Math.asin(d*p/this.a)/Math.sin(c))}return a.x=b,a.y=c,a},c.names=["Polyconic","poly"]},{"../common/adjust_lat":4,"../common/adjust_lon":5,"../common/e0fn":7,"../common/e1fn":8,"../common/e2fn":9,"../common/e3fn":10,"../common/gN":11,"../common/mlfn":14}],59:[function(a,b,c){var d=a("../common/adjust_lon"),e=a("../common/adjust_lat"),f=a("../common/pj_enfn"),g=20,h=a("../common/pj_mlfn"),i=a("../common/pj_inv_mlfn"),j=Math.PI/2,k=1e-10,l=a("../common/asinz");c.init=function(){this.sphere?(this.n=1,this.m=0,this.es=0,this.C_y=Math.sqrt((this.m+1)/this.n),this.C_x=this.C_y/(this.m+1)):this.en=f(this.es)},c.forward=function(a){var b,c,e=a.x,f=a.y;if(e=d(e-this.long0),this.sphere){if(this.m)for(var i=this.n*Math.sin(f),j=g;j;--j){var l=(this.m*f+Math.sin(f)-i)/(this.m+Math.cos(f));if(f-=l,Math.abs(l)<k)break}else f=1!==this.n?Math.asin(this.n*Math.sin(f)):f;b=this.a*this.C_x*e*(this.m+Math.cos(f)),c=this.a*this.C_y*f}else{var m=Math.sin(f),n=Math.cos(f);c=this.a*h(f,m,n,this.en),b=this.a*e*n/Math.sqrt(1-this.es*m*m)}return a.x=b,a.y=c,a},c.inverse=function(a){var b,c,f,g;return a.x-=this.x0,f=a.x/this.a,a.y-=this.y0,b=a.y/this.a,this.sphere?(b/=this.C_y,f/=this.C_x*(this.m+Math.cos(b)),this.m?b=l((this.m*b+Math.sin(b))/this.n):1!==this.n&&(b=l(Math.sin(b)/this.n)),f=d(f+this.long0),b=e(b)):(b=i(a.y/this.a,this.es,this.en),g=Math.abs(b),j>g?(g=Math.sin(b),c=this.long0+a.x*Math.sqrt(1-this.es*g*g)/(this.a*Math.cos(b)),f=d(c)):j>g-k&&(f=this.long0)),a.x=f,a.y=b,a},c.names=["Sinusoidal","sinu"]},{"../common/adjust_lat":4,"../common/adjust_lon":5,"../common/asinz":6,"../common/pj_enfn":17,"../common/pj_inv_mlfn":18,"../common/pj_mlfn":19}],60:[function(a,b,c){c.init=function(){var a=this.lat0;this.lambda0=this.long0;var b=Math.sin(a),c=this.a,d=this.rf,e=1/d,f=2*e-Math.pow(e,2),g=this.e=Math.sqrt(f);this.R=this.k0*c*Math.sqrt(1-f)/(1-f*Math.pow(b,2)),this.alpha=Math.sqrt(1+f/(1-f)*Math.pow(Math.cos(a),4)),this.b0=Math.asin(b/this.alpha);var h=Math.log(Math.tan(Math.PI/4+this.b0/2)),i=Math.log(Math.tan(Math.PI/4+a/2)),j=Math.log((1+g*b)/(1-g*b));this.K=h-this.alpha*i+this.alpha*g/2*j},c.forward=function(a){var b=Math.log(Math.tan(Math.PI/4-a.y/2)),c=this.e/2*Math.log((1+this.e*Math.sin(a.y))/(1-this.e*Math.sin(a.y))),d=-this.alpha*(b+c)+this.K,e=2*(Math.atan(Math.exp(d))-Math.PI/4),f=this.alpha*(a.x-this.lambda0),g=Math.atan(Math.sin(f)/(Math.sin(this.b0)*Math.tan(e)+Math.cos(this.b0)*Math.cos(f))),h=Math.asin(Math.cos(this.b0)*Math.sin(e)-Math.sin(this.b0)*Math.cos(e)*Math.cos(f));return a.y=this.R/2*Math.log((1+Math.sin(h))/(1-Math.sin(h)))+this.y0,a.x=this.R*g+this.x0,a},c.inverse=function(a){for(var b=a.x-this.x0,c=a.y-this.y0,d=b/this.R,e=2*(Math.atan(Math.exp(c/this.R))-Math.PI/4),f=Math.asin(Math.cos(this.b0)*Math.sin(e)+Math.sin(this.b0)*Math.cos(e)*Math.cos(d)),g=Math.atan(Math.sin(d)/(Math.cos(this.b0)*Math.cos(d)-Math.sin(this.b0)*Math.tan(e))),h=this.lambda0+g/this.alpha,i=0,j=f,k=-1e3,l=0;Math.abs(j-k)>1e-7;){if(++l>20)return;i=1/this.alpha*(Math.log(Math.tan(Math.PI/4+f/2))-this.K)+this.e*Math.log(Math.tan(Math.PI/4+Math.asin(this.e*Math.sin(j))/2)),k=j,j=2*Math.atan(Math.exp(i))-Math.PI/2}return a.x=h,a.y=j,a},c.names=["somerc"]},{}],61:[function(a,b,c){var d=Math.PI/2,e=1e-10,f=a("../common/sign"),g=a("../common/msfnz"),h=a("../common/tsfnz"),i=a("../common/phi2z"),j=a("../common/adjust_lon");c.ssfn_=function(a,b,c){return b*=c,Math.tan(.5*(d+a))*Math.pow((1-b)/(1+b),.5*c)},c.init=function(){this.coslat0=Math.cos(this.lat0),this.sinlat0=Math.sin(this.lat0),this.sphere?1===this.k0&&!isNaN(this.lat_ts)&&Math.abs(this.coslat0)<=e&&(this.k0=.5*(1+f(this.lat0)*Math.sin(this.lat_ts))):(Math.abs(this.coslat0)<=e&&(this.lat0>0?this.con=1:this.con=-1),this.cons=Math.sqrt(Math.pow(1+this.e,1+this.e)*Math.pow(1-this.e,1-this.e)),1===this.k0&&!isNaN(this.lat_ts)&&Math.abs(this.coslat0)<=e&&(this.k0=.5*this.cons*g(this.e,Math.sin(this.lat_ts),Math.cos(this.lat_ts))/h(this.e,this.con*this.lat_ts,this.con*Math.sin(this.lat_ts))),this.ms1=g(this.e,this.sinlat0,this.coslat0),this.X0=2*Math.atan(this.ssfn_(this.lat0,this.sinlat0,this.e))-d,this.cosX0=Math.cos(this.X0),this.sinX0=Math.sin(this.X0))},c.forward=function(a){var b,c,f,g,i,k,l=a.x,m=a.y,n=Math.sin(m),o=Math.cos(m),p=j(l-this.long0);return Math.abs(Math.abs(l-this.long0)-Math.PI)<=e&&Math.abs(m+this.lat0)<=e?(a.x=NaN,a.y=NaN,a):this.sphere?(b=2*this.k0/(1+this.sinlat0*n+this.coslat0*o*Math.cos(p)),a.x=this.a*b*o*Math.sin(p)+this.x0,a.y=this.a*b*(this.coslat0*n-this.sinlat0*o*Math.cos(p))+this.y0,a):(c=2*Math.atan(this.ssfn_(m,n,this.e))-d,g=Math.cos(c),f=Math.sin(c),Math.abs(this.coslat0)<=e?(i=h(this.e,m*this.con,this.con*n),k=2*this.a*this.k0*i/this.cons,a.x=this.x0+k*Math.sin(l-this.long0),a.y=this.y0-this.con*k*Math.cos(l-this.long0),a):(Math.abs(this.sinlat0)<e?(b=2*this.a*this.k0/(1+g*Math.cos(p)),a.y=b*f):(b=2*this.a*this.k0*this.ms1/(this.cosX0*(1+this.sinX0*f+this.cosX0*g*Math.cos(p))),a.y=b*(this.cosX0*f-this.sinX0*g*Math.cos(p))+this.y0),a.x=b*g*Math.sin(p)+this.x0,a))},c.inverse=function(a){a.x-=this.x0,a.y-=this.y0;var b,c,f,g,h,k=Math.sqrt(a.x*a.x+a.y*a.y);if(this.sphere){var l=2*Math.atan(k/(.5*this.a*this.k0));return b=this.long0,c=this.lat0,e>=k?(a.x=b,a.y=c,a):(c=Math.asin(Math.cos(l)*this.sinlat0+a.y*Math.sin(l)*this.coslat0/k),b=j(Math.abs(this.coslat0)<e?this.lat0>0?this.long0+Math.atan2(a.x,-1*a.y):this.long0+Math.atan2(a.x,a.y):this.long0+Math.atan2(a.x*Math.sin(l),k*this.coslat0*Math.cos(l)-a.y*this.sinlat0*Math.sin(l))),a.x=b,a.y=c,a)}if(Math.abs(this.coslat0)<=e){if(e>=k)return c=this.lat0,b=this.long0,a.x=b,a.y=c,a;a.x*=this.con,a.y*=this.con,f=k*this.cons/(2*this.a*this.k0),c=this.con*i(this.e,f),b=this.con*j(this.con*this.long0+Math.atan2(a.x,-1*a.y))}else g=2*Math.atan(k*this.cosX0/(2*this.a*this.k0*this.ms1)),b=this.long0,e>=k?h=this.X0:(h=Math.asin(Math.cos(g)*this.sinX0+a.y*Math.sin(g)*this.cosX0/k),b=j(this.long0+Math.atan2(a.x*Math.sin(g),k*this.cosX0*Math.cos(g)-a.y*this.sinX0*Math.sin(g)))),c=-1*i(this.e,Math.tan(.5*(d+h)));return a.x=b,a.y=c,a},c.names=["stere","Stereographic_South_Pole","Polar Stereographic (variant B)"]},{"../common/adjust_lon":5,"../common/msfnz":15,"../common/phi2z":16,"../common/sign":21,"../common/tsfnz":24}],62:[function(a,b,c){var d=a("./gauss"),e=a("../common/adjust_lon");c.init=function(){d.init.apply(this),this.rc&&(this.sinc0=Math.sin(this.phic0),this.cosc0=Math.cos(this.phic0),this.R2=2*this.rc,this.title||(this.title="Oblique Stereographic Alternative"))},c.forward=function(a){var b,c,f,g;return a.x=e(a.x-this.long0),d.forward.apply(this,[a]),b=Math.sin(a.y),c=Math.cos(a.y),f=Math.cos(a.x),g=this.k0*this.R2/(1+this.sinc0*b+this.cosc0*c*f),a.x=g*c*Math.sin(a.x),a.y=g*(this.cosc0*b-this.sinc0*c*f),a.x=this.a*a.x+this.x0,a.y=this.a*a.y+this.y0,a},c.inverse=function(a){var b,c,f,g,h;if(a.x=(a.x-this.x0)/this.a,a.y=(a.y-this.y0)/this.a,a.x/=this.k0,a.y/=this.k0,h=Math.sqrt(a.x*a.x+a.y*a.y)){var i=2*Math.atan2(h,this.R2);b=Math.sin(i),c=Math.cos(i),g=Math.asin(c*this.sinc0+a.y*b*this.cosc0/h),f=Math.atan2(a.x*b,h*this.cosc0*c-a.y*this.sinc0*b)}else g=this.phic0,f=0;return a.x=f,a.y=g,d.inverse.apply(this,[a]),a.x=e(a.x+this.long0),a},c.names=["Stereographic_North_Pole","Oblique_Stereographic","Polar_Stereographic","sterea","Oblique Stereographic Alternative"]},{"../common/adjust_lon":5,"./gauss":46}],63:[function(a,b,c){var d=a("../common/e0fn"),e=a("../common/e1fn"),f=a("../common/e2fn"),g=a("../common/e3fn"),h=a("../common/mlfn"),i=a("../common/adjust_lon"),j=Math.PI/2,k=1e-10,l=a("../common/sign"),m=a("../common/asinz");c.init=function(){this.e0=d(this.es),this.e1=e(this.es),this.e2=f(this.es),this.e3=g(this.es),this.ml0=this.a*h(this.e0,this.e1,this.e2,this.e3,this.lat0)},c.forward=function(a){var b,c,d,e=a.x,f=a.y,g=i(e-this.long0),j=Math.sin(f),k=Math.cos(f);if(this.sphere){var l=k*Math.sin(g);if(Math.abs(Math.abs(l)-1)<1e-10)return 93;c=.5*this.a*this.k0*Math.log((1+l)/(1-l)),b=Math.acos(k*Math.cos(g)/Math.sqrt(1-l*l)),0>f&&(b=-b),d=this.a*this.k0*(b-this.lat0)}else{var m=k*g,n=Math.pow(m,2),o=this.ep2*Math.pow(k,2),p=Math.tan(f),q=Math.pow(p,2);b=1-this.es*Math.pow(j,2);var r=this.a/Math.sqrt(b),s=this.a*h(this.e0,this.e1,this.e2,this.e3,f);c=this.k0*r*m*(1+n/6*(1-q+o+n/20*(5-18*q+Math.pow(q,2)+72*o-58*this.ep2)))+this.x0,d=this.k0*(s-this.ml0+r*p*(n*(.5+n/24*(5-q+9*o+4*Math.pow(o,2)+n/30*(61-58*q+Math.pow(q,2)+600*o-330*this.ep2)))))+this.y0}return a.x=c,a.y=d,a},c.inverse=function(a){var b,c,d,e,f,g,h=6;if(this.sphere){var n=Math.exp(a.x/(this.a*this.k0)),o=.5*(n-1/n),p=this.lat0+a.y/(this.a*this.k0),q=Math.cos(p);b=Math.sqrt((1-q*q)/(1+o*o)),f=m(b),0>p&&(f=-f),g=0===o&&0===q?this.long0:i(Math.atan2(o,q)+this.long0)}else{var r=a.x-this.x0,s=a.y-this.y0;for(b=(this.ml0+s/this.k0)/this.a,c=b,e=0;!0&&(d=(b+this.e1*Math.sin(2*c)-this.e2*Math.sin(4*c)+this.e3*Math.sin(6*c))/this.e0-c,c+=d,!(Math.abs(d)<=k));e++)if(e>=h)return 95;if(Math.abs(c)<j){var t=Math.sin(c),u=Math.cos(c),v=Math.tan(c),w=this.ep2*Math.pow(u,2),x=Math.pow(w,2),y=Math.pow(v,2),z=Math.pow(y,2);b=1-this.es*Math.pow(t,2);var A=this.a/Math.sqrt(b),B=A*(1-this.es)/b,C=r/(A*this.k0),D=Math.pow(C,2);f=c-A*v*D/B*(.5-D/24*(5+3*y+10*w-4*x-9*this.ep2-D/30*(61+90*y+298*w+45*z-252*this.ep2-3*x))),g=i(this.long0+C*(1-D/6*(1+2*y+w-D/20*(5-2*w+28*y-3*x+8*this.ep2+24*z)))/u)}else f=j*l(s),g=this.long0}return a.x=g,a.y=f,a},c.names=["Transverse_Mercator","Transverse Mercator","tmerc"]},{"../common/adjust_lon":5,"../common/asinz":6,"../common/e0fn":7,"../common/e1fn":8,"../common/e2fn":9,"../common/e3fn":10,"../common/mlfn":14,"../common/sign":21}],64:[function(a,b,c){var d=.017453292519943295,e=a("./tmerc");c.dependsOn="tmerc",c.init=function(){this.zone&&(this.lat0=0,this.long0=(6*Math.abs(this.zone)-183)*d,this.x0=5e5,this.y0=this.utmSouth?1e7:0,this.k0=.9996,e.init.apply(this),this.forward=e.forward,this.inverse=e.inverse)},c.names=["Universal Transverse Mercator System","utm"]},{"./tmerc":63}],65:[function(a,b,c){var d=a("../common/adjust_lon"),e=Math.PI/2,f=1e-10,g=a("../common/asinz");c.init=function(){this.R=this.a},c.forward=function(a){var b,c,h=a.x,i=a.y,j=d(h-this.long0);Math.abs(i)<=f&&(b=this.x0+this.R*j,c=this.y0);var k=g(2*Math.abs(i/Math.PI));(Math.abs(j)<=f||Math.abs(Math.abs(i)-e)<=f)&&(b=this.x0,c=i>=0?this.y0+Math.PI*this.R*Math.tan(.5*k):this.y0+Math.PI*this.R*-Math.tan(.5*k));var l=.5*Math.abs(Math.PI/j-j/Math.PI),m=l*l,n=Math.sin(k),o=Math.cos(k),p=o/(n+o-1),q=p*p,r=p*(2/n-1),s=r*r,t=Math.PI*this.R*(l*(p-s)+Math.sqrt(m*(p-s)*(p-s)-(s+m)*(q-s)))/(s+m);0>j&&(t=-t),b=this.x0+t;var u=m+p;return t=Math.PI*this.R*(r*u-l*Math.sqrt((s+m)*(m+1)-u*u))/(s+m),c=i>=0?this.y0+t:this.y0-t,a.x=b,a.y=c,a},c.inverse=function(a){var b,c,e,g,h,i,j,k,l,m,n,o,p;return a.x-=this.x0,a.y-=this.y0,n=Math.PI*this.R,e=a.x/n,g=a.y/n,h=e*e+g*g,i=-Math.abs(g)*(1+h),
j=i-2*g*g+e*e,k=-2*i+1+2*g*g+h*h,p=g*g/k+(2*j*j*j/k/k/k-9*i*j/k/k)/27,l=(i-j*j/3/k)/k,m=2*Math.sqrt(-l/3),n=3*p/l/m,Math.abs(n)>1&&(n=n>=0?1:-1),o=Math.acos(n)/3,c=a.y>=0?(-m*Math.cos(o+Math.PI/3)-j/3/k)*Math.PI:-(-m*Math.cos(o+Math.PI/3)-j/3/k)*Math.PI,b=Math.abs(e)<f?this.long0:d(this.long0+Math.PI*(h-1+Math.sqrt(1+2*(e*e-g*g)+h*h))/2/e),a.x=b,a.y=c,a},c.names=["Van_der_Grinten_I","VanDerGrinten","vandg"]},{"../common/adjust_lon":5,"../common/asinz":6}],66:[function(a,b,c){var d=.017453292519943295,e=57.29577951308232,f=1,g=2,h=a("./datum_transform"),i=a("./adjust_axis"),j=a("./Proj"),k=a("./common/toPoint");b.exports=function l(a,b,c){function m(a,b){return(a.datum.datum_type===f||a.datum.datum_type===g)&&"WGS84"!==b.datumCode}var n;return Array.isArray(c)&&(c=k(c)),a.datum&&b.datum&&(m(a,b)||m(b,a))&&(n=new j("WGS84"),l(a,n,c),a=n),"enu"!==a.axis&&i(a,!1,c),"longlat"===a.projName?(c.x*=d,c.y*=d):(a.to_meter&&(c.x*=a.to_meter,c.y*=a.to_meter),a.inverse(c)),a.from_greenwich&&(c.x+=a.from_greenwich),c=h(a.datum,b.datum,c),b.from_greenwich&&(c.x-=b.from_greenwich),"longlat"===b.projName?(c.x*=e,c.y*=e):(b.forward(c),b.to_meter&&(c.x/=b.to_meter,c.y/=b.to_meter)),"enu"!==b.axis&&i(b,!0,c),c}},{"./Proj":2,"./adjust_axis":3,"./common/toPoint":23,"./datum_transform":31}],67:[function(a,b,c){function d(a,b,c){a[b]=c.map(function(a){var b={};return e(a,b),b}).reduce(function(a,b){return j(a,b)},{})}function e(a,b){var c;return Array.isArray(a)?(c=a.shift(),"PARAMETER"===c&&(c=a.shift()),1===a.length?Array.isArray(a[0])?(b[c]={},e(a[0],b[c])):b[c]=a[0]:a.length?"TOWGS84"===c?b[c]=a:(b[c]={},["UNIT","PRIMEM","VERT_DATUM"].indexOf(c)>-1?(b[c]={name:a[0].toLowerCase(),convert:a[1]},3===a.length&&(b[c].auth=a[2])):"SPHEROID"===c?(b[c]={name:a[0],a:a[1],rf:a[2]},4===a.length&&(b[c].auth=a[3])):["GEOGCS","GEOCCS","DATUM","VERT_CS","COMPD_CS","LOCAL_CS","FITTED_CS","LOCAL_DATUM"].indexOf(c)>-1?(a[0]=["name",a[0]],d(b,c,a)):a.every(function(a){return Array.isArray(a)})?d(b,c,a):e(a,b[c])):b[c]=!0,void 0):void(b[a]=!0)}function f(a,b){var c=b[0],d=b[1];!(c in a)&&d in a&&(a[c]=a[d],3===b.length&&(a[c]=b[2](a[c])))}function g(a){return a*i}function h(a){function b(b){var c=a.to_meter||1;return parseFloat(b,10)*c}"GEOGCS"===a.type?a.projName="longlat":"LOCAL_CS"===a.type?(a.projName="identity",a.local=!0):"object"==typeof a.PROJECTION?a.projName=Object.keys(a.PROJECTION)[0]:a.projName=a.PROJECTION,a.UNIT&&(a.units=a.UNIT.name.toLowerCase(),"metre"===a.units&&(a.units="meter"),a.UNIT.convert&&("GEOGCS"===a.type?a.DATUM&&a.DATUM.SPHEROID&&(a.to_meter=parseFloat(a.UNIT.convert,10)*a.DATUM.SPHEROID.a):a.to_meter=parseFloat(a.UNIT.convert,10))),a.GEOGCS&&(a.GEOGCS.DATUM?a.datumCode=a.GEOGCS.DATUM.name.toLowerCase():a.datumCode=a.GEOGCS.name.toLowerCase(),"d_"===a.datumCode.slice(0,2)&&(a.datumCode=a.datumCode.slice(2)),"new_zealand_geodetic_datum_1949"!==a.datumCode&&"new_zealand_1949"!==a.datumCode||(a.datumCode="nzgd49"),"wgs_1984"===a.datumCode&&("Mercator_Auxiliary_Sphere"===a.PROJECTION&&(a.sphere=!0),a.datumCode="wgs84"),"_ferro"===a.datumCode.slice(-6)&&(a.datumCode=a.datumCode.slice(0,-6)),"_jakarta"===a.datumCode.slice(-8)&&(a.datumCode=a.datumCode.slice(0,-8)),~a.datumCode.indexOf("belge")&&(a.datumCode="rnb72"),a.GEOGCS.DATUM&&a.GEOGCS.DATUM.SPHEROID&&(a.ellps=a.GEOGCS.DATUM.SPHEROID.name.replace("_19","").replace(/[Cc]larke\_18/,"clrk"),"international"===a.ellps.toLowerCase().slice(0,13)&&(a.ellps="intl"),a.a=a.GEOGCS.DATUM.SPHEROID.a,a.rf=parseFloat(a.GEOGCS.DATUM.SPHEROID.rf,10)),~a.datumCode.indexOf("osgb_1936")&&(a.datumCode="osgb36")),a.b&&!isFinite(a.b)&&(a.b=a.a);var c=function(b){return f(a,b)},d=[["standard_parallel_1","Standard_Parallel_1"],["standard_parallel_2","Standard_Parallel_2"],["false_easting","False_Easting"],["false_northing","False_Northing"],["central_meridian","Central_Meridian"],["latitude_of_origin","Latitude_Of_Origin"],["latitude_of_origin","Central_Parallel"],["scale_factor","Scale_Factor"],["k0","scale_factor"],["latitude_of_center","Latitude_of_center"],["lat0","latitude_of_center",g],["longitude_of_center","Longitude_Of_Center"],["longc","longitude_of_center",g],["x0","false_easting",b],["y0","false_northing",b],["long0","central_meridian",g],["lat0","latitude_of_origin",g],["lat0","standard_parallel_1",g],["lat1","standard_parallel_1",g],["lat2","standard_parallel_2",g],["alpha","azimuth",g],["srsCode","name"]];d.forEach(c),a.long0||!a.longc||"Albers_Conic_Equal_Area"!==a.projName&&"Lambert_Azimuthal_Equal_Area"!==a.projName||(a.long0=a.longc),a.lat_ts||!a.lat1||"Stereographic_South_Pole"!==a.projName&&"Polar Stereographic (variant B)"!==a.projName||(a.lat0=g(a.lat1>0?90:-90),a.lat_ts=a.lat1)}var i=.017453292519943295,j=a("./extend");b.exports=function(a,b){var c=JSON.parse((","+a).replace(/\s*\,\s*([A-Z_0-9]+?)(\[)/g,',["$1",').slice(1).replace(/\s*\,\s*([A-Z_0-9]+?)\]/g,',"$1"]').replace(/,\["VERTCS".+/,"")),d=c.shift(),f=c.shift();c.unshift(["name",f]),c.unshift(["type",d]),c.unshift("output");var g={};return e(c,g),h(g.output),j(b,g.output)}},{"./extend":34}],68:[function(a,b,c){function d(a){return a*(Math.PI/180)}function e(a){return 180*(a/Math.PI)}function f(a){var b,c,e,f,g,i,j,k,l,m=a.lat,n=a.lon,o=6378137,p=.00669438,q=.9996,r=d(m),s=d(n);l=Math.floor((n+180)/6)+1,180===n&&(l=60),m>=56&&64>m&&n>=3&&12>n&&(l=32),m>=72&&84>m&&(n>=0&&9>n?l=31:n>=9&&21>n?l=33:n>=21&&33>n?l=35:n>=33&&42>n&&(l=37)),b=6*(l-1)-180+3,k=d(b),c=p/(1-p),e=o/Math.sqrt(1-p*Math.sin(r)*Math.sin(r)),f=Math.tan(r)*Math.tan(r),g=c*Math.cos(r)*Math.cos(r),i=Math.cos(r)*(s-k),j=o*((1-p/4-3*p*p/64-5*p*p*p/256)*r-(3*p/8+3*p*p/32+45*p*p*p/1024)*Math.sin(2*r)+(15*p*p/256+45*p*p*p/1024)*Math.sin(4*r)-35*p*p*p/3072*Math.sin(6*r));var t=q*e*(i+(1-f+g)*i*i*i/6+(5-18*f+f*f+72*g-58*c)*i*i*i*i*i/120)+5e5,u=q*(j+e*Math.tan(r)*(i*i/2+(5-f+9*g+4*g*g)*i*i*i*i/24+(61-58*f+f*f+600*g-330*c)*i*i*i*i*i*i/720));return 0>m&&(u+=1e7),{northing:Math.round(u),easting:Math.round(t),zoneNumber:l,zoneLetter:h(m)}}function g(a){var b=a.northing,c=a.easting,d=a.zoneLetter,f=a.zoneNumber;if(0>f||f>60)return null;var h,i,j,k,l,m,n,o,p,q,r=.9996,s=6378137,t=.00669438,u=(1-Math.sqrt(1-t))/(1+Math.sqrt(1-t)),v=c-5e5,w=b;"N">d&&(w-=1e7),o=6*(f-1)-180+3,h=t/(1-t),n=w/r,p=n/(s*(1-t/4-3*t*t/64-5*t*t*t/256)),q=p+(3*u/2-27*u*u*u/32)*Math.sin(2*p)+(21*u*u/16-55*u*u*u*u/32)*Math.sin(4*p)+151*u*u*u/96*Math.sin(6*p),i=s/Math.sqrt(1-t*Math.sin(q)*Math.sin(q)),j=Math.tan(q)*Math.tan(q),k=h*Math.cos(q)*Math.cos(q),l=s*(1-t)/Math.pow(1-t*Math.sin(q)*Math.sin(q),1.5),m=v/(i*r);var x=q-i*Math.tan(q)/l*(m*m/2-(5+3*j+10*k-4*k*k-9*h)*m*m*m*m/24+(61+90*j+298*k+45*j*j-252*h-3*k*k)*m*m*m*m*m*m/720);x=e(x);var y=(m-(1+2*j+k)*m*m*m/6+(5-2*k+28*j-3*k*k+8*h+24*j*j)*m*m*m*m*m/120)/Math.cos(q);y=o+e(y);var z;if(a.accuracy){var A=g({northing:a.northing+a.accuracy,easting:a.easting+a.accuracy,zoneLetter:a.zoneLetter,zoneNumber:a.zoneNumber});z={top:A.lat,right:A.lon,bottom:x,left:y}}else z={lat:x,lon:y};return z}function h(a){var b="Z";return 84>=a&&a>=72?b="X":72>a&&a>=64?b="W":64>a&&a>=56?b="V":56>a&&a>=48?b="U":48>a&&a>=40?b="T":40>a&&a>=32?b="S":32>a&&a>=24?b="R":24>a&&a>=16?b="Q":16>a&&a>=8?b="P":8>a&&a>=0?b="N":0>a&&a>=-8?b="M":-8>a&&a>=-16?b="L":-16>a&&a>=-24?b="K":-24>a&&a>=-32?b="J":-32>a&&a>=-40?b="H":-40>a&&a>=-48?b="G":-48>a&&a>=-56?b="F":-56>a&&a>=-64?b="E":-64>a&&a>=-72?b="D":-72>a&&a>=-80&&(b="C"),b}function i(a,b){var c="00000"+a.easting,d="00000"+a.northing;return a.zoneNumber+a.zoneLetter+j(a.easting,a.northing,a.zoneNumber)+c.substr(c.length-5,b)+d.substr(d.length-5,b)}function j(a,b,c){var d=k(c),e=Math.floor(a/1e5),f=Math.floor(b/1e5)%20;return l(e,f,d)}function k(a){var b=a%q;return 0===b&&(b=q),b}function l(a,b,c){var d=c-1,e=r.charCodeAt(d),f=s.charCodeAt(d),g=e+a-1,h=f+b,i=!1;g>x&&(g=g-x+t-1,i=!0),(g===u||u>e&&g>u||(g>u||u>e)&&i)&&g++,(g===v||v>e&&g>v||(g>v||v>e)&&i)&&(g++,g===u&&g++),g>x&&(g=g-x+t-1),h>w?(h=h-w+t-1,i=!0):i=!1,(h===u||u>f&&h>u||(h>u||u>f)&&i)&&h++,(h===v||v>f&&h>v||(h>v||v>f)&&i)&&(h++,h===u&&h++),h>w&&(h=h-w+t-1);var j=String.fromCharCode(g)+String.fromCharCode(h);return j}function m(a){if(a&&0===a.length)throw"MGRSPoint coverting from nothing";for(var b,c=a.length,d=null,e="",f=0;!/[A-Z]/.test(b=a.charAt(f));){if(f>=2)throw"MGRSPoint bad conversion from: "+a;e+=b,f++}var g=parseInt(e,10);if(0===f||f+3>c)throw"MGRSPoint bad conversion from: "+a;var h=a.charAt(f++);if("A">=h||"B"===h||"Y"===h||h>="Z"||"I"===h||"O"===h)throw"MGRSPoint zone letter "+h+" not handled: "+a;d=a.substring(f,f+=2);for(var i=k(g),j=n(d.charAt(0),i),l=o(d.charAt(1),i);l<p(h);)l+=2e6;var m=c-f;if(m%2!==0)throw"MGRSPoint has to have an even number \nof digits after the zone letter and two 100km letters - front \nhalf for easting meters, second half for \nnorthing meters"+a;var q,r,s,t,u,v=m/2,w=0,x=0;return v>0&&(q=1e5/Math.pow(10,v),r=a.substring(f,f+v),w=parseFloat(r)*q,s=a.substring(f+v),x=parseFloat(s)*q),t=w+j,u=x+l,{easting:t,northing:u,zoneLetter:h,zoneNumber:g,accuracy:q}}function n(a,b){for(var c=r.charCodeAt(b-1),d=1e5,e=!1;c!==a.charCodeAt(0);){if(c++,c===u&&c++,c===v&&c++,c>x){if(e)throw"Bad character: "+a;c=t,e=!0}d+=1e5}return d}function o(a,b){if(a>"V")throw"MGRSPoint given invalid Northing "+a;for(var c=s.charCodeAt(b-1),d=0,e=!1;c!==a.charCodeAt(0);){if(c++,c===u&&c++,c===v&&c++,c>w){if(e)throw"Bad character: "+a;c=t,e=!0}d+=1e5}return d}function p(a){var b;switch(a){case"C":b=11e5;break;case"D":b=2e6;break;case"E":b=28e5;break;case"F":b=37e5;break;case"G":b=46e5;break;case"H":b=55e5;break;case"J":b=64e5;break;case"K":b=73e5;break;case"L":b=82e5;break;case"M":b=91e5;break;case"N":b=0;break;case"P":b=8e5;break;case"Q":b=17e5;break;case"R":b=26e5;break;case"S":b=35e5;break;case"T":b=44e5;break;case"U":b=53e5;break;case"V":b=62e5;break;case"W":b=7e6;break;case"X":b=79e5;break;default:b=-1}if(b>=0)return b;throw"Invalid zone letter: "+a}var q=6,r="AJSAJS",s="AFAFAF",t=65,u=73,v=79,w=86,x=90;c.forward=function(a,b){return b=b||5,i(f({lat:a[1],lon:a[0]}),b)},c.inverse=function(a){var b=g(m(a.toUpperCase()));return b.lat&&b.lon?[b.lon,b.lat,b.lon,b.lat]:[b.left,b.bottom,b.right,b.top]},c.toPoint=function(a){var b=g(m(a.toUpperCase()));return b.lat&&b.lon?[b.lon,b.lat]:[(b.left+b.right)/2,(b.top+b.bottom)/2]}},{}],69:[function(a,b,c){b.exports={name:"proj4",version:"2.3.15",description:"Proj4js is a JavaScript library to transform point coordinates from one coordinate system to another, including datum transformations.",main:"lib/index.js",directories:{test:"test",doc:"docs"},scripts:{test:"./node_modules/istanbul/lib/cli.js test ./node_modules/mocha/bin/_mocha test/test.js"},repository:{type:"git",url:"git://github.com/proj4js/proj4js.git"},author:"",license:"MIT",jam:{main:"dist/proj4.js",include:["dist/proj4.js","README.md","AUTHORS","LICENSE.md"]},devDependencies:{"grunt-cli":"~0.1.13",grunt:"~0.4.2","grunt-contrib-connect":"~0.6.0","grunt-contrib-jshint":"~0.8.0",chai:"~1.8.1",mocha:"~1.17.1","grunt-mocha-phantomjs":"~0.4.0",browserify:"~12.0.1","grunt-browserify":"~4.0.1","grunt-contrib-uglify":"~0.11.1",curl:"git://github.com/cujojs/curl.git",istanbul:"~0.2.4",tin:"~0.4.0"},dependencies:{mgrs:"~0.0.2"}}},{}]},{},[36])(36)});
```
